# Differential gene expression and functional pathway enrichment in colon cell line CCD 841 CoN (CRL-1790) transfected with miR-mimics miR-18b, miR-142-3p, miR-155, and miR-890

**DOI:** 10.1101/747931

**Authors:** JM Robinson

## Abstract

This report is a bioinformatic analysis of samples from NCBI GEO database series GS132501. It includes previously un-reported data for miR-mimic over-expression of the miRNAs miR-18b, miR-155, miR-142-3p, and miR-890 in the ATCC CRL-1790 cell line. Data analysis was performed using Nanostring nSolver 4.0 and Advanced Analysis Module 2.0 plugin (Nanostring MAN-10030-03). Bioinformatic methods utilized include pathway scoring, differential expression (DE), and gene-set enrichment (GSE) analyses. Findings, with full supplementary data, provide a community resource for effects of dysregulation of these miRNAs in a colon ‘epithelial-like’ cell line.

## Main

Robinson et al. (2019) recently reported results from gene expression experiments utilizing a custom, 250-plex Nanostring codeset designed to investigate gene expression associated with tight junction and epithelial regulatory pathways [1]. Correspondingly, full expression data and metadata became publicly available in the NCBI GEO (**GSE132501**), and NCBI Bioproject (accession#: PRJNA525237) databases *(see Note below). The GEO submission also included a novel Platform accession describing probe panel annotations and probe sequences (**GPL26764**). In addition to data reported in Robinson, et al. (2019), the GEO series GSE132501 contains previously un-reported data for miR-mimic over-expression of the miRNAs miR-18b, miR-155, miR-142-3p, and miR-890 (in separate treatments, respectively), in the CCD 841 CoN (ATCC^®^ CRL-1790 ™) cell line.

These CRL-1790 samples (total RNA) were previously tested with the Nanostring human miRNAv3 codeset (https://www.nanostring.com/products/mirna-assays/mirna-panels), with data reported in Joseph et al. (2018) [2]. These miRNA-mimic treatments resulted in differential biological effects during expression of miR-mimics hsa-miR-18b, hsa-miR-155, hsa-miR-142-3p, and hsa-miR-890 on parameters of trans-epithelial resistance (TEER) and expression of the tight junction Occludin(OCLN) and ZO-1(TJP) transcripts. Full methods for the lipofection and RNA extraction methods for these samples are described in Joseph et al. (2018).

Results in this analysis present mRNA expression data for the 4 miR-mimic treatments. This analysis utilizes this publicly available GEO data. The data analysis represents results from Nanostring nSolver 4.0 and Advanced Analysis Module 2.0 plugin, software available free from Nanostring (Nanostring MAN-10030-03). Bioinformatic methods utilized here include pathway scoring, differential expression (DE), and gene-set enrichment (GSE) analyses. Full, html-formatted results are provided (**Sup. Results 1: HTML-formatted complete Nanostring Advanced Analysis Module results**). Samples from *C. elegans* cel-miR-67 transfected cells were used as reference samples for DE-based analyses. Cel-miR-67 is presumed to have no, or insignificant, targets and effects in human cells, other than toxicity from lipofection procedure, or results of miRNA biogenesis pathway overloading. Cel-67 mimic acts as the control for effects of the lipofection, un-transfected controls are also described here to show effects seen in cel-67 transfected cells.

In unsupervised analyses, both clustering (**Fig. 1A**) and principle components analysis (**Fig. 1C**) recover unique expression profiles associated with each respective miR-mimic treatment. miR-cel-67 and Control samples do not form discreet clusters but form a mixed cluster (**Fig. 1A**). Differential expression volcano plots show each respective miR-mimic produces a unique profile of affected mRNAs (**Fig. 1B**).

Functional biological insights can be gained from the pathway analysis results for each respective miR-mimic. A heatmap of pathway scores shows signal specific to each miR-mimic treatment. Note that, in each readout, cel-miR-67 and untreated controls show substantial similarity (**Fig. 2A**).

**Figure 1.**
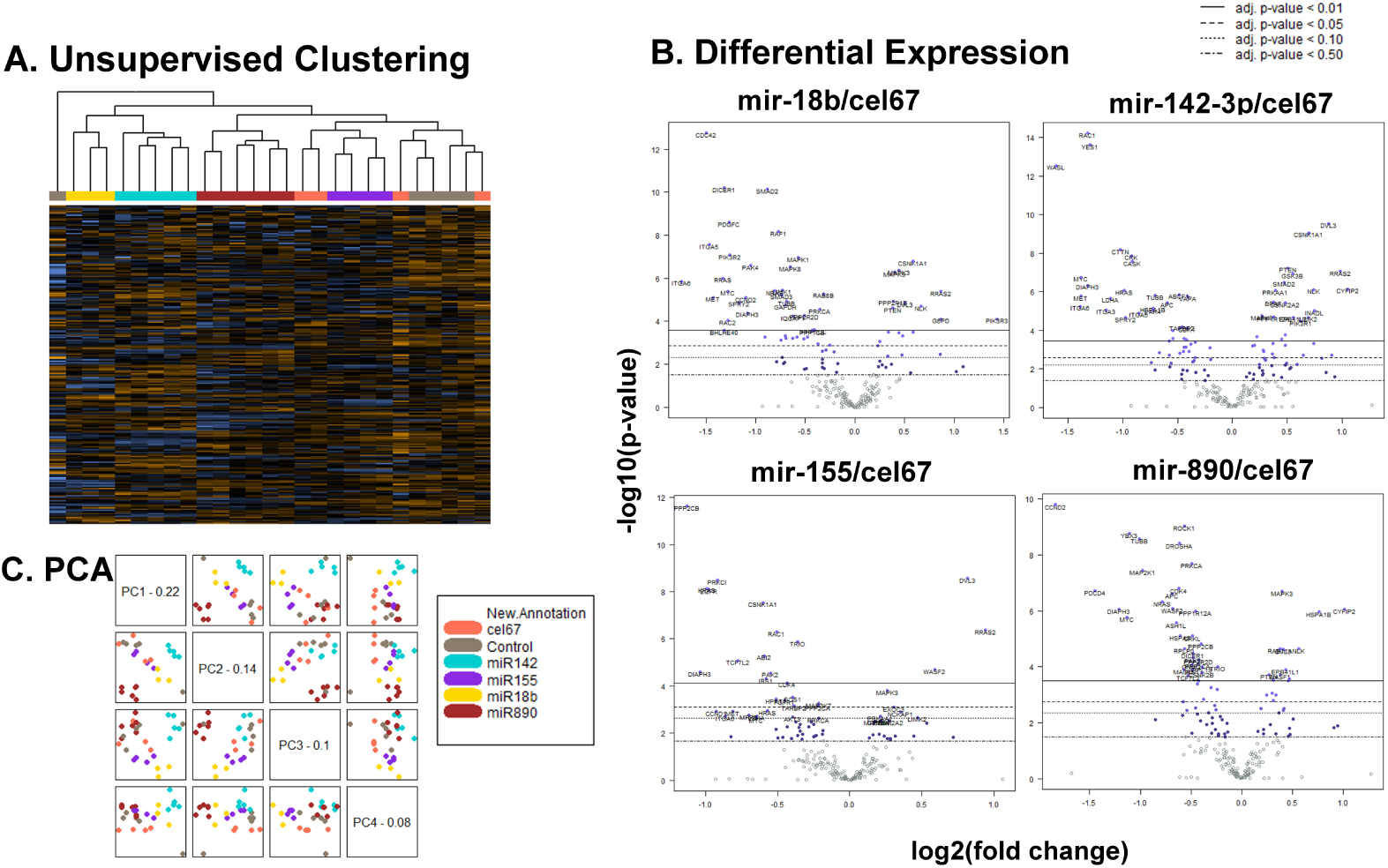

**Figure 2.**
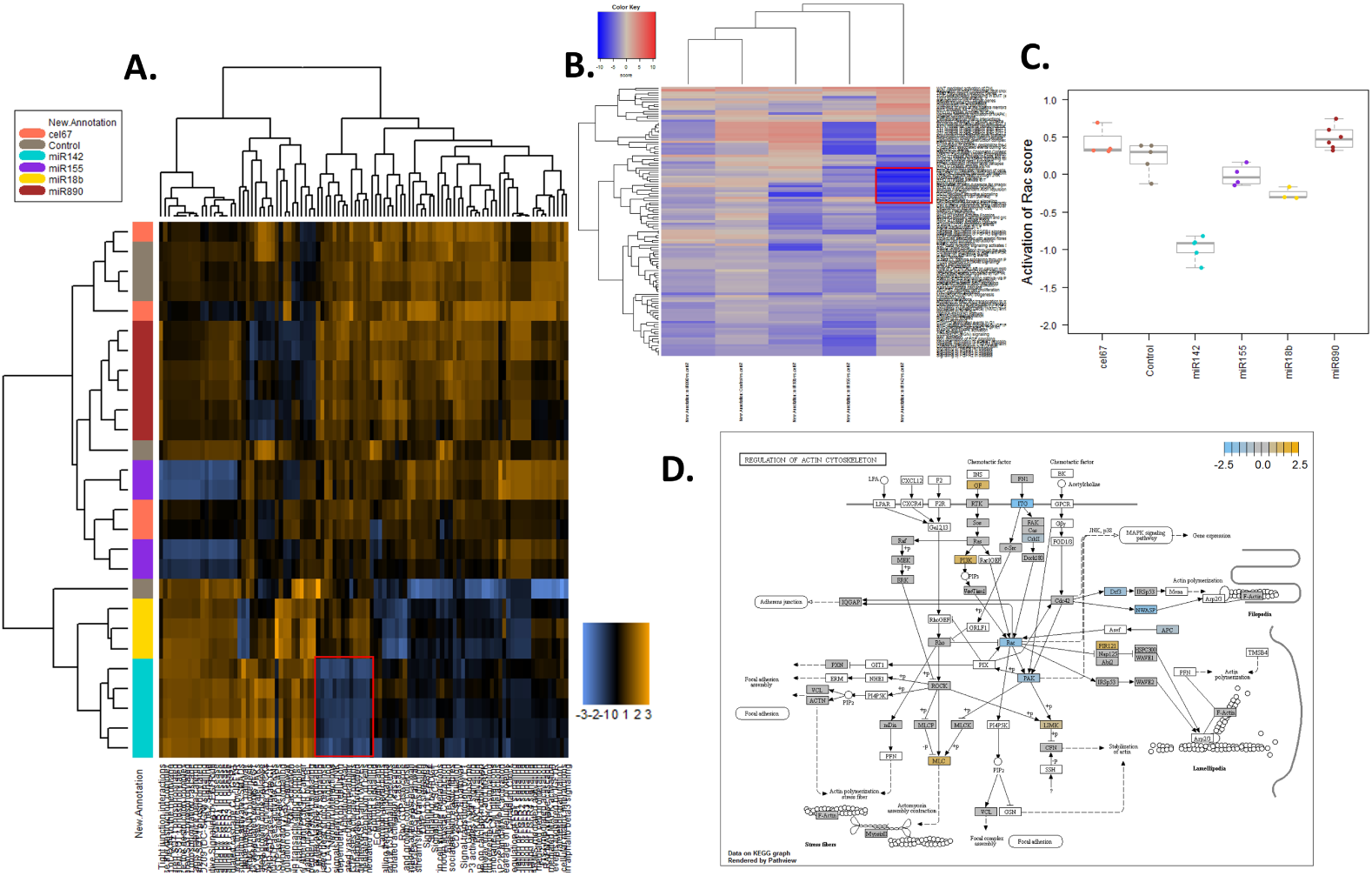
Pathway Scoring and GSE Results for miR-142-3p.

As this is writing represents a short comment on the GEO GSE132501 accession, hsa-miR-142-3p is selected to provide an vignette of potential biological insights for each miRNA of interest. miR-142-3p has been noted for its roles in cancer and in normal biological function, and known for its interactions with the RhoGTPase and actin cytoskeleton dynamics. These include roles in diverse processes of megakaryopoiesis[3], phagocytosis [4], T-cell migration [5]. miR-142-3p is often associated with tumor suppression via the Rac-Rock-Rho GTPase regulatory pathway, in cervical cancer cells [6], osteosarcoma cells [7-9], hepatocellular carcinoma [10], breast cancer stem cells [11], and in renal cell carcinoma cells [12]. miR-142 was also reported as a blood biomarker for fatty liver disease [13], and in Joseph, et al. (2018), resulted in the most significant negative effect on TEER of all miRNAs tested in that report [2].

In this data, miR-142-3p mimic transfection shows strong negative regulation of a specific block of pathway scores (**Fig. 2A**), particularly when compared against pathway scores from other transfected miRs (**Fig. 2B, boxed in red**). One of these affected pathways, *is* the “Activation of Rac” pathway (**Fig. 2C**). Mapping relative expression values on the KEGG pathway database “Regulation of Actin Cytoskeleton” map shows additional effects associated with that pathway, such as decreased expression of integrins, and increased expression of some RTK-pathway members (**Fig. 2D**). It is also interesting to note that miR-18b and miR-890 mimic treatments show the most significant suppression of the miRNA biogenesis pathway, with Dicer being one of the most highly significant differentially expressed genes in both treatments, with Drosha expression also being affected by miR-890 mimic, and multiple pathway transcripts affected by miR-142-3p (**Fig. 3A-C**).

**Figure 3.**
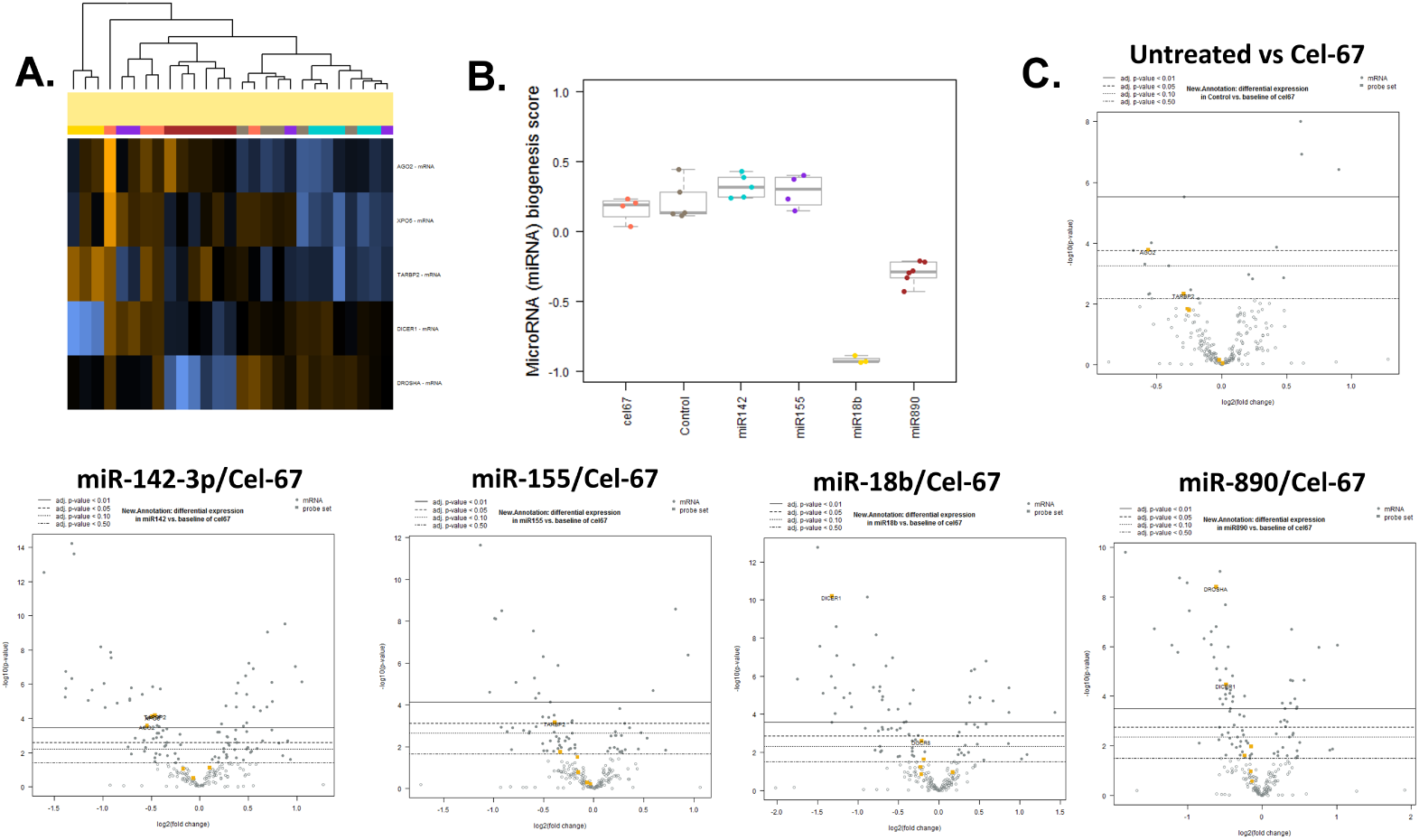
miRNA Biogenesis pathway results.

The ATCC CRL-1790 cell line was derived from human 21-week fetal colon tissue and is described as “epithelial” in terms of having an ‘epithelial-like’ morphology [14]. True epithelial identity of this cell line is questionable, however, as expressing of the key epithelial differentiation marker, cytokeratin, is not seen (https://www.atcc.org/Products/All/CRL-1790.aspx#characteristics). When gene expression is compared with the “true” epithelial lines described in the GSE132501 data, it appears more similar to fibroblasts, than to the epithelial, adenocarcinoma-derived cell lines (unpublished data).

### *Note

Two additional manuscripts associated with the NCBI GEO (**GSE132501**) series are publicly available, including (Robinson et al. 2019), and the bioRxiv pre-print, Robinson (2019) [15]. A production error was made on v1 of [15], where Fig. 1 from the current preprint was substituted for Fig.1 in [15]. This was corrected in a revised and resubmitted v2 of [15].

## Supporting information

Supplementary Results 1

