## Supplementary Results 1 for "Differential gene expression and functional pathway enrichment in colon cell line CCD 841 CoN (CRL-1790) transfected with miR-mimics miR-18b, miR-142-3p, miR-155, and miR-890": AdvAnalysisReport.html

          
   
     
     
     
     nCounter Advanced Analysis Report 
     
     
     
     
     
     
     
     
     
     
     
     
     
     
     
     
     
     
     
     
     
     
     
     
     
     
     
     
     
     
     
     
     
     
     
     
     
     
     
   
   
     
       
         
           
             
               
             
           
         
       
       
         
      
          Overview  
          Normalization  
          Pathway Score  
          Diff Expr  
          GSA  
          PathView  
          Analysis Parameters  
      
      Share   
       
     
     
      
        
          
                This module generates a series of high level plots that describe the data overall and may be useful for identifying anomalous data and/or covariates.     
              
                   Gene sets: 
                  
                      Summary  
                      Activation of BAD and translocation to mitochondria   
                      Activation of Rac  
                      Apoptotic cleavage of cellular proteins  
                      ARMS-mediated activation  
                      AXIN missense mutants destabilize the destruction complex  
                      Beta-catenin phosphorylation cascade  
                      CD209 (DC-SIGN) signaling  
                      CD28 co-stimulation  
                      CD28 dependent PI3KAkt signaling  
                      CD28 dependent Vav1 pathway  
                      Cell-extracellular matrix interactions  
                      Cell surface interactions at the vascular wall  
                      Circadian Clock  
                      Constitutive Signaling by Aberrant PI3K in Cancer  
                      Constitutive Signaling by EGFRvIII  
                      CREB phosphorylation through the activation of Ras  
                      CTLA4 inhibitory signaling  
                      Cyclin AB1 associated events during G2M transition  
                      Cyclin D associated events in G1  
                      DAP12 signaling  
                      DARPP-32 events  
                      DCC mediated attractive signaling  
                      Deactivation of the beta-catenin transactivating complex  
                      Disassembly of the destruction complex and recruitment of AXIN t...  
                      Downstream signal transduction  
                      Downstream TCR signaling  
                      ECM proteoglycans  
                      Elastic fibre formation  
                      EPH-ephrin mediated repulsion of cells  
                      EPHA-mediated growth cone collapse  
                      EPHB-mediated forward signaling  
                      Ephrin signaling  
                      ERKs are inactivated  
                      FCERI mediated MAPK activation  
                      Formation of the beta-catenin.CL.TCF transactivating complex  
                      Frs2-mediated activation  
                      G alpha (1213) signalling events  
                      G alpha (q) signalling events  
                      G beta.CL.gamma signalling through PI3Kgamma  
                      GAB1 signalosome  
                      Gastrin-CREB signalling pathway via PKC and MAPK  
                      GPVI-mediated activation cascade  
                      Integrin alphaIIb beta3 signaling  
                      Integrin cell surface interactions  
                      Interleukin receptor SHC signaling  
                      IRS-related events triggered by IGF1R  
                      Laminin interactions  
                      MAP2K and MAPK activation  
                      MAPK6MAPK4 signaling  
                      MASTL Facilitates Mitotic Progression  
                      MicroRNA (miRNA) biogenesis  
                      Misspliced GSK3beta mutants stabilize beta-catenin  
                      Molecules associated with elastic fibres  
                      Negative feedback regulation of MAPK pathway  
                      Negative regulation of FGFR1 signaling  
                      Negative regulation of FGFR2 signaling  
                      Negative regulation of FGFR3 signaling  
                      Negative regulation of FGFR4 signaling  
                      Negative regulation of MAPK pathway  
                      Nephrin interactions  
                      Non-integrin membrane-ECM interactions  
                      Nonsense Mediated Decay (NMD) enhanced by the Exon Junction Comp...  
                      NRAGE signals death through JNK  
                      p130Cas linkage to MAPK signaling for integrins  
                      PCPCE pathway  
                      PI3K Cascade  
                      PI3KAKT activation  
                      PIP3 activates AKT signaling  
                      Platelet degranulation   
                      RAF activation  
                      RAFMAP kinase cascade  
                      Regulation of actin dynamics for phagocytic cup formation  
                      Regulation of HSF1-mediated heat shock response  
                      Regulation of signaling by CBL  
                      Repression of WNT target genes  
                      Resolution of Sister Chromatid Cohesion  
                      Rho GTPase cycle  
                      RHO GTPases activate CIT  
                      RHO GTPases Activate Formins  
                      RHO GTPases activate IQGAPs  
                      RHO GTPases activate PAKs  
                      RHO GTPases activate PKNs  
                      RHO GTPases Activate ROCKs  
                      RHO GTPases Activate WASPs and WAVEs  
                      Role of LAT2NTALLAB on calcium mobilization  
                      S33 mutants of beta-catenin aren.SQT.t phosphorylated  
                      S37 mutants of beta-catenin aren.SQT.t phosphorylated  
                      S45 mutants of beta-catenin aren.SQT.t phosphorylated  
                      Sema3A PAK dependent Axon repulsion  
                      Sema4D induced cell migration and growth-cone collapse  
                      SHC-mediated cascade.CL.FGFR1  
                      SHC-related events triggered by IGF1R  
                      Signal transduction by L1  
                      Signaling by ERBB2  
                      Signaling by FGFR1 in disease  
                      Signaling by FGFR2 in disease  
                      Signaling by FGFR3 in disease  
                      Signaling by FGFR4 in disease  
                      Signaling by PDGF  
                      Signaling by SCF-KIT  
                      Smooth Muscle Contraction  
                      Spry regulation of FGF signaling  
                      Syndecan interactions  
                      Synthesis of PIPs at the plasma membrane  
                      T41 mutants of beta-catenin aren.SQT.t phosphorylated  
                      TGF-beta receptor signaling activates SMADs  
                      TGF-beta receptor signaling in EMT (epithelial to mesenchymal tr...  
                      Tie2 Signaling  
                      Tight junction interactions  
                      TP53 Regulates Metabolic Genes  
                      Truncations of AMER1 destabilize the destruction complex  
                      VEGFA-VEGFR2 Pathway  
                      VEGFR2 mediated cell proliferation  
                      VEGFR2 mediated vascular permeability  
                      WNT mediated activation of DVL  
                  
                   
                  
                    
                      
                           
                          
                               Heatmaps    PCA    Study design    Other QC     
                              
                                  Heatmaps  
                              
                               
                              
                                  New.Annotation  
                              
                               
                              
                                  Covariates  
                              
                               
                              
                                  Other QC  
                              
                              
                            
                              
                                
                                  
                                       
                                      
                                           Heatmap of Raw Data   More Plot Information      Download detected/undetected calls data                   Heatmap of Raw Data  Heatmap of the raw counts. The plot is meant to provide an overview of how robust the raw expression levels are across samples and gene sets. Datasets that entirely lack higher level expressions (e.g. counts &gt; 100) may indicate experimental failure or low input. The detected/undetected calls links to a .csv file stating whether each probe is above background, with 0/1 indicating below/above background. If the user has not specified a detection threshold, probes are called detected if they have more than double the counts of the median negative control.   
                                           Heatmap of All Data   More Plot Information                   Heatmap of All Data  Heatmap of the normalized data, scaled to give all genes equal variance, generated via unsupervised clustering. Orange indicates high expression; blue indicates low expression. This plot is meant to provide a high level exploratory view of the data.   
                                      
                                       
                                  
                                
                              
                            
                            
                              
                                
                                  
                                       
                                      
                                           Principal Components of All Data   More Plot Information                      Principal Components of All Data  Principal component analysis maps high-dimensional datasets onto a smaller number of highly informative dimensions. Here, the first four principal components of the gene expression data are plotted against each other and colored by the values of the selected covariate. This plot may be used to identify clusters in the data and to identify variables associated with prominent signal in the data. Variables that are associated with these leading principal components should be considered in downstream analyses.   
                                           Outliers   More Plot Information     
                                           
                                             
                                           
                                           
                                             
                                               
                                               Outlier on PC 1 
                                               Outlier on PC 2 
                                               Outlier on PC 3 
                                               Outlier on PC 4 
                                             
                                           
                                           
                                             
                                               20161013_sebelius2-101316_01_01.RCC 
                                               FALSE 
                                               FALSE 
                                               FALSE 
                                               TRUE 
                                             
                                           
                                           Outliers  Table identifying outliers in the first four PCs of the data.   
                                      
                                       
                                  
                                
                              
                            
                            
                              
                                
                                  
                                       
                                      
                                              More Plot Information                  Pairwise comparisons of all covariates in the analysis. The type of plot is dependent on the types of variables compared; A categorical vs. categorical covariate plot is shown as a bar chart of counts (Y axis). Continuous vs. categorical covariates generate a boxplot with whiskers denoting 1.5 IQR. Continuous vs. continuous covariates are compared via a scatter plot. Variables that are correlated with a biological variable of interest are potential confounders that may influence downstream analyses. Additionally, bar plots and histograms show the distributions of categorical and continuous variables, respectively.   
                                      
                                       
                                  
                                
                              
                            
                            
                              
                                
                                  
                                       
                                      
                                           Variance vs. Mean normalized signal plot across all targets/probes   More Plot Information      Mean and Variance statistics across all genes                Variance vs. Mean normalized signal plot across all targets/probes  Each gene's variance in the log-scaled, normalized data is plotted against its mean value across all samples. Highly variable genes are indicated by gene name. Housekeeping genes are color coded according to their use in (or omission from) normalization.   
                                           p-value distribution plots   More Plot Information                p-value distribution plots  For each covariate included in the analysis, a histogram of p-values testing each gene's univariate association with the chosen covariate is displayed. Covariates with largely flat histograms have minimal association with gene expression; covariates with histograms with significantly more mass on the left are either associated with the expression of many genes or are confounded with a covariate that is associated with the expression. Low p-values indicate strong evidence for an association.   
                                      
                                       
                                  
                                
                              
                            
                           
                            
                            
                            
                          
                           
                      
                    
                    
                      
                           
                          
                               Heatmaps    PCA     
                              
                                  Heatmaps  
                              
                               
                              
                                  New.Annotation  
                              
                              
                            
                              
                                
                                  
                                       
                                      
                                           Heatmap of Activation of BAD and translocation to mitochondria  Data   More Plot Information                      Heatmap of Activation of BAD and translocation to mitochondria  Data  Heatmap of the normalized data, scaled to give all genes equal variance, generated via unsupervised clustering. Orange indicates high expression; blue indicates low expression. This plot is meant to provide a high level exploratory view of the data.   
                                      
                                       
                                  
                                
                              
                            
                            
                              
                                
                                  
                                       
                                      
                                           Principal Components of Activation of BAD and translocation to mitochondria  Data   More Plot Information                      Principal Components of Activation of BAD and translocation to mitochondria  Data  Principal component analysis maps high-dimensional datasets onto a smaller number of highly informative dimensions. Here, the first four principal components of the gene expression data are plotted against each other and colored by the values of the selected covariate. This plot may be used to identify clusters in the data and to identify variables associated with prominent signal in the data. Variables that are associated with these leading principal components should be considered in downstream analyses.   
                                           Outliers   More Plot Information     
                                           
                                             
                                           
                                           
                                             
                                               
                                               Outlier on PC 1 
                                               Outlier on PC 2 
                                               Outlier on PC 3 
                                               Outlier on PC 4 
                                             
                                           
                                           
                                             
                                               20161013_sebelius2-101316_01_01.RCC 
                                               FALSE 
                                               FALSE 
                                               FALSE 
                                               TRUE 
                                             
                                           
                                           Outliers  Table identifying outliers in the first four PCs of the data.   
                                      
                                       
                                  
                                
                              
                            
                           
                            
                          
                           
                      
                    
                    
                      
                           
                          
                               Heatmaps    PCA     
                              
                                  Heatmaps  
                              
                               
                              
                                  New.Annotation  
                              
                              
                            
                              
                                
                                  
                                       
                                      
                                           Heatmap of Activation of Rac Data   More Plot Information                      Heatmap of Activation of Rac Data  Heatmap of the normalized data, scaled to give all genes equal variance, generated via unsupervised clustering. Orange indicates high expression; blue indicates low expression. This plot is meant to provide a high level exploratory view of the data.   
                                      
                                       
                                  
                                
                              
                            
                            
                              
                                
                                  
                                       
                                      
                                           Principal Components of Activation of Rac Data   More Plot Information                      Principal Components of Activation of Rac Data  Principal component analysis maps high-dimensional datasets onto a smaller number of highly informative dimensions. Here, the first four principal components of the gene expression data are plotted against each other and colored by the values of the selected covariate. This plot may be used to identify clusters in the data and to identify variables associated with prominent signal in the data. Variables that are associated with these leading principal components should be considered in downstream analyses.   
                                           Outliers   More Plot Information     
                                           
                                             
                                           
                                           
                                             
                                               
                                               Outlier on PC 1 
                                               Outlier on PC 2 
                                               Outlier on PC 3 
                                               Outlier on PC 4 
                                             
                                           
                                           
                                             
                                               20161013_sebelius2-101316_01_01.RCC 
                                               FALSE 
                                               FALSE 
                                               FALSE 
                                               TRUE 
                                             
                                           
                                           Outliers  Table identifying outliers in the first four PCs of the data.   
                                      
                                       
                                  
                                
                              
                            
                           
                            
                          
                           
                      
                    
                    
                      
                           
                          
                               Heatmaps    PCA     
                              
                                  Heatmaps  
                              
                               
                              
                                  New.Annotation  
                              
                              
                            
                              
                                
                                  
                                       
                                      
                                           Heatmap of Apoptotic cleavage of cellular proteins Data   More Plot Information                      Heatmap of Apoptotic cleavage of cellular proteins Data  Heatmap of the normalized data, scaled to give all genes equal variance, generated via unsupervised clustering. Orange indicates high expression; blue indicates low expression. This plot is meant to provide a high level exploratory view of the data.   
                                      
                                       
                                  
                                
                              
                            
                            
                              
                                
                                  
                                       
                                      
                                           Principal Components of Apoptotic cleavage of cellular proteins Data   More Plot Information                      Principal Components of Apoptotic cleavage of cellular proteins Data  Principal component analysis maps high-dimensional datasets onto a smaller number of highly informative dimensions. Here, the first four principal components of the gene expression data are plotted against each other and colored by the values of the selected covariate. This plot may be used to identify clusters in the data and to identify variables associated with prominent signal in the data. Variables that are associated with these leading principal components should be considered in downstream analyses.   
                                           Outliers   More Plot Information     
                                           
                                             
                                           
                                           
                                             
                                               
                                               Outlier on PC 1 
                                               Outlier on PC 2 
                                               Outlier on PC 3 
                                               Outlier on PC 4 
                                             
                                           
                                           
                                             
                                               20161013_sebelius2-101316_01_01.RCC 
                                               FALSE 
                                               FALSE 
                                               FALSE 
                                               TRUE 
                                             
                                           
                                           Outliers  Table identifying outliers in the first four PCs of the data.   
                                      
                                       
                                  
                                
                              
                            
                           
                            
                          
                           
                      
                    
                    
                      
                           
                          
                               Heatmaps    PCA     
                              
                                  Heatmaps  
                              
                               
                              
                                  New.Annotation  
                              
                              
                            
                              
                                
                                  
                                       
                                      
                                           Heatmap of ARMS-mediated activation Data   More Plot Information                      Heatmap of ARMS-mediated activation Data  Heatmap of the normalized data, scaled to give all genes equal variance, generated via unsupervised clustering. Orange indicates high expression; blue indicates low expression. This plot is meant to provide a high level exploratory view of the data.   
                                      
                                       
                                  
                                
                              
                            
                            
                              
                                
                                  
                                       
                                      
                                           Principal Components of ARMS-mediated activation Data   More Plot Information                      Principal Components of ARMS-mediated activation Data  Principal component analysis maps high-dimensional datasets onto a smaller number of highly informative dimensions. Here, the first four principal components of the gene expression data are plotted against each other and colored by the values of the selected covariate. This plot may be used to identify clusters in the data and to identify variables associated with prominent signal in the data. Variables that are associated with these leading principal components should be considered in downstream analyses.   
                                           Outliers   More Plot Information     
                                           
                                             
                                           
                                           
                                             
                                               
                                               Outlier on PC 1 
                                               Outlier on PC 2 
                                               Outlier on PC 3 
                                               Outlier on PC 4 
                                             
                                           
                                           
                                             
                                               20161013_sebelius2-101316_01_01.RCC 
                                               FALSE 
                                               FALSE 
                                               FALSE 
                                               TRUE 
                                             
                                           
                                           Outliers  Table identifying outliers in the first four PCs of the data.   
                                      
                                       
                                  
                                
                              
                            
                           
                            
                          
                           
                      
                    
                    
                      
                           
                          
                               Heatmaps    PCA     
                              
                                  Heatmaps  
                              
                               
                              
                                  New.Annotation  
                              
                              
                            
                              
                                
                                  
                                       
                                      
                                           Heatmap of AXIN missense mutants destabilize the destruction complex Data   More Plot Information                      Heatmap of AXIN missense mutants destabilize the destruction complex Data  Heatmap of the normalized data, scaled to give all genes equal variance, generated via unsupervised clustering. Orange indicates high expression; blue indicates low expression. This plot is meant to provide a high level exploratory view of the data.   
                                      
                                       
                                  
                                
                              
                            
                            
                              
                                
                                  
                                       
                                      
                                           Principal Components of AXIN missense mutants destabilize the destruction complex Data   More Plot Information                      Principal Components of AXIN missense mutants destabilize the destruction complex Data  Principal component analysis maps high-dimensional datasets onto a smaller number of highly informative dimensions. Here, the first four principal components of the gene expression data are plotted against each other and colored by the values of the selected covariate. This plot may be used to identify clusters in the data and to identify variables associated with prominent signal in the data. Variables that are associated with these leading principal components should be considered in downstream analyses.   
                                           Outliers   More Plot Information     
                                           
                                             
                                           
                                           
                                             
                                               
                                               Outlier on PC 1 
                                               Outlier on PC 2 
                                               Outlier on PC 3 
                                               Outlier on PC 4 
                                             
                                           
                                           
                                             
                                               20161013_sebelius2-101316_01_01.RCC 
                                               FALSE 
                                               FALSE 
                                               FALSE 
                                               TRUE 
                                             
                                           
                                           Outliers  Table identifying outliers in the first four PCs of the data.   
                                      
                                       
                                  
                                
                              
                            
                           
                            
                          
                           
                      
                    
                    
                      
                           
                          
                               Heatmaps    PCA     
                              
                                  Heatmaps  
                              
                               
                              
                                  New.Annotation  
                              
                              
                            
                              
                                
                                  
                                       
                                      
                                           Heatmap of Beta-catenin phosphorylation cascade Data   More Plot Information                      Heatmap of Beta-catenin phosphorylation cascade Data  Heatmap of the normalized data, scaled to give all genes equal variance, generated via unsupervised clustering. Orange indicates high expression; blue indicates low expression. This plot is meant to provide a high level exploratory view of the data.   
                                      
                                       
                                  
                                
                              
                            
                            
                              
                                
                                  
                                       
                                      
                                           Principal Components of Beta-catenin phosphorylation cascade Data   More Plot Information                      Principal Components of Beta-catenin phosphorylation cascade Data  Principal component analysis maps high-dimensional datasets onto a smaller number of highly informative dimensions. Here, the first four principal components of the gene expression data are plotted against each other and colored by the values of the selected covariate. This plot may be used to identify clusters in the data and to identify variables associated with prominent signal in the data. Variables that are associated with these leading principal components should be considered in downstream analyses.   
                                           Outliers   More Plot Information     
                                           
                                             
                                           
                                           
                                             
                                               
                                               Outlier on PC 1 
                                               Outlier on PC 2 
                                               Outlier on PC 3 
                                               Outlier on PC 4 
                                             
                                           
                                           
                                             
                                               20161013_sebelius2-101316_01_01.RCC 
                                               FALSE 
                                               FALSE 
                                               FALSE 
                                               TRUE 
                                             
                                           
                                           Outliers  Table identifying outliers in the first four PCs of the data.   
                                      
                                       
                                  
                                
                              
                            
                           
                            
                          
                           
                      
                    
                    
                      
                           
                          
                               Heatmaps    PCA     
                              
                                  Heatmaps  
                              
                               
                              
                                  New.Annotation  
                              
                              
                            
                              
                                
                                  
                                       
                                      
                                           Heatmap of CD209 (DC-SIGN) signaling Data   More Plot Information                      Heatmap of CD209 (DC-SIGN) signaling Data  Heatmap of the normalized data, scaled to give all genes equal variance, generated via unsupervised clustering. Orange indicates high expression; blue indicates low expression. This plot is meant to provide a high level exploratory view of the data.   
                                      
                                       
                                  
                                
                              
                            
                            
                              
                                
                                  
                                       
                                      
                                           Principal Components of CD209 (DC-SIGN) signaling Data   More Plot Information                      Principal Components of CD209 (DC-SIGN) signaling Data  Principal component analysis maps high-dimensional datasets onto a smaller number of highly informative dimensions. Here, the first four principal components of the gene expression data are plotted against each other and colored by the values of the selected covariate. This plot may be used to identify clusters in the data and to identify variables associated with prominent signal in the data. Variables that are associated with these leading principal components should be considered in downstream analyses.   
                                           Outliers   More Plot Information     
                                           
                                             
                                           
                                           
                                             
                                               
                                               Outlier on PC 1 
                                               Outlier on PC 2 
                                               Outlier on PC 3 
                                               Outlier on PC 4 
                                             
                                           
                                           
                                             
                                               20161013_sebelius2-101316_01_01.RCC 
                                               FALSE 
                                               FALSE 
                                               FALSE 
                                               TRUE 
                                             
                                           
                                           Outliers  Table identifying outliers in the first four PCs of the data.   
                                      
                                       
                                  
                                
                              
                            
                           
                            
                          
                           
                      
                    
                    
                      
                           
                          
                               Heatmaps    PCA     
                              
                                  Heatmaps  
                              
                               
                              
                                  New.Annotation  
                              
                              
                            
                              
                                
                                  
                                       
                                      
                                           Heatmap of CD28 co-stimulation Data   More Plot Information                      Heatmap of CD28 co-stimulation Data  Heatmap of the normalized data, scaled to give all genes equal variance, generated via unsupervised clustering. Orange indicates high expression; blue indicates low expression. This plot is meant to provide a high level exploratory view of the data.   
                                      
                                       
                                  
                                
                              
                            
                            
                              
                                
                                  
                                       
                                      
                                           Principal Components of CD28 co-stimulation Data   More Plot Information                      Principal Components of CD28 co-stimulation Data  Principal component analysis maps high-dimensional datasets onto a smaller number of highly informative dimensions. Here, the first four principal components of the gene expression data are plotted against each other and colored by the values of the selected covariate. This plot may be used to identify clusters in the data and to identify variables associated with prominent signal in the data. Variables that are associated with these leading principal components should be considered in downstream analyses.   
                                           Outliers   More Plot Information     
                                           
                                             
                                           
                                           
                                             
                                               
                                               Outlier on PC 1 
                                               Outlier on PC 2 
                                               Outlier on PC 3 
                                               Outlier on PC 4 
                                             
                                           
                                           
                                             
                                               20161013_sebelius2-101316_01_01.RCC 
                                               FALSE 
                                               FALSE 
                                               FALSE 
                                               TRUE 
                                             
                                           
                                           Outliers  Table identifying outliers in the first four PCs of the data.   
                                      
                                       
                                  
                                
                              
                            
                           
                            
                          
                           
                      
                    
                    
                      
                           
                          
                               Heatmaps    PCA     
                              
                                  Heatmaps  
                              
                               
                              
                                  New.Annotation  
                              
                              
                            
                              
                                
                                  
                                       
                                      
                                           Heatmap of CD28 dependent PI3KAkt signaling Data   More Plot Information                      Heatmap of CD28 dependent PI3KAkt signaling Data  Heatmap of the normalized data, scaled to give all genes equal variance, generated via unsupervised clustering. Orange indicates high expression; blue indicates low expression. This plot is meant to provide a high level exploratory view of the data.   
                                      
                                       
                                  
                                
                              
                            
                            
                              
                                
                                  
                                       
                                      
                                           Principal Components of CD28 dependent PI3KAkt signaling Data   More Plot Information                      Principal Components of CD28 dependent PI3KAkt signaling Data  Principal component analysis maps high-dimensional datasets onto a smaller number of highly informative dimensions. Here, the first four principal components of the gene expression data are plotted against each other and colored by the values of the selected covariate. This plot may be used to identify clusters in the data and to identify variables associated with prominent signal in the data. Variables that are associated with these leading principal components should be considered in downstream analyses.   
                                           Outliers   More Plot Information     
                                           
                                             
                                           
                                           
                                             
                                               
                                               Outlier on PC 1 
                                               Outlier on PC 2 
                                               Outlier on PC 3 
                                               Outlier on PC 4 
                                             
                                           
                                           
                                             
                                               20161013_sebelius2-101316_01_01.RCC 
                                               FALSE 
                                               FALSE 
                                               FALSE 
                                               TRUE 
                                             
                                           
                                           Outliers  Table identifying outliers in the first four PCs of the data.   
                                      
                                       
                                  
                                
                              
                            
                           
                            
                          
                           
                      
                    
                    
                      
                           
                          
                               Heatmaps    PCA     
                              
                                  Heatmaps  
                              
                               
                              
                                  New.Annotation  
                              
                              
                            
                              
                                
                                  
                                       
                                      
                                           Heatmap of CD28 dependent Vav1 pathway Data   More Plot Information                      Heatmap of CD28 dependent Vav1 pathway Data  Heatmap of the normalized data, scaled to give all genes equal variance, generated via unsupervised clustering. Orange indicates high expression; blue indicates low expression. This plot is meant to provide a high level exploratory view of the data.   
                                      
                                       
                                  
                                
                              
                            
                            
                              
                                
                                  
                                       
                                      
                                           Principal Components of CD28 dependent Vav1 pathway Data   More Plot Information                      Principal Components of CD28 dependent Vav1 pathway Data  Principal component analysis maps high-dimensional datasets onto a smaller number of highly informative dimensions. Here, the first four principal components of the gene expression data are plotted against each other and colored by the values of the selected covariate. This plot may be used to identify clusters in the data and to identify variables associated with prominent signal in the data. Variables that are associated with these leading principal components should be considered in downstream analyses.   
                                           Outliers   More Plot Information     
                                           
                                             
                                           
                                           
                                             
                                               
                                               Outlier on PC 1 
                                               Outlier on PC 2 
                                               Outlier on PC 3 
                                               Outlier on PC 4 
                                             
                                           
                                           
                                             
                                               20161013_sebelius2-101316_01_01.RCC 
                                               FALSE 
                                               FALSE 
                                               FALSE 
                                               TRUE 
                                             
                                           
                                           Outliers  Table identifying outliers in the first four PCs of the data.   
                                      
                                       
                                  
                                
                              
                            
                           
                            
                          
                           
                      
                    
                    
                      
                           
                          
                               Heatmaps    PCA     
                              
                                  Heatmaps  
                              
                               
                              
                                  New.Annotation  
                              
                              
                            
                              
                                
                                  
                                       
                                      
                                           Heatmap of Cell-extracellular matrix interactions Data   More Plot Information                      Heatmap of Cell-extracellular matrix interactions Data  Heatmap of the normalized data, scaled to give all genes equal variance, generated via unsupervised clustering. Orange indicates high expression; blue indicates low expression. This plot is meant to provide a high level exploratory view of the data.   
                                      
                                       
                                  
                                
                              
                            
                            
                              
                                
                                  
                                       
                                      
                                           Principal Components of Cell-extracellular matrix interactions Data   More Plot Information                      Principal Components of Cell-extracellular matrix interactions Data  Principal component analysis maps high-dimensional datasets onto a smaller number of highly informative dimensions. Here, the first four principal components of the gene expression data are plotted against each other and colored by the values of the selected covariate. This plot may be used to identify clusters in the data and to identify variables associated with prominent signal in the data. Variables that are associated with these leading principal components should be considered in downstream analyses.   
                                           Outliers   More Plot Information     
                                           
                                             
                                           
                                           
                                             
                                               
                                               Outlier on PC 1 
                                               Outlier on PC 2 
                                               Outlier on PC 3 
                                               Outlier on PC 4 
                                             
                                           
                                           
                                             
                                               20161013_sebelius2-101316_01_01.RCC 
                                               FALSE 
                                               FALSE 
                                               FALSE 
                                               TRUE 
                                             
                                           
                                           Outliers  Table identifying outliers in the first four PCs of the data.   
                                      
                                       
                                  
                                
                              
                            
                           
                            
                          
                           
                      
                    
                    
                      
                           
                          
                               Heatmaps    PCA     
                              
                                  Heatmaps  
                              
                               
                              
                                  New.Annotation  
                              
                              
                            
                              
                                
                                  
                                       
                                      
                                           Heatmap of Cell surface interactions at the vascular wall Data   More Plot Information                      Heatmap of Cell surface interactions at the vascular wall Data  Heatmap of the normalized data, scaled to give all genes equal variance, generated via unsupervised clustering. Orange indicates high expression; blue indicates low expression. This plot is meant to provide a high level exploratory view of the data.   
                                      
                                       
                                  
                                
                              
                            
                            
                              
                                
                                  
                                       
                                      
                                           Principal Components of Cell surface interactions at the vascular wall Data   More Plot Information                      Principal Components of Cell surface interactions at the vascular wall Data  Principal component analysis maps high-dimensional datasets onto a smaller number of highly informative dimensions. Here, the first four principal components of the gene expression data are plotted against each other and colored by the values of the selected covariate. This plot may be used to identify clusters in the data and to identify variables associated with prominent signal in the data. Variables that are associated with these leading principal components should be considered in downstream analyses.   
                                           Outliers   More Plot Information     
                                           
                                             
                                           
                                           
                                             
                                               
                                               Outlier on PC 1 
                                               Outlier on PC 2 
                                               Outlier on PC 3 
                                               Outlier on PC 4 
                                             
                                           
                                           
                                             
                                               20161013_sebelius2-101316_01_01.RCC 
                                               FALSE 
                                               FALSE 
                                               FALSE 
                                               TRUE 
                                             
                                           
                                           Outliers  Table identifying outliers in the first four PCs of the data.   
                                      
                                       
                                  
                                
                              
                            
                           
                            
                          
                           
                      
                    
                    
                      
                           
                          
                               Heatmaps    PCA     
                              
                                  Heatmaps  
                              
                               
                              
                                  New.Annotation  
                              
                              
                            
                              
                                
                                  
                                       
                                      
                                           Heatmap of Circadian Clock Data   More Plot Information                      Heatmap of Circadian Clock Data  Heatmap of the normalized data, scaled to give all genes equal variance, generated via unsupervised clustering. Orange indicates high expression; blue indicates low expression. This plot is meant to provide a high level exploratory view of the data.   
                                      
                                       
                                  
                                
                              
                            
                            
                              
                                
                                  
                                       
                                      
                                           Principal Components of Circadian Clock Data   More Plot Information                      Principal Components of Circadian Clock Data  Principal component analysis maps high-dimensional datasets onto a smaller number of highly informative dimensions. Here, the first four principal components of the gene expression data are plotted against each other and colored by the values of the selected covariate. This plot may be used to identify clusters in the data and to identify variables associated with prominent signal in the data. Variables that are associated with these leading principal components should be considered in downstream analyses.   
                                           Outliers   More Plot Information     
                                           
                                             
                                           
                                           
                                             
                                               
                                               Outlier on PC 1 
                                               Outlier on PC 2 
                                               Outlier on PC 3 
                                               Outlier on PC 4 
                                             
                                           
                                           
                                             
                                               20161013_sebelius2-101316_01_01.RCC 
                                               FALSE 
                                               FALSE 
                                               FALSE 
                                               TRUE 
                                             
                                           
                                           Outliers  Table identifying outliers in the first four PCs of the data.   
                                      
                                       
                                  
                                
                              
                            
                           
                            
                          
                           
                      
                    
                    
                      
                           
                          
                               Heatmaps    PCA     
                              
                                  Heatmaps  
                              
                               
                              
                                  New.Annotation  
                              
                              
                            
                              
                                
                                  
                                       
                                      
                                           Heatmap of Constitutive Signaling by Aberrant PI3K in Cancer Data   More Plot Information                      Heatmap of Constitutive Signaling by Aberrant PI3K in Cancer Data  Heatmap of the normalized data, scaled to give all genes equal variance, generated via unsupervised clustering. Orange indicates high expression; blue indicates low expression. This plot is meant to provide a high level exploratory view of the data.   
                                      
                                       
                                  
                                
                              
                            
                            
                              
                                
                                  
                                       
                                      
                                           Principal Components of Constitutive Signaling by Aberrant PI3K in Cancer Data   More Plot Information                      Principal Components of Constitutive Signaling by Aberrant PI3K in Cancer Data  Principal component analysis maps high-dimensional datasets onto a smaller number of highly informative dimensions. Here, the first four principal components of the gene expression data are plotted against each other and colored by the values of the selected covariate. This plot may be used to identify clusters in the data and to identify variables associated with prominent signal in the data. Variables that are associated with these leading principal components should be considered in downstream analyses.   
                                           Outliers   More Plot Information     
                                           
                                             
                                           
                                           
                                             
                                               
                                               Outlier on PC 1 
                                               Outlier on PC 2 
                                               Outlier on PC 3 
                                               Outlier on PC 4 
                                             
                                           
                                           
                                             
                                               20161013_sebelius2-101316_01_01.RCC 
                                               FALSE 
                                               FALSE 
                                               FALSE 
                                               TRUE 
                                             
                                           
                                           Outliers  Table identifying outliers in the first four PCs of the data.   
                                      
                                       
                                  
                                
                              
                            
                           
                            
                          
                           
                      
                    
                    
                      
                           
                          
                               Heatmaps    PCA     
                              
                                  Heatmaps  
                              
                               
                              
                                  New.Annotation  
                              
                              
                            
                              
                                
                                  
                                       
                                      
                                           Heatmap of Constitutive Signaling by EGFRvIII Data   More Plot Information                      Heatmap of Constitutive Signaling by EGFRvIII Data  Heatmap of the normalized data, scaled to give all genes equal variance, generated via unsupervised clustering. Orange indicates high expression; blue indicates low expression. This plot is meant to provide a high level exploratory view of the data.   
                                      
                                       
                                  
                                
                              
                            
                            
                              
                                
                                  
                                       
                                      
                                           Principal Components of Constitutive Signaling by EGFRvIII Data   More Plot Information                      Principal Components of Constitutive Signaling by EGFRvIII Data  Principal component analysis maps high-dimensional datasets onto a smaller number of highly informative dimensions. Here, the first four principal components of the gene expression data are plotted against each other and colored by the values of the selected covariate. This plot may be used to identify clusters in the data and to identify variables associated with prominent signal in the data. Variables that are associated with these leading principal components should be considered in downstream analyses.   
                                           Outliers   More Plot Information     
                                           
                                             
                                           
                                           
                                             
                                               
                                               Outlier on PC 1 
                                               Outlier on PC 2 
                                               Outlier on PC 3 
                                               Outlier on PC 4 
                                             
                                           
                                           
                                             
                                               20161013_sebelius2-101316_01_01.RCC 
                                               FALSE 
                                               FALSE 
                                               FALSE 
                                               TRUE 
                                             
                                           
                                           Outliers  Table identifying outliers in the first four PCs of the data.   
                                      
                                       
                                  
                                
                              
                            
                           
                            
                          
                           
                      
                    
                    
                      
                           
                          
                               Heatmaps    PCA     
                              
                                  Heatmaps  
                              
                               
                              
                                  New.Annotation  
                              
                              
                            
                              
                                
                                  
                                       
                                      
                                           Heatmap of CREB phosphorylation through the activation of Ras Data   More Plot Information                      Heatmap of CREB phosphorylation through the activation of Ras Data  Heatmap of the normalized data, scaled to give all genes equal variance, generated via unsupervised clustering. Orange indicates high expression; blue indicates low expression. This plot is meant to provide a high level exploratory view of the data.   
                                      
                                       
                                  
                                
                              
                            
                            
                              
                                
                                  
                                       
                                      
                                           Principal Components of CREB phosphorylation through the activation of Ras Data   More Plot Information                      Principal Components of CREB phosphorylation through the activation of Ras Data  Principal component analysis maps high-dimensional datasets onto a smaller number of highly informative dimensions. Here, the first four principal components of the gene expression data are plotted against each other and colored by the values of the selected covariate. This plot may be used to identify clusters in the data and to identify variables associated with prominent signal in the data. Variables that are associated with these leading principal components should be considered in downstream analyses.   
                                           Outliers   More Plot Information     
                                           
                                             
                                           
                                           
                                             
                                               
                                               Outlier on PC 1 
                                               Outlier on PC 2 
                                               Outlier on PC 3 
                                               Outlier on PC 4 
                                             
                                           
                                           
                                             
                                               20161013_sebelius2-101316_01_01.RCC 
                                               FALSE 
                                               FALSE 
                                               FALSE 
                                               TRUE 
                                             
                                           
                                           Outliers  Table identifying outliers in the first four PCs of the data.   
                                      
                                       
                                  
                                
                              
                            
                           
                            
                          
                           
                      
                    
                    
                      
                           
                          
                               Heatmaps    PCA     
                              
                                  Heatmaps  
                              
                               
                              
                                  New.Annotation  
                              
                              
                            
                              
                                
                                  
                                       
                                      
                                           Heatmap of CTLA4 inhibitory signaling Data   More Plot Information                      Heatmap of CTLA4 inhibitory signaling Data  Heatmap of the normalized data, scaled to give all genes equal variance, generated via unsupervised clustering. Orange indicates high expression; blue indicates low expression. This plot is meant to provide a high level exploratory view of the data.   
                                      
                                       
                                  
                                
                              
                            
                            
                              
                                
                                  
                                       
                                      
                                           Principal Components of CTLA4 inhibitory signaling Data   More Plot Information                      Principal Components of CTLA4 inhibitory signaling Data  Principal component analysis maps high-dimensional datasets onto a smaller number of highly informative dimensions. Here, the first four principal components of the gene expression data are plotted against each other and colored by the values of the selected covariate. This plot may be used to identify clusters in the data and to identify variables associated with prominent signal in the data. Variables that are associated with these leading principal components should be considered in downstream analyses.   
                                           Outliers   More Plot Information     
                                           
                                             
                                           
                                           
                                             
                                               
                                               Outlier on PC 1 
                                               Outlier on PC 2 
                                               Outlier on PC 3 
                                               Outlier on PC 4 
                                             
                                           
                                           
                                             
                                               20161013_sebelius2-101316_01_01.RCC 
                                               FALSE 
                                               FALSE 
                                               FALSE 
                                               TRUE 
                                             
                                           
                                           Outliers  Table identifying outliers in the first four PCs of the data.   
                                      
                                       
                                  
                                
                              
                            
                           
                            
                          
                           
                      
                    
                    
                      
                           
                          
                               Heatmaps    PCA     
                              
                                  Heatmaps  
                              
                               
                              
                                  New.Annotation  
                              
                              
                            
                              
                                
                                  
                                       
                                      
                                           Heatmap of Cyclin AB1 associated events during G2M transition Data   More Plot Information                      Heatmap of Cyclin AB1 associated events during G2M transition Data  Heatmap of the normalized data, scaled to give all genes equal variance, generated via unsupervised clustering. Orange indicates high expression; blue indicates low expression. This plot is meant to provide a high level exploratory view of the data.   
                                      
                                       
                                  
                                
                              
                            
                            
                              
                                
                                  
                                       
                                      
                                           Principal Components of Cyclin AB1 associated events during G2M transition Data   More Plot Information                      Principal Components of Cyclin AB1 associated events during G2M transition Data  Principal component analysis maps high-dimensional datasets onto a smaller number of highly informative dimensions. Here, the first four principal components of the gene expression data are plotted against each other and colored by the values of the selected covariate. This plot may be used to identify clusters in the data and to identify variables associated with prominent signal in the data. Variables that are associated with these leading principal components should be considered in downstream analyses.   
                                           Outliers   More Plot Information     
                                           
                                             
                                           
                                           
                                             
                                               
                                               Outlier on PC 1 
                                               Outlier on PC 2 
                                               Outlier on PC 3 
                                               Outlier on PC 4 
                                             
                                           
                                           
                                             
                                               20161013_sebelius2-101316_01_01.RCC 
                                               FALSE 
                                               FALSE 
                                               FALSE 
                                               TRUE 
                                             
                                           
                                           Outliers  Table identifying outliers in the first four PCs of the data.   
                                      
                                       
                                  
                                
                              
                            
                           
                            
                          
                           
                      
                    
                    
                      
                           
                          
                               Heatmaps    PCA     
                              
                                  Heatmaps  
                              
                               
                              
                                  New.Annotation  
                              
                              
                            
                              
                                
                                  
                                       
                                      
                                           Heatmap of Cyclin D associated events in G1 Data   More Plot Information                      Heatmap of Cyclin D associated events in G1 Data  Heatmap of the normalized data, scaled to give all genes equal variance, generated via unsupervised clustering. Orange indicates high expression; blue indicates low expression. This plot is meant to provide a high level exploratory view of the data.   
                                      
                                       
                                  
                                
                              
                            
                            
                              
                                
                                  
                                       
                                      
                                           Principal Components of Cyclin D associated events in G1 Data   More Plot Information                      Principal Components of Cyclin D associated events in G1 Data  Principal component analysis maps high-dimensional datasets onto a smaller number of highly informative dimensions. Here, the first four principal components of the gene expression data are plotted against each other and colored by the values of the selected covariate. This plot may be used to identify clusters in the data and to identify variables associated with prominent signal in the data. Variables that are associated with these leading principal components should be considered in downstream analyses.   
                                           Outliers   More Plot Information     
                                           
                                             
                                           
                                           
                                             
                                               
                                               Outlier on PC 1 
                                               Outlier on PC 2 
                                               Outlier on PC 3 
                                               Outlier on PC 4 
                                             
                                           
                                           
                                             
                                               20161013_sebelius2-101316_01_01.RCC 
                                               FALSE 
                                               FALSE 
                                               FALSE 
                                               TRUE 
                                             
                                           
                                           Outliers  Table identifying outliers in the first four PCs of the data.   
                                      
                                       
                                  
                                
                              
                            
                           
                            
                          
                           
                      
                    
                    
                      
                           
                          
                               Heatmaps    PCA     
                              
                                  Heatmaps  
                              
                               
                              
                                  New.Annotation  
                              
                              
                            
                              
                                
                                  
                                       
                                      
                                           Heatmap of DAP12 signaling Data   More Plot Information                      Heatmap of DAP12 signaling Data  Heatmap of the normalized data, scaled to give all genes equal variance, generated via unsupervised clustering. Orange indicates high expression; blue indicates low expression. This plot is meant to provide a high level exploratory view of the data.   
                                      
                                       
                                  
                                
                              
                            
                            
                              
                                
                                  
                                       
                                      
                                           Principal Components of DAP12 signaling Data   More Plot Information                      Principal Components of DAP12 signaling Data  Principal component analysis maps high-dimensional datasets onto a smaller number of highly informative dimensions. Here, the first four principal components of the gene expression data are plotted against each other and colored by the values of the selected covariate. This plot may be used to identify clusters in the data and to identify variables associated with prominent signal in the data. Variables that are associated with these leading principal components should be considered in downstream analyses.   
                                           Outliers   More Plot Information     
                                           
                                             
                                           
                                           
                                             
                                               
                                               Outlier on PC 1 
                                               Outlier on PC 2 
                                               Outlier on PC 3 
                                               Outlier on PC 4 
                                             
                                           
                                           
                                             
                                               20161013_sebelius2-101316_01_01.RCC 
                                               FALSE 
                                               FALSE 
                                               FALSE 
                                               TRUE 
                                             
                                           
                                           Outliers  Table identifying outliers in the first four PCs of the data.   
                                      
                                       
                                  
                                
                              
                            
                           
                            
                          
                           
                      
                    
                    
                      
                           
                          
                               Heatmaps    PCA     
                              
                                  Heatmaps  
                              
                               
                              
                                  New.Annotation  
                              
                              
                            
                              
                                
                                  
                                       
                                      
                                           Heatmap of DARPP-32 events Data   More Plot Information                      Heatmap of DARPP-32 events Data  Heatmap of the normalized data, scaled to give all genes equal variance, generated via unsupervised clustering. Orange indicates high expression; blue indicates low expression. This plot is meant to provide a high level exploratory view of the data.   
                                      
                                       
                                  
                                
                              
                            
                            
                              
                                
                                  
                                       
                                      
                                           Principal Components of DARPP-32 events Data   More Plot Information                      Principal Components of DARPP-32 events Data  Principal component analysis maps high-dimensional datasets onto a smaller number of highly informative dimensions. Here, the first four principal components of the gene expression data are plotted against each other and colored by the values of the selected covariate. This plot may be used to identify clusters in the data and to identify variables associated with prominent signal in the data. Variables that are associated with these leading principal components should be considered in downstream analyses.   
                                           Outliers   More Plot Information     
                                           
                                             
                                           
                                           
                                             
                                               
                                               Outlier on PC 1 
                                               Outlier on PC 2 
                                               Outlier on PC 3 
                                               Outlier on PC 4 
                                             
                                           
                                           
                                             
                                               20161013_sebelius2-101316_01_01.RCC 
                                               FALSE 
                                               FALSE 
                                               FALSE 
                                               TRUE 
                                             
                                           
                                           Outliers  Table identifying outliers in the first four PCs of the data.   
                                      
                                       
                                  
                                
                              
                            
                           
                            
                          
                           
                      
                    
                    
                      
                           
                          
                               Heatmaps    PCA     
                              
                                  Heatmaps  
                              
                               
                              
                                  New.Annotation  
                              
                              
                            
                              
                                
                                  
                                       
                                      
                                           Heatmap of DCC mediated attractive signaling Data   More Plot Information                      Heatmap of DCC mediated attractive signaling Data  Heatmap of the normalized data, scaled to give all genes equal variance, generated via unsupervised clustering. Orange indicates high expression; blue indicates low expression. This plot is meant to provide a high level exploratory view of the data.   
                                      
                                       
                                  
                                
                              
                            
                            
                              
                                
                                  
                                       
                                      
                                           Principal Components of DCC mediated attractive signaling Data   More Plot Information                      Principal Components of DCC mediated attractive signaling Data  Principal component analysis maps high-dimensional datasets onto a smaller number of highly informative dimensions. Here, the first four principal components of the gene expression data are plotted against each other and colored by the values of the selected covariate. This plot may be used to identify clusters in the data and to identify variables associated with prominent signal in the data. Variables that are associated with these leading principal components should be considered in downstream analyses.   
                                           Outliers   More Plot Information     
                                           
                                             
                                           
                                           
                                             
                                               
                                               Outlier on PC 1 
                                               Outlier on PC 2 
                                               Outlier on PC 3 
                                               Outlier on PC 4 
                                             
                                           
                                           
                                             
                                               20161013_sebelius2-101316_01_01.RCC 
                                               FALSE 
                                               FALSE 
                                               FALSE 
                                               TRUE 
                                             
                                           
                                           Outliers  Table identifying outliers in the first four PCs of the data.   
                                      
                                       
                                  
                                
                              
                            
                           
                            
                          
                           
                      
                    
                    
                      
                           
                          
                               Heatmaps    PCA     
                              
                                  Heatmaps  
                              
                               
                              
                                  New.Annotation  
                              
                              
                            
                              
                                
                                  
                                       
                                      
                                           Heatmap of Deactivation of the beta-catenin transactivating complex Data   More Plot Information                      Heatmap of Deactivation of the beta-catenin transactivating complex Data  Heatmap of the normalized data, scaled to give all genes equal variance, generated via unsupervised clustering. Orange indicates high expression; blue indicates low expression. This plot is meant to provide a high level exploratory view of the data.   
                                      
                                       
                                  
                                
                              
                            
                            
                              
                                
                                  
                                       
                                      
                                           Principal Components of Deactivation of the beta-catenin transactivating complex Data   More Plot Information                      Principal Components of Deactivation of the beta-catenin transactivating complex Data  Principal component analysis maps high-dimensional datasets onto a smaller number of highly informative dimensions. Here, the first four principal components of the gene expression data are plotted against each other and colored by the values of the selected covariate. This plot may be used to identify clusters in the data and to identify variables associated with prominent signal in the data. Variables that are associated with these leading principal components should be considered in downstream analyses.   
                                           Outliers   More Plot Information     
                                           
                                             
                                           
                                           
                                             
                                               
                                               Outlier on PC 1 
                                               Outlier on PC 2 
                                               Outlier on PC 3 
                                               Outlier on PC 4 
                                             
                                           
                                           
                                             
                                               20161013_sebelius2-101316_01_01.RCC 
                                               FALSE 
                                               FALSE 
                                               FALSE 
                                               TRUE 
                                             
                                           
                                           Outliers  Table identifying outliers in the first four PCs of the data.   
                                      
                                       
                                  
                                
                              
                            
                           
                            
                          
                           
                      
                    
                    
                      
                           
                          
                               Heatmaps    PCA     
                              
                                  Heatmaps  
                              
                               
                              
                                  New.Annotation  
                              
                              
                            
                              
                                
                                  
                                       
                                      
                                           Heatmap of Disassembly of the destruction complex and recruitment of AXIN to the membrane Data   More Plot Information                      Heatmap of Disassembly of the destruction complex and recruitment of AXIN to the membrane Data  Heatmap of the normalized data, scaled to give all genes equal variance, generated via unsupervised clustering. Orange indicates high expression; blue indicates low expression. This plot is meant to provide a high level exploratory view of the data.   
                                      
                                       
                                  
                                
                              
                            
                            
                              
                                
                                  
                                       
                                      
                                           Principal Components of Disassembly of the destruction complex and recruitment of AXIN to the membrane Data   More Plot Information                      Principal Components of Disassembly of the destruction complex and recruitment of AXIN to the membrane Data  Principal component analysis maps high-dimensional datasets onto a smaller number of highly informative dimensions. Here, the first four principal components of the gene expression data are plotted against each other and colored by the values of the selected covariate. This plot may be used to identify clusters in the data and to identify variables associated with prominent signal in the data. Variables that are associated with these leading principal components should be considered in downstream analyses.   
                                           Outliers   More Plot Information     
                                           
                                             
                                           
                                           
                                             
                                               
                                               Outlier on PC 1 
                                               Outlier on PC 2 
                                               Outlier on PC 3 
                                               Outlier on PC 4 
                                             
                                           
                                           
                                             
                                               20161013_sebelius2-101316_01_01.RCC 
                                               FALSE 
                                               FALSE 
                                               FALSE 
                                               TRUE 
                                             
                                           
                                           Outliers  Table identifying outliers in the first four PCs of the data.   
                                      
                                       
                                  
                                
                              
                            
                           
                            
                          
                           
                      
                    
                    
                      
                           
                          
                               Heatmaps    PCA     
                              
                                  Heatmaps  
                              
                               
                              
                                  New.Annotation  
                              
                              
                            
                              
                                
                                  
                                       
                                      
                                           Heatmap of Downstream signal transduction Data   More Plot Information                      Heatmap of Downstream signal transduction Data  Heatmap of the normalized data, scaled to give all genes equal variance, generated via unsupervised clustering. Orange indicates high expression; blue indicates low expression. This plot is meant to provide a high level exploratory view of the data.   
                                      
                                       
                                  
                                
                              
                            
                            
                              
                                
                                  
                                       
                                      
                                           Principal Components of Downstream signal transduction Data   More Plot Information                      Principal Components of Downstream signal transduction Data  Principal component analysis maps high-dimensional datasets onto a smaller number of highly informative dimensions. Here, the first four principal components of the gene expression data are plotted against each other and colored by the values of the selected covariate. This plot may be used to identify clusters in the data and to identify variables associated with prominent signal in the data. Variables that are associated with these leading principal components should be considered in downstream analyses.   
                                           Outliers   More Plot Information     
                                           
                                             
                                           
                                           
                                             
                                               
                                               Outlier on PC 1 
                                               Outlier on PC 2 
                                               Outlier on PC 3 
                                               Outlier on PC 4 
                                             
                                           
                                           
                                             
                                               20161013_sebelius2-101316_01_01.RCC 
                                               FALSE 
                                               FALSE 
                                               FALSE 
                                               TRUE 
                                             
                                           
                                           Outliers  Table identifying outliers in the first four PCs of the data.   
                                      
                                       
                                  
                                
                              
                            
                           
                            
                          
                           
                      
                    
                    
                      
                           
                          
                               Heatmaps    PCA     
                              
                                  Heatmaps  
                              
                               
                              
                                  New.Annotation  
                              
                              
                            
                              
                                
                                  
                                       
                                      
                                           Heatmap of Downstream TCR signaling Data   More Plot Information                      Heatmap of Downstream TCR signaling Data  Heatmap of the normalized data, scaled to give all genes equal variance, generated via unsupervised clustering. Orange indicates high expression; blue indicates low expression. This plot is meant to provide a high level exploratory view of the data.   
                                      
                                       
                                  
                                
                              
                            
                            
                              
                                
                                  
                                       
                                      
                                           Principal Components of Downstream TCR signaling Data   More Plot Information                      Principal Components of Downstream TCR signaling Data  Principal component analysis maps high-dimensional datasets onto a smaller number of highly informative dimensions. Here, the first four principal components of the gene expression data are plotted against each other and colored by the values of the selected covariate. This plot may be used to identify clusters in the data and to identify variables associated with prominent signal in the data. Variables that are associated with these leading principal components should be considered in downstream analyses.   
                                           Outliers   More Plot Information     
                                           
                                             
                                           
                                           
                                             
                                               
                                               Outlier on PC 1 
                                               Outlier on PC 2 
                                               Outlier on PC 3 
                                               Outlier on PC 4 
                                             
                                           
                                           
                                             
                                               20161013_sebelius2-101316_01_01.RCC 
                                               FALSE 
                                               FALSE 
                                               FALSE 
                                               TRUE 
                                             
                                           
                                           Outliers  Table identifying outliers in the first four PCs of the data.   
                                      
                                       
                                  
                                
                              
                            
                           
                            
                          
                           
                      
                    
                    
                      
                           
                          
                               Heatmaps    PCA     
                              
                                  Heatmaps  
                              
                               
                              
                                  New.Annotation  
                              
                              
                            
                              
                                
                                  
                                       
                                      
                                           Heatmap of ECM proteoglycans Data   More Plot Information                      Heatmap of ECM proteoglycans Data  Heatmap of the normalized data, scaled to give all genes equal variance, generated via unsupervised clustering. Orange indicates high expression; blue indicates low expression. This plot is meant to provide a high level exploratory view of the data.   
                                      
                                       
                                  
                                
                              
                            
                            
                              
                                
                                  
                                       
                                      
                                           Principal Components of ECM proteoglycans Data   More Plot Information                      Principal Components of ECM proteoglycans Data  Principal component analysis maps high-dimensional datasets onto a smaller number of highly informative dimensions. Here, the first four principal components of the gene expression data are plotted against each other and colored by the values of the selected covariate. This plot may be used to identify clusters in the data and to identify variables associated with prominent signal in the data. Variables that are associated with these leading principal components should be considered in downstream analyses.   
                                           Outliers   More Plot Information     
                                           
                                             
                                           
                                           
                                             
                                               
                                               Outlier on PC 1 
                                               Outlier on PC 2 
                                               Outlier on PC 3 
                                               Outlier on PC 4 
                                             
                                           
                                           
                                             
                                               20161013_sebelius2-101316_01_01.RCC 
                                               FALSE 
                                               FALSE 
                                               FALSE 
                                               TRUE 
                                             
                                           
                                           Outliers  Table identifying outliers in the first four PCs of the data.   
                                      
                                       
                                  
                                
                              
                            
                           
                            
                          
                           
                      
                    
                    
                      
                           
                          
                               Heatmaps    PCA     
                              
                                  Heatmaps  
                              
                               
                              
                                  New.Annotation  
                              
                              
                            
                              
                                
                                  
                                       
                                      
                                           Heatmap of Elastic fibre formation Data   More Plot Information                      Heatmap of Elastic fibre formation Data  Heatmap of the normalized data, scaled to give all genes equal variance, generated via unsupervised clustering. Orange indicates high expression; blue indicates low expression. This plot is meant to provide a high level exploratory view of the data.   
                                      
                                       
                                  
                                
                              
                            
                            
                              
                                
                                  
                                       
                                      
                                           Principal Components of Elastic fibre formation Data   More Plot Information                      Principal Components of Elastic fibre formation Data  Principal component analysis maps high-dimensional datasets onto a smaller number of highly informative dimensions. Here, the first four principal components of the gene expression data are plotted against each other and colored by the values of the selected covariate. This plot may be used to identify clusters in the data and to identify variables associated with prominent signal in the data. Variables that are associated with these leading principal components should be considered in downstream analyses.   
                                           Outliers   More Plot Information     
                                           
                                             
                                           
                                           
                                             
                                               
                                               Outlier on PC 1 
                                               Outlier on PC 2 
                                               Outlier on PC 3 
                                               Outlier on PC 4 
                                             
                                           
                                           
                                             
                                               20161013_sebelius2-101316_01_01.RCC 
                                               FALSE 
                                               FALSE 
                                               FALSE 
                                               TRUE 
                                             
                                           
                                           Outliers  Table identifying outliers in the first four PCs of the data.   
                                      
                                       
                                  
                                
                              
                            
                           
                            
                          
                           
                      
                    
                    
                      
                           
                          
                               Heatmaps    PCA     
                              
                                  Heatmaps  
                              
                               
                              
                                  New.Annotation  
                              
                              
                            
                              
                                
                                  
                                       
                                      
                                           Heatmap of EPH-ephrin mediated repulsion of cells Data   More Plot Information                      Heatmap of EPH-ephrin mediated repulsion of cells Data  Heatmap of the normalized data, scaled to give all genes equal variance, generated via unsupervised clustering. Orange indicates high expression; blue indicates low expression. This plot is meant to provide a high level exploratory view of the data.   
                                      
                                       
                                  
                                
                              
                            
                            
                              
                                
                                  
                                       
                                      
                                           Principal Components of EPH-ephrin mediated repulsion of cells Data   More Plot Information                      Principal Components of EPH-ephrin mediated repulsion of cells Data  Principal component analysis maps high-dimensional datasets onto a smaller number of highly informative dimensions. Here, the first four principal components of the gene expression data are plotted against each other and colored by the values of the selected covariate. This plot may be used to identify clusters in the data and to identify variables associated with prominent signal in the data. Variables that are associated with these leading principal components should be considered in downstream analyses.   
                                           Outliers   More Plot Information     
                                           
                                             
                                           
                                           
                                             
                                               
                                               Outlier on PC 1 
                                               Outlier on PC 2 
                                               Outlier on PC 3 
                                               Outlier on PC 4 
                                             
                                           
                                           
                                             
                                               20161013_sebelius2-101316_01_01.RCC 
                                               FALSE 
                                               FALSE 
                                               FALSE 
                                               TRUE 
                                             
                                           
                                           Outliers  Table identifying outliers in the first four PCs of the data.   
                                      
                                       
                                  
                                
                              
                            
                           
                            
                          
                           
                      
                    
                    
                      
                           
                          
                               Heatmaps    PCA     
                              
                                  Heatmaps  
                              
                               
                              
                                  New.Annotation  
                              
                              
                            
                              
                                
                                  
                                       
                                      
                                           Heatmap of EPHA-mediated growth cone collapse Data   More Plot Information                      Heatmap of EPHA-mediated growth cone collapse Data  Heatmap of the normalized data, scaled to give all genes equal variance, generated via unsupervised clustering. Orange indicates high expression; blue indicates low expression. This plot is meant to provide a high level exploratory view of the data.   
                                      
                                       
                                  
                                
                              
                            
                            
                              
                                
                                  
                                       
                                      
                                           Principal Components of EPHA-mediated growth cone collapse Data   More Plot Information                      Principal Components of EPHA-mediated growth cone collapse Data  Principal component analysis maps high-dimensional datasets onto a smaller number of highly informative dimensions. Here, the first four principal components of the gene expression data are plotted against each other and colored by the values of the selected covariate. This plot may be used to identify clusters in the data and to identify variables associated with prominent signal in the data. Variables that are associated with these leading principal components should be considered in downstream analyses.   
                                           Outliers   More Plot Information     
                                           
                                             
                                           
                                           
                                             
                                               
                                               Outlier on PC 1 
                                               Outlier on PC 2 
                                               Outlier on PC 3 
                                               Outlier on PC 4 
                                             
                                           
                                           
                                             
                                               20161013_sebelius2-101316_01_01.RCC 
                                               FALSE 
                                               FALSE 
                                               FALSE 
                                               TRUE 
                                             
                                           
                                           Outliers  Table identifying outliers in the first four PCs of the data.   
                                      
                                       
                                  
                                
                              
                            
                           
                            
                          
                           
                      
                    
                    
                      
                           
                          
                               Heatmaps    PCA     
                              
                                  Heatmaps  
                              
                               
                              
                                  New.Annotation  
                              
                              
                            
                              
                                
                                  
                                       
                                      
                                           Heatmap of EPHB-mediated forward signaling Data   More Plot Information                      Heatmap of EPHB-mediated forward signaling Data  Heatmap of the normalized data, scaled to give all genes equal variance, generated via unsupervised clustering. Orange indicates high expression; blue indicates low expression. This plot is meant to provide a high level exploratory view of the data.   
                                      
                                       
                                  
                                
                              
                            
                            
                              
                                
                                  
                                       
                                      
                                           Principal Components of EPHB-mediated forward signaling Data   More Plot Information                      Principal Components of EPHB-mediated forward signaling Data  Principal component analysis maps high-dimensional datasets onto a smaller number of highly informative dimensions. Here, the first four principal components of the gene expression data are plotted against each other and colored by the values of the selected covariate. This plot may be used to identify clusters in the data and to identify variables associated with prominent signal in the data. Variables that are associated with these leading principal components should be considered in downstream analyses.   
                                           Outliers   More Plot Information     
                                           
                                             
                                           
                                           
                                             
                                               
                                               Outlier on PC 1 
                                               Outlier on PC 2 
                                               Outlier on PC 3 
                                               Outlier on PC 4 
                                             
                                           
                                           
                                             
                                               20161013_sebelius2-101316_01_01.RCC 
                                               FALSE 
                                               FALSE 
                                               FALSE 
                                               TRUE 
                                             
                                           
                                           Outliers  Table identifying outliers in the first four PCs of the data.   
                                      
                                       
                                  
                                
                              
                            
                           
                            
                          
                           
                      
                    
                    
                      
                           
                          
                               Heatmaps    PCA     
                              
                                  Heatmaps  
                              
                               
                              
                                  New.Annotation  
                              
                              
                            
                              
                                
                                  
                                       
                                      
                                           Heatmap of Ephrin signaling Data   More Plot Information                      Heatmap of Ephrin signaling Data  Heatmap of the normalized data, scaled to give all genes equal variance, generated via unsupervised clustering. Orange indicates high expression; blue indicates low expression. This plot is meant to provide a high level exploratory view of the data.   
                                      
                                       
                                  
                                
                              
                            
                            
                              
                                
                                  
                                       
                                      
                                           Principal Components of Ephrin signaling Data   More Plot Information                      Principal Components of Ephrin signaling Data  Principal component analysis maps high-dimensional datasets onto a smaller number of highly informative dimensions. Here, the first four principal components of the gene expression data are plotted against each other and colored by the values of the selected covariate. This plot may be used to identify clusters in the data and to identify variables associated with prominent signal in the data. Variables that are associated with these leading principal components should be considered in downstream analyses.   
                                           Outliers   More Plot Information     
                                           
                                             
                                           
                                           
                                             
                                               
                                               Outlier on PC 1 
                                               Outlier on PC 2 
                                               Outlier on PC 3 
                                               Outlier on PC 4 
                                             
                                           
                                           
                                             
                                               20161013_sebelius2-101316_01_01.RCC 
                                               FALSE 
                                               FALSE 
                                               FALSE 
                                               TRUE 
                                             
                                           
                                           Outliers  Table identifying outliers in the first four PCs of the data.   
                                      
                                       
                                  
                                
                              
                            
                           
                            
                          
                           
                      
                    
                    
                      
                           
                          
                               Heatmaps    PCA     
                              
                                  Heatmaps  
                              
                               
                              
                                  New.Annotation  
                              
                              
                            
                              
                                
                                  
                                       
                                      
                                           Heatmap of ERKs are inactivated Data   More Plot Information                      Heatmap of ERKs are inactivated Data  Heatmap of the normalized data, scaled to give all genes equal variance, generated via unsupervised clustering. Orange indicates high expression; blue indicates low expression. This plot is meant to provide a high level exploratory view of the data.   
                                      
                                       
                                  
                                
                              
                            
                            
                              
                                
                                  
                                       
                                      
                                           Principal Components of ERKs are inactivated Data   More Plot Information                      Principal Components of ERKs are inactivated Data  Principal component analysis maps high-dimensional datasets onto a smaller number of highly informative dimensions. Here, the first four principal components of the gene expression data are plotted against each other and colored by the values of the selected covariate. This plot may be used to identify clusters in the data and to identify variables associated with prominent signal in the data. Variables that are associated with these leading principal components should be considered in downstream analyses.   
                                           Outliers   More Plot Information     
                                           
                                             
                                           
                                           
                                             
                                               
                                               Outlier on PC 1 
                                               Outlier on PC 2 
                                               Outlier on PC 3 
                                               Outlier on PC 4 
                                             
                                           
                                           
                                             
                                               20161013_sebelius2-101316_01_01.RCC 
                                               FALSE 
                                               FALSE 
                                               FALSE 
                                               TRUE 
                                             
                                           
                                           Outliers  Table identifying outliers in the first four PCs of the data.   
                                      
                                       
                                  
                                
                              
                            
                           
                            
                          
                           
                      
                    
                    
                      
                           
                          
                               Heatmaps    PCA     
                              
                                  Heatmaps  
                              
                               
                              
                                  New.Annotation  
                              
                              
                            
                              
                                
                                  
                                       
                                      
                                           Heatmap of FCERI mediated MAPK activation Data   More Plot Information                      Heatmap of FCERI mediated MAPK activation Data  Heatmap of the normalized data, scaled to give all genes equal variance, generated via unsupervised clustering. Orange indicates high expression; blue indicates low expression. This plot is meant to provide a high level exploratory view of the data.   
                                      
                                       
                                  
                                
                              
                            
                            
                              
                                
                                  
                                       
                                      
                                           Principal Components of FCERI mediated MAPK activation Data   More Plot Information                      Principal Components of FCERI mediated MAPK activation Data  Principal component analysis maps high-dimensional datasets onto a smaller number of highly informative dimensions. Here, the first four principal components of the gene expression data are plotted against each other and colored by the values of the selected covariate. This plot may be used to identify clusters in the data and to identify variables associated with prominent signal in the data. Variables that are associated with these leading principal components should be considered in downstream analyses.   
                                           Outliers   More Plot Information     
                                           
                                             
                                           
                                           
                                             
                                               
                                               Outlier on PC 1 
                                               Outlier on PC 2 
                                               Outlier on PC 3 
                                               Outlier on PC 4 
                                             
                                           
                                           
                                             
                                               20161013_sebelius2-101316_01_01.RCC 
                                               FALSE 
                                               FALSE 
                                               FALSE 
                                               TRUE 
                                             
                                           
                                           Outliers  Table identifying outliers in the first four PCs of the data.   
                                      
                                       
                                  
                                
                              
                            
                           
                            
                          
                           
                      
                    
                    
                      
                           
                          
                               Heatmaps    PCA     
                              
                                  Heatmaps  
                              
                               
                              
                                  New.Annotation  
                              
                              
                            
                              
                                
                                  
                                       
                                      
                                           Heatmap of Formation of the beta-catenin.CL.TCF transactivating complex Data   More Plot Information                      Heatmap of Formation of the beta-catenin.CL.TCF transactivating complex Data  Heatmap of the normalized data, scaled to give all genes equal variance, generated via unsupervised clustering. Orange indicates high expression; blue indicates low expression. This plot is meant to provide a high level exploratory view of the data.   
                                      
                                       
                                  
                                
                              
                            
                            
                              
                                
                                  
                                       
                                      
                                           Principal Components of Formation of the beta-catenin.CL.TCF transactivating complex Data   More Plot Information                      Principal Components of Formation of the beta-catenin.CL.TCF transactivating complex Data  Principal component analysis maps high-dimensional datasets onto a smaller number of highly informative dimensions. Here, the first four principal components of the gene expression data are plotted against each other and colored by the values of the selected covariate. This plot may be used to identify clusters in the data and to identify variables associated with prominent signal in the data. Variables that are associated with these leading principal components should be considered in downstream analyses.   
                                           Outliers   More Plot Information     
                                           
                                             
                                           
                                           
                                             
                                               
                                               Outlier on PC 1 
                                               Outlier on PC 2 
                                               Outlier on PC 3 
                                               Outlier on PC 4 
                                             
                                           
                                           
                                             
                                               20161013_sebelius2-101316_01_01.RCC 
                                               FALSE 
                                               FALSE 
                                               FALSE 
                                               TRUE 
                                             
                                           
                                           Outliers  Table identifying outliers in the first four PCs of the data.   
                                      
                                       
                                  
                                
                              
                            
                           
                            
                          
                           
                      
                    
                    
                      
                           
                          
                               Heatmaps    PCA     
                              
                                  Heatmaps  
                              
                               
                              
                                  New.Annotation  
                              
                              
                            
                              
                                
                                  
                                       
                                      
                                           Heatmap of Frs2-mediated activation Data   More Plot Information                      Heatmap of Frs2-mediated activation Data  Heatmap of the normalized data, scaled to give all genes equal variance, generated via unsupervised clustering. Orange indicates high expression; blue indicates low expression. This plot is meant to provide a high level exploratory view of the data.   
                                      
                                       
                                  
                                
                              
                            
                            
                              
                                
                                  
                                       
                                      
                                           Principal Components of Frs2-mediated activation Data   More Plot Information                      Principal Components of Frs2-mediated activation Data  Principal component analysis maps high-dimensional datasets onto a smaller number of highly informative dimensions. Here, the first four principal components of the gene expression data are plotted against each other and colored by the values of the selected covariate. This plot may be used to identify clusters in the data and to identify variables associated with prominent signal in the data. Variables that are associated with these leading principal components should be considered in downstream analyses.   
                                           Outliers   More Plot Information     
                                           
                                             
                                           
                                           
                                             
                                               
                                               Outlier on PC 1 
                                               Outlier on PC 2 
                                               Outlier on PC 3 
                                               Outlier on PC 4 
                                             
                                           
                                           
                                             
                                               20161013_sebelius2-101316_01_01.RCC 
                                               FALSE 
                                               FALSE 
                                               FALSE 
                                               TRUE 
                                             
                                           
                                           Outliers  Table identifying outliers in the first four PCs of the data.   
                                      
                                       
                                  
                                
                              
                            
                           
                            
                          
                           
                      
                    
                    
                      
                           
                          
                               Heatmaps    PCA     
                              
                                  Heatmaps  
                              
                               
                              
                                  New.Annotation  
                              
                              
                            
                              
                                
                                  
                                       
                                      
                                           Heatmap of G alpha (1213) signalling events Data   More Plot Information                      Heatmap of G alpha (1213) signalling events Data  Heatmap of the normalized data, scaled to give all genes equal variance, generated via unsupervised clustering. Orange indicates high expression; blue indicates low expression. This plot is meant to provide a high level exploratory view of the data.   
                                      
                                       
                                  
                                
                              
                            
                            
                              
                                
                                  
                                       
                                      
                                           Principal Components of G alpha (1213) signalling events Data   More Plot Information                      Principal Components of G alpha (1213) signalling events Data  Principal component analysis maps high-dimensional datasets onto a smaller number of highly informative dimensions. Here, the first four principal components of the gene expression data are plotted against each other and colored by the values of the selected covariate. This plot may be used to identify clusters in the data and to identify variables associated with prominent signal in the data. Variables that are associated with these leading principal components should be considered in downstream analyses.   
                                           Outliers   More Plot Information     
                                           
                                             
                                           
                                           
                                             
                                               
                                               Outlier on PC 1 
                                               Outlier on PC 2 
                                               Outlier on PC 3 
                                               Outlier on PC 4 
                                             
                                           
                                           
                                             
                                               20161013_sebelius2-101316_01_01.RCC 
                                               FALSE 
                                               FALSE 
                                               FALSE 
                                               TRUE 
                                             
                                           
                                           Outliers  Table identifying outliers in the first four PCs of the data.   
                                      
                                       
                                  
                                
                              
                            
                           
                            
                          
                           
                      
                    
                    
                      
                           
                          
                               Heatmaps    PCA     
                              
                                  Heatmaps  
                              
                               
                              
                                  New.Annotation  
                              
                              
                            
                              
                                
                                  
                                       
                                      
                                           Heatmap of G alpha (q) signalling events Data   More Plot Information                      Heatmap of G alpha (q) signalling events Data  Heatmap of the normalized data, scaled to give all genes equal variance, generated via unsupervised clustering. Orange indicates high expression; blue indicates low expression. This plot is meant to provide a high level exploratory view of the data.   
                                      
                                       
                                  
                                
                              
                            
                            
                              
                                
                                  
                                       
                                      
                                           Principal Components of G alpha (q) signalling events Data   More Plot Information                      Principal Components of G alpha (q) signalling events Data  Principal component analysis maps high-dimensional datasets onto a smaller number of highly informative dimensions. Here, the first four principal components of the gene expression data are plotted against each other and colored by the values of the selected covariate. This plot may be used to identify clusters in the data and to identify variables associated with prominent signal in the data. Variables that are associated with these leading principal components should be considered in downstream analyses.   
                                           Outliers   More Plot Information     
                                           
                                             
                                           
                                           
                                             
                                               
                                               Outlier on PC 1 
                                               Outlier on PC 2 
                                               Outlier on PC 3 
                                               Outlier on PC 4 
                                             
                                           
                                           
                                             
                                               20161013_sebelius2-101316_01_01.RCC 
                                               FALSE 
                                               FALSE 
                                               FALSE 
                                               TRUE 
                                             
                                           
                                           Outliers  Table identifying outliers in the first four PCs of the data.   
                                      
                                       
                                  
                                
                              
                            
                           
                            
                          
                           
                      
                    
                    
                      
                           
                          
                               Heatmaps    PCA     
                              
                                  Heatmaps  
                              
                               
                              
                                  New.Annotation  
                              
                              
                            
                              
                                
                                  
                                       
                                      
                                           Heatmap of G beta.CL.gamma signalling through PI3Kgamma Data   More Plot Information                      Heatmap of G beta.CL.gamma signalling through PI3Kgamma Data  Heatmap of the normalized data, scaled to give all genes equal variance, generated via unsupervised clustering. Orange indicates high expression; blue indicates low expression. This plot is meant to provide a high level exploratory view of the data.   
                                      
                                       
                                  
                                
                              
                            
                            
                              
                                
                                  
                                       
                                      
                                           Principal Components of G beta.CL.gamma signalling through PI3Kgamma Data   More Plot Information                      Principal Components of G beta.CL.gamma signalling through PI3Kgamma Data  Principal component analysis maps high-dimensional datasets onto a smaller number of highly informative dimensions. Here, the first four principal components of the gene expression data are plotted against each other and colored by the values of the selected covariate. This plot may be used to identify clusters in the data and to identify variables associated with prominent signal in the data. Variables that are associated with these leading principal components should be considered in downstream analyses.   
                                           Outliers   More Plot Information     
                                           
                                             
                                           
                                           
                                             
                                               
                                               Outlier on PC 1 
                                               Outlier on PC 2 
                                               Outlier on PC 3 
                                               Outlier on PC 4 
                                             
                                           
                                           
                                             
                                               20161013_sebelius2-101316_01_01.RCC 
                                               FALSE 
                                               FALSE 
                                               FALSE 
                                               TRUE 
                                             
                                           
                                           Outliers  Table identifying outliers in the first four PCs of the data.   
                                      
                                       
                                  
                                
                              
                            
                           
                            
                          
                           
                      
                    
                    
                      
                           
                          
                               Heatmaps    PCA     
                              
                                  Heatmaps  
                              
                               
                              
                                  New.Annotation  
                              
                              
                            
                              
                                
                                  
                                       
                                      
                                           Heatmap of GAB1 signalosome Data   More Plot Information                      Heatmap of GAB1 signalosome Data  Heatmap of the normalized data, scaled to give all genes equal variance, generated via unsupervised clustering. Orange indicates high expression; blue indicates low expression. This plot is meant to provide a high level exploratory view of the data.   
                                      
                                       
                                  
                                
                              
                            
                            
                              
                                
                                  
                                       
                                      
                                           Principal Components of GAB1 signalosome Data   More Plot Information                      Principal Components of GAB1 signalosome Data  Principal component analysis maps high-dimensional datasets onto a smaller number of highly informative dimensions. Here, the first four principal components of the gene expression data are plotted against each other and colored by the values of the selected covariate. This plot may be used to identify clusters in the data and to identify variables associated with prominent signal in the data. Variables that are associated with these leading principal components should be considered in downstream analyses.   
                                           Outliers   More Plot Information     
                                           
                                             
                                           
                                           
                                             
                                               
                                               Outlier on PC 1 
                                               Outlier on PC 2 
                                               Outlier on PC 3 
                                               Outlier on PC 4 
                                             
                                           
                                           
                                             
                                               20161013_sebelius2-101316_01_01.RCC 
                                               FALSE 
                                               FALSE 
                                               FALSE 
                                               TRUE 
                                             
                                           
                                           Outliers  Table identifying outliers in the first four PCs of the data.   
                                      
                                       
                                  
                                
                              
                            
                           
                            
                          
                           
                      
                    
                    
                      
                           
                          
                               Heatmaps    PCA     
                              
                                  Heatmaps  
                              
                               
                              
                                  New.Annotation  
                              
                              
                            
                              
                                
                                  
                                       
                                      
                                           Heatmap of Gastrin-CREB signalling pathway via PKC and MAPK Data   More Plot Information                      Heatmap of Gastrin-CREB signalling pathway via PKC and MAPK Data  Heatmap of the normalized data, scaled to give all genes equal variance, generated via unsupervised clustering. Orange indicates high expression; blue indicates low expression. This plot is meant to provide a high level exploratory view of the data.   
                                      
                                       
                                  
                                
                              
                            
                            
                              
                                
                                  
                                       
                                      
                                           Principal Components of Gastrin-CREB signalling pathway via PKC and MAPK Data   More Plot Information                      Principal Components of Gastrin-CREB signalling pathway via PKC and MAPK Data  Principal component analysis maps high-dimensional datasets onto a smaller number of highly informative dimensions. Here, the first four principal components of the gene expression data are plotted against each other and colored by the values of the selected covariate. This plot may be used to identify clusters in the data and to identify variables associated with prominent signal in the data. Variables that are associated with these leading principal components should be considered in downstream analyses.   
                                           Outliers   More Plot Information     
                                           
                                             
                                           
                                           
                                             
                                               
                                               Outlier on PC 1 
                                               Outlier on PC 2 
                                               Outlier on PC 3 
                                               Outlier on PC 4 
                                             
                                           
                                           
                                             
                                               20161013_sebelius2-101316_01_01.RCC 
                                               FALSE 
                                               FALSE 
                                               FALSE 
                                               TRUE 
                                             
                                           
                                           Outliers  Table identifying outliers in the first four PCs of the data.   
                                      
                                       
                                  
                                
                              
                            
                           
                            
                          
                           
                      
                    
                    
                      
                           
                          
                               Heatmaps    PCA     
                              
                                  Heatmaps  
                              
                               
                              
                                  New.Annotation  
                              
                              
                            
                              
                                
                                  
                                       
                                      
                                           Heatmap of GPVI-mediated activation cascade Data   More Plot Information                      Heatmap of GPVI-mediated activation cascade Data  Heatmap of the normalized data, scaled to give all genes equal variance, generated via unsupervised clustering. Orange indicates high expression; blue indicates low expression. This plot is meant to provide a high level exploratory view of the data.   
                                      
                                       
                                  
                                
                              
                            
                            
                              
                                
                                  
                                       
                                      
                                           Principal Components of GPVI-mediated activation cascade Data   More Plot Information                      Principal Components of GPVI-mediated activation cascade Data  Principal component analysis maps high-dimensional datasets onto a smaller number of highly informative dimensions. Here, the first four principal components of the gene expression data are plotted against each other and colored by the values of the selected covariate. This plot may be used to identify clusters in the data and to identify variables associated with prominent signal in the data. Variables that are associated with these leading principal components should be considered in downstream analyses.   
                                           Outliers   More Plot Information     
                                           
                                             
                                           
                                           
                                             
                                               
                                               Outlier on PC 1 
                                               Outlier on PC 2 
                                               Outlier on PC 3 
                                               Outlier on PC 4 
                                             
                                           
                                           
                                             
                                               20161013_sebelius2-101316_01_01.RCC 
                                               FALSE 
                                               FALSE 
                                               FALSE 
                                               TRUE 
                                             
                                           
                                           Outliers  Table identifying outliers in the first four PCs of the data.   
                                      
                                       
                                  
                                
                              
                            
                           
                            
                          
                           
                      
                    
                    
                      
                           
                          
                               Heatmaps    PCA     
                              
                                  Heatmaps  
                              
                               
                              
                                  New.Annotation  
                              
                              
                            
                              
                                
                                  
                                       
                                      
                                           Heatmap of Integrin alphaIIb beta3 signaling Data   More Plot Information                      Heatmap of Integrin alphaIIb beta3 signaling Data  Heatmap of the normalized data, scaled to give all genes equal variance, generated via unsupervised clustering. Orange indicates high expression; blue indicates low expression. This plot is meant to provide a high level exploratory view of the data.   
                                      
                                       
                                  
                                
                              
                            
                            
                              
                                
                                  
                                       
                                      
                                           Principal Components of Integrin alphaIIb beta3 signaling Data   More Plot Information                      Principal Components of Integrin alphaIIb beta3 signaling Data  Principal component analysis maps high-dimensional datasets onto a smaller number of highly informative dimensions. Here, the first four principal components of the gene expression data are plotted against each other and colored by the values of the selected covariate. This plot may be used to identify clusters in the data and to identify variables associated with prominent signal in the data. Variables that are associated with these leading principal components should be considered in downstream analyses.   
                                           Outliers   More Plot Information     
                                           
                                             
                                           
                                           
                                             
                                               
                                               Outlier on PC 1 
                                               Outlier on PC 2 
                                               Outlier on PC 3 
                                               Outlier on PC 4 
                                             
                                           
                                           
                                             
                                               20161013_sebelius2-101316_01_01.RCC 
                                               FALSE 
                                               FALSE 
                                               FALSE 
                                               TRUE 
                                             
                                           
                                           Outliers  Table identifying outliers in the first four PCs of the data.   
                                      
                                       
                                  
                                
                              
                            
                           
                            
                          
                           
                      
                    
                    
                      
                           
                          
                               Heatmaps    PCA     
                              
                                  Heatmaps  
                              
                               
                              
                                  New.Annotation  
                              
                              
                            
                              
                                
                                  
                                       
                                      
                                           Heatmap of Integrin cell surface interactions Data   More Plot Information                      Heatmap of Integrin cell surface interactions Data  Heatmap of the normalized data, scaled to give all genes equal variance, generated via unsupervised clustering. Orange indicates high expression; blue indicates low expression. This plot is meant to provide a high level exploratory view of the data.   
                                      
                                       
                                  
                                
                              
                            
                            
                              
                                
                                  
                                       
                                      
                                           Principal Components of Integrin cell surface interactions Data   More Plot Information                      Principal Components of Integrin cell surface interactions Data  Principal component analysis maps high-dimensional datasets onto a smaller number of highly informative dimensions. Here, the first four principal components of the gene expression data are plotted against each other and colored by the values of the selected covariate. This plot may be used to identify clusters in the data and to identify variables associated with prominent signal in the data. Variables that are associated with these leading principal components should be considered in downstream analyses.   
                                           Outliers   More Plot Information     
                                           
                                             
                                           
                                           
                                             
                                               
                                               Outlier on PC 1 
                                               Outlier on PC 2 
                                               Outlier on PC 3 
                                               Outlier on PC 4 
                                             
                                           
                                           
                                             
                                               20161013_sebelius2-101316_01_01.RCC 
                                               FALSE 
                                               FALSE 
                                               FALSE 
                                               TRUE 
                                             
                                           
                                           Outliers  Table identifying outliers in the first four PCs of the data.   
                                      
                                       
                                  
                                
                              
                            
                           
                            
                          
                           
                      
                    
                    
                      
                           
                          
                               Heatmaps    PCA     
                              
                                  Heatmaps  
                              
                               
                              
                                  New.Annotation  
                              
                              
                            
                              
                                
                                  
                                       
                                      
                                           Heatmap of Interleukin receptor SHC signaling Data   More Plot Information                      Heatmap of Interleukin receptor SHC signaling Data  Heatmap of the normalized data, scaled to give all genes equal variance, generated via unsupervised clustering. Orange indicates high expression; blue indicates low expression. This plot is meant to provide a high level exploratory view of the data.   
                                      
                                       
                                  
                                
                              
                            
                            
                              
                                
                                  
                                       
                                      
                                           Principal Components of Interleukin receptor SHC signaling Data   More Plot Information                      Principal Components of Interleukin receptor SHC signaling Data  Principal component analysis maps high-dimensional datasets onto a smaller number of highly informative dimensions. Here, the first four principal components of the gene expression data are plotted against each other and colored by the values of the selected covariate. This plot may be used to identify clusters in the data and to identify variables associated with prominent signal in the data. Variables that are associated with these leading principal components should be considered in downstream analyses.   
                                           Outliers   More Plot Information     
                                           
                                             
                                           
                                           
                                             
                                               
                                               Outlier on PC 1 
                                               Outlier on PC 2 
                                               Outlier on PC 3 
                                               Outlier on PC 4 
                                             
                                           
                                           
                                             
                                               20161013_sebelius2-101316_01_01.RCC 
                                               FALSE 
                                               FALSE 
                                               FALSE 
                                               TRUE 
                                             
                                           
                                           Outliers  Table identifying outliers in the first four PCs of the data.   
                                      
                                       
                                  
                                
                              
                            
                           
                            
                          
                           
                      
                    
                    
                      
                           
                          
                               Heatmaps    PCA     
                              
                                  Heatmaps  
                              
                               
                              
                                  New.Annotation  
                              
                              
                            
                              
                                
                                  
                                       
                                      
                                           Heatmap of IRS-related events triggered by IGF1R Data   More Plot Information                      Heatmap of IRS-related events triggered by IGF1R Data  Heatmap of the normalized data, scaled to give all genes equal variance, generated via unsupervised clustering. Orange indicates high expression; blue indicates low expression. This plot is meant to provide a high level exploratory view of the data.   
                                      
                                       
                                  
                                
                              
                            
                            
                              
                                
                                  
                                       
                                      
                                           Principal Components of IRS-related events triggered by IGF1R Data   More Plot Information                      Principal Components of IRS-related events triggered by IGF1R Data  Principal component analysis maps high-dimensional datasets onto a smaller number of highly informative dimensions. Here, the first four principal components of the gene expression data are plotted against each other and colored by the values of the selected covariate. This plot may be used to identify clusters in the data and to identify variables associated with prominent signal in the data. Variables that are associated with these leading principal components should be considered in downstream analyses.   
                                           Outliers   More Plot Information     
                                           
                                             
                                           
                                           
                                             
                                               
                                               Outlier on PC 1 
                                               Outlier on PC 2 
                                               Outlier on PC 3 
                                               Outlier on PC 4 
                                             
                                           
                                           
                                             
                                               20161013_sebelius2-101316_01_01.RCC 
                                               FALSE 
                                               FALSE 
                                               FALSE 
                                               TRUE 
                                             
                                           
                                           Outliers  Table identifying outliers in the first four PCs of the data.   
                                      
                                       
                                  
                                
                              
                            
                           
                            
                          
                           
                      
                    
                    
                      
                           
                          
                               Heatmaps    PCA     
                              
                                  Heatmaps  
                              
                               
                              
                                  New.Annotation  
                              
                              
                            
                              
                                
                                  
                                       
                                      
                                           Heatmap of Laminin interactions Data   More Plot Information                      Heatmap of Laminin interactions Data  Heatmap of the normalized data, scaled to give all genes equal variance, generated via unsupervised clustering. Orange indicates high expression; blue indicates low expression. This plot is meant to provide a high level exploratory view of the data.   
                                      
                                       
                                  
                                
                              
                            
                            
                              
                                
                                  
                                       
                                      
                                           Principal Components of Laminin interactions Data   More Plot Information                      Principal Components of Laminin interactions Data  Principal component analysis maps high-dimensional datasets onto a smaller number of highly informative dimensions. Here, the first four principal components of the gene expression data are plotted against each other and colored by the values of the selected covariate. This plot may be used to identify clusters in the data and to identify variables associated with prominent signal in the data. Variables that are associated with these leading principal components should be considered in downstream analyses.   
                                           Outliers   More Plot Information     
                                           
                                             
                                           
                                           
                                             
                                               
                                               Outlier on PC 1 
                                               Outlier on PC 2 
                                               Outlier on PC 3 
                                               Outlier on PC 4 
                                             
                                           
                                           
                                             
                                               20161013_sebelius2-101316_01_01.RCC 
                                               FALSE 
                                               FALSE 
                                               FALSE 
                                               TRUE 
                                             
                                           
                                           Outliers  Table identifying outliers in the first four PCs of the data.   
                                      
                                       
                                  
                                
                              
                            
                           
                            
                          
                           
                      
                    
                    
                      
                           
                          
                               Heatmaps    PCA     
                              
                                  Heatmaps  
                              
                               
                              
                                  New.Annotation  
                              
                              
                            
                              
                                
                                  
                                       
                                      
                                           Heatmap of MAP2K and MAPK activation Data   More Plot Information                      Heatmap of MAP2K and MAPK activation Data  Heatmap of the normalized data, scaled to give all genes equal variance, generated via unsupervised clustering. Orange indicates high expression; blue indicates low expression. This plot is meant to provide a high level exploratory view of the data.   
                                      
                                       
                                  
                                
                              
                            
                            
                              
                                
                                  
                                       
                                      
                                           Principal Components of MAP2K and MAPK activation Data   More Plot Information                      Principal Components of MAP2K and MAPK activation Data  Principal component analysis maps high-dimensional datasets onto a smaller number of highly informative dimensions. Here, the first four principal components of the gene expression data are plotted against each other and colored by the values of the selected covariate. This plot may be used to identify clusters in the data and to identify variables associated with prominent signal in the data. Variables that are associated with these leading principal components should be considered in downstream analyses.   
                                           Outliers   More Plot Information     
                                           
                                             
                                           
                                           
                                             
                                               
                                               Outlier on PC 1 
                                               Outlier on PC 2 
                                               Outlier on PC 3 
                                               Outlier on PC 4 
                                             
                                           
                                           
                                             
                                               20161013_sebelius2-101316_01_01.RCC 
                                               FALSE 
                                               FALSE 
                                               FALSE 
                                               TRUE 
                                             
                                           
                                           Outliers  Table identifying outliers in the first four PCs of the data.   
                                      
                                       
                                  
                                
                              
                            
                           
                            
                          
                           
                      
                    
                    
                      
                           
                          
                               Heatmaps    PCA     
                              
                                  Heatmaps  
                              
                               
                              
                                  New.Annotation  
                              
                              
                            
                              
                                
                                  
                                       
                                      
                                           Heatmap of MAPK6MAPK4 signaling Data   More Plot Information                      Heatmap of MAPK6MAPK4 signaling Data  Heatmap of the normalized data, scaled to give all genes equal variance, generated via unsupervised clustering. Orange indicates high expression; blue indicates low expression. This plot is meant to provide a high level exploratory view of the data.   
                                      
                                       
                                  
                                
                              
                            
                            
                              
                                
                                  
                                       
                                      
                                           Principal Components of MAPK6MAPK4 signaling Data   More Plot Information                      Principal Components of MAPK6MAPK4 signaling Data  Principal component analysis maps high-dimensional datasets onto a smaller number of highly informative dimensions. Here, the first four principal components of the gene expression data are plotted against each other and colored by the values of the selected covariate. This plot may be used to identify clusters in the data and to identify variables associated with prominent signal in the data. Variables that are associated with these leading principal components should be considered in downstream analyses.   
                                           Outliers   More Plot Information     
                                           
                                             
                                           
                                           
                                             
                                               
                                               Outlier on PC 1 
                                               Outlier on PC 2 
                                               Outlier on PC 3 
                                               Outlier on PC 4 
                                             
                                           
                                           
                                             
                                               20161013_sebelius2-101316_01_01.RCC 
                                               FALSE 
                                               FALSE 
                                               FALSE 
                                               TRUE 
                                             
                                           
                                           Outliers  Table identifying outliers in the first four PCs of the data.   
                                      
                                       
                                  
                                
                              
                            
                           
                            
                          
                           
                      
                    
                    
                      
                           
                          
                               Heatmaps    PCA     
                              
                                  Heatmaps  
                              
                               
                              
                                  New.Annotation  
                              
                              
                            
                              
                                
                                  
                                       
                                      
                                           Heatmap of MASTL Facilitates Mitotic Progression Data   More Plot Information                      Heatmap of MASTL Facilitates Mitotic Progression Data  Heatmap of the normalized data, scaled to give all genes equal variance, generated via unsupervised clustering. Orange indicates high expression; blue indicates low expression. This plot is meant to provide a high level exploratory view of the data.   
                                      
                                       
                                  
                                
                              
                            
                            
                              
                                
                                  
                                       
                                      
                                           Principal Components of MASTL Facilitates Mitotic Progression Data   More Plot Information                      Principal Components of MASTL Facilitates Mitotic Progression Data  Principal component analysis maps high-dimensional datasets onto a smaller number of highly informative dimensions. Here, the first four principal components of the gene expression data are plotted against each other and colored by the values of the selected covariate. This plot may be used to identify clusters in the data and to identify variables associated with prominent signal in the data. Variables that are associated with these leading principal components should be considered in downstream analyses.   
                                           Outliers   More Plot Information     
                                           
                                             
                                           
                                           
                                             
                                               
                                               Outlier on PC 1 
                                               Outlier on PC 2 
                                               Outlier on PC 3 
                                               Outlier on PC 4 
                                             
                                           
                                           
                                             
                                               20161013_sebelius2-101316_01_01.RCC 
                                               FALSE 
                                               FALSE 
                                               FALSE 
                                               TRUE 
                                             
                                           
                                           Outliers  Table identifying outliers in the first four PCs of the data.   
                                      
                                       
                                  
                                
                              
                            
                           
                            
                          
                           
                      
                    
                    
                      
                           
                          
                               Heatmaps    PCA     
                              
                                  Heatmaps  
                              
                               
                              
                                  New.Annotation  
                              
                              
                            
                              
                                
                                  
                                       
                                      
                                           Heatmap of MicroRNA (miRNA) biogenesis Data   More Plot Information                      Heatmap of MicroRNA (miRNA) biogenesis Data  Heatmap of the normalized data, scaled to give all genes equal variance, generated via unsupervised clustering. Orange indicates high expression; blue indicates low expression. This plot is meant to provide a high level exploratory view of the data.   
                                      
                                       
                                  
                                
                              
                            
                            
                              
                                
                                  
                                       
                                      
                                           Principal Components of MicroRNA (miRNA) biogenesis Data   More Plot Information                      Principal Components of MicroRNA (miRNA) biogenesis Data  Principal component analysis maps high-dimensional datasets onto a smaller number of highly informative dimensions. Here, the first four principal components of the gene expression data are plotted against each other and colored by the values of the selected covariate. This plot may be used to identify clusters in the data and to identify variables associated with prominent signal in the data. Variables that are associated with these leading principal components should be considered in downstream analyses.   
                                           Outliers   More Plot Information     
                                           
                                             
                                           
                                           
                                             
                                               
                                               Outlier on PC 1 
                                               Outlier on PC 2 
                                               Outlier on PC 3 
                                               Outlier on PC 4 
                                             
                                           
                                           
                                             
                                               20161013_sebelius2-101316_01_01.RCC 
                                               FALSE 
                                               FALSE 
                                               FALSE 
                                               TRUE 
                                             
                                           
                                           Outliers  Table identifying outliers in the first four PCs of the data.   
                                      
                                       
                                  
                                
                              
                            
                           
                            
                          
                           
                      
                    
                    
                      
                           
                          
                               Heatmaps    PCA     
                              
                                  Heatmaps  
                              
                               
                              
                                  New.Annotation  
                              
                              
                            
                              
                                
                                  
                                       
                                      
                                           Heatmap of Misspliced GSK3beta mutants stabilize beta-catenin Data   More Plot Information                      Heatmap of Misspliced GSK3beta mutants stabilize beta-catenin Data  Heatmap of the normalized data, scaled to give all genes equal variance, generated via unsupervised clustering. Orange indicates high expression; blue indicates low expression. This plot is meant to provide a high level exploratory view of the data.   
                                      
                                       
                                  
                                
                              
                            
                            
                              
                                
                                  
                                       
                                      
                                           Principal Components of Misspliced GSK3beta mutants stabilize beta-catenin Data   More Plot Information                      Principal Components of Misspliced GSK3beta mutants stabilize beta-catenin Data  Principal component analysis maps high-dimensional datasets onto a smaller number of highly informative dimensions. Here, the first four principal components of the gene expression data are plotted against each other and colored by the values of the selected covariate. This plot may be used to identify clusters in the data and to identify variables associated with prominent signal in the data. Variables that are associated with these leading principal components should be considered in downstream analyses.   
                                           Outliers   More Plot Information     
                                           
                                             
                                           
                                           
                                             
                                               
                                               Outlier on PC 1 
                                               Outlier on PC 2 
                                               Outlier on PC 3 
                                               Outlier on PC 4 
                                             
                                           
                                           
                                             
                                               20161013_sebelius2-101316_01_01.RCC 
                                               FALSE 
                                               FALSE 
                                               FALSE 
                                               TRUE 
                                             
                                           
                                           Outliers  Table identifying outliers in the first four PCs of the data.   
                                      
                                       
                                  
                                
                              
                            
                           
                            
                          
                           
                      
                    
                    
                      
                           
                          
                               Heatmaps    PCA     
                              
                                  Heatmaps  
                              
                               
                              
                                  New.Annotation  
                              
                              
                            
                              
                                
                                  
                                       
                                      
                                           Heatmap of Molecules associated with elastic fibres Data   More Plot Information                      Heatmap of Molecules associated with elastic fibres Data  Heatmap of the normalized data, scaled to give all genes equal variance, generated via unsupervised clustering. Orange indicates high expression; blue indicates low expression. This plot is meant to provide a high level exploratory view of the data.   
                                      
                                       
                                  
                                
                              
                            
                            
                              
                                
                                  
                                       
                                      
                                           Principal Components of Molecules associated with elastic fibres Data   More Plot Information                      Principal Components of Molecules associated with elastic fibres Data  Principal component analysis maps high-dimensional datasets onto a smaller number of highly informative dimensions. Here, the first four principal components of the gene expression data are plotted against each other and colored by the values of the selected covariate. This plot may be used to identify clusters in the data and to identify variables associated with prominent signal in the data. Variables that are associated with these leading principal components should be considered in downstream analyses.   
                                           Outliers   More Plot Information     
                                           
                                             
                                           
                                           
                                             
                                               
                                               Outlier on PC 1 
                                               Outlier on PC 2 
                                               Outlier on PC 3 
                                               Outlier on PC 4 
                                             
                                           
                                           
                                             
                                               20161013_sebelius2-101316_01_01.RCC 
                                               FALSE 
                                               FALSE 
                                               FALSE 
                                               TRUE 
                                             
                                           
                                           Outliers  Table identifying outliers in the first four PCs of the data.   
                                      
                                       
                                  
                                
                              
                            
                           
                            
                          
                           
                      
                    
                    
                      
                           
                          
                               Heatmaps    PCA     
                              
                                  Heatmaps  
                              
                               
                              
                                  New.Annotation  
                              
                              
                            
                              
                                
                                  
                                       
                                      
                                           Heatmap of Negative feedback regulation of MAPK pathway Data   More Plot Information                      Heatmap of Negative feedback regulation of MAPK pathway Data  Heatmap of the normalized data, scaled to give all genes equal variance, generated via unsupervised clustering. Orange indicates high expression; blue indicates low expression. This plot is meant to provide a high level exploratory view of the data.   
                                      
                                       
                                  
                                
                              
                            
                            
                              
                                
                                  
                                       
                                      
                                           Principal Components of Negative feedback regulation of MAPK pathway Data   More Plot Information                      Principal Components of Negative feedback regulation of MAPK pathway Data  Principal component analysis maps high-dimensional datasets onto a smaller number of highly informative dimensions. Here, the first four principal components of the gene expression data are plotted against each other and colored by the values of the selected covariate. This plot may be used to identify clusters in the data and to identify variables associated with prominent signal in the data. Variables that are associated with these leading principal components should be considered in downstream analyses.   
                                           Outliers   More Plot Information     
                                           
                                             
                                           
                                           
                                             
                                               
                                               Outlier on PC 1 
                                               Outlier on PC 2 
                                               Outlier on PC 3 
                                               Outlier on PC 4 
                                             
                                           
                                           
                                             
                                               20161013_sebelius2-101316_01_01.RCC 
                                               FALSE 
                                               FALSE 
                                               FALSE 
                                               TRUE 
                                             
                                           
                                           Outliers  Table identifying outliers in the first four PCs of the data.   
                                      
                                       
                                  
                                
                              
                            
                           
                            
                          
                           
                      
                    
                    
                      
                           
                          
                               Heatmaps    PCA     
                              
                                  Heatmaps  
                              
                               
                              
                                  New.Annotation  
                              
                              
                            
                              
                                
                                  
                                       
                                      
                                           Heatmap of Negative regulation of FGFR1 signaling Data   More Plot Information                      Heatmap of Negative regulation of FGFR1 signaling Data  Heatmap of the normalized data, scaled to give all genes equal variance, generated via unsupervised clustering. Orange indicates high expression; blue indicates low expression. This plot is meant to provide a high level exploratory view of the data.   
                                      
                                       
                                  
                                
                              
                            
                            
                              
                                
                                  
                                       
                                      
                                           Principal Components of Negative regulation of FGFR1 signaling Data   More Plot Information                      Principal Components of Negative regulation of FGFR1 signaling Data  Principal component analysis maps high-dimensional datasets onto a smaller number of highly informative dimensions. Here, the first four principal components of the gene expression data are plotted against each other and colored by the values of the selected covariate. This plot may be used to identify clusters in the data and to identify variables associated with prominent signal in the data. Variables that are associated with these leading principal components should be considered in downstream analyses.   
                                           Outliers   More Plot Information     
                                           
                                             
                                           
                                           
                                             
                                               
                                               Outlier on PC 1 
                                               Outlier on PC 2 
                                               Outlier on PC 3 
                                               Outlier on PC 4 
                                             
                                           
                                           
                                             
                                               20161013_sebelius2-101316_01_01.RCC 
                                               FALSE 
                                               FALSE 
                                               FALSE 
                                               TRUE 
                                             
                                           
                                           Outliers  Table identifying outliers in the first four PCs of the data.   
                                      
                                       
                                  
                                
                              
                            
                           
                            
                          
                           
                      
                    
                    
                      
                           
                          
                               Heatmaps    PCA     
                              
                                  Heatmaps  
                              
                               
                              
                                  New.Annotation  
                              
                              
                            
                              
                                
                                  
                                       
                                      
                                           Heatmap of Negative regulation of FGFR2 signaling Data   More Plot Information                      Heatmap of Negative regulation of FGFR2 signaling Data  Heatmap of the normalized data, scaled to give all genes equal variance, generated via unsupervised clustering. Orange indicates high expression; blue indicates low expression. This plot is meant to provide a high level exploratory view of the data.   
                                      
                                       
                                  
                                
                              
                            
                            
                              
                                
                                  
                                       
                                      
                                           Principal Components of Negative regulation of FGFR2 signaling Data   More Plot Information                      Principal Components of Negative regulation of FGFR2 signaling Data  Principal component analysis maps high-dimensional datasets onto a smaller number of highly informative dimensions. Here, the first four principal components of the gene expression data are plotted against each other and colored by the values of the selected covariate. This plot may be used to identify clusters in the data and to identify variables associated with prominent signal in the data. Variables that are associated with these leading principal components should be considered in downstream analyses.   
                                           Outliers   More Plot Information     
                                           
                                             
                                           
                                           
                                             
                                               
                                               Outlier on PC 1 
                                               Outlier on PC 2 
                                               Outlier on PC 3 
                                               Outlier on PC 4 
                                             
                                           
                                           
                                             
                                               20161013_sebelius2-101316_01_01.RCC 
                                               FALSE 
                                               FALSE 
                                               FALSE 
                                               TRUE 
                                             
                                           
                                           Outliers  Table identifying outliers in the first four PCs of the data.   
                                      
                                       
                                  
                                
                              
                            
                           
                            
                          
                           
                      
                    
                    
                      
                           
                          
                               Heatmaps    PCA     
                              
                                  Heatmaps  
                              
                               
                              
                                  New.Annotation  
                              
                              
                            
                              
                                
                                  
                                       
                                      
                                           Heatmap of Negative regulation of FGFR3 signaling Data   More Plot Information                      Heatmap of Negative regulation of FGFR3 signaling Data  Heatmap of the normalized data, scaled to give all genes equal variance, generated via unsupervised clustering. Orange indicates high expression; blue indicates low expression. This plot is meant to provide a high level exploratory view of the data.   
                                      
                                       
                                  
                                
                              
                            
                            
                              
                                
                                  
                                       
                                      
                                           Principal Components of Negative regulation of FGFR3 signaling Data   More Plot Information                      Principal Components of Negative regulation of FGFR3 signaling Data  Principal component analysis maps high-dimensional datasets onto a smaller number of highly informative dimensions. Here, the first four principal components of the gene expression data are plotted against each other and colored by the values of the selected covariate. This plot may be used to identify clusters in the data and to identify variables associated with prominent signal in the data. Variables that are associated with these leading principal components should be considered in downstream analyses.   
                                           Outliers   More Plot Information     
                                           
                                             
                                           
                                           
                                             
                                               
                                               Outlier on PC 1 
                                               Outlier on PC 2 
                                               Outlier on PC 3 
                                               Outlier on PC 4 
                                             
                                           
                                           
                                             
                                               20161013_sebelius2-101316_01_01.RCC 
                                               FALSE 
                                               FALSE 
                                               FALSE 
                                               TRUE 
                                             
                                           
                                           Outliers  Table identifying outliers in the first four PCs of the data.   
                                      
                                       
                                  
                                
                              
                            
                           
                            
                          
                           
                      
                    
                    
                      
                           
                          
                               Heatmaps    PCA     
                              
                                  Heatmaps  
                              
                               
                              
                                  New.Annotation  
                              
                              
                            
                              
                                
                                  
                                       
                                      
                                           Heatmap of Negative regulation of FGFR4 signaling Data   More Plot Information                      Heatmap of Negative regulation of FGFR4 signaling Data  Heatmap of the normalized data, scaled to give all genes equal variance, generated via unsupervised clustering. Orange indicates high expression; blue indicates low expression. This plot is meant to provide a high level exploratory view of the data.   
                                      
                                       
                                  
                                
                              
                            
                            
                              
                                
                                  
                                       
                                      
                                           Principal Components of Negative regulation of FGFR4 signaling Data   More Plot Information                      Principal Components of Negative regulation of FGFR4 signaling Data  Principal component analysis maps high-dimensional datasets onto a smaller number of highly informative dimensions. Here, the first four principal components of the gene expression data are plotted against each other and colored by the values of the selected covariate. This plot may be used to identify clusters in the data and to identify variables associated with prominent signal in the data. Variables that are associated with these leading principal components should be considered in downstream analyses.   
                                           Outliers   More Plot Information     
                                           
                                             
                                           
                                           
                                             
                                               
                                               Outlier on PC 1 
                                               Outlier on PC 2 
                                               Outlier on PC 3 
                                               Outlier on PC 4 
                                             
                                           
                                           
                                             
                                               20161013_sebelius2-101316_01_01.RCC 
                                               FALSE 
                                               FALSE 
                                               FALSE 
                                               TRUE 
                                             
                                           
                                           Outliers  Table identifying outliers in the first four PCs of the data.   
                                      
                                       
                                  
                                
                              
                            
                           
                            
                          
                           
                      
                    
                    
                      
                           
                          
                               Heatmaps    PCA     
                              
                                  Heatmaps  
                              
                               
                              
                                  New.Annotation  
                              
                              
                            
                              
                                
                                  
                                       
                                      
                                           Heatmap of Negative regulation of MAPK pathway Data   More Plot Information                      Heatmap of Negative regulation of MAPK pathway Data  Heatmap of the normalized data, scaled to give all genes equal variance, generated via unsupervised clustering. Orange indicates high expression; blue indicates low expression. This plot is meant to provide a high level exploratory view of the data.   
                                      
                                       
                                  
                                
                              
                            
                            
                              
                                
                                  
                                       
                                      
                                           Principal Components of Negative regulation of MAPK pathway Data   More Plot Information                      Principal Components of Negative regulation of MAPK pathway Data  Principal component analysis maps high-dimensional datasets onto a smaller number of highly informative dimensions. Here, the first four principal components of the gene expression data are plotted against each other and colored by the values of the selected covariate. This plot may be used to identify clusters in the data and to identify variables associated with prominent signal in the data. Variables that are associated with these leading principal components should be considered in downstream analyses.   
                                           Outliers   More Plot Information     
                                           
                                             
                                           
                                           
                                             
                                               
                                               Outlier on PC 1 
                                               Outlier on PC 2 
                                               Outlier on PC 3 
                                               Outlier on PC 4 
                                             
                                           
                                           
                                             
                                               20161013_sebelius2-101316_01_01.RCC 
                                               FALSE 
                                               FALSE 
                                               FALSE 
                                               TRUE 
                                             
                                           
                                           Outliers  Table identifying outliers in the first four PCs of the data.   
                                      
                                       
                                  
                                
                              
                            
                           
                            
                          
                           
                      
                    
                    
                      
                           
                          
                               Heatmaps    PCA     
                              
                                  Heatmaps  
                              
                               
                              
                                  New.Annotation  
                              
                              
                            
                              
                                
                                  
                                       
                                      
                                           Heatmap of Nephrin interactions Data   More Plot Information                      Heatmap of Nephrin interactions Data  Heatmap of the normalized data, scaled to give all genes equal variance, generated via unsupervised clustering. Orange indicates high expression; blue indicates low expression. This plot is meant to provide a high level exploratory view of the data.   
                                      
                                       
                                  
                                
                              
                            
                            
                              
                                
                                  
                                       
                                      
                                           Principal Components of Nephrin interactions Data   More Plot Information                      Principal Components of Nephrin interactions Data  Principal component analysis maps high-dimensional datasets onto a smaller number of highly informative dimensions. Here, the first four principal components of the gene expression data are plotted against each other and colored by the values of the selected covariate. This plot may be used to identify clusters in the data and to identify variables associated with prominent signal in the data. Variables that are associated with these leading principal components should be considered in downstream analyses.   
                                           Outliers   More Plot Information     
                                           
                                             
                                           
                                           
                                             
                                               
                                               Outlier on PC 1 
                                               Outlier on PC 2 
                                               Outlier on PC 3 
                                               Outlier on PC 4 
                                             
                                           
                                           
                                             
                                               20161013_sebelius2-101316_01_01.RCC 
                                               FALSE 
                                               FALSE 
                                               FALSE 
                                               TRUE 
                                             
                                           
                                           Outliers  Table identifying outliers in the first four PCs of the data.   
                                      
                                       
                                  
                                
                              
                            
                           
                            
                          
                           
                      
                    
                    
                      
                           
                          
                               Heatmaps    PCA     
                              
                                  Heatmaps  
                              
                               
                              
                                  New.Annotation  
                              
                              
                            
                              
                                
                                  
                                       
                                      
                                           Heatmap of Non-integrin membrane-ECM interactions Data   More Plot Information                      Heatmap of Non-integrin membrane-ECM interactions Data  Heatmap of the normalized data, scaled to give all genes equal variance, generated via unsupervised clustering. Orange indicates high expression; blue indicates low expression. This plot is meant to provide a high level exploratory view of the data.   
                                      
                                       
                                  
                                
                              
                            
                            
                              
                                
                                  
                                       
                                      
                                           Principal Components of Non-integrin membrane-ECM interactions Data   More Plot Information                      Principal Components of Non-integrin membrane-ECM interactions Data  Principal component analysis maps high-dimensional datasets onto a smaller number of highly informative dimensions. Here, the first four principal components of the gene expression data are plotted against each other and colored by the values of the selected covariate. This plot may be used to identify clusters in the data and to identify variables associated with prominent signal in the data. Variables that are associated with these leading principal components should be considered in downstream analyses.   
                                           Outliers   More Plot Information     
                                           
                                             
                                           
                                           
                                             
                                               
                                               Outlier on PC 1 
                                               Outlier on PC 2 
                                               Outlier on PC 3 
                                               Outlier on PC 4 
                                             
                                           
                                           
                                             
                                               20161013_sebelius2-101316_01_01.RCC 
                                               FALSE 
                                               FALSE 
                                               FALSE 
                                               TRUE 
                                             
                                           
                                           Outliers  Table identifying outliers in the first four PCs of the data.   
                                      
                                       
                                  
                                
                              
                            
                           
                            
                          
                           
                      
                    
                    
                      
                           
                          
                               Heatmaps    PCA     
                              
                                  Heatmaps  
                              
                               
                              
                                  New.Annotation  
                              
                              
                            
                              
                                
                                  
                                       
                                      
                                           Heatmap of Nonsense Mediated Decay (NMD) enhanced by the Exon Junction Complex (EJC) Data   More Plot Information                      Heatmap of Nonsense Mediated Decay (NMD) enhanced by the Exon Junction Complex (EJC) Data  Heatmap of the normalized data, scaled to give all genes equal variance, generated via unsupervised clustering. Orange indicates high expression; blue indicates low expression. This plot is meant to provide a high level exploratory view of the data.   
                                      
                                       
                                  
                                
                              
                            
                            
                              
                                
                                  
                                       
                                      
                                           Principal Components of Nonsense Mediated Decay (NMD) enhanced by the Exon Junction Complex (EJC) Data   More Plot Information                      Principal Components of Nonsense Mediated Decay (NMD) enhanced by the Exon Junction Complex (EJC) Data  Principal component analysis maps high-dimensional datasets onto a smaller number of highly informative dimensions. Here, the first four principal components of the gene expression data are plotted against each other and colored by the values of the selected covariate. This plot may be used to identify clusters in the data and to identify variables associated with prominent signal in the data. Variables that are associated with these leading principal components should be considered in downstream analyses.   
                                           Outliers   More Plot Information     
                                           
                                             
                                           
                                           
                                             
                                               
                                               Outlier on PC 1 
                                               Outlier on PC 2 
                                               Outlier on PC 3 
                                               Outlier on PC 4 
                                             
                                           
                                           
                                             
                                               20161013_sebelius2-101316_01_01.RCC 
                                               FALSE 
                                               FALSE 
                                               FALSE 
                                               TRUE 
                                             
                                           
                                           Outliers  Table identifying outliers in the first four PCs of the data.   
                                      
                                       
                                  
                                
                              
                            
                           
                            
                          
                           
                      
                    
                    
                      
                           
                          
                               Heatmaps    PCA     
                              
                                  Heatmaps  
                              
                               
                              
                                  New.Annotation  
                              
                              
                            
                              
                                
                                  
                                       
                                      
                                           Heatmap of NRAGE signals death through JNK Data   More Plot Information                      Heatmap of NRAGE signals death through JNK Data  Heatmap of the normalized data, scaled to give all genes equal variance, generated via unsupervised clustering. Orange indicates high expression; blue indicates low expression. This plot is meant to provide a high level exploratory view of the data.   
                                      
                                       
                                  
                                
                              
                            
                            
                              
                                
                                  
                                       
                                      
                                           Principal Components of NRAGE signals death through JNK Data   More Plot Information                      Principal Components of NRAGE signals death through JNK Data  Principal component analysis maps high-dimensional datasets onto a smaller number of highly informative dimensions. Here, the first four principal components of the gene expression data are plotted against each other and colored by the values of the selected covariate. This plot may be used to identify clusters in the data and to identify variables associated with prominent signal in the data. Variables that are associated with these leading principal components should be considered in downstream analyses.   
                                           Outliers   More Plot Information     
                                           
                                             
                                           
                                           
                                             
                                               
                                               Outlier on PC 1 
                                               Outlier on PC 2 
                                               Outlier on PC 3 
                                               Outlier on PC 4 
                                             
                                           
                                           
                                             
                                               20161013_sebelius2-101316_01_01.RCC 
                                               FALSE 
                                               FALSE 
                                               FALSE 
                                               TRUE 
                                             
                                           
                                           Outliers  Table identifying outliers in the first four PCs of the data.   
                                      
                                       
                                  
                                
                              
                            
                           
                            
                          
                           
                      
                    
                    
                      
                           
                          
                               Heatmaps    PCA     
                              
                                  Heatmaps  
                              
                               
                              
                                  New.Annotation  
                              
                              
                            
                              
                                
                                  
                                       
                                      
                                           Heatmap of p130Cas linkage to MAPK signaling for integrins Data   More Plot Information                      Heatmap of p130Cas linkage to MAPK signaling for integrins Data  Heatmap of the normalized data, scaled to give all genes equal variance, generated via unsupervised clustering. Orange indicates high expression; blue indicates low expression. This plot is meant to provide a high level exploratory view of the data.   
                                      
                                       
                                  
                                
                              
                            
                            
                              
                                
                                  
                                       
                                      
                                           Principal Components of p130Cas linkage to MAPK signaling for integrins Data   More Plot Information                      Principal Components of p130Cas linkage to MAPK signaling for integrins Data  Principal component analysis maps high-dimensional datasets onto a smaller number of highly informative dimensions. Here, the first four principal components of the gene expression data are plotted against each other and colored by the values of the selected covariate. This plot may be used to identify clusters in the data and to identify variables associated with prominent signal in the data. Variables that are associated with these leading principal components should be considered in downstream analyses.   
                                           Outliers   More Plot Information     
                                           
                                             
                                           
                                           
                                             
                                               
                                               Outlier on PC 1 
                                               Outlier on PC 2 
                                               Outlier on PC 3 
                                               Outlier on PC 4 
                                             
                                           
                                           
                                             
                                               20161013_sebelius2-101316_01_01.RCC 
                                               FALSE 
                                               FALSE 
                                               FALSE 
                                               TRUE 
                                             
                                           
                                           Outliers  Table identifying outliers in the first four PCs of the data.   
                                      
                                       
                                  
                                
                              
                            
                           
                            
                          
                           
                      
                    
                    
                      
                           
                          
                               Heatmaps    PCA     
                              
                                  Heatmaps  
                              
                               
                              
                                  New.Annotation  
                              
                              
                            
                              
                                
                                  
                                       
                                      
                                           Heatmap of PCPCE pathway Data   More Plot Information                      Heatmap of PCPCE pathway Data  Heatmap of the normalized data, scaled to give all genes equal variance, generated via unsupervised clustering. Orange indicates high expression; blue indicates low expression. This plot is meant to provide a high level exploratory view of the data.   
                                      
                                       
                                  
                                
                              
                            
                            
                              
                                
                                  
                                       
                                      
                                           Principal Components of PCPCE pathway Data   More Plot Information                      Principal Components of PCPCE pathway Data  Principal component analysis maps high-dimensional datasets onto a smaller number of highly informative dimensions. Here, the first four principal components of the gene expression data are plotted against each other and colored by the values of the selected covariate. This plot may be used to identify clusters in the data and to identify variables associated with prominent signal in the data. Variables that are associated with these leading principal components should be considered in downstream analyses.   
                                           Outliers   More Plot Information     
                                           
                                             
                                           
                                           
                                             
                                               
                                               Outlier on PC 1 
                                               Outlier on PC 2 
                                               Outlier on PC 3 
                                               Outlier on PC 4 
                                             
                                           
                                           
                                             
                                               20161013_sebelius2-101316_01_01.RCC 
                                               FALSE 
                                               FALSE 
                                               FALSE 
                                               TRUE 
                                             
                                           
                                           Outliers  Table identifying outliers in the first four PCs of the data.   
                                      
                                       
                                  
                                
                              
                            
                           
                            
                          
                           
                      
                    
                    
                      
                           
                          
                               Heatmaps    PCA     
                              
                                  Heatmaps  
                              
                               
                              
                                  New.Annotation  
                              
                              
                            
                              
                                
                                  
                                       
                                      
                                           Heatmap of PI3K Cascade Data   More Plot Information                      Heatmap of PI3K Cascade Data  Heatmap of the normalized data, scaled to give all genes equal variance, generated via unsupervised clustering. Orange indicates high expression; blue indicates low expression. This plot is meant to provide a high level exploratory view of the data.   
                                      
                                       
                                  
                                
                              
                            
                            
                              
                                
                                  
                                       
                                      
                                           Principal Components of PI3K Cascade Data   More Plot Information                      Principal Components of PI3K Cascade Data  Principal component analysis maps high-dimensional datasets onto a smaller number of highly informative dimensions. Here, the first four principal components of the gene expression data are plotted against each other and colored by the values of the selected covariate. This plot may be used to identify clusters in the data and to identify variables associated with prominent signal in the data. Variables that are associated with these leading principal components should be considered in downstream analyses.   
                                           Outliers   More Plot Information     
                                           
                                             
                                           
                                           
                                             
                                               
                                               Outlier on PC 1 
                                               Outlier on PC 2 
                                               Outlier on PC 3 
                                               Outlier on PC 4 
                                             
                                           
                                           
                                             
                                               20161013_sebelius2-101316_01_01.RCC 
                                               FALSE 
                                               FALSE 
                                               FALSE 
                                               TRUE 
                                             
                                           
                                           Outliers  Table identifying outliers in the first four PCs of the data.   
                                      
                                       
                                  
                                
                              
                            
                           
                            
                          
                           
                      
                    
                    
                      
                           
                          
                               Heatmaps    PCA     
                              
                                  Heatmaps  
                              
                               
                              
                                  New.Annotation  
                              
                              
                            
                              
                                
                                  
                                       
                                      
                                           Heatmap of PI3KAKT activation Data   More Plot Information                      Heatmap of PI3KAKT activation Data  Heatmap of the normalized data, scaled to give all genes equal variance, generated via unsupervised clustering. Orange indicates high expression; blue indicates low expression. This plot is meant to provide a high level exploratory view of the data.   
                                      
                                       
                                  
                                
                              
                            
                            
                              
                                
                                  
                                       
                                      
                                           Principal Components of PI3KAKT activation Data   More Plot Information                      Principal Components of PI3KAKT activation Data  Principal component analysis maps high-dimensional datasets onto a smaller number of highly informative dimensions. Here, the first four principal components of the gene expression data are plotted against each other and colored by the values of the selected covariate. This plot may be used to identify clusters in the data and to identify variables associated with prominent signal in the data. Variables that are associated with these leading principal components should be considered in downstream analyses.   
                                           Outliers   More Plot Information     
                                           
                                             
                                           
                                           
                                             
                                               
                                               Outlier on PC 1 
                                               Outlier on PC 2 
                                               Outlier on PC 3 
                                               Outlier on PC 4 
                                             
                                           
                                           
                                             
                                               20161013_sebelius2-101316_01_01.RCC 
                                               FALSE 
                                               FALSE 
                                               FALSE 
                                               TRUE 
                                             
                                           
                                           Outliers  Table identifying outliers in the first four PCs of the data.   
                                      
                                       
                                  
                                
                              
                            
                           
                            
                          
                           
                      
                    
                    
                      
                           
                          
                               Heatmaps    PCA     
                              
                                  Heatmaps  
                              
                               
                              
                                  New.Annotation  
                              
                              
                            
                              
                                
                                  
                                       
                                      
                                           Heatmap of PIP3 activates AKT signaling Data   More Plot Information                      Heatmap of PIP3 activates AKT signaling Data  Heatmap of the normalized data, scaled to give all genes equal variance, generated via unsupervised clustering. Orange indicates high expression; blue indicates low expression. This plot is meant to provide a high level exploratory view of the data.   
                                      
                                       
                                  
                                
                              
                            
                            
                              
                                
                                  
                                       
                                      
                                           Principal Components of PIP3 activates AKT signaling Data   More Plot Information                      Principal Components of PIP3 activates AKT signaling Data  Principal component analysis maps high-dimensional datasets onto a smaller number of highly informative dimensions. Here, the first four principal components of the gene expression data are plotted against each other and colored by the values of the selected covariate. This plot may be used to identify clusters in the data and to identify variables associated with prominent signal in the data. Variables that are associated with these leading principal components should be considered in downstream analyses.   
                                           Outliers   More Plot Information     
                                           
                                             
                                           
                                           
                                             
                                               
                                               Outlier on PC 1 
                                               Outlier on PC 2 
                                               Outlier on PC 3 
                                               Outlier on PC 4 
                                             
                                           
                                           
                                             
                                               20161013_sebelius2-101316_01_01.RCC 
                                               FALSE 
                                               FALSE 
                                               FALSE 
                                               TRUE 
                                             
                                           
                                           Outliers  Table identifying outliers in the first four PCs of the data.   
                                      
                                       
                                  
                                
                              
                            
                           
                            
                          
                           
                      
                    
                    
                      
                           
                          
                               Heatmaps    PCA     
                              
                                  Heatmaps  
                              
                               
                              
                                  New.Annotation  
                              
                              
                            
                              
                                
                                  
                                       
                                      
                                           Heatmap of Platelet degranulation  Data   More Plot Information                      Heatmap of Platelet degranulation  Data  Heatmap of the normalized data, scaled to give all genes equal variance, generated via unsupervised clustering. Orange indicates high expression; blue indicates low expression. This plot is meant to provide a high level exploratory view of the data.   
                                      
                                       
                                  
                                
                              
                            
                            
                              
                                
                                  
                                       
                                      
                                           Principal Components of Platelet degranulation  Data   More Plot Information                      Principal Components of Platelet degranulation  Data  Principal component analysis maps high-dimensional datasets onto a smaller number of highly informative dimensions. Here, the first four principal components of the gene expression data are plotted against each other and colored by the values of the selected covariate. This plot may be used to identify clusters in the data and to identify variables associated with prominent signal in the data. Variables that are associated with these leading principal components should be considered in downstream analyses.   
                                           Outliers   More Plot Information     
                                           
                                             
                                           
                                           
                                             
                                               
                                               Outlier on PC 1 
                                               Outlier on PC 2 
                                               Outlier on PC 3 
                                               Outlier on PC 4 
                                             
                                           
                                           
                                             
                                               20161013_sebelius2-101316_01_01.RCC 
                                               FALSE 
                                               FALSE 
                                               FALSE 
                                               TRUE 
                                             
                                           
                                           Outliers  Table identifying outliers in the first four PCs of the data.   
                                      
                                       
                                  
                                
                              
                            
                           
                            
                          
                           
                      
                    
                    
                      
                           
                          
                               Heatmaps    PCA     
                              
                                  Heatmaps  
                              
                               
                              
                                  New.Annotation  
                              
                              
                            
                              
                                
                                  
                                       
                                      
                                           Heatmap of RAF activation Data   More Plot Information                      Heatmap of RAF activation Data  Heatmap of the normalized data, scaled to give all genes equal variance, generated via unsupervised clustering. Orange indicates high expression; blue indicates low expression. This plot is meant to provide a high level exploratory view of the data.   
                                      
                                       
                                  
                                
                              
                            
                            
                              
                                
                                  
                                       
                                      
                                           Principal Components of RAF activation Data   More Plot Information                      Principal Components of RAF activation Data  Principal component analysis maps high-dimensional datasets onto a smaller number of highly informative dimensions. Here, the first four principal components of the gene expression data are plotted against each other and colored by the values of the selected covariate. This plot may be used to identify clusters in the data and to identify variables associated with prominent signal in the data. Variables that are associated with these leading principal components should be considered in downstream analyses.   
                                           Outliers   More Plot Information     
                                           
                                             
                                           
                                           
                                             
                                               
                                               Outlier on PC 1 
                                               Outlier on PC 2 
                                               Outlier on PC 3 
                                               Outlier on PC 4 
                                             
                                           
                                           
                                             
                                               20161013_sebelius2-101316_01_01.RCC 
                                               FALSE 
                                               FALSE 
                                               FALSE 
                                               TRUE 
                                             
                                           
                                           Outliers  Table identifying outliers in the first four PCs of the data.   
                                      
                                       
                                  
                                
                              
                            
                           
                            
                          
                           
                      
                    
                    
                      
                           
                          
                               Heatmaps    PCA     
                              
                                  Heatmaps  
                              
                               
                              
                                  New.Annotation  
                              
                              
                            
                              
                                
                                  
                                       
                                      
                                           Heatmap of RAFMAP kinase cascade Data   More Plot Information                      Heatmap of RAFMAP kinase cascade Data  Heatmap of the normalized data, scaled to give all genes equal variance, generated via unsupervised clustering. Orange indicates high expression; blue indicates low expression. This plot is meant to provide a high level exploratory view of the data.   
                                      
                                       
                                  
                                
                              
                            
                            
                              
                                
                                  
                                       
                                      
                                           Principal Components of RAFMAP kinase cascade Data   More Plot Information                      Principal Components of RAFMAP kinase cascade Data  Principal component analysis maps high-dimensional datasets onto a smaller number of highly informative dimensions. Here, the first four principal components of the gene expression data are plotted against each other and colored by the values of the selected covariate. This plot may be used to identify clusters in the data and to identify variables associated with prominent signal in the data. Variables that are associated with these leading principal components should be considered in downstream analyses.   
                                           Outliers   More Plot Information     
                                           
                                             
                                           
                                           
                                             
                                               
                                               Outlier on PC 1 
                                               Outlier on PC 2 
                                               Outlier on PC 3 
                                               Outlier on PC 4 
                                             
                                           
                                           
                                             
                                               20161013_sebelius2-101316_01_01.RCC 
                                               FALSE 
                                               FALSE 
                                               FALSE 
                                               TRUE 
                                             
                                           
                                           Outliers  Table identifying outliers in the first four PCs of the data.   
                                      
                                       
                                  
                                
                              
                            
                           
                            
                          
                           
                      
                    
                    
                      
                           
                          
                               Heatmaps    PCA     
                              
                                  Heatmaps  
                              
                               
                              
                                  New.Annotation  
                              
                              
                            
                              
                                
                                  
                                       
                                      
                                           Heatmap of Regulation of actin dynamics for phagocytic cup formation Data   More Plot Information                      Heatmap of Regulation of actin dynamics for phagocytic cup formation Data  Heatmap of the normalized data, scaled to give all genes equal variance, generated via unsupervised clustering. Orange indicates high expression; blue indicates low expression. This plot is meant to provide a high level exploratory view of the data.   
                                      
                                       
                                  
                                
                              
                            
                            
                              
                                
                                  
                                       
                                      
                                           Principal Components of Regulation of actin dynamics for phagocytic cup formation Data   More Plot Information                      Principal Components of Regulation of actin dynamics for phagocytic cup formation Data  Principal component analysis maps high-dimensional datasets onto a smaller number of highly informative dimensions. Here, the first four principal components of the gene expression data are plotted against each other and colored by the values of the selected covariate. This plot may be used to identify clusters in the data and to identify variables associated with prominent signal in the data. Variables that are associated with these leading principal components should be considered in downstream analyses.   
                                           Outliers   More Plot Information     
                                           
                                             
                                           
                                           
                                             
                                               
                                               Outlier on PC 1 
                                               Outlier on PC 2 
                                               Outlier on PC 3 
                                               Outlier on PC 4 
                                             
                                           
                                           
                                             
                                               20161013_sebelius2-101316_01_01.RCC 
                                               FALSE 
                                               FALSE 
                                               FALSE 
                                               TRUE 
                                             
                                           
                                           Outliers  Table identifying outliers in the first four PCs of the data.   
                                      
                                       
                                  
                                
                              
                            
                           
                            
                          
                           
                      
                    
                    
                      
                           
                          
                               Heatmaps    PCA     
                              
                                  Heatmaps  
                              
                               
                              
                                  New.Annotation  
                              
                              
                            
                              
                                
                                  
                                       
                                      
                                           Heatmap of Regulation of HSF1-mediated heat shock response Data   More Plot Information                      Heatmap of Regulation of HSF1-mediated heat shock response Data  Heatmap of the normalized data, scaled to give all genes equal variance, generated via unsupervised clustering. Orange indicates high expression; blue indicates low expression. This plot is meant to provide a high level exploratory view of the data.   
                                      
                                       
                                  
                                
                              
                            
                            
                              
                                
                                  
                                       
                                      
                                           Principal Components of Regulation of HSF1-mediated heat shock response Data   More Plot Information                      Principal Components of Regulation of HSF1-mediated heat shock response Data  Principal component analysis maps high-dimensional datasets onto a smaller number of highly informative dimensions. Here, the first four principal components of the gene expression data are plotted against each other and colored by the values of the selected covariate. This plot may be used to identify clusters in the data and to identify variables associated with prominent signal in the data. Variables that are associated with these leading principal components should be considered in downstream analyses.   
                                           Outliers   More Plot Information     
                                           
                                             
                                           
                                           
                                             
                                               
                                               Outlier on PC 1 
                                               Outlier on PC 2 
                                               Outlier on PC 3 
                                               Outlier on PC 4 
                                             
                                           
                                           
                                             
                                               20161013_sebelius2-101316_01_01.RCC 
                                               FALSE 
                                               FALSE 
                                               FALSE 
                                               TRUE 
                                             
                                           
                                           Outliers  Table identifying outliers in the first four PCs of the data.   
                                      
                                       
                                  
                                
                              
                            
                           
                            
                          
                           
                      
                    
                    
                      
                           
                          
                               Heatmaps    PCA     
                              
                                  Heatmaps  
                              
                               
                              
                                  New.Annotation  
                              
                              
                            
                              
                                
                                  
                                       
                                      
                                           Heatmap of Regulation of signaling by CBL Data   More Plot Information                      Heatmap of Regulation of signaling by CBL Data  Heatmap of the normalized data, scaled to give all genes equal variance, generated via unsupervised clustering. Orange indicates high expression; blue indicates low expression. This plot is meant to provide a high level exploratory view of the data.   
                                      
                                       
                                  
                                
                              
                            
                            
                              
                                
                                  
                                       
                                      
                                           Principal Components of Regulation of signaling by CBL Data   More Plot Information                      Principal Components of Regulation of signaling by CBL Data  Principal component analysis maps high-dimensional datasets onto a smaller number of highly informative dimensions. Here, the first four principal components of the gene expression data are plotted against each other and colored by the values of the selected covariate. This plot may be used to identify clusters in the data and to identify variables associated with prominent signal in the data. Variables that are associated with these leading principal components should be considered in downstream analyses.   
                                           Outliers   More Plot Information     
                                           
                                             
                                           
                                           
                                             
                                               
                                               Outlier on PC 1 
                                               Outlier on PC 2 
                                               Outlier on PC 3 
                                               Outlier on PC 4 
                                             
                                           
                                           
                                             
                                               20161013_sebelius2-101316_01_01.RCC 
                                               FALSE 
                                               FALSE 
                                               FALSE 
                                               TRUE 
                                             
                                           
                                           Outliers  Table identifying outliers in the first four PCs of the data.   
                                      
                                       
                                  
                                
                              
                            
                           
                            
                          
                           
                      
                    
                    
                      
                           
                          
                               Heatmaps    PCA     
                              
                                  Heatmaps  
                              
                               
                              
                                  New.Annotation  
                              
                              
                            
                              
                                
                                  
                                       
                                      
                                           Heatmap of Repression of WNT target genes Data   More Plot Information                      Heatmap of Repression of WNT target genes Data  Heatmap of the normalized data, scaled to give all genes equal variance, generated via unsupervised clustering. Orange indicates high expression; blue indicates low expression. This plot is meant to provide a high level exploratory view of the data.   
                                      
                                       
                                  
                                
                              
                            
                            
                              
                                
                                  
                                       
                                      
                                           Principal Components of Repression of WNT target genes Data   More Plot Information                      Principal Components of Repression of WNT target genes Data  Principal component analysis maps high-dimensional datasets onto a smaller number of highly informative dimensions. Here, the first four principal components of the gene expression data are plotted against each other and colored by the values of the selected covariate. This plot may be used to identify clusters in the data and to identify variables associated with prominent signal in the data. Variables that are associated with these leading principal components should be considered in downstream analyses.   
                                           Outliers   More Plot Information     
                                           
                                             
                                           
                                           
                                             
                                               
                                               Outlier on PC 1 
                                               Outlier on PC 2 
                                               Outlier on PC 3 
                                               Outlier on PC 4 
                                             
                                           
                                           
                                             
                                               20161013_sebelius2-101316_01_01.RCC 
                                               FALSE 
                                               FALSE 
                                               FALSE 
                                               TRUE 
                                             
                                           
                                           Outliers  Table identifying outliers in the first four PCs of the data.   
                                      
                                       
                                  
                                
                              
                            
                           
                            
                          
                           
                      
                    
                    
                      
                           
                          
                               Heatmaps    PCA     
                              
                                  Heatmaps  
                              
                               
                              
                                  New.Annotation  
                              
                              
                            
                              
                                
                                  
                                       
                                      
                                           Heatmap of Resolution of Sister Chromatid Cohesion Data   More Plot Information                      Heatmap of Resolution of Sister Chromatid Cohesion Data  Heatmap of the normalized data, scaled to give all genes equal variance, generated via unsupervised clustering. Orange indicates high expression; blue indicates low expression. This plot is meant to provide a high level exploratory view of the data.   
                                      
                                       
                                  
                                
                              
                            
                            
                              
                                
                                  
                                       
                                      
                                           Principal Components of Resolution of Sister Chromatid Cohesion Data   More Plot Information                      Principal Components of Resolution of Sister Chromatid Cohesion Data  Principal component analysis maps high-dimensional datasets onto a smaller number of highly informative dimensions. Here, the first four principal components of the gene expression data are plotted against each other and colored by the values of the selected covariate. This plot may be used to identify clusters in the data and to identify variables associated with prominent signal in the data. Variables that are associated with these leading principal components should be considered in downstream analyses.   
                                           Outliers   More Plot Information     
                                           
                                             
                                           
                                           
                                             
                                               
                                               Outlier on PC 1 
                                               Outlier on PC 2 
                                               Outlier on PC 3 
                                               Outlier on PC 4 
                                             
                                           
                                           
                                             
                                               20161013_sebelius2-101316_01_01.RCC 
                                               FALSE 
                                               FALSE 
                                               FALSE 
                                               TRUE 
                                             
                                           
                                           Outliers  Table identifying outliers in the first four PCs of the data.   
                                      
                                       
                                  
                                
                              
                            
                           
                            
                          
                           
                      
                    
                    
                      
                           
                          
                               Heatmaps    PCA     
                              
                                  Heatmaps  
                              
                               
                              
                                  New.Annotation  
                              
                              
                            
                              
                                
                                  
                                       
                                      
                                           Heatmap of Rho GTPase cycle Data   More Plot Information                      Heatmap of Rho GTPase cycle Data  Heatmap of the normalized data, scaled to give all genes equal variance, generated via unsupervised clustering. Orange indicates high expression; blue indicates low expression. This plot is meant to provide a high level exploratory view of the data.   
                                      
                                       
                                  
                                
                              
                            
                            
                              
                                
                                  
                                       
                                      
                                           Principal Components of Rho GTPase cycle Data   More Plot Information                      Principal Components of Rho GTPase cycle Data  Principal component analysis maps high-dimensional datasets onto a smaller number of highly informative dimensions. Here, the first four principal components of the gene expression data are plotted against each other and colored by the values of the selected covariate. This plot may be used to identify clusters in the data and to identify variables associated with prominent signal in the data. Variables that are associated with these leading principal components should be considered in downstream analyses.   
                                           Outliers   More Plot Information     
                                           
                                             
                                           
                                           
                                             
                                               
                                               Outlier on PC 1 
                                               Outlier on PC 2 
                                               Outlier on PC 3 
                                               Outlier on PC 4 
                                             
                                           
                                           
                                             
                                               20161013_sebelius2-101316_01_01.RCC 
                                               FALSE 
                                               FALSE 
                                               FALSE 
                                               TRUE 
                                             
                                           
                                           Outliers  Table identifying outliers in the first four PCs of the data.   
                                      
                                       
                                  
                                
                              
                            
                           
                            
                          
                           
                      
                    
                    
                      
                           
                          
                               Heatmaps    PCA     
                              
                                  Heatmaps  
                              
                               
                              
                                  New.Annotation  
                              
                              
                            
                              
                                
                                  
                                       
                                      
                                           Heatmap of RHO GTPases activate CIT Data   More Plot Information                      Heatmap of RHO GTPases activate CIT Data  Heatmap of the normalized data, scaled to give all genes equal variance, generated via unsupervised clustering. Orange indicates high expression; blue indicates low expression. This plot is meant to provide a high level exploratory view of the data.   
                                      
                                       
                                  
                                
                              
                            
                            
                              
                                
                                  
                                       
                                      
                                           Principal Components of RHO GTPases activate CIT Data   More Plot Information                      Principal Components of RHO GTPases activate CIT Data  Principal component analysis maps high-dimensional datasets onto a smaller number of highly informative dimensions. Here, the first four principal components of the gene expression data are plotted against each other and colored by the values of the selected covariate. This plot may be used to identify clusters in the data and to identify variables associated with prominent signal in the data. Variables that are associated with these leading principal components should be considered in downstream analyses.   
                                           Outliers   More Plot Information     
                                           
                                             
                                           
                                           
                                             
                                               
                                               Outlier on PC 1 
                                               Outlier on PC 2 
                                               Outlier on PC 3 
                                               Outlier on PC 4 
                                             
                                           
                                           
                                             
                                               20161013_sebelius2-101316_01_01.RCC 
                                               FALSE 
                                               FALSE 
                                               FALSE 
                                               TRUE 
                                             
                                           
                                           Outliers  Table identifying outliers in the first four PCs of the data.   
                                      
                                       
                                  
                                
                              
                            
                           
                            
                          
                           
                      
                    
                    
                      
                           
                          
                               Heatmaps    PCA     
                              
                                  Heatmaps  
                              
                               
                              
                                  New.Annotation  
                              
                              
                            
                              
                                
                                  
                                       
                                      
                                           Heatmap of RHO GTPases Activate Formins Data   More Plot Information                      Heatmap of RHO GTPases Activate Formins Data  Heatmap of the normalized data, scaled to give all genes equal variance, generated via unsupervised clustering. Orange indicates high expression; blue indicates low expression. This plot is meant to provide a high level exploratory view of the data.   
                                      
                                       
                                  
                                
                              
                            
                            
                              
                                
                                  
                                       
                                      
                                           Principal Components of RHO GTPases Activate Formins Data   More Plot Information                      Principal Components of RHO GTPases Activate Formins Data  Principal component analysis maps high-dimensional datasets onto a smaller number of highly informative dimensions. Here, the first four principal components of the gene expression data are plotted against each other and colored by the values of the selected covariate. This plot may be used to identify clusters in the data and to identify variables associated with prominent signal in the data. Variables that are associated with these leading principal components should be considered in downstream analyses.   
                                           Outliers   More Plot Information     
                                           
                                             
                                           
                                           
                                             
                                               
                                               Outlier on PC 1 
                                               Outlier on PC 2 
                                               Outlier on PC 3 
                                               Outlier on PC 4 
                                             
                                           
                                           
                                             
                                               20161013_sebelius2-101316_01_01.RCC 
                                               FALSE 
                                               FALSE 
                                               FALSE 
                                               TRUE 
                                             
                                           
                                           Outliers  Table identifying outliers in the first four PCs of the data.   
                                      
                                       
                                  
                                
                              
                            
                           
                            
                          
                           
                      
                    
                    
                      
                           
                          
                               Heatmaps    PCA     
                              
                                  Heatmaps  
                              
                               
                              
                                  New.Annotation  
                              
                              
                            
                              
                                
                                  
                                       
                                      
                                           Heatmap of RHO GTPases activate IQGAPs Data   More Plot Information                      Heatmap of RHO GTPases activate IQGAPs Data  Heatmap of the normalized data, scaled to give all genes equal variance, generated via unsupervised clustering. Orange indicates high expression; blue indicates low expression. This plot is meant to provide a high level exploratory view of the data.   
                                      
                                       
                                  
                                
                              
                            
                            
                              
                                
                                  
                                       
                                      
                                           Principal Components of RHO GTPases activate IQGAPs Data   More Plot Information                      Principal Components of RHO GTPases activate IQGAPs Data  Principal component analysis maps high-dimensional datasets onto a smaller number of highly informative dimensions. Here, the first four principal components of the gene expression data are plotted against each other and colored by the values of the selected covariate. This plot may be used to identify clusters in the data and to identify variables associated with prominent signal in the data. Variables that are associated with these leading principal components should be considered in downstream analyses.   
                                           Outliers   More Plot Information     
                                           
                                             
                                           
                                           
                                             
                                               
                                               Outlier on PC 1 
                                               Outlier on PC 2 
                                               Outlier on PC 3 
                                               Outlier on PC 4 
                                             
                                           
                                           
                                             
                                               20161013_sebelius2-101316_01_01.RCC 
                                               FALSE 
                                               FALSE 
                                               FALSE 
                                               TRUE 
                                             
                                           
                                           Outliers  Table identifying outliers in the first four PCs of the data.   
                                      
                                       
                                  
                                
                              
                            
                           
                            
                          
                           
                      
                    
                    
                      
                           
                          
                               Heatmaps    PCA     
                              
                                  Heatmaps  
                              
                               
                              
                                  New.Annotation  
                              
                              
                            
                              
                                
                                  
                                       
                                      
                                           Heatmap of RHO GTPases activate PAKs Data   More Plot Information                      Heatmap of RHO GTPases activate PAKs Data  Heatmap of the normalized data, scaled to give all genes equal variance, generated via unsupervised clustering. Orange indicates high expression; blue indicates low expression. This plot is meant to provide a high level exploratory view of the data.   
                                      
                                       
                                  
                                
                              
                            
                            
                              
                                
                                  
                                       
                                      
                                           Principal Components of RHO GTPases activate PAKs Data   More Plot Information                      Principal Components of RHO GTPases activate PAKs Data  Principal component analysis maps high-dimensional datasets onto a smaller number of highly informative dimensions. Here, the first four principal components of the gene expression data are plotted against each other and colored by the values of the selected covariate. This plot may be used to identify clusters in the data and to identify variables associated with prominent signal in the data. Variables that are associated with these leading principal components should be considered in downstream analyses.   
                                           Outliers   More Plot Information     
                                           
                                             
                                           
                                           
                                             
                                               
                                               Outlier on PC 1 
                                               Outlier on PC 2 
                                               Outlier on PC 3 
                                               Outlier on PC 4 
                                             
                                           
                                           
                                             
                                               20161013_sebelius2-101316_01_01.RCC 
                                               FALSE 
                                               FALSE 
                                               FALSE 
                                               TRUE 
                                             
                                           
                                           Outliers  Table identifying outliers in the first four PCs of the data.   
                                      
                                       
                                  
                                
                              
                            
                           
                            
                          
                           
                      
                    
                    
                      
                           
                          
                               Heatmaps    PCA     
                              
                                  Heatmaps  
                              
                               
                              
                                  New.Annotation  
                              
                              
                            
                              
                                
                                  
                                       
                                      
                                           Heatmap of RHO GTPases activate PKNs Data   More Plot Information                      Heatmap of RHO GTPases activate PKNs Data  Heatmap of the normalized data, scaled to give all genes equal variance, generated via unsupervised clustering. Orange indicates high expression; blue indicates low expression. This plot is meant to provide a high level exploratory view of the data.   
                                      
                                       
                                  
                                
                              
                            
                            
                              
                                
                                  
                                       
                                      
                                           Principal Components of RHO GTPases activate PKNs Data   More Plot Information                      Principal Components of RHO GTPases activate PKNs Data  Principal component analysis maps high-dimensional datasets onto a smaller number of highly informative dimensions. Here, the first four principal components of the gene expression data are plotted against each other and colored by the values of the selected covariate. This plot may be used to identify clusters in the data and to identify variables associated with prominent signal in the data. Variables that are associated with these leading principal components should be considered in downstream analyses.   
                                           Outliers   More Plot Information     
                                           
                                             
                                           
                                           
                                             
                                               
                                               Outlier on PC 1 
                                               Outlier on PC 2 
                                               Outlier on PC 3 
                                               Outlier on PC 4 
                                             
                                           
                                           
                                             
                                               20161013_sebelius2-101316_01_01.RCC 
                                               FALSE 
                                               FALSE 
                                               FALSE 
                                               TRUE 
                                             
                                           
                                           Outliers  Table identifying outliers in the first four PCs of the data.   
                                      
                                       
                                  
                                
                              
                            
                           
                            
                          
                           
                      
                    
                    
                      
                           
                          
                               Heatmaps    PCA     
                              
                                  Heatmaps  
                              
                               
                              
                                  New.Annotation  
                              
                              
                            
                              
                                
                                  
                                       
                                      
                                           Heatmap of RHO GTPases Activate ROCKs Data   More Plot Information                      Heatmap of RHO GTPases Activate ROCKs Data  Heatmap of the normalized data, scaled to give all genes equal variance, generated via unsupervised clustering. Orange indicates high expression; blue indicates low expression. This plot is meant to provide a high level exploratory view of the data.   
                                      
                                       
                                  
                                
                              
                            
                            
                              
                                
                                  
                                       
                                      
                                           Principal Components of RHO GTPases Activate ROCKs Data   More Plot Information                      Principal Components of RHO GTPases Activate ROCKs Data  Principal component analysis maps high-dimensional datasets onto a smaller number of highly informative dimensions. Here, the first four principal components of the gene expression data are plotted against each other and colored by the values of the selected covariate. This plot may be used to identify clusters in the data and to identify variables associated with prominent signal in the data. Variables that are associated with these leading principal components should be considered in downstream analyses.   
                                           Outliers   More Plot Information     
                                           
                                             
                                           
                                           
                                             
                                               
                                               Outlier on PC 1 
                                               Outlier on PC 2 
                                               Outlier on PC 3 
                                               Outlier on PC 4 
                                             
                                           
                                           
                                             
                                               20161013_sebelius2-101316_01_01.RCC 
                                               FALSE 
                                               FALSE 
                                               FALSE 
                                               TRUE 
                                             
                                           
                                           Outliers  Table identifying outliers in the first four PCs of the data.   
                                      
                                       
                                  
                                
                              
                            
                           
                            
                          
                           
                      
                    
                    
                      
                           
                          
                               Heatmaps    PCA     
                              
                                  Heatmaps  
                              
                               
                              
                                  New.Annotation  
                              
                              
                            
                              
                                
                                  
                                       
                                      
                                           Heatmap of RHO GTPases Activate WASPs and WAVEs Data   More Plot Information                      Heatmap of RHO GTPases Activate WASPs and WAVEs Data  Heatmap of the normalized data, scaled to give all genes equal variance, generated via unsupervised clustering. Orange indicates high expression; blue indicates low expression. This plot is meant to provide a high level exploratory view of the data.   
                                      
                                       
                                  
                                
                              
                            
                            
                              
                                
                                  
                                       
                                      
                                           Principal Components of RHO GTPases Activate WASPs and WAVEs Data   More Plot Information                      Principal Components of RHO GTPases Activate WASPs and WAVEs Data  Principal component analysis maps high-dimensional datasets onto a smaller number of highly informative dimensions. Here, the first four principal components of the gene expression data are plotted against each other and colored by the values of the selected covariate. This plot may be used to identify clusters in the data and to identify variables associated with prominent signal in the data. Variables that are associated with these leading principal components should be considered in downstream analyses.   
                                           Outliers   More Plot Information     
                                           
                                             
                                           
                                           
                                             
                                               
                                               Outlier on PC 1 
                                               Outlier on PC 2 
                                               Outlier on PC 3 
                                               Outlier on PC 4 
                                             
                                           
                                           
                                             
                                               20161013_sebelius2-101316_01_01.RCC 
                                               FALSE 
                                               FALSE 
                                               FALSE 
                                               TRUE 
                                             
                                           
                                           Outliers  Table identifying outliers in the first four PCs of the data.   
                                      
                                       
                                  
                                
                              
                            
                           
                            
                          
                           
                      
                    
                    
                      
                           
                          
                               Heatmaps    PCA     
                              
                                  Heatmaps  
                              
                               
                              
                                  New.Annotation  
                              
                              
                            
                              
                                
                                  
                                       
                                      
                                           Heatmap of Role of LAT2NTALLAB on calcium mobilization Data   More Plot Information                      Heatmap of Role of LAT2NTALLAB on calcium mobilization Data  Heatmap of the normalized data, scaled to give all genes equal variance, generated via unsupervised clustering. Orange indicates high expression; blue indicates low expression. This plot is meant to provide a high level exploratory view of the data.   
                                      
                                       
                                  
                                
                              
                            
                            
                              
                                
                                  
                                       
                                      
                                           Principal Components of Role of LAT2NTALLAB on calcium mobilization Data   More Plot Information                      Principal Components of Role of LAT2NTALLAB on calcium mobilization Data  Principal component analysis maps high-dimensional datasets onto a smaller number of highly informative dimensions. Here, the first four principal components of the gene expression data are plotted against each other and colored by the values of the selected covariate. This plot may be used to identify clusters in the data and to identify variables associated with prominent signal in the data. Variables that are associated with these leading principal components should be considered in downstream analyses.   
                                           Outliers   More Plot Information     
                                           
                                             
                                           
                                           
                                             
                                               
                                               Outlier on PC 1 
                                               Outlier on PC 2 
                                               Outlier on PC 3 
                                               Outlier on PC 4 
                                             
                                           
                                           
                                             
                                               20161013_sebelius2-101316_01_01.RCC 
                                               FALSE 
                                               FALSE 
                                               FALSE 
                                               TRUE 
                                             
                                           
                                           Outliers  Table identifying outliers in the first four PCs of the data.   
                                      
                                       
                                  
                                
                              
                            
                           
                            
                          
                           
                      
                    
                    
                      
                           
                          
                               Heatmaps    PCA     
                              
                                  Heatmaps  
                              
                               
                              
                                  New.Annotation  
                              
                              
                            
                              
                                
                                  
                                       
                                      
                                           Heatmap of S33 mutants of beta-catenin aren.SQT.t phosphorylated Data   More Plot Information                      Heatmap of S33 mutants of beta-catenin aren.SQT.t phosphorylated Data  Heatmap of the normalized data, scaled to give all genes equal variance, generated via unsupervised clustering. Orange indicates high expression; blue indicates low expression. This plot is meant to provide a high level exploratory view of the data.   
                                      
                                       
                                  
                                
                              
                            
                            
                              
                                
                                  
                                       
                                      
                                           Principal Components of S33 mutants of beta-catenin aren.SQT.t phosphorylated Data   More Plot Information                      Principal Components of S33 mutants of beta-catenin aren.SQT.t phosphorylated Data  Principal component analysis maps high-dimensional datasets onto a smaller number of highly informative dimensions. Here, the first four principal components of the gene expression data are plotted against each other and colored by the values of the selected covariate. This plot may be used to identify clusters in the data and to identify variables associated with prominent signal in the data. Variables that are associated with these leading principal components should be considered in downstream analyses.   
                                           Outliers   More Plot Information     
                                           
                                             
                                           
                                           
                                             
                                               
                                               Outlier on PC 1 
                                               Outlier on PC 2 
                                               Outlier on PC 3 
                                               Outlier on PC 4 
                                             
                                           
                                           
                                             
                                               20161013_sebelius2-101316_01_01.RCC 
                                               FALSE 
                                               FALSE 
                                               FALSE 
                                               TRUE 
                                             
                                           
                                           Outliers  Table identifying outliers in the first four PCs of the data.   
                                      
                                       
                                  
                                
                              
                            
                           
                            
                          
                           
                      
                    
                    
                      
                           
                          
                               Heatmaps    PCA     
                              
                                  Heatmaps  
                              
                               
                              
                                  New.Annotation  
                              
                              
                            
                              
                                
                                  
                                       
                                      
                                           Heatmap of S37 mutants of beta-catenin aren.SQT.t phosphorylated Data   More Plot Information                      Heatmap of S37 mutants of beta-catenin aren.SQT.t phosphorylated Data  Heatmap of the normalized data, scaled to give all genes equal variance, generated via unsupervised clustering. Orange indicates high expression; blue indicates low expression. This plot is meant to provide a high level exploratory view of the data.   
                                      
                                       
                                  
                                
                              
                            
                            
                              
                                
                                  
                                       
                                      
                                           Principal Components of S37 mutants of beta-catenin aren.SQT.t phosphorylated Data   More Plot Information                      Principal Components of S37 mutants of beta-catenin aren.SQT.t phosphorylated Data  Principal component analysis maps high-dimensional datasets onto a smaller number of highly informative dimensions. Here, the first four principal components of the gene expression data are plotted against each other and colored by the values of the selected covariate. This plot may be used to identify clusters in the data and to identify variables associated with prominent signal in the data. Variables that are associated with these leading principal components should be considered in downstream analyses.   
                                           Outliers   More Plot Information     
                                           
                                             
                                           
                                           
                                             
                                               
                                               Outlier on PC 1 
                                               Outlier on PC 2 
                                               Outlier on PC 3 
                                               Outlier on PC 4 
                                             
                                           
                                           
                                             
                                               20161013_sebelius2-101316_01_01.RCC 
                                               FALSE 
                                               FALSE 
                                               FALSE 
                                               TRUE 
                                             
                                           
                                           Outliers  Table identifying outliers in the first four PCs of the data.   
                                      
                                       
                                  
                                
                              
                            
                           
                            
                          
                           
                      
                    
                    
                      
                           
                          
                               Heatmaps    PCA     
                              
                                  Heatmaps  
                              
                               
                              
                                  New.Annotation  
                              
                              
                            
                              
                                
                                  
                                       
                                      
                                           Heatmap of S45 mutants of beta-catenin aren.SQT.t phosphorylated Data   More Plot Information                      Heatmap of S45 mutants of beta-catenin aren.SQT.t phosphorylated Data  Heatmap of the normalized data, scaled to give all genes equal variance, generated via unsupervised clustering. Orange indicates high expression; blue indicates low expression. This plot is meant to provide a high level exploratory view of the data.   
                                      
                                       
                                  
                                
                              
                            
                            
                              
                                
                                  
                                       
                                      
                                           Principal Components of S45 mutants of beta-catenin aren.SQT.t phosphorylated Data   More Plot Information                      Principal Components of S45 mutants of beta-catenin aren.SQT.t phosphorylated Data  Principal component analysis maps high-dimensional datasets onto a smaller number of highly informative dimensions. Here, the first four principal components of the gene expression data are plotted against each other and colored by the values of the selected covariate. This plot may be used to identify clusters in the data and to identify variables associated with prominent signal in the data. Variables that are associated with these leading principal components should be considered in downstream analyses.   
                                           Outliers   More Plot Information     
                                           
                                             
                                           
                                           
                                             
                                               
                                               Outlier on PC 1 
                                               Outlier on PC 2 
                                               Outlier on PC 3 
                                               Outlier on PC 4 
                                             
                                           
                                           
                                             
                                               20161013_sebelius2-101316_01_01.RCC 
                                               FALSE 
                                               FALSE 
                                               FALSE 
                                               TRUE 
                                             
                                           
                                           Outliers  Table identifying outliers in the first four PCs of the data.   
                                      
                                       
                                  
                                
                              
                            
                           
                            
                          
                           
                      
                    
                    
                      
                           
                          
                               Heatmaps    PCA     
                              
                                  Heatmaps  
                              
                               
                              
                                  New.Annotation  
                              
                              
                            
                              
                                
                                  
                                       
                                      
                                           Heatmap of Sema3A PAK dependent Axon repulsion Data   More Plot Information                      Heatmap of Sema3A PAK dependent Axon repulsion Data  Heatmap of the normalized data, scaled to give all genes equal variance, generated via unsupervised clustering. Orange indicates high expression; blue indicates low expression. This plot is meant to provide a high level exploratory view of the data.   
                                      
                                       
                                  
                                
                              
                            
                            
                              
                                
                                  
                                       
                                      
                                           Principal Components of Sema3A PAK dependent Axon repulsion Data   More Plot Information                      Principal Components of Sema3A PAK dependent Axon repulsion Data  Principal component analysis maps high-dimensional datasets onto a smaller number of highly informative dimensions. Here, the first four principal components of the gene expression data are plotted against each other and colored by the values of the selected covariate. This plot may be used to identify clusters in the data and to identify variables associated with prominent signal in the data. Variables that are associated with these leading principal components should be considered in downstream analyses.   
                                           Outliers   More Plot Information     
                                           
                                             
                                           
                                           
                                             
                                               
                                               Outlier on PC 1 
                                               Outlier on PC 2 
                                               Outlier on PC 3 
                                               Outlier on PC 4 
                                             
                                           
                                           
                                             
                                               20161013_sebelius2-101316_01_01.RCC 
                                               FALSE 
                                               FALSE 
                                               FALSE 
                                               TRUE 
                                             
                                           
                                           Outliers  Table identifying outliers in the first four PCs of the data.   
                                      
                                       
                                  
                                
                              
                            
                           
                            
                          
                           
                      
                    
                    
                      
                           
                          
                               Heatmaps    PCA     
                              
                                  Heatmaps  
                              
                               
                              
                                  New.Annotation  
                              
                              
                            
                              
                                
                                  
                                       
                                      
                                           Heatmap of Sema4D induced cell migration and growth-cone collapse Data   More Plot Information                      Heatmap of Sema4D induced cell migration and growth-cone collapse Data  Heatmap of the normalized data, scaled to give all genes equal variance, generated via unsupervised clustering. Orange indicates high expression; blue indicates low expression. This plot is meant to provide a high level exploratory view of the data.   
                                      
                                       
                                  
                                
                              
                            
                            
                              
                                
                                  
                                       
                                      
                                           Principal Components of Sema4D induced cell migration and growth-cone collapse Data   More Plot Information                      Principal Components of Sema4D induced cell migration and growth-cone collapse Data  Principal component analysis maps high-dimensional datasets onto a smaller number of highly informative dimensions. Here, the first four principal components of the gene expression data are plotted against each other and colored by the values of the selected covariate. This plot may be used to identify clusters in the data and to identify variables associated with prominent signal in the data. Variables that are associated with these leading principal components should be considered in downstream analyses.   
                                           Outliers   More Plot Information     
                                           
                                             
                                           
                                           
                                             
                                               
                                               Outlier on PC 1 
                                               Outlier on PC 2 
                                               Outlier on PC 3 
                                               Outlier on PC 4 
                                             
                                           
                                           
                                             
                                               20161013_sebelius2-101316_01_01.RCC 
                                               FALSE 
                                               FALSE 
                                               FALSE 
                                               TRUE 
                                             
                                           
                                           Outliers  Table identifying outliers in the first four PCs of the data.   
                                      
                                       
                                  
                                
                              
                            
                           
                            
                          
                           
                      
                    
                    
                      
                           
                          
                               Heatmaps    PCA     
                              
                                  Heatmaps  
                              
                               
                              
                                  New.Annotation  
                              
                              
                            
                              
                                
                                  
                                       
                                      
                                           Heatmap of SHC-mediated cascade.CL.FGFR1 Data   More Plot Information                      Heatmap of SHC-mediated cascade.CL.FGFR1 Data  Heatmap of the normalized data, scaled to give all genes equal variance, generated via unsupervised clustering. Orange indicates high expression; blue indicates low expression. This plot is meant to provide a high level exploratory view of the data.   
                                      
                                       
                                  
                                
                              
                            
                            
                              
                                
                                  
                                       
                                      
                                           Principal Components of SHC-mediated cascade.CL.FGFR1 Data   More Plot Information                      Principal Components of SHC-mediated cascade.CL.FGFR1 Data  Principal component analysis maps high-dimensional datasets onto a smaller number of highly informative dimensions. Here, the first four principal components of the gene expression data are plotted against each other and colored by the values of the selected covariate. This plot may be used to identify clusters in the data and to identify variables associated with prominent signal in the data. Variables that are associated with these leading principal components should be considered in downstream analyses.   
                                           Outliers   More Plot Information     
                                           
                                             
                                           
                                           
                                             
                                               
                                               Outlier on PC 1 
                                               Outlier on PC 2 
                                               Outlier on PC 3 
                                               Outlier on PC 4 
                                             
                                           
                                           
                                             
                                               20161013_sebelius2-101316_01_01.RCC 
                                               FALSE 
                                               FALSE 
                                               FALSE 
                                               TRUE 
                                             
                                           
                                           Outliers  Table identifying outliers in the first four PCs of the data.   
                                      
                                       
                                  
                                
                              
                            
                           
                            
                          
                           
                      
                    
                    
                      
                           
                          
                               Heatmaps    PCA     
                              
                                  Heatmaps  
                              
                               
                              
                                  New.Annotation  
                              
                              
                            
                              
                                
                                  
                                       
                                      
                                           Heatmap of SHC-related events triggered by IGF1R Data   More Plot Information                      Heatmap of SHC-related events triggered by IGF1R Data  Heatmap of the normalized data, scaled to give all genes equal variance, generated via unsupervised clustering. Orange indicates high expression; blue indicates low expression. This plot is meant to provide a high level exploratory view of the data.   
                                      
                                       
                                  
                                
                              
                            
                            
                              
                                
                                  
                                       
                                      
                                           Principal Components of SHC-related events triggered by IGF1R Data   More Plot Information                      Principal Components of SHC-related events triggered by IGF1R Data  Principal component analysis maps high-dimensional datasets onto a smaller number of highly informative dimensions. Here, the first four principal components of the gene expression data are plotted against each other and colored by the values of the selected covariate. This plot may be used to identify clusters in the data and to identify variables associated with prominent signal in the data. Variables that are associated with these leading principal components should be considered in downstream analyses.   
                                           Outliers   More Plot Information     
                                           
                                             
                                           
                                           
                                             
                                               
                                               Outlier on PC 1 
                                               Outlier on PC 2 
                                               Outlier on PC 3 
                                               Outlier on PC 4 
                                             
                                           
                                           
                                             
                                               20161013_sebelius2-101316_01_01.RCC 
                                               FALSE 
                                               FALSE 
                                               FALSE 
                                               TRUE 
                                             
                                           
                                           Outliers  Table identifying outliers in the first four PCs of the data.   
                                      
                                       
                                  
                                
                              
                            
                           
                            
                          
                           
                      
                    
                    
                      
                           
                          
                               Heatmaps    PCA     
                              
                                  Heatmaps  
                              
                               
                              
                                  New.Annotation  
                              
                              
                            
                              
                                
                                  
                                       
                                      
                                           Heatmap of Signal transduction by L1 Data   More Plot Information                      Heatmap of Signal transduction by L1 Data  Heatmap of the normalized data, scaled to give all genes equal variance, generated via unsupervised clustering. Orange indicates high expression; blue indicates low expression. This plot is meant to provide a high level exploratory view of the data.   
                                      
                                       
                                  
                                
                              
                            
                            
                              
                                
                                  
                                       
                                      
                                           Principal Components of Signal transduction by L1 Data   More Plot Information                      Principal Components of Signal transduction by L1 Data  Principal component analysis maps high-dimensional datasets onto a smaller number of highly informative dimensions. Here, the first four principal components of the gene expression data are plotted against each other and colored by the values of the selected covariate. This plot may be used to identify clusters in the data and to identify variables associated with prominent signal in the data. Variables that are associated with these leading principal components should be considered in downstream analyses.   
                                           Outliers   More Plot Information     
                                           
                                             
                                           
                                           
                                             
                                               
                                               Outlier on PC 1 
                                               Outlier on PC 2 
                                               Outlier on PC 3 
                                               Outlier on PC 4 
                                             
                                           
                                           
                                             
                                               20161013_sebelius2-101316_01_01.RCC 
                                               FALSE 
                                               FALSE 
                                               FALSE 
                                               TRUE 
                                             
                                           
                                           Outliers  Table identifying outliers in the first four PCs of the data.   
                                      
                                       
                                  
                                
                              
                            
                           
                            
                          
                           
                      
                    
                    
                      
                           
                          
                               Heatmaps    PCA     
                              
                                  Heatmaps  
                              
                               
                              
                                  New.Annotation  
                              
                              
                            
                              
                                
                                  
                                       
                                      
                                           Heatmap of Signaling by ERBB2 Data   More Plot Information                      Heatmap of Signaling by ERBB2 Data  Heatmap of the normalized data, scaled to give all genes equal variance, generated via unsupervised clustering. Orange indicates high expression; blue indicates low expression. This plot is meant to provide a high level exploratory view of the data.   
                                      
                                       
                                  
                                
                              
                            
                            
                              
                                
                                  
                                       
                                      
                                           Principal Components of Signaling by ERBB2 Data   More Plot Information                      Principal Components of Signaling by ERBB2 Data  Principal component analysis maps high-dimensional datasets onto a smaller number of highly informative dimensions. Here, the first four principal components of the gene expression data are plotted against each other and colored by the values of the selected covariate. This plot may be used to identify clusters in the data and to identify variables associated with prominent signal in the data. Variables that are associated with these leading principal components should be considered in downstream analyses.   
                                           Outliers   More Plot Information     
                                           
                                             
                                           
                                           
                                             
                                               
                                               Outlier on PC 1 
                                               Outlier on PC 2 
                                               Outlier on PC 3 
                                               Outlier on PC 4 
                                             
                                           
                                           
                                             
                                               20161013_sebelius2-101316_01_01.RCC 
                                               FALSE 
                                               FALSE 
                                               FALSE 
                                               TRUE 
                                             
                                           
                                           Outliers  Table identifying outliers in the first four PCs of the data.   
                                      
                                       
                                  
                                
                              
                            
                           
                            
                          
                           
                      
                    
                    
                      
                           
                          
                               Heatmaps    PCA     
                              
                                  Heatmaps  
                              
                               
                              
                                  New.Annotation  
                              
                              
                            
                              
                                
                                  
                                       
                                      
                                           Heatmap of Signaling by FGFR1 in disease Data   More Plot Information                      Heatmap of Signaling by FGFR1 in disease Data  Heatmap of the normalized data, scaled to give all genes equal variance, generated via unsupervised clustering. Orange indicates high expression; blue indicates low expression. This plot is meant to provide a high level exploratory view of the data.   
                                      
                                       
                                  
                                
                              
                            
                            
                              
                                
                                  
                                       
                                      
                                           Principal Components of Signaling by FGFR1 in disease Data   More Plot Information                      Principal Components of Signaling by FGFR1 in disease Data  Principal component analysis maps high-dimensional datasets onto a smaller number of highly informative dimensions. Here, the first four principal components of the gene expression data are plotted against each other and colored by the values of the selected covariate. This plot may be used to identify clusters in the data and to identify variables associated with prominent signal in the data. Variables that are associated with these leading principal components should be considered in downstream analyses.   
                                           Outliers   More Plot Information     
                                           
                                             
                                           
                                           
                                             
                                               
                                               Outlier on PC 1 
                                               Outlier on PC 2 
                                               Outlier on PC 3 
                                               Outlier on PC 4 
                                             
                                           
                                           
                                             
                                               20161013_sebelius2-101316_01_01.RCC 
                                               FALSE 
                                               FALSE 
                                               FALSE 
                                               TRUE 
                                             
                                           
                                           Outliers  Table identifying outliers in the first four PCs of the data.   
                                      
                                       
                                  
                                
                              
                            
                           
                            
                          
                           
                      
                    
                    
                      
                           
                          
                               Heatmaps    PCA     
                              
                                  Heatmaps  
                              
                               
                              
                                  New.Annotation  
                              
                              
                            
                              
                                
                                  
                                       
                                      
                                           Heatmap of Signaling by FGFR2 in disease Data   More Plot Information                      Heatmap of Signaling by FGFR2 in disease Data  Heatmap of the normalized data, scaled to give all genes equal variance, generated via unsupervised clustering. Orange indicates high expression; blue indicates low expression. This plot is meant to provide a high level exploratory view of the data.   
                                      
                                       
                                  
                                
                              
                            
                            
                              
                                
                                  
                                       
                                      
                                           Principal Components of Signaling by FGFR2 in disease Data   More Plot Information                      Principal Components of Signaling by FGFR2 in disease Data  Principal component analysis maps high-dimensional datasets onto a smaller number of highly informative dimensions. Here, the first four principal components of the gene expression data are plotted against each other and colored by the values of the selected covariate. This plot may be used to identify clusters in the data and to identify variables associated with prominent signal in the data. Variables that are associated with these leading principal components should be considered in downstream analyses.   
                                           Outliers   More Plot Information     
                                           
                                             
                                           
                                           
                                             
                                               
                                               Outlier on PC 1 
                                               Outlier on PC 2 
                                               Outlier on PC 3 
                                               Outlier on PC 4 
                                             
                                           
                                           
                                             
                                               20161013_sebelius2-101316_01_01.RCC 
                                               FALSE 
                                               FALSE 
                                               FALSE 
                                               TRUE 
                                             
                                           
                                           Outliers  Table identifying outliers in the first four PCs of the data.   
                                      
                                       
                                  
                                
                              
                            
                           
                            
                          
                           
                      
                    
                    
                      
                           
                          
                               Heatmaps    PCA     
                              
                                  Heatmaps  
                              
                               
                              
                                  New.Annotation  
                              
                              
                            
                              
                                
                                  
                                       
                                      
                                           Heatmap of Signaling by FGFR3 in disease Data   More Plot Information                      Heatmap of Signaling by FGFR3 in disease Data  Heatmap of the normalized data, scaled to give all genes equal variance, generated via unsupervised clustering. Orange indicates high expression; blue indicates low expression. This plot is meant to provide a high level exploratory view of the data.   
                                      
                                       
                                  
                                
                              
                            
                            
                              
                                
                                  
                                       
                                      
                                           Principal Components of Signaling by FGFR3 in disease Data   More Plot Information                      Principal Components of Signaling by FGFR3 in disease Data  Principal component analysis maps high-dimensional datasets onto a smaller number of highly informative dimensions. Here, the first four principal components of the gene expression data are plotted against each other and colored by the values of the selected covariate. This plot may be used to identify clusters in the data and to identify variables associated with prominent signal in the data. Variables that are associated with these leading principal components should be considered in downstream analyses.   
                                           Outliers   More Plot Information     
                                           
                                             
                                           
                                           
                                             
                                               
                                               Outlier on PC 1 
                                               Outlier on PC 2 
                                               Outlier on PC 3 
                                               Outlier on PC 4 
                                             
                                           
                                           
                                             
                                               20161013_sebelius2-101316_01_01.RCC 
                                               FALSE 
                                               FALSE 
                                               FALSE 
                                               TRUE 
                                             
                                           
                                           Outliers  Table identifying outliers in the first four PCs of the data.   
                                      
                                       
                                  
                                
                              
                            
                           
                            
                          
                           
                      
                    
                    
                      
                           
                          
                               Heatmaps    PCA     
                              
                                  Heatmaps  
                              
                               
                              
                                  New.Annotation  
                              
                              
                            
                              
                                
                                  
                                       
                                      
                                           Heatmap of Signaling by FGFR4 in disease Data   More Plot Information                      Heatmap of Signaling by FGFR4 in disease Data  Heatmap of the normalized data, scaled to give all genes equal variance, generated via unsupervised clustering. Orange indicates high expression; blue indicates low expression. This plot is meant to provide a high level exploratory view of the data.   
                                      
                                       
                                  
                                
                              
                            
                            
                              
                                
                                  
                                       
                                      
                                           Principal Components of Signaling by FGFR4 in disease Data   More Plot Information                      Principal Components of Signaling by FGFR4 in disease Data  Principal component analysis maps high-dimensional datasets onto a smaller number of highly informative dimensions. Here, the first four principal components of the gene expression data are plotted against each other and colored by the values of the selected covariate. This plot may be used to identify clusters in the data and to identify variables associated with prominent signal in the data. Variables that are associated with these leading principal components should be considered in downstream analyses.   
                                           Outliers   More Plot Information     
                                           
                                             
                                           
                                           
                                             
                                               
                                               Outlier on PC 1 
                                               Outlier on PC 2 
                                               Outlier on PC 3 
                                               Outlier on PC 4 
                                             
                                           
                                           
                                             
                                               20161013_sebelius2-101316_01_01.RCC 
                                               FALSE 
                                               FALSE 
                                               FALSE 
                                               TRUE 
                                             
                                           
                                           Outliers  Table identifying outliers in the first four PCs of the data.   
                                      
                                       
                                  
                                
                              
                            
                           
                            
                          
                           
                      
                    
                    
                      
                           
                          
                               Heatmaps    PCA     
                              
                                  Heatmaps  
                              
                               
                              
                                  New.Annotation  
                              
                              
                            
                              
                                
                                  
                                       
                                      
                                           Heatmap of Signaling by PDGF Data   More Plot Information                      Heatmap of Signaling by PDGF Data  Heatmap of the normalized data, scaled to give all genes equal variance, generated via unsupervised clustering. Orange indicates high expression; blue indicates low expression. This plot is meant to provide a high level exploratory view of the data.   
                                      
                                       
                                  
                                
                              
                            
                            
                              
                                
                                  
                                       
                                      
                                           Principal Components of Signaling by PDGF Data   More Plot Information                      Principal Components of Signaling by PDGF Data  Principal component analysis maps high-dimensional datasets onto a smaller number of highly informative dimensions. Here, the first four principal components of the gene expression data are plotted against each other and colored by the values of the selected covariate. This plot may be used to identify clusters in the data and to identify variables associated with prominent signal in the data. Variables that are associated with these leading principal components should be considered in downstream analyses.   
                                           Outliers   More Plot Information     
                                           
                                             
                                           
                                           
                                             
                                               
                                               Outlier on PC 1 
                                               Outlier on PC 2 
                                               Outlier on PC 3 
                                               Outlier on PC 4 
                                             
                                           
                                           
                                             
                                               20161013_sebelius2-101316_01_01.RCC 
                                               FALSE 
                                               FALSE 
                                               FALSE 
                                               TRUE 
                                             
                                           
                                           Outliers  Table identifying outliers in the first four PCs of the data.   
                                      
                                       
                                  
                                
                              
                            
                           
                            
                          
                           
                      
                    
                    
                      
                           
                          
                               Heatmaps    PCA     
                              
                                  Heatmaps  
                              
                               
                              
                                  New.Annotation  
                              
                              
                            
                              
                                
                                  
                                       
                                      
                                           Heatmap of Signaling by SCF-KIT Data   More Plot Information                      Heatmap of Signaling by SCF-KIT Data  Heatmap of the normalized data, scaled to give all genes equal variance, generated via unsupervised clustering. Orange indicates high expression; blue indicates low expression. This plot is meant to provide a high level exploratory view of the data.   
                                      
                                       
                                  
                                
                              
                            
                            
                              
                                
                                  
                                       
                                      
                                           Principal Components of Signaling by SCF-KIT Data   More Plot Information                      Principal Components of Signaling by SCF-KIT Data  Principal component analysis maps high-dimensional datasets onto a smaller number of highly informative dimensions. Here, the first four principal components of the gene expression data are plotted against each other and colored by the values of the selected covariate. This plot may be used to identify clusters in the data and to identify variables associated with prominent signal in the data. Variables that are associated with these leading principal components should be considered in downstream analyses.   
                                           Outliers   More Plot Information     
                                           
                                             
                                           
                                           
                                             
                                               
                                               Outlier on PC 1 
                                               Outlier on PC 2 
                                               Outlier on PC 3 
                                               Outlier on PC 4 
                                             
                                           
                                           
                                             
                                               20161013_sebelius2-101316_01_01.RCC 
                                               FALSE 
                                               FALSE 
                                               FALSE 
                                               TRUE 
                                             
                                           
                                           Outliers  Table identifying outliers in the first four PCs of the data.   
                                      
                                       
                                  
                                
                              
                            
                           
                            
                          
                           
                      
                    
                    
                      
                           
                          
                               Heatmaps    PCA     
                              
                                  Heatmaps  
                              
                               
                              
                                  New.Annotation  
                              
                              
                            
                              
                                
                                  
                                       
                                      
                                           Heatmap of Smooth Muscle Contraction Data   More Plot Information                      Heatmap of Smooth Muscle Contraction Data  Heatmap of the normalized data, scaled to give all genes equal variance, generated via unsupervised clustering. Orange indicates high expression; blue indicates low expression. This plot is meant to provide a high level exploratory view of the data.   
                                      
                                       
                                  
                                
                              
                            
                            
                              
                                
                                  
                                       
                                      
                                           Principal Components of Smooth Muscle Contraction Data   More Plot Information                      Principal Components of Smooth Muscle Contraction Data  Principal component analysis maps high-dimensional datasets onto a smaller number of highly informative dimensions. Here, the first four principal components of the gene expression data are plotted against each other and colored by the values of the selected covariate. This plot may be used to identify clusters in the data and to identify variables associated with prominent signal in the data. Variables that are associated with these leading principal components should be considered in downstream analyses.   
                                           Outliers   More Plot Information     
                                           
                                             
                                           
                                           
                                             
                                               
                                               Outlier on PC 1 
                                               Outlier on PC 2 
                                               Outlier on PC 3 
                                               Outlier on PC 4 
                                             
                                           
                                           
                                             
                                               20161013_sebelius2-101316_01_01.RCC 
                                               FALSE 
                                               FALSE 
                                               FALSE 
                                               TRUE 
                                             
                                           
                                           Outliers  Table identifying outliers in the first four PCs of the data.   
                                      
                                       
                                  
                                
                              
                            
                           
                            
                          
                           
                      
                    
                    
                      
                           
                          
                               Heatmaps    PCA     
                              
                                  Heatmaps  
                              
                               
                              
                                  New.Annotation  
                              
                              
                            
                              
                                
                                  
                                       
                                      
                                           Heatmap of Spry regulation of FGF signaling Data   More Plot Information                      Heatmap of Spry regulation of FGF signaling Data  Heatmap of the normalized data, scaled to give all genes equal variance, generated via unsupervised clustering. Orange indicates high expression; blue indicates low expression. This plot is meant to provide a high level exploratory view of the data.   
                                      
                                       
                                  
                                
                              
                            
                            
                              
                                
                                  
                                       
                                      
                                           Principal Components of Spry regulation of FGF signaling Data   More Plot Information                      Principal Components of Spry regulation of FGF signaling Data  Principal component analysis maps high-dimensional datasets onto a smaller number of highly informative dimensions. Here, the first four principal components of the gene expression data are plotted against each other and colored by the values of the selected covariate. This plot may be used to identify clusters in the data and to identify variables associated with prominent signal in the data. Variables that are associated with these leading principal components should be considered in downstream analyses.   
                                           Outliers   More Plot Information     
                                           
                                             
                                           
                                           
                                             
                                               
                                               Outlier on PC 1 
                                               Outlier on PC 2 
                                               Outlier on PC 3 
                                               Outlier on PC 4 
                                             
                                           
                                           
                                             
                                               20161013_sebelius2-101316_01_01.RCC 
                                               FALSE 
                                               FALSE 
                                               FALSE 
                                               TRUE 
                                             
                                           
                                           Outliers  Table identifying outliers in the first four PCs of the data.   
                                      
                                       
                                  
                                
                              
                            
                           
                            
                          
                           
                      
                    
                    
                      
                           
                          
                               Heatmaps    PCA     
                              
                                  Heatmaps  
                              
                               
                              
                                  New.Annotation  
                              
                              
                            
                              
                                
                                  
                                       
                                      
                                           Heatmap of Syndecan interactions Data   More Plot Information                      Heatmap of Syndecan interactions Data  Heatmap of the normalized data, scaled to give all genes equal variance, generated via unsupervised clustering. Orange indicates high expression; blue indicates low expression. This plot is meant to provide a high level exploratory view of the data.   
                                      
                                       
                                  
                                
                              
                            
                            
                              
                                
                                  
                                       
                                      
                                           Principal Components of Syndecan interactions Data   More Plot Information                      Principal Components of Syndecan interactions Data  Principal component analysis maps high-dimensional datasets onto a smaller number of highly informative dimensions. Here, the first four principal components of the gene expression data are plotted against each other and colored by the values of the selected covariate. This plot may be used to identify clusters in the data and to identify variables associated with prominent signal in the data. Variables that are associated with these leading principal components should be considered in downstream analyses.   
                                           Outliers   More Plot Information     
                                           
                                             
                                           
                                           
                                             
                                               
                                               Outlier on PC 1 
                                               Outlier on PC 2 
                                               Outlier on PC 3 
                                               Outlier on PC 4 
                                             
                                           
                                           
                                             
                                               20161013_sebelius2-101316_01_01.RCC 
                                               FALSE 
                                               FALSE 
                                               FALSE 
                                               TRUE 
                                             
                                           
                                           Outliers  Table identifying outliers in the first four PCs of the data.   
                                      
                                       
                                  
                                
                              
                            
                           
                            
                          
                           
                      
                    
                    
                      
                           
                          
                               Heatmaps    PCA     
                              
                                  Heatmaps  
                              
                               
                              
                                  New.Annotation  
                              
                              
                            
                              
                                
                                  
                                       
                                      
                                           Heatmap of Synthesis of PIPs at the plasma membrane Data   More Plot Information                      Heatmap of Synthesis of PIPs at the plasma membrane Data  Heatmap of the normalized data, scaled to give all genes equal variance, generated via unsupervised clustering. Orange indicates high expression; blue indicates low expression. This plot is meant to provide a high level exploratory view of the data.   
                                      
                                       
                                  
                                
                              
                            
                            
                              
                                
                                  
                                       
                                      
                                           Principal Components of Synthesis of PIPs at the plasma membrane Data   More Plot Information                      Principal Components of Synthesis of PIPs at the plasma membrane Data  Principal component analysis maps high-dimensional datasets onto a smaller number of highly informative dimensions. Here, the first four principal components of the gene expression data are plotted against each other and colored by the values of the selected covariate. This plot may be used to identify clusters in the data and to identify variables associated with prominent signal in the data. Variables that are associated with these leading principal components should be considered in downstream analyses.   
                                           Outliers   More Plot Information     
                                           
                                             
                                           
                                           
                                             
                                               
                                               Outlier on PC 1 
                                               Outlier on PC 2 
                                               Outlier on PC 3 
                                               Outlier on PC 4 
                                             
                                           
                                           
                                             
                                               20161013_sebelius2-101316_01_01.RCC 
                                               FALSE 
                                               FALSE 
                                               FALSE 
                                               TRUE 
                                             
                                           
                                           Outliers  Table identifying outliers in the first four PCs of the data.   
                                      
                                       
                                  
                                
                              
                            
                           
                            
                          
                           
                      
                    
                    
                      
                           
                          
                               Heatmaps    PCA     
                              
                                  Heatmaps  
                              
                               
                              
                                  New.Annotation  
                              
                              
                            
                              
                                
                                  
                                       
                                      
                                           Heatmap of T41 mutants of beta-catenin aren.SQT.t phosphorylated Data   More Plot Information                      Heatmap of T41 mutants of beta-catenin aren.SQT.t phosphorylated Data  Heatmap of the normalized data, scaled to give all genes equal variance, generated via unsupervised clustering. Orange indicates high expression; blue indicates low expression. This plot is meant to provide a high level exploratory view of the data.   
                                      
                                       
                                  
                                
                              
                            
                            
                              
                                
                                  
                                       
                                      
                                           Principal Components of T41 mutants of beta-catenin aren.SQT.t phosphorylated Data   More Plot Information                      Principal Components of T41 mutants of beta-catenin aren.SQT.t phosphorylated Data  Principal component analysis maps high-dimensional datasets onto a smaller number of highly informative dimensions. Here, the first four principal components of the gene expression data are plotted against each other and colored by the values of the selected covariate. This plot may be used to identify clusters in the data and to identify variables associated with prominent signal in the data. Variables that are associated with these leading principal components should be considered in downstream analyses.   
                                           Outliers   More Plot Information     
                                           
                                             
                                           
                                           
                                             
                                               
                                               Outlier on PC 1 
                                               Outlier on PC 2 
                                               Outlier on PC 3 
                                               Outlier on PC 4 
                                             
                                           
                                           
                                             
                                               20161013_sebelius2-101316_01_01.RCC 
                                               FALSE 
                                               FALSE 
                                               FALSE 
                                               TRUE 
                                             
                                           
                                           Outliers  Table identifying outliers in the first four PCs of the data.   
                                      
                                       
                                  
                                
                              
                            
                           
                            
                          
                           
                      
                    
                    
                      
                           
                          
                               Heatmaps    PCA     
                              
                                  Heatmaps  
                              
                               
                              
                                  New.Annotation  
                              
                              
                            
                              
                                
                                  
                                       
                                      
                                           Heatmap of TGF-beta receptor signaling activates SMADs Data   More Plot Information                      Heatmap of TGF-beta receptor signaling activates SMADs Data  Heatmap of the normalized data, scaled to give all genes equal variance, generated via unsupervised clustering. Orange indicates high expression; blue indicates low expression. This plot is meant to provide a high level exploratory view of the data.   
                                      
                                       
                                  
                                
                              
                            
                            
                              
                                
                                  
                                       
                                      
                                           Principal Components of TGF-beta receptor signaling activates SMADs Data   More Plot Information                      Principal Components of TGF-beta receptor signaling activates SMADs Data  Principal component analysis maps high-dimensional datasets onto a smaller number of highly informative dimensions. Here, the first four principal components of the gene expression data are plotted against each other and colored by the values of the selected covariate. This plot may be used to identify clusters in the data and to identify variables associated with prominent signal in the data. Variables that are associated with these leading principal components should be considered in downstream analyses.   
                                           Outliers   More Plot Information     
                                           
                                             
                                           
                                           
                                             
                                               
                                               Outlier on PC 1 
                                               Outlier on PC 2 
                                               Outlier on PC 3 
                                               Outlier on PC 4 
                                             
                                           
                                           
                                             
                                               20161013_sebelius2-101316_01_01.RCC 
                                               FALSE 
                                               FALSE 
                                               FALSE 
                                               TRUE 
                                             
                                           
                                           Outliers  Table identifying outliers in the first four PCs of the data.   
                                      
                                       
                                  
                                
                              
                            
                           
                            
                          
                           
                      
                    
                    
                      
                           
                          
                               Heatmaps    PCA     
                              
                                  Heatmaps  
                              
                               
                              
                                  New.Annotation  
                              
                              
                            
                              
                                
                                  
                                       
                                      
                                           Heatmap of TGF-beta receptor signaling in EMT (epithelial to mesenchymal transition) Data   More Plot Information                      Heatmap of TGF-beta receptor signaling in EMT (epithelial to mesenchymal transition) Data  Heatmap of the normalized data, scaled to give all genes equal variance, generated via unsupervised clustering. Orange indicates high expression; blue indicates low expression. This plot is meant to provide a high level exploratory view of the data.   
                                      
                                       
                                  
                                
                              
                            
                            
                              
                                
                                  
                                       
                                      
                                           Principal Components of TGF-beta receptor signaling in EMT (epithelial to mesenchymal transition) Data   More Plot Information                      Principal Components of TGF-beta receptor signaling in EMT (epithelial to mesenchymal transition) Data  Principal component analysis maps high-dimensional datasets onto a smaller number of highly informative dimensions. Here, the first four principal components of the gene expression data are plotted against each other and colored by the values of the selected covariate. This plot may be used to identify clusters in the data and to identify variables associated with prominent signal in the data. Variables that are associated with these leading principal components should be considered in downstream analyses.   
                                           Outliers   More Plot Information     
                                           
                                             
                                           
                                           
                                             
                                               
                                               Outlier on PC 1 
                                               Outlier on PC 2 
                                               Outlier on PC 3 
                                               Outlier on PC 4 
                                             
                                           
                                           
                                             
                                               20161013_sebelius2-101316_01_01.RCC 
                                               FALSE 
                                               FALSE 
                                               FALSE 
                                               TRUE 
                                             
                                           
                                           Outliers  Table identifying outliers in the first four PCs of the data.   
                                      
                                       
                                  
                                
                              
                            
                           
                            
                          
                           
                      
                    
                    
                      
                           
                          
                               Heatmaps    PCA     
                              
                                  Heatmaps  
                              
                               
                              
                                  New.Annotation  
                              
                              
                            
                              
                                
                                  
                                       
                                      
                                           Heatmap of Tie2 Signaling Data   More Plot Information                      Heatmap of Tie2 Signaling Data  Heatmap of the normalized data, scaled to give all genes equal variance, generated via unsupervised clustering. Orange indicates high expression; blue indicates low expression. This plot is meant to provide a high level exploratory view of the data.   
                                      
                                       
                                  
                                
                              
                            
                            
                              
                                
                                  
                                       
                                      
                                           Principal Components of Tie2 Signaling Data   More Plot Information                      Principal Components of Tie2 Signaling Data  Principal component analysis maps high-dimensional datasets onto a smaller number of highly informative dimensions. Here, the first four principal components of the gene expression data are plotted against each other and colored by the values of the selected covariate. This plot may be used to identify clusters in the data and to identify variables associated with prominent signal in the data. Variables that are associated with these leading principal components should be considered in downstream analyses.   
                                           Outliers   More Plot Information     
                                           
                                             
                                           
                                           
                                             
                                               
                                               Outlier on PC 1 
                                               Outlier on PC 2 
                                               Outlier on PC 3 
                                               Outlier on PC 4 
                                             
                                           
                                           
                                             
                                               20161013_sebelius2-101316_01_01.RCC 
                                               FALSE 
                                               FALSE 
                                               FALSE 
                                               TRUE 
                                             
                                           
                                           Outliers  Table identifying outliers in the first four PCs of the data.   
                                      
                                       
                                  
                                
                              
                            
                           
                            
                          
                           
                      
                    
                    
                      
                           
                          
                               Heatmaps    PCA     
                              
                                  Heatmaps  
                              
                               
                              
                                  New.Annotation  
                              
                              
                            
                              
                                
                                  
                                       
                                      
                                           Heatmap of Tight junction interactions Data   More Plot Information                      Heatmap of Tight junction interactions Data  Heatmap of the normalized data, scaled to give all genes equal variance, generated via unsupervised clustering. Orange indicates high expression; blue indicates low expression. This plot is meant to provide a high level exploratory view of the data.   
                                      
                                       
                                  
                                
                              
                            
                            
                              
                                
                                  
                                       
                                      
                                           Principal Components of Tight junction interactions Data   More Plot Information                      Principal Components of Tight junction interactions Data  Principal component analysis maps high-dimensional datasets onto a smaller number of highly informative dimensions. Here, the first four principal components of the gene expression data are plotted against each other and colored by the values of the selected covariate. This plot may be used to identify clusters in the data and to identify variables associated with prominent signal in the data. Variables that are associated with these leading principal components should be considered in downstream analyses.   
                                           Outliers   More Plot Information     
                                           
                                             
                                           
                                           
                                             
                                               
                                               Outlier on PC 1 
                                               Outlier on PC 2 
                                               Outlier on PC 3 
                                               Outlier on PC 4 
                                             
                                           
                                           
                                             
                                               20161013_sebelius2-101316_01_01.RCC 
                                               FALSE 
                                               FALSE 
                                               FALSE 
                                               TRUE 
                                             
                                           
                                           Outliers  Table identifying outliers in the first four PCs of the data.   
                                      
                                       
                                  
                                
                              
                            
                           
                            
                          
                           
                      
                    
                    
                      
                           
                          
                               Heatmaps    PCA     
                              
                                  Heatmaps  
                              
                               
                              
                                  New.Annotation  
                              
                              
                            
                              
                                
                                  
                                       
                                      
                                           Heatmap of TP53 Regulates Metabolic Genes Data   More Plot Information                      Heatmap of TP53 Regulates Metabolic Genes Data  Heatmap of the normalized data, scaled to give all genes equal variance, generated via unsupervised clustering. Orange indicates high expression; blue indicates low expression. This plot is meant to provide a high level exploratory view of the data.   
                                      
                                       
                                  
                                
                              
                            
                            
                              
                                
                                  
                                       
                                      
                                           Principal Components of TP53 Regulates Metabolic Genes Data   More Plot Information                      Principal Components of TP53 Regulates Metabolic Genes Data  Principal component analysis maps high-dimensional datasets onto a smaller number of highly informative dimensions. Here, the first four principal components of the gene expression data are plotted against each other and colored by the values of the selected covariate. This plot may be used to identify clusters in the data and to identify variables associated with prominent signal in the data. Variables that are associated with these leading principal components should be considered in downstream analyses.   
                                           Outliers   More Plot Information     
                                           
                                             
                                           
                                           
                                             
                                               
                                               Outlier on PC 1 
                                               Outlier on PC 2 
                                               Outlier on PC 3 
                                               Outlier on PC 4 
                                             
                                           
                                           
                                             
                                               20161013_sebelius2-101316_01_01.RCC 
                                               FALSE 
                                               FALSE 
                                               FALSE 
                                               TRUE 
                                             
                                           
                                           Outliers  Table identifying outliers in the first four PCs of the data.   
                                      
                                       
                                  
                                
                              
                            
                           
                            
                          
                           
                      
                    
                    
                      
                           
                          
                               Heatmaps    PCA     
                              
                                  Heatmaps  
                              
                               
                              
                                  New.Annotation  
                              
                              
                            
                              
                                
                                  
                                       
                                      
                                           Heatmap of Truncations of AMER1 destabilize the destruction complex Data   More Plot Information                      Heatmap of Truncations of AMER1 destabilize the destruction complex Data  Heatmap of the normalized data, scaled to give all genes equal variance, generated via unsupervised clustering. Orange indicates high expression; blue indicates low expression. This plot is meant to provide a high level exploratory view of the data.   
                                      
                                       
                                  
                                
                              
                            
                            
                              
                                
                                  
                                       
                                      
                                           Principal Components of Truncations of AMER1 destabilize the destruction complex Data   More Plot Information                      Principal Components of Truncations of AMER1 destabilize the destruction complex Data  Principal component analysis maps high-dimensional datasets onto a smaller number of highly informative dimensions. Here, the first four principal components of the gene expression data are plotted against each other and colored by the values of the selected covariate. This plot may be used to identify clusters in the data and to identify variables associated with prominent signal in the data. Variables that are associated with these leading principal components should be considered in downstream analyses.   
                                           Outliers   More Plot Information     
                                           
                                             
                                           
                                           
                                             
                                               
                                               Outlier on PC 1 
                                               Outlier on PC 2 
                                               Outlier on PC 3 
                                               Outlier on PC 4 
                                             
                                           
                                           
                                             
                                               20161013_sebelius2-101316_01_01.RCC 
                                               FALSE 
                                               FALSE 
                                               FALSE 
                                               TRUE 
                                             
                                           
                                           Outliers  Table identifying outliers in the first four PCs of the data.   
                                      
                                       
                                  
                                
                              
                            
                           
                            
                          
                           
                      
                    
                    
                      
                           
                          
                               Heatmaps    PCA     
                              
                                  Heatmaps  
                              
                               
                              
                                  New.Annotation  
                              
                              
                            
                              
                                
                                  
                                       
                                      
                                           Heatmap of VEGFA-VEGFR2 Pathway Data   More Plot Information                      Heatmap of VEGFA-VEGFR2 Pathway Data  Heatmap of the normalized data, scaled to give all genes equal variance, generated via unsupervised clustering. Orange indicates high expression; blue indicates low expression. This plot is meant to provide a high level exploratory view of the data.   
                                      
                                       
                                  
                                
                              
                            
                            
                              
                                
                                  
                                       
                                      
                                           Principal Components of VEGFA-VEGFR2 Pathway Data   More Plot Information                      Principal Components of VEGFA-VEGFR2 Pathway Data  Principal component analysis maps high-dimensional datasets onto a smaller number of highly informative dimensions. Here, the first four principal components of the gene expression data are plotted against each other and colored by the values of the selected covariate. This plot may be used to identify clusters in the data and to identify variables associated with prominent signal in the data. Variables that are associated with these leading principal components should be considered in downstream analyses.   
                                           Outliers   More Plot Information     
                                           
                                             
                                           
                                           
                                             
                                               
                                               Outlier on PC 1 
                                               Outlier on PC 2 
                                               Outlier on PC 3 
                                               Outlier on PC 4 
                                             
                                           
                                           
                                             
                                               20161013_sebelius2-101316_01_01.RCC 
                                               FALSE 
                                               FALSE 
                                               FALSE 
                                               TRUE 
                                             
                                           
                                           Outliers  Table identifying outliers in the first four PCs of the data.   
                                      
                                       
                                  
                                
                              
                            
                           
                            
                          
                           
                      
                    
                    
                      
                           
                          
                               Heatmaps    PCA     
                              
                                  Heatmaps  
                              
                               
                              
                                  New.Annotation  
                              
                              
                            
                              
                                
                                  
                                       
                                      
                                           Heatmap of VEGFR2 mediated cell proliferation Data   More Plot Information                      Heatmap of VEGFR2 mediated cell proliferation Data  Heatmap of the normalized data, scaled to give all genes equal variance, generated via unsupervised clustering. Orange indicates high expression; blue indicates low expression. This plot is meant to provide a high level exploratory view of the data.   
                                      
                                       
                                  
                                
                              
                            
                            
                              
                                
                                  
                                       
                                      
                                           Principal Components of VEGFR2 mediated cell proliferation Data   More Plot Information                      Principal Components of VEGFR2 mediated cell proliferation Data  Principal component analysis maps high-dimensional datasets onto a smaller number of highly informative dimensions. Here, the first four principal components of the gene expression data are plotted against each other and colored by the values of the selected covariate. This plot may be used to identify clusters in the data and to identify variables associated with prominent signal in the data. Variables that are associated with these leading principal components should be considered in downstream analyses.   
                                           Outliers   More Plot Information     
                                           
                                             
                                           
                                           
                                             
                                               
                                               Outlier on PC 1 
                                               Outlier on PC 2 
                                               Outlier on PC 3 
                                               Outlier on PC 4 
                                             
                                           
                                           
                                             
                                               20161013_sebelius2-101316_01_01.RCC 
                                               FALSE 
                                               FALSE 
                                               FALSE 
                                               TRUE 
                                             
                                           
                                           Outliers  Table identifying outliers in the first four PCs of the data.   
                                      
                                       
                                  
                                
                              
                            
                           
                            
                          
                           
                      
                    
                    
                      
                           
                          
                               Heatmaps    PCA     
                              
                                  Heatmaps  
                              
                               
                              
                                  New.Annotation  
                              
                              
                            
                              
                                
                                  
                                       
                                      
                                           Heatmap of VEGFR2 mediated vascular permeability Data   More Plot Information                      Heatmap of VEGFR2 mediated vascular permeability Data  Heatmap of the normalized data, scaled to give all genes equal variance, generated via unsupervised clustering. Orange indicates high expression; blue indicates low expression. This plot is meant to provide a high level exploratory view of the data.   
                                      
                                       
                                  
                                
                              
                            
                            
                              
                                
                                  
                                       
                                      
                                           Principal Components of VEGFR2 mediated vascular permeability Data   More Plot Information                      Principal Components of VEGFR2 mediated vascular permeability Data  Principal component analysis maps high-dimensional datasets onto a smaller number of highly informative dimensions. Here, the first four principal components of the gene expression data are plotted against each other and colored by the values of the selected covariate. This plot may be used to identify clusters in the data and to identify variables associated with prominent signal in the data. Variables that are associated with these leading principal components should be considered in downstream analyses.   
                                           Outliers   More Plot Information     
                                           
                                             
                                           
                                           
                                             
                                               
                                               Outlier on PC 1 
                                               Outlier on PC 2 
                                               Outlier on PC 3 
                                               Outlier on PC 4 
                                             
                                           
                                           
                                             
                                               20161013_sebelius2-101316_01_01.RCC 
                                               FALSE 
                                               FALSE 
                                               FALSE 
                                               TRUE 
                                             
                                           
                                           Outliers  Table identifying outliers in the first four PCs of the data.   
                                      
                                       
                                  
                                
                              
                            
                           
                            
                          
                           
                      
                    
                    
                      
                           
                          
                               Heatmaps    PCA     
                              
                                  Heatmaps  
                              
                               
                              
                                  New.Annotation  
                              
                              
                            
                              
                                
                                  
                                       
                                      
                                           Heatmap of WNT mediated activation of DVL Data   More Plot Information                      Heatmap of WNT mediated activation of DVL Data  Heatmap of the normalized data, scaled to give all genes equal variance, generated via unsupervised clustering. Orange indicates high expression; blue indicates low expression. This plot is meant to provide a high level exploratory view of the data.   
                                      
                                       
                                  
                                
                              
                            
                            
                              
                                
                                  
                                       
                                      
                                           Principal Components of WNT mediated activation of DVL Data   More Plot Information                      Principal Components of WNT mediated activation of DVL Data  Principal component analysis maps high-dimensional datasets onto a smaller number of highly informative dimensions. Here, the first four principal components of the gene expression data are plotted against each other and colored by the values of the selected covariate. This plot may be used to identify clusters in the data and to identify variables associated with prominent signal in the data. Variables that are associated with these leading principal components should be considered in downstream analyses.   
                                           Outliers   More Plot Information     
                                           
                                             
                                           
                                           
                                             
                                               
                                               Outlier on PC 1 
                                               Outlier on PC 2 
                                               Outlier on PC 3 
                                               Outlier on PC 4 
                                             
                                           
                                           
                                             
                                               20161013_sebelius2-101316_01_01.RCC 
                                               FALSE 
                                               FALSE 
                                               FALSE 
                                               TRUE 
                                             
                                           
                                           Outliers  Table identifying outliers in the first four PCs of the data.   
                                      
                                       
                                  
                                
                              
                            
                           
                            
                          
                           
                      
                    
                  
                 
              
               
          
        
        
          
                Displays plots that detail the impact of normalization on the data.      
              
                  Analyte: 
                  
                      mRNA  
                  
                  
                  
                    
                      
                           
                          
                               Pairwise Variance during HK Selection   More Plot Information                Pairwise Variance during HK Selection  Displays the geNorm pairwise variation statistic after successive genes are removed. This statistic cannot be computed for the final two genes, which are therefore not displayed. The ideal normalization gene set will minimize the pairwise variation statistic.   
                               Normalization Summary: mRNA   More Plot Information      mRNA Normalization Summary Data      All Normalized Data      mRNA Normalized Data      Download HK Genes                Normalization Summary: mRNA  The lower the MSE of reference profile the higher quality of normalization. The more positive normalization factor, the higher counts were registered for the sample and vice versa   
                          
                           
                      
                    
                  
                 
              
                
          
        
        
          
                Pathway scores condense each sample's gene expression profile into a small set of pathway scores. An experiment can then be explored through the lens of pathway scores instead of in the much higher-dimension lens of gene expression values. Pathway scores are fit using the first principal component of each gene set's data. They are oriented such that increasing score corresponds to mostly increasing expression (specifically, each pathway score has positive weights for at least half its genes).
        Summary plots explore the joint behavior of pathways, and Covariates plots compare pathway scores to covariates.      
              
                   Summary    Covariates     
                  
                      score  
                  
                   
                  
                      score  
                  
                  
                
                  
                    
                      
                           
                          
                               Choose a Pathway: 
                              
                                  Summary  
                                  Activation of Rac  
                                  Apoptotic cleavage of cellular proteins  
                                  ARMS-mediated activation  
                                  AXIN missense mutants destabilize the destruction complex  
                                  Beta-catenin phosphorylation cascade  
                                  CD209 (DC-SIGN) signaling  
                                  CD28 co-stimulation  
                                  CD28 dependent PI3KAkt signaling  
                                  CD28 dependent Vav1 pathway  
                                  Cell-extracellular matrix interactions  
                                  Cell surface interactions at the vascular wall  
                                  Circadian Clock  
                                  Constitutive Signaling by Aberrant PI3K in Cancer  
                                  Constitutive Signaling by EGFRvIII  
                                  CTLA4 inhibitory signaling  
                                  Cyclin D associated events in G1  
                                  DAP12 signaling  
                                  Deactivation of the beta-catenin transactivating complex  
                                  Disassembly of the destruction complex and recruitment of AXIN t...  
                                  Downstream signal transduction  
                                  Downstream TCR signaling  
                                  ECM proteoglycans  
                                  Elastic fibre formation  
                                  EPH-ephrin mediated repulsion of cells  
                                  EPHA-mediated growth cone collapse  
                                  EPHB-mediated forward signaling  
                                  Ephrin signaling  
                                  FCERI mediated MAPK activation  
                                  Formation of the beta-catenin.CL.TCF transactivating complex  
                                  Frs2-mediated activation  
                                  G alpha (1213) signalling events  
                                  G beta.CL.gamma signalling through PI3Kgamma  
                                  GAB1 signalosome  
                                  Gastrin-CREB signalling pathway via PKC and MAPK  
                                  GPVI-mediated activation cascade  
                                  Integrin alphaIIb beta3 signaling  
                                  Integrin cell surface interactions  
                                  Interleukin receptor SHC signaling  
                                  IRS-related events triggered by IGF1R  
                                  Laminin interactions  
                                  MAP2K and MAPK activation  
                                  MAPK6MAPK4 signaling  
                                  MicroRNA (miRNA) biogenesis  
                                  Misspliced GSK3beta mutants stabilize beta-catenin  
                                  Molecules associated with elastic fibres  
                                  Negative regulation of FGFR1 signaling  
                                  Negative regulation of FGFR2 signaling  
                                  Negative regulation of FGFR3 signaling  
                                  Negative regulation of FGFR4 signaling  
                                  Negative regulation of MAPK pathway  
                                  Nephrin interactions  
                                  Non-integrin membrane-ECM interactions  
                                  NRAGE signals death through JNK  
                                  PCPCE pathway  
                                  PI3K Cascade  
                                  PI3KAKT activation  
                                  PIP3 activates AKT signaling  
                                  Platelet degranulation   
                                  RAF activation  
                                  RAFMAP kinase cascade  
                                  Regulation of actin dynamics for phagocytic cup formation  
                                  Regulation of HSF1-mediated heat shock response  
                                  Regulation of signaling by CBL  
                                  Rho GTPase cycle  
                                  RHO GTPases activate CIT  
                                  RHO GTPases Activate Formins  
                                  RHO GTPases activate IQGAPs  
                                  RHO GTPases activate PAKs  
                                  RHO GTPases activate PKNs  
                                  RHO GTPases Activate ROCKs  
                                  RHO GTPases Activate WASPs and WAVEs  
                                  Role of LAT2NTALLAB on calcium mobilization  
                                  S33 mutants of beta-catenin aren.SQT.t phosphorylated  
                                  S37 mutants of beta-catenin aren.SQT.t phosphorylated  
                                  S45 mutants of beta-catenin aren.SQT.t phosphorylated  
                                  Sema3A PAK dependent Axon repulsion  
                                  Sema4D induced cell migration and growth-cone collapse  
                                  SHC-mediated cascade.CL.FGFR1  
                                  SHC-related events triggered by IGF1R  
                                  Signal transduction by L1  
                                  Signaling by ERBB2  
                                  Signaling by FGFR1 in disease  
                                  Signaling by FGFR2 in disease  
                                  Signaling by FGFR3 in disease  
                                  Signaling by FGFR4 in disease  
                                  Signaling by PDGF  
                                  Signaling by SCF-KIT  
                                  Smooth Muscle Contraction  
                                  Spry regulation of FGF signaling  
                                  Syndecan interactions  
                                  Synthesis of PIPs at the plasma membrane  
                                  T41 mutants of beta-catenin aren.SQT.t phosphorylated  
                                  TGF-beta receptor signaling activates SMADs  
                                  Tie2 Signaling  
                                  Tight junction interactions  
                                  Truncations of AMER1 destabilize the destruction complex  
                                  VEGFA-VEGFR2 Pathway  
                                  VEGFR2 mediated cell proliferation  
                                  VEGFR2 mediated vascular permeability  
                              
                               
                              
                                
                                  
                                       
                                      
                                           Heatmap of Pathway scores   More Plot Information      Download Pathway scores                      Heatmap of Pathway scores  Heatmap showing pathway score. This plot is a high level overview of how the pathway scores change across samples. Use it to understand how pathway scores cluster together and which samples exhibit similar pathway score profiles. Orange indicates high scores; blue indicates low scores. Scores are displayed on the same scale via a Z-transformation.   
                                           Heatmap of Correlation Matrix of Pathway scores   More Plot Information                Heatmap of Correlation Matrix of Pathway scores  Heatmap showing the correlation matrix of pathway scores. Orange and blue indicate positive and negative correlation, respectively.   
                                      
                                       
                                  
                                
                                
                                  
                                       
                                      
                                           Activation of Rac measurements vs. other pathway scores   More Plot Information                      Activation of Rac measurements vs. other pathway scores  Each panel plots Activation of Rac against another pathway score.  Points are colored by New.Annotation.   
                                      
                                       
                                  
                                
                                
                                  
                                       
                                      
                                           Apoptotic cleavage of cellular proteins measurements vs. other pathway scores   More Plot Information                      Apoptotic cleavage of cellular proteins measurements vs. other pathway scores  Each panel plots Apoptotic cleavage of cellular proteins against another pathway score.  Points are colored by New.Annotation.   
                                      
                                       
                                  
                                
                                
                                  
                                       
                                      
                                           ARMS-mediated activation measurements vs. other pathway scores   More Plot Information                      ARMS-mediated activation measurements vs. other pathway scores  Each panel plots ARMS-mediated activation against another pathway score.  Points are colored by New.Annotation.   
                                      
                                       
                                  
                                
                                
                                  
                                       
                                      
                                           AXIN missense mutants destabilize the destruction complex measurements vs. other pathway scores   More Plot Information                      AXIN missense mutants destabilize the destruction complex measurements vs. other pathway scores  Each panel plots AXIN missense mutants destabilize the destruction complex against another pathway score.  Points are colored by New.Annotation.   
                                      
                                       
                                  
                                
                                
                                  
                                       
                                      
                                           Beta-catenin phosphorylation cascade measurements vs. other pathway scores   More Plot Information                      Beta-catenin phosphorylation cascade measurements vs. other pathway scores  Each panel plots Beta-catenin phosphorylation cascade against another pathway score.  Points are colored by New.Annotation.   
                                      
                                       
                                  
                                
                                
                                  
                                       
                                      
                                           CD209 (DC-SIGN) signaling measurements vs. other pathway scores   More Plot Information                      CD209 (DC-SIGN) signaling measurements vs. other pathway scores  Each panel plots CD209 (DC-SIGN) signaling against another pathway score.  Points are colored by New.Annotation.   
                                      
                                       
                                  
                                
                                
                                  
                                       
                                      
                                           CD28 co-stimulation measurements vs. other pathway scores   More Plot Information                      CD28 co-stimulation measurements vs. other pathway scores  Each panel plots CD28 co-stimulation against another pathway score.  Points are colored by New.Annotation.   
                                      
                                       
                                  
                                
                                
                                  
                                       
                                      
                                           CD28 dependent PI3KAkt signaling measurements vs. other pathway scores   More Plot Information                      CD28 dependent PI3KAkt signaling measurements vs. other pathway scores  Each panel plots CD28 dependent PI3KAkt signaling against another pathway score.  Points are colored by New.Annotation.   
                                      
                                       
                                  
                                
                                
                                  
                                       
                                      
                                           CD28 dependent Vav1 pathway measurements vs. other pathway scores   More Plot Information                      CD28 dependent Vav1 pathway measurements vs. other pathway scores  Each panel plots CD28 dependent Vav1 pathway against another pathway score.  Points are colored by New.Annotation.   
                                      
                                       
                                  
                                
                                
                                  
                                       
                                      
                                           Cell-extracellular matrix interactions measurements vs. other pathway scores   More Plot Information                      Cell-extracellular matrix interactions measurements vs. other pathway scores  Each panel plots Cell-extracellular matrix interactions against another pathway score.  Points are colored by New.Annotation.   
                                      
                                       
                                  
                                
                                
                                  
                                       
                                      
                                           Cell surface interactions at the vascular wall measurements vs. other pathway scores   More Plot Information                      Cell surface interactions at the vascular wall measurements vs. other pathway scores  Each panel plots Cell surface interactions at the vascular wall against another pathway score.  Points are colored by New.Annotation.   
                                      
                                       
                                  
                                
                                
                                  
                                       
                                      
                                           Circadian Clock measurements vs. other pathway scores   More Plot Information                      Circadian Clock measurements vs. other pathway scores  Each panel plots Circadian Clock against another pathway score.  Points are colored by New.Annotation.   
                                      
                                       
                                  
                                
                                
                                  
                                       
                                      
                                           Constitutive Signaling by Aberrant PI3K in Cancer measurements vs. other pathway scores   More Plot Information                      Constitutive Signaling by Aberrant PI3K in Cancer measurements vs. other pathway scores  Each panel plots Constitutive Signaling by Aberrant PI3K in Cancer against another pathway score.  Points are colored by New.Annotation.   
                                      
                                       
                                  
                                
                                
                                  
                                       
                                      
                                           Constitutive Signaling by EGFRvIII measurements vs. other pathway scores   More Plot Information                      Constitutive Signaling by EGFRvIII measurements vs. other pathway scores  Each panel plots Constitutive Signaling by EGFRvIII against another pathway score.  Points are colored by New.Annotation.   
                                      
                                       
                                  
                                
                                
                                  
                                       
                                      
                                           CTLA4 inhibitory signaling measurements vs. other pathway scores   More Plot Information                      CTLA4 inhibitory signaling measurements vs. other pathway scores  Each panel plots CTLA4 inhibitory signaling against another pathway score.  Points are colored by New.Annotation.   
                                      
                                       
                                  
                                
                                
                                  
                                       
                                      
                                           Cyclin D associated events in G1 measurements vs. other pathway scores   More Plot Information                      Cyclin D associated events in G1 measurements vs. other pathway scores  Each panel plots Cyclin D associated events in G1 against another pathway score.  Points are colored by New.Annotation.   
                                      
                                       
                                  
                                
                                
                                  
                                       
                                      
                                           DAP12 signaling measurements vs. other pathway scores   More Plot Information                      DAP12 signaling measurements vs. other pathway scores  Each panel plots DAP12 signaling against another pathway score.  Points are colored by New.Annotation.   
                                      
                                       
                                  
                                
                                
                                  
                                       
                                      
                                           Deactivation of the beta-catenin transactivating complex measurements vs. other pathway scores   More Plot Information                      Deactivation of the beta-catenin transactivating complex measurements vs. other pathway scores  Each panel plots Deactivation of the beta-catenin transactivating complex against another pathway score.  Points are colored by New.Annotation.   
                                      
                                       
                                  
                                
                                
                                  
                                       
                                      
                                           Disassembly of the destruction complex and recruitment of AXIN to the membrane measurements vs. other pathway scores   More Plot Information                      Disassembly of the destruction complex and recruitment of AXIN to the membrane measurements vs. other pathway scores  Each panel plots Disassembly of the destruction complex and recruitment of AXIN to the membrane against another pathway score.  Points are colored by New.Annotation.   
                                      
                                       
                                  
                                
                                
                                  
                                       
                                      
                                           Downstream signal transduction measurements vs. other pathway scores   More Plot Information                      Downstream signal transduction measurements vs. other pathway scores  Each panel plots Downstream signal transduction against another pathway score.  Points are colored by New.Annotation.   
                                      
                                       
                                  
                                
                                
                                  
                                       
                                      
                                           Downstream TCR signaling measurements vs. other pathway scores   More Plot Information                      Downstream TCR signaling measurements vs. other pathway scores  Each panel plots Downstream TCR signaling against another pathway score.  Points are colored by New.Annotation.   
                                      
                                       
                                  
                                
                                
                                  
                                       
                                      
                                           ECM proteoglycans measurements vs. other pathway scores   More Plot Information                      ECM proteoglycans measurements vs. other pathway scores  Each panel plots ECM proteoglycans against another pathway score.  Points are colored by New.Annotation.   
                                      
                                       
                                  
                                
                                
                                  
                                       
                                      
                                           Elastic fibre formation measurements vs. other pathway scores   More Plot Information                      Elastic fibre formation measurements vs. other pathway scores  Each panel plots Elastic fibre formation against another pathway score.  Points are colored by New.Annotation.   
                                      
                                       
                                  
                                
                                
                                  
                                       
                                      
                                           EPH-ephrin mediated repulsion of cells measurements vs. other pathway scores   More Plot Information                      EPH-ephrin mediated repulsion of cells measurements vs. other pathway scores  Each panel plots EPH-ephrin mediated repulsion of cells against another pathway score.  Points are colored by New.Annotation.   
                                      
                                       
                                  
                                
                                
                                  
                                       
                                      
                                           EPHA-mediated growth cone collapse measurements vs. other pathway scores   More Plot Information                      EPHA-mediated growth cone collapse measurements vs. other pathway scores  Each panel plots EPHA-mediated growth cone collapse against another pathway score.  Points are colored by New.Annotation.   
                                      
                                       
                                  
                                
                                
                                  
                                       
                                      
                                           EPHB-mediated forward signaling measurements vs. other pathway scores   More Plot Information                      EPHB-mediated forward signaling measurements vs. other pathway scores  Each panel plots EPHB-mediated forward signaling against another pathway score.  Points are colored by New.Annotation.   
                                      
                                       
                                  
                                
                                
                                  
                                       
                                      
                                           Ephrin signaling measurements vs. other pathway scores   More Plot Information                      Ephrin signaling measurements vs. other pathway scores  Each panel plots Ephrin signaling against another pathway score.  Points are colored by New.Annotation.   
                                      
                                       
                                  
                                
                                
                                  
                                       
                                      
                                           FCERI mediated MAPK activation measurements vs. other pathway scores   More Plot Information                      FCERI mediated MAPK activation measurements vs. other pathway scores  Each panel plots FCERI mediated MAPK activation against another pathway score.  Points are colored by New.Annotation.   
                                      
                                       
                                  
                                
                                
                                  
                                       
                                      
                                           Formation of the beta-catenin.CL.TCF transactivating complex measurements vs. other pathway scores   More Plot Information                      Formation of the beta-catenin.CL.TCF transactivating complex measurements vs. other pathway scores  Each panel plots Formation of the beta-catenin.CL.TCF transactivating complex against another pathway score.  Points are colored by New.Annotation.   
                                      
                                       
                                  
                                
                                
                                  
                                       
                                      
                                           Frs2-mediated activation measurements vs. other pathway scores   More Plot Information                      Frs2-mediated activation measurements vs. other pathway scores  Each panel plots Frs2-mediated activation against another pathway score.  Points are colored by New.Annotation.   
                                      
                                       
                                  
                                
                                
                                  
                                       
                                      
                                           G alpha (1213) signalling events measurements vs. other pathway scores   More Plot Information                      G alpha (1213) signalling events measurements vs. other pathway scores  Each panel plots G alpha (1213) signalling events against another pathway score.  Points are colored by New.Annotation.   
                                      
                                       
                                  
                                
                                
                                  
                                       
                                      
                                           G beta.CL.gamma signalling through PI3Kgamma measurements vs. other pathway scores   More Plot Information                      G beta.CL.gamma signalling through PI3Kgamma measurements vs. other pathway scores  Each panel plots G beta.CL.gamma signalling through PI3Kgamma against another pathway score.  Points are colored by New.Annotation.   
                                      
                                       
                                  
                                
                                
                                  
                                       
                                      
                                           GAB1 signalosome measurements vs. other pathway scores   More Plot Information                      GAB1 signalosome measurements vs. other pathway scores  Each panel plots GAB1 signalosome against another pathway score.  Points are colored by New.Annotation.   
                                      
                                       
                                  
                                
                                
                                  
                                       
                                      
                                           Gastrin-CREB signalling pathway via PKC and MAPK measurements vs. other pathway scores   More Plot Information                      Gastrin-CREB signalling pathway via PKC and MAPK measurements vs. other pathway scores  Each panel plots Gastrin-CREB signalling pathway via PKC and MAPK against another pathway score.  Points are colored by New.Annotation.   
                                      
                                       
                                  
                                
                                
                                  
                                       
                                      
                                           GPVI-mediated activation cascade measurements vs. other pathway scores   More Plot Information                      GPVI-mediated activation cascade measurements vs. other pathway scores  Each panel plots GPVI-mediated activation cascade against another pathway score.  Points are colored by New.Annotation.   
                                      
                                       
                                  
                                
                                
                                  
                                       
                                      
                                           Integrin alphaIIb beta3 signaling measurements vs. other pathway scores   More Plot Information                      Integrin alphaIIb beta3 signaling measurements vs. other pathway scores  Each panel plots Integrin alphaIIb beta3 signaling against another pathway score.  Points are colored by New.Annotation.   
                                      
                                       
                                  
                                
                                
                                  
                                       
                                      
                                           Integrin cell surface interactions measurements vs. other pathway scores   More Plot Information                      Integrin cell surface interactions measurements vs. other pathway scores  Each panel plots Integrin cell surface interactions against another pathway score.  Points are colored by New.Annotation.   
                                      
                                       
                                  
                                
                                
                                  
                                       
                                      
                                           Interleukin receptor SHC signaling measurements vs. other pathway scores   More Plot Information                      Interleukin receptor SHC signaling measurements vs. other pathway scores  Each panel plots Interleukin receptor SHC signaling against another pathway score.  Points are colored by New.Annotation.   
                                      
                                       
                                  
                                
                                
                                  
                                       
                                      
                                           IRS-related events triggered by IGF1R measurements vs. other pathway scores   More Plot Information                      IRS-related events triggered by IGF1R measurements vs. other pathway scores  Each panel plots IRS-related events triggered by IGF1R against another pathway score.  Points are colored by New.Annotation.   
                                      
                                       
                                  
                                
                                
                                  
                                       
                                      
                                           Laminin interactions measurements vs. other pathway scores   More Plot Information                      Laminin interactions measurements vs. other pathway scores  Each panel plots Laminin interactions against another pathway score.  Points are colored by New.Annotation.   
                                      
                                       
                                  
                                
                                
                                  
                                       
                                      
                                           MAP2K and MAPK activation measurements vs. other pathway scores   More Plot Information                      MAP2K and MAPK activation measurements vs. other pathway scores  Each panel plots MAP2K and MAPK activation against another pathway score.  Points are colored by New.Annotation.   
                                      
                                       
                                  
                                
                                
                                  
                                       
                                      
                                           MAPK6MAPK4 signaling measurements vs. other pathway scores   More Plot Information                      MAPK6MAPK4 signaling measurements vs. other pathway scores  Each panel plots MAPK6MAPK4 signaling against another pathway score.  Points are colored by New.Annotation.   
                                      
                                       
                                  
                                
                                
                                  
                                       
                                      
                                           MicroRNA (miRNA) biogenesis measurements vs. other pathway scores   More Plot Information                      MicroRNA (miRNA) biogenesis measurements vs. other pathway scores  Each panel plots MicroRNA (miRNA) biogenesis against another pathway score.  Points are colored by New.Annotation.   
                                      
                                       
                                  
                                
                                
                                  
                                       
                                      
                                           Misspliced GSK3beta mutants stabilize beta-catenin measurements vs. other pathway scores   More Plot Information                      Misspliced GSK3beta mutants stabilize beta-catenin measurements vs. other pathway scores  Each panel plots Misspliced GSK3beta mutants stabilize beta-catenin against another pathway score.  Points are colored by New.Annotation.   
                                      
                                       
                                  
                                
                                
                                  
                                       
                                      
                                           Molecules associated with elastic fibres measurements vs. other pathway scores   More Plot Information                      Molecules associated with elastic fibres measurements vs. other pathway scores  Each panel plots Molecules associated with elastic fibres against another pathway score.  Points are colored by New.Annotation.   
                                      
                                       
                                  
                                
                                
                                  
                                       
                                      
                                           Negative regulation of FGFR1 signaling measurements vs. other pathway scores   More Plot Information                      Negative regulation of FGFR1 signaling measurements vs. other pathway scores  Each panel plots Negative regulation of FGFR1 signaling against another pathway score.  Points are colored by New.Annotation.   
                                      
                                       
                                  
                                
                                
                                  
                                       
                                      
                                           Negative regulation of FGFR2 signaling measurements vs. other pathway scores   More Plot Information                      Negative regulation of FGFR2 signaling measurements vs. other pathway scores  Each panel plots Negative regulation of FGFR2 signaling against another pathway score.  Points are colored by New.Annotation.   
                                      
                                       
                                  
                                
                                
                                  
                                       
                                      
                                           Negative regulation of FGFR3 signaling measurements vs. other pathway scores   More Plot Information                      Negative regulation of FGFR3 signaling measurements vs. other pathway scores  Each panel plots Negative regulation of FGFR3 signaling against another pathway score.  Points are colored by New.Annotation.   
                                      
                                       
                                  
                                
                                
                                  
                                       
                                      
                                           Negative regulation of FGFR4 signaling measurements vs. other pathway scores   More Plot Information                      Negative regulation of FGFR4 signaling measurements vs. other pathway scores  Each panel plots Negative regulation of FGFR4 signaling against another pathway score.  Points are colored by New.Annotation.   
                                      
                                       
                                  
                                
                                
                                  
                                       
                                      
                                           Negative regulation of MAPK pathway measurements vs. other pathway scores   More Plot Information                      Negative regulation of MAPK pathway measurements vs. other pathway scores  Each panel plots Negative regulation of MAPK pathway against another pathway score.  Points are colored by New.Annotation.   
                                      
                                       
                                  
                                
                                
                                  
                                       
                                      
                                           Nephrin interactions measurements vs. other pathway scores   More Plot Information                      Nephrin interactions measurements vs. other pathway scores  Each panel plots Nephrin interactions against another pathway score.  Points are colored by New.Annotation.   
                                      
                                       
                                  
                                
                                
                                  
                                       
                                      
                                           Non-integrin membrane-ECM interactions measurements vs. other pathway scores   More Plot Information                      Non-integrin membrane-ECM interactions measurements vs. other pathway scores  Each panel plots Non-integrin membrane-ECM interactions against another pathway score.  Points are colored by New.Annotation.   
                                      
                                       
                                  
                                
                                
                                  
                                       
                                      
                                           NRAGE signals death through JNK measurements vs. other pathway scores   More Plot Information                      NRAGE signals death through JNK measurements vs. other pathway scores  Each panel plots NRAGE signals death through JNK against another pathway score.  Points are colored by New.Annotation.   
                                      
                                       
                                  
                                
                                
                                  
                                       
                                      
                                           PCPCE pathway measurements vs. other pathway scores   More Plot Information                      PCPCE pathway measurements vs. other pathway scores  Each panel plots PCPCE pathway against another pathway score.  Points are colored by New.Annotation.   
                                      
                                       
                                  
                                
                                
                                  
                                       
                                      
                                           PI3K Cascade measurements vs. other pathway scores   More Plot Information                      PI3K Cascade measurements vs. other pathway scores  Each panel plots PI3K Cascade against another pathway score.  Points are colored by New.Annotation.   
                                      
                                       
                                  
                                
                                
                                  
                                       
                                      
                                           PI3KAKT activation measurements vs. other pathway scores   More Plot Information                      PI3KAKT activation measurements vs. other pathway scores  Each panel plots PI3KAKT activation against another pathway score.  Points are colored by New.Annotation.   
                                      
                                       
                                  
                                
                                
                                  
                                       
                                      
                                           PIP3 activates AKT signaling measurements vs. other pathway scores   More Plot Information                      PIP3 activates AKT signaling measurements vs. other pathway scores  Each panel plots PIP3 activates AKT signaling against another pathway score.  Points are colored by New.Annotation.   
                                      
                                       
                                  
                                
                                
                                  
                                       
                                      
                                           Platelet degranulation  measurements vs. other pathway scores   More Plot Information                      Platelet degranulation  measurements vs. other pathway scores  Each panel plots Platelet degranulation  against another pathway score.  Points are colored by New.Annotation.   
                                      
                                       
                                  
                                
                                
                                  
                                       
                                      
                                           RAF activation measurements vs. other pathway scores   More Plot Information                      RAF activation measurements vs. other pathway scores  Each panel plots RAF activation against another pathway score.  Points are colored by New.Annotation.   
                                      
                                       
                                  
                                
                                
                                  
                                       
                                      
                                           RAFMAP kinase cascade measurements vs. other pathway scores   More Plot Information                      RAFMAP kinase cascade measurements vs. other pathway scores  Each panel plots RAFMAP kinase cascade against another pathway score.  Points are colored by New.Annotation.   
                                      
                                       
                                  
                                
                                
                                  
                                       
                                      
                                           Regulation of actin dynamics for phagocytic cup formation measurements vs. other pathway scores   More Plot Information                      Regulation of actin dynamics for phagocytic cup formation measurements vs. other pathway scores  Each panel plots Regulation of actin dynamics for phagocytic cup formation against another pathway score.  Points are colored by New.Annotation.   
                                      
                                       
                                  
                                
                                
                                  
                                       
                                      
                                           Regulation of HSF1-mediated heat shock response measurements vs. other pathway scores   More Plot Information                      Regulation of HSF1-mediated heat shock response measurements vs. other pathway scores  Each panel plots Regulation of HSF1-mediated heat shock response against another pathway score.  Points are colored by New.Annotation.   
                                      
                                       
                                  
                                
                                
                                  
                                       
                                      
                                           Regulation of signaling by CBL measurements vs. other pathway scores   More Plot Information                      Regulation of signaling by CBL measurements vs. other pathway scores  Each panel plots Regulation of signaling by CBL against another pathway score.  Points are colored by New.Annotation.   
                                      
                                       
                                  
                                
                                
                                  
                                       
                                      
                                           Rho GTPase cycle measurements vs. other pathway scores   More Plot Information                      Rho GTPase cycle measurements vs. other pathway scores  Each panel plots Rho GTPase cycle against another pathway score.  Points are colored by New.Annotation.   
                                      
                                       
                                  
                                
                                
                                  
                                       
                                      
                                           RHO GTPases activate CIT measurements vs. other pathway scores   More Plot Information                      RHO GTPases activate CIT measurements vs. other pathway scores  Each panel plots RHO GTPases activate CIT against another pathway score.  Points are colored by New.Annotation.   
                                      
                                       
                                  
                                
                                
                                  
                                       
                                      
                                           RHO GTPases Activate Formins measurements vs. other pathway scores   More Plot Information                      RHO GTPases Activate Formins measurements vs. other pathway scores  Each panel plots RHO GTPases Activate Formins against another pathway score.  Points are colored by New.Annotation.   
                                      
                                       
                                  
                                
                                
                                  
                                       
                                      
                                           RHO GTPases activate IQGAPs measurements vs. other pathway scores   More Plot Information                      RHO GTPases activate IQGAPs measurements vs. other pathway scores  Each panel plots RHO GTPases activate IQGAPs against another pathway score.  Points are colored by New.Annotation.   
                                      
                                       
                                  
                                
                                
                                  
                                       
                                      
                                           RHO GTPases activate PAKs measurements vs. other pathway scores   More Plot Information                      RHO GTPases activate PAKs measurements vs. other pathway scores  Each panel plots RHO GTPases activate PAKs against another pathway score.  Points are colored by New.Annotation.   
                                      
                                       
                                  
                                
                                
                                  
                                       
                                      
                                           RHO GTPases activate PKNs measurements vs. other pathway scores   More Plot Information                      RHO GTPases activate PKNs measurements vs. other pathway scores  Each panel plots RHO GTPases activate PKNs against another pathway score.  Points are colored by New.Annotation.   
                                      
                                       
                                  
                                
                                
                                  
                                       
                                      
                                           RHO GTPases Activate ROCKs measurements vs. other pathway scores   More Plot Information                      RHO GTPases Activate ROCKs measurements vs. other pathway scores  Each panel plots RHO GTPases Activate ROCKs against another pathway score.  Points are colored by New.Annotation.   
                                      
                                       
                                  
                                
                                
                                  
                                       
                                      
                                           RHO GTPases Activate WASPs and WAVEs measurements vs. other pathway scores   More Plot Information                      RHO GTPases Activate WASPs and WAVEs measurements vs. other pathway scores  Each panel plots RHO GTPases Activate WASPs and WAVEs against another pathway score.  Points are colored by New.Annotation.   
                                      
                                       
                                  
                                
                                
                                  
                                       
                                      
                                           Role of LAT2NTALLAB on calcium mobilization measurements vs. other pathway scores   More Plot Information                      Role of LAT2NTALLAB on calcium mobilization measurements vs. other pathway scores  Each panel plots Role of LAT2NTALLAB on calcium mobilization against another pathway score.  Points are colored by New.Annotation.   
                                      
                                       
                                  
                                
                                
                                  
                                       
                                      
                                           S33 mutants of beta-catenin aren.SQT.t phosphorylated measurements vs. other pathway scores   More Plot Information                      S33 mutants of beta-catenin aren.SQT.t phosphorylated measurements vs. other pathway scores  Each panel plots S33 mutants of beta-catenin aren.SQT.t phosphorylated against another pathway score.  Points are colored by New.Annotation.   
                                      
                                       
                                  
                                
                                
                                  
                                       
                                      
                                           S37 mutants of beta-catenin aren.SQT.t phosphorylated measurements vs. other pathway scores   More Plot Information                      S37 mutants of beta-catenin aren.SQT.t phosphorylated measurements vs. other pathway scores  Each panel plots S37 mutants of beta-catenin aren.SQT.t phosphorylated against another pathway score.  Points are colored by New.Annotation.   
                                      
                                       
                                  
                                
                                
                                  
                                       
                                      
                                           S45 mutants of beta-catenin aren.SQT.t phosphorylated measurements vs. other pathway scores   More Plot Information                      S45 mutants of beta-catenin aren.SQT.t phosphorylated measurements vs. other pathway scores  Each panel plots S45 mutants of beta-catenin aren.SQT.t phosphorylated against another pathway score.  Points are colored by New.Annotation.   
                                      
                                       
                                  
                                
                                
                                  
                                       
                                      
                                           Sema3A PAK dependent Axon repulsion measurements vs. other pathway scores   More Plot Information                      Sema3A PAK dependent Axon repulsion measurements vs. other pathway scores  Each panel plots Sema3A PAK dependent Axon repulsion against another pathway score.  Points are colored by New.Annotation.   
                                      
                                       
                                  
                                
                                
                                  
                                       
                                      
                                           Sema4D induced cell migration and growth-cone collapse measurements vs. other pathway scores   More Plot Information                      Sema4D induced cell migration and growth-cone collapse measurements vs. other pathway scores  Each panel plots Sema4D induced cell migration and growth-cone collapse against another pathway score.  Points are colored by New.Annotation.   
                                      
                                       
                                  
                                
                                
                                  
                                       
                                      
                                           SHC-mediated cascade.CL.FGFR1 measurements vs. other pathway scores   More Plot Information                      SHC-mediated cascade.CL.FGFR1 measurements vs. other pathway scores  Each panel plots SHC-mediated cascade.CL.FGFR1 against another pathway score.  Points are colored by New.Annotation.   
                                      
                                       
                                  
                                
                                
                                  
                                       
                                      
                                           SHC-related events triggered by IGF1R measurements vs. other pathway scores   More Plot Information                      SHC-related events triggered by IGF1R measurements vs. other pathway scores  Each panel plots SHC-related events triggered by IGF1R against another pathway score.  Points are colored by New.Annotation.   
                                      
                                       
                                  
                                
                                
                                  
                                       
                                      
                                           Signal transduction by L1 measurements vs. other pathway scores   More Plot Information                      Signal transduction by L1 measurements vs. other pathway scores  Each panel plots Signal transduction by L1 against another pathway score.  Points are colored by New.Annotation.   
                                      
                                       
                                  
                                
                                
                                  
                                       
                                      
                                           Signaling by ERBB2 measurements vs. other pathway scores   More Plot Information                      Signaling by ERBB2 measurements vs. other pathway scores  Each panel plots Signaling by ERBB2 against another pathway score.  Points are colored by New.Annotation.   
                                      
                                       
                                  
                                
                                
                                  
                                       
                                      
                                           Signaling by FGFR1 in disease measurements vs. other pathway scores   More Plot Information                      Signaling by FGFR1 in disease measurements vs. other pathway scores  Each panel plots Signaling by FGFR1 in disease against another pathway score.  Points are colored by New.Annotation.   
                                      
                                       
                                  
                                
                                
                                  
                                       
                                      
                                           Signaling by FGFR2 in disease measurements vs. other pathway scores   More Plot Information                      Signaling by FGFR2 in disease measurements vs. other pathway scores  Each panel plots Signaling by FGFR2 in disease against another pathway score.  Points are colored by New.Annotation.   
                                      
                                       
                                  
                                
                                
                                  
                                       
                                      
                                           Signaling by FGFR3 in disease measurements vs. other pathway scores   More Plot Information                      Signaling by FGFR3 in disease measurements vs. other pathway scores  Each panel plots Signaling by FGFR3 in disease against another pathway score.  Points are colored by New.Annotation.   
                                      
                                       
                                  
                                
                                
                                  
                                       
                                      
                                           Signaling by FGFR4 in disease measurements vs. other pathway scores   More Plot Information                      Signaling by FGFR4 in disease measurements vs. other pathway scores  Each panel plots Signaling by FGFR4 in disease against another pathway score.  Points are colored by New.Annotation.   
                                      
                                       
                                  
                                
                                
                                  
                                       
                                      
                                           Signaling by PDGF measurements vs. other pathway scores   More Plot Information                      Signaling by PDGF measurements vs. other pathway scores  Each panel plots Signaling by PDGF against another pathway score.  Points are colored by New.Annotation.   
                                      
                                       
                                  
                                
                                
                                  
                                       
                                      
                                           Signaling by SCF-KIT measurements vs. other pathway scores   More Plot Information                      Signaling by SCF-KIT measurements vs. other pathway scores  Each panel plots Signaling by SCF-KIT against another pathway score.  Points are colored by New.Annotation.   
                                      
                                       
                                  
                                
                                
                                  
                                       
                                      
                                           Smooth Muscle Contraction measurements vs. other pathway scores   More Plot Information                      Smooth Muscle Contraction measurements vs. other pathway scores  Each panel plots Smooth Muscle Contraction against another pathway score.  Points are colored by New.Annotation.   
                                      
                                       
                                  
                                
                                
                                  
                                       
                                      
                                           Spry regulation of FGF signaling measurements vs. other pathway scores   More Plot Information                      Spry regulation of FGF signaling measurements vs. other pathway scores  Each panel plots Spry regulation of FGF signaling against another pathway score.  Points are colored by New.Annotation.   
                                      
                                       
                                  
                                
                                
                                  
                                       
                                      
                                           Syndecan interactions measurements vs. other pathway scores   More Plot Information                      Syndecan interactions measurements vs. other pathway scores  Each panel plots Syndecan interactions against another pathway score.  Points are colored by New.Annotation.   
                                      
                                       
                                  
                                
                                
                                  
                                       
                                      
                                           Synthesis of PIPs at the plasma membrane measurements vs. other pathway scores   More Plot Information                      Synthesis of PIPs at the plasma membrane measurements vs. other pathway scores  Each panel plots Synthesis of PIPs at the plasma membrane against another pathway score.  Points are colored by New.Annotation.   
                                      
                                       
                                  
                                
                                
                                  
                                       
                                      
                                           T41 mutants of beta-catenin aren.SQT.t phosphorylated measurements vs. other pathway scores   More Plot Information                      T41 mutants of beta-catenin aren.SQT.t phosphorylated measurements vs. other pathway scores  Each panel plots T41 mutants of beta-catenin aren.SQT.t phosphorylated against another pathway score.  Points are colored by New.Annotation.   
                                      
                                       
                                  
                                
                                
                                  
                                       
                                      
                                           TGF-beta receptor signaling activates SMADs measurements vs. other pathway scores   More Plot Information                      TGF-beta receptor signaling activates SMADs measurements vs. other pathway scores  Each panel plots TGF-beta receptor signaling activates SMADs against another pathway score.  Points are colored by New.Annotation.   
                                      
                                       
                                  
                                
                                
                                  
                                       
                                      
                                           Tie2 Signaling measurements vs. other pathway scores   More Plot Information                      Tie2 Signaling measurements vs. other pathway scores  Each panel plots Tie2 Signaling against another pathway score.  Points are colored by New.Annotation.   
                                      
                                       
                                  
                                
                                
                                  
                                       
                                      
                                           Tight junction interactions measurements vs. other pathway scores   More Plot Information                      Tight junction interactions measurements vs. other pathway scores  Each panel plots Tight junction interactions against another pathway score.  Points are colored by New.Annotation.   
                                      
                                       
                                  
                                
                                
                                  
                                       
                                      
                                           Truncations of AMER1 destabilize the destruction complex measurements vs. other pathway scores   More Plot Information                      Truncations of AMER1 destabilize the destruction complex measurements vs. other pathway scores  Each panel plots Truncations of AMER1 destabilize the destruction complex against another pathway score.  Points are colored by New.Annotation.   
                                      
                                       
                                  
                                
                                
                                  
                                       
                                      
                                           VEGFA-VEGFR2 Pathway measurements vs. other pathway scores   More Plot Information                      VEGFA-VEGFR2 Pathway measurements vs. other pathway scores  Each panel plots VEGFA-VEGFR2 Pathway against another pathway score.  Points are colored by New.Annotation.   
                                      
                                       
                                  
                                
                                
                                  
                                       
                                      
                                           VEGFR2 mediated cell proliferation measurements vs. other pathway scores   More Plot Information                      VEGFR2 mediated cell proliferation measurements vs. other pathway scores  Each panel plots VEGFR2 mediated cell proliferation against another pathway score.  Points are colored by New.Annotation.   
                                      
                                       
                                  
                                
                                
                                  
                                       
                                      
                                           VEGFR2 mediated vascular permeability measurements vs. other pathway scores   More Plot Information                      VEGFR2 mediated vascular permeability measurements vs. other pathway scores  Each panel plots VEGFR2 mediated vascular permeability against another pathway score.  Points are colored by New.Annotation.   
                                      
                                       
                                  
                                
                              
                             
                          
                           
                      
                    
                  
                
                
                  
                    
                      
                           
                          
                               Choose a Pathway: 
                              
                                  Summary  
                                  Activation of Rac  
                                  Apoptotic cleavage of cellular proteins  
                                  ARMS-mediated activation  
                                  AXIN missense mutants destabilize the destruction complex  
                                  Beta-catenin phosphorylation cascade  
                                  CD209 (DC-SIGN) signaling  
                                  CD28 co-stimulation  
                                  CD28 dependent PI3KAkt signaling  
                                  CD28 dependent Vav1 pathway  
                                  Cell-extracellular matrix interactions  
                                  Cell surface interactions at the vascular wall  
                                  Circadian Clock  
                                  Constitutive Signaling by Aberrant PI3K in Cancer  
                                  Constitutive Signaling by EGFRvIII  
                                  CTLA4 inhibitory signaling  
                                  Cyclin D associated events in G1  
                                  DAP12 signaling  
                                  Deactivation of the beta-catenin transactivating complex  
                                  Disassembly of the destruction complex and recruitment of AXIN t...  
                                  Downstream signal transduction  
                                  Downstream TCR signaling  
                                  ECM proteoglycans  
                                  Elastic fibre formation  
                                  EPH-ephrin mediated repulsion of cells  
                                  EPHA-mediated growth cone collapse  
                                  EPHB-mediated forward signaling  
                                  Ephrin signaling  
                                  FCERI mediated MAPK activation  
                                  Formation of the beta-catenin.CL.TCF transactivating complex  
                                  Frs2-mediated activation  
                                  G alpha (1213) signalling events  
                                  G beta.CL.gamma signalling through PI3Kgamma  
                                  GAB1 signalosome  
                                  Gastrin-CREB signalling pathway via PKC and MAPK  
                                  GPVI-mediated activation cascade  
                                  Integrin alphaIIb beta3 signaling  
                                  Integrin cell surface interactions  
                                  Interleukin receptor SHC signaling  
                                  IRS-related events triggered by IGF1R  
                                  Laminin interactions  
                                  MAP2K and MAPK activation  
                                  MAPK6MAPK4 signaling  
                                  MicroRNA (miRNA) biogenesis  
                                  Misspliced GSK3beta mutants stabilize beta-catenin  
                                  Molecules associated with elastic fibres  
                                  Negative regulation of FGFR1 signaling  
                                  Negative regulation of FGFR2 signaling  
                                  Negative regulation of FGFR3 signaling  
                                  Negative regulation of FGFR4 signaling  
                                  Negative regulation of MAPK pathway  
                                  Nephrin interactions  
                                  Non-integrin membrane-ECM interactions  
                                  NRAGE signals death through JNK  
                                  PCPCE pathway  
                                  PI3K Cascade  
                                  PI3KAKT activation  
                                  PIP3 activates AKT signaling  
                                  Platelet degranulation   
                                  RAF activation  
                                  RAFMAP kinase cascade  
                                  Regulation of actin dynamics for phagocytic cup formation  
                                  Regulation of HSF1-mediated heat shock response  
                                  Regulation of signaling by CBL  
                                  Rho GTPase cycle  
                                  RHO GTPases activate CIT  
                                  RHO GTPases Activate Formins  
                                  RHO GTPases activate IQGAPs  
                                  RHO GTPases activate PAKs  
                                  RHO GTPases activate PKNs  
                                  RHO GTPases Activate ROCKs  
                                  RHO GTPases Activate WASPs and WAVEs  
                                  Role of LAT2NTALLAB on calcium mobilization  
                                  S33 mutants of beta-catenin aren.SQT.t phosphorylated  
                                  S37 mutants of beta-catenin aren.SQT.t phosphorylated  
                                  S45 mutants of beta-catenin aren.SQT.t phosphorylated  
                                  Sema3A PAK dependent Axon repulsion  
                                  Sema4D induced cell migration and growth-cone collapse  
                                  SHC-mediated cascade.CL.FGFR1  
                                  SHC-related events triggered by IGF1R  
                                  Signal transduction by L1  
                                  Signaling by ERBB2  
                                  Signaling by FGFR1 in disease  
                                  Signaling by FGFR2 in disease  
                                  Signaling by FGFR3 in disease  
                                  Signaling by FGFR4 in disease  
                                  Signaling by PDGF  
                                  Signaling by SCF-KIT  
                                  Smooth Muscle Contraction  
                                  Spry regulation of FGF signaling  
                                  Syndecan interactions  
                                  Synthesis of PIPs at the plasma membrane  
                                  T41 mutants of beta-catenin aren.SQT.t phosphorylated  
                                  TGF-beta receptor signaling activates SMADs  
                                  Tie2 Signaling  
                                  Tight junction interactions  
                                  Truncations of AMER1 destabilize the destruction complex  
                                  VEGFA-VEGFR2 Pathway  
                                  VEGFR2 mediated cell proliferation  
                                  VEGFR2 mediated vascular permeability  
                              
                               
                              
                                
                                  
                                       
                                      
                                           Pathway scores vs. New.Annotation   More Plot Information                Pathway scores vs. New.Annotation  Plots the pathway scores against New.Annotation. The pathway scores are plotted to show how they vary across different values of New.Annotation conditions. Lines show each pathway's average score across values of New.Annotation.   
                                      
                                       
                                  
                                
                                
                                  
                                       
                                      
                                           Activation of Rac  pathway scores vs. New.Annotation   More Plot Information                Activation of Rac  pathway scores vs. New.Annotation  Plots Activation of Rac pathway scores against New.Annotation.   
                                      
                                       
                                  
                                
                                
                                  
                                       
                                      
                                           Apoptotic cleavage of cellular proteins  pathway scores vs. New.Annotation   More Plot Information                Apoptotic cleavage of cellular proteins  pathway scores vs. New.Annotation  Plots Apoptotic cleavage of cellular proteins pathway scores against New.Annotation.   
                                      
                                       
                                  
                                
                                
                                  
                                       
                                      
                                           ARMS-mediated activation  pathway scores vs. New.Annotation   More Plot Information                ARMS-mediated activation  pathway scores vs. New.Annotation  Plots ARMS-mediated activation pathway scores against New.Annotation.   
                                      
                                       
                                  
                                
                                
                                  
                                       
                                      
                                           AXIN missense mutants destabilize the destruction complex  pathway scores vs. New.Annotation   More Plot Information                AXIN missense mutants destabilize the destruction complex  pathway scores vs. New.Annotation  Plots AXIN missense mutants destabilize the destruction complex pathway scores against New.Annotation.   
                                      
                                       
                                  
                                
                                
                                  
                                       
                                      
                                           Beta-catenin phosphorylation cascade  pathway scores vs. New.Annotation   More Plot Information                Beta-catenin phosphorylation cascade  pathway scores vs. New.Annotation  Plots Beta-catenin phosphorylation cascade pathway scores against New.Annotation.   
                                      
                                       
                                  
                                
                                
                                  
                                       
                                      
                                           CD209 (DC-SIGN) signaling  pathway scores vs. New.Annotation   More Plot Information                CD209 (DC-SIGN) signaling  pathway scores vs. New.Annotation  Plots CD209 (DC-SIGN) signaling pathway scores against New.Annotation.   
                                      
                                       
                                  
                                
                                
                                  
                                       
                                      
                                           CD28 co-stimulation  pathway scores vs. New.Annotation   More Plot Information                CD28 co-stimulation  pathway scores vs. New.Annotation  Plots CD28 co-stimulation pathway scores against New.Annotation.   
                                      
                                       
                                  
                                
                                
                                  
                                       
                                      
                                           CD28 dependent PI3KAkt signaling  pathway scores vs. New.Annotation   More Plot Information                CD28 dependent PI3KAkt signaling  pathway scores vs. New.Annotation  Plots CD28 dependent PI3KAkt signaling pathway scores against New.Annotation.   
                                      
                                       
                                  
                                
                                
                                  
                                       
                                      
                                           CD28 dependent Vav1 pathway  pathway scores vs. New.Annotation   More Plot Information                CD28 dependent Vav1 pathway  pathway scores vs. New.Annotation  Plots CD28 dependent Vav1 pathway pathway scores against New.Annotation.   
                                      
                                       
                                  
                                
                                
                                  
                                       
                                      
                                           Cell-extracellular matrix interactions  pathway scores vs. New.Annotation   More Plot Information                Cell-extracellular matrix interactions  pathway scores vs. New.Annotation  Plots Cell-extracellular matrix interactions pathway scores against New.Annotation.   
                                      
                                       
                                  
                                
                                
                                  
                                       
                                      
                                           Cell surface interactions at the vascular wall  pathway scores vs. New.Annotation   More Plot Information                Cell surface interactions at the vascular wall  pathway scores vs. New.Annotation  Plots Cell surface interactions at the vascular wall pathway scores against New.Annotation.   
                                      
                                       
                                  
                                
                                
                                  
                                       
                                      
                                           Circadian Clock  pathway scores vs. New.Annotation   More Plot Information                Circadian Clock  pathway scores vs. New.Annotation  Plots Circadian Clock pathway scores against New.Annotation.   
                                      
                                       
                                  
                                
                                
                                  
                                       
                                      
                                           Constitutive Signaling by Aberrant PI3K in Cancer  pathway scores vs. New.Annotation   More Plot Information                Constitutive Signaling by Aberrant PI3K in Cancer  pathway scores vs. New.Annotation  Plots Constitutive Signaling by Aberrant PI3K in Cancer pathway scores against New.Annotation.   
                                      
                                       
                                  
                                
                                
                                  
                                       
                                      
                                           Constitutive Signaling by EGFRvIII  pathway scores vs. New.Annotation   More Plot Information                Constitutive Signaling by EGFRvIII  pathway scores vs. New.Annotation  Plots Constitutive Signaling by EGFRvIII pathway scores against New.Annotation.   
                                      
                                       
                                  
                                
                                
                                  
                                       
                                      
                                           CTLA4 inhibitory signaling  pathway scores vs. New.Annotation   More Plot Information                CTLA4 inhibitory signaling  pathway scores vs. New.Annotation  Plots CTLA4 inhibitory signaling pathway scores against New.Annotation.   
                                      
                                       
                                  
                                
                                
                                  
                                       
                                      
                                           Cyclin D associated events in G1  pathway scores vs. New.Annotation   More Plot Information                Cyclin D associated events in G1  pathway scores vs. New.Annotation  Plots Cyclin D associated events in G1 pathway scores against New.Annotation.   
                                      
                                       
                                  
                                
                                
                                  
                                       
                                      
                                           DAP12 signaling  pathway scores vs. New.Annotation   More Plot Information                DAP12 signaling  pathway scores vs. New.Annotation  Plots DAP12 signaling pathway scores against New.Annotation.   
                                      
                                       
                                  
                                
                                
                                  
                                       
                                      
                                           Deactivation of the beta-catenin transactivating complex  pathway scores vs. New.Annotation   More Plot Information                Deactivation of the beta-catenin transactivating complex  pathway scores vs. New.Annotation  Plots Deactivation of the beta-catenin transactivating complex pathway scores against New.Annotation.   
                                      
                                       
                                  
                                
                                
                                  
                                       
                                      
                                           Disassembly of the destruction complex and recruitment of AXIN to the membrane  pathway scores vs. New.Annotation   More Plot Information                Disassembly of the destruction complex and recruitment of AXIN to the membrane  pathway scores vs. New.Annotation  Plots Disassembly of the destruction complex and recruitment of AXIN to the membrane pathway scores against New.Annotation.   
                                      
                                       
                                  
                                
                                
                                  
                                       
                                      
                                           Downstream signal transduction  pathway scores vs. New.Annotation   More Plot Information                Downstream signal transduction  pathway scores vs. New.Annotation  Plots Downstream signal transduction pathway scores against New.Annotation.   
                                      
                                       
                                  
                                
                                
                                  
                                       
                                      
                                           Downstream TCR signaling  pathway scores vs. New.Annotation   More Plot Information                Downstream TCR signaling  pathway scores vs. New.Annotation  Plots Downstream TCR signaling pathway scores against New.Annotation.   
                                      
                                       
                                  
                                
                                
                                  
                                       
                                      
                                           ECM proteoglycans  pathway scores vs. New.Annotation   More Plot Information                ECM proteoglycans  pathway scores vs. New.Annotation  Plots ECM proteoglycans pathway scores against New.Annotation.   
                                      
                                       
                                  
                                
                                
                                  
                                       
                                      
                                           Elastic fibre formation  pathway scores vs. New.Annotation   More Plot Information                Elastic fibre formation  pathway scores vs. New.Annotation  Plots Elastic fibre formation pathway scores against New.Annotation.   
                                      
                                       
                                  
                                
                                
                                  
                                       
                                      
                                           EPH-ephrin mediated repulsion of cells  pathway scores vs. New.Annotation   More Plot Information                EPH-ephrin mediated repulsion of cells  pathway scores vs. New.Annotation  Plots EPH-ephrin mediated repulsion of cells pathway scores against New.Annotation.   
                                      
                                       
                                  
                                
                                
                                  
                                       
                                      
                                           EPHA-mediated growth cone collapse  pathway scores vs. New.Annotation   More Plot Information                EPHA-mediated growth cone collapse  pathway scores vs. New.Annotation  Plots EPHA-mediated growth cone collapse pathway scores against New.Annotation.   
                                      
                                       
                                  
                                
                                
                                  
                                       
                                      
                                           EPHB-mediated forward signaling  pathway scores vs. New.Annotation   More Plot Information                EPHB-mediated forward signaling  pathway scores vs. New.Annotation  Plots EPHB-mediated forward signaling pathway scores against New.Annotation.   
                                      
                                       
                                  
                                
                                
                                  
                                       
                                      
                                           Ephrin signaling  pathway scores vs. New.Annotation   More Plot Information                Ephrin signaling  pathway scores vs. New.Annotation  Plots Ephrin signaling pathway scores against New.Annotation.   
                                      
                                       
                                  
                                
                                
                                  
                                       
                                      
                                           FCERI mediated MAPK activation  pathway scores vs. New.Annotation   More Plot Information                FCERI mediated MAPK activation  pathway scores vs. New.Annotation  Plots FCERI mediated MAPK activation pathway scores against New.Annotation.   
                                      
                                       
                                  
                                
                                
                                  
                                       
                                      
                                           Formation of the beta-catenin.CL.TCF transactivating complex  pathway scores vs. New.Annotation   More Plot Information                Formation of the beta-catenin.CL.TCF transactivating complex  pathway scores vs. New.Annotation  Plots Formation of the beta-catenin.CL.TCF transactivating complex pathway scores against New.Annotation.   
                                      
                                       
                                  
                                
                                
                                  
                                       
                                      
                                           Frs2-mediated activation  pathway scores vs. New.Annotation   More Plot Information                Frs2-mediated activation  pathway scores vs. New.Annotation  Plots Frs2-mediated activation pathway scores against New.Annotation.   
                                      
                                       
                                  
                                
                                
                                  
                                       
                                      
                                           G alpha (1213) signalling events  pathway scores vs. New.Annotation   More Plot Information                G alpha (1213) signalling events  pathway scores vs. New.Annotation  Plots G alpha (1213) signalling events pathway scores against New.Annotation.   
                                      
                                       
                                  
                                
                                
                                  
                                       
                                      
                                           G beta.CL.gamma signalling through PI3Kgamma  pathway scores vs. New.Annotation   More Plot Information                G beta.CL.gamma signalling through PI3Kgamma  pathway scores vs. New.Annotation  Plots G beta.CL.gamma signalling through PI3Kgamma pathway scores against New.Annotation.   
                                      
                                       
                                  
                                
                                
                                  
                                       
                                      
                                           GAB1 signalosome  pathway scores vs. New.Annotation   More Plot Information                GAB1 signalosome  pathway scores vs. New.Annotation  Plots GAB1 signalosome pathway scores against New.Annotation.   
                                      
                                       
                                  
                                
                                
                                  
                                       
                                      
                                           Gastrin-CREB signalling pathway via PKC and MAPK  pathway scores vs. New.Annotation   More Plot Information                Gastrin-CREB signalling pathway via PKC and MAPK  pathway scores vs. New.Annotation  Plots Gastrin-CREB signalling pathway via PKC and MAPK pathway scores against New.Annotation.   
                                      
                                       
                                  
                                
                                
                                  
                                       
                                      
                                           GPVI-mediated activation cascade  pathway scores vs. New.Annotation   More Plot Information                GPVI-mediated activation cascade  pathway scores vs. New.Annotation  Plots GPVI-mediated activation cascade pathway scores against New.Annotation.   
                                      
                                       
                                  
                                
                                
                                  
                                       
                                      
                                           Integrin alphaIIb beta3 signaling  pathway scores vs. New.Annotation   More Plot Information                Integrin alphaIIb beta3 signaling  pathway scores vs. New.Annotation  Plots Integrin alphaIIb beta3 signaling pathway scores against New.Annotation.   
                                      
                                       
                                  
                                
                                
                                  
                                       
                                      
                                           Integrin cell surface interactions  pathway scores vs. New.Annotation   More Plot Information                Integrin cell surface interactions  pathway scores vs. New.Annotation  Plots Integrin cell surface interactions pathway scores against New.Annotation.   
                                      
                                       
                                  
                                
                                
                                  
                                       
                                      
                                           Interleukin receptor SHC signaling  pathway scores vs. New.Annotation   More Plot Information                Interleukin receptor SHC signaling  pathway scores vs. New.Annotation  Plots Interleukin receptor SHC signaling pathway scores against New.Annotation.   
                                      
                                       
                                  
                                
                                
                                  
                                       
                                      
                                           IRS-related events triggered by IGF1R  pathway scores vs. New.Annotation   More Plot Information                IRS-related events triggered by IGF1R  pathway scores vs. New.Annotation  Plots IRS-related events triggered by IGF1R pathway scores against New.Annotation.   
                                      
                                       
                                  
                                
                                
                                  
                                       
                                      
                                           Laminin interactions  pathway scores vs. New.Annotation   More Plot Information                Laminin interactions  pathway scores vs. New.Annotation  Plots Laminin interactions pathway scores against New.Annotation.   
                                      
                                       
                                  
                                
                                
                                  
                                       
                                      
                                           MAP2K and MAPK activation  pathway scores vs. New.Annotation   More Plot Information                MAP2K and MAPK activation  pathway scores vs. New.Annotation  Plots MAP2K and MAPK activation pathway scores against New.Annotation.   
                                      
                                       
                                  
                                
                                
                                  
                                       
                                      
                                           MAPK6MAPK4 signaling  pathway scores vs. New.Annotation   More Plot Information                MAPK6MAPK4 signaling  pathway scores vs. New.Annotation  Plots MAPK6MAPK4 signaling pathway scores against New.Annotation.   
                                      
                                       
                                  
                                
                                
                                  
                                       
                                      
                                           MicroRNA (miRNA) biogenesis  pathway scores vs. New.Annotation   More Plot Information                MicroRNA (miRNA) biogenesis  pathway scores vs. New.Annotation  Plots MicroRNA (miRNA) biogenesis pathway scores against New.Annotation.   
                                      
                                       
                                  
                                
                                
                                  
                                       
                                      
                                           Misspliced GSK3beta mutants stabilize beta-catenin  pathway scores vs. New.Annotation   More Plot Information                Misspliced GSK3beta mutants stabilize beta-catenin  pathway scores vs. New.Annotation  Plots Misspliced GSK3beta mutants stabilize beta-catenin pathway scores against New.Annotation.   
                                      
                                       
                                  
                                
                                
                                  
                                       
                                      
                                           Molecules associated with elastic fibres  pathway scores vs. New.Annotation   More Plot Information                Molecules associated with elastic fibres  pathway scores vs. New.Annotation  Plots Molecules associated with elastic fibres pathway scores against New.Annotation.   
                                      
                                       
                                  
                                
                                
                                  
                                       
                                      
                                           Negative regulation of FGFR1 signaling  pathway scores vs. New.Annotation   More Plot Information                Negative regulation of FGFR1 signaling  pathway scores vs. New.Annotation  Plots Negative regulation of FGFR1 signaling pathway scores against New.Annotation.   
                                      
                                       
                                  
                                
                                
                                  
                                       
                                      
                                           Negative regulation of FGFR2 signaling  pathway scores vs. New.Annotation   More Plot Information                Negative regulation of FGFR2 signaling  pathway scores vs. New.Annotation  Plots Negative regulation of FGFR2 signaling pathway scores against New.Annotation.   
                                      
                                       
                                  
                                
                                
                                  
                                       
                                      
                                           Negative regulation of FGFR3 signaling  pathway scores vs. New.Annotation   More Plot Information                Negative regulation of FGFR3 signaling  pathway scores vs. New.Annotation  Plots Negative regulation of FGFR3 signaling pathway scores against New.Annotation.   
                                      
                                       
                                  
                                
                                
                                  
                                       
                                      
                                           Negative regulation of FGFR4 signaling  pathway scores vs. New.Annotation   More Plot Information                Negative regulation of FGFR4 signaling  pathway scores vs. New.Annotation  Plots Negative regulation of FGFR4 signaling pathway scores against New.Annotation.   
                                      
                                       
                                  
                                
                                
                                  
                                       
                                      
                                           Negative regulation of MAPK pathway  pathway scores vs. New.Annotation   More Plot Information                Negative regulation of MAPK pathway  pathway scores vs. New.Annotation  Plots Negative regulation of MAPK pathway pathway scores against New.Annotation.   
                                      
                                       
                                  
                                
                                
                                  
                                       
                                      
                                           Nephrin interactions  pathway scores vs. New.Annotation   More Plot Information                Nephrin interactions  pathway scores vs. New.Annotation  Plots Nephrin interactions pathway scores against New.Annotation.   
                                      
                                       
                                  
                                
                                
                                  
                                       
                                      
                                           Non-integrin membrane-ECM interactions  pathway scores vs. New.Annotation   More Plot Information                Non-integrin membrane-ECM interactions  pathway scores vs. New.Annotation  Plots Non-integrin membrane-ECM interactions pathway scores against New.Annotation.   
                                      
                                       
                                  
                                
                                
                                  
                                       
                                      
                                           NRAGE signals death through JNK  pathway scores vs. New.Annotation   More Plot Information                NRAGE signals death through JNK  pathway scores vs. New.Annotation  Plots NRAGE signals death through JNK pathway scores against New.Annotation.   
                                      
                                       
                                  
                                
                                
                                  
                                       
                                      
                                           PCPCE pathway  pathway scores vs. New.Annotation   More Plot Information                PCPCE pathway  pathway scores vs. New.Annotation  Plots PCPCE pathway pathway scores against New.Annotation.   
                                      
                                       
                                  
                                
                                
                                  
                                       
                                      
                                           PI3K Cascade  pathway scores vs. New.Annotation   More Plot Information                PI3K Cascade  pathway scores vs. New.Annotation  Plots PI3K Cascade pathway scores against New.Annotation.   
                                      
                                       
                                  
                                
                                
                                  
                                       
                                      
                                           PI3KAKT activation  pathway scores vs. New.Annotation   More Plot Information                PI3KAKT activation  pathway scores vs. New.Annotation  Plots PI3KAKT activation pathway scores against New.Annotation.   
                                      
                                       
                                  
                                
                                
                                  
                                       
                                      
                                           PIP3 activates AKT signaling  pathway scores vs. New.Annotation   More Plot Information                PIP3 activates AKT signaling  pathway scores vs. New.Annotation  Plots PIP3 activates AKT signaling pathway scores against New.Annotation.   
                                      
                                       
                                  
                                
                                
                                  
                                       
                                      
                                           Platelet degranulation   pathway scores vs. New.Annotation   More Plot Information                Platelet degranulation   pathway scores vs. New.Annotation  Plots Platelet degranulation  pathway scores against New.Annotation.   
                                      
                                       
                                  
                                
                                
                                  
                                       
                                      
                                           RAF activation  pathway scores vs. New.Annotation   More Plot Information                RAF activation  pathway scores vs. New.Annotation  Plots RAF activation pathway scores against New.Annotation.   
                                      
                                       
                                  
                                
                                
                                  
                                       
                                      
                                           RAFMAP kinase cascade  pathway scores vs. New.Annotation   More Plot Information                RAFMAP kinase cascade  pathway scores vs. New.Annotation  Plots RAFMAP kinase cascade pathway scores against New.Annotation.   
                                      
                                       
                                  
                                
                                
                                  
                                       
                                      
                                           Regulation of actin dynamics for phagocytic cup formation  pathway scores vs. New.Annotation   More Plot Information                Regulation of actin dynamics for phagocytic cup formation  pathway scores vs. New.Annotation  Plots Regulation of actin dynamics for phagocytic cup formation pathway scores against New.Annotation.   
                                      
                                       
                                  
                                
                                
                                  
                                       
                                      
                                           Regulation of HSF1-mediated heat shock response  pathway scores vs. New.Annotation   More Plot Information                Regulation of HSF1-mediated heat shock response  pathway scores vs. New.Annotation  Plots Regulation of HSF1-mediated heat shock response pathway scores against New.Annotation.   
                                      
                                       
                                  
                                
                                
                                  
                                       
                                      
                                           Regulation of signaling by CBL  pathway scores vs. New.Annotation   More Plot Information                Regulation of signaling by CBL  pathway scores vs. New.Annotation  Plots Regulation of signaling by CBL pathway scores against New.Annotation.   
                                      
                                       
                                  
                                
                                
                                  
                                       
                                      
                                           Rho GTPase cycle  pathway scores vs. New.Annotation   More Plot Information                Rho GTPase cycle  pathway scores vs. New.Annotation  Plots Rho GTPase cycle pathway scores against New.Annotation.   
                                      
                                       
                                  
                                
                                
                                  
                                       
                                      
                                           RHO GTPases activate CIT  pathway scores vs. New.Annotation   More Plot Information                RHO GTPases activate CIT  pathway scores vs. New.Annotation  Plots RHO GTPases activate CIT pathway scores against New.Annotation.   
                                      
                                       
                                  
                                
                                
                                  
                                       
                                      
                                           RHO GTPases Activate Formins  pathway scores vs. New.Annotation   More Plot Information                RHO GTPases Activate Formins  pathway scores vs. New.Annotation  Plots RHO GTPases Activate Formins pathway scores against New.Annotation.   
                                      
                                       
                                  
                                
                                
                                  
                                       
                                      
                                           RHO GTPases activate IQGAPs  pathway scores vs. New.Annotation   More Plot Information                RHO GTPases activate IQGAPs  pathway scores vs. New.Annotation  Plots RHO GTPases activate IQGAPs pathway scores against New.Annotation.   
                                      
                                       
                                  
                                
                                
                                  
                                       
                                      
                                           RHO GTPases activate PAKs  pathway scores vs. New.Annotation   More Plot Information                RHO GTPases activate PAKs  pathway scores vs. New.Annotation  Plots RHO GTPases activate PAKs pathway scores against New.Annotation.   
                                      
                                       
                                  
                                
                                
                                  
                                       
                                      
                                           RHO GTPases activate PKNs  pathway scores vs. New.Annotation   More Plot Information                RHO GTPases activate PKNs  pathway scores vs. New.Annotation  Plots RHO GTPases activate PKNs pathway scores against New.Annotation.   
                                      
                                       
                                  
                                
                                
                                  
                                       
                                      
                                           RHO GTPases Activate ROCKs  pathway scores vs. New.Annotation   More Plot Information                RHO GTPases Activate ROCKs  pathway scores vs. New.Annotation  Plots RHO GTPases Activate ROCKs pathway scores against New.Annotation.   
                                      
                                       
                                  
                                
                                
                                  
                                       
                                      
                                           RHO GTPases Activate WASPs and WAVEs  pathway scores vs. New.Annotation   More Plot Information                RHO GTPases Activate WASPs and WAVEs  pathway scores vs. New.Annotation  Plots RHO GTPases Activate WASPs and WAVEs pathway scores against New.Annotation.   
                                      
                                       
                                  
                                
                                
                                  
                                       
                                      
                                           Role of LAT2NTALLAB on calcium mobilization  pathway scores vs. New.Annotation   More Plot Information                Role of LAT2NTALLAB on calcium mobilization  pathway scores vs. New.Annotation  Plots Role of LAT2NTALLAB on calcium mobilization pathway scores against New.Annotation.   
                                      
                                       
                                  
                                
                                
                                  
                                       
                                      
                                           S33 mutants of beta-catenin aren.SQT.t phosphorylated  pathway scores vs. New.Annotation   More Plot Information                S33 mutants of beta-catenin aren.SQT.t phosphorylated  pathway scores vs. New.Annotation  Plots S33 mutants of beta-catenin aren.SQT.t phosphorylated pathway scores against New.Annotation.   
                                      
                                       
                                  
                                
                                
                                  
                                       
                                      
                                           S37 mutants of beta-catenin aren.SQT.t phosphorylated  pathway scores vs. New.Annotation   More Plot Information                S37 mutants of beta-catenin aren.SQT.t phosphorylated  pathway scores vs. New.Annotation  Plots S37 mutants of beta-catenin aren.SQT.t phosphorylated pathway scores against New.Annotation.   
                                      
                                       
                                  
                                
                                
                                  
                                       
                                      
                                           S45 mutants of beta-catenin aren.SQT.t phosphorylated  pathway scores vs. New.Annotation   More Plot Information                S45 mutants of beta-catenin aren.SQT.t phosphorylated  pathway scores vs. New.Annotation  Plots S45 mutants of beta-catenin aren.SQT.t phosphorylated pathway scores against New.Annotation.   
                                      
                                       
                                  
                                
                                
                                  
                                       
                                      
                                           Sema3A PAK dependent Axon repulsion  pathway scores vs. New.Annotation   More Plot Information                Sema3A PAK dependent Axon repulsion  pathway scores vs. New.Annotation  Plots Sema3A PAK dependent Axon repulsion pathway scores against New.Annotation.   
                                      
                                       
                                  
                                
                                
                                  
                                       
                                      
                                           Sema4D induced cell migration and growth-cone collapse  pathway scores vs. New.Annotation   More Plot Information                Sema4D induced cell migration and growth-cone collapse  pathway scores vs. New.Annotation  Plots Sema4D induced cell migration and growth-cone collapse pathway scores against New.Annotation.   
                                      
                                       
                                  
                                
                                
                                  
                                       
                                      
                                           SHC-mediated cascade.CL.FGFR1  pathway scores vs. New.Annotation   More Plot Information                SHC-mediated cascade.CL.FGFR1  pathway scores vs. New.Annotation  Plots SHC-mediated cascade.CL.FGFR1 pathway scores against New.Annotation.   
                                      
                                       
                                  
                                
                                
                                  
                                       
                                      
                                           SHC-related events triggered by IGF1R  pathway scores vs. New.Annotation   More Plot Information                SHC-related events triggered by IGF1R  pathway scores vs. New.Annotation  Plots SHC-related events triggered by IGF1R pathway scores against New.Annotation.   
                                      
                                       
                                  
                                
                                
                                  
                                       
                                      
                                           Signal transduction by L1  pathway scores vs. New.Annotation   More Plot Information                Signal transduction by L1  pathway scores vs. New.Annotation  Plots Signal transduction by L1 pathway scores against New.Annotation.   
                                      
                                       
                                  
                                
                                
                                  
                                       
                                      
                                           Signaling by ERBB2  pathway scores vs. New.Annotation   More Plot Information                Signaling by ERBB2  pathway scores vs. New.Annotation  Plots Signaling by ERBB2 pathway scores against New.Annotation.   
                                      
                                       
                                  
                                
                                
                                  
                                       
                                      
                                           Signaling by FGFR1 in disease  pathway scores vs. New.Annotation   More Plot Information                Signaling by FGFR1 in disease  pathway scores vs. New.Annotation  Plots Signaling by FGFR1 in disease pathway scores against New.Annotation.   
                                      
                                       
                                  
                                
                                
                                  
                                       
                                      
                                           Signaling by FGFR2 in disease  pathway scores vs. New.Annotation   More Plot Information                Signaling by FGFR2 in disease  pathway scores vs. New.Annotation  Plots Signaling by FGFR2 in disease pathway scores against New.Annotation.   
                                      
                                       
                                  
                                
                                
                                  
                                       
                                      
                                           Signaling by FGFR3 in disease  pathway scores vs. New.Annotation   More Plot Information                Signaling by FGFR3 in disease  pathway scores vs. New.Annotation  Plots Signaling by FGFR3 in disease pathway scores against New.Annotation.   
                                      
                                       
                                  
                                
                                
                                  
                                       
                                      
                                           Signaling by FGFR4 in disease  pathway scores vs. New.Annotation   More Plot Information                Signaling by FGFR4 in disease  pathway scores vs. New.Annotation  Plots Signaling by FGFR4 in disease pathway scores against New.Annotation.   
                                      
                                       
                                  
                                
                                
                                  
                                       
                                      
                                           Signaling by PDGF  pathway scores vs. New.Annotation   More Plot Information                Signaling by PDGF  pathway scores vs. New.Annotation  Plots Signaling by PDGF pathway scores against New.Annotation.   
                                      
                                       
                                  
                                
                                
                                  
                                       
                                      
                                           Signaling by SCF-KIT  pathway scores vs. New.Annotation   More Plot Information                Signaling by SCF-KIT  pathway scores vs. New.Annotation  Plots Signaling by SCF-KIT pathway scores against New.Annotation.   
                                      
                                       
                                  
                                
                                
                                  
                                       
                                      
                                           Smooth Muscle Contraction  pathway scores vs. New.Annotation   More Plot Information                Smooth Muscle Contraction  pathway scores vs. New.Annotation  Plots Smooth Muscle Contraction pathway scores against New.Annotation.   
                                      
                                       
                                  
                                
                                
                                  
                                       
                                      
                                           Spry regulation of FGF signaling  pathway scores vs. New.Annotation   More Plot Information                Spry regulation of FGF signaling  pathway scores vs. New.Annotation  Plots Spry regulation of FGF signaling pathway scores against New.Annotation.   
                                      
                                       
                                  
                                
                                
                                  
                                       
                                      
                                           Syndecan interactions  pathway scores vs. New.Annotation   More Plot Information                Syndecan interactions  pathway scores vs. New.Annotation  Plots Syndecan interactions pathway scores against New.Annotation.   
                                      
                                       
                                  
                                
                                
                                  
                                       
                                      
                                           Synthesis of PIPs at the plasma membrane  pathway scores vs. New.Annotation   More Plot Information                Synthesis of PIPs at the plasma membrane  pathway scores vs. New.Annotation  Plots Synthesis of PIPs at the plasma membrane pathway scores against New.Annotation.   
                                      
                                       
                                  
                                
                                
                                  
                                       
                                      
                                           T41 mutants of beta-catenin aren.SQT.t phosphorylated  pathway scores vs. New.Annotation   More Plot Information                T41 mutants of beta-catenin aren.SQT.t phosphorylated  pathway scores vs. New.Annotation  Plots T41 mutants of beta-catenin aren.SQT.t phosphorylated pathway scores against New.Annotation.   
                                      
                                       
                                  
                                
                                
                                  
                                       
                                      
                                           TGF-beta receptor signaling activates SMADs  pathway scores vs. New.Annotation   More Plot Information                TGF-beta receptor signaling activates SMADs  pathway scores vs. New.Annotation  Plots TGF-beta receptor signaling activates SMADs pathway scores against New.Annotation.   
                                      
                                       
                                  
                                
                                
                                  
                                       
                                      
                                           Tie2 Signaling  pathway scores vs. New.Annotation   More Plot Information                Tie2 Signaling  pathway scores vs. New.Annotation  Plots Tie2 Signaling pathway scores against New.Annotation.   
                                      
                                       
                                  
                                
                                
                                  
                                       
                                      
                                           Tight junction interactions  pathway scores vs. New.Annotation   More Plot Information                Tight junction interactions  pathway scores vs. New.Annotation  Plots Tight junction interactions pathway scores against New.Annotation.   
                                      
                                       
                                  
                                
                                
                                  
                                       
                                      
                                           Truncations of AMER1 destabilize the destruction complex  pathway scores vs. New.Annotation   More Plot Information                Truncations of AMER1 destabilize the destruction complex  pathway scores vs. New.Annotation  Plots Truncations of AMER1 destabilize the destruction complex pathway scores against New.Annotation.   
                                      
                                       
                                  
                                
                                
                                  
                                       
                                      
                                           VEGFA-VEGFR2 Pathway  pathway scores vs. New.Annotation   More Plot Information                VEGFA-VEGFR2 Pathway  pathway scores vs. New.Annotation  Plots VEGFA-VEGFR2 Pathway pathway scores against New.Annotation.   
                                      
                                       
                                  
                                
                                
                                  
                                       
                                      
                                           VEGFR2 mediated cell proliferation  pathway scores vs. New.Annotation   More Plot Information                VEGFR2 mediated cell proliferation  pathway scores vs. New.Annotation  Plots VEGFR2 mediated cell proliferation pathway scores against New.Annotation.   
                                      
                                       
                                  
                                
                                
                                  
                                       
                                      
                                           VEGFR2 mediated vascular permeability  pathway scores vs. New.Annotation   More Plot Information                VEGFR2 mediated vascular permeability  pathway scores vs. New.Annotation  Plots VEGFR2 mediated vascular permeability pathway scores against New.Annotation.   
                                      
                                       
                                  
                                
                              
                             
                          
                           
                      
                    
                  
                
               
                
              
                
          
        
        
          
                Genes are tested for differential expression in response to each selected covariate. For each gene, a single linear 
  regression is fit using all selected covariates to predict expression. This approach eliminates confounding due to measured
  covariates and isolates the independent association of each covariate with gene expression, measuring each variable's
  association with a gene after holding all other variables constant.      
              
                  New.Annotation: 
                  
                      New.Annotation: differential expression in Control vs. baseline of cel67  
                      New.Annotation: differential expression in miR142 vs. baseline of cel67  
                      New.Annotation: differential expression in miR155 vs. baseline of cel67  
                      New.Annotation: differential expression in miR18b vs. baseline of cel67  
                      New.Annotation: differential expression in miR890 vs. baseline of cel67  
                  
                  
                  
                    
                      
                           
                          
                               Volcano Plot: New.Annotation: Control vs.cel67   More Plot Information                Volcano Plot: New.Annotation: Control vs.cel67  Volcano plot displaying each gene's -log10(p-value) and log2 fold change with the selected covariate. Highly statistically significant genes fall at the top of the plot above the horizontal lines, and highly differentially expressed genes fall to either side. Horizontal lines indicate various False Discovery Rate (FDR) thresholds or p-value thresholds if there is no adjustment to the p-values. Genes are colored if the resulting p-value is below the given FDR or p-value threshold. The 40 most statistically significant genes are labeled in the plot.   
                               DE Results: New.Annotation: Control vs.cel67   More Plot Information      Download CSV Data     
                               
                                 
                                 
                               
                               
                                 
                                   
                                   Log2 fold change 
                                   std error (log2) 
                                   Lower confidence limit (log2) 
                                   Upper confidence limit (log2) 
                                   Linear fold change 
                                   Lower confidence limit (linear) 
                                   Upper confidence limit (linear) 
                                   P-value 
                                   BY.p.value 
                                   method 
                                   Gene.sets 
                                   probe.ID 
                                 
                               
                               
                                 
                                   CSNK1A1-mRNA 
                                   0.608 
                                   0.0669 
                                   0.477 
                                   0.739 
                                   1.52 
                                   1.39 
                                   1.67 
                                   9.92e-09 
                                   1.32e-05 
                                   loglinear 
                                   AXIN missense mutants destabilize the destruction complex, Beta-catenin phosphorylation cascade, Disassembly of the destruction complex and recruitment of AXIN to the membrane, Misspliced GSK3beta mutants stabilize beta-catenin, S33 mutants of beta-catenin aren.SQT.t phosphorylated, S37 mutants of beta-catenin aren.SQT.t phosphorylated, S45 mutants of beta-catenin aren.SQT.t phosphorylated, T41 mutants of beta-catenin aren.SQT.t phosphorylated 
                                   NM_001892.4:2053 
                                 
                                 
                                   DVL3-mRNA 
                                   0.617 
                                   0.0789 
                                   0.462 
                                   0.771 
                                   1.53 
                                   1.38 
                                   1.71 
                                   1.19e-07 
                                   7.89e-05 
                                   loglinear 
                                   Disassembly of the destruction complex and recruitment of AXIN to the membrane, PCPCE pathway, RHO GTPases Activate Formins, WNT mediated activation of DVL 
                                   NM_004423.3:1012 
                                 
                                 
                                   RRAS2-mRNA 
                                   0.9 
                                   0.124 
                                   0.658 
                                   1.14 
                                   1.87 
                                   1.58 
                                   2.21 
                                   3.71e-07 
                                   0.000164 
                                   lm.nb 
                                   
                                   NM_001102669.2:1785 
                                 
                                 
                                   RAB8B-mRNA 
                                   -0.291 
                                   0.0462 
                                   -0.382 
                                   -0.201 
                                   0.817 
                                   0.768 
                                   0.87 
                                   3.01e-06 
                                   0.001 
                                   loglinear 
                                   
                                   NM_016530.2:820 
                                 
                                 
                                   NRAS-mRNA 
                                   -0.544 
                                   0.114 
                                   -0.767 
                                   -0.322 
                                   0.686 
                                   0.588 
                                   0.8 
                                   9.79e-05 
                                   0.026 
                                   lm.nb 
                                   ARMS-mediated activation, CD209 (DC-SIGN) signaling, Cell surface interactions at the vascular wall, Constitutive Signaling by EGFRvIII, DAP12 signaling, Downstream signal transduction, FCERI mediated MAPK activation, Frs2-mediated activation, Gastrin-CREB signalling pathway via PKC and MAPK, Interleukin receptor SHC signaling, IRS-related events triggered by IGF1R, MAP2K and MAPK activation, RAFMAP kinase cascade, SHC-mediated cascade.CL.FGFR1, SHC-related events triggered by IGF1R, Signaling by ERBB2, Signaling by FGFR1 in disease, Signaling by FGFR2 in disease, Signaling by FGFR3 in disease, Signaling by FGFR4 in disease, Signaling by PDGF, Signaling by SCF-KIT, Tie2 Signaling, VEGFA-VEGFR2 Pathway, VEGFR2 mediated cell proliferation 
                                   NM_002524.3:877 
                                 
                                 
                                   NCKAP1-mRNA 
                                   0.423 
                                   0.0908 
                                   0.245 
                                   0.601 
                                   1.34 
                                   1.19 
                                   1.52 
                                   0.000135 
                                   0.0286 
                                   loglinear 
                                   Regulation of actin dynamics for phagocytic cup formation, RHO GTPases Activate WASPs and WAVEs, VEGFA-VEGFR2 Pathway 
                                   NM_013436.3:2353 
                                 
                                 
                                   AGO2-mRNA 
                                   -0.568 
                                   0.124 
                                   -0.812 
                                   -0.325 
                                   0.674 
                                   0.57 
                                   0.798 
                                   0.000166 
                                   0.0286 
                                   lm.nb 
                                   DAP12 signaling, Downstream signal transduction, GAB1 signalosome, MicroRNA (miRNA) biogenesis, PI3KAKT activation, PIP3 activates AKT signaling, Role of LAT2NTALLAB on calcium mobilization, Signaling by ERBB2, Signaling by PDGF, Signaling by SCF-KIT 
                                   NM_012154.2:625 
                                 
                                 
                                   ITGA5-mRNA 
                                   -0.684 
                                   0.15 
                                   -0.978 
                                   -0.39 
                                   0.623 
                                   0.508 
                                   0.763 
                                   0.000172 
                                   0.0286 
                                   lm.nb 
                                   Cell surface interactions at the vascular wall, Elastic fibre formation, Integrin cell surface interactions, Signal transduction by L1 
                                   NM_002205.2:925 
                                 
                                 
                                   HRAS-mRNA 
                                   -0.593 
                                   0.144 
                                   -0.874 
                                   -0.311 
                                   0.663 
                                   0.545 
                                   0.806 
                                   0.000487 
                                   0.0718 
                                   loglinear 
                                   ARMS-mediated activation, CD209 (DC-SIGN) signaling, Cell surface interactions at the vascular wall, Constitutive Signaling by EGFRvIII, CREB phosphorylation through the activation of Ras, DAP12 signaling, Downstream signal transduction, FCERI mediated MAPK activation, Frs2-mediated activation, Gastrin-CREB signalling pathway via PKC and MAPK, Interleukin receptor SHC signaling, IRS-related events triggered by IGF1R, MAP2K and MAPK activation, RAFMAP kinase cascade, SHC-mediated cascade.CL.FGFR1, SHC-related events triggered by IGF1R, Signaling by ERBB2, Signaling by FGFR1 in disease, Signaling by FGFR2 in disease, Signaling by FGFR3 in disease, Signaling by FGFR4 in disease, Signaling by PDGF, Signaling by SCF-KIT, Tie2 Signaling, VEGFA-VEGFR2 Pathway, VEGFR2 mediated cell proliferation 
                                   NM_005343.2:396 
                                 
                                 
                                   EGFR-mRNA 
                                   -0.408 
                                   0.101 
                                   -0.605 
                                   -0.211 
                                   0.753 
                                   0.657 
                                   0.864 
                                   0.000564 
                                   0.0749 
                                   loglinear 
                                   ARMS-mediated activation, Constitutive Signaling by Aberrant PI3K in Cancer, Constitutive Signaling by EGFRvIII, DAP12 signaling, Downstream signal transduction, FCERI mediated MAPK activation, Frs2-mediated activation, GAB1 signalosome, Gastrin-CREB signalling pathway via PKC and MAPK, Interleukin receptor SHC signaling, IRS-related events triggered by IGF1R, PI3KAKT activation, Role of LAT2NTALLAB on calcium mobilization, Signaling by ERBB2, Signaling by PDGF, Signaling by SCF-KIT, VEGFA-VEGFR2 Pathway, VEGFR2 mediated cell proliferation 
                                   NM_201282.1:360 
                                 
                                 
                                   MAPK3-mRNA 
                                   0.205 
                                   0.0541 
                                   0.0991 
                                   0.311 
                                   1.15 
                                   1.07 
                                   1.24 
                                   0.00106 
                                   0.128 
                                   loglinear 
                                   ARMS-mediated activation, DAP12 signaling, Downstream signal transduction, ERKs are inactivated, FCERI mediated MAPK activation, Frs2-mediated activation, Gastrin-CREB signalling pathway via PKC and MAPK, Interleukin receptor SHC signaling, IRS-related events triggered by IGF1R, Negative feedback regulation of MAPK pathway, Negative regulation of FGFR3 signaling, Negative regulation of FGFR4 signaling, Negative regulation of MAPK pathway, RAFMAP kinase cascade, Regulation of HSF1-mediated heat shock response, RHO GTPases Activate WASPs and WAVEs, Signaling by ERBB2, Signaling by PDGF, Signaling by SCF-KIT, VEGFA-VEGFR2 Pathway, VEGFR2 mediated cell proliferation 
                                   NM_001040056.1:580 
                                 
                                 
                                   INADL-mRNA 
                                   0.477 
                                   0.129 
                                   0.223 
                                   0.73 
                                   1.39 
                                   1.17 
                                   1.66 
                                   0.00137 
                                   0.151 
                                   loglinear 
                                   
                                   XM_005270347.1:343 
                                 
                                 
                                   SMAD2-mRNA 
                                   0.235 
                                   0.0645 
                                   0.109 
                                   0.361 
                                   1.18 
                                   1.08 
                                   1.28 
                                   0.00151 
                                   0.155 
                                   loglinear 
                                   TGF-beta receptor signaling activates SMADs 
                                   NM_005901.5:1678 
                                 
                                 
                                   ABCF1-mRNA 
                                   -0.241 
                                   0.0732 
                                   -0.385 
                                   -0.0977 
                                   0.846 
                                   0.766 
                                   0.934 
                                   0.00345 
                                   0.328 
                                   loglinear 
                                   
                                   NM_001090.2:850 
                                 
                                 
                                   MYC-mRNA 
                                   -0.557 
                                   0.176 
                                   -0.902 
                                   -0.213 
                                   0.68 
                                   0.535 
                                   0.863 
                                   0.00464 
                                   0.381 
                                   lm.nb 
                                   Formation of the beta-catenin.CL.TCF transactivating complex 
                                   NM_002467.3:1610 
                                 
                                 
                                   TARBP2-mRNA 
                                   -0.294 
                                   0.0927 
                                   -0.475 
                                   -0.112 
                                   0.816 
                                   0.719 
                                   0.925 
                                   0.00464 
                                   0.381 
                                   loglinear 
                                   MicroRNA (miRNA) biogenesis 
                                   NM_134323.1:1060 
                                 
                                 
                                   SPRY2-mRNA 
                                   -0.566 
                                   0.18 
                                   -0.918 
                                   -0.213 
                                   0.676 
                                   0.529 
                                   0.863 
                                   0.00488 
                                   0.381 
                                   lm.nb 
                                   Negative regulation of FGFR1 signaling, Negative regulation of FGFR2 signaling, Negative regulation of FGFR3 signaling, Negative regulation of FGFR4 signaling, Spry regulation of FGF signaling 
                                   NM_005842.2:1481 
                                 
                                 
                                   AKT2-mRNA 
                                   -0.182 
                                   0.0603 
                                   -0.3 
                                   -0.0634 
                                   0.882 
                                   0.812 
                                   0.957 
                                   0.00666 
                                   0.468 
                                   loglinear 
                                   Activation of BAD and translocation to mitochondria , CD28 co-stimulation, DAP12 signaling, Deactivation of the beta-catenin transactivating complex, Downstream signal transduction, G beta.CL.gamma signalling through PI3Kgamma, GAB1 signalosome, GPVI-mediated activation cascade, IRS-related events triggered by IGF1R, PI3K Cascade, PI3KAKT activation, PIP3 activates AKT signaling, Role of LAT2NTALLAB on calcium mobilization, Signaling by ERBB2, Signaling by PDGF, Signaling by SCF-KIT, TP53 Regulates Metabolic Genes, VEGFA-VEGFR2 Pathway, VEGFR2 mediated vascular permeability 
                                   NM_001626.4:699 
                                 
                                 
                                   DIAPH3-mRNA 
                                   -0.542 
                                   0.18 
                                   -0.895 
                                   -0.189 
                                   0.687 
                                   0.538 
                                   0.877 
                                   0.0067 
                                   0.468 
                                   lm.nb 
                                   RHO GTPases Activate Formins 
                                   NM_001042517.1:1832 
                                 
                                 
                                   PPP1CB-mRNA 
                                   0.252 
                                   0.0858 
                                   0.084 
                                   0.42 
                                   1.19 
                                   1.06 
                                   1.34 
                                   0.00784 
                                   0.521 
                                   loglinear 
                                   Circadian Clock, RHO GTPases activate PKNs, RHO GTPases Activate ROCKs, TGF-beta receptor signaling activates SMADs 
                                   NM_002709.2:393 
                                 
                               
                               DE Results: New.Annotation: Control vs.cel67  Table presenting the most statistically significantly differentially expressed genes with the selected covariate. 'Estimated log fold-change' estimates a gene's differential expression. For categorical covariates, a gene is estimated to have 2^(log fold change) times its expression in baseline samples, holding all other variables in the analysis constant. If the covariate is continuous, for each unit increase in the selected covariate, a gene's expression is estimated to increase by 2^(log fold change)-fold, holding all other variables in the analysis constant. The 95% confidence interval for the log fold change is also presented, along with a p-value and an adjusted p-value or FDR if requested. Method column indicates the model used to estimate differential expression. Mixed negative binomial model uses the mle function to run the Wald test. The simplified negative binomial model uses the glm.nb function . The loglinear model uses the lm function.   
                                   
                          
                           
                      
                    
                    
                      
                           
                          
                               Volcano Plot: New.Annotation: miR142 vs.cel67   More Plot Information                Volcano Plot: New.Annotation: miR142 vs.cel67  Volcano plot displaying each gene's -log10(p-value) and log2 fold change with the selected covariate. Highly statistically significant genes fall at the top of the plot above the horizontal lines, and highly differentially expressed genes fall to either side. Horizontal lines indicate various False Discovery Rate (FDR) thresholds or p-value thresholds if there is no adjustment to the p-values. Genes are colored if the resulting p-value is below the given FDR or p-value threshold. The 40 most statistically significant genes are labeled in the plot.   
                               DE Results: New.Annotation: miR142 vs.cel67   More Plot Information      Download CSV Data     
                               
                                 
                                 
                               
                               
                                 
                                   
                                   Log2 fold change 
                                   std error (log2) 
                                   Lower confidence limit (log2) 
                                   Upper confidence limit (log2) 
                                   Linear fold change 
                                   Lower confidence limit (linear) 
                                   Upper confidence limit (linear) 
                                   P-value 
                                   BY.p.value 
                                   method 
                                   Gene.sets 
                                   probe.ID 
                                 
                               
                               
                                 
                                   RAC1-mRNA 
                                   -1.32 
                                   0.0675 
                                   -1.45 
                                   -1.19 
                                   0.401 
                                   0.366 
                                   0.439 
                                   5.98e-15 
                                   7.94e-12 
                                   loglinear 
                                   Activation of Rac, CD28 co-stimulation, CD28 dependent Vav1 pathway, DAP12 signaling, EPH-ephrin mediated repulsion of cells, Ephrin signaling, G alpha (1213) signalling events, GPVI-mediated activation cascade, MAPK6MAPK4 signaling, NRAGE signals death through JNK, PCPCE pathway, Regulation of actin dynamics for phagocytic cup formation, RHO GTPases activate CIT, RHO GTPases Activate Formins, RHO GTPases activate IQGAPs, RHO GTPases activate PAKs, RHO GTPases activate PKNs, RHO GTPases Activate WASPs and WAVEs, Sema3A PAK dependent Axon repulsion, Sema4D induced cell migration and growth-cone collapse, Signal transduction by L1, Signaling by SCF-KIT, VEGFA-VEGFR2 Pathway, VEGFR2 mediated vascular permeability 
                                   NM_198829.1:1250 
                                 
                                 
                                   YES1-mRNA 
                                   -1.3 
                                   0.0716 
                                   -1.44 
                                   -1.16 
                                   0.405 
                                   0.368 
                                   0.447 
                                   2.45e-14 
                                   1.63e-11 
                                   loglinear 
                                   CD28 co-stimulation, Cell surface interactions at the vascular wall, CTLA4 inhibitory signaling, EPH-ephrin mediated repulsion of cells, EPHA-mediated growth cone collapse, EPHB-mediated forward signaling, Regulation of signaling by CBL, Signaling by ERBB2, Signaling by SCF-KIT 
                                   NM_005433.3:265 
                                 
                                 
                                   WASL-mRNA 
                                   -1.61 
                                   0.1 
                                   -1.81 
                                   -1.41 
                                   0.328 
                                   0.286 
                                   0.375 
                                   2.83e-13 
                                   1.25e-10 
                                   loglinear 
                                   DCC mediated attractive signaling, EPHB-mediated forward signaling, Nephrin interactions, Regulation of actin dynamics for phagocytic cup formation, RHO GTPases Activate WASPs and WAVEs 
                                   NM_003941.2:400 
                                 
                                 
                                   DVL3-mRNA 
                                   0.877 
                                   0.0789 
                                   0.723 
                                   1.03 
                                   1.84 
                                   1.65 
                                   2.04 
                                   2.91e-10 
                                   9.65e-08 
                                   loglinear 
                                   Disassembly of the destruction complex and recruitment of AXIN to the membrane, PCPCE pathway, RHO GTPases Activate Formins, WNT mediated activation of DVL 
                                   NM_004423.3:1012 
                                 
                                 
                                   CSNK1A1-mRNA 
                                   0.698 
                                   0.0669 
                                   0.567 
                                   0.829 
                                   1.62 
                                   1.48 
                                   1.78 
                                   8.98e-10 
                                   2.38e-07 
                                   loglinear 
                                   AXIN missense mutants destabilize the destruction complex, Beta-catenin phosphorylation cascade, Disassembly of the destruction complex and recruitment of AXIN to the membrane, Misspliced GSK3beta mutants stabilize beta-catenin, S33 mutants of beta-catenin aren.SQT.t phosphorylated, S37 mutants of beta-catenin aren.SQT.t phosphorylated, S45 mutants of beta-catenin aren.SQT.t phosphorylated, T41 mutants of beta-catenin aren.SQT.t phosphorylated 
                                   NM_001892.4:2053 
                                 
                                 
                                   CTTN-mRNA 
                                   -1.02 
                                   0.109 
                                   -1.23 
                                   -0.802 
                                   0.494 
                                   0.426 
                                   0.573 
                                   6.67e-09 
                                   1.48e-06 
                                   loglinear 
                                   RHO GTPases activate PAKs 
                                   NM_001184740.1:240 
                                 
                                 
                                   CRK-mRNA 
                                   -0.918 
                                   0.103 
                                   -1.12 
                                   -0.716 
                                   0.529 
                                   0.46 
                                   0.609 
                                   1.38e-08 
                                   2.62e-06 
                                   loglinear 
                                   ARMS-mediated activation, Downstream signal transduction, Integrin alphaIIb beta3 signaling, p130Cas linkage to MAPK signaling for integrins, Regulation of actin dynamics for phagocytic cup formation, Regulation of signaling by CBL, Signaling by PDGF, VEGFA-VEGFR2 Pathway 
                                   NM_016823.2:1945 
                                 
                                 
                                   CASK-mRNA 
                                   -0.913 
                                   0.107 
                                   -1.12 
                                   -0.704 
                                   0.531 
                                   0.459 
                                   0.614 
                                   2.73e-08 
                                   4.54e-06 
                                   loglinear 
                                   Nephrin interactions, Non-integrin membrane-ECM interactions, Syndecan interactions 
                                   NM_001126054.2:2190 
                                 
                                 
                                   PTEN-mRNA 
                                   0.505 
                                   0.0618 
                                   0.384 
                                   0.626 
                                   1.42 
                                   1.3 
                                   1.54 
                                   5.89e-08 
                                   8.69e-06 
                                   loglinear 
                                   DAP12 signaling, Downstream signal transduction, Downstream TCR signaling, GAB1 signalosome, PI3KAKT activation, PIP3 activates AKT signaling, Role of LAT2NTALLAB on calcium mobilization, Signaling by ERBB2, Signaling by PDGF, Signaling by SCF-KIT, Synthesis of PIPs at the plasma membrane 
                                   NM_000314.4:1351 
                                 
                                 
                                   RRAS2-mRNA 
                                   0.986 
                                   0.124 
                                   0.743 
                                   1.23 
                                   1.98 
                                   1.67 
                                   2.34 
                                   8.85e-08 
                                   1.18e-05 
                                   lm.nb 
                                   
                                   NM_001102669.2:1785 
                                 
                                 
                                   GSK3B-mRNA 
                                   0.553 
                                   0.0708 
                                   0.414 
                                   0.691 
                                   1.47 
                                   1.33 
                                   1.61 
                                   1.2e-07 
                                   1.45e-05 
                                   loglinear 
                                   AXIN missense mutants destabilize the destruction complex, DAP12 signaling, Disassembly of the destruction complex and recruitment of AXIN to the membrane, Downstream signal transduction, GAB1 signalosome, Misspliced GSK3beta mutants stabilize beta-catenin, PI3KAKT activation, PIP3 activates AKT signaling, Regulation of HSF1-mediated heat shock response, Role of LAT2NTALLAB on calcium mobilization, S33 mutants of beta-catenin aren.SQT.t phosphorylated, S37 mutants of beta-catenin aren.SQT.t phosphorylated, S45 mutants of beta-catenin aren.SQT.t phosphorylated, Signaling by ERBB2, Signaling by PDGF, Signaling by SCF-KIT, T41 mutants of beta-catenin aren.SQT.t phosphorylated, Truncations of AMER1 destabilize the destruction complex 
                                   NM_002093.2:925 
                                 
                                 
                                   MYC-mRNA 
                                   -1.38 
                                   0.181 
                                   -1.74 
                                   -1.03 
                                   0.384 
                                   0.3 
                                   0.491 
                                   1.8e-07 
                                   1.99e-05 
                                   lm.nb 
                                   Formation of the beta-catenin.CL.TCF transactivating complex 
                                   NM_002467.3:1610 
                                 
                                 
                                   SMAD2-mRNA 
                                   0.473 
                                   0.0645 
                                   0.347 
                                   0.599 
                                   1.39 
                                   1.27 
                                   1.52 
                                   3.2e-07 
                                   3.27e-05 
                                   loglinear 
                                   TGF-beta receptor signaling activates SMADs 
                                   NM_005901.5:1678 
                                 
                                 
                                   DIAPH3-mRNA 
                                   -1.32 
                                   0.185 
                                   -1.69 
                                   -0.963 
                                   0.399 
                                   0.311 
                                   0.513 
                                   4.54e-07 
                                   4.3e-05 
                                   lm.nb 
                                   RHO GTPases Activate Formins 
                                   NM_001042517.1:1832 
                                 
                                 
                                   CYFIP2-mRNA 
                                   1.05 
                                   0.151 
                                   0.753 
                                   1.34 
                                   2.07 
                                   1.68 
                                   2.54 
                                   7.19e-07 
                                   6.36e-05 
                                   lm.nb 
                                   Regulation of actin dynamics for phagocytic cup formation, RHO GTPases Activate WASPs and WAVEs, VEGFA-VEGFR2 Pathway 
                                   NM_001037332.2:4990 
                                 
                                 
                                   NLK-mRNA 
                                   0.742 
                                   0.107 
                                   0.532 
                                   0.953 
                                   1.67 
                                   1.45 
                                   1.94 
                                   7.94e-07 
                                   6.43e-05 
                                   loglinear 
                                   
                                   NM_016231.4:618 
                                 
                                 
                                   PRKAA1-mRNA 
                                   0.393 
                                   0.0572 
                                   0.281 
                                   0.506 
                                   1.31 
                                   1.22 
                                   1.42 
                                   8.6e-07 
                                   6.43e-05 
                                   loglinear 
                                   IRS-related events triggered by IGF1R, PI3K Cascade, TP53 Regulates Metabolic Genes 
                                   NM_006251.5:1455 
                                 
                                 
                                   HRAS-mRNA 
                                   -0.987 
                                   0.144 
                                   -1.27 
                                   -0.705 
                                   0.504 
                                   0.415 
                                   0.613 
                                   8.72e-07 
                                   6.43e-05 
                                   loglinear 
                                   ARMS-mediated activation, CD209 (DC-SIGN) signaling, Cell surface interactions at the vascular wall, Constitutive Signaling by EGFRvIII, CREB phosphorylation through the activation of Ras, DAP12 signaling, Downstream signal transduction, FCERI mediated MAPK activation, Frs2-mediated activation, Gastrin-CREB signalling pathway via PKC and MAPK, Interleukin receptor SHC signaling, IRS-related events triggered by IGF1R, MAP2K and MAPK activation, RAFMAP kinase cascade, SHC-mediated cascade.CL.FGFR1, SHC-related events triggered by IGF1R, Signaling by ERBB2, Signaling by FGFR1 in disease, Signaling by FGFR2 in disease, Signaling by FGFR3 in disease, Signaling by FGFR4 in disease, Signaling by PDGF, Signaling by SCF-KIT, Tie2 Signaling, VEGFA-VEGFR2 Pathway, VEGFR2 mediated cell proliferation 
                                   NM_005343.2:396 
                                 
                                 
                                   ABCF1-mRNA 
                                   -0.486 
                                   0.0732 
                                   -0.63 
                                   -0.343 
                                   0.714 
                                   0.646 
                                   0.789 
                                   1.43e-06 
                                   9.98e-05 
                                   loglinear 
                                   
                                   NM_001090.2:850 
                                 
                                 
                                   TUBB-mRNA 
                                   -0.704 
                                   0.107 
                                   -0.913 
                                   -0.495 
                                   0.614 
                                   0.531 
                                   0.71 
                                   1.56e-06 
                                   0.000104 
                                   loglinear 
                                   
                                   NM_178014.2:1955 
                                 
                               
                               DE Results: New.Annotation: miR142 vs.cel67  Table presenting the most statistically significantly differentially expressed genes with the selected covariate. 'Estimated log fold-change' estimates a gene's differential expression. For categorical covariates, a gene is estimated to have 2^(log fold change) times its expression in baseline samples, holding all other variables in the analysis constant. If the covariate is continuous, for each unit increase in the selected covariate, a gene's expression is estimated to increase by 2^(log fold change)-fold, holding all other variables in the analysis constant. The 95% confidence interval for the log fold change is also presented, along with a p-value and an adjusted p-value or FDR if requested. Method column indicates the model used to estimate differential expression. Mixed negative binomial model uses the mle function to run the Wald test. The simplified negative binomial model uses the glm.nb function . The loglinear model uses the lm function.   
                                   
                          
                           
                      
                    
                    
                      
                           
                          
                               Volcano Plot: New.Annotation: miR155 vs.cel67   More Plot Information                Volcano Plot: New.Annotation: miR155 vs.cel67  Volcano plot displaying each gene's -log10(p-value) and log2 fold change with the selected covariate. Highly statistically significant genes fall at the top of the plot above the horizontal lines, and highly differentially expressed genes fall to either side. Horizontal lines indicate various False Discovery Rate (FDR) thresholds or p-value thresholds if there is no adjustment to the p-values. Genes are colored if the resulting p-value is below the given FDR or p-value threshold. The 40 most statistically significant genes are labeled in the plot.   
                               DE Results: New.Annotation: miR155 vs.cel67   More Plot Information      Download CSV Data     
                               
                                 
                                 
                               
                               
                                 
                                   
                                   Log2 fold change 
                                   std error (log2) 
                                   Lower confidence limit (log2) 
                                   Upper confidence limit (log2) 
                                   Linear fold change 
                                   Lower confidence limit (linear) 
                                   Upper confidence limit (linear) 
                                   P-value 
                                   BY.p.value 
                                   method 
                                   Gene.sets 
                                   probe.ID 
                                 
                               
                               
                                 
                                   PPP2CB-mRNA 
                                   -1.13 
                                   0.0787 
                                   -1.29 
                                   -0.979 
                                   0.456 
                                   0.41 
                                   0.507 
                                   2.34e-12 
                                   3.11e-09 
                                   loglinear 
                                   ARMS-mediated activation, AXIN missense mutants destabilize the destruction complex, Beta-catenin phosphorylation cascade, CTLA4 inhibitory signaling, Cyclin AB1 associated events during G2M transition, Cyclin D associated events in G1, DAP12 signaling, DARPP-32 events, Disassembly of the destruction complex and recruitment of AXIN to the membrane, Downstream signal transduction, ERKs are inactivated, FCERI mediated MAPK activation, Frs2-mediated activation, Gastrin-CREB signalling pathway via PKC and MAPK, Interleukin receptor SHC signaling, IRS-related events triggered by IGF1R, MASTL Facilitates Mitotic Progression, Misspliced GSK3beta mutants stabilize beta-catenin, Negative regulation of FGFR1 signaling, Negative regulation of FGFR2 signaling, Negative regulation of FGFR3 signaling, Negative regulation of FGFR4 signaling, Negative regulation of MAPK pathway, RAF activation, RAFMAP kinase cascade, Resolution of Sister Chromatid Cohesion, RHO GTPases Activate Formins, S33 mutants of beta-catenin aren.SQT.t phosphorylated, S37 mutants of beta-catenin aren.SQT.t phosphorylated, S45 mutants of beta-catenin aren.SQT.t phosphorylated, Signaling by ERBB2, Signaling by PDGF, Signaling by SCF-KIT, Spry regulation of FGF signaling, T41 mutants of beta-catenin aren.SQT.t phosphorylated, Truncations of AMER1 destabilize the destruction complex, VEGFA-VEGFR2 Pathway, VEGFR2 mediated cell proliferation 
                                   NM_001009552.1:1075 
                                 
                                 
                                   DVL3-mRNA 
                                   0.816 
                                   0.0831 
                                   0.653 
                                   0.979 
                                   1.76 
                                   1.57 
                                   1.97 
                                   2.7e-09 
                                   1.41e-06 
                                   loglinear 
                                   Disassembly of the destruction complex and recruitment of AXIN to the membrane, PCPCE pathway, RHO GTPases Activate Formins, WNT mediated activation of DVL 
                                   NM_004423.3:1012 
                                 
                                 
                                   PRKCI-mRNA 
                                   -0.919 
                                   0.0946 
                                   -1.1 
                                   -0.734 
                                   0.529 
                                   0.465 
                                   0.601 
                                   3.18e-09 
                                   1.41e-06 
                                   loglinear 
                                   Tight junction interactions 
                                   NM_002740.4:530 
                                 
                                 
                                   KRAS-mRNA 
                                   -0.99 
                                   0.107 
                                   -1.2 
                                   -0.781 
                                   0.503 
                                   0.435 
                                   0.582 
                                   7.18e-09 
                                   2.06e-06 
                                   loglinear 
                                   ARMS-mediated activation, CD209 (DC-SIGN) signaling, Cell surface interactions at the vascular wall, Constitutive Signaling by EGFRvIII, DAP12 signaling, Downstream signal transduction, FCERI mediated MAPK activation, Frs2-mediated activation, Gastrin-CREB signalling pathway via PKC and MAPK, Interleukin receptor SHC signaling, IRS-related events triggered by IGF1R, MAP2K and MAPK activation, RAFMAP kinase cascade, SHC-mediated cascade.CL.FGFR1, SHC-related events triggered by IGF1R, Signaling by ERBB2, Signaling by FGFR1 in disease, Signaling by FGFR2 in disease, Signaling by FGFR3 in disease, Signaling by FGFR4 in disease, Signaling by PDGF, Signaling by SCF-KIT, Tie2 Signaling, VEGFA-VEGFR2 Pathway, VEGFR2 mediated cell proliferation 
                                   NM_004985.3:327 
                                 
                                 
                                   EGFR-mRNA 
                                   -0.978 
                                   0.106 
                                   -1.19 
                                   -0.771 
                                   0.508 
                                   0.439 
                                   0.586 
                                   7.76e-09 
                                   2.06e-06 
                                   loglinear 
                                   ARMS-mediated activation, Constitutive Signaling by Aberrant PI3K in Cancer, Constitutive Signaling by EGFRvIII, DAP12 signaling, Downstream signal transduction, FCERI mediated MAPK activation, Frs2-mediated activation, GAB1 signalosome, Gastrin-CREB signalling pathway via PKC and MAPK, Interleukin receptor SHC signaling, IRS-related events triggered by IGF1R, PI3KAKT activation, Role of LAT2NTALLAB on calcium mobilization, Signaling by ERBB2, Signaling by PDGF, Signaling by SCF-KIT, VEGFA-VEGFR2 Pathway, VEGFR2 mediated cell proliferation 
                                   NM_201282.1:360 
                                 
                                 
                                   CSNK1A1-mRNA 
                                   -0.601 
                                   0.0705 
                                   -0.739 
                                   -0.463 
                                   0.659 
                                   0.599 
                                   0.725 
                                   2.89e-08 
                                   6.39e-06 
                                   loglinear 
                                   AXIN missense mutants destabilize the destruction complex, Beta-catenin phosphorylation cascade, Disassembly of the destruction complex and recruitment of AXIN to the membrane, Misspliced GSK3beta mutants stabilize beta-catenin, S33 mutants of beta-catenin aren.SQT.t phosphorylated, S37 mutants of beta-catenin aren.SQT.t phosphorylated, S45 mutants of beta-catenin aren.SQT.t phosphorylated, T41 mutants of beta-catenin aren.SQT.t phosphorylated 
                                   NM_001892.4:2053 
                                 
                                 
                                   RRAS2-mRNA 
                                   0.941 
                                   0.13 
                                   0.686 
                                   1.2 
                                   1.92 
                                   1.61 
                                   2.29 
                                   4.11e-07 
                                   7.8e-05 
                                   lm.nb 
                                   
                                   NM_001102669.2:1785 
                                 
                                 
                                   RAC1-mRNA 
                                   -0.507 
                                   0.0711 
                                   -0.647 
                                   -0.368 
                                   0.703 
                                   0.639 
                                   0.775 
                                   4.93e-07 
                                   8.18e-05 
                                   loglinear 
                                   Activation of Rac, CD28 co-stimulation, CD28 dependent Vav1 pathway, DAP12 signaling, EPH-ephrin mediated repulsion of cells, Ephrin signaling, G alpha (1213) signalling events, GPVI-mediated activation cascade, MAPK6MAPK4 signaling, NRAGE signals death through JNK, PCPCE pathway, Regulation of actin dynamics for phagocytic cup formation, RHO GTPases activate CIT, RHO GTPases Activate Formins, RHO GTPases activate IQGAPs, RHO GTPases activate PAKs, RHO GTPases activate PKNs, RHO GTPases Activate WASPs and WAVEs, Sema3A PAK dependent Axon repulsion, Sema4D induced cell migration and growth-cone collapse, Signal transduction by L1, Signaling by SCF-KIT, VEGFA-VEGFR2 Pathway, VEGFR2 mediated vascular permeability 
                                   NM_198829.1:1250 
                                 
                                 
                                   TRIO-mRNA 
                                   -0.36 
                                   0.0538 
                                   -0.465 
                                   -0.254 
                                   0.779 
                                   0.724 
                                   0.839 
                                   1.3e-06 
                                   0.000192 
                                   loglinear 
                                   DCC mediated attractive signaling, G alpha (1213) signalling events, G alpha (q) signalling events, Gastrin-CREB signalling pathway via PKC and MAPK, NRAGE signals death through JNK, Rho GTPase cycle 
                                   NM_007118.2:1735 
                                 
                                 
                                   ABI2-mRNA 
                                   -0.594 
                                   0.0979 
                                   -0.785 
                                   -0.402 
                                   0.663 
                                   0.58 
                                   0.757 
                                   5.12e-06 
                                   0.00068 
                                   loglinear 
                                   Regulation of actin dynamics for phagocytic cup formation, RHO GTPases Activate WASPs and WAVEs, VEGFA-VEGFR2 Pathway 
                                   NM_005759.5:1160 
                                 
                                 
                                   TCF7L2-mRNA 
                                   -0.776 
                                   0.133 
                                   -1.04 
                                   -0.516 
                                   0.584 
                                   0.488 
                                   0.699 
                                   8.27e-06 
                                   0.000998 
                                   lm.nb 
                                   Deactivation of the beta-catenin transactivating complex, Formation of the beta-catenin.CL.TCF transactivating complex, Repression of WNT target genes 
                                   NM_001146274.1:1393 
                                 
                                 
                                   WASF2-mRNA 
                                   0.592 
                                   0.108 
                                   0.379 
                                   0.805 
                                   1.51 
                                   1.3 
                                   1.75 
                                   2.04e-05 
                                   0.00226 
                                   loglinear 
                                   Regulation of actin dynamics for phagocytic cup formation, RHO GTPases Activate WASPs and WAVEs, VEGFA-VEGFR2 Pathway 
                                   NM_006990.2:485 
                                 
                                 
                                   DIAPH3-mRNA 
                                   -1.04 
                                   0.193 
                                   -1.42 
                                   -0.66 
                                   0.487 
                                   0.374 
                                   0.633 
                                   2.49e-05 
                                   0.00255 
                                   lm.nb 
                                   RHO GTPases Activate Formins 
                                   NM_001042517.1:1832 
                                 
                                 
                                   PAK2-mRNA 
                                   -0.548 
                                   0.103 
                                   -0.75 
                                   -0.347 
                                   0.684 
                                   0.595 
                                   0.786 
                                   2.77e-05 
                                   0.00263 
                                   loglinear 
                                   Activation of Rac, CD209 (DC-SIGN) signaling, CD28 co-stimulation, CD28 dependent Vav1 pathway, Ephrin signaling, RHO GTPases activate PAKs, Sema3A PAK dependent Axon repulsion, VEGFA-VEGFR2 Pathway, VEGFR2 mediated vascular permeability 
                                   NM_002577.3:3785 
                                 
                                 
                                   IRS1-mRNA 
                                   -0.579 
                                   0.114 
                                   -0.802 
                                   -0.356 
                                   0.669 
                                   0.573 
                                   0.781 
                                   4.83e-05 
                                   0.00428 
                                   loglinear 
                                   ARMS-mediated activation, Constitutive Signaling by Aberrant PI3K in Cancer, DAP12 signaling, Downstream signal transduction, FCERI mediated MAPK activation, Frs2-mediated activation, GAB1 signalosome, Gastrin-CREB signalling pathway via PKC and MAPK, Interleukin receptor SHC signaling, IRS-related events triggered by IGF1R, PI3K Cascade, PI3KAKT activation, RAFMAP kinase cascade, Role of LAT2NTALLAB on calcium mobilization, Signaling by ERBB2, Signaling by PDGF, Signaling by SCF-KIT, VEGFA-VEGFR2 Pathway, VEGFR2 mediated cell proliferation 
                                   NM_005544.2:6224 
                                 
                                 
                                   CDK4-mRNA 
                                   -0.436 
                                   0.0885 
                                   -0.609 
                                   -0.262 
                                   0.739 
                                   0.656 
                                   0.834 
                                   7.25e-05 
                                   0.00601 
                                   loglinear 
                                   Cyclin D associated events in G1 
                                   NM_000075.2:1055 
                                 
                                 
                                   MAPK3-mRNA 
                                   0.262 
                                   0.057 
                                   0.151 
                                   0.374 
                                   1.2 
                                   1.11 
                                   1.3 
                                   0.000153 
                                   0.012 
                                   loglinear 
                                   ARMS-mediated activation, DAP12 signaling, Downstream signal transduction, ERKs are inactivated, FCERI mediated MAPK activation, Frs2-mediated activation, Gastrin-CREB signalling pathway via PKC and MAPK, Interleukin receptor SHC signaling, IRS-related events triggered by IGF1R, Negative feedback regulation of MAPK pathway, Negative regulation of FGFR3 signaling, Negative regulation of FGFR4 signaling, Negative regulation of MAPK pathway, RAFMAP kinase cascade, Regulation of HSF1-mediated heat shock response, RHO GTPases Activate WASPs and WAVEs, Signaling by ERBB2, Signaling by PDGF, Signaling by SCF-KIT, VEGFA-VEGFR2 Pathway, VEGFR2 mediated cell proliferation 
                                   NM_001040056.1:580 
                                 
                                 
                                   SOS1-mRNA 
                                   -0.396 
                                   0.0917 
                                   -0.576 
                                   -0.216 
                                   0.76 
                                   0.671 
                                   0.861 
                                   0.000304 
                                   0.0224 
                                   loglinear 
                                   ARMS-mediated activation, Cell surface interactions at the vascular wall, Constitutive Signaling by EGFRvIII, DAP12 signaling, Downstream signal transduction, FCERI mediated MAPK activation, Frs2-mediated activation, Gastrin-CREB signalling pathway via PKC and MAPK, Integrin alphaIIb beta3 signaling, Interleukin receptor SHC signaling, IRS-related events triggered by IGF1R, NRAGE signals death through JNK, RAFMAP kinase cascade, Role of LAT2NTALLAB on calcium mobilization, SHC-mediated cascade.CL.FGFR1, SHC-related events triggered by IGF1R, Signaling by ERBB2, Signaling by FGFR1 in disease, Signaling by FGFR2 in disease, Signaling by FGFR3 in disease, Signaling by FGFR4 in disease, Signaling by PDGF, Signaling by SCF-KIT, Tie2 Signaling, VEGFA-VEGFR2 Pathway, VEGFR2 mediated cell proliferation 
                                   NM_005633.2:1635 
                                 
                                 
                                   HPRT1-mRNA 
                                   -0.509 
                                   0.12 
                                   -0.744 
                                   -0.273 
                                   0.703 
                                   0.597 
                                   0.827 
                                   0.000367 
                                   0.0256 
                                   lm.nb 
                                   
                                   NM_000194.1:240 
                                 
                                 
                                   FGFR1-mRNA 
                                   -0.452 
                                   0.108 
                                   -0.663 
                                   -0.241 
                                   0.731 
                                   0.631 
                                   0.846 
                                   0.000398 
                                   0.0264 
                                   lm.nb 
                                   ARMS-mediated activation, Constitutive Signaling by Aberrant PI3K in Cancer, DAP12 signaling, Downstream signal transduction, FCERI mediated MAPK activation, Frs2-mediated activation, GAB1 signalosome, Gastrin-CREB signalling pathway via PKC and MAPK, Interleukin receptor SHC signaling, IRS-related events triggered by IGF1R, PI3K Cascade, PI3KAKT activation, RAFMAP kinase cascade, Role of LAT2NTALLAB on calcium mobilization, SHC-mediated cascade.CL.FGFR1, Signaling by ERBB2, Signaling by FGFR1 in disease, Signaling by PDGF, Signaling by SCF-KIT, VEGFA-VEGFR2 Pathway, VEGFR2 mediated cell proliferation 
                                   NM_015850.2:1335 
                                 
                               
                               DE Results: New.Annotation: miR155 vs.cel67  Table presenting the most statistically significantly differentially expressed genes with the selected covariate. 'Estimated log fold-change' estimates a gene's differential expression. For categorical covariates, a gene is estimated to have 2^(log fold change) times its expression in baseline samples, holding all other variables in the analysis constant. If the covariate is continuous, for each unit increase in the selected covariate, a gene's expression is estimated to increase by 2^(log fold change)-fold, holding all other variables in the analysis constant. The 95% confidence interval for the log fold change is also presented, along with a p-value and an adjusted p-value or FDR if requested. Method column indicates the model used to estimate differential expression. Mixed negative binomial model uses the mle function to run the Wald test. The simplified negative binomial model uses the glm.nb function . The loglinear model uses the lm function.   
                                   
                          
                           
                      
                    
                    
                      
                           
                          
                               Volcano Plot: New.Annotation: miR18b vs.cel67   More Plot Information                Volcano Plot: New.Annotation: miR18b vs.cel67  Volcano plot displaying each gene's -log10(p-value) and log2 fold change with the selected covariate. Highly statistically significant genes fall at the top of the plot above the horizontal lines, and highly differentially expressed genes fall to either side. Horizontal lines indicate various False Discovery Rate (FDR) thresholds or p-value thresholds if there is no adjustment to the p-values. Genes are colored if the resulting p-value is below the given FDR or p-value threshold. The 40 most statistically significant genes are labeled in the plot.   
                               DE Results: New.Annotation: miR18b vs.cel67   More Plot Information      Download CSV Data     
                               
                                 
                                 
                               
                               
                                 
                                   
                                   Log2 fold change 
                                   std error (log2) 
                                   Lower confidence limit (log2) 
                                   Upper confidence limit (log2) 
                                   Linear fold change 
                                   Lower confidence limit (linear) 
                                   Upper confidence limit (linear) 
                                   P-value 
                                   BY.p.value 
                                   method 
                                   Gene.sets 
                                   probe.ID 
                                 
                               
                               
                                 
                                   CDC42-mRNA 
                                   -1.5 
                                   0.0913 
                                   -1.68 
                                   -1.33 
                                   0.353 
                                   0.311 
                                   0.399 
                                   1.75e-13 
                                   2.32e-10 
                                   loglinear 
                                   CD28 co-stimulation, CD28 dependent Vav1 pathway, DCC mediated attractive signaling, EPHB-mediated forward signaling, G alpha (1213) signalling events, GPVI-mediated activation cascade, Regulation of actin dynamics for phagocytic cup formation, Rho GTPase cycle, RHO GTPases Activate Formins, RHO GTPases activate IQGAPs, RHO GTPases activate PAKs, Sema4D induced cell migration and growth-cone collapse, VEGFA-VEGFR2 Pathway 
                                   NM_001039802.1:390 
                                 
                                 
                                   DICER1-mRNA 
                                   -1.32 
                                   0.109 
                                   -1.53 
                                   -1.1 
                                   0.402 
                                   0.346 
                                   0.466 
                                   6.34e-11 
                                   3.15e-08 
                                   loglinear 
                                   MicroRNA (miRNA) biogenesis 
                                   NM_177438.2:5060 
                                 
                                 
                                   SMAD2-mRNA 
                                   -0.882 
                                   0.0734 
                                   -1.03 
                                   -0.738 
                                   0.543 
                                   0.491 
                                   0.6 
                                   7.11e-11 
                                   3.15e-08 
                                   loglinear 
                                   TGF-beta receptor signaling activates SMADs 
                                   NM_005901.5:1678 
                                 
                                 
                                   PDGFC-mRNA 
                                   -1.27 
                                   0.129 
                                   -1.52 
                                   -1.01 
                                   0.416 
                                   0.349 
                                   0.495 
                                   2.49e-09 
                                   8.28e-07 
                                   lm.nb 
                                   Signaling by PDGF 
                                   NM_016205.2:2596 
                                 
                                 
                                   RAF1-mRNA 
                                   -0.772 
                                   0.0829 
                                   -0.934 
                                   -0.609 
                                   0.586 
                                   0.523 
                                   0.656 
                                   6.7e-09 
                                   1.78e-06 
                                   loglinear 
                                   ARMS-mediated activation, CD209 (DC-SIGN) signaling, CREB phosphorylation through the activation of Ras, DAP12 signaling, Downstream signal transduction, FCERI mediated MAPK activation, Frs2-mediated activation, Gastrin-CREB signalling pathway via PKC and MAPK, Interleukin receptor SHC signaling, IRS-related events triggered by IGF1R, Negative feedback regulation of MAPK pathway, Negative regulation of MAPK pathway, RAF activation, RAFMAP kinase cascade, Signaling by ERBB2, Signaling by PDGF, Signaling by SCF-KIT, VEGFA-VEGFR2 Pathway, VEGFR2 mediated cell proliferation 
                                   NM_002880.3:1444 
                                 
                                 
                                   ITGA5-mRNA 
                                   -1.47 
                                   0.171 
                                   -1.81 
                                   -1.13 
                                   0.361 
                                   0.286 
                                   0.456 
                                   2.68e-08 
                                   5.92e-06 
                                   lm.nb 
                                   Cell surface interactions at the vascular wall, Elastic fibre formation, Integrin cell surface interactions, Signal transduction by L1 
                                   NM_002205.2:925 
                                 
                                 
                                   PIK3R2-mRNA 
                                   -1.26 
                                   0.158 
                                   -1.57 
                                   -0.953 
                                   0.417 
                                   0.336 
                                   0.517 
                                   8.32e-08 
                                   1.58e-05 
                                   lm.nb 
                                   CD28 co-stimulation, CD28 dependent PI3KAkt signaling, Cell surface interactions at the vascular wall, Constitutive Signaling by Aberrant PI3K in Cancer, DAP12 signaling, Downstream signal transduction, Downstream TCR signaling, G alpha (1213) signalling events, G alpha (q) signalling events, G beta.CL.gamma signalling through PI3Kgamma, GAB1 signalosome, Gastrin-CREB signalling pathway via PKC and MAPK, GPVI-mediated activation cascade, Interleukin receptor SHC signaling, IRS-related events triggered by IGF1R, Nephrin interactions, PI3K Cascade, PI3KAKT activation, PIP3 activates AKT signaling, Regulation of signaling by CBL, Rho GTPase cycle, Role of LAT2NTALLAB on calcium mobilization, Signaling by ERBB2, Signaling by PDGF, Signaling by SCF-KIT, Synthesis of PIPs at the plasma membrane, Tie2 Signaling, VEGFA-VEGFR2 Pathway 
                                   NM_005027.2:3100 
                                 
                                 
                                   MAPK1-mRNA 
                                   -0.571 
                                   0.0726 
                                   -0.713 
                                   -0.429 
                                   0.673 
                                   0.61 
                                   0.743 
                                   1.09e-07 
                                   1.81e-05 
                                   loglinear 
                                   ARMS-mediated activation, CREB phosphorylation through the activation of Ras, DAP12 signaling, Downstream signal transduction, FCERI mediated MAPK activation, Frs2-mediated activation, Gastrin-CREB signalling pathway via PKC and MAPK, Interleukin receptor SHC signaling, IRS-related events triggered by IGF1R, MAP2K and MAPK activation, Negative feedback regulation of MAPK pathway, Negative regulation of FGFR1 signaling, Negative regulation of FGFR2 signaling, Negative regulation of FGFR3 signaling, Negative regulation of FGFR4 signaling, Negative regulation of MAPK pathway, RAFMAP kinase cascade, Regulation of HSF1-mediated heat shock response, RHO GTPases Activate WASPs and WAVEs, Signal transduction by L1, Signaling by ERBB2, Signaling by PDGF, Signaling by SCF-KIT, VEGFA-VEGFR2 Pathway, VEGFR2 mediated cell proliferation 
                                   NM_138957.2:430 
                                 
                                 
                                   CSNK1A1-mRNA 
                                   0.584 
                                   0.0761 
                                   0.435 
                                   0.733 
                                   1.5 
                                   1.35 
                                   1.66 
                                   1.6e-07 
                                   2.36e-05 
                                   loglinear 
                                   AXIN missense mutants destabilize the destruction complex, Beta-catenin phosphorylation cascade, Disassembly of the destruction complex and recruitment of AXIN to the membrane, Misspliced GSK3beta mutants stabilize beta-catenin, S33 mutants of beta-catenin aren.SQT.t phosphorylated, S37 mutants of beta-catenin aren.SQT.t phosphorylated, S45 mutants of beta-catenin aren.SQT.t phosphorylated, T41 mutants of beta-catenin aren.SQT.t phosphorylated 
                                   NM_001892.4:2053 
                                 
                                 
                                   PAK4-mRNA 
                                   -1.05 
                                   0.142 
                                   -1.33 
                                   -0.776 
                                   0.482 
                                   0.397 
                                   0.584 
                                   2.6e-07 
                                   3.45e-05 
                                   loglinear 
                                   Activation of Rac 
                                   NM_005884.3:2130 
                                 
                                 
                                   MAPK8-mRNA 
                                   -0.653 
                                   0.0885 
                                   -0.827 
                                   -0.48 
                                   0.636 
                                   0.564 
                                   0.717 
                                   2.89e-07 
                                   3.5e-05 
                                   loglinear 
                                   FCERI mediated MAPK activation 
                                   NM_002750.2:945 
                                 
                                 
                                   MAPK3-mRNA 
                                   0.442 
                                   0.0615 
                                   0.321 
                                   0.563 
                                   1.36 
                                   1.25 
                                   1.48 
                                   4.43e-07 
                                   4.91e-05 
                                   loglinear 
                                   ARMS-mediated activation, DAP12 signaling, Downstream signal transduction, ERKs are inactivated, FCERI mediated MAPK activation, Frs2-mediated activation, Gastrin-CREB signalling pathway via PKC and MAPK, Interleukin receptor SHC signaling, IRS-related events triggered by IGF1R, Negative feedback regulation of MAPK pathway, Negative regulation of FGFR3 signaling, Negative regulation of FGFR4 signaling, Negative regulation of MAPK pathway, RAFMAP kinase cascade, Regulation of HSF1-mediated heat shock response, RHO GTPases Activate WASPs and WAVEs, Signaling by ERBB2, Signaling by PDGF, Signaling by SCF-KIT, VEGFA-VEGFR2 Pathway, VEGFR2 mediated cell proliferation 
                                   NM_001040056.1:580 
                                 
                                 
                                   MAPK9-mRNA 
                                   0.397 
                                   0.056 
                                   0.288 
                                   0.507 
                                   1.32 
                                   1.22 
                                   1.42 
                                   5.25e-07 
                                   5.36e-05 
                                   loglinear 
                                   
                                   XM_005265940.1:163 
                                 
                                 
                                   RRAS-mRNA 
                                   -1.33 
                                   0.196 
                                   -1.72 
                                   -0.948 
                                   0.397 
                                   0.304 
                                   0.518 
                                   1.01e-06 
                                   9.55e-05 
                                   lm.nb 
                                   CREB phosphorylation through the activation of Ras 
                                   NM_006270.3:330 
                                 
                                 
                                   ITGA6-mRNA 
                                   -1.75 
                                   0.264 
                                   -2.27 
                                   -1.24 
                                   0.297 
                                   0.207 
                                   0.424 
                                   1.41e-06 
                                   0.000125 
                                   lm.nb 
                                   Cell surface interactions at the vascular wall, Non-integrin membrane-ECM interactions, Syndecan interactions 
                                   NM_000210.1:3065 
                                 
                                 
                                   LIMK1-mRNA 
                                   -0.731 
                                   0.117 
                                   -0.961 
                                   -0.501 
                                   0.603 
                                   0.514 
                                   0.707 
                                   3.47e-06 
                                   0.000286 
                                   loglinear 
                                   EPHB-mediated forward signaling, Regulation of actin dynamics for phagocytic cup formation, RHO GTPases activate PAKs, RHO GTPases Activate ROCKs, Sema3A PAK dependent Axon repulsion, Sema4D induced cell migration and growth-cone collapse 
                                   NM_002314.3:3133 
                                 
                                 
                                   NRAS-mRNA 
                                   -0.806 
                                   0.13 
                                   -1.06 
                                   -0.551 
                                   0.572 
                                   0.479 
                                   0.683 
                                   3.82e-06 
                                   0.000286 
                                   lm.nb 
                                   ARMS-mediated activation, CD209 (DC-SIGN) signaling, Cell surface interactions at the vascular wall, Constitutive Signaling by EGFRvIII, DAP12 signaling, Downstream signal transduction, FCERI mediated MAPK activation, Frs2-mediated activation, Gastrin-CREB signalling pathway via PKC and MAPK, Interleukin receptor SHC signaling, IRS-related events triggered by IGF1R, MAP2K and MAPK activation, RAFMAP kinase cascade, SHC-mediated cascade.CL.FGFR1, SHC-related events triggered by IGF1R, Signaling by ERBB2, Signaling by FGFR1 in disease, Signaling by FGFR2 in disease, Signaling by FGFR3 in disease, Signaling by FGFR4 in disease, Signaling by PDGF, Signaling by SCF-KIT, Tie2 Signaling, VEGFA-VEGFR2 Pathway, VEGFR2 mediated cell proliferation 
                                   NM_002524.3:877 
                                 
                                 
                                   MYC-mRNA 
                                   -1.28 
                                   0.208 
                                   -1.69 
                                   -0.875 
                                   0.411 
                                   0.31 
                                   0.545 
                                   4.04e-06 
                                   0.000286 
                                   lm.nb 
                                   Formation of the beta-catenin.CL.TCF transactivating complex 
                                   NM_002467.3:1610 
                                 
                                 
                                   RRAS2-mRNA 
                                   0.867 
                                   0.141 
                                   0.591 
                                   1.14 
                                   1.82 
                                   1.51 
                                   2.21 
                                   4.09e-06 
                                   0.000286 
                                   lm.nb 
                                   
                                   NM_001102669.2:1785 
                                 
                                 
                                   RAB8B-mRNA 
                                   -0.318 
                                   0.0526 
                                   -0.422 
                                   -0.215 
                                   0.802 
                                   0.747 
                                   0.861 
                                   5.23e-06 
                                   0.000347 
                                   loglinear 
                                   
                                   NM_016530.2:820 
                                 
                               
                               DE Results: New.Annotation: miR18b vs.cel67  Table presenting the most statistically significantly differentially expressed genes with the selected covariate. 'Estimated log fold-change' estimates a gene's differential expression. For categorical covariates, a gene is estimated to have 2^(log fold change) times its expression in baseline samples, holding all other variables in the analysis constant. If the covariate is continuous, for each unit increase in the selected covariate, a gene's expression is estimated to increase by 2^(log fold change)-fold, holding all other variables in the analysis constant. The 95% confidence interval for the log fold change is also presented, along with a p-value and an adjusted p-value or FDR if requested. Method column indicates the model used to estimate differential expression. Mixed negative binomial model uses the mle function to run the Wald test. The simplified negative binomial model uses the glm.nb function . The loglinear model uses the lm function.   
                                   
                          
                           
                      
                    
                    
                      
                           
                          
                               Volcano Plot: New.Annotation: miR890 vs.cel67   More Plot Information                Volcano Plot: New.Annotation: miR890 vs.cel67  Volcano plot displaying each gene's -log10(p-value) and log2 fold change with the selected covariate. Highly statistically significant genes fall at the top of the plot above the horizontal lines, and highly differentially expressed genes fall to either side. Horizontal lines indicate various False Discovery Rate (FDR) thresholds or p-value thresholds if there is no adjustment to the p-values. Genes are colored if the resulting p-value is below the given FDR or p-value threshold. The 40 most statistically significant genes are labeled in the plot.   
                               DE Results: New.Annotation: miR890 vs.cel67   More Plot Information      Download CSV Data     
                               
                                 
                                 
                               
                               
                                 
                                   
                                   Log2 fold change 
                                   std error (log2) 
                                   Lower confidence limit (log2) 
                                   Upper confidence limit (log2) 
                                   Linear fold change 
                                   Lower confidence limit (linear) 
                                   Upper confidence limit (linear) 
                                   P-value 
                                   BY.p.value 
                                   method 
                                   Gene.sets 
                                   probe.ID 
                                 
                               
                               
                                 
                                   CCND2-mRNA 
                                   -1.84 
                                   0.16 
                                   -2.16 
                                   -1.53 
                                   0.279 
                                   0.224 
                                   0.347 
                                   1.58e-10 
                                   2.1e-07 
                                   lm.nb 
                                   Cyclin D associated events in G1 
                                   NM_001759.3:503 
                                 
                                 
                                   ROCK1-mRNA 
                                   -0.567 
                                   0.0543 
                                   -0.674 
                                   -0.461 
                                   0.675 
                                   0.627 
                                   0.727 
                                   9.12e-10 
                                   6.05e-07 
                                   loglinear 
                                   Apoptotic cleavage of cellular proteins, EPHA-mediated growth cone collapse, EPHB-mediated forward signaling, G alpha (1213) signalling events, RHO GTPases Activate ROCKs, Sema4D induced cell migration and growth-cone collapse, VEGFA-VEGFR2 Pathway 
                                   NM_005406.1:2660 
                                 
                                 
                                   YBX3-mRNA 
                                   -1.11 
                                   0.11 
                                   -1.33 
                                   -0.895 
                                   0.463 
                                   0.399 
                                   0.538 
                                   1.68e-09 
                                   7.44e-07 
                                   lm.nb 
                                   
                                   NM_003651.3:660 
                                 
                                 
                                   TUBB-mRNA 
                                   -1.01 
                                   0.103 
                                   -1.21 
                                   -0.806 
                                   0.497 
                                   0.433 
                                   0.572 
                                   2.68e-09 
                                   8.9e-07 
                                   loglinear 
                                   
                                   NM_178014.2:1955 
                                 
                                 
                                   DROSHA-mRNA 
                                   -0.616 
                                   0.064 
                                   -0.741 
                                   -0.49 
                                   0.653 
                                   0.598 
                                   0.712 
                                   3.84e-09 
                                   1.02e-06 
                                   loglinear 
                                   MicroRNA (miRNA) biogenesis 
                                   NM_013235.4:1860 
                                 
                                 
                                   PRKCA-mRNA 
                                   -0.493 
                                   0.0565 
                                   -0.604 
                                   -0.382 
                                   0.711 
                                   0.658 
                                   0.767 
                                   2e-08 
                                   4.43e-06 
                                   loglinear 
                                   DAP12 signaling, Downstream signal transduction, Gastrin-CREB signalling pathway via PKC and MAPK, Non-integrin membrane-ECM interactions, PCPCE pathway, Signaling by ERBB2, Signaling by PDGF, Signaling by SCF-KIT, VEGFA-VEGFR2 Pathway 
                                   NM_002737.2:680 
                                 
                                 
                                   MAP2K1-mRNA 
                                   -0.977 
                                   0.116 
                                   -1.2 
                                   -0.75 
                                   0.508 
                                   0.434 
                                   0.595 
                                   3.48e-08 
                                   6.61e-06 
                                   lm.nb 
                                   ARMS-mediated activation, DAP12 signaling, Downstream signal transduction, FCERI mediated MAPK activation, Frs2-mediated activation, Gastrin-CREB signalling pathway via PKC and MAPK, Interleukin receptor SHC signaling, IRS-related events triggered by IGF1R, Negative feedback regulation of MAPK pathway, Negative regulation of MAPK pathway, RAF activation, RAFMAP kinase cascade, Signaling by ERBB2, Signaling by PDGF, Signaling by SCF-KIT, VEGFA-VEGFR2 Pathway, VEGFR2 mediated cell proliferation 
                                   NM_002755.2:970 
                                 
                                 
                                   CDK4-mRNA 
                                   -0.621 
                                   0.0808 
                                   -0.779 
                                   -0.462 
                                   0.65 
                                   0.583 
                                   0.726 
                                   1.57e-07 
                                   2.61e-05 
                                   loglinear 
                                   Cyclin D associated events in G1 
                                   NM_000075.2:1055 
                                 
                                 
                                   PDCD4-mRNA 
                                   -1.45 
                                   0.191 
                                   -1.82 
                                   -1.07 
                                   0.366 
                                   0.283 
                                   0.475 
                                   1.88e-07 
                                   2.67e-05 
                                   lm.nb 
                                   
                                   NM_014456.3:1115 
                                 
                                 
                                   MAPK3-mRNA 
                                   0.393 
                                   0.052 
                                   0.291 
                                   0.495 
                                   1.31 
                                   1.22 
                                   1.41 
                                   2.01e-07 
                                   2.67e-05 
                                   loglinear 
                                   ARMS-mediated activation, DAP12 signaling, Downstream signal transduction, ERKs are inactivated, FCERI mediated MAPK activation, Frs2-mediated activation, Gastrin-CREB signalling pathway via PKC and MAPK, Interleukin receptor SHC signaling, IRS-related events triggered by IGF1R, Negative feedback regulation of MAPK pathway, Negative regulation of FGFR3 signaling, Negative regulation of FGFR4 signaling, Negative regulation of MAPK pathway, RAFMAP kinase cascade, Regulation of HSF1-mediated heat shock response, RHO GTPases Activate WASPs and WAVEs, Signaling by ERBB2, Signaling by PDGF, Signaling by SCF-KIT, VEGFA-VEGFR2 Pathway, VEGFR2 mediated cell proliferation 
                                   NM_001040056.1:580 
                                 
                                 
                                   APC-mRNA 
                                   -0.688 
                                   0.0921 
                                   -0.869 
                                   -0.508 
                                   0.621 
                                   0.548 
                                   0.703 
                                   2.41e-07 
                                   2.92e-05 
                                   loglinear 
                                   Apoptotic cleavage of cellular proteins, AXIN missense mutants destabilize the destruction complex, Beta-catenin phosphorylation cascade, Deactivation of the beta-catenin transactivating complex, Disassembly of the destruction complex and recruitment of AXIN to the membrane, Misspliced GSK3beta mutants stabilize beta-catenin, S33 mutants of beta-catenin aren.SQT.t phosphorylated, S37 mutants of beta-catenin aren.SQT.t phosphorylated, S45 mutants of beta-catenin aren.SQT.t phosphorylated, T41 mutants of beta-catenin aren.SQT.t phosphorylated, Truncations of AMER1 destabilize the destruction complex 
                                   NM_000038.3:6850 
                                 
                                 
                                   NRAS-mRNA 
                                   -0.783 
                                   0.109 
                                   -0.997 
                                   -0.568 
                                   0.581 
                                   0.501 
                                   0.674 
                                   4.76e-07 
                                   5.27e-05 
                                   lm.nb 
                                   ARMS-mediated activation, CD209 (DC-SIGN) signaling, Cell surface interactions at the vascular wall, Constitutive Signaling by EGFRvIII, DAP12 signaling, Downstream signal transduction, FCERI mediated MAPK activation, Frs2-mediated activation, Gastrin-CREB signalling pathway via PKC and MAPK, Interleukin receptor SHC signaling, IRS-related events triggered by IGF1R, MAP2K and MAPK activation, RAFMAP kinase cascade, SHC-mediated cascade.CL.FGFR1, SHC-related events triggered by IGF1R, Signaling by ERBB2, Signaling by FGFR1 in disease, Signaling by FGFR2 in disease, Signaling by FGFR3 in disease, Signaling by FGFR4 in disease, Signaling by PDGF, Signaling by SCF-KIT, Tie2 Signaling, VEGFA-VEGFR2 Pathway, VEGFR2 mediated cell proliferation 
                                   NM_002524.3:877 
                                 
                                 
                                   WASF2-mRNA 
                                   -0.682 
                                   0.099 
                                   -0.876 
                                   -0.488 
                                   0.623 
                                   0.545 
                                   0.713 
                                   8.31e-07 
                                   7.66e-05 
                                   loglinear 
                                   Regulation of actin dynamics for phagocytic cup formation, RHO GTPases Activate WASPs and WAVEs, VEGFA-VEGFR2 Pathway 
                                   NM_006990.2:485 
                                 
                                 
                                   DIAPH3-mRNA 
                                   -1.21 
                                   0.176 
                                   -1.56 
                                   -0.867 
                                   0.431 
                                   0.339 
                                   0.548 
                                   8.57e-07 
                                   7.66e-05 
                                   lm.nb 
                                   RHO GTPases Activate Formins 
                                   NM_001042517.1:1832 
                                 
                                 
                                   CYFIP2-mRNA 
                                   1.01 
                                   0.147 
                                   0.719 
                                   1.29 
                                   2.01 
                                   1.65 
                                   2.45 
                                   8.65e-07 
                                   7.66e-05 
                                   lm.nb 
                                   Regulation of actin dynamics for phagocytic cup formation, RHO GTPases Activate WASPs and WAVEs, VEGFA-VEGFR2 Pathway 
                                   NM_001037332.2:4990 
                                 
                                 
                                   PPP1R12A-mRNA 
                                   -0.454 
                                   0.0668 
                                   -0.585 
                                   -0.323 
                                   0.73 
                                   0.667 
                                   0.799 
                                   1e-06 
                                   8.31e-05 
                                   loglinear 
                                   RHO GTPases activate PAKs, RHO GTPases activate PKNs, RHO GTPases Activate ROCKs 
                                   NM_002480.2:2935 
                                 
                                 
                                   HSPA1B-mRNA 
                                   0.761 
                                   0.112 
                                   0.541 
                                   0.981 
                                   1.69 
                                   1.45 
                                   1.97 
                                   1.08e-06 
                                   8.41e-05 
                                   lm.nb 
                                   Regulation of HSF1-mediated heat shock response 
                                   NM_005346.3:2095 
                                 
                                 
                                   MYC-mRNA 
                                   -1.13 
                                   0.172 
                                   -1.47 
                                   -0.793 
                                   0.457 
                                   0.362 
                                   0.577 
                                   1.67e-06 
                                   0.000123 
                                   lm.nb 
                                   Formation of the beta-catenin.CL.TCF transactivating complex 
                                   NM_002467.3:1610 
                                 
                                 
                                   ASH1L-mRNA 
                                   -0.645 
                                   0.102 
                                   -0.844 
                                   -0.446 
                                   0.64 
                                   0.557 
                                   0.734 
                                   2.7e-06 
                                   0.000189 
                                   loglinear 
                                   
                                   NM_018489.2:5005 
                                 
                                 
                                   HSPA8-mRNA 
                                   -0.609 
                                   0.104 
                                   -0.812 
                                   -0.406 
                                   0.656 
                                   0.569 
                                   0.755 
                                   7.71e-06 
                                   5e-04 
                                   loglinear 
                                   
                                   NM_006597.3:2111 
                                 
                               
                               DE Results: New.Annotation: miR890 vs.cel67  Table presenting the most statistically significantly differentially expressed genes with the selected covariate. 'Estimated log fold-change' estimates a gene's differential expression. For categorical covariates, a gene is estimated to have 2^(log fold change) times its expression in baseline samples, holding all other variables in the analysis constant. If the covariate is continuous, for each unit increase in the selected covariate, a gene's expression is estimated to increase by 2^(log fold change)-fold, holding all other variables in the analysis constant. The 95% confidence interval for the log fold change is also presented, along with a p-value and an adjusted p-value or FDR if requested. Method column indicates the model used to estimate differential expression. Mixed negative binomial model uses the mle function to run the Wald test. The simplified negative binomial model uses the glm.nb function . The loglinear model uses the lm function.   
                                   
                          
                           
                      
                    
                  
                 
              
                
          
        
        
          
                The results of differential expression testing are summarized at the gene set level. Each gene set's most differentially expressed genes are identified, and the extent of differential expression in each gene set is summarized using a 'global significance score'.     
              
                   Gene sets: 
                  
                      Summary  
                      Activation of BAD and translocation to mitochondria   
                      Activation of Rac  
                      Apoptotic cleavage of cellular proteins  
                      ARMS-mediated activation  
                      AXIN missense mutants destabilize the destruction complex  
                      Beta-catenin phosphorylation cascade  
                      CD209 (DC-SIGN) signaling  
                      CD28 co-stimulation  
                      CD28 dependent PI3KAkt signaling  
                      CD28 dependent Vav1 pathway  
                      Cell-extracellular matrix interactions  
                      Cell surface interactions at the vascular wall  
                      Circadian Clock  
                      Constitutive Signaling by Aberrant PI3K in Cancer  
                      Constitutive Signaling by EGFRvIII  
                      CREB phosphorylation through the activation of Ras  
                      CTLA4 inhibitory signaling  
                      Cyclin AB1 associated events during G2M transition  
                      Cyclin D associated events in G1  
                      DAP12 signaling  
                      DARPP-32 events  
                      DCC mediated attractive signaling  
                      Deactivation of the beta-catenin transactivating complex  
                      Disassembly of the destruction complex and recruitment of AXIN t...  
                      Downstream signal transduction  
                      Downstream TCR signaling  
                      ECM proteoglycans  
                      Elastic fibre formation  
                      EPH-ephrin mediated repulsion of cells  
                      EPHA-mediated growth cone collapse  
                      EPHB-mediated forward signaling  
                      Ephrin signaling  
                      ERKs are inactivated  
                      FCERI mediated MAPK activation  
                      Formation of the beta-catenin.CL.TCF transactivating complex  
                      Frs2-mediated activation  
                      G alpha (1213) signalling events  
                      G alpha (q) signalling events  
                      G beta.CL.gamma signalling through PI3Kgamma  
                      GAB1 signalosome  
                      Gastrin-CREB signalling pathway via PKC and MAPK  
                      GPVI-mediated activation cascade  
                      Integrin alphaIIb beta3 signaling  
                      Integrin cell surface interactions  
                      Interleukin receptor SHC signaling  
                      IRS-related events triggered by IGF1R  
                      Laminin interactions  
                      MAP2K and MAPK activation  
                      MAPK6MAPK4 signaling  
                      MASTL Facilitates Mitotic Progression  
                      MicroRNA (miRNA) biogenesis  
                      Misspliced GSK3beta mutants stabilize beta-catenin  
                      Molecules associated with elastic fibres  
                      Negative feedback regulation of MAPK pathway  
                      Negative regulation of FGFR1 signaling  
                      Negative regulation of FGFR2 signaling  
                      Negative regulation of FGFR3 signaling  
                      Negative regulation of FGFR4 signaling  
                      Negative regulation of MAPK pathway  
                      Nephrin interactions  
                      Non-integrin membrane-ECM interactions  
                      Nonsense Mediated Decay (NMD) enhanced by the Exon Junction Comp...  
                      NRAGE signals death through JNK  
                      p130Cas linkage to MAPK signaling for integrins  
                      PCPCE pathway  
                      PI3K Cascade  
                      PI3KAKT activation  
                      PIP3 activates AKT signaling  
                      Platelet degranulation   
                      RAF activation  
                      RAFMAP kinase cascade  
                      Regulation of actin dynamics for phagocytic cup formation  
                      Regulation of HSF1-mediated heat shock response  
                      Regulation of signaling by CBL  
                      Repression of WNT target genes  
                      Resolution of Sister Chromatid Cohesion  
                      Rho GTPase cycle  
                      RHO GTPases activate CIT  
                      RHO GTPases Activate Formins  
                      RHO GTPases activate IQGAPs  
                      RHO GTPases activate PAKs  
                      RHO GTPases activate PKNs  
                      RHO GTPases Activate ROCKs  
                      RHO GTPases Activate WASPs and WAVEs  
                      Role of LAT2NTALLAB on calcium mobilization  
                      S33 mutants of beta-catenin aren.SQT.t phosphorylated  
                      S37 mutants of beta-catenin aren.SQT.t phosphorylated  
                      S45 mutants of beta-catenin aren.SQT.t phosphorylated  
                      Sema3A PAK dependent Axon repulsion  
                      Sema4D induced cell migration and growth-cone collapse  
                      SHC-mediated cascade.CL.FGFR1  
                      SHC-related events triggered by IGF1R  
                      Signal transduction by L1  
                      Signaling by ERBB2  
                      Signaling by FGFR1 in disease  
                      Signaling by FGFR2 in disease  
                      Signaling by FGFR3 in disease  
                      Signaling by FGFR4 in disease  
                      Signaling by PDGF  
                      Signaling by SCF-KIT  
                      Smooth Muscle Contraction  
                      Spry regulation of FGF signaling  
                      Syndecan interactions  
                      Synthesis of PIPs at the plasma membrane  
                      T41 mutants of beta-catenin aren.SQT.t phosphorylated  
                      TGF-beta receptor signaling activates SMADs  
                      TGF-beta receptor signaling in EMT (epithelial to mesenchymal tr...  
                      Tie2 Signaling  
                      Tight junction interactions  
                      TP53 Regulates Metabolic Genes  
                      Truncations of AMER1 destabilize the destruction complex  
                      VEGFA-VEGFR2 Pathway  
                      VEGFR2 mediated cell proliferation  
                      VEGFR2 mediated vascular permeability  
                      WNT mediated activation of DVL  
                  
                   
                  
                    
                      
                           
                          
                               Global Significance Scores   More Plot Information                Global Significance Scores  Heatmap displaying each sample's global significance scores. Global significance statistics measure the extent of differential expression of a gene set's genes with a covariate, ignoring whether each gene within the set is up- or down-regulated. Orange denotes gene sets whose genes exhibit extensive differential expression with the covariate, blue denotes gene sets with less differential expression.   
                               Directed Global Significance Scores   More Plot Information                Directed Global Significance Scores  Heatmap displaying each sample's directed global significance scores. Directed global significance statistics measure the extent to which a gene set's genes are up- or down-regulated with the variable. Red denotes gene sets whose genes exhibit extensive over-expression with the covariate, blue denotes gene sets with extensive under-expression.   
                          
                           
                           
                          
                               Undirected and Directed global significance scores table   More Plot Information      Download CSV Data     
                               
                                 
                                 
                               
                               
                                 
                                   
                                   Undirected New.Annotation: differential expression in Control vs. baseline of cel67 
                                   Undirected New.Annotation: differential expression in miR142 vs. baseline of cel67 
                                   Undirected New.Annotation: differential expression in miR890 vs. baseline of cel67 
                                   Undirected New.Annotation: differential expression in miR18b vs. baseline of cel67 
                                   Undirected New.Annotation: differential expression in miR155 vs. baseline of cel67 
                                   Directed New.Annotation: differential expression in Control vs. baseline of cel67 
                                   Directed New.Annotation: differential expression in miR142 vs. baseline of cel67 
                                   Directed New.Annotation: differential expression in miR890 vs. baseline of cel67 
                                   Directed New.Annotation: differential expression in miR18b vs. baseline of cel67 
                                   Directed New.Annotation: differential expression in miR155 vs. baseline of cel67 
                                 
                               
                               
                                 
                                   Activation of BAD and translocation to mitochondria  
                                   1.846 
                                   2.675 
                                   1.035 
                                   1.979 
                                   2.417 
                                   -1.846 
                                   -2.373 
                                   -0.912 
                                   -1.854 
                                   -2.417 
                                 
                                 
                                   Activation of Rac 
                                   1.813 
                                   9.291 
                                   2.598 
                                   3.622 
                                   4.345 
                                   -1.813 
                                   -9.132 
                                   -2.54 
                                   -3.401 
                                   -3.992 
                                 
                                 
                                   Apoptotic cleavage of cellular proteins 
                                   1.647 
                                   3.151 
                                   4.853 
                                   1.701 
                                   1.954 
                                   1.224 
                                   -1.064 
                                   -4.26 
                                   1.2 
                                   1.507 
                                 
                                 
                                   ARMS-mediated activation 
                                   1.904 
                                   3.122 
                                   3.258 
                                   3.568 
                                   4.118 
                                   -1.169 
                                   -1.941 
                                   -2.31 
                                   -2.283 
                                   -3.796 
                                 
                                 
                                   AXIN missense mutants destabilize the destruction complex 
                                   3.678 
                                   5.447 
                                   3.782 
                                   3.877 
                                   6.254 
                                   2.931 
                                   3.63 
                                   -3.152 
                                   3.376 
                                   -5.992 
                                 
                                 
                                   Beta-catenin phosphorylation cascade 
                                   3.687 
                                   4.723 
                                   3.781 
                                   4.005 
                                   6.282 
                                   2.942 
                                   2.411 
                                   -3.153 
                                   3.522 
                                   -5.962 
                                 
                                 
                                   CD209 (DC-SIGN) signaling 
                                   2.195 
                                   3.265 
                                   3.618 
                                   4.079 
                                   3.834 
                                   -2.151 
                                   -2.121 
                                   -3.551 
                                   -3.838 
                                   -3.772 
                                 
                                 
                                   CD28 co-stimulation 
                                   1.623 
                                   7.652 
                                   2.106 
                                   5.309 
                                   3.059 
                                   -0.999 
                                   -6.986 
                                   -1.455 
                                   -4.804 
                                   -2.884 
                                 
                                 
                                   CD28 dependent PI3KAkt signaling 
                                   1.397 
                                   3.562 
                                   1.239 
                                   3.921 
                                   2.567 
                                   -0.943 
                                   2.879 
                                   -1.156 
                                   -2.546 
                                   -2.053 
                                 
                                 
                                   CD28 dependent Vav1 pathway 
                                   1.05 
                                   8.948 
                                   2.933 
                                   7.455 
                                   3.993 
                                   -0.992 
                                   -8.718 
                                   -1.531 
                                   -7.431 
                                   -3.989 
                                 
                                 
                                   Cell-extracellular matrix interactions 
                                   0.525 
                                   1.198 
                                   0.973 
                                   1.38 
                                   1.319 
                                   0.367 
                                   -0.87 
                                   -0.973 
                                   -0.77 
                                   -0.146 
                                 
                                 
                                   Cell surface interactions at the vascular wall 
                                   2.177 
                                   5.366 
                                   2.357 
                                   3.682 
                                   2.976 
                                   -1.904 
                                   -4.813 
                                   -2.129 
                                   -3.424 
                                   -2.825 
                                 
                                 
                                   Circadian Clock 
                                   1.539 
                                   3.218 
                                   3.167 
                                   3.269 
                                   1.424 
                                   0.72 
                                   -1.911 
                                   -2.791 
                                   -3.142 
                                   -0.927 
                                 
                                 
                                   Constitutive Signaling by Aberrant PI3K in Cancer 
                                   1.806 
                                   2.898 
                                   2.007 
                                   3.259 
                                   3.539 
                                   -1.367 
                                   1.671 
                                   -1.961 
                                   -2.548 
                                   -3.362 
                                 
                                 
                                   Constitutive Signaling by EGFRvIII 
                                   2.921 
                                   4.223 
                                   3.375 
                                   3.021 
                                   5.555 
                                   -2.801 
                                   -1.996 
                                   -3.283 
                                   -2.571 
                                   -5.502 
                                 
                                 
                                   CREB phosphorylation through the activation of Ras 
                                   1.91 
                                   4.368 
                                   2.868 
                                   6.537 
                                   1.949 
                                   -1.891 
                                   -1.57 
                                   -2.421 
                                   -6.537 
                                   -1.658 
                                 
                                 
                                   CTLA4 inhibitory signaling 
                                   1.726 
                                   7.23 
                                   2.535 
                                   2.956 
                                   5.713 
                                   -0.728 
                                   -7.046 
                                   -2.505 
                                   2.322 
                                   -5.673 
                                 
                                 
                                   Cyclin AB1 associated events during G2M transition 
                                   1.579 
                                   3.072 
                                   2.824 
                                   3.287 
                                   6.728 
                                   -1.572 
                                   -2.974 
                                   -2.7 
                                   2.399 
                                   -6.686 
                                 
                                 
                                   Cyclin D associated events in G1 
                                   1.552 
                                   3.209 
                                   5.153 
                                   3.574 
                                   5.473 
                                   -1.45 
                                   -2.725 
                                   -4.944 
                                   -1.509 
                                   -5.343 
                                 
                                 
                                   DAP12 signaling 
                                   1.998 
                                   4.277 
                                   3.11 
                                   3.552 
                                   3.79 
                                   -1.385 
                                   -2.548 
                                   -2.261 
                                   -2.232 
                                   -3.404 
                                 
                               
                               Undirected and Directed global significance scores table  Table displaying each sample's global significance scores and directed global significance scores as defined in the heatmaps above. The global significance score is calculated as the square root of the mean squared t-statistic for the genes in a gene set, with t-statistics coming from the linear regression underlying our differential expression analysis. The directed global significance score is calculated as the square root of the mean signed squared t-statistic for the genes in a gene set, with t-statistics coming from the linear regression underlying our differential expression analysis.   
                          
                           
                      
                    
                    
                      
                           
                          
                              New.Annotation: 
                              
                                  New.Annotation: differential expression in Control vs. baseline of cel67  
                                  New.Annotation: differential expression in miR142 vs. baseline of cel67  
                                  New.Annotation: differential expression in miR155 vs. baseline of cel67  
                                  New.Annotation: differential expression in miR18b vs. baseline of cel67  
                                  New.Annotation: differential expression in miR890 vs. baseline of cel67  
                              
                              
                              
                                
                                  
                                       
                                      
                                           Volcano Plot: New.Annotation: Control vs.cel67   More Plot Information                Volcano Plot: New.Annotation: Control vs.cel67  Volcano plot displaying each gene's -log10(p-value) and log2 fold change for the selected covariate. Highly statistically significant genes fall at the top of the plot, and highly differentially expressed genes fall to either side. Genes within the selected gene set are highlighted in orange. Horizontal lines indicate various False Discovery Rate (FDR) thresholds.   
                                           DE Results - Activation of BAD and translocation to mitochondria  genes - New.Annotation: Control vs.cel67   More Plot Information      Download CSV Data     
                                           
                                             
                                             
                                           
                                           
                                             
                                               Probe Label 
                                               Log2 fold change 
                                               std error (log2) 
                                               Lower confidence limit (log2) 
                                               Upper confidence limit (log2) 
                                               Linear fold change 
                                               Lower confidence limit (linear) 
                                               Upper confidence limit (linear) 
                                               P-value 
                                               BY.p.value 
                                               method 
                                               Gene.sets 
                                               Codeset.Name 
                                               Probe.Label 
                                               Analyte.Type 
                                               Is.Control 
                                               Control.Type 
                                               Related.Probes 
                                               Probe.Annotation 
                                               KEGG.Pathways 
                                               Official.Gene.Name 
                                               Control.Conc 
                                               volcanocol 
                                               volcanopch 
                                               tstats.all 
                                             
                                           
                                           
                                             
                                               AKT2-mRNA 
                                               -0.182 
                                               0.0603 
                                               -0.3 
                                               -0.0634 
                                               0.882 
                                               0.812 
                                               0.957 
                                               0.00666 
                                               0.468 
                                               loglinear 
                                               Activation of BAD and translocation to mitochondria , CD28 co-stimulation, DAP12 signaling, Deactivation of the beta-catenin transactivating complex, Downstream signal transduction, G beta.CL.gamma signalling through PI3Kgamma, GAB1 signalosome, GPVI-mediated activation cascade, IRS-related events triggered by IGF1R, PI3K Cascade, PI3KAKT activation, PIP3 activates AKT signaling, Role of LAT2NTALLAB on calcium mobilization, Signaling by ERBB2, Signaling by PDGF, Signaling by SCF-KIT, TP53 Regulates Metabolic Genes, VEGFA-VEGFR2 Pathway, VEGFR2 mediated vascular permeability 
                                               SEBELIUS_2_C4636 
                                               AKT2 
                                               mRNA 
                                               0 
                                               
                                               
                                               TP53 Regulates Metabolic Genes;PDE3B signalling;AKT phosphorylates targets in the cytosol;Negative regulation of the PI3KAKT network;Deactivation of the beta-catenin transactivating complex;Activation of BAD and translocation to mitochondria ;G beta.CL.gamma signalling through PI3Kgamma;AKT phosphorylates targets in the nucleus;VEGFR2 mediated vascular permeability;GPVI-mediated activation cascade;Signaling by ERBB2;PI3K Cascade;IRS-related events triggered by IGF1R;mTOR signalling;PIP3 activates AKT signaling;Downstream signal transduction;GAB1 signalosome;Signaling by SCF-KIT;PI3KAKT activation;Role of LAT2NTALLAB on calcium mobilization;DAP12 signaling;Signaling by PDGF;CD28 co-stimulation;VEGFA-VEGFR2 Pathway 
                                               hsa04010;hsa04012;hsa04014;hsa04015;hsa04022;hsa04024;hsa04062;hsa04066;hsa04068;hsa04071;hsa04072;hsa04150;hsa04151;hsa04152;hsa04210;hsa04211;hsa04213;hsa04261;hsa04370;hsa04380;hsa04510;hsa04530;hsa04550;hsa04611;hsa04620;hsa04630;hsa04660;hsa04662;hsa04664;hsa04666;hsa04668;hsa04722;hsa04725;hsa04728;hsa04910;hsa04914;hsa04915;hsa04917;hsa04919;hsa04920;hsa04922;hsa04923;hsa04931;hsa04932;hsa04933;hsa04973;hsa05142;hsa05145;hsa05152;hsa05160;hsa05161;hsa05162;hsa05164;hsa05166;hsa05169;hsa05200;hsa05205;hsa05210;hsa05211;hsa05212;hsa05213;hsa05214;hsa05215;hsa05218;hsa05220;hsa05221;hsa05222;hsa05223;hsa05230;hsa05231 
                                               AKT2 
                                               NA 
                                               slateblue4 
                                               16 
                                               -3.01824212271973 
                                             
                                             
                                               BAD-mRNA 
                                               -0.412 
                                               0.18 
                                               -0.765 
                                               -0.0586 
                                               0.752 
                                               0.589 
                                               0.96 
                                               0.0328 
                                               0.908 
                                               lm.nb 
                                               Activation of BAD and translocation to mitochondria , DAP12 signaling, Downstream signal transduction, GAB1 signalosome, NRAGE signals death through JNK, PI3KAKT activation, PIP3 activates AKT signaling, Role of LAT2NTALLAB on calcium mobilization, Signaling by ERBB2, Signaling by PDGF, Signaling by SCF-KIT 
                                               SEBELIUS_2_C4636 
                                               BAD 
                                               mRNA 
                                               0 
                                               
                                               
                                               Activation of BAD and translocation to mitochondria ;NRAGE signals death through JNK;Constitutive Signaling by AKT1 E17K in Cancer;AKT phosphorylates targets in the cytosol;PIP3 activates AKT signaling;Downstream signal transduction;GAB1 signalosome;Signaling by SCF-KIT;PI3KAKT activation;Role of LAT2NTALLAB on calcium mobilization;DAP12 signaling;Signaling by PDGF;Signaling by ERBB2 
                                               hsa04012;hsa04014;hsa04022;hsa04024;hsa04151;hsa04210;hsa04370;hsa04510;hsa04722;hsa04910;hsa04919;hsa05010;hsa05145;hsa05152;hsa05160;hsa05161;hsa05200;hsa05203;hsa05210;hsa05212;hsa05213;hsa05215;hsa05218;hsa05220;hsa05221;hsa05223 
                                               BAD 
                                               NA 
                                               azure4 
                                               1 
                                               -2.28888888888889 
                                             
                                             
                                               AKT1-mRNA 
                                               -0.242 
                                               0.151 
                                               -0.538 
                                               0.0548 
                                               0.846 
                                               0.689 
                                               1.04 
                                               0.125 
                                               1 
                                               lm.nb 
                                               Activation of BAD and translocation to mitochondria , CD28 co-stimulation, DAP12 signaling, Deactivation of the beta-catenin transactivating complex, Downstream signal transduction, G beta.CL.gamma signalling through PI3Kgamma, GAB1 signalosome, GPVI-mediated activation cascade, IRS-related events triggered by IGF1R, PI3K Cascade, PI3KAKT activation, PIP3 activates AKT signaling, Role of LAT2NTALLAB on calcium mobilization, Signaling by ERBB2, Signaling by PDGF, Signaling by SCF-KIT, TP53 Regulates Metabolic Genes, VEGFA-VEGFR2 Pathway, VEGFR2 mediated vascular permeability 
                                               SEBELIUS_2_C4636 
                                               AKT1 
                                               mRNA 
                                               0 
                                               
                                               
                                               TP53 Regulates Metabolic Genes;Deactivation of the beta-catenin transactivating complex;Activation of BAD and translocation to mitochondria ;eNOS activation;mTOR signalling;G beta.CL.gamma signalling through PI3Kgamma;KSRP (KHSRP) binds and destabilizes mRNA;AKT phosphorylates targets in the nucleus;VEGFR2 mediated vascular permeability;GPVI-mediated activation cascade;Tetrahydrobiopterin (BH4) synthesis# recycling# salvage and regulation;Butyrate Response Factor 1 (BRF1) binds and destabilizes mRNA;Signaling by ERBB2;PIP3 activates AKT signaling;Downstream signal transduction;GAB1 signalosome;Signaling by SCF-KIT;PI3KAKT activation;Role of LAT2NTALLAB on calcium mobilization;DAP12 signaling;Signaling by PDGF;CD28 co-stimulation;PI3K Cascade;IRS-related events triggered by IGF1R;VEGFA-VEGFR2 Pathway 
                                               hsa04010;hsa04012;hsa04014;hsa04015;hsa04022;hsa04024;hsa04062;hsa04066;hsa04068;hsa04071;hsa04072;hsa04150;hsa04151;hsa04152;hsa04210;hsa04211;hsa04213;hsa04261;hsa04370;hsa04380;hsa04510;hsa04530;hsa04550;hsa04611;hsa04620;hsa04630;hsa04660;hsa04662;hsa04664;hsa04666;hsa04668;hsa04722;hsa04725;hsa04728;hsa04910;hsa04914;hsa04915;hsa04917;hsa04919;hsa04920;hsa04922;hsa04923;hsa04931;hsa04932;hsa04933;hsa04973;hsa05142;hsa05145;hsa05152;hsa05160;hsa05161;hsa05162;hsa05164;hsa05166;hsa05169;hsa05200;hsa05205;hsa05210;hsa05211;hsa05212;hsa05213;hsa05214;hsa05215;hsa05218;hsa05220;hsa05221;hsa05222;hsa05223;hsa05230;hsa05231 
                                               AKT1 
                                               NA 
                                               azure4 
                                               1 
                                               -1.60264900662252 
                                             
                                             
                                               AKT3-mRNA 
                                               -0.0262 
                                               0.106 
                                               -0.234 
                                               0.182 
                                               0.982 
                                               0.85 
                                               1.13 
                                               0.808 
                                               1 
                                               lm.nb 
                                               Activation of BAD and translocation to mitochondria , CD28 co-stimulation, CD28 dependent PI3KAkt signaling, DAP12 signaling, Downstream signal transduction, G beta.CL.gamma signalling through PI3Kgamma, GAB1 signalosome, GPVI-mediated activation cascade, PI3KAKT activation, PIP3 activates AKT signaling, Role of LAT2NTALLAB on calcium mobilization, Signaling by ERBB2, Signaling by PDGF, Signaling by SCF-KIT, TP53 Regulates Metabolic Genes, VEGFA-VEGFR2 Pathway, VEGFR2 mediated vascular permeability 
                                               SEBELIUS_2_C4636 
                                               AKT3 
                                               mRNA 
                                               0 
                                               
                                               
                                               TP53 Regulates Metabolic Genes;CD28 dependent PI3KAkt signaling;AKT phosphorylates targets in the cytosol;Negative regulation of the PI3KAKT network;Activation of BAD and translocation to mitochondria ;G beta.CL.gamma signalling through PI3Kgamma;AKT phosphorylates targets in the nucleus;VEGFR2 mediated vascular permeability;GPVI-mediated activation cascade;Signaling by ERBB2;PIP3 activates AKT signaling;Downstream signal transduction;GAB1 signalosome;Signaling by SCF-KIT;PI3KAKT activation;Role of LAT2NTALLAB on calcium mobilization;DAP12 signaling;Signaling by PDGF;CD28 co-stimulation;VEGFA-VEGFR2 Pathway 
                                               hsa04010;hsa04012;hsa04014;hsa04015;hsa04022;hsa04024;hsa04062;hsa04066;hsa04068;hsa04071;hsa04072;hsa04150;hsa04151;hsa04152;hsa04210;hsa04211;hsa04213;hsa04261;hsa04370;hsa04380;hsa04510;hsa04530;hsa04550;hsa04611;hsa04620;hsa04630;hsa04660;hsa04662;hsa04664;hsa04666;hsa04668;hsa04722;hsa04725;hsa04728;hsa04910;hsa04914;hsa04915;hsa04917;hsa04919;hsa04920;hsa04922;hsa04923;hsa04931;hsa04932;hsa04933;hsa04973;hsa05142;hsa05145;hsa05152;hsa05160;hsa05161;hsa05162;hsa05164;hsa05166;hsa05169;hsa05200;hsa05205;hsa05210;hsa05211;hsa05212;hsa05213;hsa05214;hsa05215;hsa05218;hsa05220;hsa05221;hsa05222;hsa05223;hsa05230;hsa05231 
                                               AKT3 
                                               NA 
                                               azure4 
                                               1 
                                               -0.247169811320755 
                                             
                                             
                                               BCL2-mRNA 
                                               -0.875 
                                               3.57 
                                               -7.86 
                                               6.11 
                                               0.545 
                                               0.0043 
                                               69.2 
                                               0.809 
                                               1 
                                               Wald 
                                               Activation of BAD and translocation to mitochondria  
                                               SEBELIUS_2_C4636 
                                               BCL2 
                                               mRNA 
                                               0 
                                               
                                               
                                               Activation of BAD and translocation to mitochondria  
                                               hsa04064;hsa04066;hsa04071;hsa04141;hsa04151;hsa04210;hsa04261;hsa04510;hsa04630;hsa04722;hsa04725;hsa04933;hsa05145;hsa05152;hsa05161;hsa05169;hsa05200;hsa05206;hsa05210;hsa05215;hsa05222 
                                               BCL2 
                                               NA 
                                               azure4 
                                               1 
                                               -0.245098039215686 
                                             
                                           
                                           DE Results - Activation of BAD and translocation to mitochondria  genes - New.Annotation: Control vs.cel67  Table displaying each sample's global significance scores and directed global significance scores as defined in the heatmaps above. The global significance score is calculated as the square root of the mean squared t-statistic for the genes in a gene set, with t-statistics coming from the linear regression underlying our differential expression analysis. The directed global significance score is calculated as the square root of the mean signed squared t-statistic for the genes in a gene set, with t-statistics coming from the linear regression underlying our differential expression analysis.   
                                      
                                       
                                  
                                
                                
                                  
                                       
                                      
                                           Volcano Plot: New.Annotation: miR142 vs.cel67   More Plot Information                Volcano Plot: New.Annotation: miR142 vs.cel67  Volcano plot displaying each gene's -log10(p-value) and log2 fold change for the selected covariate. Highly statistically significant genes fall at the top of the plot, and highly differentially expressed genes fall to either side. Genes within the selected gene set are highlighted in orange. Horizontal lines indicate various False Discovery Rate (FDR) thresholds.   
                                           DE Results - Activation of BAD and translocation to mitochondria  genes - New.Annotation: miR142 vs.cel67   More Plot Information      Download CSV Data     
                                           
                                             
                                             
                                           
                                           
                                             
                                               Probe Label 
                                               Log2 fold change 
                                               std error (log2) 
                                               Lower confidence limit (log2) 
                                               Upper confidence limit (log2) 
                                               Linear fold change 
                                               Lower confidence limit (linear) 
                                               Upper confidence limit (linear) 
                                               P-value 
                                               BY.p.value 
                                               method 
                                               Gene.sets 
                                               Codeset.Name 
                                               Probe.Label 
                                               Analyte.Type 
                                               Is.Control 
                                               Control.Type 
                                               Related.Probes 
                                               Probe.Annotation 
                                               KEGG.Pathways 
                                               Official.Gene.Name 
                                               Control.Conc 
                                               volcanocol 
                                               volcanopch 
                                               tstats.all 
                                             
                                           
                                           
                                             
                                               BAD-mRNA 
                                               -0.668 
                                               0.182 
                                               -1.02 
                                               -0.311 
                                               0.629 
                                               0.492 
                                               0.806 
                                               0.00142 
                                               0.0294 
                                               lm.nb 
                                               Activation of BAD and translocation to mitochondria , DAP12 signaling, Downstream signal transduction, GAB1 signalosome, NRAGE signals death through JNK, PI3KAKT activation, PIP3 activates AKT signaling, Role of LAT2NTALLAB on calcium mobilization, Signaling by ERBB2, Signaling by PDGF, Signaling by SCF-KIT 
                                               SEBELIUS_2_C4636 
                                               BAD 
                                               mRNA 
                                               0 
                                               
                                               
                                               Activation of BAD and translocation to mitochondria ;NRAGE signals death through JNK;Constitutive Signaling by AKT1 E17K in Cancer;AKT phosphorylates targets in the cytosol;PIP3 activates AKT signaling;Downstream signal transduction;GAB1 signalosome;Signaling by SCF-KIT;PI3KAKT activation;Role of LAT2NTALLAB on calcium mobilization;DAP12 signaling;Signaling by PDGF;Signaling by ERBB2 
                                               hsa04012;hsa04014;hsa04022;hsa04024;hsa04151;hsa04210;hsa04370;hsa04510;hsa04722;hsa04910;hsa04919;hsa05010;hsa05145;hsa05152;hsa05160;hsa05161;hsa05200;hsa05203;hsa05210;hsa05212;hsa05213;hsa05215;hsa05218;hsa05220;hsa05221;hsa05223 
                                               BAD 
                                               NA 
                                               slateblue2 
                                               16 
                                               -3.67032967032967 
                                             
                                             
                                               AKT2-mRNA 
                                               -0.2 
                                               0.0603 
                                               -0.318 
                                               -0.0816 
                                               0.871 
                                               0.802 
                                               0.945 
                                               0.00331 
                                               0.0593 
                                               loglinear 
                                               Activation of BAD and translocation to mitochondria , CD28 co-stimulation, DAP12 signaling, Deactivation of the beta-catenin transactivating complex, Downstream signal transduction, G beta.CL.gamma signalling through PI3Kgamma, GAB1 signalosome, GPVI-mediated activation cascade, IRS-related events triggered by IGF1R, PI3K Cascade, PI3KAKT activation, PIP3 activates AKT signaling, Role of LAT2NTALLAB on calcium mobilization, Signaling by ERBB2, Signaling by PDGF, Signaling by SCF-KIT, TP53 Regulates Metabolic Genes, VEGFA-VEGFR2 Pathway, VEGFR2 mediated vascular permeability 
                                               SEBELIUS_2_C4636 
                                               AKT2 
                                               mRNA 
                                               0 
                                               
                                               
                                               TP53 Regulates Metabolic Genes;PDE3B signalling;AKT phosphorylates targets in the cytosol;Negative regulation of the PI3KAKT network;Deactivation of the beta-catenin transactivating complex;Activation of BAD and translocation to mitochondria ;G beta.CL.gamma signalling through PI3Kgamma;AKT phosphorylates targets in the nucleus;VEGFR2 mediated vascular permeability;GPVI-mediated activation cascade;Signaling by ERBB2;PI3K Cascade;IRS-related events triggered by IGF1R;mTOR signalling;PIP3 activates AKT signaling;Downstream signal transduction;GAB1 signalosome;Signaling by SCF-KIT;PI3KAKT activation;Role of LAT2NTALLAB on calcium mobilization;DAP12 signaling;Signaling by PDGF;CD28 co-stimulation;VEGFA-VEGFR2 Pathway 
                                               hsa04010;hsa04012;hsa04014;hsa04015;hsa04022;hsa04024;hsa04062;hsa04066;hsa04068;hsa04071;hsa04072;hsa04150;hsa04151;hsa04152;hsa04210;hsa04211;hsa04213;hsa04261;hsa04370;hsa04380;hsa04510;hsa04530;hsa04550;hsa04611;hsa04620;hsa04630;hsa04660;hsa04662;hsa04664;hsa04666;hsa04668;hsa04722;hsa04725;hsa04728;hsa04910;hsa04914;hsa04915;hsa04917;hsa04919;hsa04920;hsa04922;hsa04923;hsa04931;hsa04932;hsa04933;hsa04973;hsa05142;hsa05145;hsa05152;hsa05160;hsa05161;hsa05162;hsa05164;hsa05166;hsa05169;hsa05200;hsa05205;hsa05210;hsa05211;hsa05212;hsa05213;hsa05214;hsa05215;hsa05218;hsa05220;hsa05221;hsa05222;hsa05223;hsa05230;hsa05231 
                                               AKT2 
                                               NA 
                                               slateblue3 
                                               16 
                                               -3.3167495854063 
                                             
                                             
                                               AKT1-mRNA 
                                               -0.414 
                                               0.152 
                                               -0.711 
                                               -0.117 
                                               0.751 
                                               0.611 
                                               0.922 
                                               0.0125 
                                               0.18 
                                               lm.nb 
                                               Activation of BAD and translocation to mitochondria , CD28 co-stimulation, DAP12 signaling, Deactivation of the beta-catenin transactivating complex, Downstream signal transduction, G beta.CL.gamma signalling through PI3Kgamma, GAB1 signalosome, GPVI-mediated activation cascade, IRS-related events triggered by IGF1R, PI3K Cascade, PI3KAKT activation, PIP3 activates AKT signaling, Role of LAT2NTALLAB on calcium mobilization, Signaling by ERBB2, Signaling by PDGF, Signaling by SCF-KIT, TP53 Regulates Metabolic Genes, VEGFA-VEGFR2 Pathway, VEGFR2 mediated vascular permeability 
                                               SEBELIUS_2_C4636 
                                               AKT1 
                                               mRNA 
                                               0 
                                               
                                               
                                               TP53 Regulates Metabolic Genes;Deactivation of the beta-catenin transactivating complex;Activation of BAD and translocation to mitochondria ;eNOS activation;mTOR signalling;G beta.CL.gamma signalling through PI3Kgamma;KSRP (KHSRP) binds and destabilizes mRNA;AKT phosphorylates targets in the nucleus;VEGFR2 mediated vascular permeability;GPVI-mediated activation cascade;Tetrahydrobiopterin (BH4) synthesis# recycling# salvage and regulation;Butyrate Response Factor 1 (BRF1) binds and destabilizes mRNA;Signaling by ERBB2;PIP3 activates AKT signaling;Downstream signal transduction;GAB1 signalosome;Signaling by SCF-KIT;PI3KAKT activation;Role of LAT2NTALLAB on calcium mobilization;DAP12 signaling;Signaling by PDGF;CD28 co-stimulation;PI3K Cascade;IRS-related events triggered by IGF1R;VEGFA-VEGFR2 Pathway 
                                               hsa04010;hsa04012;hsa04014;hsa04015;hsa04022;hsa04024;hsa04062;hsa04066;hsa04068;hsa04071;hsa04072;hsa04150;hsa04151;hsa04152;hsa04210;hsa04211;hsa04213;hsa04261;hsa04370;hsa04380;hsa04510;hsa04530;hsa04550;hsa04611;hsa04620;hsa04630;hsa04660;hsa04662;hsa04664;hsa04666;hsa04668;hsa04722;hsa04725;hsa04728;hsa04910;hsa04914;hsa04915;hsa04917;hsa04919;hsa04920;hsa04922;hsa04923;hsa04931;hsa04932;hsa04933;hsa04973;hsa05142;hsa05145;hsa05152;hsa05160;hsa05161;hsa05162;hsa05164;hsa05166;hsa05169;hsa05200;hsa05205;hsa05210;hsa05211;hsa05212;hsa05213;hsa05214;hsa05215;hsa05218;hsa05220;hsa05221;hsa05222;hsa05223;hsa05230;hsa05231 
                                               AKT1 
                                               NA 
                                               slateblue4 
                                               16 
                                               -2.72368421052632 
                                             
                                             
                                               AKT3-mRNA 
                                               0.207 
                                               0.106 
                                               -0.000741 
                                               0.415 
                                               1.15 
                                               0.999 
                                               1.33 
                                               0.0643 
                                               0.723 
                                               lm.nb 
                                               Activation of BAD and translocation to mitochondria , CD28 co-stimulation, CD28 dependent PI3KAkt signaling, DAP12 signaling, Downstream signal transduction, G beta.CL.gamma signalling through PI3Kgamma, GAB1 signalosome, GPVI-mediated activation cascade, PI3KAKT activation, PIP3 activates AKT signaling, Role of LAT2NTALLAB on calcium mobilization, Signaling by ERBB2, Signaling by PDGF, Signaling by SCF-KIT, TP53 Regulates Metabolic Genes, VEGFA-VEGFR2 Pathway, VEGFR2 mediated vascular permeability 
                                               SEBELIUS_2_C4636 
                                               AKT3 
                                               mRNA 
                                               0 
                                               
                                               
                                               TP53 Regulates Metabolic Genes;CD28 dependent PI3KAkt signaling;AKT phosphorylates targets in the cytosol;Negative regulation of the PI3KAKT network;Activation of BAD and translocation to mitochondria ;G beta.CL.gamma signalling through PI3Kgamma;AKT phosphorylates targets in the nucleus;VEGFR2 mediated vascular permeability;GPVI-mediated activation cascade;Signaling by ERBB2;PIP3 activates AKT signaling;Downstream signal transduction;GAB1 signalosome;Signaling by SCF-KIT;PI3KAKT activation;Role of LAT2NTALLAB on calcium mobilization;DAP12 signaling;Signaling by PDGF;CD28 co-stimulation;VEGFA-VEGFR2 Pathway 
                                               hsa04010;hsa04012;hsa04014;hsa04015;hsa04022;hsa04024;hsa04062;hsa04066;hsa04068;hsa04071;hsa04072;hsa04150;hsa04151;hsa04152;hsa04210;hsa04211;hsa04213;hsa04261;hsa04370;hsa04380;hsa04510;hsa04530;hsa04550;hsa04611;hsa04620;hsa04630;hsa04660;hsa04662;hsa04664;hsa04666;hsa04668;hsa04722;hsa04725;hsa04728;hsa04910;hsa04914;hsa04915;hsa04917;hsa04919;hsa04920;hsa04922;hsa04923;hsa04931;hsa04932;hsa04933;hsa04973;hsa05142;hsa05145;hsa05152;hsa05160;hsa05161;hsa05162;hsa05164;hsa05166;hsa05169;hsa05200;hsa05205;hsa05210;hsa05211;hsa05212;hsa05213;hsa05214;hsa05215;hsa05218;hsa05220;hsa05221;hsa05222;hsa05223;hsa05230;hsa05231 
                                               AKT3 
                                               NA 
                                               azure4 
                                               1 
                                               1.95283018867925 
                                             
                                             
                                               BCL2-mRNA 
                                               -0.925 
                                               3.56 
                                               -7.9 
                                               6.05 
                                               0.527 
                                               0.00418 
                                               66.3 
                                               0.798 
                                               1 
                                               Wald 
                                               Activation of BAD and translocation to mitochondria  
                                               SEBELIUS_2_C4636 
                                               BCL2 
                                               mRNA 
                                               0 
                                               
                                               
                                               Activation of BAD and translocation to mitochondria  
                                               hsa04064;hsa04066;hsa04071;hsa04141;hsa04151;hsa04210;hsa04261;hsa04510;hsa04630;hsa04722;hsa04725;hsa04933;hsa05145;hsa05152;hsa05161;hsa05169;hsa05200;hsa05206;hsa05210;hsa05215;hsa05222 
                                               BCL2 
                                               NA 
                                               azure4 
                                               1 
                                               -0.259831460674157 
                                             
                                           
                                           DE Results - Activation of BAD and translocation to mitochondria  genes - New.Annotation: miR142 vs.cel67  Table displaying each sample's global significance scores and directed global significance scores as defined in the heatmaps above. The global significance score is calculated as the square root of the mean squared t-statistic for the genes in a gene set, with t-statistics coming from the linear regression underlying our differential expression analysis. The directed global significance score is calculated as the square root of the mean signed squared t-statistic for the genes in a gene set, with t-statistics coming from the linear regression underlying our differential expression analysis.   
                                      
                                       
                                  
                                
                                
                                  
                                       
                                      
                                           Volcano Plot: New.Annotation: miR155 vs.cel67   More Plot Information                Volcano Plot: New.Annotation: miR155 vs.cel67  Volcano plot displaying each gene's -log10(p-value) and log2 fold change for the selected covariate. Highly statistically significant genes fall at the top of the plot, and highly differentially expressed genes fall to either side. Genes within the selected gene set are highlighted in orange. Horizontal lines indicate various False Discovery Rate (FDR) thresholds.   
                                           DE Results - Activation of BAD and translocation to mitochondria  genes - New.Annotation: miR155 vs.cel67   More Plot Information      Download CSV Data     
                                           
                                             
                                             
                                           
                                           
                                             
                                               Probe Label 
                                               Log2 fold change 
                                               std error (log2) 
                                               Lower confidence limit (log2) 
                                               Upper confidence limit (log2) 
                                               Linear fold change 
                                               Lower confidence limit (linear) 
                                               Upper confidence limit (linear) 
                                               P-value 
                                               BY.p.value 
                                               method 
                                               Gene.sets 
                                               Codeset.Name 
                                               Probe.Label 
                                               Analyte.Type 
                                               Is.Control 
                                               Control.Type 
                                               Related.Probes 
                                               Probe.Annotation 
                                               KEGG.Pathways 
                                               Official.Gene.Name 
                                               Control.Conc 
                                               volcanocol 
                                               volcanopch 
                                               tstats.all 
                                             
                                           
                                           
                                             
                                               AKT3-mRNA 
                                               -0.395 
                                               0.112 
                                               -0.616 
                                               -0.175 
                                               0.76 
                                               0.653 
                                               0.886 
                                               0.00206 
                                               0.0829 
                                               lm.nb 
                                               Activation of BAD and translocation to mitochondria , CD28 co-stimulation, CD28 dependent PI3KAkt signaling, DAP12 signaling, Downstream signal transduction, G beta.CL.gamma signalling through PI3Kgamma, GAB1 signalosome, GPVI-mediated activation cascade, PI3KAKT activation, PIP3 activates AKT signaling, Role of LAT2NTALLAB on calcium mobilization, Signaling by ERBB2, Signaling by PDGF, Signaling by SCF-KIT, TP53 Regulates Metabolic Genes, VEGFA-VEGFR2 Pathway, VEGFR2 mediated vascular permeability 
                                               SEBELIUS_2_C4636 
                                               AKT3 
                                               mRNA 
                                               0 
                                               
                                               
                                               TP53 Regulates Metabolic Genes;CD28 dependent PI3KAkt signaling;AKT phosphorylates targets in the cytosol;Negative regulation of the PI3KAKT network;Activation of BAD and translocation to mitochondria ;G beta.CL.gamma signalling through PI3Kgamma;AKT phosphorylates targets in the nucleus;VEGFR2 mediated vascular permeability;GPVI-mediated activation cascade;Signaling by ERBB2;PIP3 activates AKT signaling;Downstream signal transduction;GAB1 signalosome;Signaling by SCF-KIT;PI3KAKT activation;Role of LAT2NTALLAB on calcium mobilization;DAP12 signaling;Signaling by PDGF;CD28 co-stimulation;VEGFA-VEGFR2 Pathway 
                                               hsa04010;hsa04012;hsa04014;hsa04015;hsa04022;hsa04024;hsa04062;hsa04066;hsa04068;hsa04071;hsa04072;hsa04150;hsa04151;hsa04152;hsa04210;hsa04211;hsa04213;hsa04261;hsa04370;hsa04380;hsa04510;hsa04530;hsa04550;hsa04611;hsa04620;hsa04630;hsa04660;hsa04662;hsa04664;hsa04666;hsa04668;hsa04722;hsa04725;hsa04728;hsa04910;hsa04914;hsa04915;hsa04917;hsa04919;hsa04920;hsa04922;hsa04923;hsa04931;hsa04932;hsa04933;hsa04973;hsa05142;hsa05145;hsa05152;hsa05160;hsa05161;hsa05162;hsa05164;hsa05166;hsa05169;hsa05200;hsa05205;hsa05210;hsa05211;hsa05212;hsa05213;hsa05214;hsa05215;hsa05218;hsa05220;hsa05221;hsa05222;hsa05223;hsa05230;hsa05231 
                                               AKT3 
                                               NA 
                                               slateblue3 
                                               16 
                                               -3.52678571428571 
                                             
                                             
                                               AKT2-mRNA 
                                               -0.187 
                                               0.0636 
                                               -0.311 
                                               -0.062 
                                               0.879 
                                               0.806 
                                               0.958 
                                               0.00791 
                                               0.21 
                                               loglinear 
                                               Activation of BAD and translocation to mitochondria , CD28 co-stimulation, DAP12 signaling, Deactivation of the beta-catenin transactivating complex, Downstream signal transduction, G beta.CL.gamma signalling through PI3Kgamma, GAB1 signalosome, GPVI-mediated activation cascade, IRS-related events triggered by IGF1R, PI3K Cascade, PI3KAKT activation, PIP3 activates AKT signaling, Role of LAT2NTALLAB on calcium mobilization, Signaling by ERBB2, Signaling by PDGF, Signaling by SCF-KIT, TP53 Regulates Metabolic Genes, VEGFA-VEGFR2 Pathway, VEGFR2 mediated vascular permeability 
                                               SEBELIUS_2_C4636 
                                               AKT2 
                                               mRNA 
                                               0 
                                               
                                               
                                               TP53 Regulates Metabolic Genes;PDE3B signalling;AKT phosphorylates targets in the cytosol;Negative regulation of the PI3KAKT network;Deactivation of the beta-catenin transactivating complex;Activation of BAD and translocation to mitochondria ;G beta.CL.gamma signalling through PI3Kgamma;AKT phosphorylates targets in the nucleus;VEGFR2 mediated vascular permeability;GPVI-mediated activation cascade;Signaling by ERBB2;PI3K Cascade;IRS-related events triggered by IGF1R;mTOR signalling;PIP3 activates AKT signaling;Downstream signal transduction;GAB1 signalosome;Signaling by SCF-KIT;PI3KAKT activation;Role of LAT2NTALLAB on calcium mobilization;DAP12 signaling;Signaling by PDGF;CD28 co-stimulation;VEGFA-VEGFR2 Pathway 
                                               hsa04010;hsa04012;hsa04014;hsa04015;hsa04022;hsa04024;hsa04062;hsa04066;hsa04068;hsa04071;hsa04072;hsa04150;hsa04151;hsa04152;hsa04210;hsa04211;hsa04213;hsa04261;hsa04370;hsa04380;hsa04510;hsa04530;hsa04550;hsa04611;hsa04620;hsa04630;hsa04660;hsa04662;hsa04664;hsa04666;hsa04668;hsa04722;hsa04725;hsa04728;hsa04910;hsa04914;hsa04915;hsa04917;hsa04919;hsa04920;hsa04922;hsa04923;hsa04931;hsa04932;hsa04933;hsa04973;hsa05142;hsa05145;hsa05152;hsa05160;hsa05161;hsa05162;hsa05164;hsa05166;hsa05169;hsa05200;hsa05205;hsa05210;hsa05211;hsa05212;hsa05213;hsa05214;hsa05215;hsa05218;hsa05220;hsa05221;hsa05222;hsa05223;hsa05230;hsa05231 
                                               AKT2 
                                               NA 
                                               slateblue4 
                                               16 
                                               -2.94025157232704 
                                             
                                             
                                               BAD-mRNA 
                                               -0.499 
                                               0.191 
                                               -0.874 
                                               -0.125 
                                               0.707 
                                               0.546 
                                               0.917 
                                               0.0161 
                                               0.32 
                                               lm.nb 
                                               Activation of BAD and translocation to mitochondria , DAP12 signaling, Downstream signal transduction, GAB1 signalosome, NRAGE signals death through JNK, PI3KAKT activation, PIP3 activates AKT signaling, Role of LAT2NTALLAB on calcium mobilization, Signaling by ERBB2, Signaling by PDGF, Signaling by SCF-KIT 
                                               SEBELIUS_2_C4636 
                                               BAD 
                                               mRNA 
                                               0 
                                               
                                               
                                               Activation of BAD and translocation to mitochondria ;NRAGE signals death through JNK;Constitutive Signaling by AKT1 E17K in Cancer;AKT phosphorylates targets in the cytosol;PIP3 activates AKT signaling;Downstream signal transduction;GAB1 signalosome;Signaling by SCF-KIT;PI3KAKT activation;Role of LAT2NTALLAB on calcium mobilization;DAP12 signaling;Signaling by PDGF;Signaling by ERBB2 
                                               hsa04012;hsa04014;hsa04022;hsa04024;hsa04151;hsa04210;hsa04370;hsa04510;hsa04722;hsa04910;hsa04919;hsa05010;hsa05145;hsa05152;hsa05160;hsa05161;hsa05200;hsa05203;hsa05210;hsa05212;hsa05213;hsa05215;hsa05218;hsa05220;hsa05221;hsa05223 
                                               BAD 
                                               NA 
                                               slateblue4 
                                               16 
                                               -2.61256544502618 
                                             
                                             
                                               AKT1-mRNA 
                                               -0.165 
                                               0.159 
                                               -0.478 
                                               0.147 
                                               0.892 
                                               0.718 
                                               1.11 
                                               0.311 
                                               1 
                                               lm.nb 
                                               Activation of BAD and translocation to mitochondria , CD28 co-stimulation, DAP12 signaling, Deactivation of the beta-catenin transactivating complex, Downstream signal transduction, G beta.CL.gamma signalling through PI3Kgamma, GAB1 signalosome, GPVI-mediated activation cascade, IRS-related events triggered by IGF1R, PI3K Cascade, PI3KAKT activation, PIP3 activates AKT signaling, Role of LAT2NTALLAB on calcium mobilization, Signaling by ERBB2, Signaling by PDGF, Signaling by SCF-KIT, TP53 Regulates Metabolic Genes, VEGFA-VEGFR2 Pathway, VEGFR2 mediated vascular permeability 
                                               SEBELIUS_2_C4636 
                                               AKT1 
                                               mRNA 
                                               0 
                                               
                                               
                                               TP53 Regulates Metabolic Genes;Deactivation of the beta-catenin transactivating complex;Activation of BAD and translocation to mitochondria ;eNOS activation;mTOR signalling;G beta.CL.gamma signalling through PI3Kgamma;KSRP (KHSRP) binds and destabilizes mRNA;AKT phosphorylates targets in the nucleus;VEGFR2 mediated vascular permeability;GPVI-mediated activation cascade;Tetrahydrobiopterin (BH4) synthesis# recycling# salvage and regulation;Butyrate Response Factor 1 (BRF1) binds and destabilizes mRNA;Signaling by ERBB2;PIP3 activates AKT signaling;Downstream signal transduction;GAB1 signalosome;Signaling by SCF-KIT;PI3KAKT activation;Role of LAT2NTALLAB on calcium mobilization;DAP12 signaling;Signaling by PDGF;CD28 co-stimulation;PI3K Cascade;IRS-related events triggered by IGF1R;VEGFA-VEGFR2 Pathway 
                                               hsa04010;hsa04012;hsa04014;hsa04015;hsa04022;hsa04024;hsa04062;hsa04066;hsa04068;hsa04071;hsa04072;hsa04150;hsa04151;hsa04152;hsa04210;hsa04211;hsa04213;hsa04261;hsa04370;hsa04380;hsa04510;hsa04530;hsa04550;hsa04611;hsa04620;hsa04630;hsa04660;hsa04662;hsa04664;hsa04666;hsa04668;hsa04722;hsa04725;hsa04728;hsa04910;hsa04914;hsa04915;hsa04917;hsa04919;hsa04920;hsa04922;hsa04923;hsa04931;hsa04932;hsa04933;hsa04973;hsa05142;hsa05145;hsa05152;hsa05160;hsa05161;hsa05162;hsa05164;hsa05166;hsa05169;hsa05200;hsa05205;hsa05210;hsa05211;hsa05212;hsa05213;hsa05214;hsa05215;hsa05218;hsa05220;hsa05221;hsa05222;hsa05223;hsa05230;hsa05231 
                                               AKT1 
                                               NA 
                                               azure4 
                                               1 
                                               -1.0377358490566 
                                             
                                             
                                               BCL2-mRNA 
                                               -1.73 
                                               3.77 
                                               -9.12 
                                               5.66 
                                               0.301 
                                               0.0018 
                                               50.5 
                                               0.651 
                                               1 
                                               Wald 
                                               Activation of BAD and translocation to mitochondria  
                                               SEBELIUS_2_C4636 
                                               BCL2 
                                               mRNA 
                                               0 
                                               
                                               
                                               Activation of BAD and translocation to mitochondria  
                                               hsa04064;hsa04066;hsa04071;hsa04141;hsa04151;hsa04210;hsa04261;hsa04510;hsa04630;hsa04722;hsa04725;hsa04933;hsa05145;hsa05152;hsa05161;hsa05169;hsa05200;hsa05206;hsa05210;hsa05215;hsa05222 
                                               BCL2 
                                               NA 
                                               azure4 
                                               1 
                                               -0.458885941644562 
                                             
                                           
                                           DE Results - Activation of BAD and translocation to mitochondria  genes - New.Annotation: miR155 vs.cel67  Table displaying each sample's global significance scores and directed global significance scores as defined in the heatmaps above. The global significance score is calculated as the square root of the mean squared t-statistic for the genes in a gene set, with t-statistics coming from the linear regression underlying our differential expression analysis. The directed global significance score is calculated as the square root of the mean signed squared t-statistic for the genes in a gene set, with t-statistics coming from the linear regression underlying our differential expression analysis.   
                                      
                                       
                                  
                                
                                
                                  
                                       
                                      
                                           Volcano Plot: New.Annotation: miR18b vs.cel67   More Plot Information                Volcano Plot: New.Annotation: miR18b vs.cel67  Volcano plot displaying each gene's -log10(p-value) and log2 fold change for the selected covariate. Highly statistically significant genes fall at the top of the plot, and highly differentially expressed genes fall to either side. Genes within the selected gene set are highlighted in orange. Horizontal lines indicate various False Discovery Rate (FDR) thresholds.   
                                           DE Results - Activation of BAD and translocation to mitochondria  genes - New.Annotation: miR18b vs.cel67   More Plot Information      Download CSV Data     
                                           
                                             
                                             
                                           
                                           
                                             
                                               Probe Label 
                                               Log2 fold change 
                                               std error (log2) 
                                               Lower confidence limit (log2) 
                                               Upper confidence limit (log2) 
                                               Linear fold change 
                                               Lower confidence limit (linear) 
                                               Upper confidence limit (linear) 
                                               P-value 
                                               BY.p.value 
                                               method 
                                               Gene.sets 
                                               Codeset.Name 
                                               Probe.Label 
                                               Analyte.Type 
                                               Is.Control 
                                               Control.Type 
                                               Related.Probes 
                                               Probe.Annotation 
                                               KEGG.Pathways 
                                               Official.Gene.Name 
                                               Control.Conc 
                                               volcanocol 
                                               volcanopch 
                                               tstats.all 
                                             
                                           
                                           
                                             
                                               AKT1-mRNA 
                                               -0.697 
                                               0.173 
                                               -1.04 
                                               -0.358 
                                               0.617 
                                               0.487 
                                               0.78 
                                               0.000613 
                                               0.0164 
                                               lm.nb 
                                               Activation of BAD and translocation to mitochondria , CD28 co-stimulation, DAP12 signaling, Deactivation of the beta-catenin transactivating complex, Downstream signal transduction, G beta.CL.gamma signalling through PI3Kgamma, GAB1 signalosome, GPVI-mediated activation cascade, IRS-related events triggered by IGF1R, PI3K Cascade, PI3KAKT activation, PIP3 activates AKT signaling, Role of LAT2NTALLAB on calcium mobilization, Signaling by ERBB2, Signaling by PDGF, Signaling by SCF-KIT, TP53 Regulates Metabolic Genes, VEGFA-VEGFR2 Pathway, VEGFR2 mediated vascular permeability 
                                               SEBELIUS_2_C4636 
                                               AKT1 
                                               mRNA 
                                               0 
                                               
                                               
                                               TP53 Regulates Metabolic Genes;Deactivation of the beta-catenin transactivating complex;Activation of BAD and translocation to mitochondria ;eNOS activation;mTOR signalling;G beta.CL.gamma signalling through PI3Kgamma;KSRP (KHSRP) binds and destabilizes mRNA;AKT phosphorylates targets in the nucleus;VEGFR2 mediated vascular permeability;GPVI-mediated activation cascade;Tetrahydrobiopterin (BH4) synthesis# recycling# salvage and regulation;Butyrate Response Factor 1 (BRF1) binds and destabilizes mRNA;Signaling by ERBB2;PIP3 activates AKT signaling;Downstream signal transduction;GAB1 signalosome;Signaling by SCF-KIT;PI3KAKT activation;Role of LAT2NTALLAB on calcium mobilization;DAP12 signaling;Signaling by PDGF;CD28 co-stimulation;PI3K Cascade;IRS-related events triggered by IGF1R;VEGFA-VEGFR2 Pathway 
                                               hsa04010;hsa04012;hsa04014;hsa04015;hsa04022;hsa04024;hsa04062;hsa04066;hsa04068;hsa04071;hsa04072;hsa04150;hsa04151;hsa04152;hsa04210;hsa04211;hsa04213;hsa04261;hsa04370;hsa04380;hsa04510;hsa04530;hsa04550;hsa04611;hsa04620;hsa04630;hsa04660;hsa04662;hsa04664;hsa04666;hsa04668;hsa04722;hsa04725;hsa04728;hsa04910;hsa04914;hsa04915;hsa04917;hsa04919;hsa04920;hsa04922;hsa04923;hsa04931;hsa04932;hsa04933;hsa04973;hsa05142;hsa05145;hsa05152;hsa05160;hsa05161;hsa05162;hsa05164;hsa05166;hsa05169;hsa05200;hsa05205;hsa05210;hsa05211;hsa05212;hsa05213;hsa05214;hsa05215;hsa05218;hsa05220;hsa05221;hsa05222;hsa05223;hsa05230;hsa05231 
                                               AKT1 
                                               NA 
                                               slateblue2 
                                               16 
                                               -4.02890173410405 
                                             
                                             
                                               BAD-mRNA 
                                               -0.287 
                                               0.205 
                                               -0.688 
                                               0.114 
                                               0.82 
                                               0.621 
                                               1.08 
                                               0.176 
                                               1 
                                               lm.nb 
                                               Activation of BAD and translocation to mitochondria , DAP12 signaling, Downstream signal transduction, GAB1 signalosome, NRAGE signals death through JNK, PI3KAKT activation, PIP3 activates AKT signaling, Role of LAT2NTALLAB on calcium mobilization, Signaling by ERBB2, Signaling by PDGF, Signaling by SCF-KIT 
                                               SEBELIUS_2_C4636 
                                               BAD 
                                               mRNA 
                                               0 
                                               
                                               
                                               Activation of BAD and translocation to mitochondria ;NRAGE signals death through JNK;Constitutive Signaling by AKT1 E17K in Cancer;AKT phosphorylates targets in the cytosol;PIP3 activates AKT signaling;Downstream signal transduction;GAB1 signalosome;Signaling by SCF-KIT;PI3KAKT activation;Role of LAT2NTALLAB on calcium mobilization;DAP12 signaling;Signaling by PDGF;Signaling by ERBB2 
                                               hsa04012;hsa04014;hsa04022;hsa04024;hsa04151;hsa04210;hsa04370;hsa04510;hsa04722;hsa04910;hsa04919;hsa05010;hsa05145;hsa05152;hsa05160;hsa05161;hsa05200;hsa05203;hsa05210;hsa05212;hsa05213;hsa05215;hsa05218;hsa05220;hsa05221;hsa05223 
                                               BAD 
                                               NA 
                                               azure4 
                                               1 
                                               -1.4 
                                             
                                             
                                               AKT3-mRNA 
                                               0.13 
                                               0.121 
                                               -0.107 
                                               0.366 
                                               1.09 
                                               0.929 
                                               1.29 
                                               0.294 
                                               1 
                                               lm.nb 
                                               Activation of BAD and translocation to mitochondria , CD28 co-stimulation, CD28 dependent PI3KAkt signaling, DAP12 signaling, Downstream signal transduction, G beta.CL.gamma signalling through PI3Kgamma, GAB1 signalosome, GPVI-mediated activation cascade, PI3KAKT activation, PIP3 activates AKT signaling, Role of LAT2NTALLAB on calcium mobilization, Signaling by ERBB2, Signaling by PDGF, Signaling by SCF-KIT, TP53 Regulates Metabolic Genes, VEGFA-VEGFR2 Pathway, VEGFR2 mediated vascular permeability 
                                               SEBELIUS_2_C4636 
                                               AKT3 
                                               mRNA 
                                               0 
                                               
                                               
                                               TP53 Regulates Metabolic Genes;CD28 dependent PI3KAkt signaling;AKT phosphorylates targets in the cytosol;Negative regulation of the PI3KAKT network;Activation of BAD and translocation to mitochondria ;G beta.CL.gamma signalling through PI3Kgamma;AKT phosphorylates targets in the nucleus;VEGFR2 mediated vascular permeability;GPVI-mediated activation cascade;Signaling by ERBB2;PIP3 activates AKT signaling;Downstream signal transduction;GAB1 signalosome;Signaling by SCF-KIT;PI3KAKT activation;Role of LAT2NTALLAB on calcium mobilization;DAP12 signaling;Signaling by PDGF;CD28 co-stimulation;VEGFA-VEGFR2 Pathway 
                                               hsa04010;hsa04012;hsa04014;hsa04015;hsa04022;hsa04024;hsa04062;hsa04066;hsa04068;hsa04071;hsa04072;hsa04150;hsa04151;hsa04152;hsa04210;hsa04211;hsa04213;hsa04261;hsa04370;hsa04380;hsa04510;hsa04530;hsa04550;hsa04611;hsa04620;hsa04630;hsa04660;hsa04662;hsa04664;hsa04666;hsa04668;hsa04722;hsa04725;hsa04728;hsa04910;hsa04914;hsa04915;hsa04917;hsa04919;hsa04920;hsa04922;hsa04923;hsa04931;hsa04932;hsa04933;hsa04973;hsa05142;hsa05145;hsa05152;hsa05160;hsa05161;hsa05162;hsa05164;hsa05166;hsa05169;hsa05200;hsa05205;hsa05210;hsa05211;hsa05212;hsa05213;hsa05214;hsa05215;hsa05218;hsa05220;hsa05221;hsa05222;hsa05223;hsa05230;hsa05231 
                                               AKT3 
                                               NA 
                                               azure4 
                                               1 
                                               1.07438016528926 
                                             
                                             
                                               BCL2-mRNA 
                                               -1.79 
                                               4.09 
                                               -9.8 
                                               6.23 
                                               0.29 
                                               0.00112 
                                               74.9 
                                               0.667 
                                               1 
                                               Wald 
                                               Activation of BAD and translocation to mitochondria  
                                               SEBELIUS_2_C4636 
                                               BCL2 
                                               mRNA 
                                               0 
                                               
                                               
                                               Activation of BAD and translocation to mitochondria  
                                               hsa04064;hsa04066;hsa04071;hsa04141;hsa04151;hsa04210;hsa04261;hsa04510;hsa04630;hsa04722;hsa04725;hsa04933;hsa05145;hsa05152;hsa05161;hsa05169;hsa05200;hsa05206;hsa05210;hsa05215;hsa05222 
                                               BCL2 
                                               NA 
                                               azure4 
                                               1 
                                               -0.437652811735941 
                                             
                                             
                                               AKT2-mRNA 
                                               0.0144 
                                               0.0687 
                                               -0.12 
                                               0.149 
                                               1.01 
                                               0.92 
                                               1.11 
                                               0.836 
                                               1 
                                               loglinear 
                                               Activation of BAD and translocation to mitochondria , CD28 co-stimulation, DAP12 signaling, Deactivation of the beta-catenin transactivating complex, Downstream signal transduction, G beta.CL.gamma signalling through PI3Kgamma, GAB1 signalosome, GPVI-mediated activation cascade, IRS-related events triggered by IGF1R, PI3K Cascade, PI3KAKT activation, PIP3 activates AKT signaling, Role of LAT2NTALLAB on calcium mobilization, Signaling by ERBB2, Signaling by PDGF, Signaling by SCF-KIT, TP53 Regulates Metabolic Genes, VEGFA-VEGFR2 Pathway, VEGFR2 mediated vascular permeability 
                                               SEBELIUS_2_C4636 
                                               AKT2 
                                               mRNA 
                                               0 
                                               
                                               
                                               TP53 Regulates Metabolic Genes;PDE3B signalling;AKT phosphorylates targets in the cytosol;Negative regulation of the PI3KAKT network;Deactivation of the beta-catenin transactivating complex;Activation of BAD and translocation to mitochondria ;G beta.CL.gamma signalling through PI3Kgamma;AKT phosphorylates targets in the nucleus;VEGFR2 mediated vascular permeability;GPVI-mediated activation cascade;Signaling by ERBB2;PI3K Cascade;IRS-related events triggered by IGF1R;mTOR signalling;PIP3 activates AKT signaling;Downstream signal transduction;GAB1 signalosome;Signaling by SCF-KIT;PI3KAKT activation;Role of LAT2NTALLAB on calcium mobilization;DAP12 signaling;Signaling by PDGF;CD28 co-stimulation;VEGFA-VEGFR2 Pathway 
                                               hsa04010;hsa04012;hsa04014;hsa04015;hsa04022;hsa04024;hsa04062;hsa04066;hsa04068;hsa04071;hsa04072;hsa04150;hsa04151;hsa04152;hsa04210;hsa04211;hsa04213;hsa04261;hsa04370;hsa04380;hsa04510;hsa04530;hsa04550;hsa04611;hsa04620;hsa04630;hsa04660;hsa04662;hsa04664;hsa04666;hsa04668;hsa04722;hsa04725;hsa04728;hsa04910;hsa04914;hsa04915;hsa04917;hsa04919;hsa04920;hsa04922;hsa04923;hsa04931;hsa04932;hsa04933;hsa04973;hsa05142;hsa05145;hsa05152;hsa05160;hsa05161;hsa05162;hsa05164;hsa05166;hsa05169;hsa05200;hsa05205;hsa05210;hsa05211;hsa05212;hsa05213;hsa05214;hsa05215;hsa05218;hsa05220;hsa05221;hsa05222;hsa05223;hsa05230;hsa05231 
                                               AKT2 
                                               NA 
                                               azure4 
                                               1 
                                               0.209606986899563 
                                             
                                           
                                           DE Results - Activation of BAD and translocation to mitochondria  genes - New.Annotation: miR18b vs.cel67  Table displaying each sample's global significance scores and directed global significance scores as defined in the heatmaps above. The global significance score is calculated as the square root of the mean squared t-statistic for the genes in a gene set, with t-statistics coming from the linear regression underlying our differential expression analysis. The directed global significance score is calculated as the square root of the mean signed squared t-statistic for the genes in a gene set, with t-statistics coming from the linear regression underlying our differential expression analysis.   
                                      
                                       
                                  
                                
                                
                                  
                                       
                                      
                                           Volcano Plot: New.Annotation: miR890 vs.cel67   More Plot Information                Volcano Plot: New.Annotation: miR890 vs.cel67  Volcano plot displaying each gene's -log10(p-value) and log2 fold change for the selected covariate. Highly statistically significant genes fall at the top of the plot, and highly differentially expressed genes fall to either side. Genes within the selected gene set are highlighted in orange. Horizontal lines indicate various False Discovery Rate (FDR) thresholds.   
                                           DE Results - Activation of BAD and translocation to mitochondria  genes - New.Annotation: miR890 vs.cel67   More Plot Information      Download CSV Data     
                                           
                                             
                                             
                                           
                                           
                                             
                                               Probe Label 
                                               Log2 fold change 
                                               std error (log2) 
                                               Lower confidence limit (log2) 
                                               Upper confidence limit (log2) 
                                               Linear fold change 
                                               Lower confidence limit (linear) 
                                               Upper confidence limit (linear) 
                                               P-value 
                                               BY.p.value 
                                               method 
                                               Gene.sets 
                                               Codeset.Name 
                                               Probe.Label 
                                               Analyte.Type 
                                               Is.Control 
                                               Control.Type 
                                               Related.Probes 
                                               Probe.Annotation 
                                               KEGG.Pathways 
                                               Official.Gene.Name 
                                               Control.Conc 
                                               volcanocol 
                                               volcanopch 
                                               tstats.all 
                                             
                                           
                                           
                                             
                                               AKT1-mRNA 
                                               -0.237 
                                               0.146 
                                               -0.522 
                                               0.0483 
                                               0.849 
                                               0.696 
                                               1.03 
                                               0.118 
                                               1 
                                               lm.nb 
                                               Activation of BAD and translocation to mitochondria , CD28 co-stimulation, DAP12 signaling, Deactivation of the beta-catenin transactivating complex, Downstream signal transduction, G beta.CL.gamma signalling through PI3Kgamma, GAB1 signalosome, GPVI-mediated activation cascade, IRS-related events triggered by IGF1R, PI3K Cascade, PI3KAKT activation, PIP3 activates AKT signaling, Role of LAT2NTALLAB on calcium mobilization, Signaling by ERBB2, Signaling by PDGF, Signaling by SCF-KIT, TP53 Regulates Metabolic Genes, VEGFA-VEGFR2 Pathway, VEGFR2 mediated vascular permeability 
                                               SEBELIUS_2_C4636 
                                               AKT1 
                                               mRNA 
                                               0 
                                               
                                               
                                               TP53 Regulates Metabolic Genes;Deactivation of the beta-catenin transactivating complex;Activation of BAD and translocation to mitochondria ;eNOS activation;mTOR signalling;G beta.CL.gamma signalling through PI3Kgamma;KSRP (KHSRP) binds and destabilizes mRNA;AKT phosphorylates targets in the nucleus;VEGFR2 mediated vascular permeability;GPVI-mediated activation cascade;Tetrahydrobiopterin (BH4) synthesis# recycling# salvage and regulation;Butyrate Response Factor 1 (BRF1) binds and destabilizes mRNA;Signaling by ERBB2;PIP3 activates AKT signaling;Downstream signal transduction;GAB1 signalosome;Signaling by SCF-KIT;PI3KAKT activation;Role of LAT2NTALLAB on calcium mobilization;DAP12 signaling;Signaling by PDGF;CD28 co-stimulation;PI3K Cascade;IRS-related events triggered by IGF1R;VEGFA-VEGFR2 Pathway 
                                               hsa04010;hsa04012;hsa04014;hsa04015;hsa04022;hsa04024;hsa04062;hsa04066;hsa04068;hsa04071;hsa04072;hsa04150;hsa04151;hsa04152;hsa04210;hsa04211;hsa04213;hsa04261;hsa04370;hsa04380;hsa04510;hsa04530;hsa04550;hsa04611;hsa04620;hsa04630;hsa04660;hsa04662;hsa04664;hsa04666;hsa04668;hsa04722;hsa04725;hsa04728;hsa04910;hsa04914;hsa04915;hsa04917;hsa04919;hsa04920;hsa04922;hsa04923;hsa04931;hsa04932;hsa04933;hsa04973;hsa05142;hsa05145;hsa05152;hsa05160;hsa05161;hsa05162;hsa05164;hsa05166;hsa05169;hsa05200;hsa05205;hsa05210;hsa05211;hsa05212;hsa05213;hsa05214;hsa05215;hsa05218;hsa05220;hsa05221;hsa05222;hsa05223;hsa05230;hsa05231 
                                               AKT1 
                                               NA 
                                               azure4 
                                               1 
                                               -1.62328767123288 
                                             
                                             
                                               AKT2-mRNA 
                                               -0.0777 
                                               0.0581 
                                               -0.191 
                                               0.0361 
                                               0.948 
                                               0.876 
                                               1.03 
                                               0.195 
                                               1 
                                               loglinear 
                                               Activation of BAD and translocation to mitochondria , CD28 co-stimulation, DAP12 signaling, Deactivation of the beta-catenin transactivating complex, Downstream signal transduction, G beta.CL.gamma signalling through PI3Kgamma, GAB1 signalosome, GPVI-mediated activation cascade, IRS-related events triggered by IGF1R, PI3K Cascade, PI3KAKT activation, PIP3 activates AKT signaling, Role of LAT2NTALLAB on calcium mobilization, Signaling by ERBB2, Signaling by PDGF, Signaling by SCF-KIT, TP53 Regulates Metabolic Genes, VEGFA-VEGFR2 Pathway, VEGFR2 mediated vascular permeability 
                                               SEBELIUS_2_C4636 
                                               AKT2 
                                               mRNA 
                                               0 
                                               
                                               
                                               TP53 Regulates Metabolic Genes;PDE3B signalling;AKT phosphorylates targets in the cytosol;Negative regulation of the PI3KAKT network;Deactivation of the beta-catenin transactivating complex;Activation of BAD and translocation to mitochondria ;G beta.CL.gamma signalling through PI3Kgamma;AKT phosphorylates targets in the nucleus;VEGFR2 mediated vascular permeability;GPVI-mediated activation cascade;Signaling by ERBB2;PI3K Cascade;IRS-related events triggered by IGF1R;mTOR signalling;PIP3 activates AKT signaling;Downstream signal transduction;GAB1 signalosome;Signaling by SCF-KIT;PI3KAKT activation;Role of LAT2NTALLAB on calcium mobilization;DAP12 signaling;Signaling by PDGF;CD28 co-stimulation;VEGFA-VEGFR2 Pathway 
                                               hsa04010;hsa04012;hsa04014;hsa04015;hsa04022;hsa04024;hsa04062;hsa04066;hsa04068;hsa04071;hsa04072;hsa04150;hsa04151;hsa04152;hsa04210;hsa04211;hsa04213;hsa04261;hsa04370;hsa04380;hsa04510;hsa04530;hsa04550;hsa04611;hsa04620;hsa04630;hsa04660;hsa04662;hsa04664;hsa04666;hsa04668;hsa04722;hsa04725;hsa04728;hsa04910;hsa04914;hsa04915;hsa04917;hsa04919;hsa04920;hsa04922;hsa04923;hsa04931;hsa04932;hsa04933;hsa04973;hsa05142;hsa05145;hsa05152;hsa05160;hsa05161;hsa05162;hsa05164;hsa05166;hsa05169;hsa05200;hsa05205;hsa05210;hsa05211;hsa05212;hsa05213;hsa05214;hsa05215;hsa05218;hsa05220;hsa05221;hsa05222;hsa05223;hsa05230;hsa05231 
                                               AKT2 
                                               NA 
                                               azure4 
                                               1 
                                               -1.33734939759036 
                                             
                                             
                                               AKT3-mRNA 
                                               0.079 
                                               0.102 
                                               -0.121 
                                               0.279 
                                               1.06 
                                               0.919 
                                               1.21 
                                               0.448 
                                               1 
                                               lm.nb 
                                               Activation of BAD and translocation to mitochondria , CD28 co-stimulation, CD28 dependent PI3KAkt signaling, DAP12 signaling, Downstream signal transduction, G beta.CL.gamma signalling through PI3Kgamma, GAB1 signalosome, GPVI-mediated activation cascade, PI3KAKT activation, PIP3 activates AKT signaling, Role of LAT2NTALLAB on calcium mobilization, Signaling by ERBB2, Signaling by PDGF, Signaling by SCF-KIT, TP53 Regulates Metabolic Genes, VEGFA-VEGFR2 Pathway, VEGFR2 mediated vascular permeability 
                                               SEBELIUS_2_C4636 
                                               AKT3 
                                               mRNA 
                                               0 
                                               
                                               
                                               TP53 Regulates Metabolic Genes;CD28 dependent PI3KAkt signaling;AKT phosphorylates targets in the cytosol;Negative regulation of the PI3KAKT network;Activation of BAD and translocation to mitochondria ;G beta.CL.gamma signalling through PI3Kgamma;AKT phosphorylates targets in the nucleus;VEGFR2 mediated vascular permeability;GPVI-mediated activation cascade;Signaling by ERBB2;PIP3 activates AKT signaling;Downstream signal transduction;GAB1 signalosome;Signaling by SCF-KIT;PI3KAKT activation;Role of LAT2NTALLAB on calcium mobilization;DAP12 signaling;Signaling by PDGF;CD28 co-stimulation;VEGFA-VEGFR2 Pathway 
                                               hsa04010;hsa04012;hsa04014;hsa04015;hsa04022;hsa04024;hsa04062;hsa04066;hsa04068;hsa04071;hsa04072;hsa04150;hsa04151;hsa04152;hsa04210;hsa04211;hsa04213;hsa04261;hsa04370;hsa04380;hsa04510;hsa04530;hsa04550;hsa04611;hsa04620;hsa04630;hsa04660;hsa04662;hsa04664;hsa04666;hsa04668;hsa04722;hsa04725;hsa04728;hsa04910;hsa04914;hsa04915;hsa04917;hsa04919;hsa04920;hsa04922;hsa04923;hsa04931;hsa04932;hsa04933;hsa04973;hsa05142;hsa05145;hsa05152;hsa05160;hsa05161;hsa05162;hsa05164;hsa05166;hsa05169;hsa05200;hsa05205;hsa05210;hsa05211;hsa05212;hsa05213;hsa05214;hsa05215;hsa05218;hsa05220;hsa05221;hsa05222;hsa05223;hsa05230;hsa05231 
                                               AKT3 
                                               NA 
                                               azure4 
                                               1 
                                               0.774509803921569 
                                             
                                             
                                               BCL2-mRNA 
                                               -1.68 
                                               3.44 
                                               -8.41 
                                               5.06 
                                               0.313 
                                               0.00293 
                                               33.4 
                                               0.631 
                                               1 
                                               Wald 
                                               Activation of BAD and translocation to mitochondria  
                                               SEBELIUS_2_C4636 
                                               BCL2 
                                               mRNA 
                                               0 
                                               
                                               
                                               Activation of BAD and translocation to mitochondria  
                                               hsa04064;hsa04066;hsa04071;hsa04141;hsa04151;hsa04210;hsa04261;hsa04510;hsa04630;hsa04722;hsa04725;hsa04933;hsa05145;hsa05152;hsa05161;hsa05169;hsa05200;hsa05206;hsa05210;hsa05215;hsa05222 
                                               BCL2 
                                               NA 
                                               azure4 
                                               1 
                                               -0.488372093023256 
                                             
                                             
                                               BAD-mRNA 
                                               -0.0533 
                                               0.171 
                                               -0.389 
                                               0.283 
                                               0.964 
                                               0.764 
                                               1.22 
                                               0.759 
                                               1 
                                               lm.nb 
                                               Activation of BAD and translocation to mitochondria , DAP12 signaling, Downstream signal transduction, GAB1 signalosome, NRAGE signals death through JNK, PI3KAKT activation, PIP3 activates AKT signaling, Role of LAT2NTALLAB on calcium mobilization, Signaling by ERBB2, Signaling by PDGF, Signaling by SCF-KIT 
                                               SEBELIUS_2_C4636 
                                               BAD 
                                               mRNA 
                                               0 
                                               
                                               
                                               Activation of BAD and translocation to mitochondria ;NRAGE signals death through JNK;Constitutive Signaling by AKT1 E17K in Cancer;AKT phosphorylates targets in the cytosol;PIP3 activates AKT signaling;Downstream signal transduction;GAB1 signalosome;Signaling by SCF-KIT;PI3KAKT activation;Role of LAT2NTALLAB on calcium mobilization;DAP12 signaling;Signaling by PDGF;Signaling by ERBB2 
                                               hsa04012;hsa04014;hsa04022;hsa04024;hsa04151;hsa04210;hsa04370;hsa04510;hsa04722;hsa04910;hsa04919;hsa05010;hsa05145;hsa05152;hsa05160;hsa05161;hsa05200;hsa05203;hsa05210;hsa05212;hsa05213;hsa05215;hsa05218;hsa05220;hsa05221;hsa05223 
                                               BAD 
                                               NA 
                                               azure4 
                                               1 
                                               -0.311695906432749 
                                             
                                           
                                           DE Results - Activation of BAD and translocation to mitochondria  genes - New.Annotation: miR890 vs.cel67  Table displaying each sample's global significance scores and directed global significance scores as defined in the heatmaps above. The global significance score is calculated as the square root of the mean squared t-statistic for the genes in a gene set, with t-statistics coming from the linear regression underlying our differential expression analysis. The directed global significance score is calculated as the square root of the mean signed squared t-statistic for the genes in a gene set, with t-statistics coming from the linear regression underlying our differential expression analysis.   
                                      
                                       
                                  
                                
                              
                             
                          
                           
                      
                    
                    
                      
                           
                          
                              New.Annotation: 
                              
                                  New.Annotation: differential expression in Control vs. baseline of cel67  
                                  New.Annotation: differential expression in miR142 vs. baseline of cel67  
                                  New.Annotation: differential expression in miR155 vs. baseline of cel67  
                                  New.Annotation: differential expression in miR18b vs. baseline of cel67  
                                  New.Annotation: differential expression in miR890 vs. baseline of cel67  
                              
                              
                              
                                
                                  
                                       
                                      
                                           Volcano Plot: New.Annotation: Control vs.cel67   More Plot Information                Volcano Plot: New.Annotation: Control vs.cel67  Volcano plot displaying each gene's -log10(p-value) and log2 fold change for the selected covariate. Highly statistically significant genes fall at the top of the plot, and highly differentially expressed genes fall to either side. Genes within the selected gene set are highlighted in orange. Horizontal lines indicate various False Discovery Rate (FDR) thresholds.   
                                           DE Results - Activation of Rac genes - New.Annotation: Control vs.cel67   More Plot Information      Download CSV Data     
                                           
                                             
                                             
                                           
                                           
                                             
                                               Probe Label 
                                               Log2 fold change 
                                               std error (log2) 
                                               Lower confidence limit (log2) 
                                               Upper confidence limit (log2) 
                                               Linear fold change 
                                               Lower confidence limit (linear) 
                                               Upper confidence limit (linear) 
                                               P-value 
                                               BY.p.value 
                                               method 
                                               Gene.sets 
                                               Codeset.Name 
                                               Probe.Label 
                                               Analyte.Type 
                                               Is.Control 
                                               Control.Type 
                                               Related.Probes 
                                               Probe.Annotation 
                                               KEGG.Pathways 
                                               Official.Gene.Name 
                                               Control.Conc 
                                               volcanocol 
                                               volcanopch 
                                               tstats.all 
                                             
                                           
                                           
                                             
                                               SOS2-mRNA 
                                               -0.206 
                                               0.084 
                                               -0.371 
                                               -0.0416 
                                               0.867 
                                               0.773 
                                               0.972 
                                               0.0229 
                                               0.756 
                                               loglinear 
                                               Activation of Rac, G alpha (1213) signalling events, NRAGE signals death through JNK, Rho GTPase cycle 
                                               SEBELIUS_2_C4636 
                                               SOS2 
                                               mRNA 
                                               0 
                                               
                                               
                                               Rho GTPase cycle;Activation of Rac;NRAGE signals death through JNK;G alpha (1213) signalling events 
                                               hsa04010;hsa04012;hsa04014;hsa04062;hsa04068;hsa04072;hsa04151;hsa04320;hsa04510;hsa04540;hsa04630;hsa04650;hsa04660;hsa04662;hsa04664;hsa04722;hsa04810;hsa04910;hsa04912;hsa04915;hsa04917;hsa05034;hsa05160;hsa05200;hsa05205;hsa05206;hsa05211;hsa05213;hsa05214;hsa05215;hsa05220;hsa05221;hsa05223;hsa05231 
                                               SOS2 
                                               NA 
                                               azure4 
                                               1 
                                               -2.45238095238095 
                                             
                                             
                                               PAK4-mRNA 
                                               -0.283 
                                               0.124 
                                               -0.527 
                                               -0.0392 
                                               0.822 
                                               0.694 
                                               0.973 
                                               0.0335 
                                               0.909 
                                               loglinear 
                                               Activation of Rac 
                                               SEBELIUS_2_C4636 
                                               PAK4 
                                               mRNA 
                                               0 
                                               
                                               
                                               Activation of Rac 
                                               hsa04012;hsa04014;hsa04360;hsa04510;hsa04660;hsa04810;hsa05206;hsa05211 
                                               PAK4 
                                               NA 
                                               azure4 
                                               1 
                                               -2.28225806451613 
                                             
                                             
                                               RAC1-mRNA 
                                               -0.125 
                                               0.0675 
                                               -0.257 
                                               0.00726 
                                               0.917 
                                               0.837 
                                               1.01 
                                               0.0781 
                                               1 
                                               loglinear 
                                               Activation of Rac, CD28 co-stimulation, CD28 dependent Vav1 pathway, DAP12 signaling, EPH-ephrin mediated repulsion of cells, Ephrin signaling, G alpha (1213) signalling events, GPVI-mediated activation cascade, MAPK6MAPK4 signaling, NRAGE signals death through JNK, PCPCE pathway, Regulation of actin dynamics for phagocytic cup formation, RHO GTPases activate CIT, RHO GTPases Activate Formins, RHO GTPases activate IQGAPs, RHO GTPases activate PAKs, RHO GTPases activate PKNs, RHO GTPases Activate WASPs and WAVEs, Sema3A PAK dependent Axon repulsion, Sema4D induced cell migration and growth-cone collapse, Signal transduction by L1, Signaling by SCF-KIT, VEGFA-VEGFR2 Pathway, VEGFR2 mediated vascular permeability 
                                               SEBELIUS_2_C4636 
                                               RAC1 
                                               mRNA 
                                               0 
                                               
                                               
                                               MAPK6MAPK4 signaling;Signaling by SCF-KIT;Inactivation of Cdc42 and Rac;DAP12 signaling;Ephrin signaling;EPH-ephrin mediated repulsion of cells;Activation of Rac;RHO GTPases Activate Formins;VEGFR2 mediated vascular permeability;RHO GTPases activate PAKs;Sema4D mediated inhibition of cell attachment and migration;G alpha (1213) signalling events;Regulation of actin dynamics for phagocytic cup formation;Sema4D induced cell migration and growth-cone collapse;Signal transduction by L1;RHO GTPases activate CIT;RHO GTPases Activate WASPs and WAVEs;Factors involved in megakaryocyte development and platelet production;RHO GTPases activate KTN1;CD28 dependent Vav1 pathway;DSCAM interactions;NRAGE signals death through JNK;GPVI-mediated activation cascade;Sema3A PAK dependent Axon repulsion;RHO GTPases activate PKNs;SEMA3A-Plexin repulsion signaling by inhibiting Integrin adhesion;RHO GTPases activate IQGAPs;PCPCE pathway;VEGFA-VEGFR2 Pathway;CD28 co-stimulation 
                                               hsa04010;hsa04014;hsa04015;hsa04024;hsa04062;hsa04071;hsa04145;hsa04151;hsa04310;hsa04360;hsa04370;hsa04380;hsa04510;hsa04520;hsa04620;hsa04650;hsa04662;hsa04664;hsa04666;hsa04670;hsa04722;hsa04810;hsa04932;hsa04933;hsa04972;hsa05100;hsa05120;hsa05131;hsa05132;hsa05200;hsa05203;hsa05205;hsa05210;hsa05211;hsa05212;hsa05231;hsa05416 
                                               RAC1 
                                               NA 
                                               azure4 
                                               1 
                                               -1.85185185185185 
                                             
                                             
                                               PAK1-mRNA 
                                               -0.177 
                                               0.163 
                                               -0.497 
                                               0.142 
                                               0.884 
                                               0.709 
                                               1.1 
                                               0.288 
                                               1 
                                               lm.nb 
                                               Activation of Rac, CD209 (DC-SIGN) signaling, CD28 co-stimulation, CD28 dependent Vav1 pathway, EPHB-mediated forward signaling, Ephrin signaling, FCERI mediated MAPK activation, MAPK6MAPK4 signaling, Regulation of actin dynamics for phagocytic cup formation, RHO GTPases activate PAKs, Sema3A PAK dependent Axon repulsion, VEGFA-VEGFR2 Pathway, VEGFR2 mediated vascular permeability 
                                               SEBELIUS_2_C4636 
                                               PAK1 
                                               mRNA 
                                               0 
                                               
                                               
                                               EPHB-mediated forward signaling;Ephrin signaling;MAPK6MAPK4 signaling;DSCAM interactions;Activation of Rac;VEGFR2 mediated vascular permeability;RHO GTPases activate PAKs;Sema3A PAK dependent Axon repulsion;FCERI mediated MAPK activation;Regulation of actin dynamics for phagocytic cup formation;CD209 (DC-SIGN) signaling;CD28 dependent Vav1 pathway;VEGFA-VEGFR2 Pathway;CD28 co-stimulation 
                                               hsa04010;hsa04012;hsa04014;hsa04024;hsa04062;hsa04360;hsa04510;hsa04650;hsa04660;hsa04666;hsa04810;hsa05120;hsa05205;hsa05211 
                                               PAK1 
                                               NA 
                                               azure4 
                                               1 
                                               -1.08588957055215 
                                             
                                             
                                               PAK2-mRNA 
                                               -0.0762 
                                               0.0976 
                                               -0.268 
                                               0.115 
                                               0.949 
                                               0.831 
                                               1.08 
                                               0.444 
                                               1 
                                               loglinear 
                                               Activation of Rac, CD209 (DC-SIGN) signaling, CD28 co-stimulation, CD28 dependent Vav1 pathway, Ephrin signaling, RHO GTPases activate PAKs, Sema3A PAK dependent Axon repulsion, VEGFA-VEGFR2 Pathway, VEGFR2 mediated vascular permeability 
                                               SEBELIUS_2_C4636 
                                               PAK2 
                                               mRNA 
                                               0 
                                               
                                               
                                               Nef and signal transduction;CD28 dependent Vav1 pathway;Stimulation of the cell death response by PAK-2p34;Ephrin signaling;Activation of Rac;RHO GTPases activate PAKs;VEGFR2 mediated vascular permeability;Sema3A PAK dependent Axon repulsion;CD209 (DC-SIGN) signaling;VEGFA-VEGFR2 Pathway;CD28 co-stimulation 
                                               hsa04010;hsa04012;hsa04014;hsa04360;hsa04510;hsa04660;hsa04810;hsa05211 
                                               PAK2 
                                               NA 
                                               azure4 
                                               1 
                                               -0.780737704918033 
                                             
                                           
                                           DE Results - Activation of Rac genes - New.Annotation: Control vs.cel67  Table displaying each sample's global significance scores and directed global significance scores as defined in the heatmaps above. The global significance score is calculated as the square root of the mean squared t-statistic for the genes in a gene set, with t-statistics coming from the linear regression underlying our differential expression analysis. The directed global significance score is calculated as the square root of the mean signed squared t-statistic for the genes in a gene set, with t-statistics coming from the linear regression underlying our differential expression analysis.   
                                      
                                       
                                  
                                
                                
                                  
                                       
                                      
                                           Volcano Plot: New.Annotation: miR142 vs.cel67   More Plot Information                Volcano Plot: New.Annotation: miR142 vs.cel67  Volcano plot displaying each gene's -log10(p-value) and log2 fold change for the selected covariate. Highly statistically significant genes fall at the top of the plot, and highly differentially expressed genes fall to either side. Genes within the selected gene set are highlighted in orange. Horizontal lines indicate various False Discovery Rate (FDR) thresholds.   
                                           DE Results - Activation of Rac genes - New.Annotation: miR142 vs.cel67   More Plot Information      Download CSV Data     
                                           
                                             
                                             
                                           
                                           
                                             
                                               Probe Label 
                                               Log2 fold change 
                                               std error (log2) 
                                               Lower confidence limit (log2) 
                                               Upper confidence limit (log2) 
                                               Linear fold change 
                                               Lower confidence limit (linear) 
                                               Upper confidence limit (linear) 
                                               P-value 
                                               BY.p.value 
                                               method 
                                               Gene.sets 
                                               Codeset.Name 
                                               Probe.Label 
                                               Analyte.Type 
                                               Is.Control 
                                               Control.Type 
                                               Related.Probes 
                                               Probe.Annotation 
                                               KEGG.Pathways 
                                               Official.Gene.Name 
                                               Control.Conc 
                                               volcanocol 
                                               volcanopch 
                                               tstats.all 
                                             
                                           
                                           
                                             
                                               RAC1-mRNA 
                                               -1.32 
                                               0.0675 
                                               -1.45 
                                               -1.19 
                                               0.401 
                                               0.366 
                                               0.439 
                                               5.98e-15 
                                               7.94e-12 
                                               loglinear 
                                               Activation of Rac, CD28 co-stimulation, CD28 dependent Vav1 pathway, DAP12 signaling, EPH-ephrin mediated repulsion of cells, Ephrin signaling, G alpha (1213) signalling events, GPVI-mediated activation cascade, MAPK6MAPK4 signaling, NRAGE signals death through JNK, PCPCE pathway, Regulation of actin dynamics for phagocytic cup formation, RHO GTPases activate CIT, RHO GTPases Activate Formins, RHO GTPases activate IQGAPs, RHO GTPases activate PAKs, RHO GTPases activate PKNs, RHO GTPases Activate WASPs and WAVEs, Sema3A PAK dependent Axon repulsion, Sema4D induced cell migration and growth-cone collapse, Signal transduction by L1, Signaling by SCF-KIT, VEGFA-VEGFR2 Pathway, VEGFR2 mediated vascular permeability 
                                               SEBELIUS_2_C4636 
                                               RAC1 
                                               mRNA 
                                               0 
                                               
                                               
                                               MAPK6MAPK4 signaling;Signaling by SCF-KIT;Inactivation of Cdc42 and Rac;DAP12 signaling;Ephrin signaling;EPH-ephrin mediated repulsion of cells;Activation of Rac;RHO GTPases Activate Formins;VEGFR2 mediated vascular permeability;RHO GTPases activate PAKs;Sema4D mediated inhibition of cell attachment and migration;G alpha (1213) signalling events;Regulation of actin dynamics for phagocytic cup formation;Sema4D induced cell migration and growth-cone collapse;Signal transduction by L1;RHO GTPases activate CIT;RHO GTPases Activate WASPs and WAVEs;Factors involved in megakaryocyte development and platelet production;RHO GTPases activate KTN1;CD28 dependent Vav1 pathway;DSCAM interactions;NRAGE signals death through JNK;GPVI-mediated activation cascade;Sema3A PAK dependent Axon repulsion;RHO GTPases activate PKNs;SEMA3A-Plexin repulsion signaling by inhibiting Integrin adhesion;RHO GTPases activate IQGAPs;PCPCE pathway;VEGFA-VEGFR2 Pathway;CD28 co-stimulation 
                                               hsa04010;hsa04014;hsa04015;hsa04024;hsa04062;hsa04071;hsa04145;hsa04151;hsa04310;hsa04360;hsa04370;hsa04380;hsa04510;hsa04520;hsa04620;hsa04650;hsa04662;hsa04664;hsa04666;hsa04670;hsa04722;hsa04810;hsa04932;hsa04933;hsa04972;hsa05100;hsa05120;hsa05131;hsa05132;hsa05200;hsa05203;hsa05205;hsa05210;hsa05211;hsa05212;hsa05231;hsa05416 
                                               RAC1 
                                               NA 
                                               slateblue1 
                                               16 
                                               -19.5555555555556 
                                             
                                             
                                               PAK4-mRNA 
                                               -0.724 
                                               0.124 
                                               -0.968 
                                               -0.48 
                                               0.605 
                                               0.511 
                                               0.717 
                                               8.96e-06 
                                               0.000397 
                                               loglinear 
                                               Activation of Rac 
                                               SEBELIUS_2_C4636 
                                               PAK4 
                                               mRNA 
                                               0 
                                               
                                               
                                               Activation of Rac 
                                               hsa04012;hsa04014;hsa04360;hsa04510;hsa04660;hsa04810;hsa05206;hsa05211 
                                               PAK4 
                                               NA 
                                               slateblue1 
                                               16 
                                               -5.83870967741935 
                                             
                                             
                                               PAK1-mRNA 
                                               -0.455 
                                               0.163 
                                               -0.775 
                                               -0.135 
                                               0.729 
                                               0.584 
                                               0.91 
                                               0.011 
                                               0.164 
                                               lm.nb 
                                               Activation of Rac, CD209 (DC-SIGN) signaling, CD28 co-stimulation, CD28 dependent Vav1 pathway, EPHB-mediated forward signaling, Ephrin signaling, FCERI mediated MAPK activation, MAPK6MAPK4 signaling, Regulation of actin dynamics for phagocytic cup formation, RHO GTPases activate PAKs, Sema3A PAK dependent Axon repulsion, VEGFA-VEGFR2 Pathway, VEGFR2 mediated vascular permeability 
                                               SEBELIUS_2_C4636 
                                               PAK1 
                                               mRNA 
                                               0 
                                               
                                               
                                               EPHB-mediated forward signaling;Ephrin signaling;MAPK6MAPK4 signaling;DSCAM interactions;Activation of Rac;VEGFR2 mediated vascular permeability;RHO GTPases activate PAKs;Sema3A PAK dependent Axon repulsion;FCERI mediated MAPK activation;Regulation of actin dynamics for phagocytic cup formation;CD209 (DC-SIGN) signaling;CD28 dependent Vav1 pathway;VEGFA-VEGFR2 Pathway;CD28 co-stimulation 
                                               hsa04010;hsa04012;hsa04014;hsa04024;hsa04062;hsa04360;hsa04510;hsa04650;hsa04660;hsa04666;hsa04810;hsa05120;hsa05205;hsa05211 
                                               PAK1 
                                               NA 
                                               slateblue4 
                                               16 
                                               -2.79141104294479 
                                             
                                             
                                               SOS2-mRNA 
                                               0.189 
                                               0.084 
                                               0.0249 
                                               0.354 
                                               1.14 
                                               1.02 
                                               1.28 
                                               0.0348 
                                               0.432 
                                               loglinear 
                                               Activation of Rac, G alpha (1213) signalling events, NRAGE signals death through JNK, Rho GTPase cycle 
                                               SEBELIUS_2_C4636 
                                               SOS2 
                                               mRNA 
                                               0 
                                               
                                               
                                               Rho GTPase cycle;Activation of Rac;NRAGE signals death through JNK;G alpha (1213) signalling events 
                                               hsa04010;hsa04012;hsa04014;hsa04062;hsa04068;hsa04072;hsa04151;hsa04320;hsa04510;hsa04540;hsa04630;hsa04650;hsa04660;hsa04662;hsa04664;hsa04722;hsa04810;hsa04910;hsa04912;hsa04915;hsa04917;hsa05034;hsa05160;hsa05200;hsa05205;hsa05206;hsa05211;hsa05213;hsa05214;hsa05215;hsa05220;hsa05221;hsa05223;hsa05231 
                                               SOS2 
                                               NA 
                                               slateblue4 
                                               16 
                                               2.25 
                                             
                                             
                                               PAK2-mRNA 
                                               0.147 
                                               0.0976 
                                               -0.0448 
                                               0.338 
                                               1.11 
                                               0.969 
                                               1.26 
                                               0.148 
                                               1 
                                               loglinear 
                                               Activation of Rac, CD209 (DC-SIGN) signaling, CD28 co-stimulation, CD28 dependent Vav1 pathway, Ephrin signaling, RHO GTPases activate PAKs, Sema3A PAK dependent Axon repulsion, VEGFA-VEGFR2 Pathway, VEGFR2 mediated vascular permeability 
                                               SEBELIUS_2_C4636 
                                               PAK2 
                                               mRNA 
                                               0 
                                               
                                               
                                               Nef and signal transduction;CD28 dependent Vav1 pathway;Stimulation of the cell death response by PAK-2p34;Ephrin signaling;Activation of Rac;RHO GTPases activate PAKs;VEGFR2 mediated vascular permeability;Sema3A PAK dependent Axon repulsion;CD209 (DC-SIGN) signaling;VEGFA-VEGFR2 Pathway;CD28 co-stimulation 
                                               hsa04010;hsa04012;hsa04014;hsa04360;hsa04510;hsa04660;hsa04810;hsa05211 
                                               PAK2 
                                               NA 
                                               azure4 
                                               1 
                                               1.50614754098361 
                                             
                                           
                                           DE Results - Activation of Rac genes - New.Annotation: miR142 vs.cel67  Table displaying each sample's global significance scores and directed global significance scores as defined in the heatmaps above. The global significance score is calculated as the square root of the mean squared t-statistic for the genes in a gene set, with t-statistics coming from the linear regression underlying our differential expression analysis. The directed global significance score is calculated as the square root of the mean signed squared t-statistic for the genes in a gene set, with t-statistics coming from the linear regression underlying our differential expression analysis.   
                                      
                                       
                                  
                                
                                
                                  
                                       
                                      
                                           Volcano Plot: New.Annotation: miR155 vs.cel67   More Plot Information                Volcano Plot: New.Annotation: miR155 vs.cel67  Volcano plot displaying each gene's -log10(p-value) and log2 fold change for the selected covariate. Highly statistically significant genes fall at the top of the plot, and highly differentially expressed genes fall to either side. Genes within the selected gene set are highlighted in orange. Horizontal lines indicate various False Discovery Rate (FDR) thresholds.   
                                           DE Results - Activation of Rac genes - New.Annotation: miR155 vs.cel67   More Plot Information      Download CSV Data     
                                           
                                             
                                             
                                           
                                           
                                             
                                               Probe Label 
                                               Log2 fold change 
                                               std error (log2) 
                                               Lower confidence limit (log2) 
                                               Upper confidence limit (log2) 
                                               Linear fold change 
                                               Lower confidence limit (linear) 
                                               Upper confidence limit (linear) 
                                               P-value 
                                               BY.p.value 
                                               method 
                                               Gene.sets 
                                               Codeset.Name 
                                               Probe.Label 
                                               Analyte.Type 
                                               Is.Control 
                                               Control.Type 
                                               Related.Probes 
                                               Probe.Annotation 
                                               KEGG.Pathways 
                                               Official.Gene.Name 
                                               Control.Conc 
                                               volcanocol 
                                               volcanopch 
                                               tstats.all 
                                             
                                           
                                           
                                             
                                               RAC1-mRNA 
                                               -0.507 
                                               0.0711 
                                               -0.647 
                                               -0.368 
                                               0.703 
                                               0.639 
                                               0.775 
                                               4.93e-07 
                                               8.18e-05 
                                               loglinear 
                                               Activation of Rac, CD28 co-stimulation, CD28 dependent Vav1 pathway, DAP12 signaling, EPH-ephrin mediated repulsion of cells, Ephrin signaling, G alpha (1213) signalling events, GPVI-mediated activation cascade, MAPK6MAPK4 signaling, NRAGE signals death through JNK, PCPCE pathway, Regulation of actin dynamics for phagocytic cup formation, RHO GTPases activate CIT, RHO GTPases Activate Formins, RHO GTPases activate IQGAPs, RHO GTPases activate PAKs, RHO GTPases activate PKNs, RHO GTPases Activate WASPs and WAVEs, Sema3A PAK dependent Axon repulsion, Sema4D induced cell migration and growth-cone collapse, Signal transduction by L1, Signaling by SCF-KIT, VEGFA-VEGFR2 Pathway, VEGFR2 mediated vascular permeability 
                                               SEBELIUS_2_C4636 
                                               RAC1 
                                               mRNA 
                                               0 
                                               
                                               
                                               MAPK6MAPK4 signaling;Signaling by SCF-KIT;Inactivation of Cdc42 and Rac;DAP12 signaling;Ephrin signaling;EPH-ephrin mediated repulsion of cells;Activation of Rac;RHO GTPases Activate Formins;VEGFR2 mediated vascular permeability;RHO GTPases activate PAKs;Sema4D mediated inhibition of cell attachment and migration;G alpha (1213) signalling events;Regulation of actin dynamics for phagocytic cup formation;Sema4D induced cell migration and growth-cone collapse;Signal transduction by L1;RHO GTPases activate CIT;RHO GTPases Activate WASPs and WAVEs;Factors involved in megakaryocyte development and platelet production;RHO GTPases activate KTN1;CD28 dependent Vav1 pathway;DSCAM interactions;NRAGE signals death through JNK;GPVI-mediated activation cascade;Sema3A PAK dependent Axon repulsion;RHO GTPases activate PKNs;SEMA3A-Plexin repulsion signaling by inhibiting Integrin adhesion;RHO GTPases activate IQGAPs;PCPCE pathway;VEGFA-VEGFR2 Pathway;CD28 co-stimulation 
                                               hsa04010;hsa04014;hsa04015;hsa04024;hsa04062;hsa04071;hsa04145;hsa04151;hsa04310;hsa04360;hsa04370;hsa04380;hsa04510;hsa04520;hsa04620;hsa04650;hsa04662;hsa04664;hsa04666;hsa04670;hsa04722;hsa04810;hsa04932;hsa04933;hsa04972;hsa05100;hsa05120;hsa05131;hsa05132;hsa05200;hsa05203;hsa05205;hsa05210;hsa05211;hsa05212;hsa05231;hsa05416 
                                               RAC1 
                                               NA 
                                               slateblue1 
                                               16 
                                               -7.13080168776371 
                                             
                                             
                                               PAK2-mRNA 
                                               -0.548 
                                               0.103 
                                               -0.75 
                                               -0.347 
                                               0.684 
                                               0.595 
                                               0.786 
                                               2.77e-05 
                                               0.00263 
                                               loglinear 
                                               Activation of Rac, CD209 (DC-SIGN) signaling, CD28 co-stimulation, CD28 dependent Vav1 pathway, Ephrin signaling, RHO GTPases activate PAKs, Sema3A PAK dependent Axon repulsion, VEGFA-VEGFR2 Pathway, VEGFR2 mediated vascular permeability 
                                               SEBELIUS_2_C4636 
                                               PAK2 
                                               mRNA 
                                               0 
                                               
                                               
                                               Nef and signal transduction;CD28 dependent Vav1 pathway;Stimulation of the cell death response by PAK-2p34;Ephrin signaling;Activation of Rac;RHO GTPases activate PAKs;VEGFR2 mediated vascular permeability;Sema3A PAK dependent Axon repulsion;CD209 (DC-SIGN) signaling;VEGFA-VEGFR2 Pathway;CD28 co-stimulation 
                                               hsa04010;hsa04012;hsa04014;hsa04360;hsa04510;hsa04660;hsa04810;hsa05211 
                                               PAK2 
                                               NA 
                                               slateblue1 
                                               16 
                                               -5.32038834951456 
                                             
                                             
                                               PAK4-mRNA 
                                               -0.359 
                                               0.131 
                                               -0.616 
                                               -0.102 
                                               0.78 
                                               0.652 
                                               0.932 
                                               0.0123 
                                               0.292 
                                               loglinear 
                                               Activation of Rac 
                                               SEBELIUS_2_C4636 
                                               PAK4 
                                               mRNA 
                                               0 
                                               
                                               
                                               Activation of Rac 
                                               hsa04012;hsa04014;hsa04360;hsa04510;hsa04660;hsa04810;hsa05206;hsa05211 
                                               PAK4 
                                               NA 
                                               slateblue4 
                                               16 
                                               -2.74045801526718 
                                             
                                             
                                               SOS2-mRNA 
                                               0.24 
                                               0.0885 
                                               0.067 
                                               0.414 
                                               1.18 
                                               1.05 
                                               1.33 
                                               0.0129 
                                               0.292 
                                               loglinear 
                                               Activation of Rac, G alpha (1213) signalling events, NRAGE signals death through JNK, Rho GTPase cycle 
                                               SEBELIUS_2_C4636 
                                               SOS2 
                                               mRNA 
                                               0 
                                               
                                               
                                               Rho GTPase cycle;Activation of Rac;NRAGE signals death through JNK;G alpha (1213) signalling events 
                                               hsa04010;hsa04012;hsa04014;hsa04062;hsa04068;hsa04072;hsa04151;hsa04320;hsa04510;hsa04540;hsa04630;hsa04650;hsa04660;hsa04662;hsa04664;hsa04722;hsa04810;hsa04910;hsa04912;hsa04915;hsa04917;hsa05034;hsa05160;hsa05200;hsa05205;hsa05206;hsa05211;hsa05213;hsa05214;hsa05215;hsa05220;hsa05221;hsa05223;hsa05231 
                                               SOS2 
                                               NA 
                                               slateblue4 
                                               16 
                                               2.71186440677966 
                                             
                                             
                                               PAK1-mRNA 
                                               -0.105 
                                               0.172 
                                               -0.441 
                                               0.231 
                                               0.93 
                                               0.736 
                                               1.17 
                                               0.547 
                                               1 
                                               lm.nb 
                                               Activation of Rac, CD209 (DC-SIGN) signaling, CD28 co-stimulation, CD28 dependent Vav1 pathway, EPHB-mediated forward signaling, Ephrin signaling, FCERI mediated MAPK activation, MAPK6MAPK4 signaling, Regulation of actin dynamics for phagocytic cup formation, RHO GTPases activate PAKs, Sema3A PAK dependent Axon repulsion, VEGFA-VEGFR2 Pathway, VEGFR2 mediated vascular permeability 
                                               SEBELIUS_2_C4636 
                                               PAK1 
                                               mRNA 
                                               0 
                                               
                                               
                                               EPHB-mediated forward signaling;Ephrin signaling;MAPK6MAPK4 signaling;DSCAM interactions;Activation of Rac;VEGFR2 mediated vascular permeability;RHO GTPases activate PAKs;Sema3A PAK dependent Axon repulsion;FCERI mediated MAPK activation;Regulation of actin dynamics for phagocytic cup formation;CD209 (DC-SIGN) signaling;CD28 dependent Vav1 pathway;VEGFA-VEGFR2 Pathway;CD28 co-stimulation 
                                               hsa04010;hsa04012;hsa04014;hsa04024;hsa04062;hsa04360;hsa04510;hsa04650;hsa04660;hsa04666;hsa04810;hsa05120;hsa05205;hsa05211 
                                               PAK1 
                                               NA 
                                               azure4 
                                               1 
                                               -0.61046511627907 
                                             
                                           
                                           DE Results - Activation of Rac genes - New.Annotation: miR155 vs.cel67  Table displaying each sample's global significance scores and directed global significance scores as defined in the heatmaps above. The global significance score is calculated as the square root of the mean squared t-statistic for the genes in a gene set, with t-statistics coming from the linear regression underlying our differential expression analysis. The directed global significance score is calculated as the square root of the mean signed squared t-statistic for the genes in a gene set, with t-statistics coming from the linear regression underlying our differential expression analysis.   
                                      
                                       
                                  
                                
                                
                                  
                                       
                                      
                                           Volcano Plot: New.Annotation: miR18b vs.cel67   More Plot Information                Volcano Plot: New.Annotation: miR18b vs.cel67  Volcano plot displaying each gene's -log10(p-value) and log2 fold change for the selected covariate. Highly statistically significant genes fall at the top of the plot, and highly differentially expressed genes fall to either side. Genes within the selected gene set are highlighted in orange. Horizontal lines indicate various False Discovery Rate (FDR) thresholds.   
                                           DE Results - Activation of Rac genes - New.Annotation: miR18b vs.cel67   More Plot Information      Download CSV Data     
                                           
                                             
                                             
                                           
                                           
                                             
                                               Probe Label 
                                               Log2 fold change 
                                               std error (log2) 
                                               Lower confidence limit (log2) 
                                               Upper confidence limit (log2) 
                                               Linear fold change 
                                               Lower confidence limit (linear) 
                                               Upper confidence limit (linear) 
                                               P-value 
                                               BY.p.value 
                                               method 
                                               Gene.sets 
                                               Codeset.Name 
                                               Probe.Label 
                                               Analyte.Type 
                                               Is.Control 
                                               Control.Type 
                                               Related.Probes 
                                               Probe.Annotation 
                                               KEGG.Pathways 
                                               Official.Gene.Name 
                                               Control.Conc 
                                               volcanocol 
                                               volcanopch 
                                               tstats.all 
                                             
                                           
                                           
                                             
                                               PAK4-mRNA 
                                               -1.05 
                                               0.142 
                                               -1.33 
                                               -0.776 
                                               0.482 
                                               0.397 
                                               0.584 
                                               2.6e-07 
                                               3.45e-05 
                                               loglinear 
                                               Activation of Rac 
                                               SEBELIUS_2_C4636 
                                               PAK4 
                                               mRNA 
                                               0 
                                               
                                               
                                               Activation of Rac 
                                               hsa04012;hsa04014;hsa04360;hsa04510;hsa04660;hsa04810;hsa05206;hsa05211 
                                               PAK4 
                                               NA 
                                               slateblue1 
                                               16 
                                               -7.3943661971831 
                                             
                                             
                                               PAK1-mRNA 
                                               -0.488 
                                               0.186 
                                               -0.853 
                                               -0.123 
                                               0.713 
                                               0.554 
                                               0.918 
                                               0.016 
                                               0.272 
                                               lm.nb 
                                               Activation of Rac, CD209 (DC-SIGN) signaling, CD28 co-stimulation, CD28 dependent Vav1 pathway, EPHB-mediated forward signaling, Ephrin signaling, FCERI mediated MAPK activation, MAPK6MAPK4 signaling, Regulation of actin dynamics for phagocytic cup formation, RHO GTPases activate PAKs, Sema3A PAK dependent Axon repulsion, VEGFA-VEGFR2 Pathway, VEGFR2 mediated vascular permeability 
                                               SEBELIUS_2_C4636 
                                               PAK1 
                                               mRNA 
                                               0 
                                               
                                               
                                               EPHB-mediated forward signaling;Ephrin signaling;MAPK6MAPK4 signaling;DSCAM interactions;Activation of Rac;VEGFR2 mediated vascular permeability;RHO GTPases activate PAKs;Sema3A PAK dependent Axon repulsion;FCERI mediated MAPK activation;Regulation of actin dynamics for phagocytic cup formation;CD209 (DC-SIGN) signaling;CD28 dependent Vav1 pathway;VEGFA-VEGFR2 Pathway;CD28 co-stimulation 
                                               hsa04010;hsa04012;hsa04014;hsa04024;hsa04062;hsa04360;hsa04510;hsa04650;hsa04660;hsa04666;hsa04810;hsa05120;hsa05205;hsa05211 
                                               PAK1 
                                               NA 
                                               slateblue4 
                                               16 
                                               -2.62365591397849 
                                             
                                             
                                               SOS2-mRNA 
                                               0.165 
                                               0.0956 
                                               -0.0224 
                                               0.352 
                                               1.12 
                                               0.985 
                                               1.28 
                                               0.099 
                                               1 
                                               loglinear 
                                               Activation of Rac, G alpha (1213) signalling events, NRAGE signals death through JNK, Rho GTPase cycle 
                                               SEBELIUS_2_C4636 
                                               SOS2 
                                               mRNA 
                                               0 
                                               
                                               
                                               Rho GTPase cycle;Activation of Rac;NRAGE signals death through JNK;G alpha (1213) signalling events 
                                               hsa04010;hsa04012;hsa04014;hsa04062;hsa04068;hsa04072;hsa04151;hsa04320;hsa04510;hsa04540;hsa04630;hsa04650;hsa04660;hsa04662;hsa04664;hsa04722;hsa04810;hsa04910;hsa04912;hsa04915;hsa04917;hsa05034;hsa05160;hsa05200;hsa05205;hsa05206;hsa05211;hsa05213;hsa05214;hsa05215;hsa05220;hsa05221;hsa05223;hsa05231 
                                               SOS2 
                                               NA 
                                               azure4 
                                               1 
                                               1.72594142259414 
                                             
                                             
                                               PAK2-mRNA 
                                               0.105 
                                               0.111 
                                               -0.112 
                                               0.323 
                                               1.08 
                                               0.925 
                                               1.25 
                                               0.354 
                                               1 
                                               loglinear 
                                               Activation of Rac, CD209 (DC-SIGN) signaling, CD28 co-stimulation, CD28 dependent Vav1 pathway, Ephrin signaling, RHO GTPases activate PAKs, Sema3A PAK dependent Axon repulsion, VEGFA-VEGFR2 Pathway, VEGFR2 mediated vascular permeability 
                                               SEBELIUS_2_C4636 
                                               PAK2 
                                               mRNA 
                                               0 
                                               
                                               
                                               Nef and signal transduction;CD28 dependent Vav1 pathway;Stimulation of the cell death response by PAK-2p34;Ephrin signaling;Activation of Rac;RHO GTPases activate PAKs;VEGFR2 mediated vascular permeability;Sema3A PAK dependent Axon repulsion;CD209 (DC-SIGN) signaling;VEGFA-VEGFR2 Pathway;CD28 co-stimulation 
                                               hsa04010;hsa04012;hsa04014;hsa04360;hsa04510;hsa04660;hsa04810;hsa05211 
                                               PAK2 
                                               NA 
                                               azure4 
                                               1 
                                               0.945945945945946 
                                             
                                             
                                               RAC1-mRNA 
                                               -0.0312 
                                               0.0768 
                                               -0.182 
                                               0.119 
                                               0.979 
                                               0.882 
                                               1.09 
                                               0.688 
                                               1 
                                               loglinear 
                                               Activation of Rac, CD28 co-stimulation, CD28 dependent Vav1 pathway, DAP12 signaling, EPH-ephrin mediated repulsion of cells, Ephrin signaling, G alpha (1213) signalling events, GPVI-mediated activation cascade, MAPK6MAPK4 signaling, NRAGE signals death through JNK, PCPCE pathway, Regulation of actin dynamics for phagocytic cup formation, RHO GTPases activate CIT, RHO GTPases Activate Formins, RHO GTPases activate IQGAPs, RHO GTPases activate PAKs, RHO GTPases activate PKNs, RHO GTPases Activate WASPs and WAVEs, Sema3A PAK dependent Axon repulsion, Sema4D induced cell migration and growth-cone collapse, Signal transduction by L1, Signaling by SCF-KIT, VEGFA-VEGFR2 Pathway, VEGFR2 mediated vascular permeability 
                                               SEBELIUS_2_C4636 
                                               RAC1 
                                               mRNA 
                                               0 
                                               
                                               
                                               MAPK6MAPK4 signaling;Signaling by SCF-KIT;Inactivation of Cdc42 and Rac;DAP12 signaling;Ephrin signaling;EPH-ephrin mediated repulsion of cells;Activation of Rac;RHO GTPases Activate Formins;VEGFR2 mediated vascular permeability;RHO GTPases activate PAKs;Sema4D mediated inhibition of cell attachment and migration;G alpha (1213) signalling events;Regulation of actin dynamics for phagocytic cup formation;Sema4D induced cell migration and growth-cone collapse;Signal transduction by L1;RHO GTPases activate CIT;RHO GTPases Activate WASPs and WAVEs;Factors involved in megakaryocyte development and platelet production;RHO GTPases activate KTN1;CD28 dependent Vav1 pathway;DSCAM interactions;NRAGE signals death through JNK;GPVI-mediated activation cascade;Sema3A PAK dependent Axon repulsion;RHO GTPases activate PKNs;SEMA3A-Plexin repulsion signaling by inhibiting Integrin adhesion;RHO GTPases activate IQGAPs;PCPCE pathway;VEGFA-VEGFR2 Pathway;CD28 co-stimulation 
                                               hsa04010;hsa04014;hsa04015;hsa04024;hsa04062;hsa04071;hsa04145;hsa04151;hsa04310;hsa04360;hsa04370;hsa04380;hsa04510;hsa04520;hsa04620;hsa04650;hsa04662;hsa04664;hsa04666;hsa04670;hsa04722;hsa04810;hsa04932;hsa04933;hsa04972;hsa05100;hsa05120;hsa05131;hsa05132;hsa05200;hsa05203;hsa05205;hsa05210;hsa05211;hsa05212;hsa05231;hsa05416 
                                               RAC1 
                                               NA 
                                               azure4 
                                               1 
                                               -0.40625 
                                             
                                           
                                           DE Results - Activation of Rac genes - New.Annotation: miR18b vs.cel67  Table displaying each sample's global significance scores and directed global significance scores as defined in the heatmaps above. The global significance score is calculated as the square root of the mean squared t-statistic for the genes in a gene set, with t-statistics coming from the linear regression underlying our differential expression analysis. The directed global significance score is calculated as the square root of the mean signed squared t-statistic for the genes in a gene set, with t-statistics coming from the linear regression underlying our differential expression analysis.   
                                      
                                       
                                  
                                
                                
                                  
                                       
                                      
                                           Volcano Plot: New.Annotation: miR890 vs.cel67   More Plot Information                Volcano Plot: New.Annotation: miR890 vs.cel67  Volcano plot displaying each gene's -log10(p-value) and log2 fold change for the selected covariate. Highly statistically significant genes fall at the top of the plot, and highly differentially expressed genes fall to either side. Genes within the selected gene set are highlighted in orange. Horizontal lines indicate various False Discovery Rate (FDR) thresholds.   
                                           DE Results - Activation of Rac genes - New.Annotation: miR890 vs.cel67   More Plot Information      Download CSV Data     
                                           
                                             
                                             
                                           
                                           
                                             
                                               Probe Label 
                                               Log2 fold change 
                                               std error (log2) 
                                               Lower confidence limit (log2) 
                                               Upper confidence limit (log2) 
                                               Linear fold change 
                                               Lower confidence limit (linear) 
                                               Upper confidence limit (linear) 
                                               P-value 
                                               BY.p.value 
                                               method 
                                               Gene.sets 
                                               Codeset.Name 
                                               Probe.Label 
                                               Analyte.Type 
                                               Is.Control 
                                               Control.Type 
                                               Related.Probes 
                                               Probe.Annotation 
                                               KEGG.Pathways 
                                               Official.Gene.Name 
                                               Control.Conc 
                                               volcanocol 
                                               volcanopch 
                                               tstats.all 
                                             
                                           
                                           
                                             
                                               PAK2-mRNA 
                                               -0.478 
                                               0.0939 
                                               -0.662 
                                               -0.293 
                                               0.718 
                                               0.632 
                                               0.816 
                                               4.92e-05 
                                               0.00233 
                                               loglinear 
                                               Activation of Rac, CD209 (DC-SIGN) signaling, CD28 co-stimulation, CD28 dependent Vav1 pathway, Ephrin signaling, RHO GTPases activate PAKs, Sema3A PAK dependent Axon repulsion, VEGFA-VEGFR2 Pathway, VEGFR2 mediated vascular permeability 
                                               SEBELIUS_2_C4636 
                                               PAK2 
                                               mRNA 
                                               0 
                                               
                                               
                                               Nef and signal transduction;CD28 dependent Vav1 pathway;Stimulation of the cell death response by PAK-2p34;Ephrin signaling;Activation of Rac;RHO GTPases activate PAKs;VEGFR2 mediated vascular permeability;Sema3A PAK dependent Axon repulsion;CD209 (DC-SIGN) signaling;VEGFA-VEGFR2 Pathway;CD28 co-stimulation 
                                               hsa04010;hsa04012;hsa04014;hsa04360;hsa04510;hsa04660;hsa04810;hsa05211 
                                               PAK2 
                                               NA 
                                               slateblue1 
                                               16 
                                               -5.09052183173589 
                                             
                                             
                                               SOS2-mRNA 
                                               -0.164 
                                               0.0808 
                                               -0.322 
                                               -0.00538 
                                               0.893 
                                               0.8 
                                               0.996 
                                               0.0556 
                                               0.753 
                                               loglinear 
                                               Activation of Rac, G alpha (1213) signalling events, NRAGE signals death through JNK, Rho GTPase cycle 
                                               SEBELIUS_2_C4636 
                                               SOS2 
                                               mRNA 
                                               0 
                                               
                                               
                                               Rho GTPase cycle;Activation of Rac;NRAGE signals death through JNK;G alpha (1213) signalling events 
                                               hsa04010;hsa04012;hsa04014;hsa04062;hsa04068;hsa04072;hsa04151;hsa04320;hsa04510;hsa04540;hsa04630;hsa04650;hsa04660;hsa04662;hsa04664;hsa04722;hsa04810;hsa04910;hsa04912;hsa04915;hsa04917;hsa05034;hsa05160;hsa05200;hsa05205;hsa05206;hsa05211;hsa05213;hsa05214;hsa05215;hsa05220;hsa05221;hsa05223;hsa05231 
                                               SOS2 
                                               NA 
                                               azure4 
                                               1 
                                               -2.02970297029703 
                                             
                                             
                                               PAK4-mRNA 
                                               -0.167 
                                               0.12 
                                               -0.402 
                                               0.0675 
                                               0.891 
                                               0.757 
                                               1.05 
                                               0.177 
                                               1 
                                               loglinear 
                                               Activation of Rac 
                                               SEBELIUS_2_C4636 
                                               PAK4 
                                               mRNA 
                                               0 
                                               
                                               
                                               Activation of Rac 
                                               hsa04012;hsa04014;hsa04360;hsa04510;hsa04660;hsa04810;hsa05206;hsa05211 
                                               PAK4 
                                               NA 
                                               azure4 
                                               1 
                                               -1.39166666666667 
                                             
                                             
                                               PAK1-mRNA 
                                               -0.16 
                                               0.157 
                                               -0.467 
                                               0.147 
                                               0.895 
                                               0.723 
                                               1.11 
                                               0.318 
                                               1 
                                               lm.nb 
                                               Activation of Rac, CD209 (DC-SIGN) signaling, CD28 co-stimulation, CD28 dependent Vav1 pathway, EPHB-mediated forward signaling, Ephrin signaling, FCERI mediated MAPK activation, MAPK6MAPK4 signaling, Regulation of actin dynamics for phagocytic cup formation, RHO GTPases activate PAKs, Sema3A PAK dependent Axon repulsion, VEGFA-VEGFR2 Pathway, VEGFR2 mediated vascular permeability 
                                               SEBELIUS_2_C4636 
                                               PAK1 
                                               mRNA 
                                               0 
                                               
                                               
                                               EPHB-mediated forward signaling;Ephrin signaling;MAPK6MAPK4 signaling;DSCAM interactions;Activation of Rac;VEGFR2 mediated vascular permeability;RHO GTPases activate PAKs;Sema3A PAK dependent Axon repulsion;FCERI mediated MAPK activation;Regulation of actin dynamics for phagocytic cup formation;CD209 (DC-SIGN) signaling;CD28 dependent Vav1 pathway;VEGFA-VEGFR2 Pathway;CD28 co-stimulation 
                                               hsa04010;hsa04012;hsa04014;hsa04024;hsa04062;hsa04360;hsa04510;hsa04650;hsa04660;hsa04666;hsa04810;hsa05120;hsa05205;hsa05211 
                                               PAK1 
                                               NA 
                                               azure4 
                                               1 
                                               -1.01910828025478 
                                             
                                             
                                               RAC1-mRNA 
                                               0.0558 
                                               0.0649 
                                               -0.0715 
                                               0.183 
                                               1.04 
                                               0.952 
                                               1.14 
                                               0.4 
                                               1 
                                               loglinear 
                                               Activation of Rac, CD28 co-stimulation, CD28 dependent Vav1 pathway, DAP12 signaling, EPH-ephrin mediated repulsion of cells, Ephrin signaling, G alpha (1213) signalling events, GPVI-mediated activation cascade, MAPK6MAPK4 signaling, NRAGE signals death through JNK, PCPCE pathway, Regulation of actin dynamics for phagocytic cup formation, RHO GTPases activate CIT, RHO GTPases Activate Formins, RHO GTPases activate IQGAPs, RHO GTPases activate PAKs, RHO GTPases activate PKNs, RHO GTPases Activate WASPs and WAVEs, Sema3A PAK dependent Axon repulsion, Sema4D induced cell migration and growth-cone collapse, Signal transduction by L1, Signaling by SCF-KIT, VEGFA-VEGFR2 Pathway, VEGFR2 mediated vascular permeability 
                                               SEBELIUS_2_C4636 
                                               RAC1 
                                               mRNA 
                                               0 
                                               
                                               
                                               MAPK6MAPK4 signaling;Signaling by SCF-KIT;Inactivation of Cdc42 and Rac;DAP12 signaling;Ephrin signaling;EPH-ephrin mediated repulsion of cells;Activation of Rac;RHO GTPases Activate Formins;VEGFR2 mediated vascular permeability;RHO GTPases activate PAKs;Sema4D mediated inhibition of cell attachment and migration;G alpha (1213) signalling events;Regulation of actin dynamics for phagocytic cup formation;Sema4D induced cell migration and growth-cone collapse;Signal transduction by L1;RHO GTPases activate CIT;RHO GTPases Activate WASPs and WAVEs;Factors involved in megakaryocyte development and platelet production;RHO GTPases activate KTN1;CD28 dependent Vav1 pathway;DSCAM interactions;NRAGE signals death through JNK;GPVI-mediated activation cascade;Sema3A PAK dependent Axon repulsion;RHO GTPases activate PKNs;SEMA3A-Plexin repulsion signaling by inhibiting Integrin adhesion;RHO GTPases activate IQGAPs;PCPCE pathway;VEGFA-VEGFR2 Pathway;CD28 co-stimulation 
                                               hsa04010;hsa04014;hsa04015;hsa04024;hsa04062;hsa04071;hsa04145;hsa04151;hsa04310;hsa04360;hsa04370;hsa04380;hsa04510;hsa04520;hsa04620;hsa04650;hsa04662;hsa04664;hsa04666;hsa04670;hsa04722;hsa04810;hsa04932;hsa04933;hsa04972;hsa05100;hsa05120;hsa05131;hsa05132;hsa05200;hsa05203;hsa05205;hsa05210;hsa05211;hsa05212;hsa05231;hsa05416 
                                               RAC1 
                                               NA 
                                               azure4 
                                               1 
                                               0.859784283513097 
                                             
                                           
                                           DE Results - Activation of Rac genes - New.Annotation: miR890 vs.cel67  Table displaying each sample's global significance scores and directed global significance scores as defined in the heatmaps above. The global significance score is calculated as the square root of the mean squared t-statistic for the genes in a gene set, with t-statistics coming from the linear regression underlying our differential expression analysis. The directed global significance score is calculated as the square root of the mean signed squared t-statistic for the genes in a gene set, with t-statistics coming from the linear regression underlying our differential expression analysis.   
                                      
                                       
                                  
                                
                              
                             
                          
                           
                      
                    
                    
                      
                           
                          
                              New.Annotation: 
                              
                                  New.Annotation: differential expression in Control vs. baseline of cel67  
                                  New.Annotation: differential expression in miR142 vs. baseline of cel67  
                                  New.Annotation: differential expression in miR155 vs. baseline of cel67  
                                  New.Annotation: differential expression in miR18b vs. baseline of cel67  
                                  New.Annotation: differential expression in miR890 vs. baseline of cel67  
                              
                              
                              
                                
                                  
                                       
                                      
                                           Volcano Plot: New.Annotation: Control vs.cel67   More Plot Information                Volcano Plot: New.Annotation: Control vs.cel67  Volcano plot displaying each gene's -log10(p-value) and log2 fold change for the selected covariate. Highly statistically significant genes fall at the top of the plot, and highly differentially expressed genes fall to either side. Genes within the selected gene set are highlighted in orange. Horizontal lines indicate various False Discovery Rate (FDR) thresholds.   
                                           DE Results - Apoptotic cleavage of cellular proteins genes - New.Annotation: Control vs.cel67   More Plot Information      Download CSV Data     
                                           
                                             
                                             
                                           
                                           
                                             
                                               Probe Label 
                                               Log2 fold change 
                                               std error (log2) 
                                               Lower confidence limit (log2) 
                                               Upper confidence limit (log2) 
                                               Linear fold change 
                                               Lower confidence limit (linear) 
                                               Upper confidence limit (linear) 
                                               P-value 
                                               BY.p.value 
                                               method 
                                               Gene.sets 
                                               Codeset.Name 
                                               Probe.Label 
                                               Analyte.Type 
                                               Is.Control 
                                               Control.Type 
                                               Related.Probes 
                                               Probe.Annotation 
                                               KEGG.Pathways 
                                               Official.Gene.Name 
                                               Control.Conc 
                                               volcanocol 
                                               volcanopch 
                                               tstats.all 
                                             
                                           
                                           
                                             
                                               CTNNB1-mRNA 
                                               0.141 
                                               0.056 
                                               0.0317 
                                               0.251 
                                               1.1 
                                               1.02 
                                               1.19 
                                               0.0196 
                                               0.743 
                                               loglinear 
                                               Apoptotic cleavage of cellular proteins, Beta-catenin phosphorylation cascade, Deactivation of the beta-catenin transactivating complex, Disassembly of the destruction complex and recruitment of AXIN to the membrane, Formation of the beta-catenin.CL.TCF transactivating complex, Misspliced GSK3beta mutants stabilize beta-catenin, Repression of WNT target genes, RHO GTPases activate IQGAPs, S37 mutants of beta-catenin aren.SQT.t phosphorylated, S45 mutants of beta-catenin aren.SQT.t phosphorylated, T41 mutants of beta-catenin aren.SQT.t phosphorylated, VEGFA-VEGFR2 Pathway, VEGFR2 mediated vascular permeability 
                                               SEBELIUS_2_C4636 
                                               CTNNB1 
                                               mRNA 
                                               0 
                                               
                                               
                                               Disassembly of the destruction complex and recruitment of AXIN to the membrane;S37 mutants of beta-catenin aren.SQT.t phosphorylated;T41 mutants of beta-catenin aren.SQT.t phosphorylated;S45 mutants of beta-catenin aren.SQT.t phosphorylated;Binding of TCFLEF.CL.CTNNB1 to target gene promoters;RHO GTPases activate IQGAPs;Deactivation of the beta-catenin transactivating complex;Repression of WNT target genes;Apoptotic cleavage of cell adhesion  proteins;Adherens junctions interactions;Misspliced GSK3beta mutants stabilize beta-catenin;VEGFR2 mediated vascular permeability;Beta-catenin phosphorylation cascade;Synthesis# secretion# and inactivation of Glucagon-like Peptide-1 (GLP-1);Formation of the beta-catenin.CL.TCF transactivating complex;Apoptotic cleavage of cellular proteins;VEGFA-VEGFR2 Pathway 
                                               hsa04015;hsa04310;hsa04390;hsa04510;hsa04520;hsa04530;hsa04550;hsa04670;hsa04916;hsa04919;hsa05100;hsa05130;hsa05166;hsa05200;hsa05205;hsa05210;hsa05213;hsa05215;hsa05216;hsa05217;hsa05412 
                                               CTNNB1 
                                               NA 
                                               azure4 
                                               1 
                                               2.51785714285714 
                                             
                                             
                                               TJP1-mRNA 
                                               0.17 
                                               0.077 
                                               0.0185 
                                               0.321 
                                               1.12 
                                               1.01 
                                               1.25 
                                               0.0391 
                                               0.99 
                                               loglinear 
                                               Apoptotic cleavage of cellular proteins 
                                               SEBELIUS_2_C4636 
                                               TJP1 
                                               mRNA 
                                               0 
                                               
                                               
                                               Apoptotic cleavage of cellular proteins 
                                               hsa04520;hsa04530;hsa04540;hsa05110;hsa05120;hsa05132 
                                               TJP1 
                                               NA 
                                               azure4 
                                               1 
                                               2.20779220779221 
                                             
                                             
                                               APC-mRNA 
                                               -0.21 
                                               0.0957 
                                               -0.397 
                                               -0.0221 
                                               0.865 
                                               0.759 
                                               0.985 
                                               0.0399 
                                               0.99 
                                               loglinear 
                                               Apoptotic cleavage of cellular proteins, AXIN missense mutants destabilize the destruction complex, Beta-catenin phosphorylation cascade, Deactivation of the beta-catenin transactivating complex, Disassembly of the destruction complex and recruitment of AXIN to the membrane, Misspliced GSK3beta mutants stabilize beta-catenin, S33 mutants of beta-catenin aren.SQT.t phosphorylated, S37 mutants of beta-catenin aren.SQT.t phosphorylated, S45 mutants of beta-catenin aren.SQT.t phosphorylated, T41 mutants of beta-catenin aren.SQT.t phosphorylated, Truncations of AMER1 destabilize the destruction complex 
                                               SEBELIUS_2_C4636 
                                               APC 
                                               mRNA 
                                               0 
                                               
                                               
                                               Apoptotic cleavage of cellular proteins;Misspliced GSK3beta mutants stabilize beta-catenin;APC truncation mutants have impaired AXIN binding;T41 mutants of beta-catenin aren.SQT.t phosphorylated;AXIN missense mutants destabilize the destruction complex;S45 mutants of beta-catenin aren.SQT.t phosphorylated;Deactivation of the beta-catenin transactivating complex;Disassembly of the destruction complex and recruitment of AXIN to the membrane;S37 mutants of beta-catenin aren.SQT.t phosphorylated;S33 mutants of beta-catenin aren.SQT.t phosphorylated;Beta-catenin phosphorylation cascade;Truncations of AMER1 destabilize the destruction complex 
                                               hsa04310;hsa04390;hsa04550;hsa04810;hsa05166;hsa05200;hsa05206;hsa05210;hsa05213;hsa05217 
                                               APC 
                                               NA 
                                               azure4 
                                               1 
                                               -2.19435736677116 
                                             
                                             
                                               SPTAN1-mRNA 
                                               0.217 
                                               0.111 
                                               -0.00162 
                                               0.435 
                                               1.16 
                                               0.999 
                                               1.35 
                                               0.0652 
                                               1 
                                               loglinear 
                                               Apoptotic cleavage of cellular proteins, ARMS-mediated activation, DAP12 signaling, Downstream signal transduction, FCERI mediated MAPK activation, Frs2-mediated activation, Gastrin-CREB signalling pathway via PKC and MAPK, Interleukin receptor SHC signaling, IRS-related events triggered by IGF1R, Nephrin interactions, Signaling by ERBB2, Signaling by PDGF, Signaling by SCF-KIT, VEGFA-VEGFR2 Pathway, VEGFR2 mediated cell proliferation 
                                               SEBELIUS_2_C4636 
                                               SPTAN1 
                                               mRNA 
                                               0 
                                               
                                               
                                               Nephrin interactions;Caspase-mediated cleavage of cytoskeletal proteins;Downstream signal transduction;VEGFR2 mediated cell proliferation;Signaling by SCF-KIT;Frs2-mediated activation;ARMS-mediated activation;Gastrin-CREB signalling pathway via PKC and MAPK;Interleukin receptor SHC signaling;FCERI mediated MAPK activation;DAP12 signaling;Signaling by PDGF;VEGFA-VEGFR2 Pathway;Signaling by ERBB2;IRS-related events triggered by IGF1R;Apoptotic cleavage of cellular proteins 
                                               hsa04530 
                                               SPTAN1 
                                               NA 
                                               azure4 
                                               1 
                                               1.95495495495495 
                                             
                                             
                                               ROCK1-mRNA 
                                               0.0597 
                                               0.0565 
                                               -0.051 
                                               0.17 
                                               1.04 
                                               0.965 
                                               1.13 
                                               0.303 
                                               1 
                                               loglinear 
                                               Apoptotic cleavage of cellular proteins, EPHA-mediated growth cone collapse, EPHB-mediated forward signaling, G alpha (1213) signalling events, RHO GTPases Activate ROCKs, Sema4D induced cell migration and growth-cone collapse, VEGFA-VEGFR2 Pathway 
                                               SEBELIUS_2_C4636 
                                               ROCK1 
                                               mRNA 
                                               0 
                                               
                                               
                                               EPHB-mediated forward signaling;EPHA-mediated growth cone collapse;RHO GTPases Activate ROCKs;Apoptotic cleavage of cellular proteins;G alpha (1213) signalling events;VEGFA-VEGFR2 Pathway;Sema4D induced cell migration and growth-cone collapse 
                                               hsa04022;hsa04024;hsa04062;hsa04071;hsa04270;hsa04350;hsa04360;hsa04510;hsa04611;hsa04670;hsa04810;hsa04921;hsa05130;hsa05131;hsa05132;hsa05200;hsa05205;hsa05206 
                                               ROCK1 
                                               NA 
                                               azure4 
                                               1 
                                               1.05663716814159 
                                             
                                             
                                               OCLN-mRNA 
                                               0.203 
                                               0.245 
                                               -0.278 
                                               0.683 
                                               1.15 
                                               0.825 
                                               1.61 
                                               0.417 
                                               1 
                                               lm.nb 
                                               Apoptotic cleavage of cellular proteins 
                                               SEBELIUS_2_C4636 
                                               OCLN 
                                               mRNA 
                                               0 
                                               
                                               
                                               Apoptotic cleavage of cellular proteins 
                                               hsa04514;hsa04530;hsa04670;hsa05130;hsa05160 
                                               OCLN 
                                               NA 
                                               azure4 
                                               1 
                                               0.828571428571429 
                                             
                                             
                                               TJP2-mRNA 
                                               -0.0502 
                                               0.242 
                                               -0.525 
                                               0.424 
                                               0.966 
                                               0.695 
                                               1.34 
                                               0.838 
                                               1 
                                               lm.nb 
                                               Apoptotic cleavage of cellular proteins 
                                               SEBELIUS_2_C4636 
                                               TJP2 
                                               mRNA 
                                               0 
                                               
                                               
                                               Apoptotic cleavage of cellular proteins 
                                               hsa04530;hsa05110 
                                               TJP2 
                                               NA 
                                               azure4 
                                               1 
                                               -0.207438016528926 
                                             
                                             
                                               PTK2-mRNA 
                                               0.00651 
                                               0.0697 
                                               -0.13 
                                               0.143 
                                               1 
                                               0.914 
                                               1.1 
                                               0.926 
                                               1 
                                               loglinear 
                                               Apoptotic cleavage of cellular proteins, ARMS-mediated activation, DAP12 signaling, DCC mediated attractive signaling, Downstream signal transduction, EPHA-mediated growth cone collapse, EPHB-mediated forward signaling, FCERI mediated MAPK activation, Frs2-mediated activation, Gastrin-CREB signalling pathway via PKC and MAPK, Integrin alphaIIb beta3 signaling, Interleukin receptor SHC signaling, IRS-related events triggered by IGF1R, p130Cas linkage to MAPK signaling for integrins, RAFMAP kinase cascade, Regulation of actin dynamics for phagocytic cup formation, RHO GTPases Activate WASPs and WAVEs, Signaling by ERBB2, Signaling by PDGF, Signaling by SCF-KIT, VEGFA-VEGFR2 Pathway, VEGFR2 mediated cell proliferation 
                                               SEBELIUS_2_C4636 
                                               PTK2 
                                               mRNA 
                                               0 
                                               
                                               
                                               Apoptotic cleavage of cellular proteins;RAFMAP kinase cascade;RHO GTPases Activate WASPs and WAVEs;DCC mediated attractive signaling;p130Cas linkage to MAPK signaling for integrins;EPHB-mediated forward signaling;EPHA-mediated growth cone collapse;Regulation of actin dynamics for phagocytic cup formation;Downstream signal transduction;VEGFR2 mediated cell proliferation;Signaling by SCF-KIT;Frs2-mediated activation;ARMS-mediated activation;Gastrin-CREB signalling pathway via PKC and MAPK;Interleukin receptor SHC signaling;FCERI mediated MAPK activation;DAP12 signaling;Signaling by PDGF;VEGFA-VEGFR2 Pathway;Signaling by ERBB2;IRS-related events triggered by IGF1R;Integrin alphaIIb beta3 signaling 
                                               hsa04012;hsa04062;hsa04151;hsa04360;hsa04370;hsa04510;hsa04670;hsa04810;hsa05100;hsa05146;hsa05200;hsa05202;hsa05205;hsa05222 
                                               PTK2 
                                               NA 
                                               azure4 
                                               1 
                                               0.0934002869440459 
                                             
                                           
                                           DE Results - Apoptotic cleavage of cellular proteins genes - New.Annotation: Control vs.cel67  Table displaying each sample's global significance scores and directed global significance scores as defined in the heatmaps above. The global significance score is calculated as the square root of the mean squared t-statistic for the genes in a gene set, with t-statistics coming from the linear regression underlying our differential expression analysis. The directed global significance score is calculated as the square root of the mean signed squared t-statistic for the genes in a gene set, with t-statistics coming from the linear regression underlying our differential expression analysis.   
                                      
                                       
                                  
                                
                                
                                  
                                       
                                      
                                           Volcano Plot: New.Annotation: miR142 vs.cel67   More Plot Information                Volcano Plot: New.Annotation: miR142 vs.cel67  Volcano plot displaying each gene's -log10(p-value) and log2 fold change for the selected covariate. Highly statistically significant genes fall at the top of the plot, and highly differentially expressed genes fall to either side. Genes within the selected gene set are highlighted in orange. Horizontal lines indicate various False Discovery Rate (FDR) thresholds.   
                                           DE Results - Apoptotic cleavage of cellular proteins genes - New.Annotation: miR142 vs.cel67   More Plot Information      Download CSV Data     
                                           
                                             
                                             
                                           
                                           
                                             
                                               Probe Label 
                                               Log2 fold change 
                                               std error (log2) 
                                               Lower confidence limit (log2) 
                                               Upper confidence limit (log2) 
                                               Linear fold change 
                                               Lower confidence limit (linear) 
                                               Upper confidence limit (linear) 
                                               P-value 
                                               BY.p.value 
                                               method 
                                               Gene.sets 
                                               Codeset.Name 
                                               Probe.Label 
                                               Analyte.Type 
                                               Is.Control 
                                               Control.Type 
                                               Related.Probes 
                                               Probe.Annotation 
                                               KEGG.Pathways 
                                               Official.Gene.Name 
                                               Control.Conc 
                                               volcanocol 
                                               volcanopch 
                                               tstats.all 
                                             
                                           
                                           
                                             
                                               APC-mRNA 
                                               -0.593 
                                               0.0957 
                                               -0.781 
                                               -0.406 
                                               0.663 
                                               0.582 
                                               0.755 
                                               3.8e-06 
                                               0.000196 
                                               loglinear 
                                               Apoptotic cleavage of cellular proteins, AXIN missense mutants destabilize the destruction complex, Beta-catenin phosphorylation cascade, Deactivation of the beta-catenin transactivating complex, Disassembly of the destruction complex and recruitment of AXIN to the membrane, Misspliced GSK3beta mutants stabilize beta-catenin, S33 mutants of beta-catenin aren.SQT.t phosphorylated, S37 mutants of beta-catenin aren.SQT.t phosphorylated, S45 mutants of beta-catenin aren.SQT.t phosphorylated, T41 mutants of beta-catenin aren.SQT.t phosphorylated, Truncations of AMER1 destabilize the destruction complex 
                                               SEBELIUS_2_C4636 
                                               APC 
                                               mRNA 
                                               0 
                                               
                                               
                                               Apoptotic cleavage of cellular proteins;Misspliced GSK3beta mutants stabilize beta-catenin;APC truncation mutants have impaired AXIN binding;T41 mutants of beta-catenin aren.SQT.t phosphorylated;AXIN missense mutants destabilize the destruction complex;S45 mutants of beta-catenin aren.SQT.t phosphorylated;Deactivation of the beta-catenin transactivating complex;Disassembly of the destruction complex and recruitment of AXIN to the membrane;S37 mutants of beta-catenin aren.SQT.t phosphorylated;S33 mutants of beta-catenin aren.SQT.t phosphorylated;Beta-catenin phosphorylation cascade;Truncations of AMER1 destabilize the destruction complex 
                                               hsa04310;hsa04390;hsa04550;hsa04810;hsa05166;hsa05200;hsa05206;hsa05210;hsa05213;hsa05217 
                                               APC 
                                               NA 
                                               slateblue1 
                                               16 
                                               -6.19644723092999 
                                             
                                             
                                               SPTAN1-mRNA 
                                               0.38 
                                               0.111 
                                               0.162 
                                               0.598 
                                               1.3 
                                               1.12 
                                               1.51 
                                               0.00263 
                                               0.0492 
                                               loglinear 
                                               Apoptotic cleavage of cellular proteins, ARMS-mediated activation, DAP12 signaling, Downstream signal transduction, FCERI mediated MAPK activation, Frs2-mediated activation, Gastrin-CREB signalling pathway via PKC and MAPK, Interleukin receptor SHC signaling, IRS-related events triggered by IGF1R, Nephrin interactions, Signaling by ERBB2, Signaling by PDGF, Signaling by SCF-KIT, VEGFA-VEGFR2 Pathway, VEGFR2 mediated cell proliferation 
                                               SEBELIUS_2_C4636 
                                               SPTAN1 
                                               mRNA 
                                               0 
                                               
                                               
                                               Nephrin interactions;Caspase-mediated cleavage of cytoskeletal proteins;Downstream signal transduction;VEGFR2 mediated cell proliferation;Signaling by SCF-KIT;Frs2-mediated activation;ARMS-mediated activation;Gastrin-CREB signalling pathway via PKC and MAPK;Interleukin receptor SHC signaling;FCERI mediated MAPK activation;DAP12 signaling;Signaling by PDGF;VEGFA-VEGFR2 Pathway;Signaling by ERBB2;IRS-related events triggered by IGF1R;Apoptotic cleavage of cellular proteins 
                                               hsa04530 
                                               SPTAN1 
                                               NA 
                                               slateblue3 
                                               16 
                                               3.42342342342342 
                                             
                                             
                                               OCLN-mRNA 
                                               0.814 
                                               0.241 
                                               0.342 
                                               1.29 
                                               1.76 
                                               1.27 
                                               2.44 
                                               0.00281 
                                               0.051 
                                               lm.nb 
                                               Apoptotic cleavage of cellular proteins 
                                               SEBELIUS_2_C4636 
                                               OCLN 
                                               mRNA 
                                               0 
                                               
                                               
                                               Apoptotic cleavage of cellular proteins 
                                               hsa04514;hsa04530;hsa04670;hsa05130;hsa05160 
                                               OCLN 
                                               NA 
                                               slateblue3 
                                               16 
                                               3.37759336099585 
                                             
                                             
                                               TJP1-mRNA 
                                               0.194 
                                               0.077 
                                               0.0434 
                                               0.345 
                                               1.14 
                                               1.03 
                                               1.27 
                                               0.0197 
                                               0.265 
                                               loglinear 
                                               Apoptotic cleavage of cellular proteins 
                                               SEBELIUS_2_C4636 
                                               TJP1 
                                               mRNA 
                                               0 
                                               
                                               
                                               Apoptotic cleavage of cellular proteins 
                                               hsa04520;hsa04530;hsa04540;hsa05110;hsa05120;hsa05132 
                                               TJP1 
                                               NA 
                                               slateblue4 
                                               16 
                                               2.51948051948052 
                                             
                                             
                                               ROCK1-mRNA 
                                               0.107 
                                               0.0565 
                                               -0.00401 
                                               0.217 
                                               1.08 
                                               0.997 
                                               1.16 
                                               0.0728 
                                               0.799 
                                               loglinear 
                                               Apoptotic cleavage of cellular proteins, EPHA-mediated growth cone collapse, EPHB-mediated forward signaling, G alpha (1213) signalling events, RHO GTPases Activate ROCKs, Sema4D induced cell migration and growth-cone collapse, VEGFA-VEGFR2 Pathway 
                                               SEBELIUS_2_C4636 
                                               ROCK1 
                                               mRNA 
                                               0 
                                               
                                               
                                               EPHB-mediated forward signaling;EPHA-mediated growth cone collapse;RHO GTPases Activate ROCKs;Apoptotic cleavage of cellular proteins;G alpha (1213) signalling events;VEGFA-VEGFR2 Pathway;Sema4D induced cell migration and growth-cone collapse 
                                               hsa04022;hsa04024;hsa04062;hsa04071;hsa04270;hsa04350;hsa04360;hsa04510;hsa04611;hsa04670;hsa04810;hsa04921;hsa05130;hsa05131;hsa05132;hsa05200;hsa05205;hsa05206 
                                               ROCK1 
                                               NA 
                                               azure4 
                                               1 
                                               1.89380530973451 
                                             
                                             
                                               TJP2-mRNA 
                                               -0.435 
                                               0.246 
                                               -0.917 
                                               0.046 
                                               0.739 
                                               0.53 
                                               1.03 
                                               0.0908 
                                               0.927 
                                               lm.nb 
                                               Apoptotic cleavage of cellular proteins 
                                               SEBELIUS_2_C4636 
                                               TJP2 
                                               mRNA 
                                               0 
                                               
                                               
                                               Apoptotic cleavage of cellular proteins 
                                               hsa04530;hsa05110 
                                               TJP2 
                                               NA 
                                               azure4 
                                               1 
                                               -1.76829268292683 
                                             
                                             
                                               PTK2-mRNA 
                                               -0.115 
                                               0.0697 
                                               -0.252 
                                               0.0215 
                                               0.923 
                                               0.84 
                                               1.01 
                                               0.113 
                                               1 
                                               loglinear 
                                               Apoptotic cleavage of cellular proteins, ARMS-mediated activation, DAP12 signaling, DCC mediated attractive signaling, Downstream signal transduction, EPHA-mediated growth cone collapse, EPHB-mediated forward signaling, FCERI mediated MAPK activation, Frs2-mediated activation, Gastrin-CREB signalling pathway via PKC and MAPK, Integrin alphaIIb beta3 signaling, Interleukin receptor SHC signaling, IRS-related events triggered by IGF1R, p130Cas linkage to MAPK signaling for integrins, RAFMAP kinase cascade, Regulation of actin dynamics for phagocytic cup formation, RHO GTPases Activate WASPs and WAVEs, Signaling by ERBB2, Signaling by PDGF, Signaling by SCF-KIT, VEGFA-VEGFR2 Pathway, VEGFR2 mediated cell proliferation 
                                               SEBELIUS_2_C4636 
                                               PTK2 
                                               mRNA 
                                               0 
                                               
                                               
                                               Apoptotic cleavage of cellular proteins;RAFMAP kinase cascade;RHO GTPases Activate WASPs and WAVEs;DCC mediated attractive signaling;p130Cas linkage to MAPK signaling for integrins;EPHB-mediated forward signaling;EPHA-mediated growth cone collapse;Regulation of actin dynamics for phagocytic cup formation;Downstream signal transduction;VEGFR2 mediated cell proliferation;Signaling by SCF-KIT;Frs2-mediated activation;ARMS-mediated activation;Gastrin-CREB signalling pathway via PKC and MAPK;Interleukin receptor SHC signaling;FCERI mediated MAPK activation;DAP12 signaling;Signaling by PDGF;VEGFA-VEGFR2 Pathway;Signaling by ERBB2;IRS-related events triggered by IGF1R;Integrin alphaIIb beta3 signaling 
                                               hsa04012;hsa04062;hsa04151;hsa04360;hsa04370;hsa04510;hsa04670;hsa04810;hsa05100;hsa05146;hsa05200;hsa05202;hsa05205;hsa05222 
                                               PTK2 
                                               NA 
                                               azure4 
                                               1 
                                               -1.64992826398852 
                                             
                                             
                                               CTNNB1-mRNA 
                                               0.0818 
                                               0.056 
                                               -0.0278 
                                               0.191 
                                               1.06 
                                               0.981 
                                               1.14 
                                               0.158 
                                               1 
                                               loglinear 
                                               Apoptotic cleavage of cellular proteins, Beta-catenin phosphorylation cascade, Deactivation of the beta-catenin transactivating complex, Disassembly of the destruction complex and recruitment of AXIN to the membrane, Formation of the beta-catenin.CL.TCF transactivating complex, Misspliced GSK3beta mutants stabilize beta-catenin, Repression of WNT target genes, RHO GTPases activate IQGAPs, S37 mutants of beta-catenin aren.SQT.t phosphorylated, S45 mutants of beta-catenin aren.SQT.t phosphorylated, T41 mutants of beta-catenin aren.SQT.t phosphorylated, VEGFA-VEGFR2 Pathway, VEGFR2 mediated vascular permeability 
                                               SEBELIUS_2_C4636 
                                               CTNNB1 
                                               mRNA 
                                               0 
                                               
                                               
                                               Disassembly of the destruction complex and recruitment of AXIN to the membrane;S37 mutants of beta-catenin aren.SQT.t phosphorylated;T41 mutants of beta-catenin aren.SQT.t phosphorylated;S45 mutants of beta-catenin aren.SQT.t phosphorylated;Binding of TCFLEF.CL.CTNNB1 to target gene promoters;RHO GTPases activate IQGAPs;Deactivation of the beta-catenin transactivating complex;Repression of WNT target genes;Apoptotic cleavage of cell adhesion  proteins;Adherens junctions interactions;Misspliced GSK3beta mutants stabilize beta-catenin;VEGFR2 mediated vascular permeability;Beta-catenin phosphorylation cascade;Synthesis# secretion# and inactivation of Glucagon-like Peptide-1 (GLP-1);Formation of the beta-catenin.CL.TCF transactivating complex;Apoptotic cleavage of cellular proteins;VEGFA-VEGFR2 Pathway 
                                               hsa04015;hsa04310;hsa04390;hsa04510;hsa04520;hsa04530;hsa04550;hsa04670;hsa04916;hsa04919;hsa05100;hsa05130;hsa05166;hsa05200;hsa05205;hsa05210;hsa05213;hsa05215;hsa05216;hsa05217;hsa05412 
                                               CTNNB1 
                                               NA 
                                               azure4 
                                               1 
                                               1.46071428571429 
                                             
                                           
                                           DE Results - Apoptotic cleavage of cellular proteins genes - New.Annotation: miR142 vs.cel67  Table displaying each sample's global significance scores and directed global significance scores as defined in the heatmaps above. The global significance score is calculated as the square root of the mean squared t-statistic for the genes in a gene set, with t-statistics coming from the linear regression underlying our differential expression analysis. The directed global significance score is calculated as the square root of the mean signed squared t-statistic for the genes in a gene set, with t-statistics coming from the linear regression underlying our differential expression analysis.   
                                      
                                       
                                  
                                
                                
                                  
                                       
                                      
                                           Volcano Plot: New.Annotation: miR155 vs.cel67   More Plot Information                Volcano Plot: New.Annotation: miR155 vs.cel67  Volcano plot displaying each gene's -log10(p-value) and log2 fold change for the selected covariate. Highly statistically significant genes fall at the top of the plot, and highly differentially expressed genes fall to either side. Genes within the selected gene set are highlighted in orange. Horizontal lines indicate various False Discovery Rate (FDR) thresholds.   
                                           DE Results - Apoptotic cleavage of cellular proteins genes - New.Annotation: miR155 vs.cel67   More Plot Information      Download CSV Data     
                                           
                                             
                                             
                                           
                                           
                                             
                                               Probe Label 
                                               Log2 fold change 
                                               std error (log2) 
                                               Lower confidence limit (log2) 
                                               Upper confidence limit (log2) 
                                               Linear fold change 
                                               Lower confidence limit (linear) 
                                               Upper confidence limit (linear) 
                                               P-value 
                                               BY.p.value 
                                               method 
                                               Gene.sets 
                                               Codeset.Name 
                                               Probe.Label 
                                               Analyte.Type 
                                               Is.Control 
                                               Control.Type 
                                               Related.Probes 
                                               Probe.Annotation 
                                               KEGG.Pathways 
                                               Official.Gene.Name 
                                               Control.Conc 
                                               volcanocol 
                                               volcanopch 
                                               tstats.all 
                                             
                                           
                                           
                                             
                                               CTNNB1-mRNA 
                                               0.163 
                                               0.059 
                                               0.0475 
                                               0.279 
                                               1.12 
                                               1.03 
                                               1.21 
                                               0.0116 
                                               0.28 
                                               loglinear 
                                               Apoptotic cleavage of cellular proteins, Beta-catenin phosphorylation cascade, Deactivation of the beta-catenin transactivating complex, Disassembly of the destruction complex and recruitment of AXIN to the membrane, Formation of the beta-catenin.CL.TCF transactivating complex, Misspliced GSK3beta mutants stabilize beta-catenin, Repression of WNT target genes, RHO GTPases activate IQGAPs, S37 mutants of beta-catenin aren.SQT.t phosphorylated, S45 mutants of beta-catenin aren.SQT.t phosphorylated, T41 mutants of beta-catenin aren.SQT.t phosphorylated, VEGFA-VEGFR2 Pathway, VEGFR2 mediated vascular permeability 
                                               SEBELIUS_2_C4636 
                                               CTNNB1 
                                               mRNA 
                                               0 
                                               
                                               
                                               Disassembly of the destruction complex and recruitment of AXIN to the membrane;S37 mutants of beta-catenin aren.SQT.t phosphorylated;T41 mutants of beta-catenin aren.SQT.t phosphorylated;S45 mutants of beta-catenin aren.SQT.t phosphorylated;Binding of TCFLEF.CL.CTNNB1 to target gene promoters;RHO GTPases activate IQGAPs;Deactivation of the beta-catenin transactivating complex;Repression of WNT target genes;Apoptotic cleavage of cell adhesion  proteins;Adherens junctions interactions;Misspliced GSK3beta mutants stabilize beta-catenin;VEGFR2 mediated vascular permeability;Beta-catenin phosphorylation cascade;Synthesis# secretion# and inactivation of Glucagon-like Peptide-1 (GLP-1);Formation of the beta-catenin.CL.TCF transactivating complex;Apoptotic cleavage of cellular proteins;VEGFA-VEGFR2 Pathway 
                                               hsa04015;hsa04310;hsa04390;hsa04510;hsa04520;hsa04530;hsa04550;hsa04670;hsa04916;hsa04919;hsa05100;hsa05130;hsa05166;hsa05200;hsa05205;hsa05210;hsa05213;hsa05215;hsa05216;hsa05217;hsa05412 
                                               CTNNB1 
                                               NA 
                                               slateblue4 
                                               16 
                                               2.76271186440678 
                                             
                                             
                                               SPTAN1-mRNA 
                                               0.316 
                                               0.117 
                                               0.0862 
                                               0.546 
                                               1.25 
                                               1.06 
                                               1.46 
                                               0.0136 
                                               0.295 
                                               loglinear 
                                               Apoptotic cleavage of cellular proteins, ARMS-mediated activation, DAP12 signaling, Downstream signal transduction, FCERI mediated MAPK activation, Frs2-mediated activation, Gastrin-CREB signalling pathway via PKC and MAPK, Interleukin receptor SHC signaling, IRS-related events triggered by IGF1R, Nephrin interactions, Signaling by ERBB2, Signaling by PDGF, Signaling by SCF-KIT, VEGFA-VEGFR2 Pathway, VEGFR2 mediated cell proliferation 
                                               SEBELIUS_2_C4636 
                                               SPTAN1 
                                               mRNA 
                                               0 
                                               
                                               
                                               Nephrin interactions;Caspase-mediated cleavage of cytoskeletal proteins;Downstream signal transduction;VEGFR2 mediated cell proliferation;Signaling by SCF-KIT;Frs2-mediated activation;ARMS-mediated activation;Gastrin-CREB signalling pathway via PKC and MAPK;Interleukin receptor SHC signaling;FCERI mediated MAPK activation;DAP12 signaling;Signaling by PDGF;VEGFA-VEGFR2 Pathway;Signaling by ERBB2;IRS-related events triggered by IGF1R;Apoptotic cleavage of cellular proteins 
                                               hsa04530 
                                               SPTAN1 
                                               NA 
                                               slateblue4 
                                               16 
                                               2.7008547008547 
                                             
                                             
                                               APC-mRNA 
                                               0.261 
                                               0.101 
                                               0.0632 
                                               0.459 
                                               1.2 
                                               1.04 
                                               1.37 
                                               0.0172 
                                               0.327 
                                               loglinear 
                                               Apoptotic cleavage of cellular proteins, AXIN missense mutants destabilize the destruction complex, Beta-catenin phosphorylation cascade, Deactivation of the beta-catenin transactivating complex, Disassembly of the destruction complex and recruitment of AXIN to the membrane, Misspliced GSK3beta mutants stabilize beta-catenin, S33 mutants of beta-catenin aren.SQT.t phosphorylated, S37 mutants of beta-catenin aren.SQT.t phosphorylated, S45 mutants of beta-catenin aren.SQT.t phosphorylated, T41 mutants of beta-catenin aren.SQT.t phosphorylated, Truncations of AMER1 destabilize the destruction complex 
                                               SEBELIUS_2_C4636 
                                               APC 
                                               mRNA 
                                               0 
                                               
                                               
                                               Apoptotic cleavage of cellular proteins;Misspliced GSK3beta mutants stabilize beta-catenin;APC truncation mutants have impaired AXIN binding;T41 mutants of beta-catenin aren.SQT.t phosphorylated;AXIN missense mutants destabilize the destruction complex;S45 mutants of beta-catenin aren.SQT.t phosphorylated;Deactivation of the beta-catenin transactivating complex;Disassembly of the destruction complex and recruitment of AXIN to the membrane;S37 mutants of beta-catenin aren.SQT.t phosphorylated;S33 mutants of beta-catenin aren.SQT.t phosphorylated;Beta-catenin phosphorylation cascade;Truncations of AMER1 destabilize the destruction complex 
                                               hsa04310;hsa04390;hsa04550;hsa04810;hsa05166;hsa05200;hsa05206;hsa05210;hsa05213;hsa05217 
                                               APC 
                                               NA 
                                               slateblue4 
                                               16 
                                               2.58415841584158 
                                             
                                             
                                               PTK2-mRNA 
                                               -0.182 
                                               0.0735 
                                               -0.326 
                                               -0.0377 
                                               0.882 
                                               0.798 
                                               0.974 
                                               0.022 
                                               0.4 
                                               loglinear 
                                               Apoptotic cleavage of cellular proteins, ARMS-mediated activation, DAP12 signaling, DCC mediated attractive signaling, Downstream signal transduction, EPHA-mediated growth cone collapse, EPHB-mediated forward signaling, FCERI mediated MAPK activation, Frs2-mediated activation, Gastrin-CREB signalling pathway via PKC and MAPK, Integrin alphaIIb beta3 signaling, Interleukin receptor SHC signaling, IRS-related events triggered by IGF1R, p130Cas linkage to MAPK signaling for integrins, RAFMAP kinase cascade, Regulation of actin dynamics for phagocytic cup formation, RHO GTPases Activate WASPs and WAVEs, Signaling by ERBB2, Signaling by PDGF, Signaling by SCF-KIT, VEGFA-VEGFR2 Pathway, VEGFR2 mediated cell proliferation 
                                               SEBELIUS_2_C4636 
                                               PTK2 
                                               mRNA 
                                               0 
                                               
                                               
                                               Apoptotic cleavage of cellular proteins;RAFMAP kinase cascade;RHO GTPases Activate WASPs and WAVEs;DCC mediated attractive signaling;p130Cas linkage to MAPK signaling for integrins;EPHB-mediated forward signaling;EPHA-mediated growth cone collapse;Regulation of actin dynamics for phagocytic cup formation;Downstream signal transduction;VEGFR2 mediated cell proliferation;Signaling by SCF-KIT;Frs2-mediated activation;ARMS-mediated activation;Gastrin-CREB signalling pathway via PKC and MAPK;Interleukin receptor SHC signaling;FCERI mediated MAPK activation;DAP12 signaling;Signaling by PDGF;VEGFA-VEGFR2 Pathway;Signaling by ERBB2;IRS-related events triggered by IGF1R;Integrin alphaIIb beta3 signaling 
                                               hsa04012;hsa04062;hsa04151;hsa04360;hsa04370;hsa04510;hsa04670;hsa04810;hsa05100;hsa05146;hsa05200;hsa05202;hsa05205;hsa05222 
                                               PTK2 
                                               NA 
                                               azure4 
                                               1 
                                               -2.47619047619048 
                                             
                                             
                                               ROCK1-mRNA 
                                               0.0972 
                                               0.0595 
                                               -0.0194 
                                               0.214 
                                               1.07 
                                               0.987 
                                               1.16 
                                               0.117 
                                               1 
                                               loglinear 
                                               Apoptotic cleavage of cellular proteins, EPHA-mediated growth cone collapse, EPHB-mediated forward signaling, G alpha (1213) signalling events, RHO GTPases Activate ROCKs, Sema4D induced cell migration and growth-cone collapse, VEGFA-VEGFR2 Pathway 
                                               SEBELIUS_2_C4636 
                                               ROCK1 
                                               mRNA 
                                               0 
                                               
                                               
                                               EPHB-mediated forward signaling;EPHA-mediated growth cone collapse;RHO GTPases Activate ROCKs;Apoptotic cleavage of cellular proteins;G alpha (1213) signalling events;VEGFA-VEGFR2 Pathway;Sema4D induced cell migration and growth-cone collapse 
                                               hsa04022;hsa04024;hsa04062;hsa04071;hsa04270;hsa04350;hsa04360;hsa04510;hsa04611;hsa04670;hsa04810;hsa04921;hsa05130;hsa05131;hsa05132;hsa05200;hsa05205;hsa05206 
                                               ROCK1 
                                               NA 
                                               azure4 
                                               1 
                                               1.63361344537815 
                                             
                                             
                                               TJP1-mRNA 
                                               0.0227 
                                               0.0812 
                                               -0.137 
                                               0.182 
                                               1.02 
                                               0.91 
                                               1.13 
                                               0.783 
                                               1 
                                               loglinear 
                                               Apoptotic cleavage of cellular proteins 
                                               SEBELIUS_2_C4636 
                                               TJP1 
                                               mRNA 
                                               0 
                                               
                                               
                                               Apoptotic cleavage of cellular proteins 
                                               hsa04520;hsa04530;hsa04540;hsa05110;hsa05120;hsa05132 
                                               TJP1 
                                               NA 
                                               azure4 
                                               1 
                                               0.279556650246305 
                                             
                                             
                                               OCLN-mRNA 
                                               -0.0515 
                                               0.261 
                                               -0.563 
                                               0.46 
                                               0.965 
                                               0.677 
                                               1.38 
                                               0.845 
                                               1 
                                               lm.nb 
                                               Apoptotic cleavage of cellular proteins 
                                               SEBELIUS_2_C4636 
                                               OCLN 
                                               mRNA 
                                               0 
                                               
                                               
                                               Apoptotic cleavage of cellular proteins 
                                               hsa04514;hsa04530;hsa04670;hsa05130;hsa05160 
                                               OCLN 
                                               NA 
                                               azure4 
                                               1 
                                               -0.197318007662835 
                                             
                                             
                                               TJP2-mRNA 
                                               -0.0336 
                                               0.255 
                                               -0.534 
                                               0.466 
                                               0.977 
                                               0.691 
                                               1.38 
                                               0.896 
                                               1 
                                               lm.nb 
                                               Apoptotic cleavage of cellular proteins 
                                               SEBELIUS_2_C4636 
                                               TJP2 
                                               mRNA 
                                               0 
                                               
                                               
                                               Apoptotic cleavage of cellular proteins 
                                               hsa04530;hsa05110 
                                               TJP2 
                                               NA 
                                               azure4 
                                               1 
                                               -0.131764705882353 
                                             
                                           
                                           DE Results - Apoptotic cleavage of cellular proteins genes - New.Annotation: miR155 vs.cel67  Table displaying each sample's global significance scores and directed global significance scores as defined in the heatmaps above. The global significance score is calculated as the square root of the mean squared t-statistic for the genes in a gene set, with t-statistics coming from the linear regression underlying our differential expression analysis. The directed global significance score is calculated as the square root of the mean signed squared t-statistic for the genes in a gene set, with t-statistics coming from the linear regression underlying our differential expression analysis.   
                                      
                                       
                                  
                                
                                
                                  
                                       
                                      
                                           Volcano Plot: New.Annotation: miR18b vs.cel67   More Plot Information                Volcano Plot: New.Annotation: miR18b vs.cel67  Volcano plot displaying each gene's -log10(p-value) and log2 fold change for the selected covariate. Highly statistically significant genes fall at the top of the plot, and highly differentially expressed genes fall to either side. Genes within the selected gene set are highlighted in orange. Horizontal lines indicate various False Discovery Rate (FDR) thresholds.   
                                           DE Results - Apoptotic cleavage of cellular proteins genes - New.Annotation: miR18b vs.cel67   More Plot Information      Download CSV Data     
                                           
                                             
                                             
                                           
                                           
                                             
                                               Probe Label 
                                               Log2 fold change 
                                               std error (log2) 
                                               Lower confidence limit (log2) 
                                               Upper confidence limit (log2) 
                                               Linear fold change 
                                               Lower confidence limit (linear) 
                                               Upper confidence limit (linear) 
                                               P-value 
                                               BY.p.value 
                                               method 
                                               Gene.sets 
                                               Codeset.Name 
                                               Probe.Label 
                                               Analyte.Type 
                                               Is.Control 
                                               Control.Type 
                                               Related.Probes 
                                               Probe.Annotation 
                                               KEGG.Pathways 
                                               Official.Gene.Name 
                                               Control.Conc 
                                               volcanocol 
                                               volcanopch 
                                               tstats.all 
                                             
                                           
                                           
                                             
                                               CTNNB1-mRNA 
                                               0.234 
                                               0.0637 
                                               0.109 
                                               0.359 
                                               1.18 
                                               1.08 
                                               1.28 
                                               0.0014 
                                               0.0333 
                                               loglinear 
                                               Apoptotic cleavage of cellular proteins, Beta-catenin phosphorylation cascade, Deactivation of the beta-catenin transactivating complex, Disassembly of the destruction complex and recruitment of AXIN to the membrane, Formation of the beta-catenin.CL.TCF transactivating complex, Misspliced GSK3beta mutants stabilize beta-catenin, Repression of WNT target genes, RHO GTPases activate IQGAPs, S37 mutants of beta-catenin aren.SQT.t phosphorylated, S45 mutants of beta-catenin aren.SQT.t phosphorylated, T41 mutants of beta-catenin aren.SQT.t phosphorylated, VEGFA-VEGFR2 Pathway, VEGFR2 mediated vascular permeability 
                                               SEBELIUS_2_C4636 
                                               CTNNB1 
                                               mRNA 
                                               0 
                                               
                                               
                                               Disassembly of the destruction complex and recruitment of AXIN to the membrane;S37 mutants of beta-catenin aren.SQT.t phosphorylated;T41 mutants of beta-catenin aren.SQT.t phosphorylated;S45 mutants of beta-catenin aren.SQT.t phosphorylated;Binding of TCFLEF.CL.CTNNB1 to target gene promoters;RHO GTPases activate IQGAPs;Deactivation of the beta-catenin transactivating complex;Repression of WNT target genes;Apoptotic cleavage of cell adhesion  proteins;Adherens junctions interactions;Misspliced GSK3beta mutants stabilize beta-catenin;VEGFR2 mediated vascular permeability;Beta-catenin phosphorylation cascade;Synthesis# secretion# and inactivation of Glucagon-like Peptide-1 (GLP-1);Formation of the beta-catenin.CL.TCF transactivating complex;Apoptotic cleavage of cellular proteins;VEGFA-VEGFR2 Pathway 
                                               hsa04015;hsa04310;hsa04390;hsa04510;hsa04520;hsa04530;hsa04550;hsa04670;hsa04916;hsa04919;hsa05100;hsa05130;hsa05166;hsa05200;hsa05205;hsa05210;hsa05213;hsa05215;hsa05216;hsa05217;hsa05412 
                                               CTNNB1 
                                               NA 
                                               slateblue3 
                                               16 
                                               3.6734693877551 
                                             
                                             
                                               SPTAN1-mRNA 
                                               0.215 
                                               0.127 
                                               -0.0337 
                                               0.463 
                                               1.16 
                                               0.977 
                                               1.38 
                                               0.105 
                                               1 
                                               loglinear 
                                               Apoptotic cleavage of cellular proteins, ARMS-mediated activation, DAP12 signaling, Downstream signal transduction, FCERI mediated MAPK activation, Frs2-mediated activation, Gastrin-CREB signalling pathway via PKC and MAPK, Interleukin receptor SHC signaling, IRS-related events triggered by IGF1R, Nephrin interactions, Signaling by ERBB2, Signaling by PDGF, Signaling by SCF-KIT, VEGFA-VEGFR2 Pathway, VEGFR2 mediated cell proliferation 
                                               SEBELIUS_2_C4636 
                                               SPTAN1 
                                               mRNA 
                                               0 
                                               
                                               
                                               Nephrin interactions;Caspase-mediated cleavage of cytoskeletal proteins;Downstream signal transduction;VEGFR2 mediated cell proliferation;Signaling by SCF-KIT;Frs2-mediated activation;ARMS-mediated activation;Gastrin-CREB signalling pathway via PKC and MAPK;Interleukin receptor SHC signaling;FCERI mediated MAPK activation;DAP12 signaling;Signaling by PDGF;VEGFA-VEGFR2 Pathway;Signaling by ERBB2;IRS-related events triggered by IGF1R;Apoptotic cleavage of cellular proteins 
                                               hsa04530 
                                               SPTAN1 
                                               NA 
                                               azure4 
                                               1 
                                               1.69291338582677 
                                             
                                             
                                               PTK2-mRNA 
                                               -0.129 
                                               0.0794 
                                               -0.285 
                                               0.0263 
                                               0.914 
                                               0.821 
                                               1.02 
                                               0.118 
                                               1 
                                               loglinear 
                                               Apoptotic cleavage of cellular proteins, ARMS-mediated activation, DAP12 signaling, DCC mediated attractive signaling, Downstream signal transduction, EPHA-mediated growth cone collapse, EPHB-mediated forward signaling, FCERI mediated MAPK activation, Frs2-mediated activation, Gastrin-CREB signalling pathway via PKC and MAPK, Integrin alphaIIb beta3 signaling, Interleukin receptor SHC signaling, IRS-related events triggered by IGF1R, p130Cas linkage to MAPK signaling for integrins, RAFMAP kinase cascade, Regulation of actin dynamics for phagocytic cup formation, RHO GTPases Activate WASPs and WAVEs, Signaling by ERBB2, Signaling by PDGF, Signaling by SCF-KIT, VEGFA-VEGFR2 Pathway, VEGFR2 mediated cell proliferation 
                                               SEBELIUS_2_C4636 
                                               PTK2 
                                               mRNA 
                                               0 
                                               
                                               
                                               Apoptotic cleavage of cellular proteins;RAFMAP kinase cascade;RHO GTPases Activate WASPs and WAVEs;DCC mediated attractive signaling;p130Cas linkage to MAPK signaling for integrins;EPHB-mediated forward signaling;EPHA-mediated growth cone collapse;Regulation of actin dynamics for phagocytic cup formation;Downstream signal transduction;VEGFR2 mediated cell proliferation;Signaling by SCF-KIT;Frs2-mediated activation;ARMS-mediated activation;Gastrin-CREB signalling pathway via PKC and MAPK;Interleukin receptor SHC signaling;FCERI mediated MAPK activation;DAP12 signaling;Signaling by PDGF;VEGFA-VEGFR2 Pathway;Signaling by ERBB2;IRS-related events triggered by IGF1R;Integrin alphaIIb beta3 signaling 
                                               hsa04012;hsa04062;hsa04151;hsa04360;hsa04370;hsa04510;hsa04670;hsa04810;hsa05100;hsa05146;hsa05200;hsa05202;hsa05205;hsa05222 
                                               PTK2 
                                               NA 
                                               azure4 
                                               1 
                                               -1.62468513853904 
                                             
                                             
                                               TJP1-mRNA 
                                               -0.109 
                                               0.0877 
                                               -0.28 
                                               0.0634 
                                               0.928 
                                               0.823 
                                               1.04 
                                               0.23 
                                               1 
                                               loglinear 
                                               Apoptotic cleavage of cellular proteins 
                                               SEBELIUS_2_C4636 
                                               TJP1 
                                               mRNA 
                                               0 
                                               
                                               
                                               Apoptotic cleavage of cellular proteins 
                                               hsa04520;hsa04530;hsa04540;hsa05110;hsa05120;hsa05132 
                                               TJP1 
                                               NA 
                                               azure4 
                                               1 
                                               -1.24287343215507 
                                             
                                             
                                               ROCK1-mRNA 
                                               0.0634 
                                               0.0643 
                                               -0.0626 
                                               0.189 
                                               1.04 
                                               0.958 
                                               1.14 
                                               0.335 
                                               1 
                                               loglinear 
                                               Apoptotic cleavage of cellular proteins, EPHA-mediated growth cone collapse, EPHB-mediated forward signaling, G alpha (1213) signalling events, RHO GTPases Activate ROCKs, Sema4D induced cell migration and growth-cone collapse, VEGFA-VEGFR2 Pathway 
                                               SEBELIUS_2_C4636 
                                               ROCK1 
                                               mRNA 
                                               0 
                                               
                                               
                                               EPHB-mediated forward signaling;EPHA-mediated growth cone collapse;RHO GTPases Activate ROCKs;Apoptotic cleavage of cellular proteins;G alpha (1213) signalling events;VEGFA-VEGFR2 Pathway;Sema4D induced cell migration and growth-cone collapse 
                                               hsa04022;hsa04024;hsa04062;hsa04071;hsa04270;hsa04350;hsa04360;hsa04510;hsa04611;hsa04670;hsa04810;hsa04921;hsa05130;hsa05131;hsa05132;hsa05200;hsa05205;hsa05206 
                                               ROCK1 
                                               NA 
                                               azure4 
                                               1 
                                               0.986003110419907 
                                             
                                             
                                               TJP2-mRNA 
                                               -0.263 
                                               0.278 
                                               -0.809 
                                               0.283 
                                               0.833 
                                               0.571 
                                               1.22 
                                               0.356 
                                               1 
                                               lm.nb 
                                               Apoptotic cleavage of cellular proteins 
                                               SEBELIUS_2_C4636 
                                               TJP2 
                                               mRNA 
                                               0 
                                               
                                               
                                               Apoptotic cleavage of cellular proteins 
                                               hsa04530;hsa05110 
                                               TJP2 
                                               NA 
                                               azure4 
                                               1 
                                               -0.946043165467626 
                                             
                                             
                                               APC-mRNA 
                                               -0.0927 
                                               0.109 
                                               -0.306 
                                               0.121 
                                               0.938 
                                               0.809 
                                               1.09 
                                               0.404 
                                               1 
                                               loglinear 
                                               Apoptotic cleavage of cellular proteins, AXIN missense mutants destabilize the destruction complex, Beta-catenin phosphorylation cascade, Deactivation of the beta-catenin transactivating complex, Disassembly of the destruction complex and recruitment of AXIN to the membrane, Misspliced GSK3beta mutants stabilize beta-catenin, S33 mutants of beta-catenin aren.SQT.t phosphorylated, S37 mutants of beta-catenin aren.SQT.t phosphorylated, S45 mutants of beta-catenin aren.SQT.t phosphorylated, T41 mutants of beta-catenin aren.SQT.t phosphorylated, Truncations of AMER1 destabilize the destruction complex 
                                               SEBELIUS_2_C4636 
                                               APC 
                                               mRNA 
                                               0 
                                               
                                               
                                               Apoptotic cleavage of cellular proteins;Misspliced GSK3beta mutants stabilize beta-catenin;APC truncation mutants have impaired AXIN binding;T41 mutants of beta-catenin aren.SQT.t phosphorylated;AXIN missense mutants destabilize the destruction complex;S45 mutants of beta-catenin aren.SQT.t phosphorylated;Deactivation of the beta-catenin transactivating complex;Disassembly of the destruction complex and recruitment of AXIN to the membrane;S37 mutants of beta-catenin aren.SQT.t phosphorylated;S33 mutants of beta-catenin aren.SQT.t phosphorylated;Beta-catenin phosphorylation cascade;Truncations of AMER1 destabilize the destruction complex 
                                               hsa04310;hsa04390;hsa04550;hsa04810;hsa05166;hsa05200;hsa05206;hsa05210;hsa05213;hsa05217 
                                               APC 
                                               NA 
                                               azure4 
                                               1 
                                               -0.85045871559633 
                                             
                                             
                                               OCLN-mRNA 
                                               1.07e-16 
                                               0.281 
                                               -0.551 
                                               0.551 
                                               1 
                                               0.682 
                                               1.47 
                                               1 
                                               1 
                                               lm.nb 
                                               Apoptotic cleavage of cellular proteins 
                                               SEBELIUS_2_C4636 
                                               OCLN 
                                               mRNA 
                                               0 
                                               
                                               
                                               Apoptotic cleavage of cellular proteins 
                                               hsa04514;hsa04530;hsa04670;hsa05130;hsa05160 
                                               OCLN 
                                               NA 
                                               azure4 
                                               1 
                                               3.80782918149466e-16 
                                             
                                           
                                           DE Results - Apoptotic cleavage of cellular proteins genes - New.Annotation: miR18b vs.cel67  Table displaying each sample's global significance scores and directed global significance scores as defined in the heatmaps above. The global significance score is calculated as the square root of the mean squared t-statistic for the genes in a gene set, with t-statistics coming from the linear regression underlying our differential expression analysis. The directed global significance score is calculated as the square root of the mean signed squared t-statistic for the genes in a gene set, with t-statistics coming from the linear regression underlying our differential expression analysis.   
                                      
                                       
                                  
                                
                                
                                  
                                       
                                      
                                           Volcano Plot: New.Annotation: miR890 vs.cel67   More Plot Information                Volcano Plot: New.Annotation: miR890 vs.cel67  Volcano plot displaying each gene's -log10(p-value) and log2 fold change for the selected covariate. Highly statistically significant genes fall at the top of the plot, and highly differentially expressed genes fall to either side. Genes within the selected gene set are highlighted in orange. Horizontal lines indicate various False Discovery Rate (FDR) thresholds.   
                                           DE Results - Apoptotic cleavage of cellular proteins genes - New.Annotation: miR890 vs.cel67   More Plot Information      Download CSV Data     
                                           
                                             
                                             
                                           
                                           
                                             
                                               Probe Label 
                                               Log2 fold change 
                                               std error (log2) 
                                               Lower confidence limit (log2) 
                                               Upper confidence limit (log2) 
                                               Linear fold change 
                                               Lower confidence limit (linear) 
                                               Upper confidence limit (linear) 
                                               P-value 
                                               BY.p.value 
                                               method 
                                               Gene.sets 
                                               Codeset.Name 
                                               Probe.Label 
                                               Analyte.Type 
                                               Is.Control 
                                               Control.Type 
                                               Related.Probes 
                                               Probe.Annotation 
                                               KEGG.Pathways 
                                               Official.Gene.Name 
                                               Control.Conc 
                                               volcanocol 
                                               volcanopch 
                                               tstats.all 
                                             
                                           
                                           
                                             
                                               ROCK1-mRNA 
                                               -0.567 
                                               0.0543 
                                               -0.674 
                                               -0.461 
                                               0.675 
                                               0.627 
                                               0.727 
                                               9.12e-10 
                                               6.05e-07 
                                               loglinear 
                                               Apoptotic cleavage of cellular proteins, EPHA-mediated growth cone collapse, EPHB-mediated forward signaling, G alpha (1213) signalling events, RHO GTPases Activate ROCKs, Sema4D induced cell migration and growth-cone collapse, VEGFA-VEGFR2 Pathway 
                                               SEBELIUS_2_C4636 
                                               ROCK1 
                                               mRNA 
                                               0 
                                               
                                               
                                               EPHB-mediated forward signaling;EPHA-mediated growth cone collapse;RHO GTPases Activate ROCKs;Apoptotic cleavage of cellular proteins;G alpha (1213) signalling events;VEGFA-VEGFR2 Pathway;Sema4D induced cell migration and growth-cone collapse 
                                               hsa04022;hsa04024;hsa04062;hsa04071;hsa04270;hsa04350;hsa04360;hsa04510;hsa04611;hsa04670;hsa04810;hsa04921;hsa05130;hsa05131;hsa05132;hsa05200;hsa05205;hsa05206 
                                               ROCK1 
                                               NA 
                                               slateblue1 
                                               16 
                                               -10.4419889502762 
                                             
                                             
                                               APC-mRNA 
                                               -0.688 
                                               0.0921 
                                               -0.869 
                                               -0.508 
                                               0.621 
                                               0.548 
                                               0.703 
                                               2.41e-07 
                                               2.92e-05 
                                               loglinear 
                                               Apoptotic cleavage of cellular proteins, AXIN missense mutants destabilize the destruction complex, Beta-catenin phosphorylation cascade, Deactivation of the beta-catenin transactivating complex, Disassembly of the destruction complex and recruitment of AXIN to the membrane, Misspliced GSK3beta mutants stabilize beta-catenin, S33 mutants of beta-catenin aren.SQT.t phosphorylated, S37 mutants of beta-catenin aren.SQT.t phosphorylated, S45 mutants of beta-catenin aren.SQT.t phosphorylated, T41 mutants of beta-catenin aren.SQT.t phosphorylated, Truncations of AMER1 destabilize the destruction complex 
                                               SEBELIUS_2_C4636 
                                               APC 
                                               mRNA 
                                               0 
                                               
                                               
                                               Apoptotic cleavage of cellular proteins;Misspliced GSK3beta mutants stabilize beta-catenin;APC truncation mutants have impaired AXIN binding;T41 mutants of beta-catenin aren.SQT.t phosphorylated;AXIN missense mutants destabilize the destruction complex;S45 mutants of beta-catenin aren.SQT.t phosphorylated;Deactivation of the beta-catenin transactivating complex;Disassembly of the destruction complex and recruitment of AXIN to the membrane;S37 mutants of beta-catenin aren.SQT.t phosphorylated;S33 mutants of beta-catenin aren.SQT.t phosphorylated;Beta-catenin phosphorylation cascade;Truncations of AMER1 destabilize the destruction complex 
                                               hsa04310;hsa04390;hsa04550;hsa04810;hsa05166;hsa05200;hsa05206;hsa05210;hsa05213;hsa05217 
                                               APC 
                                               NA 
                                               slateblue1 
                                               16 
                                               -7.47014115092291 
                                             
                                             
                                               SPTAN1-mRNA 
                                               0.467 
                                               0.107 
                                               0.257 
                                               0.677 
                                               1.38 
                                               1.19 
                                               1.6 
                                               0.000275 
                                               0.00892 
                                               loglinear 
                                               Apoptotic cleavage of cellular proteins, ARMS-mediated activation, DAP12 signaling, Downstream signal transduction, FCERI mediated MAPK activation, Frs2-mediated activation, Gastrin-CREB signalling pathway via PKC and MAPK, Interleukin receptor SHC signaling, IRS-related events triggered by IGF1R, Nephrin interactions, Signaling by ERBB2, Signaling by PDGF, Signaling by SCF-KIT, VEGFA-VEGFR2 Pathway, VEGFR2 mediated cell proliferation 
                                               SEBELIUS_2_C4636 
                                               SPTAN1 
                                               mRNA 
                                               0 
                                               
                                               
                                               Nephrin interactions;Caspase-mediated cleavage of cytoskeletal proteins;Downstream signal transduction;VEGFR2 mediated cell proliferation;Signaling by SCF-KIT;Frs2-mediated activation;ARMS-mediated activation;Gastrin-CREB signalling pathway via PKC and MAPK;Interleukin receptor SHC signaling;FCERI mediated MAPK activation;DAP12 signaling;Signaling by PDGF;VEGFA-VEGFR2 Pathway;Signaling by ERBB2;IRS-related events triggered by IGF1R;Apoptotic cleavage of cellular proteins 
                                               hsa04530 
                                               SPTAN1 
                                               NA 
                                               slateblue1 
                                               16 
                                               4.36448598130841 
                                             
                                             
                                               CTNNB1-mRNA 
                                               0.0834 
                                               0.0538 
                                               -0.0222 
                                               0.189 
                                               1.06 
                                               0.985 
                                               1.14 
                                               0.136 
                                               1 
                                               loglinear 
                                               Apoptotic cleavage of cellular proteins, Beta-catenin phosphorylation cascade, Deactivation of the beta-catenin transactivating complex, Disassembly of the destruction complex and recruitment of AXIN to the membrane, Formation of the beta-catenin.CL.TCF transactivating complex, Misspliced GSK3beta mutants stabilize beta-catenin, Repression of WNT target genes, RHO GTPases activate IQGAPs, S37 mutants of beta-catenin aren.SQT.t phosphorylated, S45 mutants of beta-catenin aren.SQT.t phosphorylated, T41 mutants of beta-catenin aren.SQT.t phosphorylated, VEGFA-VEGFR2 Pathway, VEGFR2 mediated vascular permeability 
                                               SEBELIUS_2_C4636 
                                               CTNNB1 
                                               mRNA 
                                               0 
                                               
                                               
                                               Disassembly of the destruction complex and recruitment of AXIN to the membrane;S37 mutants of beta-catenin aren.SQT.t phosphorylated;T41 mutants of beta-catenin aren.SQT.t phosphorylated;S45 mutants of beta-catenin aren.SQT.t phosphorylated;Binding of TCFLEF.CL.CTNNB1 to target gene promoters;RHO GTPases activate IQGAPs;Deactivation of the beta-catenin transactivating complex;Repression of WNT target genes;Apoptotic cleavage of cell adhesion  proteins;Adherens junctions interactions;Misspliced GSK3beta mutants stabilize beta-catenin;VEGFR2 mediated vascular permeability;Beta-catenin phosphorylation cascade;Synthesis# secretion# and inactivation of Glucagon-like Peptide-1 (GLP-1);Formation of the beta-catenin.CL.TCF transactivating complex;Apoptotic cleavage of cellular proteins;VEGFA-VEGFR2 Pathway 
                                               hsa04015;hsa04310;hsa04390;hsa04510;hsa04520;hsa04530;hsa04550;hsa04670;hsa04916;hsa04919;hsa05100;hsa05130;hsa05166;hsa05200;hsa05205;hsa05210;hsa05213;hsa05215;hsa05216;hsa05217;hsa05412 
                                               CTNNB1 
                                               NA 
                                               azure4 
                                               1 
                                               1.55018587360595 
                                             
                                             
                                               PTK2-mRNA 
                                               -0.0907 
                                               0.0671 
                                               -0.222 
                                               0.0408 
                                               0.939 
                                               0.857 
                                               1.03 
                                               0.191 
                                               1 
                                               loglinear 
                                               Apoptotic cleavage of cellular proteins, ARMS-mediated activation, DAP12 signaling, DCC mediated attractive signaling, Downstream signal transduction, EPHA-mediated growth cone collapse, EPHB-mediated forward signaling, FCERI mediated MAPK activation, Frs2-mediated activation, Gastrin-CREB signalling pathway via PKC and MAPK, Integrin alphaIIb beta3 signaling, Interleukin receptor SHC signaling, IRS-related events triggered by IGF1R, p130Cas linkage to MAPK signaling for integrins, RAFMAP kinase cascade, Regulation of actin dynamics for phagocytic cup formation, RHO GTPases Activate WASPs and WAVEs, Signaling by ERBB2, Signaling by PDGF, Signaling by SCF-KIT, VEGFA-VEGFR2 Pathway, VEGFR2 mediated cell proliferation 
                                               SEBELIUS_2_C4636 
                                               PTK2 
                                               mRNA 
                                               0 
                                               
                                               
                                               Apoptotic cleavage of cellular proteins;RAFMAP kinase cascade;RHO GTPases Activate WASPs and WAVEs;DCC mediated attractive signaling;p130Cas linkage to MAPK signaling for integrins;EPHB-mediated forward signaling;EPHA-mediated growth cone collapse;Regulation of actin dynamics for phagocytic cup formation;Downstream signal transduction;VEGFR2 mediated cell proliferation;Signaling by SCF-KIT;Frs2-mediated activation;ARMS-mediated activation;Gastrin-CREB signalling pathway via PKC and MAPK;Interleukin receptor SHC signaling;FCERI mediated MAPK activation;DAP12 signaling;Signaling by PDGF;VEGFA-VEGFR2 Pathway;Signaling by ERBB2;IRS-related events triggered by IGF1R;Integrin alphaIIb beta3 signaling 
                                               hsa04012;hsa04062;hsa04151;hsa04360;hsa04370;hsa04510;hsa04670;hsa04810;hsa05100;hsa05146;hsa05200;hsa05202;hsa05205;hsa05222 
                                               PTK2 
                                               NA 
                                               azure4 
                                               1 
                                               -1.35171385991058 
                                             
                                             
                                               TJP1-mRNA 
                                               0.0291 
                                               0.0741 
                                               -0.116 
                                               0.174 
                                               1.02 
                                               0.923 
                                               1.13 
                                               0.699 
                                               1 
                                               loglinear 
                                               Apoptotic cleavage of cellular proteins 
                                               SEBELIUS_2_C4636 
                                               TJP1 
                                               mRNA 
                                               0 
                                               
                                               
                                               Apoptotic cleavage of cellular proteins 
                                               hsa04520;hsa04530;hsa04540;hsa05110;hsa05120;hsa05132 
                                               TJP1 
                                               NA 
                                               azure4 
                                               1 
                                               0.392712550607287 
                                             
                                             
                                               TJP2-mRNA 
                                               -0.0614 
                                               0.233 
                                               -0.518 
                                               0.395 
                                               0.958 
                                               0.698 
                                               1.32 
                                               0.795 
                                               1 
                                               lm.nb 
                                               Apoptotic cleavage of cellular proteins 
                                               SEBELIUS_2_C4636 
                                               TJP2 
                                               mRNA 
                                               0 
                                               
                                               
                                               Apoptotic cleavage of cellular proteins 
                                               hsa04530;hsa05110 
                                               TJP2 
                                               NA 
                                               azure4 
                                               1 
                                               -0.263519313304721 
                                             
                                             
                                               OCLN-mRNA 
                                               -0.0559 
                                               0.238 
                                               -0.523 
                                               0.411 
                                               0.962 
                                               0.696 
                                               1.33 
                                               0.817 
                                               1 
                                               lm.nb 
                                               Apoptotic cleavage of cellular proteins 
                                               SEBELIUS_2_C4636 
                                               OCLN 
                                               mRNA 
                                               0 
                                               
                                               
                                               Apoptotic cleavage of cellular proteins 
                                               hsa04514;hsa04530;hsa04670;hsa05130;hsa05160 
                                               OCLN 
                                               NA 
                                               azure4 
                                               1 
                                               -0.234873949579832 
                                             
                                           
                                           DE Results - Apoptotic cleavage of cellular proteins genes - New.Annotation: miR890 vs.cel67  Table displaying each sample's global significance scores and directed global significance scores as defined in the heatmaps above. The global significance score is calculated as the square root of the mean squared t-statistic for the genes in a gene set, with t-statistics coming from the linear regression underlying our differential expression analysis. The directed global significance score is calculated as the square root of the mean signed squared t-statistic for the genes in a gene set, with t-statistics coming from the linear regression underlying our differential expression analysis.   
                                      
                                       
                                  
                                
                              
                             
                          
                           
                      
                    
                    
                      
                           
                          
                              New.Annotation: 
                              
                                  New.Annotation: differential expression in Control vs. baseline of cel67  
                                  New.Annotation: differential expression in miR142 vs. baseline of cel67  
                                  New.Annotation: differential expression in miR155 vs. baseline of cel67  
                                  New.Annotation: differential expression in miR18b vs. baseline of cel67  
                                  New.Annotation: differential expression in miR890 vs. baseline of cel67  
                              
                              
                              
                                
                                  
                                       
                                      
                                           Volcano Plot: New.Annotation: Control vs.cel67   More Plot Information                Volcano Plot: New.Annotation: Control vs.cel67  Volcano plot displaying each gene's -log10(p-value) and log2 fold change for the selected covariate. Highly statistically significant genes fall at the top of the plot, and highly differentially expressed genes fall to either side. Genes within the selected gene set are highlighted in orange. Horizontal lines indicate various False Discovery Rate (FDR) thresholds.   
                                           DE Results - ARMS-mediated activation genes - New.Annotation: Control vs.cel67   More Plot Information      Download CSV Data     
                                           
                                             
                                             
                                           
                                           
                                             
                                               Probe Label 
                                               Log2 fold change 
                                               std error (log2) 
                                               Lower confidence limit (log2) 
                                               Upper confidence limit (log2) 
                                               Linear fold change 
                                               Lower confidence limit (linear) 
                                               Upper confidence limit (linear) 
                                               P-value 
                                               BY.p.value 
                                               method 
                                               Gene.sets 
                                               Codeset.Name 
                                               Probe.Label 
                                               Analyte.Type 
                                               Is.Control 
                                               Control.Type 
                                               Related.Probes 
                                               Probe.Annotation 
                                               KEGG.Pathways 
                                               Official.Gene.Name 
                                               Control.Conc 
                                               volcanocol 
                                               volcanopch 
                                               tstats.all 
                                             
                                           
                                           
                                             
                                               NRAS-mRNA 
                                               -0.544 
                                               0.114 
                                               -0.767 
                                               -0.322 
                                               0.686 
                                               0.588 
                                               0.8 
                                               9.79e-05 
                                               0.026 
                                               lm.nb 
                                               ARMS-mediated activation, CD209 (DC-SIGN) signaling, Cell surface interactions at the vascular wall, Constitutive Signaling by EGFRvIII, DAP12 signaling, Downstream signal transduction, FCERI mediated MAPK activation, Frs2-mediated activation, Gastrin-CREB signalling pathway via PKC and MAPK, Interleukin receptor SHC signaling, IRS-related events triggered by IGF1R, MAP2K and MAPK activation, RAFMAP kinase cascade, SHC-mediated cascade.CL.FGFR1, SHC-related events triggered by IGF1R, Signaling by ERBB2, Signaling by FGFR1 in disease, Signaling by FGFR2 in disease, Signaling by FGFR3 in disease, Signaling by FGFR4 in disease, Signaling by PDGF, Signaling by SCF-KIT, Tie2 Signaling, VEGFA-VEGFR2 Pathway, VEGFR2 mediated cell proliferation 
                                               SEBELIUS_2_C4636 
                                               NRAS 
                                               mRNA 
                                               0 
                                               
                                               
                                               Signaling by FGFR2 in disease;Regulation of RAS by GAPs;SHC-mediated cascade.CL.FGFR3;Signaling by FGFR3 in disease;MAP2K and MAPK activation;SHC-related events triggered by IGF1R;Signaling by FGFR4 in disease;SHC-mediated cascade.CL.FGFR1;Tie2 Signaling;Signaling by FGFR1 in disease;Activation of RAS in B cells;SHC-mediated cascade.CL.FGFR4;Constitutive Signaling by EGFRvIII;CD209 (DC-SIGN) signaling;FCERI mediated MAPK activation;RAFMAP kinase cascade;Downstream signal transduction;VEGFR2 mediated cell proliferation;Signaling by SCF-KIT;Frs2-mediated activation;ARMS-mediated activation;Gastrin-CREB signalling pathway via PKC and MAPK;Interleukin receptor SHC signaling;DAP12 signaling;Signaling by PDGF;VEGFA-VEGFR2 Pathway;Signaling by ERBB2;IRS-related events triggered by IGF1R;Cell surface interactions at the vascular wall 
                                               hsa04010;hsa04012;hsa04014;hsa04015;hsa04062;hsa04068;hsa04071;hsa04072;hsa04151;hsa04211;hsa04213;hsa04360;hsa04370;hsa04530;hsa04540;hsa04550;hsa04650;hsa04660;hsa04662;hsa04664;hsa04720;hsa04722;hsa04725;hsa04726;hsa04730;hsa04810;hsa04910;hsa04912;hsa04915;hsa04916;hsa04917;hsa04919;hsa04921;hsa04933;hsa05034;hsa05160;hsa05161;hsa05166;hsa05200;hsa05203;hsa05205;hsa05206;hsa05211;hsa05213;hsa05214;hsa05215;hsa05216;hsa05218;hsa05219;hsa05220;hsa05221;hsa05223;hsa05230;hsa05231 
                                               NRAS 
                                               NA 
                                               slateblue2 
                                               16 
                                               -4.7719298245614 
                                             
                                             
                                               HRAS-mRNA 
                                               -0.593 
                                               0.144 
                                               -0.874 
                                               -0.311 
                                               0.663 
                                               0.545 
                                               0.806 
                                               0.000487 
                                               0.0718 
                                               loglinear 
                                               ARMS-mediated activation, CD209 (DC-SIGN) signaling, Cell surface interactions at the vascular wall, Constitutive Signaling by EGFRvIII, CREB phosphorylation through the activation of Ras, DAP12 signaling, Downstream signal transduction, FCERI mediated MAPK activation, Frs2-mediated activation, Gastrin-CREB signalling pathway via PKC and MAPK, Interleukin receptor SHC signaling, IRS-related events triggered by IGF1R, MAP2K and MAPK activation, RAFMAP kinase cascade, SHC-mediated cascade.CL.FGFR1, SHC-related events triggered by IGF1R, Signaling by ERBB2, Signaling by FGFR1 in disease, Signaling by FGFR2 in disease, Signaling by FGFR3 in disease, Signaling by FGFR4 in disease, Signaling by PDGF, Signaling by SCF-KIT, Tie2 Signaling, VEGFA-VEGFR2 Pathway, VEGFR2 mediated cell proliferation 
                                               SEBELIUS_2_C4636 
                                               HRAS 
                                               mRNA 
                                               0 
                                               
                                               
                                               Signaling by FGFR2 in disease;Regulation of RAS by GAPs;SHC-mediated cascade.CL.FGFR3;Signaling by FGFR3 in disease;MAP2K and MAPK activation;SHC-related events triggered by IGF1R;Signaling by FGFR4 in disease;SHC-mediated cascade.CL.FGFR1;Tie2 Signaling;Signaling by FGFR1 in disease;Activation of RAS in B cells;SHC-mediated cascade.CL.FGFR4;Constitutive Signaling by EGFRvIII;CD209 (DC-SIGN) signaling;Ras activation uopn Ca2.PLS. infux through NMDA receptor;FCERI mediated MAPK activation;RAFMAP kinase cascade;Downstream signal transduction;VEGFR2 mediated cell proliferation;Signaling by SCF-KIT;Frs2-mediated activation;ARMS-mediated activation;Gastrin-CREB signalling pathway via PKC and MAPK;Interleukin receptor SHC signaling;DAP12 signaling;Signaling by PDGF;VEGFA-VEGFR2 Pathway;Signaling by ERBB2;IRS-related events triggered by IGF1R;Cell surface interactions at the vascular wall;CREB phosphorylation through the activation of Ras 
                                               hsa04010;hsa04012;hsa04014;hsa04015;hsa04062;hsa04068;hsa04071;hsa04072;hsa04144;hsa04151;hsa04211;hsa04213;hsa04360;hsa04370;hsa04510;hsa04530;hsa04540;hsa04550;hsa04630;hsa04650;hsa04660;hsa04662;hsa04664;hsa04720;hsa04722;hsa04725;hsa04726;hsa04730;hsa04810;hsa04910;hsa04912;hsa04915;hsa04916;hsa04917;hsa04919;hsa04921;hsa04933;hsa05034;hsa05160;hsa05161;hsa05166;hsa05200;hsa05203;hsa05205;hsa05206;hsa05211;hsa05213;hsa05214;hsa05215;hsa05216;hsa05218;hsa05219;hsa05220;hsa05221;hsa05223;hsa05230;hsa05231 
                                               HRAS 
                                               NA 
                                               slateblue3 
                                               16 
                                               -4.11805555555556 
                                             
                                             
                                               EGFR-mRNA 
                                               -0.408 
                                               0.101 
                                               -0.605 
                                               -0.211 
                                               0.753 
                                               0.657 
                                               0.864 
                                               0.000564 
                                               0.0749 
                                               loglinear 
                                               ARMS-mediated activation, Constitutive Signaling by Aberrant PI3K in Cancer, Constitutive Signaling by EGFRvIII, DAP12 signaling, Downstream signal transduction, FCERI mediated MAPK activation, Frs2-mediated activation, GAB1 signalosome, Gastrin-CREB signalling pathway via PKC and MAPK, Interleukin receptor SHC signaling, IRS-related events triggered by IGF1R, PI3KAKT activation, Role of LAT2NTALLAB on calcium mobilization, Signaling by ERBB2, Signaling by PDGF, Signaling by SCF-KIT, VEGFA-VEGFR2 Pathway, VEGFR2 mediated cell proliferation 
                                               SEBELIUS_2_C4636 
                                               EGFR 
                                               mRNA 
                                               0 
                                               
                                               
                                               Constitutive Signaling by Aberrant PI3K in Cancer;Inhibition of Signaling by Overexpressed EGFR;PLCG1 events in ERBB2 signaling;EGFR interacts with phospholipase C-gamma;EGFR Transactivation by Gastrin;Constitutive Signaling by Ligand-Responsive EGFR Cancer Variants;Constitutive Signaling by EGFRvIII;GAB1 signalosome;Downstream signal transduction;Signaling by SCF-KIT;PI3KAKT activation;Role of LAT2NTALLAB on calcium mobilization;DAP12 signaling;Signaling by PDGF;Signaling by ERBB2;VEGFR2 mediated cell proliferation;Frs2-mediated activation;ARMS-mediated activation;Gastrin-CREB signalling pathway via PKC and MAPK;Interleukin receptor SHC signaling;FCERI mediated MAPK activation;VEGFA-VEGFR2 Pathway;IRS-related events triggered by IGF1R 
                                               hsa04010;hsa04012;hsa04014;hsa04015;hsa04020;hsa04060;hsa04066;hsa04068;hsa04072;hsa04144;hsa04151;hsa04320;hsa04510;hsa04520;hsa04540;hsa04810;hsa04912;hsa04915;hsa04921;hsa05120;hsa05160;hsa05200;hsa05205;hsa05206;hsa05212;hsa05213;hsa05214;hsa05215;hsa05218;hsa05219;hsa05223;hsa05230;hsa05231 
                                               EGFR 
                                               NA 
                                               slateblue4 
                                               16 
                                               -4.03960396039604 
                                             
                                             
                                               MAPK3-mRNA 
                                               0.205 
                                               0.0541 
                                               0.0991 
                                               0.311 
                                               1.15 
                                               1.07 
                                               1.24 
                                               0.00106 
                                               0.128 
                                               loglinear 
                                               ARMS-mediated activation, DAP12 signaling, Downstream signal transduction, ERKs are inactivated, FCERI mediated MAPK activation, Frs2-mediated activation, Gastrin-CREB signalling pathway via PKC and MAPK, Interleukin receptor SHC signaling, IRS-related events triggered by IGF1R, Negative feedback regulation of MAPK pathway, Negative regulation of FGFR3 signaling, Negative regulation of FGFR4 signaling, Negative regulation of MAPK pathway, RAFMAP kinase cascade, Regulation of HSF1-mediated heat shock response, RHO GTPases Activate WASPs and WAVEs, Signaling by ERBB2, Signaling by PDGF, Signaling by SCF-KIT, VEGFA-VEGFR2 Pathway, VEGFR2 mediated cell proliferation 
                                               SEBELIUS_2_C4636 
                                               MAPK3 
                                               mRNA 
                                               0 
                                               
                                               
                                               ERKs are inactivated;ISG15 antiviral mechanism;RHO GTPases Activate WASPs and WAVEs;Senescence-Associated Secretory Phenotype (SASP);Regulation of HSF1-mediated heat shock response;Thrombin signalling through proteinase activated receptors (PARs);Negative regulation of FGFR4 signaling;Oxidative Stress Induced Senescence;Signal attenuation;RNA Polymerase I Promoter Opening;Negative regulation of FGFR3 signaling;Negative feedback regulation of MAPK pathway;Growth hormone receptor signaling;Golgi Cisternae Pericentriolar Stack Reorganization;FCERI mediated MAPK activation;Negative regulation of MAPK pathway;RAFMAP kinase cascade;Downstream signal transduction;VEGFR2 mediated cell proliferation;Signaling by SCF-KIT;Frs2-mediated activation;ARMS-mediated activation;Gastrin-CREB signalling pathway via PKC and MAPK;Interleukin receptor SHC signaling;DAP12 signaling;Signaling by PDGF;VEGFA-VEGFR2 Pathway;Signaling by ERBB2;IRS-related events triggered by IGF1R 
                                               hsa04010;hsa04012;hsa04014;hsa04015;hsa04022;hsa04024;hsa04062;hsa04066;hsa04068;hsa04071;hsa04072;hsa04114;hsa04150;hsa04151;hsa04261;hsa04270;hsa04320;hsa04350;hsa04360;hsa04370;hsa04380;hsa04510;hsa04520;hsa04540;hsa04550;hsa04611;hsa04620;hsa04621;hsa04650;hsa04660;hsa04662;hsa04664;hsa04666;hsa04668;hsa04713;hsa04720;hsa04722;hsa04723;hsa04724;hsa04725;hsa04726;hsa04730;hsa04810;hsa04910;hsa04912;hsa04914;hsa04915;hsa04916;hsa04917;hsa04919;hsa04921;hsa04930;hsa04933;hsa04960;hsa05010;hsa05020;hsa05034;hsa05131;hsa05132;hsa05133;hsa05140;hsa05142;hsa05145;hsa05152;hsa05160;hsa05161;hsa05164;hsa05200;hsa05203;hsa05205;hsa05210;hsa05211;hsa05212;hsa05213;hsa05214;hsa05215;hsa05216;hsa05218;hsa05219;hsa05220;hsa05221;hsa05223;hsa05230;hsa05231 
                                               MAPK3 
                                               NA 
                                               slateblue4 
                                               16 
                                               3.78927911275416 
                                             
                                             
                                               PPP2CB-mRNA 
                                               -0.182 
                                               0.0747 
                                               -0.328 
                                               -0.0354 
                                               0.882 
                                               0.797 
                                               0.976 
                                               0.0239 
                                               0.756 
                                               loglinear 
                                               ARMS-mediated activation, AXIN missense mutants destabilize the destruction complex, Beta-catenin phosphorylation cascade, CTLA4 inhibitory signaling, Cyclin AB1 associated events during G2M transition, Cyclin D associated events in G1, DAP12 signaling, DARPP-32 events, Disassembly of the destruction complex and recruitment of AXIN to the membrane, Downstream signal transduction, ERKs are inactivated, FCERI mediated MAPK activation, Frs2-mediated activation, Gastrin-CREB signalling pathway via PKC and MAPK, Interleukin receptor SHC signaling, IRS-related events triggered by IGF1R, MASTL Facilitates Mitotic Progression, Misspliced GSK3beta mutants stabilize beta-catenin, Negative regulation of FGFR1 signaling, Negative regulation of FGFR2 signaling, Negative regulation of FGFR3 signaling, Negative regulation of FGFR4 signaling, Negative regulation of MAPK pathway, RAF activation, RAFMAP kinase cascade, Resolution of Sister Chromatid Cohesion, RHO GTPases Activate Formins, S33 mutants of beta-catenin aren.SQT.t phosphorylated, S37 mutants of beta-catenin aren.SQT.t phosphorylated, S45 mutants of beta-catenin aren.SQT.t phosphorylated, Signaling by ERBB2, Signaling by PDGF, Signaling by SCF-KIT, Spry regulation of FGF signaling, T41 mutants of beta-catenin aren.SQT.t phosphorylated, Truncations of AMER1 destabilize the destruction complex, VEGFA-VEGFR2 Pathway, VEGFR2 mediated cell proliferation 
                                               SEBELIUS_2_C4636 
                                               PPP2CB 
                                               mRNA 
                                               0 
                                               
                                               
                                               MASTL Facilitates Mitotic Progression;Misspliced GSK3beta mutants stabilize beta-catenin;T41 mutants of beta-catenin aren.SQT.t phosphorylated;ERKs are inactivated;Glycolysis;Negative regulation of MAPK pathway;Disassembly of the destruction complex and recruitment of AXIN to the membrane;S37 mutants of beta-catenin aren.SQT.t phosphorylated;Cyclin AB1 associated events during G2M transition;RAF activation;RHO GTPases Activate Formins;S33 mutants of beta-catenin aren.SQT.t phosphorylated;DARPP-32 events;Beta-catenin phosphorylation cascade;Spry regulation of FGF signaling;Truncations of AMER1 destabilize the destruction complex;CTLA4 inhibitory signaling;AXIN missense mutants destabilize the destruction complex;S45 mutants of beta-catenin aren.SQT.t phosphorylated;Cyclin D associated events in G1;Platelet sensitization by LDL;Resolution of Sister Chromatid Cohesion;Inhibition of replication initiation of damaged DNA by RB1E2F1;RAFMAP kinase cascade;Downstream signal transduction;VEGFR2 mediated cell proliferation;Signaling by SCF-KIT;Frs2-mediated activation;ARMS-mediated activation;Gastrin-CREB signalling pathway via PKC and MAPK;Interleukin receptor SHC signaling;FCERI mediated MAPK activation;DAP12 signaling;Signaling by PDGF;VEGFA-VEGFR2 Pathway;Signaling by ERBB2;IRS-related events triggered by IGF1R;Negative regulation of FGFR3 signaling;Negative regulation of FGFR4 signaling;Negative regulation of FGFR2 signaling;Negative regulation of FGFR1 signaling 
                                               hsa03015;hsa04071;hsa04114;hsa04151;hsa04152;hsa04261;hsa04350;hsa04390;hsa04530;hsa04728;hsa04730;hsa05142;hsa05160 
                                               PPP2CB 
                                               NA 
                                               azure4 
                                               1 
                                               -2.43641231593039 
                                             
                                             
                                               SRC-mRNA 
                                               0.261 
                                               0.114 
                                               0.0374 
                                               0.485 
                                               1.2 
                                               1.03 
                                               1.4 
                                               0.0326 
                                               0.908 
                                               loglinear 
                                               ARMS-mediated activation, CD28 co-stimulation, Cell surface interactions at the vascular wall, CTLA4 inhibitory signaling, DAP12 signaling, Downstream signal transduction, FCERI mediated MAPK activation, Frs2-mediated activation, GAB1 signalosome, Gastrin-CREB signalling pathway via PKC and MAPK, Integrin alphaIIb beta3 signaling, Interleukin receptor SHC signaling, IRS-related events triggered by IGF1R, MAP2K and MAPK activation, Negative regulation of FGFR1 signaling, Negative regulation of FGFR2 signaling, Negative regulation of FGFR3 signaling, Negative regulation of FGFR4 signaling, p130Cas linkage to MAPK signaling for integrins, RAFMAP kinase cascade, Signaling by ERBB2, Signaling by PDGF, Signaling by SCF-KIT, Spry regulation of FGF signaling, VEGFA-VEGFR2 Pathway, VEGFR2 mediated cell proliferation 
                                               SEBELIUS_2_C4636 
                                               SRC 
                                               mRNA 
                                               0 
                                               
                                               
                                               FCGR activation;Spry regulation of FGF signaling;GAB1 signalosome;MAP2K and MAPK activation;CLEC7A (Dectin-1) signaling;CD28 co-stimulation;p130Cas linkage to MAPK signaling for integrins;CTLA4 inhibitory signaling;GRB2.CL.SOS provides linkage to MAPK signaling for Integrins ;Signal regulatory protein (SIRP) family interactions;PECAM1 interactions;Thrombin signalling through proteinase activated receptors (PARs);GP1b-IX-V activation signalling;c-src mediated regulation of Cx43 function and closure of gap junctions;Signaling by ERBB2;VEGFA-VEGFR2 Pathway;Signaling by SCF-KIT;VEGFR2 mediated cell proliferation;RAFMAP kinase cascade;Downstream signal transduction;Frs2-mediated activation;ARMS-mediated activation;Gastrin-CREB signalling pathway via PKC and MAPK;Interleukin receptor SHC signaling;FCERI mediated MAPK activation;DAP12 signaling;Signaling by PDGF;IRS-related events triggered by IGF1R;Negative regulation of FGFR3 signaling;Negative regulation of FGFR4 signaling;Negative regulation of FGFR2 signaling;Negative regulation of FGFR1 signaling;Integrin alphaIIb beta3 signaling;Cell surface interactions at the vascular wall 
                                               hsa04012;hsa04015;hsa04062;hsa04144;hsa04370;hsa04510;hsa04520;hsa04530;hsa04540;hsa04611;hsa04727;hsa04750;hsa04810;hsa04912;hsa04915;hsa04917;hsa04919;hsa04921;hsa05100;hsa05120;hsa05131;hsa05152;hsa05161;hsa05203;hsa05205;hsa05219 
                                               SRC 
                                               NA 
                                               azure4 
                                               1 
                                               2.28947368421053 
                                             
                                             
                                               PPP2CA-mRNA 
                                               -0.115 
                                               0.0527 
                                               -0.219 
                                               -0.0119 
                                               0.923 
                                               0.859 
                                               0.992 
                                               0.0403 
                                               0.99 
                                               loglinear 
                                               ARMS-mediated activation, AXIN missense mutants destabilize the destruction complex, Beta-catenin phosphorylation cascade, CTLA4 inhibitory signaling, Cyclin AB1 associated events during G2M transition, Cyclin D associated events in G1, DAP12 signaling, DARPP-32 events, Disassembly of the destruction complex and recruitment of AXIN to the membrane, Downstream signal transduction, ERKs are inactivated, FCERI mediated MAPK activation, Frs2-mediated activation, Gastrin-CREB signalling pathway via PKC and MAPK, Interleukin receptor SHC signaling, IRS-related events triggered by IGF1R, MASTL Facilitates Mitotic Progression, Negative regulation of FGFR1 signaling, Negative regulation of FGFR2 signaling, Negative regulation of FGFR3 signaling, Negative regulation of FGFR4 signaling, Negative regulation of MAPK pathway, Nonsense Mediated Decay (NMD) enhanced by the Exon Junction Complex (EJC), RAF activation, RAFMAP kinase cascade, Resolution of Sister Chromatid Cohesion, RHO GTPases Activate Formins, S33 mutants of beta-catenin aren.SQT.t phosphorylated, S37 mutants of beta-catenin aren.SQT.t phosphorylated, S45 mutants of beta-catenin aren.SQT.t phosphorylated, Signaling by ERBB2, Signaling by PDGF, Signaling by SCF-KIT, Spry regulation of FGF signaling, T41 mutants of beta-catenin aren.SQT.t phosphorylated, Truncations of AMER1 destabilize the destruction complex, VEGFA-VEGFR2 Pathway, VEGFR2 mediated cell proliferation 
                                               SEBELIUS_2_C4636 
                                               PPP2CA 
                                               mRNA 
                                               0 
                                               
                                               
                                               MASTL Facilitates Mitotic Progression;T41 mutants of beta-catenin aren.SQT.t phosphorylated;ERKs are inactivated;Glycolysis;Negative regulation of MAPK pathway;Disassembly of the destruction complex and recruitment of AXIN to the membrane;S37 mutants of beta-catenin aren.SQT.t phosphorylated;Cyclin AB1 associated events during G2M transition;RAF activation;RHO GTPases Activate Formins;S33 mutants of beta-catenin aren.SQT.t phosphorylated;DARPP-32 events;Beta-catenin phosphorylation cascade;Initiation of Nuclear Envelope Reformation;Spry regulation of FGF signaling;Truncations of AMER1 destabilize the destruction complex;CTLA4 inhibitory signaling;AXIN missense mutants destabilize the destruction complex;S45 mutants of beta-catenin aren.SQT.t phosphorylated;Nonsense Mediated Decay (NMD) enhanced by the Exon Junction Complex (EJC);Cyclin D associated events in G1;Platelet sensitization by LDL;Resolution of Sister Chromatid Cohesion;Inhibition of replication initiation of damaged DNA by RB1E2F1;RAFMAP kinase cascade;Downstream signal transduction;VEGFR2 mediated cell proliferation;Signaling by SCF-KIT;Frs2-mediated activation;ARMS-mediated activation;Gastrin-CREB signalling pathway via PKC and MAPK;Interleukin receptor SHC signaling;FCERI mediated MAPK activation;DAP12 signaling;Signaling by PDGF;VEGFA-VEGFR2 Pathway;Signaling by ERBB2;IRS-related events triggered by IGF1R;Negative regulation of FGFR3 signaling;Negative regulation of FGFR4 signaling;Negative regulation of FGFR2 signaling;Negative regulation of FGFR1 signaling 
                                               hsa03015;hsa04071;hsa04114;hsa04151;hsa04152;hsa04261;hsa04350;hsa04390;hsa04530;hsa04728;hsa04730;hsa05142;hsa05160 
                                               PPP2CA 
                                               NA 
                                               azure4 
                                               1 
                                               -2.18216318785579 
                                             
                                             
                                               CRK-mRNA 
                                               -0.224 
                                               0.103 
                                               -0.425 
                                               -0.0221 
                                               0.856 
                                               0.745 
                                               0.985 
                                               0.0412 
                                               0.996 
                                               loglinear 
                                               ARMS-mediated activation, Downstream signal transduction, Integrin alphaIIb beta3 signaling, p130Cas linkage to MAPK signaling for integrins, Regulation of actin dynamics for phagocytic cup formation, Regulation of signaling by CBL, Signaling by PDGF, VEGFA-VEGFR2 Pathway 
                                               SEBELIUS_2_C4636 
                                               CRK 
                                               mRNA 
                                               0 
                                               
                                               
                                               ARMS-mediated activation;Downstream signal transduction;Regulation of signaling by CBL;p130Cas linkage to MAPK signaling for integrins;VEGFA-VEGFR2 Pathway;Regulation of actin dynamics for phagocytic cup formation;Signaling by PDGF;Integrin alphaIIb beta3 signaling 
                                               hsa04010;hsa04012;hsa04015;hsa04062;hsa04510;hsa04666;hsa04722;hsa04810;hsa04910;hsa05100;hsa05131;hsa05200;hsa05206;hsa05211;hsa05220 
                                               CRK 
                                               NA 
                                               azure4 
                                               1 
                                               -2.1747572815534 
                                             
                                             
                                               PDGFRB-mRNA 
                                               0.342 
                                               0.159 
                                               0.031 
                                               0.653 
                                               1.27 
                                               1.02 
                                               1.57 
                                               0.0429 
                                               1 
                                               lm.nb 
                                               ARMS-mediated activation, Constitutive Signaling by Aberrant PI3K in Cancer, DAP12 signaling, Downstream signal transduction, FCERI mediated MAPK activation, Frs2-mediated activation, GAB1 signalosome, Gastrin-CREB signalling pathway via PKC and MAPK, Interleukin receptor SHC signaling, IRS-related events triggered by IGF1R, PI3KAKT activation, PIP3 activates AKT signaling, RAFMAP kinase cascade, Role of LAT2NTALLAB on calcium mobilization, Signaling by ERBB2, Signaling by PDGF, Signaling by SCF-KIT, VEGFA-VEGFR2 Pathway, VEGFR2 mediated cell proliferation 
                                               SEBELIUS_2_C4636 
                                               PDGFRB 
                                               mRNA 
                                               0 
                                               
                                               
                                               PIP3 activates AKT signaling;Constitutive Signaling by Aberrant PI3K in Cancer;RAFMAP kinase cascade;Signaling by PDGF;Downstream signal transduction;GAB1 signalosome;Signaling by SCF-KIT;PI3KAKT activation;Role of LAT2NTALLAB on calcium mobilization;DAP12 signaling;Signaling by ERBB2;VEGFR2 mediated cell proliferation;Frs2-mediated activation;ARMS-mediated activation;Gastrin-CREB signalling pathway via PKC and MAPK;Interleukin receptor SHC signaling;FCERI mediated MAPK activation;VEGFA-VEGFR2 Pathway;IRS-related events triggered by IGF1R 
                                               hsa04010;hsa04014;hsa04015;hsa04020;hsa04060;hsa04072;hsa04151;hsa04510;hsa04540;hsa04810;hsa05166;hsa05200;hsa05206;hsa05214;hsa05215;hsa05218;hsa05230;hsa05231 
                                               PDGFRB 
                                               NA 
                                               azure4 
                                               1 
                                               2.15094339622642 
                                             
                                             
                                               SPTAN1-mRNA 
                                               0.217 
                                               0.111 
                                               -0.00162 
                                               0.435 
                                               1.16 
                                               0.999 
                                               1.35 
                                               0.0652 
                                               1 
                                               loglinear 
                                               Apoptotic cleavage of cellular proteins, ARMS-mediated activation, DAP12 signaling, Downstream signal transduction, FCERI mediated MAPK activation, Frs2-mediated activation, Gastrin-CREB signalling pathway via PKC and MAPK, Interleukin receptor SHC signaling, IRS-related events triggered by IGF1R, Nephrin interactions, Signaling by ERBB2, Signaling by PDGF, Signaling by SCF-KIT, VEGFA-VEGFR2 Pathway, VEGFR2 mediated cell proliferation 
                                               SEBELIUS_2_C4636 
                                               SPTAN1 
                                               mRNA 
                                               0 
                                               
                                               
                                               Nephrin interactions;Caspase-mediated cleavage of cytoskeletal proteins;Downstream signal transduction;VEGFR2 mediated cell proliferation;Signaling by SCF-KIT;Frs2-mediated activation;ARMS-mediated activation;Gastrin-CREB signalling pathway via PKC and MAPK;Interleukin receptor SHC signaling;FCERI mediated MAPK activation;DAP12 signaling;Signaling by PDGF;VEGFA-VEGFR2 Pathway;Signaling by ERBB2;IRS-related events triggered by IGF1R;Apoptotic cleavage of cellular proteins 
                                               hsa04530 
                                               SPTAN1 
                                               NA 
                                               azure4 
                                               1 
                                               1.95495495495495 
                                             
                                             
                                               VCL-mRNA 
                                               0.195 
                                               0.108 
                                               -0.0169 
                                               0.408 
                                               1.15 
                                               0.988 
                                               1.33 
                                               0.0856 
                                               1 
                                               lm.nb 
                                               ARMS-mediated activation, DAP12 signaling, Downstream signal transduction, FCERI mediated MAPK activation, Frs2-mediated activation, Gastrin-CREB signalling pathway via PKC and MAPK, Interleukin receptor SHC signaling, IRS-related events triggered by IGF1R, MAP2K and MAPK activation, Platelet degranulation , RAFMAP kinase cascade, Signaling by ERBB2, Signaling by PDGF, Signaling by SCF-KIT, Smooth Muscle Contraction, VEGFA-VEGFR2 Pathway, VEGFR2 mediated cell proliferation 
                                               SEBELIUS_2_C4636 
                                               VCL 
                                               mRNA 
                                               0 
                                               
                                               
                                               Smooth Muscle Contraction;MAP2K and MAPK activation;Platelet degranulation ;RAFMAP kinase cascade;Downstream signal transduction;VEGFR2 mediated cell proliferation;Signaling by SCF-KIT;Frs2-mediated activation;ARMS-mediated activation;Gastrin-CREB signalling pathway via PKC and MAPK;Interleukin receptor SHC signaling;FCERI mediated MAPK activation;DAP12 signaling;Signaling by PDGF;VEGFA-VEGFR2 Pathway;Signaling by ERBB2;IRS-related events triggered by IGF1R 
                                               hsa04510;hsa04520;hsa04670;hsa04810;hsa05100;hsa05131;hsa05146 
                                               VCL 
                                               NA 
                                               azure4 
                                               1 
                                               1.80555555555556 
                                             
                                             
                                               PPP2R1B-mRNA 
                                               -0.0747 
                                               0.0593 
                                               -0.191 
                                               0.0416 
                                               0.95 
                                               0.876 
                                               1.03 
                                               0.222 
                                               1 
                                               loglinear 
                                               ARMS-mediated activation, AXIN missense mutants destabilize the destruction complex, Beta-catenin phosphorylation cascade, CTLA4 inhibitory signaling, Cyclin AB1 associated events during G2M transition, Cyclin D associated events in G1, DAP12 signaling, DARPP-32 events, Disassembly of the destruction complex and recruitment of AXIN to the membrane, Downstream signal transduction, ERKs are inactivated, FCERI mediated MAPK activation, Frs2-mediated activation, Gastrin-CREB signalling pathway via PKC and MAPK, Interleukin receptor SHC signaling, IRS-related events triggered by IGF1R, MASTL Facilitates Mitotic Progression, Misspliced GSK3beta mutants stabilize beta-catenin, Negative regulation of MAPK pathway, RAF activation, RAFMAP kinase cascade, Resolution of Sister Chromatid Cohesion, RHO GTPases Activate Formins, S33 mutants of beta-catenin aren.SQT.t phosphorylated, S37 mutants of beta-catenin aren.SQT.t phosphorylated, S45 mutants of beta-catenin aren.SQT.t phosphorylated, Signaling by ERBB2, Signaling by PDGF, Signaling by SCF-KIT, T41 mutants of beta-catenin aren.SQT.t phosphorylated, Truncations of AMER1 destabilize the destruction complex, VEGFA-VEGFR2 Pathway, VEGFR2 mediated cell proliferation 
                                               SEBELIUS_2_C4636 
                                               PPP2R1B 
                                               mRNA 
                                               0 
                                               
                                               
                                               MASTL Facilitates Mitotic Progression;Misspliced GSK3beta mutants stabilize beta-catenin;T41 mutants of beta-catenin aren.SQT.t phosphorylated;ERKs are inactivated;Glycolysis;Negative regulation of MAPK pathway;Disassembly of the destruction complex and recruitment of AXIN to the membrane;S37 mutants of beta-catenin aren.SQT.t phosphorylated;Cyclin AB1 associated events during G2M transition;RAF activation;RHO GTPases Activate Formins;S33 mutants of beta-catenin aren.SQT.t phosphorylated;DARPP-32 events;Beta-catenin phosphorylation cascade;Truncations of AMER1 destabilize the destruction complex;CTLA4 inhibitory signaling;AXIN missense mutants destabilize the destruction complex;S45 mutants of beta-catenin aren.SQT.t phosphorylated;Cyclin D associated events in G1;Platelet sensitization by LDL;Resolution of Sister Chromatid Cohesion;Inhibition of replication initiation of damaged DNA by RB1E2F1;RAFMAP kinase cascade;Downstream signal transduction;VEGFR2 mediated cell proliferation;Signaling by SCF-KIT;Frs2-mediated activation;ARMS-mediated activation;Gastrin-CREB signalling pathway via PKC and MAPK;Interleukin receptor SHC signaling;FCERI mediated MAPK activation;DAP12 signaling;Signaling by PDGF;VEGFA-VEGFR2 Pathway;Signaling by ERBB2;IRS-related events triggered by IGF1R 
                                               hsa03015;hsa04071;hsa04114;hsa04151;hsa04152;hsa04261;hsa04350;hsa04390;hsa04530;hsa04728;hsa04730;hsa05142;hsa05160 
                                               PPP2R1B 
                                               NA 
                                               azure4 
                                               1 
                                               -1.25969645868465 
                                             
                                             
                                               PDGFRA-mRNA 
                                               0.414 
                                               0.374 
                                               -0.32 
                                               1.15 
                                               1.33 
                                               0.801 
                                               2.21 
                                               0.281 
                                               1 
                                               lm.nb 
                                               ARMS-mediated activation, Constitutive Signaling by Aberrant PI3K in Cancer, DAP12 signaling, Downstream signal transduction, FCERI mediated MAPK activation, Frs2-mediated activation, GAB1 signalosome, Gastrin-CREB signalling pathway via PKC and MAPK, Interleukin receptor SHC signaling, IRS-related events triggered by IGF1R, PI3KAKT activation, PIP3 activates AKT signaling, RAFMAP kinase cascade, Role of LAT2NTALLAB on calcium mobilization, Signaling by ERBB2, Signaling by PDGF, Signaling by SCF-KIT, VEGFA-VEGFR2 Pathway, VEGFR2 mediated cell proliferation 
                                               SEBELIUS_2_C4636 
                                               PDGFRA 
                                               mRNA 
                                               0 
                                               
                                               
                                               PIP3 activates AKT signaling;Constitutive Signaling by Aberrant PI3K in Cancer;RAFMAP kinase cascade;Signaling by PDGF;Downstream signal transduction;GAB1 signalosome;Signaling by SCF-KIT;PI3KAKT activation;Role of LAT2NTALLAB on calcium mobilization;DAP12 signaling;Signaling by ERBB2;VEGFR2 mediated cell proliferation;Frs2-mediated activation;ARMS-mediated activation;Gastrin-CREB signalling pathway via PKC and MAPK;Interleukin receptor SHC signaling;FCERI mediated MAPK activation;VEGFA-VEGFR2 Pathway;IRS-related events triggered by IGF1R 
                                               hsa04010;hsa04014;hsa04015;hsa04020;hsa04060;hsa04072;hsa04144;hsa04151;hsa04510;hsa04540;hsa04810;hsa05166;hsa05200;hsa05206;hsa05214;hsa05215;hsa05218;hsa05230;hsa05231 
                                               PDGFRA 
                                               NA 
                                               azure4 
                                               1 
                                               1.10695187165775 
                                             
                                             
                                               FN1-mRNA 
                                               0.272 
                                               0.255 
                                               -0.229 
                                               0.773 
                                               1.21 
                                               0.853 
                                               1.71 
                                               0.299 
                                               1 
                                               lm.nb 
                                               ARMS-mediated activation, DAP12 signaling, Downstream signal transduction, ECM proteoglycans, Elastic fibre formation, FCERI mediated MAPK activation, Frs2-mediated activation, Gastrin-CREB signalling pathway via PKC and MAPK, Integrin alphaIIb beta3 signaling, Integrin cell surface interactions, Interleukin receptor SHC signaling, IRS-related events triggered by IGF1R, MAP2K and MAPK activation, Molecules associated with elastic fibres, Non-integrin membrane-ECM interactions, p130Cas linkage to MAPK signaling for integrins, Platelet degranulation , RAFMAP kinase cascade, Signaling by ERBB2, Signaling by PDGF, Signaling by SCF-KIT, Syndecan interactions, VEGFA-VEGFR2 Pathway, VEGFR2 mediated cell proliferation 
                                               SEBELIUS_2_C4636 
                                               FN1 
                                               mRNA 
                                               0 
                                               
                                               
                                               MAP2K and MAPK activation;p130Cas linkage to MAPK signaling for integrins;Platelet degranulation ;Molecules associated with elastic fibres;Syndecan interactions;ECM proteoglycans;Fibronectin matrix formation;Integrin cell surface interactions;RAFMAP kinase cascade;Downstream signal transduction;VEGFR2 mediated cell proliferation;Signaling by SCF-KIT;Frs2-mediated activation;ARMS-mediated activation;Gastrin-CREB signalling pathway via PKC and MAPK;Interleukin receptor SHC signaling;FCERI mediated MAPK activation;DAP12 signaling;Signaling by PDGF;VEGFA-VEGFR2 Pathway;Signaling by ERBB2;IRS-related events triggered by IGF1R;Integrin alphaIIb beta3 signaling;Non-integrin membrane-ECM interactions;Elastic fibre formation 
                                               hsa04151;hsa04510;hsa04512;hsa04810;hsa04933;hsa05100;hsa05146;hsa05200;hsa05205;hsa05222 
                                               FN1 
                                               NA 
                                               azure4 
                                               1 
                                               1.06666666666667 
                                             
                                             
                                               MAP2K1-mRNA 
                                               -0.121 
                                               0.119 
                                               -0.355 
                                               0.112 
                                               0.919 
                                               0.782 
                                               1.08 
                                               0.321 
                                               1 
                                               lm.nb 
                                               ARMS-mediated activation, DAP12 signaling, Downstream signal transduction, FCERI mediated MAPK activation, Frs2-mediated activation, Gastrin-CREB signalling pathway via PKC and MAPK, Interleukin receptor SHC signaling, IRS-related events triggered by IGF1R, Negative feedback regulation of MAPK pathway, Negative regulation of MAPK pathway, RAF activation, RAFMAP kinase cascade, Signaling by ERBB2, Signaling by PDGF, Signaling by SCF-KIT, VEGFA-VEGFR2 Pathway, VEGFR2 mediated cell proliferation 
                                               SEBELIUS_2_C4636 
                                               MAP2K1 
                                               mRNA 
                                               0 
                                               
                                               
                                               RAF activation;Uptake and function of anthrax toxins;Negative feedback regulation of MAPK pathway;Interleukin-1 signaling;RAFMAP kinase cascade;Downstream signal transduction;VEGFR2 mediated cell proliferation;Signaling by SCF-KIT;Frs2-mediated activation;ARMS-mediated activation;Gastrin-CREB signalling pathway via PKC and MAPK;Interleukin receptor SHC signaling;FCERI mediated MAPK activation;DAP12 signaling;Signaling by PDGF;VEGFA-VEGFR2 Pathway;Signaling by ERBB2;IRS-related events triggered by IGF1R;Negative regulation of MAPK pathway 
                                               hsa04010;hsa04012;hsa04014;hsa04015;hsa04022;hsa04024;hsa04062;hsa04066;hsa04068;hsa04071;hsa04072;hsa04114;hsa04151;hsa04270;hsa04320;hsa04370;hsa04380;hsa04510;hsa04540;hsa04550;hsa04620;hsa04650;hsa04660;hsa04662;hsa04664;hsa04666;hsa04668;hsa04720;hsa04722;hsa04725;hsa04726;hsa04730;hsa04810;hsa04910;hsa04912;hsa04914;hsa04915;hsa04916;hsa04917;hsa04919;hsa04921;hsa05020;hsa05034;hsa05161;hsa05164;hsa05200;hsa05205;hsa05206;hsa05210;hsa05211;hsa05212;hsa05213;hsa05214;hsa05215;hsa05216;hsa05218;hsa05219;hsa05220;hsa05221;hsa05223;hsa05230;hsa05231 
                                               MAP2K1 
                                               NA 
                                               azure4 
                                               1 
                                               -1.01680672268908 
                                             
                                             
                                               IRS1-mRNA 
                                               -0.0946 
                                               0.108 
                                               -0.306 
                                               0.117 
                                               0.937 
                                               0.809 
                                               1.08 
                                               0.391 
                                               1 
                                               loglinear 
                                               ARMS-mediated activation, Constitutive Signaling by Aberrant PI3K in Cancer, DAP12 signaling, Downstream signal transduction, FCERI mediated MAPK activation, Frs2-mediated activation, GAB1 signalosome, Gastrin-CREB signalling pathway via PKC and MAPK, Interleukin receptor SHC signaling, IRS-related events triggered by IGF1R, PI3K Cascade, PI3KAKT activation, RAFMAP kinase cascade, Role of LAT2NTALLAB on calcium mobilization, Signaling by ERBB2, Signaling by PDGF, Signaling by SCF-KIT, VEGFA-VEGFR2 Pathway, VEGFR2 mediated cell proliferation 
                                               SEBELIUS_2_C4636 
                                               IRS1 
                                               mRNA 
                                               0 
                                               
                                               
                                               Constitutive Signaling by Aberrant PI3K in Cancer;RAFMAP kinase cascade;PI3K Cascade;Growth hormone receptor signaling;IRS-related events triggered by IGF1R;PI3KAKT activation;Downstream signal transduction;GAB1 signalosome;Signaling by SCF-KIT;Role of LAT2NTALLAB on calcium mobilization;DAP12 signaling;Signaling by PDGF;Signaling by ERBB2;VEGFR2 mediated cell proliferation;Frs2-mediated activation;ARMS-mediated activation;Gastrin-CREB signalling pathway via PKC and MAPK;Interleukin receptor SHC signaling;FCERI mediated MAPK activation;VEGFA-VEGFR2 Pathway 
                                               hsa04022;hsa04068;hsa04150;hsa04151;hsa04152;hsa04211;hsa04213;hsa04722;hsa04910;hsa04920;hsa04923;hsa04930;hsa04931;hsa04932;hsa04960;hsa05206 
                                               IRS1 
                                               NA 
                                               azure4 
                                               1 
                                               -0.875925925925926 
                                             
                                             
                                               FGFR1-mRNA 
                                               -0.0813 
                                               0.102 
                                               -0.28 
                                               0.118 
                                               0.945 
                                               0.823 
                                               1.09 
                                               0.432 
                                               1 
                                               lm.nb 
                                               ARMS-mediated activation, Constitutive Signaling by Aberrant PI3K in Cancer, DAP12 signaling, Downstream signal transduction, FCERI mediated MAPK activation, Frs2-mediated activation, GAB1 signalosome, Gastrin-CREB signalling pathway via PKC and MAPK, Interleukin receptor SHC signaling, IRS-related events triggered by IGF1R, PI3K Cascade, PI3KAKT activation, RAFMAP kinase cascade, Role of LAT2NTALLAB on calcium mobilization, SHC-mediated cascade.CL.FGFR1, Signaling by ERBB2, Signaling by FGFR1 in disease, Signaling by PDGF, Signaling by SCF-KIT, VEGFA-VEGFR2 Pathway, VEGFR2 mediated cell proliferation 
                                               SEBELIUS_2_C4636 
                                               FGFR1 
                                               mRNA 
                                               0 
                                               
                                               
                                               RAFMAP kinase cascade;Constitutive Signaling by Aberrant PI3K in Cancer;SHC-mediated cascade.CL.FGFR1;Phospholipase C-mediated cascade.CL. FGFR1;Signaling by FGFR1 amplification mutants;PI3K Cascade;FGFR1b ligand binding and activation;Signaling by FGFR1 in disease;Downstream signal transduction;GAB1 signalosome;Signaling by SCF-KIT;PI3KAKT activation;Role of LAT2NTALLAB on calcium mobilization;DAP12 signaling;Signaling by PDGF;Signaling by ERBB2;VEGFR2 mediated cell proliferation;Frs2-mediated activation;ARMS-mediated activation;Gastrin-CREB signalling pathway via PKC and MAPK;Interleukin receptor SHC signaling;FCERI mediated MAPK activation;VEGFA-VEGFR2 Pathway;IRS-related events triggered by IGF1R 
                                               hsa04010;hsa04014;hsa04015;hsa04151;hsa04520;hsa04550;hsa04810;hsa05200;hsa05205;hsa05215;hsa05218;hsa05230 
                                               FGFR1 
                                               NA 
                                               azure4 
                                               1 
                                               -0.797058823529412 
                                             
                                             
                                               RAF1-mRNA 
                                               -0.0567 
                                               0.0728 
                                               -0.199 
                                               0.0861 
                                               0.961 
                                               0.871 
                                               1.06 
                                               0.445 
                                               1 
                                               loglinear 
                                               ARMS-mediated activation, CD209 (DC-SIGN) signaling, CREB phosphorylation through the activation of Ras, DAP12 signaling, Downstream signal transduction, FCERI mediated MAPK activation, Frs2-mediated activation, Gastrin-CREB signalling pathway via PKC and MAPK, Interleukin receptor SHC signaling, IRS-related events triggered by IGF1R, Negative feedback regulation of MAPK pathway, Negative regulation of MAPK pathway, RAF activation, RAFMAP kinase cascade, Signaling by ERBB2, Signaling by PDGF, Signaling by SCF-KIT, VEGFA-VEGFR2 Pathway, VEGFR2 mediated cell proliferation 
                                               SEBELIUS_2_C4636 
                                               RAF1 
                                               mRNA 
                                               0 
                                               
                                               
                                               RAF activation;Stimuli-sensing channels;CREB phosphorylation through the activation of Ras;Negative feedback regulation of MAPK pathway;CD209 (DC-SIGN) signaling;RAFMAP kinase cascade;Downstream signal transduction;VEGFR2 mediated cell proliferation;Signaling by SCF-KIT;Frs2-mediated activation;ARMS-mediated activation;Gastrin-CREB signalling pathway via PKC and MAPK;Interleukin receptor SHC signaling;FCERI mediated MAPK activation;DAP12 signaling;Signaling by PDGF;VEGFA-VEGFR2 Pathway;Signaling by ERBB2;IRS-related events triggered by IGF1R;Negative regulation of MAPK pathway 
                                               hsa04010;hsa04012;hsa04014;hsa04015;hsa04022;hsa04024;hsa04062;hsa04068;hsa04071;hsa04072;hsa04151;hsa04270;hsa04370;hsa04510;hsa04540;hsa04550;hsa04630;hsa04650;hsa04660;hsa04662;hsa04664;hsa04666;hsa04720;hsa04722;hsa04726;hsa04730;hsa04810;hsa04910;hsa04912;hsa04914;hsa04915;hsa04916;hsa04917;hsa04919;hsa04921;hsa05034;hsa05152;hsa05160;hsa05161;hsa05164;hsa05200;hsa05205;hsa05206;hsa05210;hsa05211;hsa05212;hsa05213;hsa05214;hsa05215;hsa05218;hsa05219;hsa05220;hsa05221;hsa05223;hsa05230;hsa05231 
                                               RAF1 
                                               NA 
                                               azure4 
                                               1 
                                               -0.778846153846154 
                                             
                                             
                                               KRAS-mRNA 
                                               0.0772 
                                               0.101 
                                               -0.121 
                                               0.276 
                                               1.05 
                                               0.919 
                                               1.21 
                                               0.455 
                                               1 
                                               loglinear 
                                               ARMS-mediated activation, CD209 (DC-SIGN) signaling, Cell surface interactions at the vascular wall, Constitutive Signaling by EGFRvIII, DAP12 signaling, Downstream signal transduction, FCERI mediated MAPK activation, Frs2-mediated activation, Gastrin-CREB signalling pathway via PKC and MAPK, Interleukin receptor SHC signaling, IRS-related events triggered by IGF1R, MAP2K and MAPK activation, RAFMAP kinase cascade, SHC-mediated cascade.CL.FGFR1, SHC-related events triggered by IGF1R, Signaling by ERBB2, Signaling by FGFR1 in disease, Signaling by FGFR2 in disease, Signaling by FGFR3 in disease, Signaling by FGFR4 in disease, Signaling by PDGF, Signaling by SCF-KIT, Tie2 Signaling, VEGFA-VEGFR2 Pathway, VEGFR2 mediated cell proliferation 
                                               SEBELIUS_2_C4636 
                                               KRAS 
                                               mRNA 
                                               0 
                                               
                                               
                                               Signaling by FGFR2 in disease;Regulation of RAS by GAPs;SHC-mediated cascade.CL.FGFR3;Signaling by FGFR3 in disease;MAP2K and MAPK activation;SHC-related events triggered by IGF1R;Signaling by FGFR4 in disease;SHC-mediated cascade.CL.FGFR1;Tie2 Signaling;Signaling by FGFR1 in disease;Activation of RAS in B cells;SHC-mediated cascade.CL.FGFR4;Constitutive Signaling by EGFRvIII;CD209 (DC-SIGN) signaling;FCERI mediated MAPK activation;RAFMAP kinase cascade;Downstream signal transduction;VEGFR2 mediated cell proliferation;Signaling by SCF-KIT;Frs2-mediated activation;ARMS-mediated activation;Gastrin-CREB signalling pathway via PKC and MAPK;Interleukin receptor SHC signaling;DAP12 signaling;Signaling by PDGF;VEGFA-VEGFR2 Pathway;Signaling by ERBB2;IRS-related events triggered by IGF1R;Cell surface interactions at the vascular wall 
                                               hsa04010;hsa04012;hsa04014;hsa04015;hsa04062;hsa04068;hsa04071;hsa04072;hsa04151;hsa04211;hsa04213;hsa04320;hsa04360;hsa04370;hsa04530;hsa04540;hsa04550;hsa04650;hsa04660;hsa04662;hsa04664;hsa04720;hsa04722;hsa04725;hsa04726;hsa04730;hsa04810;hsa04910;hsa04912;hsa04914;hsa04915;hsa04916;hsa04917;hsa04919;hsa04921;hsa04933;hsa04960;hsa05034;hsa05160;hsa05161;hsa05166;hsa05200;hsa05203;hsa05205;hsa05206;hsa05210;hsa05211;hsa05212;hsa05213;hsa05214;hsa05215;hsa05216;hsa05218;hsa05219;hsa05220;hsa05221;hsa05223;hsa05230;hsa05231 
                                               KRAS 
                                               NA 
                                               azure4 
                                               1 
                                               0.764356435643564 
                                             
                                             
                                               ARAF-mRNA 
                                               -0.129 
                                               0.17 
                                               -0.462 
                                               0.205 
                                               0.915 
                                               0.726 
                                               1.15 
                                               0.459 
                                               1 
                                               loglinear 
                                               ARMS-mediated activation, DAP12 signaling, Downstream signal transduction, FCERI mediated MAPK activation, Frs2-mediated activation, Gastrin-CREB signalling pathway via PKC and MAPK, Interleukin receptor SHC signaling, IRS-related events triggered by IGF1R, MAP2K and MAPK activation, Negative regulation of MAPK pathway, RAF activation, RAFMAP kinase cascade, Signaling by ERBB2, Signaling by PDGF, Signaling by SCF-KIT, VEGFA-VEGFR2 Pathway, VEGFR2 mediated cell proliferation 
                                               SEBELIUS_2_C4636 
                                               ARAF 
                                               mRNA 
                                               0 
                                               
                                               
                                               MAP2K and MAPK activation;RAF activation;Negative regulation of MAPK pathway;RAFMAP kinase cascade;Downstream signal transduction;VEGFR2 mediated cell proliferation;Signaling by SCF-KIT;Frs2-mediated activation;ARMS-mediated activation;Gastrin-CREB signalling pathway via PKC and MAPK;Interleukin receptor SHC signaling;FCERI mediated MAPK activation;DAP12 signaling;Signaling by PDGF;VEGFA-VEGFR2 Pathway;Signaling by ERBB2;IRS-related events triggered by IGF1R 
                                               hsa04012;hsa04068;hsa04270;hsa04650;hsa04720;hsa04726;hsa04730;hsa04810;hsa04910;hsa04914;hsa05034;hsa05160;hsa05200;hsa05205;hsa05210;hsa05211;hsa05212;hsa05213;hsa05214;hsa05215;hsa05218;hsa05219;hsa05220;hsa05221;hsa05223 
                                               ARAF 
                                               NA 
                                               azure4 
                                               1 
                                               -0.758823529411765 
                                             
                                           
                                           DE Results - ARMS-mediated activation genes - New.Annotation: Control vs.cel67  Table displaying each sample's global significance scores and directed global significance scores as defined in the heatmaps above. The global significance score is calculated as the square root of the mean squared t-statistic for the genes in a gene set, with t-statistics coming from the linear regression underlying our differential expression analysis. The directed global significance score is calculated as the square root of the mean signed squared t-statistic for the genes in a gene set, with t-statistics coming from the linear regression underlying our differential expression analysis.   
                                      
                                       
                                  
                                
                                
                                  
                                       
                                      
                                           Volcano Plot: New.Annotation: miR142 vs.cel67   More Plot Information                Volcano Plot: New.Annotation: miR142 vs.cel67  Volcano plot displaying each gene's -log10(p-value) and log2 fold change for the selected covariate. Highly statistically significant genes fall at the top of the plot, and highly differentially expressed genes fall to either side. Genes within the selected gene set are highlighted in orange. Horizontal lines indicate various False Discovery Rate (FDR) thresholds.   
                                           DE Results - ARMS-mediated activation genes - New.Annotation: miR142 vs.cel67   More Plot Information      Download CSV Data     
                                           
                                             
                                             
                                           
                                           
                                             
                                               Probe Label 
                                               Log2 fold change 
                                               std error (log2) 
                                               Lower confidence limit (log2) 
                                               Upper confidence limit (log2) 
                                               Linear fold change 
                                               Lower confidence limit (linear) 
                                               Upper confidence limit (linear) 
                                               P-value 
                                               BY.p.value 
                                               method 
                                               Gene.sets 
                                               Codeset.Name 
                                               Probe.Label 
                                               Analyte.Type 
                                               Is.Control 
                                               Control.Type 
                                               Related.Probes 
                                               Probe.Annotation 
                                               KEGG.Pathways 
                                               Official.Gene.Name 
                                               Control.Conc 
                                               volcanocol 
                                               volcanopch 
                                               tstats.all 
                                             
                                           
                                           
                                             
                                               CRK-mRNA 
                                               -0.918 
                                               0.103 
                                               -1.12 
                                               -0.716 
                                               0.529 
                                               0.46 
                                               0.609 
                                               1.38e-08 
                                               2.62e-06 
                                               loglinear 
                                               ARMS-mediated activation, Downstream signal transduction, Integrin alphaIIb beta3 signaling, p130Cas linkage to MAPK signaling for integrins, Regulation of actin dynamics for phagocytic cup formation, Regulation of signaling by CBL, Signaling by PDGF, VEGFA-VEGFR2 Pathway 
                                               SEBELIUS_2_C4636 
                                               CRK 
                                               mRNA 
                                               0 
                                               
                                               
                                               ARMS-mediated activation;Downstream signal transduction;Regulation of signaling by CBL;p130Cas linkage to MAPK signaling for integrins;VEGFA-VEGFR2 Pathway;Regulation of actin dynamics for phagocytic cup formation;Signaling by PDGF;Integrin alphaIIb beta3 signaling 
                                               hsa04010;hsa04012;hsa04015;hsa04062;hsa04510;hsa04666;hsa04722;hsa04810;hsa04910;hsa05100;hsa05131;hsa05200;hsa05206;hsa05211;hsa05220 
                                               CRK 
                                               NA 
                                               slateblue1 
                                               16 
                                               -8.9126213592233 
                                             
                                             
                                               HRAS-mRNA 
                                               -0.987 
                                               0.144 
                                               -1.27 
                                               -0.705 
                                               0.504 
                                               0.415 
                                               0.613 
                                               8.72e-07 
                                               6.43e-05 
                                               loglinear 
                                               ARMS-mediated activation, CD209 (DC-SIGN) signaling, Cell surface interactions at the vascular wall, Constitutive Signaling by EGFRvIII, CREB phosphorylation through the activation of Ras, DAP12 signaling, Downstream signal transduction, FCERI mediated MAPK activation, Frs2-mediated activation, Gastrin-CREB signalling pathway via PKC and MAPK, Interleukin receptor SHC signaling, IRS-related events triggered by IGF1R, MAP2K and MAPK activation, RAFMAP kinase cascade, SHC-mediated cascade.CL.FGFR1, SHC-related events triggered by IGF1R, Signaling by ERBB2, Signaling by FGFR1 in disease, Signaling by FGFR2 in disease, Signaling by FGFR3 in disease, Signaling by FGFR4 in disease, Signaling by PDGF, Signaling by SCF-KIT, Tie2 Signaling, VEGFA-VEGFR2 Pathway, VEGFR2 mediated cell proliferation 
                                               SEBELIUS_2_C4636 
                                               HRAS 
                                               mRNA 
                                               0 
                                               
                                               
                                               Signaling by FGFR2 in disease;Regulation of RAS by GAPs;SHC-mediated cascade.CL.FGFR3;Signaling by FGFR3 in disease;MAP2K and MAPK activation;SHC-related events triggered by IGF1R;Signaling by FGFR4 in disease;SHC-mediated cascade.CL.FGFR1;Tie2 Signaling;Signaling by FGFR1 in disease;Activation of RAS in B cells;SHC-mediated cascade.CL.FGFR4;Constitutive Signaling by EGFRvIII;CD209 (DC-SIGN) signaling;Ras activation uopn Ca2.PLS. infux through NMDA receptor;FCERI mediated MAPK activation;RAFMAP kinase cascade;Downstream signal transduction;VEGFR2 mediated cell proliferation;Signaling by SCF-KIT;Frs2-mediated activation;ARMS-mediated activation;Gastrin-CREB signalling pathway via PKC and MAPK;Interleukin receptor SHC signaling;DAP12 signaling;Signaling by PDGF;VEGFA-VEGFR2 Pathway;Signaling by ERBB2;IRS-related events triggered by IGF1R;Cell surface interactions at the vascular wall;CREB phosphorylation through the activation of Ras 
                                               hsa04010;hsa04012;hsa04014;hsa04015;hsa04062;hsa04068;hsa04071;hsa04072;hsa04144;hsa04151;hsa04211;hsa04213;hsa04360;hsa04370;hsa04510;hsa04530;hsa04540;hsa04550;hsa04630;hsa04650;hsa04660;hsa04662;hsa04664;hsa04720;hsa04722;hsa04725;hsa04726;hsa04730;hsa04810;hsa04910;hsa04912;hsa04915;hsa04916;hsa04917;hsa04919;hsa04921;hsa04933;hsa05034;hsa05160;hsa05161;hsa05166;hsa05200;hsa05203;hsa05205;hsa05206;hsa05211;hsa05213;hsa05214;hsa05215;hsa05216;hsa05218;hsa05219;hsa05220;hsa05221;hsa05223;hsa05230;hsa05231 
                                               HRAS 
                                               NA 
                                               slateblue1 
                                               16 
                                               -6.85416666666667 
                                             
                                             
                                               BRAF-mRNA 
                                               0.375 
                                               0.0601 
                                               0.257 
                                               0.493 
                                               1.3 
                                               1.2 
                                               1.41 
                                               3.46e-06 
                                               0.000191 
                                               loglinear 
                                               ARMS-mediated activation, CREB phosphorylation through the activation of Ras, DAP12 signaling, Downstream signal transduction, FCERI mediated MAPK activation, Frs2-mediated activation, Gastrin-CREB signalling pathway via PKC and MAPK, Interleukin receptor SHC signaling, IRS-related events triggered by IGF1R, MAP2K and MAPK activation, Negative feedback regulation of MAPK pathway, Negative regulation of FGFR1 signaling, Negative regulation of FGFR2 signaling, Negative regulation of FGFR3 signaling, Negative regulation of FGFR4 signaling, Negative regulation of MAPK pathway, RAF activation, RAFMAP kinase cascade, Signaling by ERBB2, Signaling by PDGF, Signaling by SCF-KIT, Spry regulation of FGF signaling, VEGFA-VEGFR2 Pathway, VEGFR2 mediated cell proliferation 
                                               SEBELIUS_2_C4636 
                                               BRAF 
                                               mRNA 
                                               0 
                                               
                                               
                                               Spry regulation of FGF signaling;MAP2K and MAPK activation;RAF activation;CREB phosphorylation through the activation of Ras;Negative feedback regulation of MAPK pathway;ARMS-mediated activation;Negative regulation of FGFR3 signaling;Negative regulation of FGFR4 signaling;Negative regulation of FGFR2 signaling;Negative regulation of FGFR1 signaling;RAFMAP kinase cascade;Downstream signal transduction;VEGFR2 mediated cell proliferation;Signaling by SCF-KIT;Frs2-mediated activation;Gastrin-CREB signalling pathway via PKC and MAPK;Interleukin receptor SHC signaling;FCERI mediated MAPK activation;DAP12 signaling;Signaling by PDGF;VEGFA-VEGFR2 Pathway;Signaling by ERBB2;IRS-related events triggered by IGF1R;Negative regulation of MAPK pathway 
                                               hsa04010;hsa04012;hsa04015;hsa04024;hsa04062;hsa04068;hsa04150;hsa04270;hsa04510;hsa04650;hsa04720;hsa04722;hsa04726;hsa04730;hsa04810;hsa04910;hsa04914;hsa05034;hsa05160;hsa05200;hsa05205;hsa05210;hsa05211;hsa05212;hsa05213;hsa05214;hsa05215;hsa05216;hsa05218;hsa05219;hsa05220;hsa05221;hsa05223 
                                               BRAF 
                                               NA 
                                               slateblue1 
                                               16 
                                               6.2396006655574 
                                             
                                             
                                               EGFR-mRNA 
                                               -0.441 
                                               0.101 
                                               -0.638 
                                               -0.244 
                                               0.737 
                                               0.643 
                                               0.845 
                                               0.00026 
                                               0.00766 
                                               loglinear 
                                               ARMS-mediated activation, Constitutive Signaling by Aberrant PI3K in Cancer, Constitutive Signaling by EGFRvIII, DAP12 signaling, Downstream signal transduction, FCERI mediated MAPK activation, Frs2-mediated activation, GAB1 signalosome, Gastrin-CREB signalling pathway via PKC and MAPK, Interleukin receptor SHC signaling, IRS-related events triggered by IGF1R, PI3KAKT activation, Role of LAT2NTALLAB on calcium mobilization, Signaling by ERBB2, Signaling by PDGF, Signaling by SCF-KIT, VEGFA-VEGFR2 Pathway, VEGFR2 mediated cell proliferation 
                                               SEBELIUS_2_C4636 
                                               EGFR 
                                               mRNA 
                                               0 
                                               
                                               
                                               Constitutive Signaling by Aberrant PI3K in Cancer;Inhibition of Signaling by Overexpressed EGFR;PLCG1 events in ERBB2 signaling;EGFR interacts with phospholipase C-gamma;EGFR Transactivation by Gastrin;Constitutive Signaling by Ligand-Responsive EGFR Cancer Variants;Constitutive Signaling by EGFRvIII;GAB1 signalosome;Downstream signal transduction;Signaling by SCF-KIT;PI3KAKT activation;Role of LAT2NTALLAB on calcium mobilization;DAP12 signaling;Signaling by PDGF;Signaling by ERBB2;VEGFR2 mediated cell proliferation;Frs2-mediated activation;ARMS-mediated activation;Gastrin-CREB signalling pathway via PKC and MAPK;Interleukin receptor SHC signaling;FCERI mediated MAPK activation;VEGFA-VEGFR2 Pathway;IRS-related events triggered by IGF1R 
                                               hsa04010;hsa04012;hsa04014;hsa04015;hsa04020;hsa04060;hsa04066;hsa04068;hsa04072;hsa04144;hsa04151;hsa04320;hsa04510;hsa04520;hsa04540;hsa04810;hsa04912;hsa04915;hsa04921;hsa05120;hsa05160;hsa05200;hsa05205;hsa05206;hsa05212;hsa05213;hsa05214;hsa05215;hsa05218;hsa05219;hsa05223;hsa05230;hsa05231 
                                               EGFR 
                                               NA 
                                               slateblue1 
                                               16 
                                               -4.36633663366337 
                                             
                                             
                                               PPP2CA-mRNA 
                                               -0.205 
                                               0.0527 
                                               -0.309 
                                               -0.102 
                                               0.867 
                                               0.807 
                                               0.932 
                                               0.000827 
                                               0.0186 
                                               loglinear 
                                               ARMS-mediated activation, AXIN missense mutants destabilize the destruction complex, Beta-catenin phosphorylation cascade, CTLA4 inhibitory signaling, Cyclin AB1 associated events during G2M transition, Cyclin D associated events in G1, DAP12 signaling, DARPP-32 events, Disassembly of the destruction complex and recruitment of AXIN to the membrane, Downstream signal transduction, ERKs are inactivated, FCERI mediated MAPK activation, Frs2-mediated activation, Gastrin-CREB signalling pathway via PKC and MAPK, Interleukin receptor SHC signaling, IRS-related events triggered by IGF1R, MASTL Facilitates Mitotic Progression, Negative regulation of FGFR1 signaling, Negative regulation of FGFR2 signaling, Negative regulation of FGFR3 signaling, Negative regulation of FGFR4 signaling, Negative regulation of MAPK pathway, Nonsense Mediated Decay (NMD) enhanced by the Exon Junction Complex (EJC), RAF activation, RAFMAP kinase cascade, Resolution of Sister Chromatid Cohesion, RHO GTPases Activate Formins, S33 mutants of beta-catenin aren.SQT.t phosphorylated, S37 mutants of beta-catenin aren.SQT.t phosphorylated, S45 mutants of beta-catenin aren.SQT.t phosphorylated, Signaling by ERBB2, Signaling by PDGF, Signaling by SCF-KIT, Spry regulation of FGF signaling, T41 mutants of beta-catenin aren.SQT.t phosphorylated, Truncations of AMER1 destabilize the destruction complex, VEGFA-VEGFR2 Pathway, VEGFR2 mediated cell proliferation 
                                               SEBELIUS_2_C4636 
                                               PPP2CA 
                                               mRNA 
                                               0 
                                               
                                               
                                               MASTL Facilitates Mitotic Progression;T41 mutants of beta-catenin aren.SQT.t phosphorylated;ERKs are inactivated;Glycolysis;Negative regulation of MAPK pathway;Disassembly of the destruction complex and recruitment of AXIN to the membrane;S37 mutants of beta-catenin aren.SQT.t phosphorylated;Cyclin AB1 associated events during G2M transition;RAF activation;RHO GTPases Activate Formins;S33 mutants of beta-catenin aren.SQT.t phosphorylated;DARPP-32 events;Beta-catenin phosphorylation cascade;Initiation of Nuclear Envelope Reformation;Spry regulation of FGF signaling;Truncations of AMER1 destabilize the destruction complex;CTLA4 inhibitory signaling;AXIN missense mutants destabilize the destruction complex;S45 mutants of beta-catenin aren.SQT.t phosphorylated;Nonsense Mediated Decay (NMD) enhanced by the Exon Junction Complex (EJC);Cyclin D associated events in G1;Platelet sensitization by LDL;Resolution of Sister Chromatid Cohesion;Inhibition of replication initiation of damaged DNA by RB1E2F1;RAFMAP kinase cascade;Downstream signal transduction;VEGFR2 mediated cell proliferation;Signaling by SCF-KIT;Frs2-mediated activation;ARMS-mediated activation;Gastrin-CREB signalling pathway via PKC and MAPK;Interleukin receptor SHC signaling;FCERI mediated MAPK activation;DAP12 signaling;Signaling by PDGF;VEGFA-VEGFR2 Pathway;Signaling by ERBB2;IRS-related events triggered by IGF1R;Negative regulation of FGFR3 signaling;Negative regulation of FGFR4 signaling;Negative regulation of FGFR2 signaling;Negative regulation of FGFR1 signaling 
                                               hsa03015;hsa04071;hsa04114;hsa04151;hsa04152;hsa04261;hsa04350;hsa04390;hsa04530;hsa04728;hsa04730;hsa05142;hsa05160 
                                               PPP2CA 
                                               NA 
                                               slateblue2 
                                               16 
                                               -3.8899430740038 
                                             
                                             
                                               MAP2K1-mRNA 
                                               -0.444 
                                               0.12 
                                               -0.679 
                                               -0.21 
                                               0.735 
                                               0.625 
                                               0.865 
                                               0.00128 
                                               0.027 
                                               lm.nb 
                                               ARMS-mediated activation, DAP12 signaling, Downstream signal transduction, FCERI mediated MAPK activation, Frs2-mediated activation, Gastrin-CREB signalling pathway via PKC and MAPK, Interleukin receptor SHC signaling, IRS-related events triggered by IGF1R, Negative feedback regulation of MAPK pathway, Negative regulation of MAPK pathway, RAF activation, RAFMAP kinase cascade, Signaling by ERBB2, Signaling by PDGF, Signaling by SCF-KIT, VEGFA-VEGFR2 Pathway, VEGFR2 mediated cell proliferation 
                                               SEBELIUS_2_C4636 
                                               MAP2K1 
                                               mRNA 
                                               0 
                                               
                                               
                                               RAF activation;Uptake and function of anthrax toxins;Negative feedback regulation of MAPK pathway;Interleukin-1 signaling;RAFMAP kinase cascade;Downstream signal transduction;VEGFR2 mediated cell proliferation;Signaling by SCF-KIT;Frs2-mediated activation;ARMS-mediated activation;Gastrin-CREB signalling pathway via PKC and MAPK;Interleukin receptor SHC signaling;FCERI mediated MAPK activation;DAP12 signaling;Signaling by PDGF;VEGFA-VEGFR2 Pathway;Signaling by ERBB2;IRS-related events triggered by IGF1R;Negative regulation of MAPK pathway 
                                               hsa04010;hsa04012;hsa04014;hsa04015;hsa04022;hsa04024;hsa04062;hsa04066;hsa04068;hsa04071;hsa04072;hsa04114;hsa04151;hsa04270;hsa04320;hsa04370;hsa04380;hsa04510;hsa04540;hsa04550;hsa04620;hsa04650;hsa04660;hsa04662;hsa04664;hsa04666;hsa04668;hsa04720;hsa04722;hsa04725;hsa04726;hsa04730;hsa04810;hsa04910;hsa04912;hsa04914;hsa04915;hsa04916;hsa04917;hsa04919;hsa04921;hsa05020;hsa05034;hsa05161;hsa05164;hsa05200;hsa05205;hsa05206;hsa05210;hsa05211;hsa05212;hsa05213;hsa05214;hsa05215;hsa05216;hsa05218;hsa05219;hsa05220;hsa05221;hsa05223;hsa05230;hsa05231 
                                               MAP2K1 
                                               NA 
                                               slateblue2 
                                               16 
                                               -3.7 
                                             
                                             
                                               PPP2R1A-mRNA 
                                               -0.339 
                                               0.0993 
                                               -0.533 
                                               -0.144 
                                               0.791 
                                               0.691 
                                               0.905 
                                               0.00262 
                                               0.0492 
                                               loglinear 
                                               ARMS-mediated activation, AXIN missense mutants destabilize the destruction complex, Beta-catenin phosphorylation cascade, CTLA4 inhibitory signaling, Cyclin AB1 associated events during G2M transition, Cyclin D associated events in G1, DAP12 signaling, DARPP-32 events, Disassembly of the destruction complex and recruitment of AXIN to the membrane, Downstream signal transduction, FCERI mediated MAPK activation, Frs2-mediated activation, Gastrin-CREB signalling pathway via PKC and MAPK, Interleukin receptor SHC signaling, IRS-related events triggered by IGF1R, MASTL Facilitates Mitotic Progression, Negative regulation of FGFR1 signaling, Negative regulation of FGFR2 signaling, Negative regulation of FGFR3 signaling, Negative regulation of FGFR4 signaling, Nonsense Mediated Decay (NMD) enhanced by the Exon Junction Complex (EJC), RAF activation, RAFMAP kinase cascade, Resolution of Sister Chromatid Cohesion, RHO GTPases Activate Formins, S33 mutants of beta-catenin aren.SQT.t phosphorylated, S37 mutants of beta-catenin aren.SQT.t phosphorylated, S45 mutants of beta-catenin aren.SQT.t phosphorylated, Signaling by ERBB2, Signaling by PDGF, Signaling by SCF-KIT, Spry regulation of FGF signaling, Truncations of AMER1 destabilize the destruction complex, VEGFA-VEGFR2 Pathway, VEGFR2 mediated cell proliferation 
                                               SEBELIUS_2_C4636 
                                               PPP2R1A 
                                               mRNA 
                                               0 
                                               
                                               
                                               MASTL Facilitates Mitotic Progression;Regulation of PLK1 Activity at G2M Transition;Disassembly of the destruction complex and recruitment of AXIN to the membrane;S37 mutants of beta-catenin aren.SQT.t phosphorylated;Cyclin AB1 associated events during G2M transition;RAF activation;RHO GTPases Activate Formins;S33 mutants of beta-catenin aren.SQT.t phosphorylated;Loss of Nlp from mitotic centrosomes;DARPP-32 events;Beta-catenin phosphorylation cascade;Initiation of Nuclear Envelope Reformation;Recruitment of mitotic centrosome proteins and complexes;Spry regulation of FGF signaling;Truncations of AMER1 destabilize the destruction complex;CTLA4 inhibitory signaling;AXIN missense mutants destabilize the destruction complex;S45 mutants of beta-catenin aren.SQT.t phosphorylated;Anchoring of the basal body to the plasma membrane;Nonsense Mediated Decay (NMD) enhanced by the Exon Junction Complex (EJC);Cyclin D associated events in G1;Platelet sensitization by LDL;Resolution of Sister Chromatid Cohesion;Inhibition of replication initiation of damaged DNA by RB1E2F1;RAFMAP kinase cascade;Downstream signal transduction;VEGFR2 mediated cell proliferation;Signaling by SCF-KIT;Frs2-mediated activation;ARMS-mediated activation;Gastrin-CREB signalling pathway via PKC and MAPK;Interleukin receptor SHC signaling;FCERI mediated MAPK activation;DAP12 signaling;Signaling by PDGF;VEGFA-VEGFR2 Pathway;Signaling by ERBB2;IRS-related events triggered by IGF1R;Negative regulation of FGFR3 signaling;Negative regulation of FGFR4 signaling;Negative regulation of FGFR2 signaling;Negative regulation of FGFR1 signaling 
                                               hsa03015;hsa04071;hsa04114;hsa04151;hsa04152;hsa04261;hsa04350;hsa04390;hsa04530;hsa04728;hsa04730;hsa05142;hsa05160 
                                               PPP2R1A 
                                               NA 
                                               slateblue2 
                                               16 
                                               -3.41389728096677 
                                             
                                             
                                               SPTAN1-mRNA 
                                               0.38 
                                               0.111 
                                               0.162 
                                               0.598 
                                               1.3 
                                               1.12 
                                               1.51 
                                               0.00263 
                                               0.0492 
                                               loglinear 
                                               Apoptotic cleavage of cellular proteins, ARMS-mediated activation, DAP12 signaling, Downstream signal transduction, FCERI mediated MAPK activation, Frs2-mediated activation, Gastrin-CREB signalling pathway via PKC and MAPK, Interleukin receptor SHC signaling, IRS-related events triggered by IGF1R, Nephrin interactions, Signaling by ERBB2, Signaling by PDGF, Signaling by SCF-KIT, VEGFA-VEGFR2 Pathway, VEGFR2 mediated cell proliferation 
                                               SEBELIUS_2_C4636 
                                               SPTAN1 
                                               mRNA 
                                               0 
                                               
                                               
                                               Nephrin interactions;Caspase-mediated cleavage of cytoskeletal proteins;Downstream signal transduction;VEGFR2 mediated cell proliferation;Signaling by SCF-KIT;Frs2-mediated activation;ARMS-mediated activation;Gastrin-CREB signalling pathway via PKC and MAPK;Interleukin receptor SHC signaling;FCERI mediated MAPK activation;DAP12 signaling;Signaling by PDGF;VEGFA-VEGFR2 Pathway;Signaling by ERBB2;IRS-related events triggered by IGF1R;Apoptotic cleavage of cellular proteins 
                                               hsa04530 
                                               SPTAN1 
                                               NA 
                                               slateblue3 
                                               16 
                                               3.42342342342342 
                                             
                                             
                                               NRAS-mRNA 
                                               -0.363 
                                               0.113 
                                               -0.585 
                                               -0.141 
                                               0.777 
                                               0.666 
                                               0.907 
                                               0.00423 
                                               0.073 
                                               lm.nb 
                                               ARMS-mediated activation, CD209 (DC-SIGN) signaling, Cell surface interactions at the vascular wall, Constitutive Signaling by EGFRvIII, DAP12 signaling, Downstream signal transduction, FCERI mediated MAPK activation, Frs2-mediated activation, Gastrin-CREB signalling pathway via PKC and MAPK, Interleukin receptor SHC signaling, IRS-related events triggered by IGF1R, MAP2K and MAPK activation, RAFMAP kinase cascade, SHC-mediated cascade.CL.FGFR1, SHC-related events triggered by IGF1R, Signaling by ERBB2, Signaling by FGFR1 in disease, Signaling by FGFR2 in disease, Signaling by FGFR3 in disease, Signaling by FGFR4 in disease, Signaling by PDGF, Signaling by SCF-KIT, Tie2 Signaling, VEGFA-VEGFR2 Pathway, VEGFR2 mediated cell proliferation 
                                               SEBELIUS_2_C4636 
                                               NRAS 
                                               mRNA 
                                               0 
                                               
                                               
                                               Signaling by FGFR2 in disease;Regulation of RAS by GAPs;SHC-mediated cascade.CL.FGFR3;Signaling by FGFR3 in disease;MAP2K and MAPK activation;SHC-related events triggered by IGF1R;Signaling by FGFR4 in disease;SHC-mediated cascade.CL.FGFR1;Tie2 Signaling;Signaling by FGFR1 in disease;Activation of RAS in B cells;SHC-mediated cascade.CL.FGFR4;Constitutive Signaling by EGFRvIII;CD209 (DC-SIGN) signaling;FCERI mediated MAPK activation;RAFMAP kinase cascade;Downstream signal transduction;VEGFR2 mediated cell proliferation;Signaling by SCF-KIT;Frs2-mediated activation;ARMS-mediated activation;Gastrin-CREB signalling pathway via PKC and MAPK;Interleukin receptor SHC signaling;DAP12 signaling;Signaling by PDGF;VEGFA-VEGFR2 Pathway;Signaling by ERBB2;IRS-related events triggered by IGF1R;Cell surface interactions at the vascular wall 
                                               hsa04010;hsa04012;hsa04014;hsa04015;hsa04062;hsa04068;hsa04071;hsa04072;hsa04151;hsa04211;hsa04213;hsa04360;hsa04370;hsa04530;hsa04540;hsa04550;hsa04650;hsa04660;hsa04662;hsa04664;hsa04720;hsa04722;hsa04725;hsa04726;hsa04730;hsa04810;hsa04910;hsa04912;hsa04915;hsa04916;hsa04917;hsa04919;hsa04921;hsa04933;hsa05034;hsa05160;hsa05161;hsa05166;hsa05200;hsa05203;hsa05205;hsa05206;hsa05211;hsa05213;hsa05214;hsa05215;hsa05216;hsa05218;hsa05219;hsa05220;hsa05221;hsa05223;hsa05230;hsa05231 
                                               NRAS 
                                               NA 
                                               slateblue3 
                                               16 
                                               -3.21238938053097 
                                             
                                             
                                               PDGFRB-mRNA 
                                               0.484 
                                               0.159 
                                               0.172 
                                               0.795 
                                               1.4 
                                               1.13 
                                               1.73 
                                               0.00613 
                                               0.0981 
                                               lm.nb 
                                               ARMS-mediated activation, Constitutive Signaling by Aberrant PI3K in Cancer, DAP12 signaling, Downstream signal transduction, FCERI mediated MAPK activation, Frs2-mediated activation, GAB1 signalosome, Gastrin-CREB signalling pathway via PKC and MAPK, Interleukin receptor SHC signaling, IRS-related events triggered by IGF1R, PI3KAKT activation, PIP3 activates AKT signaling, RAFMAP kinase cascade, Role of LAT2NTALLAB on calcium mobilization, Signaling by ERBB2, Signaling by PDGF, Signaling by SCF-KIT, VEGFA-VEGFR2 Pathway, VEGFR2 mediated cell proliferation 
                                               SEBELIUS_2_C4636 
                                               PDGFRB 
                                               mRNA 
                                               0 
                                               
                                               
                                               PIP3 activates AKT signaling;Constitutive Signaling by Aberrant PI3K in Cancer;RAFMAP kinase cascade;Signaling by PDGF;Downstream signal transduction;GAB1 signalosome;Signaling by SCF-KIT;PI3KAKT activation;Role of LAT2NTALLAB on calcium mobilization;DAP12 signaling;Signaling by ERBB2;VEGFR2 mediated cell proliferation;Frs2-mediated activation;ARMS-mediated activation;Gastrin-CREB signalling pathway via PKC and MAPK;Interleukin receptor SHC signaling;FCERI mediated MAPK activation;VEGFA-VEGFR2 Pathway;IRS-related events triggered by IGF1R 
                                               hsa04010;hsa04014;hsa04015;hsa04020;hsa04060;hsa04072;hsa04151;hsa04510;hsa04540;hsa04810;hsa05166;hsa05200;hsa05206;hsa05214;hsa05215;hsa05218;hsa05230;hsa05231 
                                               PDGFRB 
                                               NA 
                                               slateblue4 
                                               16 
                                               3.0440251572327 
                                             
                                             
                                               KRAS-mRNA 
                                               0.276 
                                               0.101 
                                               0.0773 
                                               0.474 
                                               1.21 
                                               1.06 
                                               1.39 
                                               0.0127 
                                               0.182 
                                               loglinear 
                                               ARMS-mediated activation, CD209 (DC-SIGN) signaling, Cell surface interactions at the vascular wall, Constitutive Signaling by EGFRvIII, DAP12 signaling, Downstream signal transduction, FCERI mediated MAPK activation, Frs2-mediated activation, Gastrin-CREB signalling pathway via PKC and MAPK, Interleukin receptor SHC signaling, IRS-related events triggered by IGF1R, MAP2K and MAPK activation, RAFMAP kinase cascade, SHC-mediated cascade.CL.FGFR1, SHC-related events triggered by IGF1R, Signaling by ERBB2, Signaling by FGFR1 in disease, Signaling by FGFR2 in disease, Signaling by FGFR3 in disease, Signaling by FGFR4 in disease, Signaling by PDGF, Signaling by SCF-KIT, Tie2 Signaling, VEGFA-VEGFR2 Pathway, VEGFR2 mediated cell proliferation 
                                               SEBELIUS_2_C4636 
                                               KRAS 
                                               mRNA 
                                               0 
                                               
                                               
                                               Signaling by FGFR2 in disease;Regulation of RAS by GAPs;SHC-mediated cascade.CL.FGFR3;Signaling by FGFR3 in disease;MAP2K and MAPK activation;SHC-related events triggered by IGF1R;Signaling by FGFR4 in disease;SHC-mediated cascade.CL.FGFR1;Tie2 Signaling;Signaling by FGFR1 in disease;Activation of RAS in B cells;SHC-mediated cascade.CL.FGFR4;Constitutive Signaling by EGFRvIII;CD209 (DC-SIGN) signaling;FCERI mediated MAPK activation;RAFMAP kinase cascade;Downstream signal transduction;VEGFR2 mediated cell proliferation;Signaling by SCF-KIT;Frs2-mediated activation;ARMS-mediated activation;Gastrin-CREB signalling pathway via PKC and MAPK;Interleukin receptor SHC signaling;DAP12 signaling;Signaling by PDGF;VEGFA-VEGFR2 Pathway;Signaling by ERBB2;IRS-related events triggered by IGF1R;Cell surface interactions at the vascular wall 
                                               hsa04010;hsa04012;hsa04014;hsa04015;hsa04062;hsa04068;hsa04071;hsa04072;hsa04151;hsa04211;hsa04213;hsa04320;hsa04360;hsa04370;hsa04530;hsa04540;hsa04550;hsa04650;hsa04660;hsa04662;hsa04664;hsa04720;hsa04722;hsa04725;hsa04726;hsa04730;hsa04810;hsa04910;hsa04912;hsa04914;hsa04915;hsa04916;hsa04917;hsa04919;hsa04921;hsa04933;hsa04960;hsa05034;hsa05160;hsa05161;hsa05166;hsa05200;hsa05203;hsa05205;hsa05206;hsa05210;hsa05211;hsa05212;hsa05213;hsa05214;hsa05215;hsa05216;hsa05218;hsa05219;hsa05220;hsa05221;hsa05223;hsa05230;hsa05231 
                                               KRAS 
                                               NA 
                                               slateblue4 
                                               16 
                                               2.73267326732673 
                                             
                                             
                                               FYN-mRNA 
                                               0.275 
                                               0.11 
                                               0.0594 
                                               0.49 
                                               1.21 
                                               1.04 
                                               1.4 
                                               0.0207 
                                               0.272 
                                               loglinear 
                                               ARMS-mediated activation, CD209 (DC-SIGN) signaling, CD28 co-stimulation, CD28 dependent PI3KAkt signaling, CD28 dependent Vav1 pathway, Cell surface interactions at the vascular wall, Constitutive Signaling by Aberrant PI3K in Cancer, CTLA4 inhibitory signaling, DAP12 signaling, Downstream signal transduction, EPH-ephrin mediated repulsion of cells, FCERI mediated MAPK activation, Frs2-mediated activation, GAB1 signalosome, Gastrin-CREB signalling pathway via PKC and MAPK, GPVI-mediated activation cascade, Interleukin receptor SHC signaling, IRS-related events triggered by IGF1R, Nephrin interactions, PI3KAKT activation, RAFMAP kinase cascade, Regulation of signaling by CBL, Role of LAT2NTALLAB on calcium mobilization, Sema3A PAK dependent Axon repulsion, Signaling by ERBB2, Signaling by PDGF, Signaling by SCF-KIT, VEGFA-VEGFR2 Pathway, VEGFR2 mediated cell proliferation 
                                               SEBELIUS_2_C4636 
                                               FYN 
                                               mRNA 
                                               0 
                                               
                                               
                                               EPH-ephrin mediated repulsion of cells;Regulation of signaling by CBL;FCGR activation;Antigen activates B Cell Receptor (BCR) leading to generation of second messengers;Dectin-2 family;RAFMAP kinase cascade;Constitutive Signaling by Aberrant PI3K in Cancer;CTLA4 inhibitory signaling;CD28 dependent PI3KAkt signaling;PECAM1 interactions;CD28 dependent Vav1 pathway;GPVI-mediated activation cascade;Sema3A PAK dependent Axon repulsion;Nephrin interactions;SEMA3A-Plexin repulsion signaling by inhibiting Integrin adhesion;CRMPs in Sema3A signaling;CD209 (DC-SIGN) signaling;Downstream signal transduction;GAB1 signalosome;Signaling by SCF-KIT;PI3KAKT activation;Role of LAT2NTALLAB on calcium mobilization;DAP12 signaling;Signaling by PDGF;Signaling by ERBB2;VEGFA-VEGFR2 Pathway;VEGFR2 mediated cell proliferation;Frs2-mediated activation;ARMS-mediated activation;Gastrin-CREB signalling pathway via PKC and MAPK;Interleukin receptor SHC signaling;FCERI mediated MAPK activation;IRS-related events triggered by IGF1R;CD28 co-stimulation;Cell surface interactions at the vascular wall 
                                               hsa04071;hsa04072;hsa04360;hsa04380;hsa04510;hsa04520;hsa04611;hsa04650;hsa04660;hsa04664;hsa04725;hsa05020;hsa05130;hsa05162;hsa05416 
                                               FYN 
                                               NA 
                                               slateblue4 
                                               16 
                                               2.5 
                                             
                                             
                                               MAPK3-mRNA 
                                               0.13 
                                               0.0541 
                                               0.0237 
                                               0.236 
                                               1.09 
                                               1.02 
                                               1.18 
                                               0.0259 
                                               0.33 
                                               loglinear 
                                               ARMS-mediated activation, DAP12 signaling, Downstream signal transduction, ERKs are inactivated, FCERI mediated MAPK activation, Frs2-mediated activation, Gastrin-CREB signalling pathway via PKC and MAPK, Interleukin receptor SHC signaling, IRS-related events triggered by IGF1R, Negative feedback regulation of MAPK pathway, Negative regulation of FGFR3 signaling, Negative regulation of FGFR4 signaling, Negative regulation of MAPK pathway, RAFMAP kinase cascade, Regulation of HSF1-mediated heat shock response, RHO GTPases Activate WASPs and WAVEs, Signaling by ERBB2, Signaling by PDGF, Signaling by SCF-KIT, VEGFA-VEGFR2 Pathway, VEGFR2 mediated cell proliferation 
                                               SEBELIUS_2_C4636 
                                               MAPK3 
                                               mRNA 
                                               0 
                                               
                                               
                                               ERKs are inactivated;ISG15 antiviral mechanism;RHO GTPases Activate WASPs and WAVEs;Senescence-Associated Secretory Phenotype (SASP);Regulation of HSF1-mediated heat shock response;Thrombin signalling through proteinase activated receptors (PARs);Negative regulation of FGFR4 signaling;Oxidative Stress Induced Senescence;Signal attenuation;RNA Polymerase I Promoter Opening;Negative regulation of FGFR3 signaling;Negative feedback regulation of MAPK pathway;Growth hormone receptor signaling;Golgi Cisternae Pericentriolar Stack Reorganization;FCERI mediated MAPK activation;Negative regulation of MAPK pathway;RAFMAP kinase cascade;Downstream signal transduction;VEGFR2 mediated cell proliferation;Signaling by SCF-KIT;Frs2-mediated activation;ARMS-mediated activation;Gastrin-CREB signalling pathway via PKC and MAPK;Interleukin receptor SHC signaling;DAP12 signaling;Signaling by PDGF;VEGFA-VEGFR2 Pathway;Signaling by ERBB2;IRS-related events triggered by IGF1R 
                                               hsa04010;hsa04012;hsa04014;hsa04015;hsa04022;hsa04024;hsa04062;hsa04066;hsa04068;hsa04071;hsa04072;hsa04114;hsa04150;hsa04151;hsa04261;hsa04270;hsa04320;hsa04350;hsa04360;hsa04370;hsa04380;hsa04510;hsa04520;hsa04540;hsa04550;hsa04611;hsa04620;hsa04621;hsa04650;hsa04660;hsa04662;hsa04664;hsa04666;hsa04668;hsa04713;hsa04720;hsa04722;hsa04723;hsa04724;hsa04725;hsa04726;hsa04730;hsa04810;hsa04910;hsa04912;hsa04914;hsa04915;hsa04916;hsa04917;hsa04919;hsa04921;hsa04930;hsa04933;hsa04960;hsa05010;hsa05020;hsa05034;hsa05131;hsa05132;hsa05133;hsa05140;hsa05142;hsa05145;hsa05152;hsa05160;hsa05161;hsa05164;hsa05200;hsa05203;hsa05205;hsa05210;hsa05211;hsa05212;hsa05213;hsa05214;hsa05215;hsa05216;hsa05218;hsa05219;hsa05220;hsa05221;hsa05223;hsa05230;hsa05231 
                                               MAPK3 
                                               NA 
                                               slateblue4 
                                               16 
                                               2.40295748613678 
                                             
                                             
                                               RAF1-mRNA 
                                               -0.129 
                                               0.0728 
                                               -0.272 
                                               0.0133 
                                               0.914 
                                               0.828 
                                               1.01 
                                               0.0899 
                                               0.925 
                                               loglinear 
                                               ARMS-mediated activation, CD209 (DC-SIGN) signaling, CREB phosphorylation through the activation of Ras, DAP12 signaling, Downstream signal transduction, FCERI mediated MAPK activation, Frs2-mediated activation, Gastrin-CREB signalling pathway via PKC and MAPK, Interleukin receptor SHC signaling, IRS-related events triggered by IGF1R, Negative feedback regulation of MAPK pathway, Negative regulation of MAPK pathway, RAF activation, RAFMAP kinase cascade, Signaling by ERBB2, Signaling by PDGF, Signaling by SCF-KIT, VEGFA-VEGFR2 Pathway, VEGFR2 mediated cell proliferation 
                                               SEBELIUS_2_C4636 
                                               RAF1 
                                               mRNA 
                                               0 
                                               
                                               
                                               RAF activation;Stimuli-sensing channels;CREB phosphorylation through the activation of Ras;Negative feedback regulation of MAPK pathway;CD209 (DC-SIGN) signaling;RAFMAP kinase cascade;Downstream signal transduction;VEGFR2 mediated cell proliferation;Signaling by SCF-KIT;Frs2-mediated activation;ARMS-mediated activation;Gastrin-CREB signalling pathway via PKC and MAPK;Interleukin receptor SHC signaling;FCERI mediated MAPK activation;DAP12 signaling;Signaling by PDGF;VEGFA-VEGFR2 Pathway;Signaling by ERBB2;IRS-related events triggered by IGF1R;Negative regulation of MAPK pathway 
                                               hsa04010;hsa04012;hsa04014;hsa04015;hsa04022;hsa04024;hsa04062;hsa04068;hsa04071;hsa04072;hsa04151;hsa04270;hsa04370;hsa04510;hsa04540;hsa04550;hsa04630;hsa04650;hsa04660;hsa04662;hsa04664;hsa04666;hsa04720;hsa04722;hsa04726;hsa04730;hsa04810;hsa04910;hsa04912;hsa04914;hsa04915;hsa04916;hsa04917;hsa04919;hsa04921;hsa05034;hsa05152;hsa05160;hsa05161;hsa05164;hsa05200;hsa05205;hsa05206;hsa05210;hsa05211;hsa05212;hsa05213;hsa05214;hsa05215;hsa05218;hsa05219;hsa05220;hsa05221;hsa05223;hsa05230;hsa05231 
                                               RAF1 
                                               NA 
                                               azure4 
                                               1 
                                               -1.77197802197802 
                                             
                                             
                                               IRS1-mRNA 
                                               0.181 
                                               0.108 
                                               -0.0307 
                                               0.393 
                                               1.13 
                                               0.979 
                                               1.31 
                                               0.109 
                                               1 
                                               loglinear 
                                               ARMS-mediated activation, Constitutive Signaling by Aberrant PI3K in Cancer, DAP12 signaling, Downstream signal transduction, FCERI mediated MAPK activation, Frs2-mediated activation, GAB1 signalosome, Gastrin-CREB signalling pathway via PKC and MAPK, Interleukin receptor SHC signaling, IRS-related events triggered by IGF1R, PI3K Cascade, PI3KAKT activation, RAFMAP kinase cascade, Role of LAT2NTALLAB on calcium mobilization, Signaling by ERBB2, Signaling by PDGF, Signaling by SCF-KIT, VEGFA-VEGFR2 Pathway, VEGFR2 mediated cell proliferation 
                                               SEBELIUS_2_C4636 
                                               IRS1 
                                               mRNA 
                                               0 
                                               
                                               
                                               Constitutive Signaling by Aberrant PI3K in Cancer;RAFMAP kinase cascade;PI3K Cascade;Growth hormone receptor signaling;IRS-related events triggered by IGF1R;PI3KAKT activation;Downstream signal transduction;GAB1 signalosome;Signaling by SCF-KIT;Role of LAT2NTALLAB on calcium mobilization;DAP12 signaling;Signaling by PDGF;Signaling by ERBB2;VEGFR2 mediated cell proliferation;Frs2-mediated activation;ARMS-mediated activation;Gastrin-CREB signalling pathway via PKC and MAPK;Interleukin receptor SHC signaling;FCERI mediated MAPK activation;VEGFA-VEGFR2 Pathway 
                                               hsa04022;hsa04068;hsa04150;hsa04151;hsa04152;hsa04211;hsa04213;hsa04722;hsa04910;hsa04920;hsa04923;hsa04930;hsa04931;hsa04932;hsa04960;hsa05206 
                                               IRS1 
                                               NA 
                                               azure4 
                                               1 
                                               1.67592592592593 
                                             
                                             
                                               PTK2-mRNA 
                                               -0.115 
                                               0.0697 
                                               -0.252 
                                               0.0215 
                                               0.923 
                                               0.84 
                                               1.01 
                                               0.113 
                                               1 
                                               loglinear 
                                               Apoptotic cleavage of cellular proteins, ARMS-mediated activation, DAP12 signaling, DCC mediated attractive signaling, Downstream signal transduction, EPHA-mediated growth cone collapse, EPHB-mediated forward signaling, FCERI mediated MAPK activation, Frs2-mediated activation, Gastrin-CREB signalling pathway via PKC and MAPK, Integrin alphaIIb beta3 signaling, Interleukin receptor SHC signaling, IRS-related events triggered by IGF1R, p130Cas linkage to MAPK signaling for integrins, RAFMAP kinase cascade, Regulation of actin dynamics for phagocytic cup formation, RHO GTPases Activate WASPs and WAVEs, Signaling by ERBB2, Signaling by PDGF, Signaling by SCF-KIT, VEGFA-VEGFR2 Pathway, VEGFR2 mediated cell proliferation 
                                               SEBELIUS_2_C4636 
                                               PTK2 
                                               mRNA 
                                               0 
                                               
                                               
                                               Apoptotic cleavage of cellular proteins;RAFMAP kinase cascade;RHO GTPases Activate WASPs and WAVEs;DCC mediated attractive signaling;p130Cas linkage to MAPK signaling for integrins;EPHB-mediated forward signaling;EPHA-mediated growth cone collapse;Regulation of actin dynamics for phagocytic cup formation;Downstream signal transduction;VEGFR2 mediated cell proliferation;Signaling by SCF-KIT;Frs2-mediated activation;ARMS-mediated activation;Gastrin-CREB signalling pathway via PKC and MAPK;Interleukin receptor SHC signaling;FCERI mediated MAPK activation;DAP12 signaling;Signaling by PDGF;VEGFA-VEGFR2 Pathway;Signaling by ERBB2;IRS-related events triggered by IGF1R;Integrin alphaIIb beta3 signaling 
                                               hsa04012;hsa04062;hsa04151;hsa04360;hsa04370;hsa04510;hsa04670;hsa04810;hsa05100;hsa05146;hsa05200;hsa05202;hsa05205;hsa05222 
                                               PTK2 
                                               NA 
                                               azure4 
                                               1 
                                               -1.64992826398852 
                                             
                                             
                                               ERBB2-mRNA 
                                               0.267 
                                               0.163 
                                               -0.0519 
                                               0.586 
                                               1.2 
                                               0.965 
                                               1.5 
                                               0.116 
                                               1 
                                               lm.nb 
                                               ARMS-mediated activation, DAP12 signaling, Downstream signal transduction, FCERI mediated MAPK activation, Frs2-mediated activation, GAB1 signalosome, Gastrin-CREB signalling pathway via PKC and MAPK, Interleukin receptor SHC signaling, IRS-related events triggered by IGF1R, PI3KAKT activation, RAFMAP kinase cascade, Role of LAT2NTALLAB on calcium mobilization, Sema4D induced cell migration and growth-cone collapse, Signaling by ERBB2, Signaling by PDGF, Signaling by SCF-KIT, VEGFA-VEGFR2 Pathway, VEGFR2 mediated cell proliferation 
                                               SEBELIUS_2_C4636 
                                               ERBB2 
                                               mRNA 
                                               0 
                                               
                                               
                                               RAFMAP kinase cascade;GRB7 events in ERBB2 signaling;PLCG1 events in ERBB2 signaling;Sema4D induced cell migration and growth-cone collapse;Signaling by ERBB2;Downstream signal transduction;GAB1 signalosome;Signaling by SCF-KIT;PI3KAKT activation;Role of LAT2NTALLAB on calcium mobilization;DAP12 signaling;Signaling by PDGF;VEGFR2 mediated cell proliferation;Frs2-mediated activation;ARMS-mediated activation;Gastrin-CREB signalling pathway via PKC and MAPK;Interleukin receptor SHC signaling;FCERI mediated MAPK activation;VEGFA-VEGFR2 Pathway;IRS-related events triggered by IGF1R 
                                               hsa04012;hsa04020;hsa04066;hsa04510;hsa04520;hsa05200;hsa05205;hsa05206;hsa05212;hsa05213;hsa05215;hsa05219;hsa05223;hsa05230 
                                               ERBB2 
                                               NA 
                                               azure4 
                                               1 
                                               1.63803680981595 
                                             
                                             
                                               MAPK1-mRNA 
                                               0.103 
                                               0.0638 
                                               -0.0225 
                                               0.228 
                                               1.07 
                                               0.985 
                                               1.17 
                                               0.123 
                                               1 
                                               loglinear 
                                               ARMS-mediated activation, CREB phosphorylation through the activation of Ras, DAP12 signaling, Downstream signal transduction, FCERI mediated MAPK activation, Frs2-mediated activation, Gastrin-CREB signalling pathway via PKC and MAPK, Interleukin receptor SHC signaling, IRS-related events triggered by IGF1R, MAP2K and MAPK activation, Negative feedback regulation of MAPK pathway, Negative regulation of FGFR1 signaling, Negative regulation of FGFR2 signaling, Negative regulation of FGFR3 signaling, Negative regulation of FGFR4 signaling, Negative regulation of MAPK pathway, RAFMAP kinase cascade, Regulation of HSF1-mediated heat shock response, RHO GTPases Activate WASPs and WAVEs, Signal transduction by L1, Signaling by ERBB2, Signaling by PDGF, Signaling by SCF-KIT, VEGFA-VEGFR2 Pathway, VEGFR2 mediated cell proliferation 
                                               SEBELIUS_2_C4636 
                                               MAPK1 
                                               mRNA 
                                               0 
                                               
                                               
                                               Negative regulation of FGFR1 signaling;Signal transduction by L1;Negative regulation of FGFR2 signaling;MAP2K and MAPK activation;RHO GTPases Activate WASPs and WAVEs;Senescence-Associated Secretory Phenotype (SASP);Regulation of HSF1-mediated heat shock response;Thrombin signalling through proteinase activated receptors (PARs);Negative regulation of FGFR4 signaling;Oxidative Stress Induced Senescence;Signal attenuation;Negative regulation of FGFR3 signaling;Negative feedback regulation of MAPK pathway;Growth hormone receptor signaling;PLC beta mediated events;FCERI mediated MAPK activation;CREB phosphorylation through the activation of Ras;Negative regulation of MAPK pathway;RAFMAP kinase cascade;Downstream signal transduction;VEGFR2 mediated cell proliferation;Signaling by SCF-KIT;Frs2-mediated activation;ARMS-mediated activation;Gastrin-CREB signalling pathway via PKC and MAPK;Interleukin receptor SHC signaling;DAP12 signaling;Signaling by PDGF;VEGFA-VEGFR2 Pathway;Signaling by ERBB2;IRS-related events triggered by IGF1R 
                                               hsa04010;hsa04012;hsa04014;hsa04015;hsa04022;hsa04024;hsa04062;hsa04066;hsa04068;hsa04071;hsa04072;hsa04114;hsa04150;hsa04151;hsa04261;hsa04270;hsa04320;hsa04350;hsa04360;hsa04370;hsa04380;hsa04510;hsa04520;hsa04540;hsa04550;hsa04611;hsa04620;hsa04621;hsa04650;hsa04660;hsa04662;hsa04664;hsa04666;hsa04668;hsa04713;hsa04720;hsa04722;hsa04723;hsa04724;hsa04725;hsa04726;hsa04730;hsa04810;hsa04910;hsa04912;hsa04914;hsa04915;hsa04916;hsa04917;hsa04919;hsa04921;hsa04930;hsa04933;hsa04960;hsa05010;hsa05020;hsa05034;hsa05131;hsa05132;hsa05133;hsa05140;hsa05142;hsa05145;hsa05152;hsa05160;hsa05161;hsa05164;hsa05200;hsa05203;hsa05205;hsa05206;hsa05210;hsa05211;hsa05212;hsa05213;hsa05214;hsa05215;hsa05216;hsa05218;hsa05219;hsa05220;hsa05221;hsa05223;hsa05230;hsa05231 
                                               MAPK1 
                                               NA 
                                               azure4 
                                               1 
                                               1.61442006269592 
                                             
                                             
                                               ACTG1-mRNA 
                                               -0.23 
                                               0.154 
                                               -0.532 
                                               0.0717 
                                               0.853 
                                               0.692 
                                               1.05 
                                               0.15 
                                               1 
                                               lm.nb 
                                               ARMS-mediated activation, Cell-extracellular matrix interactions, DAP12 signaling, Downstream signal transduction, EPH-ephrin mediated repulsion of cells, EPHB-mediated forward signaling, FCERI mediated MAPK activation, Frs2-mediated activation, Gastrin-CREB signalling pathway via PKC and MAPK, Interleukin receptor SHC signaling, IRS-related events triggered by IGF1R, RAFMAP kinase cascade, Regulation of actin dynamics for phagocytic cup formation, RHO GTPases Activate Formins, RHO GTPases activate IQGAPs, RHO GTPases Activate WASPs and WAVEs, Signaling by ERBB2, Signaling by PDGF, Signaling by SCF-KIT, VEGFA-VEGFR2 Pathway, VEGFR2 mediated cell proliferation 
                                               SEBELIUS_2_C4636 
                                               ACTG1 
                                               mRNA 
                                               0 
                                               
                                               
                                               RHO GTPases Activate WASPs and WAVEs;Translocation of GLUT4 to the plasma membrane;Recycling pathway of L1;Interaction between L1 and Ankyrins;EPHB-mediated forward signaling;EPH-ephrin mediated repulsion of cells;RHO GTPases Activate Formins;RHO GTPases activate IQGAPs;Cell-extracellular matrix interactions;Regulation of actin dynamics for phagocytic cup formation;RAFMAP kinase cascade;Downstream signal transduction;VEGFR2 mediated cell proliferation;Signaling by SCF-KIT;Frs2-mediated activation;ARMS-mediated activation;Gastrin-CREB signalling pathway via PKC and MAPK;Interleukin receptor SHC signaling;FCERI mediated MAPK activation;DAP12 signaling;Signaling by PDGF;VEGFA-VEGFR2 Pathway;Signaling by ERBB2;IRS-related events triggered by IGF1R 
                                               hsa04015;hsa04145;hsa04390;hsa04510;hsa04520;hsa04530;hsa04611;hsa04670;hsa04810;hsa04919;hsa04921;hsa05100;hsa05110;hsa05130;hsa05131;hsa05132;hsa05164;hsa05205;hsa05410;hsa05412;hsa05414;hsa05416 
                                               ACTG1 
                                               NA 
                                               azure4 
                                               1 
                                               -1.49350649350649 
                                             
                                             
                                               VCL-mRNA 
                                               -0.158 
                                               0.108 
                                               -0.37 
                                               0.0548 
                                               0.896 
                                               0.774 
                                               1.04 
                                               0.161 
                                               1 
                                               lm.nb 
                                               ARMS-mediated activation, DAP12 signaling, Downstream signal transduction, FCERI mediated MAPK activation, Frs2-mediated activation, Gastrin-CREB signalling pathway via PKC and MAPK, Interleukin receptor SHC signaling, IRS-related events triggered by IGF1R, MAP2K and MAPK activation, Platelet degranulation , RAFMAP kinase cascade, Signaling by ERBB2, Signaling by PDGF, Signaling by SCF-KIT, Smooth Muscle Contraction, VEGFA-VEGFR2 Pathway, VEGFR2 mediated cell proliferation 
                                               SEBELIUS_2_C4636 
                                               VCL 
                                               mRNA 
                                               0 
                                               
                                               
                                               Smooth Muscle Contraction;MAP2K and MAPK activation;Platelet degranulation ;RAFMAP kinase cascade;Downstream signal transduction;VEGFR2 mediated cell proliferation;Signaling by SCF-KIT;Frs2-mediated activation;ARMS-mediated activation;Gastrin-CREB signalling pathway via PKC and MAPK;Interleukin receptor SHC signaling;FCERI mediated MAPK activation;DAP12 signaling;Signaling by PDGF;VEGFA-VEGFR2 Pathway;Signaling by ERBB2;IRS-related events triggered by IGF1R 
                                               hsa04510;hsa04520;hsa04670;hsa04810;hsa05100;hsa05131;hsa05146 
                                               VCL 
                                               NA 
                                               azure4 
                                               1 
                                               -1.46296296296296 
                                             
                                           
                                           DE Results - ARMS-mediated activation genes - New.Annotation: miR142 vs.cel67  Table displaying each sample's global significance scores and directed global significance scores as defined in the heatmaps above. The global significance score is calculated as the square root of the mean squared t-statistic for the genes in a gene set, with t-statistics coming from the linear regression underlying our differential expression analysis. The directed global significance score is calculated as the square root of the mean signed squared t-statistic for the genes in a gene set, with t-statistics coming from the linear regression underlying our differential expression analysis.   
                                      
                                       
                                  
                                
                                
                                  
                                       
                                      
                                           Volcano Plot: New.Annotation: miR155 vs.cel67   More Plot Information                Volcano Plot: New.Annotation: miR155 vs.cel67  Volcano plot displaying each gene's -log10(p-value) and log2 fold change for the selected covariate. Highly statistically significant genes fall at the top of the plot, and highly differentially expressed genes fall to either side. Genes within the selected gene set are highlighted in orange. Horizontal lines indicate various False Discovery Rate (FDR) thresholds.   
                                           DE Results - ARMS-mediated activation genes - New.Annotation: miR155 vs.cel67   More Plot Information      Download CSV Data     
                                           
                                             
                                             
                                           
                                           
                                             
                                               Probe Label 
                                               Log2 fold change 
                                               std error (log2) 
                                               Lower confidence limit (log2) 
                                               Upper confidence limit (log2) 
                                               Linear fold change 
                                               Lower confidence limit (linear) 
                                               Upper confidence limit (linear) 
                                               P-value 
                                               BY.p.value 
                                               method 
                                               Gene.sets 
                                               Codeset.Name 
                                               Probe.Label 
                                               Analyte.Type 
                                               Is.Control 
                                               Control.Type 
                                               Related.Probes 
                                               Probe.Annotation 
                                               KEGG.Pathways 
                                               Official.Gene.Name 
                                               Control.Conc 
                                               volcanocol 
                                               volcanopch 
                                               tstats.all 
                                             
                                           
                                           
                                             
                                               PPP2CB-mRNA 
                                               -1.13 
                                               0.0787 
                                               -1.29 
                                               -0.979 
                                               0.456 
                                               0.41 
                                               0.507 
                                               2.34e-12 
                                               3.11e-09 
                                               loglinear 
                                               ARMS-mediated activation, AXIN missense mutants destabilize the destruction complex, Beta-catenin phosphorylation cascade, CTLA4 inhibitory signaling, Cyclin AB1 associated events during G2M transition, Cyclin D associated events in G1, DAP12 signaling, DARPP-32 events, Disassembly of the destruction complex and recruitment of AXIN to the membrane, Downstream signal transduction, ERKs are inactivated, FCERI mediated MAPK activation, Frs2-mediated activation, Gastrin-CREB signalling pathway via PKC and MAPK, Interleukin receptor SHC signaling, IRS-related events triggered by IGF1R, MASTL Facilitates Mitotic Progression, Misspliced GSK3beta mutants stabilize beta-catenin, Negative regulation of FGFR1 signaling, Negative regulation of FGFR2 signaling, Negative regulation of FGFR3 signaling, Negative regulation of FGFR4 signaling, Negative regulation of MAPK pathway, RAF activation, RAFMAP kinase cascade, Resolution of Sister Chromatid Cohesion, RHO GTPases Activate Formins, S33 mutants of beta-catenin aren.SQT.t phosphorylated, S37 mutants of beta-catenin aren.SQT.t phosphorylated, S45 mutants of beta-catenin aren.SQT.t phosphorylated, Signaling by ERBB2, Signaling by PDGF, Signaling by SCF-KIT, Spry regulation of FGF signaling, T41 mutants of beta-catenin aren.SQT.t phosphorylated, Truncations of AMER1 destabilize the destruction complex, VEGFA-VEGFR2 Pathway, VEGFR2 mediated cell proliferation 
                                               SEBELIUS_2_C4636 
                                               PPP2CB 
                                               mRNA 
                                               0 
                                               
                                               
                                               MASTL Facilitates Mitotic Progression;Misspliced GSK3beta mutants stabilize beta-catenin;T41 mutants of beta-catenin aren.SQT.t phosphorylated;ERKs are inactivated;Glycolysis;Negative regulation of MAPK pathway;Disassembly of the destruction complex and recruitment of AXIN to the membrane;S37 mutants of beta-catenin aren.SQT.t phosphorylated;Cyclin AB1 associated events during G2M transition;RAF activation;RHO GTPases Activate Formins;S33 mutants of beta-catenin aren.SQT.t phosphorylated;DARPP-32 events;Beta-catenin phosphorylation cascade;Spry regulation of FGF signaling;Truncations of AMER1 destabilize the destruction complex;CTLA4 inhibitory signaling;AXIN missense mutants destabilize the destruction complex;S45 mutants of beta-catenin aren.SQT.t phosphorylated;Cyclin D associated events in G1;Platelet sensitization by LDL;Resolution of Sister Chromatid Cohesion;Inhibition of replication initiation of damaged DNA by RB1E2F1;RAFMAP kinase cascade;Downstream signal transduction;VEGFR2 mediated cell proliferation;Signaling by SCF-KIT;Frs2-mediated activation;ARMS-mediated activation;Gastrin-CREB signalling pathway via PKC and MAPK;Interleukin receptor SHC signaling;FCERI mediated MAPK activation;DAP12 signaling;Signaling by PDGF;VEGFA-VEGFR2 Pathway;Signaling by ERBB2;IRS-related events triggered by IGF1R;Negative regulation of FGFR3 signaling;Negative regulation of FGFR4 signaling;Negative regulation of FGFR2 signaling;Negative regulation of FGFR1 signaling 
                                               hsa03015;hsa04071;hsa04114;hsa04151;hsa04152;hsa04261;hsa04350;hsa04390;hsa04530;hsa04728;hsa04730;hsa05142;hsa05160 
                                               PPP2CB 
                                               NA 
                                               slateblue1 
                                               16 
                                               -14.3583227445997 
                                             
                                             
                                               KRAS-mRNA 
                                               -0.99 
                                               0.107 
                                               -1.2 
                                               -0.781 
                                               0.503 
                                               0.435 
                                               0.582 
                                               7.18e-09 
                                               2.06e-06 
                                               loglinear 
                                               ARMS-mediated activation, CD209 (DC-SIGN) signaling, Cell surface interactions at the vascular wall, Constitutive Signaling by EGFRvIII, DAP12 signaling, Downstream signal transduction, FCERI mediated MAPK activation, Frs2-mediated activation, Gastrin-CREB signalling pathway via PKC and MAPK, Interleukin receptor SHC signaling, IRS-related events triggered by IGF1R, MAP2K and MAPK activation, RAFMAP kinase cascade, SHC-mediated cascade.CL.FGFR1, SHC-related events triggered by IGF1R, Signaling by ERBB2, Signaling by FGFR1 in disease, Signaling by FGFR2 in disease, Signaling by FGFR3 in disease, Signaling by FGFR4 in disease, Signaling by PDGF, Signaling by SCF-KIT, Tie2 Signaling, VEGFA-VEGFR2 Pathway, VEGFR2 mediated cell proliferation 
                                               SEBELIUS_2_C4636 
                                               KRAS 
                                               mRNA 
                                               0 
                                               
                                               
                                               Signaling by FGFR2 in disease;Regulation of RAS by GAPs;SHC-mediated cascade.CL.FGFR3;Signaling by FGFR3 in disease;MAP2K and MAPK activation;SHC-related events triggered by IGF1R;Signaling by FGFR4 in disease;SHC-mediated cascade.CL.FGFR1;Tie2 Signaling;Signaling by FGFR1 in disease;Activation of RAS in B cells;SHC-mediated cascade.CL.FGFR4;Constitutive Signaling by EGFRvIII;CD209 (DC-SIGN) signaling;FCERI mediated MAPK activation;RAFMAP kinase cascade;Downstream signal transduction;VEGFR2 mediated cell proliferation;Signaling by SCF-KIT;Frs2-mediated activation;ARMS-mediated activation;Gastrin-CREB signalling pathway via PKC and MAPK;Interleukin receptor SHC signaling;DAP12 signaling;Signaling by PDGF;VEGFA-VEGFR2 Pathway;Signaling by ERBB2;IRS-related events triggered by IGF1R;Cell surface interactions at the vascular wall 
                                               hsa04010;hsa04012;hsa04014;hsa04015;hsa04062;hsa04068;hsa04071;hsa04072;hsa04151;hsa04211;hsa04213;hsa04320;hsa04360;hsa04370;hsa04530;hsa04540;hsa04550;hsa04650;hsa04660;hsa04662;hsa04664;hsa04720;hsa04722;hsa04725;hsa04726;hsa04730;hsa04810;hsa04910;hsa04912;hsa04914;hsa04915;hsa04916;hsa04917;hsa04919;hsa04921;hsa04933;hsa04960;hsa05034;hsa05160;hsa05161;hsa05166;hsa05200;hsa05203;hsa05205;hsa05206;hsa05210;hsa05211;hsa05212;hsa05213;hsa05214;hsa05215;hsa05216;hsa05218;hsa05219;hsa05220;hsa05221;hsa05223;hsa05230;hsa05231 
                                               KRAS 
                                               NA 
                                               slateblue1 
                                               16 
                                               -9.25233644859813 
                                             
                                             
                                               EGFR-mRNA 
                                               -0.978 
                                               0.106 
                                               -1.19 
                                               -0.771 
                                               0.508 
                                               0.439 
                                               0.586 
                                               7.76e-09 
                                               2.06e-06 
                                               loglinear 
                                               ARMS-mediated activation, Constitutive Signaling by Aberrant PI3K in Cancer, Constitutive Signaling by EGFRvIII, DAP12 signaling, Downstream signal transduction, FCERI mediated MAPK activation, Frs2-mediated activation, GAB1 signalosome, Gastrin-CREB signalling pathway via PKC and MAPK, Interleukin receptor SHC signaling, IRS-related events triggered by IGF1R, PI3KAKT activation, Role of LAT2NTALLAB on calcium mobilization, Signaling by ERBB2, Signaling by PDGF, Signaling by SCF-KIT, VEGFA-VEGFR2 Pathway, VEGFR2 mediated cell proliferation 
                                               SEBELIUS_2_C4636 
                                               EGFR 
                                               mRNA 
                                               0 
                                               
                                               
                                               Constitutive Signaling by Aberrant PI3K in Cancer;Inhibition of Signaling by Overexpressed EGFR;PLCG1 events in ERBB2 signaling;EGFR interacts with phospholipase C-gamma;EGFR Transactivation by Gastrin;Constitutive Signaling by Ligand-Responsive EGFR Cancer Variants;Constitutive Signaling by EGFRvIII;GAB1 signalosome;Downstream signal transduction;Signaling by SCF-KIT;PI3KAKT activation;Role of LAT2NTALLAB on calcium mobilization;DAP12 signaling;Signaling by PDGF;Signaling by ERBB2;VEGFR2 mediated cell proliferation;Frs2-mediated activation;ARMS-mediated activation;Gastrin-CREB signalling pathway via PKC and MAPK;Interleukin receptor SHC signaling;FCERI mediated MAPK activation;VEGFA-VEGFR2 Pathway;IRS-related events triggered by IGF1R 
                                               hsa04010;hsa04012;hsa04014;hsa04015;hsa04020;hsa04060;hsa04066;hsa04068;hsa04072;hsa04144;hsa04151;hsa04320;hsa04510;hsa04520;hsa04540;hsa04810;hsa04912;hsa04915;hsa04921;hsa05120;hsa05160;hsa05200;hsa05205;hsa05206;hsa05212;hsa05213;hsa05214;hsa05215;hsa05218;hsa05219;hsa05223;hsa05230;hsa05231 
                                               EGFR 
                                               NA 
                                               slateblue1 
                                               16 
                                               -9.22641509433962 
                                             
                                             
                                               IRS1-mRNA 
                                               -0.579 
                                               0.114 
                                               -0.802 
                                               -0.356 
                                               0.669 
                                               0.573 
                                               0.781 
                                               4.83e-05 
                                               0.00428 
                                               loglinear 
                                               ARMS-mediated activation, Constitutive Signaling by Aberrant PI3K in Cancer, DAP12 signaling, Downstream signal transduction, FCERI mediated MAPK activation, Frs2-mediated activation, GAB1 signalosome, Gastrin-CREB signalling pathway via PKC and MAPK, Interleukin receptor SHC signaling, IRS-related events triggered by IGF1R, PI3K Cascade, PI3KAKT activation, RAFMAP kinase cascade, Role of LAT2NTALLAB on calcium mobilization, Signaling by ERBB2, Signaling by PDGF, Signaling by SCF-KIT, VEGFA-VEGFR2 Pathway, VEGFR2 mediated cell proliferation 
                                               SEBELIUS_2_C4636 
                                               IRS1 
                                               mRNA 
                                               0 
                                               
                                               
                                               Constitutive Signaling by Aberrant PI3K in Cancer;RAFMAP kinase cascade;PI3K Cascade;Growth hormone receptor signaling;IRS-related events triggered by IGF1R;PI3KAKT activation;Downstream signal transduction;GAB1 signalosome;Signaling by SCF-KIT;Role of LAT2NTALLAB on calcium mobilization;DAP12 signaling;Signaling by PDGF;Signaling by ERBB2;VEGFR2 mediated cell proliferation;Frs2-mediated activation;ARMS-mediated activation;Gastrin-CREB signalling pathway via PKC and MAPK;Interleukin receptor SHC signaling;FCERI mediated MAPK activation;VEGFA-VEGFR2 Pathway 
                                               hsa04022;hsa04068;hsa04150;hsa04151;hsa04152;hsa04211;hsa04213;hsa04722;hsa04910;hsa04920;hsa04923;hsa04930;hsa04931;hsa04932;hsa04960;hsa05206 
                                               IRS1 
                                               NA 
                                               slateblue1 
                                               16 
                                               -5.07894736842105 
                                             
                                             
                                               MAPK3-mRNA 
                                               0.262 
                                               0.057 
                                               0.151 
                                               0.374 
                                               1.2 
                                               1.11 
                                               1.3 
                                               0.000153 
                                               0.012 
                                               loglinear 
                                               ARMS-mediated activation, DAP12 signaling, Downstream signal transduction, ERKs are inactivated, FCERI mediated MAPK activation, Frs2-mediated activation, Gastrin-CREB signalling pathway via PKC and MAPK, Interleukin receptor SHC signaling, IRS-related events triggered by IGF1R, Negative feedback regulation of MAPK pathway, Negative regulation of FGFR3 signaling, Negative regulation of FGFR4 signaling, Negative regulation of MAPK pathway, RAFMAP kinase cascade, Regulation of HSF1-mediated heat shock response, RHO GTPases Activate WASPs and WAVEs, Signaling by ERBB2, Signaling by PDGF, Signaling by SCF-KIT, VEGFA-VEGFR2 Pathway, VEGFR2 mediated cell proliferation 
                                               SEBELIUS_2_C4636 
                                               MAPK3 
                                               mRNA 
                                               0 
                                               
                                               
                                               ERKs are inactivated;ISG15 antiviral mechanism;RHO GTPases Activate WASPs and WAVEs;Senescence-Associated Secretory Phenotype (SASP);Regulation of HSF1-mediated heat shock response;Thrombin signalling through proteinase activated receptors (PARs);Negative regulation of FGFR4 signaling;Oxidative Stress Induced Senescence;Signal attenuation;RNA Polymerase I Promoter Opening;Negative regulation of FGFR3 signaling;Negative feedback regulation of MAPK pathway;Growth hormone receptor signaling;Golgi Cisternae Pericentriolar Stack Reorganization;FCERI mediated MAPK activation;Negative regulation of MAPK pathway;RAFMAP kinase cascade;Downstream signal transduction;VEGFR2 mediated cell proliferation;Signaling by SCF-KIT;Frs2-mediated activation;ARMS-mediated activation;Gastrin-CREB signalling pathway via PKC and MAPK;Interleukin receptor SHC signaling;DAP12 signaling;Signaling by PDGF;VEGFA-VEGFR2 Pathway;Signaling by ERBB2;IRS-related events triggered by IGF1R 
                                               hsa04010;hsa04012;hsa04014;hsa04015;hsa04022;hsa04024;hsa04062;hsa04066;hsa04068;hsa04071;hsa04072;hsa04114;hsa04150;hsa04151;hsa04261;hsa04270;hsa04320;hsa04350;hsa04360;hsa04370;hsa04380;hsa04510;hsa04520;hsa04540;hsa04550;hsa04611;hsa04620;hsa04621;hsa04650;hsa04660;hsa04662;hsa04664;hsa04666;hsa04668;hsa04713;hsa04720;hsa04722;hsa04723;hsa04724;hsa04725;hsa04726;hsa04730;hsa04810;hsa04910;hsa04912;hsa04914;hsa04915;hsa04916;hsa04917;hsa04919;hsa04921;hsa04930;hsa04933;hsa04960;hsa05010;hsa05020;hsa05034;hsa05131;hsa05132;hsa05133;hsa05140;hsa05142;hsa05145;hsa05152;hsa05160;hsa05161;hsa05164;hsa05200;hsa05203;hsa05205;hsa05210;hsa05211;hsa05212;hsa05213;hsa05214;hsa05215;hsa05216;hsa05218;hsa05219;hsa05220;hsa05221;hsa05223;hsa05230;hsa05231 
                                               MAPK3 
                                               NA 
                                               slateblue2 
                                               16 
                                               4.59649122807018 
                                             
                                             
                                               SOS1-mRNA 
                                               -0.396 
                                               0.0917 
                                               -0.576 
                                               -0.216 
                                               0.76 
                                               0.671 
                                               0.861 
                                               0.000304 
                                               0.0224 
                                               loglinear 
                                               ARMS-mediated activation, Cell surface interactions at the vascular wall, Constitutive Signaling by EGFRvIII, DAP12 signaling, Downstream signal transduction, FCERI mediated MAPK activation, Frs2-mediated activation, Gastrin-CREB signalling pathway via PKC and MAPK, Integrin alphaIIb beta3 signaling, Interleukin receptor SHC signaling, IRS-related events triggered by IGF1R, NRAGE signals death through JNK, RAFMAP kinase cascade, Role of LAT2NTALLAB on calcium mobilization, SHC-mediated cascade.CL.FGFR1, SHC-related events triggered by IGF1R, Signaling by ERBB2, Signaling by FGFR1 in disease, Signaling by FGFR2 in disease, Signaling by FGFR3 in disease, Signaling by FGFR4 in disease, Signaling by PDGF, Signaling by SCF-KIT, Tie2 Signaling, VEGFA-VEGFR2 Pathway, VEGFR2 mediated cell proliferation 
                                               SEBELIUS_2_C4636 
                                               SOS1 
                                               mRNA 
                                               0 
                                               
                                               
                                               Signaling by FGFR2 in disease;Antigen activates B Cell Receptor (BCR) leading to generation of second messengers;SHC-mediated cascade.CL.FGFR3;Signaling by FGFR3 in disease;RAFMAP kinase cascade;SHC-related events triggered by IGF1R;Signaling by FGFR4 in disease;SHC-mediated cascade.CL.FGFR1;GRB2.CL.SOS provides linkage to MAPK signaling for Integrins ;Tie2 Signaling;Role of LAT2NTALLAB on calcium mobilization;Signaling by FGFR1 in disease;Signal attenuation;NRAGE signals death through JNK;SHC-mediated cascade.CL.FGFR4;FCERI mediated Ca.PLS.2 mobilization;Constitutive Signaling by EGFRvIII;FCERI mediated MAPK activation;Signaling by ERBB2;Signaling by SCF-KIT;DAP12 signaling;Gastrin-CREB signalling pathway via PKC and MAPK;Interleukin receptor SHC signaling;IRS-related events triggered by IGF1R;Downstream signal transduction;VEGFR2 mediated cell proliferation;Frs2-mediated activation;ARMS-mediated activation;Signaling by PDGF;VEGFA-VEGFR2 Pathway;Integrin alphaIIb beta3 signaling;Cell surface interactions at the vascular wall 
                                               hsa04010;hsa04012;hsa04014;hsa04062;hsa04068;hsa04072;hsa04151;hsa04320;hsa04510;hsa04540;hsa04630;hsa04650;hsa04660;hsa04662;hsa04664;hsa04722;hsa04810;hsa04910;hsa04912;hsa04915;hsa04917;hsa05034;hsa05160;hsa05200;hsa05205;hsa05206;hsa05211;hsa05213;hsa05214;hsa05215;hsa05220;hsa05221;hsa05223;hsa05231 
                                               SOS1 
                                               NA 
                                               slateblue2 
                                               16 
                                               -4.31842966194111 
                                             
                                             
                                               FGFR1-mRNA 
                                               -0.452 
                                               0.108 
                                               -0.663 
                                               -0.241 
                                               0.731 
                                               0.631 
                                               0.846 
                                               0.000398 
                                               0.0264 
                                               lm.nb 
                                               ARMS-mediated activation, Constitutive Signaling by Aberrant PI3K in Cancer, DAP12 signaling, Downstream signal transduction, FCERI mediated MAPK activation, Frs2-mediated activation, GAB1 signalosome, Gastrin-CREB signalling pathway via PKC and MAPK, Interleukin receptor SHC signaling, IRS-related events triggered by IGF1R, PI3K Cascade, PI3KAKT activation, RAFMAP kinase cascade, Role of LAT2NTALLAB on calcium mobilization, SHC-mediated cascade.CL.FGFR1, Signaling by ERBB2, Signaling by FGFR1 in disease, Signaling by PDGF, Signaling by SCF-KIT, VEGFA-VEGFR2 Pathway, VEGFR2 mediated cell proliferation 
                                               SEBELIUS_2_C4636 
                                               FGFR1 
                                               mRNA 
                                               0 
                                               
                                               
                                               RAFMAP kinase cascade;Constitutive Signaling by Aberrant PI3K in Cancer;SHC-mediated cascade.CL.FGFR1;Phospholipase C-mediated cascade.CL. FGFR1;Signaling by FGFR1 amplification mutants;PI3K Cascade;FGFR1b ligand binding and activation;Signaling by FGFR1 in disease;Downstream signal transduction;GAB1 signalosome;Signaling by SCF-KIT;PI3KAKT activation;Role of LAT2NTALLAB on calcium mobilization;DAP12 signaling;Signaling by PDGF;Signaling by ERBB2;VEGFR2 mediated cell proliferation;Frs2-mediated activation;ARMS-mediated activation;Gastrin-CREB signalling pathway via PKC and MAPK;Interleukin receptor SHC signaling;FCERI mediated MAPK activation;VEGFA-VEGFR2 Pathway;IRS-related events triggered by IGF1R 
                                               hsa04010;hsa04014;hsa04015;hsa04151;hsa04520;hsa04550;hsa04810;hsa05200;hsa05205;hsa05215;hsa05218;hsa05230 
                                               FGFR1 
                                               NA 
                                               slateblue2 
                                               16 
                                               -4.18518518518519 
                                             
                                             
                                               PPP2CA-mRNA 
                                               -0.221 
                                               0.0556 
                                               -0.33 
                                               -0.112 
                                               0.858 
                                               0.796 
                                               0.925 
                                               0.000682 
                                               0.0404 
                                               loglinear 
                                               ARMS-mediated activation, AXIN missense mutants destabilize the destruction complex, Beta-catenin phosphorylation cascade, CTLA4 inhibitory signaling, Cyclin AB1 associated events during G2M transition, Cyclin D associated events in G1, DAP12 signaling, DARPP-32 events, Disassembly of the destruction complex and recruitment of AXIN to the membrane, Downstream signal transduction, ERKs are inactivated, FCERI mediated MAPK activation, Frs2-mediated activation, Gastrin-CREB signalling pathway via PKC and MAPK, Interleukin receptor SHC signaling, IRS-related events triggered by IGF1R, MASTL Facilitates Mitotic Progression, Negative regulation of FGFR1 signaling, Negative regulation of FGFR2 signaling, Negative regulation of FGFR3 signaling, Negative regulation of FGFR4 signaling, Negative regulation of MAPK pathway, Nonsense Mediated Decay (NMD) enhanced by the Exon Junction Complex (EJC), RAF activation, RAFMAP kinase cascade, Resolution of Sister Chromatid Cohesion, RHO GTPases Activate Formins, S33 mutants of beta-catenin aren.SQT.t phosphorylated, S37 mutants of beta-catenin aren.SQT.t phosphorylated, S45 mutants of beta-catenin aren.SQT.t phosphorylated, Signaling by ERBB2, Signaling by PDGF, Signaling by SCF-KIT, Spry regulation of FGF signaling, T41 mutants of beta-catenin aren.SQT.t phosphorylated, Truncations of AMER1 destabilize the destruction complex, VEGFA-VEGFR2 Pathway, VEGFR2 mediated cell proliferation 
                                               SEBELIUS_2_C4636 
                                               PPP2CA 
                                               mRNA 
                                               0 
                                               
                                               
                                               MASTL Facilitates Mitotic Progression;T41 mutants of beta-catenin aren.SQT.t phosphorylated;ERKs are inactivated;Glycolysis;Negative regulation of MAPK pathway;Disassembly of the destruction complex and recruitment of AXIN to the membrane;S37 mutants of beta-catenin aren.SQT.t phosphorylated;Cyclin AB1 associated events during G2M transition;RAF activation;RHO GTPases Activate Formins;S33 mutants of beta-catenin aren.SQT.t phosphorylated;DARPP-32 events;Beta-catenin phosphorylation cascade;Initiation of Nuclear Envelope Reformation;Spry regulation of FGF signaling;Truncations of AMER1 destabilize the destruction complex;CTLA4 inhibitory signaling;AXIN missense mutants destabilize the destruction complex;S45 mutants of beta-catenin aren.SQT.t phosphorylated;Nonsense Mediated Decay (NMD) enhanced by the Exon Junction Complex (EJC);Cyclin D associated events in G1;Platelet sensitization by LDL;Resolution of Sister Chromatid Cohesion;Inhibition of replication initiation of damaged DNA by RB1E2F1;RAFMAP kinase cascade;Downstream signal transduction;VEGFR2 mediated cell proliferation;Signaling by SCF-KIT;Frs2-mediated activation;ARMS-mediated activation;Gastrin-CREB signalling pathway via PKC and MAPK;Interleukin receptor SHC signaling;FCERI mediated MAPK activation;DAP12 signaling;Signaling by PDGF;VEGFA-VEGFR2 Pathway;Signaling by ERBB2;IRS-related events triggered by IGF1R;Negative regulation of FGFR3 signaling;Negative regulation of FGFR4 signaling;Negative regulation of FGFR2 signaling;Negative regulation of FGFR1 signaling 
                                               hsa03015;hsa04071;hsa04114;hsa04151;hsa04152;hsa04261;hsa04350;hsa04390;hsa04530;hsa04728;hsa04730;hsa05142;hsa05160 
                                               PPP2CA 
                                               NA 
                                               slateblue2 
                                               16 
                                               -3.97482014388489 
                                             
                                             
                                               HRAS-mRNA 
                                               -0.573 
                                               0.152 
                                               -0.87 
                                               -0.276 
                                               0.672 
                                               0.547 
                                               0.826 
                                               0.0011 
                                               0.0583 
                                               loglinear 
                                               ARMS-mediated activation, CD209 (DC-SIGN) signaling, Cell surface interactions at the vascular wall, Constitutive Signaling by EGFRvIII, CREB phosphorylation through the activation of Ras, DAP12 signaling, Downstream signal transduction, FCERI mediated MAPK activation, Frs2-mediated activation, Gastrin-CREB signalling pathway via PKC and MAPK, Interleukin receptor SHC signaling, IRS-related events triggered by IGF1R, MAP2K and MAPK activation, RAFMAP kinase cascade, SHC-mediated cascade.CL.FGFR1, SHC-related events triggered by IGF1R, Signaling by ERBB2, Signaling by FGFR1 in disease, Signaling by FGFR2 in disease, Signaling by FGFR3 in disease, Signaling by FGFR4 in disease, Signaling by PDGF, Signaling by SCF-KIT, Tie2 Signaling, VEGFA-VEGFR2 Pathway, VEGFR2 mediated cell proliferation 
                                               SEBELIUS_2_C4636 
                                               HRAS 
                                               mRNA 
                                               0 
                                               
                                               
                                               Signaling by FGFR2 in disease;Regulation of RAS by GAPs;SHC-mediated cascade.CL.FGFR3;Signaling by FGFR3 in disease;MAP2K and MAPK activation;SHC-related events triggered by IGF1R;Signaling by FGFR4 in disease;SHC-mediated cascade.CL.FGFR1;Tie2 Signaling;Signaling by FGFR1 in disease;Activation of RAS in B cells;SHC-mediated cascade.CL.FGFR4;Constitutive Signaling by EGFRvIII;CD209 (DC-SIGN) signaling;Ras activation uopn Ca2.PLS. infux through NMDA receptor;FCERI mediated MAPK activation;RAFMAP kinase cascade;Downstream signal transduction;VEGFR2 mediated cell proliferation;Signaling by SCF-KIT;Frs2-mediated activation;ARMS-mediated activation;Gastrin-CREB signalling pathway via PKC and MAPK;Interleukin receptor SHC signaling;DAP12 signaling;Signaling by PDGF;VEGFA-VEGFR2 Pathway;Signaling by ERBB2;IRS-related events triggered by IGF1R;Cell surface interactions at the vascular wall;CREB phosphorylation through the activation of Ras 
                                               hsa04010;hsa04012;hsa04014;hsa04015;hsa04062;hsa04068;hsa04071;hsa04072;hsa04144;hsa04151;hsa04211;hsa04213;hsa04360;hsa04370;hsa04510;hsa04530;hsa04540;hsa04550;hsa04630;hsa04650;hsa04660;hsa04662;hsa04664;hsa04720;hsa04722;hsa04725;hsa04726;hsa04730;hsa04810;hsa04910;hsa04912;hsa04915;hsa04916;hsa04917;hsa04919;hsa04921;hsa04933;hsa05034;hsa05160;hsa05161;hsa05166;hsa05200;hsa05203;hsa05205;hsa05206;hsa05211;hsa05213;hsa05214;hsa05215;hsa05216;hsa05218;hsa05219;hsa05220;hsa05221;hsa05223;hsa05230;hsa05231 
                                               HRAS 
                                               NA 
                                               slateblue3 
                                               16 
                                               -3.76973684210526 
                                             
                                             
                                               SPTAN1-mRNA 
                                               0.316 
                                               0.117 
                                               0.0862 
                                               0.546 
                                               1.25 
                                               1.06 
                                               1.46 
                                               0.0136 
                                               0.295 
                                               loglinear 
                                               Apoptotic cleavage of cellular proteins, ARMS-mediated activation, DAP12 signaling, Downstream signal transduction, FCERI mediated MAPK activation, Frs2-mediated activation, Gastrin-CREB signalling pathway via PKC and MAPK, Interleukin receptor SHC signaling, IRS-related events triggered by IGF1R, Nephrin interactions, Signaling by ERBB2, Signaling by PDGF, Signaling by SCF-KIT, VEGFA-VEGFR2 Pathway, VEGFR2 mediated cell proliferation 
                                               SEBELIUS_2_C4636 
                                               SPTAN1 
                                               mRNA 
                                               0 
                                               
                                               
                                               Nephrin interactions;Caspase-mediated cleavage of cytoskeletal proteins;Downstream signal transduction;VEGFR2 mediated cell proliferation;Signaling by SCF-KIT;Frs2-mediated activation;ARMS-mediated activation;Gastrin-CREB signalling pathway via PKC and MAPK;Interleukin receptor SHC signaling;FCERI mediated MAPK activation;DAP12 signaling;Signaling by PDGF;VEGFA-VEGFR2 Pathway;Signaling by ERBB2;IRS-related events triggered by IGF1R;Apoptotic cleavage of cellular proteins 
                                               hsa04530 
                                               SPTAN1 
                                               NA 
                                               slateblue4 
                                               16 
                                               2.7008547008547 
                                             
                                             
                                               PTK2-mRNA 
                                               -0.182 
                                               0.0735 
                                               -0.326 
                                               -0.0377 
                                               0.882 
                                               0.798 
                                               0.974 
                                               0.022 
                                               0.4 
                                               loglinear 
                                               Apoptotic cleavage of cellular proteins, ARMS-mediated activation, DAP12 signaling, DCC mediated attractive signaling, Downstream signal transduction, EPHA-mediated growth cone collapse, EPHB-mediated forward signaling, FCERI mediated MAPK activation, Frs2-mediated activation, Gastrin-CREB signalling pathway via PKC and MAPK, Integrin alphaIIb beta3 signaling, Interleukin receptor SHC signaling, IRS-related events triggered by IGF1R, p130Cas linkage to MAPK signaling for integrins, RAFMAP kinase cascade, Regulation of actin dynamics for phagocytic cup formation, RHO GTPases Activate WASPs and WAVEs, Signaling by ERBB2, Signaling by PDGF, Signaling by SCF-KIT, VEGFA-VEGFR2 Pathway, VEGFR2 mediated cell proliferation 
                                               SEBELIUS_2_C4636 
                                               PTK2 
                                               mRNA 
                                               0 
                                               
                                               
                                               Apoptotic cleavage of cellular proteins;RAFMAP kinase cascade;RHO GTPases Activate WASPs and WAVEs;DCC mediated attractive signaling;p130Cas linkage to MAPK signaling for integrins;EPHB-mediated forward signaling;EPHA-mediated growth cone collapse;Regulation of actin dynamics for phagocytic cup formation;Downstream signal transduction;VEGFR2 mediated cell proliferation;Signaling by SCF-KIT;Frs2-mediated activation;ARMS-mediated activation;Gastrin-CREB signalling pathway via PKC and MAPK;Interleukin receptor SHC signaling;FCERI mediated MAPK activation;DAP12 signaling;Signaling by PDGF;VEGFA-VEGFR2 Pathway;Signaling by ERBB2;IRS-related events triggered by IGF1R;Integrin alphaIIb beta3 signaling 
                                               hsa04012;hsa04062;hsa04151;hsa04360;hsa04370;hsa04510;hsa04670;hsa04810;hsa05100;hsa05146;hsa05200;hsa05202;hsa05205;hsa05222 
                                               PTK2 
                                               NA 
                                               azure4 
                                               1 
                                               -2.47619047619048 
                                             
                                             
                                               PPP2R1B-mRNA 
                                               -0.108 
                                               0.0625 
                                               -0.23 
                                               0.0148 
                                               0.928 
                                               0.852 
                                               1.01 
                                               0.0996 
                                               1 
                                               loglinear 
                                               ARMS-mediated activation, AXIN missense mutants destabilize the destruction complex, Beta-catenin phosphorylation cascade, CTLA4 inhibitory signaling, Cyclin AB1 associated events during G2M transition, Cyclin D associated events in G1, DAP12 signaling, DARPP-32 events, Disassembly of the destruction complex and recruitment of AXIN to the membrane, Downstream signal transduction, ERKs are inactivated, FCERI mediated MAPK activation, Frs2-mediated activation, Gastrin-CREB signalling pathway via PKC and MAPK, Interleukin receptor SHC signaling, IRS-related events triggered by IGF1R, MASTL Facilitates Mitotic Progression, Misspliced GSK3beta mutants stabilize beta-catenin, Negative regulation of MAPK pathway, RAF activation, RAFMAP kinase cascade, Resolution of Sister Chromatid Cohesion, RHO GTPases Activate Formins, S33 mutants of beta-catenin aren.SQT.t phosphorylated, S37 mutants of beta-catenin aren.SQT.t phosphorylated, S45 mutants of beta-catenin aren.SQT.t phosphorylated, Signaling by ERBB2, Signaling by PDGF, Signaling by SCF-KIT, T41 mutants of beta-catenin aren.SQT.t phosphorylated, Truncations of AMER1 destabilize the destruction complex, VEGFA-VEGFR2 Pathway, VEGFR2 mediated cell proliferation 
                                               SEBELIUS_2_C4636 
                                               PPP2R1B 
                                               mRNA 
                                               0 
                                               
                                               
                                               MASTL Facilitates Mitotic Progression;Misspliced GSK3beta mutants stabilize beta-catenin;T41 mutants of beta-catenin aren.SQT.t phosphorylated;ERKs are inactivated;Glycolysis;Negative regulation of MAPK pathway;Disassembly of the destruction complex and recruitment of AXIN to the membrane;S37 mutants of beta-catenin aren.SQT.t phosphorylated;Cyclin AB1 associated events during G2M transition;RAF activation;RHO GTPases Activate Formins;S33 mutants of beta-catenin aren.SQT.t phosphorylated;DARPP-32 events;Beta-catenin phosphorylation cascade;Truncations of AMER1 destabilize the destruction complex;CTLA4 inhibitory signaling;AXIN missense mutants destabilize the destruction complex;S45 mutants of beta-catenin aren.SQT.t phosphorylated;Cyclin D associated events in G1;Platelet sensitization by LDL;Resolution of Sister Chromatid Cohesion;Inhibition of replication initiation of damaged DNA by RB1E2F1;RAFMAP kinase cascade;Downstream signal transduction;VEGFR2 mediated cell proliferation;Signaling by SCF-KIT;Frs2-mediated activation;ARMS-mediated activation;Gastrin-CREB signalling pathway via PKC and MAPK;Interleukin receptor SHC signaling;FCERI mediated MAPK activation;DAP12 signaling;Signaling by PDGF;VEGFA-VEGFR2 Pathway;Signaling by ERBB2;IRS-related events triggered by IGF1R 
                                               hsa03015;hsa04071;hsa04114;hsa04151;hsa04152;hsa04261;hsa04350;hsa04390;hsa04530;hsa04728;hsa04730;hsa05142;hsa05160 
                                               PPP2R1B 
                                               NA 
                                               azure4 
                                               1 
                                               -1.728 
                                             
                                             
                                               MAP2K1-mRNA 
                                               -0.205 
                                               0.126 
                                               -0.452 
                                               0.0411 
                                               0.867 
                                               0.731 
                                               1.03 
                                               0.117 
                                               1 
                                               lm.nb 
                                               ARMS-mediated activation, DAP12 signaling, Downstream signal transduction, FCERI mediated MAPK activation, Frs2-mediated activation, Gastrin-CREB signalling pathway via PKC and MAPK, Interleukin receptor SHC signaling, IRS-related events triggered by IGF1R, Negative feedback regulation of MAPK pathway, Negative regulation of MAPK pathway, RAF activation, RAFMAP kinase cascade, Signaling by ERBB2, Signaling by PDGF, Signaling by SCF-KIT, VEGFA-VEGFR2 Pathway, VEGFR2 mediated cell proliferation 
                                               SEBELIUS_2_C4636 
                                               MAP2K1 
                                               mRNA 
                                               0 
                                               
                                               
                                               RAF activation;Uptake and function of anthrax toxins;Negative feedback regulation of MAPK pathway;Interleukin-1 signaling;RAFMAP kinase cascade;Downstream signal transduction;VEGFR2 mediated cell proliferation;Signaling by SCF-KIT;Frs2-mediated activation;ARMS-mediated activation;Gastrin-CREB signalling pathway via PKC and MAPK;Interleukin receptor SHC signaling;FCERI mediated MAPK activation;DAP12 signaling;Signaling by PDGF;VEGFA-VEGFR2 Pathway;Signaling by ERBB2;IRS-related events triggered by IGF1R;Negative regulation of MAPK pathway 
                                               hsa04010;hsa04012;hsa04014;hsa04015;hsa04022;hsa04024;hsa04062;hsa04066;hsa04068;hsa04071;hsa04072;hsa04114;hsa04151;hsa04270;hsa04320;hsa04370;hsa04380;hsa04510;hsa04540;hsa04550;hsa04620;hsa04650;hsa04660;hsa04662;hsa04664;hsa04666;hsa04668;hsa04720;hsa04722;hsa04725;hsa04726;hsa04730;hsa04810;hsa04910;hsa04912;hsa04914;hsa04915;hsa04916;hsa04917;hsa04919;hsa04921;hsa05020;hsa05034;hsa05161;hsa05164;hsa05200;hsa05205;hsa05206;hsa05210;hsa05211;hsa05212;hsa05213;hsa05214;hsa05215;hsa05216;hsa05218;hsa05219;hsa05220;hsa05221;hsa05223;hsa05230;hsa05231 
                                               MAP2K1 
                                               NA 
                                               azure4 
                                               1 
                                               -1.62698412698413 
                                             
                                             
                                               CRK-mRNA 
                                               -0.176 
                                               0.108 
                                               -0.389 
                                               0.0364 
                                               0.885 
                                               0.764 
                                               1.03 
                                               0.119 
                                               1 
                                               loglinear 
                                               ARMS-mediated activation, Downstream signal transduction, Integrin alphaIIb beta3 signaling, p130Cas linkage to MAPK signaling for integrins, Regulation of actin dynamics for phagocytic cup formation, Regulation of signaling by CBL, Signaling by PDGF, VEGFA-VEGFR2 Pathway 
                                               SEBELIUS_2_C4636 
                                               CRK 
                                               mRNA 
                                               0 
                                               
                                               
                                               ARMS-mediated activation;Downstream signal transduction;Regulation of signaling by CBL;p130Cas linkage to MAPK signaling for integrins;VEGFA-VEGFR2 Pathway;Regulation of actin dynamics for phagocytic cup formation;Signaling by PDGF;Integrin alphaIIb beta3 signaling 
                                               hsa04010;hsa04012;hsa04015;hsa04062;hsa04510;hsa04666;hsa04722;hsa04810;hsa04910;hsa05100;hsa05131;hsa05200;hsa05206;hsa05211;hsa05220 
                                               CRK 
                                               NA 
                                               azure4 
                                               1 
                                               -1.62962962962963 
                                             
                                             
                                               NRAS-mRNA 
                                               0.17 
                                               0.119 
                                               -0.0629 
                                               0.403 
                                               1.13 
                                               0.957 
                                               1.32 
                                               0.167 
                                               1 
                                               lm.nb 
                                               ARMS-mediated activation, CD209 (DC-SIGN) signaling, Cell surface interactions at the vascular wall, Constitutive Signaling by EGFRvIII, DAP12 signaling, Downstream signal transduction, FCERI mediated MAPK activation, Frs2-mediated activation, Gastrin-CREB signalling pathway via PKC and MAPK, Interleukin receptor SHC signaling, IRS-related events triggered by IGF1R, MAP2K and MAPK activation, RAFMAP kinase cascade, SHC-mediated cascade.CL.FGFR1, SHC-related events triggered by IGF1R, Signaling by ERBB2, Signaling by FGFR1 in disease, Signaling by FGFR2 in disease, Signaling by FGFR3 in disease, Signaling by FGFR4 in disease, Signaling by PDGF, Signaling by SCF-KIT, Tie2 Signaling, VEGFA-VEGFR2 Pathway, VEGFR2 mediated cell proliferation 
                                               SEBELIUS_2_C4636 
                                               NRAS 
                                               mRNA 
                                               0 
                                               
                                               
                                               Signaling by FGFR2 in disease;Regulation of RAS by GAPs;SHC-mediated cascade.CL.FGFR3;Signaling by FGFR3 in disease;MAP2K and MAPK activation;SHC-related events triggered by IGF1R;Signaling by FGFR4 in disease;SHC-mediated cascade.CL.FGFR1;Tie2 Signaling;Signaling by FGFR1 in disease;Activation of RAS in B cells;SHC-mediated cascade.CL.FGFR4;Constitutive Signaling by EGFRvIII;CD209 (DC-SIGN) signaling;FCERI mediated MAPK activation;RAFMAP kinase cascade;Downstream signal transduction;VEGFR2 mediated cell proliferation;Signaling by SCF-KIT;Frs2-mediated activation;ARMS-mediated activation;Gastrin-CREB signalling pathway via PKC and MAPK;Interleukin receptor SHC signaling;DAP12 signaling;Signaling by PDGF;VEGFA-VEGFR2 Pathway;Signaling by ERBB2;IRS-related events triggered by IGF1R;Cell surface interactions at the vascular wall 
                                               hsa04010;hsa04012;hsa04014;hsa04015;hsa04062;hsa04068;hsa04071;hsa04072;hsa04151;hsa04211;hsa04213;hsa04360;hsa04370;hsa04530;hsa04540;hsa04550;hsa04650;hsa04660;hsa04662;hsa04664;hsa04720;hsa04722;hsa04725;hsa04726;hsa04730;hsa04810;hsa04910;hsa04912;hsa04915;hsa04916;hsa04917;hsa04919;hsa04921;hsa04933;hsa05034;hsa05160;hsa05161;hsa05166;hsa05200;hsa05203;hsa05205;hsa05206;hsa05211;hsa05213;hsa05214;hsa05215;hsa05216;hsa05218;hsa05219;hsa05220;hsa05221;hsa05223;hsa05230;hsa05231 
                                               NRAS 
                                               NA 
                                               azure4 
                                               1 
                                               1.42857142857143 
                                             
                                             
                                               MAPK1-mRNA 
                                               0.0952 
                                               0.0673 
                                               -0.0366 
                                               0.227 
                                               1.07 
                                               0.975 
                                               1.17 
                                               0.172 
                                               1 
                                               loglinear 
                                               ARMS-mediated activation, CREB phosphorylation through the activation of Ras, DAP12 signaling, Downstream signal transduction, FCERI mediated MAPK activation, Frs2-mediated activation, Gastrin-CREB signalling pathway via PKC and MAPK, Interleukin receptor SHC signaling, IRS-related events triggered by IGF1R, MAP2K and MAPK activation, Negative feedback regulation of MAPK pathway, Negative regulation of FGFR1 signaling, Negative regulation of FGFR2 signaling, Negative regulation of FGFR3 signaling, Negative regulation of FGFR4 signaling, Negative regulation of MAPK pathway, RAFMAP kinase cascade, Regulation of HSF1-mediated heat shock response, RHO GTPases Activate WASPs and WAVEs, Signal transduction by L1, Signaling by ERBB2, Signaling by PDGF, Signaling by SCF-KIT, VEGFA-VEGFR2 Pathway, VEGFR2 mediated cell proliferation 
                                               SEBELIUS_2_C4636 
                                               MAPK1 
                                               mRNA 
                                               0 
                                               
                                               
                                               Negative regulation of FGFR1 signaling;Signal transduction by L1;Negative regulation of FGFR2 signaling;MAP2K and MAPK activation;RHO GTPases Activate WASPs and WAVEs;Senescence-Associated Secretory Phenotype (SASP);Regulation of HSF1-mediated heat shock response;Thrombin signalling through proteinase activated receptors (PARs);Negative regulation of FGFR4 signaling;Oxidative Stress Induced Senescence;Signal attenuation;Negative regulation of FGFR3 signaling;Negative feedback regulation of MAPK pathway;Growth hormone receptor signaling;PLC beta mediated events;FCERI mediated MAPK activation;CREB phosphorylation through the activation of Ras;Negative regulation of MAPK pathway;RAFMAP kinase cascade;Downstream signal transduction;VEGFR2 mediated cell proliferation;Signaling by SCF-KIT;Frs2-mediated activation;ARMS-mediated activation;Gastrin-CREB signalling pathway via PKC and MAPK;Interleukin receptor SHC signaling;DAP12 signaling;Signaling by PDGF;VEGFA-VEGFR2 Pathway;Signaling by ERBB2;IRS-related events triggered by IGF1R 
                                               hsa04010;hsa04012;hsa04014;hsa04015;hsa04022;hsa04024;hsa04062;hsa04066;hsa04068;hsa04071;hsa04072;hsa04114;hsa04150;hsa04151;hsa04261;hsa04270;hsa04320;hsa04350;hsa04360;hsa04370;hsa04380;hsa04510;hsa04520;hsa04540;hsa04550;hsa04611;hsa04620;hsa04621;hsa04650;hsa04660;hsa04662;hsa04664;hsa04666;hsa04668;hsa04713;hsa04720;hsa04722;hsa04723;hsa04724;hsa04725;hsa04726;hsa04730;hsa04810;hsa04910;hsa04912;hsa04914;hsa04915;hsa04916;hsa04917;hsa04919;hsa04921;hsa04930;hsa04933;hsa04960;hsa05010;hsa05020;hsa05034;hsa05131;hsa05132;hsa05133;hsa05140;hsa05142;hsa05145;hsa05152;hsa05160;hsa05161;hsa05164;hsa05200;hsa05203;hsa05205;hsa05206;hsa05210;hsa05211;hsa05212;hsa05213;hsa05214;hsa05215;hsa05216;hsa05218;hsa05219;hsa05220;hsa05221;hsa05223;hsa05230;hsa05231 
                                               MAPK1 
                                               NA 
                                               azure4 
                                               1 
                                               1.41456166419019 
                                             
                                             
                                               ACTB-mRNA 
                                               -0.255 
                                               0.201 
                                               -0.649 
                                               0.138 
                                               0.838 
                                               0.638 
                                               1.1 
                                               0.217 
                                               1 
                                               lm.nb 
                                               ARMS-mediated activation, Cell-extracellular matrix interactions, DAP12 signaling, Downstream signal transduction, EPH-ephrin mediated repulsion of cells, EPHB-mediated forward signaling, FCERI mediated MAPK activation, Frs2-mediated activation, Gastrin-CREB signalling pathway via PKC and MAPK, Interleukin receptor SHC signaling, IRS-related events triggered by IGF1R, RAFMAP kinase cascade, Regulation of actin dynamics for phagocytic cup formation, RHO GTPases Activate Formins, RHO GTPases activate IQGAPs, Signaling by ERBB2, Signaling by PDGF, Signaling by SCF-KIT, VEGFA-VEGFR2 Pathway, VEGFR2 mediated cell proliferation 
                                               SEBELIUS_2_C4636 
                                               ACTB 
                                               mRNA 
                                               0 
                                               
                                               
                                               EPHB-mediated forward signaling;Interaction between L1 and Ankyrins;DNA Damage Recognition in GG-NER;EPH-ephrin mediated repulsion of cells;Prefoldin mediated transfer of substrate  to CCTTriC;RHO GTPases Activate Formins;RHO GTPases activate IQGAPs;Cell-extracellular matrix interactions;Regulation of actin dynamics for phagocytic cup formation;Folding of actin by CCTTriC;B-WICH complex positively regulates rRNA expression;RAFMAP kinase cascade;Downstream signal transduction;VEGFR2 mediated cell proliferation;Signaling by SCF-KIT;Frs2-mediated activation;ARMS-mediated activation;Gastrin-CREB signalling pathway via PKC and MAPK;Interleukin receptor SHC signaling;FCERI mediated MAPK activation;DAP12 signaling;Signaling by PDGF;VEGFA-VEGFR2 Pathway;Signaling by ERBB2;IRS-related events triggered by IGF1R 
                                               hsa04015;hsa04145;hsa04390;hsa04510;hsa04520;hsa04530;hsa04611;hsa04670;hsa04810;hsa04919;hsa04921;hsa04971;hsa05100;hsa05110;hsa05130;hsa05131;hsa05132;hsa05164;hsa05205;hsa05410;hsa05412;hsa05414;hsa05416 
                                               ACTB 
                                               NA 
                                               azure4 
                                               1 
                                               -1.26865671641791 
                                             
                                             
                                               ERBB2-mRNA 
                                               0.205 
                                               0.172 
                                               -0.131 
                                               0.542 
                                               1.15 
                                               0.913 
                                               1.46 
                                               0.245 
                                               1 
                                               lm.nb 
                                               ARMS-mediated activation, DAP12 signaling, Downstream signal transduction, FCERI mediated MAPK activation, Frs2-mediated activation, GAB1 signalosome, Gastrin-CREB signalling pathway via PKC and MAPK, Interleukin receptor SHC signaling, IRS-related events triggered by IGF1R, PI3KAKT activation, RAFMAP kinase cascade, Role of LAT2NTALLAB on calcium mobilization, Sema4D induced cell migration and growth-cone collapse, Signaling by ERBB2, Signaling by PDGF, Signaling by SCF-KIT, VEGFA-VEGFR2 Pathway, VEGFR2 mediated cell proliferation 
                                               SEBELIUS_2_C4636 
                                               ERBB2 
                                               mRNA 
                                               0 
                                               
                                               
                                               RAFMAP kinase cascade;GRB7 events in ERBB2 signaling;PLCG1 events in ERBB2 signaling;Sema4D induced cell migration and growth-cone collapse;Signaling by ERBB2;Downstream signal transduction;GAB1 signalosome;Signaling by SCF-KIT;PI3KAKT activation;Role of LAT2NTALLAB on calcium mobilization;DAP12 signaling;Signaling by PDGF;VEGFR2 mediated cell proliferation;Frs2-mediated activation;ARMS-mediated activation;Gastrin-CREB signalling pathway via PKC and MAPK;Interleukin receptor SHC signaling;FCERI mediated MAPK activation;VEGFA-VEGFR2 Pathway;IRS-related events triggered by IGF1R 
                                               hsa04012;hsa04020;hsa04066;hsa04510;hsa04520;hsa05200;hsa05205;hsa05206;hsa05212;hsa05213;hsa05215;hsa05219;hsa05223;hsa05230 
                                               ERBB2 
                                               NA 
                                               azure4 
                                               1 
                                               1.19186046511628 
                                             
                                             
                                               PPP2R1A-mRNA 
                                               0.122 
                                               0.105 
                                               -0.0832 
                                               0.327 
                                               1.09 
                                               0.944 
                                               1.25 
                                               0.257 
                                               1 
                                               loglinear 
                                               ARMS-mediated activation, AXIN missense mutants destabilize the destruction complex, Beta-catenin phosphorylation cascade, CTLA4 inhibitory signaling, Cyclin AB1 associated events during G2M transition, Cyclin D associated events in G1, DAP12 signaling, DARPP-32 events, Disassembly of the destruction complex and recruitment of AXIN to the membrane, Downstream signal transduction, FCERI mediated MAPK activation, Frs2-mediated activation, Gastrin-CREB signalling pathway via PKC and MAPK, Interleukin receptor SHC signaling, IRS-related events triggered by IGF1R, MASTL Facilitates Mitotic Progression, Negative regulation of FGFR1 signaling, Negative regulation of FGFR2 signaling, Negative regulation of FGFR3 signaling, Negative regulation of FGFR4 signaling, Nonsense Mediated Decay (NMD) enhanced by the Exon Junction Complex (EJC), RAF activation, RAFMAP kinase cascade, Resolution of Sister Chromatid Cohesion, RHO GTPases Activate Formins, S33 mutants of beta-catenin aren.SQT.t phosphorylated, S37 mutants of beta-catenin aren.SQT.t phosphorylated, S45 mutants of beta-catenin aren.SQT.t phosphorylated, Signaling by ERBB2, Signaling by PDGF, Signaling by SCF-KIT, Spry regulation of FGF signaling, Truncations of AMER1 destabilize the destruction complex, VEGFA-VEGFR2 Pathway, VEGFR2 mediated cell proliferation 
                                               SEBELIUS_2_C4636 
                                               PPP2R1A 
                                               mRNA 
                                               0 
                                               
                                               
                                               MASTL Facilitates Mitotic Progression;Regulation of PLK1 Activity at G2M Transition;Disassembly of the destruction complex and recruitment of AXIN to the membrane;S37 mutants of beta-catenin aren.SQT.t phosphorylated;Cyclin AB1 associated events during G2M transition;RAF activation;RHO GTPases Activate Formins;S33 mutants of beta-catenin aren.SQT.t phosphorylated;Loss of Nlp from mitotic centrosomes;DARPP-32 events;Beta-catenin phosphorylation cascade;Initiation of Nuclear Envelope Reformation;Recruitment of mitotic centrosome proteins and complexes;Spry regulation of FGF signaling;Truncations of AMER1 destabilize the destruction complex;CTLA4 inhibitory signaling;AXIN missense mutants destabilize the destruction complex;S45 mutants of beta-catenin aren.SQT.t phosphorylated;Anchoring of the basal body to the plasma membrane;Nonsense Mediated Decay (NMD) enhanced by the Exon Junction Complex (EJC);Cyclin D associated events in G1;Platelet sensitization by LDL;Resolution of Sister Chromatid Cohesion;Inhibition of replication initiation of damaged DNA by RB1E2F1;RAFMAP kinase cascade;Downstream signal transduction;VEGFR2 mediated cell proliferation;Signaling by SCF-KIT;Frs2-mediated activation;ARMS-mediated activation;Gastrin-CREB signalling pathway via PKC and MAPK;Interleukin receptor SHC signaling;FCERI mediated MAPK activation;DAP12 signaling;Signaling by PDGF;VEGFA-VEGFR2 Pathway;Signaling by ERBB2;IRS-related events triggered by IGF1R;Negative regulation of FGFR3 signaling;Negative regulation of FGFR4 signaling;Negative regulation of FGFR2 signaling;Negative regulation of FGFR1 signaling 
                                               hsa03015;hsa04071;hsa04114;hsa04151;hsa04152;hsa04261;hsa04350;hsa04390;hsa04530;hsa04728;hsa04730;hsa05142;hsa05160 
                                               PPP2R1A 
                                               NA 
                                               azure4 
                                               1 
                                               1.16190476190476 
                                             
                                             
                                               FN1-mRNA 
                                               0.262 
                                               0.269 
                                               -0.266 
                                               0.79 
                                               1.2 
                                               0.832 
                                               1.73 
                                               0.342 
                                               1 
                                               lm.nb 
                                               ARMS-mediated activation, DAP12 signaling, Downstream signal transduction, ECM proteoglycans, Elastic fibre formation, FCERI mediated MAPK activation, Frs2-mediated activation, Gastrin-CREB signalling pathway via PKC and MAPK, Integrin alphaIIb beta3 signaling, Integrin cell surface interactions, Interleukin receptor SHC signaling, IRS-related events triggered by IGF1R, MAP2K and MAPK activation, Molecules associated with elastic fibres, Non-integrin membrane-ECM interactions, p130Cas linkage to MAPK signaling for integrins, Platelet degranulation , RAFMAP kinase cascade, Signaling by ERBB2, Signaling by PDGF, Signaling by SCF-KIT, Syndecan interactions, VEGFA-VEGFR2 Pathway, VEGFR2 mediated cell proliferation 
                                               SEBELIUS_2_C4636 
                                               FN1 
                                               mRNA 
                                               0 
                                               
                                               
                                               MAP2K and MAPK activation;p130Cas linkage to MAPK signaling for integrins;Platelet degranulation ;Molecules associated with elastic fibres;Syndecan interactions;ECM proteoglycans;Fibronectin matrix formation;Integrin cell surface interactions;RAFMAP kinase cascade;Downstream signal transduction;VEGFR2 mediated cell proliferation;Signaling by SCF-KIT;Frs2-mediated activation;ARMS-mediated activation;Gastrin-CREB signalling pathway via PKC and MAPK;Interleukin receptor SHC signaling;FCERI mediated MAPK activation;DAP12 signaling;Signaling by PDGF;VEGFA-VEGFR2 Pathway;Signaling by ERBB2;IRS-related events triggered by IGF1R;Integrin alphaIIb beta3 signaling;Non-integrin membrane-ECM interactions;Elastic fibre formation 
                                               hsa04151;hsa04510;hsa04512;hsa04810;hsa04933;hsa05100;hsa05146;hsa05200;hsa05205;hsa05222 
                                               FN1 
                                               NA 
                                               azure4 
                                               1 
                                               0.973977695167286 
                                             
                                           
                                           DE Results - ARMS-mediated activation genes - New.Annotation: miR155 vs.cel67  Table displaying each sample's global significance scores and directed global significance scores as defined in the heatmaps above. The global significance score is calculated as the square root of the mean squared t-statistic for the genes in a gene set, with t-statistics coming from the linear regression underlying our differential expression analysis. The directed global significance score is calculated as the square root of the mean signed squared t-statistic for the genes in a gene set, with t-statistics coming from the linear regression underlying our differential expression analysis.   
                                      
                                       
                                  
                                
                                
                                  
                                       
                                      
                                           Volcano Plot: New.Annotation: miR18b vs.cel67   More Plot Information                Volcano Plot: New.Annotation: miR18b vs.cel67  Volcano plot displaying each gene's -log10(p-value) and log2 fold change for the selected covariate. Highly statistically significant genes fall at the top of the plot, and highly differentially expressed genes fall to either side. Genes within the selected gene set are highlighted in orange. Horizontal lines indicate various False Discovery Rate (FDR) thresholds.   
                                           DE Results - ARMS-mediated activation genes - New.Annotation: miR18b vs.cel67   More Plot Information      Download CSV Data     
                                           
                                             
                                             
                                           
                                           
                                             
                                               Probe Label 
                                               Log2 fold change 
                                               std error (log2) 
                                               Lower confidence limit (log2) 
                                               Upper confidence limit (log2) 
                                               Linear fold change 
                                               Lower confidence limit (linear) 
                                               Upper confidence limit (linear) 
                                               P-value 
                                               BY.p.value 
                                               method 
                                               Gene.sets 
                                               Codeset.Name 
                                               Probe.Label 
                                               Analyte.Type 
                                               Is.Control 
                                               Control.Type 
                                               Related.Probes 
                                               Probe.Annotation 
                                               KEGG.Pathways 
                                               Official.Gene.Name 
                                               Control.Conc 
                                               volcanocol 
                                               volcanopch 
                                               tstats.all 
                                             
                                           
                                           
                                             
                                               RAF1-mRNA 
                                               -0.772 
                                               0.0829 
                                               -0.934 
                                               -0.609 
                                               0.586 
                                               0.523 
                                               0.656 
                                               6.7e-09 
                                               1.78e-06 
                                               loglinear 
                                               ARMS-mediated activation, CD209 (DC-SIGN) signaling, CREB phosphorylation through the activation of Ras, DAP12 signaling, Downstream signal transduction, FCERI mediated MAPK activation, Frs2-mediated activation, Gastrin-CREB signalling pathway via PKC and MAPK, Interleukin receptor SHC signaling, IRS-related events triggered by IGF1R, Negative feedback regulation of MAPK pathway, Negative regulation of MAPK pathway, RAF activation, RAFMAP kinase cascade, Signaling by ERBB2, Signaling by PDGF, Signaling by SCF-KIT, VEGFA-VEGFR2 Pathway, VEGFR2 mediated cell proliferation 
                                               SEBELIUS_2_C4636 
                                               RAF1 
                                               mRNA 
                                               0 
                                               
                                               
                                               RAF activation;Stimuli-sensing channels;CREB phosphorylation through the activation of Ras;Negative feedback regulation of MAPK pathway;CD209 (DC-SIGN) signaling;RAFMAP kinase cascade;Downstream signal transduction;VEGFR2 mediated cell proliferation;Signaling by SCF-KIT;Frs2-mediated activation;ARMS-mediated activation;Gastrin-CREB signalling pathway via PKC and MAPK;Interleukin receptor SHC signaling;FCERI mediated MAPK activation;DAP12 signaling;Signaling by PDGF;VEGFA-VEGFR2 Pathway;Signaling by ERBB2;IRS-related events triggered by IGF1R;Negative regulation of MAPK pathway 
                                               hsa04010;hsa04012;hsa04014;hsa04015;hsa04022;hsa04024;hsa04062;hsa04068;hsa04071;hsa04072;hsa04151;hsa04270;hsa04370;hsa04510;hsa04540;hsa04550;hsa04630;hsa04650;hsa04660;hsa04662;hsa04664;hsa04666;hsa04720;hsa04722;hsa04726;hsa04730;hsa04810;hsa04910;hsa04912;hsa04914;hsa04915;hsa04916;hsa04917;hsa04919;hsa04921;hsa05034;hsa05152;hsa05160;hsa05161;hsa05164;hsa05200;hsa05205;hsa05206;hsa05210;hsa05211;hsa05212;hsa05213;hsa05214;hsa05215;hsa05218;hsa05219;hsa05220;hsa05221;hsa05223;hsa05230;hsa05231 
                                               RAF1 
                                               NA 
                                               slateblue1 
                                               16 
                                               -9.3124246079614 
                                             
                                             
                                               MAPK1-mRNA 
                                               -0.571 
                                               0.0726 
                                               -0.713 
                                               -0.429 
                                               0.673 
                                               0.61 
                                               0.743 
                                               1.09e-07 
                                               1.81e-05 
                                               loglinear 
                                               ARMS-mediated activation, CREB phosphorylation through the activation of Ras, DAP12 signaling, Downstream signal transduction, FCERI mediated MAPK activation, Frs2-mediated activation, Gastrin-CREB signalling pathway via PKC and MAPK, Interleukin receptor SHC signaling, IRS-related events triggered by IGF1R, MAP2K and MAPK activation, Negative feedback regulation of MAPK pathway, Negative regulation of FGFR1 signaling, Negative regulation of FGFR2 signaling, Negative regulation of FGFR3 signaling, Negative regulation of FGFR4 signaling, Negative regulation of MAPK pathway, RAFMAP kinase cascade, Regulation of HSF1-mediated heat shock response, RHO GTPases Activate WASPs and WAVEs, Signal transduction by L1, Signaling by ERBB2, Signaling by PDGF, Signaling by SCF-KIT, VEGFA-VEGFR2 Pathway, VEGFR2 mediated cell proliferation 
                                               SEBELIUS_2_C4636 
                                               MAPK1 
                                               mRNA 
                                               0 
                                               
                                               
                                               Negative regulation of FGFR1 signaling;Signal transduction by L1;Negative regulation of FGFR2 signaling;MAP2K and MAPK activation;RHO GTPases Activate WASPs and WAVEs;Senescence-Associated Secretory Phenotype (SASP);Regulation of HSF1-mediated heat shock response;Thrombin signalling through proteinase activated receptors (PARs);Negative regulation of FGFR4 signaling;Oxidative Stress Induced Senescence;Signal attenuation;Negative regulation of FGFR3 signaling;Negative feedback regulation of MAPK pathway;Growth hormone receptor signaling;PLC beta mediated events;FCERI mediated MAPK activation;CREB phosphorylation through the activation of Ras;Negative regulation of MAPK pathway;RAFMAP kinase cascade;Downstream signal transduction;VEGFR2 mediated cell proliferation;Signaling by SCF-KIT;Frs2-mediated activation;ARMS-mediated activation;Gastrin-CREB signalling pathway via PKC and MAPK;Interleukin receptor SHC signaling;DAP12 signaling;Signaling by PDGF;VEGFA-VEGFR2 Pathway;Signaling by ERBB2;IRS-related events triggered by IGF1R 
                                               hsa04010;hsa04012;hsa04014;hsa04015;hsa04022;hsa04024;hsa04062;hsa04066;hsa04068;hsa04071;hsa04072;hsa04114;hsa04150;hsa04151;hsa04261;hsa04270;hsa04320;hsa04350;hsa04360;hsa04370;hsa04380;hsa04510;hsa04520;hsa04540;hsa04550;hsa04611;hsa04620;hsa04621;hsa04650;hsa04660;hsa04662;hsa04664;hsa04666;hsa04668;hsa04713;hsa04720;hsa04722;hsa04723;hsa04724;hsa04725;hsa04726;hsa04730;hsa04810;hsa04910;hsa04912;hsa04914;hsa04915;hsa04916;hsa04917;hsa04919;hsa04921;hsa04930;hsa04933;hsa04960;hsa05010;hsa05020;hsa05034;hsa05131;hsa05132;hsa05133;hsa05140;hsa05142;hsa05145;hsa05152;hsa05160;hsa05161;hsa05164;hsa05200;hsa05203;hsa05205;hsa05206;hsa05210;hsa05211;hsa05212;hsa05213;hsa05214;hsa05215;hsa05216;hsa05218;hsa05219;hsa05220;hsa05221;hsa05223;hsa05230;hsa05231 
                                               MAPK1 
                                               NA 
                                               slateblue1 
                                               16 
                                               -7.86501377410468 
                                             
                                             
                                               MAPK3-mRNA 
                                               0.442 
                                               0.0615 
                                               0.321 
                                               0.563 
                                               1.36 
                                               1.25 
                                               1.48 
                                               4.43e-07 
                                               4.91e-05 
                                               loglinear 
                                               ARMS-mediated activation, DAP12 signaling, Downstream signal transduction, ERKs are inactivated, FCERI mediated MAPK activation, Frs2-mediated activation, Gastrin-CREB signalling pathway via PKC and MAPK, Interleukin receptor SHC signaling, IRS-related events triggered by IGF1R, Negative feedback regulation of MAPK pathway, Negative regulation of FGFR3 signaling, Negative regulation of FGFR4 signaling, Negative regulation of MAPK pathway, RAFMAP kinase cascade, Regulation of HSF1-mediated heat shock response, RHO GTPases Activate WASPs and WAVEs, Signaling by ERBB2, Signaling by PDGF, Signaling by SCF-KIT, VEGFA-VEGFR2 Pathway, VEGFR2 mediated cell proliferation 
                                               SEBELIUS_2_C4636 
                                               MAPK3 
                                               mRNA 
                                               0 
                                               
                                               
                                               ERKs are inactivated;ISG15 antiviral mechanism;RHO GTPases Activate WASPs and WAVEs;Senescence-Associated Secretory Phenotype (SASP);Regulation of HSF1-mediated heat shock response;Thrombin signalling through proteinase activated receptors (PARs);Negative regulation of FGFR4 signaling;Oxidative Stress Induced Senescence;Signal attenuation;RNA Polymerase I Promoter Opening;Negative regulation of FGFR3 signaling;Negative feedback regulation of MAPK pathway;Growth hormone receptor signaling;Golgi Cisternae Pericentriolar Stack Reorganization;FCERI mediated MAPK activation;Negative regulation of MAPK pathway;RAFMAP kinase cascade;Downstream signal transduction;VEGFR2 mediated cell proliferation;Signaling by SCF-KIT;Frs2-mediated activation;ARMS-mediated activation;Gastrin-CREB signalling pathway via PKC and MAPK;Interleukin receptor SHC signaling;DAP12 signaling;Signaling by PDGF;VEGFA-VEGFR2 Pathway;Signaling by ERBB2;IRS-related events triggered by IGF1R 
                                               hsa04010;hsa04012;hsa04014;hsa04015;hsa04022;hsa04024;hsa04062;hsa04066;hsa04068;hsa04071;hsa04072;hsa04114;hsa04150;hsa04151;hsa04261;hsa04270;hsa04320;hsa04350;hsa04360;hsa04370;hsa04380;hsa04510;hsa04520;hsa04540;hsa04550;hsa04611;hsa04620;hsa04621;hsa04650;hsa04660;hsa04662;hsa04664;hsa04666;hsa04668;hsa04713;hsa04720;hsa04722;hsa04723;hsa04724;hsa04725;hsa04726;hsa04730;hsa04810;hsa04910;hsa04912;hsa04914;hsa04915;hsa04916;hsa04917;hsa04919;hsa04921;hsa04930;hsa04933;hsa04960;hsa05010;hsa05020;hsa05034;hsa05131;hsa05132;hsa05133;hsa05140;hsa05142;hsa05145;hsa05152;hsa05160;hsa05161;hsa05164;hsa05200;hsa05203;hsa05205;hsa05210;hsa05211;hsa05212;hsa05213;hsa05214;hsa05215;hsa05216;hsa05218;hsa05219;hsa05220;hsa05221;hsa05223;hsa05230;hsa05231 
                                               MAPK3 
                                               NA 
                                               slateblue1 
                                               16 
                                               7.1869918699187 
                                             
                                             
                                               NRAS-mRNA 
                                               -0.806 
                                               0.13 
                                               -1.06 
                                               -0.551 
                                               0.572 
                                               0.479 
                                               0.683 
                                               3.82e-06 
                                               0.000286 
                                               lm.nb 
                                               ARMS-mediated activation, CD209 (DC-SIGN) signaling, Cell surface interactions at the vascular wall, Constitutive Signaling by EGFRvIII, DAP12 signaling, Downstream signal transduction, FCERI mediated MAPK activation, Frs2-mediated activation, Gastrin-CREB signalling pathway via PKC and MAPK, Interleukin receptor SHC signaling, IRS-related events triggered by IGF1R, MAP2K and MAPK activation, RAFMAP kinase cascade, SHC-mediated cascade.CL.FGFR1, SHC-related events triggered by IGF1R, Signaling by ERBB2, Signaling by FGFR1 in disease, Signaling by FGFR2 in disease, Signaling by FGFR3 in disease, Signaling by FGFR4 in disease, Signaling by PDGF, Signaling by SCF-KIT, Tie2 Signaling, VEGFA-VEGFR2 Pathway, VEGFR2 mediated cell proliferation 
                                               SEBELIUS_2_C4636 
                                               NRAS 
                                               mRNA 
                                               0 
                                               
                                               
                                               Signaling by FGFR2 in disease;Regulation of RAS by GAPs;SHC-mediated cascade.CL.FGFR3;Signaling by FGFR3 in disease;MAP2K and MAPK activation;SHC-related events triggered by IGF1R;Signaling by FGFR4 in disease;SHC-mediated cascade.CL.FGFR1;Tie2 Signaling;Signaling by FGFR1 in disease;Activation of RAS in B cells;SHC-mediated cascade.CL.FGFR4;Constitutive Signaling by EGFRvIII;CD209 (DC-SIGN) signaling;FCERI mediated MAPK activation;RAFMAP kinase cascade;Downstream signal transduction;VEGFR2 mediated cell proliferation;Signaling by SCF-KIT;Frs2-mediated activation;ARMS-mediated activation;Gastrin-CREB signalling pathway via PKC and MAPK;Interleukin receptor SHC signaling;DAP12 signaling;Signaling by PDGF;VEGFA-VEGFR2 Pathway;Signaling by ERBB2;IRS-related events triggered by IGF1R;Cell surface interactions at the vascular wall 
                                               hsa04010;hsa04012;hsa04014;hsa04015;hsa04062;hsa04068;hsa04071;hsa04072;hsa04151;hsa04211;hsa04213;hsa04360;hsa04370;hsa04530;hsa04540;hsa04550;hsa04650;hsa04660;hsa04662;hsa04664;hsa04720;hsa04722;hsa04725;hsa04726;hsa04730;hsa04810;hsa04910;hsa04912;hsa04915;hsa04916;hsa04917;hsa04919;hsa04921;hsa04933;hsa05034;hsa05160;hsa05161;hsa05166;hsa05200;hsa05203;hsa05205;hsa05206;hsa05211;hsa05213;hsa05214;hsa05215;hsa05216;hsa05218;hsa05219;hsa05220;hsa05221;hsa05223;hsa05230;hsa05231 
                                               NRAS 
                                               NA 
                                               slateblue1 
                                               16 
                                               -6.2 
                                             
                                             
                                               PPP2R1B-mRNA 
                                               0.385 
                                               0.0675 
                                               0.253 
                                               0.518 
                                               1.31 
                                               1.19 
                                               1.43 
                                               1.15e-05 
                                               0.000638 
                                               loglinear 
                                               ARMS-mediated activation, AXIN missense mutants destabilize the destruction complex, Beta-catenin phosphorylation cascade, CTLA4 inhibitory signaling, Cyclin AB1 associated events during G2M transition, Cyclin D associated events in G1, DAP12 signaling, DARPP-32 events, Disassembly of the destruction complex and recruitment of AXIN to the membrane, Downstream signal transduction, ERKs are inactivated, FCERI mediated MAPK activation, Frs2-mediated activation, Gastrin-CREB signalling pathway via PKC and MAPK, Interleukin receptor SHC signaling, IRS-related events triggered by IGF1R, MASTL Facilitates Mitotic Progression, Misspliced GSK3beta mutants stabilize beta-catenin, Negative regulation of MAPK pathway, RAF activation, RAFMAP kinase cascade, Resolution of Sister Chromatid Cohesion, RHO GTPases Activate Formins, S33 mutants of beta-catenin aren.SQT.t phosphorylated, S37 mutants of beta-catenin aren.SQT.t phosphorylated, S45 mutants of beta-catenin aren.SQT.t phosphorylated, Signaling by ERBB2, Signaling by PDGF, Signaling by SCF-KIT, T41 mutants of beta-catenin aren.SQT.t phosphorylated, Truncations of AMER1 destabilize the destruction complex, VEGFA-VEGFR2 Pathway, VEGFR2 mediated cell proliferation 
                                               SEBELIUS_2_C4636 
                                               PPP2R1B 
                                               mRNA 
                                               0 
                                               
                                               
                                               MASTL Facilitates Mitotic Progression;Misspliced GSK3beta mutants stabilize beta-catenin;T41 mutants of beta-catenin aren.SQT.t phosphorylated;ERKs are inactivated;Glycolysis;Negative regulation of MAPK pathway;Disassembly of the destruction complex and recruitment of AXIN to the membrane;S37 mutants of beta-catenin aren.SQT.t phosphorylated;Cyclin AB1 associated events during G2M transition;RAF activation;RHO GTPases Activate Formins;S33 mutants of beta-catenin aren.SQT.t phosphorylated;DARPP-32 events;Beta-catenin phosphorylation cascade;Truncations of AMER1 destabilize the destruction complex;CTLA4 inhibitory signaling;AXIN missense mutants destabilize the destruction complex;S45 mutants of beta-catenin aren.SQT.t phosphorylated;Cyclin D associated events in G1;Platelet sensitization by LDL;Resolution of Sister Chromatid Cohesion;Inhibition of replication initiation of damaged DNA by RB1E2F1;RAFMAP kinase cascade;Downstream signal transduction;VEGFR2 mediated cell proliferation;Signaling by SCF-KIT;Frs2-mediated activation;ARMS-mediated activation;Gastrin-CREB signalling pathway via PKC and MAPK;Interleukin receptor SHC signaling;FCERI mediated MAPK activation;DAP12 signaling;Signaling by PDGF;VEGFA-VEGFR2 Pathway;Signaling by ERBB2;IRS-related events triggered by IGF1R 
                                               hsa03015;hsa04071;hsa04114;hsa04151;hsa04152;hsa04261;hsa04350;hsa04390;hsa04530;hsa04728;hsa04730;hsa05142;hsa05160 
                                               PPP2R1B 
                                               NA 
                                               slateblue1 
                                               16 
                                               5.7037037037037 
                                             
                                             
                                               IQGAP1-mRNA 
                                               -0.62 
                                               0.124 
                                               -0.863 
                                               -0.378 
                                               0.651 
                                               0.55 
                                               0.77 
                                               5.8e-05 
                                               0.00227 
                                               loglinear 
                                               ARMS-mediated activation, DAP12 signaling, Downstream signal transduction, FCERI mediated MAPK activation, Frs2-mediated activation, Gastrin-CREB signalling pathway via PKC and MAPK, Interleukin receptor SHC signaling, IRS-related events triggered by IGF1R, MAP2K and MAPK activation, Nephrin interactions, RAFMAP kinase cascade, RHO GTPases activate IQGAPs, Signaling by ERBB2, Signaling by PDGF, Signaling by SCF-KIT, VEGFA-VEGFR2 Pathway, VEGFR2 mediated cell proliferation 
                                               SEBELIUS_2_C4636 
                                               IQGAP1 
                                               mRNA 
                                               0 
                                               
                                               
                                               MAP2K and MAPK activation;Nephrin interactions;RHO GTPases activate IQGAPs;RAFMAP kinase cascade;Downstream signal transduction;VEGFR2 mediated cell proliferation;Signaling by SCF-KIT;Frs2-mediated activation;ARMS-mediated activation;Gastrin-CREB signalling pathway via PKC and MAPK;Interleukin receptor SHC signaling;FCERI mediated MAPK activation;DAP12 signaling;Signaling by PDGF;VEGFA-VEGFR2 Pathway;Signaling by ERBB2;IRS-related events triggered by IGF1R 
                                               hsa04520;hsa04810;hsa05205 
                                               IQGAP1 
                                               NA 
                                               slateblue1 
                                               16 
                                               -5 
                                             
                                             
                                               FGFR1-mRNA 
                                               -0.475 
                                               0.116 
                                               -0.703 
                                               -0.247 
                                               0.719 
                                               0.614 
                                               0.842 
                                               0.000528 
                                               0.0152 
                                               lm.nb 
                                               ARMS-mediated activation, Constitutive Signaling by Aberrant PI3K in Cancer, DAP12 signaling, Downstream signal transduction, FCERI mediated MAPK activation, Frs2-mediated activation, GAB1 signalosome, Gastrin-CREB signalling pathway via PKC and MAPK, Interleukin receptor SHC signaling, IRS-related events triggered by IGF1R, PI3K Cascade, PI3KAKT activation, RAFMAP kinase cascade, Role of LAT2NTALLAB on calcium mobilization, SHC-mediated cascade.CL.FGFR1, Signaling by ERBB2, Signaling by FGFR1 in disease, Signaling by PDGF, Signaling by SCF-KIT, VEGFA-VEGFR2 Pathway, VEGFR2 mediated cell proliferation 
                                               SEBELIUS_2_C4636 
                                               FGFR1 
                                               mRNA 
                                               0 
                                               
                                               
                                               RAFMAP kinase cascade;Constitutive Signaling by Aberrant PI3K in Cancer;SHC-mediated cascade.CL.FGFR1;Phospholipase C-mediated cascade.CL. FGFR1;Signaling by FGFR1 amplification mutants;PI3K Cascade;FGFR1b ligand binding and activation;Signaling by FGFR1 in disease;Downstream signal transduction;GAB1 signalosome;Signaling by SCF-KIT;PI3KAKT activation;Role of LAT2NTALLAB on calcium mobilization;DAP12 signaling;Signaling by PDGF;Signaling by ERBB2;VEGFR2 mediated cell proliferation;Frs2-mediated activation;ARMS-mediated activation;Gastrin-CREB signalling pathway via PKC and MAPK;Interleukin receptor SHC signaling;FCERI mediated MAPK activation;VEGFA-VEGFR2 Pathway;IRS-related events triggered by IGF1R 
                                               hsa04010;hsa04014;hsa04015;hsa04151;hsa04520;hsa04550;hsa04810;hsa05200;hsa05205;hsa05215;hsa05218;hsa05230 
                                               FGFR1 
                                               NA 
                                               slateblue2 
                                               16 
                                               -4.0948275862069 
                                             
                                             
                                               BRAF-mRNA 
                                               -0.255 
                                               0.0684 
                                               -0.389 
                                               -0.121 
                                               0.838 
                                               0.764 
                                               0.92 
                                               0.00126 
                                               0.0304 
                                               loglinear 
                                               ARMS-mediated activation, CREB phosphorylation through the activation of Ras, DAP12 signaling, Downstream signal transduction, FCERI mediated MAPK activation, Frs2-mediated activation, Gastrin-CREB signalling pathway via PKC and MAPK, Interleukin receptor SHC signaling, IRS-related events triggered by IGF1R, MAP2K and MAPK activation, Negative feedback regulation of MAPK pathway, Negative regulation of FGFR1 signaling, Negative regulation of FGFR2 signaling, Negative regulation of FGFR3 signaling, Negative regulation of FGFR4 signaling, Negative regulation of MAPK pathway, RAF activation, RAFMAP kinase cascade, Signaling by ERBB2, Signaling by PDGF, Signaling by SCF-KIT, Spry regulation of FGF signaling, VEGFA-VEGFR2 Pathway, VEGFR2 mediated cell proliferation 
                                               SEBELIUS_2_C4636 
                                               BRAF 
                                               mRNA 
                                               0 
                                               
                                               
                                               Spry regulation of FGF signaling;MAP2K and MAPK activation;RAF activation;CREB phosphorylation through the activation of Ras;Negative feedback regulation of MAPK pathway;ARMS-mediated activation;Negative regulation of FGFR3 signaling;Negative regulation of FGFR4 signaling;Negative regulation of FGFR2 signaling;Negative regulation of FGFR1 signaling;RAFMAP kinase cascade;Downstream signal transduction;VEGFR2 mediated cell proliferation;Signaling by SCF-KIT;Frs2-mediated activation;Gastrin-CREB signalling pathway via PKC and MAPK;Interleukin receptor SHC signaling;FCERI mediated MAPK activation;DAP12 signaling;Signaling by PDGF;VEGFA-VEGFR2 Pathway;Signaling by ERBB2;IRS-related events triggered by IGF1R;Negative regulation of MAPK pathway 
                                               hsa04010;hsa04012;hsa04015;hsa04024;hsa04062;hsa04068;hsa04150;hsa04270;hsa04510;hsa04650;hsa04720;hsa04722;hsa04726;hsa04730;hsa04810;hsa04910;hsa04914;hsa05034;hsa05160;hsa05200;hsa05205;hsa05210;hsa05211;hsa05212;hsa05213;hsa05214;hsa05215;hsa05216;hsa05218;hsa05219;hsa05220;hsa05221;hsa05223 
                                               BRAF 
                                               NA 
                                               slateblue2 
                                               16 
                                               -3.7280701754386 
                                             
                                             
                                               PPP2CB-mRNA 
                                               0.253 
                                               0.085 
                                               0.0862 
                                               0.419 
                                               1.19 
                                               1.06 
                                               1.34 
                                               0.00724 
                                               0.148 
                                               loglinear 
                                               ARMS-mediated activation, AXIN missense mutants destabilize the destruction complex, Beta-catenin phosphorylation cascade, CTLA4 inhibitory signaling, Cyclin AB1 associated events during G2M transition, Cyclin D associated events in G1, DAP12 signaling, DARPP-32 events, Disassembly of the destruction complex and recruitment of AXIN to the membrane, Downstream signal transduction, ERKs are inactivated, FCERI mediated MAPK activation, Frs2-mediated activation, Gastrin-CREB signalling pathway via PKC and MAPK, Interleukin receptor SHC signaling, IRS-related events triggered by IGF1R, MASTL Facilitates Mitotic Progression, Misspliced GSK3beta mutants stabilize beta-catenin, Negative regulation of FGFR1 signaling, Negative regulation of FGFR2 signaling, Negative regulation of FGFR3 signaling, Negative regulation of FGFR4 signaling, Negative regulation of MAPK pathway, RAF activation, RAFMAP kinase cascade, Resolution of Sister Chromatid Cohesion, RHO GTPases Activate Formins, S33 mutants of beta-catenin aren.SQT.t phosphorylated, S37 mutants of beta-catenin aren.SQT.t phosphorylated, S45 mutants of beta-catenin aren.SQT.t phosphorylated, Signaling by ERBB2, Signaling by PDGF, Signaling by SCF-KIT, Spry regulation of FGF signaling, T41 mutants of beta-catenin aren.SQT.t phosphorylated, Truncations of AMER1 destabilize the destruction complex, VEGFA-VEGFR2 Pathway, VEGFR2 mediated cell proliferation 
                                               SEBELIUS_2_C4636 
                                               PPP2CB 
                                               mRNA 
                                               0 
                                               
                                               
                                               MASTL Facilitates Mitotic Progression;Misspliced GSK3beta mutants stabilize beta-catenin;T41 mutants of beta-catenin aren.SQT.t phosphorylated;ERKs are inactivated;Glycolysis;Negative regulation of MAPK pathway;Disassembly of the destruction complex and recruitment of AXIN to the membrane;S37 mutants of beta-catenin aren.SQT.t phosphorylated;Cyclin AB1 associated events during G2M transition;RAF activation;RHO GTPases Activate Formins;S33 mutants of beta-catenin aren.SQT.t phosphorylated;DARPP-32 events;Beta-catenin phosphorylation cascade;Spry regulation of FGF signaling;Truncations of AMER1 destabilize the destruction complex;CTLA4 inhibitory signaling;AXIN missense mutants destabilize the destruction complex;S45 mutants of beta-catenin aren.SQT.t phosphorylated;Cyclin D associated events in G1;Platelet sensitization by LDL;Resolution of Sister Chromatid Cohesion;Inhibition of replication initiation of damaged DNA by RB1E2F1;RAFMAP kinase cascade;Downstream signal transduction;VEGFR2 mediated cell proliferation;Signaling by SCF-KIT;Frs2-mediated activation;ARMS-mediated activation;Gastrin-CREB signalling pathway via PKC and MAPK;Interleukin receptor SHC signaling;FCERI mediated MAPK activation;DAP12 signaling;Signaling by PDGF;VEGFA-VEGFR2 Pathway;Signaling by ERBB2;IRS-related events triggered by IGF1R;Negative regulation of FGFR3 signaling;Negative regulation of FGFR4 signaling;Negative regulation of FGFR2 signaling;Negative regulation of FGFR1 signaling 
                                               hsa03015;hsa04071;hsa04114;hsa04151;hsa04152;hsa04261;hsa04350;hsa04390;hsa04530;hsa04728;hsa04730;hsa05142;hsa05160 
                                               PPP2CB 
                                               NA 
                                               slateblue4 
                                               16 
                                               2.97647058823529 
                                             
                                             
                                               EGFR-mRNA 
                                               -0.331 
                                               0.115 
                                               -0.555 
                                               -0.106 
                                               0.795 
                                               0.681 
                                               0.929 
                                               0.0088 
                                               0.172 
                                               loglinear 
                                               ARMS-mediated activation, Constitutive Signaling by Aberrant PI3K in Cancer, Constitutive Signaling by EGFRvIII, DAP12 signaling, Downstream signal transduction, FCERI mediated MAPK activation, Frs2-mediated activation, GAB1 signalosome, Gastrin-CREB signalling pathway via PKC and MAPK, Interleukin receptor SHC signaling, IRS-related events triggered by IGF1R, PI3KAKT activation, Role of LAT2NTALLAB on calcium mobilization, Signaling by ERBB2, Signaling by PDGF, Signaling by SCF-KIT, VEGFA-VEGFR2 Pathway, VEGFR2 mediated cell proliferation 
                                               SEBELIUS_2_C4636 
                                               EGFR 
                                               mRNA 
                                               0 
                                               
                                               
                                               Constitutive Signaling by Aberrant PI3K in Cancer;Inhibition of Signaling by Overexpressed EGFR;PLCG1 events in ERBB2 signaling;EGFR interacts with phospholipase C-gamma;EGFR Transactivation by Gastrin;Constitutive Signaling by Ligand-Responsive EGFR Cancer Variants;Constitutive Signaling by EGFRvIII;GAB1 signalosome;Downstream signal transduction;Signaling by SCF-KIT;PI3KAKT activation;Role of LAT2NTALLAB on calcium mobilization;DAP12 signaling;Signaling by PDGF;Signaling by ERBB2;VEGFR2 mediated cell proliferation;Frs2-mediated activation;ARMS-mediated activation;Gastrin-CREB signalling pathway via PKC and MAPK;Interleukin receptor SHC signaling;FCERI mediated MAPK activation;VEGFA-VEGFR2 Pathway;IRS-related events triggered by IGF1R 
                                               hsa04010;hsa04012;hsa04014;hsa04015;hsa04020;hsa04060;hsa04066;hsa04068;hsa04072;hsa04144;hsa04151;hsa04320;hsa04510;hsa04520;hsa04540;hsa04810;hsa04912;hsa04915;hsa04921;hsa05120;hsa05160;hsa05200;hsa05205;hsa05206;hsa05212;hsa05213;hsa05214;hsa05215;hsa05218;hsa05219;hsa05223;hsa05230;hsa05231 
                                               EGFR 
                                               NA 
                                               slateblue4 
                                               16 
                                               -2.87826086956522 
                                             
                                             
                                               PPP2CA-mRNA 
                                               -0.173 
                                               0.06 
                                               -0.29 
                                               -0.0549 
                                               0.887 
                                               0.818 
                                               0.963 
                                               0.00906 
                                               0.174 
                                               loglinear 
                                               ARMS-mediated activation, AXIN missense mutants destabilize the destruction complex, Beta-catenin phosphorylation cascade, CTLA4 inhibitory signaling, Cyclin AB1 associated events during G2M transition, Cyclin D associated events in G1, DAP12 signaling, DARPP-32 events, Disassembly of the destruction complex and recruitment of AXIN to the membrane, Downstream signal transduction, ERKs are inactivated, FCERI mediated MAPK activation, Frs2-mediated activation, Gastrin-CREB signalling pathway via PKC and MAPK, Interleukin receptor SHC signaling, IRS-related events triggered by IGF1R, MASTL Facilitates Mitotic Progression, Negative regulation of FGFR1 signaling, Negative regulation of FGFR2 signaling, Negative regulation of FGFR3 signaling, Negative regulation of FGFR4 signaling, Negative regulation of MAPK pathway, Nonsense Mediated Decay (NMD) enhanced by the Exon Junction Complex (EJC), RAF activation, RAFMAP kinase cascade, Resolution of Sister Chromatid Cohesion, RHO GTPases Activate Formins, S33 mutants of beta-catenin aren.SQT.t phosphorylated, S37 mutants of beta-catenin aren.SQT.t phosphorylated, S45 mutants of beta-catenin aren.SQT.t phosphorylated, Signaling by ERBB2, Signaling by PDGF, Signaling by SCF-KIT, Spry regulation of FGF signaling, T41 mutants of beta-catenin aren.SQT.t phosphorylated, Truncations of AMER1 destabilize the destruction complex, VEGFA-VEGFR2 Pathway, VEGFR2 mediated cell proliferation 
                                               SEBELIUS_2_C4636 
                                               PPP2CA 
                                               mRNA 
                                               0 
                                               
                                               
                                               MASTL Facilitates Mitotic Progression;T41 mutants of beta-catenin aren.SQT.t phosphorylated;ERKs are inactivated;Glycolysis;Negative regulation of MAPK pathway;Disassembly of the destruction complex and recruitment of AXIN to the membrane;S37 mutants of beta-catenin aren.SQT.t phosphorylated;Cyclin AB1 associated events during G2M transition;RAF activation;RHO GTPases Activate Formins;S33 mutants of beta-catenin aren.SQT.t phosphorylated;DARPP-32 events;Beta-catenin phosphorylation cascade;Initiation of Nuclear Envelope Reformation;Spry regulation of FGF signaling;Truncations of AMER1 destabilize the destruction complex;CTLA4 inhibitory signaling;AXIN missense mutants destabilize the destruction complex;S45 mutants of beta-catenin aren.SQT.t phosphorylated;Nonsense Mediated Decay (NMD) enhanced by the Exon Junction Complex (EJC);Cyclin D associated events in G1;Platelet sensitization by LDL;Resolution of Sister Chromatid Cohesion;Inhibition of replication initiation of damaged DNA by RB1E2F1;RAFMAP kinase cascade;Downstream signal transduction;VEGFR2 mediated cell proliferation;Signaling by SCF-KIT;Frs2-mediated activation;ARMS-mediated activation;Gastrin-CREB signalling pathway via PKC and MAPK;Interleukin receptor SHC signaling;FCERI mediated MAPK activation;DAP12 signaling;Signaling by PDGF;VEGFA-VEGFR2 Pathway;Signaling by ERBB2;IRS-related events triggered by IGF1R;Negative regulation of FGFR3 signaling;Negative regulation of FGFR4 signaling;Negative regulation of FGFR2 signaling;Negative regulation of FGFR1 signaling 
                                               hsa03015;hsa04071;hsa04114;hsa04151;hsa04152;hsa04261;hsa04350;hsa04390;hsa04530;hsa04728;hsa04730;hsa05142;hsa05160 
                                               PPP2CA 
                                               NA 
                                               slateblue4 
                                               16 
                                               -2.88333333333333 
                                             
                                             
                                               SRC-mRNA 
                                               0.369 
                                               0.13 
                                               0.114 
                                               0.624 
                                               1.29 
                                               1.08 
                                               1.54 
                                               0.00983 
                                               0.184 
                                               loglinear 
                                               ARMS-mediated activation, CD28 co-stimulation, Cell surface interactions at the vascular wall, CTLA4 inhibitory signaling, DAP12 signaling, Downstream signal transduction, FCERI mediated MAPK activation, Frs2-mediated activation, GAB1 signalosome, Gastrin-CREB signalling pathway via PKC and MAPK, Integrin alphaIIb beta3 signaling, Interleukin receptor SHC signaling, IRS-related events triggered by IGF1R, MAP2K and MAPK activation, Negative regulation of FGFR1 signaling, Negative regulation of FGFR2 signaling, Negative regulation of FGFR3 signaling, Negative regulation of FGFR4 signaling, p130Cas linkage to MAPK signaling for integrins, RAFMAP kinase cascade, Signaling by ERBB2, Signaling by PDGF, Signaling by SCF-KIT, Spry regulation of FGF signaling, VEGFA-VEGFR2 Pathway, VEGFR2 mediated cell proliferation 
                                               SEBELIUS_2_C4636 
                                               SRC 
                                               mRNA 
                                               0 
                                               
                                               
                                               FCGR activation;Spry regulation of FGF signaling;GAB1 signalosome;MAP2K and MAPK activation;CLEC7A (Dectin-1) signaling;CD28 co-stimulation;p130Cas linkage to MAPK signaling for integrins;CTLA4 inhibitory signaling;GRB2.CL.SOS provides linkage to MAPK signaling for Integrins ;Signal regulatory protein (SIRP) family interactions;PECAM1 interactions;Thrombin signalling through proteinase activated receptors (PARs);GP1b-IX-V activation signalling;c-src mediated regulation of Cx43 function and closure of gap junctions;Signaling by ERBB2;VEGFA-VEGFR2 Pathway;Signaling by SCF-KIT;VEGFR2 mediated cell proliferation;RAFMAP kinase cascade;Downstream signal transduction;Frs2-mediated activation;ARMS-mediated activation;Gastrin-CREB signalling pathway via PKC and MAPK;Interleukin receptor SHC signaling;FCERI mediated MAPK activation;DAP12 signaling;Signaling by PDGF;IRS-related events triggered by IGF1R;Negative regulation of FGFR3 signaling;Negative regulation of FGFR4 signaling;Negative regulation of FGFR2 signaling;Negative regulation of FGFR1 signaling;Integrin alphaIIb beta3 signaling;Cell surface interactions at the vascular wall 
                                               hsa04012;hsa04015;hsa04062;hsa04144;hsa04370;hsa04510;hsa04520;hsa04530;hsa04540;hsa04611;hsa04727;hsa04750;hsa04810;hsa04912;hsa04915;hsa04917;hsa04919;hsa04921;hsa05100;hsa05120;hsa05131;hsa05152;hsa05161;hsa05203;hsa05205;hsa05219 
                                               SRC 
                                               NA 
                                               slateblue4 
                                               16 
                                               2.83846153846154 
                                             
                                             
                                               KRAS-mRNA 
                                               0.31 
                                               0.115 
                                               0.0834 
                                               0.536 
                                               1.24 
                                               1.06 
                                               1.45 
                                               0.0139 
                                               0.243 
                                               loglinear 
                                               ARMS-mediated activation, CD209 (DC-SIGN) signaling, Cell surface interactions at the vascular wall, Constitutive Signaling by EGFRvIII, DAP12 signaling, Downstream signal transduction, FCERI mediated MAPK activation, Frs2-mediated activation, Gastrin-CREB signalling pathway via PKC and MAPK, Interleukin receptor SHC signaling, IRS-related events triggered by IGF1R, MAP2K and MAPK activation, RAFMAP kinase cascade, SHC-mediated cascade.CL.FGFR1, SHC-related events triggered by IGF1R, Signaling by ERBB2, Signaling by FGFR1 in disease, Signaling by FGFR2 in disease, Signaling by FGFR3 in disease, Signaling by FGFR4 in disease, Signaling by PDGF, Signaling by SCF-KIT, Tie2 Signaling, VEGFA-VEGFR2 Pathway, VEGFR2 mediated cell proliferation 
                                               SEBELIUS_2_C4636 
                                               KRAS 
                                               mRNA 
                                               0 
                                               
                                               
                                               Signaling by FGFR2 in disease;Regulation of RAS by GAPs;SHC-mediated cascade.CL.FGFR3;Signaling by FGFR3 in disease;MAP2K and MAPK activation;SHC-related events triggered by IGF1R;Signaling by FGFR4 in disease;SHC-mediated cascade.CL.FGFR1;Tie2 Signaling;Signaling by FGFR1 in disease;Activation of RAS in B cells;SHC-mediated cascade.CL.FGFR4;Constitutive Signaling by EGFRvIII;CD209 (DC-SIGN) signaling;FCERI mediated MAPK activation;RAFMAP kinase cascade;Downstream signal transduction;VEGFR2 mediated cell proliferation;Signaling by SCF-KIT;Frs2-mediated activation;ARMS-mediated activation;Gastrin-CREB signalling pathway via PKC and MAPK;Interleukin receptor SHC signaling;DAP12 signaling;Signaling by PDGF;VEGFA-VEGFR2 Pathway;Signaling by ERBB2;IRS-related events triggered by IGF1R;Cell surface interactions at the vascular wall 
                                               hsa04010;hsa04012;hsa04014;hsa04015;hsa04062;hsa04068;hsa04071;hsa04072;hsa04151;hsa04211;hsa04213;hsa04320;hsa04360;hsa04370;hsa04530;hsa04540;hsa04550;hsa04650;hsa04660;hsa04662;hsa04664;hsa04720;hsa04722;hsa04725;hsa04726;hsa04730;hsa04810;hsa04910;hsa04912;hsa04914;hsa04915;hsa04916;hsa04917;hsa04919;hsa04921;hsa04933;hsa04960;hsa05034;hsa05160;hsa05161;hsa05166;hsa05200;hsa05203;hsa05205;hsa05206;hsa05210;hsa05211;hsa05212;hsa05213;hsa05214;hsa05215;hsa05216;hsa05218;hsa05219;hsa05220;hsa05221;hsa05223;hsa05230;hsa05231 
                                               KRAS 
                                               NA 
                                               slateblue4 
                                               16 
                                               2.69565217391304 
                                             
                                             
                                               HRAS-mRNA 
                                               -0.373 
                                               0.164 
                                               -0.694 
                                               -0.0518 
                                               0.772 
                                               0.618 
                                               0.965 
                                               0.0334 
                                               0.522 
                                               loglinear 
                                               ARMS-mediated activation, CD209 (DC-SIGN) signaling, Cell surface interactions at the vascular wall, Constitutive Signaling by EGFRvIII, CREB phosphorylation through the activation of Ras, DAP12 signaling, Downstream signal transduction, FCERI mediated MAPK activation, Frs2-mediated activation, Gastrin-CREB signalling pathway via PKC and MAPK, Interleukin receptor SHC signaling, IRS-related events triggered by IGF1R, MAP2K and MAPK activation, RAFMAP kinase cascade, SHC-mediated cascade.CL.FGFR1, SHC-related events triggered by IGF1R, Signaling by ERBB2, Signaling by FGFR1 in disease, Signaling by FGFR2 in disease, Signaling by FGFR3 in disease, Signaling by FGFR4 in disease, Signaling by PDGF, Signaling by SCF-KIT, Tie2 Signaling, VEGFA-VEGFR2 Pathway, VEGFR2 mediated cell proliferation 
                                               SEBELIUS_2_C4636 
                                               HRAS 
                                               mRNA 
                                               0 
                                               
                                               
                                               Signaling by FGFR2 in disease;Regulation of RAS by GAPs;SHC-mediated cascade.CL.FGFR3;Signaling by FGFR3 in disease;MAP2K and MAPK activation;SHC-related events triggered by IGF1R;Signaling by FGFR4 in disease;SHC-mediated cascade.CL.FGFR1;Tie2 Signaling;Signaling by FGFR1 in disease;Activation of RAS in B cells;SHC-mediated cascade.CL.FGFR4;Constitutive Signaling by EGFRvIII;CD209 (DC-SIGN) signaling;Ras activation uopn Ca2.PLS. infux through NMDA receptor;FCERI mediated MAPK activation;RAFMAP kinase cascade;Downstream signal transduction;VEGFR2 mediated cell proliferation;Signaling by SCF-KIT;Frs2-mediated activation;ARMS-mediated activation;Gastrin-CREB signalling pathway via PKC and MAPK;Interleukin receptor SHC signaling;DAP12 signaling;Signaling by PDGF;VEGFA-VEGFR2 Pathway;Signaling by ERBB2;IRS-related events triggered by IGF1R;Cell surface interactions at the vascular wall;CREB phosphorylation through the activation of Ras 
                                               hsa04010;hsa04012;hsa04014;hsa04015;hsa04062;hsa04068;hsa04071;hsa04072;hsa04144;hsa04151;hsa04211;hsa04213;hsa04360;hsa04370;hsa04510;hsa04530;hsa04540;hsa04550;hsa04630;hsa04650;hsa04660;hsa04662;hsa04664;hsa04720;hsa04722;hsa04725;hsa04726;hsa04730;hsa04810;hsa04910;hsa04912;hsa04915;hsa04916;hsa04917;hsa04919;hsa04921;hsa04933;hsa05034;hsa05160;hsa05161;hsa05166;hsa05200;hsa05203;hsa05205;hsa05206;hsa05211;hsa05213;hsa05214;hsa05215;hsa05216;hsa05218;hsa05219;hsa05220;hsa05221;hsa05223;hsa05230;hsa05231 
                                               HRAS 
                                               NA 
                                               azure4 
                                               1 
                                               -2.27439024390244 
                                             
                                             
                                               VCL-mRNA 
                                               0.225 
                                               0.123 
                                               -0.0163 
                                               0.467 
                                               1.17 
                                               0.989 
                                               1.38 
                                               0.0818 
                                               1 
                                               lm.nb 
                                               ARMS-mediated activation, DAP12 signaling, Downstream signal transduction, FCERI mediated MAPK activation, Frs2-mediated activation, Gastrin-CREB signalling pathway via PKC and MAPK, Interleukin receptor SHC signaling, IRS-related events triggered by IGF1R, MAP2K and MAPK activation, Platelet degranulation , RAFMAP kinase cascade, Signaling by ERBB2, Signaling by PDGF, Signaling by SCF-KIT, Smooth Muscle Contraction, VEGFA-VEGFR2 Pathway, VEGFR2 mediated cell proliferation 
                                               SEBELIUS_2_C4636 
                                               VCL 
                                               mRNA 
                                               0 
                                               
                                               
                                               Smooth Muscle Contraction;MAP2K and MAPK activation;Platelet degranulation ;RAFMAP kinase cascade;Downstream signal transduction;VEGFR2 mediated cell proliferation;Signaling by SCF-KIT;Frs2-mediated activation;ARMS-mediated activation;Gastrin-CREB signalling pathway via PKC and MAPK;Interleukin receptor SHC signaling;FCERI mediated MAPK activation;DAP12 signaling;Signaling by PDGF;VEGFA-VEGFR2 Pathway;Signaling by ERBB2;IRS-related events triggered by IGF1R 
                                               hsa04510;hsa04520;hsa04670;hsa04810;hsa05100;hsa05131;hsa05146 
                                               VCL 
                                               NA 
                                               azure4 
                                               1 
                                               1.82926829268293 
                                             
                                             
                                               PPP2R1A-mRNA 
                                               -0.206 
                                               0.113 
                                               -0.428 
                                               0.0153 
                                               0.867 
                                               0.743 
                                               1.01 
                                               0.0824 
                                               1 
                                               loglinear 
                                               ARMS-mediated activation, AXIN missense mutants destabilize the destruction complex, Beta-catenin phosphorylation cascade, CTLA4 inhibitory signaling, Cyclin AB1 associated events during G2M transition, Cyclin D associated events in G1, DAP12 signaling, DARPP-32 events, Disassembly of the destruction complex and recruitment of AXIN to the membrane, Downstream signal transduction, FCERI mediated MAPK activation, Frs2-mediated activation, Gastrin-CREB signalling pathway via PKC and MAPK, Interleukin receptor SHC signaling, IRS-related events triggered by IGF1R, MASTL Facilitates Mitotic Progression, Negative regulation of FGFR1 signaling, Negative regulation of FGFR2 signaling, Negative regulation of FGFR3 signaling, Negative regulation of FGFR4 signaling, Nonsense Mediated Decay (NMD) enhanced by the Exon Junction Complex (EJC), RAF activation, RAFMAP kinase cascade, Resolution of Sister Chromatid Cohesion, RHO GTPases Activate Formins, S33 mutants of beta-catenin aren.SQT.t phosphorylated, S37 mutants of beta-catenin aren.SQT.t phosphorylated, S45 mutants of beta-catenin aren.SQT.t phosphorylated, Signaling by ERBB2, Signaling by PDGF, Signaling by SCF-KIT, Spry regulation of FGF signaling, Truncations of AMER1 destabilize the destruction complex, VEGFA-VEGFR2 Pathway, VEGFR2 mediated cell proliferation 
                                               SEBELIUS_2_C4636 
                                               PPP2R1A 
                                               mRNA 
                                               0 
                                               
                                               
                                               MASTL Facilitates Mitotic Progression;Regulation of PLK1 Activity at G2M Transition;Disassembly of the destruction complex and recruitment of AXIN to the membrane;S37 mutants of beta-catenin aren.SQT.t phosphorylated;Cyclin AB1 associated events during G2M transition;RAF activation;RHO GTPases Activate Formins;S33 mutants of beta-catenin aren.SQT.t phosphorylated;Loss of Nlp from mitotic centrosomes;DARPP-32 events;Beta-catenin phosphorylation cascade;Initiation of Nuclear Envelope Reformation;Recruitment of mitotic centrosome proteins and complexes;Spry regulation of FGF signaling;Truncations of AMER1 destabilize the destruction complex;CTLA4 inhibitory signaling;AXIN missense mutants destabilize the destruction complex;S45 mutants of beta-catenin aren.SQT.t phosphorylated;Anchoring of the basal body to the plasma membrane;Nonsense Mediated Decay (NMD) enhanced by the Exon Junction Complex (EJC);Cyclin D associated events in G1;Platelet sensitization by LDL;Resolution of Sister Chromatid Cohesion;Inhibition of replication initiation of damaged DNA by RB1E2F1;RAFMAP kinase cascade;Downstream signal transduction;VEGFR2 mediated cell proliferation;Signaling by SCF-KIT;Frs2-mediated activation;ARMS-mediated activation;Gastrin-CREB signalling pathway via PKC and MAPK;Interleukin receptor SHC signaling;FCERI mediated MAPK activation;DAP12 signaling;Signaling by PDGF;VEGFA-VEGFR2 Pathway;Signaling by ERBB2;IRS-related events triggered by IGF1R;Negative regulation of FGFR3 signaling;Negative regulation of FGFR4 signaling;Negative regulation of FGFR2 signaling;Negative regulation of FGFR1 signaling 
                                               hsa03015;hsa04071;hsa04114;hsa04151;hsa04152;hsa04261;hsa04350;hsa04390;hsa04530;hsa04728;hsa04730;hsa05142;hsa05160 
                                               PPP2R1A 
                                               NA 
                                               azure4 
                                               1 
                                               -1.82300884955752 
                                             
                                             
                                               SPTAN1-mRNA 
                                               0.215 
                                               0.127 
                                               -0.0337 
                                               0.463 
                                               1.16 
                                               0.977 
                                               1.38 
                                               0.105 
                                               1 
                                               loglinear 
                                               Apoptotic cleavage of cellular proteins, ARMS-mediated activation, DAP12 signaling, Downstream signal transduction, FCERI mediated MAPK activation, Frs2-mediated activation, Gastrin-CREB signalling pathway via PKC and MAPK, Interleukin receptor SHC signaling, IRS-related events triggered by IGF1R, Nephrin interactions, Signaling by ERBB2, Signaling by PDGF, Signaling by SCF-KIT, VEGFA-VEGFR2 Pathway, VEGFR2 mediated cell proliferation 
                                               SEBELIUS_2_C4636 
                                               SPTAN1 
                                               mRNA 
                                               0 
                                               
                                               
                                               Nephrin interactions;Caspase-mediated cleavage of cytoskeletal proteins;Downstream signal transduction;VEGFR2 mediated cell proliferation;Signaling by SCF-KIT;Frs2-mediated activation;ARMS-mediated activation;Gastrin-CREB signalling pathway via PKC and MAPK;Interleukin receptor SHC signaling;FCERI mediated MAPK activation;DAP12 signaling;Signaling by PDGF;VEGFA-VEGFR2 Pathway;Signaling by ERBB2;IRS-related events triggered by IGF1R;Apoptotic cleavage of cellular proteins 
                                               hsa04530 
                                               SPTAN1 
                                               NA 
                                               azure4 
                                               1 
                                               1.69291338582677 
                                             
                                             
                                               PTK2-mRNA 
                                               -0.129 
                                               0.0794 
                                               -0.285 
                                               0.0263 
                                               0.914 
                                               0.821 
                                               1.02 
                                               0.118 
                                               1 
                                               loglinear 
                                               Apoptotic cleavage of cellular proteins, ARMS-mediated activation, DAP12 signaling, DCC mediated attractive signaling, Downstream signal transduction, EPHA-mediated growth cone collapse, EPHB-mediated forward signaling, FCERI mediated MAPK activation, Frs2-mediated activation, Gastrin-CREB signalling pathway via PKC and MAPK, Integrin alphaIIb beta3 signaling, Interleukin receptor SHC signaling, IRS-related events triggered by IGF1R, p130Cas linkage to MAPK signaling for integrins, RAFMAP kinase cascade, Regulation of actin dynamics for phagocytic cup formation, RHO GTPases Activate WASPs and WAVEs, Signaling by ERBB2, Signaling by PDGF, Signaling by SCF-KIT, VEGFA-VEGFR2 Pathway, VEGFR2 mediated cell proliferation 
                                               SEBELIUS_2_C4636 
                                               PTK2 
                                               mRNA 
                                               0 
                                               
                                               
                                               Apoptotic cleavage of cellular proteins;RAFMAP kinase cascade;RHO GTPases Activate WASPs and WAVEs;DCC mediated attractive signaling;p130Cas linkage to MAPK signaling for integrins;EPHB-mediated forward signaling;EPHA-mediated growth cone collapse;Regulation of actin dynamics for phagocytic cup formation;Downstream signal transduction;VEGFR2 mediated cell proliferation;Signaling by SCF-KIT;Frs2-mediated activation;ARMS-mediated activation;Gastrin-CREB signalling pathway via PKC and MAPK;Interleukin receptor SHC signaling;FCERI mediated MAPK activation;DAP12 signaling;Signaling by PDGF;VEGFA-VEGFR2 Pathway;Signaling by ERBB2;IRS-related events triggered by IGF1R;Integrin alphaIIb beta3 signaling 
                                               hsa04012;hsa04062;hsa04151;hsa04360;hsa04370;hsa04510;hsa04670;hsa04810;hsa05100;hsa05146;hsa05200;hsa05202;hsa05205;hsa05222 
                                               PTK2 
                                               NA 
                                               azure4 
                                               1 
                                               -1.62468513853904 
                                             
                                             
                                               PDGFRB-mRNA 
                                               -0.28 
                                               0.181 
                                               -0.634 
                                               0.0753 
                                               0.824 
                                               0.644 
                                               1.05 
                                               0.138 
                                               1 
                                               lm.nb 
                                               ARMS-mediated activation, Constitutive Signaling by Aberrant PI3K in Cancer, DAP12 signaling, Downstream signal transduction, FCERI mediated MAPK activation, Frs2-mediated activation, GAB1 signalosome, Gastrin-CREB signalling pathway via PKC and MAPK, Interleukin receptor SHC signaling, IRS-related events triggered by IGF1R, PI3KAKT activation, PIP3 activates AKT signaling, RAFMAP kinase cascade, Role of LAT2NTALLAB on calcium mobilization, Signaling by ERBB2, Signaling by PDGF, Signaling by SCF-KIT, VEGFA-VEGFR2 Pathway, VEGFR2 mediated cell proliferation 
                                               SEBELIUS_2_C4636 
                                               PDGFRB 
                                               mRNA 
                                               0 
                                               
                                               
                                               PIP3 activates AKT signaling;Constitutive Signaling by Aberrant PI3K in Cancer;RAFMAP kinase cascade;Signaling by PDGF;Downstream signal transduction;GAB1 signalosome;Signaling by SCF-KIT;PI3KAKT activation;Role of LAT2NTALLAB on calcium mobilization;DAP12 signaling;Signaling by ERBB2;VEGFR2 mediated cell proliferation;Frs2-mediated activation;ARMS-mediated activation;Gastrin-CREB signalling pathway via PKC and MAPK;Interleukin receptor SHC signaling;FCERI mediated MAPK activation;VEGFA-VEGFR2 Pathway;IRS-related events triggered by IGF1R 
                                               hsa04010;hsa04014;hsa04015;hsa04020;hsa04060;hsa04072;hsa04151;hsa04510;hsa04540;hsa04810;hsa05166;hsa05200;hsa05206;hsa05214;hsa05215;hsa05218;hsa05230;hsa05231 
                                               PDGFRB 
                                               NA 
                                               azure4 
                                               1 
                                               -1.54696132596685 
                                             
                                             
                                               ERBB2-mRNA 
                                               -0.247 
                                               0.189 
                                               -0.618 
                                               0.124 
                                               0.843 
                                               0.652 
                                               1.09 
                                               0.206 
                                               1 
                                               lm.nb 
                                               ARMS-mediated activation, DAP12 signaling, Downstream signal transduction, FCERI mediated MAPK activation, Frs2-mediated activation, GAB1 signalosome, Gastrin-CREB signalling pathway via PKC and MAPK, Interleukin receptor SHC signaling, IRS-related events triggered by IGF1R, PI3KAKT activation, RAFMAP kinase cascade, Role of LAT2NTALLAB on calcium mobilization, Sema4D induced cell migration and growth-cone collapse, Signaling by ERBB2, Signaling by PDGF, Signaling by SCF-KIT, VEGFA-VEGFR2 Pathway, VEGFR2 mediated cell proliferation 
                                               SEBELIUS_2_C4636 
                                               ERBB2 
                                               mRNA 
                                               0 
                                               
                                               
                                               RAFMAP kinase cascade;GRB7 events in ERBB2 signaling;PLCG1 events in ERBB2 signaling;Sema4D induced cell migration and growth-cone collapse;Signaling by ERBB2;Downstream signal transduction;GAB1 signalosome;Signaling by SCF-KIT;PI3KAKT activation;Role of LAT2NTALLAB on calcium mobilization;DAP12 signaling;Signaling by PDGF;VEGFR2 mediated cell proliferation;Frs2-mediated activation;ARMS-mediated activation;Gastrin-CREB signalling pathway via PKC and MAPK;Interleukin receptor SHC signaling;FCERI mediated MAPK activation;VEGFA-VEGFR2 Pathway;IRS-related events triggered by IGF1R 
                                               hsa04012;hsa04020;hsa04066;hsa04510;hsa04520;hsa05200;hsa05205;hsa05206;hsa05212;hsa05213;hsa05215;hsa05219;hsa05223;hsa05230 
                                               ERBB2 
                                               NA 
                                               azure4 
                                               1 
                                               -1.30687830687831 
                                             
                                           
                                           DE Results - ARMS-mediated activation genes - New.Annotation: miR18b vs.cel67  Table displaying each sample's global significance scores and directed global significance scores as defined in the heatmaps above. The global significance score is calculated as the square root of the mean squared t-statistic for the genes in a gene set, with t-statistics coming from the linear regression underlying our differential expression analysis. The directed global significance score is calculated as the square root of the mean signed squared t-statistic for the genes in a gene set, with t-statistics coming from the linear regression underlying our differential expression analysis.   
                                      
                                       
                                  
                                
                                
                                  
                                       
                                      
                                           Volcano Plot: New.Annotation: miR890 vs.cel67   More Plot Information                Volcano Plot: New.Annotation: miR890 vs.cel67  Volcano plot displaying each gene's -log10(p-value) and log2 fold change for the selected covariate. Highly statistically significant genes fall at the top of the plot, and highly differentially expressed genes fall to either side. Genes within the selected gene set are highlighted in orange. Horizontal lines indicate various False Discovery Rate (FDR) thresholds.   
                                           DE Results - ARMS-mediated activation genes - New.Annotation: miR890 vs.cel67   More Plot Information      Download CSV Data     
                                           
                                             
                                             
                                           
                                           
                                             
                                               Probe Label 
                                               Log2 fold change 
                                               std error (log2) 
                                               Lower confidence limit (log2) 
                                               Upper confidence limit (log2) 
                                               Linear fold change 
                                               Lower confidence limit (linear) 
                                               Upper confidence limit (linear) 
                                               P-value 
                                               BY.p.value 
                                               method 
                                               Gene.sets 
                                               Codeset.Name 
                                               Probe.Label 
                                               Analyte.Type 
                                               Is.Control 
                                               Control.Type 
                                               Related.Probes 
                                               Probe.Annotation 
                                               KEGG.Pathways 
                                               Official.Gene.Name 
                                               Control.Conc 
                                               volcanocol 
                                               volcanopch 
                                               tstats.all 
                                             
                                           
                                           
                                             
                                               MAP2K1-mRNA 
                                               -0.977 
                                               0.116 
                                               -1.2 
                                               -0.75 
                                               0.508 
                                               0.434 
                                               0.595 
                                               3.48e-08 
                                               6.61e-06 
                                               lm.nb 
                                               ARMS-mediated activation, DAP12 signaling, Downstream signal transduction, FCERI mediated MAPK activation, Frs2-mediated activation, Gastrin-CREB signalling pathway via PKC and MAPK, Interleukin receptor SHC signaling, IRS-related events triggered by IGF1R, Negative feedback regulation of MAPK pathway, Negative regulation of MAPK pathway, RAF activation, RAFMAP kinase cascade, Signaling by ERBB2, Signaling by PDGF, Signaling by SCF-KIT, VEGFA-VEGFR2 Pathway, VEGFR2 mediated cell proliferation 
                                               SEBELIUS_2_C4636 
                                               MAP2K1 
                                               mRNA 
                                               0 
                                               
                                               
                                               RAF activation;Uptake and function of anthrax toxins;Negative feedback regulation of MAPK pathway;Interleukin-1 signaling;RAFMAP kinase cascade;Downstream signal transduction;VEGFR2 mediated cell proliferation;Signaling by SCF-KIT;Frs2-mediated activation;ARMS-mediated activation;Gastrin-CREB signalling pathway via PKC and MAPK;Interleukin receptor SHC signaling;FCERI mediated MAPK activation;DAP12 signaling;Signaling by PDGF;VEGFA-VEGFR2 Pathway;Signaling by ERBB2;IRS-related events triggered by IGF1R;Negative regulation of MAPK pathway 
                                               hsa04010;hsa04012;hsa04014;hsa04015;hsa04022;hsa04024;hsa04062;hsa04066;hsa04068;hsa04071;hsa04072;hsa04114;hsa04151;hsa04270;hsa04320;hsa04370;hsa04380;hsa04510;hsa04540;hsa04550;hsa04620;hsa04650;hsa04660;hsa04662;hsa04664;hsa04666;hsa04668;hsa04720;hsa04722;hsa04725;hsa04726;hsa04730;hsa04810;hsa04910;hsa04912;hsa04914;hsa04915;hsa04916;hsa04917;hsa04919;hsa04921;hsa05020;hsa05034;hsa05161;hsa05164;hsa05200;hsa05205;hsa05206;hsa05210;hsa05211;hsa05212;hsa05213;hsa05214;hsa05215;hsa05216;hsa05218;hsa05219;hsa05220;hsa05221;hsa05223;hsa05230;hsa05231 
                                               MAP2K1 
                                               NA 
                                               slateblue1 
                                               16 
                                               -8.42241379310345 
                                             
                                             
                                               MAPK3-mRNA 
                                               0.393 
                                               0.052 
                                               0.291 
                                               0.495 
                                               1.31 
                                               1.22 
                                               1.41 
                                               2.01e-07 
                                               2.67e-05 
                                               loglinear 
                                               ARMS-mediated activation, DAP12 signaling, Downstream signal transduction, ERKs are inactivated, FCERI mediated MAPK activation, Frs2-mediated activation, Gastrin-CREB signalling pathway via PKC and MAPK, Interleukin receptor SHC signaling, IRS-related events triggered by IGF1R, Negative feedback regulation of MAPK pathway, Negative regulation of FGFR3 signaling, Negative regulation of FGFR4 signaling, Negative regulation of MAPK pathway, RAFMAP kinase cascade, Regulation of HSF1-mediated heat shock response, RHO GTPases Activate WASPs and WAVEs, Signaling by ERBB2, Signaling by PDGF, Signaling by SCF-KIT, VEGFA-VEGFR2 Pathway, VEGFR2 mediated cell proliferation 
                                               SEBELIUS_2_C4636 
                                               MAPK3 
                                               mRNA 
                                               0 
                                               
                                               
                                               ERKs are inactivated;ISG15 antiviral mechanism;RHO GTPases Activate WASPs and WAVEs;Senescence-Associated Secretory Phenotype (SASP);Regulation of HSF1-mediated heat shock response;Thrombin signalling through proteinase activated receptors (PARs);Negative regulation of FGFR4 signaling;Oxidative Stress Induced Senescence;Signal attenuation;RNA Polymerase I Promoter Opening;Negative regulation of FGFR3 signaling;Negative feedback regulation of MAPK pathway;Growth hormone receptor signaling;Golgi Cisternae Pericentriolar Stack Reorganization;FCERI mediated MAPK activation;Negative regulation of MAPK pathway;RAFMAP kinase cascade;Downstream signal transduction;VEGFR2 mediated cell proliferation;Signaling by SCF-KIT;Frs2-mediated activation;ARMS-mediated activation;Gastrin-CREB signalling pathway via PKC and MAPK;Interleukin receptor SHC signaling;DAP12 signaling;Signaling by PDGF;VEGFA-VEGFR2 Pathway;Signaling by ERBB2;IRS-related events triggered by IGF1R 
                                               hsa04010;hsa04012;hsa04014;hsa04015;hsa04022;hsa04024;hsa04062;hsa04066;hsa04068;hsa04071;hsa04072;hsa04114;hsa04150;hsa04151;hsa04261;hsa04270;hsa04320;hsa04350;hsa04360;hsa04370;hsa04380;hsa04510;hsa04520;hsa04540;hsa04550;hsa04611;hsa04620;hsa04621;hsa04650;hsa04660;hsa04662;hsa04664;hsa04666;hsa04668;hsa04713;hsa04720;hsa04722;hsa04723;hsa04724;hsa04725;hsa04726;hsa04730;hsa04810;hsa04910;hsa04912;hsa04914;hsa04915;hsa04916;hsa04917;hsa04919;hsa04921;hsa04930;hsa04933;hsa04960;hsa05010;hsa05020;hsa05034;hsa05131;hsa05132;hsa05133;hsa05140;hsa05142;hsa05145;hsa05152;hsa05160;hsa05161;hsa05164;hsa05200;hsa05203;hsa05205;hsa05210;hsa05211;hsa05212;hsa05213;hsa05214;hsa05215;hsa05216;hsa05218;hsa05219;hsa05220;hsa05221;hsa05223;hsa05230;hsa05231 
                                               MAPK3 
                                               NA 
                                               slateblue1 
                                               16 
                                               7.55769230769231 
                                             
                                             
                                               NRAS-mRNA 
                                               -0.783 
                                               0.109 
                                               -0.997 
                                               -0.568 
                                               0.581 
                                               0.501 
                                               0.674 
                                               4.76e-07 
                                               5.27e-05 
                                               lm.nb 
                                               ARMS-mediated activation, CD209 (DC-SIGN) signaling, Cell surface interactions at the vascular wall, Constitutive Signaling by EGFRvIII, DAP12 signaling, Downstream signal transduction, FCERI mediated MAPK activation, Frs2-mediated activation, Gastrin-CREB signalling pathway via PKC and MAPK, Interleukin receptor SHC signaling, IRS-related events triggered by IGF1R, MAP2K and MAPK activation, RAFMAP kinase cascade, SHC-mediated cascade.CL.FGFR1, SHC-related events triggered by IGF1R, Signaling by ERBB2, Signaling by FGFR1 in disease, Signaling by FGFR2 in disease, Signaling by FGFR3 in disease, Signaling by FGFR4 in disease, Signaling by PDGF, Signaling by SCF-KIT, Tie2 Signaling, VEGFA-VEGFR2 Pathway, VEGFR2 mediated cell proliferation 
                                               SEBELIUS_2_C4636 
                                               NRAS 
                                               mRNA 
                                               0 
                                               
                                               
                                               Signaling by FGFR2 in disease;Regulation of RAS by GAPs;SHC-mediated cascade.CL.FGFR3;Signaling by FGFR3 in disease;MAP2K and MAPK activation;SHC-related events triggered by IGF1R;Signaling by FGFR4 in disease;SHC-mediated cascade.CL.FGFR1;Tie2 Signaling;Signaling by FGFR1 in disease;Activation of RAS in B cells;SHC-mediated cascade.CL.FGFR4;Constitutive Signaling by EGFRvIII;CD209 (DC-SIGN) signaling;FCERI mediated MAPK activation;RAFMAP kinase cascade;Downstream signal transduction;VEGFR2 mediated cell proliferation;Signaling by SCF-KIT;Frs2-mediated activation;ARMS-mediated activation;Gastrin-CREB signalling pathway via PKC and MAPK;Interleukin receptor SHC signaling;DAP12 signaling;Signaling by PDGF;VEGFA-VEGFR2 Pathway;Signaling by ERBB2;IRS-related events triggered by IGF1R;Cell surface interactions at the vascular wall 
                                               hsa04010;hsa04012;hsa04014;hsa04015;hsa04062;hsa04068;hsa04071;hsa04072;hsa04151;hsa04211;hsa04213;hsa04360;hsa04370;hsa04530;hsa04540;hsa04550;hsa04650;hsa04660;hsa04662;hsa04664;hsa04720;hsa04722;hsa04725;hsa04726;hsa04730;hsa04810;hsa04910;hsa04912;hsa04915;hsa04916;hsa04917;hsa04919;hsa04921;hsa04933;hsa05034;hsa05160;hsa05161;hsa05166;hsa05200;hsa05203;hsa05205;hsa05206;hsa05211;hsa05213;hsa05214;hsa05215;hsa05216;hsa05218;hsa05219;hsa05220;hsa05221;hsa05223;hsa05230;hsa05231 
                                               NRAS 
                                               NA 
                                               slateblue1 
                                               16 
                                               -7.18348623853211 
                                             
                                             
                                               PPP2CB-mRNA 
                                               -0.402 
                                               0.0718 
                                               -0.542 
                                               -0.261 
                                               0.757 
                                               0.687 
                                               0.835 
                                               1.51e-05 
                                               0.000909 
                                               loglinear 
                                               ARMS-mediated activation, AXIN missense mutants destabilize the destruction complex, Beta-catenin phosphorylation cascade, CTLA4 inhibitory signaling, Cyclin AB1 associated events during G2M transition, Cyclin D associated events in G1, DAP12 signaling, DARPP-32 events, Disassembly of the destruction complex and recruitment of AXIN to the membrane, Downstream signal transduction, ERKs are inactivated, FCERI mediated MAPK activation, Frs2-mediated activation, Gastrin-CREB signalling pathway via PKC and MAPK, Interleukin receptor SHC signaling, IRS-related events triggered by IGF1R, MASTL Facilitates Mitotic Progression, Misspliced GSK3beta mutants stabilize beta-catenin, Negative regulation of FGFR1 signaling, Negative regulation of FGFR2 signaling, Negative regulation of FGFR3 signaling, Negative regulation of FGFR4 signaling, Negative regulation of MAPK pathway, RAF activation, RAFMAP kinase cascade, Resolution of Sister Chromatid Cohesion, RHO GTPases Activate Formins, S33 mutants of beta-catenin aren.SQT.t phosphorylated, S37 mutants of beta-catenin aren.SQT.t phosphorylated, S45 mutants of beta-catenin aren.SQT.t phosphorylated, Signaling by ERBB2, Signaling by PDGF, Signaling by SCF-KIT, Spry regulation of FGF signaling, T41 mutants of beta-catenin aren.SQT.t phosphorylated, Truncations of AMER1 destabilize the destruction complex, VEGFA-VEGFR2 Pathway, VEGFR2 mediated cell proliferation 
                                               SEBELIUS_2_C4636 
                                               PPP2CB 
                                               mRNA 
                                               0 
                                               
                                               
                                               MASTL Facilitates Mitotic Progression;Misspliced GSK3beta mutants stabilize beta-catenin;T41 mutants of beta-catenin aren.SQT.t phosphorylated;ERKs are inactivated;Glycolysis;Negative regulation of MAPK pathway;Disassembly of the destruction complex and recruitment of AXIN to the membrane;S37 mutants of beta-catenin aren.SQT.t phosphorylated;Cyclin AB1 associated events during G2M transition;RAF activation;RHO GTPases Activate Formins;S33 mutants of beta-catenin aren.SQT.t phosphorylated;DARPP-32 events;Beta-catenin phosphorylation cascade;Spry regulation of FGF signaling;Truncations of AMER1 destabilize the destruction complex;CTLA4 inhibitory signaling;AXIN missense mutants destabilize the destruction complex;S45 mutants of beta-catenin aren.SQT.t phosphorylated;Cyclin D associated events in G1;Platelet sensitization by LDL;Resolution of Sister Chromatid Cohesion;Inhibition of replication initiation of damaged DNA by RB1E2F1;RAFMAP kinase cascade;Downstream signal transduction;VEGFR2 mediated cell proliferation;Signaling by SCF-KIT;Frs2-mediated activation;ARMS-mediated activation;Gastrin-CREB signalling pathway via PKC and MAPK;Interleukin receptor SHC signaling;FCERI mediated MAPK activation;DAP12 signaling;Signaling by PDGF;VEGFA-VEGFR2 Pathway;Signaling by ERBB2;IRS-related events triggered by IGF1R;Negative regulation of FGFR3 signaling;Negative regulation of FGFR4 signaling;Negative regulation of FGFR2 signaling;Negative regulation of FGFR1 signaling 
                                               hsa03015;hsa04071;hsa04114;hsa04151;hsa04152;hsa04261;hsa04350;hsa04390;hsa04530;hsa04728;hsa04730;hsa05142;hsa05160 
                                               PPP2CB 
                                               NA 
                                               slateblue1 
                                               16 
                                               -5.59888579387187 
                                             
                                             
                                               IRS1-mRNA 
                                               -0.487 
                                               0.104 
                                               -0.691 
                                               -0.283 
                                               0.713 
                                               0.62 
                                               0.822 
                                               0.000125 
                                               0.00469 
                                               loglinear 
                                               ARMS-mediated activation, Constitutive Signaling by Aberrant PI3K in Cancer, DAP12 signaling, Downstream signal transduction, FCERI mediated MAPK activation, Frs2-mediated activation, GAB1 signalosome, Gastrin-CREB signalling pathway via PKC and MAPK, Interleukin receptor SHC signaling, IRS-related events triggered by IGF1R, PI3K Cascade, PI3KAKT activation, RAFMAP kinase cascade, Role of LAT2NTALLAB on calcium mobilization, Signaling by ERBB2, Signaling by PDGF, Signaling by SCF-KIT, VEGFA-VEGFR2 Pathway, VEGFR2 mediated cell proliferation 
                                               SEBELIUS_2_C4636 
                                               IRS1 
                                               mRNA 
                                               0 
                                               
                                               
                                               Constitutive Signaling by Aberrant PI3K in Cancer;RAFMAP kinase cascade;PI3K Cascade;Growth hormone receptor signaling;IRS-related events triggered by IGF1R;PI3KAKT activation;Downstream signal transduction;GAB1 signalosome;Signaling by SCF-KIT;Role of LAT2NTALLAB on calcium mobilization;DAP12 signaling;Signaling by PDGF;Signaling by ERBB2;VEGFR2 mediated cell proliferation;Frs2-mediated activation;ARMS-mediated activation;Gastrin-CREB signalling pathway via PKC and MAPK;Interleukin receptor SHC signaling;FCERI mediated MAPK activation;VEGFA-VEGFR2 Pathway 
                                               hsa04022;hsa04068;hsa04150;hsa04151;hsa04152;hsa04211;hsa04213;hsa04722;hsa04910;hsa04920;hsa04923;hsa04930;hsa04931;hsa04932;hsa04960;hsa05206 
                                               IRS1 
                                               NA 
                                               slateblue1 
                                               16 
                                               -4.68269230769231 
                                             
                                             
                                               SPTAN1-mRNA 
                                               0.467 
                                               0.107 
                                               0.257 
                                               0.677 
                                               1.38 
                                               1.19 
                                               1.6 
                                               0.000275 
                                               0.00892 
                                               loglinear 
                                               Apoptotic cleavage of cellular proteins, ARMS-mediated activation, DAP12 signaling, Downstream signal transduction, FCERI mediated MAPK activation, Frs2-mediated activation, Gastrin-CREB signalling pathway via PKC and MAPK, Interleukin receptor SHC signaling, IRS-related events triggered by IGF1R, Nephrin interactions, Signaling by ERBB2, Signaling by PDGF, Signaling by SCF-KIT, VEGFA-VEGFR2 Pathway, VEGFR2 mediated cell proliferation 
                                               SEBELIUS_2_C4636 
                                               SPTAN1 
                                               mRNA 
                                               0 
                                               
                                               
                                               Nephrin interactions;Caspase-mediated cleavage of cytoskeletal proteins;Downstream signal transduction;VEGFR2 mediated cell proliferation;Signaling by SCF-KIT;Frs2-mediated activation;ARMS-mediated activation;Gastrin-CREB signalling pathway via PKC and MAPK;Interleukin receptor SHC signaling;FCERI mediated MAPK activation;DAP12 signaling;Signaling by PDGF;VEGFA-VEGFR2 Pathway;Signaling by ERBB2;IRS-related events triggered by IGF1R;Apoptotic cleavage of cellular proteins 
                                               hsa04530 
                                               SPTAN1 
                                               NA 
                                               slateblue1 
                                               16 
                                               4.36448598130841 
                                             
                                             
                                               MAPK1-mRNA 
                                               -0.246 
                                               0.0614 
                                               -0.367 
                                               -0.126 
                                               0.843 
                                               0.776 
                                               0.916 
                                               0.00063 
                                               0.0178 
                                               loglinear 
                                               ARMS-mediated activation, CREB phosphorylation through the activation of Ras, DAP12 signaling, Downstream signal transduction, FCERI mediated MAPK activation, Frs2-mediated activation, Gastrin-CREB signalling pathway via PKC and MAPK, Interleukin receptor SHC signaling, IRS-related events triggered by IGF1R, MAP2K and MAPK activation, Negative feedback regulation of MAPK pathway, Negative regulation of FGFR1 signaling, Negative regulation of FGFR2 signaling, Negative regulation of FGFR3 signaling, Negative regulation of FGFR4 signaling, Negative regulation of MAPK pathway, RAFMAP kinase cascade, Regulation of HSF1-mediated heat shock response, RHO GTPases Activate WASPs and WAVEs, Signal transduction by L1, Signaling by ERBB2, Signaling by PDGF, Signaling by SCF-KIT, VEGFA-VEGFR2 Pathway, VEGFR2 mediated cell proliferation 
                                               SEBELIUS_2_C4636 
                                               MAPK1 
                                               mRNA 
                                               0 
                                               
                                               
                                               Negative regulation of FGFR1 signaling;Signal transduction by L1;Negative regulation of FGFR2 signaling;MAP2K and MAPK activation;RHO GTPases Activate WASPs and WAVEs;Senescence-Associated Secretory Phenotype (SASP);Regulation of HSF1-mediated heat shock response;Thrombin signalling through proteinase activated receptors (PARs);Negative regulation of FGFR4 signaling;Oxidative Stress Induced Senescence;Signal attenuation;Negative regulation of FGFR3 signaling;Negative feedback regulation of MAPK pathway;Growth hormone receptor signaling;PLC beta mediated events;FCERI mediated MAPK activation;CREB phosphorylation through the activation of Ras;Negative regulation of MAPK pathway;RAFMAP kinase cascade;Downstream signal transduction;VEGFR2 mediated cell proliferation;Signaling by SCF-KIT;Frs2-mediated activation;ARMS-mediated activation;Gastrin-CREB signalling pathway via PKC and MAPK;Interleukin receptor SHC signaling;DAP12 signaling;Signaling by PDGF;VEGFA-VEGFR2 Pathway;Signaling by ERBB2;IRS-related events triggered by IGF1R 
                                               hsa04010;hsa04012;hsa04014;hsa04015;hsa04022;hsa04024;hsa04062;hsa04066;hsa04068;hsa04071;hsa04072;hsa04114;hsa04150;hsa04151;hsa04261;hsa04270;hsa04320;hsa04350;hsa04360;hsa04370;hsa04380;hsa04510;hsa04520;hsa04540;hsa04550;hsa04611;hsa04620;hsa04621;hsa04650;hsa04660;hsa04662;hsa04664;hsa04666;hsa04668;hsa04713;hsa04720;hsa04722;hsa04723;hsa04724;hsa04725;hsa04726;hsa04730;hsa04810;hsa04910;hsa04912;hsa04914;hsa04915;hsa04916;hsa04917;hsa04919;hsa04921;hsa04930;hsa04933;hsa04960;hsa05010;hsa05020;hsa05034;hsa05131;hsa05132;hsa05133;hsa05140;hsa05142;hsa05145;hsa05152;hsa05160;hsa05161;hsa05164;hsa05200;hsa05203;hsa05205;hsa05206;hsa05210;hsa05211;hsa05212;hsa05213;hsa05214;hsa05215;hsa05216;hsa05218;hsa05219;hsa05220;hsa05221;hsa05223;hsa05230;hsa05231 
                                               MAPK1 
                                               NA 
                                               slateblue2 
                                               16 
                                               -4.00651465798046 
                                             
                                             
                                               HRAS-mRNA 
                                               -0.435 
                                               0.138 
                                               -0.706 
                                               -0.163 
                                               0.74 
                                               0.613 
                                               0.893 
                                               0.00495 
                                               0.104 
                                               loglinear 
                                               ARMS-mediated activation, CD209 (DC-SIGN) signaling, Cell surface interactions at the vascular wall, Constitutive Signaling by EGFRvIII, CREB phosphorylation through the activation of Ras, DAP12 signaling, Downstream signal transduction, FCERI mediated MAPK activation, Frs2-mediated activation, Gastrin-CREB signalling pathway via PKC and MAPK, Interleukin receptor SHC signaling, IRS-related events triggered by IGF1R, MAP2K and MAPK activation, RAFMAP kinase cascade, SHC-mediated cascade.CL.FGFR1, SHC-related events triggered by IGF1R, Signaling by ERBB2, Signaling by FGFR1 in disease, Signaling by FGFR2 in disease, Signaling by FGFR3 in disease, Signaling by FGFR4 in disease, Signaling by PDGF, Signaling by SCF-KIT, Tie2 Signaling, VEGFA-VEGFR2 Pathway, VEGFR2 mediated cell proliferation 
                                               SEBELIUS_2_C4636 
                                               HRAS 
                                               mRNA 
                                               0 
                                               
                                               
                                               Signaling by FGFR2 in disease;Regulation of RAS by GAPs;SHC-mediated cascade.CL.FGFR3;Signaling by FGFR3 in disease;MAP2K and MAPK activation;SHC-related events triggered by IGF1R;Signaling by FGFR4 in disease;SHC-mediated cascade.CL.FGFR1;Tie2 Signaling;Signaling by FGFR1 in disease;Activation of RAS in B cells;SHC-mediated cascade.CL.FGFR4;Constitutive Signaling by EGFRvIII;CD209 (DC-SIGN) signaling;Ras activation uopn Ca2.PLS. infux through NMDA receptor;FCERI mediated MAPK activation;RAFMAP kinase cascade;Downstream signal transduction;VEGFR2 mediated cell proliferation;Signaling by SCF-KIT;Frs2-mediated activation;ARMS-mediated activation;Gastrin-CREB signalling pathway via PKC and MAPK;Interleukin receptor SHC signaling;DAP12 signaling;Signaling by PDGF;VEGFA-VEGFR2 Pathway;Signaling by ERBB2;IRS-related events triggered by IGF1R;Cell surface interactions at the vascular wall;CREB phosphorylation through the activation of Ras 
                                               hsa04010;hsa04012;hsa04014;hsa04015;hsa04062;hsa04068;hsa04071;hsa04072;hsa04144;hsa04151;hsa04211;hsa04213;hsa04360;hsa04370;hsa04510;hsa04530;hsa04540;hsa04550;hsa04630;hsa04650;hsa04660;hsa04662;hsa04664;hsa04720;hsa04722;hsa04725;hsa04726;hsa04730;hsa04810;hsa04910;hsa04912;hsa04915;hsa04916;hsa04917;hsa04919;hsa04921;hsa04933;hsa05034;hsa05160;hsa05161;hsa05166;hsa05200;hsa05203;hsa05205;hsa05206;hsa05211;hsa05213;hsa05214;hsa05215;hsa05216;hsa05218;hsa05219;hsa05220;hsa05221;hsa05223;hsa05230;hsa05231 
                                               HRAS 
                                               NA 
                                               slateblue4 
                                               16 
                                               -3.15217391304348 
                                             
                                             
                                               RAF1-mRNA 
                                               -0.213 
                                               0.0701 
                                               -0.351 
                                               -0.0761 
                                               0.862 
                                               0.784 
                                               0.949 
                                               0.00614 
                                               0.125 
                                               loglinear 
                                               ARMS-mediated activation, CD209 (DC-SIGN) signaling, CREB phosphorylation through the activation of Ras, DAP12 signaling, Downstream signal transduction, FCERI mediated MAPK activation, Frs2-mediated activation, Gastrin-CREB signalling pathway via PKC and MAPK, Interleukin receptor SHC signaling, IRS-related events triggered by IGF1R, Negative feedback regulation of MAPK pathway, Negative regulation of MAPK pathway, RAF activation, RAFMAP kinase cascade, Signaling by ERBB2, Signaling by PDGF, Signaling by SCF-KIT, VEGFA-VEGFR2 Pathway, VEGFR2 mediated cell proliferation 
                                               SEBELIUS_2_C4636 
                                               RAF1 
                                               mRNA 
                                               0 
                                               
                                               
                                               RAF activation;Stimuli-sensing channels;CREB phosphorylation through the activation of Ras;Negative feedback regulation of MAPK pathway;CD209 (DC-SIGN) signaling;RAFMAP kinase cascade;Downstream signal transduction;VEGFR2 mediated cell proliferation;Signaling by SCF-KIT;Frs2-mediated activation;ARMS-mediated activation;Gastrin-CREB signalling pathway via PKC and MAPK;Interleukin receptor SHC signaling;FCERI mediated MAPK activation;DAP12 signaling;Signaling by PDGF;VEGFA-VEGFR2 Pathway;Signaling by ERBB2;IRS-related events triggered by IGF1R;Negative regulation of MAPK pathway 
                                               hsa04010;hsa04012;hsa04014;hsa04015;hsa04022;hsa04024;hsa04062;hsa04068;hsa04071;hsa04072;hsa04151;hsa04270;hsa04370;hsa04510;hsa04540;hsa04550;hsa04630;hsa04650;hsa04660;hsa04662;hsa04664;hsa04666;hsa04720;hsa04722;hsa04726;hsa04730;hsa04810;hsa04910;hsa04912;hsa04914;hsa04915;hsa04916;hsa04917;hsa04919;hsa04921;hsa05034;hsa05152;hsa05160;hsa05161;hsa05164;hsa05200;hsa05205;hsa05206;hsa05210;hsa05211;hsa05212;hsa05213;hsa05214;hsa05215;hsa05218;hsa05219;hsa05220;hsa05221;hsa05223;hsa05230;hsa05231 
                                               RAF1 
                                               NA 
                                               slateblue4 
                                               16 
                                               -3.03851640513552 
                                             
                                             
                                               EGFR-mRNA 
                                               -0.291 
                                               0.0968 
                                               -0.481 
                                               -0.101 
                                               0.817 
                                               0.717 
                                               0.932 
                                               0.00673 
                                               0.135 
                                               loglinear 
                                               ARMS-mediated activation, Constitutive Signaling by Aberrant PI3K in Cancer, Constitutive Signaling by EGFRvIII, DAP12 signaling, Downstream signal transduction, FCERI mediated MAPK activation, Frs2-mediated activation, GAB1 signalosome, Gastrin-CREB signalling pathway via PKC and MAPK, Interleukin receptor SHC signaling, IRS-related events triggered by IGF1R, PI3KAKT activation, Role of LAT2NTALLAB on calcium mobilization, Signaling by ERBB2, Signaling by PDGF, Signaling by SCF-KIT, VEGFA-VEGFR2 Pathway, VEGFR2 mediated cell proliferation 
                                               SEBELIUS_2_C4636 
                                               EGFR 
                                               mRNA 
                                               0 
                                               
                                               
                                               Constitutive Signaling by Aberrant PI3K in Cancer;Inhibition of Signaling by Overexpressed EGFR;PLCG1 events in ERBB2 signaling;EGFR interacts with phospholipase C-gamma;EGFR Transactivation by Gastrin;Constitutive Signaling by Ligand-Responsive EGFR Cancer Variants;Constitutive Signaling by EGFRvIII;GAB1 signalosome;Downstream signal transduction;Signaling by SCF-KIT;PI3KAKT activation;Role of LAT2NTALLAB on calcium mobilization;DAP12 signaling;Signaling by PDGF;Signaling by ERBB2;VEGFR2 mediated cell proliferation;Frs2-mediated activation;ARMS-mediated activation;Gastrin-CREB signalling pathway via PKC and MAPK;Interleukin receptor SHC signaling;FCERI mediated MAPK activation;VEGFA-VEGFR2 Pathway;IRS-related events triggered by IGF1R 
                                               hsa04010;hsa04012;hsa04014;hsa04015;hsa04020;hsa04060;hsa04066;hsa04068;hsa04072;hsa04144;hsa04151;hsa04320;hsa04510;hsa04520;hsa04540;hsa04810;hsa04912;hsa04915;hsa04921;hsa05120;hsa05160;hsa05200;hsa05205;hsa05206;hsa05212;hsa05213;hsa05214;hsa05215;hsa05218;hsa05219;hsa05223;hsa05230;hsa05231 
                                               EGFR 
                                               NA 
                                               slateblue4 
                                               16 
                                               -3.00619834710744 
                                             
                                             
                                               VCL-mRNA 
                                               -0.294 
                                               0.104 
                                               -0.499 
                                               -0.0898 
                                               0.815 
                                               0.708 
                                               0.94 
                                               0.0103 
                                               0.192 
                                               lm.nb 
                                               ARMS-mediated activation, DAP12 signaling, Downstream signal transduction, FCERI mediated MAPK activation, Frs2-mediated activation, Gastrin-CREB signalling pathway via PKC and MAPK, Interleukin receptor SHC signaling, IRS-related events triggered by IGF1R, MAP2K and MAPK activation, Platelet degranulation , RAFMAP kinase cascade, Signaling by ERBB2, Signaling by PDGF, Signaling by SCF-KIT, Smooth Muscle Contraction, VEGFA-VEGFR2 Pathway, VEGFR2 mediated cell proliferation 
                                               SEBELIUS_2_C4636 
                                               VCL 
                                               mRNA 
                                               0 
                                               
                                               
                                               Smooth Muscle Contraction;MAP2K and MAPK activation;Platelet degranulation ;RAFMAP kinase cascade;Downstream signal transduction;VEGFR2 mediated cell proliferation;Signaling by SCF-KIT;Frs2-mediated activation;ARMS-mediated activation;Gastrin-CREB signalling pathway via PKC and MAPK;Interleukin receptor SHC signaling;FCERI mediated MAPK activation;DAP12 signaling;Signaling by PDGF;VEGFA-VEGFR2 Pathway;Signaling by ERBB2;IRS-related events triggered by IGF1R 
                                               hsa04510;hsa04520;hsa04670;hsa04810;hsa05100;hsa05131;hsa05146 
                                               VCL 
                                               NA 
                                               slateblue4 
                                               16 
                                               -2.82692307692308 
                                             
                                             
                                               PPP2R1B-mRNA 
                                               -0.142 
                                               0.0571 
                                               -0.254 
                                               -0.0303 
                                               0.906 
                                               0.839 
                                               0.979 
                                               0.0212 
                                               0.348 
                                               loglinear 
                                               ARMS-mediated activation, AXIN missense mutants destabilize the destruction complex, Beta-catenin phosphorylation cascade, CTLA4 inhibitory signaling, Cyclin AB1 associated events during G2M transition, Cyclin D associated events in G1, DAP12 signaling, DARPP-32 events, Disassembly of the destruction complex and recruitment of AXIN to the membrane, Downstream signal transduction, ERKs are inactivated, FCERI mediated MAPK activation, Frs2-mediated activation, Gastrin-CREB signalling pathway via PKC and MAPK, Interleukin receptor SHC signaling, IRS-related events triggered by IGF1R, MASTL Facilitates Mitotic Progression, Misspliced GSK3beta mutants stabilize beta-catenin, Negative regulation of MAPK pathway, RAF activation, RAFMAP kinase cascade, Resolution of Sister Chromatid Cohesion, RHO GTPases Activate Formins, S33 mutants of beta-catenin aren.SQT.t phosphorylated, S37 mutants of beta-catenin aren.SQT.t phosphorylated, S45 mutants of beta-catenin aren.SQT.t phosphorylated, Signaling by ERBB2, Signaling by PDGF, Signaling by SCF-KIT, T41 mutants of beta-catenin aren.SQT.t phosphorylated, Truncations of AMER1 destabilize the destruction complex, VEGFA-VEGFR2 Pathway, VEGFR2 mediated cell proliferation 
                                               SEBELIUS_2_C4636 
                                               PPP2R1B 
                                               mRNA 
                                               0 
                                               
                                               
                                               MASTL Facilitates Mitotic Progression;Misspliced GSK3beta mutants stabilize beta-catenin;T41 mutants of beta-catenin aren.SQT.t phosphorylated;ERKs are inactivated;Glycolysis;Negative regulation of MAPK pathway;Disassembly of the destruction complex and recruitment of AXIN to the membrane;S37 mutants of beta-catenin aren.SQT.t phosphorylated;Cyclin AB1 associated events during G2M transition;RAF activation;RHO GTPases Activate Formins;S33 mutants of beta-catenin aren.SQT.t phosphorylated;DARPP-32 events;Beta-catenin phosphorylation cascade;Truncations of AMER1 destabilize the destruction complex;CTLA4 inhibitory signaling;AXIN missense mutants destabilize the destruction complex;S45 mutants of beta-catenin aren.SQT.t phosphorylated;Cyclin D associated events in G1;Platelet sensitization by LDL;Resolution of Sister Chromatid Cohesion;Inhibition of replication initiation of damaged DNA by RB1E2F1;RAFMAP kinase cascade;Downstream signal transduction;VEGFR2 mediated cell proliferation;Signaling by SCF-KIT;Frs2-mediated activation;ARMS-mediated activation;Gastrin-CREB signalling pathway via PKC and MAPK;Interleukin receptor SHC signaling;FCERI mediated MAPK activation;DAP12 signaling;Signaling by PDGF;VEGFA-VEGFR2 Pathway;Signaling by ERBB2;IRS-related events triggered by IGF1R 
                                               hsa03015;hsa04071;hsa04114;hsa04151;hsa04152;hsa04261;hsa04350;hsa04390;hsa04530;hsa04728;hsa04730;hsa05142;hsa05160 
                                               PPP2R1B 
                                               NA 
                                               slateblue4 
                                               16 
                                               -2.48686514886165 
                                             
                                             
                                               ACTG1-mRNA 
                                               -0.266 
                                               0.148 
                                               -0.556 
                                               0.0243 
                                               0.832 
                                               0.68 
                                               1.02 
                                               0.0869 
                                               1 
                                               lm.nb 
                                               ARMS-mediated activation, Cell-extracellular matrix interactions, DAP12 signaling, Downstream signal transduction, EPH-ephrin mediated repulsion of cells, EPHB-mediated forward signaling, FCERI mediated MAPK activation, Frs2-mediated activation, Gastrin-CREB signalling pathway via PKC and MAPK, Interleukin receptor SHC signaling, IRS-related events triggered by IGF1R, RAFMAP kinase cascade, Regulation of actin dynamics for phagocytic cup formation, RHO GTPases Activate Formins, RHO GTPases activate IQGAPs, RHO GTPases Activate WASPs and WAVEs, Signaling by ERBB2, Signaling by PDGF, Signaling by SCF-KIT, VEGFA-VEGFR2 Pathway, VEGFR2 mediated cell proliferation 
                                               SEBELIUS_2_C4636 
                                               ACTG1 
                                               mRNA 
                                               0 
                                               
                                               
                                               RHO GTPases Activate WASPs and WAVEs;Translocation of GLUT4 to the plasma membrane;Recycling pathway of L1;Interaction between L1 and Ankyrins;EPHB-mediated forward signaling;EPH-ephrin mediated repulsion of cells;RHO GTPases Activate Formins;RHO GTPases activate IQGAPs;Cell-extracellular matrix interactions;Regulation of actin dynamics for phagocytic cup formation;RAFMAP kinase cascade;Downstream signal transduction;VEGFR2 mediated cell proliferation;Signaling by SCF-KIT;Frs2-mediated activation;ARMS-mediated activation;Gastrin-CREB signalling pathway via PKC and MAPK;Interleukin receptor SHC signaling;FCERI mediated MAPK activation;DAP12 signaling;Signaling by PDGF;VEGFA-VEGFR2 Pathway;Signaling by ERBB2;IRS-related events triggered by IGF1R 
                                               hsa04015;hsa04145;hsa04390;hsa04510;hsa04520;hsa04530;hsa04611;hsa04670;hsa04810;hsa04919;hsa04921;hsa05100;hsa05110;hsa05130;hsa05131;hsa05132;hsa05164;hsa05205;hsa05410;hsa05412;hsa05414;hsa05416 
                                               ACTG1 
                                               NA 
                                               azure4 
                                               1 
                                               -1.7972972972973 
                                             
                                             
                                               KRAS-mRNA 
                                               0.143 
                                               0.0975 
                                               -0.0486 
                                               0.334 
                                               1.1 
                                               0.967 
                                               1.26 
                                               0.159 
                                               1 
                                               loglinear 
                                               ARMS-mediated activation, CD209 (DC-SIGN) signaling, Cell surface interactions at the vascular wall, Constitutive Signaling by EGFRvIII, DAP12 signaling, Downstream signal transduction, FCERI mediated MAPK activation, Frs2-mediated activation, Gastrin-CREB signalling pathway via PKC and MAPK, Interleukin receptor SHC signaling, IRS-related events triggered by IGF1R, MAP2K and MAPK activation, RAFMAP kinase cascade, SHC-mediated cascade.CL.FGFR1, SHC-related events triggered by IGF1R, Signaling by ERBB2, Signaling by FGFR1 in disease, Signaling by FGFR2 in disease, Signaling by FGFR3 in disease, Signaling by FGFR4 in disease, Signaling by PDGF, Signaling by SCF-KIT, Tie2 Signaling, VEGFA-VEGFR2 Pathway, VEGFR2 mediated cell proliferation 
                                               SEBELIUS_2_C4636 
                                               KRAS 
                                               mRNA 
                                               0 
                                               
                                               
                                               Signaling by FGFR2 in disease;Regulation of RAS by GAPs;SHC-mediated cascade.CL.FGFR3;Signaling by FGFR3 in disease;MAP2K and MAPK activation;SHC-related events triggered by IGF1R;Signaling by FGFR4 in disease;SHC-mediated cascade.CL.FGFR1;Tie2 Signaling;Signaling by FGFR1 in disease;Activation of RAS in B cells;SHC-mediated cascade.CL.FGFR4;Constitutive Signaling by EGFRvIII;CD209 (DC-SIGN) signaling;FCERI mediated MAPK activation;RAFMAP kinase cascade;Downstream signal transduction;VEGFR2 mediated cell proliferation;Signaling by SCF-KIT;Frs2-mediated activation;ARMS-mediated activation;Gastrin-CREB signalling pathway via PKC and MAPK;Interleukin receptor SHC signaling;DAP12 signaling;Signaling by PDGF;VEGFA-VEGFR2 Pathway;Signaling by ERBB2;IRS-related events triggered by IGF1R;Cell surface interactions at the vascular wall 
                                               hsa04010;hsa04012;hsa04014;hsa04015;hsa04062;hsa04068;hsa04071;hsa04072;hsa04151;hsa04211;hsa04213;hsa04320;hsa04360;hsa04370;hsa04530;hsa04540;hsa04550;hsa04650;hsa04660;hsa04662;hsa04664;hsa04720;hsa04722;hsa04725;hsa04726;hsa04730;hsa04810;hsa04910;hsa04912;hsa04914;hsa04915;hsa04916;hsa04917;hsa04919;hsa04921;hsa04933;hsa04960;hsa05034;hsa05160;hsa05161;hsa05166;hsa05200;hsa05203;hsa05205;hsa05206;hsa05210;hsa05211;hsa05212;hsa05213;hsa05214;hsa05215;hsa05216;hsa05218;hsa05219;hsa05220;hsa05221;hsa05223;hsa05230;hsa05231 
                                               KRAS 
                                               NA 
                                               azure4 
                                               1 
                                               1.46666666666667 
                                             
                                             
                                               SOS1-mRNA 
                                               -0.119 
                                               0.0837 
                                               -0.283 
                                               0.0453 
                                               0.921 
                                               0.822 
                                               1.03 
                                               0.171 
                                               1 
                                               loglinear 
                                               ARMS-mediated activation, Cell surface interactions at the vascular wall, Constitutive Signaling by EGFRvIII, DAP12 signaling, Downstream signal transduction, FCERI mediated MAPK activation, Frs2-mediated activation, Gastrin-CREB signalling pathway via PKC and MAPK, Integrin alphaIIb beta3 signaling, Interleukin receptor SHC signaling, IRS-related events triggered by IGF1R, NRAGE signals death through JNK, RAFMAP kinase cascade, Role of LAT2NTALLAB on calcium mobilization, SHC-mediated cascade.CL.FGFR1, SHC-related events triggered by IGF1R, Signaling by ERBB2, Signaling by FGFR1 in disease, Signaling by FGFR2 in disease, Signaling by FGFR3 in disease, Signaling by FGFR4 in disease, Signaling by PDGF, Signaling by SCF-KIT, Tie2 Signaling, VEGFA-VEGFR2 Pathway, VEGFR2 mediated cell proliferation 
                                               SEBELIUS_2_C4636 
                                               SOS1 
                                               mRNA 
                                               0 
                                               
                                               
                                               Signaling by FGFR2 in disease;Antigen activates B Cell Receptor (BCR) leading to generation of second messengers;SHC-mediated cascade.CL.FGFR3;Signaling by FGFR3 in disease;RAFMAP kinase cascade;SHC-related events triggered by IGF1R;Signaling by FGFR4 in disease;SHC-mediated cascade.CL.FGFR1;GRB2.CL.SOS provides linkage to MAPK signaling for Integrins ;Tie2 Signaling;Role of LAT2NTALLAB on calcium mobilization;Signaling by FGFR1 in disease;Signal attenuation;NRAGE signals death through JNK;SHC-mediated cascade.CL.FGFR4;FCERI mediated Ca.PLS.2 mobilization;Constitutive Signaling by EGFRvIII;FCERI mediated MAPK activation;Signaling by ERBB2;Signaling by SCF-KIT;DAP12 signaling;Gastrin-CREB signalling pathway via PKC and MAPK;Interleukin receptor SHC signaling;IRS-related events triggered by IGF1R;Downstream signal transduction;VEGFR2 mediated cell proliferation;Frs2-mediated activation;ARMS-mediated activation;Signaling by PDGF;VEGFA-VEGFR2 Pathway;Integrin alphaIIb beta3 signaling;Cell surface interactions at the vascular wall 
                                               hsa04010;hsa04012;hsa04014;hsa04062;hsa04068;hsa04072;hsa04151;hsa04320;hsa04510;hsa04540;hsa04630;hsa04650;hsa04660;hsa04662;hsa04664;hsa04722;hsa04810;hsa04910;hsa04912;hsa04915;hsa04917;hsa05034;hsa05160;hsa05200;hsa05205;hsa05206;hsa05211;hsa05213;hsa05214;hsa05215;hsa05220;hsa05221;hsa05223;hsa05231 
                                               SOS1 
                                               NA 
                                               azure4 
                                               1 
                                               -1.42174432497013 
                                             
                                             
                                               PTK2-mRNA 
                                               -0.0907 
                                               0.0671 
                                               -0.222 
                                               0.0408 
                                               0.939 
                                               0.857 
                                               1.03 
                                               0.191 
                                               1 
                                               loglinear 
                                               Apoptotic cleavage of cellular proteins, ARMS-mediated activation, DAP12 signaling, DCC mediated attractive signaling, Downstream signal transduction, EPHA-mediated growth cone collapse, EPHB-mediated forward signaling, FCERI mediated MAPK activation, Frs2-mediated activation, Gastrin-CREB signalling pathway via PKC and MAPK, Integrin alphaIIb beta3 signaling, Interleukin receptor SHC signaling, IRS-related events triggered by IGF1R, p130Cas linkage to MAPK signaling for integrins, RAFMAP kinase cascade, Regulation of actin dynamics for phagocytic cup formation, RHO GTPases Activate WASPs and WAVEs, Signaling by ERBB2, Signaling by PDGF, Signaling by SCF-KIT, VEGFA-VEGFR2 Pathway, VEGFR2 mediated cell proliferation 
                                               SEBELIUS_2_C4636 
                                               PTK2 
                                               mRNA 
                                               0 
                                               
                                               
                                               Apoptotic cleavage of cellular proteins;RAFMAP kinase cascade;RHO GTPases Activate WASPs and WAVEs;DCC mediated attractive signaling;p130Cas linkage to MAPK signaling for integrins;EPHB-mediated forward signaling;EPHA-mediated growth cone collapse;Regulation of actin dynamics for phagocytic cup formation;Downstream signal transduction;VEGFR2 mediated cell proliferation;Signaling by SCF-KIT;Frs2-mediated activation;ARMS-mediated activation;Gastrin-CREB signalling pathway via PKC and MAPK;Interleukin receptor SHC signaling;FCERI mediated MAPK activation;DAP12 signaling;Signaling by PDGF;VEGFA-VEGFR2 Pathway;Signaling by ERBB2;IRS-related events triggered by IGF1R;Integrin alphaIIb beta3 signaling 
                                               hsa04012;hsa04062;hsa04151;hsa04360;hsa04370;hsa04510;hsa04670;hsa04810;hsa05100;hsa05146;hsa05200;hsa05202;hsa05205;hsa05222 
                                               PTK2 
                                               NA 
                                               azure4 
                                               1 
                                               -1.35171385991058 
                                             
                                             
                                               CRK-mRNA 
                                               -0.133 
                                               0.099 
                                               -0.328 
                                               0.0606 
                                               0.912 
                                               0.797 
                                               1.04 
                                               0.192 
                                               1 
                                               loglinear 
                                               ARMS-mediated activation, Downstream signal transduction, Integrin alphaIIb beta3 signaling, p130Cas linkage to MAPK signaling for integrins, Regulation of actin dynamics for phagocytic cup formation, Regulation of signaling by CBL, Signaling by PDGF, VEGFA-VEGFR2 Pathway 
                                               SEBELIUS_2_C4636 
                                               CRK 
                                               mRNA 
                                               0 
                                               
                                               
                                               ARMS-mediated activation;Downstream signal transduction;Regulation of signaling by CBL;p130Cas linkage to MAPK signaling for integrins;VEGFA-VEGFR2 Pathway;Regulation of actin dynamics for phagocytic cup formation;Signaling by PDGF;Integrin alphaIIb beta3 signaling 
                                               hsa04010;hsa04012;hsa04015;hsa04062;hsa04510;hsa04666;hsa04722;hsa04810;hsa04910;hsa05100;hsa05131;hsa05200;hsa05206;hsa05211;hsa05220 
                                               CRK 
                                               NA 
                                               azure4 
                                               1 
                                               -1.34343434343434 
                                             
                                             
                                               ARAF-mRNA 
                                               -0.213 
                                               0.164 
                                               -0.534 
                                               0.109 
                                               0.863 
                                               0.691 
                                               1.08 
                                               0.209 
                                               1 
                                               loglinear 
                                               ARMS-mediated activation, DAP12 signaling, Downstream signal transduction, FCERI mediated MAPK activation, Frs2-mediated activation, Gastrin-CREB signalling pathway via PKC and MAPK, Interleukin receptor SHC signaling, IRS-related events triggered by IGF1R, MAP2K and MAPK activation, Negative regulation of MAPK pathway, RAF activation, RAFMAP kinase cascade, Signaling by ERBB2, Signaling by PDGF, Signaling by SCF-KIT, VEGFA-VEGFR2 Pathway, VEGFR2 mediated cell proliferation 
                                               SEBELIUS_2_C4636 
                                               ARAF 
                                               mRNA 
                                               0 
                                               
                                               
                                               MAP2K and MAPK activation;RAF activation;Negative regulation of MAPK pathway;RAFMAP kinase cascade;Downstream signal transduction;VEGFR2 mediated cell proliferation;Signaling by SCF-KIT;Frs2-mediated activation;ARMS-mediated activation;Gastrin-CREB signalling pathway via PKC and MAPK;Interleukin receptor SHC signaling;FCERI mediated MAPK activation;DAP12 signaling;Signaling by PDGF;VEGFA-VEGFR2 Pathway;Signaling by ERBB2;IRS-related events triggered by IGF1R 
                                               hsa04012;hsa04068;hsa04270;hsa04650;hsa04720;hsa04726;hsa04730;hsa04810;hsa04910;hsa04914;hsa05034;hsa05160;hsa05200;hsa05205;hsa05210;hsa05211;hsa05212;hsa05213;hsa05214;hsa05215;hsa05218;hsa05219;hsa05220;hsa05221;hsa05223 
                                               ARAF 
                                               NA 
                                               azure4 
                                               1 
                                               -1.29878048780488 
                                             
                                             
                                               ERBB2-mRNA 
                                               0.181 
                                               0.157 
                                               -0.127 
                                               0.489 
                                               1.13 
                                               0.916 
                                               1.4 
                                               0.261 
                                               1 
                                               lm.nb 
                                               ARMS-mediated activation, DAP12 signaling, Downstream signal transduction, FCERI mediated MAPK activation, Frs2-mediated activation, GAB1 signalosome, Gastrin-CREB signalling pathway via PKC and MAPK, Interleukin receptor SHC signaling, IRS-related events triggered by IGF1R, PI3KAKT activation, RAFMAP kinase cascade, Role of LAT2NTALLAB on calcium mobilization, Sema4D induced cell migration and growth-cone collapse, Signaling by ERBB2, Signaling by PDGF, Signaling by SCF-KIT, VEGFA-VEGFR2 Pathway, VEGFR2 mediated cell proliferation 
                                               SEBELIUS_2_C4636 
                                               ERBB2 
                                               mRNA 
                                               0 
                                               
                                               
                                               RAFMAP kinase cascade;GRB7 events in ERBB2 signaling;PLCG1 events in ERBB2 signaling;Sema4D induced cell migration and growth-cone collapse;Signaling by ERBB2;Downstream signal transduction;GAB1 signalosome;Signaling by SCF-KIT;PI3KAKT activation;Role of LAT2NTALLAB on calcium mobilization;DAP12 signaling;Signaling by PDGF;VEGFR2 mediated cell proliferation;Frs2-mediated activation;ARMS-mediated activation;Gastrin-CREB signalling pathway via PKC and MAPK;Interleukin receptor SHC signaling;FCERI mediated MAPK activation;VEGFA-VEGFR2 Pathway;IRS-related events triggered by IGF1R 
                                               hsa04012;hsa04020;hsa04066;hsa04510;hsa04520;hsa05200;hsa05205;hsa05206;hsa05212;hsa05213;hsa05215;hsa05219;hsa05223;hsa05230 
                                               ERBB2 
                                               NA 
                                               azure4 
                                               1 
                                               1.15286624203822 
                                             
                                             
                                               BRAF-mRNA 
                                               0.0635 
                                               0.0578 
                                               -0.0498 
                                               0.177 
                                               1.05 
                                               0.966 
                                               1.13 
                                               0.285 
                                               1 
                                               loglinear 
                                               ARMS-mediated activation, CREB phosphorylation through the activation of Ras, DAP12 signaling, Downstream signal transduction, FCERI mediated MAPK activation, Frs2-mediated activation, Gastrin-CREB signalling pathway via PKC and MAPK, Interleukin receptor SHC signaling, IRS-related events triggered by IGF1R, MAP2K and MAPK activation, Negative feedback regulation of MAPK pathway, Negative regulation of FGFR1 signaling, Negative regulation of FGFR2 signaling, Negative regulation of FGFR3 signaling, Negative regulation of FGFR4 signaling, Negative regulation of MAPK pathway, RAF activation, RAFMAP kinase cascade, Signaling by ERBB2, Signaling by PDGF, Signaling by SCF-KIT, Spry regulation of FGF signaling, VEGFA-VEGFR2 Pathway, VEGFR2 mediated cell proliferation 
                                               SEBELIUS_2_C4636 
                                               BRAF 
                                               mRNA 
                                               0 
                                               
                                               
                                               Spry regulation of FGF signaling;MAP2K and MAPK activation;RAF activation;CREB phosphorylation through the activation of Ras;Negative feedback regulation of MAPK pathway;ARMS-mediated activation;Negative regulation of FGFR3 signaling;Negative regulation of FGFR4 signaling;Negative regulation of FGFR2 signaling;Negative regulation of FGFR1 signaling;RAFMAP kinase cascade;Downstream signal transduction;VEGFR2 mediated cell proliferation;Signaling by SCF-KIT;Frs2-mediated activation;Gastrin-CREB signalling pathway via PKC and MAPK;Interleukin receptor SHC signaling;FCERI mediated MAPK activation;DAP12 signaling;Signaling by PDGF;VEGFA-VEGFR2 Pathway;Signaling by ERBB2;IRS-related events triggered by IGF1R;Negative regulation of MAPK pathway 
                                               hsa04010;hsa04012;hsa04015;hsa04024;hsa04062;hsa04068;hsa04150;hsa04270;hsa04510;hsa04650;hsa04720;hsa04722;hsa04726;hsa04730;hsa04810;hsa04910;hsa04914;hsa05034;hsa05160;hsa05200;hsa05205;hsa05210;hsa05211;hsa05212;hsa05213;hsa05214;hsa05215;hsa05216;hsa05218;hsa05219;hsa05220;hsa05221;hsa05223 
                                               BRAF 
                                               NA 
                                               azure4 
                                               1 
                                               1.09861591695502 
                                             
                                           
                                           DE Results - ARMS-mediated activation genes - New.Annotation: miR890 vs.cel67  Table displaying each sample's global significance scores and directed global significance scores as defined in the heatmaps above. The global significance score is calculated as the square root of the mean squared t-statistic for the genes in a gene set, with t-statistics coming from the linear regression underlying our differential expression analysis. The directed global significance score is calculated as the square root of the mean signed squared t-statistic for the genes in a gene set, with t-statistics coming from the linear regression underlying our differential expression analysis.   
                                      
                                       
                                  
                                
                              
                             
                          
                           
                      
                    
                    
                      
                           
                          
                              New.Annotation: 
                              
                                  New.Annotation: differential expression in Control vs. baseline of cel67  
                                  New.Annotation: differential expression in miR142 vs. baseline of cel67  
                                  New.Annotation: differential expression in miR155 vs. baseline of cel67  
                                  New.Annotation: differential expression in miR18b vs. baseline of cel67  
                                  New.Annotation: differential expression in miR890 vs. baseline of cel67  
                              
                              
                              
                                
                                  
                                       
                                      
                                           Volcano Plot: New.Annotation: Control vs.cel67   More Plot Information                Volcano Plot: New.Annotation: Control vs.cel67  Volcano plot displaying each gene's -log10(p-value) and log2 fold change for the selected covariate. Highly statistically significant genes fall at the top of the plot, and highly differentially expressed genes fall to either side. Genes within the selected gene set are highlighted in orange. Horizontal lines indicate various False Discovery Rate (FDR) thresholds.   
                                           DE Results - AXIN missense mutants destabilize the destruction complex genes - New.Annotation: Control vs.cel67   More Plot Information      Download CSV Data     
                                           
                                             
                                             
                                           
                                           
                                             
                                               Probe Label 
                                               Log2 fold change 
                                               std error (log2) 
                                               Lower confidence limit (log2) 
                                               Upper confidence limit (log2) 
                                               Linear fold change 
                                               Lower confidence limit (linear) 
                                               Upper confidence limit (linear) 
                                               P-value 
                                               BY.p.value 
                                               method 
                                               Gene.sets 
                                               Codeset.Name 
                                               Probe.Label 
                                               Analyte.Type 
                                               Is.Control 
                                               Control.Type 
                                               Related.Probes 
                                               Probe.Annotation 
                                               KEGG.Pathways 
                                               Official.Gene.Name 
                                               Control.Conc 
                                               volcanocol 
                                               volcanopch 
                                               tstats.all 
                                             
                                           
                                           
                                             
                                               CSNK1A1-mRNA 
                                               0.608 
                                               0.0669 
                                               0.477 
                                               0.739 
                                               1.52 
                                               1.39 
                                               1.67 
                                               9.92e-09 
                                               1.32e-05 
                                               loglinear 
                                               AXIN missense mutants destabilize the destruction complex, Beta-catenin phosphorylation cascade, Disassembly of the destruction complex and recruitment of AXIN to the membrane, Misspliced GSK3beta mutants stabilize beta-catenin, S33 mutants of beta-catenin aren.SQT.t phosphorylated, S37 mutants of beta-catenin aren.SQT.t phosphorylated, S45 mutants of beta-catenin aren.SQT.t phosphorylated, T41 mutants of beta-catenin aren.SQT.t phosphorylated 
                                               SEBELIUS_2_C4636 
                                               CSNK1A1 
                                               mRNA 
                                               0 
                                               
                                               
                                               Misspliced GSK3beta mutants stabilize beta-catenin;APC truncation mutants have impaired AXIN binding;T41 mutants of beta-catenin aren.SQT.t phosphorylated;AXIN missense mutants destabilize the destruction complex;S45 mutants of beta-catenin aren.SQT.t phosphorylated;Disassembly of the destruction complex and recruitment of AXIN to the membrane;S37 mutants of beta-catenin aren.SQT.t phosphorylated;S33 mutants of beta-catenin aren.SQT.t phosphorylated;Beta-catenin phosphorylation cascade;Degradation of GLI2 by the proteasome 
                                               hsa04310 
                                               CSNK1A1 
                                               NA 
                                               slateblue1 
                                               16 
                                               9.0881913303438 
                                             
                                             
                                               PPP2CB-mRNA 
                                               -0.182 
                                               0.0747 
                                               -0.328 
                                               -0.0354 
                                               0.882 
                                               0.797 
                                               0.976 
                                               0.0239 
                                               0.756 
                                               loglinear 
                                               ARMS-mediated activation, AXIN missense mutants destabilize the destruction complex, Beta-catenin phosphorylation cascade, CTLA4 inhibitory signaling, Cyclin AB1 associated events during G2M transition, Cyclin D associated events in G1, DAP12 signaling, DARPP-32 events, Disassembly of the destruction complex and recruitment of AXIN to the membrane, Downstream signal transduction, ERKs are inactivated, FCERI mediated MAPK activation, Frs2-mediated activation, Gastrin-CREB signalling pathway via PKC and MAPK, Interleukin receptor SHC signaling, IRS-related events triggered by IGF1R, MASTL Facilitates Mitotic Progression, Misspliced GSK3beta mutants stabilize beta-catenin, Negative regulation of FGFR1 signaling, Negative regulation of FGFR2 signaling, Negative regulation of FGFR3 signaling, Negative regulation of FGFR4 signaling, Negative regulation of MAPK pathway, RAF activation, RAFMAP kinase cascade, Resolution of Sister Chromatid Cohesion, RHO GTPases Activate Formins, S33 mutants of beta-catenin aren.SQT.t phosphorylated, S37 mutants of beta-catenin aren.SQT.t phosphorylated, S45 mutants of beta-catenin aren.SQT.t phosphorylated, Signaling by ERBB2, Signaling by PDGF, Signaling by SCF-KIT, Spry regulation of FGF signaling, T41 mutants of beta-catenin aren.SQT.t phosphorylated, Truncations of AMER1 destabilize the destruction complex, VEGFA-VEGFR2 Pathway, VEGFR2 mediated cell proliferation 
                                               SEBELIUS_2_C4636 
                                               PPP2CB 
                                               mRNA 
                                               0 
                                               
                                               
                                               MASTL Facilitates Mitotic Progression;Misspliced GSK3beta mutants stabilize beta-catenin;T41 mutants of beta-catenin aren.SQT.t phosphorylated;ERKs are inactivated;Glycolysis;Negative regulation of MAPK pathway;Disassembly of the destruction complex and recruitment of AXIN to the membrane;S37 mutants of beta-catenin aren.SQT.t phosphorylated;Cyclin AB1 associated events during G2M transition;RAF activation;RHO GTPases Activate Formins;S33 mutants of beta-catenin aren.SQT.t phosphorylated;DARPP-32 events;Beta-catenin phosphorylation cascade;Spry regulation of FGF signaling;Truncations of AMER1 destabilize the destruction complex;CTLA4 inhibitory signaling;AXIN missense mutants destabilize the destruction complex;S45 mutants of beta-catenin aren.SQT.t phosphorylated;Cyclin D associated events in G1;Platelet sensitization by LDL;Resolution of Sister Chromatid Cohesion;Inhibition of replication initiation of damaged DNA by RB1E2F1;RAFMAP kinase cascade;Downstream signal transduction;VEGFR2 mediated cell proliferation;Signaling by SCF-KIT;Frs2-mediated activation;ARMS-mediated activation;Gastrin-CREB signalling pathway via PKC and MAPK;Interleukin receptor SHC signaling;FCERI mediated MAPK activation;DAP12 signaling;Signaling by PDGF;VEGFA-VEGFR2 Pathway;Signaling by ERBB2;IRS-related events triggered by IGF1R;Negative regulation of FGFR3 signaling;Negative regulation of FGFR4 signaling;Negative regulation of FGFR2 signaling;Negative regulation of FGFR1 signaling 
                                               hsa03015;hsa04071;hsa04114;hsa04151;hsa04152;hsa04261;hsa04350;hsa04390;hsa04530;hsa04728;hsa04730;hsa05142;hsa05160 
                                               PPP2CB 
                                               NA 
                                               azure4 
                                               1 
                                               -2.43641231593039 
                                             
                                             
                                               GSK3B-mRNA 
                                               0.171 
                                               0.0708 
                                               0.0327 
                                               0.31 
                                               1.13 
                                               1.02 
                                               1.24 
                                               0.0245 
                                               0.756 
                                               loglinear 
                                               AXIN missense mutants destabilize the destruction complex, DAP12 signaling, Disassembly of the destruction complex and recruitment of AXIN to the membrane, Downstream signal transduction, GAB1 signalosome, Misspliced GSK3beta mutants stabilize beta-catenin, PI3KAKT activation, PIP3 activates AKT signaling, Regulation of HSF1-mediated heat shock response, Role of LAT2NTALLAB on calcium mobilization, S33 mutants of beta-catenin aren.SQT.t phosphorylated, S37 mutants of beta-catenin aren.SQT.t phosphorylated, S45 mutants of beta-catenin aren.SQT.t phosphorylated, Signaling by ERBB2, Signaling by PDGF, Signaling by SCF-KIT, T41 mutants of beta-catenin aren.SQT.t phosphorylated, Truncations of AMER1 destabilize the destruction complex 
                                               SEBELIUS_2_C4636 
                                               GSK3B 
                                               mRNA 
                                               0 
                                               
                                               
                                               Misspliced GSK3beta mutants stabilize beta-catenin;Constitutive Signaling by AKT1 E17K in Cancer;APC truncation mutants have impaired AXIN binding;T41 mutants of beta-catenin aren.SQT.t phosphorylated;AXIN missense mutants destabilize the destruction complex;S45 mutants of beta-catenin aren.SQT.t phosphorylated;Regulation of HSF1-mediated heat shock response;AKT phosphorylates targets in the cytosol;Disassembly of the destruction complex and recruitment of AXIN to the membrane;S37 mutants of beta-catenin aren.SQT.t phosphorylated;S33 mutants of beta-catenin aren.SQT.t phosphorylated;Truncations of AMER1 destabilize the destruction complex;Degradation of GLI2 by the proteasome;PIP3 activates AKT signaling;Downstream signal transduction;GAB1 signalosome;Signaling by SCF-KIT;PI3KAKT activation;Role of LAT2NTALLAB on calcium mobilization;DAP12 signaling;Signaling by PDGF;Signaling by ERBB2 
                                               hsa04012;hsa04062;hsa04110;hsa04151;hsa04310;hsa04360;hsa04390;hsa04510;hsa04550;hsa04660;hsa04662;hsa04722;hsa04728;hsa04910;hsa04916;hsa04917;hsa04919;hsa04931;hsa04932;hsa05010;hsa05160;hsa05162;hsa05164;hsa05166;hsa05169;hsa05200;hsa05210;hsa05213;hsa05215;hsa05217 
                                               GSK3B 
                                               NA 
                                               azure4 
                                               1 
                                               2.41525423728814 
                                             
                                             
                                               APC-mRNA 
                                               -0.21 
                                               0.0957 
                                               -0.397 
                                               -0.0221 
                                               0.865 
                                               0.759 
                                               0.985 
                                               0.0399 
                                               0.99 
                                               loglinear 
                                               Apoptotic cleavage of cellular proteins, AXIN missense mutants destabilize the destruction complex, Beta-catenin phosphorylation cascade, Deactivation of the beta-catenin transactivating complex, Disassembly of the destruction complex and recruitment of AXIN to the membrane, Misspliced GSK3beta mutants stabilize beta-catenin, S33 mutants of beta-catenin aren.SQT.t phosphorylated, S37 mutants of beta-catenin aren.SQT.t phosphorylated, S45 mutants of beta-catenin aren.SQT.t phosphorylated, T41 mutants of beta-catenin aren.SQT.t phosphorylated, Truncations of AMER1 destabilize the destruction complex 
                                               SEBELIUS_2_C4636 
                                               APC 
                                               mRNA 
                                               0 
                                               
                                               
                                               Apoptotic cleavage of cellular proteins;Misspliced GSK3beta mutants stabilize beta-catenin;APC truncation mutants have impaired AXIN binding;T41 mutants of beta-catenin aren.SQT.t phosphorylated;AXIN missense mutants destabilize the destruction complex;S45 mutants of beta-catenin aren.SQT.t phosphorylated;Deactivation of the beta-catenin transactivating complex;Disassembly of the destruction complex and recruitment of AXIN to the membrane;S37 mutants of beta-catenin aren.SQT.t phosphorylated;S33 mutants of beta-catenin aren.SQT.t phosphorylated;Beta-catenin phosphorylation cascade;Truncations of AMER1 destabilize the destruction complex 
                                               hsa04310;hsa04390;hsa04550;hsa04810;hsa05166;hsa05200;hsa05206;hsa05210;hsa05213;hsa05217 
                                               APC 
                                               NA 
                                               azure4 
                                               1 
                                               -2.19435736677116 
                                             
                                             
                                               PPP2CA-mRNA 
                                               -0.115 
                                               0.0527 
                                               -0.219 
                                               -0.0119 
                                               0.923 
                                               0.859 
                                               0.992 
                                               0.0403 
                                               0.99 
                                               loglinear 
                                               ARMS-mediated activation, AXIN missense mutants destabilize the destruction complex, Beta-catenin phosphorylation cascade, CTLA4 inhibitory signaling, Cyclin AB1 associated events during G2M transition, Cyclin D associated events in G1, DAP12 signaling, DARPP-32 events, Disassembly of the destruction complex and recruitment of AXIN to the membrane, Downstream signal transduction, ERKs are inactivated, FCERI mediated MAPK activation, Frs2-mediated activation, Gastrin-CREB signalling pathway via PKC and MAPK, Interleukin receptor SHC signaling, IRS-related events triggered by IGF1R, MASTL Facilitates Mitotic Progression, Negative regulation of FGFR1 signaling, Negative regulation of FGFR2 signaling, Negative regulation of FGFR3 signaling, Negative regulation of FGFR4 signaling, Negative regulation of MAPK pathway, Nonsense Mediated Decay (NMD) enhanced by the Exon Junction Complex (EJC), RAF activation, RAFMAP kinase cascade, Resolution of Sister Chromatid Cohesion, RHO GTPases Activate Formins, S33 mutants of beta-catenin aren.SQT.t phosphorylated, S37 mutants of beta-catenin aren.SQT.t phosphorylated, S45 mutants of beta-catenin aren.SQT.t phosphorylated, Signaling by ERBB2, Signaling by PDGF, Signaling by SCF-KIT, Spry regulation of FGF signaling, T41 mutants of beta-catenin aren.SQT.t phosphorylated, Truncations of AMER1 destabilize the destruction complex, VEGFA-VEGFR2 Pathway, VEGFR2 mediated cell proliferation 
                                               SEBELIUS_2_C4636 
                                               PPP2CA 
                                               mRNA 
                                               0 
                                               
                                               
                                               MASTL Facilitates Mitotic Progression;T41 mutants of beta-catenin aren.SQT.t phosphorylated;ERKs are inactivated;Glycolysis;Negative regulation of MAPK pathway;Disassembly of the destruction complex and recruitment of AXIN to the membrane;S37 mutants of beta-catenin aren.SQT.t phosphorylated;Cyclin AB1 associated events during G2M transition;RAF activation;RHO GTPases Activate Formins;S33 mutants of beta-catenin aren.SQT.t phosphorylated;DARPP-32 events;Beta-catenin phosphorylation cascade;Initiation of Nuclear Envelope Reformation;Spry regulation of FGF signaling;Truncations of AMER1 destabilize the destruction complex;CTLA4 inhibitory signaling;AXIN missense mutants destabilize the destruction complex;S45 mutants of beta-catenin aren.SQT.t phosphorylated;Nonsense Mediated Decay (NMD) enhanced by the Exon Junction Complex (EJC);Cyclin D associated events in G1;Platelet sensitization by LDL;Resolution of Sister Chromatid Cohesion;Inhibition of replication initiation of damaged DNA by RB1E2F1;RAFMAP kinase cascade;Downstream signal transduction;VEGFR2 mediated cell proliferation;Signaling by SCF-KIT;Frs2-mediated activation;ARMS-mediated activation;Gastrin-CREB signalling pathway via PKC and MAPK;Interleukin receptor SHC signaling;FCERI mediated MAPK activation;DAP12 signaling;Signaling by PDGF;VEGFA-VEGFR2 Pathway;Signaling by ERBB2;IRS-related events triggered by IGF1R;Negative regulation of FGFR3 signaling;Negative regulation of FGFR4 signaling;Negative regulation of FGFR2 signaling;Negative regulation of FGFR1 signaling 
                                               hsa03015;hsa04071;hsa04114;hsa04151;hsa04152;hsa04261;hsa04350;hsa04390;hsa04530;hsa04728;hsa04730;hsa05142;hsa05160 
                                               PPP2CA 
                                               NA 
                                               azure4 
                                               1 
                                               -2.18216318785579 
                                             
                                             
                                               AXIN1-mRNA 
                                               -0.348 
                                               0.214 
                                               -0.768 
                                               0.0712 
                                               0.786 
                                               0.587 
                                               1.05 
                                               0.119 
                                               1 
                                               lm.nb 
                                               AXIN missense mutants destabilize the destruction complex, Beta-catenin phosphorylation cascade, Disassembly of the destruction complex and recruitment of AXIN to the membrane, Misspliced GSK3beta mutants stabilize beta-catenin, S33 mutants of beta-catenin aren.SQT.t phosphorylated, S37 mutants of beta-catenin aren.SQT.t phosphorylated, S45 mutants of beta-catenin aren.SQT.t phosphorylated, T41 mutants of beta-catenin aren.SQT.t phosphorylated, Truncations of AMER1 destabilize the destruction complex 
                                               SEBELIUS_2_C4636 
                                               AXIN1 
                                               mRNA 
                                               0 
                                               
                                               
                                               APC truncation mutants have impaired AXIN binding;Misspliced GSK3beta mutants stabilize beta-catenin;T41 mutants of beta-catenin aren.SQT.t phosphorylated;AXIN missense mutants destabilize the destruction complex;S45 mutants of beta-catenin aren.SQT.t phosphorylated;Disassembly of the destruction complex and recruitment of AXIN to the membrane;S37 mutants of beta-catenin aren.SQT.t phosphorylated;S33 mutants of beta-catenin aren.SQT.t phosphorylated;Beta-catenin phosphorylation cascade;Truncations of AMER1 destabilize the destruction complex 
                                               hsa04310;hsa04390;hsa04550;hsa05200;hsa05210;hsa05213;hsa05217 
                                               AXIN1 
                                               NA 
                                               azure4 
                                               1 
                                               -1.62616822429907 
                                             
                                             
                                               PPP2R1B-mRNA 
                                               -0.0747 
                                               0.0593 
                                               -0.191 
                                               0.0416 
                                               0.95 
                                               0.876 
                                               1.03 
                                               0.222 
                                               1 
                                               loglinear 
                                               ARMS-mediated activation, AXIN missense mutants destabilize the destruction complex, Beta-catenin phosphorylation cascade, CTLA4 inhibitory signaling, Cyclin AB1 associated events during G2M transition, Cyclin D associated events in G1, DAP12 signaling, DARPP-32 events, Disassembly of the destruction complex and recruitment of AXIN to the membrane, Downstream signal transduction, ERKs are inactivated, FCERI mediated MAPK activation, Frs2-mediated activation, Gastrin-CREB signalling pathway via PKC and MAPK, Interleukin receptor SHC signaling, IRS-related events triggered by IGF1R, MASTL Facilitates Mitotic Progression, Misspliced GSK3beta mutants stabilize beta-catenin, Negative regulation of MAPK pathway, RAF activation, RAFMAP kinase cascade, Resolution of Sister Chromatid Cohesion, RHO GTPases Activate Formins, S33 mutants of beta-catenin aren.SQT.t phosphorylated, S37 mutants of beta-catenin aren.SQT.t phosphorylated, S45 mutants of beta-catenin aren.SQT.t phosphorylated, Signaling by ERBB2, Signaling by PDGF, Signaling by SCF-KIT, T41 mutants of beta-catenin aren.SQT.t phosphorylated, Truncations of AMER1 destabilize the destruction complex, VEGFA-VEGFR2 Pathway, VEGFR2 mediated cell proliferation 
                                               SEBELIUS_2_C4636 
                                               PPP2R1B 
                                               mRNA 
                                               0 
                                               
                                               
                                               MASTL Facilitates Mitotic Progression;Misspliced GSK3beta mutants stabilize beta-catenin;T41 mutants of beta-catenin aren.SQT.t phosphorylated;ERKs are inactivated;Glycolysis;Negative regulation of MAPK pathway;Disassembly of the destruction complex and recruitment of AXIN to the membrane;S37 mutants of beta-catenin aren.SQT.t phosphorylated;Cyclin AB1 associated events during G2M transition;RAF activation;RHO GTPases Activate Formins;S33 mutants of beta-catenin aren.SQT.t phosphorylated;DARPP-32 events;Beta-catenin phosphorylation cascade;Truncations of AMER1 destabilize the destruction complex;CTLA4 inhibitory signaling;AXIN missense mutants destabilize the destruction complex;S45 mutants of beta-catenin aren.SQT.t phosphorylated;Cyclin D associated events in G1;Platelet sensitization by LDL;Resolution of Sister Chromatid Cohesion;Inhibition of replication initiation of damaged DNA by RB1E2F1;RAFMAP kinase cascade;Downstream signal transduction;VEGFR2 mediated cell proliferation;Signaling by SCF-KIT;Frs2-mediated activation;ARMS-mediated activation;Gastrin-CREB signalling pathway via PKC and MAPK;Interleukin receptor SHC signaling;FCERI mediated MAPK activation;DAP12 signaling;Signaling by PDGF;VEGFA-VEGFR2 Pathway;Signaling by ERBB2;IRS-related events triggered by IGF1R 
                                               hsa03015;hsa04071;hsa04114;hsa04151;hsa04152;hsa04261;hsa04350;hsa04390;hsa04530;hsa04728;hsa04730;hsa05142;hsa05160 
                                               PPP2R1B 
                                               NA 
                                               azure4 
                                               1 
                                               -1.25969645868465 
                                             
                                             
                                               PPP2R1A-mRNA 
                                               0.0227 
                                               0.0993 
                                               -0.172 
                                               0.217 
                                               1.02 
                                               0.888 
                                               1.16 
                                               0.822 
                                               1 
                                               loglinear 
                                               ARMS-mediated activation, AXIN missense mutants destabilize the destruction complex, Beta-catenin phosphorylation cascade, CTLA4 inhibitory signaling, Cyclin AB1 associated events during G2M transition, Cyclin D associated events in G1, DAP12 signaling, DARPP-32 events, Disassembly of the destruction complex and recruitment of AXIN to the membrane, Downstream signal transduction, FCERI mediated MAPK activation, Frs2-mediated activation, Gastrin-CREB signalling pathway via PKC and MAPK, Interleukin receptor SHC signaling, IRS-related events triggered by IGF1R, MASTL Facilitates Mitotic Progression, Negative regulation of FGFR1 signaling, Negative regulation of FGFR2 signaling, Negative regulation of FGFR3 signaling, Negative regulation of FGFR4 signaling, Nonsense Mediated Decay (NMD) enhanced by the Exon Junction Complex (EJC), RAF activation, RAFMAP kinase cascade, Resolution of Sister Chromatid Cohesion, RHO GTPases Activate Formins, S33 mutants of beta-catenin aren.SQT.t phosphorylated, S37 mutants of beta-catenin aren.SQT.t phosphorylated, S45 mutants of beta-catenin aren.SQT.t phosphorylated, Signaling by ERBB2, Signaling by PDGF, Signaling by SCF-KIT, Spry regulation of FGF signaling, Truncations of AMER1 destabilize the destruction complex, VEGFA-VEGFR2 Pathway, VEGFR2 mediated cell proliferation 
                                               SEBELIUS_2_C4636 
                                               PPP2R1A 
                                               mRNA 
                                               0 
                                               
                                               
                                               MASTL Facilitates Mitotic Progression;Regulation of PLK1 Activity at G2M Transition;Disassembly of the destruction complex and recruitment of AXIN to the membrane;S37 mutants of beta-catenin aren.SQT.t phosphorylated;Cyclin AB1 associated events during G2M transition;RAF activation;RHO GTPases Activate Formins;S33 mutants of beta-catenin aren.SQT.t phosphorylated;Loss of Nlp from mitotic centrosomes;DARPP-32 events;Beta-catenin phosphorylation cascade;Initiation of Nuclear Envelope Reformation;Recruitment of mitotic centrosome proteins and complexes;Spry regulation of FGF signaling;Truncations of AMER1 destabilize the destruction complex;CTLA4 inhibitory signaling;AXIN missense mutants destabilize the destruction complex;S45 mutants of beta-catenin aren.SQT.t phosphorylated;Anchoring of the basal body to the plasma membrane;Nonsense Mediated Decay (NMD) enhanced by the Exon Junction Complex (EJC);Cyclin D associated events in G1;Platelet sensitization by LDL;Resolution of Sister Chromatid Cohesion;Inhibition of replication initiation of damaged DNA by RB1E2F1;RAFMAP kinase cascade;Downstream signal transduction;VEGFR2 mediated cell proliferation;Signaling by SCF-KIT;Frs2-mediated activation;ARMS-mediated activation;Gastrin-CREB signalling pathway via PKC and MAPK;Interleukin receptor SHC signaling;FCERI mediated MAPK activation;DAP12 signaling;Signaling by PDGF;VEGFA-VEGFR2 Pathway;Signaling by ERBB2;IRS-related events triggered by IGF1R;Negative regulation of FGFR3 signaling;Negative regulation of FGFR4 signaling;Negative regulation of FGFR2 signaling;Negative regulation of FGFR1 signaling 
                                               hsa03015;hsa04071;hsa04114;hsa04151;hsa04152;hsa04261;hsa04350;hsa04390;hsa04530;hsa04728;hsa04730;hsa05142;hsa05160 
                                               PPP2R1A 
                                               NA 
                                               azure4 
                                               1 
                                               0.228600201409869 
                                             
                                           
                                           DE Results - AXIN missense mutants destabilize the destruction complex genes - New.Annotation: Control vs.cel67  Table displaying each sample's global significance scores and directed global significance scores as defined in the heatmaps above. The global significance score is calculated as the square root of the mean squared t-statistic for the genes in a gene set, with t-statistics coming from the linear regression underlying our differential expression analysis. The directed global significance score is calculated as the square root of the mean signed squared t-statistic for the genes in a gene set, with t-statistics coming from the linear regression underlying our differential expression analysis.   
                                      
                                       
                                  
                                
                                
                                  
                                       
                                      
                                           Volcano Plot: New.Annotation: miR142 vs.cel67   More Plot Information                Volcano Plot: New.Annotation: miR142 vs.cel67  Volcano plot displaying each gene's -log10(p-value) and log2 fold change for the selected covariate. Highly statistically significant genes fall at the top of the plot, and highly differentially expressed genes fall to either side. Genes within the selected gene set are highlighted in orange. Horizontal lines indicate various False Discovery Rate (FDR) thresholds.   
                                           DE Results - AXIN missense mutants destabilize the destruction complex genes - New.Annotation: miR142 vs.cel67   More Plot Information      Download CSV Data     
                                           
                                             
                                             
                                           
                                           
                                             
                                               Probe Label 
                                               Log2 fold change 
                                               std error (log2) 
                                               Lower confidence limit (log2) 
                                               Upper confidence limit (log2) 
                                               Linear fold change 
                                               Lower confidence limit (linear) 
                                               Upper confidence limit (linear) 
                                               P-value 
                                               BY.p.value 
                                               method 
                                               Gene.sets 
                                               Codeset.Name 
                                               Probe.Label 
                                               Analyte.Type 
                                               Is.Control 
                                               Control.Type 
                                               Related.Probes 
                                               Probe.Annotation 
                                               KEGG.Pathways 
                                               Official.Gene.Name 
                                               Control.Conc 
                                               volcanocol 
                                               volcanopch 
                                               tstats.all 
                                             
                                           
                                           
                                             
                                               CSNK1A1-mRNA 
                                               0.698 
                                               0.0669 
                                               0.567 
                                               0.829 
                                               1.62 
                                               1.48 
                                               1.78 
                                               8.98e-10 
                                               2.38e-07 
                                               loglinear 
                                               AXIN missense mutants destabilize the destruction complex, Beta-catenin phosphorylation cascade, Disassembly of the destruction complex and recruitment of AXIN to the membrane, Misspliced GSK3beta mutants stabilize beta-catenin, S33 mutants of beta-catenin aren.SQT.t phosphorylated, S37 mutants of beta-catenin aren.SQT.t phosphorylated, S45 mutants of beta-catenin aren.SQT.t phosphorylated, T41 mutants of beta-catenin aren.SQT.t phosphorylated 
                                               SEBELIUS_2_C4636 
                                               CSNK1A1 
                                               mRNA 
                                               0 
                                               
                                               
                                               Misspliced GSK3beta mutants stabilize beta-catenin;APC truncation mutants have impaired AXIN binding;T41 mutants of beta-catenin aren.SQT.t phosphorylated;AXIN missense mutants destabilize the destruction complex;S45 mutants of beta-catenin aren.SQT.t phosphorylated;Disassembly of the destruction complex and recruitment of AXIN to the membrane;S37 mutants of beta-catenin aren.SQT.t phosphorylated;S33 mutants of beta-catenin aren.SQT.t phosphorylated;Beta-catenin phosphorylation cascade;Degradation of GLI2 by the proteasome 
                                               hsa04310 
                                               CSNK1A1 
                                               NA 
                                               slateblue1 
                                               16 
                                               10.4334828101644 
                                             
                                             
                                               GSK3B-mRNA 
                                               0.553 
                                               0.0708 
                                               0.414 
                                               0.691 
                                               1.47 
                                               1.33 
                                               1.61 
                                               1.2e-07 
                                               1.45e-05 
                                               loglinear 
                                               AXIN missense mutants destabilize the destruction complex, DAP12 signaling, Disassembly of the destruction complex and recruitment of AXIN to the membrane, Downstream signal transduction, GAB1 signalosome, Misspliced GSK3beta mutants stabilize beta-catenin, PI3KAKT activation, PIP3 activates AKT signaling, Regulation of HSF1-mediated heat shock response, Role of LAT2NTALLAB on calcium mobilization, S33 mutants of beta-catenin aren.SQT.t phosphorylated, S37 mutants of beta-catenin aren.SQT.t phosphorylated, S45 mutants of beta-catenin aren.SQT.t phosphorylated, Signaling by ERBB2, Signaling by PDGF, Signaling by SCF-KIT, T41 mutants of beta-catenin aren.SQT.t phosphorylated, Truncations of AMER1 destabilize the destruction complex 
                                               SEBELIUS_2_C4636 
                                               GSK3B 
                                               mRNA 
                                               0 
                                               
                                               
                                               Misspliced GSK3beta mutants stabilize beta-catenin;Constitutive Signaling by AKT1 E17K in Cancer;APC truncation mutants have impaired AXIN binding;T41 mutants of beta-catenin aren.SQT.t phosphorylated;AXIN missense mutants destabilize the destruction complex;S45 mutants of beta-catenin aren.SQT.t phosphorylated;Regulation of HSF1-mediated heat shock response;AKT phosphorylates targets in the cytosol;Disassembly of the destruction complex and recruitment of AXIN to the membrane;S37 mutants of beta-catenin aren.SQT.t phosphorylated;S33 mutants of beta-catenin aren.SQT.t phosphorylated;Truncations of AMER1 destabilize the destruction complex;Degradation of GLI2 by the proteasome;PIP3 activates AKT signaling;Downstream signal transduction;GAB1 signalosome;Signaling by SCF-KIT;PI3KAKT activation;Role of LAT2NTALLAB on calcium mobilization;DAP12 signaling;Signaling by PDGF;Signaling by ERBB2 
                                               hsa04012;hsa04062;hsa04110;hsa04151;hsa04310;hsa04360;hsa04390;hsa04510;hsa04550;hsa04660;hsa04662;hsa04722;hsa04728;hsa04910;hsa04916;hsa04917;hsa04919;hsa04931;hsa04932;hsa05010;hsa05160;hsa05162;hsa05164;hsa05166;hsa05169;hsa05200;hsa05210;hsa05213;hsa05215;hsa05217 
                                               GSK3B 
                                               NA 
                                               slateblue1 
                                               16 
                                               7.81073446327684 
                                             
                                             
                                               APC-mRNA 
                                               -0.593 
                                               0.0957 
                                               -0.781 
                                               -0.406 
                                               0.663 
                                               0.582 
                                               0.755 
                                               3.8e-06 
                                               0.000196 
                                               loglinear 
                                               Apoptotic cleavage of cellular proteins, AXIN missense mutants destabilize the destruction complex, Beta-catenin phosphorylation cascade, Deactivation of the beta-catenin transactivating complex, Disassembly of the destruction complex and recruitment of AXIN to the membrane, Misspliced GSK3beta mutants stabilize beta-catenin, S33 mutants of beta-catenin aren.SQT.t phosphorylated, S37 mutants of beta-catenin aren.SQT.t phosphorylated, S45 mutants of beta-catenin aren.SQT.t phosphorylated, T41 mutants of beta-catenin aren.SQT.t phosphorylated, Truncations of AMER1 destabilize the destruction complex 
                                               SEBELIUS_2_C4636 
                                               APC 
                                               mRNA 
                                               0 
                                               
                                               
                                               Apoptotic cleavage of cellular proteins;Misspliced GSK3beta mutants stabilize beta-catenin;APC truncation mutants have impaired AXIN binding;T41 mutants of beta-catenin aren.SQT.t phosphorylated;AXIN missense mutants destabilize the destruction complex;S45 mutants of beta-catenin aren.SQT.t phosphorylated;Deactivation of the beta-catenin transactivating complex;Disassembly of the destruction complex and recruitment of AXIN to the membrane;S37 mutants of beta-catenin aren.SQT.t phosphorylated;S33 mutants of beta-catenin aren.SQT.t phosphorylated;Beta-catenin phosphorylation cascade;Truncations of AMER1 destabilize the destruction complex 
                                               hsa04310;hsa04390;hsa04550;hsa04810;hsa05166;hsa05200;hsa05206;hsa05210;hsa05213;hsa05217 
                                               APC 
                                               NA 
                                               slateblue1 
                                               16 
                                               -6.19644723092999 
                                             
                                             
                                               PPP2CA-mRNA 
                                               -0.205 
                                               0.0527 
                                               -0.309 
                                               -0.102 
                                               0.867 
                                               0.807 
                                               0.932 
                                               0.000827 
                                               0.0186 
                                               loglinear 
                                               ARMS-mediated activation, AXIN missense mutants destabilize the destruction complex, Beta-catenin phosphorylation cascade, CTLA4 inhibitory signaling, Cyclin AB1 associated events during G2M transition, Cyclin D associated events in G1, DAP12 signaling, DARPP-32 events, Disassembly of the destruction complex and recruitment of AXIN to the membrane, Downstream signal transduction, ERKs are inactivated, FCERI mediated MAPK activation, Frs2-mediated activation, Gastrin-CREB signalling pathway via PKC and MAPK, Interleukin receptor SHC signaling, IRS-related events triggered by IGF1R, MASTL Facilitates Mitotic Progression, Negative regulation of FGFR1 signaling, Negative regulation of FGFR2 signaling, Negative regulation of FGFR3 signaling, Negative regulation of FGFR4 signaling, Negative regulation of MAPK pathway, Nonsense Mediated Decay (NMD) enhanced by the Exon Junction Complex (EJC), RAF activation, RAFMAP kinase cascade, Resolution of Sister Chromatid Cohesion, RHO GTPases Activate Formins, S33 mutants of beta-catenin aren.SQT.t phosphorylated, S37 mutants of beta-catenin aren.SQT.t phosphorylated, S45 mutants of beta-catenin aren.SQT.t phosphorylated, Signaling by ERBB2, Signaling by PDGF, Signaling by SCF-KIT, Spry regulation of FGF signaling, T41 mutants of beta-catenin aren.SQT.t phosphorylated, Truncations of AMER1 destabilize the destruction complex, VEGFA-VEGFR2 Pathway, VEGFR2 mediated cell proliferation 
                                               SEBELIUS_2_C4636 
                                               PPP2CA 
                                               mRNA 
                                               0 
                                               
                                               
                                               MASTL Facilitates Mitotic Progression;T41 mutants of beta-catenin aren.SQT.t phosphorylated;ERKs are inactivated;Glycolysis;Negative regulation of MAPK pathway;Disassembly of the destruction complex and recruitment of AXIN to the membrane;S37 mutants of beta-catenin aren.SQT.t phosphorylated;Cyclin AB1 associated events during G2M transition;RAF activation;RHO GTPases Activate Formins;S33 mutants of beta-catenin aren.SQT.t phosphorylated;DARPP-32 events;Beta-catenin phosphorylation cascade;Initiation of Nuclear Envelope Reformation;Spry regulation of FGF signaling;Truncations of AMER1 destabilize the destruction complex;CTLA4 inhibitory signaling;AXIN missense mutants destabilize the destruction complex;S45 mutants of beta-catenin aren.SQT.t phosphorylated;Nonsense Mediated Decay (NMD) enhanced by the Exon Junction Complex (EJC);Cyclin D associated events in G1;Platelet sensitization by LDL;Resolution of Sister Chromatid Cohesion;Inhibition of replication initiation of damaged DNA by RB1E2F1;RAFMAP kinase cascade;Downstream signal transduction;VEGFR2 mediated cell proliferation;Signaling by SCF-KIT;Frs2-mediated activation;ARMS-mediated activation;Gastrin-CREB signalling pathway via PKC and MAPK;Interleukin receptor SHC signaling;FCERI mediated MAPK activation;DAP12 signaling;Signaling by PDGF;VEGFA-VEGFR2 Pathway;Signaling by ERBB2;IRS-related events triggered by IGF1R;Negative regulation of FGFR3 signaling;Negative regulation of FGFR4 signaling;Negative regulation of FGFR2 signaling;Negative regulation of FGFR1 signaling 
                                               hsa03015;hsa04071;hsa04114;hsa04151;hsa04152;hsa04261;hsa04350;hsa04390;hsa04530;hsa04728;hsa04730;hsa05142;hsa05160 
                                               PPP2CA 
                                               NA 
                                               slateblue2 
                                               16 
                                               -3.8899430740038 
                                             
                                             
                                               PPP2R1A-mRNA 
                                               -0.339 
                                               0.0993 
                                               -0.533 
                                               -0.144 
                                               0.791 
                                               0.691 
                                               0.905 
                                               0.00262 
                                               0.0492 
                                               loglinear 
                                               ARMS-mediated activation, AXIN missense mutants destabilize the destruction complex, Beta-catenin phosphorylation cascade, CTLA4 inhibitory signaling, Cyclin AB1 associated events during G2M transition, Cyclin D associated events in G1, DAP12 signaling, DARPP-32 events, Disassembly of the destruction complex and recruitment of AXIN to the membrane, Downstream signal transduction, FCERI mediated MAPK activation, Frs2-mediated activation, Gastrin-CREB signalling pathway via PKC and MAPK, Interleukin receptor SHC signaling, IRS-related events triggered by IGF1R, MASTL Facilitates Mitotic Progression, Negative regulation of FGFR1 signaling, Negative regulation of FGFR2 signaling, Negative regulation of FGFR3 signaling, Negative regulation of FGFR4 signaling, Nonsense Mediated Decay (NMD) enhanced by the Exon Junction Complex (EJC), RAF activation, RAFMAP kinase cascade, Resolution of Sister Chromatid Cohesion, RHO GTPases Activate Formins, S33 mutants of beta-catenin aren.SQT.t phosphorylated, S37 mutants of beta-catenin aren.SQT.t phosphorylated, S45 mutants of beta-catenin aren.SQT.t phosphorylated, Signaling by ERBB2, Signaling by PDGF, Signaling by SCF-KIT, Spry regulation of FGF signaling, Truncations of AMER1 destabilize the destruction complex, VEGFA-VEGFR2 Pathway, VEGFR2 mediated cell proliferation 
                                               SEBELIUS_2_C4636 
                                               PPP2R1A 
                                               mRNA 
                                               0 
                                               
                                               
                                               MASTL Facilitates Mitotic Progression;Regulation of PLK1 Activity at G2M Transition;Disassembly of the destruction complex and recruitment of AXIN to the membrane;S37 mutants of beta-catenin aren.SQT.t phosphorylated;Cyclin AB1 associated events during G2M transition;RAF activation;RHO GTPases Activate Formins;S33 mutants of beta-catenin aren.SQT.t phosphorylated;Loss of Nlp from mitotic centrosomes;DARPP-32 events;Beta-catenin phosphorylation cascade;Initiation of Nuclear Envelope Reformation;Recruitment of mitotic centrosome proteins and complexes;Spry regulation of FGF signaling;Truncations of AMER1 destabilize the destruction complex;CTLA4 inhibitory signaling;AXIN missense mutants destabilize the destruction complex;S45 mutants of beta-catenin aren.SQT.t phosphorylated;Anchoring of the basal body to the plasma membrane;Nonsense Mediated Decay (NMD) enhanced by the Exon Junction Complex (EJC);Cyclin D associated events in G1;Platelet sensitization by LDL;Resolution of Sister Chromatid Cohesion;Inhibition of replication initiation of damaged DNA by RB1E2F1;RAFMAP kinase cascade;Downstream signal transduction;VEGFR2 mediated cell proliferation;Signaling by SCF-KIT;Frs2-mediated activation;ARMS-mediated activation;Gastrin-CREB signalling pathway via PKC and MAPK;Interleukin receptor SHC signaling;FCERI mediated MAPK activation;DAP12 signaling;Signaling by PDGF;VEGFA-VEGFR2 Pathway;Signaling by ERBB2;IRS-related events triggered by IGF1R;Negative regulation of FGFR3 signaling;Negative regulation of FGFR4 signaling;Negative regulation of FGFR2 signaling;Negative regulation of FGFR1 signaling 
                                               hsa03015;hsa04071;hsa04114;hsa04151;hsa04152;hsa04261;hsa04350;hsa04390;hsa04530;hsa04728;hsa04730;hsa05142;hsa05160 
                                               PPP2R1A 
                                               NA 
                                               slateblue2 
                                               16 
                                               -3.41389728096677 
                                             
                                             
                                               PPP2R1B-mRNA 
                                               0.0724 
                                               0.0593 
                                               -0.0439 
                                               0.189 
                                               1.05 
                                               0.97 
                                               1.14 
                                               0.236 
                                               1 
                                               loglinear 
                                               ARMS-mediated activation, AXIN missense mutants destabilize the destruction complex, Beta-catenin phosphorylation cascade, CTLA4 inhibitory signaling, Cyclin AB1 associated events during G2M transition, Cyclin D associated events in G1, DAP12 signaling, DARPP-32 events, Disassembly of the destruction complex and recruitment of AXIN to the membrane, Downstream signal transduction, ERKs are inactivated, FCERI mediated MAPK activation, Frs2-mediated activation, Gastrin-CREB signalling pathway via PKC and MAPK, Interleukin receptor SHC signaling, IRS-related events triggered by IGF1R, MASTL Facilitates Mitotic Progression, Misspliced GSK3beta mutants stabilize beta-catenin, Negative regulation of MAPK pathway, RAF activation, RAFMAP kinase cascade, Resolution of Sister Chromatid Cohesion, RHO GTPases Activate Formins, S33 mutants of beta-catenin aren.SQT.t phosphorylated, S37 mutants of beta-catenin aren.SQT.t phosphorylated, S45 mutants of beta-catenin aren.SQT.t phosphorylated, Signaling by ERBB2, Signaling by PDGF, Signaling by SCF-KIT, T41 mutants of beta-catenin aren.SQT.t phosphorylated, Truncations of AMER1 destabilize the destruction complex, VEGFA-VEGFR2 Pathway, VEGFR2 mediated cell proliferation 
                                               SEBELIUS_2_C4636 
                                               PPP2R1B 
                                               mRNA 
                                               0 
                                               
                                               
                                               MASTL Facilitates Mitotic Progression;Misspliced GSK3beta mutants stabilize beta-catenin;T41 mutants of beta-catenin aren.SQT.t phosphorylated;ERKs are inactivated;Glycolysis;Negative regulation of MAPK pathway;Disassembly of the destruction complex and recruitment of AXIN to the membrane;S37 mutants of beta-catenin aren.SQT.t phosphorylated;Cyclin AB1 associated events during G2M transition;RAF activation;RHO GTPases Activate Formins;S33 mutants of beta-catenin aren.SQT.t phosphorylated;DARPP-32 events;Beta-catenin phosphorylation cascade;Truncations of AMER1 destabilize the destruction complex;CTLA4 inhibitory signaling;AXIN missense mutants destabilize the destruction complex;S45 mutants of beta-catenin aren.SQT.t phosphorylated;Cyclin D associated events in G1;Platelet sensitization by LDL;Resolution of Sister Chromatid Cohesion;Inhibition of replication initiation of damaged DNA by RB1E2F1;RAFMAP kinase cascade;Downstream signal transduction;VEGFR2 mediated cell proliferation;Signaling by SCF-KIT;Frs2-mediated activation;ARMS-mediated activation;Gastrin-CREB signalling pathway via PKC and MAPK;Interleukin receptor SHC signaling;FCERI mediated MAPK activation;DAP12 signaling;Signaling by PDGF;VEGFA-VEGFR2 Pathway;Signaling by ERBB2;IRS-related events triggered by IGF1R 
                                               hsa03015;hsa04071;hsa04114;hsa04151;hsa04152;hsa04261;hsa04350;hsa04390;hsa04530;hsa04728;hsa04730;hsa05142;hsa05160 
                                               PPP2R1B 
                                               NA 
                                               azure4 
                                               1 
                                               1.22091062394604 
                                             
                                             
                                               AXIN1-mRNA 
                                               -0.15 
                                               0.211 
                                               -0.564 
                                               0.264 
                                               0.901 
                                               0.677 
                                               1.2 
                                               0.486 
                                               1 
                                               lm.nb 
                                               AXIN missense mutants destabilize the destruction complex, Beta-catenin phosphorylation cascade, Disassembly of the destruction complex and recruitment of AXIN to the membrane, Misspliced GSK3beta mutants stabilize beta-catenin, S33 mutants of beta-catenin aren.SQT.t phosphorylated, S37 mutants of beta-catenin aren.SQT.t phosphorylated, S45 mutants of beta-catenin aren.SQT.t phosphorylated, T41 mutants of beta-catenin aren.SQT.t phosphorylated, Truncations of AMER1 destabilize the destruction complex 
                                               SEBELIUS_2_C4636 
                                               AXIN1 
                                               mRNA 
                                               0 
                                               
                                               
                                               APC truncation mutants have impaired AXIN binding;Misspliced GSK3beta mutants stabilize beta-catenin;T41 mutants of beta-catenin aren.SQT.t phosphorylated;AXIN missense mutants destabilize the destruction complex;S45 mutants of beta-catenin aren.SQT.t phosphorylated;Disassembly of the destruction complex and recruitment of AXIN to the membrane;S37 mutants of beta-catenin aren.SQT.t phosphorylated;S33 mutants of beta-catenin aren.SQT.t phosphorylated;Beta-catenin phosphorylation cascade;Truncations of AMER1 destabilize the destruction complex 
                                               hsa04310;hsa04390;hsa04550;hsa05200;hsa05210;hsa05213;hsa05217 
                                               AXIN1 
                                               NA 
                                               azure4 
                                               1 
                                               -0.710900473933649 
                                             
                                             
                                               PPP2CB-mRNA 
                                               -0.0393 
                                               0.0747 
                                               -0.186 
                                               0.107 
                                               0.973 
                                               0.879 
                                               1.08 
                                               0.604 
                                               1 
                                               loglinear 
                                               ARMS-mediated activation, AXIN missense mutants destabilize the destruction complex, Beta-catenin phosphorylation cascade, CTLA4 inhibitory signaling, Cyclin AB1 associated events during G2M transition, Cyclin D associated events in G1, DAP12 signaling, DARPP-32 events, Disassembly of the destruction complex and recruitment of AXIN to the membrane, Downstream signal transduction, ERKs are inactivated, FCERI mediated MAPK activation, Frs2-mediated activation, Gastrin-CREB signalling pathway via PKC and MAPK, Interleukin receptor SHC signaling, IRS-related events triggered by IGF1R, MASTL Facilitates Mitotic Progression, Misspliced GSK3beta mutants stabilize beta-catenin, Negative regulation of FGFR1 signaling, Negative regulation of FGFR2 signaling, Negative regulation of FGFR3 signaling, Negative regulation of FGFR4 signaling, Negative regulation of MAPK pathway, RAF activation, RAFMAP kinase cascade, Resolution of Sister Chromatid Cohesion, RHO GTPases Activate Formins, S33 mutants of beta-catenin aren.SQT.t phosphorylated, S37 mutants of beta-catenin aren.SQT.t phosphorylated, S45 mutants of beta-catenin aren.SQT.t phosphorylated, Signaling by ERBB2, Signaling by PDGF, Signaling by SCF-KIT, Spry regulation of FGF signaling, T41 mutants of beta-catenin aren.SQT.t phosphorylated, Truncations of AMER1 destabilize the destruction complex, VEGFA-VEGFR2 Pathway, VEGFR2 mediated cell proliferation 
                                               SEBELIUS_2_C4636 
                                               PPP2CB 
                                               mRNA 
                                               0 
                                               
                                               
                                               MASTL Facilitates Mitotic Progression;Misspliced GSK3beta mutants stabilize beta-catenin;T41 mutants of beta-catenin aren.SQT.t phosphorylated;ERKs are inactivated;Glycolysis;Negative regulation of MAPK pathway;Disassembly of the destruction complex and recruitment of AXIN to the membrane;S37 mutants of beta-catenin aren.SQT.t phosphorylated;Cyclin AB1 associated events during G2M transition;RAF activation;RHO GTPases Activate Formins;S33 mutants of beta-catenin aren.SQT.t phosphorylated;DARPP-32 events;Beta-catenin phosphorylation cascade;Spry regulation of FGF signaling;Truncations of AMER1 destabilize the destruction complex;CTLA4 inhibitory signaling;AXIN missense mutants destabilize the destruction complex;S45 mutants of beta-catenin aren.SQT.t phosphorylated;Cyclin D associated events in G1;Platelet sensitization by LDL;Resolution of Sister Chromatid Cohesion;Inhibition of replication initiation of damaged DNA by RB1E2F1;RAFMAP kinase cascade;Downstream signal transduction;VEGFR2 mediated cell proliferation;Signaling by SCF-KIT;Frs2-mediated activation;ARMS-mediated activation;Gastrin-CREB signalling pathway via PKC and MAPK;Interleukin receptor SHC signaling;FCERI mediated MAPK activation;DAP12 signaling;Signaling by PDGF;VEGFA-VEGFR2 Pathway;Signaling by ERBB2;IRS-related events triggered by IGF1R;Negative regulation of FGFR3 signaling;Negative regulation of FGFR4 signaling;Negative regulation of FGFR2 signaling;Negative regulation of FGFR1 signaling 
                                               hsa03015;hsa04071;hsa04114;hsa04151;hsa04152;hsa04261;hsa04350;hsa04390;hsa04530;hsa04728;hsa04730;hsa05142;hsa05160 
                                               PPP2CB 
                                               NA 
                                               azure4 
                                               1 
                                               -0.526104417670683 
                                             
                                           
                                           DE Results - AXIN missense mutants destabilize the destruction complex genes - New.Annotation: miR142 vs.cel67  Table displaying each sample's global significance scores and directed global significance scores as defined in the heatmaps above. The global significance score is calculated as the square root of the mean squared t-statistic for the genes in a gene set, with t-statistics coming from the linear regression underlying our differential expression analysis. The directed global significance score is calculated as the square root of the mean signed squared t-statistic for the genes in a gene set, with t-statistics coming from the linear regression underlying our differential expression analysis.   
                                      
                                       
                                  
                                
                                
                                  
                                       
                                      
                                           Volcano Plot: New.Annotation: miR155 vs.cel67   More Plot Information                Volcano Plot: New.Annotation: miR155 vs.cel67  Volcano plot displaying each gene's -log10(p-value) and log2 fold change for the selected covariate. Highly statistically significant genes fall at the top of the plot, and highly differentially expressed genes fall to either side. Genes within the selected gene set are highlighted in orange. Horizontal lines indicate various False Discovery Rate (FDR) thresholds.   
                                           DE Results - AXIN missense mutants destabilize the destruction complex genes - New.Annotation: miR155 vs.cel67   More Plot Information      Download CSV Data     
                                           
                                             
                                             
                                           
                                           
                                             
                                               Probe Label 
                                               Log2 fold change 
                                               std error (log2) 
                                               Lower confidence limit (log2) 
                                               Upper confidence limit (log2) 
                                               Linear fold change 
                                               Lower confidence limit (linear) 
                                               Upper confidence limit (linear) 
                                               P-value 
                                               BY.p.value 
                                               method 
                                               Gene.sets 
                                               Codeset.Name 
                                               Probe.Label 
                                               Analyte.Type 
                                               Is.Control 
                                               Control.Type 
                                               Related.Probes 
                                               Probe.Annotation 
                                               KEGG.Pathways 
                                               Official.Gene.Name 
                                               Control.Conc 
                                               volcanocol 
                                               volcanopch 
                                               tstats.all 
                                             
                                           
                                           
                                             
                                               PPP2CB-mRNA 
                                               -1.13 
                                               0.0787 
                                               -1.29 
                                               -0.979 
                                               0.456 
                                               0.41 
                                               0.507 
                                               2.34e-12 
                                               3.11e-09 
                                               loglinear 
                                               ARMS-mediated activation, AXIN missense mutants destabilize the destruction complex, Beta-catenin phosphorylation cascade, CTLA4 inhibitory signaling, Cyclin AB1 associated events during G2M transition, Cyclin D associated events in G1, DAP12 signaling, DARPP-32 events, Disassembly of the destruction complex and recruitment of AXIN to the membrane, Downstream signal transduction, ERKs are inactivated, FCERI mediated MAPK activation, Frs2-mediated activation, Gastrin-CREB signalling pathway via PKC and MAPK, Interleukin receptor SHC signaling, IRS-related events triggered by IGF1R, MASTL Facilitates Mitotic Progression, Misspliced GSK3beta mutants stabilize beta-catenin, Negative regulation of FGFR1 signaling, Negative regulation of FGFR2 signaling, Negative regulation of FGFR3 signaling, Negative regulation of FGFR4 signaling, Negative regulation of MAPK pathway, RAF activation, RAFMAP kinase cascade, Resolution of Sister Chromatid Cohesion, RHO GTPases Activate Formins, S33 mutants of beta-catenin aren.SQT.t phosphorylated, S37 mutants of beta-catenin aren.SQT.t phosphorylated, S45 mutants of beta-catenin aren.SQT.t phosphorylated, Signaling by ERBB2, Signaling by PDGF, Signaling by SCF-KIT, Spry regulation of FGF signaling, T41 mutants of beta-catenin aren.SQT.t phosphorylated, Truncations of AMER1 destabilize the destruction complex, VEGFA-VEGFR2 Pathway, VEGFR2 mediated cell proliferation 
                                               SEBELIUS_2_C4636 
                                               PPP2CB 
                                               mRNA 
                                               0 
                                               
                                               
                                               MASTL Facilitates Mitotic Progression;Misspliced GSK3beta mutants stabilize beta-catenin;T41 mutants of beta-catenin aren.SQT.t phosphorylated;ERKs are inactivated;Glycolysis;Negative regulation of MAPK pathway;Disassembly of the destruction complex and recruitment of AXIN to the membrane;S37 mutants of beta-catenin aren.SQT.t phosphorylated;Cyclin AB1 associated events during G2M transition;RAF activation;RHO GTPases Activate Formins;S33 mutants of beta-catenin aren.SQT.t phosphorylated;DARPP-32 events;Beta-catenin phosphorylation cascade;Spry regulation of FGF signaling;Truncations of AMER1 destabilize the destruction complex;CTLA4 inhibitory signaling;AXIN missense mutants destabilize the destruction complex;S45 mutants of beta-catenin aren.SQT.t phosphorylated;Cyclin D associated events in G1;Platelet sensitization by LDL;Resolution of Sister Chromatid Cohesion;Inhibition of replication initiation of damaged DNA by RB1E2F1;RAFMAP kinase cascade;Downstream signal transduction;VEGFR2 mediated cell proliferation;Signaling by SCF-KIT;Frs2-mediated activation;ARMS-mediated activation;Gastrin-CREB signalling pathway via PKC and MAPK;Interleukin receptor SHC signaling;FCERI mediated MAPK activation;DAP12 signaling;Signaling by PDGF;VEGFA-VEGFR2 Pathway;Signaling by ERBB2;IRS-related events triggered by IGF1R;Negative regulation of FGFR3 signaling;Negative regulation of FGFR4 signaling;Negative regulation of FGFR2 signaling;Negative regulation of FGFR1 signaling 
                                               hsa03015;hsa04071;hsa04114;hsa04151;hsa04152;hsa04261;hsa04350;hsa04390;hsa04530;hsa04728;hsa04730;hsa05142;hsa05160 
                                               PPP2CB 
                                               NA 
                                               slateblue1 
                                               16 
                                               -14.3583227445997 
                                             
                                             
                                               CSNK1A1-mRNA 
                                               -0.601 
                                               0.0705 
                                               -0.739 
                                               -0.463 
                                               0.659 
                                               0.599 
                                               0.725 
                                               2.89e-08 
                                               6.39e-06 
                                               loglinear 
                                               AXIN missense mutants destabilize the destruction complex, Beta-catenin phosphorylation cascade, Disassembly of the destruction complex and recruitment of AXIN to the membrane, Misspliced GSK3beta mutants stabilize beta-catenin, S33 mutants of beta-catenin aren.SQT.t phosphorylated, S37 mutants of beta-catenin aren.SQT.t phosphorylated, S45 mutants of beta-catenin aren.SQT.t phosphorylated, T41 mutants of beta-catenin aren.SQT.t phosphorylated 
                                               SEBELIUS_2_C4636 
                                               CSNK1A1 
                                               mRNA 
                                               0 
                                               
                                               
                                               Misspliced GSK3beta mutants stabilize beta-catenin;APC truncation mutants have impaired AXIN binding;T41 mutants of beta-catenin aren.SQT.t phosphorylated;AXIN missense mutants destabilize the destruction complex;S45 mutants of beta-catenin aren.SQT.t phosphorylated;Disassembly of the destruction complex and recruitment of AXIN to the membrane;S37 mutants of beta-catenin aren.SQT.t phosphorylated;S33 mutants of beta-catenin aren.SQT.t phosphorylated;Beta-catenin phosphorylation cascade;Degradation of GLI2 by the proteasome 
                                               hsa04310 
                                               CSNK1A1 
                                               NA 
                                               slateblue1 
                                               16 
                                               -8.52482269503546 
                                             
                                             
                                               PPP2CA-mRNA 
                                               -0.221 
                                               0.0556 
                                               -0.33 
                                               -0.112 
                                               0.858 
                                               0.796 
                                               0.925 
                                               0.000682 
                                               0.0404 
                                               loglinear 
                                               ARMS-mediated activation, AXIN missense mutants destabilize the destruction complex, Beta-catenin phosphorylation cascade, CTLA4 inhibitory signaling, Cyclin AB1 associated events during G2M transition, Cyclin D associated events in G1, DAP12 signaling, DARPP-32 events, Disassembly of the destruction complex and recruitment of AXIN to the membrane, Downstream signal transduction, ERKs are inactivated, FCERI mediated MAPK activation, Frs2-mediated activation, Gastrin-CREB signalling pathway via PKC and MAPK, Interleukin receptor SHC signaling, IRS-related events triggered by IGF1R, MASTL Facilitates Mitotic Progression, Negative regulation of FGFR1 signaling, Negative regulation of FGFR2 signaling, Negative regulation of FGFR3 signaling, Negative regulation of FGFR4 signaling, Negative regulation of MAPK pathway, Nonsense Mediated Decay (NMD) enhanced by the Exon Junction Complex (EJC), RAF activation, RAFMAP kinase cascade, Resolution of Sister Chromatid Cohesion, RHO GTPases Activate Formins, S33 mutants of beta-catenin aren.SQT.t phosphorylated, S37 mutants of beta-catenin aren.SQT.t phosphorylated, S45 mutants of beta-catenin aren.SQT.t phosphorylated, Signaling by ERBB2, Signaling by PDGF, Signaling by SCF-KIT, Spry regulation of FGF signaling, T41 mutants of beta-catenin aren.SQT.t phosphorylated, Truncations of AMER1 destabilize the destruction complex, VEGFA-VEGFR2 Pathway, VEGFR2 mediated cell proliferation 
                                               SEBELIUS_2_C4636 
                                               PPP2CA 
                                               mRNA 
                                               0 
                                               
                                               
                                               MASTL Facilitates Mitotic Progression;T41 mutants of beta-catenin aren.SQT.t phosphorylated;ERKs are inactivated;Glycolysis;Negative regulation of MAPK pathway;Disassembly of the destruction complex and recruitment of AXIN to the membrane;S37 mutants of beta-catenin aren.SQT.t phosphorylated;Cyclin AB1 associated events during G2M transition;RAF activation;RHO GTPases Activate Formins;S33 mutants of beta-catenin aren.SQT.t phosphorylated;DARPP-32 events;Beta-catenin phosphorylation cascade;Initiation of Nuclear Envelope Reformation;Spry regulation of FGF signaling;Truncations of AMER1 destabilize the destruction complex;CTLA4 inhibitory signaling;AXIN missense mutants destabilize the destruction complex;S45 mutants of beta-catenin aren.SQT.t phosphorylated;Nonsense Mediated Decay (NMD) enhanced by the Exon Junction Complex (EJC);Cyclin D associated events in G1;Platelet sensitization by LDL;Resolution of Sister Chromatid Cohesion;Inhibition of replication initiation of damaged DNA by RB1E2F1;RAFMAP kinase cascade;Downstream signal transduction;VEGFR2 mediated cell proliferation;Signaling by SCF-KIT;Frs2-mediated activation;ARMS-mediated activation;Gastrin-CREB signalling pathway via PKC and MAPK;Interleukin receptor SHC signaling;FCERI mediated MAPK activation;DAP12 signaling;Signaling by PDGF;VEGFA-VEGFR2 Pathway;Signaling by ERBB2;IRS-related events triggered by IGF1R;Negative regulation of FGFR3 signaling;Negative regulation of FGFR4 signaling;Negative regulation of FGFR2 signaling;Negative regulation of FGFR1 signaling 
                                               hsa03015;hsa04071;hsa04114;hsa04151;hsa04152;hsa04261;hsa04350;hsa04390;hsa04530;hsa04728;hsa04730;hsa05142;hsa05160 
                                               PPP2CA 
                                               NA 
                                               slateblue2 
                                               16 
                                               -3.97482014388489 
                                             
                                             
                                               APC-mRNA 
                                               0.261 
                                               0.101 
                                               0.0632 
                                               0.459 
                                               1.2 
                                               1.04 
                                               1.37 
                                               0.0172 
                                               0.327 
                                               loglinear 
                                               Apoptotic cleavage of cellular proteins, AXIN missense mutants destabilize the destruction complex, Beta-catenin phosphorylation cascade, Deactivation of the beta-catenin transactivating complex, Disassembly of the destruction complex and recruitment of AXIN to the membrane, Misspliced GSK3beta mutants stabilize beta-catenin, S33 mutants of beta-catenin aren.SQT.t phosphorylated, S37 mutants of beta-catenin aren.SQT.t phosphorylated, S45 mutants of beta-catenin aren.SQT.t phosphorylated, T41 mutants of beta-catenin aren.SQT.t phosphorylated, Truncations of AMER1 destabilize the destruction complex 
                                               SEBELIUS_2_C4636 
                                               APC 
                                               mRNA 
                                               0 
                                               
                                               
                                               Apoptotic cleavage of cellular proteins;Misspliced GSK3beta mutants stabilize beta-catenin;APC truncation mutants have impaired AXIN binding;T41 mutants of beta-catenin aren.SQT.t phosphorylated;AXIN missense mutants destabilize the destruction complex;S45 mutants of beta-catenin aren.SQT.t phosphorylated;Deactivation of the beta-catenin transactivating complex;Disassembly of the destruction complex and recruitment of AXIN to the membrane;S37 mutants of beta-catenin aren.SQT.t phosphorylated;S33 mutants of beta-catenin aren.SQT.t phosphorylated;Beta-catenin phosphorylation cascade;Truncations of AMER1 destabilize the destruction complex 
                                               hsa04310;hsa04390;hsa04550;hsa04810;hsa05166;hsa05200;hsa05206;hsa05210;hsa05213;hsa05217 
                                               APC 
                                               NA 
                                               slateblue4 
                                               16 
                                               2.58415841584158 
                                             
                                             
                                               GSK3B-mRNA 
                                               0.164 
                                               0.0746 
                                               0.0182 
                                               0.311 
                                               1.12 
                                               1.01 
                                               1.24 
                                               0.0388 
                                               0.678 
                                               loglinear 
                                               AXIN missense mutants destabilize the destruction complex, DAP12 signaling, Disassembly of the destruction complex and recruitment of AXIN to the membrane, Downstream signal transduction, GAB1 signalosome, Misspliced GSK3beta mutants stabilize beta-catenin, PI3KAKT activation, PIP3 activates AKT signaling, Regulation of HSF1-mediated heat shock response, Role of LAT2NTALLAB on calcium mobilization, S33 mutants of beta-catenin aren.SQT.t phosphorylated, S37 mutants of beta-catenin aren.SQT.t phosphorylated, S45 mutants of beta-catenin aren.SQT.t phosphorylated, Signaling by ERBB2, Signaling by PDGF, Signaling by SCF-KIT, T41 mutants of beta-catenin aren.SQT.t phosphorylated, Truncations of AMER1 destabilize the destruction complex 
                                               SEBELIUS_2_C4636 
                                               GSK3B 
                                               mRNA 
                                               0 
                                               
                                               
                                               Misspliced GSK3beta mutants stabilize beta-catenin;Constitutive Signaling by AKT1 E17K in Cancer;APC truncation mutants have impaired AXIN binding;T41 mutants of beta-catenin aren.SQT.t phosphorylated;AXIN missense mutants destabilize the destruction complex;S45 mutants of beta-catenin aren.SQT.t phosphorylated;Regulation of HSF1-mediated heat shock response;AKT phosphorylates targets in the cytosol;Disassembly of the destruction complex and recruitment of AXIN to the membrane;S37 mutants of beta-catenin aren.SQT.t phosphorylated;S33 mutants of beta-catenin aren.SQT.t phosphorylated;Truncations of AMER1 destabilize the destruction complex;Degradation of GLI2 by the proteasome;PIP3 activates AKT signaling;Downstream signal transduction;GAB1 signalosome;Signaling by SCF-KIT;PI3KAKT activation;Role of LAT2NTALLAB on calcium mobilization;DAP12 signaling;Signaling by PDGF;Signaling by ERBB2 
                                               hsa04012;hsa04062;hsa04110;hsa04151;hsa04310;hsa04360;hsa04390;hsa04510;hsa04550;hsa04660;hsa04662;hsa04722;hsa04728;hsa04910;hsa04916;hsa04917;hsa04919;hsa04931;hsa04932;hsa05010;hsa05160;hsa05162;hsa05164;hsa05166;hsa05169;hsa05200;hsa05210;hsa05213;hsa05215;hsa05217 
                                               GSK3B 
                                               NA 
                                               azure4 
                                               1 
                                               2.19839142091153 
                                             
                                             
                                               PPP2R1B-mRNA 
                                               -0.108 
                                               0.0625 
                                               -0.23 
                                               0.0148 
                                               0.928 
                                               0.852 
                                               1.01 
                                               0.0996 
                                               1 
                                               loglinear 
                                               ARMS-mediated activation, AXIN missense mutants destabilize the destruction complex, Beta-catenin phosphorylation cascade, CTLA4 inhibitory signaling, Cyclin AB1 associated events during G2M transition, Cyclin D associated events in G1, DAP12 signaling, DARPP-32 events, Disassembly of the destruction complex and recruitment of AXIN to the membrane, Downstream signal transduction, ERKs are inactivated, FCERI mediated MAPK activation, Frs2-mediated activation, Gastrin-CREB signalling pathway via PKC and MAPK, Interleukin receptor SHC signaling, IRS-related events triggered by IGF1R, MASTL Facilitates Mitotic Progression, Misspliced GSK3beta mutants stabilize beta-catenin, Negative regulation of MAPK pathway, RAF activation, RAFMAP kinase cascade, Resolution of Sister Chromatid Cohesion, RHO GTPases Activate Formins, S33 mutants of beta-catenin aren.SQT.t phosphorylated, S37 mutants of beta-catenin aren.SQT.t phosphorylated, S45 mutants of beta-catenin aren.SQT.t phosphorylated, Signaling by ERBB2, Signaling by PDGF, Signaling by SCF-KIT, T41 mutants of beta-catenin aren.SQT.t phosphorylated, Truncations of AMER1 destabilize the destruction complex, VEGFA-VEGFR2 Pathway, VEGFR2 mediated cell proliferation 
                                               SEBELIUS_2_C4636 
                                               PPP2R1B 
                                               mRNA 
                                               0 
                                               
                                               
                                               MASTL Facilitates Mitotic Progression;Misspliced GSK3beta mutants stabilize beta-catenin;T41 mutants of beta-catenin aren.SQT.t phosphorylated;ERKs are inactivated;Glycolysis;Negative regulation of MAPK pathway;Disassembly of the destruction complex and recruitment of AXIN to the membrane;S37 mutants of beta-catenin aren.SQT.t phosphorylated;Cyclin AB1 associated events during G2M transition;RAF activation;RHO GTPases Activate Formins;S33 mutants of beta-catenin aren.SQT.t phosphorylated;DARPP-32 events;Beta-catenin phosphorylation cascade;Truncations of AMER1 destabilize the destruction complex;CTLA4 inhibitory signaling;AXIN missense mutants destabilize the destruction complex;S45 mutants of beta-catenin aren.SQT.t phosphorylated;Cyclin D associated events in G1;Platelet sensitization by LDL;Resolution of Sister Chromatid Cohesion;Inhibition of replication initiation of damaged DNA by RB1E2F1;RAFMAP kinase cascade;Downstream signal transduction;VEGFR2 mediated cell proliferation;Signaling by SCF-KIT;Frs2-mediated activation;ARMS-mediated activation;Gastrin-CREB signalling pathway via PKC and MAPK;Interleukin receptor SHC signaling;FCERI mediated MAPK activation;DAP12 signaling;Signaling by PDGF;VEGFA-VEGFR2 Pathway;Signaling by ERBB2;IRS-related events triggered by IGF1R 
                                               hsa03015;hsa04071;hsa04114;hsa04151;hsa04152;hsa04261;hsa04350;hsa04390;hsa04530;hsa04728;hsa04730;hsa05142;hsa05160 
                                               PPP2R1B 
                                               NA 
                                               azure4 
                                               1 
                                               -1.728 
                                             
                                             
                                               AXIN1-mRNA 
                                               -0.353 
                                               0.226 
                                               -0.797 
                                               0.0904 
                                               0.783 
                                               0.576 
                                               1.06 
                                               0.134 
                                               1 
                                               lm.nb 
                                               AXIN missense mutants destabilize the destruction complex, Beta-catenin phosphorylation cascade, Disassembly of the destruction complex and recruitment of AXIN to the membrane, Misspliced GSK3beta mutants stabilize beta-catenin, S33 mutants of beta-catenin aren.SQT.t phosphorylated, S37 mutants of beta-catenin aren.SQT.t phosphorylated, S45 mutants of beta-catenin aren.SQT.t phosphorylated, T41 mutants of beta-catenin aren.SQT.t phosphorylated, Truncations of AMER1 destabilize the destruction complex 
                                               SEBELIUS_2_C4636 
                                               AXIN1 
                                               mRNA 
                                               0 
                                               
                                               
                                               APC truncation mutants have impaired AXIN binding;Misspliced GSK3beta mutants stabilize beta-catenin;T41 mutants of beta-catenin aren.SQT.t phosphorylated;AXIN missense mutants destabilize the destruction complex;S45 mutants of beta-catenin aren.SQT.t phosphorylated;Disassembly of the destruction complex and recruitment of AXIN to the membrane;S37 mutants of beta-catenin aren.SQT.t phosphorylated;S33 mutants of beta-catenin aren.SQT.t phosphorylated;Beta-catenin phosphorylation cascade;Truncations of AMER1 destabilize the destruction complex 
                                               hsa04310;hsa04390;hsa04550;hsa05200;hsa05210;hsa05213;hsa05217 
                                               AXIN1 
                                               NA 
                                               azure4 
                                               1 
                                               -1.56194690265487 
                                             
                                             
                                               PPP2R1A-mRNA 
                                               0.122 
                                               0.105 
                                               -0.0832 
                                               0.327 
                                               1.09 
                                               0.944 
                                               1.25 
                                               0.257 
                                               1 
                                               loglinear 
                                               ARMS-mediated activation, AXIN missense mutants destabilize the destruction complex, Beta-catenin phosphorylation cascade, CTLA4 inhibitory signaling, Cyclin AB1 associated events during G2M transition, Cyclin D associated events in G1, DAP12 signaling, DARPP-32 events, Disassembly of the destruction complex and recruitment of AXIN to the membrane, Downstream signal transduction, FCERI mediated MAPK activation, Frs2-mediated activation, Gastrin-CREB signalling pathway via PKC and MAPK, Interleukin receptor SHC signaling, IRS-related events triggered by IGF1R, MASTL Facilitates Mitotic Progression, Negative regulation of FGFR1 signaling, Negative regulation of FGFR2 signaling, Negative regulation of FGFR3 signaling, Negative regulation of FGFR4 signaling, Nonsense Mediated Decay (NMD) enhanced by the Exon Junction Complex (EJC), RAF activation, RAFMAP kinase cascade, Resolution of Sister Chromatid Cohesion, RHO GTPases Activate Formins, S33 mutants of beta-catenin aren.SQT.t phosphorylated, S37 mutants of beta-catenin aren.SQT.t phosphorylated, S45 mutants of beta-catenin aren.SQT.t phosphorylated, Signaling by ERBB2, Signaling by PDGF, Signaling by SCF-KIT, Spry regulation of FGF signaling, Truncations of AMER1 destabilize the destruction complex, VEGFA-VEGFR2 Pathway, VEGFR2 mediated cell proliferation 
                                               SEBELIUS_2_C4636 
                                               PPP2R1A 
                                               mRNA 
                                               0 
                                               
                                               
                                               MASTL Facilitates Mitotic Progression;Regulation of PLK1 Activity at G2M Transition;Disassembly of the destruction complex and recruitment of AXIN to the membrane;S37 mutants of beta-catenin aren.SQT.t phosphorylated;Cyclin AB1 associated events during G2M transition;RAF activation;RHO GTPases Activate Formins;S33 mutants of beta-catenin aren.SQT.t phosphorylated;Loss of Nlp from mitotic centrosomes;DARPP-32 events;Beta-catenin phosphorylation cascade;Initiation of Nuclear Envelope Reformation;Recruitment of mitotic centrosome proteins and complexes;Spry regulation of FGF signaling;Truncations of AMER1 destabilize the destruction complex;CTLA4 inhibitory signaling;AXIN missense mutants destabilize the destruction complex;S45 mutants of beta-catenin aren.SQT.t phosphorylated;Anchoring of the basal body to the plasma membrane;Nonsense Mediated Decay (NMD) enhanced by the Exon Junction Complex (EJC);Cyclin D associated events in G1;Platelet sensitization by LDL;Resolution of Sister Chromatid Cohesion;Inhibition of replication initiation of damaged DNA by RB1E2F1;RAFMAP kinase cascade;Downstream signal transduction;VEGFR2 mediated cell proliferation;Signaling by SCF-KIT;Frs2-mediated activation;ARMS-mediated activation;Gastrin-CREB signalling pathway via PKC and MAPK;Interleukin receptor SHC signaling;FCERI mediated MAPK activation;DAP12 signaling;Signaling by PDGF;VEGFA-VEGFR2 Pathway;Signaling by ERBB2;IRS-related events triggered by IGF1R;Negative regulation of FGFR3 signaling;Negative regulation of FGFR4 signaling;Negative regulation of FGFR2 signaling;Negative regulation of FGFR1 signaling 
                                               hsa03015;hsa04071;hsa04114;hsa04151;hsa04152;hsa04261;hsa04350;hsa04390;hsa04530;hsa04728;hsa04730;hsa05142;hsa05160 
                                               PPP2R1A 
                                               NA 
                                               azure4 
                                               1 
                                               1.16190476190476 
                                             
                                           
                                           DE Results - AXIN missense mutants destabilize the destruction complex genes - New.Annotation: miR155 vs.cel67  Table displaying each sample's global significance scores and directed global significance scores as defined in the heatmaps above. The global significance score is calculated as the square root of the mean squared t-statistic for the genes in a gene set, with t-statistics coming from the linear regression underlying our differential expression analysis. The directed global significance score is calculated as the square root of the mean signed squared t-statistic for the genes in a gene set, with t-statistics coming from the linear regression underlying our differential expression analysis.   
                                      
                                       
                                  
                                
                                
                                  
                                       
                                      
                                           Volcano Plot: New.Annotation: miR18b vs.cel67   More Plot Information                Volcano Plot: New.Annotation: miR18b vs.cel67  Volcano plot displaying each gene's -log10(p-value) and log2 fold change for the selected covariate. Highly statistically significant genes fall at the top of the plot, and highly differentially expressed genes fall to either side. Genes within the selected gene set are highlighted in orange. Horizontal lines indicate various False Discovery Rate (FDR) thresholds.   
                                           DE Results - AXIN missense mutants destabilize the destruction complex genes - New.Annotation: miR18b vs.cel67   More Plot Information      Download CSV Data     
                                           
                                             
                                             
                                           
                                           
                                             
                                               Probe Label 
                                               Log2 fold change 
                                               std error (log2) 
                                               Lower confidence limit (log2) 
                                               Upper confidence limit (log2) 
                                               Linear fold change 
                                               Lower confidence limit (linear) 
                                               Upper confidence limit (linear) 
                                               P-value 
                                               BY.p.value 
                                               method 
                                               Gene.sets 
                                               Codeset.Name 
                                               Probe.Label 
                                               Analyte.Type 
                                               Is.Control 
                                               Control.Type 
                                               Related.Probes 
                                               Probe.Annotation 
                                               KEGG.Pathways 
                                               Official.Gene.Name 
                                               Control.Conc 
                                               volcanocol 
                                               volcanopch 
                                               tstats.all 
                                             
                                           
                                           
                                             
                                               CSNK1A1-mRNA 
                                               0.584 
                                               0.0761 
                                               0.435 
                                               0.733 
                                               1.5 
                                               1.35 
                                               1.66 
                                               1.6e-07 
                                               2.36e-05 
                                               loglinear 
                                               AXIN missense mutants destabilize the destruction complex, Beta-catenin phosphorylation cascade, Disassembly of the destruction complex and recruitment of AXIN to the membrane, Misspliced GSK3beta mutants stabilize beta-catenin, S33 mutants of beta-catenin aren.SQT.t phosphorylated, S37 mutants of beta-catenin aren.SQT.t phosphorylated, S45 mutants of beta-catenin aren.SQT.t phosphorylated, T41 mutants of beta-catenin aren.SQT.t phosphorylated 
                                               SEBELIUS_2_C4636 
                                               CSNK1A1 
                                               mRNA 
                                               0 
                                               
                                               
                                               Misspliced GSK3beta mutants stabilize beta-catenin;APC truncation mutants have impaired AXIN binding;T41 mutants of beta-catenin aren.SQT.t phosphorylated;AXIN missense mutants destabilize the destruction complex;S45 mutants of beta-catenin aren.SQT.t phosphorylated;Disassembly of the destruction complex and recruitment of AXIN to the membrane;S37 mutants of beta-catenin aren.SQT.t phosphorylated;S33 mutants of beta-catenin aren.SQT.t phosphorylated;Beta-catenin phosphorylation cascade;Degradation of GLI2 by the proteasome 
                                               hsa04310 
                                               CSNK1A1 
                                               NA 
                                               slateblue1 
                                               16 
                                               7.67411300919842 
                                             
                                             
                                               PPP2R1B-mRNA 
                                               0.385 
                                               0.0675 
                                               0.253 
                                               0.518 
                                               1.31 
                                               1.19 
                                               1.43 
                                               1.15e-05 
                                               0.000638 
                                               loglinear 
                                               ARMS-mediated activation, AXIN missense mutants destabilize the destruction complex, Beta-catenin phosphorylation cascade, CTLA4 inhibitory signaling, Cyclin AB1 associated events during G2M transition, Cyclin D associated events in G1, DAP12 signaling, DARPP-32 events, Disassembly of the destruction complex and recruitment of AXIN to the membrane, Downstream signal transduction, ERKs are inactivated, FCERI mediated MAPK activation, Frs2-mediated activation, Gastrin-CREB signalling pathway via PKC and MAPK, Interleukin receptor SHC signaling, IRS-related events triggered by IGF1R, MASTL Facilitates Mitotic Progression, Misspliced GSK3beta mutants stabilize beta-catenin, Negative regulation of MAPK pathway, RAF activation, RAFMAP kinase cascade, Resolution of Sister Chromatid Cohesion, RHO GTPases Activate Formins, S33 mutants of beta-catenin aren.SQT.t phosphorylated, S37 mutants of beta-catenin aren.SQT.t phosphorylated, S45 mutants of beta-catenin aren.SQT.t phosphorylated, Signaling by ERBB2, Signaling by PDGF, Signaling by SCF-KIT, T41 mutants of beta-catenin aren.SQT.t phosphorylated, Truncations of AMER1 destabilize the destruction complex, VEGFA-VEGFR2 Pathway, VEGFR2 mediated cell proliferation 
                                               SEBELIUS_2_C4636 
                                               PPP2R1B 
                                               mRNA 
                                               0 
                                               
                                               
                                               MASTL Facilitates Mitotic Progression;Misspliced GSK3beta mutants stabilize beta-catenin;T41 mutants of beta-catenin aren.SQT.t phosphorylated;ERKs are inactivated;Glycolysis;Negative regulation of MAPK pathway;Disassembly of the destruction complex and recruitment of AXIN to the membrane;S37 mutants of beta-catenin aren.SQT.t phosphorylated;Cyclin AB1 associated events during G2M transition;RAF activation;RHO GTPases Activate Formins;S33 mutants of beta-catenin aren.SQT.t phosphorylated;DARPP-32 events;Beta-catenin phosphorylation cascade;Truncations of AMER1 destabilize the destruction complex;CTLA4 inhibitory signaling;AXIN missense mutants destabilize the destruction complex;S45 mutants of beta-catenin aren.SQT.t phosphorylated;Cyclin D associated events in G1;Platelet sensitization by LDL;Resolution of Sister Chromatid Cohesion;Inhibition of replication initiation of damaged DNA by RB1E2F1;RAFMAP kinase cascade;Downstream signal transduction;VEGFR2 mediated cell proliferation;Signaling by SCF-KIT;Frs2-mediated activation;ARMS-mediated activation;Gastrin-CREB signalling pathway via PKC and MAPK;Interleukin receptor SHC signaling;FCERI mediated MAPK activation;DAP12 signaling;Signaling by PDGF;VEGFA-VEGFR2 Pathway;Signaling by ERBB2;IRS-related events triggered by IGF1R 
                                               hsa03015;hsa04071;hsa04114;hsa04151;hsa04152;hsa04261;hsa04350;hsa04390;hsa04530;hsa04728;hsa04730;hsa05142;hsa05160 
                                               PPP2R1B 
                                               NA 
                                               slateblue1 
                                               16 
                                               5.7037037037037 
                                             
                                             
                                               PPP2CB-mRNA 
                                               0.253 
                                               0.085 
                                               0.0862 
                                               0.419 
                                               1.19 
                                               1.06 
                                               1.34 
                                               0.00724 
                                               0.148 
                                               loglinear 
                                               ARMS-mediated activation, AXIN missense mutants destabilize the destruction complex, Beta-catenin phosphorylation cascade, CTLA4 inhibitory signaling, Cyclin AB1 associated events during G2M transition, Cyclin D associated events in G1, DAP12 signaling, DARPP-32 events, Disassembly of the destruction complex and recruitment of AXIN to the membrane, Downstream signal transduction, ERKs are inactivated, FCERI mediated MAPK activation, Frs2-mediated activation, Gastrin-CREB signalling pathway via PKC and MAPK, Interleukin receptor SHC signaling, IRS-related events triggered by IGF1R, MASTL Facilitates Mitotic Progression, Misspliced GSK3beta mutants stabilize beta-catenin, Negative regulation of FGFR1 signaling, Negative regulation of FGFR2 signaling, Negative regulation of FGFR3 signaling, Negative regulation of FGFR4 signaling, Negative regulation of MAPK pathway, RAF activation, RAFMAP kinase cascade, Resolution of Sister Chromatid Cohesion, RHO GTPases Activate Formins, S33 mutants of beta-catenin aren.SQT.t phosphorylated, S37 mutants of beta-catenin aren.SQT.t phosphorylated, S45 mutants of beta-catenin aren.SQT.t phosphorylated, Signaling by ERBB2, Signaling by PDGF, Signaling by SCF-KIT, Spry regulation of FGF signaling, T41 mutants of beta-catenin aren.SQT.t phosphorylated, Truncations of AMER1 destabilize the destruction complex, VEGFA-VEGFR2 Pathway, VEGFR2 mediated cell proliferation 
                                               SEBELIUS_2_C4636 
                                               PPP2CB 
                                               mRNA 
                                               0 
                                               
                                               
                                               MASTL Facilitates Mitotic Progression;Misspliced GSK3beta mutants stabilize beta-catenin;T41 mutants of beta-catenin aren.SQT.t phosphorylated;ERKs are inactivated;Glycolysis;Negative regulation of MAPK pathway;Disassembly of the destruction complex and recruitment of AXIN to the membrane;S37 mutants of beta-catenin aren.SQT.t phosphorylated;Cyclin AB1 associated events during G2M transition;RAF activation;RHO GTPases Activate Formins;S33 mutants of beta-catenin aren.SQT.t phosphorylated;DARPP-32 events;Beta-catenin phosphorylation cascade;Spry regulation of FGF signaling;Truncations of AMER1 destabilize the destruction complex;CTLA4 inhibitory signaling;AXIN missense mutants destabilize the destruction complex;S45 mutants of beta-catenin aren.SQT.t phosphorylated;Cyclin D associated events in G1;Platelet sensitization by LDL;Resolution of Sister Chromatid Cohesion;Inhibition of replication initiation of damaged DNA by RB1E2F1;RAFMAP kinase cascade;Downstream signal transduction;VEGFR2 mediated cell proliferation;Signaling by SCF-KIT;Frs2-mediated activation;ARMS-mediated activation;Gastrin-CREB signalling pathway via PKC and MAPK;Interleukin receptor SHC signaling;FCERI mediated MAPK activation;DAP12 signaling;Signaling by PDGF;VEGFA-VEGFR2 Pathway;Signaling by ERBB2;IRS-related events triggered by IGF1R;Negative regulation of FGFR3 signaling;Negative regulation of FGFR4 signaling;Negative regulation of FGFR2 signaling;Negative regulation of FGFR1 signaling 
                                               hsa03015;hsa04071;hsa04114;hsa04151;hsa04152;hsa04261;hsa04350;hsa04390;hsa04530;hsa04728;hsa04730;hsa05142;hsa05160 
                                               PPP2CB 
                                               NA 
                                               slateblue4 
                                               16 
                                               2.97647058823529 
                                             
                                             
                                               PPP2CA-mRNA 
                                               -0.173 
                                               0.06 
                                               -0.29 
                                               -0.0549 
                                               0.887 
                                               0.818 
                                               0.963 
                                               0.00906 
                                               0.174 
                                               loglinear 
                                               ARMS-mediated activation, AXIN missense mutants destabilize the destruction complex, Beta-catenin phosphorylation cascade, CTLA4 inhibitory signaling, Cyclin AB1 associated events during G2M transition, Cyclin D associated events in G1, DAP12 signaling, DARPP-32 events, Disassembly of the destruction complex and recruitment of AXIN to the membrane, Downstream signal transduction, ERKs are inactivated, FCERI mediated MAPK activation, Frs2-mediated activation, Gastrin-CREB signalling pathway via PKC and MAPK, Interleukin receptor SHC signaling, IRS-related events triggered by IGF1R, MASTL Facilitates Mitotic Progression, Negative regulation of FGFR1 signaling, Negative regulation of FGFR2 signaling, Negative regulation of FGFR3 signaling, Negative regulation of FGFR4 signaling, Negative regulation of MAPK pathway, Nonsense Mediated Decay (NMD) enhanced by the Exon Junction Complex (EJC), RAF activation, RAFMAP kinase cascade, Resolution of Sister Chromatid Cohesion, RHO GTPases Activate Formins, S33 mutants of beta-catenin aren.SQT.t phosphorylated, S37 mutants of beta-catenin aren.SQT.t phosphorylated, S45 mutants of beta-catenin aren.SQT.t phosphorylated, Signaling by ERBB2, Signaling by PDGF, Signaling by SCF-KIT, Spry regulation of FGF signaling, T41 mutants of beta-catenin aren.SQT.t phosphorylated, Truncations of AMER1 destabilize the destruction complex, VEGFA-VEGFR2 Pathway, VEGFR2 mediated cell proliferation 
                                               SEBELIUS_2_C4636 
                                               PPP2CA 
                                               mRNA 
                                               0 
                                               
                                               
                                               MASTL Facilitates Mitotic Progression;T41 mutants of beta-catenin aren.SQT.t phosphorylated;ERKs are inactivated;Glycolysis;Negative regulation of MAPK pathway;Disassembly of the destruction complex and recruitment of AXIN to the membrane;S37 mutants of beta-catenin aren.SQT.t phosphorylated;Cyclin AB1 associated events during G2M transition;RAF activation;RHO GTPases Activate Formins;S33 mutants of beta-catenin aren.SQT.t phosphorylated;DARPP-32 events;Beta-catenin phosphorylation cascade;Initiation of Nuclear Envelope Reformation;Spry regulation of FGF signaling;Truncations of AMER1 destabilize the destruction complex;CTLA4 inhibitory signaling;AXIN missense mutants destabilize the destruction complex;S45 mutants of beta-catenin aren.SQT.t phosphorylated;Nonsense Mediated Decay (NMD) enhanced by the Exon Junction Complex (EJC);Cyclin D associated events in G1;Platelet sensitization by LDL;Resolution of Sister Chromatid Cohesion;Inhibition of replication initiation of damaged DNA by RB1E2F1;RAFMAP kinase cascade;Downstream signal transduction;VEGFR2 mediated cell proliferation;Signaling by SCF-KIT;Frs2-mediated activation;ARMS-mediated activation;Gastrin-CREB signalling pathway via PKC and MAPK;Interleukin receptor SHC signaling;FCERI mediated MAPK activation;DAP12 signaling;Signaling by PDGF;VEGFA-VEGFR2 Pathway;Signaling by ERBB2;IRS-related events triggered by IGF1R;Negative regulation of FGFR3 signaling;Negative regulation of FGFR4 signaling;Negative regulation of FGFR2 signaling;Negative regulation of FGFR1 signaling 
                                               hsa03015;hsa04071;hsa04114;hsa04151;hsa04152;hsa04261;hsa04350;hsa04390;hsa04530;hsa04728;hsa04730;hsa05142;hsa05160 
                                               PPP2CA 
                                               NA 
                                               slateblue4 
                                               16 
                                               -2.88333333333333 
                                             
                                             
                                               GSK3B-mRNA 
                                               0.188 
                                               0.0806 
                                               0.0299 
                                               0.346 
                                               1.14 
                                               1.02 
                                               1.27 
                                               0.0298 
                                               0.476 
                                               loglinear 
                                               AXIN missense mutants destabilize the destruction complex, DAP12 signaling, Disassembly of the destruction complex and recruitment of AXIN to the membrane, Downstream signal transduction, GAB1 signalosome, Misspliced GSK3beta mutants stabilize beta-catenin, PI3KAKT activation, PIP3 activates AKT signaling, Regulation of HSF1-mediated heat shock response, Role of LAT2NTALLAB on calcium mobilization, S33 mutants of beta-catenin aren.SQT.t phosphorylated, S37 mutants of beta-catenin aren.SQT.t phosphorylated, S45 mutants of beta-catenin aren.SQT.t phosphorylated, Signaling by ERBB2, Signaling by PDGF, Signaling by SCF-KIT, T41 mutants of beta-catenin aren.SQT.t phosphorylated, Truncations of AMER1 destabilize the destruction complex 
                                               SEBELIUS_2_C4636 
                                               GSK3B 
                                               mRNA 
                                               0 
                                               
                                               
                                               Misspliced GSK3beta mutants stabilize beta-catenin;Constitutive Signaling by AKT1 E17K in Cancer;APC truncation mutants have impaired AXIN binding;T41 mutants of beta-catenin aren.SQT.t phosphorylated;AXIN missense mutants destabilize the destruction complex;S45 mutants of beta-catenin aren.SQT.t phosphorylated;Regulation of HSF1-mediated heat shock response;AKT phosphorylates targets in the cytosol;Disassembly of the destruction complex and recruitment of AXIN to the membrane;S37 mutants of beta-catenin aren.SQT.t phosphorylated;S33 mutants of beta-catenin aren.SQT.t phosphorylated;Truncations of AMER1 destabilize the destruction complex;Degradation of GLI2 by the proteasome;PIP3 activates AKT signaling;Downstream signal transduction;GAB1 signalosome;Signaling by SCF-KIT;PI3KAKT activation;Role of LAT2NTALLAB on calcium mobilization;DAP12 signaling;Signaling by PDGF;Signaling by ERBB2 
                                               hsa04012;hsa04062;hsa04110;hsa04151;hsa04310;hsa04360;hsa04390;hsa04510;hsa04550;hsa04660;hsa04662;hsa04722;hsa04728;hsa04910;hsa04916;hsa04917;hsa04919;hsa04931;hsa04932;hsa05010;hsa05160;hsa05162;hsa05164;hsa05166;hsa05169;hsa05200;hsa05210;hsa05213;hsa05215;hsa05217 
                                               GSK3B 
                                               NA 
                                               azure4 
                                               1 
                                               2.33250620347395 
                                             
                                             
                                               PPP2R1A-mRNA 
                                               -0.206 
                                               0.113 
                                               -0.428 
                                               0.0153 
                                               0.867 
                                               0.743 
                                               1.01 
                                               0.0824 
                                               1 
                                               loglinear 
                                               ARMS-mediated activation, AXIN missense mutants destabilize the destruction complex, Beta-catenin phosphorylation cascade, CTLA4 inhibitory signaling, Cyclin AB1 associated events during G2M transition, Cyclin D associated events in G1, DAP12 signaling, DARPP-32 events, Disassembly of the destruction complex and recruitment of AXIN to the membrane, Downstream signal transduction, FCERI mediated MAPK activation, Frs2-mediated activation, Gastrin-CREB signalling pathway via PKC and MAPK, Interleukin receptor SHC signaling, IRS-related events triggered by IGF1R, MASTL Facilitates Mitotic Progression, Negative regulation of FGFR1 signaling, Negative regulation of FGFR2 signaling, Negative regulation of FGFR3 signaling, Negative regulation of FGFR4 signaling, Nonsense Mediated Decay (NMD) enhanced by the Exon Junction Complex (EJC), RAF activation, RAFMAP kinase cascade, Resolution of Sister Chromatid Cohesion, RHO GTPases Activate Formins, S33 mutants of beta-catenin aren.SQT.t phosphorylated, S37 mutants of beta-catenin aren.SQT.t phosphorylated, S45 mutants of beta-catenin aren.SQT.t phosphorylated, Signaling by ERBB2, Signaling by PDGF, Signaling by SCF-KIT, Spry regulation of FGF signaling, Truncations of AMER1 destabilize the destruction complex, VEGFA-VEGFR2 Pathway, VEGFR2 mediated cell proliferation 
                                               SEBELIUS_2_C4636 
                                               PPP2R1A 
                                               mRNA 
                                               0 
                                               
                                               
                                               MASTL Facilitates Mitotic Progression;Regulation of PLK1 Activity at G2M Transition;Disassembly of the destruction complex and recruitment of AXIN to the membrane;S37 mutants of beta-catenin aren.SQT.t phosphorylated;Cyclin AB1 associated events during G2M transition;RAF activation;RHO GTPases Activate Formins;S33 mutants of beta-catenin aren.SQT.t phosphorylated;Loss of Nlp from mitotic centrosomes;DARPP-32 events;Beta-catenin phosphorylation cascade;Initiation of Nuclear Envelope Reformation;Recruitment of mitotic centrosome proteins and complexes;Spry regulation of FGF signaling;Truncations of AMER1 destabilize the destruction complex;CTLA4 inhibitory signaling;AXIN missense mutants destabilize the destruction complex;S45 mutants of beta-catenin aren.SQT.t phosphorylated;Anchoring of the basal body to the plasma membrane;Nonsense Mediated Decay (NMD) enhanced by the Exon Junction Complex (EJC);Cyclin D associated events in G1;Platelet sensitization by LDL;Resolution of Sister Chromatid Cohesion;Inhibition of replication initiation of damaged DNA by RB1E2F1;RAFMAP kinase cascade;Downstream signal transduction;VEGFR2 mediated cell proliferation;Signaling by SCF-KIT;Frs2-mediated activation;ARMS-mediated activation;Gastrin-CREB signalling pathway via PKC and MAPK;Interleukin receptor SHC signaling;FCERI mediated MAPK activation;DAP12 signaling;Signaling by PDGF;VEGFA-VEGFR2 Pathway;Signaling by ERBB2;IRS-related events triggered by IGF1R;Negative regulation of FGFR3 signaling;Negative regulation of FGFR4 signaling;Negative regulation of FGFR2 signaling;Negative regulation of FGFR1 signaling 
                                               hsa03015;hsa04071;hsa04114;hsa04151;hsa04152;hsa04261;hsa04350;hsa04390;hsa04530;hsa04728;hsa04730;hsa05142;hsa05160 
                                               PPP2R1A 
                                               NA 
                                               azure4 
                                               1 
                                               -1.82300884955752 
                                             
                                             
                                               AXIN1-mRNA 
                                               -0.362 
                                               0.246 
                                               -0.843 
                                               0.12 
                                               0.778 
                                               0.558 
                                               1.09 
                                               0.156 
                                               1 
                                               lm.nb 
                                               AXIN missense mutants destabilize the destruction complex, Beta-catenin phosphorylation cascade, Disassembly of the destruction complex and recruitment of AXIN to the membrane, Misspliced GSK3beta mutants stabilize beta-catenin, S33 mutants of beta-catenin aren.SQT.t phosphorylated, S37 mutants of beta-catenin aren.SQT.t phosphorylated, S45 mutants of beta-catenin aren.SQT.t phosphorylated, T41 mutants of beta-catenin aren.SQT.t phosphorylated, Truncations of AMER1 destabilize the destruction complex 
                                               SEBELIUS_2_C4636 
                                               AXIN1 
                                               mRNA 
                                               0 
                                               
                                               
                                               APC truncation mutants have impaired AXIN binding;Misspliced GSK3beta mutants stabilize beta-catenin;T41 mutants of beta-catenin aren.SQT.t phosphorylated;AXIN missense mutants destabilize the destruction complex;S45 mutants of beta-catenin aren.SQT.t phosphorylated;Disassembly of the destruction complex and recruitment of AXIN to the membrane;S37 mutants of beta-catenin aren.SQT.t phosphorylated;S33 mutants of beta-catenin aren.SQT.t phosphorylated;Beta-catenin phosphorylation cascade;Truncations of AMER1 destabilize the destruction complex 
                                               hsa04310;hsa04390;hsa04550;hsa05200;hsa05210;hsa05213;hsa05217 
                                               AXIN1 
                                               NA 
                                               azure4 
                                               1 
                                               -1.47154471544715 
                                             
                                             
                                               APC-mRNA 
                                               -0.0927 
                                               0.109 
                                               -0.306 
                                               0.121 
                                               0.938 
                                               0.809 
                                               1.09 
                                               0.404 
                                               1 
                                               loglinear 
                                               Apoptotic cleavage of cellular proteins, AXIN missense mutants destabilize the destruction complex, Beta-catenin phosphorylation cascade, Deactivation of the beta-catenin transactivating complex, Disassembly of the destruction complex and recruitment of AXIN to the membrane, Misspliced GSK3beta mutants stabilize beta-catenin, S33 mutants of beta-catenin aren.SQT.t phosphorylated, S37 mutants of beta-catenin aren.SQT.t phosphorylated, S45 mutants of beta-catenin aren.SQT.t phosphorylated, T41 mutants of beta-catenin aren.SQT.t phosphorylated, Truncations of AMER1 destabilize the destruction complex 
                                               SEBELIUS_2_C4636 
                                               APC 
                                               mRNA 
                                               0 
                                               
                                               
                                               Apoptotic cleavage of cellular proteins;Misspliced GSK3beta mutants stabilize beta-catenin;APC truncation mutants have impaired AXIN binding;T41 mutants of beta-catenin aren.SQT.t phosphorylated;AXIN missense mutants destabilize the destruction complex;S45 mutants of beta-catenin aren.SQT.t phosphorylated;Deactivation of the beta-catenin transactivating complex;Disassembly of the destruction complex and recruitment of AXIN to the membrane;S37 mutants of beta-catenin aren.SQT.t phosphorylated;S33 mutants of beta-catenin aren.SQT.t phosphorylated;Beta-catenin phosphorylation cascade;Truncations of AMER1 destabilize the destruction complex 
                                               hsa04310;hsa04390;hsa04550;hsa04810;hsa05166;hsa05200;hsa05206;hsa05210;hsa05213;hsa05217 
                                               APC 
                                               NA 
                                               azure4 
                                               1 
                                               -0.85045871559633 
                                             
                                           
                                           DE Results - AXIN missense mutants destabilize the destruction complex genes - New.Annotation: miR18b vs.cel67  Table displaying each sample's global significance scores and directed global significance scores as defined in the heatmaps above. The global significance score is calculated as the square root of the mean squared t-statistic for the genes in a gene set, with t-statistics coming from the linear regression underlying our differential expression analysis. The directed global significance score is calculated as the square root of the mean signed squared t-statistic for the genes in a gene set, with t-statistics coming from the linear regression underlying our differential expression analysis.   
                                      
                                       
                                  
                                
                                
                                  
                                       
                                      
                                           Volcano Plot: New.Annotation: miR890 vs.cel67   More Plot Information                Volcano Plot: New.Annotation: miR890 vs.cel67  Volcano plot displaying each gene's -log10(p-value) and log2 fold change for the selected covariate. Highly statistically significant genes fall at the top of the plot, and highly differentially expressed genes fall to either side. Genes within the selected gene set are highlighted in orange. Horizontal lines indicate various False Discovery Rate (FDR) thresholds.   
                                           DE Results - AXIN missense mutants destabilize the destruction complex genes - New.Annotation: miR890 vs.cel67   More Plot Information      Download CSV Data     
                                           
                                             
                                             
                                           
                                           
                                             
                                               Probe Label 
                                               Log2 fold change 
                                               std error (log2) 
                                               Lower confidence limit (log2) 
                                               Upper confidence limit (log2) 
                                               Linear fold change 
                                               Lower confidence limit (linear) 
                                               Upper confidence limit (linear) 
                                               P-value 
                                               BY.p.value 
                                               method 
                                               Gene.sets 
                                               Codeset.Name 
                                               Probe.Label 
                                               Analyte.Type 
                                               Is.Control 
                                               Control.Type 
                                               Related.Probes 
                                               Probe.Annotation 
                                               KEGG.Pathways 
                                               Official.Gene.Name 
                                               Control.Conc 
                                               volcanocol 
                                               volcanopch 
                                               tstats.all 
                                             
                                           
                                           
                                             
                                               APC-mRNA 
                                               -0.688 
                                               0.0921 
                                               -0.869 
                                               -0.508 
                                               0.621 
                                               0.548 
                                               0.703 
                                               2.41e-07 
                                               2.92e-05 
                                               loglinear 
                                               Apoptotic cleavage of cellular proteins, AXIN missense mutants destabilize the destruction complex, Beta-catenin phosphorylation cascade, Deactivation of the beta-catenin transactivating complex, Disassembly of the destruction complex and recruitment of AXIN to the membrane, Misspliced GSK3beta mutants stabilize beta-catenin, S33 mutants of beta-catenin aren.SQT.t phosphorylated, S37 mutants of beta-catenin aren.SQT.t phosphorylated, S45 mutants of beta-catenin aren.SQT.t phosphorylated, T41 mutants of beta-catenin aren.SQT.t phosphorylated, Truncations of AMER1 destabilize the destruction complex 
                                               SEBELIUS_2_C4636 
                                               APC 
                                               mRNA 
                                               0 
                                               
                                               
                                               Apoptotic cleavage of cellular proteins;Misspliced GSK3beta mutants stabilize beta-catenin;APC truncation mutants have impaired AXIN binding;T41 mutants of beta-catenin aren.SQT.t phosphorylated;AXIN missense mutants destabilize the destruction complex;S45 mutants of beta-catenin aren.SQT.t phosphorylated;Deactivation of the beta-catenin transactivating complex;Disassembly of the destruction complex and recruitment of AXIN to the membrane;S37 mutants of beta-catenin aren.SQT.t phosphorylated;S33 mutants of beta-catenin aren.SQT.t phosphorylated;Beta-catenin phosphorylation cascade;Truncations of AMER1 destabilize the destruction complex 
                                               hsa04310;hsa04390;hsa04550;hsa04810;hsa05166;hsa05200;hsa05206;hsa05210;hsa05213;hsa05217 
                                               APC 
                                               NA 
                                               slateblue1 
                                               16 
                                               -7.47014115092291 
                                             
                                             
                                               PPP2CB-mRNA 
                                               -0.402 
                                               0.0718 
                                               -0.542 
                                               -0.261 
                                               0.757 
                                               0.687 
                                               0.835 
                                               1.51e-05 
                                               0.000909 
                                               loglinear 
                                               ARMS-mediated activation, AXIN missense mutants destabilize the destruction complex, Beta-catenin phosphorylation cascade, CTLA4 inhibitory signaling, Cyclin AB1 associated events during G2M transition, Cyclin D associated events in G1, DAP12 signaling, DARPP-32 events, Disassembly of the destruction complex and recruitment of AXIN to the membrane, Downstream signal transduction, ERKs are inactivated, FCERI mediated MAPK activation, Frs2-mediated activation, Gastrin-CREB signalling pathway via PKC and MAPK, Interleukin receptor SHC signaling, IRS-related events triggered by IGF1R, MASTL Facilitates Mitotic Progression, Misspliced GSK3beta mutants stabilize beta-catenin, Negative regulation of FGFR1 signaling, Negative regulation of FGFR2 signaling, Negative regulation of FGFR3 signaling, Negative regulation of FGFR4 signaling, Negative regulation of MAPK pathway, RAF activation, RAFMAP kinase cascade, Resolution of Sister Chromatid Cohesion, RHO GTPases Activate Formins, S33 mutants of beta-catenin aren.SQT.t phosphorylated, S37 mutants of beta-catenin aren.SQT.t phosphorylated, S45 mutants of beta-catenin aren.SQT.t phosphorylated, Signaling by ERBB2, Signaling by PDGF, Signaling by SCF-KIT, Spry regulation of FGF signaling, T41 mutants of beta-catenin aren.SQT.t phosphorylated, Truncations of AMER1 destabilize the destruction complex, VEGFA-VEGFR2 Pathway, VEGFR2 mediated cell proliferation 
                                               SEBELIUS_2_C4636 
                                               PPP2CB 
                                               mRNA 
                                               0 
                                               
                                               
                                               MASTL Facilitates Mitotic Progression;Misspliced GSK3beta mutants stabilize beta-catenin;T41 mutants of beta-catenin aren.SQT.t phosphorylated;ERKs are inactivated;Glycolysis;Negative regulation of MAPK pathway;Disassembly of the destruction complex and recruitment of AXIN to the membrane;S37 mutants of beta-catenin aren.SQT.t phosphorylated;Cyclin AB1 associated events during G2M transition;RAF activation;RHO GTPases Activate Formins;S33 mutants of beta-catenin aren.SQT.t phosphorylated;DARPP-32 events;Beta-catenin phosphorylation cascade;Spry regulation of FGF signaling;Truncations of AMER1 destabilize the destruction complex;CTLA4 inhibitory signaling;AXIN missense mutants destabilize the destruction complex;S45 mutants of beta-catenin aren.SQT.t phosphorylated;Cyclin D associated events in G1;Platelet sensitization by LDL;Resolution of Sister Chromatid Cohesion;Inhibition of replication initiation of damaged DNA by RB1E2F1;RAFMAP kinase cascade;Downstream signal transduction;VEGFR2 mediated cell proliferation;Signaling by SCF-KIT;Frs2-mediated activation;ARMS-mediated activation;Gastrin-CREB signalling pathway via PKC and MAPK;Interleukin receptor SHC signaling;FCERI mediated MAPK activation;DAP12 signaling;Signaling by PDGF;VEGFA-VEGFR2 Pathway;Signaling by ERBB2;IRS-related events triggered by IGF1R;Negative regulation of FGFR3 signaling;Negative regulation of FGFR4 signaling;Negative regulation of FGFR2 signaling;Negative regulation of FGFR1 signaling 
                                               hsa03015;hsa04071;hsa04114;hsa04151;hsa04152;hsa04261;hsa04350;hsa04390;hsa04530;hsa04728;hsa04730;hsa05142;hsa05160 
                                               PPP2CB 
                                               NA 
                                               slateblue1 
                                               16 
                                               -5.59888579387187 
                                             
                                             
                                               CSNK1A1-mRNA 
                                               0.245 
                                               0.0643 
                                               0.119 
                                               0.371 
                                               1.18 
                                               1.09 
                                               1.29 
                                               0.00104 
                                               0.0276 
                                               loglinear 
                                               AXIN missense mutants destabilize the destruction complex, Beta-catenin phosphorylation cascade, Disassembly of the destruction complex and recruitment of AXIN to the membrane, Misspliced GSK3beta mutants stabilize beta-catenin, S33 mutants of beta-catenin aren.SQT.t phosphorylated, S37 mutants of beta-catenin aren.SQT.t phosphorylated, S45 mutants of beta-catenin aren.SQT.t phosphorylated, T41 mutants of beta-catenin aren.SQT.t phosphorylated 
                                               SEBELIUS_2_C4636 
                                               CSNK1A1 
                                               mRNA 
                                               0 
                                               
                                               
                                               Misspliced GSK3beta mutants stabilize beta-catenin;APC truncation mutants have impaired AXIN binding;T41 mutants of beta-catenin aren.SQT.t phosphorylated;AXIN missense mutants destabilize the destruction complex;S45 mutants of beta-catenin aren.SQT.t phosphorylated;Disassembly of the destruction complex and recruitment of AXIN to the membrane;S37 mutants of beta-catenin aren.SQT.t phosphorylated;S33 mutants of beta-catenin aren.SQT.t phosphorylated;Beta-catenin phosphorylation cascade;Degradation of GLI2 by the proteasome 
                                               hsa04310 
                                               CSNK1A1 
                                               NA 
                                               slateblue2 
                                               16 
                                               3.81026438569207 
                                             
                                             
                                               PPP2R1B-mRNA 
                                               -0.142 
                                               0.0571 
                                               -0.254 
                                               -0.0303 
                                               0.906 
                                               0.839 
                                               0.979 
                                               0.0212 
                                               0.348 
                                               loglinear 
                                               ARMS-mediated activation, AXIN missense mutants destabilize the destruction complex, Beta-catenin phosphorylation cascade, CTLA4 inhibitory signaling, Cyclin AB1 associated events during G2M transition, Cyclin D associated events in G1, DAP12 signaling, DARPP-32 events, Disassembly of the destruction complex and recruitment of AXIN to the membrane, Downstream signal transduction, ERKs are inactivated, FCERI mediated MAPK activation, Frs2-mediated activation, Gastrin-CREB signalling pathway via PKC and MAPK, Interleukin receptor SHC signaling, IRS-related events triggered by IGF1R, MASTL Facilitates Mitotic Progression, Misspliced GSK3beta mutants stabilize beta-catenin, Negative regulation of MAPK pathway, RAF activation, RAFMAP kinase cascade, Resolution of Sister Chromatid Cohesion, RHO GTPases Activate Formins, S33 mutants of beta-catenin aren.SQT.t phosphorylated, S37 mutants of beta-catenin aren.SQT.t phosphorylated, S45 mutants of beta-catenin aren.SQT.t phosphorylated, Signaling by ERBB2, Signaling by PDGF, Signaling by SCF-KIT, T41 mutants of beta-catenin aren.SQT.t phosphorylated, Truncations of AMER1 destabilize the destruction complex, VEGFA-VEGFR2 Pathway, VEGFR2 mediated cell proliferation 
                                               SEBELIUS_2_C4636 
                                               PPP2R1B 
                                               mRNA 
                                               0 
                                               
                                               
                                               MASTL Facilitates Mitotic Progression;Misspliced GSK3beta mutants stabilize beta-catenin;T41 mutants of beta-catenin aren.SQT.t phosphorylated;ERKs are inactivated;Glycolysis;Negative regulation of MAPK pathway;Disassembly of the destruction complex and recruitment of AXIN to the membrane;S37 mutants of beta-catenin aren.SQT.t phosphorylated;Cyclin AB1 associated events during G2M transition;RAF activation;RHO GTPases Activate Formins;S33 mutants of beta-catenin aren.SQT.t phosphorylated;DARPP-32 events;Beta-catenin phosphorylation cascade;Truncations of AMER1 destabilize the destruction complex;CTLA4 inhibitory signaling;AXIN missense mutants destabilize the destruction complex;S45 mutants of beta-catenin aren.SQT.t phosphorylated;Cyclin D associated events in G1;Platelet sensitization by LDL;Resolution of Sister Chromatid Cohesion;Inhibition of replication initiation of damaged DNA by RB1E2F1;RAFMAP kinase cascade;Downstream signal transduction;VEGFR2 mediated cell proliferation;Signaling by SCF-KIT;Frs2-mediated activation;ARMS-mediated activation;Gastrin-CREB signalling pathway via PKC and MAPK;Interleukin receptor SHC signaling;FCERI mediated MAPK activation;DAP12 signaling;Signaling by PDGF;VEGFA-VEGFR2 Pathway;Signaling by ERBB2;IRS-related events triggered by IGF1R 
                                               hsa03015;hsa04071;hsa04114;hsa04151;hsa04152;hsa04261;hsa04350;hsa04390;hsa04530;hsa04728;hsa04730;hsa05142;hsa05160 
                                               PPP2R1B 
                                               NA 
                                               slateblue4 
                                               16 
                                               -2.48686514886165 
                                             
                                             
                                               AXIN1-mRNA 
                                               -0.356 
                                               0.206 
                                               -0.759 
                                               0.0469 
                                               0.781 
                                               0.591 
                                               1.03 
                                               0.098 
                                               1 
                                               lm.nb 
                                               AXIN missense mutants destabilize the destruction complex, Beta-catenin phosphorylation cascade, Disassembly of the destruction complex and recruitment of AXIN to the membrane, Misspliced GSK3beta mutants stabilize beta-catenin, S33 mutants of beta-catenin aren.SQT.t phosphorylated, S37 mutants of beta-catenin aren.SQT.t phosphorylated, S45 mutants of beta-catenin aren.SQT.t phosphorylated, T41 mutants of beta-catenin aren.SQT.t phosphorylated, Truncations of AMER1 destabilize the destruction complex 
                                               SEBELIUS_2_C4636 
                                               AXIN1 
                                               mRNA 
                                               0 
                                               
                                               
                                               APC truncation mutants have impaired AXIN binding;Misspliced GSK3beta mutants stabilize beta-catenin;T41 mutants of beta-catenin aren.SQT.t phosphorylated;AXIN missense mutants destabilize the destruction complex;S45 mutants of beta-catenin aren.SQT.t phosphorylated;Disassembly of the destruction complex and recruitment of AXIN to the membrane;S37 mutants of beta-catenin aren.SQT.t phosphorylated;S33 mutants of beta-catenin aren.SQT.t phosphorylated;Beta-catenin phosphorylation cascade;Truncations of AMER1 destabilize the destruction complex 
                                               hsa04310;hsa04390;hsa04550;hsa05200;hsa05210;hsa05213;hsa05217 
                                               AXIN1 
                                               NA 
                                               azure4 
                                               1 
                                               -1.72815533980583 
                                             
                                             
                                               GSK3B-mRNA 
                                               0.106 
                                               0.0681 
                                               -0.027 
                                               0.24 
                                               1.08 
                                               0.981 
                                               1.18 
                                               0.133 
                                               1 
                                               loglinear 
                                               AXIN missense mutants destabilize the destruction complex, DAP12 signaling, Disassembly of the destruction complex and recruitment of AXIN to the membrane, Downstream signal transduction, GAB1 signalosome, Misspliced GSK3beta mutants stabilize beta-catenin, PI3KAKT activation, PIP3 activates AKT signaling, Regulation of HSF1-mediated heat shock response, Role of LAT2NTALLAB on calcium mobilization, S33 mutants of beta-catenin aren.SQT.t phosphorylated, S37 mutants of beta-catenin aren.SQT.t phosphorylated, S45 mutants of beta-catenin aren.SQT.t phosphorylated, Signaling by ERBB2, Signaling by PDGF, Signaling by SCF-KIT, T41 mutants of beta-catenin aren.SQT.t phosphorylated, Truncations of AMER1 destabilize the destruction complex 
                                               SEBELIUS_2_C4636 
                                               GSK3B 
                                               mRNA 
                                               0 
                                               
                                               
                                               Misspliced GSK3beta mutants stabilize beta-catenin;Constitutive Signaling by AKT1 E17K in Cancer;APC truncation mutants have impaired AXIN binding;T41 mutants of beta-catenin aren.SQT.t phosphorylated;AXIN missense mutants destabilize the destruction complex;S45 mutants of beta-catenin aren.SQT.t phosphorylated;Regulation of HSF1-mediated heat shock response;AKT phosphorylates targets in the cytosol;Disassembly of the destruction complex and recruitment of AXIN to the membrane;S37 mutants of beta-catenin aren.SQT.t phosphorylated;S33 mutants of beta-catenin aren.SQT.t phosphorylated;Truncations of AMER1 destabilize the destruction complex;Degradation of GLI2 by the proteasome;PIP3 activates AKT signaling;Downstream signal transduction;GAB1 signalosome;Signaling by SCF-KIT;PI3KAKT activation;Role of LAT2NTALLAB on calcium mobilization;DAP12 signaling;Signaling by PDGF;Signaling by ERBB2 
                                               hsa04012;hsa04062;hsa04110;hsa04151;hsa04310;hsa04360;hsa04390;hsa04510;hsa04550;hsa04660;hsa04662;hsa04722;hsa04728;hsa04910;hsa04916;hsa04917;hsa04919;hsa04931;hsa04932;hsa05010;hsa05160;hsa05162;hsa05164;hsa05166;hsa05169;hsa05200;hsa05210;hsa05213;hsa05215;hsa05217 
                                               GSK3B 
                                               NA 
                                               azure4 
                                               1 
                                               1.55653450807636 
                                             
                                             
                                               PPP2CA-mRNA 
                                               -0.0403 
                                               0.0507 
                                               -0.14 
                                               0.0591 
                                               0.972 
                                               0.908 
                                               1.04 
                                               0.436 
                                               1 
                                               loglinear 
                                               ARMS-mediated activation, AXIN missense mutants destabilize the destruction complex, Beta-catenin phosphorylation cascade, CTLA4 inhibitory signaling, Cyclin AB1 associated events during G2M transition, Cyclin D associated events in G1, DAP12 signaling, DARPP-32 events, Disassembly of the destruction complex and recruitment of AXIN to the membrane, Downstream signal transduction, ERKs are inactivated, FCERI mediated MAPK activation, Frs2-mediated activation, Gastrin-CREB signalling pathway via PKC and MAPK, Interleukin receptor SHC signaling, IRS-related events triggered by IGF1R, MASTL Facilitates Mitotic Progression, Negative regulation of FGFR1 signaling, Negative regulation of FGFR2 signaling, Negative regulation of FGFR3 signaling, Negative regulation of FGFR4 signaling, Negative regulation of MAPK pathway, Nonsense Mediated Decay (NMD) enhanced by the Exon Junction Complex (EJC), RAF activation, RAFMAP kinase cascade, Resolution of Sister Chromatid Cohesion, RHO GTPases Activate Formins, S33 mutants of beta-catenin aren.SQT.t phosphorylated, S37 mutants of beta-catenin aren.SQT.t phosphorylated, S45 mutants of beta-catenin aren.SQT.t phosphorylated, Signaling by ERBB2, Signaling by PDGF, Signaling by SCF-KIT, Spry regulation of FGF signaling, T41 mutants of beta-catenin aren.SQT.t phosphorylated, Truncations of AMER1 destabilize the destruction complex, VEGFA-VEGFR2 Pathway, VEGFR2 mediated cell proliferation 
                                               SEBELIUS_2_C4636 
                                               PPP2CA 
                                               mRNA 
                                               0 
                                               
                                               
                                               MASTL Facilitates Mitotic Progression;T41 mutants of beta-catenin aren.SQT.t phosphorylated;ERKs are inactivated;Glycolysis;Negative regulation of MAPK pathway;Disassembly of the destruction complex and recruitment of AXIN to the membrane;S37 mutants of beta-catenin aren.SQT.t phosphorylated;Cyclin AB1 associated events during G2M transition;RAF activation;RHO GTPases Activate Formins;S33 mutants of beta-catenin aren.SQT.t phosphorylated;DARPP-32 events;Beta-catenin phosphorylation cascade;Initiation of Nuclear Envelope Reformation;Spry regulation of FGF signaling;Truncations of AMER1 destabilize the destruction complex;CTLA4 inhibitory signaling;AXIN missense mutants destabilize the destruction complex;S45 mutants of beta-catenin aren.SQT.t phosphorylated;Nonsense Mediated Decay (NMD) enhanced by the Exon Junction Complex (EJC);Cyclin D associated events in G1;Platelet sensitization by LDL;Resolution of Sister Chromatid Cohesion;Inhibition of replication initiation of damaged DNA by RB1E2F1;RAFMAP kinase cascade;Downstream signal transduction;VEGFR2 mediated cell proliferation;Signaling by SCF-KIT;Frs2-mediated activation;ARMS-mediated activation;Gastrin-CREB signalling pathway via PKC and MAPK;Interleukin receptor SHC signaling;FCERI mediated MAPK activation;DAP12 signaling;Signaling by PDGF;VEGFA-VEGFR2 Pathway;Signaling by ERBB2;IRS-related events triggered by IGF1R;Negative regulation of FGFR3 signaling;Negative regulation of FGFR4 signaling;Negative regulation of FGFR2 signaling;Negative regulation of FGFR1 signaling 
                                               hsa03015;hsa04071;hsa04114;hsa04151;hsa04152;hsa04261;hsa04350;hsa04390;hsa04530;hsa04728;hsa04730;hsa05142;hsa05160 
                                               PPP2CA 
                                               NA 
                                               azure4 
                                               1 
                                               -0.794871794871795 
                                             
                                             
                                               PPP2R1A-mRNA 
                                               0.069 
                                               0.0955 
                                               -0.118 
                                               0.256 
                                               1.05 
                                               0.921 
                                               1.19 
                                               0.478 
                                               1 
                                               loglinear 
                                               ARMS-mediated activation, AXIN missense mutants destabilize the destruction complex, Beta-catenin phosphorylation cascade, CTLA4 inhibitory signaling, Cyclin AB1 associated events during G2M transition, Cyclin D associated events in G1, DAP12 signaling, DARPP-32 events, Disassembly of the destruction complex and recruitment of AXIN to the membrane, Downstream signal transduction, FCERI mediated MAPK activation, Frs2-mediated activation, Gastrin-CREB signalling pathway via PKC and MAPK, Interleukin receptor SHC signaling, IRS-related events triggered by IGF1R, MASTL Facilitates Mitotic Progression, Negative regulation of FGFR1 signaling, Negative regulation of FGFR2 signaling, Negative regulation of FGFR3 signaling, Negative regulation of FGFR4 signaling, Nonsense Mediated Decay (NMD) enhanced by the Exon Junction Complex (EJC), RAF activation, RAFMAP kinase cascade, Resolution of Sister Chromatid Cohesion, RHO GTPases Activate Formins, S33 mutants of beta-catenin aren.SQT.t phosphorylated, S37 mutants of beta-catenin aren.SQT.t phosphorylated, S45 mutants of beta-catenin aren.SQT.t phosphorylated, Signaling by ERBB2, Signaling by PDGF, Signaling by SCF-KIT, Spry regulation of FGF signaling, Truncations of AMER1 destabilize the destruction complex, VEGFA-VEGFR2 Pathway, VEGFR2 mediated cell proliferation 
                                               SEBELIUS_2_C4636 
                                               PPP2R1A 
                                               mRNA 
                                               0 
                                               
                                               
                                               MASTL Facilitates Mitotic Progression;Regulation of PLK1 Activity at G2M Transition;Disassembly of the destruction complex and recruitment of AXIN to the membrane;S37 mutants of beta-catenin aren.SQT.t phosphorylated;Cyclin AB1 associated events during G2M transition;RAF activation;RHO GTPases Activate Formins;S33 mutants of beta-catenin aren.SQT.t phosphorylated;Loss of Nlp from mitotic centrosomes;DARPP-32 events;Beta-catenin phosphorylation cascade;Initiation of Nuclear Envelope Reformation;Recruitment of mitotic centrosome proteins and complexes;Spry regulation of FGF signaling;Truncations of AMER1 destabilize the destruction complex;CTLA4 inhibitory signaling;AXIN missense mutants destabilize the destruction complex;S45 mutants of beta-catenin aren.SQT.t phosphorylated;Anchoring of the basal body to the plasma membrane;Nonsense Mediated Decay (NMD) enhanced by the Exon Junction Complex (EJC);Cyclin D associated events in G1;Platelet sensitization by LDL;Resolution of Sister Chromatid Cohesion;Inhibition of replication initiation of damaged DNA by RB1E2F1;RAFMAP kinase cascade;Downstream signal transduction;VEGFR2 mediated cell proliferation;Signaling by SCF-KIT;Frs2-mediated activation;ARMS-mediated activation;Gastrin-CREB signalling pathway via PKC and MAPK;Interleukin receptor SHC signaling;FCERI mediated MAPK activation;DAP12 signaling;Signaling by PDGF;VEGFA-VEGFR2 Pathway;Signaling by ERBB2;IRS-related events triggered by IGF1R;Negative regulation of FGFR3 signaling;Negative regulation of FGFR4 signaling;Negative regulation of FGFR2 signaling;Negative regulation of FGFR1 signaling 
                                               hsa03015;hsa04071;hsa04114;hsa04151;hsa04152;hsa04261;hsa04350;hsa04390;hsa04530;hsa04728;hsa04730;hsa05142;hsa05160 
                                               PPP2R1A 
                                               NA 
                                               azure4 
                                               1 
                                               0.722513089005236 
                                             
                                           
                                           DE Results - AXIN missense mutants destabilize the destruction complex genes - New.Annotation: miR890 vs.cel67  Table displaying each sample's global significance scores and directed global significance scores as defined in the heatmaps above. The global significance score is calculated as the square root of the mean squared t-statistic for the genes in a gene set, with t-statistics coming from the linear regression underlying our differential expression analysis. The directed global significance score is calculated as the square root of the mean signed squared t-statistic for the genes in a gene set, with t-statistics coming from the linear regression underlying our differential expression analysis.   
                                      
                                       
                                  
                                
                              
                             
                          
                           
                      
                    
                    
                      
                           
                          
                              New.Annotation: 
                              
                                  New.Annotation: differential expression in Control vs. baseline of cel67  
                                  New.Annotation: differential expression in miR142 vs. baseline of cel67  
                                  New.Annotation: differential expression in miR155 vs. baseline of cel67  
                                  New.Annotation: differential expression in miR18b vs. baseline of cel67  
                                  New.Annotation: differential expression in miR890 vs. baseline of cel67  
                              
                              
                              
                                
                                  
                                       
                                      
                                           Volcano Plot: New.Annotation: Control vs.cel67   More Plot Information                Volcano Plot: New.Annotation: Control vs.cel67  Volcano plot displaying each gene's -log10(p-value) and log2 fold change for the selected covariate. Highly statistically significant genes fall at the top of the plot, and highly differentially expressed genes fall to either side. Genes within the selected gene set are highlighted in orange. Horizontal lines indicate various False Discovery Rate (FDR) thresholds.   
                                           DE Results - Beta-catenin phosphorylation cascade genes - New.Annotation: Control vs.cel67   More Plot Information      Download CSV Data     
                                           
                                             
                                             
                                           
                                           
                                             
                                               Probe Label 
                                               Log2 fold change 
                                               std error (log2) 
                                               Lower confidence limit (log2) 
                                               Upper confidence limit (log2) 
                                               Linear fold change 
                                               Lower confidence limit (linear) 
                                               Upper confidence limit (linear) 
                                               P-value 
                                               BY.p.value 
                                               method 
                                               Gene.sets 
                                               Codeset.Name 
                                               Probe.Label 
                                               Analyte.Type 
                                               Is.Control 
                                               Control.Type 
                                               Related.Probes 
                                               Probe.Annotation 
                                               KEGG.Pathways 
                                               Official.Gene.Name 
                                               Control.Conc 
                                               volcanocol 
                                               volcanopch 
                                               tstats.all 
                                             
                                           
                                           
                                             
                                               CSNK1A1-mRNA 
                                               0.608 
                                               0.0669 
                                               0.477 
                                               0.739 
                                               1.52 
                                               1.39 
                                               1.67 
                                               9.92e-09 
                                               1.32e-05 
                                               loglinear 
                                               AXIN missense mutants destabilize the destruction complex, Beta-catenin phosphorylation cascade, Disassembly of the destruction complex and recruitment of AXIN to the membrane, Misspliced GSK3beta mutants stabilize beta-catenin, S33 mutants of beta-catenin aren.SQT.t phosphorylated, S37 mutants of beta-catenin aren.SQT.t phosphorylated, S45 mutants of beta-catenin aren.SQT.t phosphorylated, T41 mutants of beta-catenin aren.SQT.t phosphorylated 
                                               SEBELIUS_2_C4636 
                                               CSNK1A1 
                                               mRNA 
                                               0 
                                               
                                               
                                               Misspliced GSK3beta mutants stabilize beta-catenin;APC truncation mutants have impaired AXIN binding;T41 mutants of beta-catenin aren.SQT.t phosphorylated;AXIN missense mutants destabilize the destruction complex;S45 mutants of beta-catenin aren.SQT.t phosphorylated;Disassembly of the destruction complex and recruitment of AXIN to the membrane;S37 mutants of beta-catenin aren.SQT.t phosphorylated;S33 mutants of beta-catenin aren.SQT.t phosphorylated;Beta-catenin phosphorylation cascade;Degradation of GLI2 by the proteasome 
                                               hsa04310 
                                               CSNK1A1 
                                               NA 
                                               slateblue1 
                                               16 
                                               9.0881913303438 
                                             
                                             
                                               CTNNB1-mRNA 
                                               0.141 
                                               0.056 
                                               0.0317 
                                               0.251 
                                               1.1 
                                               1.02 
                                               1.19 
                                               0.0196 
                                               0.743 
                                               loglinear 
                                               Apoptotic cleavage of cellular proteins, Beta-catenin phosphorylation cascade, Deactivation of the beta-catenin transactivating complex, Disassembly of the destruction complex and recruitment of AXIN to the membrane, Formation of the beta-catenin.CL.TCF transactivating complex, Misspliced GSK3beta mutants stabilize beta-catenin, Repression of WNT target genes, RHO GTPases activate IQGAPs, S37 mutants of beta-catenin aren.SQT.t phosphorylated, S45 mutants of beta-catenin aren.SQT.t phosphorylated, T41 mutants of beta-catenin aren.SQT.t phosphorylated, VEGFA-VEGFR2 Pathway, VEGFR2 mediated vascular permeability 
                                               SEBELIUS_2_C4636 
                                               CTNNB1 
                                               mRNA 
                                               0 
                                               
                                               
                                               Disassembly of the destruction complex and recruitment of AXIN to the membrane;S37 mutants of beta-catenin aren.SQT.t phosphorylated;T41 mutants of beta-catenin aren.SQT.t phosphorylated;S45 mutants of beta-catenin aren.SQT.t phosphorylated;Binding of TCFLEF.CL.CTNNB1 to target gene promoters;RHO GTPases activate IQGAPs;Deactivation of the beta-catenin transactivating complex;Repression of WNT target genes;Apoptotic cleavage of cell adhesion  proteins;Adherens junctions interactions;Misspliced GSK3beta mutants stabilize beta-catenin;VEGFR2 mediated vascular permeability;Beta-catenin phosphorylation cascade;Synthesis# secretion# and inactivation of Glucagon-like Peptide-1 (GLP-1);Formation of the beta-catenin.CL.TCF transactivating complex;Apoptotic cleavage of cellular proteins;VEGFA-VEGFR2 Pathway 
                                               hsa04015;hsa04310;hsa04390;hsa04510;hsa04520;hsa04530;hsa04550;hsa04670;hsa04916;hsa04919;hsa05100;hsa05130;hsa05166;hsa05200;hsa05205;hsa05210;hsa05213;hsa05215;hsa05216;hsa05217;hsa05412 
                                               CTNNB1 
                                               NA 
                                               azure4 
                                               1 
                                               2.51785714285714 
                                             
                                             
                                               PPP2CB-mRNA 
                                               -0.182 
                                               0.0747 
                                               -0.328 
                                               -0.0354 
                                               0.882 
                                               0.797 
                                               0.976 
                                               0.0239 
                                               0.756 
                                               loglinear 
                                               ARMS-mediated activation, AXIN missense mutants destabilize the destruction complex, Beta-catenin phosphorylation cascade, CTLA4 inhibitory signaling, Cyclin AB1 associated events during G2M transition, Cyclin D associated events in G1, DAP12 signaling, DARPP-32 events, Disassembly of the destruction complex and recruitment of AXIN to the membrane, Downstream signal transduction, ERKs are inactivated, FCERI mediated MAPK activation, Frs2-mediated activation, Gastrin-CREB signalling pathway via PKC and MAPK, Interleukin receptor SHC signaling, IRS-related events triggered by IGF1R, MASTL Facilitates Mitotic Progression, Misspliced GSK3beta mutants stabilize beta-catenin, Negative regulation of FGFR1 signaling, Negative regulation of FGFR2 signaling, Negative regulation of FGFR3 signaling, Negative regulation of FGFR4 signaling, Negative regulation of MAPK pathway, RAF activation, RAFMAP kinase cascade, Resolution of Sister Chromatid Cohesion, RHO GTPases Activate Formins, S33 mutants of beta-catenin aren.SQT.t phosphorylated, S37 mutants of beta-catenin aren.SQT.t phosphorylated, S45 mutants of beta-catenin aren.SQT.t phosphorylated, Signaling by ERBB2, Signaling by PDGF, Signaling by SCF-KIT, Spry regulation of FGF signaling, T41 mutants of beta-catenin aren.SQT.t phosphorylated, Truncations of AMER1 destabilize the destruction complex, VEGFA-VEGFR2 Pathway, VEGFR2 mediated cell proliferation 
                                               SEBELIUS_2_C4636 
                                               PPP2CB 
                                               mRNA 
                                               0 
                                               
                                               
                                               MASTL Facilitates Mitotic Progression;Misspliced GSK3beta mutants stabilize beta-catenin;T41 mutants of beta-catenin aren.SQT.t phosphorylated;ERKs are inactivated;Glycolysis;Negative regulation of MAPK pathway;Disassembly of the destruction complex and recruitment of AXIN to the membrane;S37 mutants of beta-catenin aren.SQT.t phosphorylated;Cyclin AB1 associated events during G2M transition;RAF activation;RHO GTPases Activate Formins;S33 mutants of beta-catenin aren.SQT.t phosphorylated;DARPP-32 events;Beta-catenin phosphorylation cascade;Spry regulation of FGF signaling;Truncations of AMER1 destabilize the destruction complex;CTLA4 inhibitory signaling;AXIN missense mutants destabilize the destruction complex;S45 mutants of beta-catenin aren.SQT.t phosphorylated;Cyclin D associated events in G1;Platelet sensitization by LDL;Resolution of Sister Chromatid Cohesion;Inhibition of replication initiation of damaged DNA by RB1E2F1;RAFMAP kinase cascade;Downstream signal transduction;VEGFR2 mediated cell proliferation;Signaling by SCF-KIT;Frs2-mediated activation;ARMS-mediated activation;Gastrin-CREB signalling pathway via PKC and MAPK;Interleukin receptor SHC signaling;FCERI mediated MAPK activation;DAP12 signaling;Signaling by PDGF;VEGFA-VEGFR2 Pathway;Signaling by ERBB2;IRS-related events triggered by IGF1R;Negative regulation of FGFR3 signaling;Negative regulation of FGFR4 signaling;Negative regulation of FGFR2 signaling;Negative regulation of FGFR1 signaling 
                                               hsa03015;hsa04071;hsa04114;hsa04151;hsa04152;hsa04261;hsa04350;hsa04390;hsa04530;hsa04728;hsa04730;hsa05142;hsa05160 
                                               PPP2CB 
                                               NA 
                                               azure4 
                                               1 
                                               -2.43641231593039 
                                             
                                             
                                               APC-mRNA 
                                               -0.21 
                                               0.0957 
                                               -0.397 
                                               -0.0221 
                                               0.865 
                                               0.759 
                                               0.985 
                                               0.0399 
                                               0.99 
                                               loglinear 
                                               Apoptotic cleavage of cellular proteins, AXIN missense mutants destabilize the destruction complex, Beta-catenin phosphorylation cascade, Deactivation of the beta-catenin transactivating complex, Disassembly of the destruction complex and recruitment of AXIN to the membrane, Misspliced GSK3beta mutants stabilize beta-catenin, S33 mutants of beta-catenin aren.SQT.t phosphorylated, S37 mutants of beta-catenin aren.SQT.t phosphorylated, S45 mutants of beta-catenin aren.SQT.t phosphorylated, T41 mutants of beta-catenin aren.SQT.t phosphorylated, Truncations of AMER1 destabilize the destruction complex 
                                               SEBELIUS_2_C4636 
                                               APC 
                                               mRNA 
                                               0 
                                               
                                               
                                               Apoptotic cleavage of cellular proteins;Misspliced GSK3beta mutants stabilize beta-catenin;APC truncation mutants have impaired AXIN binding;T41 mutants of beta-catenin aren.SQT.t phosphorylated;AXIN missense mutants destabilize the destruction complex;S45 mutants of beta-catenin aren.SQT.t phosphorylated;Deactivation of the beta-catenin transactivating complex;Disassembly of the destruction complex and recruitment of AXIN to the membrane;S37 mutants of beta-catenin aren.SQT.t phosphorylated;S33 mutants of beta-catenin aren.SQT.t phosphorylated;Beta-catenin phosphorylation cascade;Truncations of AMER1 destabilize the destruction complex 
                                               hsa04310;hsa04390;hsa04550;hsa04810;hsa05166;hsa05200;hsa05206;hsa05210;hsa05213;hsa05217 
                                               APC 
                                               NA 
                                               azure4 
                                               1 
                                               -2.19435736677116 
                                             
                                             
                                               PPP2CA-mRNA 
                                               -0.115 
                                               0.0527 
                                               -0.219 
                                               -0.0119 
                                               0.923 
                                               0.859 
                                               0.992 
                                               0.0403 
                                               0.99 
                                               loglinear 
                                               ARMS-mediated activation, AXIN missense mutants destabilize the destruction complex, Beta-catenin phosphorylation cascade, CTLA4 inhibitory signaling, Cyclin AB1 associated events during G2M transition, Cyclin D associated events in G1, DAP12 signaling, DARPP-32 events, Disassembly of the destruction complex and recruitment of AXIN to the membrane, Downstream signal transduction, ERKs are inactivated, FCERI mediated MAPK activation, Frs2-mediated activation, Gastrin-CREB signalling pathway via PKC and MAPK, Interleukin receptor SHC signaling, IRS-related events triggered by IGF1R, MASTL Facilitates Mitotic Progression, Negative regulation of FGFR1 signaling, Negative regulation of FGFR2 signaling, Negative regulation of FGFR3 signaling, Negative regulation of FGFR4 signaling, Negative regulation of MAPK pathway, Nonsense Mediated Decay (NMD) enhanced by the Exon Junction Complex (EJC), RAF activation, RAFMAP kinase cascade, Resolution of Sister Chromatid Cohesion, RHO GTPases Activate Formins, S33 mutants of beta-catenin aren.SQT.t phosphorylated, S37 mutants of beta-catenin aren.SQT.t phosphorylated, S45 mutants of beta-catenin aren.SQT.t phosphorylated, Signaling by ERBB2, Signaling by PDGF, Signaling by SCF-KIT, Spry regulation of FGF signaling, T41 mutants of beta-catenin aren.SQT.t phosphorylated, Truncations of AMER1 destabilize the destruction complex, VEGFA-VEGFR2 Pathway, VEGFR2 mediated cell proliferation 
                                               SEBELIUS_2_C4636 
                                               PPP2CA 
                                               mRNA 
                                               0 
                                               
                                               
                                               MASTL Facilitates Mitotic Progression;T41 mutants of beta-catenin aren.SQT.t phosphorylated;ERKs are inactivated;Glycolysis;Negative regulation of MAPK pathway;Disassembly of the destruction complex and recruitment of AXIN to the membrane;S37 mutants of beta-catenin aren.SQT.t phosphorylated;Cyclin AB1 associated events during G2M transition;RAF activation;RHO GTPases Activate Formins;S33 mutants of beta-catenin aren.SQT.t phosphorylated;DARPP-32 events;Beta-catenin phosphorylation cascade;Initiation of Nuclear Envelope Reformation;Spry regulation of FGF signaling;Truncations of AMER1 destabilize the destruction complex;CTLA4 inhibitory signaling;AXIN missense mutants destabilize the destruction complex;S45 mutants of beta-catenin aren.SQT.t phosphorylated;Nonsense Mediated Decay (NMD) enhanced by the Exon Junction Complex (EJC);Cyclin D associated events in G1;Platelet sensitization by LDL;Resolution of Sister Chromatid Cohesion;Inhibition of replication initiation of damaged DNA by RB1E2F1;RAFMAP kinase cascade;Downstream signal transduction;VEGFR2 mediated cell proliferation;Signaling by SCF-KIT;Frs2-mediated activation;ARMS-mediated activation;Gastrin-CREB signalling pathway via PKC and MAPK;Interleukin receptor SHC signaling;FCERI mediated MAPK activation;DAP12 signaling;Signaling by PDGF;VEGFA-VEGFR2 Pathway;Signaling by ERBB2;IRS-related events triggered by IGF1R;Negative regulation of FGFR3 signaling;Negative regulation of FGFR4 signaling;Negative regulation of FGFR2 signaling;Negative regulation of FGFR1 signaling 
                                               hsa03015;hsa04071;hsa04114;hsa04151;hsa04152;hsa04261;hsa04350;hsa04390;hsa04530;hsa04728;hsa04730;hsa05142;hsa05160 
                                               PPP2CA 
                                               NA 
                                               azure4 
                                               1 
                                               -2.18216318785579 
                                             
                                             
                                               AXIN1-mRNA 
                                               -0.348 
                                               0.214 
                                               -0.768 
                                               0.0712 
                                               0.786 
                                               0.587 
                                               1.05 
                                               0.119 
                                               1 
                                               lm.nb 
                                               AXIN missense mutants destabilize the destruction complex, Beta-catenin phosphorylation cascade, Disassembly of the destruction complex and recruitment of AXIN to the membrane, Misspliced GSK3beta mutants stabilize beta-catenin, S33 mutants of beta-catenin aren.SQT.t phosphorylated, S37 mutants of beta-catenin aren.SQT.t phosphorylated, S45 mutants of beta-catenin aren.SQT.t phosphorylated, T41 mutants of beta-catenin aren.SQT.t phosphorylated, Truncations of AMER1 destabilize the destruction complex 
                                               SEBELIUS_2_C4636 
                                               AXIN1 
                                               mRNA 
                                               0 
                                               
                                               
                                               APC truncation mutants have impaired AXIN binding;Misspliced GSK3beta mutants stabilize beta-catenin;T41 mutants of beta-catenin aren.SQT.t phosphorylated;AXIN missense mutants destabilize the destruction complex;S45 mutants of beta-catenin aren.SQT.t phosphorylated;Disassembly of the destruction complex and recruitment of AXIN to the membrane;S37 mutants of beta-catenin aren.SQT.t phosphorylated;S33 mutants of beta-catenin aren.SQT.t phosphorylated;Beta-catenin phosphorylation cascade;Truncations of AMER1 destabilize the destruction complex 
                                               hsa04310;hsa04390;hsa04550;hsa05200;hsa05210;hsa05213;hsa05217 
                                               AXIN1 
                                               NA 
                                               azure4 
                                               1 
                                               -1.62616822429907 
                                             
                                             
                                               PPP2R1B-mRNA 
                                               -0.0747 
                                               0.0593 
                                               -0.191 
                                               0.0416 
                                               0.95 
                                               0.876 
                                               1.03 
                                               0.222 
                                               1 
                                               loglinear 
                                               ARMS-mediated activation, AXIN missense mutants destabilize the destruction complex, Beta-catenin phosphorylation cascade, CTLA4 inhibitory signaling, Cyclin AB1 associated events during G2M transition, Cyclin D associated events in G1, DAP12 signaling, DARPP-32 events, Disassembly of the destruction complex and recruitment of AXIN to the membrane, Downstream signal transduction, ERKs are inactivated, FCERI mediated MAPK activation, Frs2-mediated activation, Gastrin-CREB signalling pathway via PKC and MAPK, Interleukin receptor SHC signaling, IRS-related events triggered by IGF1R, MASTL Facilitates Mitotic Progression, Misspliced GSK3beta mutants stabilize beta-catenin, Negative regulation of MAPK pathway, RAF activation, RAFMAP kinase cascade, Resolution of Sister Chromatid Cohesion, RHO GTPases Activate Formins, S33 mutants of beta-catenin aren.SQT.t phosphorylated, S37 mutants of beta-catenin aren.SQT.t phosphorylated, S45 mutants of beta-catenin aren.SQT.t phosphorylated, Signaling by ERBB2, Signaling by PDGF, Signaling by SCF-KIT, T41 mutants of beta-catenin aren.SQT.t phosphorylated, Truncations of AMER1 destabilize the destruction complex, VEGFA-VEGFR2 Pathway, VEGFR2 mediated cell proliferation 
                                               SEBELIUS_2_C4636 
                                               PPP2R1B 
                                               mRNA 
                                               0 
                                               
                                               
                                               MASTL Facilitates Mitotic Progression;Misspliced GSK3beta mutants stabilize beta-catenin;T41 mutants of beta-catenin aren.SQT.t phosphorylated;ERKs are inactivated;Glycolysis;Negative regulation of MAPK pathway;Disassembly of the destruction complex and recruitment of AXIN to the membrane;S37 mutants of beta-catenin aren.SQT.t phosphorylated;Cyclin AB1 associated events during G2M transition;RAF activation;RHO GTPases Activate Formins;S33 mutants of beta-catenin aren.SQT.t phosphorylated;DARPP-32 events;Beta-catenin phosphorylation cascade;Truncations of AMER1 destabilize the destruction complex;CTLA4 inhibitory signaling;AXIN missense mutants destabilize the destruction complex;S45 mutants of beta-catenin aren.SQT.t phosphorylated;Cyclin D associated events in G1;Platelet sensitization by LDL;Resolution of Sister Chromatid Cohesion;Inhibition of replication initiation of damaged DNA by RB1E2F1;RAFMAP kinase cascade;Downstream signal transduction;VEGFR2 mediated cell proliferation;Signaling by SCF-KIT;Frs2-mediated activation;ARMS-mediated activation;Gastrin-CREB signalling pathway via PKC and MAPK;Interleukin receptor SHC signaling;FCERI mediated MAPK activation;DAP12 signaling;Signaling by PDGF;VEGFA-VEGFR2 Pathway;Signaling by ERBB2;IRS-related events triggered by IGF1R 
                                               hsa03015;hsa04071;hsa04114;hsa04151;hsa04152;hsa04261;hsa04350;hsa04390;hsa04530;hsa04728;hsa04730;hsa05142;hsa05160 
                                               PPP2R1B 
                                               NA 
                                               azure4 
                                               1 
                                               -1.25969645868465 
                                             
                                             
                                               PPP2R1A-mRNA 
                                               0.0227 
                                               0.0993 
                                               -0.172 
                                               0.217 
                                               1.02 
                                               0.888 
                                               1.16 
                                               0.822 
                                               1 
                                               loglinear 
                                               ARMS-mediated activation, AXIN missense mutants destabilize the destruction complex, Beta-catenin phosphorylation cascade, CTLA4 inhibitory signaling, Cyclin AB1 associated events during G2M transition, Cyclin D associated events in G1, DAP12 signaling, DARPP-32 events, Disassembly of the destruction complex and recruitment of AXIN to the membrane, Downstream signal transduction, FCERI mediated MAPK activation, Frs2-mediated activation, Gastrin-CREB signalling pathway via PKC and MAPK, Interleukin receptor SHC signaling, IRS-related events triggered by IGF1R, MASTL Facilitates Mitotic Progression, Negative regulation of FGFR1 signaling, Negative regulation of FGFR2 signaling, Negative regulation of FGFR3 signaling, Negative regulation of FGFR4 signaling, Nonsense Mediated Decay (NMD) enhanced by the Exon Junction Complex (EJC), RAF activation, RAFMAP kinase cascade, Resolution of Sister Chromatid Cohesion, RHO GTPases Activate Formins, S33 mutants of beta-catenin aren.SQT.t phosphorylated, S37 mutants of beta-catenin aren.SQT.t phosphorylated, S45 mutants of beta-catenin aren.SQT.t phosphorylated, Signaling by ERBB2, Signaling by PDGF, Signaling by SCF-KIT, Spry regulation of FGF signaling, Truncations of AMER1 destabilize the destruction complex, VEGFA-VEGFR2 Pathway, VEGFR2 mediated cell proliferation 
                                               SEBELIUS_2_C4636 
                                               PPP2R1A 
                                               mRNA 
                                               0 
                                               
                                               
                                               MASTL Facilitates Mitotic Progression;Regulation of PLK1 Activity at G2M Transition;Disassembly of the destruction complex and recruitment of AXIN to the membrane;S37 mutants of beta-catenin aren.SQT.t phosphorylated;Cyclin AB1 associated events during G2M transition;RAF activation;RHO GTPases Activate Formins;S33 mutants of beta-catenin aren.SQT.t phosphorylated;Loss of Nlp from mitotic centrosomes;DARPP-32 events;Beta-catenin phosphorylation cascade;Initiation of Nuclear Envelope Reformation;Recruitment of mitotic centrosome proteins and complexes;Spry regulation of FGF signaling;Truncations of AMER1 destabilize the destruction complex;CTLA4 inhibitory signaling;AXIN missense mutants destabilize the destruction complex;S45 mutants of beta-catenin aren.SQT.t phosphorylated;Anchoring of the basal body to the plasma membrane;Nonsense Mediated Decay (NMD) enhanced by the Exon Junction Complex (EJC);Cyclin D associated events in G1;Platelet sensitization by LDL;Resolution of Sister Chromatid Cohesion;Inhibition of replication initiation of damaged DNA by RB1E2F1;RAFMAP kinase cascade;Downstream signal transduction;VEGFR2 mediated cell proliferation;Signaling by SCF-KIT;Frs2-mediated activation;ARMS-mediated activation;Gastrin-CREB signalling pathway via PKC and MAPK;Interleukin receptor SHC signaling;FCERI mediated MAPK activation;DAP12 signaling;Signaling by PDGF;VEGFA-VEGFR2 Pathway;Signaling by ERBB2;IRS-related events triggered by IGF1R;Negative regulation of FGFR3 signaling;Negative regulation of FGFR4 signaling;Negative regulation of FGFR2 signaling;Negative regulation of FGFR1 signaling 
                                               hsa03015;hsa04071;hsa04114;hsa04151;hsa04152;hsa04261;hsa04350;hsa04390;hsa04530;hsa04728;hsa04730;hsa05142;hsa05160 
                                               PPP2R1A 
                                               NA 
                                               azure4 
                                               1 
                                               0.228600201409869 
                                             
                                           
                                           DE Results - Beta-catenin phosphorylation cascade genes - New.Annotation: Control vs.cel67  Table displaying each sample's global significance scores and directed global significance scores as defined in the heatmaps above. The global significance score is calculated as the square root of the mean squared t-statistic for the genes in a gene set, with t-statistics coming from the linear regression underlying our differential expression analysis. The directed global significance score is calculated as the square root of the mean signed squared t-statistic for the genes in a gene set, with t-statistics coming from the linear regression underlying our differential expression analysis.   
                                      
                                       
                                  
                                
                                
                                  
                                       
                                      
                                           Volcano Plot: New.Annotation: miR142 vs.cel67   More Plot Information                Volcano Plot: New.Annotation: miR142 vs.cel67  Volcano plot displaying each gene's -log10(p-value) and log2 fold change for the selected covariate. Highly statistically significant genes fall at the top of the plot, and highly differentially expressed genes fall to either side. Genes within the selected gene set are highlighted in orange. Horizontal lines indicate various False Discovery Rate (FDR) thresholds.   
                                           DE Results - Beta-catenin phosphorylation cascade genes - New.Annotation: miR142 vs.cel67   More Plot Information      Download CSV Data     
                                           
                                             
                                             
                                           
                                           
                                             
                                               Probe Label 
                                               Log2 fold change 
                                               std error (log2) 
                                               Lower confidence limit (log2) 
                                               Upper confidence limit (log2) 
                                               Linear fold change 
                                               Lower confidence limit (linear) 
                                               Upper confidence limit (linear) 
                                               P-value 
                                               BY.p.value 
                                               method 
                                               Gene.sets 
                                               Codeset.Name 
                                               Probe.Label 
                                               Analyte.Type 
                                               Is.Control 
                                               Control.Type 
                                               Related.Probes 
                                               Probe.Annotation 
                                               KEGG.Pathways 
                                               Official.Gene.Name 
                                               Control.Conc 
                                               volcanocol 
                                               volcanopch 
                                               tstats.all 
                                             
                                           
                                           
                                             
                                               CSNK1A1-mRNA 
                                               0.698 
                                               0.0669 
                                               0.567 
                                               0.829 
                                               1.62 
                                               1.48 
                                               1.78 
                                               8.98e-10 
                                               2.38e-07 
                                               loglinear 
                                               AXIN missense mutants destabilize the destruction complex, Beta-catenin phosphorylation cascade, Disassembly of the destruction complex and recruitment of AXIN to the membrane, Misspliced GSK3beta mutants stabilize beta-catenin, S33 mutants of beta-catenin aren.SQT.t phosphorylated, S37 mutants of beta-catenin aren.SQT.t phosphorylated, S45 mutants of beta-catenin aren.SQT.t phosphorylated, T41 mutants of beta-catenin aren.SQT.t phosphorylated 
                                               SEBELIUS_2_C4636 
                                               CSNK1A1 
                                               mRNA 
                                               0 
                                               
                                               
                                               Misspliced GSK3beta mutants stabilize beta-catenin;APC truncation mutants have impaired AXIN binding;T41 mutants of beta-catenin aren.SQT.t phosphorylated;AXIN missense mutants destabilize the destruction complex;S45 mutants of beta-catenin aren.SQT.t phosphorylated;Disassembly of the destruction complex and recruitment of AXIN to the membrane;S37 mutants of beta-catenin aren.SQT.t phosphorylated;S33 mutants of beta-catenin aren.SQT.t phosphorylated;Beta-catenin phosphorylation cascade;Degradation of GLI2 by the proteasome 
                                               hsa04310 
                                               CSNK1A1 
                                               NA 
                                               slateblue1 
                                               16 
                                               10.4334828101644 
                                             
                                             
                                               APC-mRNA 
                                               -0.593 
                                               0.0957 
                                               -0.781 
                                               -0.406 
                                               0.663 
                                               0.582 
                                               0.755 
                                               3.8e-06 
                                               0.000196 
                                               loglinear 
                                               Apoptotic cleavage of cellular proteins, AXIN missense mutants destabilize the destruction complex, Beta-catenin phosphorylation cascade, Deactivation of the beta-catenin transactivating complex, Disassembly of the destruction complex and recruitment of AXIN to the membrane, Misspliced GSK3beta mutants stabilize beta-catenin, S33 mutants of beta-catenin aren.SQT.t phosphorylated, S37 mutants of beta-catenin aren.SQT.t phosphorylated, S45 mutants of beta-catenin aren.SQT.t phosphorylated, T41 mutants of beta-catenin aren.SQT.t phosphorylated, Truncations of AMER1 destabilize the destruction complex 
                                               SEBELIUS_2_C4636 
                                               APC 
                                               mRNA 
                                               0 
                                               
                                               
                                               Apoptotic cleavage of cellular proteins;Misspliced GSK3beta mutants stabilize beta-catenin;APC truncation mutants have impaired AXIN binding;T41 mutants of beta-catenin aren.SQT.t phosphorylated;AXIN missense mutants destabilize the destruction complex;S45 mutants of beta-catenin aren.SQT.t phosphorylated;Deactivation of the beta-catenin transactivating complex;Disassembly of the destruction complex and recruitment of AXIN to the membrane;S37 mutants of beta-catenin aren.SQT.t phosphorylated;S33 mutants of beta-catenin aren.SQT.t phosphorylated;Beta-catenin phosphorylation cascade;Truncations of AMER1 destabilize the destruction complex 
                                               hsa04310;hsa04390;hsa04550;hsa04810;hsa05166;hsa05200;hsa05206;hsa05210;hsa05213;hsa05217 
                                               APC 
                                               NA 
                                               slateblue1 
                                               16 
                                               -6.19644723092999 
                                             
                                             
                                               PPP2CA-mRNA 
                                               -0.205 
                                               0.0527 
                                               -0.309 
                                               -0.102 
                                               0.867 
                                               0.807 
                                               0.932 
                                               0.000827 
                                               0.0186 
                                               loglinear 
                                               ARMS-mediated activation, AXIN missense mutants destabilize the destruction complex, Beta-catenin phosphorylation cascade, CTLA4 inhibitory signaling, Cyclin AB1 associated events during G2M transition, Cyclin D associated events in G1, DAP12 signaling, DARPP-32 events, Disassembly of the destruction complex and recruitment of AXIN to the membrane, Downstream signal transduction, ERKs are inactivated, FCERI mediated MAPK activation, Frs2-mediated activation, Gastrin-CREB signalling pathway via PKC and MAPK, Interleukin receptor SHC signaling, IRS-related events triggered by IGF1R, MASTL Facilitates Mitotic Progression, Negative regulation of FGFR1 signaling, Negative regulation of FGFR2 signaling, Negative regulation of FGFR3 signaling, Negative regulation of FGFR4 signaling, Negative regulation of MAPK pathway, Nonsense Mediated Decay (NMD) enhanced by the Exon Junction Complex (EJC), RAF activation, RAFMAP kinase cascade, Resolution of Sister Chromatid Cohesion, RHO GTPases Activate Formins, S33 mutants of beta-catenin aren.SQT.t phosphorylated, S37 mutants of beta-catenin aren.SQT.t phosphorylated, S45 mutants of beta-catenin aren.SQT.t phosphorylated, Signaling by ERBB2, Signaling by PDGF, Signaling by SCF-KIT, Spry regulation of FGF signaling, T41 mutants of beta-catenin aren.SQT.t phosphorylated, Truncations of AMER1 destabilize the destruction complex, VEGFA-VEGFR2 Pathway, VEGFR2 mediated cell proliferation 
                                               SEBELIUS_2_C4636 
                                               PPP2CA 
                                               mRNA 
                                               0 
                                               
                                               
                                               MASTL Facilitates Mitotic Progression;T41 mutants of beta-catenin aren.SQT.t phosphorylated;ERKs are inactivated;Glycolysis;Negative regulation of MAPK pathway;Disassembly of the destruction complex and recruitment of AXIN to the membrane;S37 mutants of beta-catenin aren.SQT.t phosphorylated;Cyclin AB1 associated events during G2M transition;RAF activation;RHO GTPases Activate Formins;S33 mutants of beta-catenin aren.SQT.t phosphorylated;DARPP-32 events;Beta-catenin phosphorylation cascade;Initiation of Nuclear Envelope Reformation;Spry regulation of FGF signaling;Truncations of AMER1 destabilize the destruction complex;CTLA4 inhibitory signaling;AXIN missense mutants destabilize the destruction complex;S45 mutants of beta-catenin aren.SQT.t phosphorylated;Nonsense Mediated Decay (NMD) enhanced by the Exon Junction Complex (EJC);Cyclin D associated events in G1;Platelet sensitization by LDL;Resolution of Sister Chromatid Cohesion;Inhibition of replication initiation of damaged DNA by RB1E2F1;RAFMAP kinase cascade;Downstream signal transduction;VEGFR2 mediated cell proliferation;Signaling by SCF-KIT;Frs2-mediated activation;ARMS-mediated activation;Gastrin-CREB signalling pathway via PKC and MAPK;Interleukin receptor SHC signaling;FCERI mediated MAPK activation;DAP12 signaling;Signaling by PDGF;VEGFA-VEGFR2 Pathway;Signaling by ERBB2;IRS-related events triggered by IGF1R;Negative regulation of FGFR3 signaling;Negative regulation of FGFR4 signaling;Negative regulation of FGFR2 signaling;Negative regulation of FGFR1 signaling 
                                               hsa03015;hsa04071;hsa04114;hsa04151;hsa04152;hsa04261;hsa04350;hsa04390;hsa04530;hsa04728;hsa04730;hsa05142;hsa05160 
                                               PPP2CA 
                                               NA 
                                               slateblue2 
                                               16 
                                               -3.8899430740038 
                                             
                                             
                                               PPP2R1A-mRNA 
                                               -0.339 
                                               0.0993 
                                               -0.533 
                                               -0.144 
                                               0.791 
                                               0.691 
                                               0.905 
                                               0.00262 
                                               0.0492 
                                               loglinear 
                                               ARMS-mediated activation, AXIN missense mutants destabilize the destruction complex, Beta-catenin phosphorylation cascade, CTLA4 inhibitory signaling, Cyclin AB1 associated events during G2M transition, Cyclin D associated events in G1, DAP12 signaling, DARPP-32 events, Disassembly of the destruction complex and recruitment of AXIN to the membrane, Downstream signal transduction, FCERI mediated MAPK activation, Frs2-mediated activation, Gastrin-CREB signalling pathway via PKC and MAPK, Interleukin receptor SHC signaling, IRS-related events triggered by IGF1R, MASTL Facilitates Mitotic Progression, Negative regulation of FGFR1 signaling, Negative regulation of FGFR2 signaling, Negative regulation of FGFR3 signaling, Negative regulation of FGFR4 signaling, Nonsense Mediated Decay (NMD) enhanced by the Exon Junction Complex (EJC), RAF activation, RAFMAP kinase cascade, Resolution of Sister Chromatid Cohesion, RHO GTPases Activate Formins, S33 mutants of beta-catenin aren.SQT.t phosphorylated, S37 mutants of beta-catenin aren.SQT.t phosphorylated, S45 mutants of beta-catenin aren.SQT.t phosphorylated, Signaling by ERBB2, Signaling by PDGF, Signaling by SCF-KIT, Spry regulation of FGF signaling, Truncations of AMER1 destabilize the destruction complex, VEGFA-VEGFR2 Pathway, VEGFR2 mediated cell proliferation 
                                               SEBELIUS_2_C4636 
                                               PPP2R1A 
                                               mRNA 
                                               0 
                                               
                                               
                                               MASTL Facilitates Mitotic Progression;Regulation of PLK1 Activity at G2M Transition;Disassembly of the destruction complex and recruitment of AXIN to the membrane;S37 mutants of beta-catenin aren.SQT.t phosphorylated;Cyclin AB1 associated events during G2M transition;RAF activation;RHO GTPases Activate Formins;S33 mutants of beta-catenin aren.SQT.t phosphorylated;Loss of Nlp from mitotic centrosomes;DARPP-32 events;Beta-catenin phosphorylation cascade;Initiation of Nuclear Envelope Reformation;Recruitment of mitotic centrosome proteins and complexes;Spry regulation of FGF signaling;Truncations of AMER1 destabilize the destruction complex;CTLA4 inhibitory signaling;AXIN missense mutants destabilize the destruction complex;S45 mutants of beta-catenin aren.SQT.t phosphorylated;Anchoring of the basal body to the plasma membrane;Nonsense Mediated Decay (NMD) enhanced by the Exon Junction Complex (EJC);Cyclin D associated events in G1;Platelet sensitization by LDL;Resolution of Sister Chromatid Cohesion;Inhibition of replication initiation of damaged DNA by RB1E2F1;RAFMAP kinase cascade;Downstream signal transduction;VEGFR2 mediated cell proliferation;Signaling by SCF-KIT;Frs2-mediated activation;ARMS-mediated activation;Gastrin-CREB signalling pathway via PKC and MAPK;Interleukin receptor SHC signaling;FCERI mediated MAPK activation;DAP12 signaling;Signaling by PDGF;VEGFA-VEGFR2 Pathway;Signaling by ERBB2;IRS-related events triggered by IGF1R;Negative regulation of FGFR3 signaling;Negative regulation of FGFR4 signaling;Negative regulation of FGFR2 signaling;Negative regulation of FGFR1 signaling 
                                               hsa03015;hsa04071;hsa04114;hsa04151;hsa04152;hsa04261;hsa04350;hsa04390;hsa04530;hsa04728;hsa04730;hsa05142;hsa05160 
                                               PPP2R1A 
                                               NA 
                                               slateblue2 
                                               16 
                                               -3.41389728096677 
                                             
                                             
                                               CTNNB1-mRNA 
                                               0.0818 
                                               0.056 
                                               -0.0278 
                                               0.191 
                                               1.06 
                                               0.981 
                                               1.14 
                                               0.158 
                                               1 
                                               loglinear 
                                               Apoptotic cleavage of cellular proteins, Beta-catenin phosphorylation cascade, Deactivation of the beta-catenin transactivating complex, Disassembly of the destruction complex and recruitment of AXIN to the membrane, Formation of the beta-catenin.CL.TCF transactivating complex, Misspliced GSK3beta mutants stabilize beta-catenin, Repression of WNT target genes, RHO GTPases activate IQGAPs, S37 mutants of beta-catenin aren.SQT.t phosphorylated, S45 mutants of beta-catenin aren.SQT.t phosphorylated, T41 mutants of beta-catenin aren.SQT.t phosphorylated, VEGFA-VEGFR2 Pathway, VEGFR2 mediated vascular permeability 
                                               SEBELIUS_2_C4636 
                                               CTNNB1 
                                               mRNA 
                                               0 
                                               
                                               
                                               Disassembly of the destruction complex and recruitment of AXIN to the membrane;S37 mutants of beta-catenin aren.SQT.t phosphorylated;T41 mutants of beta-catenin aren.SQT.t phosphorylated;S45 mutants of beta-catenin aren.SQT.t phosphorylated;Binding of TCFLEF.CL.CTNNB1 to target gene promoters;RHO GTPases activate IQGAPs;Deactivation of the beta-catenin transactivating complex;Repression of WNT target genes;Apoptotic cleavage of cell adhesion  proteins;Adherens junctions interactions;Misspliced GSK3beta mutants stabilize beta-catenin;VEGFR2 mediated vascular permeability;Beta-catenin phosphorylation cascade;Synthesis# secretion# and inactivation of Glucagon-like Peptide-1 (GLP-1);Formation of the beta-catenin.CL.TCF transactivating complex;Apoptotic cleavage of cellular proteins;VEGFA-VEGFR2 Pathway 
                                               hsa04015;hsa04310;hsa04390;hsa04510;hsa04520;hsa04530;hsa04550;hsa04670;hsa04916;hsa04919;hsa05100;hsa05130;hsa05166;hsa05200;hsa05205;hsa05210;hsa05213;hsa05215;hsa05216;hsa05217;hsa05412 
                                               CTNNB1 
                                               NA 
                                               azure4 
                                               1 
                                               1.46071428571429 
                                             
                                             
                                               PPP2R1B-mRNA 
                                               0.0724 
                                               0.0593 
                                               -0.0439 
                                               0.189 
                                               1.05 
                                               0.97 
                                               1.14 
                                               0.236 
                                               1 
                                               loglinear 
                                               ARMS-mediated activation, AXIN missense mutants destabilize the destruction complex, Beta-catenin phosphorylation cascade, CTLA4 inhibitory signaling, Cyclin AB1 associated events during G2M transition, Cyclin D associated events in G1, DAP12 signaling, DARPP-32 events, Disassembly of the destruction complex and recruitment of AXIN to the membrane, Downstream signal transduction, ERKs are inactivated, FCERI mediated MAPK activation, Frs2-mediated activation, Gastrin-CREB signalling pathway via PKC and MAPK, Interleukin receptor SHC signaling, IRS-related events triggered by IGF1R, MASTL Facilitates Mitotic Progression, Misspliced GSK3beta mutants stabilize beta-catenin, Negative regulation of MAPK pathway, RAF activation, RAFMAP kinase cascade, Resolution of Sister Chromatid Cohesion, RHO GTPases Activate Formins, S33 mutants of beta-catenin aren.SQT.t phosphorylated, S37 mutants of beta-catenin aren.SQT.t phosphorylated, S45 mutants of beta-catenin aren.SQT.t phosphorylated, Signaling by ERBB2, Signaling by PDGF, Signaling by SCF-KIT, T41 mutants of beta-catenin aren.SQT.t phosphorylated, Truncations of AMER1 destabilize the destruction complex, VEGFA-VEGFR2 Pathway, VEGFR2 mediated cell proliferation 
                                               SEBELIUS_2_C4636 
                                               PPP2R1B 
                                               mRNA 
                                               0 
                                               
                                               
                                               MASTL Facilitates Mitotic Progression;Misspliced GSK3beta mutants stabilize beta-catenin;T41 mutants of beta-catenin aren.SQT.t phosphorylated;ERKs are inactivated;Glycolysis;Negative regulation of MAPK pathway;Disassembly of the destruction complex and recruitment of AXIN to the membrane;S37 mutants of beta-catenin aren.SQT.t phosphorylated;Cyclin AB1 associated events during G2M transition;RAF activation;RHO GTPases Activate Formins;S33 mutants of beta-catenin aren.SQT.t phosphorylated;DARPP-32 events;Beta-catenin phosphorylation cascade;Truncations of AMER1 destabilize the destruction complex;CTLA4 inhibitory signaling;AXIN missense mutants destabilize the destruction complex;S45 mutants of beta-catenin aren.SQT.t phosphorylated;Cyclin D associated events in G1;Platelet sensitization by LDL;Resolution of Sister Chromatid Cohesion;Inhibition of replication initiation of damaged DNA by RB1E2F1;RAFMAP kinase cascade;Downstream signal transduction;VEGFR2 mediated cell proliferation;Signaling by SCF-KIT;Frs2-mediated activation;ARMS-mediated activation;Gastrin-CREB signalling pathway via PKC and MAPK;Interleukin receptor SHC signaling;FCERI mediated MAPK activation;DAP12 signaling;Signaling by PDGF;VEGFA-VEGFR2 Pathway;Signaling by ERBB2;IRS-related events triggered by IGF1R 
                                               hsa03015;hsa04071;hsa04114;hsa04151;hsa04152;hsa04261;hsa04350;hsa04390;hsa04530;hsa04728;hsa04730;hsa05142;hsa05160 
                                               PPP2R1B 
                                               NA 
                                               azure4 
                                               1 
                                               1.22091062394604 
                                             
                                             
                                               AXIN1-mRNA 
                                               -0.15 
                                               0.211 
                                               -0.564 
                                               0.264 
                                               0.901 
                                               0.677 
                                               1.2 
                                               0.486 
                                               1 
                                               lm.nb 
                                               AXIN missense mutants destabilize the destruction complex, Beta-catenin phosphorylation cascade, Disassembly of the destruction complex and recruitment of AXIN to the membrane, Misspliced GSK3beta mutants stabilize beta-catenin, S33 mutants of beta-catenin aren.SQT.t phosphorylated, S37 mutants of beta-catenin aren.SQT.t phosphorylated, S45 mutants of beta-catenin aren.SQT.t phosphorylated, T41 mutants of beta-catenin aren.SQT.t phosphorylated, Truncations of AMER1 destabilize the destruction complex 
                                               SEBELIUS_2_C4636 
                                               AXIN1 
                                               mRNA 
                                               0 
                                               
                                               
                                               APC truncation mutants have impaired AXIN binding;Misspliced GSK3beta mutants stabilize beta-catenin;T41 mutants of beta-catenin aren.SQT.t phosphorylated;AXIN missense mutants destabilize the destruction complex;S45 mutants of beta-catenin aren.SQT.t phosphorylated;Disassembly of the destruction complex and recruitment of AXIN to the membrane;S37 mutants of beta-catenin aren.SQT.t phosphorylated;S33 mutants of beta-catenin aren.SQT.t phosphorylated;Beta-catenin phosphorylation cascade;Truncations of AMER1 destabilize the destruction complex 
                                               hsa04310;hsa04390;hsa04550;hsa05200;hsa05210;hsa05213;hsa05217 
                                               AXIN1 
                                               NA 
                                               azure4 
                                               1 
                                               -0.710900473933649 
                                             
                                             
                                               PPP2CB-mRNA 
                                               -0.0393 
                                               0.0747 
                                               -0.186 
                                               0.107 
                                               0.973 
                                               0.879 
                                               1.08 
                                               0.604 
                                               1 
                                               loglinear 
                                               ARMS-mediated activation, AXIN missense mutants destabilize the destruction complex, Beta-catenin phosphorylation cascade, CTLA4 inhibitory signaling, Cyclin AB1 associated events during G2M transition, Cyclin D associated events in G1, DAP12 signaling, DARPP-32 events, Disassembly of the destruction complex and recruitment of AXIN to the membrane, Downstream signal transduction, ERKs are inactivated, FCERI mediated MAPK activation, Frs2-mediated activation, Gastrin-CREB signalling pathway via PKC and MAPK, Interleukin receptor SHC signaling, IRS-related events triggered by IGF1R, MASTL Facilitates Mitotic Progression, Misspliced GSK3beta mutants stabilize beta-catenin, Negative regulation of FGFR1 signaling, Negative regulation of FGFR2 signaling, Negative regulation of FGFR3 signaling, Negative regulation of FGFR4 signaling, Negative regulation of MAPK pathway, RAF activation, RAFMAP kinase cascade, Resolution of Sister Chromatid Cohesion, RHO GTPases Activate Formins, S33 mutants of beta-catenin aren.SQT.t phosphorylated, S37 mutants of beta-catenin aren.SQT.t phosphorylated, S45 mutants of beta-catenin aren.SQT.t phosphorylated, Signaling by ERBB2, Signaling by PDGF, Signaling by SCF-KIT, Spry regulation of FGF signaling, T41 mutants of beta-catenin aren.SQT.t phosphorylated, Truncations of AMER1 destabilize the destruction complex, VEGFA-VEGFR2 Pathway, VEGFR2 mediated cell proliferation 
                                               SEBELIUS_2_C4636 
                                               PPP2CB 
                                               mRNA 
                                               0 
                                               
                                               
                                               MASTL Facilitates Mitotic Progression;Misspliced GSK3beta mutants stabilize beta-catenin;T41 mutants of beta-catenin aren.SQT.t phosphorylated;ERKs are inactivated;Glycolysis;Negative regulation of MAPK pathway;Disassembly of the destruction complex and recruitment of AXIN to the membrane;S37 mutants of beta-catenin aren.SQT.t phosphorylated;Cyclin AB1 associated events during G2M transition;RAF activation;RHO GTPases Activate Formins;S33 mutants of beta-catenin aren.SQT.t phosphorylated;DARPP-32 events;Beta-catenin phosphorylation cascade;Spry regulation of FGF signaling;Truncations of AMER1 destabilize the destruction complex;CTLA4 inhibitory signaling;AXIN missense mutants destabilize the destruction complex;S45 mutants of beta-catenin aren.SQT.t phosphorylated;Cyclin D associated events in G1;Platelet sensitization by LDL;Resolution of Sister Chromatid Cohesion;Inhibition of replication initiation of damaged DNA by RB1E2F1;RAFMAP kinase cascade;Downstream signal transduction;VEGFR2 mediated cell proliferation;Signaling by SCF-KIT;Frs2-mediated activation;ARMS-mediated activation;Gastrin-CREB signalling pathway via PKC and MAPK;Interleukin receptor SHC signaling;FCERI mediated MAPK activation;DAP12 signaling;Signaling by PDGF;VEGFA-VEGFR2 Pathway;Signaling by ERBB2;IRS-related events triggered by IGF1R;Negative regulation of FGFR3 signaling;Negative regulation of FGFR4 signaling;Negative regulation of FGFR2 signaling;Negative regulation of FGFR1 signaling 
                                               hsa03015;hsa04071;hsa04114;hsa04151;hsa04152;hsa04261;hsa04350;hsa04390;hsa04530;hsa04728;hsa04730;hsa05142;hsa05160 
                                               PPP2CB 
                                               NA 
                                               azure4 
                                               1 
                                               -0.526104417670683 
                                             
                                           
                                           DE Results - Beta-catenin phosphorylation cascade genes - New.Annotation: miR142 vs.cel67  Table displaying each sample's global significance scores and directed global significance scores as defined in the heatmaps above. The global significance score is calculated as the square root of the mean squared t-statistic for the genes in a gene set, with t-statistics coming from the linear regression underlying our differential expression analysis. The directed global significance score is calculated as the square root of the mean signed squared t-statistic for the genes in a gene set, with t-statistics coming from the linear regression underlying our differential expression analysis.   
                                      
                                       
                                  
                                
                                
                                  
                                       
                                      
                                           Volcano Plot: New.Annotation: miR155 vs.cel67   More Plot Information                Volcano Plot: New.Annotation: miR155 vs.cel67  Volcano plot displaying each gene's -log10(p-value) and log2 fold change for the selected covariate. Highly statistically significant genes fall at the top of the plot, and highly differentially expressed genes fall to either side. Genes within the selected gene set are highlighted in orange. Horizontal lines indicate various False Discovery Rate (FDR) thresholds.   
                                           DE Results - Beta-catenin phosphorylation cascade genes - New.Annotation: miR155 vs.cel67   More Plot Information      Download CSV Data     
                                           
                                             
                                             
                                           
                                           
                                             
                                               Probe Label 
                                               Log2 fold change 
                                               std error (log2) 
                                               Lower confidence limit (log2) 
                                               Upper confidence limit (log2) 
                                               Linear fold change 
                                               Lower confidence limit (linear) 
                                               Upper confidence limit (linear) 
                                               P-value 
                                               BY.p.value 
                                               method 
                                               Gene.sets 
                                               Codeset.Name 
                                               Probe.Label 
                                               Analyte.Type 
                                               Is.Control 
                                               Control.Type 
                                               Related.Probes 
                                               Probe.Annotation 
                                               KEGG.Pathways 
                                               Official.Gene.Name 
                                               Control.Conc 
                                               volcanocol 
                                               volcanopch 
                                               tstats.all 
                                             
                                           
                                           
                                             
                                               PPP2CB-mRNA 
                                               -1.13 
                                               0.0787 
                                               -1.29 
                                               -0.979 
                                               0.456 
                                               0.41 
                                               0.507 
                                               2.34e-12 
                                               3.11e-09 
                                               loglinear 
                                               ARMS-mediated activation, AXIN missense mutants destabilize the destruction complex, Beta-catenin phosphorylation cascade, CTLA4 inhibitory signaling, Cyclin AB1 associated events during G2M transition, Cyclin D associated events in G1, DAP12 signaling, DARPP-32 events, Disassembly of the destruction complex and recruitment of AXIN to the membrane, Downstream signal transduction, ERKs are inactivated, FCERI mediated MAPK activation, Frs2-mediated activation, Gastrin-CREB signalling pathway via PKC and MAPK, Interleukin receptor SHC signaling, IRS-related events triggered by IGF1R, MASTL Facilitates Mitotic Progression, Misspliced GSK3beta mutants stabilize beta-catenin, Negative regulation of FGFR1 signaling, Negative regulation of FGFR2 signaling, Negative regulation of FGFR3 signaling, Negative regulation of FGFR4 signaling, Negative regulation of MAPK pathway, RAF activation, RAFMAP kinase cascade, Resolution of Sister Chromatid Cohesion, RHO GTPases Activate Formins, S33 mutants of beta-catenin aren.SQT.t phosphorylated, S37 mutants of beta-catenin aren.SQT.t phosphorylated, S45 mutants of beta-catenin aren.SQT.t phosphorylated, Signaling by ERBB2, Signaling by PDGF, Signaling by SCF-KIT, Spry regulation of FGF signaling, T41 mutants of beta-catenin aren.SQT.t phosphorylated, Truncations of AMER1 destabilize the destruction complex, VEGFA-VEGFR2 Pathway, VEGFR2 mediated cell proliferation 
                                               SEBELIUS_2_C4636 
                                               PPP2CB 
                                               mRNA 
                                               0 
                                               
                                               
                                               MASTL Facilitates Mitotic Progression;Misspliced GSK3beta mutants stabilize beta-catenin;T41 mutants of beta-catenin aren.SQT.t phosphorylated;ERKs are inactivated;Glycolysis;Negative regulation of MAPK pathway;Disassembly of the destruction complex and recruitment of AXIN to the membrane;S37 mutants of beta-catenin aren.SQT.t phosphorylated;Cyclin AB1 associated events during G2M transition;RAF activation;RHO GTPases Activate Formins;S33 mutants of beta-catenin aren.SQT.t phosphorylated;DARPP-32 events;Beta-catenin phosphorylation cascade;Spry regulation of FGF signaling;Truncations of AMER1 destabilize the destruction complex;CTLA4 inhibitory signaling;AXIN missense mutants destabilize the destruction complex;S45 mutants of beta-catenin aren.SQT.t phosphorylated;Cyclin D associated events in G1;Platelet sensitization by LDL;Resolution of Sister Chromatid Cohesion;Inhibition of replication initiation of damaged DNA by RB1E2F1;RAFMAP kinase cascade;Downstream signal transduction;VEGFR2 mediated cell proliferation;Signaling by SCF-KIT;Frs2-mediated activation;ARMS-mediated activation;Gastrin-CREB signalling pathway via PKC and MAPK;Interleukin receptor SHC signaling;FCERI mediated MAPK activation;DAP12 signaling;Signaling by PDGF;VEGFA-VEGFR2 Pathway;Signaling by ERBB2;IRS-related events triggered by IGF1R;Negative regulation of FGFR3 signaling;Negative regulation of FGFR4 signaling;Negative regulation of FGFR2 signaling;Negative regulation of FGFR1 signaling 
                                               hsa03015;hsa04071;hsa04114;hsa04151;hsa04152;hsa04261;hsa04350;hsa04390;hsa04530;hsa04728;hsa04730;hsa05142;hsa05160 
                                               PPP2CB 
                                               NA 
                                               slateblue1 
                                               16 
                                               -14.3583227445997 
                                             
                                             
                                               CSNK1A1-mRNA 
                                               -0.601 
                                               0.0705 
                                               -0.739 
                                               -0.463 
                                               0.659 
                                               0.599 
                                               0.725 
                                               2.89e-08 
                                               6.39e-06 
                                               loglinear 
                                               AXIN missense mutants destabilize the destruction complex, Beta-catenin phosphorylation cascade, Disassembly of the destruction complex and recruitment of AXIN to the membrane, Misspliced GSK3beta mutants stabilize beta-catenin, S33 mutants of beta-catenin aren.SQT.t phosphorylated, S37 mutants of beta-catenin aren.SQT.t phosphorylated, S45 mutants of beta-catenin aren.SQT.t phosphorylated, T41 mutants of beta-catenin aren.SQT.t phosphorylated 
                                               SEBELIUS_2_C4636 
                                               CSNK1A1 
                                               mRNA 
                                               0 
                                               
                                               
                                               Misspliced GSK3beta mutants stabilize beta-catenin;APC truncation mutants have impaired AXIN binding;T41 mutants of beta-catenin aren.SQT.t phosphorylated;AXIN missense mutants destabilize the destruction complex;S45 mutants of beta-catenin aren.SQT.t phosphorylated;Disassembly of the destruction complex and recruitment of AXIN to the membrane;S37 mutants of beta-catenin aren.SQT.t phosphorylated;S33 mutants of beta-catenin aren.SQT.t phosphorylated;Beta-catenin phosphorylation cascade;Degradation of GLI2 by the proteasome 
                                               hsa04310 
                                               CSNK1A1 
                                               NA 
                                               slateblue1 
                                               16 
                                               -8.52482269503546 
                                             
                                             
                                               PPP2CA-mRNA 
                                               -0.221 
                                               0.0556 
                                               -0.33 
                                               -0.112 
                                               0.858 
                                               0.796 
                                               0.925 
                                               0.000682 
                                               0.0404 
                                               loglinear 
                                               ARMS-mediated activation, AXIN missense mutants destabilize the destruction complex, Beta-catenin phosphorylation cascade, CTLA4 inhibitory signaling, Cyclin AB1 associated events during G2M transition, Cyclin D associated events in G1, DAP12 signaling, DARPP-32 events, Disassembly of the destruction complex and recruitment of AXIN to the membrane, Downstream signal transduction, ERKs are inactivated, FCERI mediated MAPK activation, Frs2-mediated activation, Gastrin-CREB signalling pathway via PKC and MAPK, Interleukin receptor SHC signaling, IRS-related events triggered by IGF1R, MASTL Facilitates Mitotic Progression, Negative regulation of FGFR1 signaling, Negative regulation of FGFR2 signaling, Negative regulation of FGFR3 signaling, Negative regulation of FGFR4 signaling, Negative regulation of MAPK pathway, Nonsense Mediated Decay (NMD) enhanced by the Exon Junction Complex (EJC), RAF activation, RAFMAP kinase cascade, Resolution of Sister Chromatid Cohesion, RHO GTPases Activate Formins, S33 mutants of beta-catenin aren.SQT.t phosphorylated, S37 mutants of beta-catenin aren.SQT.t phosphorylated, S45 mutants of beta-catenin aren.SQT.t phosphorylated, Signaling by ERBB2, Signaling by PDGF, Signaling by SCF-KIT, Spry regulation of FGF signaling, T41 mutants of beta-catenin aren.SQT.t phosphorylated, Truncations of AMER1 destabilize the destruction complex, VEGFA-VEGFR2 Pathway, VEGFR2 mediated cell proliferation 
                                               SEBELIUS_2_C4636 
                                               PPP2CA 
                                               mRNA 
                                               0 
                                               
                                               
                                               MASTL Facilitates Mitotic Progression;T41 mutants of beta-catenin aren.SQT.t phosphorylated;ERKs are inactivated;Glycolysis;Negative regulation of MAPK pathway;Disassembly of the destruction complex and recruitment of AXIN to the membrane;S37 mutants of beta-catenin aren.SQT.t phosphorylated;Cyclin AB1 associated events during G2M transition;RAF activation;RHO GTPases Activate Formins;S33 mutants of beta-catenin aren.SQT.t phosphorylated;DARPP-32 events;Beta-catenin phosphorylation cascade;Initiation of Nuclear Envelope Reformation;Spry regulation of FGF signaling;Truncations of AMER1 destabilize the destruction complex;CTLA4 inhibitory signaling;AXIN missense mutants destabilize the destruction complex;S45 mutants of beta-catenin aren.SQT.t phosphorylated;Nonsense Mediated Decay (NMD) enhanced by the Exon Junction Complex (EJC);Cyclin D associated events in G1;Platelet sensitization by LDL;Resolution of Sister Chromatid Cohesion;Inhibition of replication initiation of damaged DNA by RB1E2F1;RAFMAP kinase cascade;Downstream signal transduction;VEGFR2 mediated cell proliferation;Signaling by SCF-KIT;Frs2-mediated activation;ARMS-mediated activation;Gastrin-CREB signalling pathway via PKC and MAPK;Interleukin receptor SHC signaling;FCERI mediated MAPK activation;DAP12 signaling;Signaling by PDGF;VEGFA-VEGFR2 Pathway;Signaling by ERBB2;IRS-related events triggered by IGF1R;Negative regulation of FGFR3 signaling;Negative regulation of FGFR4 signaling;Negative regulation of FGFR2 signaling;Negative regulation of FGFR1 signaling 
                                               hsa03015;hsa04071;hsa04114;hsa04151;hsa04152;hsa04261;hsa04350;hsa04390;hsa04530;hsa04728;hsa04730;hsa05142;hsa05160 
                                               PPP2CA 
                                               NA 
                                               slateblue2 
                                               16 
                                               -3.97482014388489 
                                             
                                             
                                               CTNNB1-mRNA 
                                               0.163 
                                               0.059 
                                               0.0475 
                                               0.279 
                                               1.12 
                                               1.03 
                                               1.21 
                                               0.0116 
                                               0.28 
                                               loglinear 
                                               Apoptotic cleavage of cellular proteins, Beta-catenin phosphorylation cascade, Deactivation of the beta-catenin transactivating complex, Disassembly of the destruction complex and recruitment of AXIN to the membrane, Formation of the beta-catenin.CL.TCF transactivating complex, Misspliced GSK3beta mutants stabilize beta-catenin, Repression of WNT target genes, RHO GTPases activate IQGAPs, S37 mutants of beta-catenin aren.SQT.t phosphorylated, S45 mutants of beta-catenin aren.SQT.t phosphorylated, T41 mutants of beta-catenin aren.SQT.t phosphorylated, VEGFA-VEGFR2 Pathway, VEGFR2 mediated vascular permeability 
                                               SEBELIUS_2_C4636 
                                               CTNNB1 
                                               mRNA 
                                               0 
                                               
                                               
                                               Disassembly of the destruction complex and recruitment of AXIN to the membrane;S37 mutants of beta-catenin aren.SQT.t phosphorylated;T41 mutants of beta-catenin aren.SQT.t phosphorylated;S45 mutants of beta-catenin aren.SQT.t phosphorylated;Binding of TCFLEF.CL.CTNNB1 to target gene promoters;RHO GTPases activate IQGAPs;Deactivation of the beta-catenin transactivating complex;Repression of WNT target genes;Apoptotic cleavage of cell adhesion  proteins;Adherens junctions interactions;Misspliced GSK3beta mutants stabilize beta-catenin;VEGFR2 mediated vascular permeability;Beta-catenin phosphorylation cascade;Synthesis# secretion# and inactivation of Glucagon-like Peptide-1 (GLP-1);Formation of the beta-catenin.CL.TCF transactivating complex;Apoptotic cleavage of cellular proteins;VEGFA-VEGFR2 Pathway 
                                               hsa04015;hsa04310;hsa04390;hsa04510;hsa04520;hsa04530;hsa04550;hsa04670;hsa04916;hsa04919;hsa05100;hsa05130;hsa05166;hsa05200;hsa05205;hsa05210;hsa05213;hsa05215;hsa05216;hsa05217;hsa05412 
                                               CTNNB1 
                                               NA 
                                               slateblue4 
                                               16 
                                               2.76271186440678 
                                             
                                             
                                               APC-mRNA 
                                               0.261 
                                               0.101 
                                               0.0632 
                                               0.459 
                                               1.2 
                                               1.04 
                                               1.37 
                                               0.0172 
                                               0.327 
                                               loglinear 
                                               Apoptotic cleavage of cellular proteins, AXIN missense mutants destabilize the destruction complex, Beta-catenin phosphorylation cascade, Deactivation of the beta-catenin transactivating complex, Disassembly of the destruction complex and recruitment of AXIN to the membrane, Misspliced GSK3beta mutants stabilize beta-catenin, S33 mutants of beta-catenin aren.SQT.t phosphorylated, S37 mutants of beta-catenin aren.SQT.t phosphorylated, S45 mutants of beta-catenin aren.SQT.t phosphorylated, T41 mutants of beta-catenin aren.SQT.t phosphorylated, Truncations of AMER1 destabilize the destruction complex 
                                               SEBELIUS_2_C4636 
                                               APC 
                                               mRNA 
                                               0 
                                               
                                               
                                               Apoptotic cleavage of cellular proteins;Misspliced GSK3beta mutants stabilize beta-catenin;APC truncation mutants have impaired AXIN binding;T41 mutants of beta-catenin aren.SQT.t phosphorylated;AXIN missense mutants destabilize the destruction complex;S45 mutants of beta-catenin aren.SQT.t phosphorylated;Deactivation of the beta-catenin transactivating complex;Disassembly of the destruction complex and recruitment of AXIN to the membrane;S37 mutants of beta-catenin aren.SQT.t phosphorylated;S33 mutants of beta-catenin aren.SQT.t phosphorylated;Beta-catenin phosphorylation cascade;Truncations of AMER1 destabilize the destruction complex 
                                               hsa04310;hsa04390;hsa04550;hsa04810;hsa05166;hsa05200;hsa05206;hsa05210;hsa05213;hsa05217 
                                               APC 
                                               NA 
                                               slateblue4 
                                               16 
                                               2.58415841584158 
                                             
                                             
                                               PPP2R1B-mRNA 
                                               -0.108 
                                               0.0625 
                                               -0.23 
                                               0.0148 
                                               0.928 
                                               0.852 
                                               1.01 
                                               0.0996 
                                               1 
                                               loglinear 
                                               ARMS-mediated activation, AXIN missense mutants destabilize the destruction complex, Beta-catenin phosphorylation cascade, CTLA4 inhibitory signaling, Cyclin AB1 associated events during G2M transition, Cyclin D associated events in G1, DAP12 signaling, DARPP-32 events, Disassembly of the destruction complex and recruitment of AXIN to the membrane, Downstream signal transduction, ERKs are inactivated, FCERI mediated MAPK activation, Frs2-mediated activation, Gastrin-CREB signalling pathway via PKC and MAPK, Interleukin receptor SHC signaling, IRS-related events triggered by IGF1R, MASTL Facilitates Mitotic Progression, Misspliced GSK3beta mutants stabilize beta-catenin, Negative regulation of MAPK pathway, RAF activation, RAFMAP kinase cascade, Resolution of Sister Chromatid Cohesion, RHO GTPases Activate Formins, S33 mutants of beta-catenin aren.SQT.t phosphorylated, S37 mutants of beta-catenin aren.SQT.t phosphorylated, S45 mutants of beta-catenin aren.SQT.t phosphorylated, Signaling by ERBB2, Signaling by PDGF, Signaling by SCF-KIT, T41 mutants of beta-catenin aren.SQT.t phosphorylated, Truncations of AMER1 destabilize the destruction complex, VEGFA-VEGFR2 Pathway, VEGFR2 mediated cell proliferation 
                                               SEBELIUS_2_C4636 
                                               PPP2R1B 
                                               mRNA 
                                               0 
                                               
                                               
                                               MASTL Facilitates Mitotic Progression;Misspliced GSK3beta mutants stabilize beta-catenin;T41 mutants of beta-catenin aren.SQT.t phosphorylated;ERKs are inactivated;Glycolysis;Negative regulation of MAPK pathway;Disassembly of the destruction complex and recruitment of AXIN to the membrane;S37 mutants of beta-catenin aren.SQT.t phosphorylated;Cyclin AB1 associated events during G2M transition;RAF activation;RHO GTPases Activate Formins;S33 mutants of beta-catenin aren.SQT.t phosphorylated;DARPP-32 events;Beta-catenin phosphorylation cascade;Truncations of AMER1 destabilize the destruction complex;CTLA4 inhibitory signaling;AXIN missense mutants destabilize the destruction complex;S45 mutants of beta-catenin aren.SQT.t phosphorylated;Cyclin D associated events in G1;Platelet sensitization by LDL;Resolution of Sister Chromatid Cohesion;Inhibition of replication initiation of damaged DNA by RB1E2F1;RAFMAP kinase cascade;Downstream signal transduction;VEGFR2 mediated cell proliferation;Signaling by SCF-KIT;Frs2-mediated activation;ARMS-mediated activation;Gastrin-CREB signalling pathway via PKC and MAPK;Interleukin receptor SHC signaling;FCERI mediated MAPK activation;DAP12 signaling;Signaling by PDGF;VEGFA-VEGFR2 Pathway;Signaling by ERBB2;IRS-related events triggered by IGF1R 
                                               hsa03015;hsa04071;hsa04114;hsa04151;hsa04152;hsa04261;hsa04350;hsa04390;hsa04530;hsa04728;hsa04730;hsa05142;hsa05160 
                                               PPP2R1B 
                                               NA 
                                               azure4 
                                               1 
                                               -1.728 
                                             
                                             
                                               AXIN1-mRNA 
                                               -0.353 
                                               0.226 
                                               -0.797 
                                               0.0904 
                                               0.783 
                                               0.576 
                                               1.06 
                                               0.134 
                                               1 
                                               lm.nb 
                                               AXIN missense mutants destabilize the destruction complex, Beta-catenin phosphorylation cascade, Disassembly of the destruction complex and recruitment of AXIN to the membrane, Misspliced GSK3beta mutants stabilize beta-catenin, S33 mutants of beta-catenin aren.SQT.t phosphorylated, S37 mutants of beta-catenin aren.SQT.t phosphorylated, S45 mutants of beta-catenin aren.SQT.t phosphorylated, T41 mutants of beta-catenin aren.SQT.t phosphorylated, Truncations of AMER1 destabilize the destruction complex 
                                               SEBELIUS_2_C4636 
                                               AXIN1 
                                               mRNA 
                                               0 
                                               
                                               
                                               APC truncation mutants have impaired AXIN binding;Misspliced GSK3beta mutants stabilize beta-catenin;T41 mutants of beta-catenin aren.SQT.t phosphorylated;AXIN missense mutants destabilize the destruction complex;S45 mutants of beta-catenin aren.SQT.t phosphorylated;Disassembly of the destruction complex and recruitment of AXIN to the membrane;S37 mutants of beta-catenin aren.SQT.t phosphorylated;S33 mutants of beta-catenin aren.SQT.t phosphorylated;Beta-catenin phosphorylation cascade;Truncations of AMER1 destabilize the destruction complex 
                                               hsa04310;hsa04390;hsa04550;hsa05200;hsa05210;hsa05213;hsa05217 
                                               AXIN1 
                                               NA 
                                               azure4 
                                               1 
                                               -1.56194690265487 
                                             
                                             
                                               PPP2R1A-mRNA 
                                               0.122 
                                               0.105 
                                               -0.0832 
                                               0.327 
                                               1.09 
                                               0.944 
                                               1.25 
                                               0.257 
                                               1 
                                               loglinear 
                                               ARMS-mediated activation, AXIN missense mutants destabilize the destruction complex, Beta-catenin phosphorylation cascade, CTLA4 inhibitory signaling, Cyclin AB1 associated events during G2M transition, Cyclin D associated events in G1, DAP12 signaling, DARPP-32 events, Disassembly of the destruction complex and recruitment of AXIN to the membrane, Downstream signal transduction, FCERI mediated MAPK activation, Frs2-mediated activation, Gastrin-CREB signalling pathway via PKC and MAPK, Interleukin receptor SHC signaling, IRS-related events triggered by IGF1R, MASTL Facilitates Mitotic Progression, Negative regulation of FGFR1 signaling, Negative regulation of FGFR2 signaling, Negative regulation of FGFR3 signaling, Negative regulation of FGFR4 signaling, Nonsense Mediated Decay (NMD) enhanced by the Exon Junction Complex (EJC), RAF activation, RAFMAP kinase cascade, Resolution of Sister Chromatid Cohesion, RHO GTPases Activate Formins, S33 mutants of beta-catenin aren.SQT.t phosphorylated, S37 mutants of beta-catenin aren.SQT.t phosphorylated, S45 mutants of beta-catenin aren.SQT.t phosphorylated, Signaling by ERBB2, Signaling by PDGF, Signaling by SCF-KIT, Spry regulation of FGF signaling, Truncations of AMER1 destabilize the destruction complex, VEGFA-VEGFR2 Pathway, VEGFR2 mediated cell proliferation 
                                               SEBELIUS_2_C4636 
                                               PPP2R1A 
                                               mRNA 
                                               0 
                                               
                                               
                                               MASTL Facilitates Mitotic Progression;Regulation of PLK1 Activity at G2M Transition;Disassembly of the destruction complex and recruitment of AXIN to the membrane;S37 mutants of beta-catenin aren.SQT.t phosphorylated;Cyclin AB1 associated events during G2M transition;RAF activation;RHO GTPases Activate Formins;S33 mutants of beta-catenin aren.SQT.t phosphorylated;Loss of Nlp from mitotic centrosomes;DARPP-32 events;Beta-catenin phosphorylation cascade;Initiation of Nuclear Envelope Reformation;Recruitment of mitotic centrosome proteins and complexes;Spry regulation of FGF signaling;Truncations of AMER1 destabilize the destruction complex;CTLA4 inhibitory signaling;AXIN missense mutants destabilize the destruction complex;S45 mutants of beta-catenin aren.SQT.t phosphorylated;Anchoring of the basal body to the plasma membrane;Nonsense Mediated Decay (NMD) enhanced by the Exon Junction Complex (EJC);Cyclin D associated events in G1;Platelet sensitization by LDL;Resolution of Sister Chromatid Cohesion;Inhibition of replication initiation of damaged DNA by RB1E2F1;RAFMAP kinase cascade;Downstream signal transduction;VEGFR2 mediated cell proliferation;Signaling by SCF-KIT;Frs2-mediated activation;ARMS-mediated activation;Gastrin-CREB signalling pathway via PKC and MAPK;Interleukin receptor SHC signaling;FCERI mediated MAPK activation;DAP12 signaling;Signaling by PDGF;VEGFA-VEGFR2 Pathway;Signaling by ERBB2;IRS-related events triggered by IGF1R;Negative regulation of FGFR3 signaling;Negative regulation of FGFR4 signaling;Negative regulation of FGFR2 signaling;Negative regulation of FGFR1 signaling 
                                               hsa03015;hsa04071;hsa04114;hsa04151;hsa04152;hsa04261;hsa04350;hsa04390;hsa04530;hsa04728;hsa04730;hsa05142;hsa05160 
                                               PPP2R1A 
                                               NA 
                                               azure4 
                                               1 
                                               1.16190476190476 
                                             
                                           
                                           DE Results - Beta-catenin phosphorylation cascade genes - New.Annotation: miR155 vs.cel67  Table displaying each sample's global significance scores and directed global significance scores as defined in the heatmaps above. The global significance score is calculated as the square root of the mean squared t-statistic for the genes in a gene set, with t-statistics coming from the linear regression underlying our differential expression analysis. The directed global significance score is calculated as the square root of the mean signed squared t-statistic for the genes in a gene set, with t-statistics coming from the linear regression underlying our differential expression analysis.   
                                      
                                       
                                  
                                
                                
                                  
                                       
                                      
                                           Volcano Plot: New.Annotation: miR18b vs.cel67   More Plot Information                Volcano Plot: New.Annotation: miR18b vs.cel67  Volcano plot displaying each gene's -log10(p-value) and log2 fold change for the selected covariate. Highly statistically significant genes fall at the top of the plot, and highly differentially expressed genes fall to either side. Genes within the selected gene set are highlighted in orange. Horizontal lines indicate various False Discovery Rate (FDR) thresholds.   
                                           DE Results - Beta-catenin phosphorylation cascade genes - New.Annotation: miR18b vs.cel67   More Plot Information      Download CSV Data     
                                           
                                             
                                             
                                           
                                           
                                             
                                               Probe Label 
                                               Log2 fold change 
                                               std error (log2) 
                                               Lower confidence limit (log2) 
                                               Upper confidence limit (log2) 
                                               Linear fold change 
                                               Lower confidence limit (linear) 
                                               Upper confidence limit (linear) 
                                               P-value 
                                               BY.p.value 
                                               method 
                                               Gene.sets 
                                               Codeset.Name 
                                               Probe.Label 
                                               Analyte.Type 
                                               Is.Control 
                                               Control.Type 
                                               Related.Probes 
                                               Probe.Annotation 
                                               KEGG.Pathways 
                                               Official.Gene.Name 
                                               Control.Conc 
                                               volcanocol 
                                               volcanopch 
                                               tstats.all 
                                             
                                           
                                           
                                             
                                               CSNK1A1-mRNA 
                                               0.584 
                                               0.0761 
                                               0.435 
                                               0.733 
                                               1.5 
                                               1.35 
                                               1.66 
                                               1.6e-07 
                                               2.36e-05 
                                               loglinear 
                                               AXIN missense mutants destabilize the destruction complex, Beta-catenin phosphorylation cascade, Disassembly of the destruction complex and recruitment of AXIN to the membrane, Misspliced GSK3beta mutants stabilize beta-catenin, S33 mutants of beta-catenin aren.SQT.t phosphorylated, S37 mutants of beta-catenin aren.SQT.t phosphorylated, S45 mutants of beta-catenin aren.SQT.t phosphorylated, T41 mutants of beta-catenin aren.SQT.t phosphorylated 
                                               SEBELIUS_2_C4636 
                                               CSNK1A1 
                                               mRNA 
                                               0 
                                               
                                               
                                               Misspliced GSK3beta mutants stabilize beta-catenin;APC truncation mutants have impaired AXIN binding;T41 mutants of beta-catenin aren.SQT.t phosphorylated;AXIN missense mutants destabilize the destruction complex;S45 mutants of beta-catenin aren.SQT.t phosphorylated;Disassembly of the destruction complex and recruitment of AXIN to the membrane;S37 mutants of beta-catenin aren.SQT.t phosphorylated;S33 mutants of beta-catenin aren.SQT.t phosphorylated;Beta-catenin phosphorylation cascade;Degradation of GLI2 by the proteasome 
                                               hsa04310 
                                               CSNK1A1 
                                               NA 
                                               slateblue1 
                                               16 
                                               7.67411300919842 
                                             
                                             
                                               PPP2R1B-mRNA 
                                               0.385 
                                               0.0675 
                                               0.253 
                                               0.518 
                                               1.31 
                                               1.19 
                                               1.43 
                                               1.15e-05 
                                               0.000638 
                                               loglinear 
                                               ARMS-mediated activation, AXIN missense mutants destabilize the destruction complex, Beta-catenin phosphorylation cascade, CTLA4 inhibitory signaling, Cyclin AB1 associated events during G2M transition, Cyclin D associated events in G1, DAP12 signaling, DARPP-32 events, Disassembly of the destruction complex and recruitment of AXIN to the membrane, Downstream signal transduction, ERKs are inactivated, FCERI mediated MAPK activation, Frs2-mediated activation, Gastrin-CREB signalling pathway via PKC and MAPK, Interleukin receptor SHC signaling, IRS-related events triggered by IGF1R, MASTL Facilitates Mitotic Progression, Misspliced GSK3beta mutants stabilize beta-catenin, Negative regulation of MAPK pathway, RAF activation, RAFMAP kinase cascade, Resolution of Sister Chromatid Cohesion, RHO GTPases Activate Formins, S33 mutants of beta-catenin aren.SQT.t phosphorylated, S37 mutants of beta-catenin aren.SQT.t phosphorylated, S45 mutants of beta-catenin aren.SQT.t phosphorylated, Signaling by ERBB2, Signaling by PDGF, Signaling by SCF-KIT, T41 mutants of beta-catenin aren.SQT.t phosphorylated, Truncations of AMER1 destabilize the destruction complex, VEGFA-VEGFR2 Pathway, VEGFR2 mediated cell proliferation 
                                               SEBELIUS_2_C4636 
                                               PPP2R1B 
                                               mRNA 
                                               0 
                                               
                                               
                                               MASTL Facilitates Mitotic Progression;Misspliced GSK3beta mutants stabilize beta-catenin;T41 mutants of beta-catenin aren.SQT.t phosphorylated;ERKs are inactivated;Glycolysis;Negative regulation of MAPK pathway;Disassembly of the destruction complex and recruitment of AXIN to the membrane;S37 mutants of beta-catenin aren.SQT.t phosphorylated;Cyclin AB1 associated events during G2M transition;RAF activation;RHO GTPases Activate Formins;S33 mutants of beta-catenin aren.SQT.t phosphorylated;DARPP-32 events;Beta-catenin phosphorylation cascade;Truncations of AMER1 destabilize the destruction complex;CTLA4 inhibitory signaling;AXIN missense mutants destabilize the destruction complex;S45 mutants of beta-catenin aren.SQT.t phosphorylated;Cyclin D associated events in G1;Platelet sensitization by LDL;Resolution of Sister Chromatid Cohesion;Inhibition of replication initiation of damaged DNA by RB1E2F1;RAFMAP kinase cascade;Downstream signal transduction;VEGFR2 mediated cell proliferation;Signaling by SCF-KIT;Frs2-mediated activation;ARMS-mediated activation;Gastrin-CREB signalling pathway via PKC and MAPK;Interleukin receptor SHC signaling;FCERI mediated MAPK activation;DAP12 signaling;Signaling by PDGF;VEGFA-VEGFR2 Pathway;Signaling by ERBB2;IRS-related events triggered by IGF1R 
                                               hsa03015;hsa04071;hsa04114;hsa04151;hsa04152;hsa04261;hsa04350;hsa04390;hsa04530;hsa04728;hsa04730;hsa05142;hsa05160 
                                               PPP2R1B 
                                               NA 
                                               slateblue1 
                                               16 
                                               5.7037037037037 
                                             
                                             
                                               CTNNB1-mRNA 
                                               0.234 
                                               0.0637 
                                               0.109 
                                               0.359 
                                               1.18 
                                               1.08 
                                               1.28 
                                               0.0014 
                                               0.0333 
                                               loglinear 
                                               Apoptotic cleavage of cellular proteins, Beta-catenin phosphorylation cascade, Deactivation of the beta-catenin transactivating complex, Disassembly of the destruction complex and recruitment of AXIN to the membrane, Formation of the beta-catenin.CL.TCF transactivating complex, Misspliced GSK3beta mutants stabilize beta-catenin, Repression of WNT target genes, RHO GTPases activate IQGAPs, S37 mutants of beta-catenin aren.SQT.t phosphorylated, S45 mutants of beta-catenin aren.SQT.t phosphorylated, T41 mutants of beta-catenin aren.SQT.t phosphorylated, VEGFA-VEGFR2 Pathway, VEGFR2 mediated vascular permeability 
                                               SEBELIUS_2_C4636 
                                               CTNNB1 
                                               mRNA 
                                               0 
                                               
                                               
                                               Disassembly of the destruction complex and recruitment of AXIN to the membrane;S37 mutants of beta-catenin aren.SQT.t phosphorylated;T41 mutants of beta-catenin aren.SQT.t phosphorylated;S45 mutants of beta-catenin aren.SQT.t phosphorylated;Binding of TCFLEF.CL.CTNNB1 to target gene promoters;RHO GTPases activate IQGAPs;Deactivation of the beta-catenin transactivating complex;Repression of WNT target genes;Apoptotic cleavage of cell adhesion  proteins;Adherens junctions interactions;Misspliced GSK3beta mutants stabilize beta-catenin;VEGFR2 mediated vascular permeability;Beta-catenin phosphorylation cascade;Synthesis# secretion# and inactivation of Glucagon-like Peptide-1 (GLP-1);Formation of the beta-catenin.CL.TCF transactivating complex;Apoptotic cleavage of cellular proteins;VEGFA-VEGFR2 Pathway 
                                               hsa04015;hsa04310;hsa04390;hsa04510;hsa04520;hsa04530;hsa04550;hsa04670;hsa04916;hsa04919;hsa05100;hsa05130;hsa05166;hsa05200;hsa05205;hsa05210;hsa05213;hsa05215;hsa05216;hsa05217;hsa05412 
                                               CTNNB1 
                                               NA 
                                               slateblue3 
                                               16 
                                               3.6734693877551 
                                             
                                             
                                               PPP2CB-mRNA 
                                               0.253 
                                               0.085 
                                               0.0862 
                                               0.419 
                                               1.19 
                                               1.06 
                                               1.34 
                                               0.00724 
                                               0.148 
                                               loglinear 
                                               ARMS-mediated activation, AXIN missense mutants destabilize the destruction complex, Beta-catenin phosphorylation cascade, CTLA4 inhibitory signaling, Cyclin AB1 associated events during G2M transition, Cyclin D associated events in G1, DAP12 signaling, DARPP-32 events, Disassembly of the destruction complex and recruitment of AXIN to the membrane, Downstream signal transduction, ERKs are inactivated, FCERI mediated MAPK activation, Frs2-mediated activation, Gastrin-CREB signalling pathway via PKC and MAPK, Interleukin receptor SHC signaling, IRS-related events triggered by IGF1R, MASTL Facilitates Mitotic Progression, Misspliced GSK3beta mutants stabilize beta-catenin, Negative regulation of FGFR1 signaling, Negative regulation of FGFR2 signaling, Negative regulation of FGFR3 signaling, Negative regulation of FGFR4 signaling, Negative regulation of MAPK pathway, RAF activation, RAFMAP kinase cascade, Resolution of Sister Chromatid Cohesion, RHO GTPases Activate Formins, S33 mutants of beta-catenin aren.SQT.t phosphorylated, S37 mutants of beta-catenin aren.SQT.t phosphorylated, S45 mutants of beta-catenin aren.SQT.t phosphorylated, Signaling by ERBB2, Signaling by PDGF, Signaling by SCF-KIT, Spry regulation of FGF signaling, T41 mutants of beta-catenin aren.SQT.t phosphorylated, Truncations of AMER1 destabilize the destruction complex, VEGFA-VEGFR2 Pathway, VEGFR2 mediated cell proliferation 
                                               SEBELIUS_2_C4636 
                                               PPP2CB 
                                               mRNA 
                                               0 
                                               
                                               
                                               MASTL Facilitates Mitotic Progression;Misspliced GSK3beta mutants stabilize beta-catenin;T41 mutants of beta-catenin aren.SQT.t phosphorylated;ERKs are inactivated;Glycolysis;Negative regulation of MAPK pathway;Disassembly of the destruction complex and recruitment of AXIN to the membrane;S37 mutants of beta-catenin aren.SQT.t phosphorylated;Cyclin AB1 associated events during G2M transition;RAF activation;RHO GTPases Activate Formins;S33 mutants of beta-catenin aren.SQT.t phosphorylated;DARPP-32 events;Beta-catenin phosphorylation cascade;Spry regulation of FGF signaling;Truncations of AMER1 destabilize the destruction complex;CTLA4 inhibitory signaling;AXIN missense mutants destabilize the destruction complex;S45 mutants of beta-catenin aren.SQT.t phosphorylated;Cyclin D associated events in G1;Platelet sensitization by LDL;Resolution of Sister Chromatid Cohesion;Inhibition of replication initiation of damaged DNA by RB1E2F1;RAFMAP kinase cascade;Downstream signal transduction;VEGFR2 mediated cell proliferation;Signaling by SCF-KIT;Frs2-mediated activation;ARMS-mediated activation;Gastrin-CREB signalling pathway via PKC and MAPK;Interleukin receptor SHC signaling;FCERI mediated MAPK activation;DAP12 signaling;Signaling by PDGF;VEGFA-VEGFR2 Pathway;Signaling by ERBB2;IRS-related events triggered by IGF1R;Negative regulation of FGFR3 signaling;Negative regulation of FGFR4 signaling;Negative regulation of FGFR2 signaling;Negative regulation of FGFR1 signaling 
                                               hsa03015;hsa04071;hsa04114;hsa04151;hsa04152;hsa04261;hsa04350;hsa04390;hsa04530;hsa04728;hsa04730;hsa05142;hsa05160 
                                               PPP2CB 
                                               NA 
                                               slateblue4 
                                               16 
                                               2.97647058823529 
                                             
                                             
                                               PPP2CA-mRNA 
                                               -0.173 
                                               0.06 
                                               -0.29 
                                               -0.0549 
                                               0.887 
                                               0.818 
                                               0.963 
                                               0.00906 
                                               0.174 
                                               loglinear 
                                               ARMS-mediated activation, AXIN missense mutants destabilize the destruction complex, Beta-catenin phosphorylation cascade, CTLA4 inhibitory signaling, Cyclin AB1 associated events during G2M transition, Cyclin D associated events in G1, DAP12 signaling, DARPP-32 events, Disassembly of the destruction complex and recruitment of AXIN to the membrane, Downstream signal transduction, ERKs are inactivated, FCERI mediated MAPK activation, Frs2-mediated activation, Gastrin-CREB signalling pathway via PKC and MAPK, Interleukin receptor SHC signaling, IRS-related events triggered by IGF1R, MASTL Facilitates Mitotic Progression, Negative regulation of FGFR1 signaling, Negative regulation of FGFR2 signaling, Negative regulation of FGFR3 signaling, Negative regulation of FGFR4 signaling, Negative regulation of MAPK pathway, Nonsense Mediated Decay (NMD) enhanced by the Exon Junction Complex (EJC), RAF activation, RAFMAP kinase cascade, Resolution of Sister Chromatid Cohesion, RHO GTPases Activate Formins, S33 mutants of beta-catenin aren.SQT.t phosphorylated, S37 mutants of beta-catenin aren.SQT.t phosphorylated, S45 mutants of beta-catenin aren.SQT.t phosphorylated, Signaling by ERBB2, Signaling by PDGF, Signaling by SCF-KIT, Spry regulation of FGF signaling, T41 mutants of beta-catenin aren.SQT.t phosphorylated, Truncations of AMER1 destabilize the destruction complex, VEGFA-VEGFR2 Pathway, VEGFR2 mediated cell proliferation 
                                               SEBELIUS_2_C4636 
                                               PPP2CA 
                                               mRNA 
                                               0 
                                               
                                               
                                               MASTL Facilitates Mitotic Progression;T41 mutants of beta-catenin aren.SQT.t phosphorylated;ERKs are inactivated;Glycolysis;Negative regulation of MAPK pathway;Disassembly of the destruction complex and recruitment of AXIN to the membrane;S37 mutants of beta-catenin aren.SQT.t phosphorylated;Cyclin AB1 associated events during G2M transition;RAF activation;RHO GTPases Activate Formins;S33 mutants of beta-catenin aren.SQT.t phosphorylated;DARPP-32 events;Beta-catenin phosphorylation cascade;Initiation of Nuclear Envelope Reformation;Spry regulation of FGF signaling;Truncations of AMER1 destabilize the destruction complex;CTLA4 inhibitory signaling;AXIN missense mutants destabilize the destruction complex;S45 mutants of beta-catenin aren.SQT.t phosphorylated;Nonsense Mediated Decay (NMD) enhanced by the Exon Junction Complex (EJC);Cyclin D associated events in G1;Platelet sensitization by LDL;Resolution of Sister Chromatid Cohesion;Inhibition of replication initiation of damaged DNA by RB1E2F1;RAFMAP kinase cascade;Downstream signal transduction;VEGFR2 mediated cell proliferation;Signaling by SCF-KIT;Frs2-mediated activation;ARMS-mediated activation;Gastrin-CREB signalling pathway via PKC and MAPK;Interleukin receptor SHC signaling;FCERI mediated MAPK activation;DAP12 signaling;Signaling by PDGF;VEGFA-VEGFR2 Pathway;Signaling by ERBB2;IRS-related events triggered by IGF1R;Negative regulation of FGFR3 signaling;Negative regulation of FGFR4 signaling;Negative regulation of FGFR2 signaling;Negative regulation of FGFR1 signaling 
                                               hsa03015;hsa04071;hsa04114;hsa04151;hsa04152;hsa04261;hsa04350;hsa04390;hsa04530;hsa04728;hsa04730;hsa05142;hsa05160 
                                               PPP2CA 
                                               NA 
                                               slateblue4 
                                               16 
                                               -2.88333333333333 
                                             
                                             
                                               PPP2R1A-mRNA 
                                               -0.206 
                                               0.113 
                                               -0.428 
                                               0.0153 
                                               0.867 
                                               0.743 
                                               1.01 
                                               0.0824 
                                               1 
                                               loglinear 
                                               ARMS-mediated activation, AXIN missense mutants destabilize the destruction complex, Beta-catenin phosphorylation cascade, CTLA4 inhibitory signaling, Cyclin AB1 associated events during G2M transition, Cyclin D associated events in G1, DAP12 signaling, DARPP-32 events, Disassembly of the destruction complex and recruitment of AXIN to the membrane, Downstream signal transduction, FCERI mediated MAPK activation, Frs2-mediated activation, Gastrin-CREB signalling pathway via PKC and MAPK, Interleukin receptor SHC signaling, IRS-related events triggered by IGF1R, MASTL Facilitates Mitotic Progression, Negative regulation of FGFR1 signaling, Negative regulation of FGFR2 signaling, Negative regulation of FGFR3 signaling, Negative regulation of FGFR4 signaling, Nonsense Mediated Decay (NMD) enhanced by the Exon Junction Complex (EJC), RAF activation, RAFMAP kinase cascade, Resolution of Sister Chromatid Cohesion, RHO GTPases Activate Formins, S33 mutants of beta-catenin aren.SQT.t phosphorylated, S37 mutants of beta-catenin aren.SQT.t phosphorylated, S45 mutants of beta-catenin aren.SQT.t phosphorylated, Signaling by ERBB2, Signaling by PDGF, Signaling by SCF-KIT, Spry regulation of FGF signaling, Truncations of AMER1 destabilize the destruction complex, VEGFA-VEGFR2 Pathway, VEGFR2 mediated cell proliferation 
                                               SEBELIUS_2_C4636 
                                               PPP2R1A 
                                               mRNA 
                                               0 
                                               
                                               
                                               MASTL Facilitates Mitotic Progression;Regulation of PLK1 Activity at G2M Transition;Disassembly of the destruction complex and recruitment of AXIN to the membrane;S37 mutants of beta-catenin aren.SQT.t phosphorylated;Cyclin AB1 associated events during G2M transition;RAF activation;RHO GTPases Activate Formins;S33 mutants of beta-catenin aren.SQT.t phosphorylated;Loss of Nlp from mitotic centrosomes;DARPP-32 events;Beta-catenin phosphorylation cascade;Initiation of Nuclear Envelope Reformation;Recruitment of mitotic centrosome proteins and complexes;Spry regulation of FGF signaling;Truncations of AMER1 destabilize the destruction complex;CTLA4 inhibitory signaling;AXIN missense mutants destabilize the destruction complex;S45 mutants of beta-catenin aren.SQT.t phosphorylated;Anchoring of the basal body to the plasma membrane;Nonsense Mediated Decay (NMD) enhanced by the Exon Junction Complex (EJC);Cyclin D associated events in G1;Platelet sensitization by LDL;Resolution of Sister Chromatid Cohesion;Inhibition of replication initiation of damaged DNA by RB1E2F1;RAFMAP kinase cascade;Downstream signal transduction;VEGFR2 mediated cell proliferation;Signaling by SCF-KIT;Frs2-mediated activation;ARMS-mediated activation;Gastrin-CREB signalling pathway via PKC and MAPK;Interleukin receptor SHC signaling;FCERI mediated MAPK activation;DAP12 signaling;Signaling by PDGF;VEGFA-VEGFR2 Pathway;Signaling by ERBB2;IRS-related events triggered by IGF1R;Negative regulation of FGFR3 signaling;Negative regulation of FGFR4 signaling;Negative regulation of FGFR2 signaling;Negative regulation of FGFR1 signaling 
                                               hsa03015;hsa04071;hsa04114;hsa04151;hsa04152;hsa04261;hsa04350;hsa04390;hsa04530;hsa04728;hsa04730;hsa05142;hsa05160 
                                               PPP2R1A 
                                               NA 
                                               azure4 
                                               1 
                                               -1.82300884955752 
                                             
                                             
                                               AXIN1-mRNA 
                                               -0.362 
                                               0.246 
                                               -0.843 
                                               0.12 
                                               0.778 
                                               0.558 
                                               1.09 
                                               0.156 
                                               1 
                                               lm.nb 
                                               AXIN missense mutants destabilize the destruction complex, Beta-catenin phosphorylation cascade, Disassembly of the destruction complex and recruitment of AXIN to the membrane, Misspliced GSK3beta mutants stabilize beta-catenin, S33 mutants of beta-catenin aren.SQT.t phosphorylated, S37 mutants of beta-catenin aren.SQT.t phosphorylated, S45 mutants of beta-catenin aren.SQT.t phosphorylated, T41 mutants of beta-catenin aren.SQT.t phosphorylated, Truncations of AMER1 destabilize the destruction complex 
                                               SEBELIUS_2_C4636 
                                               AXIN1 
                                               mRNA 
                                               0 
                                               
                                               
                                               APC truncation mutants have impaired AXIN binding;Misspliced GSK3beta mutants stabilize beta-catenin;T41 mutants of beta-catenin aren.SQT.t phosphorylated;AXIN missense mutants destabilize the destruction complex;S45 mutants of beta-catenin aren.SQT.t phosphorylated;Disassembly of the destruction complex and recruitment of AXIN to the membrane;S37 mutants of beta-catenin aren.SQT.t phosphorylated;S33 mutants of beta-catenin aren.SQT.t phosphorylated;Beta-catenin phosphorylation cascade;Truncations of AMER1 destabilize the destruction complex 
                                               hsa04310;hsa04390;hsa04550;hsa05200;hsa05210;hsa05213;hsa05217 
                                               AXIN1 
                                               NA 
                                               azure4 
                                               1 
                                               -1.47154471544715 
                                             
                                             
                                               APC-mRNA 
                                               -0.0927 
                                               0.109 
                                               -0.306 
                                               0.121 
                                               0.938 
                                               0.809 
                                               1.09 
                                               0.404 
                                               1 
                                               loglinear 
                                               Apoptotic cleavage of cellular proteins, AXIN missense mutants destabilize the destruction complex, Beta-catenin phosphorylation cascade, Deactivation of the beta-catenin transactivating complex, Disassembly of the destruction complex and recruitment of AXIN to the membrane, Misspliced GSK3beta mutants stabilize beta-catenin, S33 mutants of beta-catenin aren.SQT.t phosphorylated, S37 mutants of beta-catenin aren.SQT.t phosphorylated, S45 mutants of beta-catenin aren.SQT.t phosphorylated, T41 mutants of beta-catenin aren.SQT.t phosphorylated, Truncations of AMER1 destabilize the destruction complex 
                                               SEBELIUS_2_C4636 
                                               APC 
                                               mRNA 
                                               0 
                                               
                                               
                                               Apoptotic cleavage of cellular proteins;Misspliced GSK3beta mutants stabilize beta-catenin;APC truncation mutants have impaired AXIN binding;T41 mutants of beta-catenin aren.SQT.t phosphorylated;AXIN missense mutants destabilize the destruction complex;S45 mutants of beta-catenin aren.SQT.t phosphorylated;Deactivation of the beta-catenin transactivating complex;Disassembly of the destruction complex and recruitment of AXIN to the membrane;S37 mutants of beta-catenin aren.SQT.t phosphorylated;S33 mutants of beta-catenin aren.SQT.t phosphorylated;Beta-catenin phosphorylation cascade;Truncations of AMER1 destabilize the destruction complex 
                                               hsa04310;hsa04390;hsa04550;hsa04810;hsa05166;hsa05200;hsa05206;hsa05210;hsa05213;hsa05217 
                                               APC 
                                               NA 
                                               azure4 
                                               1 
                                               -0.85045871559633 
                                             
                                           
                                           DE Results - Beta-catenin phosphorylation cascade genes - New.Annotation: miR18b vs.cel67  Table displaying each sample's global significance scores and directed global significance scores as defined in the heatmaps above. The global significance score is calculated as the square root of the mean squared t-statistic for the genes in a gene set, with t-statistics coming from the linear regression underlying our differential expression analysis. The directed global significance score is calculated as the square root of the mean signed squared t-statistic for the genes in a gene set, with t-statistics coming from the linear regression underlying our differential expression analysis.   
                                      
                                       
                                  
                                
                                
                                  
                                       
                                      
                                           Volcano Plot: New.Annotation: miR890 vs.cel67   More Plot Information                Volcano Plot: New.Annotation: miR890 vs.cel67  Volcano plot displaying each gene's -log10(p-value) and log2 fold change for the selected covariate. Highly statistically significant genes fall at the top of the plot, and highly differentially expressed genes fall to either side. Genes within the selected gene set are highlighted in orange. Horizontal lines indicate various False Discovery Rate (FDR) thresholds.   
                                           DE Results - Beta-catenin phosphorylation cascade genes - New.Annotation: miR890 vs.cel67   More Plot Information      Download CSV Data     
                                           
                                             
                                             
                                           
                                           
                                             
                                               Probe Label 
                                               Log2 fold change 
                                               std error (log2) 
                                               Lower confidence limit (log2) 
                                               Upper confidence limit (log2) 
                                               Linear fold change 
                                               Lower confidence limit (linear) 
                                               Upper confidence limit (linear) 
                                               P-value 
                                               BY.p.value 
                                               method 
                                               Gene.sets 
                                               Codeset.Name 
                                               Probe.Label 
                                               Analyte.Type 
                                               Is.Control 
                                               Control.Type 
                                               Related.Probes 
                                               Probe.Annotation 
                                               KEGG.Pathways 
                                               Official.Gene.Name 
                                               Control.Conc 
                                               volcanocol 
                                               volcanopch 
                                               tstats.all 
                                             
                                           
                                           
                                             
                                               APC-mRNA 
                                               -0.688 
                                               0.0921 
                                               -0.869 
                                               -0.508 
                                               0.621 
                                               0.548 
                                               0.703 
                                               2.41e-07 
                                               2.92e-05 
                                               loglinear 
                                               Apoptotic cleavage of cellular proteins, AXIN missense mutants destabilize the destruction complex, Beta-catenin phosphorylation cascade, Deactivation of the beta-catenin transactivating complex, Disassembly of the destruction complex and recruitment of AXIN to the membrane, Misspliced GSK3beta mutants stabilize beta-catenin, S33 mutants of beta-catenin aren.SQT.t phosphorylated, S37 mutants of beta-catenin aren.SQT.t phosphorylated, S45 mutants of beta-catenin aren.SQT.t phosphorylated, T41 mutants of beta-catenin aren.SQT.t phosphorylated, Truncations of AMER1 destabilize the destruction complex 
                                               SEBELIUS_2_C4636 
                                               APC 
                                               mRNA 
                                               0 
                                               
                                               
                                               Apoptotic cleavage of cellular proteins;Misspliced GSK3beta mutants stabilize beta-catenin;APC truncation mutants have impaired AXIN binding;T41 mutants of beta-catenin aren.SQT.t phosphorylated;AXIN missense mutants destabilize the destruction complex;S45 mutants of beta-catenin aren.SQT.t phosphorylated;Deactivation of the beta-catenin transactivating complex;Disassembly of the destruction complex and recruitment of AXIN to the membrane;S37 mutants of beta-catenin aren.SQT.t phosphorylated;S33 mutants of beta-catenin aren.SQT.t phosphorylated;Beta-catenin phosphorylation cascade;Truncations of AMER1 destabilize the destruction complex 
                                               hsa04310;hsa04390;hsa04550;hsa04810;hsa05166;hsa05200;hsa05206;hsa05210;hsa05213;hsa05217 
                                               APC 
                                               NA 
                                               slateblue1 
                                               16 
                                               -7.47014115092291 
                                             
                                             
                                               PPP2CB-mRNA 
                                               -0.402 
                                               0.0718 
                                               -0.542 
                                               -0.261 
                                               0.757 
                                               0.687 
                                               0.835 
                                               1.51e-05 
                                               0.000909 
                                               loglinear 
                                               ARMS-mediated activation, AXIN missense mutants destabilize the destruction complex, Beta-catenin phosphorylation cascade, CTLA4 inhibitory signaling, Cyclin AB1 associated events during G2M transition, Cyclin D associated events in G1, DAP12 signaling, DARPP-32 events, Disassembly of the destruction complex and recruitment of AXIN to the membrane, Downstream signal transduction, ERKs are inactivated, FCERI mediated MAPK activation, Frs2-mediated activation, Gastrin-CREB signalling pathway via PKC and MAPK, Interleukin receptor SHC signaling, IRS-related events triggered by IGF1R, MASTL Facilitates Mitotic Progression, Misspliced GSK3beta mutants stabilize beta-catenin, Negative regulation of FGFR1 signaling, Negative regulation of FGFR2 signaling, Negative regulation of FGFR3 signaling, Negative regulation of FGFR4 signaling, Negative regulation of MAPK pathway, RAF activation, RAFMAP kinase cascade, Resolution of Sister Chromatid Cohesion, RHO GTPases Activate Formins, S33 mutants of beta-catenin aren.SQT.t phosphorylated, S37 mutants of beta-catenin aren.SQT.t phosphorylated, S45 mutants of beta-catenin aren.SQT.t phosphorylated, Signaling by ERBB2, Signaling by PDGF, Signaling by SCF-KIT, Spry regulation of FGF signaling, T41 mutants of beta-catenin aren.SQT.t phosphorylated, Truncations of AMER1 destabilize the destruction complex, VEGFA-VEGFR2 Pathway, VEGFR2 mediated cell proliferation 
                                               SEBELIUS_2_C4636 
                                               PPP2CB 
                                               mRNA 
                                               0 
                                               
                                               
                                               MASTL Facilitates Mitotic Progression;Misspliced GSK3beta mutants stabilize beta-catenin;T41 mutants of beta-catenin aren.SQT.t phosphorylated;ERKs are inactivated;Glycolysis;Negative regulation of MAPK pathway;Disassembly of the destruction complex and recruitment of AXIN to the membrane;S37 mutants of beta-catenin aren.SQT.t phosphorylated;Cyclin AB1 associated events during G2M transition;RAF activation;RHO GTPases Activate Formins;S33 mutants of beta-catenin aren.SQT.t phosphorylated;DARPP-32 events;Beta-catenin phosphorylation cascade;Spry regulation of FGF signaling;Truncations of AMER1 destabilize the destruction complex;CTLA4 inhibitory signaling;AXIN missense mutants destabilize the destruction complex;S45 mutants of beta-catenin aren.SQT.t phosphorylated;Cyclin D associated events in G1;Platelet sensitization by LDL;Resolution of Sister Chromatid Cohesion;Inhibition of replication initiation of damaged DNA by RB1E2F1;RAFMAP kinase cascade;Downstream signal transduction;VEGFR2 mediated cell proliferation;Signaling by SCF-KIT;Frs2-mediated activation;ARMS-mediated activation;Gastrin-CREB signalling pathway via PKC and MAPK;Interleukin receptor SHC signaling;FCERI mediated MAPK activation;DAP12 signaling;Signaling by PDGF;VEGFA-VEGFR2 Pathway;Signaling by ERBB2;IRS-related events triggered by IGF1R;Negative regulation of FGFR3 signaling;Negative regulation of FGFR4 signaling;Negative regulation of FGFR2 signaling;Negative regulation of FGFR1 signaling 
                                               hsa03015;hsa04071;hsa04114;hsa04151;hsa04152;hsa04261;hsa04350;hsa04390;hsa04530;hsa04728;hsa04730;hsa05142;hsa05160 
                                               PPP2CB 
                                               NA 
                                               slateblue1 
                                               16 
                                               -5.59888579387187 
                                             
                                             
                                               CSNK1A1-mRNA 
                                               0.245 
                                               0.0643 
                                               0.119 
                                               0.371 
                                               1.18 
                                               1.09 
                                               1.29 
                                               0.00104 
                                               0.0276 
                                               loglinear 
                                               AXIN missense mutants destabilize the destruction complex, Beta-catenin phosphorylation cascade, Disassembly of the destruction complex and recruitment of AXIN to the membrane, Misspliced GSK3beta mutants stabilize beta-catenin, S33 mutants of beta-catenin aren.SQT.t phosphorylated, S37 mutants of beta-catenin aren.SQT.t phosphorylated, S45 mutants of beta-catenin aren.SQT.t phosphorylated, T41 mutants of beta-catenin aren.SQT.t phosphorylated 
                                               SEBELIUS_2_C4636 
                                               CSNK1A1 
                                               mRNA 
                                               0 
                                               
                                               
                                               Misspliced GSK3beta mutants stabilize beta-catenin;APC truncation mutants have impaired AXIN binding;T41 mutants of beta-catenin aren.SQT.t phosphorylated;AXIN missense mutants destabilize the destruction complex;S45 mutants of beta-catenin aren.SQT.t phosphorylated;Disassembly of the destruction complex and recruitment of AXIN to the membrane;S37 mutants of beta-catenin aren.SQT.t phosphorylated;S33 mutants of beta-catenin aren.SQT.t phosphorylated;Beta-catenin phosphorylation cascade;Degradation of GLI2 by the proteasome 
                                               hsa04310 
                                               CSNK1A1 
                                               NA 
                                               slateblue2 
                                               16 
                                               3.81026438569207 
                                             
                                             
                                               PPP2R1B-mRNA 
                                               -0.142 
                                               0.0571 
                                               -0.254 
                                               -0.0303 
                                               0.906 
                                               0.839 
                                               0.979 
                                               0.0212 
                                               0.348 
                                               loglinear 
                                               ARMS-mediated activation, AXIN missense mutants destabilize the destruction complex, Beta-catenin phosphorylation cascade, CTLA4 inhibitory signaling, Cyclin AB1 associated events during G2M transition, Cyclin D associated events in G1, DAP12 signaling, DARPP-32 events, Disassembly of the destruction complex and recruitment of AXIN to the membrane, Downstream signal transduction, ERKs are inactivated, FCERI mediated MAPK activation, Frs2-mediated activation, Gastrin-CREB signalling pathway via PKC and MAPK, Interleukin receptor SHC signaling, IRS-related events triggered by IGF1R, MASTL Facilitates Mitotic Progression, Misspliced GSK3beta mutants stabilize beta-catenin, Negative regulation of MAPK pathway, RAF activation, RAFMAP kinase cascade, Resolution of Sister Chromatid Cohesion, RHO GTPases Activate Formins, S33 mutants of beta-catenin aren.SQT.t phosphorylated, S37 mutants of beta-catenin aren.SQT.t phosphorylated, S45 mutants of beta-catenin aren.SQT.t phosphorylated, Signaling by ERBB2, Signaling by PDGF, Signaling by SCF-KIT, T41 mutants of beta-catenin aren.SQT.t phosphorylated, Truncations of AMER1 destabilize the destruction complex, VEGFA-VEGFR2 Pathway, VEGFR2 mediated cell proliferation 
                                               SEBELIUS_2_C4636 
                                               PPP2R1B 
                                               mRNA 
                                               0 
                                               
                                               
                                               MASTL Facilitates Mitotic Progression;Misspliced GSK3beta mutants stabilize beta-catenin;T41 mutants of beta-catenin aren.SQT.t phosphorylated;ERKs are inactivated;Glycolysis;Negative regulation of MAPK pathway;Disassembly of the destruction complex and recruitment of AXIN to the membrane;S37 mutants of beta-catenin aren.SQT.t phosphorylated;Cyclin AB1 associated events during G2M transition;RAF activation;RHO GTPases Activate Formins;S33 mutants of beta-catenin aren.SQT.t phosphorylated;DARPP-32 events;Beta-catenin phosphorylation cascade;Truncations of AMER1 destabilize the destruction complex;CTLA4 inhibitory signaling;AXIN missense mutants destabilize the destruction complex;S45 mutants of beta-catenin aren.SQT.t phosphorylated;Cyclin D associated events in G1;Platelet sensitization by LDL;Resolution of Sister Chromatid Cohesion;Inhibition of replication initiation of damaged DNA by RB1E2F1;RAFMAP kinase cascade;Downstream signal transduction;VEGFR2 mediated cell proliferation;Signaling by SCF-KIT;Frs2-mediated activation;ARMS-mediated activation;Gastrin-CREB signalling pathway via PKC and MAPK;Interleukin receptor SHC signaling;FCERI mediated MAPK activation;DAP12 signaling;Signaling by PDGF;VEGFA-VEGFR2 Pathway;Signaling by ERBB2;IRS-related events triggered by IGF1R 
                                               hsa03015;hsa04071;hsa04114;hsa04151;hsa04152;hsa04261;hsa04350;hsa04390;hsa04530;hsa04728;hsa04730;hsa05142;hsa05160 
                                               PPP2R1B 
                                               NA 
                                               slateblue4 
                                               16 
                                               -2.48686514886165 
                                             
                                             
                                               AXIN1-mRNA 
                                               -0.356 
                                               0.206 
                                               -0.759 
                                               0.0469 
                                               0.781 
                                               0.591 
                                               1.03 
                                               0.098 
                                               1 
                                               lm.nb 
                                               AXIN missense mutants destabilize the destruction complex, Beta-catenin phosphorylation cascade, Disassembly of the destruction complex and recruitment of AXIN to the membrane, Misspliced GSK3beta mutants stabilize beta-catenin, S33 mutants of beta-catenin aren.SQT.t phosphorylated, S37 mutants of beta-catenin aren.SQT.t phosphorylated, S45 mutants of beta-catenin aren.SQT.t phosphorylated, T41 mutants of beta-catenin aren.SQT.t phosphorylated, Truncations of AMER1 destabilize the destruction complex 
                                               SEBELIUS_2_C4636 
                                               AXIN1 
                                               mRNA 
                                               0 
                                               
                                               
                                               APC truncation mutants have impaired AXIN binding;Misspliced GSK3beta mutants stabilize beta-catenin;T41 mutants of beta-catenin aren.SQT.t phosphorylated;AXIN missense mutants destabilize the destruction complex;S45 mutants of beta-catenin aren.SQT.t phosphorylated;Disassembly of the destruction complex and recruitment of AXIN to the membrane;S37 mutants of beta-catenin aren.SQT.t phosphorylated;S33 mutants of beta-catenin aren.SQT.t phosphorylated;Beta-catenin phosphorylation cascade;Truncations of AMER1 destabilize the destruction complex 
                                               hsa04310;hsa04390;hsa04550;hsa05200;hsa05210;hsa05213;hsa05217 
                                               AXIN1 
                                               NA 
                                               azure4 
                                               1 
                                               -1.72815533980583 
                                             
                                             
                                               CTNNB1-mRNA 
                                               0.0834 
                                               0.0538 
                                               -0.0222 
                                               0.189 
                                               1.06 
                                               0.985 
                                               1.14 
                                               0.136 
                                               1 
                                               loglinear 
                                               Apoptotic cleavage of cellular proteins, Beta-catenin phosphorylation cascade, Deactivation of the beta-catenin transactivating complex, Disassembly of the destruction complex and recruitment of AXIN to the membrane, Formation of the beta-catenin.CL.TCF transactivating complex, Misspliced GSK3beta mutants stabilize beta-catenin, Repression of WNT target genes, RHO GTPases activate IQGAPs, S37 mutants of beta-catenin aren.SQT.t phosphorylated, S45 mutants of beta-catenin aren.SQT.t phosphorylated, T41 mutants of beta-catenin aren.SQT.t phosphorylated, VEGFA-VEGFR2 Pathway, VEGFR2 mediated vascular permeability 
                                               SEBELIUS_2_C4636 
                                               CTNNB1 
                                               mRNA 
                                               0 
                                               
                                               
                                               Disassembly of the destruction complex and recruitment of AXIN to the membrane;S37 mutants of beta-catenin aren.SQT.t phosphorylated;T41 mutants of beta-catenin aren.SQT.t phosphorylated;S45 mutants of beta-catenin aren.SQT.t phosphorylated;Binding of TCFLEF.CL.CTNNB1 to target gene promoters;RHO GTPases activate IQGAPs;Deactivation of the beta-catenin transactivating complex;Repression of WNT target genes;Apoptotic cleavage of cell adhesion  proteins;Adherens junctions interactions;Misspliced GSK3beta mutants stabilize beta-catenin;VEGFR2 mediated vascular permeability;Beta-catenin phosphorylation cascade;Synthesis# secretion# and inactivation of Glucagon-like Peptide-1 (GLP-1);Formation of the beta-catenin.CL.TCF transactivating complex;Apoptotic cleavage of cellular proteins;VEGFA-VEGFR2 Pathway 
                                               hsa04015;hsa04310;hsa04390;hsa04510;hsa04520;hsa04530;hsa04550;hsa04670;hsa04916;hsa04919;hsa05100;hsa05130;hsa05166;hsa05200;hsa05205;hsa05210;hsa05213;hsa05215;hsa05216;hsa05217;hsa05412 
                                               CTNNB1 
                                               NA 
                                               azure4 
                                               1 
                                               1.55018587360595 
                                             
                                             
                                               PPP2CA-mRNA 
                                               -0.0403 
                                               0.0507 
                                               -0.14 
                                               0.0591 
                                               0.972 
                                               0.908 
                                               1.04 
                                               0.436 
                                               1 
                                               loglinear 
                                               ARMS-mediated activation, AXIN missense mutants destabilize the destruction complex, Beta-catenin phosphorylation cascade, CTLA4 inhibitory signaling, Cyclin AB1 associated events during G2M transition, Cyclin D associated events in G1, DAP12 signaling, DARPP-32 events, Disassembly of the destruction complex and recruitment of AXIN to the membrane, Downstream signal transduction, ERKs are inactivated, FCERI mediated MAPK activation, Frs2-mediated activation, Gastrin-CREB signalling pathway via PKC and MAPK, Interleukin receptor SHC signaling, IRS-related events triggered by IGF1R, MASTL Facilitates Mitotic Progression, Negative regulation of FGFR1 signaling, Negative regulation of FGFR2 signaling, Negative regulation of FGFR3 signaling, Negative regulation of FGFR4 signaling, Negative regulation of MAPK pathway, Nonsense Mediated Decay (NMD) enhanced by the Exon Junction Complex (EJC), RAF activation, RAFMAP kinase cascade, Resolution of Sister Chromatid Cohesion, RHO GTPases Activate Formins, S33 mutants of beta-catenin aren.SQT.t phosphorylated, S37 mutants of beta-catenin aren.SQT.t phosphorylated, S45 mutants of beta-catenin aren.SQT.t phosphorylated, Signaling by ERBB2, Signaling by PDGF, Signaling by SCF-KIT, Spry regulation of FGF signaling, T41 mutants of beta-catenin aren.SQT.t phosphorylated, Truncations of AMER1 destabilize the destruction complex, VEGFA-VEGFR2 Pathway, VEGFR2 mediated cell proliferation 
                                               SEBELIUS_2_C4636 
                                               PPP2CA 
                                               mRNA 
                                               0 
                                               
                                               
                                               MASTL Facilitates Mitotic Progression;T41 mutants of beta-catenin aren.SQT.t phosphorylated;ERKs are inactivated;Glycolysis;Negative regulation of MAPK pathway;Disassembly of the destruction complex and recruitment of AXIN to the membrane;S37 mutants of beta-catenin aren.SQT.t phosphorylated;Cyclin AB1 associated events during G2M transition;RAF activation;RHO GTPases Activate Formins;S33 mutants of beta-catenin aren.SQT.t phosphorylated;DARPP-32 events;Beta-catenin phosphorylation cascade;Initiation of Nuclear Envelope Reformation;Spry regulation of FGF signaling;Truncations of AMER1 destabilize the destruction complex;CTLA4 inhibitory signaling;AXIN missense mutants destabilize the destruction complex;S45 mutants of beta-catenin aren.SQT.t phosphorylated;Nonsense Mediated Decay (NMD) enhanced by the Exon Junction Complex (EJC);Cyclin D associated events in G1;Platelet sensitization by LDL;Resolution of Sister Chromatid Cohesion;Inhibition of replication initiation of damaged DNA by RB1E2F1;RAFMAP kinase cascade;Downstream signal transduction;VEGFR2 mediated cell proliferation;Signaling by SCF-KIT;Frs2-mediated activation;ARMS-mediated activation;Gastrin-CREB signalling pathway via PKC and MAPK;Interleukin receptor SHC signaling;FCERI mediated MAPK activation;DAP12 signaling;Signaling by PDGF;VEGFA-VEGFR2 Pathway;Signaling by ERBB2;IRS-related events triggered by IGF1R;Negative regulation of FGFR3 signaling;Negative regulation of FGFR4 signaling;Negative regulation of FGFR2 signaling;Negative regulation of FGFR1 signaling 
                                               hsa03015;hsa04071;hsa04114;hsa04151;hsa04152;hsa04261;hsa04350;hsa04390;hsa04530;hsa04728;hsa04730;hsa05142;hsa05160 
                                               PPP2CA 
                                               NA 
                                               azure4 
                                               1 
                                               -0.794871794871795 
                                             
                                             
                                               PPP2R1A-mRNA 
                                               0.069 
                                               0.0955 
                                               -0.118 
                                               0.256 
                                               1.05 
                                               0.921 
                                               1.19 
                                               0.478 
                                               1 
                                               loglinear 
                                               ARMS-mediated activation, AXIN missense mutants destabilize the destruction complex, Beta-catenin phosphorylation cascade, CTLA4 inhibitory signaling, Cyclin AB1 associated events during G2M transition, Cyclin D associated events in G1, DAP12 signaling, DARPP-32 events, Disassembly of the destruction complex and recruitment of AXIN to the membrane, Downstream signal transduction, FCERI mediated MAPK activation, Frs2-mediated activation, Gastrin-CREB signalling pathway via PKC and MAPK, Interleukin receptor SHC signaling, IRS-related events triggered by IGF1R, MASTL Facilitates Mitotic Progression, Negative regulation of FGFR1 signaling, Negative regulation of FGFR2 signaling, Negative regulation of FGFR3 signaling, Negative regulation of FGFR4 signaling, Nonsense Mediated Decay (NMD) enhanced by the Exon Junction Complex (EJC), RAF activation, RAFMAP kinase cascade, Resolution of Sister Chromatid Cohesion, RHO GTPases Activate Formins, S33 mutants of beta-catenin aren.SQT.t phosphorylated, S37 mutants of beta-catenin aren.SQT.t phosphorylated, S45 mutants of beta-catenin aren.SQT.t phosphorylated, Signaling by ERBB2, Signaling by PDGF, Signaling by SCF-KIT, Spry regulation of FGF signaling, Truncations of AMER1 destabilize the destruction complex, VEGFA-VEGFR2 Pathway, VEGFR2 mediated cell proliferation 
                                               SEBELIUS_2_C4636 
                                               PPP2R1A 
                                               mRNA 
                                               0 
                                               
                                               
                                               MASTL Facilitates Mitotic Progression;Regulation of PLK1 Activity at G2M Transition;Disassembly of the destruction complex and recruitment of AXIN to the membrane;S37 mutants of beta-catenin aren.SQT.t phosphorylated;Cyclin AB1 associated events during G2M transition;RAF activation;RHO GTPases Activate Formins;S33 mutants of beta-catenin aren.SQT.t phosphorylated;Loss of Nlp from mitotic centrosomes;DARPP-32 events;Beta-catenin phosphorylation cascade;Initiation of Nuclear Envelope Reformation;Recruitment of mitotic centrosome proteins and complexes;Spry regulation of FGF signaling;Truncations of AMER1 destabilize the destruction complex;CTLA4 inhibitory signaling;AXIN missense mutants destabilize the destruction complex;S45 mutants of beta-catenin aren.SQT.t phosphorylated;Anchoring of the basal body to the plasma membrane;Nonsense Mediated Decay (NMD) enhanced by the Exon Junction Complex (EJC);Cyclin D associated events in G1;Platelet sensitization by LDL;Resolution of Sister Chromatid Cohesion;Inhibition of replication initiation of damaged DNA by RB1E2F1;RAFMAP kinase cascade;Downstream signal transduction;VEGFR2 mediated cell proliferation;Signaling by SCF-KIT;Frs2-mediated activation;ARMS-mediated activation;Gastrin-CREB signalling pathway via PKC and MAPK;Interleukin receptor SHC signaling;FCERI mediated MAPK activation;DAP12 signaling;Signaling by PDGF;VEGFA-VEGFR2 Pathway;Signaling by ERBB2;IRS-related events triggered by IGF1R;Negative regulation of FGFR3 signaling;Negative regulation of FGFR4 signaling;Negative regulation of FGFR2 signaling;Negative regulation of FGFR1 signaling 
                                               hsa03015;hsa04071;hsa04114;hsa04151;hsa04152;hsa04261;hsa04350;hsa04390;hsa04530;hsa04728;hsa04730;hsa05142;hsa05160 
                                               PPP2R1A 
                                               NA 
                                               azure4 
                                               1 
                                               0.722513089005236 
                                             
                                           
                                           DE Results - Beta-catenin phosphorylation cascade genes - New.Annotation: miR890 vs.cel67  Table displaying each sample's global significance scores and directed global significance scores as defined in the heatmaps above. The global significance score is calculated as the square root of the mean squared t-statistic for the genes in a gene set, with t-statistics coming from the linear regression underlying our differential expression analysis. The directed global significance score is calculated as the square root of the mean signed squared t-statistic for the genes in a gene set, with t-statistics coming from the linear regression underlying our differential expression analysis.   
                                      
                                       
                                  
                                
                              
                             
                          
                           
                      
                    
                    
                      
                           
                          
                              New.Annotation: 
                              
                                  New.Annotation: differential expression in Control vs. baseline of cel67  
                                  New.Annotation: differential expression in miR142 vs. baseline of cel67  
                                  New.Annotation: differential expression in miR155 vs. baseline of cel67  
                                  New.Annotation: differential expression in miR18b vs. baseline of cel67  
                                  New.Annotation: differential expression in miR890 vs. baseline of cel67  
                              
                              
                              
                                
                                  
                                       
                                      
                                           Volcano Plot: New.Annotation: Control vs.cel67   More Plot Information                Volcano Plot: New.Annotation: Control vs.cel67  Volcano plot displaying each gene's -log10(p-value) and log2 fold change for the selected covariate. Highly statistically significant genes fall at the top of the plot, and highly differentially expressed genes fall to either side. Genes within the selected gene set are highlighted in orange. Horizontal lines indicate various False Discovery Rate (FDR) thresholds.   
                                           DE Results - CD209 (DC-SIGN) signaling genes - New.Annotation: Control vs.cel67   More Plot Information      Download CSV Data     
                                           
                                             
                                             
                                           
                                           
                                             
                                               Probe Label 
                                               Log2 fold change 
                                               std error (log2) 
                                               Lower confidence limit (log2) 
                                               Upper confidence limit (log2) 
                                               Linear fold change 
                                               Lower confidence limit (linear) 
                                               Upper confidence limit (linear) 
                                               P-value 
                                               BY.p.value 
                                               method 
                                               Gene.sets 
                                               Codeset.Name 
                                               Probe.Label 
                                               Analyte.Type 
                                               Is.Control 
                                               Control.Type 
                                               Related.Probes 
                                               Probe.Annotation 
                                               KEGG.Pathways 
                                               Official.Gene.Name 
                                               Control.Conc 
                                               volcanocol 
                                               volcanopch 
                                               tstats.all 
                                             
                                           
                                           
                                             
                                               NRAS-mRNA 
                                               -0.544 
                                               0.114 
                                               -0.767 
                                               -0.322 
                                               0.686 
                                               0.588 
                                               0.8 
                                               9.79e-05 
                                               0.026 
                                               lm.nb 
                                               ARMS-mediated activation, CD209 (DC-SIGN) signaling, Cell surface interactions at the vascular wall, Constitutive Signaling by EGFRvIII, DAP12 signaling, Downstream signal transduction, FCERI mediated MAPK activation, Frs2-mediated activation, Gastrin-CREB signalling pathway via PKC and MAPK, Interleukin receptor SHC signaling, IRS-related events triggered by IGF1R, MAP2K and MAPK activation, RAFMAP kinase cascade, SHC-mediated cascade.CL.FGFR1, SHC-related events triggered by IGF1R, Signaling by ERBB2, Signaling by FGFR1 in disease, Signaling by FGFR2 in disease, Signaling by FGFR3 in disease, Signaling by FGFR4 in disease, Signaling by PDGF, Signaling by SCF-KIT, Tie2 Signaling, VEGFA-VEGFR2 Pathway, VEGFR2 mediated cell proliferation 
                                               SEBELIUS_2_C4636 
                                               NRAS 
                                               mRNA 
                                               0 
                                               
                                               
                                               Signaling by FGFR2 in disease;Regulation of RAS by GAPs;SHC-mediated cascade.CL.FGFR3;Signaling by FGFR3 in disease;MAP2K and MAPK activation;SHC-related events triggered by IGF1R;Signaling by FGFR4 in disease;SHC-mediated cascade.CL.FGFR1;Tie2 Signaling;Signaling by FGFR1 in disease;Activation of RAS in B cells;SHC-mediated cascade.CL.FGFR4;Constitutive Signaling by EGFRvIII;CD209 (DC-SIGN) signaling;FCERI mediated MAPK activation;RAFMAP kinase cascade;Downstream signal transduction;VEGFR2 mediated cell proliferation;Signaling by SCF-KIT;Frs2-mediated activation;ARMS-mediated activation;Gastrin-CREB signalling pathway via PKC and MAPK;Interleukin receptor SHC signaling;DAP12 signaling;Signaling by PDGF;VEGFA-VEGFR2 Pathway;Signaling by ERBB2;IRS-related events triggered by IGF1R;Cell surface interactions at the vascular wall 
                                               hsa04010;hsa04012;hsa04014;hsa04015;hsa04062;hsa04068;hsa04071;hsa04072;hsa04151;hsa04211;hsa04213;hsa04360;hsa04370;hsa04530;hsa04540;hsa04550;hsa04650;hsa04660;hsa04662;hsa04664;hsa04720;hsa04722;hsa04725;hsa04726;hsa04730;hsa04810;hsa04910;hsa04912;hsa04915;hsa04916;hsa04917;hsa04919;hsa04921;hsa04933;hsa05034;hsa05160;hsa05161;hsa05166;hsa05200;hsa05203;hsa05205;hsa05206;hsa05211;hsa05213;hsa05214;hsa05215;hsa05216;hsa05218;hsa05219;hsa05220;hsa05221;hsa05223;hsa05230;hsa05231 
                                               NRAS 
                                               NA 
                                               slateblue2 
                                               16 
                                               -4.7719298245614 
                                             
                                             
                                               HRAS-mRNA 
                                               -0.593 
                                               0.144 
                                               -0.874 
                                               -0.311 
                                               0.663 
                                               0.545 
                                               0.806 
                                               0.000487 
                                               0.0718 
                                               loglinear 
                                               ARMS-mediated activation, CD209 (DC-SIGN) signaling, Cell surface interactions at the vascular wall, Constitutive Signaling by EGFRvIII, CREB phosphorylation through the activation of Ras, DAP12 signaling, Downstream signal transduction, FCERI mediated MAPK activation, Frs2-mediated activation, Gastrin-CREB signalling pathway via PKC and MAPK, Interleukin receptor SHC signaling, IRS-related events triggered by IGF1R, MAP2K and MAPK activation, RAFMAP kinase cascade, SHC-mediated cascade.CL.FGFR1, SHC-related events triggered by IGF1R, Signaling by ERBB2, Signaling by FGFR1 in disease, Signaling by FGFR2 in disease, Signaling by FGFR3 in disease, Signaling by FGFR4 in disease, Signaling by PDGF, Signaling by SCF-KIT, Tie2 Signaling, VEGFA-VEGFR2 Pathway, VEGFR2 mediated cell proliferation 
                                               SEBELIUS_2_C4636 
                                               HRAS 
                                               mRNA 
                                               0 
                                               
                                               
                                               Signaling by FGFR2 in disease;Regulation of RAS by GAPs;SHC-mediated cascade.CL.FGFR3;Signaling by FGFR3 in disease;MAP2K and MAPK activation;SHC-related events triggered by IGF1R;Signaling by FGFR4 in disease;SHC-mediated cascade.CL.FGFR1;Tie2 Signaling;Signaling by FGFR1 in disease;Activation of RAS in B cells;SHC-mediated cascade.CL.FGFR4;Constitutive Signaling by EGFRvIII;CD209 (DC-SIGN) signaling;Ras activation uopn Ca2.PLS. infux through NMDA receptor;FCERI mediated MAPK activation;RAFMAP kinase cascade;Downstream signal transduction;VEGFR2 mediated cell proliferation;Signaling by SCF-KIT;Frs2-mediated activation;ARMS-mediated activation;Gastrin-CREB signalling pathway via PKC and MAPK;Interleukin receptor SHC signaling;DAP12 signaling;Signaling by PDGF;VEGFA-VEGFR2 Pathway;Signaling by ERBB2;IRS-related events triggered by IGF1R;Cell surface interactions at the vascular wall;CREB phosphorylation through the activation of Ras 
                                               hsa04010;hsa04012;hsa04014;hsa04015;hsa04062;hsa04068;hsa04071;hsa04072;hsa04144;hsa04151;hsa04211;hsa04213;hsa04360;hsa04370;hsa04510;hsa04530;hsa04540;hsa04550;hsa04630;hsa04650;hsa04660;hsa04662;hsa04664;hsa04720;hsa04722;hsa04725;hsa04726;hsa04730;hsa04810;hsa04910;hsa04912;hsa04915;hsa04916;hsa04917;hsa04919;hsa04921;hsa04933;hsa05034;hsa05160;hsa05161;hsa05166;hsa05200;hsa05203;hsa05205;hsa05206;hsa05211;hsa05213;hsa05214;hsa05215;hsa05216;hsa05218;hsa05219;hsa05220;hsa05221;hsa05223;hsa05230;hsa05231 
                                               HRAS 
                                               NA 
                                               slateblue3 
                                               16 
                                               -4.11805555555556 
                                             
                                             
                                               PAK1-mRNA 
                                               -0.177 
                                               0.163 
                                               -0.497 
                                               0.142 
                                               0.884 
                                               0.709 
                                               1.1 
                                               0.288 
                                               1 
                                               lm.nb 
                                               Activation of Rac, CD209 (DC-SIGN) signaling, CD28 co-stimulation, CD28 dependent Vav1 pathway, EPHB-mediated forward signaling, Ephrin signaling, FCERI mediated MAPK activation, MAPK6MAPK4 signaling, Regulation of actin dynamics for phagocytic cup formation, RHO GTPases activate PAKs, Sema3A PAK dependent Axon repulsion, VEGFA-VEGFR2 Pathway, VEGFR2 mediated vascular permeability 
                                               SEBELIUS_2_C4636 
                                               PAK1 
                                               mRNA 
                                               0 
                                               
                                               
                                               EPHB-mediated forward signaling;Ephrin signaling;MAPK6MAPK4 signaling;DSCAM interactions;Activation of Rac;VEGFR2 mediated vascular permeability;RHO GTPases activate PAKs;Sema3A PAK dependent Axon repulsion;FCERI mediated MAPK activation;Regulation of actin dynamics for phagocytic cup formation;CD209 (DC-SIGN) signaling;CD28 dependent Vav1 pathway;VEGFA-VEGFR2 Pathway;CD28 co-stimulation 
                                               hsa04010;hsa04012;hsa04014;hsa04024;hsa04062;hsa04360;hsa04510;hsa04650;hsa04660;hsa04666;hsa04810;hsa05120;hsa05205;hsa05211 
                                               PAK1 
                                               NA 
                                               azure4 
                                               1 
                                               -1.08588957055215 
                                             
                                             
                                               PAK2-mRNA 
                                               -0.0762 
                                               0.0976 
                                               -0.268 
                                               0.115 
                                               0.949 
                                               0.831 
                                               1.08 
                                               0.444 
                                               1 
                                               loglinear 
                                               Activation of Rac, CD209 (DC-SIGN) signaling, CD28 co-stimulation, CD28 dependent Vav1 pathway, Ephrin signaling, RHO GTPases activate PAKs, Sema3A PAK dependent Axon repulsion, VEGFA-VEGFR2 Pathway, VEGFR2 mediated vascular permeability 
                                               SEBELIUS_2_C4636 
                                               PAK2 
                                               mRNA 
                                               0 
                                               
                                               
                                               Nef and signal transduction;CD28 dependent Vav1 pathway;Stimulation of the cell death response by PAK-2p34;Ephrin signaling;Activation of Rac;RHO GTPases activate PAKs;VEGFR2 mediated vascular permeability;Sema3A PAK dependent Axon repulsion;CD209 (DC-SIGN) signaling;VEGFA-VEGFR2 Pathway;CD28 co-stimulation 
                                               hsa04010;hsa04012;hsa04014;hsa04360;hsa04510;hsa04660;hsa04810;hsa05211 
                                               PAK2 
                                               NA 
                                               azure4 
                                               1 
                                               -0.780737704918033 
                                             
                                             
                                               RAF1-mRNA 
                                               -0.0567 
                                               0.0728 
                                               -0.199 
                                               0.0861 
                                               0.961 
                                               0.871 
                                               1.06 
                                               0.445 
                                               1 
                                               loglinear 
                                               ARMS-mediated activation, CD209 (DC-SIGN) signaling, CREB phosphorylation through the activation of Ras, DAP12 signaling, Downstream signal transduction, FCERI mediated MAPK activation, Frs2-mediated activation, Gastrin-CREB signalling pathway via PKC and MAPK, Interleukin receptor SHC signaling, IRS-related events triggered by IGF1R, Negative feedback regulation of MAPK pathway, Negative regulation of MAPK pathway, RAF activation, RAFMAP kinase cascade, Signaling by ERBB2, Signaling by PDGF, Signaling by SCF-KIT, VEGFA-VEGFR2 Pathway, VEGFR2 mediated cell proliferation 
                                               SEBELIUS_2_C4636 
                                               RAF1 
                                               mRNA 
                                               0 
                                               
                                               
                                               RAF activation;Stimuli-sensing channels;CREB phosphorylation through the activation of Ras;Negative feedback regulation of MAPK pathway;CD209 (DC-SIGN) signaling;RAFMAP kinase cascade;Downstream signal transduction;VEGFR2 mediated cell proliferation;Signaling by SCF-KIT;Frs2-mediated activation;ARMS-mediated activation;Gastrin-CREB signalling pathway via PKC and MAPK;Interleukin receptor SHC signaling;FCERI mediated MAPK activation;DAP12 signaling;Signaling by PDGF;VEGFA-VEGFR2 Pathway;Signaling by ERBB2;IRS-related events triggered by IGF1R;Negative regulation of MAPK pathway 
                                               hsa04010;hsa04012;hsa04014;hsa04015;hsa04022;hsa04024;hsa04062;hsa04068;hsa04071;hsa04072;hsa04151;hsa04270;hsa04370;hsa04510;hsa04540;hsa04550;hsa04630;hsa04650;hsa04660;hsa04662;hsa04664;hsa04666;hsa04720;hsa04722;hsa04726;hsa04730;hsa04810;hsa04910;hsa04912;hsa04914;hsa04915;hsa04916;hsa04917;hsa04919;hsa04921;hsa05034;hsa05152;hsa05160;hsa05161;hsa05164;hsa05200;hsa05205;hsa05206;hsa05210;hsa05211;hsa05212;hsa05213;hsa05214;hsa05215;hsa05218;hsa05219;hsa05220;hsa05221;hsa05223;hsa05230;hsa05231 
                                               RAF1 
                                               NA 
                                               azure4 
                                               1 
                                               -0.778846153846154 
                                             
                                             
                                               KRAS-mRNA 
                                               0.0772 
                                               0.101 
                                               -0.121 
                                               0.276 
                                               1.05 
                                               0.919 
                                               1.21 
                                               0.455 
                                               1 
                                               loglinear 
                                               ARMS-mediated activation, CD209 (DC-SIGN) signaling, Cell surface interactions at the vascular wall, Constitutive Signaling by EGFRvIII, DAP12 signaling, Downstream signal transduction, FCERI mediated MAPK activation, Frs2-mediated activation, Gastrin-CREB signalling pathway via PKC and MAPK, Interleukin receptor SHC signaling, IRS-related events triggered by IGF1R, MAP2K and MAPK activation, RAFMAP kinase cascade, SHC-mediated cascade.CL.FGFR1, SHC-related events triggered by IGF1R, Signaling by ERBB2, Signaling by FGFR1 in disease, Signaling by FGFR2 in disease, Signaling by FGFR3 in disease, Signaling by FGFR4 in disease, Signaling by PDGF, Signaling by SCF-KIT, Tie2 Signaling, VEGFA-VEGFR2 Pathway, VEGFR2 mediated cell proliferation 
                                               SEBELIUS_2_C4636 
                                               KRAS 
                                               mRNA 
                                               0 
                                               
                                               
                                               Signaling by FGFR2 in disease;Regulation of RAS by GAPs;SHC-mediated cascade.CL.FGFR3;Signaling by FGFR3 in disease;MAP2K and MAPK activation;SHC-related events triggered by IGF1R;Signaling by FGFR4 in disease;SHC-mediated cascade.CL.FGFR1;Tie2 Signaling;Signaling by FGFR1 in disease;Activation of RAS in B cells;SHC-mediated cascade.CL.FGFR4;Constitutive Signaling by EGFRvIII;CD209 (DC-SIGN) signaling;FCERI mediated MAPK activation;RAFMAP kinase cascade;Downstream signal transduction;VEGFR2 mediated cell proliferation;Signaling by SCF-KIT;Frs2-mediated activation;ARMS-mediated activation;Gastrin-CREB signalling pathway via PKC and MAPK;Interleukin receptor SHC signaling;DAP12 signaling;Signaling by PDGF;VEGFA-VEGFR2 Pathway;Signaling by ERBB2;IRS-related events triggered by IGF1R;Cell surface interactions at the vascular wall 
                                               hsa04010;hsa04012;hsa04014;hsa04015;hsa04062;hsa04068;hsa04071;hsa04072;hsa04151;hsa04211;hsa04213;hsa04320;hsa04360;hsa04370;hsa04530;hsa04540;hsa04550;hsa04650;hsa04660;hsa04662;hsa04664;hsa04720;hsa04722;hsa04725;hsa04726;hsa04730;hsa04810;hsa04910;hsa04912;hsa04914;hsa04915;hsa04916;hsa04917;hsa04919;hsa04921;hsa04933;hsa04960;hsa05034;hsa05160;hsa05161;hsa05166;hsa05200;hsa05203;hsa05205;hsa05206;hsa05210;hsa05211;hsa05212;hsa05213;hsa05214;hsa05215;hsa05216;hsa05218;hsa05219;hsa05220;hsa05221;hsa05223;hsa05230;hsa05231 
                                               KRAS 
                                               NA 
                                               azure4 
                                               1 
                                               0.764356435643564 
                                             
                                             
                                               CREBBP-mRNA 
                                               -0.0445 
                                               0.0805 
                                               -0.202 
                                               0.113 
                                               0.97 
                                               0.869 
                                               1.08 
                                               0.586 
                                               1 
                                               loglinear 
                                               CD209 (DC-SIGN) signaling, Circadian Clock 
                                               SEBELIUS_2_C4636 
                                               CREBBP 
                                               mRNA 
                                               0 
                                               
                                               
                                               YAP1- and WWTR1 (TAZ)-stimulated gene expression;Activation of gene expression by SREBF (SREBP);Transcriptional activation of mitochondrial biogenesis;Constitutive Signaling by NOTCH1 HD.PLS.PEST Domain Mutants;Constitutive Signaling by NOTCH1 PEST Domain Mutants;CD209 (DC-SIGN) signaling;Circadian Clock 
                                               hsa04024;hsa04066;hsa04068;hsa04110;hsa04310;hsa04330;hsa04350;hsa04520;hsa04630;hsa04720;hsa04916;hsa04919;hsa04922;hsa05016;hsa05152;hsa05161;hsa05164;hsa05166;hsa05168;hsa05169;hsa05200;hsa05203;hsa05206;hsa05211;hsa05215 
                                               CREBBP 
                                               NA 
                                               azure4 
                                               1 
                                               -0.552795031055901 
                                             
                                             
                                               FYN-mRNA 
                                               0.0581 
                                               0.11 
                                               -0.157 
                                               0.273 
                                               1.04 
                                               0.897 
                                               1.21 
                                               0.602 
                                               1 
                                               loglinear 
                                               ARMS-mediated activation, CD209 (DC-SIGN) signaling, CD28 co-stimulation, CD28 dependent PI3KAkt signaling, CD28 dependent Vav1 pathway, Cell surface interactions at the vascular wall, Constitutive Signaling by Aberrant PI3K in Cancer, CTLA4 inhibitory signaling, DAP12 signaling, Downstream signal transduction, EPH-ephrin mediated repulsion of cells, FCERI mediated MAPK activation, Frs2-mediated activation, GAB1 signalosome, Gastrin-CREB signalling pathway via PKC and MAPK, GPVI-mediated activation cascade, Interleukin receptor SHC signaling, IRS-related events triggered by IGF1R, Nephrin interactions, PI3KAKT activation, RAFMAP kinase cascade, Regulation of signaling by CBL, Role of LAT2NTALLAB on calcium mobilization, Sema3A PAK dependent Axon repulsion, Signaling by ERBB2, Signaling by PDGF, Signaling by SCF-KIT, VEGFA-VEGFR2 Pathway, VEGFR2 mediated cell proliferation 
                                               SEBELIUS_2_C4636 
                                               FYN 
                                               mRNA 
                                               0 
                                               
                                               
                                               EPH-ephrin mediated repulsion of cells;Regulation of signaling by CBL;FCGR activation;Antigen activates B Cell Receptor (BCR) leading to generation of second messengers;Dectin-2 family;RAFMAP kinase cascade;Constitutive Signaling by Aberrant PI3K in Cancer;CTLA4 inhibitory signaling;CD28 dependent PI3KAkt signaling;PECAM1 interactions;CD28 dependent Vav1 pathway;GPVI-mediated activation cascade;Sema3A PAK dependent Axon repulsion;Nephrin interactions;SEMA3A-Plexin repulsion signaling by inhibiting Integrin adhesion;CRMPs in Sema3A signaling;CD209 (DC-SIGN) signaling;Downstream signal transduction;GAB1 signalosome;Signaling by SCF-KIT;PI3KAKT activation;Role of LAT2NTALLAB on calcium mobilization;DAP12 signaling;Signaling by PDGF;Signaling by ERBB2;VEGFA-VEGFR2 Pathway;VEGFR2 mediated cell proliferation;Frs2-mediated activation;ARMS-mediated activation;Gastrin-CREB signalling pathway via PKC and MAPK;Interleukin receptor SHC signaling;FCERI mediated MAPK activation;IRS-related events triggered by IGF1R;CD28 co-stimulation;Cell surface interactions at the vascular wall 
                                               hsa04071;hsa04072;hsa04360;hsa04380;hsa04510;hsa04520;hsa04611;hsa04650;hsa04660;hsa04664;hsa04725;hsa05020;hsa05130;hsa05162;hsa05416 
                                               FYN 
                                               NA 
                                               azure4 
                                               1 
                                               0.528181818181818 
                                             
                                             
                                               EP300-mRNA 
                                               -0.028 
                                               0.106 
                                               -0.236 
                                               0.18 
                                               0.981 
                                               0.849 
                                               1.13 
                                               0.795 
                                               1 
                                               loglinear 
                                               CD209 (DC-SIGN) signaling, Circadian Clock 
                                               SEBELIUS_2_C4636 
                                               EP300 
                                               mRNA 
                                               0 
                                               
                                               
                                               LRR FLII-interacting protein 1 (LRRFIP1) activates type I IFN production;TRAF6 mediated IRF7 activation;Constitutive Signaling by NOTCH1 PEST Domain Mutants;Polo-like kinase mediated events;B-WICH complex positively regulates rRNA expression;CD209 (DC-SIGN) signaling;Circadian Clock 
                                               hsa04024;hsa04066;hsa04068;hsa04110;hsa04310;hsa04330;hsa04350;hsa04520;hsa04630;hsa04720;hsa04916;hsa04919;hsa04922;hsa05016;hsa05152;hsa05161;hsa05164;hsa05166;hsa05168;hsa05169;hsa05200;hsa05203;hsa05206;hsa05211;hsa05215 
                                               EP300 
                                               NA 
                                               azure4 
                                               1 
                                               -0.264150943396226 
                                             
                                           
                                           DE Results - CD209 (DC-SIGN) signaling genes - New.Annotation: Control vs.cel67  Table displaying each sample's global significance scores and directed global significance scores as defined in the heatmaps above. The global significance score is calculated as the square root of the mean squared t-statistic for the genes in a gene set, with t-statistics coming from the linear regression underlying our differential expression analysis. The directed global significance score is calculated as the square root of the mean signed squared t-statistic for the genes in a gene set, with t-statistics coming from the linear regression underlying our differential expression analysis.   
                                      
                                       
                                  
                                
                                
                                  
                                       
                                      
                                           Volcano Plot: New.Annotation: miR142 vs.cel67   More Plot Information                Volcano Plot: New.Annotation: miR142 vs.cel67  Volcano plot displaying each gene's -log10(p-value) and log2 fold change for the selected covariate. Highly statistically significant genes fall at the top of the plot, and highly differentially expressed genes fall to either side. Genes within the selected gene set are highlighted in orange. Horizontal lines indicate various False Discovery Rate (FDR) thresholds.   
                                           DE Results - CD209 (DC-SIGN) signaling genes - New.Annotation: miR142 vs.cel67   More Plot Information      Download CSV Data     
                                           
                                             
                                             
                                           
                                           
                                             
                                               Probe Label 
                                               Log2 fold change 
                                               std error (log2) 
                                               Lower confidence limit (log2) 
                                               Upper confidence limit (log2) 
                                               Linear fold change 
                                               Lower confidence limit (linear) 
                                               Upper confidence limit (linear) 
                                               P-value 
                                               BY.p.value 
                                               method 
                                               Gene.sets 
                                               Codeset.Name 
                                               Probe.Label 
                                               Analyte.Type 
                                               Is.Control 
                                               Control.Type 
                                               Related.Probes 
                                               Probe.Annotation 
                                               KEGG.Pathways 
                                               Official.Gene.Name 
                                               Control.Conc 
                                               volcanocol 
                                               volcanopch 
                                               tstats.all 
                                             
                                           
                                           
                                             
                                               HRAS-mRNA 
                                               -0.987 
                                               0.144 
                                               -1.27 
                                               -0.705 
                                               0.504 
                                               0.415 
                                               0.613 
                                               8.72e-07 
                                               6.43e-05 
                                               loglinear 
                                               ARMS-mediated activation, CD209 (DC-SIGN) signaling, Cell surface interactions at the vascular wall, Constitutive Signaling by EGFRvIII, CREB phosphorylation through the activation of Ras, DAP12 signaling, Downstream signal transduction, FCERI mediated MAPK activation, Frs2-mediated activation, Gastrin-CREB signalling pathway via PKC and MAPK, Interleukin receptor SHC signaling, IRS-related events triggered by IGF1R, MAP2K and MAPK activation, RAFMAP kinase cascade, SHC-mediated cascade.CL.FGFR1, SHC-related events triggered by IGF1R, Signaling by ERBB2, Signaling by FGFR1 in disease, Signaling by FGFR2 in disease, Signaling by FGFR3 in disease, Signaling by FGFR4 in disease, Signaling by PDGF, Signaling by SCF-KIT, Tie2 Signaling, VEGFA-VEGFR2 Pathway, VEGFR2 mediated cell proliferation 
                                               SEBELIUS_2_C4636 
                                               HRAS 
                                               mRNA 
                                               0 
                                               
                                               
                                               Signaling by FGFR2 in disease;Regulation of RAS by GAPs;SHC-mediated cascade.CL.FGFR3;Signaling by FGFR3 in disease;MAP2K and MAPK activation;SHC-related events triggered by IGF1R;Signaling by FGFR4 in disease;SHC-mediated cascade.CL.FGFR1;Tie2 Signaling;Signaling by FGFR1 in disease;Activation of RAS in B cells;SHC-mediated cascade.CL.FGFR4;Constitutive Signaling by EGFRvIII;CD209 (DC-SIGN) signaling;Ras activation uopn Ca2.PLS. infux through NMDA receptor;FCERI mediated MAPK activation;RAFMAP kinase cascade;Downstream signal transduction;VEGFR2 mediated cell proliferation;Signaling by SCF-KIT;Frs2-mediated activation;ARMS-mediated activation;Gastrin-CREB signalling pathway via PKC and MAPK;Interleukin receptor SHC signaling;DAP12 signaling;Signaling by PDGF;VEGFA-VEGFR2 Pathway;Signaling by ERBB2;IRS-related events triggered by IGF1R;Cell surface interactions at the vascular wall;CREB phosphorylation through the activation of Ras 
                                               hsa04010;hsa04012;hsa04014;hsa04015;hsa04062;hsa04068;hsa04071;hsa04072;hsa04144;hsa04151;hsa04211;hsa04213;hsa04360;hsa04370;hsa04510;hsa04530;hsa04540;hsa04550;hsa04630;hsa04650;hsa04660;hsa04662;hsa04664;hsa04720;hsa04722;hsa04725;hsa04726;hsa04730;hsa04810;hsa04910;hsa04912;hsa04915;hsa04916;hsa04917;hsa04919;hsa04921;hsa04933;hsa05034;hsa05160;hsa05161;hsa05166;hsa05200;hsa05203;hsa05205;hsa05206;hsa05211;hsa05213;hsa05214;hsa05215;hsa05216;hsa05218;hsa05219;hsa05220;hsa05221;hsa05223;hsa05230;hsa05231 
                                               HRAS 
                                               NA 
                                               slateblue1 
                                               16 
                                               -6.85416666666667 
                                             
                                             
                                               NRAS-mRNA 
                                               -0.363 
                                               0.113 
                                               -0.585 
                                               -0.141 
                                               0.777 
                                               0.666 
                                               0.907 
                                               0.00423 
                                               0.073 
                                               lm.nb 
                                               ARMS-mediated activation, CD209 (DC-SIGN) signaling, Cell surface interactions at the vascular wall, Constitutive Signaling by EGFRvIII, DAP12 signaling, Downstream signal transduction, FCERI mediated MAPK activation, Frs2-mediated activation, Gastrin-CREB signalling pathway via PKC and MAPK, Interleukin receptor SHC signaling, IRS-related events triggered by IGF1R, MAP2K and MAPK activation, RAFMAP kinase cascade, SHC-mediated cascade.CL.FGFR1, SHC-related events triggered by IGF1R, Signaling by ERBB2, Signaling by FGFR1 in disease, Signaling by FGFR2 in disease, Signaling by FGFR3 in disease, Signaling by FGFR4 in disease, Signaling by PDGF, Signaling by SCF-KIT, Tie2 Signaling, VEGFA-VEGFR2 Pathway, VEGFR2 mediated cell proliferation 
                                               SEBELIUS_2_C4636 
                                               NRAS 
                                               mRNA 
                                               0 
                                               
                                               
                                               Signaling by FGFR2 in disease;Regulation of RAS by GAPs;SHC-mediated cascade.CL.FGFR3;Signaling by FGFR3 in disease;MAP2K and MAPK activation;SHC-related events triggered by IGF1R;Signaling by FGFR4 in disease;SHC-mediated cascade.CL.FGFR1;Tie2 Signaling;Signaling by FGFR1 in disease;Activation of RAS in B cells;SHC-mediated cascade.CL.FGFR4;Constitutive Signaling by EGFRvIII;CD209 (DC-SIGN) signaling;FCERI mediated MAPK activation;RAFMAP kinase cascade;Downstream signal transduction;VEGFR2 mediated cell proliferation;Signaling by SCF-KIT;Frs2-mediated activation;ARMS-mediated activation;Gastrin-CREB signalling pathway via PKC and MAPK;Interleukin receptor SHC signaling;DAP12 signaling;Signaling by PDGF;VEGFA-VEGFR2 Pathway;Signaling by ERBB2;IRS-related events triggered by IGF1R;Cell surface interactions at the vascular wall 
                                               hsa04010;hsa04012;hsa04014;hsa04015;hsa04062;hsa04068;hsa04071;hsa04072;hsa04151;hsa04211;hsa04213;hsa04360;hsa04370;hsa04530;hsa04540;hsa04550;hsa04650;hsa04660;hsa04662;hsa04664;hsa04720;hsa04722;hsa04725;hsa04726;hsa04730;hsa04810;hsa04910;hsa04912;hsa04915;hsa04916;hsa04917;hsa04919;hsa04921;hsa04933;hsa05034;hsa05160;hsa05161;hsa05166;hsa05200;hsa05203;hsa05205;hsa05206;hsa05211;hsa05213;hsa05214;hsa05215;hsa05216;hsa05218;hsa05219;hsa05220;hsa05221;hsa05223;hsa05230;hsa05231 
                                               NRAS 
                                               NA 
                                               slateblue3 
                                               16 
                                               -3.21238938053097 
                                             
                                             
                                               CREBBP-mRNA 
                                               0.247 
                                               0.0805 
                                               0.089 
                                               0.405 
                                               1.19 
                                               1.06 
                                               1.32 
                                               0.00588 
                                               0.0952 
                                               loglinear 
                                               CD209 (DC-SIGN) signaling, Circadian Clock 
                                               SEBELIUS_2_C4636 
                                               CREBBP 
                                               mRNA 
                                               0 
                                               
                                               
                                               YAP1- and WWTR1 (TAZ)-stimulated gene expression;Activation of gene expression by SREBF (SREBP);Transcriptional activation of mitochondrial biogenesis;Constitutive Signaling by NOTCH1 HD.PLS.PEST Domain Mutants;Constitutive Signaling by NOTCH1 PEST Domain Mutants;CD209 (DC-SIGN) signaling;Circadian Clock 
                                               hsa04024;hsa04066;hsa04068;hsa04110;hsa04310;hsa04330;hsa04350;hsa04520;hsa04630;hsa04720;hsa04916;hsa04919;hsa04922;hsa05016;hsa05152;hsa05161;hsa05164;hsa05166;hsa05168;hsa05169;hsa05200;hsa05203;hsa05206;hsa05211;hsa05215 
                                               CREBBP 
                                               NA 
                                               slateblue3 
                                               16 
                                               3.06832298136646 
                                             
                                             
                                               PAK1-mRNA 
                                               -0.455 
                                               0.163 
                                               -0.775 
                                               -0.135 
                                               0.729 
                                               0.584 
                                               0.91 
                                               0.011 
                                               0.164 
                                               lm.nb 
                                               Activation of Rac, CD209 (DC-SIGN) signaling, CD28 co-stimulation, CD28 dependent Vav1 pathway, EPHB-mediated forward signaling, Ephrin signaling, FCERI mediated MAPK activation, MAPK6MAPK4 signaling, Regulation of actin dynamics for phagocytic cup formation, RHO GTPases activate PAKs, Sema3A PAK dependent Axon repulsion, VEGFA-VEGFR2 Pathway, VEGFR2 mediated vascular permeability 
                                               SEBELIUS_2_C4636 
                                               PAK1 
                                               mRNA 
                                               0 
                                               
                                               
                                               EPHB-mediated forward signaling;Ephrin signaling;MAPK6MAPK4 signaling;DSCAM interactions;Activation of Rac;VEGFR2 mediated vascular permeability;RHO GTPases activate PAKs;Sema3A PAK dependent Axon repulsion;FCERI mediated MAPK activation;Regulation of actin dynamics for phagocytic cup formation;CD209 (DC-SIGN) signaling;CD28 dependent Vav1 pathway;VEGFA-VEGFR2 Pathway;CD28 co-stimulation 
                                               hsa04010;hsa04012;hsa04014;hsa04024;hsa04062;hsa04360;hsa04510;hsa04650;hsa04660;hsa04666;hsa04810;hsa05120;hsa05205;hsa05211 
                                               PAK1 
                                               NA 
                                               slateblue4 
                                               16 
                                               -2.79141104294479 
                                             
                                             
                                               KRAS-mRNA 
                                               0.276 
                                               0.101 
                                               0.0773 
                                               0.474 
                                               1.21 
                                               1.06 
                                               1.39 
                                               0.0127 
                                               0.182 
                                               loglinear 
                                               ARMS-mediated activation, CD209 (DC-SIGN) signaling, Cell surface interactions at the vascular wall, Constitutive Signaling by EGFRvIII, DAP12 signaling, Downstream signal transduction, FCERI mediated MAPK activation, Frs2-mediated activation, Gastrin-CREB signalling pathway via PKC and MAPK, Interleukin receptor SHC signaling, IRS-related events triggered by IGF1R, MAP2K and MAPK activation, RAFMAP kinase cascade, SHC-mediated cascade.CL.FGFR1, SHC-related events triggered by IGF1R, Signaling by ERBB2, Signaling by FGFR1 in disease, Signaling by FGFR2 in disease, Signaling by FGFR3 in disease, Signaling by FGFR4 in disease, Signaling by PDGF, Signaling by SCF-KIT, Tie2 Signaling, VEGFA-VEGFR2 Pathway, VEGFR2 mediated cell proliferation 
                                               SEBELIUS_2_C4636 
                                               KRAS 
                                               mRNA 
                                               0 
                                               
                                               
                                               Signaling by FGFR2 in disease;Regulation of RAS by GAPs;SHC-mediated cascade.CL.FGFR3;Signaling by FGFR3 in disease;MAP2K and MAPK activation;SHC-related events triggered by IGF1R;Signaling by FGFR4 in disease;SHC-mediated cascade.CL.FGFR1;Tie2 Signaling;Signaling by FGFR1 in disease;Activation of RAS in B cells;SHC-mediated cascade.CL.FGFR4;Constitutive Signaling by EGFRvIII;CD209 (DC-SIGN) signaling;FCERI mediated MAPK activation;RAFMAP kinase cascade;Downstream signal transduction;VEGFR2 mediated cell proliferation;Signaling by SCF-KIT;Frs2-mediated activation;ARMS-mediated activation;Gastrin-CREB signalling pathway via PKC and MAPK;Interleukin receptor SHC signaling;DAP12 signaling;Signaling by PDGF;VEGFA-VEGFR2 Pathway;Signaling by ERBB2;IRS-related events triggered by IGF1R;Cell surface interactions at the vascular wall 
                                               hsa04010;hsa04012;hsa04014;hsa04015;hsa04062;hsa04068;hsa04071;hsa04072;hsa04151;hsa04211;hsa04213;hsa04320;hsa04360;hsa04370;hsa04530;hsa04540;hsa04550;hsa04650;hsa04660;hsa04662;hsa04664;hsa04720;hsa04722;hsa04725;hsa04726;hsa04730;hsa04810;hsa04910;hsa04912;hsa04914;hsa04915;hsa04916;hsa04917;hsa04919;hsa04921;hsa04933;hsa04960;hsa05034;hsa05160;hsa05161;hsa05166;hsa05200;hsa05203;hsa05205;hsa05206;hsa05210;hsa05211;hsa05212;hsa05213;hsa05214;hsa05215;hsa05216;hsa05218;hsa05219;hsa05220;hsa05221;hsa05223;hsa05230;hsa05231 
                                               KRAS 
                                               NA 
                                               slateblue4 
                                               16 
                                               2.73267326732673 
                                             
                                             
                                               FYN-mRNA 
                                               0.275 
                                               0.11 
                                               0.0594 
                                               0.49 
                                               1.21 
                                               1.04 
                                               1.4 
                                               0.0207 
                                               0.272 
                                               loglinear 
                                               ARMS-mediated activation, CD209 (DC-SIGN) signaling, CD28 co-stimulation, CD28 dependent PI3KAkt signaling, CD28 dependent Vav1 pathway, Cell surface interactions at the vascular wall, Constitutive Signaling by Aberrant PI3K in Cancer, CTLA4 inhibitory signaling, DAP12 signaling, Downstream signal transduction, EPH-ephrin mediated repulsion of cells, FCERI mediated MAPK activation, Frs2-mediated activation, GAB1 signalosome, Gastrin-CREB signalling pathway via PKC and MAPK, GPVI-mediated activation cascade, Interleukin receptor SHC signaling, IRS-related events triggered by IGF1R, Nephrin interactions, PI3KAKT activation, RAFMAP kinase cascade, Regulation of signaling by CBL, Role of LAT2NTALLAB on calcium mobilization, Sema3A PAK dependent Axon repulsion, Signaling by ERBB2, Signaling by PDGF, Signaling by SCF-KIT, VEGFA-VEGFR2 Pathway, VEGFR2 mediated cell proliferation 
                                               SEBELIUS_2_C4636 
                                               FYN 
                                               mRNA 
                                               0 
                                               
                                               
                                               EPH-ephrin mediated repulsion of cells;Regulation of signaling by CBL;FCGR activation;Antigen activates B Cell Receptor (BCR) leading to generation of second messengers;Dectin-2 family;RAFMAP kinase cascade;Constitutive Signaling by Aberrant PI3K in Cancer;CTLA4 inhibitory signaling;CD28 dependent PI3KAkt signaling;PECAM1 interactions;CD28 dependent Vav1 pathway;GPVI-mediated activation cascade;Sema3A PAK dependent Axon repulsion;Nephrin interactions;SEMA3A-Plexin repulsion signaling by inhibiting Integrin adhesion;CRMPs in Sema3A signaling;CD209 (DC-SIGN) signaling;Downstream signal transduction;GAB1 signalosome;Signaling by SCF-KIT;PI3KAKT activation;Role of LAT2NTALLAB on calcium mobilization;DAP12 signaling;Signaling by PDGF;Signaling by ERBB2;VEGFA-VEGFR2 Pathway;VEGFR2 mediated cell proliferation;Frs2-mediated activation;ARMS-mediated activation;Gastrin-CREB signalling pathway via PKC and MAPK;Interleukin receptor SHC signaling;FCERI mediated MAPK activation;IRS-related events triggered by IGF1R;CD28 co-stimulation;Cell surface interactions at the vascular wall 
                                               hsa04071;hsa04072;hsa04360;hsa04380;hsa04510;hsa04520;hsa04611;hsa04650;hsa04660;hsa04664;hsa04725;hsa05020;hsa05130;hsa05162;hsa05416 
                                               FYN 
                                               NA 
                                               slateblue4 
                                               16 
                                               2.5 
                                             
                                             
                                               RAF1-mRNA 
                                               -0.129 
                                               0.0728 
                                               -0.272 
                                               0.0133 
                                               0.914 
                                               0.828 
                                               1.01 
                                               0.0899 
                                               0.925 
                                               loglinear 
                                               ARMS-mediated activation, CD209 (DC-SIGN) signaling, CREB phosphorylation through the activation of Ras, DAP12 signaling, Downstream signal transduction, FCERI mediated MAPK activation, Frs2-mediated activation, Gastrin-CREB signalling pathway via PKC and MAPK, Interleukin receptor SHC signaling, IRS-related events triggered by IGF1R, Negative feedback regulation of MAPK pathway, Negative regulation of MAPK pathway, RAF activation, RAFMAP kinase cascade, Signaling by ERBB2, Signaling by PDGF, Signaling by SCF-KIT, VEGFA-VEGFR2 Pathway, VEGFR2 mediated cell proliferation 
                                               SEBELIUS_2_C4636 
                                               RAF1 
                                               mRNA 
                                               0 
                                               
                                               
                                               RAF activation;Stimuli-sensing channels;CREB phosphorylation through the activation of Ras;Negative feedback regulation of MAPK pathway;CD209 (DC-SIGN) signaling;RAFMAP kinase cascade;Downstream signal transduction;VEGFR2 mediated cell proliferation;Signaling by SCF-KIT;Frs2-mediated activation;ARMS-mediated activation;Gastrin-CREB signalling pathway via PKC and MAPK;Interleukin receptor SHC signaling;FCERI mediated MAPK activation;DAP12 signaling;Signaling by PDGF;VEGFA-VEGFR2 Pathway;Signaling by ERBB2;IRS-related events triggered by IGF1R;Negative regulation of MAPK pathway 
                                               hsa04010;hsa04012;hsa04014;hsa04015;hsa04022;hsa04024;hsa04062;hsa04068;hsa04071;hsa04072;hsa04151;hsa04270;hsa04370;hsa04510;hsa04540;hsa04550;hsa04630;hsa04650;hsa04660;hsa04662;hsa04664;hsa04666;hsa04720;hsa04722;hsa04726;hsa04730;hsa04810;hsa04910;hsa04912;hsa04914;hsa04915;hsa04916;hsa04917;hsa04919;hsa04921;hsa05034;hsa05152;hsa05160;hsa05161;hsa05164;hsa05200;hsa05205;hsa05206;hsa05210;hsa05211;hsa05212;hsa05213;hsa05214;hsa05215;hsa05218;hsa05219;hsa05220;hsa05221;hsa05223;hsa05230;hsa05231 
                                               RAF1 
                                               NA 
                                               azure4 
                                               1 
                                               -1.77197802197802 
                                             
                                             
                                               EP300-mRNA 
                                               0.162 
                                               0.106 
                                               -0.0462 
                                               0.37 
                                               1.12 
                                               0.968 
                                               1.29 
                                               0.142 
                                               1 
                                               loglinear 
                                               CD209 (DC-SIGN) signaling, Circadian Clock 
                                               SEBELIUS_2_C4636 
                                               EP300 
                                               mRNA 
                                               0 
                                               
                                               
                                               LRR FLII-interacting protein 1 (LRRFIP1) activates type I IFN production;TRAF6 mediated IRF7 activation;Constitutive Signaling by NOTCH1 PEST Domain Mutants;Polo-like kinase mediated events;B-WICH complex positively regulates rRNA expression;CD209 (DC-SIGN) signaling;Circadian Clock 
                                               hsa04024;hsa04066;hsa04068;hsa04110;hsa04310;hsa04330;hsa04350;hsa04520;hsa04630;hsa04720;hsa04916;hsa04919;hsa04922;hsa05016;hsa05152;hsa05161;hsa05164;hsa05166;hsa05168;hsa05169;hsa05200;hsa05203;hsa05206;hsa05211;hsa05215 
                                               EP300 
                                               NA 
                                               azure4 
                                               1 
                                               1.52830188679245 
                                             
                                             
                                               PAK2-mRNA 
                                               0.147 
                                               0.0976 
                                               -0.0448 
                                               0.338 
                                               1.11 
                                               0.969 
                                               1.26 
                                               0.148 
                                               1 
                                               loglinear 
                                               Activation of Rac, CD209 (DC-SIGN) signaling, CD28 co-stimulation, CD28 dependent Vav1 pathway, Ephrin signaling, RHO GTPases activate PAKs, Sema3A PAK dependent Axon repulsion, VEGFA-VEGFR2 Pathway, VEGFR2 mediated vascular permeability 
                                               SEBELIUS_2_C4636 
                                               PAK2 
                                               mRNA 
                                               0 
                                               
                                               
                                               Nef and signal transduction;CD28 dependent Vav1 pathway;Stimulation of the cell death response by PAK-2p34;Ephrin signaling;Activation of Rac;RHO GTPases activate PAKs;VEGFR2 mediated vascular permeability;Sema3A PAK dependent Axon repulsion;CD209 (DC-SIGN) signaling;VEGFA-VEGFR2 Pathway;CD28 co-stimulation 
                                               hsa04010;hsa04012;hsa04014;hsa04360;hsa04510;hsa04660;hsa04810;hsa05211 
                                               PAK2 
                                               NA 
                                               azure4 
                                               1 
                                               1.50614754098361 
                                             
                                           
                                           DE Results - CD209 (DC-SIGN) signaling genes - New.Annotation: miR142 vs.cel67  Table displaying each sample's global significance scores and directed global significance scores as defined in the heatmaps above. The global significance score is calculated as the square root of the mean squared t-statistic for the genes in a gene set, with t-statistics coming from the linear regression underlying our differential expression analysis. The directed global significance score is calculated as the square root of the mean signed squared t-statistic for the genes in a gene set, with t-statistics coming from the linear regression underlying our differential expression analysis.   
                                      
                                       
                                  
                                
                                
                                  
                                       
                                      
                                           Volcano Plot: New.Annotation: miR155 vs.cel67   More Plot Information                Volcano Plot: New.Annotation: miR155 vs.cel67  Volcano plot displaying each gene's -log10(p-value) and log2 fold change for the selected covariate. Highly statistically significant genes fall at the top of the plot, and highly differentially expressed genes fall to either side. Genes within the selected gene set are highlighted in orange. Horizontal lines indicate various False Discovery Rate (FDR) thresholds.   
                                           DE Results - CD209 (DC-SIGN) signaling genes - New.Annotation: miR155 vs.cel67   More Plot Information      Download CSV Data     
                                           
                                             
                                             
                                           
                                           
                                             
                                               Probe Label 
                                               Log2 fold change 
                                               std error (log2) 
                                               Lower confidence limit (log2) 
                                               Upper confidence limit (log2) 
                                               Linear fold change 
                                               Lower confidence limit (linear) 
                                               Upper confidence limit (linear) 
                                               P-value 
                                               BY.p.value 
                                               method 
                                               Gene.sets 
                                               Codeset.Name 
                                               Probe.Label 
                                               Analyte.Type 
                                               Is.Control 
                                               Control.Type 
                                               Related.Probes 
                                               Probe.Annotation 
                                               KEGG.Pathways 
                                               Official.Gene.Name 
                                               Control.Conc 
                                               volcanocol 
                                               volcanopch 
                                               tstats.all 
                                             
                                           
                                           
                                             
                                               KRAS-mRNA 
                                               -0.99 
                                               0.107 
                                               -1.2 
                                               -0.781 
                                               0.503 
                                               0.435 
                                               0.582 
                                               7.18e-09 
                                               2.06e-06 
                                               loglinear 
                                               ARMS-mediated activation, CD209 (DC-SIGN) signaling, Cell surface interactions at the vascular wall, Constitutive Signaling by EGFRvIII, DAP12 signaling, Downstream signal transduction, FCERI mediated MAPK activation, Frs2-mediated activation, Gastrin-CREB signalling pathway via PKC and MAPK, Interleukin receptor SHC signaling, IRS-related events triggered by IGF1R, MAP2K and MAPK activation, RAFMAP kinase cascade, SHC-mediated cascade.CL.FGFR1, SHC-related events triggered by IGF1R, Signaling by ERBB2, Signaling by FGFR1 in disease, Signaling by FGFR2 in disease, Signaling by FGFR3 in disease, Signaling by FGFR4 in disease, Signaling by PDGF, Signaling by SCF-KIT, Tie2 Signaling, VEGFA-VEGFR2 Pathway, VEGFR2 mediated cell proliferation 
                                               SEBELIUS_2_C4636 
                                               KRAS 
                                               mRNA 
                                               0 
                                               
                                               
                                               Signaling by FGFR2 in disease;Regulation of RAS by GAPs;SHC-mediated cascade.CL.FGFR3;Signaling by FGFR3 in disease;MAP2K and MAPK activation;SHC-related events triggered by IGF1R;Signaling by FGFR4 in disease;SHC-mediated cascade.CL.FGFR1;Tie2 Signaling;Signaling by FGFR1 in disease;Activation of RAS in B cells;SHC-mediated cascade.CL.FGFR4;Constitutive Signaling by EGFRvIII;CD209 (DC-SIGN) signaling;FCERI mediated MAPK activation;RAFMAP kinase cascade;Downstream signal transduction;VEGFR2 mediated cell proliferation;Signaling by SCF-KIT;Frs2-mediated activation;ARMS-mediated activation;Gastrin-CREB signalling pathway via PKC and MAPK;Interleukin receptor SHC signaling;DAP12 signaling;Signaling by PDGF;VEGFA-VEGFR2 Pathway;Signaling by ERBB2;IRS-related events triggered by IGF1R;Cell surface interactions at the vascular wall 
                                               hsa04010;hsa04012;hsa04014;hsa04015;hsa04062;hsa04068;hsa04071;hsa04072;hsa04151;hsa04211;hsa04213;hsa04320;hsa04360;hsa04370;hsa04530;hsa04540;hsa04550;hsa04650;hsa04660;hsa04662;hsa04664;hsa04720;hsa04722;hsa04725;hsa04726;hsa04730;hsa04810;hsa04910;hsa04912;hsa04914;hsa04915;hsa04916;hsa04917;hsa04919;hsa04921;hsa04933;hsa04960;hsa05034;hsa05160;hsa05161;hsa05166;hsa05200;hsa05203;hsa05205;hsa05206;hsa05210;hsa05211;hsa05212;hsa05213;hsa05214;hsa05215;hsa05216;hsa05218;hsa05219;hsa05220;hsa05221;hsa05223;hsa05230;hsa05231 
                                               KRAS 
                                               NA 
                                               slateblue1 
                                               16 
                                               -9.25233644859813 
                                             
                                             
                                               PAK2-mRNA 
                                               -0.548 
                                               0.103 
                                               -0.75 
                                               -0.347 
                                               0.684 
                                               0.595 
                                               0.786 
                                               2.77e-05 
                                               0.00263 
                                               loglinear 
                                               Activation of Rac, CD209 (DC-SIGN) signaling, CD28 co-stimulation, CD28 dependent Vav1 pathway, Ephrin signaling, RHO GTPases activate PAKs, Sema3A PAK dependent Axon repulsion, VEGFA-VEGFR2 Pathway, VEGFR2 mediated vascular permeability 
                                               SEBELIUS_2_C4636 
                                               PAK2 
                                               mRNA 
                                               0 
                                               
                                               
                                               Nef and signal transduction;CD28 dependent Vav1 pathway;Stimulation of the cell death response by PAK-2p34;Ephrin signaling;Activation of Rac;RHO GTPases activate PAKs;VEGFR2 mediated vascular permeability;Sema3A PAK dependent Axon repulsion;CD209 (DC-SIGN) signaling;VEGFA-VEGFR2 Pathway;CD28 co-stimulation 
                                               hsa04010;hsa04012;hsa04014;hsa04360;hsa04510;hsa04660;hsa04810;hsa05211 
                                               PAK2 
                                               NA 
                                               slateblue1 
                                               16 
                                               -5.32038834951456 
                                             
                                             
                                               HRAS-mRNA 
                                               -0.573 
                                               0.152 
                                               -0.87 
                                               -0.276 
                                               0.672 
                                               0.547 
                                               0.826 
                                               0.0011 
                                               0.0583 
                                               loglinear 
                                               ARMS-mediated activation, CD209 (DC-SIGN) signaling, Cell surface interactions at the vascular wall, Constitutive Signaling by EGFRvIII, CREB phosphorylation through the activation of Ras, DAP12 signaling, Downstream signal transduction, FCERI mediated MAPK activation, Frs2-mediated activation, Gastrin-CREB signalling pathway via PKC and MAPK, Interleukin receptor SHC signaling, IRS-related events triggered by IGF1R, MAP2K and MAPK activation, RAFMAP kinase cascade, SHC-mediated cascade.CL.FGFR1, SHC-related events triggered by IGF1R, Signaling by ERBB2, Signaling by FGFR1 in disease, Signaling by FGFR2 in disease, Signaling by FGFR3 in disease, Signaling by FGFR4 in disease, Signaling by PDGF, Signaling by SCF-KIT, Tie2 Signaling, VEGFA-VEGFR2 Pathway, VEGFR2 mediated cell proliferation 
                                               SEBELIUS_2_C4636 
                                               HRAS 
                                               mRNA 
                                               0 
                                               
                                               
                                               Signaling by FGFR2 in disease;Regulation of RAS by GAPs;SHC-mediated cascade.CL.FGFR3;Signaling by FGFR3 in disease;MAP2K and MAPK activation;SHC-related events triggered by IGF1R;Signaling by FGFR4 in disease;SHC-mediated cascade.CL.FGFR1;Tie2 Signaling;Signaling by FGFR1 in disease;Activation of RAS in B cells;SHC-mediated cascade.CL.FGFR4;Constitutive Signaling by EGFRvIII;CD209 (DC-SIGN) signaling;Ras activation uopn Ca2.PLS. infux through NMDA receptor;FCERI mediated MAPK activation;RAFMAP kinase cascade;Downstream signal transduction;VEGFR2 mediated cell proliferation;Signaling by SCF-KIT;Frs2-mediated activation;ARMS-mediated activation;Gastrin-CREB signalling pathway via PKC and MAPK;Interleukin receptor SHC signaling;DAP12 signaling;Signaling by PDGF;VEGFA-VEGFR2 Pathway;Signaling by ERBB2;IRS-related events triggered by IGF1R;Cell surface interactions at the vascular wall;CREB phosphorylation through the activation of Ras 
                                               hsa04010;hsa04012;hsa04014;hsa04015;hsa04062;hsa04068;hsa04071;hsa04072;hsa04144;hsa04151;hsa04211;hsa04213;hsa04360;hsa04370;hsa04510;hsa04530;hsa04540;hsa04550;hsa04630;hsa04650;hsa04660;hsa04662;hsa04664;hsa04720;hsa04722;hsa04725;hsa04726;hsa04730;hsa04810;hsa04910;hsa04912;hsa04915;hsa04916;hsa04917;hsa04919;hsa04921;hsa04933;hsa05034;hsa05160;hsa05161;hsa05166;hsa05200;hsa05203;hsa05205;hsa05206;hsa05211;hsa05213;hsa05214;hsa05215;hsa05216;hsa05218;hsa05219;hsa05220;hsa05221;hsa05223;hsa05230;hsa05231 
                                               HRAS 
                                               NA 
                                               slateblue3 
                                               16 
                                               -3.76973684210526 
                                             
                                             
                                               NRAS-mRNA 
                                               0.17 
                                               0.119 
                                               -0.0629 
                                               0.403 
                                               1.13 
                                               0.957 
                                               1.32 
                                               0.167 
                                               1 
                                               lm.nb 
                                               ARMS-mediated activation, CD209 (DC-SIGN) signaling, Cell surface interactions at the vascular wall, Constitutive Signaling by EGFRvIII, DAP12 signaling, Downstream signal transduction, FCERI mediated MAPK activation, Frs2-mediated activation, Gastrin-CREB signalling pathway via PKC and MAPK, Interleukin receptor SHC signaling, IRS-related events triggered by IGF1R, MAP2K and MAPK activation, RAFMAP kinase cascade, SHC-mediated cascade.CL.FGFR1, SHC-related events triggered by IGF1R, Signaling by ERBB2, Signaling by FGFR1 in disease, Signaling by FGFR2 in disease, Signaling by FGFR3 in disease, Signaling by FGFR4 in disease, Signaling by PDGF, Signaling by SCF-KIT, Tie2 Signaling, VEGFA-VEGFR2 Pathway, VEGFR2 mediated cell proliferation 
                                               SEBELIUS_2_C4636 
                                               NRAS 
                                               mRNA 
                                               0 
                                               
                                               
                                               Signaling by FGFR2 in disease;Regulation of RAS by GAPs;SHC-mediated cascade.CL.FGFR3;Signaling by FGFR3 in disease;MAP2K and MAPK activation;SHC-related events triggered by IGF1R;Signaling by FGFR4 in disease;SHC-mediated cascade.CL.FGFR1;Tie2 Signaling;Signaling by FGFR1 in disease;Activation of RAS in B cells;SHC-mediated cascade.CL.FGFR4;Constitutive Signaling by EGFRvIII;CD209 (DC-SIGN) signaling;FCERI mediated MAPK activation;RAFMAP kinase cascade;Downstream signal transduction;VEGFR2 mediated cell proliferation;Signaling by SCF-KIT;Frs2-mediated activation;ARMS-mediated activation;Gastrin-CREB signalling pathway via PKC and MAPK;Interleukin receptor SHC signaling;DAP12 signaling;Signaling by PDGF;VEGFA-VEGFR2 Pathway;Signaling by ERBB2;IRS-related events triggered by IGF1R;Cell surface interactions at the vascular wall 
                                               hsa04010;hsa04012;hsa04014;hsa04015;hsa04062;hsa04068;hsa04071;hsa04072;hsa04151;hsa04211;hsa04213;hsa04360;hsa04370;hsa04530;hsa04540;hsa04550;hsa04650;hsa04660;hsa04662;hsa04664;hsa04720;hsa04722;hsa04725;hsa04726;hsa04730;hsa04810;hsa04910;hsa04912;hsa04915;hsa04916;hsa04917;hsa04919;hsa04921;hsa04933;hsa05034;hsa05160;hsa05161;hsa05166;hsa05200;hsa05203;hsa05205;hsa05206;hsa05211;hsa05213;hsa05214;hsa05215;hsa05216;hsa05218;hsa05219;hsa05220;hsa05221;hsa05223;hsa05230;hsa05231 
                                               NRAS 
                                               NA 
                                               azure4 
                                               1 
                                               1.42857142857143 
                                             
                                             
                                               EP300-mRNA 
                                               -0.117 
                                               0.112 
                                               -0.337 
                                               0.103 
                                               0.922 
                                               0.792 
                                               1.07 
                                               0.308 
                                               1 
                                               loglinear 
                                               CD209 (DC-SIGN) signaling, Circadian Clock 
                                               SEBELIUS_2_C4636 
                                               EP300 
                                               mRNA 
                                               0 
                                               
                                               
                                               LRR FLII-interacting protein 1 (LRRFIP1) activates type I IFN production;TRAF6 mediated IRF7 activation;Constitutive Signaling by NOTCH1 PEST Domain Mutants;Polo-like kinase mediated events;B-WICH complex positively regulates rRNA expression;CD209 (DC-SIGN) signaling;Circadian Clock 
                                               hsa04024;hsa04066;hsa04068;hsa04110;hsa04310;hsa04330;hsa04350;hsa04520;hsa04630;hsa04720;hsa04916;hsa04919;hsa04922;hsa05016;hsa05152;hsa05161;hsa05164;hsa05166;hsa05168;hsa05169;hsa05200;hsa05203;hsa05206;hsa05211;hsa05215 
                                               EP300 
                                               NA 
                                               azure4 
                                               1 
                                               -1.04464285714286 
                                             
                                             
                                               RAF1-mRNA 
                                               -0.0523 
                                               0.0768 
                                               -0.203 
                                               0.0982 
                                               0.964 
                                               0.869 
                                               1.07 
                                               0.503 
                                               1 
                                               loglinear 
                                               ARMS-mediated activation, CD209 (DC-SIGN) signaling, CREB phosphorylation through the activation of Ras, DAP12 signaling, Downstream signal transduction, FCERI mediated MAPK activation, Frs2-mediated activation, Gastrin-CREB signalling pathway via PKC and MAPK, Interleukin receptor SHC signaling, IRS-related events triggered by IGF1R, Negative feedback regulation of MAPK pathway, Negative regulation of MAPK pathway, RAF activation, RAFMAP kinase cascade, Signaling by ERBB2, Signaling by PDGF, Signaling by SCF-KIT, VEGFA-VEGFR2 Pathway, VEGFR2 mediated cell proliferation 
                                               SEBELIUS_2_C4636 
                                               RAF1 
                                               mRNA 
                                               0 
                                               
                                               
                                               RAF activation;Stimuli-sensing channels;CREB phosphorylation through the activation of Ras;Negative feedback regulation of MAPK pathway;CD209 (DC-SIGN) signaling;RAFMAP kinase cascade;Downstream signal transduction;VEGFR2 mediated cell proliferation;Signaling by SCF-KIT;Frs2-mediated activation;ARMS-mediated activation;Gastrin-CREB signalling pathway via PKC and MAPK;Interleukin receptor SHC signaling;FCERI mediated MAPK activation;DAP12 signaling;Signaling by PDGF;VEGFA-VEGFR2 Pathway;Signaling by ERBB2;IRS-related events triggered by IGF1R;Negative regulation of MAPK pathway 
                                               hsa04010;hsa04012;hsa04014;hsa04015;hsa04022;hsa04024;hsa04062;hsa04068;hsa04071;hsa04072;hsa04151;hsa04270;hsa04370;hsa04510;hsa04540;hsa04550;hsa04630;hsa04650;hsa04660;hsa04662;hsa04664;hsa04666;hsa04720;hsa04722;hsa04726;hsa04730;hsa04810;hsa04910;hsa04912;hsa04914;hsa04915;hsa04916;hsa04917;hsa04919;hsa04921;hsa05034;hsa05152;hsa05160;hsa05161;hsa05164;hsa05200;hsa05205;hsa05206;hsa05210;hsa05211;hsa05212;hsa05213;hsa05214;hsa05215;hsa05218;hsa05219;hsa05220;hsa05221;hsa05223;hsa05230;hsa05231 
                                               RAF1 
                                               NA 
                                               azure4 
                                               1 
                                               -0.680989583333333 
                                             
                                             
                                               PAK1-mRNA 
                                               -0.105 
                                               0.172 
                                               -0.441 
                                               0.231 
                                               0.93 
                                               0.736 
                                               1.17 
                                               0.547 
                                               1 
                                               lm.nb 
                                               Activation of Rac, CD209 (DC-SIGN) signaling, CD28 co-stimulation, CD28 dependent Vav1 pathway, EPHB-mediated forward signaling, Ephrin signaling, FCERI mediated MAPK activation, MAPK6MAPK4 signaling, Regulation of actin dynamics for phagocytic cup formation, RHO GTPases activate PAKs, Sema3A PAK dependent Axon repulsion, VEGFA-VEGFR2 Pathway, VEGFR2 mediated vascular permeability 
                                               SEBELIUS_2_C4636 
                                               PAK1 
                                               mRNA 
                                               0 
                                               
                                               
                                               EPHB-mediated forward signaling;Ephrin signaling;MAPK6MAPK4 signaling;DSCAM interactions;Activation of Rac;VEGFR2 mediated vascular permeability;RHO GTPases activate PAKs;Sema3A PAK dependent Axon repulsion;FCERI mediated MAPK activation;Regulation of actin dynamics for phagocytic cup formation;CD209 (DC-SIGN) signaling;CD28 dependent Vav1 pathway;VEGFA-VEGFR2 Pathway;CD28 co-stimulation 
                                               hsa04010;hsa04012;hsa04014;hsa04024;hsa04062;hsa04360;hsa04510;hsa04650;hsa04660;hsa04666;hsa04810;hsa05120;hsa05205;hsa05211 
                                               PAK1 
                                               NA 
                                               azure4 
                                               1 
                                               -0.61046511627907 
                                             
                                             
                                               CREBBP-mRNA 
                                               -0.0319 
                                               0.0849 
                                               -0.198 
                                               0.134 
                                               0.978 
                                               0.872 
                                               1.1 
                                               0.711 
                                               1 
                                               loglinear 
                                               CD209 (DC-SIGN) signaling, Circadian Clock 
                                               SEBELIUS_2_C4636 
                                               CREBBP 
                                               mRNA 
                                               0 
                                               
                                               
                                               YAP1- and WWTR1 (TAZ)-stimulated gene expression;Activation of gene expression by SREBF (SREBP);Transcriptional activation of mitochondrial biogenesis;Constitutive Signaling by NOTCH1 HD.PLS.PEST Domain Mutants;Constitutive Signaling by NOTCH1 PEST Domain Mutants;CD209 (DC-SIGN) signaling;Circadian Clock 
                                               hsa04024;hsa04066;hsa04068;hsa04110;hsa04310;hsa04330;hsa04350;hsa04520;hsa04630;hsa04720;hsa04916;hsa04919;hsa04922;hsa05016;hsa05152;hsa05161;hsa05164;hsa05166;hsa05168;hsa05169;hsa05200;hsa05203;hsa05206;hsa05211;hsa05215 
                                               CREBBP 
                                               NA 
                                               azure4 
                                               1 
                                               -0.375736160188457 
                                             
                                             
                                               FYN-mRNA 
                                               0.0321 
                                               0.116 
                                               -0.195 
                                               0.259 
                                               1.02 
                                               0.874 
                                               1.2 
                                               0.784 
                                               1 
                                               loglinear 
                                               ARMS-mediated activation, CD209 (DC-SIGN) signaling, CD28 co-stimulation, CD28 dependent PI3KAkt signaling, CD28 dependent Vav1 pathway, Cell surface interactions at the vascular wall, Constitutive Signaling by Aberrant PI3K in Cancer, CTLA4 inhibitory signaling, DAP12 signaling, Downstream signal transduction, EPH-ephrin mediated repulsion of cells, FCERI mediated MAPK activation, Frs2-mediated activation, GAB1 signalosome, Gastrin-CREB signalling pathway via PKC and MAPK, GPVI-mediated activation cascade, Interleukin receptor SHC signaling, IRS-related events triggered by IGF1R, Nephrin interactions, PI3KAKT activation, RAFMAP kinase cascade, Regulation of signaling by CBL, Role of LAT2NTALLAB on calcium mobilization, Sema3A PAK dependent Axon repulsion, Signaling by ERBB2, Signaling by PDGF, Signaling by SCF-KIT, VEGFA-VEGFR2 Pathway, VEGFR2 mediated cell proliferation 
                                               SEBELIUS_2_C4636 
                                               FYN 
                                               mRNA 
                                               0 
                                               
                                               
                                               EPH-ephrin mediated repulsion of cells;Regulation of signaling by CBL;FCGR activation;Antigen activates B Cell Receptor (BCR) leading to generation of second messengers;Dectin-2 family;RAFMAP kinase cascade;Constitutive Signaling by Aberrant PI3K in Cancer;CTLA4 inhibitory signaling;CD28 dependent PI3KAkt signaling;PECAM1 interactions;CD28 dependent Vav1 pathway;GPVI-mediated activation cascade;Sema3A PAK dependent Axon repulsion;Nephrin interactions;SEMA3A-Plexin repulsion signaling by inhibiting Integrin adhesion;CRMPs in Sema3A signaling;CD209 (DC-SIGN) signaling;Downstream signal transduction;GAB1 signalosome;Signaling by SCF-KIT;PI3KAKT activation;Role of LAT2NTALLAB on calcium mobilization;DAP12 signaling;Signaling by PDGF;Signaling by ERBB2;VEGFA-VEGFR2 Pathway;VEGFR2 mediated cell proliferation;Frs2-mediated activation;ARMS-mediated activation;Gastrin-CREB signalling pathway via PKC and MAPK;Interleukin receptor SHC signaling;FCERI mediated MAPK activation;IRS-related events triggered by IGF1R;CD28 co-stimulation;Cell surface interactions at the vascular wall 
                                               hsa04071;hsa04072;hsa04360;hsa04380;hsa04510;hsa04520;hsa04611;hsa04650;hsa04660;hsa04664;hsa04725;hsa05020;hsa05130;hsa05162;hsa05416 
                                               FYN 
                                               NA 
                                               azure4 
                                               1 
                                               0.276724137931034 
                                             
                                           
                                           DE Results - CD209 (DC-SIGN) signaling genes - New.Annotation: miR155 vs.cel67  Table displaying each sample's global significance scores and directed global significance scores as defined in the heatmaps above. The global significance score is calculated as the square root of the mean squared t-statistic for the genes in a gene set, with t-statistics coming from the linear regression underlying our differential expression analysis. The directed global significance score is calculated as the square root of the mean signed squared t-statistic for the genes in a gene set, with t-statistics coming from the linear regression underlying our differential expression analysis.   
                                      
                                       
                                  
                                
                                
                                  
                                       
                                      
                                           Volcano Plot: New.Annotation: miR18b vs.cel67   More Plot Information                Volcano Plot: New.Annotation: miR18b vs.cel67  Volcano plot displaying each gene's -log10(p-value) and log2 fold change for the selected covariate. Highly statistically significant genes fall at the top of the plot, and highly differentially expressed genes fall to either side. Genes within the selected gene set are highlighted in orange. Horizontal lines indicate various False Discovery Rate (FDR) thresholds.   
                                           DE Results - CD209 (DC-SIGN) signaling genes - New.Annotation: miR18b vs.cel67   More Plot Information      Download CSV Data     
                                           
                                             
                                             
                                           
                                           
                                             
                                               Probe Label 
                                               Log2 fold change 
                                               std error (log2) 
                                               Lower confidence limit (log2) 
                                               Upper confidence limit (log2) 
                                               Linear fold change 
                                               Lower confidence limit (linear) 
                                               Upper confidence limit (linear) 
                                               P-value 
                                               BY.p.value 
                                               method 
                                               Gene.sets 
                                               Codeset.Name 
                                               Probe.Label 
                                               Analyte.Type 
                                               Is.Control 
                                               Control.Type 
                                               Related.Probes 
                                               Probe.Annotation 
                                               KEGG.Pathways 
                                               Official.Gene.Name 
                                               Control.Conc 
                                               volcanocol 
                                               volcanopch 
                                               tstats.all 
                                             
                                           
                                           
                                             
                                               RAF1-mRNA 
                                               -0.772 
                                               0.0829 
                                               -0.934 
                                               -0.609 
                                               0.586 
                                               0.523 
                                               0.656 
                                               6.7e-09 
                                               1.78e-06 
                                               loglinear 
                                               ARMS-mediated activation, CD209 (DC-SIGN) signaling, CREB phosphorylation through the activation of Ras, DAP12 signaling, Downstream signal transduction, FCERI mediated MAPK activation, Frs2-mediated activation, Gastrin-CREB signalling pathway via PKC and MAPK, Interleukin receptor SHC signaling, IRS-related events triggered by IGF1R, Negative feedback regulation of MAPK pathway, Negative regulation of MAPK pathway, RAF activation, RAFMAP kinase cascade, Signaling by ERBB2, Signaling by PDGF, Signaling by SCF-KIT, VEGFA-VEGFR2 Pathway, VEGFR2 mediated cell proliferation 
                                               SEBELIUS_2_C4636 
                                               RAF1 
                                               mRNA 
                                               0 
                                               
                                               
                                               RAF activation;Stimuli-sensing channels;CREB phosphorylation through the activation of Ras;Negative feedback regulation of MAPK pathway;CD209 (DC-SIGN) signaling;RAFMAP kinase cascade;Downstream signal transduction;VEGFR2 mediated cell proliferation;Signaling by SCF-KIT;Frs2-mediated activation;ARMS-mediated activation;Gastrin-CREB signalling pathway via PKC and MAPK;Interleukin receptor SHC signaling;FCERI mediated MAPK activation;DAP12 signaling;Signaling by PDGF;VEGFA-VEGFR2 Pathway;Signaling by ERBB2;IRS-related events triggered by IGF1R;Negative regulation of MAPK pathway 
                                               hsa04010;hsa04012;hsa04014;hsa04015;hsa04022;hsa04024;hsa04062;hsa04068;hsa04071;hsa04072;hsa04151;hsa04270;hsa04370;hsa04510;hsa04540;hsa04550;hsa04630;hsa04650;hsa04660;hsa04662;hsa04664;hsa04666;hsa04720;hsa04722;hsa04726;hsa04730;hsa04810;hsa04910;hsa04912;hsa04914;hsa04915;hsa04916;hsa04917;hsa04919;hsa04921;hsa05034;hsa05152;hsa05160;hsa05161;hsa05164;hsa05200;hsa05205;hsa05206;hsa05210;hsa05211;hsa05212;hsa05213;hsa05214;hsa05215;hsa05218;hsa05219;hsa05220;hsa05221;hsa05223;hsa05230;hsa05231 
                                               RAF1 
                                               NA 
                                               slateblue1 
                                               16 
                                               -9.3124246079614 
                                             
                                             
                                               NRAS-mRNA 
                                               -0.806 
                                               0.13 
                                               -1.06 
                                               -0.551 
                                               0.572 
                                               0.479 
                                               0.683 
                                               3.82e-06 
                                               0.000286 
                                               lm.nb 
                                               ARMS-mediated activation, CD209 (DC-SIGN) signaling, Cell surface interactions at the vascular wall, Constitutive Signaling by EGFRvIII, DAP12 signaling, Downstream signal transduction, FCERI mediated MAPK activation, Frs2-mediated activation, Gastrin-CREB signalling pathway via PKC and MAPK, Interleukin receptor SHC signaling, IRS-related events triggered by IGF1R, MAP2K and MAPK activation, RAFMAP kinase cascade, SHC-mediated cascade.CL.FGFR1, SHC-related events triggered by IGF1R, Signaling by ERBB2, Signaling by FGFR1 in disease, Signaling by FGFR2 in disease, Signaling by FGFR3 in disease, Signaling by FGFR4 in disease, Signaling by PDGF, Signaling by SCF-KIT, Tie2 Signaling, VEGFA-VEGFR2 Pathway, VEGFR2 mediated cell proliferation 
                                               SEBELIUS_2_C4636 
                                               NRAS 
                                               mRNA 
                                               0 
                                               
                                               
                                               Signaling by FGFR2 in disease;Regulation of RAS by GAPs;SHC-mediated cascade.CL.FGFR3;Signaling by FGFR3 in disease;MAP2K and MAPK activation;SHC-related events triggered by IGF1R;Signaling by FGFR4 in disease;SHC-mediated cascade.CL.FGFR1;Tie2 Signaling;Signaling by FGFR1 in disease;Activation of RAS in B cells;SHC-mediated cascade.CL.FGFR4;Constitutive Signaling by EGFRvIII;CD209 (DC-SIGN) signaling;FCERI mediated MAPK activation;RAFMAP kinase cascade;Downstream signal transduction;VEGFR2 mediated cell proliferation;Signaling by SCF-KIT;Frs2-mediated activation;ARMS-mediated activation;Gastrin-CREB signalling pathway via PKC and MAPK;Interleukin receptor SHC signaling;DAP12 signaling;Signaling by PDGF;VEGFA-VEGFR2 Pathway;Signaling by ERBB2;IRS-related events triggered by IGF1R;Cell surface interactions at the vascular wall 
                                               hsa04010;hsa04012;hsa04014;hsa04015;hsa04062;hsa04068;hsa04071;hsa04072;hsa04151;hsa04211;hsa04213;hsa04360;hsa04370;hsa04530;hsa04540;hsa04550;hsa04650;hsa04660;hsa04662;hsa04664;hsa04720;hsa04722;hsa04725;hsa04726;hsa04730;hsa04810;hsa04910;hsa04912;hsa04915;hsa04916;hsa04917;hsa04919;hsa04921;hsa04933;hsa05034;hsa05160;hsa05161;hsa05166;hsa05200;hsa05203;hsa05205;hsa05206;hsa05211;hsa05213;hsa05214;hsa05215;hsa05216;hsa05218;hsa05219;hsa05220;hsa05221;hsa05223;hsa05230;hsa05231 
                                               NRAS 
                                               NA 
                                               slateblue1 
                                               16 
                                               -6.2 
                                             
                                             
                                               KRAS-mRNA 
                                               0.31 
                                               0.115 
                                               0.0834 
                                               0.536 
                                               1.24 
                                               1.06 
                                               1.45 
                                               0.0139 
                                               0.243 
                                               loglinear 
                                               ARMS-mediated activation, CD209 (DC-SIGN) signaling, Cell surface interactions at the vascular wall, Constitutive Signaling by EGFRvIII, DAP12 signaling, Downstream signal transduction, FCERI mediated MAPK activation, Frs2-mediated activation, Gastrin-CREB signalling pathway via PKC and MAPK, Interleukin receptor SHC signaling, IRS-related events triggered by IGF1R, MAP2K and MAPK activation, RAFMAP kinase cascade, SHC-mediated cascade.CL.FGFR1, SHC-related events triggered by IGF1R, Signaling by ERBB2, Signaling by FGFR1 in disease, Signaling by FGFR2 in disease, Signaling by FGFR3 in disease, Signaling by FGFR4 in disease, Signaling by PDGF, Signaling by SCF-KIT, Tie2 Signaling, VEGFA-VEGFR2 Pathway, VEGFR2 mediated cell proliferation 
                                               SEBELIUS_2_C4636 
                                               KRAS 
                                               mRNA 
                                               0 
                                               
                                               
                                               Signaling by FGFR2 in disease;Regulation of RAS by GAPs;SHC-mediated cascade.CL.FGFR3;Signaling by FGFR3 in disease;MAP2K and MAPK activation;SHC-related events triggered by IGF1R;Signaling by FGFR4 in disease;SHC-mediated cascade.CL.FGFR1;Tie2 Signaling;Signaling by FGFR1 in disease;Activation of RAS in B cells;SHC-mediated cascade.CL.FGFR4;Constitutive Signaling by EGFRvIII;CD209 (DC-SIGN) signaling;FCERI mediated MAPK activation;RAFMAP kinase cascade;Downstream signal transduction;VEGFR2 mediated cell proliferation;Signaling by SCF-KIT;Frs2-mediated activation;ARMS-mediated activation;Gastrin-CREB signalling pathway via PKC and MAPK;Interleukin receptor SHC signaling;DAP12 signaling;Signaling by PDGF;VEGFA-VEGFR2 Pathway;Signaling by ERBB2;IRS-related events triggered by IGF1R;Cell surface interactions at the vascular wall 
                                               hsa04010;hsa04012;hsa04014;hsa04015;hsa04062;hsa04068;hsa04071;hsa04072;hsa04151;hsa04211;hsa04213;hsa04320;hsa04360;hsa04370;hsa04530;hsa04540;hsa04550;hsa04650;hsa04660;hsa04662;hsa04664;hsa04720;hsa04722;hsa04725;hsa04726;hsa04730;hsa04810;hsa04910;hsa04912;hsa04914;hsa04915;hsa04916;hsa04917;hsa04919;hsa04921;hsa04933;hsa04960;hsa05034;hsa05160;hsa05161;hsa05166;hsa05200;hsa05203;hsa05205;hsa05206;hsa05210;hsa05211;hsa05212;hsa05213;hsa05214;hsa05215;hsa05216;hsa05218;hsa05219;hsa05220;hsa05221;hsa05223;hsa05230;hsa05231 
                                               KRAS 
                                               NA 
                                               slateblue4 
                                               16 
                                               2.69565217391304 
                                             
                                             
                                               PAK1-mRNA 
                                               -0.488 
                                               0.186 
                                               -0.853 
                                               -0.123 
                                               0.713 
                                               0.554 
                                               0.918 
                                               0.016 
                                               0.272 
                                               lm.nb 
                                               Activation of Rac, CD209 (DC-SIGN) signaling, CD28 co-stimulation, CD28 dependent Vav1 pathway, EPHB-mediated forward signaling, Ephrin signaling, FCERI mediated MAPK activation, MAPK6MAPK4 signaling, Regulation of actin dynamics for phagocytic cup formation, RHO GTPases activate PAKs, Sema3A PAK dependent Axon repulsion, VEGFA-VEGFR2 Pathway, VEGFR2 mediated vascular permeability 
                                               SEBELIUS_2_C4636 
                                               PAK1 
                                               mRNA 
                                               0 
                                               
                                               
                                               EPHB-mediated forward signaling;Ephrin signaling;MAPK6MAPK4 signaling;DSCAM interactions;Activation of Rac;VEGFR2 mediated vascular permeability;RHO GTPases activate PAKs;Sema3A PAK dependent Axon repulsion;FCERI mediated MAPK activation;Regulation of actin dynamics for phagocytic cup formation;CD209 (DC-SIGN) signaling;CD28 dependent Vav1 pathway;VEGFA-VEGFR2 Pathway;CD28 co-stimulation 
                                               hsa04010;hsa04012;hsa04014;hsa04024;hsa04062;hsa04360;hsa04510;hsa04650;hsa04660;hsa04666;hsa04810;hsa05120;hsa05205;hsa05211 
                                               PAK1 
                                               NA 
                                               slateblue4 
                                               16 
                                               -2.62365591397849 
                                             
                                             
                                               HRAS-mRNA 
                                               -0.373 
                                               0.164 
                                               -0.694 
                                               -0.0518 
                                               0.772 
                                               0.618 
                                               0.965 
                                               0.0334 
                                               0.522 
                                               loglinear 
                                               ARMS-mediated activation, CD209 (DC-SIGN) signaling, Cell surface interactions at the vascular wall, Constitutive Signaling by EGFRvIII, CREB phosphorylation through the activation of Ras, DAP12 signaling, Downstream signal transduction, FCERI mediated MAPK activation, Frs2-mediated activation, Gastrin-CREB signalling pathway via PKC and MAPK, Interleukin receptor SHC signaling, IRS-related events triggered by IGF1R, MAP2K and MAPK activation, RAFMAP kinase cascade, SHC-mediated cascade.CL.FGFR1, SHC-related events triggered by IGF1R, Signaling by ERBB2, Signaling by FGFR1 in disease, Signaling by FGFR2 in disease, Signaling by FGFR3 in disease, Signaling by FGFR4 in disease, Signaling by PDGF, Signaling by SCF-KIT, Tie2 Signaling, VEGFA-VEGFR2 Pathway, VEGFR2 mediated cell proliferation 
                                               SEBELIUS_2_C4636 
                                               HRAS 
                                               mRNA 
                                               0 
                                               
                                               
                                               Signaling by FGFR2 in disease;Regulation of RAS by GAPs;SHC-mediated cascade.CL.FGFR3;Signaling by FGFR3 in disease;MAP2K and MAPK activation;SHC-related events triggered by IGF1R;Signaling by FGFR4 in disease;SHC-mediated cascade.CL.FGFR1;Tie2 Signaling;Signaling by FGFR1 in disease;Activation of RAS in B cells;SHC-mediated cascade.CL.FGFR4;Constitutive Signaling by EGFRvIII;CD209 (DC-SIGN) signaling;Ras activation uopn Ca2.PLS. infux through NMDA receptor;FCERI mediated MAPK activation;RAFMAP kinase cascade;Downstream signal transduction;VEGFR2 mediated cell proliferation;Signaling by SCF-KIT;Frs2-mediated activation;ARMS-mediated activation;Gastrin-CREB signalling pathway via PKC and MAPK;Interleukin receptor SHC signaling;DAP12 signaling;Signaling by PDGF;VEGFA-VEGFR2 Pathway;Signaling by ERBB2;IRS-related events triggered by IGF1R;Cell surface interactions at the vascular wall;CREB phosphorylation through the activation of Ras 
                                               hsa04010;hsa04012;hsa04014;hsa04015;hsa04062;hsa04068;hsa04071;hsa04072;hsa04144;hsa04151;hsa04211;hsa04213;hsa04360;hsa04370;hsa04510;hsa04530;hsa04540;hsa04550;hsa04630;hsa04650;hsa04660;hsa04662;hsa04664;hsa04720;hsa04722;hsa04725;hsa04726;hsa04730;hsa04810;hsa04910;hsa04912;hsa04915;hsa04916;hsa04917;hsa04919;hsa04921;hsa04933;hsa05034;hsa05160;hsa05161;hsa05166;hsa05200;hsa05203;hsa05205;hsa05206;hsa05211;hsa05213;hsa05214;hsa05215;hsa05216;hsa05218;hsa05219;hsa05220;hsa05221;hsa05223;hsa05230;hsa05231 
                                               HRAS 
                                               NA 
                                               azure4 
                                               1 
                                               -2.27439024390244 
                                             
                                             
                                               EP300-mRNA 
                                               -0.24 
                                               0.121 
                                               -0.477 
                                               -0.00233 
                                               0.847 
                                               0.719 
                                               0.998 
                                               0.061 
                                               0.853 
                                               loglinear 
                                               CD209 (DC-SIGN) signaling, Circadian Clock 
                                               SEBELIUS_2_C4636 
                                               EP300 
                                               mRNA 
                                               0 
                                               
                                               
                                               LRR FLII-interacting protein 1 (LRRFIP1) activates type I IFN production;TRAF6 mediated IRF7 activation;Constitutive Signaling by NOTCH1 PEST Domain Mutants;Polo-like kinase mediated events;B-WICH complex positively regulates rRNA expression;CD209 (DC-SIGN) signaling;Circadian Clock 
                                               hsa04024;hsa04066;hsa04068;hsa04110;hsa04310;hsa04330;hsa04350;hsa04520;hsa04630;hsa04720;hsa04916;hsa04919;hsa04922;hsa05016;hsa05152;hsa05161;hsa05164;hsa05166;hsa05168;hsa05169;hsa05200;hsa05203;hsa05206;hsa05211;hsa05215 
                                               EP300 
                                               NA 
                                               azure4 
                                               1 
                                               -1.98347107438017 
                                             
                                             
                                               PAK2-mRNA 
                                               0.105 
                                               0.111 
                                               -0.112 
                                               0.323 
                                               1.08 
                                               0.925 
                                               1.25 
                                               0.354 
                                               1 
                                               loglinear 
                                               Activation of Rac, CD209 (DC-SIGN) signaling, CD28 co-stimulation, CD28 dependent Vav1 pathway, Ephrin signaling, RHO GTPases activate PAKs, Sema3A PAK dependent Axon repulsion, VEGFA-VEGFR2 Pathway, VEGFR2 mediated vascular permeability 
                                               SEBELIUS_2_C4636 
                                               PAK2 
                                               mRNA 
                                               0 
                                               
                                               
                                               Nef and signal transduction;CD28 dependent Vav1 pathway;Stimulation of the cell death response by PAK-2p34;Ephrin signaling;Activation of Rac;RHO GTPases activate PAKs;VEGFR2 mediated vascular permeability;Sema3A PAK dependent Axon repulsion;CD209 (DC-SIGN) signaling;VEGFA-VEGFR2 Pathway;CD28 co-stimulation 
                                               hsa04010;hsa04012;hsa04014;hsa04360;hsa04510;hsa04660;hsa04810;hsa05211 
                                               PAK2 
                                               NA 
                                               azure4 
                                               1 
                                               0.945945945945946 
                                             
                                             
                                               CREBBP-mRNA 
                                               0.0606 
                                               0.0917 
                                               -0.119 
                                               0.24 
                                               1.04 
                                               0.921 
                                               1.18 
                                               0.516 
                                               1 
                                               loglinear 
                                               CD209 (DC-SIGN) signaling, Circadian Clock 
                                               SEBELIUS_2_C4636 
                                               CREBBP 
                                               mRNA 
                                               0 
                                               
                                               
                                               YAP1- and WWTR1 (TAZ)-stimulated gene expression;Activation of gene expression by SREBF (SREBP);Transcriptional activation of mitochondrial biogenesis;Constitutive Signaling by NOTCH1 HD.PLS.PEST Domain Mutants;Constitutive Signaling by NOTCH1 PEST Domain Mutants;CD209 (DC-SIGN) signaling;Circadian Clock 
                                               hsa04024;hsa04066;hsa04068;hsa04110;hsa04310;hsa04330;hsa04350;hsa04520;hsa04630;hsa04720;hsa04916;hsa04919;hsa04922;hsa05016;hsa05152;hsa05161;hsa05164;hsa05166;hsa05168;hsa05169;hsa05200;hsa05203;hsa05206;hsa05211;hsa05215 
                                               CREBBP 
                                               NA 
                                               azure4 
                                               1 
                                               0.660850599781898 
                                             
                                             
                                               FYN-mRNA 
                                               -0.000154 
                                               0.125 
                                               -0.245 
                                               0.245 
                                               1 
                                               0.844 
                                               1.18 
                                               0.999 
                                               1 
                                               loglinear 
                                               ARMS-mediated activation, CD209 (DC-SIGN) signaling, CD28 co-stimulation, CD28 dependent PI3KAkt signaling, CD28 dependent Vav1 pathway, Cell surface interactions at the vascular wall, Constitutive Signaling by Aberrant PI3K in Cancer, CTLA4 inhibitory signaling, DAP12 signaling, Downstream signal transduction, EPH-ephrin mediated repulsion of cells, FCERI mediated MAPK activation, Frs2-mediated activation, GAB1 signalosome, Gastrin-CREB signalling pathway via PKC and MAPK, GPVI-mediated activation cascade, Interleukin receptor SHC signaling, IRS-related events triggered by IGF1R, Nephrin interactions, PI3KAKT activation, RAFMAP kinase cascade, Regulation of signaling by CBL, Role of LAT2NTALLAB on calcium mobilization, Sema3A PAK dependent Axon repulsion, Signaling by ERBB2, Signaling by PDGF, Signaling by SCF-KIT, VEGFA-VEGFR2 Pathway, VEGFR2 mediated cell proliferation 
                                               SEBELIUS_2_C4636 
                                               FYN 
                                               mRNA 
                                               0 
                                               
                                               
                                               EPH-ephrin mediated repulsion of cells;Regulation of signaling by CBL;FCGR activation;Antigen activates B Cell Receptor (BCR) leading to generation of second messengers;Dectin-2 family;RAFMAP kinase cascade;Constitutive Signaling by Aberrant PI3K in Cancer;CTLA4 inhibitory signaling;CD28 dependent PI3KAkt signaling;PECAM1 interactions;CD28 dependent Vav1 pathway;GPVI-mediated activation cascade;Sema3A PAK dependent Axon repulsion;Nephrin interactions;SEMA3A-Plexin repulsion signaling by inhibiting Integrin adhesion;CRMPs in Sema3A signaling;CD209 (DC-SIGN) signaling;Downstream signal transduction;GAB1 signalosome;Signaling by SCF-KIT;PI3KAKT activation;Role of LAT2NTALLAB on calcium mobilization;DAP12 signaling;Signaling by PDGF;Signaling by ERBB2;VEGFA-VEGFR2 Pathway;VEGFR2 mediated cell proliferation;Frs2-mediated activation;ARMS-mediated activation;Gastrin-CREB signalling pathway via PKC and MAPK;Interleukin receptor SHC signaling;FCERI mediated MAPK activation;IRS-related events triggered by IGF1R;CD28 co-stimulation;Cell surface interactions at the vascular wall 
                                               hsa04071;hsa04072;hsa04360;hsa04380;hsa04510;hsa04520;hsa04611;hsa04650;hsa04660;hsa04664;hsa04725;hsa05020;hsa05130;hsa05162;hsa05416 
                                               FYN 
                                               NA 
                                               azure4 
                                               1 
                                               -0.001232 
                                             
                                           
                                           DE Results - CD209 (DC-SIGN) signaling genes - New.Annotation: miR18b vs.cel67  Table displaying each sample's global significance scores and directed global significance scores as defined in the heatmaps above. The global significance score is calculated as the square root of the mean squared t-statistic for the genes in a gene set, with t-statistics coming from the linear regression underlying our differential expression analysis. The directed global significance score is calculated as the square root of the mean signed squared t-statistic for the genes in a gene set, with t-statistics coming from the linear regression underlying our differential expression analysis.   
                                      
                                       
                                  
                                
                                
                                  
                                       
                                      
                                           Volcano Plot: New.Annotation: miR890 vs.cel67   More Plot Information                Volcano Plot: New.Annotation: miR890 vs.cel67  Volcano plot displaying each gene's -log10(p-value) and log2 fold change for the selected covariate. Highly statistically significant genes fall at the top of the plot, and highly differentially expressed genes fall to either side. Genes within the selected gene set are highlighted in orange. Horizontal lines indicate various False Discovery Rate (FDR) thresholds.   
                                           DE Results - CD209 (DC-SIGN) signaling genes - New.Annotation: miR890 vs.cel67   More Plot Information      Download CSV Data     
                                           
                                             
                                             
                                           
                                           
                                             
                                               Probe Label 
                                               Log2 fold change 
                                               std error (log2) 
                                               Lower confidence limit (log2) 
                                               Upper confidence limit (log2) 
                                               Linear fold change 
                                               Lower confidence limit (linear) 
                                               Upper confidence limit (linear) 
                                               P-value 
                                               BY.p.value 
                                               method 
                                               Gene.sets 
                                               Codeset.Name 
                                               Probe.Label 
                                               Analyte.Type 
                                               Is.Control 
                                               Control.Type 
                                               Related.Probes 
                                               Probe.Annotation 
                                               KEGG.Pathways 
                                               Official.Gene.Name 
                                               Control.Conc 
                                               volcanocol 
                                               volcanopch 
                                               tstats.all 
                                             
                                           
                                           
                                             
                                               NRAS-mRNA 
                                               -0.783 
                                               0.109 
                                               -0.997 
                                               -0.568 
                                               0.581 
                                               0.501 
                                               0.674 
                                               4.76e-07 
                                               5.27e-05 
                                               lm.nb 
                                               ARMS-mediated activation, CD209 (DC-SIGN) signaling, Cell surface interactions at the vascular wall, Constitutive Signaling by EGFRvIII, DAP12 signaling, Downstream signal transduction, FCERI mediated MAPK activation, Frs2-mediated activation, Gastrin-CREB signalling pathway via PKC and MAPK, Interleukin receptor SHC signaling, IRS-related events triggered by IGF1R, MAP2K and MAPK activation, RAFMAP kinase cascade, SHC-mediated cascade.CL.FGFR1, SHC-related events triggered by IGF1R, Signaling by ERBB2, Signaling by FGFR1 in disease, Signaling by FGFR2 in disease, Signaling by FGFR3 in disease, Signaling by FGFR4 in disease, Signaling by PDGF, Signaling by SCF-KIT, Tie2 Signaling, VEGFA-VEGFR2 Pathway, VEGFR2 mediated cell proliferation 
                                               SEBELIUS_2_C4636 
                                               NRAS 
                                               mRNA 
                                               0 
                                               
                                               
                                               Signaling by FGFR2 in disease;Regulation of RAS by GAPs;SHC-mediated cascade.CL.FGFR3;Signaling by FGFR3 in disease;MAP2K and MAPK activation;SHC-related events triggered by IGF1R;Signaling by FGFR4 in disease;SHC-mediated cascade.CL.FGFR1;Tie2 Signaling;Signaling by FGFR1 in disease;Activation of RAS in B cells;SHC-mediated cascade.CL.FGFR4;Constitutive Signaling by EGFRvIII;CD209 (DC-SIGN) signaling;FCERI mediated MAPK activation;RAFMAP kinase cascade;Downstream signal transduction;VEGFR2 mediated cell proliferation;Signaling by SCF-KIT;Frs2-mediated activation;ARMS-mediated activation;Gastrin-CREB signalling pathway via PKC and MAPK;Interleukin receptor SHC signaling;DAP12 signaling;Signaling by PDGF;VEGFA-VEGFR2 Pathway;Signaling by ERBB2;IRS-related events triggered by IGF1R;Cell surface interactions at the vascular wall 
                                               hsa04010;hsa04012;hsa04014;hsa04015;hsa04062;hsa04068;hsa04071;hsa04072;hsa04151;hsa04211;hsa04213;hsa04360;hsa04370;hsa04530;hsa04540;hsa04550;hsa04650;hsa04660;hsa04662;hsa04664;hsa04720;hsa04722;hsa04725;hsa04726;hsa04730;hsa04810;hsa04910;hsa04912;hsa04915;hsa04916;hsa04917;hsa04919;hsa04921;hsa04933;hsa05034;hsa05160;hsa05161;hsa05166;hsa05200;hsa05203;hsa05205;hsa05206;hsa05211;hsa05213;hsa05214;hsa05215;hsa05216;hsa05218;hsa05219;hsa05220;hsa05221;hsa05223;hsa05230;hsa05231 
                                               NRAS 
                                               NA 
                                               slateblue1 
                                               16 
                                               -7.18348623853211 
                                             
                                             
                                               PAK2-mRNA 
                                               -0.478 
                                               0.0939 
                                               -0.662 
                                               -0.293 
                                               0.718 
                                               0.632 
                                               0.816 
                                               4.92e-05 
                                               0.00233 
                                               loglinear 
                                               Activation of Rac, CD209 (DC-SIGN) signaling, CD28 co-stimulation, CD28 dependent Vav1 pathway, Ephrin signaling, RHO GTPases activate PAKs, Sema3A PAK dependent Axon repulsion, VEGFA-VEGFR2 Pathway, VEGFR2 mediated vascular permeability 
                                               SEBELIUS_2_C4636 
                                               PAK2 
                                               mRNA 
                                               0 
                                               
                                               
                                               Nef and signal transduction;CD28 dependent Vav1 pathway;Stimulation of the cell death response by PAK-2p34;Ephrin signaling;Activation of Rac;RHO GTPases activate PAKs;VEGFR2 mediated vascular permeability;Sema3A PAK dependent Axon repulsion;CD209 (DC-SIGN) signaling;VEGFA-VEGFR2 Pathway;CD28 co-stimulation 
                                               hsa04010;hsa04012;hsa04014;hsa04360;hsa04510;hsa04660;hsa04810;hsa05211 
                                               PAK2 
                                               NA 
                                               slateblue1 
                                               16 
                                               -5.09052183173589 
                                             
                                             
                                               CREBBP-mRNA 
                                               -0.255 
                                               0.0775 
                                               -0.407 
                                               -0.103 
                                               0.838 
                                               0.754 
                                               0.931 
                                               0.00351 
                                               0.0801 
                                               loglinear 
                                               CD209 (DC-SIGN) signaling, Circadian Clock 
                                               SEBELIUS_2_C4636 
                                               CREBBP 
                                               mRNA 
                                               0 
                                               
                                               
                                               YAP1- and WWTR1 (TAZ)-stimulated gene expression;Activation of gene expression by SREBF (SREBP);Transcriptional activation of mitochondrial biogenesis;Constitutive Signaling by NOTCH1 HD.PLS.PEST Domain Mutants;Constitutive Signaling by NOTCH1 PEST Domain Mutants;CD209 (DC-SIGN) signaling;Circadian Clock 
                                               hsa04024;hsa04066;hsa04068;hsa04110;hsa04310;hsa04330;hsa04350;hsa04520;hsa04630;hsa04720;hsa04916;hsa04919;hsa04922;hsa05016;hsa05152;hsa05161;hsa05164;hsa05166;hsa05168;hsa05169;hsa05200;hsa05203;hsa05206;hsa05211;hsa05215 
                                               CREBBP 
                                               NA 
                                               slateblue3 
                                               16 
                                               -3.29032258064516 
                                             
                                             
                                               HRAS-mRNA 
                                               -0.435 
                                               0.138 
                                               -0.706 
                                               -0.163 
                                               0.74 
                                               0.613 
                                               0.893 
                                               0.00495 
                                               0.104 
                                               loglinear 
                                               ARMS-mediated activation, CD209 (DC-SIGN) signaling, Cell surface interactions at the vascular wall, Constitutive Signaling by EGFRvIII, CREB phosphorylation through the activation of Ras, DAP12 signaling, Downstream signal transduction, FCERI mediated MAPK activation, Frs2-mediated activation, Gastrin-CREB signalling pathway via PKC and MAPK, Interleukin receptor SHC signaling, IRS-related events triggered by IGF1R, MAP2K and MAPK activation, RAFMAP kinase cascade, SHC-mediated cascade.CL.FGFR1, SHC-related events triggered by IGF1R, Signaling by ERBB2, Signaling by FGFR1 in disease, Signaling by FGFR2 in disease, Signaling by FGFR3 in disease, Signaling by FGFR4 in disease, Signaling by PDGF, Signaling by SCF-KIT, Tie2 Signaling, VEGFA-VEGFR2 Pathway, VEGFR2 mediated cell proliferation 
                                               SEBELIUS_2_C4636 
                                               HRAS 
                                               mRNA 
                                               0 
                                               
                                               
                                               Signaling by FGFR2 in disease;Regulation of RAS by GAPs;SHC-mediated cascade.CL.FGFR3;Signaling by FGFR3 in disease;MAP2K and MAPK activation;SHC-related events triggered by IGF1R;Signaling by FGFR4 in disease;SHC-mediated cascade.CL.FGFR1;Tie2 Signaling;Signaling by FGFR1 in disease;Activation of RAS in B cells;SHC-mediated cascade.CL.FGFR4;Constitutive Signaling by EGFRvIII;CD209 (DC-SIGN) signaling;Ras activation uopn Ca2.PLS. infux through NMDA receptor;FCERI mediated MAPK activation;RAFMAP kinase cascade;Downstream signal transduction;VEGFR2 mediated cell proliferation;Signaling by SCF-KIT;Frs2-mediated activation;ARMS-mediated activation;Gastrin-CREB signalling pathway via PKC and MAPK;Interleukin receptor SHC signaling;DAP12 signaling;Signaling by PDGF;VEGFA-VEGFR2 Pathway;Signaling by ERBB2;IRS-related events triggered by IGF1R;Cell surface interactions at the vascular wall;CREB phosphorylation through the activation of Ras 
                                               hsa04010;hsa04012;hsa04014;hsa04015;hsa04062;hsa04068;hsa04071;hsa04072;hsa04144;hsa04151;hsa04211;hsa04213;hsa04360;hsa04370;hsa04510;hsa04530;hsa04540;hsa04550;hsa04630;hsa04650;hsa04660;hsa04662;hsa04664;hsa04720;hsa04722;hsa04725;hsa04726;hsa04730;hsa04810;hsa04910;hsa04912;hsa04915;hsa04916;hsa04917;hsa04919;hsa04921;hsa04933;hsa05034;hsa05160;hsa05161;hsa05166;hsa05200;hsa05203;hsa05205;hsa05206;hsa05211;hsa05213;hsa05214;hsa05215;hsa05216;hsa05218;hsa05219;hsa05220;hsa05221;hsa05223;hsa05230;hsa05231 
                                               HRAS 
                                               NA 
                                               slateblue4 
                                               16 
                                               -3.15217391304348 
                                             
                                             
                                               RAF1-mRNA 
                                               -0.213 
                                               0.0701 
                                               -0.351 
                                               -0.0761 
                                               0.862 
                                               0.784 
                                               0.949 
                                               0.00614 
                                               0.125 
                                               loglinear 
                                               ARMS-mediated activation, CD209 (DC-SIGN) signaling, CREB phosphorylation through the activation of Ras, DAP12 signaling, Downstream signal transduction, FCERI mediated MAPK activation, Frs2-mediated activation, Gastrin-CREB signalling pathway via PKC and MAPK, Interleukin receptor SHC signaling, IRS-related events triggered by IGF1R, Negative feedback regulation of MAPK pathway, Negative regulation of MAPK pathway, RAF activation, RAFMAP kinase cascade, Signaling by ERBB2, Signaling by PDGF, Signaling by SCF-KIT, VEGFA-VEGFR2 Pathway, VEGFR2 mediated cell proliferation 
                                               SEBELIUS_2_C4636 
                                               RAF1 
                                               mRNA 
                                               0 
                                               
                                               
                                               RAF activation;Stimuli-sensing channels;CREB phosphorylation through the activation of Ras;Negative feedback regulation of MAPK pathway;CD209 (DC-SIGN) signaling;RAFMAP kinase cascade;Downstream signal transduction;VEGFR2 mediated cell proliferation;Signaling by SCF-KIT;Frs2-mediated activation;ARMS-mediated activation;Gastrin-CREB signalling pathway via PKC and MAPK;Interleukin receptor SHC signaling;FCERI mediated MAPK activation;DAP12 signaling;Signaling by PDGF;VEGFA-VEGFR2 Pathway;Signaling by ERBB2;IRS-related events triggered by IGF1R;Negative regulation of MAPK pathway 
                                               hsa04010;hsa04012;hsa04014;hsa04015;hsa04022;hsa04024;hsa04062;hsa04068;hsa04071;hsa04072;hsa04151;hsa04270;hsa04370;hsa04510;hsa04540;hsa04550;hsa04630;hsa04650;hsa04660;hsa04662;hsa04664;hsa04666;hsa04720;hsa04722;hsa04726;hsa04730;hsa04810;hsa04910;hsa04912;hsa04914;hsa04915;hsa04916;hsa04917;hsa04919;hsa04921;hsa05034;hsa05152;hsa05160;hsa05161;hsa05164;hsa05200;hsa05205;hsa05206;hsa05210;hsa05211;hsa05212;hsa05213;hsa05214;hsa05215;hsa05218;hsa05219;hsa05220;hsa05221;hsa05223;hsa05230;hsa05231 
                                               RAF1 
                                               NA 
                                               slateblue4 
                                               16 
                                               -3.03851640513552 
                                             
                                             
                                               EP300-mRNA 
                                               -0.264 
                                               0.102 
                                               -0.465 
                                               -0.0636 
                                               0.833 
                                               0.725 
                                               0.957 
                                               0.0174 
                                               0.293 
                                               loglinear 
                                               CD209 (DC-SIGN) signaling, Circadian Clock 
                                               SEBELIUS_2_C4636 
                                               EP300 
                                               mRNA 
                                               0 
                                               
                                               
                                               LRR FLII-interacting protein 1 (LRRFIP1) activates type I IFN production;TRAF6 mediated IRF7 activation;Constitutive Signaling by NOTCH1 PEST Domain Mutants;Polo-like kinase mediated events;B-WICH complex positively regulates rRNA expression;CD209 (DC-SIGN) signaling;Circadian Clock 
                                               hsa04024;hsa04066;hsa04068;hsa04110;hsa04310;hsa04330;hsa04350;hsa04520;hsa04630;hsa04720;hsa04916;hsa04919;hsa04922;hsa05016;hsa05152;hsa05161;hsa05164;hsa05166;hsa05168;hsa05169;hsa05200;hsa05203;hsa05206;hsa05211;hsa05215 
                                               EP300 
                                               NA 
                                               slateblue4 
                                               16 
                                               -2.58823529411765 
                                             
                                             
                                               KRAS-mRNA 
                                               0.143 
                                               0.0975 
                                               -0.0486 
                                               0.334 
                                               1.1 
                                               0.967 
                                               1.26 
                                               0.159 
                                               1 
                                               loglinear 
                                               ARMS-mediated activation, CD209 (DC-SIGN) signaling, Cell surface interactions at the vascular wall, Constitutive Signaling by EGFRvIII, DAP12 signaling, Downstream signal transduction, FCERI mediated MAPK activation, Frs2-mediated activation, Gastrin-CREB signalling pathway via PKC and MAPK, Interleukin receptor SHC signaling, IRS-related events triggered by IGF1R, MAP2K and MAPK activation, RAFMAP kinase cascade, SHC-mediated cascade.CL.FGFR1, SHC-related events triggered by IGF1R, Signaling by ERBB2, Signaling by FGFR1 in disease, Signaling by FGFR2 in disease, Signaling by FGFR3 in disease, Signaling by FGFR4 in disease, Signaling by PDGF, Signaling by SCF-KIT, Tie2 Signaling, VEGFA-VEGFR2 Pathway, VEGFR2 mediated cell proliferation 
                                               SEBELIUS_2_C4636 
                                               KRAS 
                                               mRNA 
                                               0 
                                               
                                               
                                               Signaling by FGFR2 in disease;Regulation of RAS by GAPs;SHC-mediated cascade.CL.FGFR3;Signaling by FGFR3 in disease;MAP2K and MAPK activation;SHC-related events triggered by IGF1R;Signaling by FGFR4 in disease;SHC-mediated cascade.CL.FGFR1;Tie2 Signaling;Signaling by FGFR1 in disease;Activation of RAS in B cells;SHC-mediated cascade.CL.FGFR4;Constitutive Signaling by EGFRvIII;CD209 (DC-SIGN) signaling;FCERI mediated MAPK activation;RAFMAP kinase cascade;Downstream signal transduction;VEGFR2 mediated cell proliferation;Signaling by SCF-KIT;Frs2-mediated activation;ARMS-mediated activation;Gastrin-CREB signalling pathway via PKC and MAPK;Interleukin receptor SHC signaling;DAP12 signaling;Signaling by PDGF;VEGFA-VEGFR2 Pathway;Signaling by ERBB2;IRS-related events triggered by IGF1R;Cell surface interactions at the vascular wall 
                                               hsa04010;hsa04012;hsa04014;hsa04015;hsa04062;hsa04068;hsa04071;hsa04072;hsa04151;hsa04211;hsa04213;hsa04320;hsa04360;hsa04370;hsa04530;hsa04540;hsa04550;hsa04650;hsa04660;hsa04662;hsa04664;hsa04720;hsa04722;hsa04725;hsa04726;hsa04730;hsa04810;hsa04910;hsa04912;hsa04914;hsa04915;hsa04916;hsa04917;hsa04919;hsa04921;hsa04933;hsa04960;hsa05034;hsa05160;hsa05161;hsa05166;hsa05200;hsa05203;hsa05205;hsa05206;hsa05210;hsa05211;hsa05212;hsa05213;hsa05214;hsa05215;hsa05216;hsa05218;hsa05219;hsa05220;hsa05221;hsa05223;hsa05230;hsa05231 
                                               KRAS 
                                               NA 
                                               azure4 
                                               1 
                                               1.46666666666667 
                                             
                                             
                                               PAK1-mRNA 
                                               -0.16 
                                               0.157 
                                               -0.467 
                                               0.147 
                                               0.895 
                                               0.723 
                                               1.11 
                                               0.318 
                                               1 
                                               lm.nb 
                                               Activation of Rac, CD209 (DC-SIGN) signaling, CD28 co-stimulation, CD28 dependent Vav1 pathway, EPHB-mediated forward signaling, Ephrin signaling, FCERI mediated MAPK activation, MAPK6MAPK4 signaling, Regulation of actin dynamics for phagocytic cup formation, RHO GTPases activate PAKs, Sema3A PAK dependent Axon repulsion, VEGFA-VEGFR2 Pathway, VEGFR2 mediated vascular permeability 
                                               SEBELIUS_2_C4636 
                                               PAK1 
                                               mRNA 
                                               0 
                                               
                                               
                                               EPHB-mediated forward signaling;Ephrin signaling;MAPK6MAPK4 signaling;DSCAM interactions;Activation of Rac;VEGFR2 mediated vascular permeability;RHO GTPases activate PAKs;Sema3A PAK dependent Axon repulsion;FCERI mediated MAPK activation;Regulation of actin dynamics for phagocytic cup formation;CD209 (DC-SIGN) signaling;CD28 dependent Vav1 pathway;VEGFA-VEGFR2 Pathway;CD28 co-stimulation 
                                               hsa04010;hsa04012;hsa04014;hsa04024;hsa04062;hsa04360;hsa04510;hsa04650;hsa04660;hsa04666;hsa04810;hsa05120;hsa05205;hsa05211 
                                               PAK1 
                                               NA 
                                               azure4 
                                               1 
                                               -1.01910828025478 
                                             
                                             
                                               FYN-mRNA 
                                               -0.0678 
                                               0.106 
                                               -0.275 
                                               0.139 
                                               0.954 
                                               0.827 
                                               1.1 
                                               0.528 
                                               1 
                                               loglinear 
                                               ARMS-mediated activation, CD209 (DC-SIGN) signaling, CD28 co-stimulation, CD28 dependent PI3KAkt signaling, CD28 dependent Vav1 pathway, Cell surface interactions at the vascular wall, Constitutive Signaling by Aberrant PI3K in Cancer, CTLA4 inhibitory signaling, DAP12 signaling, Downstream signal transduction, EPH-ephrin mediated repulsion of cells, FCERI mediated MAPK activation, Frs2-mediated activation, GAB1 signalosome, Gastrin-CREB signalling pathway via PKC and MAPK, GPVI-mediated activation cascade, Interleukin receptor SHC signaling, IRS-related events triggered by IGF1R, Nephrin interactions, PI3KAKT activation, RAFMAP kinase cascade, Regulation of signaling by CBL, Role of LAT2NTALLAB on calcium mobilization, Sema3A PAK dependent Axon repulsion, Signaling by ERBB2, Signaling by PDGF, Signaling by SCF-KIT, VEGFA-VEGFR2 Pathway, VEGFR2 mediated cell proliferation 
                                               SEBELIUS_2_C4636 
                                               FYN 
                                               mRNA 
                                               0 
                                               
                                               
                                               EPH-ephrin mediated repulsion of cells;Regulation of signaling by CBL;FCGR activation;Antigen activates B Cell Receptor (BCR) leading to generation of second messengers;Dectin-2 family;RAFMAP kinase cascade;Constitutive Signaling by Aberrant PI3K in Cancer;CTLA4 inhibitory signaling;CD28 dependent PI3KAkt signaling;PECAM1 interactions;CD28 dependent Vav1 pathway;GPVI-mediated activation cascade;Sema3A PAK dependent Axon repulsion;Nephrin interactions;SEMA3A-Plexin repulsion signaling by inhibiting Integrin adhesion;CRMPs in Sema3A signaling;CD209 (DC-SIGN) signaling;Downstream signal transduction;GAB1 signalosome;Signaling by SCF-KIT;PI3KAKT activation;Role of LAT2NTALLAB on calcium mobilization;DAP12 signaling;Signaling by PDGF;Signaling by ERBB2;VEGFA-VEGFR2 Pathway;VEGFR2 mediated cell proliferation;Frs2-mediated activation;ARMS-mediated activation;Gastrin-CREB signalling pathway via PKC and MAPK;Interleukin receptor SHC signaling;FCERI mediated MAPK activation;IRS-related events triggered by IGF1R;CD28 co-stimulation;Cell surface interactions at the vascular wall 
                                               hsa04071;hsa04072;hsa04360;hsa04380;hsa04510;hsa04520;hsa04611;hsa04650;hsa04660;hsa04664;hsa04725;hsa05020;hsa05130;hsa05162;hsa05416 
                                               FYN 
                                               NA 
                                               azure4 
                                               1 
                                               -0.639622641509434 
                                             
                                           
                                           DE Results - CD209 (DC-SIGN) signaling genes - New.Annotation: miR890 vs.cel67  Table displaying each sample's global significance scores and directed global significance scores as defined in the heatmaps above. The global significance score is calculated as the square root of the mean squared t-statistic for the genes in a gene set, with t-statistics coming from the linear regression underlying our differential expression analysis. The directed global significance score is calculated as the square root of the mean signed squared t-statistic for the genes in a gene set, with t-statistics coming from the linear regression underlying our differential expression analysis.   
                                      
                                       
                                  
                                
                              
                             
                          
                           
                      
                    
                    
                      
                           
                          
                              New.Annotation: 
                              
                                  New.Annotation: differential expression in Control vs. baseline of cel67  
                                  New.Annotation: differential expression in miR142 vs. baseline of cel67  
                                  New.Annotation: differential expression in miR155 vs. baseline of cel67  
                                  New.Annotation: differential expression in miR18b vs. baseline of cel67  
                                  New.Annotation: differential expression in miR890 vs. baseline of cel67  
                              
                              
                              
                                
                                  
                                       
                                      
                                           Volcano Plot: New.Annotation: Control vs.cel67   More Plot Information                Volcano Plot: New.Annotation: Control vs.cel67  Volcano plot displaying each gene's -log10(p-value) and log2 fold change for the selected covariate. Highly statistically significant genes fall at the top of the plot, and highly differentially expressed genes fall to either side. Genes within the selected gene set are highlighted in orange. Horizontal lines indicate various False Discovery Rate (FDR) thresholds.   
                                           DE Results - CD28 co-stimulation genes - New.Annotation: Control vs.cel67   More Plot Information      Download CSV Data     
                                           
                                             
                                             
                                           
                                           
                                             
                                               Probe Label 
                                               Log2 fold change 
                                               std error (log2) 
                                               Lower confidence limit (log2) 
                                               Upper confidence limit (log2) 
                                               Linear fold change 
                                               Lower confidence limit (linear) 
                                               Upper confidence limit (linear) 
                                               P-value 
                                               BY.p.value 
                                               method 
                                               Gene.sets 
                                               Codeset.Name 
                                               Probe.Label 
                                               Analyte.Type 
                                               Is.Control 
                                               Control.Type 
                                               Related.Probes 
                                               Probe.Annotation 
                                               KEGG.Pathways 
                                               Official.Gene.Name 
                                               Control.Conc 
                                               volcanocol 
                                               volcanopch 
                                               tstats.all 
                                             
                                           
                                           
                                             
                                               AKT2-mRNA 
                                               -0.182 
                                               0.0603 
                                               -0.3 
                                               -0.0634 
                                               0.882 
                                               0.812 
                                               0.957 
                                               0.00666 
                                               0.468 
                                               loglinear 
                                               Activation of BAD and translocation to mitochondria , CD28 co-stimulation, DAP12 signaling, Deactivation of the beta-catenin transactivating complex, Downstream signal transduction, G beta.CL.gamma signalling through PI3Kgamma, GAB1 signalosome, GPVI-mediated activation cascade, IRS-related events triggered by IGF1R, PI3K Cascade, PI3KAKT activation, PIP3 activates AKT signaling, Role of LAT2NTALLAB on calcium mobilization, Signaling by ERBB2, Signaling by PDGF, Signaling by SCF-KIT, TP53 Regulates Metabolic Genes, VEGFA-VEGFR2 Pathway, VEGFR2 mediated vascular permeability 
                                               SEBELIUS_2_C4636 
                                               AKT2 
                                               mRNA 
                                               0 
                                               
                                               
                                               TP53 Regulates Metabolic Genes;PDE3B signalling;AKT phosphorylates targets in the cytosol;Negative regulation of the PI3KAKT network;Deactivation of the beta-catenin transactivating complex;Activation of BAD and translocation to mitochondria ;G beta.CL.gamma signalling through PI3Kgamma;AKT phosphorylates targets in the nucleus;VEGFR2 mediated vascular permeability;GPVI-mediated activation cascade;Signaling by ERBB2;PI3K Cascade;IRS-related events triggered by IGF1R;mTOR signalling;PIP3 activates AKT signaling;Downstream signal transduction;GAB1 signalosome;Signaling by SCF-KIT;PI3KAKT activation;Role of LAT2NTALLAB on calcium mobilization;DAP12 signaling;Signaling by PDGF;CD28 co-stimulation;VEGFA-VEGFR2 Pathway 
                                               hsa04010;hsa04012;hsa04014;hsa04015;hsa04022;hsa04024;hsa04062;hsa04066;hsa04068;hsa04071;hsa04072;hsa04150;hsa04151;hsa04152;hsa04210;hsa04211;hsa04213;hsa04261;hsa04370;hsa04380;hsa04510;hsa04530;hsa04550;hsa04611;hsa04620;hsa04630;hsa04660;hsa04662;hsa04664;hsa04666;hsa04668;hsa04722;hsa04725;hsa04728;hsa04910;hsa04914;hsa04915;hsa04917;hsa04919;hsa04920;hsa04922;hsa04923;hsa04931;hsa04932;hsa04933;hsa04973;hsa05142;hsa05145;hsa05152;hsa05160;hsa05161;hsa05162;hsa05164;hsa05166;hsa05169;hsa05200;hsa05205;hsa05210;hsa05211;hsa05212;hsa05213;hsa05214;hsa05215;hsa05218;hsa05220;hsa05221;hsa05222;hsa05223;hsa05230;hsa05231 
                                               AKT2 
                                               NA 
                                               slateblue4 
                                               16 
                                               -3.01824212271973 
                                             
                                             
                                               PIK3R2-mRNA 
                                               -0.355 
                                               0.132 
                                               -0.615 
                                               -0.096 
                                               0.782 
                                               0.653 
                                               0.936 
                                               0.0139 
                                               0.682 
                                               lm.nb 
                                               CD28 co-stimulation, CD28 dependent PI3KAkt signaling, Cell surface interactions at the vascular wall, Constitutive Signaling by Aberrant PI3K in Cancer, DAP12 signaling, Downstream signal transduction, Downstream TCR signaling, G alpha (1213) signalling events, G alpha (q) signalling events, G beta.CL.gamma signalling through PI3Kgamma, GAB1 signalosome, Gastrin-CREB signalling pathway via PKC and MAPK, GPVI-mediated activation cascade, Interleukin receptor SHC signaling, IRS-related events triggered by IGF1R, Nephrin interactions, PI3K Cascade, PI3KAKT activation, PIP3 activates AKT signaling, Regulation of signaling by CBL, Rho GTPase cycle, Role of LAT2NTALLAB on calcium mobilization, Signaling by ERBB2, Signaling by PDGF, Signaling by SCF-KIT, Synthesis of PIPs at the plasma membrane, Tie2 Signaling, VEGFA-VEGFR2 Pathway 
                                               SEBELIUS_2_C4636 
                                               PIK3R2 
                                               mRNA 
                                               0 
                                               
                                               
                                               G alpha (q) signalling events;Downstream TCR signaling;PIP3 activates AKT signaling;VEGFA-VEGFR2 Pathway;Synthesis of PIPs at the plasma membrane;Rho GTPase cycle;Interleukin receptor SHC signaling;G beta.CL.gamma signalling through PI3Kgamma;Regulation of signaling by CBL;Role of phospholipids in phagocytosis;G alpha (1213) signalling events;PI3K Cascade;Constitutive Signaling by Aberrant PI3K in Cancer;CD28 dependent PI3KAkt signaling;Tie2 Signaling;GPVI-mediated activation cascade;Nephrin interactions;Gastrin-CREB signalling pathway via PKC and MAPK;Downstream signal transduction;GAB1 signalosome;Signaling by SCF-KIT;PI3KAKT activation;Role of LAT2NTALLAB on calcium mobilization;DAP12 signaling;Signaling by PDGF;Signaling by ERBB2;IRS-related events triggered by IGF1R;CD28 co-stimulation;Cell surface interactions at the vascular wall 
                                               hsa04012;hsa04014;hsa04015;hsa04022;hsa04024;hsa04062;hsa04066;hsa04068;hsa04070;hsa04071;hsa04072;hsa04150;hsa04151;hsa04152;hsa04210;hsa04211;hsa04213;hsa04261;hsa04370;hsa04380;hsa04510;hsa04550;hsa04611;hsa04620;hsa04630;hsa04650;hsa04660;hsa04662;hsa04664;hsa04666;hsa04668;hsa04670;hsa04722;hsa04725;hsa04750;hsa04810;hsa04910;hsa04914;hsa04915;hsa04917;hsa04919;hsa04921;hsa04923;hsa04930;hsa04931;hsa04932;hsa04933;hsa04960;hsa04973;hsa05100;hsa05142;hsa05145;hsa05146;hsa05160;hsa05161;hsa05162;hsa05164;hsa05166;hsa05169;hsa05200;hsa05203;hsa05205;hsa05206;hsa05210;hsa05211;hsa05212;hsa05213;hsa05214;hsa05215;hsa05218;hsa05220;hsa05221;hsa05222;hsa05223;hsa05230;hsa05231 
                                               PIK3R2 
                                               NA 
                                               azure4 
                                               1 
                                               -2.68939393939394 
                                             
                                             
                                               SRC-mRNA 
                                               0.261 
                                               0.114 
                                               0.0374 
                                               0.485 
                                               1.2 
                                               1.03 
                                               1.4 
                                               0.0326 
                                               0.908 
                                               loglinear 
                                               ARMS-mediated activation, CD28 co-stimulation, Cell surface interactions at the vascular wall, CTLA4 inhibitory signaling, DAP12 signaling, Downstream signal transduction, FCERI mediated MAPK activation, Frs2-mediated activation, GAB1 signalosome, Gastrin-CREB signalling pathway via PKC and MAPK, Integrin alphaIIb beta3 signaling, Interleukin receptor SHC signaling, IRS-related events triggered by IGF1R, MAP2K and MAPK activation, Negative regulation of FGFR1 signaling, Negative regulation of FGFR2 signaling, Negative regulation of FGFR3 signaling, Negative regulation of FGFR4 signaling, p130Cas linkage to MAPK signaling for integrins, RAFMAP kinase cascade, Signaling by ERBB2, Signaling by PDGF, Signaling by SCF-KIT, Spry regulation of FGF signaling, VEGFA-VEGFR2 Pathway, VEGFR2 mediated cell proliferation 
                                               SEBELIUS_2_C4636 
                                               SRC 
                                               mRNA 
                                               0 
                                               
                                               
                                               FCGR activation;Spry regulation of FGF signaling;GAB1 signalosome;MAP2K and MAPK activation;CLEC7A (Dectin-1) signaling;CD28 co-stimulation;p130Cas linkage to MAPK signaling for integrins;CTLA4 inhibitory signaling;GRB2.CL.SOS provides linkage to MAPK signaling for Integrins ;Signal regulatory protein (SIRP) family interactions;PECAM1 interactions;Thrombin signalling through proteinase activated receptors (PARs);GP1b-IX-V activation signalling;c-src mediated regulation of Cx43 function and closure of gap junctions;Signaling by ERBB2;VEGFA-VEGFR2 Pathway;Signaling by SCF-KIT;VEGFR2 mediated cell proliferation;RAFMAP kinase cascade;Downstream signal transduction;Frs2-mediated activation;ARMS-mediated activation;Gastrin-CREB signalling pathway via PKC and MAPK;Interleukin receptor SHC signaling;FCERI mediated MAPK activation;DAP12 signaling;Signaling by PDGF;IRS-related events triggered by IGF1R;Negative regulation of FGFR3 signaling;Negative regulation of FGFR4 signaling;Negative regulation of FGFR2 signaling;Negative regulation of FGFR1 signaling;Integrin alphaIIb beta3 signaling;Cell surface interactions at the vascular wall 
                                               hsa04012;hsa04015;hsa04062;hsa04144;hsa04370;hsa04510;hsa04520;hsa04530;hsa04540;hsa04611;hsa04727;hsa04750;hsa04810;hsa04912;hsa04915;hsa04917;hsa04919;hsa04921;hsa05100;hsa05120;hsa05131;hsa05152;hsa05161;hsa05203;hsa05205;hsa05219 
                                               SRC 
                                               NA 
                                               azure4 
                                               1 
                                               2.28947368421053 
                                             
                                             
                                               RAC1-mRNA 
                                               -0.125 
                                               0.0675 
                                               -0.257 
                                               0.00726 
                                               0.917 
                                               0.837 
                                               1.01 
                                               0.0781 
                                               1 
                                               loglinear 
                                               Activation of Rac, CD28 co-stimulation, CD28 dependent Vav1 pathway, DAP12 signaling, EPH-ephrin mediated repulsion of cells, Ephrin signaling, G alpha (1213) signalling events, GPVI-mediated activation cascade, MAPK6MAPK4 signaling, NRAGE signals death through JNK, PCPCE pathway, Regulation of actin dynamics for phagocytic cup formation, RHO GTPases activate CIT, RHO GTPases Activate Formins, RHO GTPases activate IQGAPs, RHO GTPases activate PAKs, RHO GTPases activate PKNs, RHO GTPases Activate WASPs and WAVEs, Sema3A PAK dependent Axon repulsion, Sema4D induced cell migration and growth-cone collapse, Signal transduction by L1, Signaling by SCF-KIT, VEGFA-VEGFR2 Pathway, VEGFR2 mediated vascular permeability 
                                               SEBELIUS_2_C4636 
                                               RAC1 
                                               mRNA 
                                               0 
                                               
                                               
                                               MAPK6MAPK4 signaling;Signaling by SCF-KIT;Inactivation of Cdc42 and Rac;DAP12 signaling;Ephrin signaling;EPH-ephrin mediated repulsion of cells;Activation of Rac;RHO GTPases Activate Formins;VEGFR2 mediated vascular permeability;RHO GTPases activate PAKs;Sema4D mediated inhibition of cell attachment and migration;G alpha (1213) signalling events;Regulation of actin dynamics for phagocytic cup formation;Sema4D induced cell migration and growth-cone collapse;Signal transduction by L1;RHO GTPases activate CIT;RHO GTPases Activate WASPs and WAVEs;Factors involved in megakaryocyte development and platelet production;RHO GTPases activate KTN1;CD28 dependent Vav1 pathway;DSCAM interactions;NRAGE signals death through JNK;GPVI-mediated activation cascade;Sema3A PAK dependent Axon repulsion;RHO GTPases activate PKNs;SEMA3A-Plexin repulsion signaling by inhibiting Integrin adhesion;RHO GTPases activate IQGAPs;PCPCE pathway;VEGFA-VEGFR2 Pathway;CD28 co-stimulation 
                                               hsa04010;hsa04014;hsa04015;hsa04024;hsa04062;hsa04071;hsa04145;hsa04151;hsa04310;hsa04360;hsa04370;hsa04380;hsa04510;hsa04520;hsa04620;hsa04650;hsa04662;hsa04664;hsa04666;hsa04670;hsa04722;hsa04810;hsa04932;hsa04933;hsa04972;hsa05100;hsa05120;hsa05131;hsa05132;hsa05200;hsa05203;hsa05205;hsa05210;hsa05211;hsa05212;hsa05231;hsa05416 
                                               RAC1 
                                               NA 
                                               azure4 
                                               1 
                                               -1.85185185185185 
                                             
                                             
                                               YES1-mRNA 
                                               0.124 
                                               0.0716 
                                               -0.0159 
                                               0.265 
                                               1.09 
                                               0.989 
                                               1.2 
                                               0.0968 
                                               1 
                                               loglinear 
                                               CD28 co-stimulation, Cell surface interactions at the vascular wall, CTLA4 inhibitory signaling, EPH-ephrin mediated repulsion of cells, EPHA-mediated growth cone collapse, EPHB-mediated forward signaling, Regulation of signaling by CBL, Signaling by ERBB2, Signaling by SCF-KIT 
                                               SEBELIUS_2_C4636 
                                               YES1 
                                               mRNA 
                                               0 
                                               
                                               
                                               Signaling by ERBB2;EPHB-mediated forward signaling;EPHA-mediated growth cone collapse;Regulation of KIT signaling;EPH-ephrin mediated repulsion of cells;Regulation of signaling by CBL;CD28 co-stimulation;CTLA4 inhibitory signaling;Signaling by SCF-KIT;Cell surface interactions at the vascular wall 
                                               hsa04520;hsa04530 
                                               YES1 
                                               NA 
                                               azure4 
                                               1 
                                               1.73184357541899 
                                             
                                             
                                               AKT1-mRNA 
                                               -0.242 
                                               0.151 
                                               -0.538 
                                               0.0548 
                                               0.846 
                                               0.689 
                                               1.04 
                                               0.125 
                                               1 
                                               lm.nb 
                                               Activation of BAD and translocation to mitochondria , CD28 co-stimulation, DAP12 signaling, Deactivation of the beta-catenin transactivating complex, Downstream signal transduction, G beta.CL.gamma signalling through PI3Kgamma, GAB1 signalosome, GPVI-mediated activation cascade, IRS-related events triggered by IGF1R, PI3K Cascade, PI3KAKT activation, PIP3 activates AKT signaling, Role of LAT2NTALLAB on calcium mobilization, Signaling by ERBB2, Signaling by PDGF, Signaling by SCF-KIT, TP53 Regulates Metabolic Genes, VEGFA-VEGFR2 Pathway, VEGFR2 mediated vascular permeability 
                                               SEBELIUS_2_C4636 
                                               AKT1 
                                               mRNA 
                                               0 
                                               
                                               
                                               TP53 Regulates Metabolic Genes;Deactivation of the beta-catenin transactivating complex;Activation of BAD and translocation to mitochondria ;eNOS activation;mTOR signalling;G beta.CL.gamma signalling through PI3Kgamma;KSRP (KHSRP) binds and destabilizes mRNA;AKT phosphorylates targets in the nucleus;VEGFR2 mediated vascular permeability;GPVI-mediated activation cascade;Tetrahydrobiopterin (BH4) synthesis# recycling# salvage and regulation;Butyrate Response Factor 1 (BRF1) binds and destabilizes mRNA;Signaling by ERBB2;PIP3 activates AKT signaling;Downstream signal transduction;GAB1 signalosome;Signaling by SCF-KIT;PI3KAKT activation;Role of LAT2NTALLAB on calcium mobilization;DAP12 signaling;Signaling by PDGF;CD28 co-stimulation;PI3K Cascade;IRS-related events triggered by IGF1R;VEGFA-VEGFR2 Pathway 
                                               hsa04010;hsa04012;hsa04014;hsa04015;hsa04022;hsa04024;hsa04062;hsa04066;hsa04068;hsa04071;hsa04072;hsa04150;hsa04151;hsa04152;hsa04210;hsa04211;hsa04213;hsa04261;hsa04370;hsa04380;hsa04510;hsa04530;hsa04550;hsa04611;hsa04620;hsa04630;hsa04660;hsa04662;hsa04664;hsa04666;hsa04668;hsa04722;hsa04725;hsa04728;hsa04910;hsa04914;hsa04915;hsa04917;hsa04919;hsa04920;hsa04922;hsa04923;hsa04931;hsa04932;hsa04933;hsa04973;hsa05142;hsa05145;hsa05152;hsa05160;hsa05161;hsa05162;hsa05164;hsa05166;hsa05169;hsa05200;hsa05205;hsa05210;hsa05211;hsa05212;hsa05213;hsa05214;hsa05215;hsa05218;hsa05220;hsa05221;hsa05222;hsa05223;hsa05230;hsa05231 
                                               AKT1 
                                               NA 
                                               azure4 
                                               1 
                                               -1.60264900662252 
                                             
                                             
                                               PIK3R1-mRNA 
                                               0.161 
                                               0.119 
                                               -0.0725 
                                               0.395 
                                               1.12 
                                               0.951 
                                               1.31 
                                               0.191 
                                               1 
                                               loglinear 
                                               CD28 co-stimulation, CD28 dependent PI3KAkt signaling, Cell surface interactions at the vascular wall, Constitutive Signaling by Aberrant PI3K in Cancer, Constitutive Signaling by EGFRvIII, DAP12 signaling, Downstream signal transduction, Downstream TCR signaling, G alpha (q) signalling events, G beta.CL.gamma signalling through PI3Kgamma, GAB1 signalosome, Gastrin-CREB signalling pathway via PKC and MAPK, GPVI-mediated activation cascade, Interleukin receptor SHC signaling, IRS-related events triggered by IGF1R, Nephrin interactions, PI3KAKT activation, PIP3 activates AKT signaling, Regulation of signaling by CBL, Role of LAT2NTALLAB on calcium mobilization, Signaling by ERBB2, Signaling by FGFR1 in disease, Signaling by FGFR2 in disease, Signaling by FGFR3 in disease, Signaling by FGFR4 in disease, Signaling by PDGF, Signaling by SCF-KIT, Synthesis of PIPs at the plasma membrane, Tie2 Signaling, VEGFA-VEGFR2 Pathway 
                                               SEBELIUS_2_C4636 
                                               PIK3R1 
                                               mRNA 
                                               0 
                                               
                                               
                                               G alpha (q) signalling events;Downstream TCR signaling;Signaling by FGFR2 in disease;PIP3 activates AKT signaling;VEGFA-VEGFR2 Pathway;Synthesis of PIPs at the plasma membrane;Constitutive Signaling by Ligand-Responsive EGFR Cancer Variants;Interleukin receptor SHC signaling;G beta.CL.gamma signalling through PI3Kgamma;Regulation of signaling by CBL;Role of phospholipids in phagocytosis;Signaling by FGFR3 in disease;Constitutive Signaling by Aberrant PI3K in Cancer;Signaling by FGFR4 in disease;CD28 dependent PI3KAkt signaling;Tie2 Signaling;Signaling by FGFR1 fusion mutants;GP1b-IX-V activation signalling;GPVI-mediated activation cascade;Nephrin interactions;Constitutive Signaling by EGFRvIII;Gastrin-CREB signalling pathway via PKC and MAPK;Downstream signal transduction;GAB1 signalosome;Signaling by SCF-KIT;PI3KAKT activation;Role of LAT2NTALLAB on calcium mobilization;DAP12 signaling;Signaling by PDGF;Signaling by ERBB2;IRS-related events triggered by IGF1R;CD28 co-stimulation;Cell surface interactions at the vascular wall;Signaling by FGFR1 in disease 
                                               hsa04012;hsa04014;hsa04015;hsa04022;hsa04024;hsa04062;hsa04066;hsa04068;hsa04070;hsa04071;hsa04072;hsa04150;hsa04151;hsa04152;hsa04210;hsa04211;hsa04213;hsa04261;hsa04370;hsa04380;hsa04510;hsa04550;hsa04611;hsa04620;hsa04630;hsa04650;hsa04660;hsa04662;hsa04664;hsa04666;hsa04668;hsa04670;hsa04722;hsa04725;hsa04750;hsa04810;hsa04910;hsa04914;hsa04915;hsa04917;hsa04919;hsa04921;hsa04923;hsa04930;hsa04931;hsa04932;hsa04933;hsa04960;hsa04973;hsa05100;hsa05142;hsa05145;hsa05146;hsa05160;hsa05161;hsa05162;hsa05164;hsa05166;hsa05169;hsa05200;hsa05203;hsa05205;hsa05210;hsa05211;hsa05212;hsa05213;hsa05214;hsa05215;hsa05218;hsa05220;hsa05221;hsa05222;hsa05223;hsa05230;hsa05231 
                                               PIK3R1 
                                               NA 
                                               azure4 
                                               1 
                                               1.35294117647059 
                                             
                                             
                                               PIK3CA-mRNA 
                                               -0.113 
                                               0.102 
                                               -0.312 
                                               0.0861 
                                               0.925 
                                               0.806 
                                               1.06 
                                               0.279 
                                               1 
                                               loglinear 
                                               CD28 co-stimulation, CD28 dependent PI3KAkt signaling, Cell surface interactions at the vascular wall, Constitutive Signaling by Aberrant PI3K in Cancer, Constitutive Signaling by EGFRvIII, DAP12 signaling, Downstream signal transduction, Downstream TCR signaling, G alpha (1213) signalling events, G alpha (q) signalling events, G beta.CL.gamma signalling through PI3Kgamma, GAB1 signalosome, Gastrin-CREB signalling pathway via PKC and MAPK, GPVI-mediated activation cascade, Interleukin receptor SHC signaling, IRS-related events triggered by IGF1R, Nephrin interactions, PI3K Cascade, PI3KAKT activation, PIP3 activates AKT signaling, Regulation of signaling by CBL, Role of LAT2NTALLAB on calcium mobilization, Signaling by ERBB2, Signaling by FGFR1 in disease, Signaling by FGFR2 in disease, Signaling by FGFR3 in disease, Signaling by FGFR4 in disease, Signaling by PDGF, Signaling by SCF-KIT, Synthesis of PIPs at the plasma membrane, Tie2 Signaling, VEGFA-VEGFR2 Pathway 
                                               SEBELIUS_2_C4636 
                                               PIK3CA 
                                               mRNA 
                                               0 
                                               
                                               
                                               G alpha (q) signalling events;Downstream TCR signaling;Signaling by FGFR2 in disease;PIP3 activates AKT signaling;VEGFA-VEGFR2 Pathway;Synthesis of PIPs at the plasma membrane;Constitutive Signaling by Ligand-Responsive EGFR Cancer Variants;Interleukin receptor SHC signaling;G beta.CL.gamma signalling through PI3Kgamma;Regulation of signaling by CBL;Role of phospholipids in phagocytosis;G alpha (1213) signalling events;PI3K Cascade;Signaling by FGFR3 in disease;Constitutive Signaling by Aberrant PI3K in Cancer;Signaling by FGFR4 in disease;CD28 dependent PI3KAkt signaling;Tie2 Signaling;GPVI-mediated activation cascade;Nephrin interactions;Constitutive Signaling by EGFRvIII;Gastrin-CREB signalling pathway via PKC and MAPK;Downstream signal transduction;GAB1 signalosome;Signaling by SCF-KIT;PI3KAKT activation;Role of LAT2NTALLAB on calcium mobilization;DAP12 signaling;Signaling by PDGF;Signaling by ERBB2;IRS-related events triggered by IGF1R;CD28 co-stimulation;Cell surface interactions at the vascular wall;Signaling by FGFR1 in disease 
                                               hsa00562;hsa04012;hsa04014;hsa04015;hsa04022;hsa04024;hsa04062;hsa04066;hsa04068;hsa04070;hsa04071;hsa04072;hsa04150;hsa04151;hsa04152;hsa04210;hsa04211;hsa04213;hsa04261;hsa04370;hsa04380;hsa04510;hsa04550;hsa04611;hsa04620;hsa04630;hsa04650;hsa04660;hsa04662;hsa04664;hsa04666;hsa04668;hsa04670;hsa04722;hsa04725;hsa04750;hsa04810;hsa04910;hsa04914;hsa04915;hsa04917;hsa04919;hsa04921;hsa04923;hsa04930;hsa04931;hsa04932;hsa04933;hsa04960;hsa04973;hsa05100;hsa05142;hsa05145;hsa05146;hsa05160;hsa05161;hsa05162;hsa05164;hsa05166;hsa05169;hsa05200;hsa05203;hsa05205;hsa05206;hsa05210;hsa05211;hsa05212;hsa05213;hsa05214;hsa05215;hsa05218;hsa05220;hsa05221;hsa05222;hsa05223;hsa05230;hsa05231 
                                               PIK3CA 
                                               NA 
                                               azure4 
                                               1 
                                               -1.1078431372549 
                                             
                                             
                                               PAK1-mRNA 
                                               -0.177 
                                               0.163 
                                               -0.497 
                                               0.142 
                                               0.884 
                                               0.709 
                                               1.1 
                                               0.288 
                                               1 
                                               lm.nb 
                                               Activation of Rac, CD209 (DC-SIGN) signaling, CD28 co-stimulation, CD28 dependent Vav1 pathway, EPHB-mediated forward signaling, Ephrin signaling, FCERI mediated MAPK activation, MAPK6MAPK4 signaling, Regulation of actin dynamics for phagocytic cup formation, RHO GTPases activate PAKs, Sema3A PAK dependent Axon repulsion, VEGFA-VEGFR2 Pathway, VEGFR2 mediated vascular permeability 
                                               SEBELIUS_2_C4636 
                                               PAK1 
                                               mRNA 
                                               0 
                                               
                                               
                                               EPHB-mediated forward signaling;Ephrin signaling;MAPK6MAPK4 signaling;DSCAM interactions;Activation of Rac;VEGFR2 mediated vascular permeability;RHO GTPases activate PAKs;Sema3A PAK dependent Axon repulsion;FCERI mediated MAPK activation;Regulation of actin dynamics for phagocytic cup formation;CD209 (DC-SIGN) signaling;CD28 dependent Vav1 pathway;VEGFA-VEGFR2 Pathway;CD28 co-stimulation 
                                               hsa04010;hsa04012;hsa04014;hsa04024;hsa04062;hsa04360;hsa04510;hsa04650;hsa04660;hsa04666;hsa04810;hsa05120;hsa05205;hsa05211 
                                               PAK1 
                                               NA 
                                               azure4 
                                               1 
                                               -1.08588957055215 
                                             
                                             
                                               PIK3R3-mRNA 
                                               0.268 
                                               0.258 
                                               -0.238 
                                               0.775 
                                               1.2 
                                               0.848 
                                               1.71 
                                               0.311 
                                               1 
                                               lm.nb 
                                               CD28 co-stimulation, CD28 dependent PI3KAkt signaling, Constitutive Signaling by Aberrant PI3K in Cancer, DAP12 signaling, Downstream signal transduction, G alpha (1213) signalling events, G alpha (q) signalling events, G beta.CL.gamma signalling through PI3Kgamma, GAB1 signalosome, Gastrin-CREB signalling pathway via PKC and MAPK, GPVI-mediated activation cascade, Interleukin receptor SHC signaling, PI3KAKT activation, PIP3 activates AKT signaling, Regulation of signaling by CBL, Role of LAT2NTALLAB on calcium mobilization, Signaling by ERBB2, Signaling by PDGF, Signaling by SCF-KIT, Synthesis of PIPs at the plasma membrane 
                                               SEBELIUS_2_C4636 
                                               PIK3R3 
                                               mRNA 
                                               0 
                                               
                                               
                                               G alpha (q) signalling events;PIP3 activates AKT signaling;Constitutive Signaling by Aberrant PI3K in Cancer;CD28 dependent PI3KAkt signaling;Synthesis of PIPs at the plasma membrane;Interleukin receptor SHC signaling;G beta.CL.gamma signalling through PI3Kgamma;Regulation of signaling by CBL;GPVI-mediated activation cascade;G alpha (1213) signalling events;Gastrin-CREB signalling pathway via PKC and MAPK;Downstream signal transduction;GAB1 signalosome;Signaling by SCF-KIT;PI3KAKT activation;Role of LAT2NTALLAB on calcium mobilization;DAP12 signaling;Signaling by PDGF;Signaling by ERBB2;CD28 co-stimulation 
                                               hsa04012;hsa04014;hsa04015;hsa04022;hsa04024;hsa04062;hsa04066;hsa04068;hsa04070;hsa04071;hsa04072;hsa04150;hsa04151;hsa04152;hsa04210;hsa04211;hsa04213;hsa04261;hsa04370;hsa04380;hsa04510;hsa04550;hsa04611;hsa04620;hsa04630;hsa04650;hsa04660;hsa04662;hsa04664;hsa04666;hsa04668;hsa04670;hsa04722;hsa04725;hsa04750;hsa04810;hsa04910;hsa04914;hsa04915;hsa04917;hsa04919;hsa04921;hsa04923;hsa04930;hsa04931;hsa04932;hsa04933;hsa04960;hsa04973;hsa05100;hsa05142;hsa05145;hsa05146;hsa05160;hsa05161;hsa05162;hsa05164;hsa05166;hsa05169;hsa05200;hsa05203;hsa05205;hsa05210;hsa05211;hsa05212;hsa05213;hsa05214;hsa05215;hsa05218;hsa05220;hsa05221;hsa05222;hsa05223;hsa05230;hsa05231 
                                               PIK3R3 
                                               NA 
                                               azure4 
                                               1 
                                               1.03875968992248 
                                             
                                             
                                               PAK2-mRNA 
                                               -0.0762 
                                               0.0976 
                                               -0.268 
                                               0.115 
                                               0.949 
                                               0.831 
                                               1.08 
                                               0.444 
                                               1 
                                               loglinear 
                                               Activation of Rac, CD209 (DC-SIGN) signaling, CD28 co-stimulation, CD28 dependent Vav1 pathway, Ephrin signaling, RHO GTPases activate PAKs, Sema3A PAK dependent Axon repulsion, VEGFA-VEGFR2 Pathway, VEGFR2 mediated vascular permeability 
                                               SEBELIUS_2_C4636 
                                               PAK2 
                                               mRNA 
                                               0 
                                               
                                               
                                               Nef and signal transduction;CD28 dependent Vav1 pathway;Stimulation of the cell death response by PAK-2p34;Ephrin signaling;Activation of Rac;RHO GTPases activate PAKs;VEGFR2 mediated vascular permeability;Sema3A PAK dependent Axon repulsion;CD209 (DC-SIGN) signaling;VEGFA-VEGFR2 Pathway;CD28 co-stimulation 
                                               hsa04010;hsa04012;hsa04014;hsa04360;hsa04510;hsa04660;hsa04810;hsa05211 
                                               PAK2 
                                               NA 
                                               azure4 
                                               1 
                                               -0.780737704918033 
                                             
                                             
                                               FYN-mRNA 
                                               0.0581 
                                               0.11 
                                               -0.157 
                                               0.273 
                                               1.04 
                                               0.897 
                                               1.21 
                                               0.602 
                                               1 
                                               loglinear 
                                               ARMS-mediated activation, CD209 (DC-SIGN) signaling, CD28 co-stimulation, CD28 dependent PI3KAkt signaling, CD28 dependent Vav1 pathway, Cell surface interactions at the vascular wall, Constitutive Signaling by Aberrant PI3K in Cancer, CTLA4 inhibitory signaling, DAP12 signaling, Downstream signal transduction, EPH-ephrin mediated repulsion of cells, FCERI mediated MAPK activation, Frs2-mediated activation, GAB1 signalosome, Gastrin-CREB signalling pathway via PKC and MAPK, GPVI-mediated activation cascade, Interleukin receptor SHC signaling, IRS-related events triggered by IGF1R, Nephrin interactions, PI3KAKT activation, RAFMAP kinase cascade, Regulation of signaling by CBL, Role of LAT2NTALLAB on calcium mobilization, Sema3A PAK dependent Axon repulsion, Signaling by ERBB2, Signaling by PDGF, Signaling by SCF-KIT, VEGFA-VEGFR2 Pathway, VEGFR2 mediated cell proliferation 
                                               SEBELIUS_2_C4636 
                                               FYN 
                                               mRNA 
                                               0 
                                               
                                               
                                               EPH-ephrin mediated repulsion of cells;Regulation of signaling by CBL;FCGR activation;Antigen activates B Cell Receptor (BCR) leading to generation of second messengers;Dectin-2 family;RAFMAP kinase cascade;Constitutive Signaling by Aberrant PI3K in Cancer;CTLA4 inhibitory signaling;CD28 dependent PI3KAkt signaling;PECAM1 interactions;CD28 dependent Vav1 pathway;GPVI-mediated activation cascade;Sema3A PAK dependent Axon repulsion;Nephrin interactions;SEMA3A-Plexin repulsion signaling by inhibiting Integrin adhesion;CRMPs in Sema3A signaling;CD209 (DC-SIGN) signaling;Downstream signal transduction;GAB1 signalosome;Signaling by SCF-KIT;PI3KAKT activation;Role of LAT2NTALLAB on calcium mobilization;DAP12 signaling;Signaling by PDGF;Signaling by ERBB2;VEGFA-VEGFR2 Pathway;VEGFR2 mediated cell proliferation;Frs2-mediated activation;ARMS-mediated activation;Gastrin-CREB signalling pathway via PKC and MAPK;Interleukin receptor SHC signaling;FCERI mediated MAPK activation;IRS-related events triggered by IGF1R;CD28 co-stimulation;Cell surface interactions at the vascular wall 
                                               hsa04071;hsa04072;hsa04360;hsa04380;hsa04510;hsa04520;hsa04611;hsa04650;hsa04660;hsa04664;hsa04725;hsa05020;hsa05130;hsa05162;hsa05416 
                                               FYN 
                                               NA 
                                               azure4 
                                               1 
                                               0.528181818181818 
                                             
                                             
                                               AKT3-mRNA 
                                               -0.0262 
                                               0.106 
                                               -0.234 
                                               0.182 
                                               0.982 
                                               0.85 
                                               1.13 
                                               0.808 
                                               1 
                                               lm.nb 
                                               Activation of BAD and translocation to mitochondria , CD28 co-stimulation, CD28 dependent PI3KAkt signaling, DAP12 signaling, Downstream signal transduction, G beta.CL.gamma signalling through PI3Kgamma, GAB1 signalosome, GPVI-mediated activation cascade, PI3KAKT activation, PIP3 activates AKT signaling, Role of LAT2NTALLAB on calcium mobilization, Signaling by ERBB2, Signaling by PDGF, Signaling by SCF-KIT, TP53 Regulates Metabolic Genes, VEGFA-VEGFR2 Pathway, VEGFR2 mediated vascular permeability 
                                               SEBELIUS_2_C4636 
                                               AKT3 
                                               mRNA 
                                               0 
                                               
                                               
                                               TP53 Regulates Metabolic Genes;CD28 dependent PI3KAkt signaling;AKT phosphorylates targets in the cytosol;Negative regulation of the PI3KAKT network;Activation of BAD and translocation to mitochondria ;G beta.CL.gamma signalling through PI3Kgamma;AKT phosphorylates targets in the nucleus;VEGFR2 mediated vascular permeability;GPVI-mediated activation cascade;Signaling by ERBB2;PIP3 activates AKT signaling;Downstream signal transduction;GAB1 signalosome;Signaling by SCF-KIT;PI3KAKT activation;Role of LAT2NTALLAB on calcium mobilization;DAP12 signaling;Signaling by PDGF;CD28 co-stimulation;VEGFA-VEGFR2 Pathway 
                                               hsa04010;hsa04012;hsa04014;hsa04015;hsa04022;hsa04024;hsa04062;hsa04066;hsa04068;hsa04071;hsa04072;hsa04150;hsa04151;hsa04152;hsa04210;hsa04211;hsa04213;hsa04261;hsa04370;hsa04380;hsa04510;hsa04530;hsa04550;hsa04611;hsa04620;hsa04630;hsa04660;hsa04662;hsa04664;hsa04666;hsa04668;hsa04722;hsa04725;hsa04728;hsa04910;hsa04914;hsa04915;hsa04917;hsa04919;hsa04920;hsa04922;hsa04923;hsa04931;hsa04932;hsa04933;hsa04973;hsa05142;hsa05145;hsa05152;hsa05160;hsa05161;hsa05162;hsa05164;hsa05166;hsa05169;hsa05200;hsa05205;hsa05210;hsa05211;hsa05212;hsa05213;hsa05214;hsa05215;hsa05218;hsa05220;hsa05221;hsa05222;hsa05223;hsa05230;hsa05231 
                                               AKT3 
                                               NA 
                                               azure4 
                                               1 
                                               -0.247169811320755 
                                             
                                             
                                               CDC42-mRNA 
                                               0.011 
                                               0.0802 
                                               -0.146 
                                               0.168 
                                               1.01 
                                               0.904 
                                               1.12 
                                               0.892 
                                               1 
                                               loglinear 
                                               CD28 co-stimulation, CD28 dependent Vav1 pathway, DCC mediated attractive signaling, EPHB-mediated forward signaling, G alpha (1213) signalling events, GPVI-mediated activation cascade, Regulation of actin dynamics for phagocytic cup formation, Rho GTPase cycle, RHO GTPases Activate Formins, RHO GTPases activate IQGAPs, RHO GTPases activate PAKs, Sema4D induced cell migration and growth-cone collapse, VEGFA-VEGFR2 Pathway 
                                               SEBELIUS_2_C4636 
                                               CDC42 
                                               mRNA 
                                               0 
                                               
                                               
                                               DCC mediated attractive signaling;Factors involved in megakaryocyte development and platelet production;VEGFA-VEGFR2 Pathway;RHO GTPases activate KTN1;CD28 dependent Vav1 pathway;Inactivation of Cdc42 and Rac;Rho GTPase cycle;EPHB-mediated forward signaling;RHO GTPases Activate Formins;RHO GTPases activate PAKs;GPVI-mediated activation cascade;G alpha (1213) signalling events;RHO GTPases activate IQGAPs;Sema4D induced cell migration and growth-cone collapse;Regulation of actin dynamics for phagocytic cup formation;CD28 co-stimulation 
                                               hsa04010;hsa04014;hsa04015;hsa04062;hsa04144;hsa04360;hsa04370;hsa04510;hsa04520;hsa04530;hsa04660;hsa04666;hsa04670;hsa04722;hsa04810;hsa04912;hsa04932;hsa04933;hsa05100;hsa05120;hsa05130;hsa05131;hsa05132;hsa05200;hsa05203;hsa05205;hsa05211;hsa05212 
                                               CDC42 
                                               NA 
                                               azure4 
                                               1 
                                               0.13715710723192 
                                             
                                           
                                           DE Results - CD28 co-stimulation genes - New.Annotation: Control vs.cel67  Table displaying each sample's global significance scores and directed global significance scores as defined in the heatmaps above. The global significance score is calculated as the square root of the mean squared t-statistic for the genes in a gene set, with t-statistics coming from the linear regression underlying our differential expression analysis. The directed global significance score is calculated as the square root of the mean signed squared t-statistic for the genes in a gene set, with t-statistics coming from the linear regression underlying our differential expression analysis.   
                                      
                                       
                                  
                                
                                
                                  
                                       
                                      
                                           Volcano Plot: New.Annotation: miR142 vs.cel67   More Plot Information                Volcano Plot: New.Annotation: miR142 vs.cel67  Volcano plot displaying each gene's -log10(p-value) and log2 fold change for the selected covariate. Highly statistically significant genes fall at the top of the plot, and highly differentially expressed genes fall to either side. Genes within the selected gene set are highlighted in orange. Horizontal lines indicate various False Discovery Rate (FDR) thresholds.   
                                           DE Results - CD28 co-stimulation genes - New.Annotation: miR142 vs.cel67   More Plot Information      Download CSV Data     
                                           
                                             
                                             
                                           
                                           
                                             
                                               Probe Label 
                                               Log2 fold change 
                                               std error (log2) 
                                               Lower confidence limit (log2) 
                                               Upper confidence limit (log2) 
                                               Linear fold change 
                                               Lower confidence limit (linear) 
                                               Upper confidence limit (linear) 
                                               P-value 
                                               BY.p.value 
                                               method 
                                               Gene.sets 
                                               Codeset.Name 
                                               Probe.Label 
                                               Analyte.Type 
                                               Is.Control 
                                               Control.Type 
                                               Related.Probes 
                                               Probe.Annotation 
                                               KEGG.Pathways 
                                               Official.Gene.Name 
                                               Control.Conc 
                                               volcanocol 
                                               volcanopch 
                                               tstats.all 
                                             
                                           
                                           
                                             
                                               RAC1-mRNA 
                                               -1.32 
                                               0.0675 
                                               -1.45 
                                               -1.19 
                                               0.401 
                                               0.366 
                                               0.439 
                                               5.98e-15 
                                               7.94e-12 
                                               loglinear 
                                               Activation of Rac, CD28 co-stimulation, CD28 dependent Vav1 pathway, DAP12 signaling, EPH-ephrin mediated repulsion of cells, Ephrin signaling, G alpha (1213) signalling events, GPVI-mediated activation cascade, MAPK6MAPK4 signaling, NRAGE signals death through JNK, PCPCE pathway, Regulation of actin dynamics for phagocytic cup formation, RHO GTPases activate CIT, RHO GTPases Activate Formins, RHO GTPases activate IQGAPs, RHO GTPases activate PAKs, RHO GTPases activate PKNs, RHO GTPases Activate WASPs and WAVEs, Sema3A PAK dependent Axon repulsion, Sema4D induced cell migration and growth-cone collapse, Signal transduction by L1, Signaling by SCF-KIT, VEGFA-VEGFR2 Pathway, VEGFR2 mediated vascular permeability 
                                               SEBELIUS_2_C4636 
                                               RAC1 
                                               mRNA 
                                               0 
                                               
                                               
                                               MAPK6MAPK4 signaling;Signaling by SCF-KIT;Inactivation of Cdc42 and Rac;DAP12 signaling;Ephrin signaling;EPH-ephrin mediated repulsion of cells;Activation of Rac;RHO GTPases Activate Formins;VEGFR2 mediated vascular permeability;RHO GTPases activate PAKs;Sema4D mediated inhibition of cell attachment and migration;G alpha (1213) signalling events;Regulation of actin dynamics for phagocytic cup formation;Sema4D induced cell migration and growth-cone collapse;Signal transduction by L1;RHO GTPases activate CIT;RHO GTPases Activate WASPs and WAVEs;Factors involved in megakaryocyte development and platelet production;RHO GTPases activate KTN1;CD28 dependent Vav1 pathway;DSCAM interactions;NRAGE signals death through JNK;GPVI-mediated activation cascade;Sema3A PAK dependent Axon repulsion;RHO GTPases activate PKNs;SEMA3A-Plexin repulsion signaling by inhibiting Integrin adhesion;RHO GTPases activate IQGAPs;PCPCE pathway;VEGFA-VEGFR2 Pathway;CD28 co-stimulation 
                                               hsa04010;hsa04014;hsa04015;hsa04024;hsa04062;hsa04071;hsa04145;hsa04151;hsa04310;hsa04360;hsa04370;hsa04380;hsa04510;hsa04520;hsa04620;hsa04650;hsa04662;hsa04664;hsa04666;hsa04670;hsa04722;hsa04810;hsa04932;hsa04933;hsa04972;hsa05100;hsa05120;hsa05131;hsa05132;hsa05200;hsa05203;hsa05205;hsa05210;hsa05211;hsa05212;hsa05231;hsa05416 
                                               RAC1 
                                               NA 
                                               slateblue1 
                                               16 
                                               -19.5555555555556 
                                             
                                             
                                               YES1-mRNA 
                                               -1.3 
                                               0.0716 
                                               -1.44 
                                               -1.16 
                                               0.405 
                                               0.368 
                                               0.447 
                                               2.45e-14 
                                               1.63e-11 
                                               loglinear 
                                               CD28 co-stimulation, Cell surface interactions at the vascular wall, CTLA4 inhibitory signaling, EPH-ephrin mediated repulsion of cells, EPHA-mediated growth cone collapse, EPHB-mediated forward signaling, Regulation of signaling by CBL, Signaling by ERBB2, Signaling by SCF-KIT 
                                               SEBELIUS_2_C4636 
                                               YES1 
                                               mRNA 
                                               0 
                                               
                                               
                                               Signaling by ERBB2;EPHB-mediated forward signaling;EPHA-mediated growth cone collapse;Regulation of KIT signaling;EPH-ephrin mediated repulsion of cells;Regulation of signaling by CBL;CD28 co-stimulation;CTLA4 inhibitory signaling;Signaling by SCF-KIT;Cell surface interactions at the vascular wall 
                                               hsa04520;hsa04530 
                                               YES1 
                                               NA 
                                               slateblue1 
                                               16 
                                               -18.1564245810056 
                                             
                                             
                                               PIK3R1-mRNA 
                                               0.624 
                                               0.119 
                                               0.39 
                                               0.858 
                                               1.54 
                                               1.31 
                                               1.81 
                                               3.45e-05 
                                               0.00121 
                                               loglinear 
                                               CD28 co-stimulation, CD28 dependent PI3KAkt signaling, Cell surface interactions at the vascular wall, Constitutive Signaling by Aberrant PI3K in Cancer, Constitutive Signaling by EGFRvIII, DAP12 signaling, Downstream signal transduction, Downstream TCR signaling, G alpha (q) signalling events, G beta.CL.gamma signalling through PI3Kgamma, GAB1 signalosome, Gastrin-CREB signalling pathway via PKC and MAPK, GPVI-mediated activation cascade, Interleukin receptor SHC signaling, IRS-related events triggered by IGF1R, Nephrin interactions, PI3KAKT activation, PIP3 activates AKT signaling, Regulation of signaling by CBL, Role of LAT2NTALLAB on calcium mobilization, Signaling by ERBB2, Signaling by FGFR1 in disease, Signaling by FGFR2 in disease, Signaling by FGFR3 in disease, Signaling by FGFR4 in disease, Signaling by PDGF, Signaling by SCF-KIT, Synthesis of PIPs at the plasma membrane, Tie2 Signaling, VEGFA-VEGFR2 Pathway 
                                               SEBELIUS_2_C4636 
                                               PIK3R1 
                                               mRNA 
                                               0 
                                               
                                               
                                               G alpha (q) signalling events;Downstream TCR signaling;Signaling by FGFR2 in disease;PIP3 activates AKT signaling;VEGFA-VEGFR2 Pathway;Synthesis of PIPs at the plasma membrane;Constitutive Signaling by Ligand-Responsive EGFR Cancer Variants;Interleukin receptor SHC signaling;G beta.CL.gamma signalling through PI3Kgamma;Regulation of signaling by CBL;Role of phospholipids in phagocytosis;Signaling by FGFR3 in disease;Constitutive Signaling by Aberrant PI3K in Cancer;Signaling by FGFR4 in disease;CD28 dependent PI3KAkt signaling;Tie2 Signaling;Signaling by FGFR1 fusion mutants;GP1b-IX-V activation signalling;GPVI-mediated activation cascade;Nephrin interactions;Constitutive Signaling by EGFRvIII;Gastrin-CREB signalling pathway via PKC and MAPK;Downstream signal transduction;GAB1 signalosome;Signaling by SCF-KIT;PI3KAKT activation;Role of LAT2NTALLAB on calcium mobilization;DAP12 signaling;Signaling by PDGF;Signaling by ERBB2;IRS-related events triggered by IGF1R;CD28 co-stimulation;Cell surface interactions at the vascular wall;Signaling by FGFR1 in disease 
                                               hsa04012;hsa04014;hsa04015;hsa04022;hsa04024;hsa04062;hsa04066;hsa04068;hsa04070;hsa04071;hsa04072;hsa04150;hsa04151;hsa04152;hsa04210;hsa04211;hsa04213;hsa04261;hsa04370;hsa04380;hsa04510;hsa04550;hsa04611;hsa04620;hsa04630;hsa04650;hsa04660;hsa04662;hsa04664;hsa04666;hsa04668;hsa04670;hsa04722;hsa04725;hsa04750;hsa04810;hsa04910;hsa04914;hsa04915;hsa04917;hsa04919;hsa04921;hsa04923;hsa04930;hsa04931;hsa04932;hsa04933;hsa04960;hsa04973;hsa05100;hsa05142;hsa05145;hsa05146;hsa05160;hsa05161;hsa05162;hsa05164;hsa05166;hsa05169;hsa05200;hsa05203;hsa05205;hsa05210;hsa05211;hsa05212;hsa05213;hsa05214;hsa05215;hsa05218;hsa05220;hsa05221;hsa05222;hsa05223;hsa05230;hsa05231 
                                               PIK3R1 
                                               NA 
                                               slateblue1 
                                               16 
                                               5.2436974789916 
                                             
                                             
                                               PIK3R2-mRNA 
                                               -0.483 
                                               0.133 
                                               -0.743 
                                               -0.222 
                                               0.716 
                                               0.598 
                                               0.857 
                                               0.00156 
                                               0.032 
                                               lm.nb 
                                               CD28 co-stimulation, CD28 dependent PI3KAkt signaling, Cell surface interactions at the vascular wall, Constitutive Signaling by Aberrant PI3K in Cancer, DAP12 signaling, Downstream signal transduction, Downstream TCR signaling, G alpha (1213) signalling events, G alpha (q) signalling events, G beta.CL.gamma signalling through PI3Kgamma, GAB1 signalosome, Gastrin-CREB signalling pathway via PKC and MAPK, GPVI-mediated activation cascade, Interleukin receptor SHC signaling, IRS-related events triggered by IGF1R, Nephrin interactions, PI3K Cascade, PI3KAKT activation, PIP3 activates AKT signaling, Regulation of signaling by CBL, Rho GTPase cycle, Role of LAT2NTALLAB on calcium mobilization, Signaling by ERBB2, Signaling by PDGF, Signaling by SCF-KIT, Synthesis of PIPs at the plasma membrane, Tie2 Signaling, VEGFA-VEGFR2 Pathway 
                                               SEBELIUS_2_C4636 
                                               PIK3R2 
                                               mRNA 
                                               0 
                                               
                                               
                                               G alpha (q) signalling events;Downstream TCR signaling;PIP3 activates AKT signaling;VEGFA-VEGFR2 Pathway;Synthesis of PIPs at the plasma membrane;Rho GTPase cycle;Interleukin receptor SHC signaling;G beta.CL.gamma signalling through PI3Kgamma;Regulation of signaling by CBL;Role of phospholipids in phagocytosis;G alpha (1213) signalling events;PI3K Cascade;Constitutive Signaling by Aberrant PI3K in Cancer;CD28 dependent PI3KAkt signaling;Tie2 Signaling;GPVI-mediated activation cascade;Nephrin interactions;Gastrin-CREB signalling pathway via PKC and MAPK;Downstream signal transduction;GAB1 signalosome;Signaling by SCF-KIT;PI3KAKT activation;Role of LAT2NTALLAB on calcium mobilization;DAP12 signaling;Signaling by PDGF;Signaling by ERBB2;IRS-related events triggered by IGF1R;CD28 co-stimulation;Cell surface interactions at the vascular wall 
                                               hsa04012;hsa04014;hsa04015;hsa04022;hsa04024;hsa04062;hsa04066;hsa04068;hsa04070;hsa04071;hsa04072;hsa04150;hsa04151;hsa04152;hsa04210;hsa04211;hsa04213;hsa04261;hsa04370;hsa04380;hsa04510;hsa04550;hsa04611;hsa04620;hsa04630;hsa04650;hsa04660;hsa04662;hsa04664;hsa04666;hsa04668;hsa04670;hsa04722;hsa04725;hsa04750;hsa04810;hsa04910;hsa04914;hsa04915;hsa04917;hsa04919;hsa04921;hsa04923;hsa04930;hsa04931;hsa04932;hsa04933;hsa04960;hsa04973;hsa05100;hsa05142;hsa05145;hsa05146;hsa05160;hsa05161;hsa05162;hsa05164;hsa05166;hsa05169;hsa05200;hsa05203;hsa05205;hsa05206;hsa05210;hsa05211;hsa05212;hsa05213;hsa05214;hsa05215;hsa05218;hsa05220;hsa05221;hsa05222;hsa05223;hsa05230;hsa05231 
                                               PIK3R2 
                                               NA 
                                               slateblue2 
                                               16 
                                               -3.63157894736842 
                                             
                                             
                                               PIK3CA-mRNA 
                                               0.366 
                                               0.102 
                                               0.166 
                                               0.565 
                                               1.29 
                                               1.12 
                                               1.48 
                                               0.00169 
                                               0.034 
                                               loglinear 
                                               CD28 co-stimulation, CD28 dependent PI3KAkt signaling, Cell surface interactions at the vascular wall, Constitutive Signaling by Aberrant PI3K in Cancer, Constitutive Signaling by EGFRvIII, DAP12 signaling, Downstream signal transduction, Downstream TCR signaling, G alpha (1213) signalling events, G alpha (q) signalling events, G beta.CL.gamma signalling through PI3Kgamma, GAB1 signalosome, Gastrin-CREB signalling pathway via PKC and MAPK, GPVI-mediated activation cascade, Interleukin receptor SHC signaling, IRS-related events triggered by IGF1R, Nephrin interactions, PI3K Cascade, PI3KAKT activation, PIP3 activates AKT signaling, Regulation of signaling by CBL, Role of LAT2NTALLAB on calcium mobilization, Signaling by ERBB2, Signaling by FGFR1 in disease, Signaling by FGFR2 in disease, Signaling by FGFR3 in disease, Signaling by FGFR4 in disease, Signaling by PDGF, Signaling by SCF-KIT, Synthesis of PIPs at the plasma membrane, Tie2 Signaling, VEGFA-VEGFR2 Pathway 
                                               SEBELIUS_2_C4636 
                                               PIK3CA 
                                               mRNA 
                                               0 
                                               
                                               
                                               G alpha (q) signalling events;Downstream TCR signaling;Signaling by FGFR2 in disease;PIP3 activates AKT signaling;VEGFA-VEGFR2 Pathway;Synthesis of PIPs at the plasma membrane;Constitutive Signaling by Ligand-Responsive EGFR Cancer Variants;Interleukin receptor SHC signaling;G beta.CL.gamma signalling through PI3Kgamma;Regulation of signaling by CBL;Role of phospholipids in phagocytosis;G alpha (1213) signalling events;PI3K Cascade;Signaling by FGFR3 in disease;Constitutive Signaling by Aberrant PI3K in Cancer;Signaling by FGFR4 in disease;CD28 dependent PI3KAkt signaling;Tie2 Signaling;GPVI-mediated activation cascade;Nephrin interactions;Constitutive Signaling by EGFRvIII;Gastrin-CREB signalling pathway via PKC and MAPK;Downstream signal transduction;GAB1 signalosome;Signaling by SCF-KIT;PI3KAKT activation;Role of LAT2NTALLAB on calcium mobilization;DAP12 signaling;Signaling by PDGF;Signaling by ERBB2;IRS-related events triggered by IGF1R;CD28 co-stimulation;Cell surface interactions at the vascular wall;Signaling by FGFR1 in disease 
                                               hsa00562;hsa04012;hsa04014;hsa04015;hsa04022;hsa04024;hsa04062;hsa04066;hsa04068;hsa04070;hsa04071;hsa04072;hsa04150;hsa04151;hsa04152;hsa04210;hsa04211;hsa04213;hsa04261;hsa04370;hsa04380;hsa04510;hsa04550;hsa04611;hsa04620;hsa04630;hsa04650;hsa04660;hsa04662;hsa04664;hsa04666;hsa04668;hsa04670;hsa04722;hsa04725;hsa04750;hsa04810;hsa04910;hsa04914;hsa04915;hsa04917;hsa04919;hsa04921;hsa04923;hsa04930;hsa04931;hsa04932;hsa04933;hsa04960;hsa04973;hsa05100;hsa05142;hsa05145;hsa05146;hsa05160;hsa05161;hsa05162;hsa05164;hsa05166;hsa05169;hsa05200;hsa05203;hsa05205;hsa05206;hsa05210;hsa05211;hsa05212;hsa05213;hsa05214;hsa05215;hsa05218;hsa05220;hsa05221;hsa05222;hsa05223;hsa05230;hsa05231 
                                               PIK3CA 
                                               NA 
                                               slateblue2 
                                               16 
                                               3.58823529411765 
                                             
                                             
                                               PIK3R3-mRNA 
                                               0.908 
                                               0.257 
                                               0.404 
                                               1.41 
                                               1.88 
                                               1.32 
                                               2.66 
                                               0.002 
                                               0.0386 
                                               lm.nb 
                                               CD28 co-stimulation, CD28 dependent PI3KAkt signaling, Constitutive Signaling by Aberrant PI3K in Cancer, DAP12 signaling, Downstream signal transduction, G alpha (1213) signalling events, G alpha (q) signalling events, G beta.CL.gamma signalling through PI3Kgamma, GAB1 signalosome, Gastrin-CREB signalling pathway via PKC and MAPK, GPVI-mediated activation cascade, Interleukin receptor SHC signaling, PI3KAKT activation, PIP3 activates AKT signaling, Regulation of signaling by CBL, Role of LAT2NTALLAB on calcium mobilization, Signaling by ERBB2, Signaling by PDGF, Signaling by SCF-KIT, Synthesis of PIPs at the plasma membrane 
                                               SEBELIUS_2_C4636 
                                               PIK3R3 
                                               mRNA 
                                               0 
                                               
                                               
                                               G alpha (q) signalling events;PIP3 activates AKT signaling;Constitutive Signaling by Aberrant PI3K in Cancer;CD28 dependent PI3KAkt signaling;Synthesis of PIPs at the plasma membrane;Interleukin receptor SHC signaling;G beta.CL.gamma signalling through PI3Kgamma;Regulation of signaling by CBL;GPVI-mediated activation cascade;G alpha (1213) signalling events;Gastrin-CREB signalling pathway via PKC and MAPK;Downstream signal transduction;GAB1 signalosome;Signaling by SCF-KIT;PI3KAKT activation;Role of LAT2NTALLAB on calcium mobilization;DAP12 signaling;Signaling by PDGF;Signaling by ERBB2;CD28 co-stimulation 
                                               hsa04012;hsa04014;hsa04015;hsa04022;hsa04024;hsa04062;hsa04066;hsa04068;hsa04070;hsa04071;hsa04072;hsa04150;hsa04151;hsa04152;hsa04210;hsa04211;hsa04213;hsa04261;hsa04370;hsa04380;hsa04510;hsa04550;hsa04611;hsa04620;hsa04630;hsa04650;hsa04660;hsa04662;hsa04664;hsa04666;hsa04668;hsa04670;hsa04722;hsa04725;hsa04750;hsa04810;hsa04910;hsa04914;hsa04915;hsa04917;hsa04919;hsa04921;hsa04923;hsa04930;hsa04931;hsa04932;hsa04933;hsa04960;hsa04973;hsa05100;hsa05142;hsa05145;hsa05146;hsa05160;hsa05161;hsa05162;hsa05164;hsa05166;hsa05169;hsa05200;hsa05203;hsa05205;hsa05210;hsa05211;hsa05212;hsa05213;hsa05214;hsa05215;hsa05218;hsa05220;hsa05221;hsa05222;hsa05223;hsa05230;hsa05231 
                                               PIK3R3 
                                               NA 
                                               slateblue2 
                                               16 
                                               3.53307392996109 
                                             
                                             
                                               AKT2-mRNA 
                                               -0.2 
                                               0.0603 
                                               -0.318 
                                               -0.0816 
                                               0.871 
                                               0.802 
                                               0.945 
                                               0.00331 
                                               0.0593 
                                               loglinear 
                                               Activation of BAD and translocation to mitochondria , CD28 co-stimulation, DAP12 signaling, Deactivation of the beta-catenin transactivating complex, Downstream signal transduction, G beta.CL.gamma signalling through PI3Kgamma, GAB1 signalosome, GPVI-mediated activation cascade, IRS-related events triggered by IGF1R, PI3K Cascade, PI3KAKT activation, PIP3 activates AKT signaling, Role of LAT2NTALLAB on calcium mobilization, Signaling by ERBB2, Signaling by PDGF, Signaling by SCF-KIT, TP53 Regulates Metabolic Genes, VEGFA-VEGFR2 Pathway, VEGFR2 mediated vascular permeability 
                                               SEBELIUS_2_C4636 
                                               AKT2 
                                               mRNA 
                                               0 
                                               
                                               
                                               TP53 Regulates Metabolic Genes;PDE3B signalling;AKT phosphorylates targets in the cytosol;Negative regulation of the PI3KAKT network;Deactivation of the beta-catenin transactivating complex;Activation of BAD and translocation to mitochondria ;G beta.CL.gamma signalling through PI3Kgamma;AKT phosphorylates targets in the nucleus;VEGFR2 mediated vascular permeability;GPVI-mediated activation cascade;Signaling by ERBB2;PI3K Cascade;IRS-related events triggered by IGF1R;mTOR signalling;PIP3 activates AKT signaling;Downstream signal transduction;GAB1 signalosome;Signaling by SCF-KIT;PI3KAKT activation;Role of LAT2NTALLAB on calcium mobilization;DAP12 signaling;Signaling by PDGF;CD28 co-stimulation;VEGFA-VEGFR2 Pathway 
                                               hsa04010;hsa04012;hsa04014;hsa04015;hsa04022;hsa04024;hsa04062;hsa04066;hsa04068;hsa04071;hsa04072;hsa04150;hsa04151;hsa04152;hsa04210;hsa04211;hsa04213;hsa04261;hsa04370;hsa04380;hsa04510;hsa04530;hsa04550;hsa04611;hsa04620;hsa04630;hsa04660;hsa04662;hsa04664;hsa04666;hsa04668;hsa04722;hsa04725;hsa04728;hsa04910;hsa04914;hsa04915;hsa04917;hsa04919;hsa04920;hsa04922;hsa04923;hsa04931;hsa04932;hsa04933;hsa04973;hsa05142;hsa05145;hsa05152;hsa05160;hsa05161;hsa05162;hsa05164;hsa05166;hsa05169;hsa05200;hsa05205;hsa05210;hsa05211;hsa05212;hsa05213;hsa05214;hsa05215;hsa05218;hsa05220;hsa05221;hsa05222;hsa05223;hsa05230;hsa05231 
                                               AKT2 
                                               NA 
                                               slateblue3 
                                               16 
                                               -3.3167495854063 
                                             
                                             
                                               PAK1-mRNA 
                                               -0.455 
                                               0.163 
                                               -0.775 
                                               -0.135 
                                               0.729 
                                               0.584 
                                               0.91 
                                               0.011 
                                               0.164 
                                               lm.nb 
                                               Activation of Rac, CD209 (DC-SIGN) signaling, CD28 co-stimulation, CD28 dependent Vav1 pathway, EPHB-mediated forward signaling, Ephrin signaling, FCERI mediated MAPK activation, MAPK6MAPK4 signaling, Regulation of actin dynamics for phagocytic cup formation, RHO GTPases activate PAKs, Sema3A PAK dependent Axon repulsion, VEGFA-VEGFR2 Pathway, VEGFR2 mediated vascular permeability 
                                               SEBELIUS_2_C4636 
                                               PAK1 
                                               mRNA 
                                               0 
                                               
                                               
                                               EPHB-mediated forward signaling;Ephrin signaling;MAPK6MAPK4 signaling;DSCAM interactions;Activation of Rac;VEGFR2 mediated vascular permeability;RHO GTPases activate PAKs;Sema3A PAK dependent Axon repulsion;FCERI mediated MAPK activation;Regulation of actin dynamics for phagocytic cup formation;CD209 (DC-SIGN) signaling;CD28 dependent Vav1 pathway;VEGFA-VEGFR2 Pathway;CD28 co-stimulation 
                                               hsa04010;hsa04012;hsa04014;hsa04024;hsa04062;hsa04360;hsa04510;hsa04650;hsa04660;hsa04666;hsa04810;hsa05120;hsa05205;hsa05211 
                                               PAK1 
                                               NA 
                                               slateblue4 
                                               16 
                                               -2.79141104294479 
                                             
                                             
                                               AKT1-mRNA 
                                               -0.414 
                                               0.152 
                                               -0.711 
                                               -0.117 
                                               0.751 
                                               0.611 
                                               0.922 
                                               0.0125 
                                               0.18 
                                               lm.nb 
                                               Activation of BAD and translocation to mitochondria , CD28 co-stimulation, DAP12 signaling, Deactivation of the beta-catenin transactivating complex, Downstream signal transduction, G beta.CL.gamma signalling through PI3Kgamma, GAB1 signalosome, GPVI-mediated activation cascade, IRS-related events triggered by IGF1R, PI3K Cascade, PI3KAKT activation, PIP3 activates AKT signaling, Role of LAT2NTALLAB on calcium mobilization, Signaling by ERBB2, Signaling by PDGF, Signaling by SCF-KIT, TP53 Regulates Metabolic Genes, VEGFA-VEGFR2 Pathway, VEGFR2 mediated vascular permeability 
                                               SEBELIUS_2_C4636 
                                               AKT1 
                                               mRNA 
                                               0 
                                               
                                               
                                               TP53 Regulates Metabolic Genes;Deactivation of the beta-catenin transactivating complex;Activation of BAD and translocation to mitochondria ;eNOS activation;mTOR signalling;G beta.CL.gamma signalling through PI3Kgamma;KSRP (KHSRP) binds and destabilizes mRNA;AKT phosphorylates targets in the nucleus;VEGFR2 mediated vascular permeability;GPVI-mediated activation cascade;Tetrahydrobiopterin (BH4) synthesis# recycling# salvage and regulation;Butyrate Response Factor 1 (BRF1) binds and destabilizes mRNA;Signaling by ERBB2;PIP3 activates AKT signaling;Downstream signal transduction;GAB1 signalosome;Signaling by SCF-KIT;PI3KAKT activation;Role of LAT2NTALLAB on calcium mobilization;DAP12 signaling;Signaling by PDGF;CD28 co-stimulation;PI3K Cascade;IRS-related events triggered by IGF1R;VEGFA-VEGFR2 Pathway 
                                               hsa04010;hsa04012;hsa04014;hsa04015;hsa04022;hsa04024;hsa04062;hsa04066;hsa04068;hsa04071;hsa04072;hsa04150;hsa04151;hsa04152;hsa04210;hsa04211;hsa04213;hsa04261;hsa04370;hsa04380;hsa04510;hsa04530;hsa04550;hsa04611;hsa04620;hsa04630;hsa04660;hsa04662;hsa04664;hsa04666;hsa04668;hsa04722;hsa04725;hsa04728;hsa04910;hsa04914;hsa04915;hsa04917;hsa04919;hsa04920;hsa04922;hsa04923;hsa04931;hsa04932;hsa04933;hsa04973;hsa05142;hsa05145;hsa05152;hsa05160;hsa05161;hsa05162;hsa05164;hsa05166;hsa05169;hsa05200;hsa05205;hsa05210;hsa05211;hsa05212;hsa05213;hsa05214;hsa05215;hsa05218;hsa05220;hsa05221;hsa05222;hsa05223;hsa05230;hsa05231 
                                               AKT1 
                                               NA 
                                               slateblue4 
                                               16 
                                               -2.72368421052632 
                                             
                                             
                                               FYN-mRNA 
                                               0.275 
                                               0.11 
                                               0.0594 
                                               0.49 
                                               1.21 
                                               1.04 
                                               1.4 
                                               0.0207 
                                               0.272 
                                               loglinear 
                                               ARMS-mediated activation, CD209 (DC-SIGN) signaling, CD28 co-stimulation, CD28 dependent PI3KAkt signaling, CD28 dependent Vav1 pathway, Cell surface interactions at the vascular wall, Constitutive Signaling by Aberrant PI3K in Cancer, CTLA4 inhibitory signaling, DAP12 signaling, Downstream signal transduction, EPH-ephrin mediated repulsion of cells, FCERI mediated MAPK activation, Frs2-mediated activation, GAB1 signalosome, Gastrin-CREB signalling pathway via PKC and MAPK, GPVI-mediated activation cascade, Interleukin receptor SHC signaling, IRS-related events triggered by IGF1R, Nephrin interactions, PI3KAKT activation, RAFMAP kinase cascade, Regulation of signaling by CBL, Role of LAT2NTALLAB on calcium mobilization, Sema3A PAK dependent Axon repulsion, Signaling by ERBB2, Signaling by PDGF, Signaling by SCF-KIT, VEGFA-VEGFR2 Pathway, VEGFR2 mediated cell proliferation 
                                               SEBELIUS_2_C4636 
                                               FYN 
                                               mRNA 
                                               0 
                                               
                                               
                                               EPH-ephrin mediated repulsion of cells;Regulation of signaling by CBL;FCGR activation;Antigen activates B Cell Receptor (BCR) leading to generation of second messengers;Dectin-2 family;RAFMAP kinase cascade;Constitutive Signaling by Aberrant PI3K in Cancer;CTLA4 inhibitory signaling;CD28 dependent PI3KAkt signaling;PECAM1 interactions;CD28 dependent Vav1 pathway;GPVI-mediated activation cascade;Sema3A PAK dependent Axon repulsion;Nephrin interactions;SEMA3A-Plexin repulsion signaling by inhibiting Integrin adhesion;CRMPs in Sema3A signaling;CD209 (DC-SIGN) signaling;Downstream signal transduction;GAB1 signalosome;Signaling by SCF-KIT;PI3KAKT activation;Role of LAT2NTALLAB on calcium mobilization;DAP12 signaling;Signaling by PDGF;Signaling by ERBB2;VEGFA-VEGFR2 Pathway;VEGFR2 mediated cell proliferation;Frs2-mediated activation;ARMS-mediated activation;Gastrin-CREB signalling pathway via PKC and MAPK;Interleukin receptor SHC signaling;FCERI mediated MAPK activation;IRS-related events triggered by IGF1R;CD28 co-stimulation;Cell surface interactions at the vascular wall 
                                               hsa04071;hsa04072;hsa04360;hsa04380;hsa04510;hsa04520;hsa04611;hsa04650;hsa04660;hsa04664;hsa04725;hsa05020;hsa05130;hsa05162;hsa05416 
                                               FYN 
                                               NA 
                                               slateblue4 
                                               16 
                                               2.5 
                                             
                                             
                                               AKT3-mRNA 
                                               0.207 
                                               0.106 
                                               -0.000741 
                                               0.415 
                                               1.15 
                                               0.999 
                                               1.33 
                                               0.0643 
                                               0.723 
                                               lm.nb 
                                               Activation of BAD and translocation to mitochondria , CD28 co-stimulation, CD28 dependent PI3KAkt signaling, DAP12 signaling, Downstream signal transduction, G beta.CL.gamma signalling through PI3Kgamma, GAB1 signalosome, GPVI-mediated activation cascade, PI3KAKT activation, PIP3 activates AKT signaling, Role of LAT2NTALLAB on calcium mobilization, Signaling by ERBB2, Signaling by PDGF, Signaling by SCF-KIT, TP53 Regulates Metabolic Genes, VEGFA-VEGFR2 Pathway, VEGFR2 mediated vascular permeability 
                                               SEBELIUS_2_C4636 
                                               AKT3 
                                               mRNA 
                                               0 
                                               
                                               
                                               TP53 Regulates Metabolic Genes;CD28 dependent PI3KAkt signaling;AKT phosphorylates targets in the cytosol;Negative regulation of the PI3KAKT network;Activation of BAD and translocation to mitochondria ;G beta.CL.gamma signalling through PI3Kgamma;AKT phosphorylates targets in the nucleus;VEGFR2 mediated vascular permeability;GPVI-mediated activation cascade;Signaling by ERBB2;PIP3 activates AKT signaling;Downstream signal transduction;GAB1 signalosome;Signaling by SCF-KIT;PI3KAKT activation;Role of LAT2NTALLAB on calcium mobilization;DAP12 signaling;Signaling by PDGF;CD28 co-stimulation;VEGFA-VEGFR2 Pathway 
                                               hsa04010;hsa04012;hsa04014;hsa04015;hsa04022;hsa04024;hsa04062;hsa04066;hsa04068;hsa04071;hsa04072;hsa04150;hsa04151;hsa04152;hsa04210;hsa04211;hsa04213;hsa04261;hsa04370;hsa04380;hsa04510;hsa04530;hsa04550;hsa04611;hsa04620;hsa04630;hsa04660;hsa04662;hsa04664;hsa04666;hsa04668;hsa04722;hsa04725;hsa04728;hsa04910;hsa04914;hsa04915;hsa04917;hsa04919;hsa04920;hsa04922;hsa04923;hsa04931;hsa04932;hsa04933;hsa04973;hsa05142;hsa05145;hsa05152;hsa05160;hsa05161;hsa05162;hsa05164;hsa05166;hsa05169;hsa05200;hsa05205;hsa05210;hsa05211;hsa05212;hsa05213;hsa05214;hsa05215;hsa05218;hsa05220;hsa05221;hsa05222;hsa05223;hsa05230;hsa05231 
                                               AKT3 
                                               NA 
                                               azure4 
                                               1 
                                               1.95283018867925 
                                             
                                             
                                               PAK2-mRNA 
                                               0.147 
                                               0.0976 
                                               -0.0448 
                                               0.338 
                                               1.11 
                                               0.969 
                                               1.26 
                                               0.148 
                                               1 
                                               loglinear 
                                               Activation of Rac, CD209 (DC-SIGN) signaling, CD28 co-stimulation, CD28 dependent Vav1 pathway, Ephrin signaling, RHO GTPases activate PAKs, Sema3A PAK dependent Axon repulsion, VEGFA-VEGFR2 Pathway, VEGFR2 mediated vascular permeability 
                                               SEBELIUS_2_C4636 
                                               PAK2 
                                               mRNA 
                                               0 
                                               
                                               
                                               Nef and signal transduction;CD28 dependent Vav1 pathway;Stimulation of the cell death response by PAK-2p34;Ephrin signaling;Activation of Rac;RHO GTPases activate PAKs;VEGFR2 mediated vascular permeability;Sema3A PAK dependent Axon repulsion;CD209 (DC-SIGN) signaling;VEGFA-VEGFR2 Pathway;CD28 co-stimulation 
                                               hsa04010;hsa04012;hsa04014;hsa04360;hsa04510;hsa04660;hsa04810;hsa05211 
                                               PAK2 
                                               NA 
                                               azure4 
                                               1 
                                               1.50614754098361 
                                             
                                             
                                               CDC42-mRNA 
                                               0.103 
                                               0.0802 
                                               -0.0547 
                                               0.26 
                                               1.07 
                                               0.963 
                                               1.2 
                                               0.215 
                                               1 
                                               loglinear 
                                               CD28 co-stimulation, CD28 dependent Vav1 pathway, DCC mediated attractive signaling, EPHB-mediated forward signaling, G alpha (1213) signalling events, GPVI-mediated activation cascade, Regulation of actin dynamics for phagocytic cup formation, Rho GTPase cycle, RHO GTPases Activate Formins, RHO GTPases activate IQGAPs, RHO GTPases activate PAKs, Sema4D induced cell migration and growth-cone collapse, VEGFA-VEGFR2 Pathway 
                                               SEBELIUS_2_C4636 
                                               CDC42 
                                               mRNA 
                                               0 
                                               
                                               
                                               DCC mediated attractive signaling;Factors involved in megakaryocyte development and platelet production;VEGFA-VEGFR2 Pathway;RHO GTPases activate KTN1;CD28 dependent Vav1 pathway;Inactivation of Cdc42 and Rac;Rho GTPase cycle;EPHB-mediated forward signaling;RHO GTPases Activate Formins;RHO GTPases activate PAKs;GPVI-mediated activation cascade;G alpha (1213) signalling events;RHO GTPases activate IQGAPs;Sema4D induced cell migration and growth-cone collapse;Regulation of actin dynamics for phagocytic cup formation;CD28 co-stimulation 
                                               hsa04010;hsa04014;hsa04015;hsa04062;hsa04144;hsa04360;hsa04370;hsa04510;hsa04520;hsa04530;hsa04660;hsa04666;hsa04670;hsa04722;hsa04810;hsa04912;hsa04932;hsa04933;hsa05100;hsa05120;hsa05130;hsa05131;hsa05132;hsa05200;hsa05203;hsa05205;hsa05211;hsa05212 
                                               CDC42 
                                               NA 
                                               azure4 
                                               1 
                                               1.28428927680798 
                                             
                                             
                                               SRC-mRNA 
                                               0.138 
                                               0.114 
                                               -0.086 
                                               0.362 
                                               1.1 
                                               0.942 
                                               1.28 
                                               0.241 
                                               1 
                                               loglinear 
                                               ARMS-mediated activation, CD28 co-stimulation, Cell surface interactions at the vascular wall, CTLA4 inhibitory signaling, DAP12 signaling, Downstream signal transduction, FCERI mediated MAPK activation, Frs2-mediated activation, GAB1 signalosome, Gastrin-CREB signalling pathway via PKC and MAPK, Integrin alphaIIb beta3 signaling, Interleukin receptor SHC signaling, IRS-related events triggered by IGF1R, MAP2K and MAPK activation, Negative regulation of FGFR1 signaling, Negative regulation of FGFR2 signaling, Negative regulation of FGFR3 signaling, Negative regulation of FGFR4 signaling, p130Cas linkage to MAPK signaling for integrins, RAFMAP kinase cascade, Signaling by ERBB2, Signaling by PDGF, Signaling by SCF-KIT, Spry regulation of FGF signaling, VEGFA-VEGFR2 Pathway, VEGFR2 mediated cell proliferation 
                                               SEBELIUS_2_C4636 
                                               SRC 
                                               mRNA 
                                               0 
                                               
                                               
                                               FCGR activation;Spry regulation of FGF signaling;GAB1 signalosome;MAP2K and MAPK activation;CLEC7A (Dectin-1) signaling;CD28 co-stimulation;p130Cas linkage to MAPK signaling for integrins;CTLA4 inhibitory signaling;GRB2.CL.SOS provides linkage to MAPK signaling for Integrins ;Signal regulatory protein (SIRP) family interactions;PECAM1 interactions;Thrombin signalling through proteinase activated receptors (PARs);GP1b-IX-V activation signalling;c-src mediated regulation of Cx43 function and closure of gap junctions;Signaling by ERBB2;VEGFA-VEGFR2 Pathway;Signaling by SCF-KIT;VEGFR2 mediated cell proliferation;RAFMAP kinase cascade;Downstream signal transduction;Frs2-mediated activation;ARMS-mediated activation;Gastrin-CREB signalling pathway via PKC and MAPK;Interleukin receptor SHC signaling;FCERI mediated MAPK activation;DAP12 signaling;Signaling by PDGF;IRS-related events triggered by IGF1R;Negative regulation of FGFR3 signaling;Negative regulation of FGFR4 signaling;Negative regulation of FGFR2 signaling;Negative regulation of FGFR1 signaling;Integrin alphaIIb beta3 signaling;Cell surface interactions at the vascular wall 
                                               hsa04012;hsa04015;hsa04062;hsa04144;hsa04370;hsa04510;hsa04520;hsa04530;hsa04540;hsa04611;hsa04727;hsa04750;hsa04810;hsa04912;hsa04915;hsa04917;hsa04919;hsa04921;hsa05100;hsa05120;hsa05131;hsa05152;hsa05161;hsa05203;hsa05205;hsa05219 
                                               SRC 
                                               NA 
                                               azure4 
                                               1 
                                               1.21052631578947 
                                             
                                           
                                           DE Results - CD28 co-stimulation genes - New.Annotation: miR142 vs.cel67  Table displaying each sample's global significance scores and directed global significance scores as defined in the heatmaps above. The global significance score is calculated as the square root of the mean squared t-statistic for the genes in a gene set, with t-statistics coming from the linear regression underlying our differential expression analysis. The directed global significance score is calculated as the square root of the mean signed squared t-statistic for the genes in a gene set, with t-statistics coming from the linear regression underlying our differential expression analysis.   
                                      
                                       
                                  
                                
                                
                                  
                                       
                                      
                                           Volcano Plot: New.Annotation: miR155 vs.cel67   More Plot Information                Volcano Plot: New.Annotation: miR155 vs.cel67  Volcano plot displaying each gene's -log10(p-value) and log2 fold change for the selected covariate. Highly statistically significant genes fall at the top of the plot, and highly differentially expressed genes fall to either side. Genes within the selected gene set are highlighted in orange. Horizontal lines indicate various False Discovery Rate (FDR) thresholds.   
                                           DE Results - CD28 co-stimulation genes - New.Annotation: miR155 vs.cel67   More Plot Information      Download CSV Data     
                                           
                                             
                                             
                                           
                                           
                                             
                                               Probe Label 
                                               Log2 fold change 
                                               std error (log2) 
                                               Lower confidence limit (log2) 
                                               Upper confidence limit (log2) 
                                               Linear fold change 
                                               Lower confidence limit (linear) 
                                               Upper confidence limit (linear) 
                                               P-value 
                                               BY.p.value 
                                               method 
                                               Gene.sets 
                                               Codeset.Name 
                                               Probe.Label 
                                               Analyte.Type 
                                               Is.Control 
                                               Control.Type 
                                               Related.Probes 
                                               Probe.Annotation 
                                               KEGG.Pathways 
                                               Official.Gene.Name 
                                               Control.Conc 
                                               volcanocol 
                                               volcanopch 
                                               tstats.all 
                                             
                                           
                                           
                                             
                                               RAC1-mRNA 
                                               -0.507 
                                               0.0711 
                                               -0.647 
                                               -0.368 
                                               0.703 
                                               0.639 
                                               0.775 
                                               4.93e-07 
                                               8.18e-05 
                                               loglinear 
                                               Activation of Rac, CD28 co-stimulation, CD28 dependent Vav1 pathway, DAP12 signaling, EPH-ephrin mediated repulsion of cells, Ephrin signaling, G alpha (1213) signalling events, GPVI-mediated activation cascade, MAPK6MAPK4 signaling, NRAGE signals death through JNK, PCPCE pathway, Regulation of actin dynamics for phagocytic cup formation, RHO GTPases activate CIT, RHO GTPases Activate Formins, RHO GTPases activate IQGAPs, RHO GTPases activate PAKs, RHO GTPases activate PKNs, RHO GTPases Activate WASPs and WAVEs, Sema3A PAK dependent Axon repulsion, Sema4D induced cell migration and growth-cone collapse, Signal transduction by L1, Signaling by SCF-KIT, VEGFA-VEGFR2 Pathway, VEGFR2 mediated vascular permeability 
                                               SEBELIUS_2_C4636 
                                               RAC1 
                                               mRNA 
                                               0 
                                               
                                               
                                               MAPK6MAPK4 signaling;Signaling by SCF-KIT;Inactivation of Cdc42 and Rac;DAP12 signaling;Ephrin signaling;EPH-ephrin mediated repulsion of cells;Activation of Rac;RHO GTPases Activate Formins;VEGFR2 mediated vascular permeability;RHO GTPases activate PAKs;Sema4D mediated inhibition of cell attachment and migration;G alpha (1213) signalling events;Regulation of actin dynamics for phagocytic cup formation;Sema4D induced cell migration and growth-cone collapse;Signal transduction by L1;RHO GTPases activate CIT;RHO GTPases Activate WASPs and WAVEs;Factors involved in megakaryocyte development and platelet production;RHO GTPases activate KTN1;CD28 dependent Vav1 pathway;DSCAM interactions;NRAGE signals death through JNK;GPVI-mediated activation cascade;Sema3A PAK dependent Axon repulsion;RHO GTPases activate PKNs;SEMA3A-Plexin repulsion signaling by inhibiting Integrin adhesion;RHO GTPases activate IQGAPs;PCPCE pathway;VEGFA-VEGFR2 Pathway;CD28 co-stimulation 
                                               hsa04010;hsa04014;hsa04015;hsa04024;hsa04062;hsa04071;hsa04145;hsa04151;hsa04310;hsa04360;hsa04370;hsa04380;hsa04510;hsa04520;hsa04620;hsa04650;hsa04662;hsa04664;hsa04666;hsa04670;hsa04722;hsa04810;hsa04932;hsa04933;hsa04972;hsa05100;hsa05120;hsa05131;hsa05132;hsa05200;hsa05203;hsa05205;hsa05210;hsa05211;hsa05212;hsa05231;hsa05416 
                                               RAC1 
                                               NA 
                                               slateblue1 
                                               16 
                                               -7.13080168776371 
                                             
                                             
                                               PAK2-mRNA 
                                               -0.548 
                                               0.103 
                                               -0.75 
                                               -0.347 
                                               0.684 
                                               0.595 
                                               0.786 
                                               2.77e-05 
                                               0.00263 
                                               loglinear 
                                               Activation of Rac, CD209 (DC-SIGN) signaling, CD28 co-stimulation, CD28 dependent Vav1 pathway, Ephrin signaling, RHO GTPases activate PAKs, Sema3A PAK dependent Axon repulsion, VEGFA-VEGFR2 Pathway, VEGFR2 mediated vascular permeability 
                                               SEBELIUS_2_C4636 
                                               PAK2 
                                               mRNA 
                                               0 
                                               
                                               
                                               Nef and signal transduction;CD28 dependent Vav1 pathway;Stimulation of the cell death response by PAK-2p34;Ephrin signaling;Activation of Rac;RHO GTPases activate PAKs;VEGFR2 mediated vascular permeability;Sema3A PAK dependent Axon repulsion;CD209 (DC-SIGN) signaling;VEGFA-VEGFR2 Pathway;CD28 co-stimulation 
                                               hsa04010;hsa04012;hsa04014;hsa04360;hsa04510;hsa04660;hsa04810;hsa05211 
                                               PAK2 
                                               NA 
                                               slateblue1 
                                               16 
                                               -5.32038834951456 
                                             
                                             
                                               AKT3-mRNA 
                                               -0.395 
                                               0.112 
                                               -0.616 
                                               -0.175 
                                               0.76 
                                               0.653 
                                               0.886 
                                               0.00206 
                                               0.0829 
                                               lm.nb 
                                               Activation of BAD and translocation to mitochondria , CD28 co-stimulation, CD28 dependent PI3KAkt signaling, DAP12 signaling, Downstream signal transduction, G beta.CL.gamma signalling through PI3Kgamma, GAB1 signalosome, GPVI-mediated activation cascade, PI3KAKT activation, PIP3 activates AKT signaling, Role of LAT2NTALLAB on calcium mobilization, Signaling by ERBB2, Signaling by PDGF, Signaling by SCF-KIT, TP53 Regulates Metabolic Genes, VEGFA-VEGFR2 Pathway, VEGFR2 mediated vascular permeability 
                                               SEBELIUS_2_C4636 
                                               AKT3 
                                               mRNA 
                                               0 
                                               
                                               
                                               TP53 Regulates Metabolic Genes;CD28 dependent PI3KAkt signaling;AKT phosphorylates targets in the cytosol;Negative regulation of the PI3KAKT network;Activation of BAD and translocation to mitochondria ;G beta.CL.gamma signalling through PI3Kgamma;AKT phosphorylates targets in the nucleus;VEGFR2 mediated vascular permeability;GPVI-mediated activation cascade;Signaling by ERBB2;PIP3 activates AKT signaling;Downstream signal transduction;GAB1 signalosome;Signaling by SCF-KIT;PI3KAKT activation;Role of LAT2NTALLAB on calcium mobilization;DAP12 signaling;Signaling by PDGF;CD28 co-stimulation;VEGFA-VEGFR2 Pathway 
                                               hsa04010;hsa04012;hsa04014;hsa04015;hsa04022;hsa04024;hsa04062;hsa04066;hsa04068;hsa04071;hsa04072;hsa04150;hsa04151;hsa04152;hsa04210;hsa04211;hsa04213;hsa04261;hsa04370;hsa04380;hsa04510;hsa04530;hsa04550;hsa04611;hsa04620;hsa04630;hsa04660;hsa04662;hsa04664;hsa04666;hsa04668;hsa04722;hsa04725;hsa04728;hsa04910;hsa04914;hsa04915;hsa04917;hsa04919;hsa04920;hsa04922;hsa04923;hsa04931;hsa04932;hsa04933;hsa04973;hsa05142;hsa05145;hsa05152;hsa05160;hsa05161;hsa05162;hsa05164;hsa05166;hsa05169;hsa05200;hsa05205;hsa05210;hsa05211;hsa05212;hsa05213;hsa05214;hsa05215;hsa05218;hsa05220;hsa05221;hsa05222;hsa05223;hsa05230;hsa05231 
                                               AKT3 
                                               NA 
                                               slateblue3 
                                               16 
                                               -3.52678571428571 
                                             
                                             
                                               PIK3CA-mRNA 
                                               -0.344 
                                               0.107 
                                               -0.554 
                                               -0.134 
                                               0.788 
                                               0.681 
                                               0.911 
                                               0.00418 
                                               0.125 
                                               loglinear 
                                               CD28 co-stimulation, CD28 dependent PI3KAkt signaling, Cell surface interactions at the vascular wall, Constitutive Signaling by Aberrant PI3K in Cancer, Constitutive Signaling by EGFRvIII, DAP12 signaling, Downstream signal transduction, Downstream TCR signaling, G alpha (1213) signalling events, G alpha (q) signalling events, G beta.CL.gamma signalling through PI3Kgamma, GAB1 signalosome, Gastrin-CREB signalling pathway via PKC and MAPK, GPVI-mediated activation cascade, Interleukin receptor SHC signaling, IRS-related events triggered by IGF1R, Nephrin interactions, PI3K Cascade, PI3KAKT activation, PIP3 activates AKT signaling, Regulation of signaling by CBL, Role of LAT2NTALLAB on calcium mobilization, Signaling by ERBB2, Signaling by FGFR1 in disease, Signaling by FGFR2 in disease, Signaling by FGFR3 in disease, Signaling by FGFR4 in disease, Signaling by PDGF, Signaling by SCF-KIT, Synthesis of PIPs at the plasma membrane, Tie2 Signaling, VEGFA-VEGFR2 Pathway 
                                               SEBELIUS_2_C4636 
                                               PIK3CA 
                                               mRNA 
                                               0 
                                               
                                               
                                               G alpha (q) signalling events;Downstream TCR signaling;Signaling by FGFR2 in disease;PIP3 activates AKT signaling;VEGFA-VEGFR2 Pathway;Synthesis of PIPs at the plasma membrane;Constitutive Signaling by Ligand-Responsive EGFR Cancer Variants;Interleukin receptor SHC signaling;G beta.CL.gamma signalling through PI3Kgamma;Regulation of signaling by CBL;Role of phospholipids in phagocytosis;G alpha (1213) signalling events;PI3K Cascade;Signaling by FGFR3 in disease;Constitutive Signaling by Aberrant PI3K in Cancer;Signaling by FGFR4 in disease;CD28 dependent PI3KAkt signaling;Tie2 Signaling;GPVI-mediated activation cascade;Nephrin interactions;Constitutive Signaling by EGFRvIII;Gastrin-CREB signalling pathway via PKC and MAPK;Downstream signal transduction;GAB1 signalosome;Signaling by SCF-KIT;PI3KAKT activation;Role of LAT2NTALLAB on calcium mobilization;DAP12 signaling;Signaling by PDGF;Signaling by ERBB2;IRS-related events triggered by IGF1R;CD28 co-stimulation;Cell surface interactions at the vascular wall;Signaling by FGFR1 in disease 
                                               hsa00562;hsa04012;hsa04014;hsa04015;hsa04022;hsa04024;hsa04062;hsa04066;hsa04068;hsa04070;hsa04071;hsa04072;hsa04150;hsa04151;hsa04152;hsa04210;hsa04211;hsa04213;hsa04261;hsa04370;hsa04380;hsa04510;hsa04550;hsa04611;hsa04620;hsa04630;hsa04650;hsa04660;hsa04662;hsa04664;hsa04666;hsa04668;hsa04670;hsa04722;hsa04725;hsa04750;hsa04810;hsa04910;hsa04914;hsa04915;hsa04917;hsa04919;hsa04921;hsa04923;hsa04930;hsa04931;hsa04932;hsa04933;hsa04960;hsa04973;hsa05100;hsa05142;hsa05145;hsa05146;hsa05160;hsa05161;hsa05162;hsa05164;hsa05166;hsa05169;hsa05200;hsa05203;hsa05205;hsa05206;hsa05210;hsa05211;hsa05212;hsa05213;hsa05214;hsa05215;hsa05218;hsa05220;hsa05221;hsa05222;hsa05223;hsa05230;hsa05231 
                                               PIK3CA 
                                               NA 
                                               slateblue4 
                                               16 
                                               -3.21495327102804 
                                             
                                             
                                               PIK3R2-mRNA 
                                               -0.434 
                                               0.14 
                                               -0.709 
                                               -0.16 
                                               0.74 
                                               0.612 
                                               0.895 
                                               0.00541 
                                               0.153 
                                               lm.nb 
                                               CD28 co-stimulation, CD28 dependent PI3KAkt signaling, Cell surface interactions at the vascular wall, Constitutive Signaling by Aberrant PI3K in Cancer, DAP12 signaling, Downstream signal transduction, Downstream TCR signaling, G alpha (1213) signalling events, G alpha (q) signalling events, G beta.CL.gamma signalling through PI3Kgamma, GAB1 signalosome, Gastrin-CREB signalling pathway via PKC and MAPK, GPVI-mediated activation cascade, Interleukin receptor SHC signaling, IRS-related events triggered by IGF1R, Nephrin interactions, PI3K Cascade, PI3KAKT activation, PIP3 activates AKT signaling, Regulation of signaling by CBL, Rho GTPase cycle, Role of LAT2NTALLAB on calcium mobilization, Signaling by ERBB2, Signaling by PDGF, Signaling by SCF-KIT, Synthesis of PIPs at the plasma membrane, Tie2 Signaling, VEGFA-VEGFR2 Pathway 
                                               SEBELIUS_2_C4636 
                                               PIK3R2 
                                               mRNA 
                                               0 
                                               
                                               
                                               G alpha (q) signalling events;Downstream TCR signaling;PIP3 activates AKT signaling;VEGFA-VEGFR2 Pathway;Synthesis of PIPs at the plasma membrane;Rho GTPase cycle;Interleukin receptor SHC signaling;G beta.CL.gamma signalling through PI3Kgamma;Regulation of signaling by CBL;Role of phospholipids in phagocytosis;G alpha (1213) signalling events;PI3K Cascade;Constitutive Signaling by Aberrant PI3K in Cancer;CD28 dependent PI3KAkt signaling;Tie2 Signaling;GPVI-mediated activation cascade;Nephrin interactions;Gastrin-CREB signalling pathway via PKC and MAPK;Downstream signal transduction;GAB1 signalosome;Signaling by SCF-KIT;PI3KAKT activation;Role of LAT2NTALLAB on calcium mobilization;DAP12 signaling;Signaling by PDGF;Signaling by ERBB2;IRS-related events triggered by IGF1R;CD28 co-stimulation;Cell surface interactions at the vascular wall 
                                               hsa04012;hsa04014;hsa04015;hsa04022;hsa04024;hsa04062;hsa04066;hsa04068;hsa04070;hsa04071;hsa04072;hsa04150;hsa04151;hsa04152;hsa04210;hsa04211;hsa04213;hsa04261;hsa04370;hsa04380;hsa04510;hsa04550;hsa04611;hsa04620;hsa04630;hsa04650;hsa04660;hsa04662;hsa04664;hsa04666;hsa04668;hsa04670;hsa04722;hsa04725;hsa04750;hsa04810;hsa04910;hsa04914;hsa04915;hsa04917;hsa04919;hsa04921;hsa04923;hsa04930;hsa04931;hsa04932;hsa04933;hsa04960;hsa04973;hsa05100;hsa05142;hsa05145;hsa05146;hsa05160;hsa05161;hsa05162;hsa05164;hsa05166;hsa05169;hsa05200;hsa05203;hsa05205;hsa05206;hsa05210;hsa05211;hsa05212;hsa05213;hsa05214;hsa05215;hsa05218;hsa05220;hsa05221;hsa05222;hsa05223;hsa05230;hsa05231 
                                               PIK3R2 
                                               NA 
                                               slateblue4 
                                               16 
                                               -3.1 
                                             
                                             
                                               AKT2-mRNA 
                                               -0.187 
                                               0.0636 
                                               -0.311 
                                               -0.062 
                                               0.879 
                                               0.806 
                                               0.958 
                                               0.00791 
                                               0.21 
                                               loglinear 
                                               Activation of BAD and translocation to mitochondria , CD28 co-stimulation, DAP12 signaling, Deactivation of the beta-catenin transactivating complex, Downstream signal transduction, G beta.CL.gamma signalling through PI3Kgamma, GAB1 signalosome, GPVI-mediated activation cascade, IRS-related events triggered by IGF1R, PI3K Cascade, PI3KAKT activation, PIP3 activates AKT signaling, Role of LAT2NTALLAB on calcium mobilization, Signaling by ERBB2, Signaling by PDGF, Signaling by SCF-KIT, TP53 Regulates Metabolic Genes, VEGFA-VEGFR2 Pathway, VEGFR2 mediated vascular permeability 
                                               SEBELIUS_2_C4636 
                                               AKT2 
                                               mRNA 
                                               0 
                                               
                                               
                                               TP53 Regulates Metabolic Genes;PDE3B signalling;AKT phosphorylates targets in the cytosol;Negative regulation of the PI3KAKT network;Deactivation of the beta-catenin transactivating complex;Activation of BAD and translocation to mitochondria ;G beta.CL.gamma signalling through PI3Kgamma;AKT phosphorylates targets in the nucleus;VEGFR2 mediated vascular permeability;GPVI-mediated activation cascade;Signaling by ERBB2;PI3K Cascade;IRS-related events triggered by IGF1R;mTOR signalling;PIP3 activates AKT signaling;Downstream signal transduction;GAB1 signalosome;Signaling by SCF-KIT;PI3KAKT activation;Role of LAT2NTALLAB on calcium mobilization;DAP12 signaling;Signaling by PDGF;CD28 co-stimulation;VEGFA-VEGFR2 Pathway 
                                               hsa04010;hsa04012;hsa04014;hsa04015;hsa04022;hsa04024;hsa04062;hsa04066;hsa04068;hsa04071;hsa04072;hsa04150;hsa04151;hsa04152;hsa04210;hsa04211;hsa04213;hsa04261;hsa04370;hsa04380;hsa04510;hsa04530;hsa04550;hsa04611;hsa04620;hsa04630;hsa04660;hsa04662;hsa04664;hsa04666;hsa04668;hsa04722;hsa04725;hsa04728;hsa04910;hsa04914;hsa04915;hsa04917;hsa04919;hsa04920;hsa04922;hsa04923;hsa04931;hsa04932;hsa04933;hsa04973;hsa05142;hsa05145;hsa05152;hsa05160;hsa05161;hsa05162;hsa05164;hsa05166;hsa05169;hsa05200;hsa05205;hsa05210;hsa05211;hsa05212;hsa05213;hsa05214;hsa05215;hsa05218;hsa05220;hsa05221;hsa05222;hsa05223;hsa05230;hsa05231 
                                               AKT2 
                                               NA 
                                               slateblue4 
                                               16 
                                               -2.94025157232704 
                                             
                                             
                                               PIK3R3-mRNA 
                                               0.72 
                                               0.271 
                                               0.188 
                                               1.25 
                                               1.65 
                                               1.14 
                                               2.38 
                                               0.0149 
                                               0.309 
                                               lm.nb 
                                               CD28 co-stimulation, CD28 dependent PI3KAkt signaling, Constitutive Signaling by Aberrant PI3K in Cancer, DAP12 signaling, Downstream signal transduction, G alpha (1213) signalling events, G alpha (q) signalling events, G beta.CL.gamma signalling through PI3Kgamma, GAB1 signalosome, Gastrin-CREB signalling pathway via PKC and MAPK, GPVI-mediated activation cascade, Interleukin receptor SHC signaling, PI3KAKT activation, PIP3 activates AKT signaling, Regulation of signaling by CBL, Role of LAT2NTALLAB on calcium mobilization, Signaling by ERBB2, Signaling by PDGF, Signaling by SCF-KIT, Synthesis of PIPs at the plasma membrane 
                                               SEBELIUS_2_C4636 
                                               PIK3R3 
                                               mRNA 
                                               0 
                                               
                                               
                                               G alpha (q) signalling events;PIP3 activates AKT signaling;Constitutive Signaling by Aberrant PI3K in Cancer;CD28 dependent PI3KAkt signaling;Synthesis of PIPs at the plasma membrane;Interleukin receptor SHC signaling;G beta.CL.gamma signalling through PI3Kgamma;Regulation of signaling by CBL;GPVI-mediated activation cascade;G alpha (1213) signalling events;Gastrin-CREB signalling pathway via PKC and MAPK;Downstream signal transduction;GAB1 signalosome;Signaling by SCF-KIT;PI3KAKT activation;Role of LAT2NTALLAB on calcium mobilization;DAP12 signaling;Signaling by PDGF;Signaling by ERBB2;CD28 co-stimulation 
                                               hsa04012;hsa04014;hsa04015;hsa04022;hsa04024;hsa04062;hsa04066;hsa04068;hsa04070;hsa04071;hsa04072;hsa04150;hsa04151;hsa04152;hsa04210;hsa04211;hsa04213;hsa04261;hsa04370;hsa04380;hsa04510;hsa04550;hsa04611;hsa04620;hsa04630;hsa04650;hsa04660;hsa04662;hsa04664;hsa04666;hsa04668;hsa04670;hsa04722;hsa04725;hsa04750;hsa04810;hsa04910;hsa04914;hsa04915;hsa04917;hsa04919;hsa04921;hsa04923;hsa04930;hsa04931;hsa04932;hsa04933;hsa04960;hsa04973;hsa05100;hsa05142;hsa05145;hsa05146;hsa05160;hsa05161;hsa05162;hsa05164;hsa05166;hsa05169;hsa05200;hsa05203;hsa05205;hsa05210;hsa05211;hsa05212;hsa05213;hsa05214;hsa05215;hsa05218;hsa05220;hsa05221;hsa05222;hsa05223;hsa05230;hsa05231 
                                               PIK3R3 
                                               NA 
                                               slateblue4 
                                               16 
                                               2.65682656826568 
                                             
                                             
                                               YES1-mRNA 
                                               -0.105 
                                               0.0754 
                                               -0.253 
                                               0.0432 
                                               0.93 
                                               0.839 
                                               1.03 
                                               0.18 
                                               1 
                                               loglinear 
                                               CD28 co-stimulation, Cell surface interactions at the vascular wall, CTLA4 inhibitory signaling, EPH-ephrin mediated repulsion of cells, EPHA-mediated growth cone collapse, EPHB-mediated forward signaling, Regulation of signaling by CBL, Signaling by ERBB2, Signaling by SCF-KIT 
                                               SEBELIUS_2_C4636 
                                               YES1 
                                               mRNA 
                                               0 
                                               
                                               
                                               Signaling by ERBB2;EPHB-mediated forward signaling;EPHA-mediated growth cone collapse;Regulation of KIT signaling;EPH-ephrin mediated repulsion of cells;Regulation of signaling by CBL;CD28 co-stimulation;CTLA4 inhibitory signaling;Signaling by SCF-KIT;Cell surface interactions at the vascular wall 
                                               hsa04520;hsa04530 
                                               YES1 
                                               NA 
                                               azure4 
                                               1 
                                               -1.39257294429708 
                                             
                                             
                                               AKT1-mRNA 
                                               -0.165 
                                               0.159 
                                               -0.478 
                                               0.147 
                                               0.892 
                                               0.718 
                                               1.11 
                                               0.311 
                                               1 
                                               lm.nb 
                                               Activation of BAD and translocation to mitochondria , CD28 co-stimulation, DAP12 signaling, Deactivation of the beta-catenin transactivating complex, Downstream signal transduction, G beta.CL.gamma signalling through PI3Kgamma, GAB1 signalosome, GPVI-mediated activation cascade, IRS-related events triggered by IGF1R, PI3K Cascade, PI3KAKT activation, PIP3 activates AKT signaling, Role of LAT2NTALLAB on calcium mobilization, Signaling by ERBB2, Signaling by PDGF, Signaling by SCF-KIT, TP53 Regulates Metabolic Genes, VEGFA-VEGFR2 Pathway, VEGFR2 mediated vascular permeability 
                                               SEBELIUS_2_C4636 
                                               AKT1 
                                               mRNA 
                                               0 
                                               
                                               
                                               TP53 Regulates Metabolic Genes;Deactivation of the beta-catenin transactivating complex;Activation of BAD and translocation to mitochondria ;eNOS activation;mTOR signalling;G beta.CL.gamma signalling through PI3Kgamma;KSRP (KHSRP) binds and destabilizes mRNA;AKT phosphorylates targets in the nucleus;VEGFR2 mediated vascular permeability;GPVI-mediated activation cascade;Tetrahydrobiopterin (BH4) synthesis# recycling# salvage and regulation;Butyrate Response Factor 1 (BRF1) binds and destabilizes mRNA;Signaling by ERBB2;PIP3 activates AKT signaling;Downstream signal transduction;GAB1 signalosome;Signaling by SCF-KIT;PI3KAKT activation;Role of LAT2NTALLAB on calcium mobilization;DAP12 signaling;Signaling by PDGF;CD28 co-stimulation;PI3K Cascade;IRS-related events triggered by IGF1R;VEGFA-VEGFR2 Pathway 
                                               hsa04010;hsa04012;hsa04014;hsa04015;hsa04022;hsa04024;hsa04062;hsa04066;hsa04068;hsa04071;hsa04072;hsa04150;hsa04151;hsa04152;hsa04210;hsa04211;hsa04213;hsa04261;hsa04370;hsa04380;hsa04510;hsa04530;hsa04550;hsa04611;hsa04620;hsa04630;hsa04660;hsa04662;hsa04664;hsa04666;hsa04668;hsa04722;hsa04725;hsa04728;hsa04910;hsa04914;hsa04915;hsa04917;hsa04919;hsa04920;hsa04922;hsa04923;hsa04931;hsa04932;hsa04933;hsa04973;hsa05142;hsa05145;hsa05152;hsa05160;hsa05161;hsa05162;hsa05164;hsa05166;hsa05169;hsa05200;hsa05205;hsa05210;hsa05211;hsa05212;hsa05213;hsa05214;hsa05215;hsa05218;hsa05220;hsa05221;hsa05222;hsa05223;hsa05230;hsa05231 
                                               AKT1 
                                               NA 
                                               azure4 
                                               1 
                                               -1.0377358490566 
                                             
                                             
                                               PAK1-mRNA 
                                               -0.105 
                                               0.172 
                                               -0.441 
                                               0.231 
                                               0.93 
                                               0.736 
                                               1.17 
                                               0.547 
                                               1 
                                               lm.nb 
                                               Activation of Rac, CD209 (DC-SIGN) signaling, CD28 co-stimulation, CD28 dependent Vav1 pathway, EPHB-mediated forward signaling, Ephrin signaling, FCERI mediated MAPK activation, MAPK6MAPK4 signaling, Regulation of actin dynamics for phagocytic cup formation, RHO GTPases activate PAKs, Sema3A PAK dependent Axon repulsion, VEGFA-VEGFR2 Pathway, VEGFR2 mediated vascular permeability 
                                               SEBELIUS_2_C4636 
                                               PAK1 
                                               mRNA 
                                               0 
                                               
                                               
                                               EPHB-mediated forward signaling;Ephrin signaling;MAPK6MAPK4 signaling;DSCAM interactions;Activation of Rac;VEGFR2 mediated vascular permeability;RHO GTPases activate PAKs;Sema3A PAK dependent Axon repulsion;FCERI mediated MAPK activation;Regulation of actin dynamics for phagocytic cup formation;CD209 (DC-SIGN) signaling;CD28 dependent Vav1 pathway;VEGFA-VEGFR2 Pathway;CD28 co-stimulation 
                                               hsa04010;hsa04012;hsa04014;hsa04024;hsa04062;hsa04360;hsa04510;hsa04650;hsa04660;hsa04666;hsa04810;hsa05120;hsa05205;hsa05211 
                                               PAK1 
                                               NA 
                                               azure4 
                                               1 
                                               -0.61046511627907 
                                             
                                             
                                               SRC-mRNA 
                                               0.0464 
                                               0.12 
                                               -0.19 
                                               0.282 
                                               1.03 
                                               0.877 
                                               1.22 
                                               0.704 
                                               1 
                                               loglinear 
                                               ARMS-mediated activation, CD28 co-stimulation, Cell surface interactions at the vascular wall, CTLA4 inhibitory signaling, DAP12 signaling, Downstream signal transduction, FCERI mediated MAPK activation, Frs2-mediated activation, GAB1 signalosome, Gastrin-CREB signalling pathway via PKC and MAPK, Integrin alphaIIb beta3 signaling, Interleukin receptor SHC signaling, IRS-related events triggered by IGF1R, MAP2K and MAPK activation, Negative regulation of FGFR1 signaling, Negative regulation of FGFR2 signaling, Negative regulation of FGFR3 signaling, Negative regulation of FGFR4 signaling, p130Cas linkage to MAPK signaling for integrins, RAFMAP kinase cascade, Signaling by ERBB2, Signaling by PDGF, Signaling by SCF-KIT, Spry regulation of FGF signaling, VEGFA-VEGFR2 Pathway, VEGFR2 mediated cell proliferation 
                                               SEBELIUS_2_C4636 
                                               SRC 
                                               mRNA 
                                               0 
                                               
                                               
                                               FCGR activation;Spry regulation of FGF signaling;GAB1 signalosome;MAP2K and MAPK activation;CLEC7A (Dectin-1) signaling;CD28 co-stimulation;p130Cas linkage to MAPK signaling for integrins;CTLA4 inhibitory signaling;GRB2.CL.SOS provides linkage to MAPK signaling for Integrins ;Signal regulatory protein (SIRP) family interactions;PECAM1 interactions;Thrombin signalling through proteinase activated receptors (PARs);GP1b-IX-V activation signalling;c-src mediated regulation of Cx43 function and closure of gap junctions;Signaling by ERBB2;VEGFA-VEGFR2 Pathway;Signaling by SCF-KIT;VEGFR2 mediated cell proliferation;RAFMAP kinase cascade;Downstream signal transduction;Frs2-mediated activation;ARMS-mediated activation;Gastrin-CREB signalling pathway via PKC and MAPK;Interleukin receptor SHC signaling;FCERI mediated MAPK activation;DAP12 signaling;Signaling by PDGF;IRS-related events triggered by IGF1R;Negative regulation of FGFR3 signaling;Negative regulation of FGFR4 signaling;Negative regulation of FGFR2 signaling;Negative regulation of FGFR1 signaling;Integrin alphaIIb beta3 signaling;Cell surface interactions at the vascular wall 
                                               hsa04012;hsa04015;hsa04062;hsa04144;hsa04370;hsa04510;hsa04520;hsa04530;hsa04540;hsa04611;hsa04727;hsa04750;hsa04810;hsa04912;hsa04915;hsa04917;hsa04919;hsa04921;hsa05100;hsa05120;hsa05131;hsa05152;hsa05161;hsa05203;hsa05205;hsa05219 
                                               SRC 
                                               NA 
                                               azure4 
                                               1 
                                               0.386666666666667 
                                             
                                             
                                               CDC42-mRNA 
                                               -0.0271 
                                               0.0845 
                                               -0.193 
                                               0.139 
                                               0.981 
                                               0.875 
                                               1.1 
                                               0.752 
                                               1 
                                               loglinear 
                                               CD28 co-stimulation, CD28 dependent Vav1 pathway, DCC mediated attractive signaling, EPHB-mediated forward signaling, G alpha (1213) signalling events, GPVI-mediated activation cascade, Regulation of actin dynamics for phagocytic cup formation, Rho GTPase cycle, RHO GTPases Activate Formins, RHO GTPases activate IQGAPs, RHO GTPases activate PAKs, Sema4D induced cell migration and growth-cone collapse, VEGFA-VEGFR2 Pathway 
                                               SEBELIUS_2_C4636 
                                               CDC42 
                                               mRNA 
                                               0 
                                               
                                               
                                               DCC mediated attractive signaling;Factors involved in megakaryocyte development and platelet production;VEGFA-VEGFR2 Pathway;RHO GTPases activate KTN1;CD28 dependent Vav1 pathway;Inactivation of Cdc42 and Rac;Rho GTPase cycle;EPHB-mediated forward signaling;RHO GTPases Activate Formins;RHO GTPases activate PAKs;GPVI-mediated activation cascade;G alpha (1213) signalling events;RHO GTPases activate IQGAPs;Sema4D induced cell migration and growth-cone collapse;Regulation of actin dynamics for phagocytic cup formation;CD28 co-stimulation 
                                               hsa04010;hsa04014;hsa04015;hsa04062;hsa04144;hsa04360;hsa04370;hsa04510;hsa04520;hsa04530;hsa04660;hsa04666;hsa04670;hsa04722;hsa04810;hsa04912;hsa04932;hsa04933;hsa05100;hsa05120;hsa05130;hsa05131;hsa05132;hsa05200;hsa05203;hsa05205;hsa05211;hsa05212 
                                               CDC42 
                                               NA 
                                               azure4 
                                               1 
                                               -0.320710059171598 
                                             
                                             
                                               FYN-mRNA 
                                               0.0321 
                                               0.116 
                                               -0.195 
                                               0.259 
                                               1.02 
                                               0.874 
                                               1.2 
                                               0.784 
                                               1 
                                               loglinear 
                                               ARMS-mediated activation, CD209 (DC-SIGN) signaling, CD28 co-stimulation, CD28 dependent PI3KAkt signaling, CD28 dependent Vav1 pathway, Cell surface interactions at the vascular wall, Constitutive Signaling by Aberrant PI3K in Cancer, CTLA4 inhibitory signaling, DAP12 signaling, Downstream signal transduction, EPH-ephrin mediated repulsion of cells, FCERI mediated MAPK activation, Frs2-mediated activation, GAB1 signalosome, Gastrin-CREB signalling pathway via PKC and MAPK, GPVI-mediated activation cascade, Interleukin receptor SHC signaling, IRS-related events triggered by IGF1R, Nephrin interactions, PI3KAKT activation, RAFMAP kinase cascade, Regulation of signaling by CBL, Role of LAT2NTALLAB on calcium mobilization, Sema3A PAK dependent Axon repulsion, Signaling by ERBB2, Signaling by PDGF, Signaling by SCF-KIT, VEGFA-VEGFR2 Pathway, VEGFR2 mediated cell proliferation 
                                               SEBELIUS_2_C4636 
                                               FYN 
                                               mRNA 
                                               0 
                                               
                                               
                                               EPH-ephrin mediated repulsion of cells;Regulation of signaling by CBL;FCGR activation;Antigen activates B Cell Receptor (BCR) leading to generation of second messengers;Dectin-2 family;RAFMAP kinase cascade;Constitutive Signaling by Aberrant PI3K in Cancer;CTLA4 inhibitory signaling;CD28 dependent PI3KAkt signaling;PECAM1 interactions;CD28 dependent Vav1 pathway;GPVI-mediated activation cascade;Sema3A PAK dependent Axon repulsion;Nephrin interactions;SEMA3A-Plexin repulsion signaling by inhibiting Integrin adhesion;CRMPs in Sema3A signaling;CD209 (DC-SIGN) signaling;Downstream signal transduction;GAB1 signalosome;Signaling by SCF-KIT;PI3KAKT activation;Role of LAT2NTALLAB on calcium mobilization;DAP12 signaling;Signaling by PDGF;Signaling by ERBB2;VEGFA-VEGFR2 Pathway;VEGFR2 mediated cell proliferation;Frs2-mediated activation;ARMS-mediated activation;Gastrin-CREB signalling pathway via PKC and MAPK;Interleukin receptor SHC signaling;FCERI mediated MAPK activation;IRS-related events triggered by IGF1R;CD28 co-stimulation;Cell surface interactions at the vascular wall 
                                               hsa04071;hsa04072;hsa04360;hsa04380;hsa04510;hsa04520;hsa04611;hsa04650;hsa04660;hsa04664;hsa04725;hsa05020;hsa05130;hsa05162;hsa05416 
                                               FYN 
                                               NA 
                                               azure4 
                                               1 
                                               0.276724137931034 
                                             
                                             
                                               PIK3R1-mRNA 
                                               -0.0225 
                                               0.126 
                                               -0.269 
                                               0.224 
                                               0.985 
                                               0.83 
                                               1.17 
                                               0.86 
                                               1 
                                               loglinear 
                                               CD28 co-stimulation, CD28 dependent PI3KAkt signaling, Cell surface interactions at the vascular wall, Constitutive Signaling by Aberrant PI3K in Cancer, Constitutive Signaling by EGFRvIII, DAP12 signaling, Downstream signal transduction, Downstream TCR signaling, G alpha (q) signalling events, G beta.CL.gamma signalling through PI3Kgamma, GAB1 signalosome, Gastrin-CREB signalling pathway via PKC and MAPK, GPVI-mediated activation cascade, Interleukin receptor SHC signaling, IRS-related events triggered by IGF1R, Nephrin interactions, PI3KAKT activation, PIP3 activates AKT signaling, Regulation of signaling by CBL, Role of LAT2NTALLAB on calcium mobilization, Signaling by ERBB2, Signaling by FGFR1 in disease, Signaling by FGFR2 in disease, Signaling by FGFR3 in disease, Signaling by FGFR4 in disease, Signaling by PDGF, Signaling by SCF-KIT, Synthesis of PIPs at the plasma membrane, Tie2 Signaling, VEGFA-VEGFR2 Pathway 
                                               SEBELIUS_2_C4636 
                                               PIK3R1 
                                               mRNA 
                                               0 
                                               
                                               
                                               G alpha (q) signalling events;Downstream TCR signaling;Signaling by FGFR2 in disease;PIP3 activates AKT signaling;VEGFA-VEGFR2 Pathway;Synthesis of PIPs at the plasma membrane;Constitutive Signaling by Ligand-Responsive EGFR Cancer Variants;Interleukin receptor SHC signaling;G beta.CL.gamma signalling through PI3Kgamma;Regulation of signaling by CBL;Role of phospholipids in phagocytosis;Signaling by FGFR3 in disease;Constitutive Signaling by Aberrant PI3K in Cancer;Signaling by FGFR4 in disease;CD28 dependent PI3KAkt signaling;Tie2 Signaling;Signaling by FGFR1 fusion mutants;GP1b-IX-V activation signalling;GPVI-mediated activation cascade;Nephrin interactions;Constitutive Signaling by EGFRvIII;Gastrin-CREB signalling pathway via PKC and MAPK;Downstream signal transduction;GAB1 signalosome;Signaling by SCF-KIT;PI3KAKT activation;Role of LAT2NTALLAB on calcium mobilization;DAP12 signaling;Signaling by PDGF;Signaling by ERBB2;IRS-related events triggered by IGF1R;CD28 co-stimulation;Cell surface interactions at the vascular wall;Signaling by FGFR1 in disease 
                                               hsa04012;hsa04014;hsa04015;hsa04022;hsa04024;hsa04062;hsa04066;hsa04068;hsa04070;hsa04071;hsa04072;hsa04150;hsa04151;hsa04152;hsa04210;hsa04211;hsa04213;hsa04261;hsa04370;hsa04380;hsa04510;hsa04550;hsa04611;hsa04620;hsa04630;hsa04650;hsa04660;hsa04662;hsa04664;hsa04666;hsa04668;hsa04670;hsa04722;hsa04725;hsa04750;hsa04810;hsa04910;hsa04914;hsa04915;hsa04917;hsa04919;hsa04921;hsa04923;hsa04930;hsa04931;hsa04932;hsa04933;hsa04960;hsa04973;hsa05100;hsa05142;hsa05145;hsa05146;hsa05160;hsa05161;hsa05162;hsa05164;hsa05166;hsa05169;hsa05200;hsa05203;hsa05205;hsa05210;hsa05211;hsa05212;hsa05213;hsa05214;hsa05215;hsa05218;hsa05220;hsa05221;hsa05222;hsa05223;hsa05230;hsa05231 
                                               PIK3R1 
                                               NA 
                                               azure4 
                                               1 
                                               -0.178571428571429 
                                             
                                           
                                           DE Results - CD28 co-stimulation genes - New.Annotation: miR155 vs.cel67  Table displaying each sample's global significance scores and directed global significance scores as defined in the heatmaps above. The global significance score is calculated as the square root of the mean squared t-statistic for the genes in a gene set, with t-statistics coming from the linear regression underlying our differential expression analysis. The directed global significance score is calculated as the square root of the mean signed squared t-statistic for the genes in a gene set, with t-statistics coming from the linear regression underlying our differential expression analysis.   
                                      
                                       
                                  
                                
                                
                                  
                                       
                                      
                                           Volcano Plot: New.Annotation: miR18b vs.cel67   More Plot Information                Volcano Plot: New.Annotation: miR18b vs.cel67  Volcano plot displaying each gene's -log10(p-value) and log2 fold change for the selected covariate. Highly statistically significant genes fall at the top of the plot, and highly differentially expressed genes fall to either side. Genes within the selected gene set are highlighted in orange. Horizontal lines indicate various False Discovery Rate (FDR) thresholds.   
                                           DE Results - CD28 co-stimulation genes - New.Annotation: miR18b vs.cel67   More Plot Information      Download CSV Data     
                                           
                                             
                                             
                                           
                                           
                                             
                                               Probe Label 
                                               Log2 fold change 
                                               std error (log2) 
                                               Lower confidence limit (log2) 
                                               Upper confidence limit (log2) 
                                               Linear fold change 
                                               Lower confidence limit (linear) 
                                               Upper confidence limit (linear) 
                                               P-value 
                                               BY.p.value 
                                               method 
                                               Gene.sets 
                                               Codeset.Name 
                                               Probe.Label 
                                               Analyte.Type 
                                               Is.Control 
                                               Control.Type 
                                               Related.Probes 
                                               Probe.Annotation 
                                               KEGG.Pathways 
                                               Official.Gene.Name 
                                               Control.Conc 
                                               volcanocol 
                                               volcanopch 
                                               tstats.all 
                                             
                                           
                                           
                                             
                                               CDC42-mRNA 
                                               -1.5 
                                               0.0913 
                                               -1.68 
                                               -1.33 
                                               0.353 
                                               0.311 
                                               0.399 
                                               1.75e-13 
                                               2.32e-10 
                                               loglinear 
                                               CD28 co-stimulation, CD28 dependent Vav1 pathway, DCC mediated attractive signaling, EPHB-mediated forward signaling, G alpha (1213) signalling events, GPVI-mediated activation cascade, Regulation of actin dynamics for phagocytic cup formation, Rho GTPase cycle, RHO GTPases Activate Formins, RHO GTPases activate IQGAPs, RHO GTPases activate PAKs, Sema4D induced cell migration and growth-cone collapse, VEGFA-VEGFR2 Pathway 
                                               SEBELIUS_2_C4636 
                                               CDC42 
                                               mRNA 
                                               0 
                                               
                                               
                                               DCC mediated attractive signaling;Factors involved in megakaryocyte development and platelet production;VEGFA-VEGFR2 Pathway;RHO GTPases activate KTN1;CD28 dependent Vav1 pathway;Inactivation of Cdc42 and Rac;Rho GTPase cycle;EPHB-mediated forward signaling;RHO GTPases Activate Formins;RHO GTPases activate PAKs;GPVI-mediated activation cascade;G alpha (1213) signalling events;RHO GTPases activate IQGAPs;Sema4D induced cell migration and growth-cone collapse;Regulation of actin dynamics for phagocytic cup formation;CD28 co-stimulation 
                                               hsa04010;hsa04014;hsa04015;hsa04062;hsa04144;hsa04360;hsa04370;hsa04510;hsa04520;hsa04530;hsa04660;hsa04666;hsa04670;hsa04722;hsa04810;hsa04912;hsa04932;hsa04933;hsa05100;hsa05120;hsa05130;hsa05131;hsa05132;hsa05200;hsa05203;hsa05205;hsa05211;hsa05212 
                                               CDC42 
                                               NA 
                                               slateblue1 
                                               16 
                                               -16.4293537787514 
                                             
                                             
                                               PIK3R2-mRNA 
                                               -1.26 
                                               0.158 
                                               -1.57 
                                               -0.953 
                                               0.417 
                                               0.336 
                                               0.517 
                                               8.32e-08 
                                               1.58e-05 
                                               lm.nb 
                                               CD28 co-stimulation, CD28 dependent PI3KAkt signaling, Cell surface interactions at the vascular wall, Constitutive Signaling by Aberrant PI3K in Cancer, DAP12 signaling, Downstream signal transduction, Downstream TCR signaling, G alpha (1213) signalling events, G alpha (q) signalling events, G beta.CL.gamma signalling through PI3Kgamma, GAB1 signalosome, Gastrin-CREB signalling pathway via PKC and MAPK, GPVI-mediated activation cascade, Interleukin receptor SHC signaling, IRS-related events triggered by IGF1R, Nephrin interactions, PI3K Cascade, PI3KAKT activation, PIP3 activates AKT signaling, Regulation of signaling by CBL, Rho GTPase cycle, Role of LAT2NTALLAB on calcium mobilization, Signaling by ERBB2, Signaling by PDGF, Signaling by SCF-KIT, Synthesis of PIPs at the plasma membrane, Tie2 Signaling, VEGFA-VEGFR2 Pathway 
                                               SEBELIUS_2_C4636 
                                               PIK3R2 
                                               mRNA 
                                               0 
                                               
                                               
                                               G alpha (q) signalling events;Downstream TCR signaling;PIP3 activates AKT signaling;VEGFA-VEGFR2 Pathway;Synthesis of PIPs at the plasma membrane;Rho GTPase cycle;Interleukin receptor SHC signaling;G beta.CL.gamma signalling through PI3Kgamma;Regulation of signaling by CBL;Role of phospholipids in phagocytosis;G alpha (1213) signalling events;PI3K Cascade;Constitutive Signaling by Aberrant PI3K in Cancer;CD28 dependent PI3KAkt signaling;Tie2 Signaling;GPVI-mediated activation cascade;Nephrin interactions;Gastrin-CREB signalling pathway via PKC and MAPK;Downstream signal transduction;GAB1 signalosome;Signaling by SCF-KIT;PI3KAKT activation;Role of LAT2NTALLAB on calcium mobilization;DAP12 signaling;Signaling by PDGF;Signaling by ERBB2;IRS-related events triggered by IGF1R;CD28 co-stimulation;Cell surface interactions at the vascular wall 
                                               hsa04012;hsa04014;hsa04015;hsa04022;hsa04024;hsa04062;hsa04066;hsa04068;hsa04070;hsa04071;hsa04072;hsa04150;hsa04151;hsa04152;hsa04210;hsa04211;hsa04213;hsa04261;hsa04370;hsa04380;hsa04510;hsa04550;hsa04611;hsa04620;hsa04630;hsa04650;hsa04660;hsa04662;hsa04664;hsa04666;hsa04668;hsa04670;hsa04722;hsa04725;hsa04750;hsa04810;hsa04910;hsa04914;hsa04915;hsa04917;hsa04919;hsa04921;hsa04923;hsa04930;hsa04931;hsa04932;hsa04933;hsa04960;hsa04973;hsa05100;hsa05142;hsa05145;hsa05146;hsa05160;hsa05161;hsa05162;hsa05164;hsa05166;hsa05169;hsa05200;hsa05203;hsa05205;hsa05206;hsa05210;hsa05211;hsa05212;hsa05213;hsa05214;hsa05215;hsa05218;hsa05220;hsa05221;hsa05222;hsa05223;hsa05230;hsa05231 
                                               PIK3R2 
                                               NA 
                                               slateblue1 
                                               16 
                                               -7.9746835443038 
                                             
                                             
                                               PIK3R3-mRNA 
                                               1.43 
                                               0.292 
                                               0.854 
                                               2 
                                               2.69 
                                               1.81 
                                               4 
                                               7.9e-05 
                                               0.00291 
                                               lm.nb 
                                               CD28 co-stimulation, CD28 dependent PI3KAkt signaling, Constitutive Signaling by Aberrant PI3K in Cancer, DAP12 signaling, Downstream signal transduction, G alpha (1213) signalling events, G alpha (q) signalling events, G beta.CL.gamma signalling through PI3Kgamma, GAB1 signalosome, Gastrin-CREB signalling pathway via PKC and MAPK, GPVI-mediated activation cascade, Interleukin receptor SHC signaling, PI3KAKT activation, PIP3 activates AKT signaling, Regulation of signaling by CBL, Role of LAT2NTALLAB on calcium mobilization, Signaling by ERBB2, Signaling by PDGF, Signaling by SCF-KIT, Synthesis of PIPs at the plasma membrane 
                                               SEBELIUS_2_C4636 
                                               PIK3R3 
                                               mRNA 
                                               0 
                                               
                                               
                                               G alpha (q) signalling events;PIP3 activates AKT signaling;Constitutive Signaling by Aberrant PI3K in Cancer;CD28 dependent PI3KAkt signaling;Synthesis of PIPs at the plasma membrane;Interleukin receptor SHC signaling;G beta.CL.gamma signalling through PI3Kgamma;Regulation of signaling by CBL;GPVI-mediated activation cascade;G alpha (1213) signalling events;Gastrin-CREB signalling pathway via PKC and MAPK;Downstream signal transduction;GAB1 signalosome;Signaling by SCF-KIT;PI3KAKT activation;Role of LAT2NTALLAB on calcium mobilization;DAP12 signaling;Signaling by PDGF;Signaling by ERBB2;CD28 co-stimulation 
                                               hsa04012;hsa04014;hsa04015;hsa04022;hsa04024;hsa04062;hsa04066;hsa04068;hsa04070;hsa04071;hsa04072;hsa04150;hsa04151;hsa04152;hsa04210;hsa04211;hsa04213;hsa04261;hsa04370;hsa04380;hsa04510;hsa04550;hsa04611;hsa04620;hsa04630;hsa04650;hsa04660;hsa04662;hsa04664;hsa04666;hsa04668;hsa04670;hsa04722;hsa04725;hsa04750;hsa04810;hsa04910;hsa04914;hsa04915;hsa04917;hsa04919;hsa04921;hsa04923;hsa04930;hsa04931;hsa04932;hsa04933;hsa04960;hsa04973;hsa05100;hsa05142;hsa05145;hsa05146;hsa05160;hsa05161;hsa05162;hsa05164;hsa05166;hsa05169;hsa05200;hsa05203;hsa05205;hsa05210;hsa05211;hsa05212;hsa05213;hsa05214;hsa05215;hsa05218;hsa05220;hsa05221;hsa05222;hsa05223;hsa05230;hsa05231 
                                               PIK3R3 
                                               NA 
                                               slateblue1 
                                               16 
                                               4.8972602739726 
                                             
                                             
                                               AKT1-mRNA 
                                               -0.697 
                                               0.173 
                                               -1.04 
                                               -0.358 
                                               0.617 
                                               0.487 
                                               0.78 
                                               0.000613 
                                               0.0164 
                                               lm.nb 
                                               Activation of BAD and translocation to mitochondria , CD28 co-stimulation, DAP12 signaling, Deactivation of the beta-catenin transactivating complex, Downstream signal transduction, G beta.CL.gamma signalling through PI3Kgamma, GAB1 signalosome, GPVI-mediated activation cascade, IRS-related events triggered by IGF1R, PI3K Cascade, PI3KAKT activation, PIP3 activates AKT signaling, Role of LAT2NTALLAB on calcium mobilization, Signaling by ERBB2, Signaling by PDGF, Signaling by SCF-KIT, TP53 Regulates Metabolic Genes, VEGFA-VEGFR2 Pathway, VEGFR2 mediated vascular permeability 
                                               SEBELIUS_2_C4636 
                                               AKT1 
                                               mRNA 
                                               0 
                                               
                                               
                                               TP53 Regulates Metabolic Genes;Deactivation of the beta-catenin transactivating complex;Activation of BAD and translocation to mitochondria ;eNOS activation;mTOR signalling;G beta.CL.gamma signalling through PI3Kgamma;KSRP (KHSRP) binds and destabilizes mRNA;AKT phosphorylates targets in the nucleus;VEGFR2 mediated vascular permeability;GPVI-mediated activation cascade;Tetrahydrobiopterin (BH4) synthesis# recycling# salvage and regulation;Butyrate Response Factor 1 (BRF1) binds and destabilizes mRNA;Signaling by ERBB2;PIP3 activates AKT signaling;Downstream signal transduction;GAB1 signalosome;Signaling by SCF-KIT;PI3KAKT activation;Role of LAT2NTALLAB on calcium mobilization;DAP12 signaling;Signaling by PDGF;CD28 co-stimulation;PI3K Cascade;IRS-related events triggered by IGF1R;VEGFA-VEGFR2 Pathway 
                                               hsa04010;hsa04012;hsa04014;hsa04015;hsa04022;hsa04024;hsa04062;hsa04066;hsa04068;hsa04071;hsa04072;hsa04150;hsa04151;hsa04152;hsa04210;hsa04211;hsa04213;hsa04261;hsa04370;hsa04380;hsa04510;hsa04530;hsa04550;hsa04611;hsa04620;hsa04630;hsa04660;hsa04662;hsa04664;hsa04666;hsa04668;hsa04722;hsa04725;hsa04728;hsa04910;hsa04914;hsa04915;hsa04917;hsa04919;hsa04920;hsa04922;hsa04923;hsa04931;hsa04932;hsa04933;hsa04973;hsa05142;hsa05145;hsa05152;hsa05160;hsa05161;hsa05162;hsa05164;hsa05166;hsa05169;hsa05200;hsa05205;hsa05210;hsa05211;hsa05212;hsa05213;hsa05214;hsa05215;hsa05218;hsa05220;hsa05221;hsa05222;hsa05223;hsa05230;hsa05231 
                                               AKT1 
                                               NA 
                                               slateblue2 
                                               16 
                                               -4.02890173410405 
                                             
                                             
                                               SRC-mRNA 
                                               0.369 
                                               0.13 
                                               0.114 
                                               0.624 
                                               1.29 
                                               1.08 
                                               1.54 
                                               0.00983 
                                               0.184 
                                               loglinear 
                                               ARMS-mediated activation, CD28 co-stimulation, Cell surface interactions at the vascular wall, CTLA4 inhibitory signaling, DAP12 signaling, Downstream signal transduction, FCERI mediated MAPK activation, Frs2-mediated activation, GAB1 signalosome, Gastrin-CREB signalling pathway via PKC and MAPK, Integrin alphaIIb beta3 signaling, Interleukin receptor SHC signaling, IRS-related events triggered by IGF1R, MAP2K and MAPK activation, Negative regulation of FGFR1 signaling, Negative regulation of FGFR2 signaling, Negative regulation of FGFR3 signaling, Negative regulation of FGFR4 signaling, p130Cas linkage to MAPK signaling for integrins, RAFMAP kinase cascade, Signaling by ERBB2, Signaling by PDGF, Signaling by SCF-KIT, Spry regulation of FGF signaling, VEGFA-VEGFR2 Pathway, VEGFR2 mediated cell proliferation 
                                               SEBELIUS_2_C4636 
                                               SRC 
                                               mRNA 
                                               0 
                                               
                                               
                                               FCGR activation;Spry regulation of FGF signaling;GAB1 signalosome;MAP2K and MAPK activation;CLEC7A (Dectin-1) signaling;CD28 co-stimulation;p130Cas linkage to MAPK signaling for integrins;CTLA4 inhibitory signaling;GRB2.CL.SOS provides linkage to MAPK signaling for Integrins ;Signal regulatory protein (SIRP) family interactions;PECAM1 interactions;Thrombin signalling through proteinase activated receptors (PARs);GP1b-IX-V activation signalling;c-src mediated regulation of Cx43 function and closure of gap junctions;Signaling by ERBB2;VEGFA-VEGFR2 Pathway;Signaling by SCF-KIT;VEGFR2 mediated cell proliferation;RAFMAP kinase cascade;Downstream signal transduction;Frs2-mediated activation;ARMS-mediated activation;Gastrin-CREB signalling pathway via PKC and MAPK;Interleukin receptor SHC signaling;FCERI mediated MAPK activation;DAP12 signaling;Signaling by PDGF;IRS-related events triggered by IGF1R;Negative regulation of FGFR3 signaling;Negative regulation of FGFR4 signaling;Negative regulation of FGFR2 signaling;Negative regulation of FGFR1 signaling;Integrin alphaIIb beta3 signaling;Cell surface interactions at the vascular wall 
                                               hsa04012;hsa04015;hsa04062;hsa04144;hsa04370;hsa04510;hsa04520;hsa04530;hsa04540;hsa04611;hsa04727;hsa04750;hsa04810;hsa04912;hsa04915;hsa04917;hsa04919;hsa04921;hsa05100;hsa05120;hsa05131;hsa05152;hsa05161;hsa05203;hsa05205;hsa05219 
                                               SRC 
                                               NA 
                                               slateblue4 
                                               16 
                                               2.83846153846154 
                                             
                                             
                                               PAK1-mRNA 
                                               -0.488 
                                               0.186 
                                               -0.853 
                                               -0.123 
                                               0.713 
                                               0.554 
                                               0.918 
                                               0.016 
                                               0.272 
                                               lm.nb 
                                               Activation of Rac, CD209 (DC-SIGN) signaling, CD28 co-stimulation, CD28 dependent Vav1 pathway, EPHB-mediated forward signaling, Ephrin signaling, FCERI mediated MAPK activation, MAPK6MAPK4 signaling, Regulation of actin dynamics for phagocytic cup formation, RHO GTPases activate PAKs, Sema3A PAK dependent Axon repulsion, VEGFA-VEGFR2 Pathway, VEGFR2 mediated vascular permeability 
                                               SEBELIUS_2_C4636 
                                               PAK1 
                                               mRNA 
                                               0 
                                               
                                               
                                               EPHB-mediated forward signaling;Ephrin signaling;MAPK6MAPK4 signaling;DSCAM interactions;Activation of Rac;VEGFR2 mediated vascular permeability;RHO GTPases activate PAKs;Sema3A PAK dependent Axon repulsion;FCERI mediated MAPK activation;Regulation of actin dynamics for phagocytic cup formation;CD209 (DC-SIGN) signaling;CD28 dependent Vav1 pathway;VEGFA-VEGFR2 Pathway;CD28 co-stimulation 
                                               hsa04010;hsa04012;hsa04014;hsa04024;hsa04062;hsa04360;hsa04510;hsa04650;hsa04660;hsa04666;hsa04810;hsa05120;hsa05205;hsa05211 
                                               PAK1 
                                               NA 
                                               slateblue4 
                                               16 
                                               -2.62365591397849 
                                             
                                             
                                               PIK3CA-mRNA
