## Supplementary Results 1 for "Differential gene expression and functional pathway enrichment in colon cell line CCD 841 CoN (CRL-1790) transfected with miR-mimics miR-18b, miR-142-3p, miR-155, and miR-890": panel2_2.js

```
document.write('      	  <li>Volcano Plot - New.AnnotationControl<br><img src="results//DE//volcano plotNew.AnnotationControl.png"></img></li>');
document.write('      	  <li>Volcano Plot - New.AnnotationmiR142<br><img src="results//DE//volcano plotNew.AnnotationmiR142.png"></img></li>');
document.write('      	  <li>Volcano Plot - New.AnnotationmiR890<br><img src="results//DE//volcano plotNew.AnnotationmiR890.png"></img></li>');
document.write('      	  <li>Volcano Plot - New.AnnotationmiR18b<br><img src="results//DE//volcano plotNew.AnnotationmiR18b.png"></img></li>');
document.write('      	  <li>Volcano Plot - New.AnnotationmiR155<br><img src="results//DE//volcano plotNew.AnnotationmiR155.png"></img></li>');
```
