## Supplementary Results 1 for "Differential gene expression and functional pathway enrichment in colon cell line CCD 841 CoN (CRL-1790) transfected with miR-mimics miR-18b, miR-142-3p, miR-155, and miR-890": status.js

```
document.write('<p>Loading R libraries</p>');document.write('<p>Loaded  5 / 60  libraries</p>');document.write('<p>Loaded  10 / 60  libraries</p>');document.write('<p>Loaded  15 / 60  libraries</p>');document.write('<p>Loaded  20 / 60  libraries</p>');document.write('<p>Loaded  25 / 60  libraries</p>');document.write('<p>Loaded  30 / 60  libraries</p>');document.write('<p>Loaded  35 / 60  libraries</p>');document.write('<p>Loaded  40 / 60  libraries</p>');document.write('<p>Loaded  45 / 60  libraries</p>');document.write('<p>Loaded  50 / 60  libraries</p>');document.write('<p>Loaded  55 / 60  libraries</p>');document.write('<p>Loaded  60 / 60  libraries</p>');document.write('<p>Finished loading R libraries</p>');document.write('<p>Normalized mRNA data</p>');document.write('<p>Created mRNA normalized data files</p>');document.write('<p>Starting Overview</p>');document.write('<p>Creating Overview plots</p>');document.write('<p>Creating Overview heatmaps for gene set  1 / 115 </p>');document.write('<p>Creating Overview heatmaps for gene set  2 / 115 </p>');document.write('<p>Creating Overview heatmaps for gene set  3 / 115 </p>');document.write('<p>Creating Overview heatmaps for gene set  4 / 115 </p>');document.write('<p>Creating Overview heatmaps for gene set  5 / 115 </p>');document.write('<p>Creating Overview heatmaps for gene set  6 / 115 </p>');document.write('<p>Creating Overview heatmaps for gene set  7 / 115 </p>');document.write('<p>Creating Overview heatmaps for gene set  8 / 115 </p>');document.write('<p>Creating Overview heatmaps for gene set  9 / 115 </p>');document.write('<p>Creating Overview heatmaps for gene set  10 / 115 </p>');document.write('<p>Creating Overview heatmaps for gene set  11 / 115 </p>');document.write('<p>Creating Overview heatmaps for gene set  12 / 115 </p>');document.write('<p>Creating Overview heatmaps for gene set  13 / 115 </p>');document.write('<p>Creating Overview heatmaps for gene set  14 / 115 </p>');document.write('<p>Creating Overview heatmaps for gene set  15 / 115 </p>');document.write('<p>Creating Overview heatmaps for gene set  16 / 115 </p>');document.write('<p>Creating Overview heatmaps for gene set  17 / 115 </p>');document.write('<p>Creating Overview heatmaps for gene set  18 / 115 </p>');document.write('<p>Creating Overview heatmaps for gene set  19 / 115 </p>');document.write('<p>Creating Overview heatmaps for gene set  20 / 115 </p>');document.write('<p>Creating Overview heatmaps for gene set  21 / 115 </p>');document.write('<p>Creating Overview heatmaps for gene set  22 / 115 </p>');document.write('<p>Creating Overview heatmaps for gene set  23 / 115 </p>');document.write('<p>Creating Overview heatmaps for gene set  24 / 115 </p>');document.write('<p>Creating Overview heatmaps for gene set  25 / 115 </p>');document.write('<p>Creating Overview heatmaps for gene set  26 / 115 </p>');document.write('<p>Creating Overview heatmaps for gene set  27 / 115 </p>');document.write('<p>Creating Overview heatmaps for gene set  28 / 115 </p>');document.write('<p>Creating Overview heatmaps for gene set  29 / 115 </p>');document.write('<p>Creating Overview heatmaps for gene set  30 / 115 </p>');document.write('<p>Creating Overview heatmaps for gene set  31 / 115 </p>');document.write('<p>Creating Overview heatmaps for gene set  32 / 115 </p>');document.write('<p>Creating Overview heatmaps for gene set  33 / 115 </p>');document.write('<p>Creating Overview heatmaps for gene set  34 / 115 </p>');document.write('<p>Creating Overview heatmaps for gene set  35 / 115 </p>');document.write('<p>Creating Overview heatmaps for gene set  36 / 115 </p>');document.write('<p>Creating Overview heatmaps for gene set  37 / 115 </p>');document.write('<p>Creating Overview heatmaps for gene set  38 / 115 </p>');document.write('<p>Creating Overview heatmaps for gene set  39 / 115 </p>');document.write('<p>Creating Overview heatmaps for gene set  40 / 115 </p>');document.write('<p>Creating Overview heatmaps for gene set  41 / 115 </p>');document.write('<p>Creating Overview heatmaps for gene set  42 / 115 </p>');document.write('<p>Creating Overview heatmaps for gene set  43 / 115 </p>');document.write('<p>Creating Overview heatmaps for gene set  44 / 115 </p>');document.write('<p>Creating Overview heatmaps for gene set  45 / 115 </p>');document.write('<p>Creating Overview heatmaps for gene set  46 / 115 </p>');document.write('<p>Creating Overview heatmaps for gene set  47 / 115 </p>');document.write('<p>Creating Overview heatmaps for gene set  48 / 115 </p>');document.write('<p>Creating Overview heatmaps for gene set  49 / 115 </p>');document.write('<p>Creating Overview heatmaps for gene set  50 / 115 </p>');document.write('<p>Creating Overview heatmaps for gene set  51 / 115 </p>');document.write('<p>Creating Overview heatmaps for gene set  52 / 115 </p>');document.write('<p>Creating Overview heatmaps for gene set  53 / 115 </p>');document.write('<p>Creating Overview heatmaps for gene set  54 / 115 </p>');document.write('<p>Creating Overview heatmaps for gene set  55 / 115 </p>');document.write('<p>Creating Overview heatmaps for gene set  56 / 115 </p>');document.write('<p>Creating Overview heatmaps for gene set  57 / 115 </p>');document.write('<p>Creating Overview heatmaps for gene set  58 / 115 </p>');document.write('<p>Creating Overview heatmaps for gene set  59 / 115 </p>');document.write('<p>Creating Overview heatmaps for gene set  60 / 115 </p>');document.write('<p>Creating Overview heatmaps for gene set  61 / 115 </p>');document.write('<p>Creating Overview heatmaps for gene set  62 / 115 </p>');document.write('<p>Creating Overview heatmaps for gene set  63 / 115 </p>');document.write('<p>Creating Overview heatmaps for gene set  64 / 115 </p>');document.write('<p>Creating Overview heatmaps for gene set  65 / 115 </p>');document.write('<p>Creating Overview heatmaps for gene set  66 / 115 </p>');document.write('<p>Creating Overview heatmaps for gene set  67 / 115 </p>');document.write('<p>Creating Overview heatmaps for gene set  68 / 115 </p>');document.write('<p>Creating Overview heatmaps for gene set  69 / 115 </p>');document.write('<p>Creating Overview heatmaps for gene set  70 / 115 </p>');document.write('<p>Creating Overview heatmaps for gene set  71 / 115 </p>');document.write('<p>Creating Overview heatmaps for gene set  72 / 115 </p>');document.write('<p>Creating Overview heatmaps for gene set  73 / 115 </p>');document.write('<p>Creating Overview heatmaps for gene set  74 / 115 </p>');document.write('<p>Creating Overview heatmaps for gene set  75 / 115 </p>');document.write('<p>Creating Overview heatmaps for gene set  76 / 115 </p>');document.write('<p>Creating Overview heatmaps for gene set  77 / 115 </p>');document.write('<p>Creating Overview heatmaps for gene set  78 / 115 </p>');document.write('<p>Creating Overview heatmaps for gene set  79 / 115 </p>');document.write('<p>Creating Overview heatmaps for gene set  80 / 115 </p>');document.write('<p>Creating Overview heatmaps for gene set  81 / 115 </p>');document.write('<p>Creating Overview heatmaps for gene set  82 / 115 </p>');document.write('<p>Creating Overview heatmaps for gene set  83 / 115 </p>');document.write('<p>Creating Overview heatmaps for gene set  84 / 115 </p>');document.write('<p>Creating Overview heatmaps for gene set  85 / 115 </p>');document.write('<p>Creating Overview heatmaps for gene set  86 / 115 </p>');document.write('<p>Creating Overview heatmaps for gene set  87 / 115 </p>');document.write('<p>Creating Overview heatmaps for gene set  88 / 115 </p>');document.write('<p>Creating Overview heatmaps for gene set  89 / 115 </p>');document.write('<p>Creating Overview heatmaps for gene set  90 / 115 </p>');document.write('<p>Creating Overview heatmaps for gene set  91 / 115 </p>');document.write('<p>Creating Overview heatmaps for gene set  92 / 115 </p>');document.write('<p>Creating Overview heatmaps for gene set  93 / 115 </p>');document.write('<p>Creating Overview heatmaps for gene set  94 / 115 </p>');document.write('<p>Creating Overview heatmaps for gene set  95 / 115 </p>');document.write('<p>Creating Overview heatmaps for gene set  96 / 115 </p>');document.write('<p>Creating Overview heatmaps for gene set  97 / 115 </p>');document.write('<p>Creating Overview heatmaps for gene set  98 / 115 </p>');document.write('<p>Creating Overview heatmaps for gene set  99 / 115 </p>');document.write('<p>Creating Overview heatmaps for gene set  100 / 115 </p>');document.write('<p>Creating Overview heatmaps for gene set  101 / 115 </p>');document.write('<p>Creating Overview heatmaps for gene set  102 / 115 </p>');document.write('<p>Creating Overview heatmaps for gene set  103 / 115 </p>');document.write('<p>Creating Overview heatmaps for gene set  104 / 115 </p>');document.write('<p>Creating Overview heatmaps for gene set  105 / 115 </p>');document.write('<p>Creating Overview heatmaps for gene set  106 / 115 </p>');document.write('<p>Creating Overview heatmaps for gene set  107 / 115 </p>');document.write('<p>Creating Overview heatmaps for gene set  108 / 115 </p>');document.write('<p>Creating Overview heatmaps for gene set  109 / 115 </p>');document.write('<p>Creating Overview heatmaps for gene set  110 / 115 </p>');document.write('<p>Creating Overview heatmaps for gene set  111 / 115 </p>');document.write('<p>Creating Overview heatmaps for gene set  112 / 115 </p>');document.write('<p>Creating Overview heatmaps for gene set  113 / 115 </p>');document.write('<p>Creating Overview heatmaps for gene set  114 / 115 </p>');document.write('<p>Creating Overview heatmaps for gene set  115 / 115 </p>');document.write('<p>Creating Overview histograms  1 / 1 </p>');document.write('<p>Creating Overview HTML infrastructure</p>');document.write('<p>Finished Overview</p>');document.write('<p>Starting differential expression (DE) analysis</p>');document.write('<p>Creating differential expression (DE) analysis results files</p>');document.write('<p>Finished differential expression (DE) analysis</p>');document.write('<p>Starting gene set analysis (GSA)</p>');document.write('<p>Creating gene set analysis (GSA) plots for gene set 1/115</p>');document.write('<p>Creating gene set analysis (GSA) plots for gene set 2/115</p>');document.write('<p>Creating gene set analysis (GSA) plots for gene set 3/115</p>');document.write('<p>Creating gene set analysis (GSA) plots for gene set 4/115</p>');document.write('<p>Creating gene set analysis (GSA) plots for gene set 5/115</p>');document.write('<p>Creating gene set analysis (GSA) plots for gene set 6/115</p>');document.write('<p>Creating gene set analysis (GSA) plots for gene set 7/115</p>');document.write('<p>Creating gene set analysis (GSA) plots for gene set 8/115</p>');document.write('<p>Creating gene set analysis (GSA) plots for gene set 9/115</p>');document.write('<p>Creating gene set analysis (GSA) plots for gene set 10/115</p>');document.write('<p>Creating gene set analysis (GSA) plots for gene set 11/115</p>');document.write('<p>Creating gene set analysis (GSA) plots for gene set 12/115</p>');document.write('<p>Creating gene set analysis (GSA) plots for gene set 13/115</p>');document.write('<p>Creating gene set analysis (GSA) plots for gene set 14/115</p>');document.write('<p>Creating gene set analysis (GSA) plots for gene set 15/115</p>');document.write('<p>Creating gene set analysis (GSA) plots for gene set 16/115</p>');document.write('<p>Creating gene set analysis (GSA) plots for gene set 17/115</p>');document.write('<p>Creating gene set analysis (GSA) plots for gene set 18/115</p>');document.write('<p>Creating gene set analysis (GSA) plots for gene set 19/115</p>');document.write('<p>Creating gene set analysis (GSA) plots for gene set 20/115</p>');document.write('<p>Creating gene set analysis (GSA) plots for gene set 21/115</p>');document.write('<p>Creating gene set analysis (GSA) plots for gene set 22/115</p>');document.write('<p>Creating gene set analysis (GSA) plots for gene set 23/115</p>');document.write('<p>Creating gene set analysis (GSA) plots for gene set 24/115</p>');document.write('<p>Creating gene set analysis (GSA) plots for gene set 25/115</p>');document.write('<p>Creating gene set analysis (GSA) plots for gene set 26/115</p>');document.write('<p>Creating gene set analysis (GSA) plots for gene set 27/115</p>');document.write('<p>Creating gene set analysis (GSA) plots for gene set 28/115</p>');document.write('<p>Creating gene set analysis (GSA) plots for gene set 29/115</p>');document.write('<p>Creating gene set analysis (GSA) plots for gene set 30/115</p>');document.write('<p>Creating gene set analysis (GSA) plots for gene set 31/115</p>');document.write('<p>Creating gene set analysis (GSA) plots for gene set 32/115</p>');document.write('<p>Creating gene set analysis (GSA) plots for gene set 33/115</p>');document.write('<p>Creating gene set analysis (GSA) plots for gene set 34/115</p>');document.write('<p>Creating gene set analysis (GSA) plots for gene set 35/115</p>');document.write('<p>Creating gene set analysis (GSA) plots for gene set 36/115</p>');document.write('<p>Creating gene set analysis (GSA) plots for gene set 37/115</p>');document.write('<p>Creating gene set analysis (GSA) plots for gene set 38/115</p>');document.write('<p>Creating gene set analysis (GSA) plots for gene set 39/115</p>');document.write('<p>Creating gene set analysis (GSA) plots for gene set 40/115</p>');document.write('<p>Creating gene set analysis (GSA) plots for gene set 41/115</p>');document.write('<p>Creating gene set analysis (GSA) plots for gene set 42/115</p>');document.write('<p>Creating gene set analysis (GSA) plots for gene set 43/115</p>');document.write('<p>Creating gene set analysis (GSA) plots for gene set 44/115</p>');document.write('<p>Creating gene set analysis (GSA) plots for gene set 45/115</p>');document.write('<p>Creating gene set analysis (GSA) plots for gene set 46/115</p>');document.write('<p>Creating gene set analysis (GSA) plots for gene set 47/115</p>');document.write('<p>Creating gene set analysis (GSA) plots for gene set 48/115</p>');document.write('<p>Creating gene set analysis (GSA) plots for gene set 49/115</p>');document.write('<p>Creating gene set analysis (GSA) plots for gene set 50/115</p>');document.write('<p>Creating gene set analysis (GSA) plots for gene set 51/115</p>');document.write('<p>Creating gene set analysis (GSA) plots for gene set 52/115</p>');document.write('<p>Creating gene set analysis (GSA) plots for gene set 53/115</p>');document.write('<p>Creating gene set analysis (GSA) plots for gene set 54/115</p>');document.write('<p>Creating gene set analysis (GSA) plots for gene set 55/115</p>');document.write('<p>Creating gene set analysis (GSA) plots for gene set 56/115</p>');document.write('<p>Creating gene set analysis (GSA) plots for gene set 57/115</p>');document.write('<p>Creating gene set analysis (GSA) plots for gene set 58/115</p>');document.write('<p>Creating gene set analysis (GSA) plots for gene set 59/115</p>');document.write('<p>Creating gene set analysis (GSA) plots for gene set 60/115</p>');document.write('<p>Creating gene set analysis (GSA) plots for gene set 61/115</p>');document.write('<p>Creating gene set analysis (GSA) plots for gene set 62/115</p>');document.write('<p>Creating gene set analysis (GSA) plots for gene set 63/115</p>');document.write('<p>Creating gene set analysis (GSA) plots for gene set 64/115</p>');document.write('<p>Creating gene set analysis (GSA) plots for gene set 65/115</p>');document.write('<p>Creating gene set analysis (GSA) plots for gene set 66/115</p>');document.write('<p>Creating gene set analysis (GSA) plots for gene set 67/115</p>');document.write('<p>Creating gene set analysis (GSA) plots for gene set 68/115</p>');document.write('<p>Creating gene set analysis (GSA) plots for gene set 69/115</p>');document.write('<p>Creating gene set analysis (GSA) plots for gene set 70/115</p>');document.write('<p>Creating gene set analysis (GSA) plots for gene set 71/115</p>');document.write('<p>Creating gene set analysis (GSA) plots for gene set 72/115</p>');document.write('<p>Creating gene set analysis (GSA) plots for gene set 73/115</p>');document.write('<p>Creating gene set analysis (GSA) plots for gene set 74/115</p>');document.write('<p>Creating gene set analysis (GSA) plots for gene set 75/115</p>');document.write('<p>Creating gene set analysis (GSA) plots for gene set 76/115</p>');document.write('<p>Creating gene set analysis (GSA) plots for gene set 77/115</p>');document.write('<p>Creating gene set analysis (GSA) plots for gene set 78/115</p>');document.write('<p>Creating gene set analysis (GSA) plots for gene set 79/115</p>');document.write('<p>Creating gene set analysis (GSA) plots for gene set 80/115</p>');document.write('<p>Creating gene set analysis (GSA) plots for gene set 81/115</p>');document.write('<p>Creating gene set analysis (GSA) plots for gene set 82/115</p>');document.write('<p>Creating gene set analysis (GSA) plots for gene set 83/115</p>');document.write('<p>Creating gene set analysis (GSA) plots for gene set 84/115</p>');document.write('<p>Creating gene set analysis (GSA) plots for gene set 85/115</p>');document.write('<p>Creating gene set analysis (GSA) plots for gene set 86/115</p>');document.write('<p>Creating gene set analysis (GSA) plots for gene set 87/115</p>');document.write('<p>Creating gene set analysis (GSA) plots for gene set 88/115</p>');document.write('<p>Creating gene set analysis (GSA) plots for gene set 89/115</p>');document.write('<p>Creating gene set analysis (GSA) plots for gene set 90/115</p>');document.write('<p>Creating gene set analysis (GSA) plots for gene set 91/115</p>');document.write('<p>Creating gene set analysis (GSA) plots for gene set 92/115</p>');document.write('<p>Creating gene set analysis (GSA) plots for gene set 93/115</p>');document.write('<p>Creating gene set analysis (GSA) plots for gene set 94/115</p>');document.write('<p>Creating gene set analysis (GSA) plots for gene set 95/115</p>');document.write('<p>Creating gene set analysis (GSA) plots for gene set 96/115</p>');document.write('<p>Creating gene set analysis (GSA) plots for gene set 97/115</p>');document.write('<p>Creating gene set analysis (GSA) plots for gene set 98/115</p>');document.write('<p>Creating gene set analysis (GSA) plots for gene set 99/115</p>');document.write('<p>Creating gene set analysis (GSA) plots for gene set 100/115</p>');document.write('<p>Creating gene set analysis (GSA) plots for gene set 101/115</p>');document.write('<p>Creating gene set analysis (GSA) plots for gene set 102/115</p>');document.write('<p>Creating gene set analysis (GSA) plots for gene set 103/115</p>');document.write('<p>Creating gene set analysis (GSA) plots for gene set 104/115</p>');document.write('<p>Creating gene set analysis (GSA) plots for gene set 105/115</p>');document.write('<p>Creating gene set analysis (GSA) plots for gene set 106/115</p>');document.write('<p>Creating gene set analysis (GSA) plots for gene set 107/115</p>');document.write('<p>Creating gene set analysis (GSA) plots for gene set 108/115</p>');document.write('<p>Creating gene set analysis (GSA) plots for gene set 109/115</p>');document.write('<p>Creating gene set analysis (GSA) plots for gene set 110/115</p>');document.write('<p>Creating gene set analysis (GSA) plots for gene set 111/115</p>');document.write('<p>Creating gene set analysis (GSA) plots for gene set 112/115</p>');document.write('<p>Creating gene set analysis (GSA) plots for gene set 113/115</p>');document.write('<p>Creating gene set analysis (GSA) plots for gene set 114/115</p>');document.write('<p>Creating gene set analysis (GSA) plots for gene set 115/115</p>');document.write('<p>Creating gene set analysis (GSA) results files</p>');document.write('<p>Creating gene set analysis (GSA) HTML infrastructure</p>');document.write('<p>Finished gene set analysis (GSA)</p>');document.write('<p>Starting pathview analysis</p>');document.write('<p>Downloading KEGG files for hsa04810  pathway  1 / 20  </p>');document.write('<p>Downloading KEGG files for hsa04510  pathway  2 / 20  </p>');document.write('<p>Downloading KEGG files for hsa04530  pathway  3 / 20  </p>');document.write('<p>Downloading KEGG files for hsa05200  pathway  4 / 20  </p>');document.write('<p>Downloading KEGG files for hsa05205  pathway  5 / 20  </p>');document.write('<p>Downloading KEGG files for hsa04151  pathway  6 / 20  </p>');document.write('<p>Downloading KEGG files for hsa04015  pathway  7 / 20  </p>');document.write('<p>Downloading KEGG files for hsa04670  pathway  8 / 20  </p>');document.write('<p>Downloading KEGG files for hsa04010  pathway  9 / 20  </p>');document.write('<p>Downloading KEGG files for hsa04062  pathway  10 / 20  </p>');document.write('<p>Downloading KEGG files for hsa04014  pathway  11 / 20  </p>');document.write('<p>Downloading KEGG files for hsa05166  pathway  12 / 20  </p>');document.write('<p>Downloading KEGG files for hsa04520  pathway  13 / 20  </p>');document.write('<p>Downloading KEGG files for hsa05210  pathway  14 / 20  </p>');document.write('<p>Downloading KEGG files for hsa04024  pathway  15 / 20  </p>');document.write('<p>Downloading KEGG files for hsa04071  pathway  16 / 20  </p>');document.write('<p>Downloading KEGG files for hsa05215  pathway  17 / 20  </p>');document.write('<p>Downloading KEGG files for hsa05206  pathway  18 / 20  </p>');document.write('<p>Downloading KEGG files for hsa05160  pathway  19 / 20  </p>');document.write('<p>Downloading KEGG files for hsa04310  pathway  20 / 20  </p>');document.write('<p>Finished pathview analysis</p>');document.write('<p>Starting pathway scoring analysis</p>');document.write('<p>Activation of BAD and translocation to mitochondria , CREB phosphorylation through the activation of Ras, Cyclin AB1 associated events during G2M transition, DARPP-32 events, DCC mediated attractive signaling, ERKs are inactivated, G alpha (q) signalling events, MASTL Facilitates Mitotic Progression, Negative feedback regulation of MAPK pathway, Nonsense Mediated Decay (NMD) enhanced by the Exon Junction Complex (EJC), p130Cas linkage to MAPK signaling for integrins, Repression of WNT target genes, Resolution of Sister Chromatid Cohesion, TGF-beta receptor signaling in EMT (epithelial to mesenchymal transition), TP53 Regulates Metabolic Genes, WNT mediated activation of DVL pathways dropped from Pathway Scoring Analysis due to low number of probes.</p>');document.write('<p>Finished pathway scoring analysis</p>');document.write('<p>Writing analysis report</p>');document.write('<p>Writing Overview results </p>');document.write('<p>Writing normalization results </p>');document.write('<p>Writing DE analysis results </p>');document.write('<p>Writing gene set analysis results</p>');document.write('<p>Writing Pathview results </p>');document.write('<p>Writing pathway scoring results</p>');
```
