## Supplementary Results 1 for "Differential gene expression and functional pathway enrichment in colon cell line CCD 841 CoN (CRL-1790) transfected with miR-mimics miR-18b, miR-142-3p, miR-155, and miR-890": advanced.analysis.js

```
	heatmapData_UQh1YWZvRyUX_kmCGIztRmQ = "Dimname,20161013_sebelius2-101316_01_01.RCC,20180703_1790-jr1_SA44miR890_02.RCC,20180712_1790-jr2_SA38cel67_02.RCC,20161013_sebelius2-101316_01_01.RCC,20180703_1790-jr1_SA54amiR155_06.RCC,20180703_1790-jr1_SA60miR142_09.RCC,20180712_1790-jr2_SA49amiR142_06.RCC,20180703_1790-jr1_SA50bmiR18b_04.RCC,20180703_1790-jr1_SA62miR18b_11.RCC,20180712_1790-jr2_SA62miR18b_11.RCC,20180703_1790-jr1_SA51miR890_05.RCC,20180703_1790-jr1_SA63miR890_12.RCC,20180712_1790-jr2_SA44miR890_04.RCC,20161013_sebelius2-101316_01_01.RCC,20180703_1790-jr1_SA54bmiR155_07.RCC,20180703_1790-jr1_SA61miR155_10.RCC,20180703_1790-jr1_SA40miR142_01.RCC,20180712_1790-jr2_SA40miR142_03.RCC,20180712_1790-jr2_SA51bmiR890_07.RCC,20180712_1790-jr2_SA63miR890_12.RCC,20180703_1790-jr1_SA48bcel67_03.RCC,20180712_1790-jr2_SA48bcel67_05.RCC,20180703_1790-jr1_SA57Control_08.RCC,20180712_1790-jr2_SA59cel67_09.RCC,20180712_1790-jr2_SA61miR155_10.RCC,20180712_1790-jr2_SA36Control_01.RCC,20180712_1790-jr2_SA57Control_08.RCC\nGAPDH - mRNA,53852,42735,34121,33083,33628,31906,37068,29397,32161,48083,54245,62604,63022,54230,54171,55930,45202,49475,91158,89249,69219,74419,73351,85097,83933,98371,109748\nCGN - mRNA,6,1,1,3,2,2,1,6,2,3,1,1,4,1,3,5,3,5,11,4,2,4,2,3,3,5,7\nNEG_C(0) - mRNA,5,1,2,2,1,2,1,4,1,6,1,3,5,3,4,3,1,4,8,3,5,2,1,3,1,2,7\nNEG_D(0) - mRNA,1,1,3,3,1,1,3,1,3,1,1,1,4,2,1,2,2,6,8,3,4,2,1,4,1,4,2\nPIK3R5 - mRNA,6,1,2,1,1,1,1,1,1,1,2,2,1,2,1,3,1,4,9,1,1,1,2,2,3,8,5\nNEG_G(0) - mRNA,3,2,2,1,1,3,2,2,1,4,2,1,5,1,2,3,2,1,5,5,1,1,2,2,3,6,3\nNEG_H(0) - mRNA,1,1,1,1,1,1,2,1,4,1,1,1,1,2,2,2,1,1,1,4,2,3,1,4,6,2,2\nNEG_B(0) - mRNA,1,5,3,2,1,1,3,2,1,1,1,1,3,7,1,1,2,2,4,1,3,5,3,4,1,6,3\nEPB41L3 - mRNA,5,2,2,4,2,1,5,1,2,2,2,6,4,5,2,4,4,3,6,4,2,6,6,2,6,8,5\nACTN3 - mRNA,11,3,2,4,1,2,3,2,1,3,2,2,2,2,5,1,5,1,3,3,3,4,2,3,8,6,2\nALPI - mRNA,8,3,3,3,4,1,4,6,2,5,4,4,10,6,9,4,2,12,7,11,3,9,2,10,7,8,10\nWAS - mRNA,6,1,5,4,5,6,8,7,1,5,5,1,10,9,8,7,5,8,14,4,8,6,6,7,7,7,12\nNEG_A(0) - mRNA,4,4,4,3,5,3,7,3,3,4,2,1,6,11,5,6,3,8,8,1,5,6,6,10,6,5,8\nMYH14 - mRNA,1,2,4,6,4,3,1,2,3,2,2,3,4,9,8,4,6,4,6,5,4,4,8,4,4,5,12\nPRKCB - mRNA,10,4,3,8,2,2,3,3,6,5,3,3,3,4,8,5,2,7,5,4,1,6,12,5,4,8,10\nVAV3 b - mRNA,7,3,5,2,4,3,6,3,6,5,4,5,2,6,3,5,3,4,4,7,3,5,4,10,6,13,6\nCDX2 - mRNA,6,3,5,1,1,1,2,1,3,5,5,3,7,2,2,6,2,5,4,9,5,5,2,7,7,9,7\nCLDN3 - mRNA,4,1,5,1,4,1,6,3,3,9,2,8,1,6,6,2,3,2,6,3,6,3,7,5,3,7,7\nCXCR4 - mRNA,7,2,5,4,5,2,1,4,1,3,3,3,2,6,3,5,3,7,7,7,6,4,2,9,3,5,7\nVAV3 - mRNA,2,1,2,8,3,2,4,3,3,5,1,4,4,7,4,3,4,11,4,4,6,1,4,8,7,8,7\nTJP3 - mRNA,2,2,4,2,1,1,2,4,1,5,4,2,2,6,3,2,3,10,3,3,5,9,3,5,7,6,6\nCA2 - mRNA,3,13,3,5,5,1,3,1,5,3,14,4,25,4,7,5,3,5,13,6,4,6,4,5,3,8,12\nLYZ - mRNA,13,8,8,10,12,3,8,11,5,7,6,7,12,11,8,6,8,9,11,9,10,9,16,10,7,12,14\nLLGL2 - mRNA,13,7,5,1,7,2,10,5,7,5,7,3,11,7,11,9,12,8,15,10,5,7,14,6,14,10,17\nPIK3CG - mRNA,13,4,3,5,1,1,3,5,3,4,6,5,8,7,2,12,4,11,6,4,7,4,8,12,14,7,15\nIL1R2 - mRNA,8,5,2,6,4,3,6,4,1,10,5,7,9,7,6,10,5,12,9,12,15,9,9,10,12,9,22\nCRB3 - mRNA,7,8,4,6,5,1,5,1,6,7,11,10,7,15,9,11,5,7,8,9,4,4,8,8,16,12,16\nHPSE - mRNA,7,5,3,3,3,1,6,4,2,4,1,6,7,6,6,9,8,6,12,10,8,8,4,8,10,14,16\nITGB4 - mRNA,10,8,5,4,3,4,3,4,5,7,3,16,8,10,9,9,5,8,14,7,7,9,5,6,13,18,19\nCLDN8 - mRNA,7,7,9,9,7,2,7,7,7,11,5,5,5,4,6,8,4,10,12,10,17,8,9,13,7,19,14\nNEG_F(0) - mRNA,10,4,9,8,5,3,1,7,4,8,7,5,8,8,4,3,5,7,11,10,11,6,4,3,13,14,8\nCA1 - mRNA,4,1,3,4,2,1,5,4,6,6,12,4,5,12,7,3,1,3,7,5,6,6,12,5,4,15,8\nCLDN4 - mRNA,7,4,3,4,6,1,2,4,4,6,6,8,5,4,8,4,3,6,11,11,2,7,11,18,6,16,16\nMYH11 - mRNA,8,1,4,5,4,3,5,1,5,12,6,8,3,3,4,7,9,8,6,9,7,5,4,12,8,11,9\nNEG_E(0) - mRNA,7,1,6,3,4,5,5,5,8,9,3,5,4,5,4,2,4,8,9,7,2,7,6,9,7,7,16\nJAM2 - mRNA,50,4,6,11,6,6,10,9,2,11,9,10,17,13,21,4,15,17,8,7,11,9,4,8,11,15,11\nITGA4 - mRNA,6,7,12,13,5,6,7,4,3,9,4,11,16,15,11,14,8,12,24,21,17,26,10,16,23,28,22\nCLDN7 - mRNA,6,4,4,9,6,9,17,5,10,6,8,3,13,10,4,9,11,15,17,10,9,14,10,17,12,24,12\nTCF7 - mRNA,14,12,6,9,9,1,14,9,13,14,16,8,18,16,13,10,13,13,16,8,16,17,10,19,12,24,22\nBCL2 - mRNA,13,9,28,13,10,7,16,7,6,21,10,4,17,28,14,5,21,30,24,14,27,28,17,40,20,37,24\nPRKCZ - mRNA,30,15,25,29,10,18,23,14,15,19,19,14,35,33,19,21,30,45,32,30,22,23,16,34,23,38,44\nPOS_F(0.125) - mRNA,21,16,29,20,6,17,25,14,17,21,20,19,23,34,15,16,23,28,25,17,17,16,13,38,23,17,40\nEPB41 - mRNA,26,16,14,19,3,5,20,12,11,20,16,12,34,24,13,11,24,33,38,33,16,20,8,11,18,36,27\nITGA7 - mRNA,17,8,5,9,4,9,4,6,12,12,17,21,21,20,16,14,17,23,32,41,5,16,21,16,22,41,35\nMYL12A - mRNA,22,11,14,12,6,5,13,12,10,15,11,20,22,34,11,20,11,20,26,34,8,21,23,22,34,31,35\nAPC2 - mRNA,18,11,6,21,18,9,18,17,13,17,16,22,24,10,34,25,10,27,32,25,22,32,12,16,47,29,22\nCLDN15 - mRNA,36,5,9,16,13,13,21,13,12,18,6,9,11,15,22,18,21,36,17,14,25,24,14,13,24,32,22\nHCLS1 - mRNA,27,15,12,17,14,11,11,11,14,22,29,11,10,19,19,18,20,29,23,26,13,24,18,24,35,31,28\nAXIN2 - mRNA,94,9,12,28,11,16,27,16,16,23,20,18,26,22,32,17,31,35,45,23,42,38,27,33,29,40,46\nF11R - mRNA,55,25,8,22,15,21,32,14,14,19,45,40,45,15,17,6,27,32,75,78,19,16,24,26,8,36,33\nPDGFA - mRNA,13,22,36,22,20,12,27,24,14,46,28,32,32,30,31,29,29,49,38,38,44,42,41,44,49,83,64\nPOS_E(0.5) - mRNA,42,35,47,27,32,35,35,31,37,55,30,33,53,38,38,30,52,68,54,44,33,54,42,63,60,57,55\nHSPA1A - mRNA,53,28,30,31,24,21,44,44,28,64,39,23,62,43,48,27,42,83,76,45,50,64,32,55,64,101,71\nAXIN1 - mRNA,48,27,49,32,36,30,54,30,41,59,42,27,65,43,43,45,41,71,88,62,45,70,36,58,79,97,79\nMYL12A c - mRNA,72,47,61,76,39,30,67,40,61,74,50,57,84,119,64,73,61,111,98,99,52,65,74,62,100,180,122\nMAGI1 - mRNA,32,56,41,43,37,62,52,37,39,59,46,64,74,63,59,49,96,111,61,58,66,59,95,88,81,174,134\nTJP2 - mRNA,49,48,30,37,53,45,50,34,55,103,81,62,68,63,68,92,57,55,136,106,87,98,100,129,129,110,159\nRAC3 - mRNA,41,73,38,25,35,27,32,36,21,45,55,53,81,41,46,46,35,50,105,102,60,63,52,70,88,113,103\nRAC2 - mRNA,56,40,35,32,31,26,43,22,17,43,77,34,61,46,46,61,52,77,151,75,57,87,49,84,94,108,98\nBAIAP2 - mRNA,58,35,36,26,26,28,36,42,46,79,56,51,66,39,51,72,55,74,82,58,66,60,49,56,101,125,95\nARAF - mRNA,55,52,32,43,64,41,68,60,42,89,52,65,67,58,87,76,68,79,106,56,89,96,74,87,116,125,107\nLEF1 - mRNA,142,65,56,83,66,54,106,111,71,83,95,54,85,74,108,81,112,146,182,67,153,159,41,89,105,162,70\nCLDN23 - mRNA,219,58,48,96,51,40,78,57,50,61,65,37,96,79,64,46,115,130,126,71,132,153,58,68,70,132,77\nMYL5 - mRNA,173,40,43,94,44,72,110,65,49,101,55,59,68,53,106,77,123,112,111,91,93,99,81,73,95,127,121\nOCLN - mRNA,151,36,28,71,36,60,99,36,67,60,48,42,76,44,91,44,88,117,110,76,81,95,50,55,92,107,82\nNLK - mRNA,118,79,49,96,74,80,138,94,96,169,114,109,135,86,117,86,110,139,165,163,94,108,88,82,145,150,134\nCYFIP2 - mRNA,50,77,33,82,56,73,83,57,46,87,115,103,127,71,96,68,114,138,218,98,85,75,62,48,125,116,98\nMYL12A b - mRNA,107,71,53,74,45,41,82,67,59,104,95,88,107,83,71,58,90,96,172,147,77,95,86,82,98,182,130\nINADL - mRNA,110,53,35,65,49,68,102,48,40,77,66,60,107,70,70,67,104,112,113,103,71,99,98,69,104,146,138\nMAGI3 - mRNA,188,71,59,72,79,54,89,80,74,103,85,76,103,102,156,114,117,157,170,115,190,203,116,150,164,218,161\nTCF7L1 - mRNA,183,96,95,123,108,74,157,132,115,197,112,74,160,108,173,123,181,251,199,117,170,180,107,133,236,266,186\nPIK3R1 - mRNA,164,67,76,147,72,83,146,106,101,149,105,87,125,143,142,108,176,202,142,135,134,147,111,108,191,236,152\nBHLHE40 - mRNA,70,90,91,62,82,51,33,35,45,68,95,125,160,144,163,222,99,91,157,167,112,113,130,173,320,274,216\nBAIAP2L1 - mRNA,102,109,103,121,72,124,121,54,78,122,132,139,163,205,110,118,143,181,224,245,133,142,184,182,230,342,327\nDVL1 - mRNA,103,109,107,92,107,69,78,105,69,151,146,137,176,133,164,170,119,152,252,199,177,217,164,207,242,299,259\nBAD - mRNA,146,116,129,68,96,60,85,113,84,211,130,140,210,101,132,104,108,158,298,203,128,192,100,212,215,267,241\nDIAPH3 - mRNA,132,77,100,73,103,54,114,103,85,174,102,99,103,174,159,101,105,117,200,196,282,367,191,403,222,306,367\nMYC - mRNA,128,68,131,73,95,67,72,68,86,131,107,118,108,141,169,166,90,90,167,169,216,245,201,398,282,280,296\nMET - mRNA,162,89,80,52,56,63,48,61,65,79,124,166,130,158,103,145,67,89,215,235,200,232,229,323,220,308,299\nPIK3CB - mRNA,132,100,83,112,117,119,161,172,180,259,155,128,140,150,185,180,145,217,251,160,252,248,217,251,277,288,313\nXPO5 - mRNA,165,141,104,111,110,86,115,134,156,209,165,168,186,181,231,215,159,173,288,212,238,255,190,285,309,279,285\nHRAS - mRNA,109,116,119,86,118,62,99,140,142,200,159,148,175,164,155,164,121,123,297,206,243,265,156,307,233,274,274\nPAK4 - mRNA,175,121,115,87,126,62,130,88,83,135,204,179,197,153,184,175,152,174,320,302,271,256,197,288,301,358,307\nERBB2 - mRNA,179,109,59,86,118,98,172,97,95,168,133,158,218,103,169,184,176,264,259,246,205,241,164,155,311,313,268\nWASF1 - mRNA,238,133,82,134,90,92,151,150,139,218,197,172,221,152,176,114,192,231,316,234,175,194,173,174,174,246,229\nGUSB - mRNA,302,115,114,180,104,105,213,164,118,219,127,114,195,169,201,121,229,339,240,177,238,285,148,168,248,384,239\nDNMT3A - mRNA,322,164,114,188,150,105,239,154,138,234,167,142,268,189,250,172,234,322,317,242,228,230,131,123,262,310,186\nPOS_D(2) - mRNA,210,214,344,214,182,151,249,188,192,296,191,241,297,290,295,183,258,379,342,261,227,253,259,338,300,334,348\nPRKCD - mRNA,248,131,131,154,136,92,207,152,147,236,212,183,281,186,227,196,213,280,383,294,213,330,208,300,395,405,366\nDVL2 - mRNA,347,128,127,161,165,97,235,182,158,261,173,146,232,186,286,215,228,276,303,217,264,349,186,223,337,383,305\nCDK4 - mRNA,288,137,146,172,150,107,173,190,173,243,167,131,213,234,247,194,216,218,328,219,309,367,229,319,318,388,338\nTARBP2 - mRNA,219,152,133,138,128,103,120,222,199,334,206,184,205,166,227,174,178,211,393,296,252,303,215,299,296,321,316\nEPB41L1 - mRNA,174,211,119,160,136,122,182,150,155,242,216,225,343,168,226,215,216,283,419,364,238,270,177,200,304,364,307\nEXOC3 - mRNA,300,176,95,177,168,132,192,165,158,216,196,215,259,189,306,225,224,253,389,305,247,247,186,212,347,323,316\nVAV2 - mRNA,282,230,127,183,243,154,249,281,237,369,277,255,349,272,421,331,308,354,473,330,392,393,311,398,512,509,447\nAGO2 - mRNA,268,231,199,168,192,132,233,233,270,423,298,322,473,233,339,311,273,342,521,428,330,445,250,598,547,504,394\nPIK3CD - mRNA,172,237,194,169,199,180,225,171,151,231,407,284,356,238,316,244,264,291,629,420,346,324,273,430,381,475,453\nBCAR1 - mRNA,118,234,216,146,182,143,247,189,180,334,260,208,377,216,326,261,274,403,475,332,309,378,289,419,392,710,467\nPIK3R2 - mRNA,243,230,214,175,206,126,238,123,118,197,259,335,412,217,303,259,252,377,545,505,386,464,275,442,459,665,509\nAPC - mRNA,234,131,152,132,234,112,153,237,192,348,173,157,211,225,410,398,188,267,300,232,350,351,285,371,531,480,410\nLIMK2 - mRNA,219,147,92,207,140,168,246,156,195,346,194,197,281,198,244,262,244,337,382,362,190,232,245,223,490,402,375\nABI2 - mRNA,314,157,142,229,107,148,260,223,217,375,239,185,314,271,195,162,288,388,448,314,298,322,259,247,280,513,415\nCSNK2A2 - mRNA,326,193,121,210,199,180,279,216,211,324,236,231,291,231,338,284,322,374,402,294,324,288,233,226,403,422,308\nSRC - mRNA,416,179,150,216,174,131,229,236,254,415,207,193,346,280,290,265,268,347,452,293,293,338,253,241,433,528,415\nSYMPK - mRNA,371,221,139,230,196,181,254,223,239,348,284,255,285,262,352,308,332,352,428,343,346,345,280,304,493,526,448\nFYN - mRNA,378,169,138,237,187,173,299,220,188,348,268,169,305,247,320,268,298,374,490,328,304,352,261,321,460,455,413\nTCF7L2 - mRNA,553,145,137,292,109,216,380,246,251,403,225,150,207,282,242,169,409,496,394,258,398,419,262,319,278,561,453\nIGF1R - mRNA,555,250,166,315,135,141,277,235,249,488,245,301,478,373,267,221,364,475,464,508,246,352,209,236,377,671,395\nMLLT4 - mRNA,594,274,137,265,223,163,260,300,228,329,373,323,431,286,386,264,347,411,644,407,341,391,315,330,377,512,401\nJUN - mRNA,877,242,184,171,187,148,218,191,222,374,277,317,409,231,331,300,241,359,476,475,326,453,280,537,515,541,487\nASH1L - mRNA,684,195,217,239,285,150,293,380,331,494,283,218,302,402,492,417,319,407,499,301,511,588,375,452,698,774,592\nPPP1CC - mRNA,483,251,215,261,252,186,291,260,241,410,280,307,409,369,420,364,296,422,613,514,466,581,372,536,613,712,548\nPPP2R2D - mRNA,460,259,259,313,277,189,350,270,231,360,285,282,391,405,470,322,393,452,523,395,452,558,337,523,592,692,578\nMAPK7 - mRNA,482,271,170,270,235,176,280,230,183,309,334,337,377,295,454,361,395,425,550,511,407,437,275,379,537,647,442\nBRAF - mRNA,423,257,201,300,244,247,372,241,226,320,326,323,430,302,386,372,451,512,517,448,429,407,320,349,575,626,487\nNUMA1 - mRNA,477,248,179,220,299,196,365,312,256,386,303,251,361,251,499,394,422,507,562,317,446,470,346,375,573,733,461\nDGCR8 - mRNA,414,221,204,293,245,210,311,270,224,372,292,309,378,357,456,338,363,474,480,439,412,441,329,414,538,610,514\nTBP - mRNA,470,224,162,270,241,195,270,282,233,370,296,234,312,359,478,365,366,418,462,348,404,454,343,367,493,601,552\nPDCD4 - mRNA,1018,96,223,385,212,281,508,341,420,626,128,119,141,324,395,350,591,573,226,177,436,532,353,288,565,599,508\nMAPK10 - mRNA,1013,216,236,426,213,201,370,235,207,326,255,169,308,418,533,325,664,675,419,247,473,491,216,215,445,713,360\nMRAS - mRNA,272,187,225,329,213,216,274,231,331,537,158,241,349,408,384,408,458,522,306,304,287,358,429,313,654,803,612\nCCND2 - mRNA,265,69,214,416,280,425,489,142,177,256,121,113,135,303,591,535,566,664,237,154,485,540,477,581,864,707,752\nHSPA1B - mRNA,542,340,165,161,206,82,144,313,210,373,460,395,629,358,405,263,191,241,987,662,354,439,300,322,430,683,537\nRRAS - mRNA,257,310,290,229,223,164,222,141,98,196,467,405,597,333,328,325,272,452,792,638,327,446,369,411,623,736,579\nTGFB2 - mRNA,356,433,300,201,220,148,177,311,339,484,440,445,760,620,380,371,278,313,782,637,371,400,402,383,611,1050,648\nWASL - mRNA,499,353,277,140,380,78,102,406,331,537,458,409,572,465,605,555,175,197,779,613,588,615,415,466,809,824,600\nSOS2 - mRNA,500,278,260,372,361,331,463,364,439,721,401,416,468,378,624,638,521,627,614,617,573,605,358,532,933,749,662\nCCND3 - mRNA,208,376,212,320,393,292,517,403,510,833,538,542,593,319,490,606,442,565,944,793,439,543,541,558,1051,930,787\nIRS1 - mRNA,672,306,344,515,264,334,415,397,386,644,322,354,525,533,503,356,709,769,581,509,600,739,440,579,547,915,733\nKRAS - mRNA,531,451,307,549,177,336,498,490,515,836,457,588,646,629,357,283,626,727,856,802,603,638,528,610,382,972,863\nPIK3R3 - mRNA,836,136,112,333,362,355,620,401,746,1062,239,227,206,228,501,511,456,500,337,303,416,471,310,284,754,355,423\nIL1R1 b - mRNA,1084,393,141,281,336,219,443,713,466,689,803,355,690,195,598,322,456,540,1298,459,472,513,150,264,524,503,283\nPDGFD - mRNA,1966,409,538,1476,482,539,1457,804,477,783,729,272,740,850,1130,448,1730,2137,1375,415,1205,1393,298,418,753,1395,483\nSPRY2 - mRNA,450,643,354,317,461,303,378,371,293,457,908,1020,947,564,861,891,435,505,1440,1426,887,924,718,1696,1324,1012,1122\nITGA3 - mRNA,436,443,600,336,608,281,373,392,446,751,600,697,811,694,868,1093,438,623,1060,1026,925,1127,927,1617,1912,1603,1530\nCASK - mRNA,1362,427,346,313,588,252,334,521,499,834,603,556,617,698,905,920,448,466,1021,830,1073,1114,807,909,1322,1230,1237\nYES1 - mRNA,1256,594,480,287,514,164,282,669,699,1011,762,675,834,954,1040,853,381,433,1222,946,1064,1126,855,912,1249,1569,1299\nCSNK2A1 - mRNA,777,512,376,443,534,415,580,573,427,732,741,663,866,608,933,779,682,879,1281,839,793,895,699,783,1311,1380,1236\nDIAPH1 - mRNA,515,505,320,474,477,439,578,508,503,763,702,810,798,672,654,755,694,858,1168,1050,715,819,776,1021,1129,1223,1150\nABCF1 - mRNA,732,512,409,444,473,315,474,566,482,867,688,645,824,637,765,769,604,722,1247,982,934,1015,730,1129,1150,1236,1121\nPTK2 - mRNA,876,491,372,554,420,364,553,525,527,817,592,669,761,750,711,655,639,776,984,992,849,941,695,890,1185,1191,1085\nAKT2 - mRNA,737,477,381,433,431,326,560,599,561,867,616,641,751,503,753,640,600,810,1081,914,833,943,628,834,1039,1176,1049\nPXN - mRNA,333,376,293,379,598,388,541,387,377,612,610,639,699,515,853,931,563,693,1066,892,716,834,832,1061,1526,1239,1318\nAKT1 - mRNA,557,536,447,376,563,416,607,466,417,681,676,723,879,520,834,839,642,893,1237,997,921,1152,887,1264,1424,1572,1463\nMPP5 - mRNA,519,557,395,411,304,398,479,568,737,1064,698,934,778,650,522,594,567,583,1128,1373,888,902,1023,1177,847,1078,1497\nCTTN - mRNA,533,520,335,236,418,196,256,506,563,904,678,729,721,554,667,699,355,355,1076,1052,731,837,684,854,982,1134,1028\nPAK1 - mRNA,484,400,282,330,387,328,424,349,397,633,441,707,654,582,649,757,480,549,825,987,729,814,669,1102,1121,1030,1066\nHPRT1 - mRNA,581,389,325,379,345,303,403,421,372,632,623,573,598,565,572,542,468,560,1077,889,809,900,776,1042,851,992,1154\nCCNG2 - mRNA,1280,512,368,647,393,448,799,459,476,671,554,448,761,512,793,503,923,1133,971,713,709,785,407,489,814,986,621\nDVL3 - mRNA,818,346,202,521,457,411,602,474,427,712,552,490,627,548,874,768,743,935,918,626,547,585,577,393,1127,1114,888\nMAPK8 - mRNA,662,401,326,489,433,419,529,366,276,445,513,521,633,628,807,699,654,760,944,782,681,767,517,762,1033,1049,762\nRAF1 - mRNA,653,382,369,444,438,336,442,297,271,482,468,515,613,619,671,588,572,673,851,754,676,794,565,745,1069,1057,948\nDROSHA - mRNA,719,293,271,423,371,325,480,443,407,624,388,309,445,576,672,543,559,668,700,470,796,798,617,707,833,1016,863\nGSK3B - mRNA,784,397,279,546,398,419,614,457,467,622,521,458,611,532,687,549,731,743,797,657,583,643,497,520,822,860,807\nSOS1 - mRNA,825,390,321,468,322,299,497,445,440,645,525,528,621,534,618,472,690,770,863,637,769,791,542,590,595,1037,772\nEP300 - mRNA,931,373,295,468,433,355,607,409,418,637,522,406,639,537,690,535,701,880,910,561,760,857,494,705,886,1097,837\nYBX3 - mRNA,513,256,386,415,539,431,500,585,623,880,391,313,403,576,805,747,629,849,627,398,922,946,744,1025,1186,1216,1076\nSMAD4 - mRNA,849,443,377,412,534,277,519,602,505,713,586,475,682,582,944,742,668,856,866,614,787,895,607,667,1003,1125,902\nCRKL - mRNA,729,416,368,556,497,416,584,622,509,814,511,386,649,624,783,725,710,794,954,627,883,980,676,900,1091,1261,1119\nPIK3CA - mRNA,812,465,373,679,380,449,945,519,473,773,548,504,776,668,711,533,775,1089,1056,849,764,980,518,822,893,1194,976\nTP53 - mRNA,842,450,399,688,478,395,700,706,566,844,631,412,814,762,961,603,825,1121,1217,616,807,969,567,520,990,1507,951\nMYH10 - mRNA,1247,586,604,740,479,325,638,660,612,993,446,431,936,1214,943,709,1115,1392,821,631,1006,1030,717,678,1111,2313,1158\nPOLR2A - mRNA,755,403,357,546,603,515,800,641,651,1042,558,513,670,538,925,776,888,1135,1004,662,967,1088,769,922,1286,1554,1187\nCREBBP - mRNA,1085,543,453,644,576,541,836,745,683,1006,704,618,816,708,1114,816,1088,1235,1109,801,1104,1141,786,816,1180,1488,1118\nPRKCA - mRNA,1041,558,496,742,581,645,785,618,655,943,697,609,819,922,1053,915,1026,1186,1043,983,1238,1410,882,1147,1422,1503,1326\nLIMK1 - mRNA,857,591,555,682,500,550,821,398,472,675,665,763,991,811,903,917,805,1098,1125,1155,946,1068,769,1145,1394,1340,1265\nBRK1 - mRNA,1100,485,510,557,370,355,692,497,545,1138,674,656,1066,770,669,653,562,1030,1619,1331,664,982,682,1062,1408,1455,1455\nSMAD2 - mRNA,1133,605,449,781,449,551,941,369,320,449,768,745,960,914,845,671,1001,1235,1412,1053,865,982,832,796,1026,1456,1230\nZEB1 - mRNA,877,714,738,803,599,422,729,768,625,1124,889,823,1552,1189,1169,889,865,1227,1788,1325,1118,1385,786,1102,1541,2208,1475\nITGAV - mRNA,915,974,771,544,472,315,514,783,606,906,1066,801,1420,1049,1036,767,791,1017,1746,1106,1133,1290,841,970,1113,2286,1231\nCRK - mRNA,726,578,575,372,543,237,402,801,739,1211,800,821,1039,780,1005,873,532,701,1341,1135,1018,1144,896,1147,1438,1494,1336\nPPP2R2A - mRNA,853,685,480,505,672,413,505,779,696,942,947,1082,1075,898,1158,1016,777,863,1545,1294,1074,1158,1085,1241,1458,1541,1449\nGNAI3 - mRNA,878,669,465,628,644,414,618,660,692,1089,925,1033,1116,873,1168,1083,776,811,1630,1592,994,1129,913,1200,1725,1496,1456\nPPP2R1B - mRNA,1169,583,454,713,591,540,767,908,902,1449,723,693,945,750,978,808,875,1097,1317,1050,1006,1160,781,1099,1342,1419,1232\nMAPK9 - mRNA,932,619,434,763,648,573,857,903,839,1359,842,769,948,913,1103,1034,995,1144,1277,1066,1040,1075,956,961,1477,1535,1421\nEXOC4 - mRNA,1161,638,415,617,612,456,710,720,664,901,856,850,959,784,1091,870,968,1053,1469,1107,1001,1052,811,817,1223,1383,1288\nMAPK3 - mRNA,999,705,381,629,630,462,671,815,790,1299,986,909,1025,748,1030,1001,871,934,1573,1354,925,1081,883,821,1439,1542,1419\nPARD3 - mRNA,1328,656,390,852,618,642,1159,868,798,1060,1023,797,916,809,1099,810,1247,1303,1466,1040,1227,1209,814,895,1082,1470,1087\nRAB8B - mRNA,923,728,538,765,672,648,982,643,614,962,917,955,1126,702,1112,1020,1136,1289,1565,1307,1219,1252,892,1084,1518,1383,1207\nCTBP1 - mRNA,1035,757,552,511,696,534,847,729,626,925,927,878,1213,673,1234,886,959,1135,1552,1227,1257,1287,892,1187,1411,1696,1293\nPRKCI - mRNA,1055,676,469,811,255,605,839,727,830,1313,827,1034,1037,909,592,433,986,1213,1582,1493,986,1050,929,820,699,1592,1507\nEGFR - mRNA,840,592,520,560,371,446,645,833,616,975,906,770,1101,764,802,543,812,1035,1520,1030,1209,1417,862,1493,801,1514,1269\nPOS_C(8) - mRNA,894,984,1536,770,752,735,1049,804,803,1131,920,900,1277,941,1175,838,953,1607,1378,982,988,1175,975,1486,1099,1304,1279\nALDH18A1 - mRNA,865,714,607,763,709,580,797,634,646,971,837,844,1095,1076,1342,1214,1022,1291,1512,1151,1131,1255,1088,1071,1726,2079,1740\nVAPA - mRNA,1147,868,643,674,767,523,652,942,846,1283,1169,1036,1209,1041,1360,1161,960,969,1842,1448,1430,1406,1114,1342,1573,1839,1781\nCTBP2 - mRNA,1453,851,637,848,748,587,938,916,804,1276,1124,906,1208,1056,1396,1161,1284,1242,1791,1259,1479,1427,1041,1226,1564,1939,1489\nMAP3K7 - mRNA,1309,696,590,963,616,697,1034,895,947,1360,1021,904,1058,1171,1101,981,1210,1308,1650,1261,1322,1362,1135,1124,1429,1933,1723\nNRAS - mRNA,651,535,740,633,1011,597,895,588,568,908,744,622,815,806,1527,1539,942,1218,1247,962,1370,1540,899,1595,2454,1822,1527\nDICER1 - mRNA,1236,486,560,754,623,589,744,338,347,566,714,726,865,865,1186,1171,951,1126,1342,1139,1162,1407,870,1486,1910,1478,1376\nRRAS2 - mRNA,800,515,322,889,669,669,911,710,856,1385,613,565,817,1093,1181,1264,994,1352,1025,796,608,743,1163,686,1918,1733,1940\nMAP2K1 - mRNA,832,410,603,592,644,521,575,923,851,1262,598,508,612,991,1046,1077,863,995,947,686,1154,1212,1185,1669,1681,1804,1735\nROCK1 - mRNA,1262,578,602,879,837,650,965,928,941,1328,741,608,876,1162,1393,1200,1178,1438,1237,844,1302,1355,1134,1293,1865,1885,1776\nPPP1R12A - mRNA,1087,495,598,1017,772,696,998,970,985,1496,651,681,854,1126,1323,1193,1255,1474,1088,905,1097,1269,945,1019,1776,1860,1508\nEPB41L2 - mRNA,2403,1018,1093,2182,1511,1290,2480,2049,2080,3321,1433,1057,1646,2368,2741,2359,2756,3536,2715,1633,2226,2655,2048,1876,3925,4039,3407\nHSPB1 - mRNA,865,906,1395,1195,1079,1208,1708,1294,1676,2853,932,989,1686,1680,1762,2050,2012,3380,1957,1444,1519,1871,2165,1912,3789,4912,3599\nWASF2 - mRNA,2207,799,710,1567,1760,1191,1714,1314,1109,1438,1029,751,1145,2023,2909,2383,2232,2344,1605,1007,2091,2109,1837,1715,3295,3149,2660\nTUBB - mRNA,2055,939,1086,1000,1471,917,1176,1354,1181,1797,1229,1089,1398,1723,2622,2324,1596,2027,2092,1458,3057,3241,2499,3445,3318,3578,3613\nTGFB1 - mRNA,1335,1411,1167,851,1277,756,1057,1201,1298,1888,1335,1618,2123,1573,2021,2144,1612,2034,2160,2026,1865,1964,2150,2118,3078,4359,2927\nSMAD3 - mRNA,1414,1217,1120,1060,1097,1003,1235,989,1007,1551,1520,1536,1792,1584,1842,1766,1552,1851,2593,2366,2484,2760,2296,2398,2653,3277,3273\nMAPK1 - mRNA,1841,1049,858,1331,1274,1147,1617,1022,893,1426,1450,1263,1678,1759,2151,2037,1899,2135,2515,1940,2095,2318,1830,2220,3138,3156,2890\nRAC1 - mRNA,2638,2042,1460,835,1206,532,875,2182,2068,3350,2578,2606,3301,2332,2454,2022,1156,1507,4667,3753,3172,3512,2538,3360,3265,4644,4059\nIQGAP1 - mRNA,1975,1927,1206,1852,1694,1354,1812,1425,1279,1883,2411,2500,2847,2648,2953,2680,2403,2620,3982,3647,2992,3164,2652,3385,4353,4339,4019\nPPP2CB - mRNA,2246,1475,1412,1728,785,1469,2193,2406,2537,3794,1900,1838,2229,2242,1712,1193,2567,3012,3119,2391,2989,3235,2441,3220,1734,4102,3655\nRHOA - mRNA,1608,1404,1363,1280,1240,1096,1441,1620,1854,3407,1554,2003,2632,1804,1875,1951,1583,2621,3243,3131,1956,2544,2031,2800,3695,3910,3716\nPPP2CA - mRNA,2297,1544,1217,1541,1335,1130,1693,1651,1655,2513,2067,2112,2555,2162,2393,2106,1957,2411,3726,3076,2616,2936,2138,3019,3312,3692,3399\nCSNK2B - mRNA,2307,1132,1247,1437,1319,1061,1748,1593,1606,2882,1636,1408,1930,1971,2157,2063,1684,2317,2974,2262,2298,2925,2135,2777,3432,3786,3599\nGSTM3 - mRNA,2066,1812,1124,1414,1140,1846,2128,1991,2505,3391,2344,2386,2516,1700,1991,2060,2329,2286,3669,3087,2315,2294,2228,1950,2877,2828,3230\nTJP1 - mRNA,3026,1256,927,1668,1267,1223,1727,1460,1298,2015,2034,1613,2014,1965,2262,1976,2080,2282,3270,2299,2291,2511,1841,2028,2959,3317,2962\nRAB11A - mRNA,2211,1585,835,1289,1196,1087,1423,1544,1467,2187,2101,2167,2210,1617,2042,1891,1745,1797,3396,3119,2082,2120,1975,2157,2798,2712,2880\nCD9 - mRNA,911,1351,926,868,1034,824,845,733,989,1854,1762,2206,2334,1521,1637,1821,1000,1356,3467,3455,1372,1753,1881,2329,3178,2777,3223\nG6PD - mRNA,968,1158,821,1149,1341,969,1463,1311,1959,3515,1239,1415,1980,1288,1852,1918,1452,1876,2438,2338,1346,1640,1424,1393,3393,2786,2349\nITGA6 - mRNA,809,822,668,462,954,564,677,554,500,703,1629,1930,1231,1155,1259,1192,810,943,2665,2463,2357,2576,1740,3786,1762,2403,2694\nPAK2 - mRNA,1178,779,656,998,616,933,1148,1225,1123,1618,958,766,1127,1218,1254,896,1488,1728,1578,1049,1645,1679,1298,1609,1344,2437,1966\nDOCK1 - mRNA,2061,754,693,1350,750,805,1331,1134,1081,1746,1024,866,1220,1370,1365,1074,1619,1795,1790,1408,1592,1783,1233,1412,1750,2431,1924\nAKT3 - mRNA,1423,1188,838,1059,650,811,1091,1200,1122,1534,1289,1045,1714,1334,1323,996,1593,1835,1988,1448,1488,1485,1159,1283,1390,2422,1572\nPTEN - mRNA,1581,1180,627,1287,1050,963,1342,1253,1309,1771,1347,1350,1596,1416,1680,1461,1802,1920,2196,1720,1528,1486,1186,1372,2018,2338,1825\nPSEN1 - mRNA,1292,779,645,836,853,729,1039,1093,1175,1906,1167,1247,1296,1076,1586,1339,1156,1416,2172,1817,1407,1698,1254,1581,2316,2088,2049\nJAM3 - mRNA,1039,1052,711,786,880,501,813,879,740,1152,1215,1157,1596,1149,1589,1394,1064,1239,2099,1509,1241,1355,1005,1099,2137,2342,1570\nPDGFC - mRNA,1316,1244,986,1354,955,803,1468,556,426,639,1480,1393,1882,1571,1692,1473,1797,2010,2494,1868,1462,1660,1174,1424,2014,2569,1829\nCLTC - mRNA,1386,904,719,1045,1033,726,1134,934,985,1662,1343,1389,1611,1650,1597,1609,1207,1492,2438,2305,1530,1858,1679,1942,2706,2641,2744\nFGFR1 - mRNA,1247,972,865,994,668,809,1230,854,810,1418,1204,1355,1760,1292,1313,1143,1416,1973,2417,1963,1509,1915,1417,1914,2003,2846,2437\nROCK2 - mRNA,1722,1307,807,1101,1329,908,1296,1348,1482,1921,1673,1600,1938,1592,2218,1803,1671,1866,2516,1968,2199,2192,1630,1801,2552,2793,2353\nPRKAA1 - mRNA,1853,1055,758,1378,1164,1123,1563,1373,1385,2005,1495,1369,1628,1581,2026,1757,1878,2091,2499,1836,1881,1979,1542,1407,2520,2500,2465\nPPP1CB - mRNA,2169,1127,820,1360,999,857,1594,855,804,1394,1384,1546,2008,1680,1685,1645,1572,2070,2702,2424,1571,2021,1514,1588,2659,2914,2674\nSPTAN1 - mRNA,1302,1320,756,1186,1130,954,1345,1130,1237,1936,1506,1726,2214,1501,1768,1717,1741,2205,2575,2576,1411,1685,1522,1555,2841,3192,2454\nGNAI1 - mRNA,3278,1358,650,1702,1326,1294,2454,3510,2202,3331,3065,1607,2178,1434,2527,1404,2320,2611,5481,2202,2660,3017,1405,1728,2197,2405,2254\nIL1R1 - mRNA,3994,1279,417,952,953,655,1268,1915,1370,1873,2298,1211,1904,795,1751,1054,1382,1457,3622,1555,1408,1441,505,754,1451,1218,788\nPDGFRA - mRNA,19098,2865,2137,4877,2671,2324,5783,4345,2836,5013,4639,2493,5319,3883,5899,2972,5636,8260,9272,4031,5991,7996,2262,2701,5239,7044,3995\nCTNNB1 - mRNA,7357,3528,2372,3565,3818,2892,4164,4660,4454,6519,5159,4608,5488,4530,6233,5790,5396,5709,8159,6301,6070,6345,5216,5327,8093,8465,7742\nCTNNA1 - mRNA,4874,3729,2604,3741,3920,2878,4140,4411,4321,6727,4795,4872,6179,4919,6578,5927,5277,6335,8180,7053,5612,6460,5062,5962,9733,9332,8161\nRAB13 - mRNA,7994,3504,2858,4014,2720,2994,4273,4659,3692,5308,5195,3661,4885,4474,5782,3958,5618,6024,8339,4727,6008,6324,4090,4261,5609,8490,5945\nCSNK1A1 - mRNA,6116,2682,1714,3977,1340,2715,4126,3474,3767,5853,3298,3213,4387,4828,2862,1963,4829,5998,5891,4717,3597,4189,4344,3350,3168,7655,6737\nITGA5 - mRNA,1911,2671,2602,1754,2949,1512,1839,1212,1222,2158,4033,3944,4444,2629,4405,4997,2416,3157,7286,5926,4153,4962,2961,6470,8587,5461,4987\nITGB5 - mRNA,1941,4258,2772,2890,2859,2254,2639,1878,1940,3339,3912,5174,7428,5120,4270,4941,4003,5087,7139,7786,3662,4297,4236,5715,7882,10094,7025\nCDC42 - mRNA,4730,4087,2519,3479,3007,3019,3979,1505,1182,1779,5094,5731,5767,4286,5186,4984,5079,5250,8283,7584,5296,5492,5148,5469,6976,7445,7322\nACTN4 - mRNA,1567,2387,2018,2517,2473,1979,3141,2347,2756,5036,2577,3052,4140,3621,3461,3676,3557,4823,4922,4688,2989,3884,4171,3830,6462,8804,6985\nVCL - mRNA,4026,2039,2264,2745,2508,1686,2640,3318,3448,6205,2905,2607,3761,4409,4574,4505,3508,4765,5748,4245,4017,5007,4197,4659,7802,8451,7542\nACTN1 - mRNA,2184,2750,2244,2317,2945,2213,2666,2798,3070,5092,3356,3705,4103,3434,4663,5322,3574,4369,5615,5128,4215,4523,4978,5337,8372,8726,7529\nPPP1CA - mRNA,2981,2005,2106,2146,2859,1760,2413,3445,3394,5014,2846,2578,2933,3339,4409,4111,2960,3333,4574,3761,4284,4643,3910,4830,6267,6133,5734\nTRIO - mRNA,3236,1988,1801,2400,1705,1889,2575,2469,2249,3378,2502,2369,3064,2895,3290,2429,3485,3750,4282,3460,3889,4038,3006,3332,3724,5577,4555\nNCKAP1 - mRNA,2764,2333,1259,1648,2030,1112,1512,2625,2550,3577,2875,2640,3379,3232,3531,3234,2328,2685,4363,3389,2887,2949,3346,2551,4602,5830,4733\nADAM9 - mRNA,1863,2442,1691,2130,1480,1862,2175,1891,2103,3355,2864,3120,3715,3168,2625,3087,2622,3109,5128,4736,3232,3721,3508,3549,4761,5363,5641\nPPP2R1A - mRNA,2727,1889,1448,1616,2144,1360,1859,1850,1947,3053,2593,2887,3207,2620,3201,3241,2102,2516,4667,4333,3097,3443,3117,3759,5330,4828,4920\nPDGFRB - mRNA,2680,2236,1830,2864,1789,1654,2579,1750,1610,2463,2152,2125,3767,3892,3470,2959,3929,4504,3730,3083,2771,3125,2689,2202,4606,7189,4076\nPOS_B(32) - mRNA,3270,3272,4702,3001,2762,2537,3481,2783,2967,3787,3116,3192,4131,3274,4024,2717,3346,5219,4671,3328,3662,3672,3265,4911,3690,4083,4326\nPOS_A(128) - mRNA,10357,10573,18305,9528,9118,8184,12963,9249,9384,14093,10154,10483,15119,10507,13334,8819,10777,19476,17029,11494,12037,13353,11050,18228,13855,15877,15407\nCFL1 - mRNA,8326,6577,6142,7305,7552,5733,7791,8648,9626,15935,9137,9056,10458,11636,11638,12231,8593,10251,16588,13648,11091,13073,12801,13520,19986,19765,21122\nFLNA - mRNA,6587,7281,6073,7662,7249,5773,8682,8612,9205,14541,7915,8030,11504,10461,12325,11484,12194,15066,13757,10670,10931,12186,10798,10117,18138,25020,17195\nITGA1 - mRNA,3101,7588,6840,12149,4380,6104,7810,7171,7758,11514,5258,5574,11373,14382,9012,8952,15611,16874,8546,7987,6942,7337,7294,5763,12599,23285,10630\nMYL12B - mRNA,5328,7237,5100,5826,5504,5200,6511,5709,6494,9757,8196,9520,10764,8240,8523,9656,8097,9229,13915,13733,7902,8619,9546,8901,14443,14461,14417\nMYLK - mRNA,5492,4561,5986,5953,4060,4249,6274,4336,4040,6092,6232,4690,6962,9858,8651,6419,9448,11364,9816,5840,10320,10865,8958,7352,9497,20026,13136\nGNAI2 - mRNA,5783,3608,3883,4868,4849,3884,4967,6176,6443,9762,4791,4543,5520,6963,8101,8256,6276,7201,7882,6593,8050,8804,7947,8486,12828,12452,12417\nMYH9 - mRNA,5310,6037,5399,5324,5509,4221,5988,7532,7287,10790,6322,6264,8971,7863,9737,8724,8736,9563,10339,8240,10196,10659,8436,9344,12562,16990,12167\nACTG1 - mRNA,20965,20218,21078,22372,22404,20208,26933,29401,35918,56317,27367,31616,31926,35440,38155,38375,30042,35881,49346,45704,41684,47456,43223,43901,59851,68087,67691\nFN1 - mRNA,13100,16017,15458,16275,13823,12925,10663,11611,15168,22846,11231,20350,25240,27790,23179,33049,22865,26979,19181,27624,19056,20682,28544,27758,49731,51521,44067\nACTB - mRNA,13009,20118,18241,13916,17040,15746,20988,23652,27296,41112,24531,27057,29930,21283,27020,26634,24984,32445,40956,33183,30769,33264,34811,35954,40554,57682,50254\nHSPA8 - mRNA,23617,14585,16251,17265,18728,14983,18705,22835,19332,28568,21088,16190,21391,27612,32034,28764,23439,25595,34483,22316,34484,35889,30467,37751,42235,46721,45494\nLDHA - mRNA,16867,10527,9163,6464,8126,6487,5324,11798,11703,16159,14009,13555,14491,17346,13959,15668,10576,11566,21250,17005,22309,22432,30493,32072,21560,33433,42172\nCCND1 - mRNA,9755,14189,7894,7303,5683,7990,8095,8835,8039,12389,20065,25458,21554,18900,10290,10974,10204,11658,33798,36990,17860,19629,29665,31382,17094,34900,45147\nMYL9 - mRNA,8378,11694,9257,12515,8749,9108,11540,11247,13945,20817,11287,13141,16216,17207,14541,17344,17670,18147,18178,18102,12508,12896,15974,12508,24828,32417,23362\nITGB1 - mRNA,13291,10133,10898,12349,7903,8547,10690,9503,10992,19122,12519,13519,19345,18803,13909,15560,14160,20866,25013,21549,15274,20303,17556,22184,28215,33232,31283\nCD44 - mRNA,5156,11001,9577,8101,10090,10861,13332,12389,15681,28118,16189,18576,19090,10873,14625,15813,11356,17488,31203,27496,17000,21791,19684,26584,28362,26826,32810\nRPL19 - mRNA,19433,11768,13143,14674,13124,13385,19587,14977,15105,23469,16510,15138,17131,17411,21096,19344,20452,22405,27595,21594,24429,26643,21291,24681,29367,35334,32462\nRPLP0 - mRNA,20751,9681,14376,14696,12186,11755,20060,12996,13446,23558,13596,13010,16691,18039,19202,17137,18269,24631,26775,20282,21839,27736,19530,25373,30316,34220,34019\n";
	colAnnotation_UQh1YWZvRyUX_kmCGIztRmQ = "File.Name,Description,Batch.ID,Sample.Name,Cartridge.Id,Lane.Number,Analyte.Types,System.Type,Gene.RLF,Scanner.Id,Stage.Position,Scanned.Date,Gender,QC.Flag,FOV.Count,FOV.Counted,Binding.Density,Is.Reference,Imaging.QC.Flag,Imaging.QC,Binding.Density.QC.Flag,Binding.Density.QC,Positive.Control.QC.Flag,Positive.Control.QC,Limit.of.Detection.QC.Flag.of.mRNA,Limit.of.Detection.QC.of.mRNA,New.Annotation,\n20161013_sebelius2-101316_01_01.RCC,CRL1790 Control,SA,01,sebelius2-101316,1,mRNA,NONE,SEBELIUS_2_C4636,1507C0300,3,2016-10-13 00:00:00.0,Unknown,false,280,279,0.74,false,false,0.996429,false,0.74,false,0.995739,false,10.502747,Control,\n20161013_sebelius2-101316_01_03.RCC,CRL1790 Control,SA,01,sebelius2-101316,3,mRNA,NONE,SEBELIUS_2_C4636,1507C0300,3,2016-10-13 00:00:00.0,Unknown,false,280,278,0.84,false,false,0.992857,false,0.84,false,0.985546,false,11.920769,Control,\n20161013_sebelius2-101316_01_04.RCC,CRL1790 miR142,SA,01,sebelius2-101316,4,mRNA,NONE,SEBELIUS_2_C4636,1507C0300,3,2016-10-13 00:00:00.0,Unknown,false,280,273,0.63,false,false,0.975,false,0.63,false,0.987008,false,7.339143,miR142,\n20180703_1790-jr1_SA40miR142_01.RCC,CRL1790 miR142,SA,SA40miR142,1790-jr1,1,mRNA,NONE,SEBELIUS_2_C4636,1507C0300,3,2018-07-03 00:00:00.0,Unknown,false,555,524,0.44,false,false,0.944144,false,0.44,false,0.996285,false,6.5,miR142,\n20180703_1790-jr1_SA44miR890_02.RCC,CRL1790 miR890,SA,SA44miR890,1790-jr1,2,mRNA,NONE,SEBELIUS_2_C4636,1507C0300,3,2018-07-03 00:00:00.0,Unknown,false,555,536,0.35,false,false,0.965766,false,0.35,false,0.990552,false,6.375,miR890,\n20180703_1790-jr1_SA48bcel67_03.RCC,CRL1790 cel67,SA,SA48bcel67,1790-jr1,3,mRNA,NONE,SEBELIUS_2_C4636,1507C0300,3,2018-07-03 00:00:00.0,Unknown,false,555,522,0.5,false,false,0.940541,false,0.5,false,0.990499,false,10.398527,cel67,\n20180703_1790-jr1_SA50bmiR18b_04.RCC,CRL1790 miR18b,SA,SA50bmiR18b,1790-jr1,4,mRNA,NONE,SEBELIUS_2_C4636,1507C0300,3,2018-07-03 00:00:00.0,Unknown,false,555,495,0.4,false,false,0.891892,false,0.4,false,0.992382,false,7.32534,miR18b,\n20180703_1790-jr1_SA51miR890_05.RCC,CRL1790 miR890,SA,SA51miR890,1790-jr1,5,mRNA,NONE,SEBELIUS_2_C4636,1507C0300,3,2018-07-03 00:00:00.0,Unknown,false,555,526,0.4,false,false,0.947748,false,0.4,false,0.989921,false,6.355745,miR890,\n20180703_1790-jr1_SA54amiR155_06.RCC,CRL1790 miR155,SA,SA54amiR155,1790-jr1,6,mRNA,NONE,SEBELIUS_2_C4636,1507C0300,3,2018-07-03 00:00:00.0,Unknown,false,555,481,0.35,false,false,0.866667,false,0.35,false,0.994494,false,6.375,miR155,\n20180703_1790-jr1_SA54bmiR155_07.RCC,CRL1790 miR155,SA,SA54bmiR155,1790-jr1,7,mRNA,NONE,SEBELIUS_2_C4636,1507C0300,3,2018-07-03 00:00:00.0,Unknown,false,555,527,0.47,false,false,0.94955,false,0.47,false,0.986179,false,6.875,miR155,\n20180703_1790-jr1_SA57Control_08.RCC,CRL1790 Control,SA,SA57Control,1790-jr1,8,mRNA,NONE,SEBELIUS_2_C4636,1507C0300,3,2018-07-03 00:00:00.0,Unknown,false,555,520,0.5,false,false,0.936937,false,0.5,false,0.992037,false,7.27618,Control,\n20180703_1790-jr1_SA60miR142_09.RCC,CRL1790 miR142,SA,SA60miR142,1790-jr1,9,mRNA,NONE,SEBELIUS_2_C4636,1507C0300,3,2018-07-03 00:00:00.0,Unknown,false,555,506,0.32,false,false,0.911712,false,0.32,false,0.996168,false,6.375,miR142,\n20180703_1790-jr1_SA61miR155_10.RCC,CRL1790 miR155,SA,SA61miR155,1790-jr1,10,mRNA,NONE,SEBELIUS_2_C4636,1507C0300,3,2018-07-03 00:00:00.0,Unknown,false,555,473,0.51,false,false,0.852252,false,0.51,false,0.990233,false,6.75,miR155,\n20180703_1790-jr1_SA62miR18b_11.RCC,CRL1790 miR18b,SA,SA62miR18b,1790-jr1,11,mRNA,NONE,SEBELIUS_2_C4636,1507C0300,3,2018-07-03 00:00:00.0,Unknown,false,555,492,0.39,false,false,0.886486,false,0.39,false,0.995342,false,7.838203,miR18b,\n20180703_1790-jr1_SA63miR890_12.RCC,CRL1790 miR890,SA,SA63miR890,1790-jr1,12,mRNA,NONE,SEBELIUS_2_C4636,1507C0300,3,2018-07-03 00:00:00.0,Unknown,false,555,477,0.46,false,false,0.859459,false,0.46,false,0.988295,false,6.25,miR890,\n20180712_1790-jr2_SA36Control_01.RCC,CRL1790 Control,SA,SA36Control,1790-jr2,1,mRNA,NONE,SEBELIUS_2_C4636,1507C0300,4,2018-07-03 00:00:00.0,Unknown,false,555,529,0.95,false,false,0.953153,false,0.95,false,0.993858,false,13.365773,Control,\n20180712_1790-jr2_SA38cel67_02.RCC,CRL1790 cel67,SA,SA38cel67,1790-jr2,2,mRNA,NONE,SEBELIUS_2_C4636,1507C0300,4,2018-07-03 00:00:00.0,Unknown,false,555,534,0.47,false,false,0.962162,false,0.47,false,0.988335,false,8.959881,cel67,\n20180712_1790-jr2_SA40miR142_03.RCC,CRL1790 miR142,SA,SA40miR142,1790-jr2,3,mRNA,NONE,SEBELIUS_2_C4636,1507C0300,4,2018-07-03 00:00:00.0,Unknown,false,555,530,0.6,false,false,0.954955,false,0.6,false,0.994833,false,10.666523,miR142,\n20180712_1790-jr2_SA44miR890_04.RCC,CRL1790 miR890,SA,SA44miR890,1790-jr2,4,mRNA,NONE,SEBELIUS_2_C4636,1507C0300,4,2018-07-03 00:00:00.0,Unknown,false,555,528,0.61,false,false,0.951351,false,0.61,false,0.994373,false,8.640393,miR890,\n20180712_1790-jr2_SA48bcel67_05.RCC,CRL1790 cel67,SA,SA48bcel67,1790-jr2,5,mRNA,NONE,SEBELIUS_2_C4636,1507C0300,4,2018-07-03 00:00:00.0,Unknown,false,555,531,0.64,false,false,0.956757,false,0.64,false,0.996108,false,8.535574,cel67,\n20180712_1790-jr2_SA49amiR142_06.RCC,CRL1790 miR142,SA,SA49amiR142,1790-jr2,6,mRNA,NONE,SEBELIUS_2_C4636,1507C0300,4,2018-07-03 00:00:00.0,Unknown,false,555,534,0.58,false,false,0.962162,false,0.58,false,0.990119,false,7.140393,miR142,\n20180712_1790-jr2_SA51bmiR890_07.RCC,CRL1790 miR890,SA,SA51bmiR890,1790-jr2,7,mRNA,NONE,SEBELIUS_2_C4636,1507C0300,4,2018-07-03 00:00:00.0,Unknown,false,555,527,0.73,false,false,0.94955,false,0.73,false,0.992824,false,13.141959,miR890,\n20180712_1790-jr2_SA57Control_08.RCC,CRL1790 Control,SA,SA57Control,1790-jr2,8,mRNA,NONE,SEBELIUS_2_C4636,1507C0300,4,2018-07-03 00:00:00.0,Unknown,false,555,530,0.83,false,false,0.954955,false,0.83,false,0.992413,false,15.653154,Control,\n20180712_1790-jr2_SA59cel67_09.RCC,CRL1790 cel67,SA,SA59cel67,1790-jr2,9,mRNA,NONE,SEBELIUS_2_C4636,1507C0300,4,2018-07-03 00:00:00.0,Unknown,false,555,530,0.65,false,false,0.954955,false,0.65,false,0.995539,false,10.772942,cel67,\n20180712_1790-jr2_SA61miR155_10.RCC,CRL1790 miR155,SA,SA61miR155,1790-jr2,10,mRNA,NONE,SEBELIUS_2_C4636,1507C0300,4,2018-07-03 00:00:00.0,Unknown,false,555,527,0.79,false,false,0.94955,false,0.79,false,0.997276,false,13.082381,miR155,\n20180712_1790-jr2_SA62miR18b_11.RCC,CRL1790 miR18b,SA,SA62miR18b,1790-jr2,11,mRNA,NONE,SEBELIUS_2_C4636,1507C0300,4,2018-07-03 00:00:00.0,Unknown,false,555,533,0.63,false,false,0.96036,false,0.63,false,0.996031,false,10.641959,miR18b,\n20180712_1790-jr2_SA63miR890_12.RCC,CRL1790 miR890,SA,SA63miR890,1790-jr2,12,mRNA,NONE,SEBELIUS_2_C4636,1507C0300,4,2018-07-03 00:00:00.0,Unknown,false,555,524,0.66,false,false,0.944144,false,0.66,false,0.993428,false,10.367889,miR890,\n";
	rowAnnotation_UQh1YWZvRyUX_kmCGIztRmQ = "Dimname,ProbeID,Codeset.Name,Probe.Label,Analyte.Type,Is.Control,Control.Type,Related.Probes,Probe.Annotation,KEGG.Pathways,Official.Gene.Name,Control.Conc,probeLabel.Type,below.threshold\nPIK3R5 - mRNA,NM_001142633.1:3335,SEBELIUS_2_C4636,PIK3R5,mRNA,0,,,Synthesis of PIPs at the plasma membrane;G beta.CL.gamma signalling through PI3Kgamma;GPVI-mediated activation cascade,hsa04012;hsa04014;hsa04015;hsa04022;hsa04024;hsa04062;hsa04066;hsa04068;hsa04070;hsa04071;hsa04072;hsa04150;hsa04151;hsa04152;hsa04210;hsa04211;hsa04213;hsa04261;hsa04370;hsa04380;hsa04510;hsa04550;hsa04611;hsa04620;hsa04630;hsa04650;hsa04660;hsa04662;hsa04664;hsa04666;hsa04668;hsa04670;hsa04722;hsa04725;hsa04750;hsa04810;hsa04910;hsa04914;hsa04915;hsa04917;hsa04919;hsa04921;hsa04923;hsa04930;hsa04931;hsa04932;hsa04933;hsa04960;hsa04973;hsa05100;hsa05142;hsa05145;hsa05146;hsa05160;hsa05161;hsa05162;hsa05164;hsa05166;hsa05169;hsa05200;hsa05203;hsa05205;hsa05210;hsa05211;hsa05212;hsa05213;hsa05214;hsa05215;hsa05218;hsa05220;hsa05221;hsa05222;hsa05223;hsa05230;hsa05231,PIK3R5,NA,PIK3R5 - mRNA,True\nCDX2 - mRNA,NM_001265.2:1812,SEBELIUS_2_C4636,CDX2,mRNA,0,,,,,CDX2,NA,CDX2 - mRNA,True\nTJP3 - mRNA,NM_014428.1:1925,SEBELIUS_2_C4636,TJP3,mRNA,0,,,,hsa04530,TJP3,NA,TJP3 - mRNA,True\nCXCR4 - mRNA,NM_003467.2:1335,SEBELIUS_2_C4636,CXCR4,mRNA,0,,,,hsa04060;hsa04062;hsa04144;hsa04360;hsa04670;hsa05200,CXCR4,NA,CXCR4 - mRNA,True\nCGN - mRNA,NM_020770.2:1112,SEBELIUS_2_C4636,CGN,mRNA,0,,,TGF-beta receptor signaling in EMT (epithelial to mesenchymal transition),hsa04530,CGN,NA,CGN - mRNA,True\nACTN3 - mRNA,NM_001104.1:296,SEBELIUS_2_C4636,ACTN3,mRNA,0,,,Nephrin interactions,hsa04510;hsa04520;hsa04530;hsa04670;hsa04810;hsa05146;hsa05203;hsa05412,ACTN3,NA,ACTN3 - mRNA,True\nCA2 - mRNA,NM_000067.2:575,SEBELIUS_2_C4636,CA2,mRNA,0,,,,hsa04971;hsa04972,CA2,NA,CA2 - mRNA,True\nCRB3 - mRNA,NM_139161.3:300,SEBELIUS_2_C4636,CRB3,mRNA,0,,,Tight junction interactions,hsa04530,CRB3,NA,CRB3 - mRNA,True\nCLDN4 - mRNA,NM_001305.3:1242,SEBELIUS_2_C4636,CLDN4,mRNA,0,,,Tight junction interactions,hsa04514;hsa04530;hsa04670;hsa05160,CLDN4,NA,CLDN4 - mRNA,True\nALPI - mRNA,NM_001631.3:1270,SEBELIUS_2_C4636,ALPI,mRNA,0,,,,hsa01100,ALPI,NA,ALPI - mRNA,True\nPIK3CG - mRNA,NM_002649.2:2125,SEBELIUS_2_C4636,PIK3CG,mRNA,0,,,Synthesis of PIPs at the plasma membrane;G beta.CL.gamma signalling through PI3Kgamma;GPVI-mediated activation cascade,hsa00562;hsa04012;hsa04014;hsa04015;hsa04022;hsa04024;hsa04062;hsa04066;hsa04068;hsa04070;hsa04071;hsa04072;hsa04150;hsa04151;hsa04152;hsa04210;hsa04211;hsa04213;hsa04261;hsa04370;hsa04380;hsa04510;hsa04550;hsa04611;hsa04620;hsa04630;hsa04650;hsa04660;hsa04662;hsa04664;hsa04666;hsa04668;hsa04670;hsa04722;hsa04725;hsa04750;hsa04810;hsa04910;hsa04914;hsa04915;hsa04917;hsa04919;hsa04921;hsa04923;hsa04930;hsa04931;hsa04932;hsa04933;hsa04960;hsa04973;hsa05100;hsa05142;hsa05145;hsa05146;hsa05160;hsa05161;hsa05162;hsa05164;hsa05166;hsa05169;hsa05200;hsa05203;hsa05205;hsa05210;hsa05211;hsa05212;hsa05213;hsa05214;hsa05215;hsa05218;hsa05220;hsa05221;hsa05222;hsa05223;hsa05230;hsa05231,PIK3CG,NA,PIK3CG - mRNA,True\nIL1R2 - mRNA,NM_001261419.1:186,SEBELIUS_2_C4636,IL1R2,mRNA,0,,,,hsa04010;hsa04060;hsa04640;hsa05146;hsa05166;hsa05202,IL1R2,NA,IL1R2 - mRNA,True\nHPSE - mRNA,NM_006665.3:1625,SEBELIUS_2_C4636,HPSE,mRNA,0,,,,hsa01100;hsa05205,HPSE,NA,HPSE - mRNA,True\nITGB4 - mRNA,NM_001005731.1:4151,SEBELIUS_2_C4636,ITGB4,mRNA,0,,,Syndecan interactions;Assembly of collagen fibrils and other multimeric structures;Non-integrin membrane-ECM interactions,hsa04151;hsa04510;hsa04512;hsa04810;hsa05410;hsa05412;hsa05414,ITGB4,NA,ITGB4 - mRNA,True\nWAS - mRNA,NM_000377.2:90,SEBELIUS_2_C4636,WAS,mRNA,0,,,RHO GTPases Activate WASPs and WAVEs;Regulation of actin dynamics for phagocytic cup formation,hsa04062;hsa04144;hsa04520;hsa04666;hsa04810;hsa05100;hsa05130;hsa05131;hsa05132;hsa05231,WAS,NA,WAS - mRNA,True\nVAV3 - mRNA,NM_006113.4:1632,SEBELIUS_2_C4636,VAV3,mRNA,0,,NM_001079874.1:2680,Rho GTPase cycle;DAP12 signaling;EPH-ephrin mediated repulsion of cells;VEGFR2 mediated vascular permeability;NRAGE signals death through JNK;GPVI-mediated activation cascade;G alpha (1213) signalling events;FCERI mediated MAPK activation;Regulation of actin dynamics for phagocytic cup formation;VEGFA-VEGFR2 Pathway,hsa04024;hsa04062;hsa04510;hsa04650;hsa04660;hsa04662;hsa04664;hsa04666;hsa04670;hsa04810,VAV3,NA,VAV3 - mRNA,True\nMYH14 - mRNA,NM_001077186.1:6304,SEBELIUS_2_C4636,MYH14,mRNA,0,,,EPHA-mediated growth cone collapse;RHO GTPases Activate ROCKs;RHO GTPases activate CIT;RHO GTPases activate PAKs;RHO GTPases activate PKNs;Sema4D induced cell migration and growth-cone collapse,hsa04530;hsa04810;hsa05132,MYH14,NA,MYH14 - mRNA,True\nEPB41L3 - mRNA,NM_012307.2:3758,SEBELIUS_2_C4636,EPB41L3,mRNA,0,,,,hsa04530,EPB41L3,NA,EPB41L3 - mRNA,True\nPRKCB - mRNA,NM_212535.1:1750,SEBELIUS_2_C4636,PRKCB,mRNA,0,,,Disinhibition of SNARE formation;Trafficking of AMPA receptors;VEGFA-VEGFR2 Pathway;PCPCE pathway,hsa04010;hsa04012;hsa04014;hsa04015;hsa04020;hsa04062;hsa04064;hsa04066;hsa04070;hsa04071;hsa04150;hsa04270;hsa04310;hsa04370;hsa04510;hsa04530;hsa04540;hsa04650;hsa04662;hsa04666;hsa04670;hsa04713;hsa04720;hsa04723;hsa04724;hsa04725;hsa04726;hsa04727;hsa04728;hsa04730;hsa04750;hsa04912;hsa04916;hsa04919;hsa04921;hsa04931;hsa04933;hsa04960;hsa04971;hsa04972;hsa04973;hsa05031;hsa05032;hsa05140;hsa05146;hsa05161;hsa05164;hsa05200;hsa05205;hsa05206;hsa05214;hsa05223;hsa05231,PRKCB,NA,PRKCB - mRNA,True\nVAV3 b - mRNA,NM_001079874.1:2680,SEBELIUS_2_C4636,VAV3 b,mRNA,0,,NM_006113.4:1632,Rho GTPase cycle;DAP12 signaling;EPH-ephrin mediated repulsion of cells;VEGFR2 mediated vascular permeability;NRAGE signals death through JNK;GPVI-mediated activation cascade;G alpha (1213) signalling events;FCERI mediated MAPK activation;Regulation of actin dynamics for phagocytic cup formation;VEGFA-VEGFR2 Pathway,hsa04024;hsa04062;hsa04510;hsa04650;hsa04660;hsa04662;hsa04664;hsa04666;hsa04670;hsa04810,VAV3,NA,VAV3 b - mRNA,True\nMYH11 - mRNA,NM_001040113.1:920,SEBELIUS_2_C4636,MYH11,mRNA,0,,,EPHA-mediated growth cone collapse;RHO GTPases Activate ROCKs;Smooth Muscle Contraction;RHO GTPases activate CIT;RHO GTPases activate PAKs;RHO GTPases activate PKNs;Sema4D induced cell migration and growth-cone collapse,hsa04270;hsa04530,MYH11,NA,MYH11 - mRNA,True\nCLDN3 - mRNA,NM_001306.3:606,SEBELIUS_2_C4636,CLDN3,mRNA,0,,,Tight junction interactions,hsa04514;hsa04530;hsa04670;hsa05160,CLDN3,NA,CLDN3 - mRNA,True\nCA1 - mRNA,NM_001738.2:895,SEBELIUS_2_C4636,CA1,mRNA,0,,,,,CA1,NA,CA1 - mRNA,True\nPRKCZ - mRNA,NM_002744.4:771,SEBELIUS_2_C4636,PRKCZ,mRNA,0,,,TGF-beta receptor signaling in EMT (epithelial to mesenchymal transition);VEGFR2 mediated cell proliferation;GPVI-mediated activation cascade;VEGFA-VEGFR2 Pathway,hsa04015;hsa04062;hsa04071;hsa04144;hsa04390;hsa04530;hsa04611;hsa04910;hsa04930;hsa04931;hsa04933,PRKCZ,NA,PRKCZ - mRNA,False\nAXIN2 - mRNA,NM_004655.3:1035,SEBELIUS_2_C4636,AXIN2,mRNA,0,,,Binding of TCFLEF.CL.CTNNB1 to target gene promoters;Formation of the beta-catenin.CL.TCF transactivating complex,hsa04310;hsa04390;hsa04550;hsa05200;hsa05210;hsa05213;hsa05217,AXIN2,NA,AXIN2 - mRNA,False\nCLDN15 - mRNA,NM_014343.1:1129,SEBELIUS_2_C4636,CLDN15,mRNA,0,,,Tight junction interactions,hsa04514;hsa04530;hsa04670;hsa05160,CLDN15,NA,CLDN15 - mRNA,False\nAPC2 - mRNA,NM_005883.2:8120,SEBELIUS_2_C4636,APC2,mRNA,0,,,,hsa04310;hsa04390;hsa04550;hsa04810;hsa05166;hsa05200;hsa05206;hsa05210;hsa05213;hsa05217,APC2,NA,APC2 - mRNA,False\nHCLS1 - mRNA,NM_005335.4:1846,SEBELIUS_2_C4636,HCLS1,mRNA,0,,,,hsa04530;hsa05100;hsa05130;hsa05131;hsa05205,HCLS1,NA,HCLS1 - mRNA,False\nPDGFA - mRNA,NM_002607.5:2460,SEBELIUS_2_C4636,PDGFA,mRNA,0,,,Non-integrin membrane-ECM interactions;PIP3 activates AKT signaling;Constitutive Signaling by Aberrant PI3K in Cancer;RAFMAP kinase cascade;Platelet degranulation ;Signaling by PDGF;Downstream signal transduction;GAB1 signalosome;Signaling by SCF-KIT;PI3KAKT activation;Role of LAT2NTALLAB on calcium mobilization;DAP12 signaling;Signaling by ERBB2;VEGFR2 mediated cell proliferation;Frs2-mediated activation;ARMS-mediated activation;Gastrin-CREB signalling pathway via PKC and MAPK;Interleukin receptor SHC signaling;FCERI mediated MAPK activation;VEGFA-VEGFR2 Pathway;IRS-related events triggered by IGF1R,hsa04010;hsa04014;hsa04015;hsa04060;hsa04072;hsa04151;hsa04510;hsa04540;hsa04810;hsa05166;hsa05200;hsa05202;hsa05206;hsa05214;hsa05215;hsa05218;hsa05231,PDGFA,NA,PDGFA - mRNA,False\nF11R - mRNA,NM_144503.1:3650,SEBELIUS_2_C4636,F11R,mRNA,0,,,TGF-beta receptor signaling in EMT (epithelial to mesenchymal transition);Tight junction interactions;Integrin cell surface interactions;Cell surface interactions at the vascular wall,hsa04514;hsa04530;hsa04670;hsa05120,F11R,NA,F11R - mRNA,False\nEPB41 - mRNA,NM_001166005.1:1330,SEBELIUS_2_C4636,EPB41,mRNA,0,,,,hsa04530,EPB41,NA,EPB41 - mRNA,False\nITGA7 - mRNA,NM_002206.1:1170,SEBELIUS_2_C4636,ITGA7,mRNA,0,,,Laminin interactions;ECM proteoglycans;Integrin cell surface interactions,hsa04151;hsa04510;hsa04512;hsa04810;hsa05410;hsa05412;hsa05414,ITGA7,NA,ITGA7 - mRNA,False\nMYL12A - mRNA,NM_006471.2:78,SEBELIUS_2_C4636,MYL12A,mRNA,0,,NM_001303049.1:9;NM_001303048.1:27,Smooth Muscle Contraction;Ephrin signaling,hsa04510;hsa04530;hsa04611;hsa04670;hsa04810,MYL12A,NA,MYL12A - mRNA,False\nBCL2 - mRNA,NM_000657.2:5,SEBELIUS_2_C4636,BCL2,mRNA,0,,,Activation of BAD and translocation to mitochondria ,hsa04064;hsa04066;hsa04071;hsa04141;hsa04151;hsa04210;hsa04261;hsa04510;hsa04630;hsa04722;hsa04725;hsa04933;hsa05145;hsa05152;hsa05161;hsa05169;hsa05200;hsa05206;hsa05210;hsa05215;hsa05222,BCL2,NA,BCL2 - mRNA,False\nITGA4 - mRNA,NM_000885.4:975,SEBELIUS_2_C4636,ITGA4,mRNA,0,,,Integrin cell surface interactions;Cell surface interactions at the vascular wall,hsa04151;hsa04510;hsa04512;hsa04514;hsa04640;hsa04670;hsa04810;hsa05140;hsa05410;hsa05412;hsa05414,ITGA4,NA,ITGA4 - mRNA,False\nJAM2 - mRNA,NM_001270407.1:840,SEBELIUS_2_C4636,JAM2,mRNA,0,,,Integrin cell surface interactions;Cell surface interactions at the vascular wall,hsa04514;hsa04530;hsa04670;hsa05120,JAM2,NA,JAM2 - mRNA,True\nLLGL2 - mRNA,NM_001015002.1:1065,SEBELIUS_2_C4636,LLGL2,mRNA,0,,,,hsa04390;hsa04530,LLGL2,NA,LLGL2 - mRNA,True\nTCF7 - mRNA,NM_201633.2:519,SEBELIUS_2_C4636,TCF7,mRNA,0,,,,,,NA,TCF7 - mRNA,False\nCLDN7 - mRNA,NM_001307.3:175,SEBELIUS_2_C4636,CLDN7,mRNA,0,,,Tight junction interactions,hsa04514;hsa04530;hsa04670;hsa05160,CLDN7,NA,CLDN7 - mRNA,True\nCLDN8 - mRNA,NM_199328.2:805,SEBELIUS_2_C4636,CLDN8,mRNA,0,,,Tight junction interactions,hsa04514;hsa04530;hsa04670;hsa05160,CLDN8,NA,CLDN8 - mRNA,True\nLYZ - mRNA,NM_000239.2:305,SEBELIUS_2_C4636,LYZ,mRNA,0,,,,,LYZ,NA,LYZ - mRNA,True\nGAPDH - mRNA,NM_002046.3:972,SEBELIUS_2_C4636,GAPDH,mRNA,0,,,Glycolysis,hsa01100;hsa04066;hsa05010,GAPDH,NA,GAPDH - mRNA,False\nACTG1 - mRNA,NM_001614.2:1615,SEBELIUS_2_C4636,ACTG1,mRNA,0,,,RHO GTPases Activate WASPs and WAVEs;Translocation of GLUT4 to the plasma membrane;Recycling pathway of L1;Interaction between L1 and Ankyrins;EPHB-mediated forward signaling;EPH-ephrin mediated repulsion of cells;RHO GTPases Activate Formins;RHO GTPases activate IQGAPs;Cell-extracellular matrix interactions;Regulation of actin dynamics for phagocytic cup formation;RAFMAP kinase cascade;Downstream signal transduction;VEGFR2 mediated cell proliferation;Signaling by SCF-KIT;Frs2-mediated activation;ARMS-mediated activation;Gastrin-CREB signalling pathway via PKC and MAPK;Interleukin receptor SHC signaling;FCERI mediated MAPK activation;DAP12 signaling;Signaling by PDGF;VEGFA-VEGFR2 Pathway;Signaling by ERBB2;IRS-related events triggered by IGF1R,hsa04015;hsa04145;hsa04390;hsa04510;hsa04520;hsa04530;hsa04611;hsa04670;hsa04810;hsa04919;hsa04921;hsa05100;hsa05110;hsa05130;hsa05131;hsa05132;hsa05164;hsa05205;hsa05410;hsa05412;hsa05414;hsa05416,ACTG1,NA,ACTG1 - mRNA,False\nACTB - mRNA,NM_001101.2:1010,SEBELIUS_2_C4636,ACTB,mRNA,0,,,EPHB-mediated forward signaling;Interaction between L1 and Ankyrins;DNA Damage Recognition in GG-NER;EPH-ephrin mediated repulsion of cells;Prefoldin mediated transfer of substrate  to CCTTriC;RHO GTPases Activate Formins;RHO GTPases activate IQGAPs;Cell-extracellular matrix interactions;Regulation of actin dynamics for phagocytic cup formation;Folding of actin by CCTTriC;B-WICH complex positively regulates rRNA expression;RAFMAP kinase cascade;Downstream signal transduction;VEGFR2 mediated cell proliferation;Signaling by SCF-KIT;Frs2-mediated activation;ARMS-mediated activation;Gastrin-CREB signalling pathway via PKC and MAPK;Interleukin receptor SHC signaling;FCERI mediated MAPK activation;DAP12 signaling;Signaling by PDGF;VEGFA-VEGFR2 Pathway;Signaling by ERBB2;IRS-related events triggered by IGF1R,hsa04015;hsa04145;hsa04390;hsa04510;hsa04520;hsa04530;hsa04611;hsa04670;hsa04810;hsa04919;hsa04921;hsa04971;hsa05100;hsa05110;hsa05130;hsa05131;hsa05132;hsa05164;hsa05205;hsa05410;hsa05412;hsa05414;hsa05416,ACTB,NA,ACTB - mRNA,False\nHSPA8 - mRNA,NM_006597.3:2111,SEBELIUS_2_C4636,HSPA8,mRNA,0,,,AUF1 (hnRNP D0) binds and destabilizes mRNA;GABA synthesis# release# reuptake and degradation,hsa04010;hsa04141;hsa04144;hsa04213;hsa04915;hsa05145;hsa05162;hsa05164;hsa05169,HSPA8,NA,HSPA8 - mRNA,False\nCFL1 - mRNA,NM_005507.2:225,SEBELIUS_2_C4636,CFL1,mRNA,0,,,EPHB-mediated forward signaling;Sema3A PAK dependent Axon repulsion;Platelet degranulation ;Regulation of actin dynamics for phagocytic cup formation,hsa04360;hsa04666;hsa04810;hsa05133,CFL1,NA,CFL1 - mRNA,False\nFLNA - mRNA,NM_001456.3:7335,SEBELIUS_2_C4636,FLNA,mRNA,0,,,RHO GTPases activate PAKs;Platelet degranulation ;Cell-extracellular matrix interactions,hsa04010;hsa04510;hsa05132;hsa05205,FLNA,NA,FLNA - mRNA,False\nMYL12B - mRNA,NM_001144945.1:749,SEBELIUS_2_C4636,MYL12B,mRNA,0,,,EPHA-mediated growth cone collapse;RHO GTPases Activate ROCKs;Smooth Muscle Contraction;RHO GTPases activate CIT;RHO GTPases activate PAKs;RHO GTPases activate PKNs;Sema4D induced cell migration and growth-cone collapse,hsa04510;hsa04530;hsa04611;hsa04670;hsa04810,MYL12B,NA,MYL12B - mRNA,False\nMYH9 - mRNA,NM_002473.3:3153,SEBELIUS_2_C4636,MYH9,mRNA,0,,,EPHA-mediated growth cone collapse;RHO GTPases Activate ROCKs;RHO GTPases activate CIT;RHO GTPases activate PAKs;RHO GTPases activate PKNs;Regulation of actin dynamics for phagocytic cup formation;Sema4D induced cell migration and growth-cone collapse,hsa04530;hsa04810;hsa05132,MYH9,NA,MYH9 - mRNA,False\nITGA1 - mRNA,NM_181501.1:1875,SEBELIUS_2_C4636,ITGA1,mRNA,0,,,Laminin interactions;Other semaphorin interactions;Integrin cell surface interactions,hsa04151;hsa04510;hsa04512;hsa04640;hsa04810;hsa05410;hsa05412;hsa05414,ITGA1,NA,ITGA1 - mRNA,False\nMYLK - mRNA,NM_053032.2:710,SEBELIUS_2_C4636,MYLK,mRNA,0,,,Smooth Muscle Contraction;RHO GTPases activate PAKs,hsa04020;hsa04022;hsa04270;hsa04510;hsa04611;hsa04810;hsa04921;hsa04971,MYLK,NA,MYLK - mRNA,False\nGNAI2 - mRNA,NM_002070.2:1000,SEBELIUS_2_C4636,GNAI2,mRNA,0,,,ADP signalling through P2Y purinoceptor 12;Adrenaline#noradrenaline inhibits insulin secretion,hsa04015;hsa04022;hsa04024;hsa04062;hsa04071;hsa04261;hsa04360;hsa04530;hsa04540;hsa04611;hsa04670;hsa04713;hsa04723;hsa04724;hsa04725;hsa04726;hsa04727;hsa04728;hsa04730;hsa04914;hsa04915;hsa04916;hsa04921;hsa04923;hsa04971;hsa05032;hsa05034;hsa05133;hsa05142;hsa05145;hsa05200,GNAI2,NA,GNAI2 - mRNA,False\nLDHA - mRNA,NM_001165414.1:1690,SEBELIUS_2_C4636,LDHA,mRNA,0,,,,hsa01100;hsa04066;hsa04922;hsa05230,LDHA,NA,LDHA - mRNA,False\nCCND1 - mRNA,NM_053056.2:2725,SEBELIUS_2_C4636,CCND1,mRNA,0,,,Cyclin D associated events in G1;Pre-NOTCH Transcription and Translation,hsa04068;hsa04110;hsa04115;hsa04151;hsa04152;hsa04310;hsa04390;hsa04510;hsa04630;hsa04917;hsa04919;hsa04921;hsa04933;hsa05161;hsa05162;hsa05166;hsa05200;hsa05203;hsa05205;hsa05206;hsa05210;hsa05212;hsa05213;hsa05214;hsa05215;hsa05216;hsa05218;hsa05219;hsa05220;hsa05221;hsa05222;hsa05223;hsa05416,CCND1,NA,CCND1 - mRNA,False\nCD44 - mRNA,NM_001001392.1:429,SEBELIUS_2_C4636,CD44,mRNA,0,,,Hyaluronan uptake and degradation;Cell surface interactions at the vascular wall,hsa04512;hsa04640;hsa05131;hsa05169;hsa05205;hsa05206,CD44,NA,CD44 - mRNA,False\nMYL9 - mRNA,NM_181526.1:133,SEBELIUS_2_C4636,MYL9,mRNA,0,,,EPHA-mediated growth cone collapse;RHO GTPases Activate ROCKs;Smooth Muscle Contraction;RHO GTPases activate CIT;RHO GTPases activate PAKs;RHO GTPases activate PKNs;Sema4D induced cell migration and growth-cone collapse,hsa04022;hsa04024;hsa04270;hsa04510;hsa04530;hsa04670;hsa04810;hsa04921,MYL9,NA,MYL9 - mRNA,False\nITGB1 - mRNA,NM_033666.2:2000,SEBELIUS_2_C4636,ITGB1,mRNA,0,,,Laminin interactions;RHO GTPases Activate Formins;ECM proteoglycans;Fibronectin matrix formation;Integrin cell surface interactions;Immunoregulatory interactions between a Lymphoid and a non-Lymphoid cell;Localization of the PINCH-ILK-PARVIN complex to focal adhesions;Elastic fibre formation;Cell surface interactions at the vascular wall;Non-integrin membrane-ECM interactions;Cell-extracellular matrix interactions,hsa04015;hsa04145;hsa04151;hsa04360;hsa04510;hsa04512;hsa04514;hsa04611;hsa04670;hsa04810;hsa05100;hsa05130;hsa05131;hsa05133;hsa05140;hsa05145;hsa05200;hsa05205;hsa05222;hsa05410;hsa05412;hsa05414,ITGB1,NA,ITGB1 - mRNA,False\nRPLP0 - mRNA,NM_001002.3:250,SEBELIUS_2_C4636,RPLP0,mRNA,0,,,Nonsense Mediated Decay (NMD) enhanced by the Exon Junction Complex (EJC),,RPLP0,NA,RPLP0 - mRNA,False\nFN1 - mRNA,NM_212482.1:1776,SEBELIUS_2_C4636,FN1,mRNA,0,,,MAP2K and MAPK activation;p130Cas linkage to MAPK signaling for integrins;Platelet degranulation ;Molecules associated with elastic fibres;Syndecan interactions;ECM proteoglycans;Fibronectin matrix formation;Integrin cell surface interactions;RAFMAP kinase cascade;Downstream signal transduction;VEGFR2 mediated cell proliferation;Signaling by SCF-KIT;Frs2-mediated activation;ARMS-mediated activation;Gastrin-CREB signalling pathway via PKC and MAPK;Interleukin receptor SHC signaling;FCERI mediated MAPK activation;DAP12 signaling;Signaling by PDGF;VEGFA-VEGFR2 Pathway;Signaling by ERBB2;IRS-related events triggered by IGF1R;Integrin alphaIIb beta3 signaling;Non-integrin membrane-ECM interactions;Elastic fibre formation,hsa04151;hsa04510;hsa04512;hsa04810;hsa04933;hsa05100;hsa05146;hsa05200;hsa05205;hsa05222,FN1,NA,FN1 - mRNA,False\nCYFIP2 - mRNA,NM_001037332.2:4990,SEBELIUS_2_C4636,CYFIP2,mRNA,0,,,RHO GTPases Activate WASPs and WAVEs;VEGFA-VEGFR2 Pathway;Regulation of actin dynamics for phagocytic cup formation,hsa04810,CYFIP2,NA,CYFIP2 - mRNA,False\nMYL12A b - mRNA,NM_001303048.1:27,SEBELIUS_2_C4636,MYL12A b,mRNA,0,,NM_006471.2:78;NM_001303049.1:9,Smooth Muscle Contraction;Ephrin signaling,hsa04510;hsa04530;hsa04611;hsa04670;hsa04810,MYL12A,NA,MYL12A b - mRNA,False\nNLK - mRNA,NM_016231.4:618,SEBELIUS_2_C4636,NLK,mRNA,0,,,Ca2.PLS. pathway,hsa04010;hsa04068;hsa04310;hsa04520,NLK,NA,NLK - mRNA,False\nLEF1 - mRNA,NM_016269.3:1165,SEBELIUS_2_C4636,LEF1,mRNA,0,,,Ca2.PLS. pathway;Binding of TCFLEF.CL.CTNNB1 to target gene promoters;Deactivation of the beta-catenin transactivating complex;Repression of WNT target genes;Formation of the beta-catenin.CL.TCF transactivating complex,hsa04310;hsa04390;hsa04520;hsa04916;hsa05200;hsa05210;hsa05213;hsa05215;hsa05216;hsa05217;hsa05221;hsa05412,LEF1,NA,LEF1 - mRNA,False\nCLDN23 - mRNA,NM_194284.2:1768,SEBELIUS_2_C4636,CLDN23,mRNA,0,,,Tight junction interactions,hsa04514;hsa04530;hsa04670;hsa05160,CLDN23,NA,CLDN23 - mRNA,False\nOCLN - mRNA,NM_002538.3:5130,SEBELIUS_2_C4636,OCLN,mRNA,0,,,Apoptotic cleavage of cellular proteins,hsa04514;hsa04530;hsa04670;hsa05130;hsa05160,OCLN,NA,OCLN - mRNA,False\nMYL5 - mRNA,NM_002477.1:398,SEBELIUS_2_C4636,MYL5,mRNA,0,,,Smooth Muscle Contraction,hsa04510;hsa04530;hsa04670;hsa04810,MYL5,NA,MYL5 - mRNA,False\nINADL - mRNA,XM_005270347.1:343,SEBELIUS_2_C4636,INADL,mRNA,0,,,,hsa04390;hsa04530,INADL,NA,INADL - mRNA,False\nTJP2 - mRNA,NM_004817.2:2785,SEBELIUS_2_C4636,TJP2,mRNA,0,,,Apoptotic cleavage of cellular proteins,hsa04530;hsa05110,TJP2,NA,TJP2 - mRNA,False\nARAF - mRNA,NM_001654.3:245,SEBELIUS_2_C4636,ARAF,mRNA,0,,,MAP2K and MAPK activation;RAF activation;Negative regulation of MAPK pathway;RAFMAP kinase cascade;Downstream signal transduction;VEGFR2 mediated cell proliferation;Signaling by SCF-KIT;Frs2-mediated activation;ARMS-mediated activation;Gastrin-CREB signalling pathway via PKC and MAPK;Interleukin receptor SHC signaling;FCERI mediated MAPK activation;DAP12 signaling;Signaling by PDGF;VEGFA-VEGFR2 Pathway;Signaling by ERBB2;IRS-related events triggered by IGF1R,hsa04012;hsa04068;hsa04270;hsa04650;hsa04720;hsa04726;hsa04730;hsa04810;hsa04910;hsa04914;hsa05034;hsa05160;hsa05200;hsa05205;hsa05210;hsa05211;hsa05212;hsa05213;hsa05214;hsa05215;hsa05218;hsa05219;hsa05220;hsa05221;hsa05223,ARAF,NA,ARAF - mRNA,False\nMYL12A c - mRNA,NM_001303049.1:9,SEBELIUS_2_C4636,MYL12A c,mRNA,0,,NM_006471.2:78;NM_001303048.1:27,EPHA-mediated growth cone collapse;RHO GTPases Activate ROCKs;Smooth Muscle Contraction;RHO GTPases activate CIT;RHO GTPases activate PAKs;RHO GTPases activate PKNs;Sema4D induced cell migration and growth-cone collapse;Ephrin signaling,hsa04510;hsa04530;hsa04611;hsa04670;hsa04810,MYL12A,NA,MYL12A c - mRNA,False\nMAGI1 - mRNA,NM_004742.2:325,SEBELIUS_2_C4636,MAGI1,mRNA,0,,,,hsa04015;hsa04530,MAGI1,NA,MAGI1 - mRNA,False\nHSPA1A - mRNA,NM_005345.5:98,SEBELIUS_2_C4636,HSPA1A,mRNA,0,,,Regulation of HSF1-mediated heat shock response;AUF1 (hnRNP D0) binds and destabilizes mRNA,hsa04010;hsa04141;hsa04144;hsa04213;hsa04915;hsa05020;hsa05145;hsa05162;hsa05164;hsa05169,HSPA1A,NA,HSPA1A - mRNA,False\nAXIN1 - mRNA,NM_181050.1:2742,SEBELIUS_2_C4636,AXIN1,mRNA,0,,,APC truncation mutants have impaired AXIN binding;Misspliced GSK3beta mutants stabilize beta-catenin;T41 mutants of beta-catenin aren.SQT.t phosphorylated;AXIN missense mutants destabilize the destruction complex;S45 mutants of beta-catenin aren.SQT.t phosphorylated;Disassembly of the destruction complex and recruitment of AXIN to the membrane;S37 mutants of beta-catenin aren.SQT.t phosphorylated;S33 mutants of beta-catenin aren.SQT.t phosphorylated;Beta-catenin phosphorylation cascade;Truncations of AMER1 destabilize the destruction complex,hsa04310;hsa04390;hsa04550;hsa05200;hsa05210;hsa05213;hsa05217,AXIN1,NA,AXIN1 - mRNA,False\nBAIAP2 - mRNA,NM_017450.2:653,SEBELIUS_2_C4636,BAIAP2,mRNA,0,,,RHO GTPases Activate WASPs and WAVEs;VEGFA-VEGFR2 Pathway;Regulation of actin dynamics for phagocytic cup formation,hsa04520;hsa04810,BAIAP2,NA,BAIAP2 - mRNA,False\nRAC3 - mRNA,NM_005052.2:702,SEBELIUS_2_C4636,RAC3,mRNA,0,,,Rho GTPase cycle;PCPCE pathway,hsa04010;hsa04014;hsa04015;hsa04024;hsa04071;hsa04310;hsa04360;hsa04370;hsa04510;hsa04520;hsa04650;hsa04662;hsa04664;hsa04810;hsa05200;hsa05210;hsa05212;hsa05231;hsa05416,RAC3,NA,RAC3 - mRNA,False\nRAC2 - mRNA,NM_002872.4:341,SEBELIUS_2_C4636,RAC2,mRNA,0,,,Rho GTPase cycle;GPVI-mediated activation cascade;G alpha (1213) signalling events;PCPCE pathway;Sema4D induced cell migration and growth-cone collapse,hsa04010;hsa04014;hsa04015;hsa04024;hsa04062;hsa04071;hsa04310;hsa04360;hsa04370;hsa04510;hsa04520;hsa04650;hsa04662;hsa04664;hsa04666;hsa04670;hsa04810;hsa05200;hsa05210;hsa05212;hsa05231;hsa05416,RAC2,NA,RAC2 - mRNA,False\nWASF1 - mRNA,NM_001024934.1:1425,SEBELIUS_2_C4636,WASF1,mRNA,0,,,RHO GTPases Activate WASPs and WAVEs;VEGFA-VEGFR2 Pathway;Regulation of actin dynamics for phagocytic cup formation,hsa04520;hsa04666;hsa04810;hsa05100;hsa05131;hsa05132;hsa05231,WASF1,NA,WASF1 - mRNA,False\nGUSB - mRNA,NM_000181.1:1350,SEBELIUS_2_C4636,GUSB,mRNA,0,,,,hsa01100,GUSB,NA,GUSB - mRNA,False\nDNMT3A - mRNA,NM_022552.4:2056,SEBELIUS_2_C4636,DNMT3A,mRNA,0,,,,hsa01100;hsa05206,DNMT3A,NA,DNMT3A - mRNA,False\nAPC - mRNA,NM_000038.3:6850,SEBELIUS_2_C4636,APC,mRNA,0,,,Apoptotic cleavage of cellular proteins;Misspliced GSK3beta mutants stabilize beta-catenin;APC truncation mutants have impaired AXIN binding;T41 mutants of beta-catenin aren.SQT.t phosphorylated;AXIN missense mutants destabilize the destruction complex;S45 mutants of beta-catenin aren.SQT.t phosphorylated;Deactivation of the beta-catenin transactivating complex;Disassembly of the destruction complex and recruitment of AXIN to the membrane;S37 mutants of beta-catenin aren.SQT.t phosphorylated;S33 mutants of beta-catenin aren.SQT.t phosphorylated;Beta-catenin phosphorylation cascade;Truncations of AMER1 destabilize the destruction complex,hsa04310;hsa04390;hsa04550;hsa04810;hsa05166;hsa05200;hsa05206;hsa05210;hsa05213;hsa05217,APC,NA,APC - mRNA,False\nLIMK2 - mRNA,NM_001031801.1:1590,SEBELIUS_2_C4636,LIMK2,mRNA,0,,,EPHB-mediated forward signaling;RHO GTPases Activate ROCKs;Sema4D induced cell migration and growth-cone collapse,hsa04360;hsa04666;hsa04810,LIMK2,NA,LIMK2 - mRNA,False\nEPB41L1 - mRNA,NM_001258330.1:900,SEBELIUS_2_C4636,EPB41L1,mRNA,0,,,,hsa04530,EPB41L1,NA,EPB41L1 - mRNA,False\nEXOC3 - mRNA,NM_007277.4:1820,SEBELIUS_2_C4636,EXOC3,mRNA,0,,,Translocation of GLUT4 to the plasma membrane,hsa04530,EXOC3,NA,EXOC3 - mRNA,False\nTARBP2 - mRNA,NM_134323.1:1060,SEBELIUS_2_C4636,TARBP2,mRNA,0,,,MicroRNA (miRNA) biogenesis,,TARBP2,NA,TARBP2 - mRNA,False\nDVL2 - mRNA,NM_004422.2:1025,SEBELIUS_2_C4636,DVL2,mRNA,0,,,Disassembly of the destruction complex and recruitment of AXIN to the membrane;WNT mediated activation of DVL;Negative regulation of TCF-dependent signaling by DVL-interacting proteins;PCPCE pathway,hsa04310;hsa04330;hsa04390;hsa04550;hsa04916;hsa05166;hsa05200;hsa05217,DVL2,NA,DVL2 - mRNA,False\nCDK4 - mRNA,NM_000075.2:1055,SEBELIUS_2_C4636,CDK4,mRNA,0,,,Cyclin D associated events in G1;Oncogene Induced Senescence;Ubiquitin-dependent degradation of Cyclin D1;RMTs methylate histone arginines,hsa04110;hsa04115;hsa04151;hsa04530;hsa04660;hsa04933;hsa05161;hsa05162;hsa05166;hsa05200;hsa05203;hsa05212;hsa05214;hsa05218;hsa05219;hsa05220;hsa05222;hsa05223,CDK4,NA,CDK4 - mRNA,False\nMAGI3 - mRNA,NM_152900.2:6685,SEBELIUS_2_C4636,MAGI3,mRNA,0,,,,hsa04015;hsa04530,MAGI3,NA,MAGI3 - mRNA,False\nTCF7L1 - mRNA,NM_031283.1:2215,SEBELIUS_2_C4636,TCF7L1,mRNA,0,,,Ca2.PLS. pathway;Binding of TCFLEF.CL.CTNNB1 to target gene promoters;Deactivation of the beta-catenin transactivating complex;Repression of WNT target genes;Formation of the beta-catenin.CL.TCF transactivating complex,hsa04310;hsa04390;hsa04520;hsa04916;hsa05200;hsa05210;hsa05213;hsa05215;hsa05216;hsa05217;hsa05221;hsa05412,TCF7L1,NA,TCF7L1 - mRNA,False\nPIK3R1 - mRNA,NM_181504.2:1105,SEBELIUS_2_C4636,PIK3R1,mRNA,0,,,G alpha (q) signalling events;Downstream TCR signaling;Signaling by FGFR2 in disease;PIP3 activates AKT signaling;VEGFA-VEGFR2 Pathway;Synthesis of PIPs at the plasma membrane;Constitutive Signaling by Ligand-Responsive EGFR Cancer Variants;Interleukin receptor SHC signaling;G beta.CL.gamma signalling through PI3Kgamma;Regulation of signaling by CBL;Role of phospholipids in phagocytosis;Signaling by FGFR3 in disease;Constitutive Signaling by Aberrant PI3K in Cancer;Signaling by FGFR4 in disease;CD28 dependent PI3KAkt signaling;Tie2 Signaling;Signaling by FGFR1 fusion mutants;GP1b-IX-V activation signalling;GPVI-mediated activation cascade;Nephrin interactions;Constitutive Signaling by EGFRvIII;Gastrin-CREB signalling pathway via PKC and MAPK;Downstream signal transduction;GAB1 signalosome;Signaling by SCF-KIT;PI3KAKT activation;Role of LAT2NTALLAB on calcium mobilization;DAP12 signaling;Signaling by PDGF;Signaling by ERBB2;IRS-related events triggered by IGF1R;CD28 co-stimulation;Cell surface interactions at the vascular wall;Signaling by FGFR1 in disease,hsa04012;hsa04014;hsa04015;hsa04022;hsa04024;hsa04062;hsa04066;hsa04068;hsa04070;hsa04071;hsa04072;hsa04150;hsa04151;hsa04152;hsa04210;hsa04211;hsa04213;hsa04261;hsa04370;hsa04380;hsa04510;hsa04550;hsa04611;hsa04620;hsa04630;hsa04650;hsa04660;hsa04662;hsa04664;hsa04666;hsa04668;hsa04670;hsa04722;hsa04725;hsa04750;hsa04810;hsa04910;hsa04914;hsa04915;hsa04917;hsa04919;hsa04921;hsa04923;hsa04930;hsa04931;hsa04932;hsa04933;hsa04960;hsa04973;hsa05100;hsa05142;hsa05145;hsa05146;hsa05160;hsa05161;hsa05162;hsa05164;hsa05166;hsa05169;hsa05200;hsa05203;hsa05205;hsa05210;hsa05211;hsa05212;hsa05213;hsa05214;hsa05215;hsa05218;hsa05220;hsa05221;hsa05222;hsa05223;hsa05230;hsa05231,PIK3R1,NA,PIK3R1 - mRNA,False\nPAK4 - mRNA,NM_005884.3:2130,SEBELIUS_2_C4636,PAK4,mRNA,0,,,Activation of Rac,hsa04012;hsa04014;hsa04360;hsa04510;hsa04660;hsa04810;hsa05206;hsa05211,PAK4,NA,PAK4 - mRNA,False\nXPO5 - mRNA,NM_020750.2:375,SEBELIUS_2_C4636,XPO5,mRNA,0,,,MicroRNA (miRNA) biogenesis,,XPO5,NA,XPO5 - mRNA,False\nHRAS - mRNA,NM_005343.2:396,SEBELIUS_2_C4636,HRAS,mRNA,0,,,Signaling by FGFR2 in disease;Regulation of RAS by GAPs;SHC-mediated cascade.CL.FGFR3;Signaling by FGFR3 in disease;MAP2K and MAPK activation;SHC-related events triggered by IGF1R;Signaling by FGFR4 in disease;SHC-mediated cascade.CL.FGFR1;Tie2 Signaling;Signaling by FGFR1 in disease;Activation of RAS in B cells;SHC-mediated cascade.CL.FGFR4;Constitutive Signaling by EGFRvIII;CD209 (DC-SIGN) signaling;Ras activation uopn Ca2.PLS. infux through NMDA receptor;FCERI mediated MAPK activation;RAFMAP kinase cascade;Downstream signal transduction;VEGFR2 mediated cell proliferation;Signaling by SCF-KIT;Frs2-mediated activation;ARMS-mediated activation;Gastrin-CREB signalling pathway via PKC and MAPK;Interleukin receptor SHC signaling;DAP12 signaling;Signaling by PDGF;VEGFA-VEGFR2 Pathway;Signaling by ERBB2;IRS-related events triggered by IGF1R;Cell surface interactions at the vascular wall;CREB phosphorylation through the activation of Ras,hsa04010;hsa04012;hsa04014;hsa04015;hsa04062;hsa04068;hsa04071;hsa04072;hsa04144;hsa04151;hsa04211;hsa04213;hsa04360;hsa04370;hsa04510;hsa04530;hsa04540;hsa04550;hsa04630;hsa04650;hsa04660;hsa04662;hsa04664;hsa04720;hsa04722;hsa04725;hsa04726;hsa04730;hsa04810;hsa04910;hsa04912;hsa04915;hsa04916;hsa04917;hsa04919;hsa04921;hsa04933;hsa05034;hsa05160;hsa05161;hsa05166;hsa05200;hsa05203;hsa05205;hsa05206;hsa05211;hsa05213;hsa05214;hsa05215;hsa05216;hsa05218;hsa05219;hsa05220;hsa05221;hsa05223;hsa05230;hsa05231,HRAS,NA,HRAS - mRNA,False\nERBB2 - mRNA,NM_001005862.1:1255,SEBELIUS_2_C4636,ERBB2,mRNA,0,,,RAFMAP kinase cascade;GRB7 events in ERBB2 signaling;PLCG1 events in ERBB2 signaling;Sema4D induced cell migration and growth-cone collapse;Signaling by ERBB2;Downstream signal transduction;GAB1 signalosome;Signaling by SCF-KIT;PI3KAKT activation;Role of LAT2NTALLAB on calcium mobilization;DAP12 signaling;Signaling by PDGF;VEGFR2 mediated cell proliferation;Frs2-mediated activation;ARMS-mediated activation;Gastrin-CREB signalling pathway via PKC and MAPK;Interleukin receptor SHC signaling;FCERI mediated MAPK activation;VEGFA-VEGFR2 Pathway;IRS-related events triggered by IGF1R,hsa04012;hsa04020;hsa04066;hsa04510;hsa04520;hsa05200;hsa05205;hsa05206;hsa05212;hsa05213;hsa05215;hsa05219;hsa05223;hsa05230,ERBB2,NA,ERBB2 - mRNA,False\nPIK3CB - mRNA,NM_006219.1:2945,SEBELIUS_2_C4636,PIK3CB,mRNA,0,,,Downstream TCR signaling;PIP3 activates AKT signaling;Constitutive Signaling by Aberrant PI3K in Cancer;Tie2 Signaling;VEGFA-VEGFR2 Pathway;Synthesis of PIPs at the plasma membrane;Interleukin receptor SHC signaling;G beta.CL.gamma signalling through PI3Kgamma;Regulation of signaling by CBL;GPVI-mediated activation cascade;Role of phospholipids in phagocytosis;Nephrin interactions;PI3K Cascade;Downstream signal transduction;GAB1 signalosome;Signaling by SCF-KIT;PI3KAKT activation;Role of LAT2NTALLAB on calcium mobilization;DAP12 signaling;Signaling by PDGF;Signaling by ERBB2;Cell surface interactions at the vascular wall;IRS-related events triggered by IGF1R,hsa00562;hsa04012;hsa04014;hsa04015;hsa04022;hsa04024;hsa04062;hsa04066;hsa04068;hsa04070;hsa04071;hsa04072;hsa04150;hsa04151;hsa04152;hsa04210;hsa04211;hsa04213;hsa04261;hsa04370;hsa04380;hsa04510;hsa04550;hsa04611;hsa04620;hsa04630;hsa04650;hsa04660;hsa04662;hsa04664;hsa04666;hsa04668;hsa04670;hsa04722;hsa04725;hsa04750;hsa04810;hsa04910;hsa04914;hsa04915;hsa04917;hsa04919;hsa04921;hsa04923;hsa04930;hsa04931;hsa04932;hsa04933;hsa04960;hsa04973;hsa05100;hsa05142;hsa05145;hsa05146;hsa05160;hsa05161;hsa05162;hsa05164;hsa05166;hsa05169;hsa05200;hsa05203;hsa05205;hsa05210;hsa05211;hsa05212;hsa05213;hsa05214;hsa05215;hsa05218;hsa05220;hsa05221;hsa05222;hsa05223;hsa05230;hsa05231,PIK3CB,NA,PIK3CB - mRNA,False\nBAIAP2L1 - mRNA,NM_018842.3:259,SEBELIUS_2_C4636,BAIAP2L1,mRNA,0,,,,,BAIAP2L1,NA,BAIAP2L1 - mRNA,False\nDVL1 - mRNA,NM_004421.2:789,SEBELIUS_2_C4636,DVL1,mRNA,0,,,Disassembly of the destruction complex and recruitment of AXIN to the membrane;WNT mediated activation of DVL;RHO GTPases Activate Formins;PCPCE pathway,hsa04310;hsa04330;hsa04390;hsa04550;hsa04916;hsa05166;hsa05200;hsa05217,DVL1,NA,DVL1 - mRNA,False\nBAD - mRNA,NM_004322.3:963,SEBELIUS_2_C4636,BAD,mRNA,0,,,Activation of BAD and translocation to mitochondria ;NRAGE signals death through JNK;Constitutive Signaling by AKT1 E17K in Cancer;AKT phosphorylates targets in the cytosol;PIP3 activates AKT signaling;Downstream signal transduction;GAB1 signalosome;Signaling by SCF-KIT;PI3KAKT activation;Role of LAT2NTALLAB on calcium mobilization;DAP12 signaling;Signaling by PDGF;Signaling by ERBB2,hsa04012;hsa04014;hsa04022;hsa04024;hsa04151;hsa04210;hsa04370;hsa04510;hsa04722;hsa04910;hsa04919;hsa05010;hsa05145;hsa05152;hsa05160;hsa05161;hsa05200;hsa05203;hsa05210;hsa05212;hsa05213;hsa05215;hsa05218;hsa05220;hsa05221;hsa05223,BAD,NA,BAD - mRNA,False\nBHLHE40 - mRNA,NM_003670.1:560,SEBELIUS_2_C4636,BHLHE40,mRNA,0,,,Circadian Clock,,BHLHE40,NA,BHLHE40 - mRNA,False\nMET - mRNA,NM_001127500.1:1925,SEBELIUS_2_C4636,MET,mRNA,0,,,,hsa04014;hsa04015;hsa04060;hsa04144;hsa04151;hsa04360;hsa04510;hsa04520;hsa05100;hsa05120;hsa05200;hsa05202;hsa05205;hsa05206;hsa05211;hsa05218;hsa05230,MET,NA,MET - mRNA,False\nMYC - mRNA,NM_002467.3:1610,SEBELIUS_2_C4636,MYC,mRNA,0,,,Constitutive Signaling by NOTCH1 PEST Domain Mutants;SMAD2SMAD3.CL.SMAD4 heterotrimer regulates transcription;Formation of the beta-catenin.CL.TCF transactivating complex,hsa04010;hsa04012;hsa04110;hsa04151;hsa04310;hsa04350;hsa04390;hsa04550;hsa04630;hsa04919;hsa05161;hsa05166;hsa05169;hsa05200;hsa05202;hsa05205;hsa05206;hsa05210;hsa05213;hsa05216;hsa05219;hsa05220;hsa05221;hsa05222;hsa05230,MYC,NA,MYC - mRNA,False\nDIAPH3 - mRNA,NM_001042517.1:1832,SEBELIUS_2_C4636,DIAPH3,mRNA,0,,,RHO GTPases Activate Formins,hsa04810,DIAPH3,NA,DIAPH3 - mRNA,False\nCCND2 - mRNA,NM_001759.3:503,SEBELIUS_2_C4636,CCND2,mRNA,0,,,Cyclin D associated events in G1,hsa04068;hsa04110;hsa04115;hsa04151;hsa04310;hsa04390;hsa04510;hsa04630;hsa04917;hsa05162;hsa05166;hsa05202;hsa05203;hsa05206,CCND2,NA,CCND2 - mRNA,False\nPDCD4 - mRNA,NM_014456.3:1115,SEBELIUS_2_C4636,PDCD4,mRNA,0,,,,hsa05205;hsa05206,PDCD4,NA,PDCD4 - mRNA,False\nPIK3R3 - mRNA,NM_003629.3:1800,SEBELIUS_2_C4636,PIK3R3,mRNA,0,,,G alpha (q) signalling events;PIP3 activates AKT signaling;Constitutive Signaling by Aberrant PI3K in Cancer;CD28 dependent PI3KAkt signaling;Synthesis of PIPs at the plasma membrane;Interleukin receptor SHC signaling;G beta.CL.gamma signalling through PI3Kgamma;Regulation of signaling by CBL;GPVI-mediated activation cascade;G alpha (1213) signalling events;Gastrin-CREB signalling pathway via PKC and MAPK;Downstream signal transduction;GAB1 signalosome;Signaling by SCF-KIT;PI3KAKT activation;Role of LAT2NTALLAB on calcium mobilization;DAP12 signaling;Signaling by PDGF;Signaling by ERBB2;CD28 co-stimulation,hsa04012;hsa04014;hsa04015;hsa04022;hsa04024;hsa04062;hsa04066;hsa04068;hsa04070;hsa04071;hsa04072;hsa04150;hsa04151;hsa04152;hsa04210;hsa04211;hsa04213;hsa04261;hsa04370;hsa04380;hsa04510;hsa04550;hsa04611;hsa04620;hsa04630;hsa04650;hsa04660;hsa04662;hsa04664;hsa04666;hsa04668;hsa04670;hsa04722;hsa04725;hsa04750;hsa04810;hsa04910;hsa04914;hsa04915;hsa04917;hsa04919;hsa04921;hsa04923;hsa04930;hsa04931;hsa04932;hsa04933;hsa04960;hsa04973;hsa05100;hsa05142;hsa05145;hsa05146;hsa05160;hsa05161;hsa05162;hsa05164;hsa05166;hsa05169;hsa05200;hsa05203;hsa05205;hsa05210;hsa05211;hsa05212;hsa05213;hsa05214;hsa05215;hsa05218;hsa05220;hsa05221;hsa05222;hsa05223;hsa05230;hsa05231,PIK3R3,NA,PIK3R3 - mRNA,False\nWASL - mRNA,NM_003941.2:400,SEBELIUS_2_C4636,WASL,mRNA,0,,,EPHB-mediated forward signaling;RHO GTPases Activate WASPs and WAVEs;DCC mediated attractive signaling;Nephrin interactions;Regulation of actin dynamics for phagocytic cup formation,hsa04062;hsa04144;hsa04520;hsa04666;hsa04810;hsa05100;hsa05130;hsa05131;hsa05132;hsa05231,WASL,NA,WASL - mRNA,False\nTGFB2 - mRNA,NM_003238.2:1125,SEBELIUS_2_C4636,TGFB2,mRNA,0,,,Molecules associated with elastic fibres;ECM proteoglycans;Platelet degranulation ;Elastic fibre formation,hsa04010;hsa04060;hsa04068;hsa04110;hsa04144;hsa04350;hsa04380;hsa04390;hsa04933;hsa05140;hsa05142;hsa05145;hsa05146;hsa05152;hsa05161;hsa05166;hsa05200;hsa05205;hsa05206;hsa05210;hsa05211;hsa05212;hsa05220;hsa05321;hsa05410;hsa05414,TGFB2,NA,TGFB2 - mRNA,False\nHSPA1B - mRNA,NM_005346.3:2095,SEBELIUS_2_C4636,HSPA1B,mRNA,0,,,Regulation of HSF1-mediated heat shock response;AUF1 (hnRNP D0) binds and destabilizes mRNA,hsa04010;hsa04141;hsa04144;hsa04213;hsa04915;hsa05145;hsa05162;hsa05164;hsa05169,HSPA1B,NA,HSPA1B - mRNA,False\nVAV2 - mRNA,NM_001134398.1:1615,SEBELIUS_2_C4636,VAV2,mRNA,0,,,Rho GTPase cycle;DAP12 signaling;Signal transduction by L1;EPH-ephrin mediated repulsion of cells;VEGFR2 mediated vascular permeability;NRAGE signals death through JNK;GPVI-mediated activation cascade;G alpha (1213) signalling events;FCERI mediated MAPK activation;Regulation of actin dynamics for phagocytic cup formation;VEGFA-VEGFR2 Pathway,hsa04015;hsa04024;hsa04062;hsa04510;hsa04650;hsa04660;hsa04662;hsa04664;hsa04666;hsa04670;hsa04810;hsa05205,VAV2,NA,VAV2 - mRNA,False\nAGO2 - mRNA,NM_012154.2:625,SEBELIUS_2_C4636,AGO2,mRNA,0,,,Pre-NOTCH Transcription and Translation;PIP3 activates AKT signaling;Post-transcriptional silencing by small RNAs;MicroRNA (miRNA) biogenesis;Small interfering RNA (siRNA) biogenesis;Downstream signal transduction;GAB1 signalosome;Signaling by SCF-KIT;PI3KAKT activation;Role of LAT2NTALLAB on calcium mobilization;DAP12 signaling;Signaling by PDGF;Signaling by ERBB2,,AGO2,NA,AGO2 - mRNA,False\nMLLT4 - mRNA,NM_005936.2:1560,SEBELIUS_2_C4636,MLLT4,mRNA,0,,,Adherens junctions interactions,hsa04014;hsa04015;hsa04024;hsa04520;hsa04530;hsa04670,MLLT4,NA,MLLT4 - mRNA,False\nJUN - mRNA,NM_002228.3:140,SEBELIUS_2_C4636,JUN,mRNA,0,,,Activation of the AP-1 family of transcription factors;MAPK6MAPK4 signaling;Pre-NOTCH Transcription and Translation;Oxidative Stress Induced Senescence;FCERI mediated MAPK activation;Senescence-Associated Secretory Phenotype (SASP),hsa04010;hsa04012;hsa04024;hsa04310;hsa04380;hsa04510;hsa04620;hsa04660;hsa04662;hsa04668;hsa04722;hsa04912;hsa04915;hsa04921;hsa04932;hsa04933;hsa05031;hsa05120;hsa05132;hsa05133;hsa05140;hsa05142;hsa05161;hsa05164;hsa05166;hsa05168;hsa05169;hsa05200;hsa05203;hsa05210;hsa05211;hsa05231;hsa05321,JUN,NA,JUN - mRNA,False\nRRAS - mRNA,NM_006270.3:330,SEBELIUS_2_C4636,RRAS,mRNA,0,,,CREB phosphorylation through the activation of Ras,hsa04010;hsa04014;hsa04015;hsa04024;hsa04072;hsa04530;hsa04810;hsa05166;hsa05205,RRAS,NA,RRAS - mRNA,False\nBCAR1 - mRNA,NM_001170717.1:1341,SEBELIUS_2_C4636,BCAR1,mRNA,0,,,Downstream signal transduction;p130Cas linkage to MAPK signaling for integrins;VEGFA-VEGFR2 Pathway;Signaling by PDGF;Integrin alphaIIb beta3 signaling,hsa04015;hsa04062;hsa04510;hsa04670;hsa04810;hsa05100,BCAR1,NA,BCAR1 - mRNA,False\nPIK3R2 - mRNA,NM_005027.2:3100,SEBELIUS_2_C4636,PIK3R2,mRNA,0,,,G alpha (q) signalling events;Downstream TCR signaling;PIP3 activates AKT signaling;VEGFA-VEGFR2 Pathway;Synthesis of PIPs at the plasma membrane;Rho GTPase cycle;Interleukin receptor SHC signaling;G beta.CL.gamma signalling through PI3Kgamma;Regulation of signaling by CBL;Role of phospholipids in phagocytosis;G alpha (1213) signalling events;PI3K Cascade;Constitutive Signaling by Aberrant PI3K in Cancer;CD28 dependent PI3KAkt signaling;Tie2 Signaling;GPVI-mediated activation cascade;Nephrin interactions;Gastrin-CREB signalling pathway via PKC and MAPK;Downstream signal transduction;GAB1 signalosome;Signaling by SCF-KIT;PI3KAKT activation;Role of LAT2NTALLAB on calcium mobilization;DAP12 signaling;Signaling by PDGF;Signaling by ERBB2;IRS-related events triggered by IGF1R;CD28 co-stimulation;Cell surface interactions at the vascular wall,hsa04012;hsa04014;hsa04015;hsa04022;hsa04024;hsa04062;hsa04066;hsa04068;hsa04070;hsa04071;hsa04072;hsa04150;hsa04151;hsa04152;hsa04210;hsa04211;hsa04213;hsa04261;hsa04370;hsa04380;hsa04510;hsa04550;hsa04611;hsa04620;hsa04630;hsa04650;hsa04660;hsa04662;hsa04664;hsa04666;hsa04668;hsa04670;hsa04722;hsa04725;hsa04750;hsa04810;hsa04910;hsa04914;hsa04915;hsa04917;hsa04919;hsa04921;hsa04923;hsa04930;hsa04931;hsa04932;hsa04933;hsa04960;hsa04973;hsa05100;hsa05142;hsa05145;hsa05146;hsa05160;hsa05161;hsa05162;hsa05164;hsa05166;hsa05169;hsa05200;hsa05203;hsa05205;hsa05206;hsa05210;hsa05211;hsa05212;hsa05213;hsa05214;hsa05215;hsa05218;hsa05220;hsa05221;hsa05222;hsa05223;hsa05230;hsa05231,PIK3R2,NA,PIK3R2 - mRNA,False\nPIK3CD - mRNA,NM_005026.3:2978,SEBELIUS_2_C4636,PIK3CD,mRNA,0,,,Synthesis of PIPs at the plasma membrane;Interleukin receptor SHC signaling;G beta.CL.gamma signalling through PI3Kgamma;PIP3 activates AKT signaling;Constitutive Signaling by Aberrant PI3K in Cancer;Regulation of signaling by CBL;GPVI-mediated activation cascade;Downstream signal transduction;GAB1 signalosome;Signaling by SCF-KIT;PI3KAKT activation;Role of LAT2NTALLAB on calcium mobilization;DAP12 signaling;Signaling by PDGF;Signaling by ERBB2,hsa00562;hsa04012;hsa04014;hsa04015;hsa04022;hsa04024;hsa04062;hsa04066;hsa04068;hsa04070;hsa04071;hsa04072;hsa04150;hsa04151;hsa04152;hsa04210;hsa04211;hsa04213;hsa04261;hsa04370;hsa04380;hsa04510;hsa04550;hsa04611;hsa04620;hsa04630;hsa04650;hsa04660;hsa04662;hsa04664;hsa04666;hsa04668;hsa04670;hsa04722;hsa04725;hsa04750;hsa04810;hsa04910;hsa04914;hsa04915;hsa04917;hsa04919;hsa04921;hsa04923;hsa04930;hsa04931;hsa04932;hsa04933;hsa04960;hsa04973;hsa05100;hsa05142;hsa05145;hsa05146;hsa05160;hsa05161;hsa05162;hsa05164;hsa05166;hsa05169;hsa05200;hsa05203;hsa05205;hsa05210;hsa05211;hsa05212;hsa05213;hsa05214;hsa05215;hsa05218;hsa05220;hsa05221;hsa05222;hsa05223;hsa05230;hsa05231,PIK3CD,NA,PIK3CD - mRNA,False\nABI2 - mRNA,NM_005759.5:1160,SEBELIUS_2_C4636,ABI2,mRNA,0,,,RHO GTPases Activate WASPs and WAVEs;VEGFA-VEGFR2 Pathway;Regulation of actin dynamics for phagocytic cup formation,hsa04810,ABI2,NA,ABI2 - mRNA,False\nTCF7L2 - mRNA,NM_001146274.1:1393,SEBELIUS_2_C4636,TCF7L2,mRNA,0,,,Ca2.PLS. pathway;Deactivation of the beta-catenin transactivating complex;Repression of WNT target genes;Synthesis# secretion# and inactivation of Glucagon-like Peptide-1 (GLP-1);Formation of the beta-catenin.CL.TCF transactivating complex,hsa04310;hsa04390;hsa04520;hsa04916;hsa05200;hsa05210;hsa05213;hsa05215;hsa05216;hsa05217;hsa05221;hsa05412,TCF7L2,NA,TCF7L2 - mRNA,False\nIGF1R - mRNA,NM_000875.3:10810,SEBELIUS_2_C4636,IGF1R,mRNA,0,,,IRS-related events triggered by IGF1R;SHC-related events triggered by IGF1R,hsa04014;hsa04015;hsa04066;hsa04068;hsa04114;hsa04144;hsa04151;hsa04152;hsa04211;hsa04213;hsa04510;hsa04520;hsa04550;hsa04730;hsa04914;hsa05200;hsa05202;hsa05205;hsa05214;hsa05215;hsa05218,IGF1R,NA,IGF1R - mRNA,False\nSRC - mRNA,NM_005417.3:1410,SEBELIUS_2_C4636,SRC,mRNA,0,,,FCGR activation;Spry regulation of FGF signaling;GAB1 signalosome;MAP2K and MAPK activation;CLEC7A (Dectin-1) signaling;CD28 co-stimulation;p130Cas linkage to MAPK signaling for integrins;CTLA4 inhibitory signaling;GRB2.CL.SOS provides linkage to MAPK signaling for Integrins ;Signal regulatory protein (SIRP) family interactions;PECAM1 interactions;Thrombin signalling through proteinase activated receptors (PARs);GP1b-IX-V activation signalling;c-src mediated regulation of Cx43 function and closure of gap junctions;Signaling by ERBB2;VEGFA-VEGFR2 Pathway;Signaling by SCF-KIT;VEGFR2 mediated cell proliferation;RAFMAP kinase cascade;Downstream signal transduction;Frs2-mediated activation;ARMS-mediated activation;Gastrin-CREB signalling pathway via PKC and MAPK;Interleukin receptor SHC signaling;FCERI mediated MAPK activation;DAP12 signaling;Signaling by PDGF;IRS-related events triggered by IGF1R;Negative regulation of FGFR3 signaling;Negative regulation of FGFR4 signaling;Negative regulation of FGFR2 signaling;Negative regulation of FGFR1 signaling;Integrin alphaIIb beta3 signaling;Cell surface interactions at the vascular wall,hsa04012;hsa04015;hsa04062;hsa04144;hsa04370;hsa04510;hsa04520;hsa04530;hsa04540;hsa04611;hsa04727;hsa04750;hsa04810;hsa04912;hsa04915;hsa04917;hsa04919;hsa04921;hsa05100;hsa05120;hsa05131;hsa05152;hsa05161;hsa05203;hsa05205;hsa05219,SRC,NA,SRC - mRNA,False\nCSNK2A2 - mRNA,NM_001896.2:1350,SEBELIUS_2_C4636,CSNK2A2,mRNA,0,,,WNT mediated activation of DVL;Signal transduction by L1,hsa04064;hsa04310;hsa04520;hsa04530;hsa05162;hsa05168;hsa05169,CSNK2A2,NA,CSNK2A2 - mRNA,False\nFYN - mRNA,NM_002037.3:765,SEBELIUS_2_C4636,FYN,mRNA,0,,,EPH-ephrin mediated repulsion of cells;Regulation of signaling by CBL;FCGR activation;Antigen activates B Cell Receptor (BCR) leading to generation of second messengers;Dectin-2 family;RAFMAP kinase cascade;Constitutive Signaling by Aberrant PI3K in Cancer;CTLA4 inhibitory signaling;CD28 dependent PI3KAkt signaling;PECAM1 interactions;CD28 dependent Vav1 pathway;GPVI-mediated activation cascade;Sema3A PAK dependent Axon repulsion;Nephrin interactions;SEMA3A-Plexin repulsion signaling by inhibiting Integrin adhesion;CRMPs in Sema3A signaling;CD209 (DC-SIGN) signaling;Downstream signal transduction;GAB1 signalosome;Signaling by SCF-KIT;PI3KAKT activation;Role of LAT2NTALLAB on calcium mobilization;DAP12 signaling;Signaling by PDGF;Signaling by ERBB2;VEGFA-VEGFR2 Pathway;VEGFR2 mediated cell proliferation;Frs2-mediated activation;ARMS-mediated activation;Gastrin-CREB signalling pathway via PKC and MAPK;Interleukin receptor SHC signaling;FCERI mediated MAPK activation;IRS-related events triggered by IGF1R;CD28 co-stimulation;Cell surface interactions at the vascular wall,hsa04071;hsa04072;hsa04360;hsa04380;hsa04510;hsa04520;hsa04611;hsa04650;hsa04660;hsa04664;hsa04725;hsa05020;hsa05130;hsa05162;hsa05416,FYN,NA,FYN - mRNA,False\nMAPK10 - mRNA,NM_138981.2:450,SEBELIUS_2_C4636,MAPK10,mRNA,0,,,Activation of the AP-1 family of transcription factors;Oxidative Stress Induced Senescence;FCERI mediated MAPK activation,hsa04010;hsa04012;hsa04014;hsa04024;hsa04068;hsa04071;hsa04141;hsa04310;hsa04380;hsa04510;hsa04620;hsa04621;hsa04664;hsa04668;hsa04722;hsa04723;hsa04728;hsa04750;hsa04910;hsa04912;hsa04914;hsa04917;hsa04920;hsa04930;hsa04931;hsa04932;hsa04933;hsa05120;hsa05131;hsa05132;hsa05133;hsa05142;hsa05145;hsa05152;hsa05160;hsa05161;hsa05164;hsa05168;hsa05169;hsa05200;hsa05210;hsa05212;hsa05231,MAPK10,NA,MAPK10 - mRNA,False\nMRAS - mRNA,NM_012219.3:4210,SEBELIUS_2_C4636,MRAS,mRNA,0,,,,hsa04010;hsa04014;hsa04015;hsa04072;hsa04530;hsa04810;hsa05166;hsa05205,MRAS,NA,MRAS - mRNA,False\nASH1L - mRNA,NM_018489.2:5005,SEBELIUS_2_C4636,ASH1L,mRNA,0,,,,hsa04530,ASH1L,NA,ASH1L - mRNA,False\nMAPK7 - mRNA,NM_139033.1:2850,SEBELIUS_2_C4636,MAPK7,mRNA,0,,,ERKs are inactivated;Gastrin-CREB signalling pathway via PKC and MAPK;Senescence-Associated Secretory Phenotype (SASP),hsa04010;hsa04540;hsa04722;hsa04912;hsa04921;hsa05206,MAPK7,NA,MAPK7 - mRNA,False\nBRAF - mRNA,NM_004333.3:565,SEBELIUS_2_C4636,BRAF,mRNA,0,,,Spry regulation of FGF signaling;MAP2K and MAPK activation;RAF activation;CREB phosphorylation through the activation of Ras;Negative feedback regulation of MAPK pathway;ARMS-mediated activation;Negative regulation of FGFR3 signaling;Negative regulation of FGFR4 signaling;Negative regulation of FGFR2 signaling;Negative regulation of FGFR1 signaling;RAFMAP kinase cascade;Downstream signal transduction;VEGFR2 mediated cell proliferation;Signaling by SCF-KIT;Frs2-mediated activation;Gastrin-CREB signalling pathway via PKC and MAPK;Interleukin receptor SHC signaling;FCERI mediated MAPK activation;DAP12 signaling;Signaling by PDGF;VEGFA-VEGFR2 Pathway;Signaling by ERBB2;IRS-related events triggered by IGF1R;Negative regulation of MAPK pathway,hsa04010;hsa04012;hsa04015;hsa04024;hsa04062;hsa04068;hsa04150;hsa04270;hsa04510;hsa04650;hsa04720;hsa04722;hsa04726;hsa04730;hsa04810;hsa04910;hsa04914;hsa05034;hsa05160;hsa05200;hsa05205;hsa05210;hsa05211;hsa05212;hsa05213;hsa05214;hsa05215;hsa05216;hsa05218;hsa05219;hsa05220;hsa05221;hsa05223,BRAF,NA,BRAF - mRNA,False\nPPP2R2D - mRNA,NM_001291310.1:2320,SEBELIUS_2_C4636,PPP2R2D,mRNA,0,,,MASTL Facilitates Mitotic Progression,hsa03015;hsa04071;hsa04151;hsa04152;hsa04261;hsa04390;hsa04530;hsa04728;hsa05142;hsa05160,PPP2R2D,NA,PPP2R2D - mRNA,False\nNUMA1 - mRNA,NM_006185.2:575,SEBELIUS_2_C4636,NUMA1,mRNA,0,,,,,NUMA1,NA,NUMA1 - mRNA,False\nTBP - mRNA,NM_001172085.1:587,SEBELIUS_2_C4636,TBP,mRNA,0,,,,hsa05016;hsa05166;hsa05168;hsa05169;hsa05203,TBP,NA,TBP - mRNA,False\nSMAD2 - mRNA,NM_005901.5:1678,SEBELIUS_2_C4636,SMAD2,mRNA,0,,,Transcriptional regulation of pluripotent stem cells;TGF-beta receptor signaling activates SMADs,hsa04068;hsa04110;hsa04144;hsa04350;hsa04390;hsa04520;hsa04550;hsa04933;hsa05142;hsa05166;hsa05200;hsa05205;hsa05210;hsa05212;hsa05321,SMAD2,NA,SMAD2 - mRNA,False\nDICER1 - mRNA,NM_177438.2:5060,SEBELIUS_2_C4636,DICER1,mRNA,0,,,MicroRNA (miRNA) biogenesis,hsa05206,DICER1,NA,DICER1 - mRNA,False\nLIMK1 - mRNA,NM_002314.3:3133,SEBELIUS_2_C4636,LIMK1,mRNA,0,,,EPHB-mediated forward signaling;RHO GTPases Activate ROCKs;RHO GTPases activate PAKs;Sema3A PAK dependent Axon repulsion;Regulation of actin dynamics for phagocytic cup formation;Sema4D induced cell migration and growth-cone collapse,hsa04360;hsa04666;hsa04810,LIMK1,NA,LIMK1 - mRNA,False\nPARD3 - mRNA,NM_019619.2:1060,SEBELIUS_2_C4636,PARD3,mRNA,0,,,TGF-beta receptor signaling in EMT (epithelial to mesenchymal transition);Tight junction interactions,hsa04015;hsa04062;hsa04144;hsa04390;hsa04520;hsa04530,PARD3,NA,PARD3 - mRNA,False\nCREBBP - mRNA,NM_004380.2:1301,SEBELIUS_2_C4636,CREBBP,mRNA,0,,,YAP1- and WWTR1 (TAZ)-stimulated gene expression;Activation of gene expression by SREBF (SREBP);Transcriptional activation of mitochondrial biogenesis;Constitutive Signaling by NOTCH1 HD.PLS.PEST Domain Mutants;Constitutive Signaling by NOTCH1 PEST Domain Mutants;CD209 (DC-SIGN) signaling;Circadian Clock,hsa04024;hsa04066;hsa04068;hsa04110;hsa04310;hsa04330;hsa04350;hsa04520;hsa04630;hsa04720;hsa04916;hsa04919;hsa04922;hsa05016;hsa05152;hsa05161;hsa05164;hsa05166;hsa05168;hsa05169;hsa05200;hsa05203;hsa05206;hsa05211;hsa05215,CREBBP,NA,CREBBP - mRNA,False\nPRKCA - mRNA,NM_002737.2:680,SEBELIUS_2_C4636,PRKCA,mRNA,0,,,WNT5A-dependent internalization of FZD4;Disinhibition of SNARE formation;Trafficking of AMPA receptors;Downstream signal transduction;EGFR interacts with phospholipase C-gamma;Phospholipase C-mediated cascade.CL. FGFR1;PLCG1 events in ERBB2 signaling;DAP12 signaling;PLC beta mediated events;Signaling by PDGF;Signaling by ERBB2;Signaling by SCF-KIT;Non-integrin membrane-ECM interactions;Gastrin-CREB signalling pathway via PKC and MAPK;VEGFA-VEGFR2 Pathway;PCPCE pathway,hsa04010;hsa04012;hsa04014;hsa04015;hsa04020;hsa04066;hsa04070;hsa04071;hsa04072;hsa04150;hsa04151;hsa04261;hsa04270;hsa04310;hsa04370;hsa04510;hsa04530;hsa04540;hsa04650;hsa04664;hsa04666;hsa04670;hsa04713;hsa04720;hsa04723;hsa04724;hsa04725;hsa04726;hsa04727;hsa04728;hsa04730;hsa04750;hsa04912;hsa04916;hsa04919;hsa04921;hsa04933;hsa04960;hsa04971;hsa04972;hsa05031;hsa05032;hsa05110;hsa05130;hsa05146;hsa05161;hsa05164;hsa05200;hsa05205;hsa05206;hsa05214;hsa05223;hsa05231,PRKCA,NA,PRKCA - mRNA,False\nRAB8B - mRNA,NM_016530.2:820,SEBELIUS_2_C4636,RAB8B,mRNA,0,,,,,RAB8B,NA,RAB8B - mRNA,False\nCTBP1 - mRNA,NM_001328.2:425,SEBELIUS_2_C4636,CTBP1,mRNA,0,,,Deactivation of the beta-catenin transactivating complex;Repression of WNT target genes,hsa04310;hsa04330;hsa05200;hsa05220,CTBP1,NA,CTBP1 - mRNA,False\nMAPK3 - mRNA,NM_001040056.1:580,SEBELIUS_2_C4636,MAPK3,mRNA,0,,,ERKs are inactivated;ISG15 antiviral mechanism;RHO GTPases Activate WASPs and WAVEs;Senescence-Associated Secretory Phenotype (SASP);Regulation of HSF1-mediated heat shock response;Thrombin signalling through proteinase activated receptors (PARs);Negative regulation of FGFR4 signaling;Oxidative Stress Induced Senescence;Signal attenuation;RNA Polymerase I Promoter Opening;Negative regulation of FGFR3 signaling;Negative feedback regulation of MAPK pathway;Growth hormone receptor signaling;Golgi Cisternae Pericentriolar Stack Reorganization;FCERI mediated MAPK activation;Negative regulation of MAPK pathway;RAFMAP kinase cascade;Downstream signal transduction;VEGFR2 mediated cell proliferation;Signaling by SCF-KIT;Frs2-mediated activation;ARMS-mediated activation;Gastrin-CREB signalling pathway via PKC and MAPK;Interleukin receptor SHC signaling;DAP12 signaling;Signaling by PDGF;VEGFA-VEGFR2 Pathway;Signaling by ERBB2;IRS-related events triggered by IGF1R,hsa04010;hsa04012;hsa04014;hsa04015;hsa04022;hsa04024;hsa04062;hsa04066;hsa04068;hsa04071;hsa04072;hsa04114;hsa04150;hsa04151;hsa04261;hsa04270;hsa04320;hsa04350;hsa04360;hsa04370;hsa04380;hsa04510;hsa04520;hsa04540;hsa04550;hsa04611;hsa04620;hsa04621;hsa04650;hsa04660;hsa04662;hsa04664;hsa04666;hsa04668;hsa04713;hsa04720;hsa04722;hsa04723;hsa04724;hsa04725;hsa04726;hsa04730;hsa04810;hsa04910;hsa04912;hsa04914;hsa04915;hsa04916;hsa04917;hsa04919;hsa04921;hsa04930;hsa04933;hsa04960;hsa05010;hsa05020;hsa05034;hsa05131;hsa05132;hsa05133;hsa05140;hsa05142;hsa05145;hsa05152;hsa05160;hsa05161;hsa05164;hsa05200;hsa05203;hsa05205;hsa05210;hsa05211;hsa05212;hsa05213;hsa05214;hsa05215;hsa05216;hsa05218;hsa05219;hsa05220;hsa05221;hsa05223;hsa05230;hsa05231,MAPK3,NA,MAPK3 - mRNA,False\nPPP2R1B - mRNA,NM_002716.4:1445,SEBELIUS_2_C4636,PPP2R1B,mRNA,0,,,MASTL Facilitates Mitotic Progression;Misspliced GSK3beta mutants stabilize beta-catenin;T41 mutants of beta-catenin aren.SQT.t phosphorylated;ERKs are inactivated;Glycolysis;Negative regulation of MAPK pathway;Disassembly of the destruction complex and recruitment of AXIN to the membrane;S37 mutants of beta-catenin aren.SQT.t phosphorylated;Cyclin AB1 associated events during G2M transition;RAF activation;RHO GTPases Activate Formins;S33 mutants of beta-catenin aren.SQT.t phosphorylated;DARPP-32 events;Beta-catenin phosphorylation cascade;Truncations of AMER1 destabilize the destruction complex;CTLA4 inhibitory signaling;AXIN missense mutants destabilize the destruction complex;S45 mutants of beta-catenin aren.SQT.t phosphorylated;Cyclin D associated events in G1;Platelet sensitization by LDL;Resolution of Sister Chromatid Cohesion;Inhibition of replication initiation of damaged DNA by RB1E2F1;RAFMAP kinase cascade;Downstream signal transduction;VEGFR2 mediated cell proliferation;Signaling by SCF-KIT;Frs2-mediated activation;ARMS-mediated activation;Gastrin-CREB signalling pathway via PKC and MAPK;Interleukin receptor SHC signaling;FCERI mediated MAPK activation;DAP12 signaling;Signaling by PDGF;VEGFA-VEGFR2 Pathway;Signaling by ERBB2;IRS-related events triggered by IGF1R,hsa03015;hsa04071;hsa04114;hsa04151;hsa04152;hsa04261;hsa04350;hsa04390;hsa04530;hsa04728;hsa04730;hsa05142;hsa05160,PPP2R1B,NA,PPP2R1B - mRNA,False\nMAPK9 - mRNA,XM_005265940.1:163,SEBELIUS_2_C4636,MAPK9,mRNA,0,,,,hsa04010;hsa04012;hsa04014;hsa04024;hsa04068;hsa04071;hsa04141;hsa04310;hsa04380;hsa04510;hsa04620;hsa04621;hsa04660;hsa04664;hsa04668;hsa04722;hsa04723;hsa04728;hsa04750;hsa04910;hsa04912;hsa04914;hsa04917;hsa04920;hsa04930;hsa04931;hsa04932;hsa04933;hsa05120;hsa05131;hsa05132;hsa05133;hsa05142;hsa05145;hsa05152;hsa05160;hsa05161;hsa05164;hsa05168;hsa05169;hsa05200;hsa05210;hsa05212;hsa05231,MAPK9,NA,MAPK9 - mRNA,False\nPRKCI - mRNA,NM_002740.4:530,SEBELIUS_2_C4636,PRKCI,mRNA,0,,,Tight junction interactions,hsa04015;hsa04144;hsa04390;hsa04530;hsa04611;hsa04910,PRKCI,NA,PRKCI - mRNA,False\nEGFR - mRNA,NM_201282.1:360,SEBELIUS_2_C4636,EGFR,mRNA,0,,,Constitutive Signaling by Aberrant PI3K in Cancer;Inhibition of Signaling by Overexpressed EGFR;PLCG1 events in ERBB2 signaling;EGFR interacts with phospholipase C-gamma;EGFR Transactivation by Gastrin;Constitutive Signaling by Ligand-Responsive EGFR Cancer Variants;Constitutive Signaling by EGFRvIII;GAB1 signalosome;Downstream signal transduction;Signaling by SCF-KIT;PI3KAKT activation;Role of LAT2NTALLAB on calcium mobilization;DAP12 signaling;Signaling by PDGF;Signaling by ERBB2;VEGFR2 mediated cell proliferation;Frs2-mediated activation;ARMS-mediated activation;Gastrin-CREB signalling pathway via PKC and MAPK;Interleukin receptor SHC signaling;FCERI mediated MAPK activation;VEGFA-VEGFR2 Pathway;IRS-related events triggered by IGF1R,hsa04010;hsa04012;hsa04014;hsa04015;hsa04020;hsa04060;hsa04066;hsa04068;hsa04072;hsa04144;hsa04151;hsa04320;hsa04510;hsa04520;hsa04540;hsa04810;hsa04912;hsa04915;hsa04921;hsa05120;hsa05160;hsa05200;hsa05205;hsa05206;hsa05212;hsa05213;hsa05214;hsa05215;hsa05218;hsa05219;hsa05223;hsa05230;hsa05231,EGFR,NA,EGFR - mRNA,False\nAKT3 - mRNA,NM_005465.4:287,SEBELIUS_2_C4636,AKT3,mRNA,0,,,TP53 Regulates Metabolic Genes;CD28 dependent PI3KAkt signaling;AKT phosphorylates targets in the cytosol;Negative regulation of the PI3KAKT network;Activation of BAD and translocation to mitochondria ;G beta.CL.gamma signalling through PI3Kgamma;AKT phosphorylates targets in the nucleus;VEGFR2 mediated vascular permeability;GPVI-mediated activation cascade;Signaling by ERBB2;PIP3 activates AKT signaling;Downstream signal transduction;GAB1 signalosome;Signaling by SCF-KIT;PI3KAKT activation;Role of LAT2NTALLAB on calcium mobilization;DAP12 signaling;Signaling by PDGF;CD28 co-stimulation;VEGFA-VEGFR2 Pathway,hsa04010;hsa04012;hsa04014;hsa04015;hsa04022;hsa04024;hsa04062;hsa04066;hsa04068;hsa04071;hsa04072;hsa04150;hsa04151;hsa04152;hsa04210;hsa04211;hsa04213;hsa04261;hsa04370;hsa04380;hsa04510;hsa04530;hsa04550;hsa04611;hsa04620;hsa04630;hsa04660;hsa04662;hsa04664;hsa04666;hsa04668;hsa04722;hsa04725;hsa04728;hsa04910;hsa04914;hsa04915;hsa04917;hsa04919;hsa04920;hsa04922;hsa04923;hsa04931;hsa04932;hsa04933;hsa04973;hsa05142;hsa05145;hsa05152;hsa05160;hsa05161;hsa05162;hsa05164;hsa05166;hsa05169;hsa05200;hsa05205;hsa05210;hsa05211;hsa05212;hsa05213;hsa05214;hsa05215;hsa05218;hsa05220;hsa05221;hsa05222;hsa05223;hsa05230;hsa05231,AKT3,NA,AKT3 - mRNA,False\nPAK2 - mRNA,NM_002577.3:3785,SEBELIUS_2_C4636,PAK2,mRNA,0,,,Nef and signal transduction;CD28 dependent Vav1 pathway;Stimulation of the cell death response by PAK-2p34;Ephrin signaling;Activation of Rac;RHO GTPases activate PAKs;VEGFR2 mediated vascular permeability;Sema3A PAK dependent Axon repulsion;CD209 (DC-SIGN) signaling;VEGFA-VEGFR2 Pathway;CD28 co-stimulation,hsa04010;hsa04012;hsa04014;hsa04360;hsa04510;hsa04660;hsa04810;hsa05211,PAK2,NA,PAK2 - mRNA,False\nDOCK1 - mRNA,NM_001380.3:840,SEBELIUS_2_C4636,DOCK1,mRNA,0,,,DCC mediated attractive signaling;Factors involved in megakaryocyte development and platelet production;VEGFA-VEGFR2 Pathway;Regulation of actin dynamics for phagocytic cup formation,hsa04510;hsa04810;hsa05100;hsa05131,DOCK1,NA,DOCK1 - mRNA,False\nVAPA - mRNA,NM_194434.2:763,SEBELIUS_2_C4636,VAPA,mRNA,0,,,,hsa04530,VAPA,NA,VAPA - mRNA,False\nJAM3 - mRNA,NM_032801.3:890,SEBELIUS_2_C4636,JAM3,mRNA,0,,,Integrin cell surface interactions;Cell surface interactions at the vascular wall,hsa04514;hsa04530;hsa04670;hsa05120,JAM3,NA,JAM3 - mRNA,False\nZEB1 - mRNA,NM_001128128.1:1450,SEBELIUS_2_C4636,ZEB1,mRNA,0,,,,hsa05202;hsa05206,ZEB1,NA,ZEB1 - mRNA,False\nITGAV - mRNA,NM_002210.2:2615,SEBELIUS_2_C4636,ITGAV,mRNA,0,,,Molecules associated with elastic fibres;Syndecan interactions;Signal transduction by L1;Laminin interactions;ECM proteoglycans;Integrin cell surface interactions;VEGFA-VEGFR2 Pathway;Elastic fibre formation;Non-integrin membrane-ECM interactions;Cell surface interactions at the vascular wall,hsa04145;hsa04151;hsa04510;hsa04512;hsa04514;hsa04810;hsa04919;hsa05200;hsa05205;hsa05222;hsa05410;hsa05412;hsa05414,ITGAV,NA,ITGAV - mRNA,False\nRRAS2 - mRNA,NM_001102669.2:1785,SEBELIUS_2_C4636,RRAS2,mRNA,0,,,,hsa04010;hsa04014;hsa04024;hsa04072;hsa04530;hsa04810;hsa05166;hsa05205,RRAS2,NA,RRAS2 - mRNA,False\nNRAS - mRNA,NM_002524.3:877,SEBELIUS_2_C4636,NRAS,mRNA,0,,,Signaling by FGFR2 in disease;Regulation of RAS by GAPs;SHC-mediated cascade.CL.FGFR3;Signaling by FGFR3 in disease;MAP2K and MAPK activation;SHC-related events triggered by IGF1R;Signaling by FGFR4 in disease;SHC-mediated cascade.CL.FGFR1;Tie2 Signaling;Signaling by FGFR1 in disease;Activation of RAS in B cells;SHC-mediated cascade.CL.FGFR4;Constitutive Signaling by EGFRvIII;CD209 (DC-SIGN) signaling;FCERI mediated MAPK activation;RAFMAP kinase cascade;Downstream signal transduction;VEGFR2 mediated cell proliferation;Signaling by SCF-KIT;Frs2-mediated activation;ARMS-mediated activation;Gastrin-CREB signalling pathway via PKC and MAPK;Interleukin receptor SHC signaling;DAP12 signaling;Signaling by PDGF;VEGFA-VEGFR2 Pathway;Signaling by ERBB2;IRS-related events triggered by IGF1R;Cell surface interactions at the vascular wall,hsa04010;hsa04012;hsa04014;hsa04015;hsa04062;hsa04068;hsa04071;hsa04072;hsa04151;hsa04211;hsa04213;hsa04360;hsa04370;hsa04530;hsa04540;hsa04550;hsa04650;hsa04660;hsa04662;hsa04664;hsa04720;hsa04722;hsa04725;hsa04726;hsa04730;hsa04810;hsa04910;hsa04912;hsa04915;hsa04916;hsa04917;hsa04919;hsa04921;hsa04933;hsa05034;hsa05160;hsa05161;hsa05166;hsa05200;hsa05203;hsa05205;hsa05206;hsa05211;hsa05213;hsa05214;hsa05215;hsa05216;hsa05218;hsa05219;hsa05220;hsa05221;hsa05223;hsa05230;hsa05231,NRAS,NA,NRAS - mRNA,False\nMAP2K1 - mRNA,NM_002755.2:970,SEBELIUS_2_C4636,MAP2K1,mRNA,0,,,RAF activation;Uptake and function of anthrax toxins;Negative feedback regulation of MAPK pathway;Interleukin-1 signaling;RAFMAP kinase cascade;Downstream signal transduction;VEGFR2 mediated cell proliferation;Signaling by SCF-KIT;Frs2-mediated activation;ARMS-mediated activation;Gastrin-CREB signalling pathway via PKC and MAPK;Interleukin receptor SHC signaling;FCERI mediated MAPK activation;DAP12 signaling;Signaling by PDGF;VEGFA-VEGFR2 Pathway;Signaling by ERBB2;IRS-related events triggered by IGF1R;Negative regulation of MAPK pathway,hsa04010;hsa04012;hsa04014;hsa04015;hsa04022;hsa04024;hsa04062;hsa04066;hsa04068;hsa04071;hsa04072;hsa04114;hsa04151;hsa04270;hsa04320;hsa04370;hsa04380;hsa04510;hsa04540;hsa04550;hsa04620;hsa04650;hsa04660;hsa04662;hsa04664;hsa04666;hsa04668;hsa04720;hsa04722;hsa04725;hsa04726;hsa04730;hsa04810;hsa04910;hsa04912;hsa04914;hsa04915;hsa04916;hsa04917;hsa04919;hsa04921;hsa05020;hsa05034;hsa05161;hsa05164;hsa05200;hsa05205;hsa05206;hsa05210;hsa05211;hsa05212;hsa05213;hsa05214;hsa05215;hsa05216;hsa05218;hsa05219;hsa05220;hsa05221;hsa05223;hsa05230;hsa05231,MAP2K1,NA,MAP2K1 - mRNA,False\nROCK1 - mRNA,NM_005406.1:2660,SEBELIUS_2_C4636,ROCK1,mRNA,0,,,EPHB-mediated forward signaling;EPHA-mediated growth cone collapse;RHO GTPases Activate ROCKs;Apoptotic cleavage of cellular proteins;G alpha (1213) signalling events;VEGFA-VEGFR2 Pathway;Sema4D induced cell migration and growth-cone collapse,hsa04022;hsa04024;hsa04062;hsa04071;hsa04270;hsa04350;hsa04360;hsa04510;hsa04611;hsa04670;hsa04810;hsa04921;hsa05130;hsa05131;hsa05132;hsa05200;hsa05205;hsa05206,ROCK1,NA,ROCK1 - mRNA,False\nPPP1R12A - mRNA,NM_002480.2:2935,SEBELIUS_2_C4636,PPP1R12A,mRNA,0,,,RHO GTPases Activate ROCKs;RHO GTPases activate PAKs;RHO GTPases activate PKNs,hsa04022;hsa04024;hsa04270;hsa04510;hsa04611;hsa04810;hsa04921;hsa05205,PPP1R12A,NA,PPP1R12A - mRNA,False\nPDGFD - mRNA,NM_025208.4:1120,SEBELIUS_2_C4636,PDGFD,mRNA,0,,,Signaling by PDGF,hsa04014;hsa04015;hsa04072;hsa04151;hsa04510;hsa04540;hsa04810;hsa05215;hsa05218;hsa05231,PDGFD,NA,PDGFD - mRNA,False\nIL1R1 - mRNA,NM_000877.3:582,SEBELIUS_2_C4636,IL1R1,mRNA,0,,,,hsa04010;hsa04060;hsa04064;hsa04210;hsa04380;hsa04640;hsa04750;hsa05146;hsa05166,IL1R1,NA,IL1R1 - mRNA,False\nYBX3 - mRNA,NM_003651.3:660,SEBELIUS_2_C4636,YBX3,mRNA,0,,,,hsa04530,YBX3,NA,YBX3 - mRNA,False\nMYH10 - mRNA,NM_005964.1:7092,SEBELIUS_2_C4636,MYH10,mRNA,0,,,EPHA-mediated growth cone collapse;RHO GTPases Activate ROCKs;RHO GTPases activate CIT;RHO GTPases activate PAKs;RHO GTPases activate PKNs;Sema4D induced cell migration and growth-cone collapse,hsa04530;hsa04810;hsa05132,MYH10,NA,MYH10 - mRNA,False\nSMAD4 - mRNA,NM_005359.3:1370,SEBELIUS_2_C4636,SMAD4,mRNA,0,,,Signaling by NODAL,hsa04068;hsa04110;hsa04310;hsa04350;hsa04390;hsa04520;hsa04550;hsa04933;hsa05161;hsa05166;hsa05200;hsa05210;hsa05212;hsa05220,SMAD4,NA,SMAD4 - mRNA,False\nCRKL - mRNA,NM_005207.3:1372,SEBELIUS_2_C4636,CRKL,mRNA,0,,,Downstream signal transduction;Regulation of signaling by CBL;Frs2-mediated activation;Signaling by PDGF,hsa04010;hsa04012;hsa04015;hsa04062;hsa04510;hsa04666;hsa04722;hsa04810;hsa04910;hsa05100;hsa05131;hsa05200;hsa05206;hsa05211;hsa05220,CRKL,NA,CRKL - mRNA,False\nPOLR2A - mRNA,NM_000937.2:3775,SEBELIUS_2_C4636,POLR2A,mRNA,0,,,RNA Polymerase II Transcription Initiation,hsa01100;hsa05016;hsa05168;hsa05169,POLR2A,NA,POLR2A - mRNA,False\nPIK3CA - mRNA,NM_006218.2:2445,SEBELIUS_2_C4636,PIK3CA,mRNA,0,,,G alpha (q) signalling events;Downstream TCR signaling;Signaling by FGFR2 in disease;PIP3 activates AKT signaling;VEGFA-VEGFR2 Pathway;Synthesis of PIPs at the plasma membrane;Constitutive Signaling by Ligand-Responsive EGFR Cancer Variants;Interleukin receptor SHC signaling;G beta.CL.gamma signalling through PI3Kgamma;Regulation of signaling by CBL;Role of phospholipids in phagocytosis;G alpha (1213) signalling events;PI3K Cascade;Signaling by FGFR3 in disease;Constitutive Signaling by Aberrant PI3K in Cancer;Signaling by FGFR4 in disease;CD28 dependent PI3KAkt signaling;Tie2 Signaling;GPVI-mediated activation cascade;Nephrin interactions;Constitutive Signaling by EGFRvIII;Gastrin-CREB signalling pathway via PKC and MAPK;Downstream signal transduction;GAB1 signalosome;Signaling by SCF-KIT;PI3KAKT activation;Role of LAT2NTALLAB on calcium mobilization;DAP12 signaling;Signaling by PDGF;Signaling by ERBB2;IRS-related events triggered by IGF1R;CD28 co-stimulation;Cell surface interactions at the vascular wall;Signaling by FGFR1 in disease,hsa00562;hsa04012;hsa04014;hsa04015;hsa04022;hsa04024;hsa04062;hsa04066;hsa04068;hsa04070;hsa04071;hsa04072;hsa04150;hsa04151;hsa04152;hsa04210;hsa04211;hsa04213;hsa04261;hsa04370;hsa04380;hsa04510;hsa04550;hsa04611;hsa04620;hsa04630;hsa04650;hsa04660;hsa04662;hsa04664;hsa04666;hsa04668;hsa04670;hsa04722;hsa04725;hsa04750;hsa04810;hsa04910;hsa04914;hsa04915;hsa04917;hsa04919;hsa04921;hsa04923;hsa04930;hsa04931;hsa04932;hsa04933;hsa04960;hsa04973;hsa05100;hsa05142;hsa05145;hsa05146;hsa05160;hsa05161;hsa05162;hsa05164;hsa05166;hsa05169;hsa05200;hsa05203;hsa05205;hsa05206;hsa05210;hsa05211;hsa05212;hsa05213;hsa05214;hsa05215;hsa05218;hsa05220;hsa05221;hsa05222;hsa05223;hsa05230;hsa05231,PIK3CA,NA,PIK3CA - mRNA,False\nTP53 - mRNA,NM_000546.2:1330,SEBELIUS_2_C4636,TP53,mRNA,0,,,Activation of PUMA and translocation to mitochondria;Recruitment and ATM-mediated phosphorylation of repair and signaling proteins at DNA double strand breaks;Factors involved in megakaryocyte development and platelet production;Oncogene Induced Senescence,hsa04010;hsa04071;hsa04110;hsa04115;hsa04151;hsa04210;hsa04211;hsa04310;hsa04722;hsa04919;hsa05016;hsa05160;hsa05161;hsa05162;hsa05166;hsa05168;hsa05169;hsa05200;hsa05202;hsa05203;hsa05205;hsa05206;hsa05210;hsa05212;hsa05213;hsa05214;hsa05215;hsa05216;hsa05217;hsa05218;hsa05219;hsa05220;hsa05222;hsa05223;hsa05230,TP53,NA,TP53 - mRNA,False\nCTTN - mRNA,NM_001184740.1:240,SEBELIUS_2_C4636,CTTN,mRNA,0,,,RHO GTPases activate PAKs,hsa04530;hsa05100;hsa05130;hsa05131;hsa05205,CTTN,NA,CTTN - mRNA,False\nPAK1 - mRNA,NM_002576.3:975,SEBELIUS_2_C4636,PAK1,mRNA,0,,,EPHB-mediated forward signaling;Ephrin signaling;MAPK6MAPK4 signaling;DSCAM interactions;Activation of Rac;VEGFR2 mediated vascular permeability;RHO GTPases activate PAKs;Sema3A PAK dependent Axon repulsion;FCERI mediated MAPK activation;Regulation of actin dynamics for phagocytic cup formation;CD209 (DC-SIGN) signaling;CD28 dependent Vav1 pathway;VEGFA-VEGFR2 Pathway;CD28 co-stimulation,hsa04010;hsa04012;hsa04014;hsa04024;hsa04062;hsa04360;hsa04510;hsa04650;hsa04660;hsa04666;hsa04810;hsa05120;hsa05205;hsa05211,PAK1,NA,PAK1 - mRNA,False\nHPRT1 - mRNA,NM_000194.1:240,SEBELIUS_2_C4636,HPRT1,mRNA,0,,,,hsa01100,HPRT1,NA,HPRT1 - mRNA,False\nSPRY2 - mRNA,NM_005842.2:1481,SEBELIUS_2_C4636,SPRY2,mRNA,0,,,Spry regulation of FGF signaling;Negative regulation of FGFR3 signaling;Negative regulation of FGFR4 signaling;Negative regulation of FGFR2 signaling;Negative regulation of FGFR1 signaling,hsa05206,SPRY2,NA,SPRY2 - mRNA,False\nPXN - mRNA,NM_001080855.1:2495,SEBELIUS_2_C4636,PXN,mRNA,0,,,Smooth Muscle Contraction;GAB1 signalosome;VEGFA-VEGFR2 Pathway;Cell-extracellular matrix interactions,hsa04062;hsa04370;hsa04510;hsa04670;hsa04810;hsa05100;hsa05203;hsa05205,PXN,NA,PXN - mRNA,False\nITGA3 - mRNA,NM_002204.2:4550,SEBELIUS_2_C4636,ITGA3,mRNA,0,,,Laminin interactions;Integrin cell surface interactions;Cell surface interactions at the vascular wall,hsa04151;hsa04510;hsa04512;hsa04640;hsa04810;hsa05200;hsa05222;hsa05410;hsa05412;hsa05414,ITGA3,NA,ITGA3 - mRNA,False\nCASK - mRNA,NM_001126054.2:2190,SEBELIUS_2_C4636,CASK,mRNA,0,,,Syndecan interactions;Nephrin interactions;Non-integrin membrane-ECM interactions,hsa04530,CASK,NA,CASK - mRNA,False\nYES1 - mRNA,NM_005433.3:265,SEBELIUS_2_C4636,YES1,mRNA,0,,,Signaling by ERBB2;EPHB-mediated forward signaling;EPHA-mediated growth cone collapse;Regulation of KIT signaling;EPH-ephrin mediated repulsion of cells;Regulation of signaling by CBL;CD28 co-stimulation;CTLA4 inhibitory signaling;Signaling by SCF-KIT;Cell surface interactions at the vascular wall,hsa04520;hsa04530,YES1,NA,YES1 - mRNA,False\nCRK - mRNA,NM_016823.2:1945,SEBELIUS_2_C4636,CRK,mRNA,0,,,ARMS-mediated activation;Downstream signal transduction;Regulation of signaling by CBL;p130Cas linkage to MAPK signaling for integrins;VEGFA-VEGFR2 Pathway;Regulation of actin dynamics for phagocytic cup formation;Signaling by PDGF;Integrin alphaIIb beta3 signaling,hsa04010;hsa04012;hsa04015;hsa04062;hsa04510;hsa04666;hsa04722;hsa04810;hsa04910;hsa05100;hsa05131;hsa05200;hsa05206;hsa05211;hsa05220,CRK,NA,CRK - mRNA,False\nPPP2R2A - mRNA,NM_001177591.1:430,SEBELIUS_2_C4636,PPP2R2A,mRNA,0,,,Nonsense Mediated Decay (NMD) enhanced by the Exon Junction Complex (EJC);Cyclin D associated events in G1;Cyclin AB1 associated events during G2M transition,hsa03015;hsa04071;hsa04151;hsa04152;hsa04261;hsa04390;hsa04530;hsa04728;hsa05142;hsa05160,PPP2R2A,NA,PPP2R2A - mRNA,False\nGNAI3 - mRNA,NM_006496.1:1150,SEBELIUS_2_C4636,GNAI3,mRNA,0,,,G alpha (i) signalling events;ADP signalling through P2Y purinoceptor 12,hsa04015;hsa04022;hsa04024;hsa04062;hsa04071;hsa04261;hsa04360;hsa04530;hsa04540;hsa04611;hsa04670;hsa04713;hsa04723;hsa04724;hsa04725;hsa04726;hsa04727;hsa04728;hsa04730;hsa04914;hsa04915;hsa04916;hsa04921;hsa04923;hsa04971;hsa05032;hsa05034;hsa05133;hsa05142;hsa05145;hsa05200,GNAI3,NA,GNAI3 - mRNA,False\nMPP5 - mRNA,NM_022474.2:1110,SEBELIUS_2_C4636,MPP5,mRNA,0,,,Tight junction interactions,hsa04390;hsa04530,MPP5,NA,MPP5 - mRNA,False\nBRK1 - mRNA,NM_018462.4:710,SEBELIUS_2_C4636,BRK1,mRNA,0,,,RHO GTPases Activate WASPs and WAVEs;VEGFA-VEGFR2 Pathway;Regulation of actin dynamics for phagocytic cup formation,hsa04810,BRK1,NA,BRK1 - mRNA,False\nAKT1 - mRNA,NM_001014432.1:1275,SEBELIUS_2_C4636,AKT1,mRNA,0,,,TP53 Regulates Metabolic Genes;Deactivation of the beta-catenin transactivating complex;Activation of BAD and translocation to mitochondria ;eNOS activation;mTOR signalling;G beta.CL.gamma signalling through PI3Kgamma;KSRP (KHSRP) binds and destabilizes mRNA;AKT phosphorylates targets in the nucleus;VEGFR2 mediated vascular permeability;GPVI-mediated activation cascade;Tetrahydrobiopterin (BH4) synthesis# recycling# salvage and regulation;Butyrate Response Factor 1 (BRF1) binds and destabilizes mRNA;Signaling by ERBB2;PIP3 activates AKT signaling;Downstream signal transduction;GAB1 signalosome;Signaling by SCF-KIT;PI3KAKT activation;Role of LAT2NTALLAB on calcium mobilization;DAP12 signaling;Signaling by PDGF;CD28 co-stimulation;PI3K Cascade;IRS-related events triggered by IGF1R;VEGFA-VEGFR2 Pathway,hsa04010;hsa04012;hsa04014;hsa04015;hsa04022;hsa04024;hsa04062;hsa04066;hsa04068;hsa04071;hsa04072;hsa04150;hsa04151;hsa04152;hsa04210;hsa04211;hsa04213;hsa04261;hsa04370;hsa04380;hsa04510;hsa04530;hsa04550;hsa04611;hsa04620;hsa04630;hsa04660;hsa04662;hsa04664;hsa04666;hsa04668;hsa04722;hsa04725;hsa04728;hsa04910;hsa04914;hsa04915;hsa04917;hsa04919;hsa04920;hsa04922;hsa04923;hsa04931;hsa04932;hsa04933;hsa04973;hsa05142;hsa05145;hsa05152;hsa05160;hsa05161;hsa05162;hsa05164;hsa05166;hsa05169;hsa05200;hsa05205;hsa05210;hsa05211;hsa05212;hsa05213;hsa05214;hsa05215;hsa05218;hsa05220;hsa05221;hsa05222;hsa05223;hsa05230;hsa05231,AKT1,NA,AKT1 - mRNA,False\nDIAPH1 - mRNA,NM_005219.4:1398,SEBELIUS_2_C4636,DIAPH1,mRNA,0,,,RHO GTPases Activate Formins,hsa04510;hsa04810;hsa04933;hsa05131,DIAPH1,NA,DIAPH1 - mRNA,False\nABCF1 - mRNA,NM_001090.2:850,SEBELIUS_2_C4636,ABCF1,mRNA,0,,,,,ABCF1,NA,ABCF1 - mRNA,False\nIL1R1 b - mRNA,NM_001320984.1:2649,SEBELIUS_2_C4636,IL1R1 b,mRNA,0,,,,,,NA,IL1R1 b - mRNA,False\nSOS2 - mRNA,NM_006939.2:3845,SEBELIUS_2_C4636,SOS2,mRNA,0,,,Rho GTPase cycle;Activation of Rac;NRAGE signals death through JNK;G alpha (1213) signalling events,hsa04010;hsa04012;hsa04014;hsa04062;hsa04068;hsa04072;hsa04151;hsa04320;hsa04510;hsa04540;hsa04630;hsa04650;hsa04660;hsa04662;hsa04664;hsa04722;hsa04810;hsa04910;hsa04912;hsa04915;hsa04917;hsa05034;hsa05160;hsa05200;hsa05205;hsa05206;hsa05211;hsa05213;hsa05214;hsa05215;hsa05220;hsa05221;hsa05223;hsa05231,SOS2,NA,SOS2 - mRNA,False\nIRS1 - mRNA,NM_005544.2:6224,SEBELIUS_2_C4636,IRS1,mRNA,0,,,Constitutive Signaling by Aberrant PI3K in Cancer;RAFMAP kinase cascade;PI3K Cascade;Growth hormone receptor signaling;IRS-related events triggered by IGF1R;PI3KAKT activation;Downstream signal transduction;GAB1 signalosome;Signaling by SCF-KIT;Role of LAT2NTALLAB on calcium mobilization;DAP12 signaling;Signaling by PDGF;Signaling by ERBB2;VEGFR2 mediated cell proliferation;Frs2-mediated activation;ARMS-mediated activation;Gastrin-CREB signalling pathway via PKC and MAPK;Interleukin receptor SHC signaling;FCERI mediated MAPK activation;VEGFA-VEGFR2 Pathway,hsa04022;hsa04068;hsa04150;hsa04151;hsa04152;hsa04211;hsa04213;hsa04722;hsa04910;hsa04920;hsa04923;hsa04930;hsa04931;hsa04932;hsa04960;hsa05206,IRS1,NA,IRS1 - mRNA,False\nDROSHA - mRNA,NM_013235.4:1860,SEBELIUS_2_C4636,DROSHA,mRNA,0,,,MicroRNA (miRNA) biogenesis,hsa05205,DROSHA,NA,DROSHA - mRNA,False\nCCNG2 - mRNA,NM_004354.2:336,SEBELIUS_2_C4636,CCNG2,mRNA,0,,,,hsa04068;hsa04115,CCNG2,NA,CCNG2 - mRNA,False\nDVL3 - mRNA,NM_004423.3:1012,SEBELIUS_2_C4636,DVL3,mRNA,0,,,Disassembly of the destruction complex and recruitment of AXIN to the membrane;WNT mediated activation of DVL;RHO GTPases Activate Formins;PCPCE pathway,hsa04310;hsa04330;hsa04390;hsa04550;hsa04916;hsa05166;hsa05200;hsa05217,DVL3,NA,DVL3 - mRNA,False\nMAPK8 - mRNA,NM_002750.2:945,SEBELIUS_2_C4636,MAPK8,mRNA,0,,,Activation of the AP-1 family of transcription factors;FCERI mediated MAPK activation;Activation of BMF and translocation to mitochondria;JNK (c-Jun kinases) phosphorylation and  activation mediated by activated human TAK1,hsa04010;hsa04012;hsa04014;hsa04024;hsa04068;hsa04071;hsa04141;hsa04310;hsa04380;hsa04510;hsa04620;hsa04621;hsa04664;hsa04668;hsa04722;hsa04723;hsa04728;hsa04750;hsa04910;hsa04912;hsa04914;hsa04917;hsa04920;hsa04930;hsa04931;hsa04932;hsa04933;hsa05120;hsa05131;hsa05132;hsa05133;hsa05142;hsa05145;hsa05152;hsa05160;hsa05161;hsa05164;hsa05166;hsa05168;hsa05169;hsa05200;hsa05210;hsa05212;hsa05231,MAPK8,NA,MAPK8 - mRNA,False\nSOS1 - mRNA,NM_005633.2:1635,SEBELIUS_2_C4636,SOS1,mRNA,0,,,Signaling by FGFR2 in disease;Antigen activates B Cell Receptor (BCR) leading to generation of second messengers;SHC-mediated cascade.CL.FGFR3;Signaling by FGFR3 in disease;RAFMAP kinase cascade;SHC-related events triggered by IGF1R;Signaling by FGFR4 in disease;SHC-mediated cascade.CL.FGFR1;GRB2.CL.SOS provides linkage to MAPK signaling for Integrins ;Tie2 Signaling;Role of LAT2NTALLAB on calcium mobilization;Signaling by FGFR1 in disease;Signal attenuation;NRAGE signals death through JNK;SHC-mediated cascade.CL.FGFR4;FCERI mediated Ca.PLS.2 mobilization;Constitutive Signaling by EGFRvIII;FCERI mediated MAPK activation;Signaling by ERBB2;Signaling by SCF-KIT;DAP12 signaling;Gastrin-CREB signalling pathway via PKC and MAPK;Interleukin receptor SHC signaling;IRS-related events triggered by IGF1R;Downstream signal transduction;VEGFR2 mediated cell proliferation;Frs2-mediated activation;ARMS-mediated activation;Signaling by PDGF;VEGFA-VEGFR2 Pathway;Integrin alphaIIb beta3 signaling;Cell surface interactions at the vascular wall,hsa04010;hsa04012;hsa04014;hsa04062;hsa04068;hsa04072;hsa04151;hsa04320;hsa04510;hsa04540;hsa04630;hsa04650;hsa04660;hsa04662;hsa04664;hsa04722;hsa04810;hsa04910;hsa04912;hsa04915;hsa04917;hsa05034;hsa05160;hsa05200;hsa05205;hsa05206;hsa05211;hsa05213;hsa05214;hsa05215;hsa05220;hsa05221;hsa05223;hsa05231,SOS1,NA,SOS1 - mRNA,False\nGSK3B - mRNA,NM_002093.2:925,SEBELIUS_2_C4636,GSK3B,mRNA,0,,,Misspliced GSK3beta mutants stabilize beta-catenin;Constitutive Signaling by AKT1 E17K in Cancer;APC truncation mutants have impaired AXIN binding;T41 mutants of beta-catenin aren.SQT.t phosphorylated;AXIN missense mutants destabilize the destruction complex;S45 mutants of beta-catenin aren.SQT.t phosphorylated;Regulation of HSF1-mediated heat shock response;AKT phosphorylates targets in the cytosol;Disassembly of the destruction complex and recruitment of AXIN to the membrane;S37 mutants of beta-catenin aren.SQT.t phosphorylated;S33 mutants of beta-catenin aren.SQT.t phosphorylated;Truncations of AMER1 destabilize the destruction complex;Degradation of GLI2 by the proteasome;PIP3 activates AKT signaling;Downstream signal transduction;GAB1 signalosome;Signaling by SCF-KIT;PI3KAKT activation;Role of LAT2NTALLAB on calcium mobilization;DAP12 signaling;Signaling by PDGF;Signaling by ERBB2,hsa04012;hsa04062;hsa04110;hsa04151;hsa04310;hsa04360;hsa04390;hsa04510;hsa04550;hsa04660;hsa04662;hsa04722;hsa04728;hsa04910;hsa04916;hsa04917;hsa04919;hsa04931;hsa04932;hsa05010;hsa05160;hsa05162;hsa05164;hsa05166;hsa05169;hsa05200;hsa05210;hsa05213;hsa05215;hsa05217,GSK3B,NA,GSK3B - mRNA,False\nEP300 - mRNA,NM_001429.2:715,SEBELIUS_2_C4636,EP300,mRNA,0,,,LRR FLII-interacting protein 1 (LRRFIP1) activates type I IFN production;TRAF6 mediated IRF7 activation;Constitutive Signaling by NOTCH1 PEST Domain Mutants;Polo-like kinase mediated events;B-WICH complex positively regulates rRNA expression;CD209 (DC-SIGN) signaling;Circadian Clock,hsa04024;hsa04066;hsa04068;hsa04110;hsa04310;hsa04330;hsa04350;hsa04520;hsa04630;hsa04720;hsa04916;hsa04919;hsa04922;hsa05016;hsa05152;hsa05161;hsa05164;hsa05166;hsa05168;hsa05169;hsa05200;hsa05203;hsa05206;hsa05211;hsa05215,EP300,NA,EP300 - mRNA,False\nCCND3 - mRNA,NM_001760.2:1215,SEBELIUS_2_C4636,CCND3,mRNA,0,,,Cyclin D associated events in G1;MAPK6MAPK4 signaling;Transcriptional regulation of white adipocyte differentiation,hsa04110;hsa04115;hsa04151;hsa04310;hsa04390;hsa04510;hsa04630;hsa05162;hsa05166;hsa05203,CCND3,NA,CCND3 - mRNA,False\nKRAS - mRNA,NM_004985.3:327,SEBELIUS_2_C4636,KRAS,mRNA,0,,,Signaling by FGFR2 in disease;Regulation of RAS by GAPs;SHC-mediated cascade.CL.FGFR3;Signaling by FGFR3 in disease;MAP2K and MAPK activation;SHC-related events triggered by IGF1R;Signaling by FGFR4 in disease;SHC-mediated cascade.CL.FGFR1;Tie2 Signaling;Signaling by FGFR1 in disease;Activation of RAS in B cells;SHC-mediated cascade.CL.FGFR4;Constitutive Signaling by EGFRvIII;CD209 (DC-SIGN) signaling;FCERI mediated MAPK activation;RAFMAP kinase cascade;Downstream signal transduction;VEGFR2 mediated cell proliferation;Signaling by SCF-KIT;Frs2-mediated activation;ARMS-mediated activation;Gastrin-CREB signalling pathway via PKC and MAPK;Interleukin receptor SHC signaling;DAP12 signaling;Signaling by PDGF;VEGFA-VEGFR2 Pathway;Signaling by ERBB2;IRS-related events triggered by IGF1R;Cell surface interactions at the vascular wall,hsa04010;hsa04012;hsa04014;hsa04015;hsa04062;hsa04068;hsa04071;hsa04072;hsa04151;hsa04211;hsa04213;hsa04320;hsa04360;hsa04370;hsa04530;hsa04540;hsa04550;hsa04650;hsa04660;hsa04662;hsa04664;hsa04720;hsa04722;hsa04725;hsa04726;hsa04730;hsa04810;hsa04910;hsa04912;hsa04914;hsa04915;hsa04916;hsa04917;hsa04919;hsa04921;hsa04933;hsa04960;hsa05034;hsa05160;hsa05161;hsa05166;hsa05200;hsa05203;hsa05205;hsa05206;hsa05210;hsa05211;hsa05212;hsa05213;hsa05214;hsa05215;hsa05216;hsa05218;hsa05219;hsa05220;hsa05221;hsa05223;hsa05230;hsa05231,KRAS,NA,KRAS - mRNA,False\nCSNK1A1 - mRNA,NM_001892.4:2053,SEBELIUS_2_C4636,CSNK1A1,mRNA,0,,,Misspliced GSK3beta mutants stabilize beta-catenin;APC truncation mutants have impaired AXIN binding;T41 mutants of beta-catenin aren.SQT.t phosphorylated;AXIN missense mutants destabilize the destruction complex;S45 mutants of beta-catenin aren.SQT.t phosphorylated;Disassembly of the destruction complex and recruitment of AXIN to the membrane;S37 mutants of beta-catenin aren.SQT.t phosphorylated;S33 mutants of beta-catenin aren.SQT.t phosphorylated;Beta-catenin phosphorylation cascade;Degradation of GLI2 by the proteasome,hsa04310,CSNK1A1,NA,CSNK1A1 - mRNA,False\nRAB13 - mRNA,NM_002870.3:332,SEBELIUS_2_C4636,RAB13,mRNA,0,,,Translocation of GLUT4 to the plasma membrane,hsa04530,RAB13,NA,RAB13 - mRNA,False\nPDGFRA - mRNA,NM_006206.3:1925,SEBELIUS_2_C4636,PDGFRA,mRNA,0,,,PIP3 activates AKT signaling;Constitutive Signaling by Aberrant PI3K in Cancer;RAFMAP kinase cascade;Signaling by PDGF;Downstream signal transduction;GAB1 signalosome;Signaling by SCF-KIT;PI3KAKT activation;Role of LAT2NTALLAB on calcium mobilization;DAP12 signaling;Signaling by ERBB2;VEGFR2 mediated cell proliferation;Frs2-mediated activation;ARMS-mediated activation;Gastrin-CREB signalling pathway via PKC and MAPK;Interleukin receptor SHC signaling;FCERI mediated MAPK activation;VEGFA-VEGFR2 Pathway;IRS-related events triggered by IGF1R,hsa04010;hsa04014;hsa04015;hsa04020;hsa04060;hsa04072;hsa04144;hsa04151;hsa04510;hsa04540;hsa04810;hsa05166;hsa05200;hsa05206;hsa05214;hsa05215;hsa05218;hsa05230;hsa05231,PDGFRA,NA,PDGFRA - mRNA,False\nPPP1CA - mRNA,NM_002708.3:505,SEBELIUS_2_C4636,PPP1CA,mRNA,0,,,DARPP-32 events;Downregulation of TGF-beta receptor signaling;Circadian Clock;TGF-beta receptor signaling activates SMADs,hsa03015;hsa04022;hsa04024;hsa04114;hsa04261;hsa04270;hsa04390;hsa04510;hsa04611;hsa04720;hsa04728;hsa04750;hsa04810;hsa04910;hsa04921;hsa04931;hsa05031;hsa05034;hsa05168;hsa05205,PPP1CA,NA,PPP1CA - mRNA,False\nVCL - mRNA,NM_014000.2:4780,SEBELIUS_2_C4636,VCL,mRNA,0,,,Smooth Muscle Contraction;MAP2K and MAPK activation;Platelet degranulation ;RAFMAP kinase cascade;Downstream signal transduction;VEGFR2 mediated cell proliferation;Signaling by SCF-KIT;Frs2-mediated activation;ARMS-mediated activation;Gastrin-CREB signalling pathway via PKC and MAPK;Interleukin receptor SHC signaling;FCERI mediated MAPK activation;DAP12 signaling;Signaling by PDGF;VEGFA-VEGFR2 Pathway;Signaling by ERBB2;IRS-related events triggered by IGF1R,hsa04510;hsa04520;hsa04670;hsa04810;hsa05100;hsa05131;hsa05146,VCL,NA,VCL - mRNA,False\nACTN4 - mRNA,NM_004924.4:1768,SEBELIUS_2_C4636,ACTN4,mRNA,0,,,Nephrin interactions;Platelet degranulation ,hsa04510;hsa04520;hsa04530;hsa04670;hsa04810;hsa05146;hsa05203;hsa05412,ACTN4,NA,ACTN4 - mRNA,False\nACTN1 - mRNA,NM_001102.3:2760,SEBELIUS_2_C4636,ACTN1,mRNA,0,,,Syndecan interactions;Nephrin interactions;Platelet degranulation ;Non-integrin membrane-ECM interactions;Cell-extracellular matrix interactions,hsa04510;hsa04520;hsa04530;hsa04670;hsa04810;hsa05146;hsa05203;hsa05412,ACTN1,NA,ACTN1 - mRNA,False\nITGA5 - mRNA,NM_002205.2:925,SEBELIUS_2_C4636,ITGA5,mRNA,0,,,Signal transduction by L1;Elastic fibre formation;Integrin cell surface interactions;Cell surface interactions at the vascular wall,hsa04145;hsa04151;hsa04510;hsa04512;hsa04640;hsa04810;hsa05100;hsa05131;hsa05133;hsa05205;hsa05206;hsa05410;hsa05412;hsa05414,ITGA5,NA,ITGA5 - mRNA,False\nITGB5 - mRNA,NM_002213.3:2560,SEBELIUS_2_C4636,ITGB5,mRNA,0,,,Molecules associated with elastic fibres;Syndecan interactions;Smooth Muscle Contraction;ECM proteoglycans;Integrin cell surface interactions;Elastic fibre formation;Non-integrin membrane-ECM interactions,hsa04145;hsa04151;hsa04510;hsa04512;hsa04810;hsa05205;hsa05410;hsa05412;hsa05414,ITGB5,NA,ITGB5 - mRNA,False\nCDC42 - mRNA,NM_001039802.1:390,SEBELIUS_2_C4636,CDC42,mRNA,0,,,DCC mediated attractive signaling;Factors involved in megakaryocyte development and platelet production;VEGFA-VEGFR2 Pathway;RHO GTPases activate KTN1;CD28 dependent Vav1 pathway;Inactivation of Cdc42 and Rac;Rho GTPase cycle;EPHB-mediated forward signaling;RHO GTPases Activate Formins;RHO GTPases activate PAKs;GPVI-mediated activation cascade;G alpha (1213) signalling events;RHO GTPases activate IQGAPs;Sema4D induced cell migration and growth-cone collapse;Regulation of actin dynamics for phagocytic cup formation;CD28 co-stimulation,hsa04010;hsa04014;hsa04015;hsa04062;hsa04144;hsa04360;hsa04370;hsa04510;hsa04520;hsa04530;hsa04660;hsa04666;hsa04670;hsa04722;hsa04810;hsa04912;hsa04932;hsa04933;hsa05100;hsa05120;hsa05130;hsa05131;hsa05132;hsa05200;hsa05203;hsa05205;hsa05211;hsa05212,CDC42,NA,CDC42 - mRNA,False\nITGA6 - mRNA,NM_000210.1:3065,SEBELIUS_2_C4636,ITGA6,mRNA,0,,,Syndecan interactions;Assembly of collagen fibrils and other multimeric structures;Basigin interactions;Non-integrin membrane-ECM interactions;Cell surface interactions at the vascular wall,hsa04151;hsa04510;hsa04512;hsa04514;hsa04640;hsa04810;hsa05145;hsa05200;hsa05222;hsa05410;hsa05412;hsa05414,ITGA6,NA,ITGA6 - mRNA,False\nPDGFC - mRNA,NM_016205.2:2596,SEBELIUS_2_C4636,PDGFC,mRNA,0,,,Signaling by PDGF,hsa04014;hsa04015;hsa04060;hsa04072;hsa04151;hsa04510;hsa04540;hsa04810;hsa05215;hsa05218;hsa05231,PDGFC,NA,PDGFC - mRNA,False\nCD9 - mRNA,NM_001769.2:405,SEBELIUS_2_C4636,CD9,mRNA,0,,,Platelet degranulation ,hsa04640,CD9,NA,CD9 - mRNA,False\nPSEN1 - mRNA,NM_000021.2:900,SEBELIUS_2_C4636,PSEN1,mRNA,0,,,,hsa04310;hsa04330;hsa04722;hsa05010,PSEN1,NA,PSEN1 - mRNA,False\nCLTC - mRNA,NM_004859.2:290,SEBELIUS_2_C4636,CLTC,mRNA,0,,,MHC class II antigen presentation;PCPCE pathway,hsa04144;hsa05016;hsa05100,CLTC,NA,CLTC - mRNA,False\nFGFR1 - mRNA,NM_015850.2:1335,SEBELIUS_2_C4636,FGFR1,mRNA,0,,,RAFMAP kinase cascade;Constitutive Signaling by Aberrant PI3K in Cancer;SHC-mediated cascade.CL.FGFR1;Phospholipase C-mediated cascade.CL. FGFR1;Signaling by FGFR1 amplification mutants;PI3K Cascade;FGFR1b ligand binding and activation;Signaling by FGFR1 in disease;Downstream signal transduction;GAB1 signalosome;Signaling by SCF-KIT;PI3KAKT activation;Role of LAT2NTALLAB on calcium mobilization;DAP12 signaling;Signaling by PDGF;Signaling by ERBB2;VEGFR2 mediated cell proliferation;Frs2-mediated activation;ARMS-mediated activation;Gastrin-CREB signalling pathway via PKC and MAPK;Interleukin receptor SHC signaling;FCERI mediated MAPK activation;VEGFA-VEGFR2 Pathway;IRS-related events triggered by IGF1R,hsa04010;hsa04014;hsa04015;hsa04151;hsa04520;hsa04550;hsa04810;hsa05200;hsa05205;hsa05215;hsa05218;hsa05230,FGFR1,NA,FGFR1 - mRNA,False\nPTEN - mRNA,NM_000314.4:1351,SEBELIUS_2_C4636,PTEN,mRNA,0,,,Synthesis of PIPs at the plasma membrane;Downstream TCR signaling;PIP3 activates AKT signaling;Downstream signal transduction;GAB1 signalosome;Signaling by SCF-KIT;PI3KAKT activation;Role of LAT2NTALLAB on calcium mobilization;DAP12 signaling;Signaling by PDGF;Signaling by ERBB2,hsa00562;hsa04068;hsa04070;hsa04071;hsa04115;hsa04150;hsa04151;hsa04510;hsa04530;hsa04931;hsa05161;hsa05200;hsa05206;hsa05213;hsa05214;hsa05215;hsa05218;hsa05222;hsa05230,PTEN,NA,PTEN - mRNA,False\nPRKAA1 - mRNA,NM_006251.5:1455,SEBELIUS_2_C4636,PRKAA1,mRNA,0,,,TP53 Regulates Metabolic Genes;PI3K Cascade;IRS-related events triggered by IGF1R,hsa04068;hsa04150;hsa04151;hsa04152;hsa04211;hsa04213;hsa04910;hsa04920;hsa04921;hsa04922;hsa04931;hsa04932;hsa05410,PRKAA1,NA,PRKAA1 - mRNA,False\nPPP1CB - mRNA,NM_002709.2:393,SEBELIUS_2_C4636,PPP1CB,mRNA,0,,,RHO GTPases Activate ROCKs;Downregulation of TGF-beta receptor signaling;RHO GTPases activate PKNs;Circadian Clock;Regulation of PLK1 Activity at G2M Transition;TGF-beta receptor signaling activates SMADs,hsa03015;hsa04022;hsa04024;hsa04114;hsa04261;hsa04270;hsa04390;hsa04510;hsa04611;hsa04720;hsa04728;hsa04750;hsa04810;hsa04910;hsa04921;hsa04931;hsa05031;hsa05034;hsa05168;hsa05205,PPP1CB,NA,PPP1CB - mRNA,False\nSPTAN1 - mRNA,NM_003127.2:7610,SEBELIUS_2_C4636,SPTAN1,mRNA,0,,,Nephrin interactions;Caspase-mediated cleavage of cytoskeletal proteins;Downstream signal transduction;VEGFR2 mediated cell proliferation;Signaling by SCF-KIT;Frs2-mediated activation;ARMS-mediated activation;Gastrin-CREB signalling pathway via PKC and MAPK;Interleukin receptor SHC signaling;FCERI mediated MAPK activation;DAP12 signaling;Signaling by PDGF;VEGFA-VEGFR2 Pathway;Signaling by ERBB2;IRS-related events triggered by IGF1R;Apoptotic cleavage of cellular proteins,hsa04530,SPTAN1,NA,SPTAN1 - mRNA,False\nEPB41L2 - mRNA,NM_001199389.1:1880,SEBELIUS_2_C4636,EPB41L2,mRNA,0,,,,hsa04530,EPB41L2,NA,EPB41L2 - mRNA,False\nWASF2 - mRNA,NM_006990.2:485,SEBELIUS_2_C4636,WASF2,mRNA,0,,,RHO GTPases Activate WASPs and WAVEs;VEGFA-VEGFR2 Pathway;Regulation of actin dynamics for phagocytic cup formation,hsa04520;hsa04666;hsa04810;hsa05100;hsa05131;hsa05132;hsa05231,WASF2,NA,WASF2 - mRNA,False\nTGFB1 - mRNA,NM_000660.3:1260,SEBELIUS_2_C4636,TGFB1,mRNA,0,,,TGF-beta receptor signaling in EMT (epithelial to mesenchymal transition);TGFBR1 LBD Mutants in Cancer;Influenza Virus Induced Apoptosis;Molecules associated with elastic fibres;Syndecan interactions;ECM proteoglycans;Transcriptional regulation of white adipocyte differentiation;TGF-beta receptor signaling activates SMADs;Elastic fibre formation;Non-integrin membrane-ECM interactions,hsa04010;hsa04060;hsa04068;hsa04110;hsa04144;hsa04350;hsa04380;hsa04390;hsa04932;hsa04933;hsa05140;hsa05142;hsa05145;hsa05146;hsa05152;hsa05161;hsa05166;hsa05200;hsa05205;hsa05210;hsa05211;hsa05212;hsa05220;hsa05321;hsa05410;hsa05414,TGFB1,NA,TGFB1 - mRNA,False\nTUBB - mRNA,NM_178014.2:1955,SEBELIUS_2_C4636,TUBB,mRNA,0,,,,hsa04145;hsa04540;hsa05130,TUBB,NA,TUBB - mRNA,False\nSMAD3 - mRNA,NM_005902.3:4220,SEBELIUS_2_C4636,SMAD3,mRNA,0,,,Signaling by NODAL;TGF-beta receptor signaling activates SMADs,hsa04068;hsa04110;hsa04144;hsa04310;hsa04350;hsa04390;hsa04520;hsa04550;hsa04933;hsa05142;hsa05161;hsa05166;hsa05200;hsa05210;hsa05212;hsa05220;hsa05321,SMAD3,NA,SMAD3 - mRNA,False\nMAPK1 - mRNA,NM_138957.2:430,SEBELIUS_2_C4636,MAPK1,mRNA,0,,,Negative regulation of FGFR1 signaling;Signal transduction by L1;Negative regulation of FGFR2 signaling;MAP2K and MAPK activation;RHO GTPases Activate WASPs and WAVEs;Senescence-Associated Secretory Phenotype (SASP);Regulation of HSF1-mediated heat shock response;Thrombin signalling through proteinase activated receptors (PARs);Negative regulation of FGFR4 signaling;Oxidative Stress Induced Senescence;Signal attenuation;Negative regulation of FGFR3 signaling;Negative feedback regulation of MAPK pathway;Growth hormone receptor signaling;PLC beta mediated events;FCERI mediated MAPK activation;CREB phosphorylation through the activation of Ras;Negative regulation of MAPK pathway;RAFMAP kinase cascade;Downstream signal transduction;VEGFR2 mediated cell proliferation;Signaling by SCF-KIT;Frs2-mediated activation;ARMS-mediated activation;Gastrin-CREB signalling pathway via PKC and MAPK;Interleukin receptor SHC signaling;DAP12 signaling;Signaling by PDGF;VEGFA-VEGFR2 Pathway;Signaling by ERBB2;IRS-related events triggered by IGF1R,hsa04010;hsa04012;hsa04014;hsa04015;hsa04022;hsa04024;hsa04062;hsa04066;hsa04068;hsa04071;hsa04072;hsa04114;hsa04150;hsa04151;hsa04261;hsa04270;hsa04320;hsa04350;hsa04360;hsa04370;hsa04380;hsa04510;hsa04520;hsa04540;hsa04550;hsa04611;hsa04620;hsa04621;hsa04650;hsa04660;hsa04662;hsa04664;hsa04666;hsa04668;hsa04713;hsa04720;hsa04722;hsa04723;hsa04724;hsa04725;hsa04726;hsa04730;hsa04810;hsa04910;hsa04912;hsa04914;hsa04915;hsa04916;hsa04917;hsa04919;hsa04921;hsa04930;hsa04933;hsa04960;hsa05010;hsa05020;hsa05034;hsa05131;hsa05132;hsa05133;hsa05140;hsa05142;hsa05145;hsa05152;hsa05160;hsa05161;hsa05164;hsa05200;hsa05203;hsa05205;hsa05206;hsa05210;hsa05211;hsa05212;hsa05213;hsa05214;hsa05215;hsa05216;hsa05218;hsa05219;hsa05220;hsa05221;hsa05223;hsa05230;hsa05231,MAPK1,NA,MAPK1 - mRNA,False\nHSPB1 - mRNA,NM_001540.3:374,SEBELIUS_2_C4636,HSPB1,mRNA,0,,,MAPK6MAPK4 signaling;VEGFA-VEGFR2 Pathway,hsa04010;hsa04370;hsa05146;hsa05169,HSPB1,NA,HSPB1 - mRNA,False\nG6PD - mRNA,NM_000402.2:1155,SEBELIUS_2_C4636,G6PD,mRNA,0,,,TP53 Regulates Metabolic Genes,hsa01100;hsa05230,G6PD,NA,G6PD - mRNA,False\nPDGFRB - mRNA,NM_002609.3:840,SEBELIUS_2_C4636,PDGFRB,mRNA,0,,,PIP3 activates AKT signaling;Constitutive Signaling by Aberrant PI3K in Cancer;RAFMAP kinase cascade;Signaling by PDGF;Downstream signal transduction;GAB1 signalosome;Signaling by SCF-KIT;PI3KAKT activation;Role of LAT2NTALLAB on calcium mobilization;DAP12 signaling;Signaling by ERBB2;VEGFR2 mediated cell proliferation;Frs2-mediated activation;ARMS-mediated activation;Gastrin-CREB signalling pathway via PKC and MAPK;Interleukin receptor SHC signaling;FCERI mediated MAPK activation;VEGFA-VEGFR2 Pathway;IRS-related events triggered by IGF1R,hsa04010;hsa04014;hsa04015;hsa04020;hsa04060;hsa04072;hsa04151;hsa04510;hsa04540;hsa04810;hsa05166;hsa05200;hsa05206;hsa05214;hsa05215;hsa05218;hsa05230;hsa05231,PDGFRB,NA,PDGFRB - mRNA,False\nIQGAP1 - mRNA,NM_003870.3:820,SEBELIUS_2_C4636,IQGAP1,mRNA,0,,,MAP2K and MAPK activation;Nephrin interactions;RHO GTPases activate IQGAPs;RAFMAP kinase cascade;Downstream signal transduction;VEGFR2 mediated cell proliferation;Signaling by SCF-KIT;Frs2-mediated activation;ARMS-mediated activation;Gastrin-CREB signalling pathway via PKC and MAPK;Interleukin receptor SHC signaling;FCERI mediated MAPK activation;DAP12 signaling;Signaling by PDGF;VEGFA-VEGFR2 Pathway;Signaling by ERBB2;IRS-related events triggered by IGF1R,hsa04520;hsa04810;hsa05205,IQGAP1,NA,IQGAP1 - mRNA,False\nADAM9 - mRNA,NM_001005845.1:695,SEBELIUS_2_C4636,ADAM9,mRNA,0,,,,,ADAM9,NA,ADAM9 - mRNA,False\nNCKAP1 - mRNA,NM_013436.3:2353,SEBELIUS_2_C4636,NCKAP1,mRNA,0,,,RHO GTPases Activate WASPs and WAVEs;VEGFA-VEGFR2 Pathway;Regulation of actin dynamics for phagocytic cup formation,hsa04810,NCKAP1,NA,NCKAP1 - mRNA,False\nPPP2R1A - mRNA,NM_014225.3:1440,SEBELIUS_2_C4636,PPP2R1A,mRNA,0,,,MASTL Facilitates Mitotic Progression;Regulation of PLK1 Activity at G2M Transition;Disassembly of the destruction complex and recruitment of AXIN to the membrane;S37 mutants of beta-catenin aren.SQT.t phosphorylated;Cyclin AB1 associated events during G2M transition;RAF activation;RHO GTPases Activate Formins;S33 mutants of beta-catenin aren.SQT.t phosphorylated;Loss of Nlp from mitotic centrosomes;DARPP-32 events;Beta-catenin phosphorylation cascade;Initiation of Nuclear Envelope Reformation;Recruitment of mitotic centrosome proteins and complexes;Spry regulation of FGF signaling;Truncations of AMER1 destabilize the destruction complex;CTLA4 inhibitory signaling;AXIN missense mutants destabilize the destruction complex;S45 mutants of beta-catenin aren.SQT.t phosphorylated;Anchoring of the basal body to the plasma membrane;Nonsense Mediated Decay (NMD) enhanced by the Exon Junction Complex (EJC);Cyclin D associated events in G1;Platelet sensitization by LDL;Resolution of Sister Chromatid Cohesion;Inhibition of replication initiation of damaged DNA by RB1E2F1;RAFMAP kinase cascade;Downstream signal transduction;VEGFR2 mediated cell proliferation;Signaling by SCF-KIT;Frs2-mediated activation;ARMS-mediated activation;Gastrin-CREB signalling pathway via PKC and MAPK;Interleukin receptor SHC signaling;FCERI mediated MAPK activation;DAP12 signaling;Signaling by PDGF;VEGFA-VEGFR2 Pathway;Signaling by ERBB2;IRS-related events triggered by IGF1R;Negative regulation of FGFR3 signaling;Negative regulation of FGFR4 signaling;Negative regulation of FGFR2 signaling;Negative regulation of FGFR1 signaling,hsa03015;hsa04071;hsa04114;hsa04151;hsa04152;hsa04261;hsa04350;hsa04390;hsa04530;hsa04728;hsa04730;hsa05142;hsa05160,PPP2R1A,NA,PPP2R1A - mRNA,False\nRAC1 - mRNA,NM_198829.1:1250,SEBELIUS_2_C4636,RAC1,mRNA,0,,,MAPK6MAPK4 signaling;Signaling by SCF-KIT;Inactivation of Cdc42 and Rac;DAP12 signaling;Ephrin signaling;EPH-ephrin mediated repulsion of cells;Activation of Rac;RHO GTPases Activate Formins;VEGFR2 mediated vascular permeability;RHO GTPases activate PAKs;Sema4D mediated inhibition of cell attachment and migration;G alpha (1213) signalling events;Regulation of actin dynamics for phagocytic cup formation;Sema4D induced cell migration and growth-cone collapse;Signal transduction by L1;RHO GTPases activate CIT;RHO GTPases Activate WASPs and WAVEs;Factors involved in megakaryocyte development and platelet production;RHO GTPases activate KTN1;CD28 dependent Vav1 pathway;DSCAM interactions;NRAGE signals death through JNK;GPVI-mediated activation cascade;Sema3A PAK dependent Axon repulsion;RHO GTPases activate PKNs;SEMA3A-Plexin repulsion signaling by inhibiting Integrin adhesion;RHO GTPases activate IQGAPs;PCPCE pathway;VEGFA-VEGFR2 Pathway;CD28 co-stimulation,hsa04010;hsa04014;hsa04015;hsa04024;hsa04062;hsa04071;hsa04145;hsa04151;hsa04310;hsa04360;hsa04370;hsa04380;hsa04510;hsa04520;hsa04620;hsa04650;hsa04662;hsa04664;hsa04666;hsa04670;hsa04722;hsa04810;hsa04932;hsa04933;hsa04972;hsa05100;hsa05120;hsa05131;hsa05132;hsa05200;hsa05203;hsa05205;hsa05210;hsa05211;hsa05212;hsa05231;hsa05416,RAC1,NA,RAC1 - mRNA,False\nGNAI1 - mRNA,NM_002069.4:670,SEBELIUS_2_C4636,GNAI1,mRNA,0,,,ADP signalling through P2Y purinoceptor 12;Adrenaline#noradrenaline inhibits insulin secretion,hsa04015;hsa04022;hsa04024;hsa04062;hsa04071;hsa04261;hsa04360;hsa04530;hsa04540;hsa04611;hsa04670;hsa04713;hsa04723;hsa04724;hsa04725;hsa04726;hsa04727;hsa04728;hsa04730;hsa04914;hsa04915;hsa04916;hsa04921;hsa04923;hsa04971;hsa05032;hsa05034;hsa05133;hsa05142;hsa05145;hsa05200,GNAI1,NA,GNAI1 - mRNA,False\nPPP2CB - mRNA,NM_001009552.1:1075,SEBELIUS_2_C4636,PPP2CB,mRNA,0,,,MASTL Facilitates Mitotic Progression;Misspliced GSK3beta mutants stabilize beta-catenin;T41 mutants of beta-catenin aren.SQT.t phosphorylated;ERKs are inactivated;Glycolysis;Negative regulation of MAPK pathway;Disassembly of the destruction complex and recruitment of AXIN to the membrane;S37 mutants of beta-catenin aren.SQT.t phosphorylated;Cyclin AB1 associated events during G2M transition;RAF activation;RHO GTPases Activate Formins;S33 mutants of beta-catenin aren.SQT.t phosphorylated;DARPP-32 events;Beta-catenin phosphorylation cascade;Spry regulation of FGF signaling;Truncations of AMER1 destabilize the destruction complex;CTLA4 inhibitory signaling;AXIN missense mutants destabilize the destruction complex;S45 mutants of beta-catenin aren.SQT.t phosphorylated;Cyclin D associated events in G1;Platelet sensitization by LDL;Resolution of Sister Chromatid Cohesion;Inhibition of replication initiation of damaged DNA by RB1E2F1;RAFMAP kinase cascade;Downstream signal transduction;VEGFR2 mediated cell proliferation;Signaling by SCF-KIT;Frs2-mediated activation;ARMS-mediated activation;Gastrin-CREB signalling pathway via PKC and MAPK;Interleukin receptor SHC signaling;FCERI mediated MAPK activation;DAP12 signaling;Signaling by PDGF;VEGFA-VEGFR2 Pathway;Signaling by ERBB2;IRS-related events triggered by IGF1R;Negative regulation of FGFR3 signaling;Negative regulation of FGFR4 signaling;Negative regulation of FGFR2 signaling;Negative regulation of FGFR1 signaling,hsa03015;hsa04071;hsa04114;hsa04151;hsa04152;hsa04261;hsa04350;hsa04390;hsa04530;hsa04728;hsa04730;hsa05142;hsa05160,PPP2CB,NA,PPP2CB - mRNA,False\nGSTM3 - mRNA,NM_000849.4:602,SEBELIUS_2_C4636,GSTM3,mRNA,0,,,,,GSTM3,NA,GSTM3 - mRNA,False\nRHOA - mRNA,NM_001664.2:1230,SEBELIUS_2_C4636,RHOA,mRNA,0,,,RHO GTPases activate KTN1;EPHB-mediated forward signaling;Rho GTPase cycle;EPHA-mediated growth cone collapse;RHO GTPases Activate ROCKs;Axonal growth inhibition (RHOA activation);PI3KAKT activation;G beta.CL.gamma signalling through PI3Kgamma;RHO GTPases Activate Formins;RHO GTPases activate PKNs;GPVI-mediated activation cascade;G alpha (1213) signalling events;Sema4D mediated inhibition of cell attachment and migration;PCPCE pathway;Sema4D induced cell migration and growth-cone collapse,hsa04014;hsa04015;hsa04022;hsa04024;hsa04062;hsa04071;hsa04072;hsa04144;hsa04270;hsa04310;hsa04350;hsa04360;hsa04510;hsa04520;hsa04530;hsa04611;hsa04660;hsa04670;hsa04722;hsa04810;hsa04921;hsa04972;hsa05100;hsa05130;hsa05133;hsa05152;hsa05200;hsa05203;hsa05205;hsa05206;hsa05210,RHOA,NA,RHOA - mRNA,False\nRAB11A - mRNA,NM_004663.3:710,SEBELIUS_2_C4636,RAB11A,mRNA,0,,,Anchoring of the basal body to the plasma membrane,hsa04144;hsa04972,RAB11A,NA,RAB11A - mRNA,False\n";
	heatmapData_UyySR3RU_9hQ_hWdumMiueg = "Dimname,20161013_sebelius2-101316_01_01.RCC,20180703_1790-jr1_SA60miR142_09.RCC,20161013_sebelius2-101316_01_04.RCC,20180712_1790-jr2_SA40miR142_03.RCC,20180703_1790-jr1_SA40miR142_01.RCC,20180712_1790-jr2_SA49amiR142_06.RCC,20180703_1790-jr1_SA50bmiR18b_04.RCC,20180703_1790-jr1_SA62miR18b_11.RCC,20180712_1790-jr2_SA62miR18b_11.RCC,20180703_1790-jr1_SA63miR890_12.RCC,20180712_1790-jr2_SA63miR890_12.RCC,20180703_1790-jr1_SA44miR890_02.RCC,20180712_1790-jr2_SA44miR890_04.RCC,20180703_1790-jr1_SA51miR890_05.RCC,20180712_1790-jr2_SA51bmiR890_07.RCC,20161013_sebelius2-101316_01_03.RCC,20180712_1790-jr2_SA36Control_01.RCC,20180712_1790-jr2_SA38cel67_02.RCC,20180712_1790-jr2_SA59cel67_09.RCC,20180703_1790-jr1_SA57Control_08.RCC,20180712_1790-jr2_SA57Control_08.RCC,20180703_1790-jr1_SA48bcel67_03.RCC,20180712_1790-jr2_SA48bcel67_05.RCC,20180703_1790-jr1_SA61miR155_10.RCC,20180712_1790-jr2_SA61miR155_10.RCC,20180703_1790-jr1_SA54amiR155_06.RCC,20180703_1790-jr1_SA54bmiR155_07.RCC\nPIK3R5 - mRNA,2.35949571944106,0.891349145029288,0.469916668321527,1.78336651708132,-0.0105179049980162,0.301219147757543,0.312147895756265,0.363976105784351,-0.216642731515099,1.13325512034551,-0.336406073261212,0.491420450008309,-0.126838625911843,1.09981304891791,2.55222395130172,1.11498819078674,2.23746219806384,1.95913407695416,0.829952977998165,1.06706191171516,1.75443478151005,-0.255654297375793,-0.342676149847659,1.564131922885,0.946337736366942,0.555616221032077,-0.251827405654147\nCDX2 - mRNA,2.35949571944106,0.891349145029288,0.469916668321527,2.10529461196868,0.989482095001984,1.30121914775754,0.312147895756265,1.94893860650551,2.10528536337226,1.71821762106667,2.8335189281811,2.07638295072947,2.68051629614576,2.42174114380527,1.38229894985941,1.11498819078674,2.40738719950615,3.28106217184152,2.63730790005577,1.06706191171516,2.23986160868029,2.06627379751157,1.9792519450397,2.564131922885,2.16873015770339,0.555616221032077,0.748172594345853\nTJP3 - mRNA,0.774533218719899,0.891349145029288,1.46991666832153,3.10529461196868,1.57444459572314,1.30121914775754,2.31214789575627,0.363976105784351,2.10528536337226,1.13325512034551,1.24855642745994,1.49142045000831,0.873161374088157,2.09981304891791,0.967261450580565,2.6999506915079,1.822424698785,2.95913407695416,2.15188107288553,1.65202441243631,2.01746918734384,2.06627379751157,2.82724885159465,0.979169422163848,2.16873015770339,0.555616221032077,1.33313509506701\nCXCR4 - mRNA,2.5818881407775,1.89134914502929,2.46991666832153,2.59072143913892,1.57444459572314,0.301219147757543,2.31214789575627,0.363976105784351,1.36831976920606,1.71821762106667,2.47094884879639,1.49142045000831,0.873161374088157,1.68477554963906,2.18965387191701,2.6999506915079,1.5593902929512,3.28106217184152,2.99987797944048,1.06706191171516,2.23986160868029,2.32930820334536,1.65732385015234,2.30109751705121,0.946337736366942,2.87754431591944,1.33313509506701\nCGN - mRNA,2.35949571944106,1.89134914502929,2.05487916904268,2.10529461196868,1.57444459572314,0.301219147757543,2.89711039647742,1.36397610578435,1.36831976920606,0.13325512034551,1.66359392673879,0.491420450008309,1.87316137408816,0.0998130489179072,2.84173056849671,0.114988190786745,1.5593902929512,0.959134076954157,1.41491547871932,1.06706191171516,2.23986160868029,0.744345702624207,1.65732385015234,2.30109751705121,0.946337736366942,1.55561622103208,1.33313509506701\nACTN3 - mRNA,3.2339648373572,1.89134914502929,2.46991666832153,-0.216633482918684,2.31141018988935,1.8861816484787,1.31214789575627,0.363976105784351,1.36831976920606,1.13325512034551,1.24855642745994,2.07638295072947,0.873161374088157,1.09981304891791,0.967261450580565,1.11498819078674,1.822424698785,1.95913407695416,1.41491547871932,1.06706191171516,0.432506686622686,1.32930820334536,1.65732385015234,-0.0208305778361524,2.36137523564579,0.555616221032077,2.07010068923321\nCA2 - mRNA,1.35949571944106,0.891349145029288,2.79184476320889,2.10529461196868,1.57444459572314,1.8861816484787,0.312147895756265,2.68590420067171,1.36831976920606,2.13325512034551,2.24855642745994,4.1918601681494,4.51701756386288,3.90716797097551,3.0827386680005,2.11498819078674,2.23746219806384,2.54409657767531,2.15188107288553,2.06706191171516,3.01746918734384,1.74434570262421,2.2422863508735,2.30109751705121,0.946337736366942,2.87754431591944,2.55552751640346\nCRB3 - mRNA,2.5818881407775,0.891349145029288,3.05487916904268,2.59072143913892,2.31141018988935,2.6231472426449,0.312147895756265,2.94893860650551,2.5907121905425,3.45518321523287,2.8335189281811,3.49142045000831,2.68051629614576,3.5592446675552,2.38229894985941,4.02187878639526,2.822424698785,2.95913407695416,2.82995297799816,3.06706191171516,3.43250668662269,1.74434570262421,1.65732385015234,3.43860104080114,3.36137523564579,2.87754431591944,2.91809759578816\nCLDN4 - mRNA,2.5818881407775,0.891349145029288,2.46991666832153,2.36832901780247,1.57444459572314,1.30121914775754,2.31214789575627,2.36397610578435,2.36831976920606,3.13325512034551,3.12302554537608,2.49142045000831,2.19508946897552,2.68477554963906,2.84173056849671,2.11498819078674,3.23746219806384,2.54409657767531,3.99987797944048,3.52649353035245,3.43250668662269,0.744345702624207,2.46467877220995,1.97916942216385,1.94633773636694,3.14057872175323,2.74817259434585\nALPI - mRNA,2.7745332187199,0.891349145029288,2.05487916904268,3.36832901780247,0.989482095001984,2.30121914775754,2.89711039647742,1.36397610578435,2.10528536337226,2.13325512034551,3.12302554537608,2.07638295072947,3.19508946897552,2.09981304891791,2.18965387191701,2.6999506915079,2.23746219806384,2.54409657767531,3.15188107288553,1.06706191171516,2.75443478151005,1.32930820334536,2.82724885159465,1.97916942216385,2.16873015770339,2.55561622103208,2.91809759578816\nPIK3CG - mRNA,3.47497293686099,0.891349145029288,2.79184476320889,3.24279813571861,1.98948209500198,1.8861816484787,2.63407599064363,1.94893860650551,1.7833572684849,2.45518321523287,1.66359392673879,2.49142045000831,2.87316137408816,2.68477554963906,1.96726145058057,2.92234311284435,2.04481712012145,2.54409657767531,3.41491547871932,3.06706191171516,3.3393972822312,2.55170062468181,1.65732385015234,3.564131922885,3.16873015770339,0.555616221032077,0.748172594345853\nIL1R2 - mRNA,2.7745332187199,2.47631164575044,3.05487916904268,3.36832901780247,2.31141018988935,2.8861816484787,2.31214789575627,0.363976105784351,3.10528536337226,2.94061004240311,3.24855642745994,2.81334854489567,3.04308637553047,2.42174114380527,2.55222395130172,2.92234311284435,2.40738719950615,1.95913407695416,3.15188107288553,3.23698691315747,3.89193830525998,3.65123629823273,2.82724885159465,3.30109751705121,2.94633773636694,2.55561622103208,2.33313509506701\nHPSE - mRNA,2.5818881407775,0.891349145029288,2.05487916904268,2.36832901780247,2.98948209500198,2.8861816484787,2.31214789575627,1.36397610578435,1.7833572684849,2.71821762106667,2.98552202162615,2.81334854489567,2.68051629614576,0.0998130489179072,2.96726145058057,2.6999506915079,3.04481712012145,2.54409657767531,2.82995297799816,2.06706191171516,3.43250668662269,2.74434570262421,2.65732385015234,3.14909442360616,2.68330333053315,2.14057872175323,2.33313509506701\nITGB4 - mRNA,3.09646131360726,2.89134914502929,2.46991666832153,2.78336651708132,2.31141018988935,1.8861816484787,2.31214789575627,2.68590420067171,2.5907121905425,4.13325512034551,2.47094884879639,3.49142045000831,2.87316137408816,1.68477554963906,3.18965387191701,3.43691628567411,3.40738719950615,3.28106217184152,2.41491547871932,2.38899000660252,3.68043420006627,2.55170062468181,2.82724885159465,3.14909442360616,3.06181495378688,2.14057872175323,2.91809759578816\nWAS - mRNA,2.35949571944106,3.47631164575044,2.46991666832153,2.78336651708132,2.31141018988935,3.30121914775754,3.11950281781387,0.363976105784351,2.10528536337226,0.13325512034551,1.66359392673879,0.491420450008309,3.19508946897552,2.42174114380527,3.18965387191701,3.28491319222906,2.04481712012145,3.28106217184152,2.63730790005577,2.65202441243631,3.01746918734384,2.74434570262421,2.2422863508735,2.78652434422145,2.16873015770339,2.87754431591944,2.74817259434585\nVAV3 - mRNA,0.774533218719899,1.89134914502929,3.46991666832153,3.24279813571861,1.98948209500198,2.30121914775754,1.89711039647742,1.94893860650551,2.10528536337226,2.13325512034551,1.66359392673879,0.491420450008309,1.87316137408816,0.0998130489179072,1.38229894985941,2.92234311284435,2.23746219806384,1.95913407695416,2.82995297799816,2.06706191171516,2.23986160868029,2.32930820334536,-0.342676149847659,1.564131922885,2.16873015770339,2.14057872175323,1.74817259434585\nMYH14 - mRNA,-0.225466781280101,2.47631164575044,3.05487916904268,1.78336651708132,2.57444459572314,0.301219147757543,1.31214789575627,1.94893860650551,0.783357268484901,1.71821762106667,1.98552202162615,1.49142045000831,1.87316137408816,1.09981304891791,1.96726145058057,3.28491319222906,1.5593902929512,2.95913407695416,1.82995297799816,3.06706191171516,3.01746918734384,1.74434570262421,1.65732385015234,1.97916942216385,1.36137523564579,2.55561622103208,2.74817259434585\nEPB41L3 - mRNA,2.09646131360726,0.891349145029288,2.46991666832153,1.36832901780247,1.98948209500198,2.6231472426449,0.312147895756265,1.36397610578435,0.783357268484901,2.71821762106667,1.66359392673879,1.49142045000831,1.87316137408816,1.09981304891791,1.96726145058057,2.43691628567411,2.23746219806384,1.95913407695416,0.829952977998165,2.65202441243631,1.75443478151005,0.744345702624207,2.2422863508735,1.97916942216385,1.94633773636694,1.55561622103208,0.748172594345853\nPRKCB - mRNA,3.09646131360726,1.89134914502929,3.46991666832153,2.59072143913892,0.989482095001984,1.8861816484787,1.89711039647742,2.94893860650551,2.10528536337226,1.71821762106667,1.66359392673879,2.49142045000831,1.45812387480931,1.68477554963906,1.70422704474677,2.11498819078674,2.23746219806384,2.54409657767531,2.15188107288553,3.65202441243631,2.75443478151005,-0.255654297375793,2.2422863508735,2.30109751705121,1.36137523564579,1.55561622103208,2.74817259434585\nVAV3 b - mRNA,2.5818881407775,2.47631164575044,1.46991666832153,1.78336651708132,1.57444459572314,2.8861816484787,1.89711039647742,2.94893860650551,2.10528536337226,2.45518321523287,2.47094884879639,2.07638295072947,0.873161374088157,2.09981304891791,1.38229894985941,2.6999506915079,2.93790191620493,3.28106217184152,3.15188107288553,2.06706191171516,2.01746918734384,1.32930820334536,1.9792519450397,2.30109751705121,1.94633773636694,2.55561622103208,1.33313509506701\nMYH11 - mRNA,2.7745332187199,2.47631164575044,2.79184476320889,2.78336651708132,3.1594070964443,2.6231472426449,0.312147895756265,2.68590420067171,3.36831976920606,3.13325512034551,2.8335189281811,0.491420450008309,1.45812387480931,2.68477554963906,1.96726145058057,1.6999506915079,2.69689381670114,2.95913407695416,3.41491547871932,2.06706191171516,2.602431688065,2.55170062468181,1.9792519450397,2.78652434422145,2.36137523564579,2.55561622103208,1.74817259434585\nCLDN3 - mRNA,1.7745332187199,0.891349145029288,0.469916668321527,0.783366517081316,1.57444459572314,2.8861816484787,1.89711039647742,1.94893860650551,2.95328226992721,3.13325512034551,1.24855642745994,0.491420450008309,-0.126838625911843,1.09981304891791,1.96726145058057,2.6999506915079,2.04481712012145,3.28106217184152,2.15188107288553,2.87441683377276,2.23986160868029,2.32930820334536,1.2422863508735,0.979169422163848,0.946337736366942,2.55561622103208,2.33313509506701\nCA1 - mRNA,1.7745332187199,0.891349145029288,2.46991666832153,1.36832901780247,-0.0105179049980162,2.6231472426449,2.31214789575627,2.94893860650551,2.36831976920606,2.13325512034551,1.98552202162615,0.491420450008309,2.19508946897552,3.68477554963906,2.18965387191701,3.6999506915079,3.14435279367236,2.54409657767531,2.15188107288553,3.65202441243631,2.43250668662269,2.32930820334536,2.2422863508735,1.564131922885,1.36137523564579,1.55561622103208,2.55552751640346\nPRKCZ - mRNA,4.68142381432842,5.0612741464716,5.3278976634491,5.27521961341099,4.8963726906105,4.82478110381456,4.11950281781387,4.27086670139287,4.03128478192849,3.94061004240311,4.57048452234731,4.39831104561683,5.00244439103312,4.34774056236149,4.38229894985941,5.1593823101452,4.48538971150743,5.60299026672888,4.9174158192485,4.06706191171516,4.89193830525998,4.2037773212615,4.18088580620935,4.37148684494261,3.8849371917028,3.87754431591944,3.99610010778944\nAXIN2 - mRNA,6.32912207039754,4.89134914502929,5.27727159037913,4.91264953402628,4.94367840538886,5.05610664992101,4.31214789575627,4.36397610578435,4.30691922454191,4.30318012178782,4.1871558827958,3.66134545145062,4.57360109222925,4.42174114380527,4.87415204618908,4.57441980942404,4.5593902929512,4.54409657767531,4.87434709735662,4.82194941387863,4.9560686426797,5.13666312540297,4.90525136359593,4.06663226341419,4.21935623077336,4.01504783966937,4.74817259434585\nCLDN15 - mRNA,4.94445822016221,4.59178886317038,4.46991666832153,4.95329151852363,4.38179951778074,4.6935365705363,4.01258761389736,3.94893860650551,3.95328226992721,3.30318012178782,3.47094884879639,2.81334854489567,3.33259299272545,2.68477554963906,3.46976179110975,4.02187878639526,4.23746219806384,4.12905907839647,3.53039269613926,3.87441683377276,3.89193830525998,4.38820189239893,4.2422863508735,4.14909442360616,3.94633773636694,4.25605593917317,4.20760421298315\nAPC2 - mRNA,3.94445822016221,4.0612741464716,4.86223409110029,4.53825401924478,3.31141018988935,4.47114414919985,4.3996107370066,4.06441582392544,3.87082010973524,4.59268673898281,4.30745011651351,3.95085206864561,4.45812387480931,4.09981304891791,4.38229894985941,3.43691628567411,4.09544319319141,3.54409657767531,3.82995297799816,3.65202441243631,3.89193830525998,4.2037773212615,4.65732385015234,4.62302561193857,4.91596408732342,4.72554122247439,4.83563543559619\nHCLS1 - mRNA,4.52942072088337,4.35078076366659,4.55737950957187,4.64134751220889,4.31141018988935,3.76065076639484,3.77157951439356,4.17133102784195,4.2427888871222,3.59268673898281,4.36403364487988,4.39831104561683,3.19508946897552,4.95779404404548,3.90586090591642,4.36291570423033,4.19165850845072,4.54409657767531,4.41491547871932,4.23698691315747,4.23986160868029,3.4447854207653,4.2422863508735,4.14909442360616,4.49065825259075,4.36297114308968,3.99610010778944\nPDGFA - mRNA,3.47497293686099,4.47631164575044,4.92934828695882,5.39807636119652,4.84746309012956,5.05610664992101,4.89711039647742,4.17133102784195,5.30691922454191,5.13325512034551,4.91152144018237,4.95085206864561,4.87316137408816,4.90716797097551,4.63022646330299,5.02187878639526,5.61250162941077,6.12905907839647,5.28938459663546,5.42461391633324,5.43250668662269,5.2037773212615,5.0496412729311,4.83715041729142,4.97608507976099,4.87754431591944,4.70236890473273\nF11R - mRNA,5.55589293224456,5.28366656780805,4.92934828695882,4.78336651708132,4.74436959716545,5.30121914775754,4.11950281781387,4.17133102784195,4.03128478192849,5.45518321523287,5.94899614560104,5.13527663978303,5.36501447041783,5.59166614524758,5.61111764035529,4.02187878639526,4.40738719950615,3.95913407695416,4.53039269613926,4.65202441243631,4.47690080598114,3.99227321606779,3.65732385015234,2.564131922885,2.36137523564579,4.4625068166406,3.83563543559619\nEPB41 - mRNA,4.47497293686099,3.21327723991665,4.71784418176511,4.82776063643977,4.57444459572314,4.62314724264491,3.89711039647742,3.82340772442165,4.10528536337226,3.71821762106667,4.70798804609724,4.49142045000831,4.9606242153385,4.09981304891791,4.63022646330299,4.6999506915079,4.40738719950615,4.76648899901176,3.28938459663546,3.06706191171516,4.18739418878615,3.74434570262421,3.9792519450397,3.43860104080114,3.5313002370881,2.14057872175323,3.44861231248694\nITGA7 - mRNA,3.86199605997024,4.0612741464716,3.63984166976384,4.30692847313833,4.07694493625232,2.30121914775754,2.89711039647742,3.94893860650551,3.36831976920606,4.52557254312427,5.02114593135687,3.49142045000831,4.26547879686692,4.18727589016825,4.38229894985941,4.43691628567411,4.59501420268193,3.28106217184152,3.82995297799816,4.45937933449392,4.56178970356765,2.06627379751157,3.65732385015234,3.78652434422145,3.82080685428308,2.55561622103208,3.74817259434585\nMYL12A - mRNA,4.2339648373572,3.21327723991665,4.05487916904268,4.10529461196868,3.44891371363928,4.00165886589864,3.89711039647742,3.68590420067171,3.69024786409342,4.45518321523287,4.75105676798913,3.95085206864561,4.33259299272545,3.5592446675552,4.0827386680005,5.20245103203708,4.19165850845072,4.76648899901176,4.28938459663546,4.59062386777217,4.56178970356765,2.74434570262421,4.0496412729311,4.30109751705121,4.44883807689613,3.14057872175323,3.20760421298315\nBCL2 - mRNA,3.47497293686099,3.69870406708689,4.17035638646262,4.69025711268983,4.38179951778074,4.30121914775754,3.11950281781387,2.94893860650551,4.17567469126366,2.13325512034551,3.47094884879639,3.66134545145062,3.9606242153385,3.42174114380527,3.96726145058057,4.92234311284435,4.44691556369279,5.76648899901176,5.15188107288553,4.1545247529655,4.01746918734384,4.49923320478768,4.46467877220995,2.30109751705121,3.68330333053315,3.87754431591944,3.55552751640346\nITGA4 - mRNA,2.35949571944106,3.47631164575044,4.17035638646262,3.36832901780247,2.98948209500198,3.10857406981515,2.31214789575627,1.94893860650551,2.95328226992721,3.59268673898281,4.05591134951755,3.29877537206591,3.87316137408816,2.09981304891791,3.96726145058057,4.02187878639526,4.04481712012145,4.54409657767531,3.82995297799816,3.38899000660252,3.89193830525998,3.83180854387455,4.35776356829343,3.78652434422145,3.8849371917028,2.87754431591944,3.20760421298315\nJAM2 - mRNA,5.41838940849462,3.47631164575044,3.92934828695882,3.87082935833165,3.8963726906105,3.6231472426449,3.48207289719858,1.36397610578435,3.2427888871222,3.45518321523287,2.47094884879639,2.49142045000831,3.9606242153385,3.26973805036022,2.38229894985941,3.81542790892784,3.14435279367236,3.54409657767531,2.82995297799816,2.06706191171516,2.89193830525998,3.2037773212615,2.82724885159465,1.97916942216385,2.82080685428308,3.14057872175323,4.14049001712461\nLLGL2 - mRNA,3.47497293686099,1.89134914502929,0.469916668321527,2.78336651708132,3.57444459572314,3.6231472426449,2.63407599064363,3.17133102784196,2.10528536337226,1.71821762106667,2.98552202162615,3.29877537206591,3.33259299272545,2.90716797097551,3.28918954546793,2.92234311284435,2.5593902929512,3.28106217184152,2.41491547871932,3.87441683377276,3.51996952787302,2.06627379751157,2.46467877220995,3.14909442360616,3.16873015770339,3.36297114308968,3.20760421298315\nTCF7 - mRNA,3.5818881407775,0.891349145029288,3.63984166976384,3.48380623522241,3.68992181314308,4.10857406981515,3.48207289719858,4.06441582392544,3.5907121905425,3.13325512034551,2.66359392673879,4.07638295072947,4.04308637553047,4.09981304891791,3.38229894985941,4.11498819078674,3.822424698785,3.54409657767531,4.07788049144175,3.38899000660252,3.89193830525998,3.74434570262421,3.74478669140268,3.30109751705121,2.94633773636694,3.72554122247439,3.44861231248694\nCLDN7 - mRNA,2.35949571944106,4.0612741464716,3.63984166976384,3.69025711268983,3.44891371363928,4.38868198900788,2.63407599064363,3.68590420067171,2.36831976920606,1.71821762106667,2.98552202162615,2.49142045000831,3.57360109222925,3.09981304891791,3.46976179110975,3.43691628567411,3.822424698785,2.95913407695416,3.9174158192485,3.38899000660252,3.01746918734384,2.91427070406652,3.46467877220995,3.14909442360616,2.94633773636694,3.14057872175323,1.74817259434585\nCLDN8 - mRNA,2.5818881407775,1.89134914502929,3.63984166976384,3.10529461196868,1.98948209500198,3.10857406981515,3.11950281781387,3.17133102784196,3.2427888871222,2.45518321523287,2.98552202162615,3.29877537206591,2.19508946897552,2.42174114380527,2.96726145058057,2.11498819078674,3.48538971150743,4.12905907839647,3.53039269613926,3.23698691315747,3.23986160868029,3.83180854387455,2.65732385015234,2.97916942216385,2.16873015770339,3.36297114308968,2.33313509506701\nLYZ - mRNA,3.47497293686099,2.47631164575044,3.79184476320889,2.95329151852363,2.98948209500198,3.30121914775754,3.77157951439356,2.68590420067171,2.5907121905425,2.94061004240311,2.8335189281811,3.49142045000831,3.45812387480931,2.68477554963906,2.84173056849671,3.57441980942404,2.822424698785,3.95913407695416,3.15188107288553,4.06706191171516,3.23986160868029,3.06627379751157,2.82724885159465,2.564131922885,2.16873015770339,4.14057872175323,2.74817259434585\nGAPDH - mRNA,15.4912455240324,15.8528892763522,15.483719113362,15.3777786039036,15.4535810818434,15.4791058041063,15.1555292090763,15.3370007510736,15.3365965596845,16.0672223390944,16.1091423106891,15.8745509519407,15.8167292925741,15.8270155926117,15.8583806017333,15.8417917406893,15.8234076459972,16.0175063855133,16.2067736297478,16.2295909256732,16.1763418100104,15.823226181356,15.8407072341841,15.7505041320368,15.7183257628794,15.5929915787953,15.4734056971895\nACTG1 - mRNA,14.1302284284537,15.1939880688168,14.9193232833071,14.9142989952195,14.8641753358069,15.0183064643158,15.1557255007927,15.4963955049448,15.5646401318604,15.0816223519302,15.1436267413227,14.7947731199346,14.8356055629923,14.8399627220545,14.972944473583,15.2280791742291,15.2925539452209,15.3225844388977,15.2519191601467,15.4665735007808,15.479183096154,15.0915518075653,15.191626732607,15.2070485521961,15.2304629673429,15.0070849339794,14.9677571004844\nACTB - mRNA,13.4417556648224,14.834046907916,14.2343736319414,14.7690750626518,14.5981989490226,14.6584962224004,14.8418224575209,15.1003780371026,15.1106292052901,14.8569693863248,14.6817506292749,14.7876197185232,14.7424660324736,14.6821314746594,14.7040861496794,14.4924020942989,15.0532957654247,15.114031278961,14.9638176416793,15.1543175495755,15.0494574992894,14.6535356386665,14.6789978930107,14.6801509162101,14.6689318353235,14.6122539361453,14.4699126485589\nHSPA8 - mRNA,14.302071313216,14.7623880436888,14.5454793820623,14.4269409029601,14.5061054945754,14.4923554931957,14.791107065022,14.6026793852121,14.585469687071,14.1160704855354,14.1093847627869,14.3236182157939,14.2578776817375,14.463947703926,14.455896622886,14.867995959732,14.7492457311035,14.9473749531464,15.0341802185663,14.9620217401501,14.9058953534388,14.8179852129074,14.7885779546072,14.7911461165341,14.7275266662228,14.7485254401441,14.7154889287315\nCFL1 - mRNA,12.7979410618601,13.3764237087279,13.3045852260124,13.1068435502317,13.0584282746508,13.2288119471571,13.3902987034909,13.5966968137492,13.7432686671501,13.2779133631774,13.3999958580571,13.1746344049427,13.225480728935,13.257317888844,13.4001512817654,13.6212957718871,13.5081225327708,13.54362687393,13.5527605091677,13.7110308076374,13.7989655132488,13.1814475315326,13.3316264793645,13.5574241643736,13.6480773750413,13.4382592703939,13.2547281250972\nFLNA - mRNA,12.459939053506,13.3864546550371,13.3734219787454,13.662375331074,13.5633659256585,13.3850308550104,13.3842804999235,13.5321781120216,13.6111961363725,13.1044393927502,13.0448664824506,13.3213413438806,13.3630093345275,13.0501866839425,13.1301772240853,13.4677213402846,13.848256367117,13.5273277308042,13.134446907781,13.4655384131644,13.5022081820368,13.1604834712025,13.2302608751696,13.466507037371,13.5081029999946,13.3791824946774,13.3374726204985\nMYL12B - mRNA,12.1539115857912,13.2356450529451,12.9782066544619,12.9553251361584,12.97265385719,12.9698825709328,12.7911702427514,13.0288677750704,13.0355791814233,13.3500009785408,13.4089531256807,13.3125965062782,13.2670880497286,13.1005173179292,13.1466522380784,13.1234168128573,13.0573418978938,13.2754156087004,12.9497046901996,13.2877425327674,13.2479900549255,12.6923478336019,12.730628628406,13.2163793829189,13.179458055176,12.9818809757342,12.8053182112565\nMYH9 - mRNA,12.1490293644114,12.934718258987,12.8482115242334,13.0066140774988,13.0822392359218,12.849077653816,13.1909651803705,13.1950853811414,13.1807645128603,12.7461236176366,12.6720225488094,13.051036535821,13.0042144704978,12.7259783688224,12.7181079826818,13.0558523298363,13.2898604301354,13.3576105784034,13.0197775368782,13.1094052915088,13.0031925547937,13.0600613606464,13.0371083242047,13.0699434768046,12.9781537896459,12.9831909685972,12.9974342209457\nITGA1 - mRNA,11.3730510293325,13.4668883918638,14.0384666164827,13.8258809032215,13.919757430114,13.2323259807869,13.1201064980127,13.2854451663158,13.2744587654014,12.5777524074493,12.6270319250851,13.3809244134198,13.34648661598,12.460111475536,12.4433325515405,13.926964885157,13.7445754597715,13.6989146867274,12.3225572823848,12.8995563960513,12.8083606630412,12.5054813524529,12.4982984199742,13.1071637431402,12.9823968447701,12.6523313755206,12.8858041925813\nMYLK - mRNA,12.197649128867,12.9442567733426,13.0093178473496,13.2555496332709,13.1952753443991,12.9163889616011,12.3942969371101,12.3441156834235,12.3560574949722,12.3286273277482,12.1753465805062,12.6465549048661,12.6384474728118,12.7052925669796,12.6432184835223,13.382067456458,13.5270488631446,13.5065106404523,12.6738740292872,13.1960228625709,13.1137450984005,13.0775010529348,13.064724404221,12.6273022678165,12.5746313738937,12.5428802331046,12.8268237876711\nGNAI2 - mRNA,12.2721356055034,12.8146766304485,12.7190301210353,12.597348068696,12.6051117319701,12.5793781791948,12.9046049330243,13.0174929852058,13.0363183048786,12.2826847104023,12.3503132927606,12.3084040732637,12.3036139257537,12.3259241470503,12.3266449839972,12.8804814988524,12.841552059533,12.8820900688626,12.8808219412889,13.0232565408059,13.0325357205741,12.7191187705135,12.7612672800439,12.9903966775871,13.0083838740874,12.7990877594284,12.7320568865105\nLDHA - mRNA,13.8164489939457,13.5546848685076,13.1281281510733,13.2809689038648,13.3579885565097,12.6795140036694,13.8384025895849,13.8785668895301,13.7634075676785,13.8597926130706,13.717265312334,13.8532271831316,13.6960309099721,13.8738794044694,13.7574741706592,14.197303584985,14.2664473938589,14.120738380957,14.7989796801489,14.9632523856008,14.7965045089811,14.1896839284597,14.110594484163,13.9147028351146,13.7574447932857,13.5439458702632,13.5170805668243\nCCND1 - mRNA,13.0264593762163,13.8553289328446,13.3041901835562,13.2923992033878,13.3063292786045,13.2840345129475,13.4211623151952,13.3367764410088,13.3801293905359,14.7690865843574,14.8384416067501,14.2839057455328,14.268829383725,14.3922065845853,14.4269492068201,14.3210868047825,14.3284016140482,13.9056748808473,14.7676026568943,14.9235360740625,14.8948491905578,13.8687901626328,13.9180229062354,13.4009712826337,13.4225776426908,13.0280532210834,13.0771279561272\nCD44 - mRNA,12.1065697670816,14.2982184664524,13.4538009604862,13.8774442027532,13.4606492287264,14.0038247498678,13.9089200178073,14.3007060504654,14.5625636289065,14.3144073772111,14.4105280631111,13.9167675014411,14.0936908563795,14.0825393010999,14.3116960725046,13.5234506236711,13.9488065293276,14.1844921634087,14.5282235556902,14.3317977128453,14.4343546600285,13.7975928285366,14.0687686336791,13.9279928996145,14.1530468857256,13.8562547750258,13.5843215985913\nMYL9 - mRNA,12.8069233875864,14.0442677211659,14.0812873388972,13.9308099631518,14.098496514441,13.7955747512904,13.7694005067655,14.1314364197,14.1288318205634,13.8150425655207,13.8074554086066,14.0049013259874,13.8582917475551,13.5621875080009,13.5322048078845,14.1856961591497,14.2219251618251,14.1354630835718,13.4405164819232,14.0304999100614,13.9443828535138,13.3549092065492,13.3119598786803,14.0613184635177,13.961055666227,13.6505186337835,13.5760114622334\nITGB1 - mRNA,13.4726952537312,13.9525515520743,14.062023267539,14.1322329598379,13.7790157399723,13.6851933803654,13.5263152098498,13.7881423946025,14.0063030729839,13.8559559393555,14.0589272276437,13.7981941946512,14.1128344824187,13.7116447549025,13.9926894307327,14.3136634311664,14.2577476989085,14.3709098525307,14.2671848790376,14.1667384665748,14.3655979380753,13.6431360110054,13.9667291475233,13.9047238620374,14.1455499647775,13.5038009138627,13.5119036736531\nRPLP0 - mRNA,14.1154264605255,14.4123360635697,14.3130525794329,14.3715540979859,14.1465921411099,14.5932531332571,13.9779279240857,14.0788655400159,14.3072867121366,13.8005884619507,13.9715062292317,13.7323608132164,13.8999441464341,13.8307076954596,14.0908979043844,14.253819934704,14.3000143374406,14.7705087704641,14.4609594654101,14.3204662404893,14.4865597992885,14.1589648794538,14.4167959713552,14.0439963753447,14.2491670269874,14.1285532460494,13.9771415577486\nFN1 - mRNA,13.4518124100321,14.5492258051189,14.4602865912619,14.5029157711668,14.4703353936963,13.6815449200213,13.8153525052967,14.2527193546826,14.2630112406558,14.4459962943865,14.4172285455922,14.458736784394,14.4965856639581,13.555011818433,13.6096892665237,14.8772664048747,14.8903351734152,14.8752001278534,14.5905689811544,14.8679618116355,14.8599177501158,13.9623034004883,13.9934119342408,14.9914884208935,14.9632330566021,14.3103993578003,14.2487033001119\nCYFIP2 - mRNA,5.41838940849462,7.08117370390931,6.82746867293961,6.89189097385948,6.82237210916673,6.67625857910447,6.14503790992101,5.88753806184136,6.22630076433363,6.81975564752873,6.278303770854,6.75820699070321,6.86184606086032,6.94530309986228,7.15048327463634,6.26473531029143,6.09544319319141,6.00352819631261,5.41491547871932,6.02125822210203,6.04721653073789,6.15373663876191,5.88614254064822,6.06663226341419,6.32715952030787,6.36297114308968,6.33313509506701\nMYL12A b - mRNA,6.51600020512105,6.24890114964737,6.67937003395048,6.36832901780247,6.48133519133166,6.65877115237563,6.37823708621404,6.24661915514619,6.48379698662599,6.59268673898281,6.86326627157515,6.64116756951299,6.6146283604893,6.66966865724886,6.80856370456151,6.49002762213367,6.74525683826254,6.68705453151736,6.18750498261625,6.49332666641726,6.45487449965114,6.01113224331911,6.22717945848329,5.83715041729142,5.97608507976099,6.04746931736175,5.89791971385054\nNLK - mRNA,6.65717626808174,7.21327723991665,7.05487916904268,6.90230758980482,6.77084180852664,7.40974360453571,6.8667367474339,6.94893860650551,7.18423670476708,6.90143944512244,7.01232208096986,6.79520119818541,6.94997697113899,6.93270306308265,6.74862116410523,6.54125294548884,6.46628088855972,6.57384392106937,6.18750498261625,6.52649353035245,6.49859587708046,6.29893455430184,6.41221135231581,6.40543417686595,6.54128432566072,6.76506958666103,6.61853731392926\nLEF1 - mRNA,6.92428033822458,6.64623664719276,6.84495609966845,6.97319107596133,6.79683701705959,7.02913960232074,7.10656376210637,6.51372322528903,6.15839669983183,5.88814262250898,5.72968311719656,6.51378826303676,6.28255231022586,6.66966865724886,6.89009359005811,6.3244415564157,6.57731220094847,6.76648899901176,6.30568640896456,5.42461391633324,5.56178970356765,7.00173354531686,6.9702068054367,6.31901942504847,6.07562075331191,6.60001034039053,6.50306009650932\nCLDN23 - mRNA,7.54932027832107,6.21327723991665,7.05487916904268,6.80573433010977,6.83497214594636,6.58662136661979,6.14503790992101,6.00783229555908,5.71409460604779,5.34270848597446,5.81334104624347,6.34940144513588,6.45812387480931,6.12218086194636,6.35957887335933,6.41876893896385,6.28185631742229,6.54409657767531,5.9174158192485,5.92504290684273,5.69929322731759,6.78873982198266,6.91471169284499,5.50273137822086,5.49065825259075,6.22804156300357,5.74817259434585\nOCLN - mRNA,7.01293795804498,6.79823974063781,6.61966378782621,6.65373123666472,6.44891371363928,6.93057576783715,5.48207289719858,6.43006529624212,5.69024786409342,5.52557254312427,5.91152144018237,5.66134545145062,6.12108888753174,5.68477554963906,6.16365866338407,5.57441980942404,5.97892918446499,5.76648899901176,5.61131269152282,5.71091810148988,5.79005869124077,6.08419570550883,6.22717945848329,5.43860104080114,5.8849371917028,5.72554122247439,6.25596723454455\nMYL5 - mRNA,7.20916144635662,7.0612741464716,7.02450551999917,6.59072143913892,6.93199660034122,7.0825788612822,6.33451570878472,5.97868594989956,6.4415687512367,6.01589816970735,6.17138856693748,5.81334854489567,5.9606242153385,5.88117276244257,6.17671481620952,5.84290864534994,6.22614688483601,6.38539883165626,6.01977753687818,6.40691191459978,6.35136992389728,6.28350451373224,6.28668047023195,6.24595596285875,5.93123084397673,6.01504783966937,6.47609304890905\nINADL - mRNA,6.55589293224456,6.97881198627963,6.49228448134998,6.59072143913892,6.68992181314308,6.97364448972904,5.89711039647742,5.68590420067171,6.0501438091798,6.04014571595403,6.35009445392201,6.21934090457151,6.6146283604893,6.14420716827636,6.2024779122746,6.24427120773171,6.42728675694386,6.08841709389912,5.93847743477633,6.68177175583037,6.54103114340086,5.89409282212889,6.28668047023195,6.04525861262162,6.06181495378688,6.17032606514729,5.87745561129082\nTJP2 - mRNA,5.38924306283511,6.38320224135896,5.67937003395048,5.56472623060598,5.82237210916673,5.94507533753227,5.3996107370066,6.14533581930901,6.46985779566812,6.08745143073238,6.39151438130199,6.07638295072947,5.9606242153385,6.43966305180253,6.46976179110975,6.09226811428666,6.0188219115885,5.86602467256268,6.84118023342142,6.71091810148988,6.74538964190704,6.18728919847293,6.27203369426755,6.50273137822086,6.37260249106904,6.28353667559528,5.83563543559619\nARAF - mRNA,5.55589293224456,6.24890114964737,5.89618142302363,6.08714726525842,6.07694493625232,6.38868198900788,6.21903849136478,5.75629352856311,6.2590906994513,6.15562293337396,5.47094884879639,6.1918601681494,5.93925056454593,5.800252767059,6.11021940442261,5.97296918591432,6.20324648272593,5.95913407695416,6.27289647384689,6.27651527734411,6.17397367302383,6.2200791335906,6.2422863508735,6.22709693560743,6.21935623077336,6.55561622103208,6.19111609019458\nMYL12A c - mRNA,5.94445822016221,5.79823974063781,6.71784418176511,6.57778238343142,5.92021943256487,6.36730833821531,5.63407599064363,6.29471344334724,5.99281063411385,5.96614513451025,6.2929505468184,6.04600930168595,6.26547879686692,5.74366923869263,5.99700879397462,7.00980595409469,6.72931529439352,6.88987141451704,5.78414928838504,6.27651527734411,6.36324402418557,5.4447854207653,5.6796916631808,6.16899398104387,6.00523142542051,5.84101843989433,5.74817259434585\nMAGI1 - mRNA,4.7745332187199,6.84554545541616,5.89618142302363,6.57778238343142,6.57444459572314,6.00165886589863,5.52160126138522,5.6493783246466,5.66600031784674,6.13325512034551,5.52157492186636,6.29877537206591,6.08261473971711,5.62337500497492,5.3130362874223,6.09226811428666,6.68040569391257,6.31668608157224,6.28938459663546,6.63691752004611,6.49859587708046,5.78873982198266,5.53996689951418,5.59387926627906,5.70122523853041,5.76506958666103,5.63081564370769\nHSPA1A - mRNA,5.5024536732831,5.28366656780805,5.4241129787084,6.15840594842824,5.38179951778074,5.76065076639484,5.77157951439356,5.17133102784195,5.7833572684849,4.65681707640252,5.15544702306846,5.29877537206591,5.82735768447503,5.38521526778016,5.63022646330299,5.54125294548884,5.89567368081564,5.86602467256268,5.61131269152282,5.06706191171516,5.58225380612737,5.38820189239893,5.65732385015234,4.73405692432732,5.36137523564579,5.14057872175323,5.33313509506701\nAXIN1 - mRNA,5.35949571944106,5.79823974063781,5.46991666832153,5.933113636586,5.34703409962007,6.05610664992101,5.21903849136478,5.72152811040243,5.66600031784674,4.88814262250898,5.61779023712566,5.24630795217178,5.89552918711661,5.49213047169667,5.84173056849671,5.54125294548884,5.83737504025097,6.57384392106937,5.68793397312574,5.23698691315747,5.73628743479979,5.23619879895388,5.78660686709731,5.47102251849352,5.66515598382289,5.72554122247439,5.17443734904795\nBAIAP2 - mRNA,5.63251421384747,5.69870406708689,5.17035638646262,5.99281988271027,5.77084180852664,5.47114414919985,5.70446531853503,5.88753806184136,6.087138016662,5.805680462317,5.52157492186636,5.62070346695328,5.91755549344661,5.90716797097551,5.73985095447749,5.40039040964899,6.20324648272593,6.12905907839647,5.63730790005577,5.68177175583037,6.00236229495363,5.78873982198266,5.56421444576086,6.14909442360616,6.01958671839758,5.25605593917317,5.42059793631735\nRAC3 - mRNA,5.13208522333798,5.64623664719276,5.11377285809625,5.42722270685604,5.11876511194695,5.30121914775754,5.48207289719858,4.75629352856311,5.27521036481458,5.86117557490871,6.33601926871028,6.68124500888833,6.21301137697278,5.88117276244257,6.09654446752553,5.47254019540483,6.05764116047903,6.20706159039774,5.95923599494313,5.76750162985625,6.1190072138059,5.65123629823273,5.63460377365226,5.50273137822086,5.82080685428308,5.68489923797704,5.27173455040287\nRAC2 - mRNA,5.5818881407775,5.59178886317038,5.46991666832153,6.05015305777622,5.68992181314308,5.72748390245964,4.77157951439356,4.45143894703469,5.209622023187,5.22071796159585,5.89241261723467,5.81334854489567,5.80389871165104,6.36659958961281,6.62070368918449,5.63855014684376,5.99234970022731,6.08841709389912,6.22227040077693,5.68177175583037,6.04721653073789,5.57723571678895,6.10026734600107,5.90990675972673,5.91596408732342,5.50981253141895,5.27173455040287\nWASF1 - mRNA,7.66935098202784,7.4149111010863,7.5360058587793,7.63511555849737,7.57444459572314,7.53962388708262,7.54096658625215,7.48291717850786,7.55154159326183,7.55951987504761,7.53395864632219,7.5467028855095,7.66106393347959,7.72186486837428,7.68607969803651,7.36291570423033,7.17997670340308,7.31668608157224,7.27289647384689,7.50169013935188,7.27171047471963,7.19555681445654,7.25723669233947,6.81205943632859,6.80431873149451,7.04746931736175,7.20760421298315\nGUSB - mRNA,8.01293795804498,7.60559466269541,7.9617697646512,8.18850798021766,7.82868588309893,8.03592876798338,7.66969990037435,7.24661915514619,7.55814432808607,6.96614513451025,7.13119947682179,7.33691050095268,7.48049168783777,7.08849773569007,7.28918954546793,7.51586762706893,7.822424698785,7.7920240911189,7.22227040077693,7.27651527734411,7.33337349460343,7.63916346593215,7.81214195920445,6.89803265943844,7.31557154603266,7.25605593917317,7.39922428552478\nDNMT3A - mRNA,8.10545009683452,7.60559466269541,8.02450551999917,8.11428339519593,7.85984681458539,8.20208595573829,7.57893443645117,7.47250056256252,7.65372198806831,7.28300223985019,7.58245716401338,7.84897245462639,7.93925056454593,7.48351734139196,7.69063797999882,7.67723061500782,7.51358660333808,7.7920240911189,6.7724674833374,7.10048491325261,6.97166549773072,7.57723571678895,7.50281390109672,7.40543417686595,7.39479823718324,7.78443491152796,7.71395687900794\nAPC - mRNA,7.6448979383033,7.69870406708689,7.51431078767998,7.84406244876887,7.54407094667962,7.55860699045019,8.20089114465452,7.94893860650551,8.22630076433363,7.42787586923714,7.52157492186636,7.52484345154576,7.59426056279534,7.53444127655463,7.61111764035529,7.92876938200378,8.14435279367236,8.20706159039774,8.36522835461897,8.22188002076726,8.11198678612813,8.19555681445654,8.1126510704569,8.6157940427075,8.41394328644994,8.42598094061548,8.4276526938513\nLIMK2 - mRNA,7.54932027832107,8.28366656780805,8.16340362582085,8.17997129826317,7.92021943256487,8.24373365309678,7.59755011461851,7.97130641953396,8.21798549612163,7.75530693980189,8.16343981382199,7.69109279484467,8.00758769430908,7.69972589110504,7.95972777789516,7.74434481086635,7.88851388924277,7.48269603301117,7.63085287791847,8.00369985071773,7.98325347200593,7.31420131095515,7.51530484527991,8.0125924237013,8.29801317464836,7.68489923797704,7.67890993190874\nEPB41L1 - mRNA,7.21747671456863,7.82208648259217,7.79184476320889,7.9280247599132,7.74436959716545,7.80901378795624,7.54096658625215,7.64010051105859,7.7022205057595,7.94703631156255,8.17138856693748,8.2125196387155,8.29522614026097,7.85470055108138,8.09310538355876,7.50730561356551,7.74525683826254,7.8539518402621,7.47380916777289,7.53466746179816,7.69460153199286,7.63916346593215,7.73413944720317,7.72736227175331,7.60930274908937,7.64307906228242,7.56835155676104\nEXOC3 - mRNA,8.00335190921578,7.93574326438774,7.93752221840453,7.76636009177563,7.79683701705959,7.8861816484787,7.67847011000208,7.66775685396145,7.53824477064837,7.88144796993497,7.91625935918904,7.95085206864561,7.88996966177471,7.71452289303312,7.9859252948456,7.67723061500782,7.57285255275777,7.52898968528511,7.55787343256136,7.60622072282319,7.73628743479979,7.69271293420888,7.60569108173702,7.79295061338089,7.80016708822405,7.94793364381084,8.0055604370385\nTARBP2 - mRNA,7.54932027832107,7.57784967221251,7.5784411250997,7.5044657057885,7.46521552596838,7.20810974336606,8.10656376210637,8.000600726328,8.16706156095895,7.65681707640252,7.87304729236774,7.73934796345189,7.5526414735936,7.78631357610113,8.00068445211802,7.49002762213367,7.56389168518614,8.01441651245535,8.05395465219627,7.81525476130462,7.73628743479979,7.72162562612412,7.90049783362529,7.42211291801258,7.57082860127474,7.55561622103208,7.57472108163677\nDVL2 - mRNA,8.21332507129816,7.49126198721642,7.80083354643614,7.89189097385948,7.82237210916673,8.17773609432254,7.81994253595496,7.66775685396145,7.81126326505479,7.32307967922553,7.42514515918327,7.49142045000831,7.73114236921573,7.53444127655463,7.62547293333236,7.65414700189478,7.8186627799888,7.94781876372632,7.63085287791847,7.60622072282319,7.68517211907293,7.78873982198266,8.10440707636199,7.72736227175331,7.75798001682764,7.92193843527789,7.90804393112424\nCDK4 - mRNA,7.94445822016221,7.63281613143044,7.89618142302363,7.55155084185824,7.74436959716545,7.73584737539427,7.88200350408721,7.79860433342108,7.70816977209068,7.16667812188296,7.43838098633996,7.58945253296884,7.607870994314,7.48351734139196,7.73985095447749,7.98535291037015,7.83737504025097,8.14895863583418,8.14736559176303,7.9062656998121,7.83338612290487,8.01580873052858,8.17696010299555,7.57908226435098,7.67425819093014,7.78443491152796,7.69653982593053\nMAGI3 - mRNA,7.32912207039754,6.64623664719276,6.63984166976384,7.07798726597294,6.85984681458539,6.77695257872394,6.63407599064363,6.5734294714133,6.46985779566812,6.38118263378909,6.50908397768316,6.64116756951299,6.55966190127138,6.50920398505561,6.79168988599711,6.78741353275824,7.00564652284077,6.841777126316,7.05877166849405,6.92504290684273,6.7634235647373,7.31420131095515,7.32265976733752,6.81205943632859,6.71892724026387,6.85939696920918,7.0335748132081\nTCF7L1 - mRNA,7.29023305700394,7.10080251065824,7.41243117366077,7.75491007103209,7.48932798208519,7.59583989664917,7.35654201511472,7.20946615672873,7.40540908794128,6.34270848597446,6.53395864632219,7.07638295072947,7.19508946897552,6.90716797097551,7.01892357040306,6.86987569295021,7.29274463356503,7.52898968528511,6.88523541349935,6.8085288981163,6.97166549773072,7.15373663876191,7.14917694648202,6.92168392750309,7.24401828500763,7.31050372319554,7.18280082198258\nPIK3R1 - mRNA,7.13208522333798,7.26638857637621,7.66958901315789,7.44157799983311,7.44891371363928,7.49104370663756,7.04006835031946,7.02218758853615,7.00252578894706,6.57619861619424,6.74040952378962,6.55750964046608,6.83894565875024,6.81405856658403,6.53204606936409,7.27485952756513,7.12010524742568,7.20706159039774,6.58484048016163,6.86147777806526,6.68043420006627,6.81043489308198,6.85699619498871,6.73405692432732,6.93880406368153,6.72554122247439,6.89791971385054\nPAK4 - mRNA,7.22574433055223,6.84554545541616,6.91286016417026,7.22631001293004,7.23740960844557,7.323586960786,6.77157951439356,6.73901553713128,6.86017286553573,7.61707089760977,7.90199866606387,7.4102836872829,7.49521319354453,7.7722383908894,7.70422704474677,7.3723760334794,7.7212779753281,7.80462412789853,7.99987797944048,7.68911373117153,7.69460153199286,7.82649474397808,7.65732385015234,7.43038053399618,7.59499491240549,7.53289614453199,7.27173455040287\nXPO5 - mRNA,7.14085543296572,7.31761389973139,7.26433253467163,7.21799474471804,7.30236505028634,7.14670919870192,7.37823708621404,7.6493783246466,7.49071640056578,7.52557254312427,7.39151438130199,7.6309718024071,7.41232018519619,7.46613526316372,7.55222395130172,7.61483407786995,7.36158350989303,7.65957379509525,7.98477108705027,7.63691752004611,7.58732479567479,7.63916346593215,7.6516772870112,7.72736227175331,7.63283826355016,7.33697593455674,7.59992163576191\nHRAS - mRNA,6.54271754349682,6.84554545541616,6.89618142302363,6.72588102242056,6.90834533227658,6.93057576783715,7.44143091270123,7.51372322528903,7.42721345825962,7.34270848597446,7.35009445392201,7.34940144513588,7.32437248592049,7.41269600420226,7.59661807066018,7.47254019540483,7.33549428102437,7.8539518402621,8.09204782336834,7.35246413057741,7.53053876958321,7.66915820622999,7.7071723996029,7.33672142678193,7.22556138030007,7.43825927039392,7.02429699962009\nERBB2 - mRNA,7.25834899598416,7.5060589891445,6.89618142302363,7.82776063643977,7.44891371363928,7.72748390245964,6.91206073794339,6.9338317141153,7.17567469126366,7.43703586852261,7.60610843207803,7.25960477478524,7.64134569886508,7.1550954844191,7.39910723754596,6.80148871796996,7.52748104499646,6.841777126316,7.1060773832724,7.42461391633324,7.49859587708046,7.42382580212965,7.5702131863823,7.50273137822086,7.64214600577639,7.43825927039392,7.14905203062804\nPIK3CB - mRNA,6.81892733807835,7.78616690833723,7.27727159037913,7.5449177495258,7.16939118501692,7.63213602587216,7.73841265045836,7.85582920211403,7.80016555617146,7.13325512034551,6.98552202162615,7.13527663978303,7.00244439103312,7.37593745419214,7.35384250381018,7.34380688128263,7.40738719950615,7.33417350830108,7.80149653194894,7.82861314415964,7.7225255335553,7.72162562612412,7.61152016053922,7.47102251849352,7.47511740169498,7.42598094061548,7.27955405486216\nBAIAP2L1 - mRNA,6.44695856069139,7.84554545541616,7.38877990559612,7.28321240416452,7.14935343178037,7.22008238503214,6.06703539791973,6.6493783246466,6.71409460604779,7.25219619306902,7.60023186574136,7.25960477478524,7.22188952831923,7.14420716827636,7.18965387191701,7.79446829029219,7.65531471294974,7.64563460413738,7.33774761819686,7.59062386777217,7.78565351212077,6.7996281381254,6.80707096965702,6.86181247152569,7.20686528659016,6.72554122247439,6.52953230787051\nDVL1 - mRNA,6.46103374590312,6.99987360180746,6.99347862437854,7.0312940305249,6.88429985830993,6.58662136661979,7.02639341342239,6.47250056256252,7.02176200780998,7.23128720330604,7.30021854728244,7.25960477478524,7.33259299272545,7.28963760779792,7.35957887335933,7.17027062628793,7.46146387226195,7.7006010633553,7.52343993549749,7.42461391633324,7.44931497430924,7.2119512527072,7.41887508259682,7.38856035830155,7.28023847292038,7.29708320743322,7.10572459896394\nBAD - mRNA,6.96435777759992,6.79823974063781,6.55737950957187,7.08714726525842,6.74436959716545,6.71061008389524,7.13232685817145,6.75629352856311,7.50445645719209,7.26253813729048,7.32892984392396,7.34940144513588,7.58740689175428,7.12218086194636,7.60146747032157,6.77319967353854,7.2981581297514,7.97036133237741,7.55787343256136,6.71091810148988,7.34539602285265,6.74434570262421,7.2422863508735,6.67960914030494,7.10956808523525,7.14057872175323,6.79256671370431\nBHLHE40 - mRNA,5.90381623566487,6.56377448700078,6.4241129787084,6.29116115728001,6.61883871508159,5.345613267116,5.44143091270123,5.85582920211403,5.87082010973524,7.0990394050076,7.04729821921284,6.98327354633798,7.19508946897552,6.66966865724886,6.67691969875104,7.28491319222906,7.33549428102437,7.46692871715285,7.26458120563489,7.08942972474361,7.18739418878615,6.55170062468181,6.47750281256753,7.77358528851395,7.68330333053315,6.91316822565016,7.09690074857693\nMET - mRNA,7.11438322160452,6.8686290685292,6.17035638646262,6.25909994804771,6.05557128545976,5.8861816484787,6.24288523331915,6.38634391881281,6.087138016662,7.50829455169243,7.54011087330379,6.96715388097471,6.89552918711661,7.05400935930478,7.13049179944887,7.41876893896385,7.50424873875874,7.28106217184152,8.16534333269209,7.9062656998121,7.65650836082079,7.38820189239893,7.51530484527991,7.15907851217878,7.14273494917045,6.36297114308968,6.43467312152907\nMYC - mRNA,6.7745332187199,6.95743833548706,6.65974122720155,6.27521961341099,6.48133519133166,6.47114414919985,6.3996107370066,6.79024086048645,6.81678027002235,7.01589816970735,7.06447336302097,6.57888329125865,6.62804887625163,6.84128003531905,6.76600324233346,7.25453954318554,7.36674521500881,7.99255707849161,8.46657759854181,7.71811360289409,7.64196005225164,7.49923320478768,7.59396178915491,7.35420885351077,7.50092658804458,7.12547182936302,7.14905203062804\nDIAPH3 - mRNA,6.81892733807835,6.64623664719276,6.65974122720155,6.65373123666472,6.70372761266811,7.13410916192228,6.99864842293948,6.77336704192205,7.22630076433363,6.76261174042512,7.278303770854,6.75820699070321,6.55966190127138,6.7722383908894,7.02615513963413,7.55793168663547,7.49485004075649,7.60299026672888,8.48458900652613,7.64449073975091,7.9521429394659,7.883897055023,8.17696010299555,6.63738090491564,7.15579110199589,7.2421167482153,7.06105554963021\nCCND2 - mRNA,7.82438176817046,9.62266817605435,9.17035638646262,9.15840594842824,9.13414033783387,9.23490980270978,7.46189501526095,7.83158165586735,7.7833572684849,6.9534340827607,6.93038046743369,6.59994490678648,6.94997697113899,7.0186762861925,7.27104219875767,8.3581621742597,8.70302860287324,8.7006010633553,9.01234733140269,8.96490736772067,8.98709553830032,8.6661866396987,8.73413944720317,9.04256450345236,9.11626273780925,8.68489923797704,8.95518691452338\nPDCD4 - mRNA,9.76605506479559,9.02577546525021,9.05863130390379,8.94575784583822,9.19649641517952,9.28990383452971,8.72577582478044,9.07822162345047,9.07337611541752,7.02807288365345,7.13119947682179,7.07638295072947,7.01271272648695,7.09981304891791,7.2024779122746,8.45483819367137,8.46387439085263,8.76003397687446,7.99987797944048,8.53058628498634,8.42119137339485,8.51253002740113,8.71260628565353,8.43038053399618,8.50348229294834,8.28353667559528,8.37388143741032\nPIK3R3 - mRNA,9.48189235080078,9.36302435942133,8.84929503539279,8.7491508017434,8.82237210916673,9.57734355303178,8.95960632221118,9.90700792603959,9.83592531928905,7.95980360763642,7.90676791021174,7.57888329125865,7.55966190127138,8.00067985689866,7.77890373104127,7.94787820495149,7.70913741245589,7.76648899901176,7.97970009750285,8.3431863169894,8.15702053974264,8.4447854207653,8.53690709976513,8.97634890310147,8.91979594891445,9.05546210811528,8.71683938754106\nWASL - mRNA,8.73742922405716,7.17675136389154,7.59919968526649,7.40541833653769,7.44069320683431,6.97364448972904,8.97748381294144,8.73466351259157,8.85213554647031,8.80921215328726,8.92333719042957,8.95494482327949,9.03303271086655,8.93901683701485,8.98777846792108,8.97607509678214,8.92396272524706,9.07287624300335,8.69413912265244,8.76402943794944,8.66132537711857,8.94401804746057,8.92176645037894,9.09551338340132,9.02137112807576,9.12547182936303,8.98896392650781\nTGFB2 - mRNA,8.2502666496863,8.10080251065824,8.12096835950046,8.07338536401393,8.10842316772549,7.76882469784054,8.59291866588687,8.7691175689207,8.7022205057595,8.93091664619927,8.97874348899509,9.24964366473503,9.44301698241911,8.88117276244257,8.99332374716676,9.39111259606098,9.27363581061733,9.18795276745004,8.41115355992312,8.71811360289409,8.77235668950731,8.27962107924501,8.30118003992707,8.51444479878465,8.61640380546452,8.33697593455674,8.3180282026768\nHSPA1B - mRNA,8.85668226007377,7.24890114964737,7.80083354643614,7.69625585331128,7.56691092303773,7.47114414919985,8.60216674268888,8.07822162345047,8.32638908874014,8.75896396340998,9.03428133354601,8.90081138614601,9.17007758096745,8.94530309986228,9.32920522431581,8.598803968051,8.65320396635393,8.32545629119997,8.16086985611278,8.29588060221104,8.5012849646081,8.2119512527072,8.4354009796877,8.01808841145615,8.10956808523525,8.2421167482153,8.40995069211784\nVAV2 - mRNA,7.91408457111869,8.15813568572419,7.98561650660557,8.25097206716431,8.25626863569689,8.26122107982562,8.44657421597719,8.25271935468261,8.3108342745453,8.12760855720437,8.0299161409846,8.33691050095268,8.32024460029781,8.2135552149671,8.2679953231988,8.20245103203708,8.22898404413954,7.94781876372632,8.46657759854181,8.34783268184576,8.236637707806,8.35905554673941,8.27570935241095,8.34985682897107,8.36137523564579,8.48042872463786,8.46584901741225\nAGO2 - mRNA,7.84062240917767,7.93574326438774,7.86223409110029,8.20121903196721,8.08223923592184,8.16540529241182,8.17633404041054,8.44079170283518,8.50787112160485,8.46417199846013,8.40506091313993,8.34316949142437,8.75885774742755,8.31898156938007,8.40743851213792,7.97917433544103,8.21474212156376,8.59575869749781,9.05395465219627,8.03284619637724,8.05455850607906,8.11066791687002,8.4549853760061,8.25994019229445,8.45677225843834,8.14057872175323,8.1533140574822\nMLLT4 - mRNA,8.98885233952067,8.24007729926037,8.51976521777209,8.466361100763,8.42827394758024,8.323586960786,8.54096658625215,8.19686611994909,8.14530104222014,8.46864547503943,8.33247891100504,8.58945253296884,8.62470543317726,8.64284486917314,8.71321582797403,8.27485952756513,8.23746219806384,8.05716615991468,8.19627519224398,8.36626993010244,8.07996511307761,8.15797363164838,8.26834864745969,8.0235635415223,7.91979594891445,8.35651612095238,8.34062963161393\nJUN - mRNA,9.55096625116463,8.10080251065824,7.88776918320743,8.27120655090437,7.90237143123195,8.06940347253447,7.88957672379201,8.15839197213446,8.33025172837254,8.44159415048492,8.5553776299571,8.4102836872829,8.54911840702991,8.2135552149671,8.27711671316735,7.9667372322028,8.31694698189066,8.48269603301117,8.89873125598358,8.19634492866012,8.36028464870503,8.09307385685528,8.48069109019858,8.20798811265973,8.36980385771637,8.10251068091971,8.11886000115307\nRRAS - mRNA,7.78015776791378,8.24890114964737,8.30912045641847,8.6035454794965,8.07694493625232,8.09563501410765,7.45169924815506,6.97868594989956,7.39806711260011,8.7950332181175,8.98100654050366,8.76754485528255,9.09474849535296,8.96709178862757,9.01165556993902,8.49436655785801,8.76102415412085,9.13904316696909,8.51294756167985,8.59453891777555,8.60992622461192,8.09749252812229,8.45822375007265,8.32346533007966,8.64446358866979,8.35651612095238,8.10572459896394\nBCAR1 - mRNA,6.65717626808174,8.05122048180768,7.65974122720155,8.43800254560928,8.08751417796251,8.24958637934222,7.87439031997734,7.85582920211403,8.16706156095895,7.8336948384866,8.03863335808571,8.36178516959171,8.43158208735682,8.12218086194636,8.27408265307772,7.86987569295021,8.70913741245589,8.71402157911763,8.54075941169752,8.24198759421584,8.29978542633235,8.01580873052858,8.21956627437341,8.00707541873373,7.97608507976099,8.06341086123077,8.09690074857693\nPIK3R2 - mRNA,7.69934572232568,7.8686290685292,7.92112778015386,8.34178723034998,7.9667620185019,8.19603691106549,7.2546624010955,7.24661915514619,7.40540908794128,8.52127240569065,8.64373350437794,8.33691050095268,8.55966190127138,8.11662133660446,8.4724113695237,7.87653942323122,8.61467272845239,8.7006010633553,8.6178555373896,8.17034972012718,8.42402853269838,8.33680273989229,8.51530484527991,7.9959777098504,8.20372557905959,8.2421167482153,7.9913465778188\nPIK3CD - mRNA,7.200797973422,8.38320224135896,7.87079610460371,7.9682418599896,8.03387621436044,8.11500033897458,7.73000041064216,7.60238084510943,7.63510630990096,8.28300223985019,8.37783944440491,8.38016369890657,8.34889480505456,8.76869803318415,8.6792151567387,8.00980595409469,8.12924590128215,8.55904691914128,8.57814582758762,8.15981905263501,8.25587392666892,8.17897393026093,7.99717385303697,7.90990675972673,7.93502242313911,8.19224084157572,8.05195334252296\nABI2 - mRNA,8.06915396761153,8.10080251065824,8.30912045641847,8.38327935926844,8.1594070964443,8.323586960786,8.11304779567657,8.12552733822883,8.33410405386814,7.66463658086182,7.95821467563041,7.78604119889994,8.16778212297978,8.00067985689866,8.18965387191701,8.19713723214062,8.2402772136709,8.10888119645884,7.77832020958284,8.08387019940171,8.12947421285697,7.96351422308637,7.98824072826696,7.31901942504847,7.49065825259075,7.29708320743322,7.35550290809546\nTCF7L2 - mRNA,8.88566888895461,8.64623664719276,8.65974122720155,8.73756282746819,8.66543912794373,8.87107475608849,8.2546624010955,8.33551965973512,8.43799329701287,7.36207381084139,7.67482118216204,7.67132954002324,7.56664833158748,7.91359424013494,8.00435076931578,8.25453954318554,8.36931915867263,8.05716615991468,8.14736559176303,8.10048491325261,8.25587392666892,8.38097032316786,8.36813028385169,7.38004885844603,7.48031630836929,7.323800545809,7.66703583162045\nIGF1R - mRNA,8.89087717995737,8.03090049742808,8.76912468670881,8.67515022029963,8.49727673520068,8.41496131380673,8.18866484232126,8.32397803785243,8.71409460604779,8.36687479710521,8.65227861351095,8.4572047346704,8.77402818206891,8.03645098792048,8.24027994498698,8.65802001104198,8.62763115426402,8.33417350830108,7.71259602736001,7.77442104379604,8.05821552968715,7.68686020796345,8.11675546878964,7.76707198155528,7.91979594891445,7.63243181808291,7.80886852603341\nSRC - mRNA,8.47497293686099,7.92477214656674,8.224804170485,8.22215836965958,8.05557128545976,8.14042293585449,8.19479094511811,8.35266079255652,8.48032479471919,7.72571215761359,7.85835078116104,7.97523622727257,8.30778960172488,7.79330000641723,8.2024779122746,8.24427120773171,8.2818563174223,8.18795276745004,7.74284231422813,8.05005548640947,8.12947421285697,7.93910255704646,8.05820328643452,8.02901797161441,8.11959845037251,7.9985597168808,7.92808168436079\nCSNK2A2 - mRNA,8.12326137295098,8.38320224135896,8.18416218598765,8.33026097696895,8.3203989731166,8.42534045958673,8.06703539791973,8.08507529449153,8.12320727136952,7.98500416176157,7.86326627157515,8.08387748727639,8.05803671699644,7.98245609827975,8.03335064103834,7.9667372322028,7.95856138677103,7.87799731422875,7.65013194041335,7.93124805636944,7.69929322731759,8.08419570550883,7.82724885159465,8.12891654166853,8.01601126417375,8.19224084157572,8.14905203062804\nFYN - mRNA,8.33677564294097,8.32597737266601,8.35865991721979,8.33026097696895,8.20865061546415,8.52522082195565,8.09350760928093,7.91856495746199,8.22630076433363,7.53413455662769,8.02114593135687,7.89229988629049,8.12582680653841,8.16590223937568,8.31893688886198,8.06335542237142,8.0671849331499,8.06765853373233,8.15638246512047,8.09496790828504,8.12250465804213,7.99227321606779,8.11675546878964,8.04525861262162,8.20686528659016,8.10251068091971,8.07010068923321\nMAPK10 - mRNA,9.75895167752104,8.54240083620822,9.20462628854736,9.18211020901951,9.36452152634891,8.83260060827386,8.18866484232126,8.05746306328368,8.13208542271598,7.53413455662769,7.61196115832347,8.24630795217178,8.13994791478306,8.09416648577676,8.09310538355876,8.82234732286763,8.71522046450773,8.841777126316,7.57814582758763,7.82194941387863,7.92435978295236,8.6300420759636,8.59690306446703,8.32346533007966,8.15903676149955,8.29032584125791,8.80616431710503\nMRAS - mRNA,7.86199605997024,8.64623664719276,8.83186044205677,8.8112725136512,8.82868588309893,8.39925123071807,8.16389693717232,8.73466351259157,8.85213554647031,8.04614445657547,7.91152144018237,8.03831490989594,8.32024460029781,7.40359379709501,7.63968679255206,8.78741353275824,8.88671837558116,8.77291526817119,8.11997182493078,8.8118957492147,8.68989452931534,7.90925262929989,8.1411396274166,8.65159476413534,8.71452206114387,8.29032584125791,8.33313509506701\nASH1L - mRNA,9.1923857336058,8.12016783552517,8.37078347630228,8.45225150134756,8.30689470876685,8.49597600217979,8.88200350408721,8.73466351259157,8.73172450006958,7.90143944512244,7.89721360349849,8.09875076375792,8.11156611341324,8.24447129174979,8.34519495519667,8.76603988196567,8.83365195420825,8.72068530939864,8.65013194041335,8.6178086970984,8.64196005225164,8.74152518356183,8.8569961949887,8.68307299560851,8.80845846185544,8.71043433008418,8.69068709968509\nMAPK7 - mRNA,8.68742255494986,8.35078076366658,8.54673226537236,8.51468554810638,8.61519093806645,8.43050216470251,8.15763794670064,7.87967594406839,8.05482029638928,8.52985990152737,8.66077340767641,8.57356949136218,8.43158208735682,8.48351734139196,8.48558675827143,8.31955933503595,8.57508410005635,8.36852501309186,8.39600701616926,8.17034972012718,8.22040924601412,8.41323068689045,8.42881331965294,8.47502444905102,8.4301535136312,8.43213316759708,8.57472108163677\nBRAF - mRNA,8.49904707183985,8.83971637661397,8.69873535881741,8.78336651708132,8.80646571825736,8.84037795886557,8.22503723198623,8.18415506819954,8.10528536337226,8.46864547503943,8.47094884879639,8.49704499920219,8.62135422367762,8.44854120314898,8.39631942017434,8.35339293011182,8.52748104499646,8.61018576813309,8.27703620420782,8.38899000660252,8.36028464870503,8.48917954012375,8.32620883441859,8.51832823327188,8.52879338147752,8.48635355859496,8.34062963161393\nPPP2R2D - mRNA,8.62002326966427,8.45359156925036,8.75993551525415,8.6035454794965,8.60786759726059,8.75243025958987,8.3889634928071,8.21572514720041,8.27521036481457,8.2728064727443,8.28930276980325,8.50822873769486,8.48418617139551,8.25463115797001,8.41296608610635,8.77676628855873,8.67209042570057,8.97594236464071,8.86062011424511,8.46366669289702,8.60743236912336,8.56452466503939,8.78144516198153,8.31008630027847,8.57082860127474,8.66935838708127,8.62468954091085\nNUMA1 - mRNA,8.67237867472541,8.5060589891445,8.25127638184619,8.76920845408466,8.71058128370917,8.81297180152492,8.59755011461851,8.36397610578435,8.37581430575298,8.10479867429628,7.9719329568782,8.44561676039518,8.36901640097533,8.34298703239086,8.51672527008034,8.08653174473752,8.75513158619765,8.44294985421841,8.38069976338141,8.50169013935188,8.28112962705202,8.54524560254451,8.53384079671734,8.60122124162022,8.52376656440269,8.77961789523018,8.71106859968311\nTBP - mRNA,8.6510501652849,8.4986794587789,8.54673226537236,8.4907256491622,8.50518193328603,8.37803474480837,8.45169924815506,8.22816225043863,8.31473872900121,8.00361983992891,8.10653742258752,8.29877537206591,8.15856359295041,8.30926641454686,8.23404799127547,8.6028282246098,8.46868337877503,8.29898407983878,8.34958923084138,8.48912667788797,8.54103114340085,8.402557185376,8.48387233744326,8.49092207593123,8.3068190720237,8.46850555726204,8.6490394023266\nSMAD2 - mRNA,9.92046536454041,9.99725765360045,10.0790954064635,10.0536618435534,9.95670835383798,10.1792700604861,8.83962490181666,8.68590420067171,8.59392890322605,9.67435173569503,9.7038836477645,9.73221178217027,9.78005196969668,9.68477554963906,9.84582332313059,9.95103854584482,9.74525683826254,9.7697057116953,9.46657759854181,9.76750162985625,9.69694928684929,9.50090202514829,9.59690306446703,9.36933837836403,9.36419025125284,9.36618785577322,9.4709801255154\nDICER1 - mRNA,10.0459962466243,10.0934729688597,10.0283373815902,9.92035762916155,9.88278362586255,9.84037795886557,8.71302733203845,8.80276795836261,8.92801551131678,9.63708085834126,9.8171459584469,9.41623295361409,9.62971769661224,9.57959331294701,9.77246790605959,9.87154451331083,9.76689275220999,10.0884170938991,10.3671713785368,9.83193350245125,9.85877144132478,9.92674005602874,10.1157304633889,10.1726947826651,10.2607321585689,9.83870457405608,9.96006088889186\nLIMK1 - mRNA,9.5176846128324,9.99463695344131,9.8835445973457,9.88402885608651,9.64232706800396,9.98245755953535,8.94877251629991,9.24661915514619,9.18210096042309,9.70879436718004,9.83727106304221,9.69843477018584,9.82590262127465,9.47702357930646,9.51800823596381,9.77854629500402,9.62547948340898,10.0754780381916,9.99108486098247,9.65390169967698,9.73742835620436,9.6300420759636,9.7180197818399,9.8199473457589,9.80639008151421,9.52140050569416,9.56675477182671\nPARD3 - mRNA,10.1495726500668,10.2177786321516,10.2046262885474,10.1309878856494,10.2737278448317,10.479883998764,10.0736991282007,10.0042210420067,9.83320581793546,9.77169103433598,9.68596173976724,9.84897245462639,9.7123651621851,10.0984034786632,9.89996833799322,9.77498408321672,9.75906263778757,9.56646439070377,9.63569685014978,9.7359468959814,9.51864291164999,10.0052652362871,9.89692237940146,9.64094751993584,9.4408600194726,9.82707924893645,9.85014826529508\nCREBBP - mRNA,9.85801254605174,9.9708339288561,9.80083354643615,10.0536618435534,10.0769449362523,10.0085782798384,9.85324451110579,9.77971787407444,9.75777185829043,9.40471814824988,9.3092523591475,9.57622883781267,9.54558671605965,9.5592446675552,9.49734260002112,9.58259374086974,9.77662100917187,9.78250131700039,9.50237831996966,9.68544741397376,9.55921115946588,9.85287015940238,9.81340692644102,9.65159476413534,9.56594637989499,9.72554122247439,9.86970611168588\nPRKCA - mRNA,9.79828757201932,10.2245044953399,10.0051920449423,9.99525481162732,9.99229711060904,9.91776799153653,9.58361092366064,9.71932720220916,9.66447122916,9.38355353825184,9.60464153307937,9.6155417618375,9.55088101572917,9.54482789478633,9.40882239237918,9.96361113121608,9.79109149198021,9.91333038734103,9.99360265401399,9.85169675727268,9.80537174673527,10.0181413018385,10.1188032974385,9.81679735533525,9.8350809852652,9.73801057443661,9.78846231537157\nRAB8B - mRNA,9.62472005636567,10.2311991479139,10.0492326059015,10.115403065443,10.1392292145067,10.2407983620722,9.64082282308421,9.62607095115453,9.69325035225494,10.0326120432686,10.0156373525342,9.99921509020701,10.0101524861684,9.94059097251296,9.99424589167939,9.57031541109131,9.67104763921434,10.0305964395108,9.91210201935204,9.86796181163546,9.66971664737771,9.99582811324442,9.94734269708496,9.97352285902271,9.92933131106125,9.94793364381084,9.86711366706936\nCTBP1 - mRNA,9.78994827110659,9.95204507671684,9.46709614925915,9.93184309925959,9.89486910002011,10.0274373070897,9.8219229000832,9.65399495271697,9.63666682388858,9.91133224988087,9.92451346040169,10.055569939994,10.1175252092087,9.95623857754344,9.98221179204654,9.50945088539706,9.96538265262704,10.0676585337323,10.0430571976401,9.86796181163546,9.76901324643294,10.0401146370447,9.98712018837304,9.77033231071887,9.82387750824305,9.99855971688081,10.0172992734953\nMAPK3 - mRNA,9.73887408651232,9.74309818644535,9.76683287520082,9.65064525679098,9.75601100360085,9.69138810395773,9.98280414487471,9.98968494884882,10.1265429839328,9.96139160453962,10.0666059503138,9.95289989729446,9.87456956848097,10.0452568852958,10.0016019052751,9.66188265067438,9.82804924797888,9.53278126444748,9.51119138977597,9.85333153936362,9.90316556068324,9.59765525802788,9.73547465788699,9.94639568099984,9.85222611238608,9.85482423941936,9.75660121641643\nPPP2R1B - mRNA,9.96559243325155,9.96816474208012,9.94767493476542,9.88271432748451,9.76262130172167,9.88430191526048,10.1386963830472,10.1809597290397,10.2841991480418,9.56996666248272,9.69976753929227,9.67877252320881,9.75733189319659,9.59766488586902,9.74533858011592,9.66573497616999,9.70812107212439,9.78568256424507,9.9319286489474,9.67624064985714,9.69929322731759,9.71876029242973,9.83723294016728,9.63738090491564,9.75154419184597,9.76263054120961,9.68186324929809\nMAPK9 - mRNA,9.63871936337418,10.0537404737862,10.0454559151561,9.9432378538597,9.948034810433,10.04437054187,10.1307300732371,10.0765031062242,10.1916870092523,9.72009490830734,9.72158564949796,9.76521604922257,9.76190462298642,9.8174894719843,9.70084175956213,9.94945924077097,9.82148513832138,9.72068530939864,9.73834559877191,9.96792871969591,9.90519752586547,9.76671351565266,9.72744479462916,9.99318989247878,9.88982934641058,9.8954662239167,9.85538966993724\nPRKCI - mRNA,9.81756050231445,10.1321404771912,10.1334747725388,10.0277303522018,9.9349259313799,10.0137461481974,9.81795944967586,10.0609436320186,10.1420084693778,10.1472755906604,10.2075923768733,9.89229988629049,9.89136155290138,9.79155656808918,10.0098328343322,9.94312467498085,9.87408681860749,9.83257818946953,9.50943307750361,9.92659669809781,9.98997038822051,9.68978953900212,9.69349746270583,8.73739263689057,8.81052388102122,8.54996965789093,8.9576259599748\nEGFR - mRNA,9.48877873638602,9.69224904494959,9.59919968526649,9.798781569468,9.65481801218716,9.63437449806816,10.0143205811218,9.63076264647925,9.71261567712187,9.72196975592777,9.67202254880937,9.70087381563726,9.97776012765253,9.92318028896414,9.95215455819036,9.69241701882249,9.80161168804957,9.98150188998261,10.3739514281327,9.81860597080426,9.74198304046379,9.98394423187333,10.1259478930704,9.06397780996821,9.0070336680545,9.09089159765288,9.39563102080077\nAKT3 - mRNA,10.2492531652464,10.5549072492466,10.5184035423139,10.6249308648119,10.6270126465273,10.3926545340811,10.5409665862521,10.4958330663931,10.3664400359878,10.1625423473138,10.163439813822,10.7057395708091,10.6163127682007,10.4318495972796,10.3394009914217,10.4965311419713,10.4794453477582,10.6699405106535,10.1552584330878,10.2457267627216,10.0508921888813,10.2835045137322,10.1935710658405,9.93917135423193,9.80224440325666,9.89991212894789,10.1177699406245\nPAK2 - mRNA,9.97665704255036,10.757082415881,10.4328126736588,10.5382540192448,10.52864090611,10.4661260744332,10.5707139296462,10.497118318185,10.4433531609149,9.71445570227047,9.69839288931606,10.09689996807,10.0114331742604,10.0036948946541,10.0061804398729,10.3652866086931,10.4883527337872,10.3166860815722,10.4819015887216,10.4091365797143,10.3735542929633,10.4282175712468,10.3707103650894,9.78652434422145,9.75369265842455,9.82240276172698,10.0404942271479\nDOCK1 - mRNA,10.7836620082592,10.5441941180313,10.8686603602597,10.5931346457917,10.6503693653046,10.6795140036694,10.4593528206985,10.442126913519,10.5531951121143,9.89147833507223,10.1230255453761,10.049841163277,10.1258268065384,10.0998130489179,10.188042822011,10.5349483686346,10.4847963760926,10.3958456190914,10.2934773512693,10.335018996118,10.3423997703927,10.3809703231679,10.4574148377773,10.0479477001493,10.1345144423655,10.1063630064153,10.1628578301531\nVAPA - mRNA,9.93818289473572,9.92201628127623,9.86652144950339,9.7037193724964,9.8963726906105,9.64994730198862,10.191731145369,10.0884899589043,10.1086627235746,10.1500634080321,10.163439813822,10.2529716824528,10.1127599033373,10.2908722634496,10.2293562959507,10.1387425440862,10.0821679624758,10.2878090042821,10.2201219341983,10.1885954290552,10.2309784877243,10.22614513429,10.1147047292249,10.1603216790294,9.98067819106147,10.138698988535,10.1575635304836\nJAM3 - mRNA,9.79551315762411,9.8600159382245,10.0883021705801,10.0583269892219,10.0447645305032,9.96833068983257,10.0918672508997,9.89535756630066,9.95328226992721,10.309428269453,10.2229710172655,10.5303394393006,10.5134063103105,10.3465536474111,10.4177854011516,10.2811512734329,10.4309875585651,10.4328398265736,9.9319286489474,10.0400416977814,10.0490555304017,10.0216331027546,10.0614009863936,10.4241842680323,10.4227464283056,10.3369759345567,10.3820760036944\nZEB1 - mRNA,9.55096625116463,9.61244833373647,10.1191728458388,10.0442860507442,9.74603841752607,9.81099415208448,9.89711039647742,9.6516884853338,9.91778358870583,9.81800374076714,10.0353705710767,9.97120071403741,10.4730742162753,9.89585265774768,10.1864299710427,10.3305211905324,10.345986654842,10.4866110830146,9.93586148656932,9.68544741397376,9.95900592575925,9.8710501754674,10.0929941110889,9.77520903099362,9.95102638216057,9.78202841382086,9.93923180887751\nITGAV - mRNA,9.6121611518913,9.19055716341657,9.55737950957187,9.77347048093882,9.61701597947478,9.30684369695142,9.92501639304731,9.6071500892573,9.60672450853114,9.77891355275422,9.7747295969735,10.4191984120907,10.3448365884802,10.1578047716771,10.1521367934889,10.149787153364,10.3960718862783,10.5497211268692,9.75179391507266,9.7830239019703,9.69812173310714,9.89027784844472,9.99047920046296,9.56225218966678,9.48161311298775,9.43825927039392,9.76498088203241\nRRAS2 - mRNA,9.41838940849462,10.2772115456707,10.2659562771513,10.1842459533635,9.94658413656427,10.1325263915596,9.78382311014831,10.1054430921855,10.2190275294215,9.27536217764806,9.30021854728244,9.49984907207889,9.54735364223384,9.35955631260869,9.38370714425222,10.2090658764587,9.99651813722401,9.29005095506878,9.25201774417098,10.2506972931884,10.3543476236972,8.99227321606779,9.19454225069094,10.2829501703409,10.2667622406639,9.94147862167354,9.95396584374301\nNRAS - mRNA,9.12104695188553,10.1129362662941,9.77597835774987,10.0336649349876,9.86906534461477,10.1069630199092,9.51182024059263,9.51372322528903,9.60990575577581,9.41402589047611,9.57348701050883,9.55481553129682,9.5438176232066,9.63897186002594,9.66654469968908,9.76962421931471,10.0687694418659,10.4905155374705,10.4692936866504,9.87923921722961,10.0089910334195,10.1643058804721,10.2460384857346,10.5669469384921,10.6222947693087,10.5371835029351,10.3246569411427\nMAP2K1 - mRNA,9.47497293686099,9.9164887073078,9.67937003395048,9.74191923251233,9.74269884418094,9.46863729358928,10.162334733402,10.0969914274703,10.0848534634675,9.12193980711768,9.0856586929116,9.17090054951375,9.13054921678081,9.32381472311601,9.26951956532779,10.0677294379732,10.0544458213192,10.1951482688542,10.5347212173608,10.2777332555008,10.1932266340883,9.91677321126969,9.90049783362529,10.0519719567081,10.076479244882,9.88653309914669,9.77883973059279\nROCK1 - mRNA,10.0760294137024,10.2356450529451,10.2496360234649,10.2732144775206,10.1916059188324,10.215604279913,10.1701288908838,10.2420270185129,10.1583966998318,9.38118263378909,9.38469311544597,9.66634613250899,9.64794843368933,9.63314278122374,9.65492873483578,10.2973825441913,10.1178110062199,10.1927537537139,10.1664595378084,10.2142668366574,10.2269225529728,10.0908594357898,10.0614009863936,10.2079881126597,10.2263351507884,10.2647000335824,10.1921521369471\nPPP1R12A - mRNA,9.86066944374721,10.334292640878,10.460020632179,10.3088873261764,10.2829537438401,10.2641151530948,10.2339888328308,10.3079560201281,10.3302517283725,9.54476610835758,9.48536790870935,9.44270516497528,9.61125363370865,9.44632678208354,9.46976179110975,10.251979302867,10.0985491040592,10.1831357511523,9.82289131416398,9.95123243082359,9.99092739989135,9.8436935130274,9.96680020399345,10.1995477498591,10.1557911019959,10.1480732583002,10.1177699406245\nPDGFD - mRNA,10.7155808250605,9.96549060778179,10.9973936743819,10.8447377097411,10.7460384175261,10.8100043098221,9.96319958693519,9.26182156178986,9.39622576577594,8.22071796159585,8.36056145297307,9.16737748295006,9.40454283460447,9.60958805324484,9.80751485315879,9.84630722181181,9.68351160478039,10.0305964395108,8.53731211007905,8.28623043217732,8.34838606545846,9.97916313374153,10.1013033927536,8.78652434422145,8.91788129031771,9.46850555726204,9.8902796516484\nIL1R1 - mRNA,11.7381518361639,10.2467002414541,10.3647344316295,10.2921516791458,10.4220239953902,10.609558177897,11.2152765725686,10.7839362836322,10.6544924527279,10.3752382700398,10.2662927917568,10.8122209988899,10.7679791373961,11.265976131564,11.2048697808096,9.74979924095846,9.48776061597017,9.66303765039882,9.38837369126683,9.04720148935431,9.05455850607906,10.2037773212615,10.1501784703271,10.0208285738011,9.86420703971253,10.451948624942,10.5221359627794\nYBX3 - mRNA,8.77734823432695,9.64289320411839,9.16688419455581,9.51298726063435,9.28639830188127,9.26700343241963,9.50444071022703,9.64706445880835,9.56471698200956,8.42327396727813,8.30021854728244,8.49142045000831,8.52779740261612,8.71083784622526,8.67462058266145,9.28491319222906,9.48538971150743,9.55159111422224,9.83136117239097,9.60622072282319,9.50396904917931,9.59296864305355,9.54302022349174,9.52413385495309,9.57326353019179,9.62975768378458,9.40101756734783\nMYH10 - mRNA,10.0587789685496,9.2356450529451,10.0012981288378,10.22631001293,10.1123100898097,9.61863176152241,9.67847011000208,9.621363948477,9.73900717601327,8.88479917943461,8.96509012172134,9.68617730443056,9.74352609367156,8.90071294883821,9.06353736163721,10.3605408970424,10.4130117487,10.1975388162792,9.23509444113451,9.55289122041706,9.60992622461192,9.71876029242973,9.66575247222292,9.44881123940336,9.47901833703488,9.45949806676826,9.62928655502095\nSMAD4 - mRNA,9.50415396227294,9.00509131107848,9.15641719550475,9.52483350348246,9.37318638747604,9.32080987611542,9.54576757251597,9.34411568342351,9.26111553492879,9.02503882356382,8.92568877210897,9.28258333856333,9.28678930311233,9.29456990334016,9.14052216458613,9.29986353369503,9.37317148416824,9.51755479022282,9.21149592918275,9.31261461797084,9.24949030987807,9.36456552813169,9.46306772230396,9.51444479878465,9.33148112625797,9.61631215271963,9.63081564370769\nCRKL - mRNA,9.28430822304684,9.59178886317038,9.58885774104503,9.41636171422427,9.46115730939403,9.49104370663756,9.59291866588687,9.35549795186005,9.45224225275115,8.72571215761359,8.95591555954083,9.1918601681494,9.2152360420873,9.09699252985553,9.28014440586492,9.40039040964899,9.53781475839206,9.48269603301117,9.6437341692152,9.46794134799734,9.56050100759908,9.53061533027267,9.59396178915491,9.48100660706614,9.45281062196939,9.51271826259436,9.36104109163689\nPOLR2A - mRNA,9.33486605293234,9.89977776709987,9.56267380924138,9.93184309925959,9.78389796135209,9.94507533753227,9.63632844237501,9.71048983894999,9.80849683076341,9.13607013595256,9.03428133354601,9.14605647853628,9.26117865943329,9.22393436074709,9.35384250381018,9.18645055334337,9.83923298647155,9.43891434098326,9.6785759184275,9.65390169967698,9.64561090626459,9.66171778210105,9.74478669140268,9.57908226435097,9.69005016297373,9.79163041293216,9.60148214974953\nPIK3CA - mRNA,9.43986913590508,9.70192077977043,9.87718443256626,9.87215475579822,9.58753459516358,10.185389666866,9.33173862411415,9.24967247912375,9.37768187240975,9.11053504384543,9.39321467029183,9.3525073560037,9.47307421627528,9.19784513187843,9.42669306921786,9.4986924832608,9.45904931932865,9.50216589720939,9.51294756167985,9.08387019940171,9.36324402418557,9.32177453065996,9.59396178915491,9.03716114492302,9.16389160076701,9.12547182936303,9.22187834396527\nTP53 - mRNA,9.49220964178629,9.51705798809375,9.89618142302363,9.91393707988674,9.67773240413516,9.75243025958987,9.77567226902745,9.50863434861623,9.50445645719209,8.81975564752873,8.93038046743369,9.30520164122535,9.5420463583544,9.40130924390046,9.63141240257314,9.68863537828007,9.79492589966167,9.5993790131765,8.85232079102662,9.21426683665739,9.32580821748325,9.40077056590199,9.57767670556742,9.21518361406393,9.31265995061276,9.45648302901282,9.6565652151196\nCTTN - mRNA,8.83252494147908,8.5060589891445,8.35255971768337,8.25504173147336,8.46115730939403,8.30121914775754,9.29514147045058,9.50096721786458,9.60353623090009,9.64303012467245,9.70251291603109,9.51378826303676,9.36701682332898,9.50495451205425,9.45376131241603,9.22873035683593,9.38466712300607,9.34715136229929,9.56804523761866,9.48491442660105,9.43813123581656,9.25807329857664,9.36640766270268,9.42831806753928,9.30095444996048,9.26297535311296,9.12971554553044\nPAK1 - mRNA,8.69339645599449,9.24890114964737,8.83623888256734,8.88402885608651,8.8963726906105,9.02913960232074,8.75923112196592,8.99697130292731,9.08941895791324,9.59882152515491,9.61050020119518,9.13527663978303,9.22630819958624,8.88444789447543,9.07054925899259,9.29986353369503,9.24589082013442,9.09868542935295,9.93586148656932,9.45292431235662,9.49049840938186,9.25412070695114,9.32620883441859,9.54331891214958,9.49194579845121,9.15180597717649,9.09024726234499\nHPRT1 - mRNA,8.95692757212443,9.13452312850224,9.03597070649262,8.91264953402628,8.85984681458539,8.95585517628551,9.02982431882266,8.90313491689238,9.087138016662,9.29564644910242,9.45963353556856,9.0950467949945,9.09716304828626,9.38290140194191,9.45510148440362,9.25709524808929,9.19165850845072,9.30342998486997,9.85509254027667,9.66697475390228,9.60493419526817,9.40434159505418,9.47110504136938,9.06131846351772,9.09439055733175,8.98606877269761,8.90804393112424\nSPRY2 - mRNA,8.58831440993694,9.13452312850224,8.77825569846094,8.76350609472047,8.75435368573807,8.86346157197861,8.84742327237707,8.5587329602066,8.61940762354297,10.1276085572044,10.1413521931827,9.82009537733626,9.76038198955654,9.92636153620882,9.87415204618908,9.25453954318554,9.22045577275815,9.42673962703715,10.5578734325614,9.55490194553821,9.56436364723148,9.53713599692527,9.5090728915684,9.77845104368577,9.732062642453,9.40423916146142,9.4980420217427\nPXN - mRNA,8.15391158579116,9.49126198721642,9.03597070649262,9.22007805921853,9.12647320708221,9.38070393158436,8.90833765190068,8.92239681905301,9.04074511117755,9.4529272412925,9.46449382665909,9.04600930168595,9.32231001946359,9.35247848136816,9.44029067261859,9.12341681285733,9.51242267020444,9.15389093137641,9.88116191891293,9.76750162985625,9.79664134163074,9.22816147988846,9.361227423597,9.84180677972264,9.93691448248032,9.77961789523018,9.48457452566414\nITGA3 - mRNA,8.54271754349683,9.02577546525021,8.86223409110029,9.06645487010532,8.76426915460316,8.84425096801278,8.92685773987147,9.16487600570466,9.33602636599917,9.57826996621393,9.66640894234584,9.28258333856333,9.53671947830543,9.32863173941379,9.43214749930997,9.55378004336501,9.88402090821839,10.18795276745,10.4890569414718,9.92348744034069,10.0118226242027,9.59765525802788,9.79559565032456,10.0732471078358,10.2622420436265,9.80354373447566,9.50972382679033\nCASK - mRNA,10.186044206732,8.86862906852921,8.75993551525415,8.6475526617356,8.79683701705959,8.68492344023159,9.33728745803477,9.32687211112161,9.48726084192956,9.25219619306902,9.36056145297307,9.2295127096288,9.14228805323757,9.33582724081799,9.37806610073721,9.5620714169964,9.50190479829044,9.39376230459088,9.65808946219227,9.72348677499294,9.70513647159906,9.81178006338073,9.77885736749237,9.82465947310822,9.72988169715348,9.75528856586844,9.56994657631642\nYES1 - mRNA,10.0691539676115,8.24890114964737,8.63482359499721,8.54158973180804,8.56312928249531,8.44077050015634,9.69801029639773,9.81312475115979,9.76492455038792,9.5319988122837,9.54929030007818,9.70573957080908,9.57706494753282,9.67346023641123,9.63732751967814,10.0128336467923,9.85309183503194,9.86602467256268,9.66284299216291,9.80684252148842,9.77569240207057,9.7996281381254,9.79431496223257,9.71557135348214,9.64793299725374,9.56124077022596,9.77054040737431\nCRK - mRNA,9.27835895671565,8.78009239392755,9.00907547942956,9.23663715109194,9.04476453050317,8.95227083893647,9.95780632816498,9.8934066599305,10.0253404181792,9.81449353212331,9.81207050891707,9.66634613250899,9.89414131299237,9.74366923869263,9.77139247176388,9.72231850453636,9.78242663085308,10.1265522227859,9.99360265401399,9.87441683377276,9.81621097909674,9.7358675486999,9.81719518693073,9.74900726579329,9.85122319608508,9.64042460883644,9.72115238041214\nPPP2R2A - mRNA,9.51093515003819,9.58134711644873,9.45005624596068,9.53658326626027,9.59125288340969,9.2813587253967,9.91762741381793,9.80691960163308,9.66294051809768,10.2127399041723,10.0012158287313,9.9113806278562,9.94328231856498,9.98703366438629,9.97569007265115,9.92555982552789,9.82711334457863,9.86602467256268,10.1072403781285,10.150541239047,9.93334856617962,9.81312398060962,9.83474338814158,9.96785410893601,9.87115023997272,9.94793364381084,9.92559213233509\nGNAI3 - mRNA,9.55261034825526,9.58483610252861,9.76453741721315,9.44692462129859,9.58939493718911,9.57268217566192,9.67847011000208,9.79860433342108,9.87214550720181,10.1458796592106,10.3002185472824,9.87728285064977,9.99728268591734,9.95312260432158,10.0529551989779,9.88482603441619,9.78435665795148,9.82022098294955,10.058771668494,9.90153296169938,9.94030132682138,9.70144774418649,9.79815362092534,10.0599869497722,10.1137558821987,9.88653309914669,9.93799715322587\nMPP5 - mRNA,8.79412394707778,9.52797376557294,9.15291125200321,8.97071859028181,9.13668701994421,9.20510099349372,9.46189501526095,9.88949691487942,9.83863970398609,10.0005338600552,10.0867098368859,9.61295396734834,9.47678771907435,9.54689627512756,9.5218503022582,9.45928409870256,9.31160366081635,9.58484292001862,10.0308515830366,10.0656523414605,9.9803651926811,9.53876156897431,9.47430747340772,9.19348854296461,9.08759339497798,8.80354373447566,8.77607859091574\nBRK1 - mRNA,9.87782102713192,9.36302435942133,9.59145018566156,9.7917951391519,9.12390841522291,9.73584737539427,9.26924993731855,9.45408852544864,9.93564211079148,9.49080712496359,10.0418887826507,9.4132613870828,9.93115309684733,9.49641783009977,10.043186220162,9.70370282636901,9.74426563585949,9.95348751381302,9.88252102880232,9.48068984073933,9.93931012441833,9.11938513397113,9.59690306446703,9.33010860371028,9.82080685428308,9.08699768154839,9.13403499498731\nAKT1 - mRNA,8.89606673605993,9.59178886317038,9.02450551999917,9.58588288220254,9.31591158212429,9.54677185401322,9.17633404041054,9.06787967922901,9.19486825649697,9.63110695729663,9.62504362113698,9.55750964046608,9.65288072923156,9.50069248520009,9.65492873483578,9.1373560038152,9.85584770032245,9.76326509813748,10.1337337261753,9.85985220601622,9.94722074076117,9.59140304871554,9.82724885159465,9.69169642260367,9.83710866661218,9.69260733311231,9.45207616779052\nDIAPH1 - mRNA,8.78296184079048,9.66942627456465,9.35865991721979,9.52820035458086,9.42827394758024,9.47614483025822,9.30083258252843,9.33839069558988,9.35889651531943,9.7950332181175,9.69976753929227,9.47156002764747,9.5134063103105,9.55514026922247,9.57212350873943,9.5073056135655,9.49367088659123,9.28106217184152,9.82572012887597,9.66697475390228,9.59992483245442,9.22614513428996,9.33504349179335,9.53950225637629,9.50220500641879,9.45346167703759,9.10131941984394\nABCF1 - mRNA,9.29023305700394,9.19055716341657,9.26433253467163,9.27922154396849,9.22788683432706,9.1899623966558,9.45680613858815,9.27686544201431,9.54324545170673,9.46641047065613,9.60317314105348,9.49142045000831,9.55966190127138,9.52607780362001,9.66654469968908,9.43013775304305,9.50892522596822,9.63509110989591,9.97078274877117,9.57881456548254,9.56307724942811,9.61162444233387,9.64458786222488,9.56600921012567,9.52879338147752,9.44131259437147,9.32748853192587\nIL1R1 b - mRNA,9.85668226007377,8.66613620463046,8.60434298854245,8.86018211413215,8.82237210916673,9.09238203631256,9.78990616220015,9.22816225043863,9.21171744118919,8.60493033473755,8.50594427015259,9.10980595226692,9.30361392575369,9.74906922643522,9.72437361785855,7.72231850453636,8.21187678786937,8.09868542935295,7.87434709735662,7.29588060221104,7.57716492945457,8.62698875198605,8.66013886575939,8.31008630027847,8.39479823718324,8.94793364381084,8.97217426854396\nSOS2 - mRNA,8.74031750338199,9.26203655183651,9.00907547942956,9.07568814988336,9.01462165728049,9.15608753101778,8.81994253595496,9.14205323531971,9.27721271772572,8.8336948384866,8.93272060588821,8.61036152273182,8.74352609367156,8.74727147537283,8.64439379522959,8.67723061500782,8.78628410652259,8.98150188998261,8.88523541349935,8.55087768897941,8.8031940934299,8.90673703138111,8.8981151823143,9.29658203592872,9.22710850649755,9.05147124791925,9.0335748132081\nIRS1 - mRNA,9.16685064149866,9.27505343750334,9.47834529039211,9.37020630504314,9.4591239122415,8.99818667399183,8.94514309289922,8.95643314305243,9.11427414659952,8.60086067042851,8.65511577281448,8.74880829270096,8.90933498664164,8.43072992703252,8.56469330326394,9.17297991354592,9.07509013123525,9.38539883165626,9.0073725159874,8.84842162523982,8.9501760747565,8.97316439312009,9.18675440429849,8.45490285313025,8.45677225843834,8.60001034039053,8.72258718415138\nDROSHA - mRNA,9.2643811791592,9.2356450529451,9.19443052144148,9.16707080955537,9.11618656784517,9.20810974336606,9.10331078431128,9.0328610900506,9.06875948734715,8.40471814824988,8.54011087330379,8.68617730443056,8.67082289994192,8.69972589110503,8.83351006169174,9.28491319222906,9.22614688483601,9.04128311830803,9.29551938280756,9.33618859086457,9.18572343580164,9.38097032316786,9.29756878637469,9.06397780996821,9.06354792101133,9.09089159765288,9.14049001712461\nCCNG2 - mRNA,10.0964613136073,9.69870406708689,9.80753857031404,9.92929866290183,9.83966893264776,9.94327084068552,9.15449823917007,9.25879386909229,9.17352622468508,8.94061004240311,9.14135219318268,9.49142045000831,9.4449140175917,9.2135552149671,9.3056264352786,9.11498819078674,9.18290603444175,9.48269603301117,8.7636436329504,8.7359468959814,8.71095614484317,9.21398751986372,9.27387269393133,8.95358401196937,9.03026021991203,9.17400172329068,9.37934965004983\nDVL3 - mRNA,9.45049025166165,9.57434372871097,9.49505623060004,9.65218907185631,9.52670049554058,9.53483882451724,9.20089114465452,9.10206836540484,9.2590906994513,9.06989305934808,8.95361277367141,8.92604867764503,9.1654830068902,9.20833750569608,9.22464929327322,9.21302027374727,9.35899571540387,8.61734555970595,8.44833848025677,9.23948942036064,9.22692255297279,8.83974272541676,8.84961666462311,9.564131922885,9.49964703581801,9.39166657609015,9.51966206384645\nMAPK8 - mRNA,9.14522062552712,9.60215557872864,9.40360732327376,9.35322212541226,9.34262892050007,9.34834305987157,8.82784773404031,8.47250056256252,8.58101879433866,9.15839468262402,9.27461872404614,9.13887887646323,9.1792230635165,9.10262806452496,9.26494199922125,9.40960893967837,9.27226116064111,9.30786223118523,9.40360016549149,9.08108238203009,9.00615387411601,9.15585669063628,9.24040661765527,9.42831806753928,9.37399977451085,9.3138394357588,9.40459745762363\nSOS1 - mRNA,9.46278352785308,9.11535081922739,9.34028138790493,9.37208115266358,9.41993464666752,9.25832118931983,9.10980942161003,9.14533581930901,9.11651261879552,9.17764923970396,8.97874348899509,9.09875076375792,9.15161083230864,9.13598666147139,9.13551569903836,9.1756841224743,9.25566237687707,9.28556356407646,9.03452412224737,9.14921095306903,9.02496372389077,9.33118549058603,9.28485773462513,8.86181247152569,8.57812109384109,8.88653309914669,9.01963562225023\nGSK3B - mRNA,9.38924306283511,9.60215557872864,9.56267380924138,9.32058491761991,9.50320969095442,9.56331399312772,9.14819825081434,9.23125484549401,9.0641280386155,8.97245890844245,9.02334348706112,9.12441564715127,9.12818994390689,9.12495261119641,9.02073486384988,9.17027062628793,8.9856550476533,9.08325538878334,8.85232079102662,9.02416395327744,9.08893154990047,8.9316977758247,8.98599877748029,9.07983176116905,9.04436981932747,9.19224084157573,9.17233888316395\nEP300 - mRNA,9.63717057627869,9.36302435942133,9.34028138790493,9.56472623060598,9.44275272901261,9.54677185401322,8.98810492869801,9.07133523786523,9.0985068307412,8.79859103753069,8.79545088734758,9.03445227026355,9.19283349503515,9.12771904548779,9.21202168494547,9.18376646877216,9.33681000846703,9.16370522120336,9.29143242528432,9.01542914329984,9.14159049917303,9.31420131095516,9.40047524426484,9.04256450345236,9.1525381242008,9.3138394357588,9.17862514601138\nCCND3 - mRNA,7.47497293686099,9.08117370390931,8.79184476320889,8.92547357438387,8.77738465439342,9.31523961807248,8.96678392428423,9.35832954264321,9.48552995385045,9.21540416169938,9.29477098244277,9.04600930168595,9.08504966863416,9.17127541147453,9.26494199922125,8.43240080455161,9.09854910405924,8.68705453151736,8.95407428982735,9.14654669554197,9.05272651213017,8.52242283215956,8.7421322379567,9.2223434056368,9.39892218960796,9.17400172329068,8.68481053334842\nKRAS - mRNA,8.82710126952405,9.28366656780805,9.57057900732673,9.28917807100091,9.2795009419346,9.26122107982562,9.24878583475884,9.37240472785493,9.49071640056578,9.33292746518187,9.31105235319371,9.30840407326369,9.20855172878208,8.93586340397598,9.12376593626056,9.41190439766603,9.16227470166962,9.22122892232434,9.08261841044841,9.11145603107361,9.18572343580164,8.98035989452429,8.97473646391721,8.12382766499573,7.93880406368154,8.02322177111508,8.22795285837495\nCSNK1A1 - mRNA,12.3529059100807,12.298085627721,12.4273815151267,12.3336323239501,12.2269908431746,12.311747253644,12.0745299344667,12.2431764237317,12.2983178318151,11.7829603858169,11.8672478122684,11.8805139719128,11.9721803651074,11.7871887323554,11.906595787522,12.3521981515418,12.139648860451,11.7022854710667,11.5398983582307,12.1518702995195,12.15039727061,11.5569241467596,11.6897140190189,10.91801387963,10.9907318557254,10.9436335063772,11.2309805510725\nRAB13 - mRNA,12.7392350761403,12.4392076510877,12.4407415696841,12.3398725717532,12.4453230041284,12.3622527494387,12.4979525108647,12.2141629434301,12.1573099238551,11.9712771786501,11.8703030760974,12.2662075096095,12.1273042209439,12.4427210827095,12.4079576228029,12.2423377318459,12.2890110365042,12.43992427805,11.8869293141764,12.0649470395443,11.9699677812557,12.2970148001385,12.2839455025107,11.9297253192114,11.8149031033276,11.9650071571698,12.2455254878229\nPDGFRA - mRNA,13.995667161077,12.0737434984338,12.7216949253574,12.7952925833881,12.4499379913116,12.798821534541,12.397288357458,11.8336179230239,12.0748157872672,11.416922287837,11.6405159945899,11.9757398736526,12.250100695708,12.2794111797677,12.5609638008659,12.0379441826952,12.019641391895,12.020505269614,11.2292309025071,11.210445125705,11.396486474438,12.2929268207893,12.622386606897,11.5163878227024,11.7164509823148,11.9387804816672,12.2744272881745\nPPP1CA - mRNA,11.3161138787111,11.6727088585539,11.537351029078,11.4859721191916,11.5208635555183,11.5378313479733,12.0624361633479,12.092746955327,12.0751035493188,11.4652916687072,11.5404945177791,11.4608069713506,11.3913227298451,11.5745329954444,11.5415396002711,11.8201885688499,11.8198394337045,11.9994237979799,12.0677604517213,12.0000148039099,11.9178328758632,11.8090884673745,11.838165415719,11.9844430787278,11.9749345134341,12.0369111256632,11.8544083559616\nVCL - mRNA,11.7496646756412,11.6107379659714,11.8925071022141,12.0016270158786,11.7659151274467,11.6675413620034,12.0082460667121,12.1155201648735,12.3825727635026,11.4814299878811,11.7151427651792,11.4850665100249,11.7500619651285,11.6041354972098,11.8711432963167,12.2212239524025,12.2823685471599,12.103792319786,12.0157575931066,12.1022046596011,12.3132381217643,11.7162484486697,11.9470545902361,12.1164808128647,12.2910035186703,11.8479378538341,11.9074132447575\nACTN4 - mRNA,10.3883226831925,11.8419050420768,11.7674061694825,12.0190816118432,11.7859273540638,11.9182273754095,11.5087500222794,11.7923362784886,12.0814198362039,11.70879436718,11.858350781161,11.7124033010901,11.8885764264748,11.4312898707775,11.6473278923176,11.9371606522629,12.3414056279554,11.9378445360605,11.7330816548105,12.0932395086044,12.2025510869195,11.2897928843023,11.5806513355715,11.8230904734529,12.0191402709111,11.827662745422,11.5051458205113\nACTN1 - mRNA,10.8672903596398,12.0031368808303,11.6479589971863,11.8764539075255,11.7928060140435,11.6816802128465,11.762328142921,11.9479990460419,12.0973739723863,11.9885129475387,11.9877744733575,11.9166363533077,11.8756248129807,11.8123400493577,11.8373692571474,11.8606625147889,12.3285669564403,12.090991037563,12.2117662739865,12.3484124266962,12.310749230492,11.7856626184537,11.8003881307838,12.3569222171861,12.3927318319015,12.0796681397499,11.9352153086857\nITGA5 - mRNA,10.6746452816974,11.4535915692504,11.2463497007663,11.4077050623943,11.2278868343271,11.1459249121695,10.5553218792292,10.6190046756031,10.8588364179729,12.0786989567234,12.1964368377001,11.8745847106434,11.9908044754772,12.0774507393238,12.2132102296022,11.4752866174048,11.6524116395381,12.304539323672,12.489502974878,11.5989306868927,11.7164631742878,11.764283859049,11.9340298700448,12.2660159248997,12.4293137120818,12.0816263280938,11.8530988981536\nITGB5 - mRNA,10.6971176214336,12.0296209452015,11.9667704457096,12.0959658959439,11.9563479953895,11.6669947830838,11.18712924341,11.2858170428588,11.488557646548,12.4703194590303,12.5902605560861,12.5473806844606,12.7319194740236,12.0335037038701,12.1838052364958,12.4369162856741,12.5386725693623,12.3958456190914,12.3104907611,12.1155487857075,12.2107891966183,11.5827617784215,11.7264379119263,12.2497567626114,12.3057212697836,12.0369111256632,11.8081929488537\nCDC42 - mRNA,11.9821576869467,12.4512041865429,12.2343736319414,12.1414682245222,12.2998108536188,12.2594093314513,10.8676956674033,10.5709904259619,10.5802080637521,12.6178263012613,12.552337175637,12.488246978029,12.3667666811453,12.4143962931921,12.3982366231686,12.1804043254633,12.0995183315333,12.2577694836884,12.2470143247585,12.3968582499359,12.2705287449272,12.1150331094314,12.0804397602994,12.2622577751878,12.1295595604227,12.1097253736061,12.0885790851991\nITGA6 - mRNA,9.43452911114988,10.0309004974281,9.32166570973759,9.66448047775641,9.65126019277397,9.70423117133254,9.42589006180545,9.32976039044644,9.24073814755744,11.047640252501,10.9297948392377,10.17441503369,10.1387764205726,10.7695839374434,10.7622187675059,10.2886653270902,10.4680831311937,10.3428383694282,11.7164116737019,10.8319335024512,10.828040822085,10.9470817458744,10.988240728267,10.198337942626,10.1443734445662,10.4534616770376,10.0462351620649\nPDGFC - mRNA,10.1364769924551,10.5406053225466,10.8729286918965,10.7563463031476,10.8008567885119,10.8208554006008,9.43108896847977,9.09868572601019,9.1030293894319,10.5772346629468,10.5308726664484,10.7721912201389,10.7512122868167,10.6311945094342,10.6665446996891,10.7324556560809,10.5644533729647,10.9045779133321,10.3056864089646,10.2642786048252,10.2693460463714,10.2580732985766,10.3542913763866,10.5037111371997,10.3372232036526,10.4549731439552,10.4726864474658\nCD9 - mRNA,9.60584046252195,10.5778496722125,10.231467900766,10.1885079802177,9.95526637966407,10.0240266789271,9.82981728389008,10.3138028165435,10.6397827971104,11.2404721959369,11.4180639220144,10.891232409334,11.0617502197955,10.8828112578383,11.1417710710622,10.6857926285112,10.6767736598827,10.8140024602144,11.015447902209,10.9443460452384,11.0866951596822,10.166410468797,10.4329341309119,10.8096846291407,10.9952786449943,10.569636691347,10.42501120082\nPSEN1 - mRNA,10.1099235734138,10.4011241493562,10.1772758004024,10.2509720671643,10.1644077775027,10.3221990866618,10.4062255814282,10.5624211472367,10.6796896723948,10.4175008701752,10.4909366309729,10.09689996807,10.2130113769728,10.2884018946253,10.4671073376638,10.1864505533434,10.2653681946337,10.2922894272648,10.4565746303565,10.3593835445172,10.4332109556339,10.2027523158607,10.3869445937054,10.3661096674882,10.538794773635,10.2920181523504,10.3793496500498\nCLTC - mRNA,10.2112447608571,10.395174883025,10.4992038952898,10.3263983373366,10.226692055757,10.4484240726998,10.1794266354659,10.3079560201281,10.4820619352552,10.5730860043269,10.8341449620555,10.3115994124235,10.5269021527947,10.4910566383454,10.6337813604796,10.8032384999199,10.6043307842309,10.4489820373935,10.7532804634174,10.7804484266522,10.8545714527955,10.3236616402042,10.516858636535,10.6311180328873,10.7633213596223,10.5682407598971,10.3893211917571\nFGFR1 - mRNA,10.0587789685496,10.5513450374593,10.4270187098838,10.7295417578049,10.457087645085,10.5656617479841,10.0502401553768,10.0257542035563,10.2529990857244,10.5373322565867,10.602438384205,10.4162329536141,10.6545210876128,10.3334327256776,10.621300707625,10.4503785454807,10.7121821445903,10.7156903994782,10.7323280924842,10.5356859546332,10.683397222346,10.3037227931509,10.5604525269647,10.1377791103783,10.3293219414585,9.93932051350613,10.1068237952387\nPTEN - mRNA,10.4011548710783,10.8027411328727,10.7997130065422,10.6902571126898,10.8048653908155,10.6913881039577,10.6033185950781,10.7182254877296,10.5737057652368,10.5319988122837,10.4117867763282,10.6959915942575,10.5134063103105,10.4953471843802,10.4829612888646,10.5825937408697,10.4285214125955,10.2514557097562,10.252017744171,10.2789502062612,10.2661874352774,10.32177453066,10.1945422506909,10.4919098849673,10.3400856947521,10.5917898335856,10.462418112012\nPRKAA1 - mRNA,10.6301803847472,11.0244913574299,10.8982768410258,10.8133438636709,10.8644634426558,10.9113212107572,10.7352638059034,10.7996463667209,10.7527437898272,10.5521618516034,10.5059442701526,10.5344477336029,10.5420463583544,10.6457428180034,10.6694341359461,10.7416098431451,10.5251745776133,10.5251881151252,10.2883595912347,10.6576489616302,10.699878617888,10.6216298361474,10.6078797471999,10.7580678981722,10.6605832540331,10.7404915639404,10.732591053147\nPPP1CB - mRNA,10.8573475563922,10.6345005391418,10.8793076044592,10.798781569468,10.6078675972606,10.939655061748,10.0519285055295,10.0150277969633,10.2283721143533,10.7275797242704,10.9067679102117,10.6296922501805,10.8447049280389,10.5344412765546,10.7821109091851,10.8292337084529,10.7462473601284,10.6386141764596,10.4629481751411,10.6312114017009,10.817290436716,10.3618131679184,10.6381774565321,10.6630412907865,10.7380432975035,10.5199570888245,10.4667054704151\nSPTAN1 - mRNA,10.1210469518855,10.7891946010348,10.6818049628675,10.8899294575262,10.7551825826531,10.6946096052015,10.4542549530588,10.6366058907607,10.7022205057595,10.8864718695245,10.9945108048534,10.8577426642541,10.9856008808697,10.6563191035898,10.7126556668174,10.6666964524074,10.8777071342862,10.5213765011752,10.4326518430161,10.6388145552187,10.6934262202856,10.2068479752215,10.3758567262216,10.724843746166,10.8335583518353,10.6977232783346,10.5360751537373\nEPB41L2 - mRNA,11.0051541518498,11.2245044953399,11.5613520546451,11.5712690764727,11.4178422677063,11.5773435530318,11.3128521647675,11.3863439188128,11.4807592759876,10.1790147817282,10.3369030023005,10.482942296084,10.5579099945098,10.5846359431801,10.7890354325511,11.3244415564157,11.2172446287669,11.0532117626261,10.7033970905135,11.0670619117152,11.166792918719,10.8645835799662,11.0318199958439,11.1831291254033,11.2998521743121,11.1169041661969,11.1686592071716\nWASF2 - mRNA,10.8824041329995,11.1093068428934,11.0837061327941,10.9781233715036,11.1136034068312,11.04437054187,10.6718974560786,10.4790197559461,10.2732052289242,9.68592421785978,9.63944189474557,10.1334721429363,10.0342932570725,10.1068403158119,10.0306565318691,11.097268795345,10.8581402401993,10.4308092913462,10.5739458390583,10.9101978228265,10.8097172170112,10.7743230492138,10.699667229946,11.1977326583541,11.0474379855412,11.3369759345567,11.2544801754462\nTGFB1 - mRNA,10.1571572452948,10.4535915692504,10.2029319900075,10.7734704809388,10.64411812353,10.3469788091402,10.5421683314619,10.7060507737835,10.6660003178467,10.7932510127755,10.6480123855399,10.9539227226056,10.9250500299935,10.4824370754928,10.4591145469102,10.7342911462024,11.3272436864178,11.1477229226615,10.8784398519905,11.137182856192,10.9477137170314,10.6093056177668,10.596903064467,11.0452586126216,10.949152751974,10.8741590307348,10.7290262007256\nTUBB - mRNA,10.7794558972889,10.7321270686243,10.4357009529836,10.7684968905482,10.6297270312243,10.5008914925939,10.7151599193313,10.5697693551815,10.5947319619948,10.2220433590624,10.1733689310657,10.3664017976621,10.3223100194636,10.3630822492906,10.4129660861064,10.8656951770117,11.0423998701157,11.0439424647585,11.5802412455898,11.3541970978019,11.2514882260551,11.3222465395112,11.3195471554702,11.1615637755684,11.0574734066017,11.0781977522869,11.1046245645676\nSMAD3 - mRNA,10.2400996235293,10.8614550356415,10.5197652177721,10.6374556969519,10.5893949371891,10.5715144742296,10.2619746065154,10.3398240737911,10.382340239521,10.7182176210667,10.8718282850786,10.740533902722,10.6805162961458,10.6696686572489,10.7227054407127,10.7443448108664,10.9156221556066,11.0884170938991,11.0575689214124,11.2319688383908,11.1089045736674,11.0227951608447,11.0877764018179,10.7654390498123,10.7347841958745,10.6549640314353,10.5952299404372\nMAPK1 - mRNA,10.6208071300698,11.0549988210451,10.8482115242334,10.8433868715892,10.8805062851515,10.9603231112312,10.3093273766939,10.1664924709056,10.2611155349288,10.4358940441331,10.5854348638133,10.5262194125856,10.585688374528,10.6016502338202,10.6786416345523,10.8955279582587,10.8613436880773,10.7039679144537,10.9462969392356,10.9046898448866,10.9293604640107,10.7770802312109,10.8359887011588,10.9713996871297,10.9770048726139,10.8707657832884,10.8189644037689\nHSPB1 - mRNA,9.53108954124399,11.1297538843544,10.6927115711896,11.5061740482509,10.9638966848075,11.039311407378,10.6497697977488,11.0747825394837,11.2616213000666,10.0830818311046,10.159448953626,10.3147876900545,10.5925501950302,9.96399919357219,10.3167269904862,10.8292337084529,11.499557043434,11.4051834836707,10.7308197859789,11.1472132213292,11.2458870735474,10.3132518571263,10.5269176933931,10.9805776165567,11.2489766601545,10.6310953705201,10.5311708032663\nG6PD - mRNA,9.69339645599449,10.8117020004444,10.6360797509676,10.6568106295967,10.4933078329977,10.815933201896,10.668599865978,11.2998777880519,11.5626662424448,10.5998414580648,10.8546531412704,10.6688399879975,10.8244460890551,10.3747735210585,10.6337813604796,10.4459050689014,10.6814417406651,10.640372488732,10.2739325205994,10.5427953426816,10.6303376846349,10.1388083972345,10.3368039496578,10.884556427182,11.0897209503568,10.9447097429366,10.6030409776061\nPDGFRB - mRNA,11.162550504065,11.5830926642006,11.9537324455858,11.9203576291615,11.9294285474607,11.6338152055468,11.085287102476,11.0168210787863,11.0495581809839,11.1865022462579,11.253713100846,11.6181249228515,11.7523616920355,11.1712754114745,11.247258865002,12.0412841855679,12.0490375864396,11.7967620101256,10.9345517315625,11.459915951588,11.4254450227885,11.1805366981056,11.2669643245892,11.5100634033578,11.5306739314386,11.360553893084,11.5088925418115\nIQGAP1 - mRNA,10.7221701566717,11.2943611686043,11.3247850515818,11.1387176135061,11.2201030281319,11.1245863878038,10.7888940996957,10.684776654666,10.6621745530991,11.420967499895,11.4960884110749,11.403561306304,11.3483881518304,11.3352289845204,11.3415764555799,11.485675597594,11.3206090688806,11.1951482688542,11.5548930964605,11.4399269718277,11.4051275486814,11.2912401625118,11.2848577346251,11.367186707509,11.4491695404337,11.2818343803643,11.2761382354798\nADAM9 - mRNA,10.6379451776615,11.7539865025881,11.5265543834347,11.3856014184224,11.3459340652237,11.388018833381,11.197081543706,11.4022092405161,11.4954543195724,11.7405854340951,11.8730472923677,11.7452679349957,11.7323078695141,11.5836288261822,11.7064794964782,11.7443448108664,11.6262867360874,11.6827950213641,11.6231498370591,11.8434949441599,11.8942419082275,11.402557185376,11.5187985252781,11.571159189779,11.5784241492021,11.0869976815484,11.1062743017867\nNCKAP1 - mRNA,11.2070751191082,11.0102902177528,11.1564171955047,11.1740728899541,11.1743574379103,10.8634615719786,11.6702496031971,11.6802576375306,11.5878916714801,11.4995773345913,11.3902378482228,11.6793910419925,11.595542009252,11.589159289637,11.4734037082359,11.7731996735385,11.7467423661517,11.2571966446732,11.1468001616006,11.7752836417535,11.641045893361,11.2397010948602,11.1833339572141,11.6382733856375,11.5294205038699,11.5428802331046,11.5340337001055\nPPP2R1A - mRNA,11.1876322036352,11.300740081167,11.1281281510733,11.0802827239606,11.0270290489642,11.161530202677,11.1654574511599,11.2910132745046,11.3593691426917,11.6286105125815,11.7447444480107,11.3748274363967,11.5201700125298,11.4402195397712,11.5705787018236,11.4703392872116,11.4746721588189,11.4589799640374,11.7060861780412,11.6730043513405,11.6969492868493,11.3410013702989,11.4067743157223,11.6413927274817,11.7412950532923,11.6217054114898,11.3924795558914\nRAC1 - mRNA,11.1397620679714,9.94663158053048,10.1755490556829,10.3408302186791,10.1644077775027,10.0743583544772,11.4035832820799,11.3779965760993,11.4933026487174,11.4808764889136,11.5374225016258,11.4871876008861,11.5618487973107,11.4318495972796,11.5705787018236,11.3023402639872,11.4186144549294,11.4708867307215,11.5441984956643,11.3765382655563,11.4194153113204,11.3755227583282,11.4354009796877,10.9607367040669,11.0342425120796,10.7916304129322,11.0090921280088\nGNAI1 - mRNA,11.4531333578194,11.2289710470218,11.2029319900075,11.1337532593942,11.1693911850169,11.5621386814204,12.0894032109482,11.4685748593487,11.4850969071593,10.7834093340206,10.7681926803032,10.898688214253,10.9619496128051,11.6814844074961,11.8025223692081,10.6008174994886,10.4692833767212,10.30342998487,10.5848404801616,10.5234163268234,10.5707784867949,11.1215562330128,11.216222831299,10.4344966424684,10.4626943900691,10.9284812811447,11.0513825432906\nPPP2CB - mRNA,10.9076754311205,11.4119678255856,11.224804170485,11.3398725717532,11.3153496755774,11.3999092444311,11.5445688229323,11.6728839098483,11.6728612318964,10.9771761716345,10.8869923414504,11.0179196891449,10.9953422753479,10.9915967521362,10.9891667872336,11.2455587535922,11.2395739745437,11.4226584502253,11.4827979510001,11.3203184914963,11.2681623774625,11.2897928843023,11.3168738470322,10.1995477498591,10.1212634188676,10.1721650648111,10.489639580747\nGSTM3 - mRNA,10.787157757585,11.7415359826751,10.9354830731309,10.9419762052958,11.1749770192129,11.3565015832587,11.2714254014768,11.6545709938669,11.5108523375103,11.3536334480407,11.2555836943539,11.3147876900545,11.1700775809674,11.2945699033402,11.223470139108,10.8463072218118,10.7030286028732,11.0935603971751,10.7592113866351,11.1885954290552,11.089825136204,10.9211421807718,10.8209735261682,10.9875980442344,10.8517247413851,10.7104343300842,10.7074501000664\nRHOA - mRNA,10.4255849098988,10.9893812279898,10.7918447632089,11.1392681554097,10.6179276351392,10.7940737679323,10.9739259935283,11.2204016344099,11.5176435005812,11.1012018261582,11.2760017198775,10.9467476703129,11.2351051478234,10.7015838373256,11.0454122583152,10.9319718140421,11.1704150902586,11.3717039237594,11.2811640898305,11.0550364360113,11.2920414730053,10.6780363575764,10.9702068054367,10.9091674847729,11.2127338846793,10.8317406263063,10.6208474746165\nRAB11A - mRNA,10.8850165285361,10.9774853700566,10.8019532166832,10.5947412105913,10.758493416099,10.775939094284,10.9046049330243,10.8826292614577,10.878094773533,11.2147385584392,11.2704617641129,11.1216875750351,10.982992028367,11.136673495591,11.1119196934124,10.7740921542604,10.6426036612002,10.6647664643156,10.9047634391586,11.014698849667,10.9243597829524,10.7681000559236,10.7071723996029,10.8641030701136,10.8115554828105,10.7796178952302,10.7439397452237\n";
	colAnnotation_UyySR3RU_9hQ_hWdumMiueg = "File.Name,Description,Batch.ID,Sample.Name,Cartridge.Id,Lane.Number,Analyte.Types,System.Type,Gene.RLF,Scanner.Id,Stage.Position,Scanned.Date,Gender,QC.Flag,FOV.Count,FOV.Counted,Binding.Density,Is.Reference,Imaging.QC.Flag,Imaging.QC,Binding.Density.QC.Flag,Binding.Density.QC,Positive.Control.QC.Flag,Positive.Control.QC,Limit.of.Detection.QC.Flag.of.mRNA,Limit.of.Detection.QC.of.mRNA,New.Annotation,\n20161013_sebelius2-101316_01_01.RCC,CRL1790 Control,SA,01,sebelius2-101316,1,mRNA,NONE,SEBELIUS_2_C4636,1507C0300,3,2016-10-13 00:00:00.0,Unknown,false,280,279,0.74,false,false,0.996429,false,0.74,false,0.995739,false,10.502747,Control,\n20161013_sebelius2-101316_01_03.RCC,CRL1790 Control,SA,01,sebelius2-101316,3,mRNA,NONE,SEBELIUS_2_C4636,1507C0300,3,2016-10-13 00:00:00.0,Unknown,false,280,278,0.84,false,false,0.992857,false,0.84,false,0.985546,false,11.920769,Control,\n20161013_sebelius2-101316_01_04.RCC,CRL1790 miR142,SA,01,sebelius2-101316,4,mRNA,NONE,SEBELIUS_2_C4636,1507C0300,3,2016-10-13 00:00:00.0,Unknown,false,280,273,0.63,false,false,0.975,false,0.63,false,0.987008,false,7.339143,miR142,\n20180703_1790-jr1_SA40miR142_01.RCC,CRL1790 miR142,SA,SA40miR142,1790-jr1,1,mRNA,NONE,SEBELIUS_2_C4636,1507C0300,3,2018-07-03 00:00:00.0,Unknown,false,555,524,0.44,false,false,0.944144,false,0.44,false,0.996285,false,6.5,miR142,\n20180703_1790-jr1_SA44miR890_02.RCC,CRL1790 miR890,SA,SA44miR890,1790-jr1,2,mRNA,NONE,SEBELIUS_2_C4636,1507C0300,3,2018-07-03 00:00:00.0,Unknown,false,555,536,0.35,false,false,0.965766,false,0.35,false,0.990552,false,6.375,miR890,\n20180703_1790-jr1_SA48bcel67_03.RCC,CRL1790 cel67,SA,SA48bcel67,1790-jr1,3,mRNA,NONE,SEBELIUS_2_C4636,1507C0300,3,2018-07-03 00:00:00.0,Unknown,false,555,522,0.5,false,false,0.940541,false,0.5,false,0.990499,false,10.398527,cel67,\n20180703_1790-jr1_SA50bmiR18b_04.RCC,CRL1790 miR18b,SA,SA50bmiR18b,1790-jr1,4,mRNA,NONE,SEBELIUS_2_C4636,1507C0300,3,2018-07-03 00:00:00.0,Unknown,false,555,495,0.4,false,false,0.891892,false,0.4,false,0.992382,false,7.32534,miR18b,\n20180703_1790-jr1_SA51miR890_05.RCC,CRL1790 miR890,SA,SA51miR890,1790-jr1,5,mRNA,NONE,SEBELIUS_2_C4636,1507C0300,3,2018-07-03 00:00:00.0,Unknown,false,555,526,0.4,false,false,0.947748,false,0.4,false,0.989921,false,6.355745,miR890,\n20180703_1790-jr1_SA54amiR155_06.RCC,CRL1790 miR155,SA,SA54amiR155,1790-jr1,6,mRNA,NONE,SEBELIUS_2_C4636,1507C0300,3,2018-07-03 00:00:00.0,Unknown,false,555,481,0.35,false,false,0.866667,false,0.35,false,0.994494,false,6.375,miR155,\n20180703_1790-jr1_SA54bmiR155_07.RCC,CRL1790 miR155,SA,SA54bmiR155,1790-jr1,7,mRNA,NONE,SEBELIUS_2_C4636,1507C0300,3,2018-07-03 00:00:00.0,Unknown,false,555,527,0.47,false,false,0.94955,false,0.47,false,0.986179,false,6.875,miR155,\n20180703_1790-jr1_SA57Control_08.RCC,CRL1790 Control,SA,SA57Control,1790-jr1,8,mRNA,NONE,SEBELIUS_2_C4636,1507C0300,3,2018-07-03 00:00:00.0,Unknown,false,555,520,0.5,false,false,0.936937,false,0.5,false,0.992037,false,7.27618,Control,\n20180703_1790-jr1_SA60miR142_09.RCC,CRL1790 miR142,SA,SA60miR142,1790-jr1,9,mRNA,NONE,SEBELIUS_2_C4636,1507C0300,3,2018-07-03 00:00:00.0,Unknown,false,555,506,0.32,false,false,0.911712,false,0.32,false,0.996168,false,6.375,miR142,\n20180703_1790-jr1_SA61miR155_10.RCC,CRL1790 miR155,SA,SA61miR155,1790-jr1,10,mRNA,NONE,SEBELIUS_2_C4636,1507C0300,3,2018-07-03 00:00:00.0,Unknown,false,555,473,0.51,false,false,0.852252,false,0.51,false,0.990233,false,6.75,miR155,\n20180703_1790-jr1_SA62miR18b_11.RCC,CRL1790 miR18b,SA,SA62miR18b,1790-jr1,11,mRNA,NONE,SEBELIUS_2_C4636,1507C0300,3,2018-07-03 00:00:00.0,Unknown,false,555,492,0.39,false,false,0.886486,false,0.39,false,0.995342,false,7.838203,miR18b,\n20180703_1790-jr1_SA63miR890_12.RCC,CRL1790 miR890,SA,SA63miR890,1790-jr1,12,mRNA,NONE,SEBELIUS_2_C4636,1507C0300,3,2018-07-03 00:00:00.0,Unknown,false,555,477,0.46,false,false,0.859459,false,0.46,false,0.988295,false,6.25,miR890,\n20180712_1790-jr2_SA36Control_01.RCC,CRL1790 Control,SA,SA36Control,1790-jr2,1,mRNA,NONE,SEBELIUS_2_C4636,1507C0300,4,2018-07-03 00:00:00.0,Unknown,false,555,529,0.95,false,false,0.953153,false,0.95,false,0.993858,false,13.365773,Control,\n20180712_1790-jr2_SA38cel67_02.RCC,CRL1790 cel67,SA,SA38cel67,1790-jr2,2,mRNA,NONE,SEBELIUS_2_C4636,1507C0300,4,2018-07-03 00:00:00.0,Unknown,false,555,534,0.47,false,false,0.962162,false,0.47,false,0.988335,false,8.959881,cel67,\n20180712_1790-jr2_SA40miR142_03.RCC,CRL1790 miR142,SA,SA40miR142,1790-jr2,3,mRNA,NONE,SEBELIUS_2_C4636,1507C0300,4,2018-07-03 00:00:00.0,Unknown,false,555,530,0.6,false,false,0.954955,false,0.6,false,0.994833,false,10.666523,miR142,\n20180712_1790-jr2_SA44miR890_04.RCC,CRL1790 miR890,SA,SA44miR890,1790-jr2,4,mRNA,NONE,SEBELIUS_2_C4636,1507C0300,4,2018-07-03 00:00:00.0,Unknown,false,555,528,0.61,false,false,0.951351,false,0.61,false,0.994373,false,8.640393,miR890,\n20180712_1790-jr2_SA48bcel67_05.RCC,CRL1790 cel67,SA,SA48bcel67,1790-jr2,5,mRNA,NONE,SEBELIUS_2_C4636,1507C0300,4,2018-07-03 00:00:00.0,Unknown,false,555,531,0.64,false,false,0.956757,false,0.64,false,0.996108,false,8.535574,cel67,\n20180712_1790-jr2_SA49amiR142_06.RCC,CRL1790 miR142,SA,SA49amiR142,1790-jr2,6,mRNA,NONE,SEBELIUS_2_C4636,1507C0300,4,2018-07-03 00:00:00.0,Unknown,false,555,534,0.58,false,false,0.962162,false,0.58,false,0.990119,false,7.140393,miR142,\n20180712_1790-jr2_SA51bmiR890_07.RCC,CRL1790 miR890,SA,SA51bmiR890,1790-jr2,7,mRNA,NONE,SEBELIUS_2_C4636,1507C0300,4,2018-07-03 00:00:00.0,Unknown,false,555,527,0.73,false,false,0.94955,false,0.73,false,0.992824,false,13.141959,miR890,\n20180712_1790-jr2_SA57Control_08.RCC,CRL1790 Control,SA,SA57Control,1790-jr2,8,mRNA,NONE,SEBELIUS_2_C4636,1507C0300,4,2018-07-03 00:00:00.0,Unknown,false,555,530,0.83,false,false,0.954955,false,0.83,false,0.992413,false,15.653154,Control,\n20180712_1790-jr2_SA59cel67_09.RCC,CRL1790 cel67,SA,SA59cel67,1790-jr2,9,mRNA,NONE,SEBELIUS_2_C4636,1507C0300,4,2018-07-03 00:00:00.0,Unknown,false,555,530,0.65,false,false,0.954955,false,0.65,false,0.995539,false,10.772942,cel67,\n20180712_1790-jr2_SA61miR155_10.RCC,CRL1790 miR155,SA,SA61miR155,1790-jr2,10,mRNA,NONE,SEBELIUS_2_C4636,1507C0300,4,2018-07-03 00:00:00.0,Unknown,false,555,527,0.79,false,false,0.94955,false,0.79,false,0.997276,false,13.082381,miR155,\n20180712_1790-jr2_SA62miR18b_11.RCC,CRL1790 miR18b,SA,SA62miR18b,1790-jr2,11,mRNA,NONE,SEBELIUS_2_C4636,1507C0300,4,2018-07-03 00:00:00.0,Unknown,false,555,533,0.63,false,false,0.96036,false,0.63,false,0.996031,false,10.641959,miR18b,\n20180712_1790-jr2_SA63miR890_12.RCC,CRL1790 miR890,SA,SA63miR890,1790-jr2,12,mRNA,NONE,SEBELIUS_2_C4636,1507C0300,4,2018-07-03 00:00:00.0,Unknown,false,555,524,0.66,false,false,0.944144,false,0.66,false,0.993428,false,10.367889,miR890,\n";
	rowAnnotation_UyySR3RU_9hQ_hWdumMiueg = "Dimname,ProbeID,Codeset.Name,Probe.Label,Analyte.Type,Is.Control,Control.Type,Related.Probes,Probe.Annotation,KEGG.Pathways,Official.Gene.Name,Control.Conc,probeLabel.Type,below.threshold\nPIK3R5 - mRNA,NM_001142633.1:3335,SEBELIUS_2_C4636,PIK3R5,mRNA,0,,,Synthesis of PIPs at the plasma membrane;G beta.CL.gamma signalling through PI3Kgamma;GPVI-mediated activation cascade,hsa04012;hsa04014;hsa04015;hsa04022;hsa04024;hsa04062;hsa04066;hsa04068;hsa04070;hsa04071;hsa04072;hsa04150;hsa04151;hsa04152;hsa04210;hsa04211;hsa04213;hsa04261;hsa04370;hsa04380;hsa04510;hsa04550;hsa04611;hsa04620;hsa04630;hsa04650;hsa04660;hsa04662;hsa04664;hsa04666;hsa04668;hsa04670;hsa04722;hsa04725;hsa04750;hsa04810;hsa04910;hsa04914;hsa04915;hsa04917;hsa04919;hsa04921;hsa04923;hsa04930;hsa04931;hsa04932;hsa04933;hsa04960;hsa04973;hsa05100;hsa05142;hsa05145;hsa05146;hsa05160;hsa05161;hsa05162;hsa05164;hsa05166;hsa05169;hsa05200;hsa05203;hsa05205;hsa05210;hsa05211;hsa05212;hsa05213;hsa05214;hsa05215;hsa05218;hsa05220;hsa05221;hsa05222;hsa05223;hsa05230;hsa05231,PIK3R5,NA,PIK3R5 - mRNA,True\nCDX2 - mRNA,NM_001265.2:1812,SEBELIUS_2_C4636,CDX2,mRNA,0,,,,,CDX2,NA,CDX2 - mRNA,True\nTJP3 - mRNA,NM_014428.1:1925,SEBELIUS_2_C4636,TJP3,mRNA,0,,,,hsa04530,TJP3,NA,TJP3 - mRNA,True\nCXCR4 - mRNA,NM_003467.2:1335,SEBELIUS_2_C4636,CXCR4,mRNA,0,,,,hsa04060;hsa04062;hsa04144;hsa04360;hsa04670;hsa05200,CXCR4,NA,CXCR4 - mRNA,True\nCGN - mRNA,NM_020770.2:1112,SEBELIUS_2_C4636,CGN,mRNA,0,,,TGF-beta receptor signaling in EMT (epithelial to mesenchymal transition),hsa04530,CGN,NA,CGN - mRNA,True\nACTN3 - mRNA,NM_001104.1:296,SEBELIUS_2_C4636,ACTN3,mRNA,0,,,Nephrin interactions,hsa04510;hsa04520;hsa04530;hsa04670;hsa04810;hsa05146;hsa05203;hsa05412,ACTN3,NA,ACTN3 - mRNA,True\nCA2 - mRNA,NM_000067.2:575,SEBELIUS_2_C4636,CA2,mRNA,0,,,,hsa04971;hsa04972,CA2,NA,CA2 - mRNA,True\nCRB3 - mRNA,NM_139161.3:300,SEBELIUS_2_C4636,CRB3,mRNA,0,,,Tight junction interactions,hsa04530,CRB3,NA,CRB3 - mRNA,True\nCLDN4 - mRNA,NM_001305.3:1242,SEBELIUS_2_C4636,CLDN4,mRNA,0,,,Tight junction interactions,hsa04514;hsa04530;hsa04670;hsa05160,CLDN4,NA,CLDN4 - mRNA,True\nALPI - mRNA,NM_001631.3:1270,SEBELIUS_2_C4636,ALPI,mRNA,0,,,,hsa01100,ALPI,NA,ALPI - mRNA,True\nPIK3CG - mRNA,NM_002649.2:2125,SEBELIUS_2_C4636,PIK3CG,mRNA,0,,,Synthesis of PIPs at the plasma membrane;G beta.CL.gamma signalling through PI3Kgamma;GPVI-mediated activation cascade,hsa00562;hsa04012;hsa04014;hsa04015;hsa04022;hsa04024;hsa04062;hsa04066;hsa04068;hsa04070;hsa04071;hsa04072;hsa04150;hsa04151;hsa04152;hsa04210;hsa04211;hsa04213;hsa04261;hsa04370;hsa04380;hsa04510;hsa04550;hsa04611;hsa04620;hsa04630;hsa04650;hsa04660;hsa04662;hsa04664;hsa04666;hsa04668;hsa04670;hsa04722;hsa04725;hsa04750;hsa04810;hsa04910;hsa04914;hsa04915;hsa04917;hsa04919;hsa04921;hsa04923;hsa04930;hsa04931;hsa04932;hsa04933;hsa04960;hsa04973;hsa05100;hsa05142;hsa05145;hsa05146;hsa05160;hsa05161;hsa05162;hsa05164;hsa05166;hsa05169;hsa05200;hsa05203;hsa05205;hsa05210;hsa05211;hsa05212;hsa05213;hsa05214;hsa05215;hsa05218;hsa05220;hsa05221;hsa05222;hsa05223;hsa05230;hsa05231,PIK3CG,NA,PIK3CG - mRNA,True\nIL1R2 - mRNA,NM_001261419.1:186,SEBELIUS_2_C4636,IL1R2,mRNA,0,,,,hsa04010;hsa04060;hsa04640;hsa05146;hsa05166;hsa05202,IL1R2,NA,IL1R2 - mRNA,True\nHPSE - mRNA,NM_006665.3:1625,SEBELIUS_2_C4636,HPSE,mRNA,0,,,,hsa01100;hsa05205,HPSE,NA,HPSE - mRNA,True\nITGB4 - mRNA,NM_001005731.1:4151,SEBELIUS_2_C4636,ITGB4,mRNA,0,,,Syndecan interactions;Assembly of collagen fibrils and other multimeric structures;Non-integrin membrane-ECM interactions,hsa04151;hsa04510;hsa04512;hsa04810;hsa05410;hsa05412;hsa05414,ITGB4,NA,ITGB4 - mRNA,True\nWAS - mRNA,NM_000377.2:90,SEBELIUS_2_C4636,WAS,mRNA,0,,,RHO GTPases Activate WASPs and WAVEs;Regulation of actin dynamics for phagocytic cup formation,hsa04062;hsa04144;hsa04520;hsa04666;hsa04810;hsa05100;hsa05130;hsa05131;hsa05132;hsa05231,WAS,NA,WAS - mRNA,True\nVAV3 - mRNA,NM_006113.4:1632,SEBELIUS_2_C4636,VAV3,mRNA,0,,NM_001079874.1:2680,Rho GTPase cycle;DAP12 signaling;EPH-ephrin mediated repulsion of cells;VEGFR2 mediated vascular permeability;NRAGE signals death through JNK;GPVI-mediated activation cascade;G alpha (1213) signalling events;FCERI mediated MAPK activation;Regulation of actin dynamics for phagocytic cup formation;VEGFA-VEGFR2 Pathway,hsa04024;hsa04062;hsa04510;hsa04650;hsa04660;hsa04662;hsa04664;hsa04666;hsa04670;hsa04810,VAV3,NA,VAV3 - mRNA,True\nMYH14 - mRNA,NM_001077186.1:6304,SEBELIUS_2_C4636,MYH14,mRNA,0,,,EPHA-mediated growth cone collapse;RHO GTPases Activate ROCKs;RHO GTPases activate CIT;RHO GTPases activate PAKs;RHO GTPases activate PKNs;Sema4D induced cell migration and growth-cone collapse,hsa04530;hsa04810;hsa05132,MYH14,NA,MYH14 - mRNA,True\nEPB41L3 - mRNA,NM_012307.2:3758,SEBELIUS_2_C4636,EPB41L3,mRNA,0,,,,hsa04530,EPB41L3,NA,EPB41L3 - mRNA,True\nPRKCB - mRNA,NM_212535.1:1750,SEBELIUS_2_C4636,PRKCB,mRNA,0,,,Disinhibition of SNARE formation;Trafficking of AMPA receptors;VEGFA-VEGFR2 Pathway;PCPCE pathway,hsa04010;hsa04012;hsa04014;hsa04015;hsa04020;hsa04062;hsa04064;hsa04066;hsa04070;hsa04071;hsa04150;hsa04270;hsa04310;hsa04370;hsa04510;hsa04530;hsa04540;hsa04650;hsa04662;hsa04666;hsa04670;hsa04713;hsa04720;hsa04723;hsa04724;hsa04725;hsa04726;hsa04727;hsa04728;hsa04730;hsa04750;hsa04912;hsa04916;hsa04919;hsa04921;hsa04931;hsa04933;hsa04960;hsa04971;hsa04972;hsa04973;hsa05031;hsa05032;hsa05140;hsa05146;hsa05161;hsa05164;hsa05200;hsa05205;hsa05206;hsa05214;hsa05223;hsa05231,PRKCB,NA,PRKCB - mRNA,True\nVAV3 b - mRNA,NM_001079874.1:2680,SEBELIUS_2_C4636,VAV3 b,mRNA,0,,NM_006113.4:1632,Rho GTPase cycle;DAP12 signaling;EPH-ephrin mediated repulsion of cells;VEGFR2 mediated vascular permeability;NRAGE signals death through JNK;GPVI-mediated activation cascade;G alpha (1213) signalling events;FCERI mediated MAPK activation;Regulation of actin dynamics for phagocytic cup formation;VEGFA-VEGFR2 Pathway,hsa04024;hsa04062;hsa04510;hsa04650;hsa04660;hsa04662;hsa04664;hsa04666;hsa04670;hsa04810,VAV3,NA,VAV3 b - mRNA,True\nMYH11 - mRNA,NM_001040113.1:920,SEBELIUS_2_C4636,MYH11,mRNA,0,,,EPHA-mediated growth cone collapse;RHO GTPases Activate ROCKs;Smooth Muscle Contraction;RHO GTPases activate CIT;RHO GTPases activate PAKs;RHO GTPases activate PKNs;Sema4D induced cell migration and growth-cone collapse,hsa04270;hsa04530,MYH11,NA,MYH11 - mRNA,True\nCLDN3 - mRNA,NM_001306.3:606,SEBELIUS_2_C4636,CLDN3,mRNA,0,,,Tight junction interactions,hsa04514;hsa04530;hsa04670;hsa05160,CLDN3,NA,CLDN3 - mRNA,True\nCA1 - mRNA,NM_001738.2:895,SEBELIUS_2_C4636,CA1,mRNA,0,,,,,CA1,NA,CA1 - mRNA,True\nPRKCZ - mRNA,NM_002744.4:771,SEBELIUS_2_C4636,PRKCZ,mRNA,0,,,TGF-beta receptor signaling in EMT (epithelial to mesenchymal transition);VEGFR2 mediated cell proliferation;GPVI-mediated activation cascade;VEGFA-VEGFR2 Pathway,hsa04015;hsa04062;hsa04071;hsa04144;hsa04390;hsa04530;hsa04611;hsa04910;hsa04930;hsa04931;hsa04933,PRKCZ,NA,PRKCZ - mRNA,False\nAXIN2 - mRNA,NM_004655.3:1035,SEBELIUS_2_C4636,AXIN2,mRNA,0,,,Binding of TCFLEF.CL.CTNNB1 to target gene promoters;Formation of the beta-catenin.CL.TCF transactivating complex,hsa04310;hsa04390;hsa04550;hsa05200;hsa05210;hsa05213;hsa05217,AXIN2,NA,AXIN2 - mRNA,False\nCLDN15 - mRNA,NM_014343.1:1129,SEBELIUS_2_C4636,CLDN15,mRNA,0,,,Tight junction interactions,hsa04514;hsa04530;hsa04670;hsa05160,CLDN15,NA,CLDN15 - mRNA,False\nAPC2 - mRNA,NM_005883.2:8120,SEBELIUS_2_C4636,APC2,mRNA,0,,,,hsa04310;hsa04390;hsa04550;hsa04810;hsa05166;hsa05200;hsa05206;hsa05210;hsa05213;hsa05217,APC2,NA,APC2 - mRNA,False\nHCLS1 - mRNA,NM_005335.4:1846,SEBELIUS_2_C4636,HCLS1,mRNA,0,,,,hsa04530;hsa05100;hsa05130;hsa05131;hsa05205,HCLS1,NA,HCLS1 - mRNA,False\nPDGFA - mRNA,NM_002607.5:2460,SEBELIUS_2_C4636,PDGFA,mRNA,0,,,Non-integrin membrane-ECM interactions;PIP3 activates AKT signaling;Constitutive Signaling by Aberrant PI3K in Cancer;RAFMAP kinase cascade;Platelet degranulation ;Signaling by PDGF;Downstream signal transduction;GAB1 signalosome;Signaling by SCF-KIT;PI3KAKT activation;Role of LAT2NTALLAB on calcium mobilization;DAP12 signaling;Signaling by ERBB2;VEGFR2 mediated cell proliferation;Frs2-mediated activation;ARMS-mediated activation;Gastrin-CREB signalling pathway via PKC and MAPK;Interleukin receptor SHC signaling;FCERI mediated MAPK activation;VEGFA-VEGFR2 Pathway;IRS-related events triggered by IGF1R,hsa04010;hsa04014;hsa04015;hsa04060;hsa04072;hsa04151;hsa04510;hsa04540;hsa04810;hsa05166;hsa05200;hsa05202;hsa05206;hsa05214;hsa05215;hsa05218;hsa05231,PDGFA,NA,PDGFA - mRNA,False\nF11R - mRNA,NM_144503.1:3650,SEBELIUS_2_C4636,F11R,mRNA,0,,,TGF-beta receptor signaling in EMT (epithelial to mesenchymal transition);Tight junction interactions;Integrin cell surface interactions;Cell surface interactions at the vascular wall,hsa04514;hsa04530;hsa04670;hsa05120,F11R,NA,F11R - mRNA,False\nEPB41 - mRNA,NM_001166005.1:1330,SEBELIUS_2_C4636,EPB41,mRNA,0,,,,hsa04530,EPB41,NA,EPB41 - mRNA,False\nITGA7 - mRNA,NM_002206.1:1170,SEBELIUS_2_C4636,ITGA7,mRNA,0,,,Laminin interactions;ECM proteoglycans;Integrin cell surface interactions,hsa04151;hsa04510;hsa04512;hsa04810;hsa05410;hsa05412;hsa05414,ITGA7,NA,ITGA7 - mRNA,False\nMYL12A - mRNA,NM_006471.2:78,SEBELIUS_2_C4636,MYL12A,mRNA,0,,NM_001303049.1:9;NM_001303048.1:27,Smooth Muscle Contraction;Ephrin signaling,hsa04510;hsa04530;hsa04611;hsa04670;hsa04810,MYL12A,NA,MYL12A - mRNA,False\nBCL2 - mRNA,NM_000657.2:5,SEBELIUS_2_C4636,BCL2,mRNA,0,,,Activation of BAD and translocation to mitochondria ,hsa04064;hsa04066;hsa04071;hsa04141;hsa04151;hsa04210;hsa04261;hsa04510;hsa04630;hsa04722;hsa04725;hsa04933;hsa05145;hsa05152;hsa05161;hsa05169;hsa05200;hsa05206;hsa05210;hsa05215;hsa05222,BCL2,NA,BCL2 - mRNA,False\nITGA4 - mRNA,NM_000885.4:975,SEBELIUS_2_C4636,ITGA4,mRNA,0,,,Integrin cell surface interactions;Cell surface interactions at the vascular wall,hsa04151;hsa04510;hsa04512;hsa04514;hsa04640;hsa04670;hsa04810;hsa05140;hsa05410;hsa05412;hsa05414,ITGA4,NA,ITGA4 - mRNA,False\nJAM2 - mRNA,NM_001270407.1:840,SEBELIUS_2_C4636,JAM2,mRNA,0,,,Integrin cell surface interactions;Cell surface interactions at the vascular wall,hsa04514;hsa04530;hsa04670;hsa05120,JAM2,NA,JAM2 - mRNA,True\nLLGL2 - mRNA,NM_001015002.1:1065,SEBELIUS_2_C4636,LLGL2,mRNA,0,,,,hsa04390;hsa04530,LLGL2,NA,LLGL2 - mRNA,True\nTCF7 - mRNA,NM_201633.2:519,SEBELIUS_2_C4636,TCF7,mRNA,0,,,,,,NA,TCF7 - mRNA,False\nCLDN7 - mRNA,NM_001307.3:175,SEBELIUS_2_C4636,CLDN7,mRNA,0,,,Tight junction interactions,hsa04514;hsa04530;hsa04670;hsa05160,CLDN7,NA,CLDN7 - mRNA,True\nCLDN8 - mRNA,NM_199328.2:805,SEBELIUS_2_C4636,CLDN8,mRNA,0,,,Tight junction interactions,hsa04514;hsa04530;hsa04670;hsa05160,CLDN8,NA,CLDN8 - mRNA,True\nLYZ - mRNA,NM_000239.2:305,SEBELIUS_2_C4636,LYZ,mRNA,0,,,,,LYZ,NA,LYZ - mRNA,True\nGAPDH - mRNA,NM_002046.3:972,SEBELIUS_2_C4636,GAPDH,mRNA,0,,,Glycolysis,hsa01100;hsa04066;hsa05010,GAPDH,NA,GAPDH - mRNA,False\nACTG1 - mRNA,NM_001614.2:1615,SEBELIUS_2_C4636,ACTG1,mRNA,0,,,RHO GTPases Activate WASPs and WAVEs;Translocation of GLUT4 to the plasma membrane;Recycling pathway of L1;Interaction between L1 and Ankyrins;EPHB-mediated forward signaling;EPH-ephrin mediated repulsion of cells;RHO GTPases Activate Formins;RHO GTPases activate IQGAPs;Cell-extracellular matrix interactions;Regulation of actin dynamics for phagocytic cup formation;RAFMAP kinase cascade;Downstream signal transduction;VEGFR2 mediated cell proliferation;Signaling by SCF-KIT;Frs2-mediated activation;ARMS-mediated activation;Gastrin-CREB signalling pathway via PKC and MAPK;Interleukin receptor SHC signaling;FCERI mediated MAPK activation;DAP12 signaling;Signaling by PDGF;VEGFA-VEGFR2 Pathway;Signaling by ERBB2;IRS-related events triggered by IGF1R,hsa04015;hsa04145;hsa04390;hsa04510;hsa04520;hsa04530;hsa04611;hsa04670;hsa04810;hsa04919;hsa04921;hsa05100;hsa05110;hsa05130;hsa05131;hsa05132;hsa05164;hsa05205;hsa05410;hsa05412;hsa05414;hsa05416,ACTG1,NA,ACTG1 - mRNA,False\nACTB - mRNA,NM_001101.2:1010,SEBELIUS_2_C4636,ACTB,mRNA,0,,,EPHB-mediated forward signaling;Interaction between L1 and Ankyrins;DNA Damage Recognition in GG-NER;EPH-ephrin mediated repulsion of cells;Prefoldin mediated transfer of substrate  to CCTTriC;RHO GTPases Activate Formins;RHO GTPases activate IQGAPs;Cell-extracellular matrix interactions;Regulation of actin dynamics for phagocytic cup formation;Folding of actin by CCTTriC;B-WICH complex positively regulates rRNA expression;RAFMAP kinase cascade;Downstream signal transduction;VEGFR2 mediated cell proliferation;Signaling by SCF-KIT;Frs2-mediated activation;ARMS-mediated activation;Gastrin-CREB signalling pathway via PKC and MAPK;Interleukin receptor SHC signaling;FCERI mediated MAPK activation;DAP12 signaling;Signaling by PDGF;VEGFA-VEGFR2 Pathway;Signaling by ERBB2;IRS-related events triggered by IGF1R,hsa04015;hsa04145;hsa04390;hsa04510;hsa04520;hsa04530;hsa04611;hsa04670;hsa04810;hsa04919;hsa04921;hsa04971;hsa05100;hsa05110;hsa05130;hsa05131;hsa05132;hsa05164;hsa05205;hsa05410;hsa05412;hsa05414;hsa05416,ACTB,NA,ACTB - mRNA,False\nHSPA8 - mRNA,NM_006597.3:2111,SEBELIUS_2_C4636,HSPA8,mRNA,0,,,AUF1 (hnRNP D0) binds and destabilizes mRNA;GABA synthesis# release# reuptake and degradation,hsa04010;hsa04141;hsa04144;hsa04213;hsa04915;hsa05145;hsa05162;hsa05164;hsa05169,HSPA8,NA,HSPA8 - mRNA,False\nCFL1 - mRNA,NM_005507.2:225,SEBELIUS_2_C4636,CFL1,mRNA,0,,,EPHB-mediated forward signaling;Sema3A PAK dependent Axon repulsion;Platelet degranulation ;Regulation of actin dynamics for phagocytic cup formation,hsa04360;hsa04666;hsa04810;hsa05133,CFL1,NA,CFL1 - mRNA,False\nFLNA - mRNA,NM_001456.3:7335,SEBELIUS_2_C4636,FLNA,mRNA,0,,,RHO GTPases activate PAKs;Platelet degranulation ;Cell-extracellular matrix interactions,hsa04010;hsa04510;hsa05132;hsa05205,FLNA,NA,FLNA - mRNA,False\nMYL12B - mRNA,NM_001144945.1:749,SEBELIUS_2_C4636,MYL12B,mRNA,0,,,EPHA-mediated growth cone collapse;RHO GTPases Activate ROCKs;Smooth Muscle Contraction;RHO GTPases activate CIT;RHO GTPases activate PAKs;RHO GTPases activate PKNs;Sema4D induced cell migration and growth-cone collapse,hsa04510;hsa04530;hsa04611;hsa04670;hsa04810,MYL12B,NA,MYL12B - mRNA,False\nMYH9 - mRNA,NM_002473.3:3153,SEBELIUS_2_C4636,MYH9,mRNA,0,,,EPHA-mediated growth cone collapse;RHO GTPases Activate ROCKs;RHO GTPases activate CIT;RHO GTPases activate PAKs;RHO GTPases activate PKNs;Regulation of actin dynamics for phagocytic cup formation;Sema4D induced cell migration and growth-cone collapse,hsa04530;hsa04810;hsa05132,MYH9,NA,MYH9 - mRNA,False\nITGA1 - mRNA,NM_181501.1:1875,SEBELIUS_2_C4636,ITGA1,mRNA,0,,,Laminin interactions;Other semaphorin interactions;Integrin cell surface interactions,hsa04151;hsa04510;hsa04512;hsa04640;hsa04810;hsa05410;hsa05412;hsa05414,ITGA1,NA,ITGA1 - mRNA,False\nMYLK - mRNA,NM_053032.2:710,SEBELIUS_2_C4636,MYLK,mRNA,0,,,Smooth Muscle Contraction;RHO GTPases activate PAKs,hsa04020;hsa04022;hsa04270;hsa04510;hsa04611;hsa04810;hsa04921;hsa04971,MYLK,NA,MYLK - mRNA,False\nGNAI2 - mRNA,NM_002070.2:1000,SEBELIUS_2_C4636,GNAI2,mRNA,0,,,ADP signalling through P2Y purinoceptor 12;Adrenaline#noradrenaline inhibits insulin secretion,hsa04015;hsa04022;hsa04024;hsa04062;hsa04071;hsa04261;hsa04360;hsa04530;hsa04540;hsa04611;hsa04670;hsa04713;hsa04723;hsa04724;hsa04725;hsa04726;hsa04727;hsa04728;hsa04730;hsa04914;hsa04915;hsa04916;hsa04921;hsa04923;hsa04971;hsa05032;hsa05034;hsa05133;hsa05142;hsa05145;hsa05200,GNAI2,NA,GNAI2 - mRNA,False\nLDHA - mRNA,NM_001165414.1:1690,SEBELIUS_2_C4636,LDHA,mRNA,0,,,,hsa01100;hsa04066;hsa04922;hsa05230,LDHA,NA,LDHA - mRNA,False\nCCND1 - mRNA,NM_053056.2:2725,SEBELIUS_2_C4636,CCND1,mRNA,0,,,Cyclin D associated events in G1;Pre-NOTCH Transcription and Translation,hsa04068;hsa04110;hsa04115;hsa04151;hsa04152;hsa04310;hsa04390;hsa04510;hsa04630;hsa04917;hsa04919;hsa04921;hsa04933;hsa05161;hsa05162;hsa05166;hsa05200;hsa05203;hsa05205;hsa05206;hsa05210;hsa05212;hsa05213;hsa05214;hsa05215;hsa05216;hsa05218;hsa05219;hsa05220;hsa05221;hsa05222;hsa05223;hsa05416,CCND1,NA,CCND1 - mRNA,False\nCD44 - mRNA,NM_001001392.1:429,SEBELIUS_2_C4636,CD44,mRNA,0,,,Hyaluronan uptake and degradation;Cell surface interactions at the vascular wall,hsa04512;hsa04640;hsa05131;hsa05169;hsa05205;hsa05206,CD44,NA,CD44 - mRNA,False\nMYL9 - mRNA,NM_181526.1:133,SEBELIUS_2_C4636,MYL9,mRNA,0,,,EPHA-mediated growth cone collapse;RHO GTPases Activate ROCKs;Smooth Muscle Contraction;RHO GTPases activate CIT;RHO GTPases activate PAKs;RHO GTPases activate PKNs;Sema4D induced cell migration and growth-cone collapse,hsa04022;hsa04024;hsa04270;hsa04510;hsa04530;hsa04670;hsa04810;hsa04921,MYL9,NA,MYL9 - mRNA,False\nITGB1 - mRNA,NM_033666.2:2000,SEBELIUS_2_C4636,ITGB1,mRNA,0,,,Laminin interactions;RHO GTPases Activate Formins;ECM proteoglycans;Fibronectin matrix formation;Integrin cell surface interactions;Immunoregulatory interactions between a Lymphoid and a non-Lymphoid cell;Localization of the PINCH-ILK-PARVIN complex to focal adhesions;Elastic fibre formation;Cell surface interactions at the vascular wall;Non-integrin membrane-ECM interactions;Cell-extracellular matrix interactions,hsa04015;hsa04145;hsa04151;hsa04360;hsa04510;hsa04512;hsa04514;hsa04611;hsa04670;hsa04810;hsa05100;hsa05130;hsa05131;hsa05133;hsa05140;hsa05145;hsa05200;hsa05205;hsa05222;hsa05410;hsa05412;hsa05414,ITGB1,NA,ITGB1 - mRNA,False\nRPLP0 - mRNA,NM_001002.3:250,SEBELIUS_2_C4636,RPLP0,mRNA,0,,,Nonsense Mediated Decay (NMD) enhanced by the Exon Junction Complex (EJC),,RPLP0,NA,RPLP0 - mRNA,False\nFN1 - mRNA,NM_212482.1:1776,SEBELIUS_2_C4636,FN1,mRNA,0,,,MAP2K and MAPK activation;p130Cas linkage to MAPK signaling for integrins;Platelet degranulation ;Molecules associated with elastic fibres;Syndecan interactions;ECM proteoglycans;Fibronectin matrix formation;Integrin cell surface interactions;RAFMAP kinase cascade;Downstream signal transduction;VEGFR2 mediated cell proliferation;Signaling by SCF-KIT;Frs2-mediated activation;ARMS-mediated activation;Gastrin-CREB signalling pathway via PKC and MAPK;Interleukin receptor SHC signaling;FCERI mediated MAPK activation;DAP12 signaling;Signaling by PDGF;VEGFA-VEGFR2 Pathway;Signaling by ERBB2;IRS-related events triggered by IGF1R;Integrin alphaIIb beta3 signaling;Non-integrin membrane-ECM interactions;Elastic fibre formation,hsa04151;hsa04510;hsa04512;hsa04810;hsa04933;hsa05100;hsa05146;hsa05200;hsa05205;hsa05222,FN1,NA,FN1 - mRNA,False\nCYFIP2 - mRNA,NM_001037332.2:4990,SEBELIUS_2_C4636,CYFIP2,mRNA,0,,,RHO GTPases Activate WASPs and WAVEs;VEGFA-VEGFR2 Pathway;Regulation of actin dynamics for phagocytic cup formation,hsa04810,CYFIP2,NA,CYFIP2 - mRNA,False\nMYL12A b - mRNA,NM_001303048.1:27,SEBELIUS_2_C4636,MYL12A b,mRNA,0,,NM_006471.2:78;NM_001303049.1:9,Smooth Muscle Contraction;Ephrin signaling,hsa04510;hsa04530;hsa04611;hsa04670;hsa04810,MYL12A,NA,MYL12A b - mRNA,False\nNLK - mRNA,NM_016231.4:618,SEBELIUS_2_C4636,NLK,mRNA,0,,,Ca2.PLS. pathway,hsa04010;hsa04068;hsa04310;hsa04520,NLK,NA,NLK - mRNA,False\nLEF1 - mRNA,NM_016269.3:1165,SEBELIUS_2_C4636,LEF1,mRNA,0,,,Ca2.PLS. pathway;Binding of TCFLEF.CL.CTNNB1 to target gene promoters;Deactivation of the beta-catenin transactivating complex;Repression of WNT target genes;Formation of the beta-catenin.CL.TCF transactivating complex,hsa04310;hsa04390;hsa04520;hsa04916;hsa05200;hsa05210;hsa05213;hsa05215;hsa05216;hsa05217;hsa05221;hsa05412,LEF1,NA,LEF1 - mRNA,False\nCLDN23 - mRNA,NM_194284.2:1768,SEBELIUS_2_C4636,CLDN23,mRNA,0,,,Tight junction interactions,hsa04514;hsa04530;hsa04670;hsa05160,CLDN23,NA,CLDN23 - mRNA,False\nOCLN - mRNA,NM_002538.3:5130,SEBELIUS_2_C4636,OCLN,mRNA,0,,,Apoptotic cleavage of cellular proteins,hsa04514;hsa04530;hsa04670;hsa05130;hsa05160,OCLN,NA,OCLN - mRNA,False\nMYL5 - mRNA,NM_002477.1:398,SEBELIUS_2_C4636,MYL5,mRNA,0,,,Smooth Muscle Contraction,hsa04510;hsa04530;hsa04670;hsa04810,MYL5,NA,MYL5 - mRNA,False\nINADL - mRNA,XM_005270347.1:343,SEBELIUS_2_C4636,INADL,mRNA,0,,,,hsa04390;hsa04530,INADL,NA,INADL - mRNA,False\nTJP2 - mRNA,NM_004817.2:2785,SEBELIUS_2_C4636,TJP2,mRNA,0,,,Apoptotic cleavage of cellular proteins,hsa04530;hsa05110,TJP2,NA,TJP2 - mRNA,False\nARAF - mRNA,NM_001654.3:245,SEBELIUS_2_C4636,ARAF,mRNA,0,,,MAP2K and MAPK activation;RAF activation;Negative regulation of MAPK pathway;RAFMAP kinase cascade;Downstream signal transduction;VEGFR2 mediated cell proliferation;Signaling by SCF-KIT;Frs2-mediated activation;ARMS-mediated activation;Gastrin-CREB signalling pathway via PKC and MAPK;Interleukin receptor SHC signaling;FCERI mediated MAPK activation;DAP12 signaling;Signaling by PDGF;VEGFA-VEGFR2 Pathway;Signaling by ERBB2;IRS-related events triggered by IGF1R,hsa04012;hsa04068;hsa04270;hsa04650;hsa04720;hsa04726;hsa04730;hsa04810;hsa04910;hsa04914;hsa05034;hsa05160;hsa05200;hsa05205;hsa05210;hsa05211;hsa05212;hsa05213;hsa05214;hsa05215;hsa05218;hsa05219;hsa05220;hsa05221;hsa05223,ARAF,NA,ARAF - mRNA,False\nMYL12A c - mRNA,NM_001303049.1:9,SEBELIUS_2_C4636,MYL12A c,mRNA,0,,NM_006471.2:78;NM_001303048.1:27,EPHA-mediated growth cone collapse;RHO GTPases Activate ROCKs;Smooth Muscle Contraction;RHO GTPases activate CIT;RHO GTPases activate PAKs;RHO GTPases activate PKNs;Sema4D induced cell migration and growth-cone collapse;Ephrin signaling,hsa04510;hsa04530;hsa04611;hsa04670;hsa04810,MYL12A,NA,MYL12A c - mRNA,False\nMAGI1 - mRNA,NM_004742.2:325,SEBELIUS_2_C4636,MAGI1,mRNA,0,,,,hsa04015;hsa04530,MAGI1,NA,MAGI1 - mRNA,False\nHSPA1A - mRNA,NM_005345.5:98,SEBELIUS_2_C4636,HSPA1A,mRNA,0,,,Regulation of HSF1-mediated heat shock response;AUF1 (hnRNP D0) binds and destabilizes mRNA,hsa04010;hsa04141;hsa04144;hsa04213;hsa04915;hsa05020;hsa05145;hsa05162;hsa05164;hsa05169,HSPA1A,NA,HSPA1A - mRNA,False\nAXIN1 - mRNA,NM_181050.1:2742,SEBELIUS_2_C4636,AXIN1,mRNA,0,,,APC truncation mutants have impaired AXIN binding;Misspliced GSK3beta mutants stabilize beta-catenin;T41 mutants of beta-catenin aren.SQT.t phosphorylated;AXIN missense mutants destabilize the destruction complex;S45 mutants of beta-catenin aren.SQT.t phosphorylated;Disassembly of the destruction complex and recruitment of AXIN to the membrane;S37 mutants of beta-catenin aren.SQT.t phosphorylated;S33 mutants of beta-catenin aren.SQT.t phosphorylated;Beta-catenin phosphorylation cascade;Truncations of AMER1 destabilize the destruction complex,hsa04310;hsa04390;hsa04550;hsa05200;hsa05210;hsa05213;hsa05217,AXIN1,NA,AXIN1 - mRNA,False\nBAIAP2 - mRNA,NM_017450.2:653,SEBELIUS_2_C4636,BAIAP2,mRNA,0,,,RHO GTPases Activate WASPs and WAVEs;VEGFA-VEGFR2 Pathway;Regulation of actin dynamics for phagocytic cup formation,hsa04520;hsa04810,BAIAP2,NA,BAIAP2 - mRNA,False\nRAC3 - mRNA,NM_005052.2:702,SEBELIUS_2_C4636,RAC3,mRNA,0,,,Rho GTPase cycle;PCPCE pathway,hsa04010;hsa04014;hsa04015;hsa04024;hsa04071;hsa04310;hsa04360;hsa04370;hsa04510;hsa04520;hsa04650;hsa04662;hsa04664;hsa04810;hsa05200;hsa05210;hsa05212;hsa05231;hsa05416,RAC3,NA,RAC3 - mRNA,False\nRAC2 - mRNA,NM_002872.4:341,SEBELIUS_2_C4636,RAC2,mRNA,0,,,Rho GTPase cycle;GPVI-mediated activation cascade;G alpha (1213) signalling events;PCPCE pathway;Sema4D induced cell migration and growth-cone collapse,hsa04010;hsa04014;hsa04015;hsa04024;hsa04062;hsa04071;hsa04310;hsa04360;hsa04370;hsa04510;hsa04520;hsa04650;hsa04662;hsa04664;hsa04666;hsa04670;hsa04810;hsa05200;hsa05210;hsa05212;hsa05231;hsa05416,RAC2,NA,RAC2 - mRNA,False\nWASF1 - mRNA,NM_001024934.1:1425,SEBELIUS_2_C4636,WASF1,mRNA,0,,,RHO GTPases Activate WASPs and WAVEs;VEGFA-VEGFR2 Pathway;Regulation of actin dynamics for phagocytic cup formation,hsa04520;hsa04666;hsa04810;hsa05100;hsa05131;hsa05132;hsa05231,WASF1,NA,WASF1 - mRNA,False\nGUSB - mRNA,NM_000181.1:1350,SEBELIUS_2_C4636,GUSB,mRNA,0,,,,hsa01100,GUSB,NA,GUSB - mRNA,False\nDNMT3A - mRNA,NM_022552.4:2056,SEBELIUS_2_C4636,DNMT3A,mRNA,0,,,,hsa01100;hsa05206,DNMT3A,NA,DNMT3A - mRNA,False\nAPC - mRNA,NM_000038.3:6850,SEBELIUS_2_C4636,APC,mRNA,0,,,Apoptotic cleavage of cellular proteins;Misspliced GSK3beta mutants stabilize beta-catenin;APC truncation mutants have impaired AXIN binding;T41 mutants of beta-catenin aren.SQT.t phosphorylated;AXIN missense mutants destabilize the destruction complex;S45 mutants of beta-catenin aren.SQT.t phosphorylated;Deactivation of the beta-catenin transactivating complex;Disassembly of the destruction complex and recruitment of AXIN to the membrane;S37 mutants of beta-catenin aren.SQT.t phosphorylated;S33 mutants of beta-catenin aren.SQT.t phosphorylated;Beta-catenin phosphorylation cascade;Truncations of AMER1 destabilize the destruction complex,hsa04310;hsa04390;hsa04550;hsa04810;hsa05166;hsa05200;hsa05206;hsa05210;hsa05213;hsa05217,APC,NA,APC - mRNA,False\nLIMK2 - mRNA,NM_001031801.1:1590,SEBELIUS_2_C4636,LIMK2,mRNA,0,,,EPHB-mediated forward signaling;RHO GTPases Activate ROCKs;Sema4D induced cell migration and growth-cone collapse,hsa04360;hsa04666;hsa04810,LIMK2,NA,LIMK2 - mRNA,False\nEPB41L1 - mRNA,NM_001258330.1:900,SEBELIUS_2_C4636,EPB41L1,mRNA,0,,,,hsa04530,EPB41L1,NA,EPB41L1 - mRNA,False\nEXOC3 - mRNA,NM_007277.4:1820,SEBELIUS_2_C4636,EXOC3,mRNA,0,,,Translocation of GLUT4 to the plasma membrane,hsa04530,EXOC3,NA,EXOC3 - mRNA,False\nTARBP2 - mRNA,NM_134323.1:1060,SEBELIUS_2_C4636,TARBP2,mRNA,0,,,MicroRNA (miRNA) biogenesis,,TARBP2,NA,TARBP2 - mRNA,False\nDVL2 - mRNA,NM_004422.2:1025,SEBELIUS_2_C4636,DVL2,mRNA,0,,,Disassembly of the destruction complex and recruitment of AXIN to the membrane;WNT mediated activation of DVL;Negative regulation of TCF-dependent signaling by DVL-interacting proteins;PCPCE pathway,hsa04310;hsa04330;hsa04390;hsa04550;hsa04916;hsa05166;hsa05200;hsa05217,DVL2,NA,DVL2 - mRNA,False\nCDK4 - mRNA,NM_000075.2:1055,SEBELIUS_2_C4636,CDK4,mRNA,0,,,Cyclin D associated events in G1;Oncogene Induced Senescence;Ubiquitin-dependent degradation of Cyclin D1;RMTs methylate histone arginines,hsa04110;hsa04115;hsa04151;hsa04530;hsa04660;hsa04933;hsa05161;hsa05162;hsa05166;hsa05200;hsa05203;hsa05212;hsa05214;hsa05218;hsa05219;hsa05220;hsa05222;hsa05223,CDK4,NA,CDK4 - mRNA,False\nMAGI3 - mRNA,NM_152900.2:6685,SEBELIUS_2_C4636,MAGI3,mRNA,0,,,,hsa04015;hsa04530,MAGI3,NA,MAGI3 - mRNA,False\nTCF7L1 - mRNA,NM_031283.1:2215,SEBELIUS_2_C4636,TCF7L1,mRNA,0,,,Ca2.PLS. pathway;Binding of TCFLEF.CL.CTNNB1 to target gene promoters;Deactivation of the beta-catenin transactivating complex;Repression of WNT target genes;Formation of the beta-catenin.CL.TCF transactivating complex,hsa04310;hsa04390;hsa04520;hsa04916;hsa05200;hsa05210;hsa05213;hsa05215;hsa05216;hsa05217;hsa05221;hsa05412,TCF7L1,NA,TCF7L1 - mRNA,False\nPIK3R1 - mRNA,NM_181504.2:1105,SEBELIUS_2_C4636,PIK3R1,mRNA,0,,,G alpha (q) signalling events;Downstream TCR signaling;Signaling by FGFR2 in disease;PIP3 activates AKT signaling;VEGFA-VEGFR2 Pathway;Synthesis of PIPs at the plasma membrane;Constitutive Signaling by Ligand-Responsive EGFR Cancer Variants;Interleukin receptor SHC signaling;G beta.CL.gamma signalling through PI3Kgamma;Regulation of signaling by CBL;Role of phospholipids in phagocytosis;Signaling by FGFR3 in disease;Constitutive Signaling by Aberrant PI3K in Cancer;Signaling by FGFR4 in disease;CD28 dependent PI3KAkt signaling;Tie2 Signaling;Signaling by FGFR1 fusion mutants;GP1b-IX-V activation signalling;GPVI-mediated activation cascade;Nephrin interactions;Constitutive Signaling by EGFRvIII;Gastrin-CREB signalling pathway via PKC and MAPK;Downstream signal transduction;GAB1 signalosome;Signaling by SCF-KIT;PI3KAKT activation;Role of LAT2NTALLAB on calcium mobilization;DAP12 signaling;Signaling by PDGF;Signaling by ERBB2;IRS-related events triggered by IGF1R;CD28 co-stimulation;Cell surface interactions at the vascular wall;Signaling by FGFR1 in disease,hsa04012;hsa04014;hsa04015;hsa04022;hsa04024;hsa04062;hsa04066;hsa04068;hsa04070;hsa04071;hsa04072;hsa04150;hsa04151;hsa04152;hsa04210;hsa04211;hsa04213;hsa04261;hsa04370;hsa04380;hsa04510;hsa04550;hsa04611;hsa04620;hsa04630;hsa04650;hsa04660;hsa04662;hsa04664;hsa04666;hsa04668;hsa04670;hsa04722;hsa04725;hsa04750;hsa04810;hsa04910;hsa04914;hsa04915;hsa04917;hsa04919;hsa04921;hsa04923;hsa04930;hsa04931;hsa04932;hsa04933;hsa04960;hsa04973;hsa05100;hsa05142;hsa05145;hsa05146;hsa05160;hsa05161;hsa05162;hsa05164;hsa05166;hsa05169;hsa05200;hsa05203;hsa05205;hsa05210;hsa05211;hsa05212;hsa05213;hsa05214;hsa05215;hsa05218;hsa05220;hsa05221;hsa05222;hsa05223;hsa05230;hsa05231,PIK3R1,NA,PIK3R1 - mRNA,False\nPAK4 - mRNA,NM_005884.3:2130,SEBELIUS_2_C4636,PAK4,mRNA,0,,,Activation of Rac,hsa04012;hsa04014;hsa04360;hsa04510;hsa04660;hsa04810;hsa05206;hsa05211,PAK4,NA,PAK4 - mRNA,False\nXPO5 - mRNA,NM_020750.2:375,SEBELIUS_2_C4636,XPO5,mRNA,0,,,MicroRNA (miRNA) biogenesis,,XPO5,NA,XPO5 - mRNA,False\nHRAS - mRNA,NM_005343.2:396,SEBELIUS_2_C4636,HRAS,mRNA,0,,,Signaling by FGFR2 in disease;Regulation of RAS by GAPs;SHC-mediated cascade.CL.FGFR3;Signaling by FGFR3 in disease;MAP2K and MAPK activation;SHC-related events triggered by IGF1R;Signaling by FGFR4 in disease;SHC-mediated cascade.CL.FGFR1;Tie2 Signaling;Signaling by FGFR1 in disease;Activation of RAS in B cells;SHC-mediated cascade.CL.FGFR4;Constitutive Signaling by EGFRvIII;CD209 (DC-SIGN) signaling;Ras activation uopn Ca2.PLS. infux through NMDA receptor;FCERI mediated MAPK activation;RAFMAP kinase cascade;Downstream signal transduction;VEGFR2 mediated cell proliferation;Signaling by SCF-KIT;Frs2-mediated activation;ARMS-mediated activation;Gastrin-CREB signalling pathway via PKC and MAPK;Interleukin receptor SHC signaling;DAP12 signaling;Signaling by PDGF;VEGFA-VEGFR2 Pathway;Signaling by ERBB2;IRS-related events triggered by IGF1R;Cell surface interactions at the vascular wall;CREB phosphorylation through the activation of Ras,hsa04010;hsa04012;hsa04014;hsa04015;hsa04062;hsa04068;hsa04071;hsa04072;hsa04144;hsa04151;hsa04211;hsa04213;hsa04360;hsa04370;hsa04510;hsa04530;hsa04540;hsa04550;hsa04630;hsa04650;hsa04660;hsa04662;hsa04664;hsa04720;hsa04722;hsa04725;hsa04726;hsa04730;hsa04810;hsa04910;hsa04912;hsa04915;hsa04916;hsa04917;hsa04919;hsa04921;hsa04933;hsa05034;hsa05160;hsa05161;hsa05166;hsa05200;hsa05203;hsa05205;hsa05206;hsa05211;hsa05213;hsa05214;hsa05215;hsa05216;hsa05218;hsa05219;hsa05220;hsa05221;hsa05223;hsa05230;hsa05231,HRAS,NA,HRAS - mRNA,False\nERBB2 - mRNA,NM_001005862.1:1255,SEBELIUS_2_C4636,ERBB2,mRNA,0,,,RAFMAP kinase cascade;GRB7 events in ERBB2 signaling;PLCG1 events in ERBB2 signaling;Sema4D induced cell migration and growth-cone collapse;Signaling by ERBB2;Downstream signal transduction;GAB1 signalosome;Signaling by SCF-KIT;PI3KAKT activation;Role of LAT2NTALLAB on calcium mobilization;DAP12 signaling;Signaling by PDGF;VEGFR2 mediated cell proliferation;Frs2-mediated activation;ARMS-mediated activation;Gastrin-CREB signalling pathway via PKC and MAPK;Interleukin receptor SHC signaling;FCERI mediated MAPK activation;VEGFA-VEGFR2 Pathway;IRS-related events triggered by IGF1R,hsa04012;hsa04020;hsa04066;hsa04510;hsa04520;hsa05200;hsa05205;hsa05206;hsa05212;hsa05213;hsa05215;hsa05219;hsa05223;hsa05230,ERBB2,NA,ERBB2 - mRNA,False\nPIK3CB - mRNA,NM_006219.1:2945,SEBELIUS_2_C4636,PIK3CB,mRNA,0,,,Downstream TCR signaling;PIP3 activates AKT signaling;Constitutive Signaling by Aberrant PI3K in Cancer;Tie2 Signaling;VEGFA-VEGFR2 Pathway;Synthesis of PIPs at the plasma membrane;Interleukin receptor SHC signaling;G beta.CL.gamma signalling through PI3Kgamma;Regulation of signaling by CBL;GPVI-mediated activation cascade;Role of phospholipids in phagocytosis;Nephrin interactions;PI3K Cascade;Downstream signal transduction;GAB1 signalosome;Signaling by SCF-KIT;PI3KAKT activation;Role of LAT2NTALLAB on calcium mobilization;DAP12 signaling;Signaling by PDGF;Signaling by ERBB2;Cell surface interactions at the vascular wall;IRS-related events triggered by IGF1R,hsa00562;hsa04012;hsa04014;hsa04015;hsa04022;hsa04024;hsa04062;hsa04066;hsa04068;hsa04070;hsa04071;hsa04072;hsa04150;hsa04151;hsa04152;hsa04210;hsa04211;hsa04213;hsa04261;hsa04370;hsa04380;hsa04510;hsa04550;hsa04611;hsa04620;hsa04630;hsa04650;hsa04660;hsa04662;hsa04664;hsa04666;hsa04668;hsa04670;hsa04722;hsa04725;hsa04750;hsa04810;hsa04910;hsa04914;hsa04915;hsa04917;hsa04919;hsa04921;hsa04923;hsa04930;hsa04931;hsa04932;hsa04933;hsa04960;hsa04973;hsa05100;hsa05142;hsa05145;hsa05146;hsa05160;hsa05161;hsa05162;hsa05164;hsa05166;hsa05169;hsa05200;hsa05203;hsa05205;hsa05210;hsa05211;hsa05212;hsa05213;hsa05214;hsa05215;hsa05218;hsa05220;hsa05221;hsa05222;hsa05223;hsa05230;hsa05231,PIK3CB,NA,PIK3CB - mRNA,False\nBAIAP2L1 - mRNA,NM_018842.3:259,SEBELIUS_2_C4636,BAIAP2L1,mRNA,0,,,,,BAIAP2L1,NA,BAIAP2L1 - mRNA,False\nDVL1 - mRNA,NM_004421.2:789,SEBELIUS_2_C4636,DVL1,mRNA,0,,,Disassembly of the destruction complex and recruitment of AXIN to the membrane;WNT mediated activation of DVL;RHO GTPases Activate Formins;PCPCE pathway,hsa04310;hsa04330;hsa04390;hsa04550;hsa04916;hsa05166;hsa05200;hsa05217,DVL1,NA,DVL1 - mRNA,False\nBAD - mRNA,NM_004322.3:963,SEBELIUS_2_C4636,BAD,mRNA,0,,,Activation of BAD and translocation to mitochondria ;NRAGE signals death through JNK;Constitutive Signaling by AKT1 E17K in Cancer;AKT phosphorylates targets in the cytosol;PIP3 activates AKT signaling;Downstream signal transduction;GAB1 signalosome;Signaling by SCF-KIT;PI3KAKT activation;Role of LAT2NTALLAB on calcium mobilization;DAP12 signaling;Signaling by PDGF;Signaling by ERBB2,hsa04012;hsa04014;hsa04022;hsa04024;hsa04151;hsa04210;hsa04370;hsa04510;hsa04722;hsa04910;hsa04919;hsa05010;hsa05145;hsa05152;hsa05160;hsa05161;hsa05200;hsa05203;hsa05210;hsa05212;hsa05213;hsa05215;hsa05218;hsa05220;hsa05221;hsa05223,BAD,NA,BAD - mRNA,False\nBHLHE40 - mRNA,NM_003670.1:560,SEBELIUS_2_C4636,BHLHE40,mRNA,0,,,Circadian Clock,,BHLHE40,NA,BHLHE40 - mRNA,False\nMET - mRNA,NM_001127500.1:1925,SEBELIUS_2_C4636,MET,mRNA,0,,,,hsa04014;hsa04015;hsa04060;hsa04144;hsa04151;hsa04360;hsa04510;hsa04520;hsa05100;hsa05120;hsa05200;hsa05202;hsa05205;hsa05206;hsa05211;hsa05218;hsa05230,MET,NA,MET - mRNA,False\nMYC - mRNA,NM_002467.3:1610,SEBELIUS_2_C4636,MYC,mRNA,0,,,Constitutive Signaling by NOTCH1 PEST Domain Mutants;SMAD2SMAD3.CL.SMAD4 heterotrimer regulates transcription;Formation of the beta-catenin.CL.TCF transactivating complex,hsa04010;hsa04012;hsa04110;hsa04151;hsa04310;hsa04350;hsa04390;hsa04550;hsa04630;hsa04919;hsa05161;hsa05166;hsa05169;hsa05200;hsa05202;hsa05205;hsa05206;hsa05210;hsa05213;hsa05216;hsa05219;hsa05220;hsa05221;hsa05222;hsa05230,MYC,NA,MYC - mRNA,False\nDIAPH3 - mRNA,NM_001042517.1:1832,SEBELIUS_2_C4636,DIAPH3,mRNA,0,,,RHO GTPases Activate Formins,hsa04810,DIAPH3,NA,DIAPH3 - mRNA,False\nCCND2 - mRNA,NM_001759.3:503,SEBELIUS_2_C4636,CCND2,mRNA,0,,,Cyclin D associated events in G1,hsa04068;hsa04110;hsa04115;hsa04151;hsa04310;hsa04390;hsa04510;hsa04630;hsa04917;hsa05162;hsa05166;hsa05202;hsa05203;hsa05206,CCND2,NA,CCND2 - mRNA,False\nPDCD4 - mRNA,NM_014456.3:1115,SEBELIUS_2_C4636,PDCD4,mRNA,0,,,,hsa05205;hsa05206,PDCD4,NA,PDCD4 - mRNA,False\nPIK3R3 - mRNA,NM_003629.3:1800,SEBELIUS_2_C4636,PIK3R3,mRNA,0,,,G alpha (q) signalling events;PIP3 activates AKT signaling;Constitutive Signaling by Aberrant PI3K in Cancer;CD28 dependent PI3KAkt signaling;Synthesis of PIPs at the plasma membrane;Interleukin receptor SHC signaling;G beta.CL.gamma signalling through PI3Kgamma;Regulation of signaling by CBL;GPVI-mediated activation cascade;G alpha (1213) signalling events;Gastrin-CREB signalling pathway via PKC and MAPK;Downstream signal transduction;GAB1 signalosome;Signaling by SCF-KIT;PI3KAKT activation;Role of LAT2NTALLAB on calcium mobilization;DAP12 signaling;Signaling by PDGF;Signaling by ERBB2;CD28 co-stimulation,hsa04012;hsa04014;hsa04015;hsa04022;hsa04024;hsa04062;hsa04066;hsa04068;hsa04070;hsa04071;hsa04072;hsa04150;hsa04151;hsa04152;hsa04210;hsa04211;hsa04213;hsa04261;hsa04370;hsa04380;hsa04510;hsa04550;hsa04611;hsa04620;hsa04630;hsa04650;hsa04660;hsa04662;hsa04664;hsa04666;hsa04668;hsa04670;hsa04722;hsa04725;hsa04750;hsa04810;hsa04910;hsa04914;hsa04915;hsa04917;hsa04919;hsa04921;hsa04923;hsa04930;hsa04931;hsa04932;hsa04933;hsa04960;hsa04973;hsa05100;hsa05142;hsa05145;hsa05146;hsa05160;hsa05161;hsa05162;hsa05164;hsa05166;hsa05169;hsa05200;hsa05203;hsa05205;hsa05210;hsa05211;hsa05212;hsa05213;hsa05214;hsa05215;hsa05218;hsa05220;hsa05221;hsa05222;hsa05223;hsa05230;hsa05231,PIK3R3,NA,PIK3R3 - mRNA,False\nWASL - mRNA,NM_003941.2:400,SEBELIUS_2_C4636,WASL,mRNA,0,,,EPHB-mediated forward signaling;RHO GTPases Activate WASPs and WAVEs;DCC mediated attractive signaling;Nephrin interactions;Regulation of actin dynamics for phagocytic cup formation,hsa04062;hsa04144;hsa04520;hsa04666;hsa04810;hsa05100;hsa05130;hsa05131;hsa05132;hsa05231,WASL,NA,WASL - mRNA,False\nTGFB2 - mRNA,NM_003238.2:1125,SEBELIUS_2_C4636,TGFB2,mRNA,0,,,Molecules associated with elastic fibres;ECM proteoglycans;Platelet degranulation ;Elastic fibre formation,hsa04010;hsa04060;hsa04068;hsa04110;hsa04144;hsa04350;hsa04380;hsa04390;hsa04933;hsa05140;hsa05142;hsa05145;hsa05146;hsa05152;hsa05161;hsa05166;hsa05200;hsa05205;hsa05206;hsa05210;hsa05211;hsa05212;hsa05220;hsa05321;hsa05410;hsa05414,TGFB2,NA,TGFB2 - mRNA,False\nHSPA1B - mRNA,NM_005346.3:2095,SEBELIUS_2_C4636,HSPA1B,mRNA,0,,,Regulation of HSF1-mediated heat shock response;AUF1 (hnRNP D0) binds and destabilizes mRNA,hsa04010;hsa04141;hsa04144;hsa04213;hsa04915;hsa05145;hsa05162;hsa05164;hsa05169,HSPA1B,NA,HSPA1B - mRNA,False\nVAV2 - mRNA,NM_001134398.1:1615,SEBELIUS_2_C4636,VAV2,mRNA,0,,,Rho GTPase cycle;DAP12 signaling;Signal transduction by L1;EPH-ephrin mediated repulsion of cells;VEGFR2 mediated vascular permeability;NRAGE signals death through JNK;GPVI-mediated activation cascade;G alpha (1213) signalling events;FCERI mediated MAPK activation;Regulation of actin dynamics for phagocytic cup formation;VEGFA-VEGFR2 Pathway,hsa04015;hsa04024;hsa04062;hsa04510;hsa04650;hsa04660;hsa04662;hsa04664;hsa04666;hsa04670;hsa04810;hsa05205,VAV2,NA,VAV2 - mRNA,False\nAGO2 - mRNA,NM_012154.2:625,SEBELIUS_2_C4636,AGO2,mRNA,0,,,Pre-NOTCH Transcription and Translation;PIP3 activates AKT signaling;Post-transcriptional silencing by small RNAs;MicroRNA (miRNA) biogenesis;Small interfering RNA (siRNA) biogenesis;Downstream signal transduction;GAB1 signalosome;Signaling by SCF-KIT;PI3KAKT activation;Role of LAT2NTALLAB on calcium mobilization;DAP12 signaling;Signaling by PDGF;Signaling by ERBB2,,AGO2,NA,AGO2 - mRNA,False\nMLLT4 - mRNA,NM_005936.2:1560,SEBELIUS_2_C4636,MLLT4,mRNA,0,,,Adherens junctions interactions,hsa04014;hsa04015;hsa04024;hsa04520;hsa04530;hsa04670,MLLT4,NA,MLLT4 - mRNA,False\nJUN - mRNA,NM_002228.3:140,SEBELIUS_2_C4636,JUN,mRNA,0,,,Activation of the AP-1 family of transcription factors;MAPK6MAPK4 signaling;Pre-NOTCH Transcription and Translation;Oxidative Stress Induced Senescence;FCERI mediated MAPK activation;Senescence-Associated Secretory Phenotype (SASP),hsa04010;hsa04012;hsa04024;hsa04310;hsa04380;hsa04510;hsa04620;hsa04660;hsa04662;hsa04668;hsa04722;hsa04912;hsa04915;hsa04921;hsa04932;hsa04933;hsa05031;hsa05120;hsa05132;hsa05133;hsa05140;hsa05142;hsa05161;hsa05164;hsa05166;hsa05168;hsa05169;hsa05200;hsa05203;hsa05210;hsa05211;hsa05231;hsa05321,JUN,NA,JUN - mRNA,False\nRRAS - mRNA,NM_006270.3:330,SEBELIUS_2_C4636,RRAS,mRNA,0,,,CREB phosphorylation through the activation of Ras,hsa04010;hsa04014;hsa04015;hsa04024;hsa04072;hsa04530;hsa04810;hsa05166;hsa05205,RRAS,NA,RRAS - mRNA,False\nBCAR1 - mRNA,NM_001170717.1:1341,SEBELIUS_2_C4636,BCAR1,mRNA,0,,,Downstream signal transduction;p130Cas linkage to MAPK signaling for integrins;VEGFA-VEGFR2 Pathway;Signaling by PDGF;Integrin alphaIIb beta3 signaling,hsa04015;hsa04062;hsa04510;hsa04670;hsa04810;hsa05100,BCAR1,NA,BCAR1 - mRNA,False\nPIK3R2 - mRNA,NM_005027.2:3100,SEBELIUS_2_C4636,PIK3R2,mRNA,0,,,G alpha (q) signalling events;Downstream TCR signaling;PIP3 activates AKT signaling;VEGFA-VEGFR2 Pathway;Synthesis of PIPs at the plasma membrane;Rho GTPase cycle;Interleukin receptor SHC signaling;G beta.CL.gamma signalling through PI3Kgamma;Regulation of signaling by CBL;Role of phospholipids in phagocytosis;G alpha (1213) signalling events;PI3K Cascade;Constitutive Signaling by Aberrant PI3K in Cancer;CD28 dependent PI3KAkt signaling;Tie2 Signaling;GPVI-mediated activation cascade;Nephrin interactions;Gastrin-CREB signalling pathway via PKC and MAPK;Downstream signal transduction;GAB1 signalosome;Signaling by SCF-KIT;PI3KAKT activation;Role of LAT2NTALLAB on calcium mobilization;DAP12 signaling;Signaling by PDGF;Signaling by ERBB2;IRS-related events triggered by IGF1R;CD28 co-stimulation;Cell surface interactions at the vascular wall,hsa04012;hsa04014;hsa04015;hsa04022;hsa04024;hsa04062;hsa04066;hsa04068;hsa04070;hsa04071;hsa04072;hsa04150;hsa04151;hsa04152;hsa04210;hsa04211;hsa04213;hsa04261;hsa04370;hsa04380;hsa04510;hsa04550;hsa04611;hsa04620;hsa04630;hsa04650;hsa04660;hsa04662;hsa04664;hsa04666;hsa04668;hsa04670;hsa04722;hsa04725;hsa04750;hsa04810;hsa04910;hsa04914;hsa04915;hsa04917;hsa04919;hsa04921;hsa04923;hsa04930;hsa04931;hsa04932;hsa04933;hsa04960;hsa04973;hsa05100;hsa05142;hsa05145;hsa05146;hsa05160;hsa05161;hsa05162;hsa05164;hsa05166;hsa05169;hsa05200;hsa05203;hsa05205;hsa05206;hsa05210;hsa05211;hsa05212;hsa05213;hsa05214;hsa05215;hsa05218;hsa05220;hsa05221;hsa05222;hsa05223;hsa05230;hsa05231,PIK3R2,NA,PIK3R2 - mRNA,False\nPIK3CD - mRNA,NM_005026.3:2978,SEBELIUS_2_C4636,PIK3CD,mRNA,0,,,Synthesis of PIPs at the plasma membrane;Interleukin receptor SHC signaling;G beta.CL.gamma signalling through PI3Kgamma;PIP3 activates AKT signaling;Constitutive Signaling by Aberrant PI3K in Cancer;Regulation of signaling by CBL;GPVI-mediated activation cascade;Downstream signal transduction;GAB1 signalosome;Signaling by SCF-KIT;PI3KAKT activation;Role of LAT2NTALLAB on calcium mobilization;DAP12 signaling;Signaling by PDGF;Signaling by ERBB2,hsa00562;hsa04012;hsa04014;hsa04015;hsa04022;hsa04024;hsa04062;hsa04066;hsa04068;hsa04070;hsa04071;hsa04072;hsa04150;hsa04151;hsa04152;hsa04210;hsa04211;hsa04213;hsa04261;hsa04370;hsa04380;hsa04510;hsa04550;hsa04611;hsa04620;hsa04630;hsa04650;hsa04660;hsa04662;hsa04664;hsa04666;hsa04668;hsa04670;hsa04722;hsa04725;hsa04750;hsa04810;hsa04910;hsa04914;hsa04915;hsa04917;hsa04919;hsa04921;hsa04923;hsa04930;hsa04931;hsa04932;hsa04933;hsa04960;hsa04973;hsa05100;hsa05142;hsa05145;hsa05146;hsa05160;hsa05161;hsa05162;hsa05164;hsa05166;hsa05169;hsa05200;hsa05203;hsa05205;hsa05210;hsa05211;hsa05212;hsa05213;hsa05214;hsa05215;hsa05218;hsa05220;hsa05221;hsa05222;hsa05223;hsa05230;hsa05231,PIK3CD,NA,PIK3CD - mRNA,False\nABI2 - mRNA,NM_005759.5:1160,SEBELIUS_2_C4636,ABI2,mRNA,0,,,RHO GTPases Activate WASPs and WAVEs;VEGFA-VEGFR2 Pathway;Regulation of actin dynamics for phagocytic cup formation,hsa04810,ABI2,NA,ABI2 - mRNA,False\nTCF7L2 - mRNA,NM_001146274.1:1393,SEBELIUS_2_C4636,TCF7L2,mRNA,0,,,Ca2.PLS. pathway;Deactivation of the beta-catenin transactivating complex;Repression of WNT target genes;Synthesis# secretion# and inactivation of Glucagon-like Peptide-1 (GLP-1);Formation of the beta-catenin.CL.TCF transactivating complex,hsa04310;hsa04390;hsa04520;hsa04916;hsa05200;hsa05210;hsa05213;hsa05215;hsa05216;hsa05217;hsa05221;hsa05412,TCF7L2,NA,TCF7L2 - mRNA,False\nIGF1R - mRNA,NM_000875.3:10810,SEBELIUS_2_C4636,IGF1R,mRNA,0,,,IRS-related events triggered by IGF1R;SHC-related events triggered by IGF1R,hsa04014;hsa04015;hsa04066;hsa04068;hsa04114;hsa04144;hsa04151;hsa04152;hsa04211;hsa04213;hsa04510;hsa04520;hsa04550;hsa04730;hsa04914;hsa05200;hsa05202;hsa05205;hsa05214;hsa05215;hsa05218,IGF1R,NA,IGF1R - mRNA,False\nSRC - mRNA,NM_005417.3:1410,SEBELIUS_2_C4636,SRC,mRNA,0,,,FCGR activation;Spry regulation of FGF signaling;GAB1 signalosome;MAP2K and MAPK activation;CLEC7A (Dectin-1) signaling;CD28 co-stimulation;p130Cas linkage to MAPK signaling for integrins;CTLA4 inhibitory signaling;GRB2.CL.SOS provides linkage to MAPK signaling for Integrins ;Signal regulatory protein (SIRP) family interactions;PECAM1 interactions;Thrombin signalling through proteinase activated receptors (PARs);GP1b-IX-V activation signalling;c-src mediated regulation of Cx43 function and closure of gap junctions;Signaling by ERBB2;VEGFA-VEGFR2 Pathway;Signaling by SCF-KIT;VEGFR2 mediated cell proliferation;RAFMAP kinase cascade;Downstream signal transduction;Frs2-mediated activation;ARMS-mediated activation;Gastrin-CREB signalling pathway via PKC and MAPK;Interleukin receptor SHC signaling;FCERI mediated MAPK activation;DAP12 signaling;Signaling by PDGF;IRS-related events triggered by IGF1R;Negative regulation of FGFR3 signaling;Negative regulation of FGFR4 signaling;Negative regulation of FGFR2 signaling;Negative regulation of FGFR1 signaling;Integrin alphaIIb beta3 signaling;Cell surface interactions at the vascular wall,hsa04012;hsa04015;hsa04062;hsa04144;hsa04370;hsa04510;hsa04520;hsa04530;hsa04540;hsa04611;hsa04727;hsa04750;hsa04810;hsa04912;hsa04915;hsa04917;hsa04919;hsa04921;hsa05100;hsa05120;hsa05131;hsa05152;hsa05161;hsa05203;hsa05205;hsa05219,SRC,NA,SRC - mRNA,False\nCSNK2A2 - mRNA,NM_001896.2:1350,SEBELIUS_2_C4636,CSNK2A2,mRNA,0,,,WNT mediated activation of DVL;Signal transduction by L1,hsa04064;hsa04310;hsa04520;hsa04530;hsa05162;hsa05168;hsa05169,CSNK2A2,NA,CSNK2A2 - mRNA,False\nFYN - mRNA,NM_002037.3:765,SEBELIUS_2_C4636,FYN,mRNA,0,,,EPH-ephrin mediated repulsion of cells;Regulation of signaling by CBL;FCGR activation;Antigen activates B Cell Receptor (BCR) leading to generation of second messengers;Dectin-2 family;RAFMAP kinase cascade;Constitutive Signaling by Aberrant PI3K in Cancer;CTLA4 inhibitory signaling;CD28 dependent PI3KAkt signaling;PECAM1 interactions;CD28 dependent Vav1 pathway;GPVI-mediated activation cascade;Sema3A PAK dependent Axon repulsion;Nephrin interactions;SEMA3A-Plexin repulsion signaling by inhibiting Integrin adhesion;CRMPs in Sema3A signaling;CD209 (DC-SIGN) signaling;Downstream signal transduction;GAB1 signalosome;Signaling by SCF-KIT;PI3KAKT activation;Role of LAT2NTALLAB on calcium mobilization;DAP12 signaling;Signaling by PDGF;Signaling by ERBB2;VEGFA-VEGFR2 Pathway;VEGFR2 mediated cell proliferation;Frs2-mediated activation;ARMS-mediated activation;Gastrin-CREB signalling pathway via PKC and MAPK;Interleukin receptor SHC signaling;FCERI mediated MAPK activation;IRS-related events triggered by IGF1R;CD28 co-stimulation;Cell surface interactions at the vascular wall,hsa04071;hsa04072;hsa04360;hsa04380;hsa04510;hsa04520;hsa04611;hsa04650;hsa04660;hsa04664;hsa04725;hsa05020;hsa05130;hsa05162;hsa05416,FYN,NA,FYN - mRNA,False\nMAPK10 - mRNA,NM_138981.2:450,SEBELIUS_2_C4636,MAPK10,mRNA,0,,,Activation of the AP-1 family of transcription factors;Oxidative Stress Induced Senescence;FCERI mediated MAPK activation,hsa04010;hsa04012;hsa04014;hsa04024;hsa04068;hsa04071;hsa04141;hsa04310;hsa04380;hsa04510;hsa04620;hsa04621;hsa04664;hsa04668;hsa04722;hsa04723;hsa04728;hsa04750;hsa04910;hsa04912;hsa04914;hsa04917;hsa04920;hsa04930;hsa04931;hsa04932;hsa04933;hsa05120;hsa05131;hsa05132;hsa05133;hsa05142;hsa05145;hsa05152;hsa05160;hsa05161;hsa05164;hsa05168;hsa05169;hsa05200;hsa05210;hsa05212;hsa05231,MAPK10,NA,MAPK10 - mRNA,False\nMRAS - mRNA,NM_012219.3:4210,SEBELIUS_2_C4636,MRAS,mRNA,0,,,,hsa04010;hsa04014;hsa04015;hsa04072;hsa04530;hsa04810;hsa05166;hsa05205,MRAS,NA,MRAS - mRNA,False\nASH1L - mRNA,NM_018489.2:5005,SEBELIUS_2_C4636,ASH1L,mRNA,0,,,,hsa04530,ASH1L,NA,ASH1L - mRNA,False\nMAPK7 - mRNA,NM_139033.1:2850,SEBELIUS_2_C4636,MAPK7,mRNA,0,,,ERKs are inactivated;Gastrin-CREB signalling pathway via PKC and MAPK;Senescence-Associated Secretory Phenotype (SASP),hsa04010;hsa04540;hsa04722;hsa04912;hsa04921;hsa05206,MAPK7,NA,MAPK7 - mRNA,False\nBRAF - mRNA,NM_004333.3:565,SEBELIUS_2_C4636,BRAF,mRNA,0,,,Spry regulation of FGF signaling;MAP2K and MAPK activation;RAF activation;CREB phosphorylation through the activation of Ras;Negative feedback regulation of MAPK pathway;ARMS-mediated activation;Negative regulation of FGFR3 signaling;Negative regulation of FGFR4 signaling;Negative regulation of FGFR2 signaling;Negative regulation of FGFR1 signaling;RAFMAP kinase cascade;Downstream signal transduction;VEGFR2 mediated cell proliferation;Signaling by SCF-KIT;Frs2-mediated activation;Gastrin-CREB signalling pathway via PKC and MAPK;Interleukin receptor SHC signaling;FCERI mediated MAPK activation;DAP12 signaling;Signaling by PDGF;VEGFA-VEGFR2 Pathway;Signaling by ERBB2;IRS-related events triggered by IGF1R;Negative regulation of MAPK pathway,hsa04010;hsa04012;hsa04015;hsa04024;hsa04062;hsa04068;hsa04150;hsa04270;hsa04510;hsa04650;hsa04720;hsa04722;hsa04726;hsa04730;hsa04810;hsa04910;hsa04914;hsa05034;hsa05160;hsa05200;hsa05205;hsa05210;hsa05211;hsa05212;hsa05213;hsa05214;hsa05215;hsa05216;hsa05218;hsa05219;hsa05220;hsa05221;hsa05223,BRAF,NA,BRAF - mRNA,False\nPPP2R2D - mRNA,NM_001291310.1:2320,SEBELIUS_2_C4636,PPP2R2D,mRNA,0,,,MASTL Facilitates Mitotic Progression,hsa03015;hsa04071;hsa04151;hsa04152;hsa04261;hsa04390;hsa04530;hsa04728;hsa05142;hsa05160,PPP2R2D,NA,PPP2R2D - mRNA,False\nNUMA1 - mRNA,NM_006185.2:575,SEBELIUS_2_C4636,NUMA1,mRNA,0,,,,,NUMA1,NA,NUMA1 - mRNA,False\nTBP - mRNA,NM_001172085.1:587,SEBELIUS_2_C4636,TBP,mRNA,0,,,,hsa05016;hsa05166;hsa05168;hsa05169;hsa05203,TBP,NA,TBP - mRNA,False\nSMAD2 - mRNA,NM_005901.5:1678,SEBELIUS_2_C4636,SMAD2,mRNA,0,,,Transcriptional regulation of pluripotent stem cells;TGF-beta receptor signaling activates SMADs,hsa04068;hsa04110;hsa04144;hsa04350;hsa04390;hsa04520;hsa04550;hsa04933;hsa05142;hsa05166;hsa05200;hsa05205;hsa05210;hsa05212;hsa05321,SMAD2,NA,SMAD2 - mRNA,False\nDICER1 - mRNA,NM_177438.2:5060,SEBELIUS_2_C4636,DICER1,mRNA,0,,,MicroRNA (miRNA) biogenesis,hsa05206,DICER1,NA,DICER1 - mRNA,False\nLIMK1 - mRNA,NM_002314.3:3133,SEBELIUS_2_C4636,LIMK1,mRNA,0,,,EPHB-mediated forward signaling;RHO GTPases Activate ROCKs;RHO GTPases activate PAKs;Sema3A PAK dependent Axon repulsion;Regulation of actin dynamics for phagocytic cup formation;Sema4D induced cell migration and growth-cone collapse,hsa04360;hsa04666;hsa04810,LIMK1,NA,LIMK1 - mRNA,False\nPARD3 - mRNA,NM_019619.2:1060,SEBELIUS_2_C4636,PARD3,mRNA,0,,,TGF-beta receptor signaling in EMT (epithelial to mesenchymal transition);Tight junction interactions,hsa04015;hsa04062;hsa04144;hsa04390;hsa04520;hsa04530,PARD3,NA,PARD3 - mRNA,False\nCREBBP - mRNA,NM_004380.2:1301,SEBELIUS_2_C4636,CREBBP,mRNA,0,,,YAP1- and WWTR1 (TAZ)-stimulated gene expression;Activation of gene expression by SREBF (SREBP);Transcriptional activation of mitochondrial biogenesis;Constitutive Signaling by NOTCH1 HD.PLS.PEST Domain Mutants;Constitutive Signaling by NOTCH1 PEST Domain Mutants;CD209 (DC-SIGN) signaling;Circadian Clock,hsa04024;hsa04066;hsa04068;hsa04110;hsa04310;hsa04330;hsa04350;hsa04520;hsa04630;hsa04720;hsa04916;hsa04919;hsa04922;hsa05016;hsa05152;hsa05161;hsa05164;hsa05166;hsa05168;hsa05169;hsa05200;hsa05203;hsa05206;hsa05211;hsa05215,CREBBP,NA,CREBBP - mRNA,False\nPRKCA - mRNA,NM_002737.2:680,SEBELIUS_2_C4636,PRKCA,mRNA,0,,,WNT5A-dependent internalization of FZD4;Disinhibition of SNARE formation;Trafficking of AMPA receptors;Downstream signal transduction;EGFR interacts with phospholipase C-gamma;Phospholipase C-mediated cascade.CL. FGFR1;PLCG1 events in ERBB2 signaling;DAP12 signaling;PLC beta mediated events;Signaling by PDGF;Signaling by ERBB2;Signaling by SCF-KIT;Non-integrin membrane-ECM interactions;Gastrin-CREB signalling pathway via PKC and MAPK;VEGFA-VEGFR2 Pathway;PCPCE pathway,hsa04010;hsa04012;hsa04014;hsa04015;hsa04020;hsa04066;hsa04070;hsa04071;hsa04072;hsa04150;hsa04151;hsa04261;hsa04270;hsa04310;hsa04370;hsa04510;hsa04530;hsa04540;hsa04650;hsa04664;hsa04666;hsa04670;hsa04713;hsa04720;hsa04723;hsa04724;hsa04725;hsa04726;hsa04727;hsa04728;hsa04730;hsa04750;hsa04912;hsa04916;hsa04919;hsa04921;hsa04933;hsa04960;hsa04971;hsa04972;hsa05031;hsa05032;hsa05110;hsa05130;hsa05146;hsa05161;hsa05164;hsa05200;hsa05205;hsa05206;hsa05214;hsa05223;hsa05231,PRKCA,NA,PRKCA - mRNA,False\nRAB8B - mRNA,NM_016530.2:820,SEBELIUS_2_C4636,RAB8B,mRNA,0,,,,,RAB8B,NA,RAB8B - mRNA,False\nCTBP1 - mRNA,NM_001328.2:425,SEBELIUS_2_C4636,CTBP1,mRNA,0,,,Deactivation of the beta-catenin transactivating complex;Repression of WNT target genes,hsa04310;hsa04330;hsa05200;hsa05220,CTBP1,NA,CTBP1 - mRNA,False\nMAPK3 - mRNA,NM_001040056.1:580,SEBELIUS_2_C4636,MAPK3,mRNA,0,,,ERKs are inactivated;ISG15 antiviral mechanism;RHO GTPases Activate WASPs and WAVEs;Senescence-Associated Secretory Phenotype (SASP);Regulation of HSF1-mediated heat shock response;Thrombin signalling through proteinase activated receptors (PARs);Negative regulation of FGFR4 signaling;Oxidative Stress Induced Senescence;Signal attenuation;RNA Polymerase I Promoter Opening;Negative regulation of FGFR3 signaling;Negative feedback regulation of MAPK pathway;Growth hormone receptor signaling;Golgi Cisternae Pericentriolar Stack Reorganization;FCERI mediated MAPK activation;Negative regulation of MAPK pathway;RAFMAP kinase cascade;Downstream signal transduction;VEGFR2 mediated cell proliferation;Signaling by SCF-KIT;Frs2-mediated activation;ARMS-mediated activation;Gastrin-CREB signalling pathway via PKC and MAPK;Interleukin receptor SHC signaling;DAP12 signaling;Signaling by PDGF;VEGFA-VEGFR2 Pathway;Signaling by ERBB2;IRS-related events triggered by IGF1R,hsa04010;hsa04012;hsa04014;hsa04015;hsa04022;hsa04024;hsa04062;hsa04066;hsa04068;hsa04071;hsa04072;hsa04114;hsa04150;hsa04151;hsa04261;hsa04270;hsa04320;hsa04350;hsa04360;hsa04370;hsa04380;hsa04510;hsa04520;hsa04540;hsa04550;hsa04611;hsa04620;hsa04621;hsa04650;hsa04660;hsa04662;hsa04664;hsa04666;hsa04668;hsa04713;hsa04720;hsa04722;hsa04723;hsa04724;hsa04725;hsa04726;hsa04730;hsa04810;hsa04910;hsa04912;hsa04914;hsa04915;hsa04916;hsa04917;hsa04919;hsa04921;hsa04930;hsa04933;hsa04960;hsa05010;hsa05020;hsa05034;hsa05131;hsa05132;hsa05133;hsa05140;hsa05142;hsa05145;hsa05152;hsa05160;hsa05161;hsa05164;hsa05200;hsa05203;hsa05205;hsa05210;hsa05211;hsa05212;hsa05213;hsa05214;hsa05215;hsa05216;hsa05218;hsa05219;hsa05220;hsa05221;hsa05223;hsa05230;hsa05231,MAPK3,NA,MAPK3 - mRNA,False\nPPP2R1B - mRNA,NM_002716.4:1445,SEBELIUS_2_C4636,PPP2R1B,mRNA,0,,,MASTL Facilitates Mitotic Progression;Misspliced GSK3beta mutants stabilize beta-catenin;T41 mutants of beta-catenin aren.SQT.t phosphorylated;ERKs are inactivated;Glycolysis;Negative regulation of MAPK pathway;Disassembly of the destruction complex and recruitment of AXIN to the membrane;S37 mutants of beta-catenin aren.SQT.t phosphorylated;Cyclin AB1 associated events during G2M transition;RAF activation;RHO GTPases Activate Formins;S33 mutants of beta-catenin aren.SQT.t phosphorylated;DARPP-32 events;Beta-catenin phosphorylation cascade;Truncations of AMER1 destabilize the destruction complex;CTLA4 inhibitory signaling;AXIN missense mutants destabilize the destruction complex;S45 mutants of beta-catenin aren.SQT.t phosphorylated;Cyclin D associated events in G1;Platelet sensitization by LDL;Resolution of Sister Chromatid Cohesion;Inhibition of replication initiation of damaged DNA by RB1E2F1;RAFMAP kinase cascade;Downstream signal transduction;VEGFR2 mediated cell proliferation;Signaling by SCF-KIT;Frs2-mediated activation;ARMS-mediated activation;Gastrin-CREB signalling pathway via PKC and MAPK;Interleukin receptor SHC signaling;FCERI mediated MAPK activation;DAP12 signaling;Signaling by PDGF;VEGFA-VEGFR2 Pathway;Signaling by ERBB2;IRS-related events triggered by IGF1R,hsa03015;hsa04071;hsa04114;hsa04151;hsa04152;hsa04261;hsa04350;hsa04390;hsa04530;hsa04728;hsa04730;hsa05142;hsa05160,PPP2R1B,NA,PPP2R1B - mRNA,False\nMAPK9 - mRNA,XM_005265940.1:163,SEBELIUS_2_C4636,MAPK9,mRNA,0,,,,hsa04010;hsa04012;hsa04014;hsa04024;hsa04068;hsa04071;hsa04141;hsa04310;hsa04380;hsa04510;hsa04620;hsa04621;hsa04660;hsa04664;hsa04668;hsa04722;hsa04723;hsa04728;hsa04750;hsa04910;hsa04912;hsa04914;hsa04917;hsa04920;hsa04930;hsa04931;hsa04932;hsa04933;hsa05120;hsa05131;hsa05132;hsa05133;hsa05142;hsa05145;hsa05152;hsa05160;hsa05161;hsa05164;hsa05168;hsa05169;hsa05200;hsa05210;hsa05212;hsa05231,MAPK9,NA,MAPK9 - mRNA,False\nPRKCI - mRNA,NM_002740.4:530,SEBELIUS_2_C4636,PRKCI,mRNA,0,,,Tight junction interactions,hsa04015;hsa04144;hsa04390;hsa04530;hsa04611;hsa04910,PRKCI,NA,PRKCI - mRNA,False\nEGFR - mRNA,NM_201282.1:360,SEBELIUS_2_C4636,EGFR,mRNA,0,,,Constitutive Signaling by Aberrant PI3K in Cancer;Inhibition of Signaling by Overexpressed EGFR;PLCG1 events in ERBB2 signaling;EGFR interacts with phospholipase C-gamma;EGFR Transactivation by Gastrin;Constitutive Signaling by Ligand-Responsive EGFR Cancer Variants;Constitutive Signaling by EGFRvIII;GAB1 signalosome;Downstream signal transduction;Signaling by SCF-KIT;PI3KAKT activation;Role of LAT2NTALLAB on calcium mobilization;DAP12 signaling;Signaling by PDGF;Signaling by ERBB2;VEGFR2 mediated cell proliferation;Frs2-mediated activation;ARMS-mediated activation;Gastrin-CREB signalling pathway via PKC and MAPK;Interleukin receptor SHC signaling;FCERI mediated MAPK activation;VEGFA-VEGFR2 Pathway;IRS-related events triggered by IGF1R,hsa04010;hsa04012;hsa04014;hsa04015;hsa04020;hsa04060;hsa04066;hsa04068;hsa04072;hsa04144;hsa04151;hsa04320;hsa04510;hsa04520;hsa04540;hsa04810;hsa04912;hsa04915;hsa04921;hsa05120;hsa05160;hsa05200;hsa05205;hsa05206;hsa05212;hsa05213;hsa05214;hsa05215;hsa05218;hsa05219;hsa05223;hsa05230;hsa05231,EGFR,NA,EGFR - mRNA,False\nAKT3 - mRNA,NM_005465.4:287,SEBELIUS_2_C4636,AKT3,mRNA,0,,,TP53 Regulates Metabolic Genes;CD28 dependent PI3KAkt signaling;AKT phosphorylates targets in the cytosol;Negative regulation of the PI3KAKT network;Activation of BAD and translocation to mitochondria ;G beta.CL.gamma signalling through PI3Kgamma;AKT phosphorylates targets in the nucleus;VEGFR2 mediated vascular permeability;GPVI-mediated activation cascade;Signaling by ERBB2;PIP3 activates AKT signaling;Downstream signal transduction;GAB1 signalosome;Signaling by SCF-KIT;PI3KAKT activation;Role of LAT2NTALLAB on calcium mobilization;DAP12 signaling;Signaling by PDGF;CD28 co-stimulation;VEGFA-VEGFR2 Pathway,hsa04010;hsa04012;hsa04014;hsa04015;hsa04022;hsa04024;hsa04062;hsa04066;hsa04068;hsa04071;hsa04072;hsa04150;hsa04151;hsa04152;hsa04210;hsa04211;hsa04213;hsa04261;hsa04370;hsa04380;hsa04510;hsa04530;hsa04550;hsa04611;hsa04620;hsa04630;hsa04660;hsa04662;hsa04664;hsa04666;hsa04668;hsa04722;hsa04725;hsa04728;hsa04910;hsa04914;hsa04915;hsa04917;hsa04919;hsa04920;hsa04922;hsa04923;hsa04931;hsa04932;hsa04933;hsa04973;hsa05142;hsa05145;hsa05152;hsa05160;hsa05161;hsa05162;hsa05164;hsa05166;hsa05169;hsa05200;hsa05205;hsa05210;hsa05211;hsa05212;hsa05213;hsa05214;hsa05215;hsa05218;hsa05220;hsa05221;hsa05222;hsa05223;hsa05230;hsa05231,AKT3,NA,AKT3 - mRNA,False\nPAK2 - mRNA,NM_002577.3:3785,SEBELIUS_2_C4636,PAK2,mRNA,0,,,Nef and signal transduction;CD28 dependent Vav1 pathway;Stimulation of the cell death response by PAK-2p34;Ephrin signaling;Activation of Rac;RHO GTPases activate PAKs;VEGFR2 mediated vascular permeability;Sema3A PAK dependent Axon repulsion;CD209 (DC-SIGN) signaling;VEGFA-VEGFR2 Pathway;CD28 co-stimulation,hsa04010;hsa04012;hsa04014;hsa04360;hsa04510;hsa04660;hsa04810;hsa05211,PAK2,NA,PAK2 - mRNA,False\nDOCK1 - mRNA,NM_001380.3:840,SEBELIUS_2_C4636,DOCK1,mRNA,0,,,DCC mediated attractive signaling;Factors involved in megakaryocyte development and platelet production;VEGFA-VEGFR2 Pathway;Regulation of actin dynamics for phagocytic cup formation,hsa04510;hsa04810;hsa05100;hsa05131,DOCK1,NA,DOCK1 - mRNA,False\nVAPA - mRNA,NM_194434.2:763,SEBELIUS_2_C4636,VAPA,mRNA,0,,,,hsa04530,VAPA,NA,VAPA - mRNA,False\nJAM3 - mRNA,NM_032801.3:890,SEBELIUS_2_C4636,JAM3,mRNA,0,,,Integrin cell surface interactions;Cell surface interactions at the vascular wall,hsa04514;hsa04530;hsa04670;hsa05120,JAM3,NA,JAM3 - mRNA,False\nZEB1 - mRNA,NM_001128128.1:1450,SEBELIUS_2_C4636,ZEB1,mRNA,0,,,,hsa05202;hsa05206,ZEB1,NA,ZEB1 - mRNA,False\nITGAV - mRNA,NM_002210.2:2615,SEBELIUS_2_C4636,ITGAV,mRNA,0,,,Molecules associated with elastic fibres;Syndecan interactions;Signal transduction by L1;Laminin interactions;ECM proteoglycans;Integrin cell surface interactions;VEGFA-VEGFR2 Pathway;Elastic fibre formation;Non-integrin membrane-ECM interactions;Cell surface interactions at the vascular wall,hsa04145;hsa04151;hsa04510;hsa04512;hsa04514;hsa04810;hsa04919;hsa05200;hsa05205;hsa05222;hsa05410;hsa05412;hsa05414,ITGAV,NA,ITGAV - mRNA,False\nRRAS2 - mRNA,NM_001102669.2:1785,SEBELIUS_2_C4636,RRAS2,mRNA,0,,,,hsa04010;hsa04014;hsa04024;hsa04072;hsa04530;hsa04810;hsa05166;hsa05205,RRAS2,NA,RRAS2 - mRNA,False\nNRAS - mRNA,NM_002524.3:877,SEBELIUS_2_C4636,NRAS,mRNA,0,,,Signaling by FGFR2 in disease;Regulation of RAS by GAPs;SHC-mediated cascade.CL.FGFR3;Signaling by FGFR3 in disease;MAP2K and MAPK activation;SHC-related events triggered by IGF1R;Signaling by FGFR4 in disease;SHC-mediated cascade.CL.FGFR1;Tie2 Signaling;Signaling by FGFR1 in disease;Activation of RAS in B cells;SHC-mediated cascade.CL.FGFR4;Constitutive Signaling by EGFRvIII;CD209 (DC-SIGN) signaling;FCERI mediated MAPK activation;RAFMAP kinase cascade;Downstream signal transduction;VEGFR2 mediated cell proliferation;Signaling by SCF-KIT;Frs2-mediated activation;ARMS-mediated activation;Gastrin-CREB signalling pathway via PKC and MAPK;Interleukin receptor SHC signaling;DAP12 signaling;Signaling by PDGF;VEGFA-VEGFR2 Pathway;Signaling by ERBB2;IRS-related events triggered by IGF1R;Cell surface interactions at the vascular wall,hsa04010;hsa04012;hsa04014;hsa04015;hsa04062;hsa04068;hsa04071;hsa04072;hsa04151;hsa04211;hsa04213;hsa04360;hsa04370;hsa04530;hsa04540;hsa04550;hsa04650;hsa04660;hsa04662;hsa04664;hsa04720;hsa04722;hsa04725;hsa04726;hsa04730;hsa04810;hsa04910;hsa04912;hsa04915;hsa04916;hsa04917;hsa04919;hsa04921;hsa04933;hsa05034;hsa05160;hsa05161;hsa05166;hsa05200;hsa05203;hsa05205;hsa05206;hsa05211;hsa05213;hsa05214;hsa05215;hsa05216;hsa05218;hsa05219;hsa05220;hsa05221;hsa05223;hsa05230;hsa05231,NRAS,NA,NRAS - mRNA,False\nMAP2K1 - mRNA,NM_002755.2:970,SEBELIUS_2_C4636,MAP2K1,mRNA,0,,,RAF activation;Uptake and function of anthrax toxins;Negative feedback regulation of MAPK pathway;Interleukin-1 signaling;RAFMAP kinase cascade;Downstream signal transduction;VEGFR2 mediated cell proliferation;Signaling by SCF-KIT;Frs2-mediated activation;ARMS-mediated activation;Gastrin-CREB signalling pathway via PKC and MAPK;Interleukin receptor SHC signaling;FCERI mediated MAPK activation;DAP12 signaling;Signaling by PDGF;VEGFA-VEGFR2 Pathway;Signaling by ERBB2;IRS-related events triggered by IGF1R;Negative regulation of MAPK pathway,hsa04010;hsa04012;hsa04014;hsa04015;hsa04022;hsa04024;hsa04062;hsa04066;hsa04068;hsa04071;hsa04072;hsa04114;hsa04151;hsa04270;hsa04320;hsa04370;hsa04380;hsa04510;hsa04540;hsa04550;hsa04620;hsa04650;hsa04660;hsa04662;hsa04664;hsa04666;hsa04668;hsa04720;hsa04722;hsa04725;hsa04726;hsa04730;hsa04810;hsa04910;hsa04912;hsa04914;hsa04915;hsa04916;hsa04917;hsa04919;hsa04921;hsa05020;hsa05034;hsa05161;hsa05164;hsa05200;hsa05205;hsa05206;hsa05210;hsa05211;hsa05212;hsa05213;hsa05214;hsa05215;hsa05216;hsa05218;hsa05219;hsa05220;hsa05221;hsa05223;hsa05230;hsa05231,MAP2K1,NA,MAP2K1 - mRNA,False\nROCK1 - mRNA,NM_005406.1:2660,SEBELIUS_2_C4636,ROCK1,mRNA,0,,,EPHB-mediated forward signaling;EPHA-mediated growth cone collapse;RHO GTPases Activate ROCKs;Apoptotic cleavage of cellular proteins;G alpha (1213) signalling events;VEGFA-VEGFR2 Pathway;Sema4D induced cell migration and growth-cone collapse,hsa04022;hsa04024;hsa04062;hsa04071;hsa04270;hsa04350;hsa04360;hsa04510;hsa04611;hsa04670;hsa04810;hsa04921;hsa05130;hsa05131;hsa05132;hsa05200;hsa05205;hsa05206,ROCK1,NA,ROCK1 - mRNA,False\nPPP1R12A - mRNA,NM_002480.2:2935,SEBELIUS_2_C4636,PPP1R12A,mRNA,0,,,RHO GTPases Activate ROCKs;RHO GTPases activate PAKs;RHO GTPases activate PKNs,hsa04022;hsa04024;hsa04270;hsa04510;hsa04611;hsa04810;hsa04921;hsa05205,PPP1R12A,NA,PPP1R12A - mRNA,False\nPDGFD - mRNA,NM_025208.4:1120,SEBELIUS_2_C4636,PDGFD,mRNA,0,,,Signaling by PDGF,hsa04014;hsa04015;hsa04072;hsa04151;hsa04510;hsa04540;hsa04810;hsa05215;hsa05218;hsa05231,PDGFD,NA,PDGFD - mRNA,False\nIL1R1 - mRNA,NM_000877.3:582,SEBELIUS_2_C4636,IL1R1,mRNA,0,,,,hsa04010;hsa04060;hsa04064;hsa04210;hsa04380;hsa04640;hsa04750;hsa05146;hsa05166,IL1R1,NA,IL1R1 - mRNA,False\nYBX3 - mRNA,NM_003651.3:660,SEBELIUS_2_C4636,YBX3,mRNA,0,,,,hsa04530,YBX3,NA,YBX3 - mRNA,False\nMYH10 - mRNA,NM_005964.1:7092,SEBELIUS_2_C4636,MYH10,mRNA,0,,,EPHA-mediated growth cone collapse;RHO GTPases Activate ROCKs;RHO GTPases activate CIT;RHO GTPases activate PAKs;RHO GTPases activate PKNs;Sema4D induced cell migration and growth-cone collapse,hsa04530;hsa04810;hsa05132,MYH10,NA,MYH10 - mRNA,False\nSMAD4 - mRNA,NM_005359.3:1370,SEBELIUS_2_C4636,SMAD4,mRNA,0,,,Signaling by NODAL,hsa04068;hsa04110;hsa04310;hsa04350;hsa04390;hsa04520;hsa04550;hsa04933;hsa05161;hsa05166;hsa05200;hsa05210;hsa05212;hsa05220,SMAD4,NA,SMAD4 - mRNA,False\nCRKL - mRNA,NM_005207.3:1372,SEBELIUS_2_C4636,CRKL,mRNA,0,,,Downstream signal transduction;Regulation of signaling by CBL;Frs2-mediated activation;Signaling by PDGF,hsa04010;hsa04012;hsa04015;hsa04062;hsa04510;hsa04666;hsa04722;hsa04810;hsa04910;hsa05100;hsa05131;hsa05200;hsa05206;hsa05211;hsa05220,CRKL,NA,CRKL - mRNA,False\nPOLR2A - mRNA,NM_000937.2:3775,SEBELIUS_2_C4636,POLR2A,mRNA,0,,,RNA Polymerase II Transcription Initiation,hsa01100;hsa05016;hsa05168;hsa05169,POLR2A,NA,POLR2A - mRNA,False\nPIK3CA - mRNA,NM_006218.2:2445,SEBELIUS_2_C4636,PIK3CA,mRNA,0,,,G alpha (q) signalling events;Downstream TCR signaling;Signaling by FGFR2 in disease;PIP3 activates AKT signaling;VEGFA-VEGFR2 Pathway;Synthesis of PIPs at the plasma membrane;Constitutive Signaling by Ligand-Responsive EGFR Cancer Variants;Interleukin receptor SHC signaling;G beta.CL.gamma signalling through PI3Kgamma;Regulation of signaling by CBL;Role of phospholipids in phagocytosis;G alpha (1213) signalling events;PI3K Cascade;Signaling by FGFR3 in disease;Constitutive Signaling by Aberrant PI3K in Cancer;Signaling by FGFR4 in disease;CD28 dependent PI3KAkt signaling;Tie2 Signaling;GPVI-mediated activation cascade;Nephrin interactions;Constitutive Signaling by EGFRvIII;Gastrin-CREB signalling pathway via PKC and MAPK;Downstream signal transduction;GAB1 signalosome;Signaling by SCF-KIT;PI3KAKT activation;Role of LAT2NTALLAB on calcium mobilization;DAP12 signaling;Signaling by PDGF;Signaling by ERBB2;IRS-related events triggered by IGF1R;CD28 co-stimulation;Cell surface interactions at the vascular wall;Signaling by FGFR1 in disease,hsa00562;hsa04012;hsa04014;hsa04015;hsa04022;hsa04024;hsa04062;hsa04066;hsa04068;hsa04070;hsa04071;hsa04072;hsa04150;hsa04151;hsa04152;hsa04210;hsa04211;hsa04213;hsa04261;hsa04370;hsa04380;hsa04510;hsa04550;hsa04611;hsa04620;hsa04630;hsa04650;hsa04660;hsa04662;hsa04664;hsa04666;hsa04668;hsa04670;hsa04722;hsa04725;hsa04750;hsa04810;hsa04910;hsa04914;hsa04915;hsa04917;hsa04919;hsa04921;hsa04923;hsa04930;hsa04931;hsa04932;hsa04933;hsa04960;hsa04973;hsa05100;hsa05142;hsa05145;hsa05146;hsa05160;hsa05161;hsa05162;hsa05164;hsa05166;hsa05169;hsa05200;hsa05203;hsa05205;hsa05206;hsa05210;hsa05211;hsa05212;hsa05213;hsa05214;hsa05215;hsa05218;hsa05220;hsa05221;hsa05222;hsa05223;hsa05230;hsa05231,PIK3CA,NA,PIK3CA - mRNA,False\nTP53 - mRNA,NM_000546.2:1330,SEBELIUS_2_C4636,TP53,mRNA,0,,,Activation of PUMA and translocation to mitochondria;Recruitment and ATM-mediated phosphorylation of repair and signaling proteins at DNA double strand breaks;Factors involved in megakaryocyte development and platelet production;Oncogene Induced Senescence,hsa04010;hsa04071;hsa04110;hsa04115;hsa04151;hsa04210;hsa04211;hsa04310;hsa04722;hsa04919;hsa05016;hsa05160;hsa05161;hsa05162;hsa05166;hsa05168;hsa05169;hsa05200;hsa05202;hsa05203;hsa05205;hsa05206;hsa05210;hsa05212;hsa05213;hsa05214;hsa05215;hsa05216;hsa05217;hsa05218;hsa05219;hsa05220;hsa05222;hsa05223;hsa05230,TP53,NA,TP53 - mRNA,False\nCTTN - mRNA,NM_001184740.1:240,SEBELIUS_2_C4636,CTTN,mRNA,0,,,RHO GTPases activate PAKs,hsa04530;hsa05100;hsa05130;hsa05131;hsa05205,CTTN,NA,CTTN - mRNA,False\nPAK1 - mRNA,NM_002576.3:975,SEBELIUS_2_C4636,PAK1,mRNA,0,,,EPHB-mediated forward signaling;Ephrin signaling;MAPK6MAPK4 signaling;DSCAM interactions;Activation of Rac;VEGFR2 mediated vascular permeability;RHO GTPases activate PAKs;Sema3A PAK dependent Axon repulsion;FCERI mediated MAPK activation;Regulation of actin dynamics for phagocytic cup formation;CD209 (DC-SIGN) signaling;CD28 dependent Vav1 pathway;VEGFA-VEGFR2 Pathway;CD28 co-stimulation,hsa04010;hsa04012;hsa04014;hsa04024;hsa04062;hsa04360;hsa04510;hsa04650;hsa04660;hsa04666;hsa04810;hsa05120;hsa05205;hsa05211,PAK1,NA,PAK1 - mRNA,False\nHPRT1 - mRNA,NM_000194.1:240,SEBELIUS_2_C4636,HPRT1,mRNA,0,,,,hsa01100,HPRT1,NA,HPRT1 - mRNA,False\nSPRY2 - mRNA,NM_005842.2:1481,SEBELIUS_2_C4636,SPRY2,mRNA,0,,,Spry regulation of FGF signaling;Negative regulation of FGFR3 signaling;Negative regulation of FGFR4 signaling;Negative regulation of FGFR2 signaling;Negative regulation of FGFR1 signaling,hsa05206,SPRY2,NA,SPRY2 - mRNA,False\nPXN - mRNA,NM_001080855.1:2495,SEBELIUS_2_C4636,PXN,mRNA,0,,,Smooth Muscle Contraction;GAB1 signalosome;VEGFA-VEGFR2 Pathway;Cell-extracellular matrix interactions,hsa04062;hsa04370;hsa04510;hsa04670;hsa04810;hsa05100;hsa05203;hsa05205,PXN,NA,PXN - mRNA,False\nITGA3 - mRNA,NM_002204.2:4550,SEBELIUS_2_C4636,ITGA3,mRNA,0,,,Laminin interactions;Integrin cell surface interactions;Cell surface interactions at the vascular wall,hsa04151;hsa04510;hsa04512;hsa04640;hsa04810;hsa05200;hsa05222;hsa05410;hsa05412;hsa05414,ITGA3,NA,ITGA3 - mRNA,False\nCASK - mRNA,NM_001126054.2:2190,SEBELIUS_2_C4636,CASK,mRNA,0,,,Syndecan interactions;Nephrin interactions;Non-integrin membrane-ECM interactions,hsa04530,CASK,NA,CASK - mRNA,False\nYES1 - mRNA,NM_005433.3:265,SEBELIUS_2_C4636,YES1,mRNA,0,,,Signaling by ERBB2;EPHB-mediated forward signaling;EPHA-mediated growth cone collapse;Regulation of KIT signaling;EPH-ephrin mediated repulsion of cells;Regulation of signaling by CBL;CD28 co-stimulation;CTLA4 inhibitory signaling;Signaling by SCF-KIT;Cell surface interactions at the vascular wall,hsa04520;hsa04530,YES1,NA,YES1 - mRNA,False\nCRK - mRNA,NM_016823.2:1945,SEBELIUS_2_C4636,CRK,mRNA,0,,,ARMS-mediated activation;Downstream signal transduction;Regulation of signaling by CBL;p130Cas linkage to MAPK signaling for integrins;VEGFA-VEGFR2 Pathway;Regulation of actin dynamics for phagocytic cup formation;Signaling by PDGF;Integrin alphaIIb beta3 signaling,hsa04010;hsa04012;hsa04015;hsa04062;hsa04510;hsa04666;hsa04722;hsa04810;hsa04910;hsa05100;hsa05131;hsa05200;hsa05206;hsa05211;hsa05220,CRK,NA,CRK - mRNA,False\nPPP2R2A - mRNA,NM_001177591.1:430,SEBELIUS_2_C4636,PPP2R2A,mRNA,0,,,Nonsense Mediated Decay (NMD) enhanced by the Exon Junction Complex (EJC);Cyclin D associated events in G1;Cyclin AB1 associated events during G2M transition,hsa03015;hsa04071;hsa04151;hsa04152;hsa04261;hsa04390;hsa04530;hsa04728;hsa05142;hsa05160,PPP2R2A,NA,PPP2R2A - mRNA,False\nGNAI3 - mRNA,NM_006496.1:1150,SEBELIUS_2_C4636,GNAI3,mRNA,0,,,G alpha (i) signalling events;ADP signalling through P2Y purinoceptor 12,hsa04015;hsa04022;hsa04024;hsa04062;hsa04071;hsa04261;hsa04360;hsa04530;hsa04540;hsa04611;hsa04670;hsa04713;hsa04723;hsa04724;hsa04725;hsa04726;hsa04727;hsa04728;hsa04730;hsa04914;hsa04915;hsa04916;hsa04921;hsa04923;hsa04971;hsa05032;hsa05034;hsa05133;hsa05142;hsa05145;hsa05200,GNAI3,NA,GNAI3 - mRNA,False\nMPP5 - mRNA,NM_022474.2:1110,SEBELIUS_2_C4636,MPP5,mRNA,0,,,Tight junction interactions,hsa04390;hsa04530,MPP5,NA,MPP5 - mRNA,False\nBRK1 - mRNA,NM_018462.4:710,SEBELIUS_2_C4636,BRK1,mRNA,0,,,RHO GTPases Activate WASPs and WAVEs;VEGFA-VEGFR2 Pathway;Regulation of actin dynamics for phagocytic cup formation,hsa04810,BRK1,NA,BRK1 - mRNA,False\nAKT1 - mRNA,NM_001014432.1:1275,SEBELIUS_2_C4636,AKT1,mRNA,0,,,TP53 Regulates Metabolic Genes;Deactivation of the beta-catenin transactivating complex;Activation of BAD and translocation to mitochondria ;eNOS activation;mTOR signalling;G beta.CL.gamma signalling through PI3Kgamma;KSRP (KHSRP) binds and destabilizes mRNA;AKT phosphorylates targets in the nucleus;VEGFR2 mediated vascular permeability;GPVI-mediated activation cascade;Tetrahydrobiopterin (BH4) synthesis# recycling# salvage and regulation;Butyrate Response Factor 1 (BRF1) binds and destabilizes mRNA;Signaling by ERBB2;PIP3 activates AKT signaling;Downstream signal transduction;GAB1 signalosome;Signaling by SCF-KIT;PI3KAKT activation;Role of LAT2NTALLAB on calcium mobilization;DAP12 signaling;Signaling by PDGF;CD28 co-stimulation;PI3K Cascade;IRS-related events triggered by IGF1R;VEGFA-VEGFR2 Pathway,hsa04010;hsa04012;hsa04014;hsa04015;hsa04022;hsa04024;hsa04062;hsa04066;hsa04068;hsa04071;hsa04072;hsa04150;hsa04151;hsa04152;hsa04210;hsa04211;hsa04213;hsa04261;hsa04370;hsa04380;hsa04510;hsa04530;hsa04550;hsa04611;hsa04620;hsa04630;hsa04660;hsa04662;hsa04664;hsa04666;hsa04668;hsa04722;hsa04725;hsa04728;hsa04910;hsa04914;hsa04915;hsa04917;hsa04919;hsa04920;hsa04922;hsa04923;hsa04931;hsa04932;hsa04933;hsa04973;hsa05142;hsa05145;hsa05152;hsa05160;hsa05161;hsa05162;hsa05164;hsa05166;hsa05169;hsa05200;hsa05205;hsa05210;hsa05211;hsa05212;hsa05213;hsa05214;hsa05215;hsa05218;hsa05220;hsa05221;hsa05222;hsa05223;hsa05230;hsa05231,AKT1,NA,AKT1 - mRNA,False\nDIAPH1 - mRNA,NM_005219.4:1398,SEBELIUS_2_C4636,DIAPH1,mRNA,0,,,RHO GTPases Activate Formins,hsa04510;hsa04810;hsa04933;hsa05131,DIAPH1,NA,DIAPH1 - mRNA,False\nABCF1 - mRNA,NM_001090.2:850,SEBELIUS_2_C4636,ABCF1,mRNA,0,,,,,ABCF1,NA,ABCF1 - mRNA,False\nIL1R1 b - mRNA,NM_001320984.1:2649,SEBELIUS_2_C4636,IL1R1 b,mRNA,0,,,,,,NA,IL1R1 b - mRNA,False\nSOS2 - mRNA,NM_006939.2:3845,SEBELIUS_2_C4636,SOS2,mRNA,0,,,Rho GTPase cycle;Activation of Rac;NRAGE signals death through JNK;G alpha (1213) signalling events,hsa04010;hsa04012;hsa04014;hsa04062;hsa04068;hsa04072;hsa04151;hsa04320;hsa04510;hsa04540;hsa04630;hsa04650;hsa04660;hsa04662;hsa04664;hsa04722;hsa04810;hsa04910;hsa04912;hsa04915;hsa04917;hsa05034;hsa05160;hsa05200;hsa05205;hsa05206;hsa05211;hsa05213;hsa05214;hsa05215;hsa05220;hsa05221;hsa05223;hsa05231,SOS2,NA,SOS2 - mRNA,False\nIRS1 - mRNA,NM_005544.2:6224,SEBELIUS_2_C4636,IRS1,mRNA,0,,,Constitutive Signaling by Aberrant PI3K in Cancer;RAFMAP kinase cascade;PI3K Cascade;Growth hormone receptor signaling;IRS-related events triggered by IGF1R;PI3KAKT activation;Downstream signal transduction;GAB1 signalosome;Signaling by SCF-KIT;Role of LAT2NTALLAB on calcium mobilization;DAP12 signaling;Signaling by PDGF;Signaling by ERBB2;VEGFR2 mediated cell proliferation;Frs2-mediated activation;ARMS-mediated activation;Gastrin-CREB signalling pathway via PKC and MAPK;Interleukin receptor SHC signaling;FCERI mediated MAPK activation;VEGFA-VEGFR2 Pathway,hsa04022;hsa04068;hsa04150;hsa04151;hsa04152;hsa04211;hsa04213;hsa04722;hsa04910;hsa04920;hsa04923;hsa04930;hsa04931;hsa04932;hsa04960;hsa05206,IRS1,NA,IRS1 - mRNA,False\nDROSHA - mRNA,NM_013235.4:1860,SEBELIUS_2_C4636,DROSHA,mRNA,0,,,MicroRNA (miRNA) biogenesis,hsa05205,DROSHA,NA,DROSHA - mRNA,False\nCCNG2 - mRNA,NM_004354.2:336,SEBELIUS_2_C4636,CCNG2,mRNA,0,,,,hsa04068;hsa04115,CCNG2,NA,CCNG2 - mRNA,False\nDVL3 - mRNA,NM_004423.3:1012,SEBELIUS_2_C4636,DVL3,mRNA,0,,,Disassembly of the destruction complex and recruitment of AXIN to the membrane;WNT mediated activation of DVL;RHO GTPases Activate Formins;PCPCE pathway,hsa04310;hsa04330;hsa04390;hsa04550;hsa04916;hsa05166;hsa05200;hsa05217,DVL3,NA,DVL3 - mRNA,False\nMAPK8 - mRNA,NM_002750.2:945,SEBELIUS_2_C4636,MAPK8,mRNA,0,,,Activation of the AP-1 family of transcription factors;FCERI mediated MAPK activation;Activation of BMF and translocation to mitochondria;JNK (c-Jun kinases) phosphorylation and  activation mediated by activated human TAK1,hsa04010;hsa04012;hsa04014;hsa04024;hsa04068;hsa04071;hsa04141;hsa04310;hsa04380;hsa04510;hsa04620;hsa04621;hsa04664;hsa04668;hsa04722;hsa04723;hsa04728;hsa04750;hsa04910;hsa04912;hsa04914;hsa04917;hsa04920;hsa04930;hsa04931;hsa04932;hsa04933;hsa05120;hsa05131;hsa05132;hsa05133;hsa05142;hsa05145;hsa05152;hsa05160;hsa05161;hsa05164;hsa05166;hsa05168;hsa05169;hsa05200;hsa05210;hsa05212;hsa05231,MAPK8,NA,MAPK8 - mRNA,False\nSOS1 - mRNA,NM_005633.2:1635,SEBELIUS_2_C4636,SOS1,mRNA,0,,,Signaling by FGFR2 in disease;Antigen activates B Cell Receptor (BCR) leading to generation of second messengers;SHC-mediated cascade.CL.FGFR3;Signaling by FGFR3 in disease;RAFMAP kinase cascade;SHC-related events triggered by IGF1R;Signaling by FGFR4 in disease;SHC-mediated cascade.CL.FGFR1;GRB2.CL.SOS provides linkage to MAPK signaling for Integrins ;Tie2 Signaling;Role of LAT2NTALLAB on calcium mobilization;Signaling by FGFR1 in disease;Signal attenuation;NRAGE signals death through JNK;SHC-mediated cascade.CL.FGFR4;FCERI mediated Ca.PLS.2 mobilization;Constitutive Signaling by EGFRvIII;FCERI mediated MAPK activation;Signaling by ERBB2;Signaling by SCF-KIT;DAP12 signaling;Gastrin-CREB signalling pathway via PKC and MAPK;Interleukin receptor SHC signaling;IRS-related events triggered by IGF1R;Downstream signal transduction;VEGFR2 mediated cell proliferation;Frs2-mediated activation;ARMS-mediated activation;Signaling by PDGF;VEGFA-VEGFR2 Pathway;Integrin alphaIIb beta3 signaling;Cell surface interactions at the vascular wall,hsa04010;hsa04012;hsa04014;hsa04062;hsa04068;hsa04072;hsa04151;hsa04320;hsa04510;hsa04540;hsa04630;hsa04650;hsa04660;hsa04662;hsa04664;hsa04722;hsa04810;hsa04910;hsa04912;hsa04915;hsa04917;hsa05034;hsa05160;hsa05200;hsa05205;hsa05206;hsa05211;hsa05213;hsa05214;hsa05215;hsa05220;hsa05221;hsa05223;hsa05231,SOS1,NA,SOS1 - mRNA,False\nGSK3B - mRNA,NM_002093.2:925,SEBELIUS_2_C4636,GSK3B,mRNA,0,,,Misspliced GSK3beta mutants stabilize beta-catenin;Constitutive Signaling by AKT1 E17K in Cancer;APC truncation mutants have impaired AXIN binding;T41 mutants of beta-catenin aren.SQT.t phosphorylated;AXIN missense mutants destabilize the destruction complex;S45 mutants of beta-catenin aren.SQT.t phosphorylated;Regulation of HSF1-mediated heat shock response;AKT phosphorylates targets in the cytosol;Disassembly of the destruction complex and recruitment of AXIN to the membrane;S37 mutants of beta-catenin aren.SQT.t phosphorylated;S33 mutants of beta-catenin aren.SQT.t phosphorylated;Truncations of AMER1 destabilize the destruction complex;Degradation of GLI2 by the proteasome;PIP3 activates AKT signaling;Downstream signal transduction;GAB1 signalosome;Signaling by SCF-KIT;PI3KAKT activation;Role of LAT2NTALLAB on calcium mobilization;DAP12 signaling;Signaling by PDGF;Signaling by ERBB2,hsa04012;hsa04062;hsa04110;hsa04151;hsa04310;hsa04360;hsa04390;hsa04510;hsa04550;hsa04660;hsa04662;hsa04722;hsa04728;hsa04910;hsa04916;hsa04917;hsa04919;hsa04931;hsa04932;hsa05010;hsa05160;hsa05162;hsa05164;hsa05166;hsa05169;hsa05200;hsa05210;hsa05213;hsa05215;hsa05217,GSK3B,NA,GSK3B - mRNA,False\nEP300 - mRNA,NM_001429.2:715,SEBELIUS_2_C4636,EP300,mRNA,0,,,LRR FLII-interacting protein 1 (LRRFIP1) activates type I IFN production;TRAF6 mediated IRF7 activation;Constitutive Signaling by NOTCH1 PEST Domain Mutants;Polo-like kinase mediated events;B-WICH complex positively regulates rRNA expression;CD209 (DC-SIGN) signaling;Circadian Clock,hsa04024;hsa04066;hsa04068;hsa04110;hsa04310;hsa04330;hsa04350;hsa04520;hsa04630;hsa04720;hsa04916;hsa04919;hsa04922;hsa05016;hsa05152;hsa05161;hsa05164;hsa05166;hsa05168;hsa05169;hsa05200;hsa05203;hsa05206;hsa05211;hsa05215,EP300,NA,EP300 - mRNA,False\nCCND3 - mRNA,NM_001760.2:1215,SEBELIUS_2_C4636,CCND3,mRNA,0,,,Cyclin D associated events in G1;MAPK6MAPK4 signaling;Transcriptional regulation of white adipocyte differentiation,hsa04110;hsa04115;hsa04151;hsa04310;hsa04390;hsa04510;hsa04630;hsa05162;hsa05166;hsa05203,CCND3,NA,CCND3 - mRNA,False\nKRAS - mRNA,NM_004985.3:327,SEBELIUS_2_C4636,KRAS,mRNA,0,,,Signaling by FGFR2 in disease;Regulation of RAS by GAPs;SHC-mediated cascade.CL.FGFR3;Signaling by FGFR3 in disease;MAP2K and MAPK activation;SHC-related events triggered by IGF1R;Signaling by FGFR4 in disease;SHC-mediated cascade.CL.FGFR1;Tie2 Signaling;Signaling by FGFR1 in disease;Activation of RAS in B cells;SHC-mediated cascade.CL.FGFR4;Constitutive Signaling by EGFRvIII;CD209 (DC-SIGN) signaling;FCERI mediated MAPK activation;RAFMAP kinase cascade;Downstream signal transduction;VEGFR2 mediated cell proliferation;Signaling by SCF-KIT;Frs2-mediated activation;ARMS-mediated activation;Gastrin-CREB signalling pathway via PKC and MAPK;Interleukin receptor SHC signaling;DAP12 signaling;Signaling by PDGF;VEGFA-VEGFR2 Pathway;Signaling by ERBB2;IRS-related events triggered by IGF1R;Cell surface interactions at the vascular wall,hsa04010;hsa04012;hsa04014;hsa04015;hsa04062;hsa04068;hsa04071;hsa04072;hsa04151;hsa04211;hsa04213;hsa04320;hsa04360;hsa04370;hsa04530;hsa04540;hsa04550;hsa04650;hsa04660;hsa04662;hsa04664;hsa04720;hsa04722;hsa04725;hsa04726;hsa04730;hsa04810;hsa04910;hsa04912;hsa04914;hsa04915;hsa04916;hsa04917;hsa04919;hsa04921;hsa04933;hsa04960;hsa05034;hsa05160;hsa05161;hsa05166;hsa05200;hsa05203;hsa05205;hsa05206;hsa05210;hsa05211;hsa05212;hsa05213;hsa05214;hsa05215;hsa05216;hsa05218;hsa05219;hsa05220;hsa05221;hsa05223;hsa05230;hsa05231,KRAS,NA,KRAS - mRNA,False\nCSNK1A1 - mRNA,NM_001892.4:2053,SEBELIUS_2_C4636,CSNK1A1,mRNA,0,,,Misspliced GSK3beta mutants stabilize beta-catenin;APC truncation mutants have impaired AXIN binding;T41 mutants of beta-catenin aren.SQT.t phosphorylated;AXIN missense mutants destabilize the destruction complex;S45 mutants of beta-catenin aren.SQT.t phosphorylated;Disassembly of the destruction complex and recruitment of AXIN to the membrane;S37 mutants of beta-catenin aren.SQT.t phosphorylated;S33 mutants of beta-catenin aren.SQT.t phosphorylated;Beta-catenin phosphorylation cascade;Degradation of GLI2 by the proteasome,hsa04310,CSNK1A1,NA,CSNK1A1 - mRNA,False\nRAB13 - mRNA,NM_002870.3:332,SEBELIUS_2_C4636,RAB13,mRNA,0,,,Translocation of GLUT4 to the plasma membrane,hsa04530,RAB13,NA,RAB13 - mRNA,False\nPDGFRA - mRNA,NM_006206.3:1925,SEBELIUS_2_C4636,PDGFRA,mRNA,0,,,PIP3 activates AKT signaling;Constitutive Signaling by Aberrant PI3K in Cancer;RAFMAP kinase cascade;Signaling by PDGF;Downstream signal transduction;GAB1 signalosome;Signaling by SCF-KIT;PI3KAKT activation;Role of LAT2NTALLAB on calcium mobilization;DAP12 signaling;Signaling by ERBB2;VEGFR2 mediated cell proliferation;Frs2-mediated activation;ARMS-mediated activation;Gastrin-CREB signalling pathway via PKC and MAPK;Interleukin receptor SHC signaling;FCERI mediated MAPK activation;VEGFA-VEGFR2 Pathway;IRS-related events triggered by IGF1R,hsa04010;hsa04014;hsa04015;hsa04020;hsa04060;hsa04072;hsa04144;hsa04151;hsa04510;hsa04540;hsa04810;hsa05166;hsa05200;hsa05206;hsa05214;hsa05215;hsa05218;hsa05230;hsa05231,PDGFRA,NA,PDGFRA - mRNA,False\nPPP1CA - mRNA,NM_002708.3:505,SEBELIUS_2_C4636,PPP1CA,mRNA,0,,,DARPP-32 events;Downregulation of TGF-beta receptor signaling;Circadian Clock;TGF-beta receptor signaling activates SMADs,hsa03015;hsa04022;hsa04024;hsa04114;hsa04261;hsa04270;hsa04390;hsa04510;hsa04611;hsa04720;hsa04728;hsa04750;hsa04810;hsa04910;hsa04921;hsa04931;hsa05031;hsa05034;hsa05168;hsa05205,PPP1CA,NA,PPP1CA - mRNA,False\nVCL - mRNA,NM_014000.2:4780,SEBELIUS_2_C4636,VCL,mRNA,0,,,Smooth Muscle Contraction;MAP2K and MAPK activation;Platelet degranulation ;RAFMAP kinase cascade;Downstream signal transduction;VEGFR2 mediated cell proliferation;Signaling by SCF-KIT;Frs2-mediated activation;ARMS-mediated activation;Gastrin-CREB signalling pathway via PKC and MAPK;Interleukin receptor SHC signaling;FCERI mediated MAPK activation;DAP12 signaling;Signaling by PDGF;VEGFA-VEGFR2 Pathway;Signaling by ERBB2;IRS-related events triggered by IGF1R,hsa04510;hsa04520;hsa04670;hsa04810;hsa05100;hsa05131;hsa05146,VCL,NA,VCL - mRNA,False\nACTN4 - mRNA,NM_004924.4:1768,SEBELIUS_2_C4636,ACTN4,mRNA,0,,,Nephrin interactions;Platelet degranulation ,hsa04510;hsa04520;hsa04530;hsa04670;hsa04810;hsa05146;hsa05203;hsa05412,ACTN4,NA,ACTN4 - mRNA,False\nACTN1 - mRNA,NM_001102.3:2760,SEBELIUS_2_C4636,ACTN1,mRNA,0,,,Syndecan interactions;Nephrin interactions;Platelet degranulation ;Non-integrin membrane-ECM interactions;Cell-extracellular matrix interactions,hsa04510;hsa04520;hsa04530;hsa04670;hsa04810;hsa05146;hsa05203;hsa05412,ACTN1,NA,ACTN1 - mRNA,False\nITGA5 - mRNA,NM_002205.2:925,SEBELIUS_2_C4636,ITGA5,mRNA,0,,,Signal transduction by L1;Elastic fibre formation;Integrin cell surface interactions;Cell surface interactions at the vascular wall,hsa04145;hsa04151;hsa04510;hsa04512;hsa04640;hsa04810;hsa05100;hsa05131;hsa05133;hsa05205;hsa05206;hsa05410;hsa05412;hsa05414,ITGA5,NA,ITGA5 - mRNA,False\nITGB5 - mRNA,NM_002213.3:2560,SEBELIUS_2_C4636,ITGB5,mRNA,0,,,Molecules associated with elastic fibres;Syndecan interactions;Smooth Muscle Contraction;ECM proteoglycans;Integrin cell surface interactions;Elastic fibre formation;Non-integrin membrane-ECM interactions,hsa04145;hsa04151;hsa04510;hsa04512;hsa04810;hsa05205;hsa05410;hsa05412;hsa05414,ITGB5,NA,ITGB5 - mRNA,False\nCDC42 - mRNA,NM_001039802.1:390,SEBELIUS_2_C4636,CDC42,mRNA,0,,,DCC mediated attractive signaling;Factors involved in megakaryocyte development and platelet production;VEGFA-VEGFR2 Pathway;RHO GTPases activate KTN1;CD28 dependent Vav1 pathway;Inactivation of Cdc42 and Rac;Rho GTPase cycle;EPHB-mediated forward signaling;RHO GTPases Activate Formins;RHO GTPases activate PAKs;GPVI-mediated activation cascade;G alpha (1213) signalling events;RHO GTPases activate IQGAPs;Sema4D induced cell migration and growth-cone collapse;Regulation of actin dynamics for phagocytic cup formation;CD28 co-stimulation,hsa04010;hsa04014;hsa04015;hsa04062;hsa04144;hsa04360;hsa04370;hsa04510;hsa04520;hsa04530;hsa04660;hsa04666;hsa04670;hsa04722;hsa04810;hsa04912;hsa04932;hsa04933;hsa05100;hsa05120;hsa05130;hsa05131;hsa05132;hsa05200;hsa05203;hsa05205;hsa05211;hsa05212,CDC42,NA,CDC42 - mRNA,False\nITGA6 - mRNA,NM_000210.1:3065,SEBELIUS_2_C4636,ITGA6,mRNA,0,,,Syndecan interactions;Assembly of collagen fibrils and other multimeric structures;Basigin interactions;Non-integrin membrane-ECM interactions;Cell surface interactions at the vascular wall,hsa04151;hsa04510;hsa04512;hsa04514;hsa04640;hsa04810;hsa05145;hsa05200;hsa05222;hsa05410;hsa05412;hsa05414,ITGA6,NA,ITGA6 - mRNA,False\nPDGFC - mRNA,NM_016205.2:2596,SEBELIUS_2_C4636,PDGFC,mRNA,0,,,Signaling by PDGF,hsa04014;hsa04015;hsa04060;hsa04072;hsa04151;hsa04510;hsa04540;hsa04810;hsa05215;hsa05218;hsa05231,PDGFC,NA,PDGFC - mRNA,False\nCD9 - mRNA,NM_001769.2:405,SEBELIUS_2_C4636,CD9,mRNA,0,,,Platelet degranulation ,hsa04640,CD9,NA,CD9 - mRNA,False\nPSEN1 - mRNA,NM_000021.2:900,SEBELIUS_2_C4636,PSEN1,mRNA,0,,,,hsa04310;hsa04330;hsa04722;hsa05010,PSEN1,NA,PSEN1 - mRNA,False\nCLTC - mRNA,NM_004859.2:290,SEBELIUS_2_C4636,CLTC,mRNA,0,,,MHC class II antigen presentation;PCPCE pathway,hsa04144;hsa05016;hsa05100,CLTC,NA,CLTC - mRNA,False\nFGFR1 - mRNA,NM_015850.2:1335,SEBELIUS_2_C4636,FGFR1,mRNA,0,,,RAFMAP kinase cascade;Constitutive Signaling by Aberrant PI3K in Cancer;SHC-mediated cascade.CL.FGFR1;Phospholipase C-mediated cascade.CL. FGFR1;Signaling by FGFR1 amplification mutants;PI3K Cascade;FGFR1b ligand binding and activation;Signaling by FGFR1 in disease;Downstream signal transduction;GAB1 signalosome;Signaling by SCF-KIT;PI3KAKT activation;Role of LAT2NTALLAB on calcium mobilization;DAP12 signaling;Signaling by PDGF;Signaling by ERBB2;VEGFR2 mediated cell proliferation;Frs2-mediated activation;ARMS-mediated activation;Gastrin-CREB signalling pathway via PKC and MAPK;Interleukin receptor SHC signaling;FCERI mediated MAPK activation;VEGFA-VEGFR2 Pathway;IRS-related events triggered by IGF1R,hsa04010;hsa04014;hsa04015;hsa04151;hsa04520;hsa04550;hsa04810;hsa05200;hsa05205;hsa05215;hsa05218;hsa05230,FGFR1,NA,FGFR1 - mRNA,False\nPTEN - mRNA,NM_000314.4:1351,SEBELIUS_2_C4636,PTEN,mRNA,0,,,Synthesis of PIPs at the plasma membrane;Downstream TCR signaling;PIP3 activates AKT signaling;Downstream signal transduction;GAB1 signalosome;Signaling by SCF-KIT;PI3KAKT activation;Role of LAT2NTALLAB on calcium mobilization;DAP12 signaling;Signaling by PDGF;Signaling by ERBB2,hsa00562;hsa04068;hsa04070;hsa04071;hsa04115;hsa04150;hsa04151;hsa04510;hsa04530;hsa04931;hsa05161;hsa05200;hsa05206;hsa05213;hsa05214;hsa05215;hsa05218;hsa05222;hsa05230,PTEN,NA,PTEN - mRNA,False\nPRKAA1 - mRNA,NM_006251.5:1455,SEBELIUS_2_C4636,PRKAA1,mRNA,0,,,TP53 Regulates Metabolic Genes;PI3K Cascade;IRS-related events triggered by IGF1R,hsa04068;hsa04150;hsa04151;hsa04152;hsa04211;hsa04213;hsa04910;hsa04920;hsa04921;hsa04922;hsa04931;hsa04932;hsa05410,PRKAA1,NA,PRKAA1 - mRNA,False\nPPP1CB - mRNA,NM_002709.2:393,SEBELIUS_2_C4636,PPP1CB,mRNA,0,,,RHO GTPases Activate ROCKs;Downregulation of TGF-beta receptor signaling;RHO GTPases activate PKNs;Circadian Clock;Regulation of PLK1 Activity at G2M Transition;TGF-beta receptor signaling activates SMADs,hsa03015;hsa04022;hsa04024;hsa04114;hsa04261;hsa04270;hsa04390;hsa04510;hsa04611;hsa04720;hsa04728;hsa04750;hsa04810;hsa04910;hsa04921;hsa04931;hsa05031;hsa05034;hsa05168;hsa05205,PPP1CB,NA,PPP1CB - mRNA,False\nSPTAN1 - mRNA,NM_003127.2:7610,SEBELIUS_2_C4636,SPTAN1,mRNA,0,,,Nephrin interactions;Caspase-mediated cleavage of cytoskeletal proteins;Downstream signal transduction;VEGFR2 mediated cell proliferation;Signaling by SCF-KIT;Frs2-mediated activation;ARMS-mediated activation;Gastrin-CREB signalling pathway via PKC and MAPK;Interleukin receptor SHC signaling;FCERI mediated MAPK activation;DAP12 signaling;Signaling by PDGF;VEGFA-VEGFR2 Pathway;Signaling by ERBB2;IRS-related events triggered by IGF1R;Apoptotic cleavage of cellular proteins,hsa04530,SPTAN1,NA,SPTAN1 - mRNA,False\nEPB41L2 - mRNA,NM_001199389.1:1880,SEBELIUS_2_C4636,EPB41L2,mRNA,0,,,,hsa04530,EPB41L2,NA,EPB41L2 - mRNA,False\nWASF2 - mRNA,NM_006990.2:485,SEBELIUS_2_C4636,WASF2,mRNA,0,,,RHO GTPases Activate WASPs and WAVEs;VEGFA-VEGFR2 Pathway;Regulation of actin dynamics for phagocytic cup formation,hsa04520;hsa04666;hsa04810;hsa05100;hsa05131;hsa05132;hsa05231,WASF2,NA,WASF2 - mRNA,False\nTGFB1 - mRNA,NM_000660.3:1260,SEBELIUS_2_C4636,TGFB1,mRNA,0,,,TGF-beta receptor signaling in EMT (epithelial to mesenchymal transition);TGFBR1 LBD Mutants in Cancer;Influenza Virus Induced Apoptosis;Molecules associated with elastic fibres;Syndecan interactions;ECM proteoglycans;Transcriptional regulation of white adipocyte differentiation;TGF-beta receptor signaling activates SMADs;Elastic fibre formation;Non-integrin membrane-ECM interactions,hsa04010;hsa04060;hsa04068;hsa04110;hsa04144;hsa04350;hsa04380;hsa04390;hsa04932;hsa04933;hsa05140;hsa05142;hsa05145;hsa05146;hsa05152;hsa05161;hsa05166;hsa05200;hsa05205;hsa05210;hsa05211;hsa05212;hsa05220;hsa05321;hsa05410;hsa05414,TGFB1,NA,TGFB1 - mRNA,False\nTUBB - mRNA,NM_178014.2:1955,SEBELIUS_2_C4636,TUBB,mRNA,0,,,,hsa04145;hsa04540;hsa05130,TUBB,NA,TUBB - mRNA,False\nSMAD3 - mRNA,NM_005902.3:4220,SEBELIUS_2_C4636,SMAD3,mRNA,0,,,Signaling by NODAL;TGF-beta receptor signaling activates SMADs,hsa04068;hsa04110;hsa04144;hsa04310;hsa04350;hsa04390;hsa04520;hsa04550;hsa04933;hsa05142;hsa05161;hsa05166;hsa05200;hsa05210;hsa05212;hsa05220;hsa05321,SMAD3,NA,SMAD3 - mRNA,False\nMAPK1 - mRNA,NM_138957.2:430,SEBELIUS_2_C4636,MAPK1,mRNA,0,,,Negative regulation of FGFR1 signaling;Signal transduction by L1;Negative regulation of FGFR2 signaling;MAP2K and MAPK activation;RHO GTPases Activate WASPs and WAVEs;Senescence-Associated Secretory Phenotype (SASP);Regulation of HSF1-mediated heat shock response;Thrombin signalling through proteinase activated receptors (PARs);Negative regulation of FGFR4 signaling;Oxidative Stress Induced Senescence;Signal attenuation;Negative regulation of FGFR3 signaling;Negative feedback regulation of MAPK pathway;Growth hormone receptor signaling;PLC beta mediated events;FCERI mediated MAPK activation;CREB phosphorylation through the activation of Ras;Negative regulation of MAPK pathway;RAFMAP kinase cascade;Downstream signal transduction;VEGFR2 mediated cell proliferation;Signaling by SCF-KIT;Frs2-mediated activation;ARMS-mediated activation;Gastrin-CREB signalling pathway via PKC and MAPK;Interleukin receptor SHC signaling;DAP12 signaling;Signaling by PDGF;VEGFA-VEGFR2 Pathway;Signaling by ERBB2;IRS-related events triggered by IGF1R,hsa04010;hsa04012;hsa04014;hsa04015;hsa04022;hsa04024;hsa04062;hsa04066;hsa04068;hsa04071;hsa04072;hsa04114;hsa04150;hsa04151;hsa04261;hsa04270;hsa04320;hsa04350;hsa04360;hsa04370;hsa04380;hsa04510;hsa04520;hsa04540;hsa04550;hsa04611;hsa04620;hsa04621;hsa04650;hsa04660;hsa04662;hsa04664;hsa04666;hsa04668;hsa04713;hsa04720;hsa04722;hsa04723;hsa04724;hsa04725;hsa04726;hsa04730;hsa04810;hsa04910;hsa04912;hsa04914;hsa04915;hsa04916;hsa04917;hsa04919;hsa04921;hsa04930;hsa04933;hsa04960;hsa05010;hsa05020;hsa05034;hsa05131;hsa05132;hsa05133;hsa05140;hsa05142;hsa05145;hsa05152;hsa05160;hsa05161;hsa05164;hsa05200;hsa05203;hsa05205;hsa05206;hsa05210;hsa05211;hsa05212;hsa05213;hsa05214;hsa05215;hsa05216;hsa05218;hsa05219;hsa05220;hsa05221;hsa05223;hsa05230;hsa05231,MAPK1,NA,MAPK1 - mRNA,False\nHSPB1 - mRNA,NM_001540.3:374,SEBELIUS_2_C4636,HSPB1,mRNA,0,,,MAPK6MAPK4 signaling;VEGFA-VEGFR2 Pathway,hsa04010;hsa04370;hsa05146;hsa05169,HSPB1,NA,HSPB1 - mRNA,False\nG6PD - mRNA,NM_000402.2:1155,SEBELIUS_2_C4636,G6PD,mRNA,0,,,TP53 Regulates Metabolic Genes,hsa01100;hsa05230,G6PD,NA,G6PD - mRNA,False\nPDGFRB - mRNA,NM_002609.3:840,SEBELIUS_2_C4636,PDGFRB,mRNA,0,,,PIP3 activates AKT signaling;Constitutive Signaling by Aberrant PI3K in Cancer;RAFMAP kinase cascade;Signaling by PDGF;Downstream signal transduction;GAB1 signalosome;Signaling by SCF-KIT;PI3KAKT activation;Role of LAT2NTALLAB on calcium mobilization;DAP12 signaling;Signaling by ERBB2;VEGFR2 mediated cell proliferation;Frs2-mediated activation;ARMS-mediated activation;Gastrin-CREB signalling pathway via PKC and MAPK;Interleukin receptor SHC signaling;FCERI mediated MAPK activation;VEGFA-VEGFR2 Pathway;IRS-related events triggered by IGF1R,hsa04010;hsa04014;hsa04015;hsa04020;hsa04060;hsa04072;hsa04151;hsa04510;hsa04540;hsa04810;hsa05166;hsa05200;hsa05206;hsa05214;hsa05215;hsa05218;hsa05230;hsa05231,PDGFRB,NA,PDGFRB - mRNA,False\nIQGAP1 - mRNA,NM_003870.3:820,SEBELIUS_2_C4636,IQGAP1,mRNA,0,,,MAP2K and MAPK activation;Nephrin interactions;RHO GTPases activate IQGAPs;RAFMAP kinase cascade;Downstream signal transduction;VEGFR2 mediated cell proliferation;Signaling by SCF-KIT;Frs2-mediated activation;ARMS-mediated activation;Gastrin-CREB signalling pathway via PKC and MAPK;Interleukin receptor SHC signaling;FCERI mediated MAPK activation;DAP12 signaling;Signaling by PDGF;VEGFA-VEGFR2 Pathway;Signaling by ERBB2;IRS-related events triggered by IGF1R,hsa04520;hsa04810;hsa05205,IQGAP1,NA,IQGAP1 - mRNA,False\nADAM9 - mRNA,NM_001005845.1:695,SEBELIUS_2_C4636,ADAM9,mRNA,0,,,,,ADAM9,NA,ADAM9 - mRNA,False\nNCKAP1 - mRNA,NM_013436.3:2353,SEBELIUS_2_C4636,NCKAP1,mRNA,0,,,RHO GTPases Activate WASPs and WAVEs;VEGFA-VEGFR2 Pathway;Regulation of actin dynamics for phagocytic cup formation,hsa04810,NCKAP1,NA,NCKAP1 - mRNA,False\nPPP2R1A - mRNA,NM_014225.3:1440,SEBELIUS_2_C4636,PPP2R1A,mRNA,0,,,MASTL Facilitates Mitotic Progression;Regulation of PLK1 Activity at G2M Transition;Disassembly of the destruction complex and recruitment of AXIN to the membrane;S37 mutants of beta-catenin aren.SQT.t phosphorylated;Cyclin AB1 associated events during G2M transition;RAF activation;RHO GTPases Activate Formins;S33 mutants of beta-catenin aren.SQT.t phosphorylated;Loss of Nlp from mitotic centrosomes;DARPP-32 events;Beta-catenin phosphorylation cascade;Initiation of Nuclear Envelope Reformation;Recruitment of mitotic centrosome proteins and complexes;Spry regulation of FGF signaling;Truncations of AMER1 destabilize the destruction complex;CTLA4 inhibitory signaling;AXIN missense mutants destabilize the destruction complex;S45 mutants of beta-catenin aren.SQT.t phosphorylated;Anchoring of the basal body to the plasma membrane;Nonsense Mediated Decay (NMD) enhanced by the Exon Junction Complex (EJC);Cyclin D associated events in G1;Platelet sensitization by LDL;Resolution of Sister Chromatid Cohesion;Inhibition of replication initiation of damaged DNA by RB1E2F1;RAFMAP kinase cascade;Downstream signal transduction;VEGFR2 mediated cell proliferation;Signaling by SCF-KIT;Frs2-mediated activation;ARMS-mediated activation;Gastrin-CREB signalling pathway via PKC and MAPK;Interleukin receptor SHC signaling;FCERI mediated MAPK activation;DAP12 signaling;Signaling by PDGF;VEGFA-VEGFR2 Pathway;Signaling by ERBB2;IRS-related events triggered by IGF1R;Negative regulation of FGFR3 signaling;Negative regulation of FGFR4 signaling;Negative regulation of FGFR2 signaling;Negative regulation of FGFR1 signaling,hsa03015;hsa04071;hsa04114;hsa04151;hsa04152;hsa04261;hsa04350;hsa04390;hsa04530;hsa04728;hsa04730;hsa05142;hsa05160,PPP2R1A,NA,PPP2R1A - mRNA,False\nRAC1 - mRNA,NM_198829.1:1250,SEBELIUS_2_C4636,RAC1,mRNA,0,,,MAPK6MAPK4 signaling;Signaling by SCF-KIT;Inactivation of Cdc42 and Rac;DAP12 signaling;Ephrin signaling;EPH-ephrin mediated repulsion of cells;Activation of Rac;RHO GTPases Activate Formins;VEGFR2 mediated vascular permeability;RHO GTPases activate PAKs;Sema4D mediated inhibition of cell attachment and migration;G alpha (1213) signalling events;Regulation of actin dynamics for phagocytic cup formation;Sema4D induced cell migration and growth-cone collapse;Signal transduction by L1;RHO GTPases activate CIT;RHO GTPases Activate WASPs and WAVEs;Factors involved in megakaryocyte development and platelet production;RHO GTPases activate KTN1;CD28 dependent Vav1 pathway;DSCAM interactions;NRAGE signals death through JNK;GPVI-mediated activation cascade;Sema3A PAK dependent Axon repulsion;RHO GTPases activate PKNs;SEMA3A-Plexin repulsion signaling by inhibiting Integrin adhesion;RHO GTPases activate IQGAPs;PCPCE pathway;VEGFA-VEGFR2 Pathway;CD28 co-stimulation,hsa04010;hsa04014;hsa04015;hsa04024;hsa04062;hsa04071;hsa04145;hsa04151;hsa04310;hsa04360;hsa04370;hsa04380;hsa04510;hsa04520;hsa04620;hsa04650;hsa04662;hsa04664;hsa04666;hsa04670;hsa04722;hsa04810;hsa04932;hsa04933;hsa04972;hsa05100;hsa05120;hsa05131;hsa05132;hsa05200;hsa05203;hsa05205;hsa05210;hsa05211;hsa05212;hsa05231;hsa05416,RAC1,NA,RAC1 - mRNA,False\nGNAI1 - mRNA,NM_002069.4:670,SEBELIUS_2_C4636,GNAI1,mRNA,0,,,ADP signalling through P2Y purinoceptor 12;Adrenaline#noradrenaline inhibits insulin secretion,hsa04015;hsa04022;hsa04024;hsa04062;hsa04071;hsa04261;hsa04360;hsa04530;hsa04540;hsa04611;hsa04670;hsa04713;hsa04723;hsa04724;hsa04725;hsa04726;hsa04727;hsa04728;hsa04730;hsa04914;hsa04915;hsa04916;hsa04921;hsa04923;hsa04971;hsa05032;hsa05034;hsa05133;hsa05142;hsa05145;hsa05200,GNAI1,NA,GNAI1 - mRNA,False\nPPP2CB - mRNA,NM_001009552.1:1075,SEBELIUS_2_C4636,PPP2CB,mRNA,0,,,MASTL Facilitates Mitotic Progression;Misspliced GSK3beta mutants stabilize beta-catenin;T41 mutants of beta-catenin aren.SQT.t phosphorylated;ERKs are inactivated;Glycolysis;Negative regulation of MAPK pathway;Disassembly of the destruction complex and recruitment of AXIN to the membrane;S37 mutants of beta-catenin aren.SQT.t phosphorylated;Cyclin AB1 associated events during G2M transition;RAF activation;RHO GTPases Activate Formins;S33 mutants of beta-catenin aren.SQT.t phosphorylated;DARPP-32 events;Beta-catenin phosphorylation cascade;Spry regulation of FGF signaling;Truncations of AMER1 destabilize the destruction complex;CTLA4 inhibitory signaling;AXIN missense mutants destabilize the destruction complex;S45 mutants of beta-catenin aren.SQT.t phosphorylated;Cyclin D associated events in G1;Platelet sensitization by LDL;Resolution of Sister Chromatid Cohesion;Inhibition of replication initiation of damaged DNA by RB1E2F1;RAFMAP kinase cascade;Downstream signal transduction;VEGFR2 mediated cell proliferation;Signaling by SCF-KIT;Frs2-mediated activation;ARMS-mediated activation;Gastrin-CREB signalling pathway via PKC and MAPK;Interleukin receptor SHC signaling;FCERI mediated MAPK activation;DAP12 signaling;Signaling by PDGF;VEGFA-VEGFR2 Pathway;Signaling by ERBB2;IRS-related events triggered by IGF1R;Negative regulation of FGFR3 signaling;Negative regulation of FGFR4 signaling;Negative regulation of FGFR2 signaling;Negative regulation of FGFR1 signaling,hsa03015;hsa04071;hsa04114;hsa04151;hsa04152;hsa04261;hsa04350;hsa04390;hsa04530;hsa04728;hsa04730;hsa05142;hsa05160,PPP2CB,NA,PPP2CB - mRNA,False\nGSTM3 - mRNA,NM_000849.4:602,SEBELIUS_2_C4636,GSTM3,mRNA,0,,,,,GSTM3,NA,GSTM3 - mRNA,False\nRHOA - mRNA,NM_001664.2:1230,SEBELIUS_2_C4636,RHOA,mRNA,0,,,RHO GTPases activate KTN1;EPHB-mediated forward signaling;Rho GTPase cycle;EPHA-mediated growth cone collapse;RHO GTPases Activate ROCKs;Axonal growth inhibition (RHOA activation);PI3KAKT activation;G beta.CL.gamma signalling through PI3Kgamma;RHO GTPases Activate Formins;RHO GTPases activate PKNs;GPVI-mediated activation cascade;G alpha (1213) signalling events;Sema4D mediated inhibition of cell attachment and migration;PCPCE pathway;Sema4D induced cell migration and growth-cone collapse,hsa04014;hsa04015;hsa04022;hsa04024;hsa04062;hsa04071;hsa04072;hsa04144;hsa04270;hsa04310;hsa04350;hsa04360;hsa04510;hsa04520;hsa04530;hsa04611;hsa04660;hsa04670;hsa04722;hsa04810;hsa04921;hsa04972;hsa05100;hsa05130;hsa05133;hsa05152;hsa05200;hsa05203;hsa05205;hsa05206;hsa05210,RHOA,NA,RHOA - mRNA,False\nRAB11A - mRNA,NM_004663.3:710,SEBELIUS_2_C4636,RAB11A,mRNA,0,,,Anchoring of the basal body to the plasma membrane,hsa04144;hsa04972,RAB11A,NA,RAB11A - mRNA,False\n";
```
