## Supplementary figures and images for "Differential gene expression and functional pathway enrichment in colon cell line CCD 841 CoN (CRL-1790) transfected with miR-mimics miR-18b, miR-142-3p, miR-155, and miR-890"

### color legend - New.Annotation.png

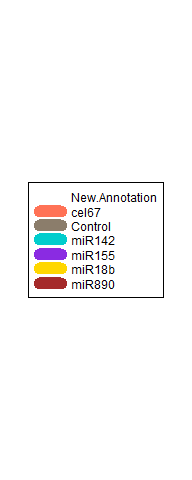

### color legend.png

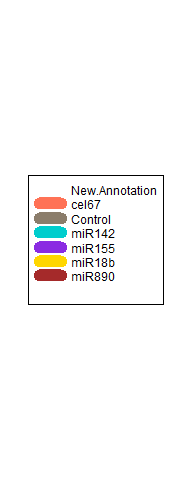

### logo_nanostring.png

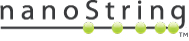

### logo_nanostring_Flat_189x40.png

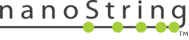

### logo_nanostring_white_Flat.png

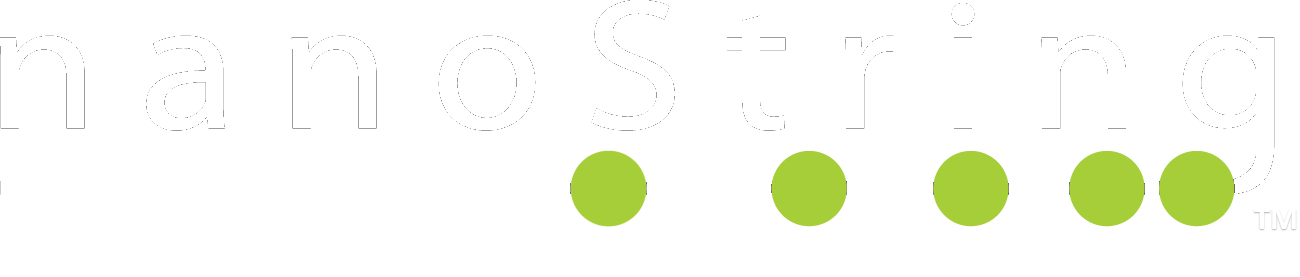

### logo_nanostring_white_Flat_189x40.png

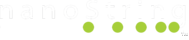

### nanostring_icon.png

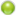

### ui-bg_flat_0_aaaaaa_40x100.png

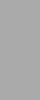

### ui-bg_flat_75_ffffff_40x100.png

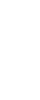

### ui-icons_2e83ff_256x240.png

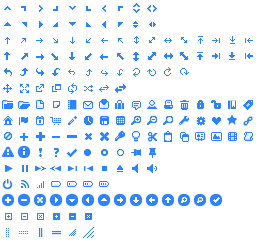

### ui-icons_222222_256x240.png

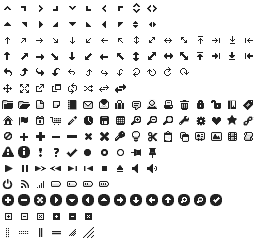

### ui-icons_454545_256x240.png

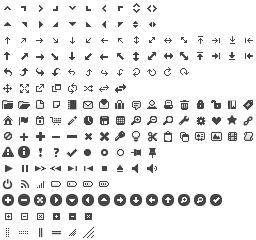

### ui-icons_888888_256x240.png

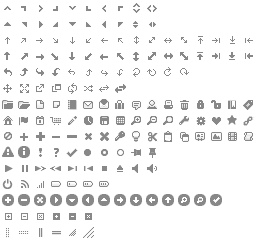

### ui-icons_cd0a0a_256x240.png

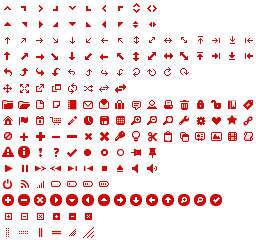

### volcano plotNew.AnnotationControl.png

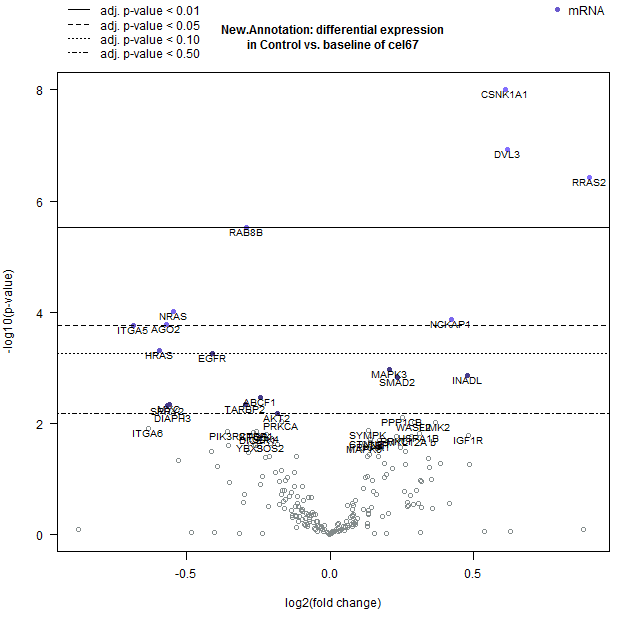

### volcano plotNew.AnnotationmiR18b.png

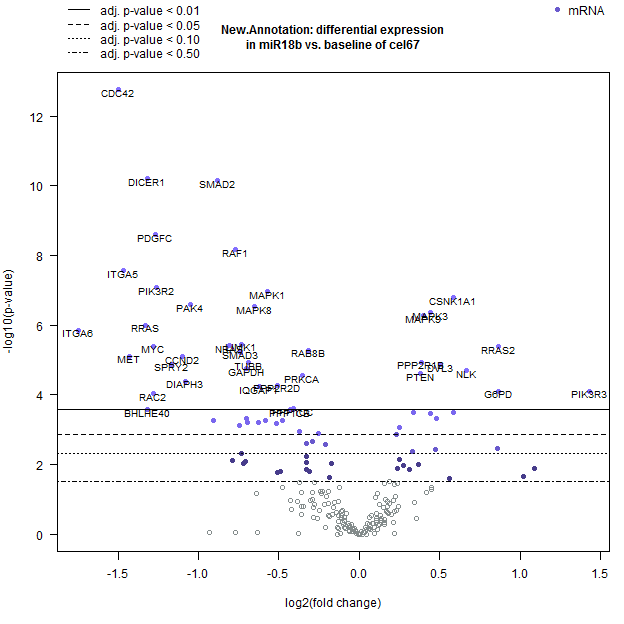

### volcano plotNew.AnnotationmiR142.png

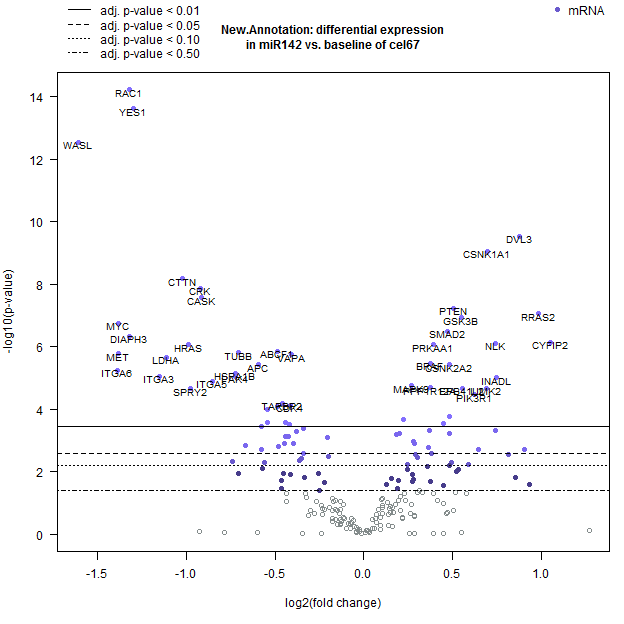

### volcano plotNew.AnnotationmiR155.png

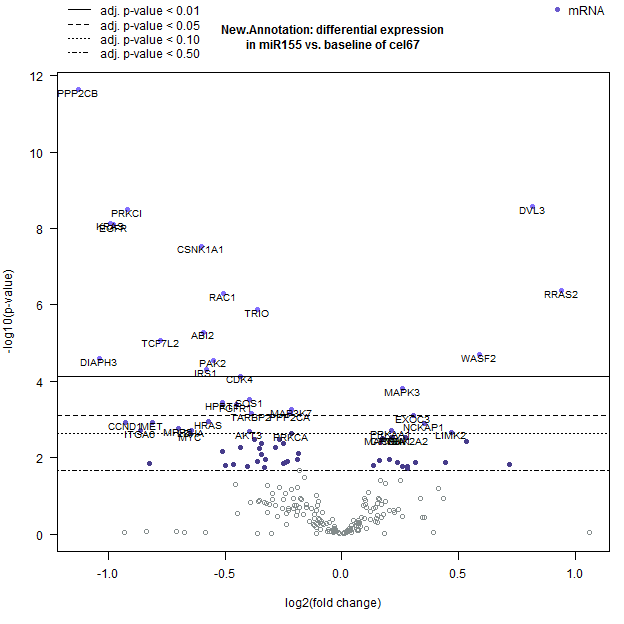

### volcano plotNew.AnnotationmiR890.png

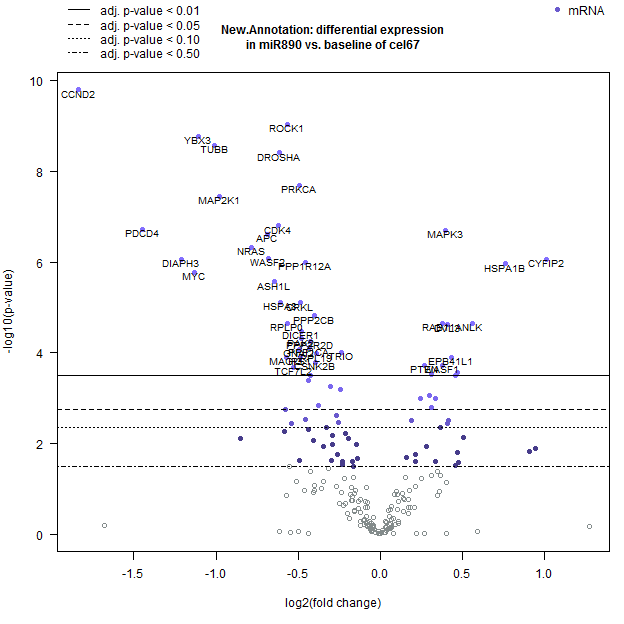
